# Effects of water, sanitation, handwashing and nutritional interventions on soil-transmitted helminth infections in young children: a cluster-randomized controlled trial in rural Bangladesh

**DOI:** 10.1101/512509

**Authors:** Ayse Ercumen, Jade Benjamin-Chung, Benjamin F. Arnold, Audrie Lin, Alan E. Hubbard, Christine Stewart, Zahidur Rahman, Sarker Masud Parvez, Leanne Unicomb, Mahbubur Rahman, Rashidul Haque, John M. Colford, Stephen P. Luby

## Abstract

**Background:** Soil transmitted helminths (STH) infect >1.5 billion people. Mass drug administration (MDA) reduces infection; however, drug resistance is emerging and reinfection occurs rapidly. We conducted a randomized controlled trial in Bangladesh (WASH Benefits, NCT01590095) to assess whether water, sanitation, hygiene and nutrition interventions, alone and combined, reduce STH in a setting with ongoing MDA.

**Methodology/Principal Findings:** We randomized clusters of pregnant women into water treatment, sanitation, handwashing, combined water+sanitation+handwashing (WSH), nutrition, nutrition+WSH (N+WSH) or control arms. After 2.5 years of intervention, we enumerated STH infections in children aged 2-12 years with Kato-Katz. We estimated intention-to-treat intervention effects on infection prevalence and intensity. Participants and field staff were not blinded; laboratory technicians and data analysts were blinded.

In 2012-2013, we randomized 5551 women in 720 clusters. In 2015-2016, we enrolled 7795 children of 4102 available women for STH follow-up and collected stool from 7187. Prevalence among controls was 36.8% for *A. lumbricoides*, 9.2% for hookworm and 7.5% for *T. trichiura*. Most infections were low-intensity. Compared to controls, the water intervention reduced hookworm (prevalence ratio [PR]=0.69 (0.50, 0.95), prevalence difference [PD]=−2.83 (−5.16, −0.50)) but did not affect other STH. Sanitation improvements reduced *T. trichiura* (PR=0.71 (0.52, 0.98), PD=−2.17 (−4.03, −0.38)), had a similar borderline effect on hookworm and no effect on *A. lumbricoides*. Handwashing and nutrition interventions did not reduce any STH. WSH and N+WSH reduced hookworm prevalence by 29-33% (2-3 percentage points) and marginally reduced *A. lumbricoides*. Effects on infection intensity were similar.

**Conclusions/Significance:** In a low-intensity infection setting with MDA, we found modest but sustained hookworm reduction from water treatment, sanitation and combined WSH interventions. Interventions more effectively reduced STH species with no persistent environmental reservoirs. Our findings highlight waterborne transmission for hookworm and suggest that water treatment and sanitation improvements can augment MDA programs to interrupt STH transmission.

**Author summary:** Soil-transmitted helminths (STH) infect >1.5 billion people worldwide. Mass-administration of deworming drugs is the cornerstone of global strategy for STH control but treated individuals often rapidly get reinfected and there is also concern about emerging drug resistance. Interventions to treat drinking water, wash hands at critical times and isolate human feces from the environment through improved sanitation could reduce STH transmission by reducing the spread of ova from the feces of infected individuals into the environment and subsequently to new hosts, while nutrition improvements could reduce host susceptibility to infection. Existing evidence on the effect of these interventions on STH is scarce. In a setting with ongoing mass-drug administration, we assessed the effect of individual and combined water, sanitation, handwashing and nutrition interventions on STH infection in children. Approximately 2.5 years after delivering interventions, we found reductions in STH infection from water treatment and sanitation interventions; there was no reduction from the handwashing and nutrition interventions. While the reductions were modest in magnitude compared to cure rates achieved by deworming drugs, they indicated sustained reduction in environmental transmission. The reductions were more pronounced for STH species that do not have persistent environmental reservoirs. These findings suggest that water treatment and sanitation interventions can augment mass-drug administration programs in striving toward elimination of STH.

## Introduction

Soil transmitted helminths (STH), specifically *Ascaris lumbricoides* (roundworm), *Trichuris trichiura* (whipworm), and *Necator americanus* and *Ancylostoma duodenale* (hookworms), infect >1.5 billion people worldwide [1]. Deworming with mass drug administration (MDA) is the cornerstone of global policy for STH control and effectively reduces infection [2]. However, developing drug resistance threatens the effectiveness of MDA programs given the wide-scale use, inadequate monitoring and limited number of effective anthelminthics, and frequent anthelminthic resistance in livestock [3]. Additionally, without environmental interventions to interrupt transmission, rapid reinfection is common; a systematic review demonstrated that prevalence reverts to 94% of pre-treatment levels for *A. lumbricoides*, 82% for *T. trichiura* and 57% for hookworm within 12 months post-treatment [4].

Water, sanitation and hygiene improvements could potentially complement MDA programs in reducing STH transmission. Two systematic reviews found reduced STH infection associated with improved water, sanitation and hygiene conditions in observational studies [5,6]; however, there are few randomized assessments of the effect of water, sanitation and hygiene interventions on STH [3]. School-based hygiene education trials have had mixed effects on STH [7,8] while handwashing with soap and fingernail clipping reduced parasite infections in children in a trial in Ethiopia [9]. Two trials in India found no STH reduction from sanitation improvements, potentially because they did not attain sufficiently high latrine usage [10,11]. It is also possible that persistent environmental reservoirs of STH ova sustain infections given the prolonged survival of some STH species in soil [12]. While sanitation improvements should reduce immediate fecal input into the environment, their protective effect against STH infections may not be apparent until pre-existing ova in the environment are naturally inactivated [13]. Combined water, sanitation and hygiene improvements targeting multiple transmission routes might achieve a larger impact by complementing the primary barrier of sanitation with the secondary barriers of water treatment and handwashing [14,15]. School-based provision of combined water, sanitation and hygiene hardware reduced reinfection with *A. lumbricoides* but not other STH in a Kenyan trial [16]. Another trial on the effect of water, sanitation and hygiene improvements on parasite infection has recently been completed in Timor-Leste [17,18].

The effect of nutrition on STH infections is also poorly understood. Impaired immune function from nutritional deficiencies could increase host susceptibility to STH infection or exacerbate infection severity while nutritional supplements could also increase infection severity as excess nutrients are available for pathogens [19,20]. A systematic review found mixed impact of nutritional supplements on STH infection, concluding that the evidence is scarce and low-quality [19]. Implementing nutrition interventions alongside water, sanitation and hygiene improvements could achieve synergistic benefits against STH.

We conducted a cluster-randomized trial (WASH Benefits, NCT01590095) in Bangladesh to assess the impact of individual and combined water, sanitation, handwashing (WSH) and nutrition interventions on child diarrhea and growth (primary and secondary outcomes) [21]. The trial found that all interventions except for the individual water intervention reduced reported diarrhea, and all interventions with a nutrition component improved linear growth [22]. Here, we report trial findings on STH infections (pre-specified tertiary outcomes) and test the hypotheses whether (1) individual and combined WSH and nutrition interventions reduce STH, (2) combined WSH interventions reduce STH more than individual WSH interventions, and (3) combined nutrition and WSH interventions reduce STH more than nutrition or WSH interventions alone. This work provides a novel investigation of the effect of improved WSH and nutrition on STH in a population with ongoing MDA to inform policy dialogue on whether these can complement MDA programs.

## Methods

### Study setting

The trial was conducted in the Gazipur, Mymensingh, Tangail and Kishoreganj districts of central rural Bangladesh, selected because they had low groundwater arsenic and iron (to not interfere with the trial’s chlorine-based water intervention) and no other water, sanitation, hygiene or nutrition programs. Since 2008, the Bangladesh Ministry of Health has implemented a school-based MDA program that provides deworming to school-aged children while pre-school-aged children receive deworming through the Expanded Program on Immunization (EPI). A 2010 evaluation of the national MDA campaign in two districts (not included in the WASH Benefits trial) found that 63-73% of school-attending children, 11-14% of non-school-attending school-aged children and 60% of pre-school-aged children received deworming [23]. WASH Benefits activities were implemented independently from the MDA and EPI programs. Caregivers reported that 65% of both school-aged and pre-school-aged children enrolled in the trial had been dewormed in the six months prior to our data collection; the percentage of dewormed children was balanced (61-68%) across trial arms.The school-based MDA program offers a single dose of mebendazole biannually while the EPI uses albendazole [23]. In a single dose, both drugs are effective for *A. lumbricoides* but have lower cure rates for *T. trichiura;* for hookworm, albendazole has a high cure rate while mebendazole has a modest cure rate [2,24].

### Randomization and masking

The WASH Benefits trial enrolled pregnant women in their first or second trimester intending to stay in their village for 24 months post-enrollment, with the objective of following the birth cohort (“index children”) born to them. Field staff screened the study area for pregnant women and collected their global positioning system (GPS) coordinates. Eight neighboring eligible women were grouped into clusters using their GPS coordinates. Cluster dimensions were chosen such that one field worker could visit all participants in a cluster in one day. A minimum 1-km buffer was enforced between clusters to minimize spillovers of infections and/or intervention behaviors between study arms. Every eight adjacent clusters enrolled formed a geographic block. An off-site investigator (BFA) used a random number generator to block-randomize clusters into study arms, providing geographically pair-matched randomization. Participants and field staff were not blinded as interventions entailed distinct hardware; blinded technicians enumerated STH outcomes and blinded analysts (AE, JBC) independently replicated data management and analysis. Details of the study design have been previously described [21]. The study protocol and a CONSORT checklist of trial procedures has been provided (Text S1-S2).

### Interventions

Study arms included (1) water treatment: chlorination with sodium dichloroisocyanurate (NaDCC) tablets coupled with safe storage in a narrow-mouth lidded vessel with spigot, (2) sanitation improvements: upgrades to concrete-lined double-pit latrines and provision of child potties and sani-scoops for feces disposal, (3) handwashing promotion: handwashing stations with a water reservoir and a bottle of soapy water mixture at the food preparation and latrine areas, (4) combined water treatment, sanitation and handwashing (WSH), (5) nutrition improvements including exclusive breastfeeding promotion (birth to 6 months), lipid-based nutrient supplements (6-24 months), and age-appropriate maternal, infant, and young child nutrition recommendations (pregnancy to 24 months), (6) nutrition plus combined WSH (N+WSH), and (7) a double-sized control arm with no intervention (Fig 1).

**Fig 1.**
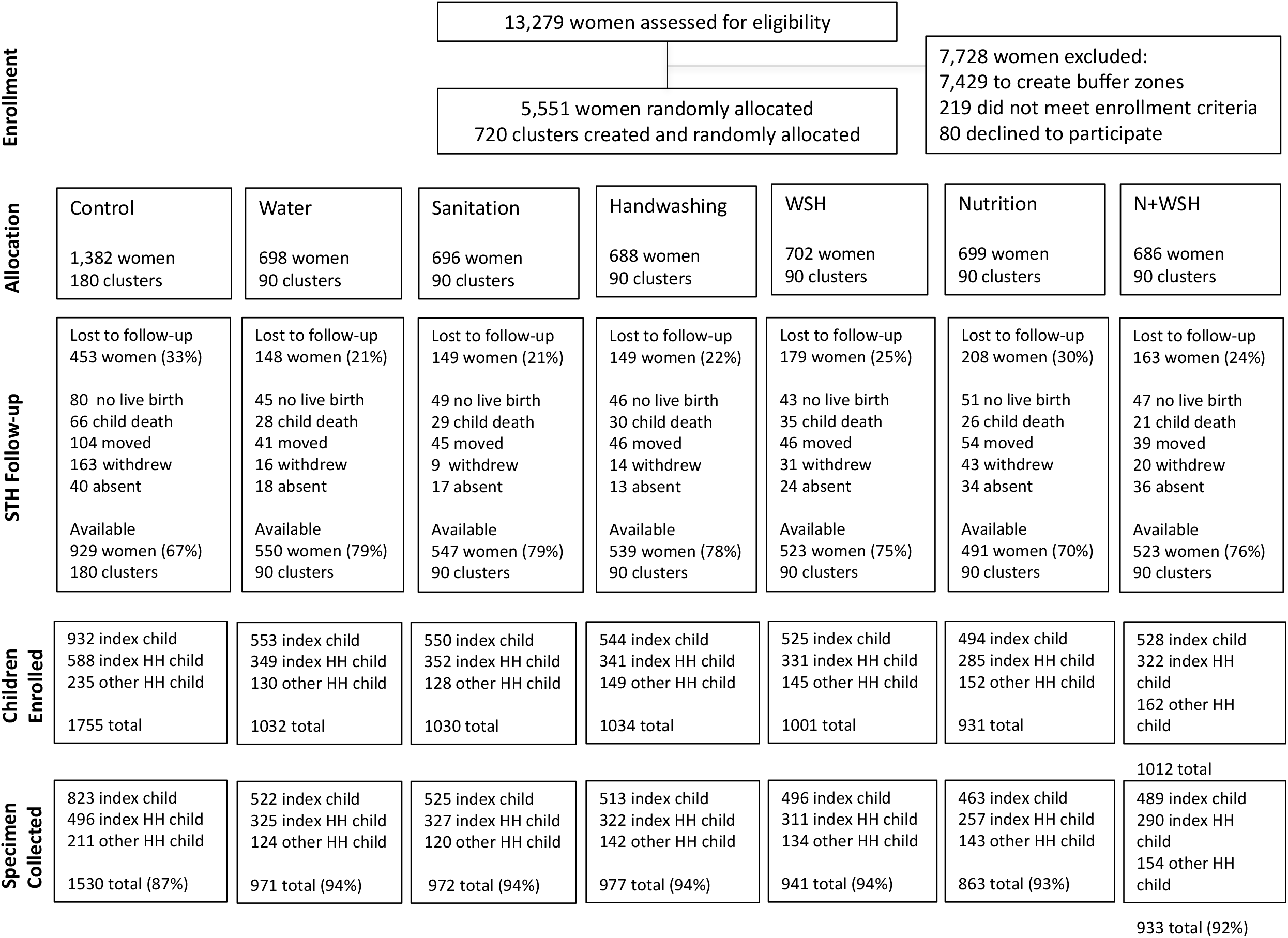
Flowchart of study participation. Legend: Index child refers to the birth cohort born to the enrolled women. Index HH refers to the household where the index child lived. Other HH refers to other households in the compound that contained the index household.

The WSH interventions were delivered around the time of index children’s birth and aimed to reduce their early-life exposure to fecal pathogens. Bangladeshi households are clustered in compounds shared by extended families; in our study, the household where the index child lived (“index household”) was surrounded by an average of 2.5 households per compound. The interventions targeted the compound environment as we expected this to be the primary exposure domain for young children [25]. Interventions were delivered at index child, index household and compound levels (Fig 2). The nutrition intervention targeted index children only. The water and handwashing interventions were delivered to the index household. The sanitation intervention provided upgraded latrines, potties and scoops to all households in the compound; as the shared compound courtyard serves as play space for children, we aimed to improve sanitary conditions in this environment with compound-level latrine coverage. Enrolled compounds made up roughly 10% of a given geographical area because of the eligibility criterion of having a pregnant woman; as such, we did not provide community-level latrine coverage.

**Fig 2.**
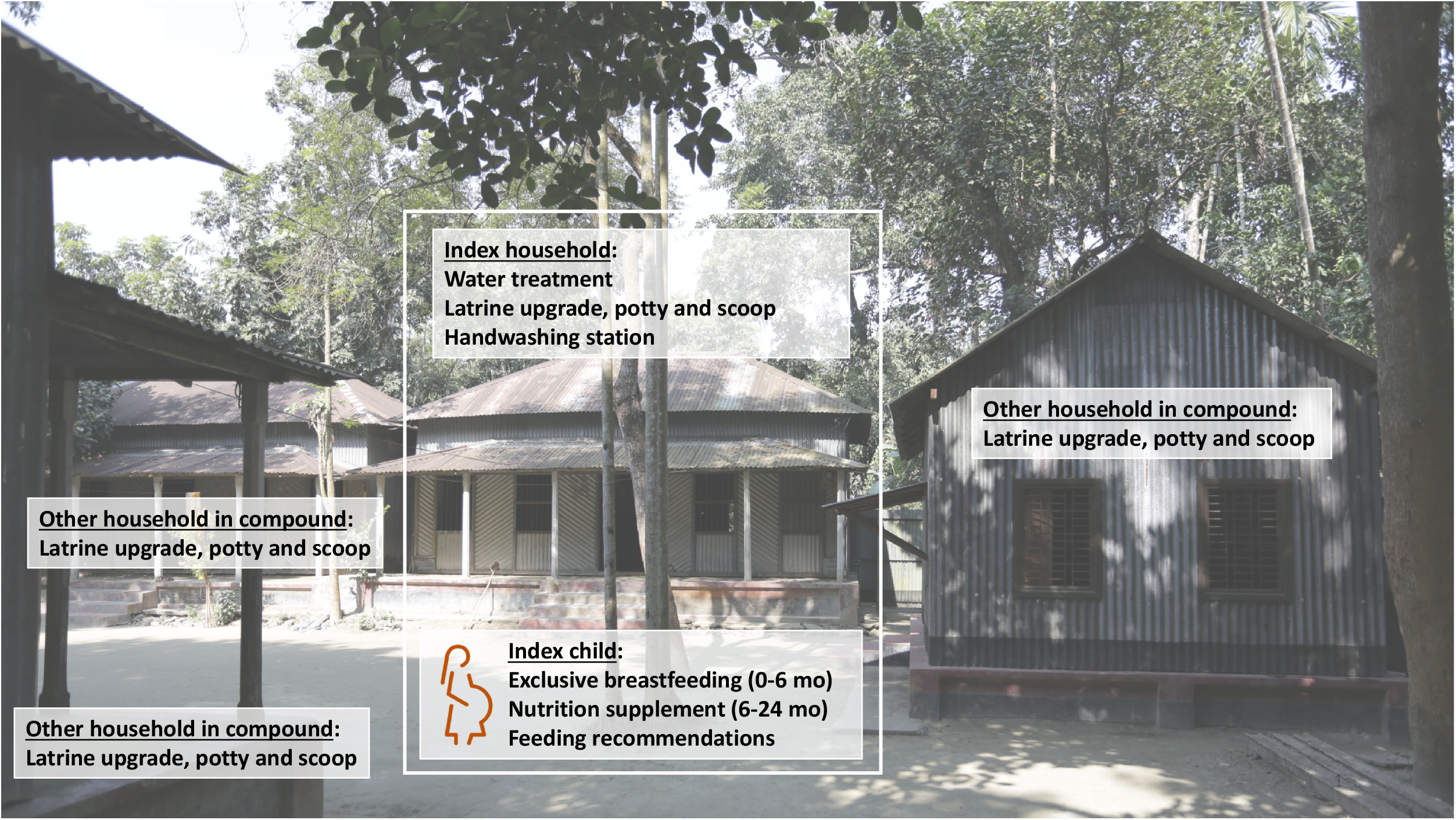
Interventions implemented at index child, index household, and compound levels. Legend: Index child refers to the birth cohort born to enrolled pregnant women. Index household refers to the household where the index child lived. Each enrolled compound contained a single index household and an average of 2.5 households total.

Local women hired and trained as community health promoters visited intervention arm participants on average six times per month to deliver intervention products for free, replenish the supply of consumables (chlorine tablets, soapy water solution, nutrient supplements), resolve hardware problems and encourage adherence to the targeted WSH and nutrition behaviors; health promoters did not visit control arm participants (Text S3). All interventions had high user adherence throughout the study as measured by objective indicators (Text S3). Further details of the interventions and adherence measurements have been previously reported [26–28].

### Outcome assessment

We assessed STH infections in children living in WASH Benefits compounds approximately 2.5 years after intervention initiation. The following children were eligible to enroll in the STH assessment: (1) all index children (aged 30 months on average at follow-up), (2) one child per compound who was aged 18-27 months at trial enrollment (representing expected index child age at follow-up) and (3) one child per compound aged 5-12 years at follow-up (representing school-aged children). Non-index children were enrolled in the preferential order of sibling of index child, child living in index household, or child living in another household in the compound. Households with no live birth or index child death were excluded from intervention promotion and subsequently follow-up.

To measure STH outcomes, field staff distributed sterile containers to primary caregivers of enrolled children, instructed them to collect stool from the following morning’s defecation event, and retrieved the containers on the morning of defecation. If any enrolled child was absent or failed to provide a specimen, field staff returned to the household twice before classifying them as lost to follow-up. After the completion of stool collection in a given compound, all compound members were offered a single dose of albendazole.

Specimens without preservatives were transported on ice to the field laboratory of the International Centre for Diarrhoeal Disease Research, Bangladesh (icddr,b) and analyzed on the same day. Laboratory staff were trained at the icddr,b parasitology laboratory using the Vestergaard Frandsen protocol to perform double-slide Kato-Katz and enumerate ova of *A. lumbricoides*, hookworm (*N. americanus* and *A. duodenale*) and *T. trichiura*. Two slides were prepared from each stool sample and enumerated within 30 minutes of slide preparation [29]. 10% of slides were counted by two technicians (within the 30 minute-window since slide preparation), and 5% were counted by a senior parasitologist (by sending the slides to the icddr,b parasitology laboratory in Dhaka 0-4 days following the original count at the field laboratory) for quality assurance. Two independent technicians double-entered slide counts into a database.

### Ethics

Primary caregivers of children provided written informed consent. Children aged 7-12 years provided written assent. The protocol was approved by human subjects committees at University of California, Berkeley (2011-09-3652), Stanford University (25863), and the icddr,b (PR-11063). A data safety monitoring committee at icddr,b oversaw procedures.

### Registration

WASH Benefits was registered at ClinicalTrials.gov (NCT01590095) in April 2012 before trial enrolment began in May 2012; this registration lists the trial’s primary and secondary outcomes (diarrhea, child growth). The trial design was published in June 2013 before the STH follow-up began in May 2015 and lists STH under tertiary outcomes in Appendix 3 [21]. The pre-specified analysis plan for the STH outcomes was registered at Open Science Framework (OSF, https://osf.io/v2c8p/) in August 2016 before data analysis began.

### Statistical analysis

#### Outcomes

Our pre-specified outcome measures were the infection prevalence, infection intensity and moderate/heavy infection prevalence for each STH species and for any of the three (“any STH”). For each species, we classified stool samples with any ova as positive. We quantified infection intensity in eggs per gram (epg) by multiplying the sum of egg counts from the two duplicated slides by 12. We defined moderate/heavy intensity infections based on WHO categories (≥5,000 epg for *A. lumbricoides*, ≥1,000 epg for hookworm, and ≥2,000 epg for *T. trichiura*) [30]. We assessed the interrater agreement between two independent technicians and between a given technician and the senior parasitologist by calculating the kappa statistic for slides classified as positive [31].

#### Sample size

WASH Benefits was designed to detect effects on child length and diarrhea with a planned sample size of 5040 pregnant women [21]. We assumed that two children per pregnant woman would be eligible for the STH assessment and 70% of children would provide stool. We estimated STH prevalence and intra-class correlation coefficients (ICC) from the literature. With a two-sided *a* of 0.05, we had 80% power to detect the following relative reductions in prevalence between any intervention arm vs. control: 41% for *A. lumbricoides*, 50% for hookworm, 39% for *T. trichiura*, and 18% for any STH.

#### Statistical parameters and estimation strategy

We compared STH outcomes in (1) individual and combined water, sanitation, handwashing and nutrition arms vs. controls (primary hypothesis), (2) combined vs. single WSH intervention arms, and (3) N+WSH vs. WSH and nutrition arms. We estimated prevalence ratios (PR), prevalence differences (PD) and fecal egg count reductions (FECR, defined as the epg ratio minus one) between arms. We estimated FECRs using geometric and arithmetic means; while geometric means prevent extreme data points from skewing means, arithmetic means are more sensitive to high infection intensities thought to correlate with higher morbidity burden and transmission [1]. We estimated the unadjusted parameters using targeted maximum likelihood estimation (TMLE) with influence-curve based standard errors treating clusters as independent units of analysis [32]. Randomization led to extremely good covariate balance [22], and our primary analysis relied on unadjusted estimates. Secondary analyses adjusted for pre-specified covariates using data-adaptive machine learning (see analysis plan). Analyses were intention-to-treat as user uptake of interventions was high. All analyses were conducted using R (version 3.3.2).

#### Subgroup analyses

Our primary analysis included all enrolled children (index children, other children living in index household, and children living in other households in compound). As different interventions were implemented at index child, index household and compound levels, we also conducted subgroup analyses for these three categories of children. The subgroup analysis for index children was pre-specified and the analysis for other children living in the index household vs. the rest of the compound was added post-hoc. We conducted additional pre-specified subgroup analyses by child age, deworming status, household size, wealth, housing materials, and baseline sanitation conditions (see analysis plan for details of subgroup analyses).

#### Missing outcomes

Individuals that were lost at follow-up or failed to submit a specimen were classified as missing. To assess if missingness was differential by study arm and/or covariates, we compared the percentage of missing observations between arms and the enrollment characteristics of those with available vs. missing specimens. We also assessed covariate balance between arms at follow-up. We conducted a complete-case analysis and an inverse probability of censoring-weighted (IPCW) analysis re-weighting the measured population to reflect the original enrolled population (see analysis plan) [32].

## Results

### Enrolment

Fieldworkers identified 13279 pregnant women in the study area (Fig 1). Between May 2012-July 2013, we enrolled and randomized 5551 women in 720 clusters; the rest were excluded to create between-cluster buffers (n=7429), were ineligible (n=219) or refused (n=80). At the STH follow-up in May 2015-2016, 1449 women (26%) were lost because of no live birth (n=361), index child death (n=235), relocation (n=375), absence (n=182) or withdrawal (n=296) (Fig 1). The control arm had higher attrition (33%) than intervention arms combined (24%) as they had more withdrawals (12% vs. 3%). Among 4102 (74%) available women, we enrolled 7795 children in the STH assessment and successfully collected stool from 7187 (92%) (Fig 1). Stool recovery was somewhat lower in controls (87%) than in intervention arms (94%). Enrolment covariates were balanced between arms at follow-up (Table 1) and those with vs. without specimens (Table S1). We enrolled an average of 1.8 children per compound and 5.4 compounds per cluster.

**Table 1.** Enrolment characteristics by intervention group at follow-up

| No. of women: | Control<br>(N=929) | Water<br>(N=550) | Sanitation<br>(N=547) | Handwashing<br>(N=539) | WSH<br>(N=523) | Nutrition<br>(N=491) | N+WSH<br>(N=523) |
| --- | --- | --- | --- | --- | --- | --- | --- |
| **Maternal** |  |  |  |  |  |  |  |
| Age, mean (range) | 24 (15-43) | 24 (15-43) | 24 (15-41) | 24 (15-60) | 25 (15-44) | 24 (15-45) | 24 (14-43) |
| Years of education, mean (range) | 6 (0-15) | 6 (0-14) | 6 (0-17) | 6 (0-16) | 6 (0-14) | 6 (0-16) | 6 (0-14) |
| **Paternal** |  |  |  |  |  |  |  |
| Years of education, mean (range) | 5 (0-16) | 5 (0-16) | 5 (0-17) | 5 (0-16) | 5 (0-16) | 5 (0-16) | 5 (0-16) |
| Works in agriculture, % (n) | 31.4 (292) | 31.5 (173) | 30.7 (168) | 37.5 (202) | 31.0 (162) | 33.6 (165) | 31.2 (163) |
| **Household** |  |  |  |  |  |  |  |
| Number of persons, mean (range) | 5 (2-17) | 5 (2-23) | 5 (2-17) | 5 (2-22) | 5 (1-14) | 5 (2-18) | 5 (2-14) |
| Has electricity, % (n) | 57.9 (538) | 62.7 (345) | 60.5 (331) | 59.7 (322) | 63.1 (330) | 61.3 (301) | 60.6 (317) |
| Has a cement floor, % (n) | 10.0 (93) | 12.0 (66) | 12.1 (66) | 8.0 (43) | 10.7 (56) | 8.6 (42) | 12.1 (63) |
| Acres of agricultural land owned, mean (range) | 0.1 (0.0-2.5) | 0.1 (0.0-2.4) | 0.1 (0.0-3.2) | 0.1 (0.0-2.6) | 0.2 (0.0-3.1) | 0.2 (0.0-2.8) | 0.2 (0.0-8.9) |
| **Drinking water** |  |  |  |  |  |  |  |
| Shallow tubewell primary water source, % (n) | 76.5 (711) | 73.5 (404) | 75.1 (411) | 70.3 (379) | 79.0 (413) | 75.2 (369) | 74.0 (387) |
| Stored water observed at home, % (n) | 46.6 (433) | 51.1 (281) | 47.4 (259) | 48.8 (263) | 41.5 (217) | 41.8 (205) | 48.0 (251) |
| Reported treating water yesterday, % (n) | 0.3 (3) | 0.2 (1) | 0.0 (0) | 0.2 (1) | 0.0 (0) | 0.0 (0) | 0.4 (2) |
| **Sanitation** |  |  |  |  |  |  |  |
| Daily defecating in the open, % (n) |  |  |  |  |  |  |  |
| Adult men | 7.2 (67) | 5.3 (29) | 6.6 (36) | 9.8 (53) | 6.5 (34) | 7.3 (36) | 7.5 (39) |
| Adult women | 4.7 (44) | 2.6 (14) | 4.2 (23) | 5.2 (28) | 4.0 (21) | 5.30 (26) | 3.8 (20) |
| Children: 8-<15 years (N=1743) | 10.2 (38) | 9.5 (21) | 8.9 (22) | 15.5 (37) | 8.3 (19) | 8.3 (17) | 9.5 (22) |
| Children: 3-<8 years (N=2179) | 40.0 (197) | 36.0 (111) | 37.2 (109) | 37.7 (110) | 35.5 (99) | 35.1 (85) | 36.3 (99) |
| Children: 0-<3 years (N=848) | 80.5 (157) | 85.6 (89) | 81.1 (86) | 85.5 (100) | 78.0 (92) | 82.5 (85) | 88.6 (93) |
| Latrine, % (n) |  |  |  |  |  |  |  |
| Owned | 53.4 (496) | 52.9 (291) | 53.4 (292) | 54.6 (294) | 53.0 (277) | 54.2 (266) | 54.1 (283) |
| Concrete slab | 90.4 (840) | 92.7 (510) | 88.3 (483) | 89.6 (483) | 90.1 (471) | 90.0 (440) | 90.3 (472) |
| Functional water seal | 25.3 (235) | 26.4 (145) | 26.0 (142) | 25.4 (137) | 21.0 (110) | 26.5 (130) | 22.6 (118) |
| Visible stool on slab or floor | 48.6 (451) | 44.9 (247) | 44.8 (245) | 43.8 (236) | 52.6 (275) | 46.2 (227) | 49.3 (258) |
| Owned a potty, % (n) | 3.4 (32) | 3.8 (21) | 3.8 (21) | 5.0 (27) | 3.6 (19) | 5.3 (26) | 4.8 (25) |
| Human feces observed in, % (n) |  |  |  |  |  |  |  |
| House | 9.0 (84) | 9.6 (53) | 7.7 (42) | 10.6 (57) | 7.1 (37) | 7.1 (35) | 6.9 (36) |
| Child's play area | 1.4 (13) | 1.1 (6) | 0.9 (5) | 1.1 (6) | 0.8 (4) | 0.6 (3) | 1.2 (6) |
| **Handwashing** |  |  |  |  |  |  |  |
| Has within 6 steps of latrine, % (n) |  |  |  |  |  |  |  |
| Water | 12.8 (119) | 12.0 (66) | 11.9 (65) | 8.5 (46) | 8.2 (43) | 8.6 (42) | 11.7 (61) |
| Soap | 5.5 (51) | 6.9 (38) | 7.5 (41) | 4.8 (26) | 4.6 (24) | 4.5 (22) | 5.9 (31) |
| Has within 6 steps of kitchen, % (n) |  |  |  |  |  |  |  |
| Water | 8.5 (79) | 6.6 (36) | 7.3 (40) | 5.8 (31) | 8.2 (43) | 9.1 (45) | 8.8 (46) |
| Soap | 2.4 (22) | 2.2 (12) | 2.0 (11) | 2.0 (11) | 2.1 (11) | 3.9 (19) | 3.3 (17) |

### Infection prevalence and intensity

STH prevalence among all children in the control arm was 36.8% (n=563) for *A. lumbricoides*, 9.2% (n=142) for hookworm, 7.5% (n=115) for *T. trichiura*, and 43.4% (n=664) for any STH. The geometric mean egg count among controls was 5.2 epg for *A. lumbricoides*, 0.6 epg for hookworm and 0.4 epg for *T. trichiura*. Most infections were low-intensity; moderate/heavy infection prevalence among controls was 4.2% (n=65) for *A. lumbricoides*, 0.1% (n=2) for hookworm and 0.4% (n=6) for *T. trichiura*. The ICC for any STH infection was 18% for children within the same compound and 8% for children within the same cluster of compounds.

### Interventions vs. control

Among all enrolled children, the single water intervention reduced hookworm prevalence by 31% (PR=0.69 (0.50, 0.95); PD=−2.83 (−5.16, −0.50)) from a control prevalence of 9.2% but had no effect on other STH (Fig 3, Table S2). The sanitation intervention reduced *T. trichiura* prevalence by 29% (PR=0.71 (0.52, 0.98); PD=−2.17 (−4.10, −0.24)) from a control prevalence of 7.5% and achieved a similar borderline reduction on hookworm but had no effect on *A. lumbricoides*. Single handwashing or nutrition interventions did not reduce the prevalence of any STH; there was a borderline increase in *A. lumbricoides* prevalence in these arms (Fig 3, Table S2). Combined WSH reduced hookworm prevalence by 29% (PR=0.71 (0.52, 0.99); PD=−2.63 (−4.95, −0.31)) and N+WSH by 33% (PR=0.67 (0.50, 0.91); PD=−3.00 (−5.14, −0.85)). WSH and N+WSH also marginally reduced *A. lumbricoides* by 7-10% but had no effect on *T. trichiura* (Fig 3, Table S2).

**Fig 3.**
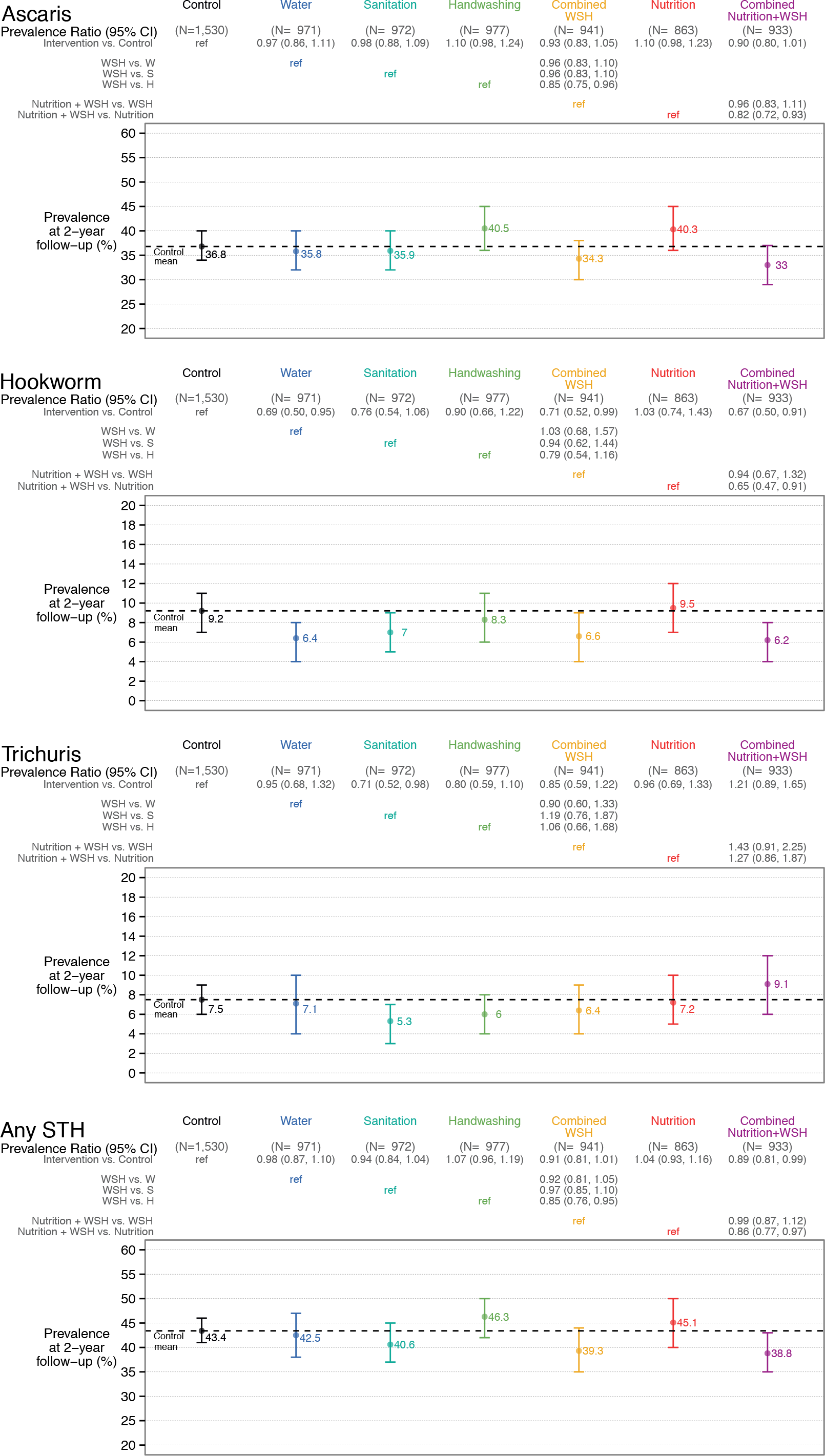
Prevalence ratio for *A. lumbricoides*, hookworm, *T. trichiura* and any STH infection in children aged 2-12 years measured with double-slide Kato-Katz 2.5 years after intervention initiation.

### Combined vs. single interventions

Compared with single water, sanitation and handwashing interventions, combined WSH reduced *A. lumbricoides* more than handwashing alone but this was likely because of the increased *A. lumbricoides* prevalence in the handwashing arm. We found no other benefit from combined WSH vs. its individual components (Fig 3, Table S3). Combined N+WSH reduced *A. lumbricoides* and hookworm prevalence compared to nutrition alone but did not achieve any reduction compared to WSH (Fig 3, Table S4).

### Other effects

Effects on any STH recapitulated *A. lumbricoides* results due to the high prevalence of *A. lumbricoides* compared to the other two species (Fig 3, Tables S2-S4). Interventions did not affect the prevalence of moderate/heavy infections (Tables S5-S7) but we had low power for these rare outcomes. Effects on infection intensity were similar to effects on prevalence, except for a modest reduction in *T. trichiura* intensity from handwashing (Fig 4, Tables S8-S10). Arithmetic means yielded similar results with wider confidence intervals (Tables S8-S10). Unadjusted, adjusted and IPCW estimates were similar for all outcomes (Tables S2-S10).

**Fig 4.**
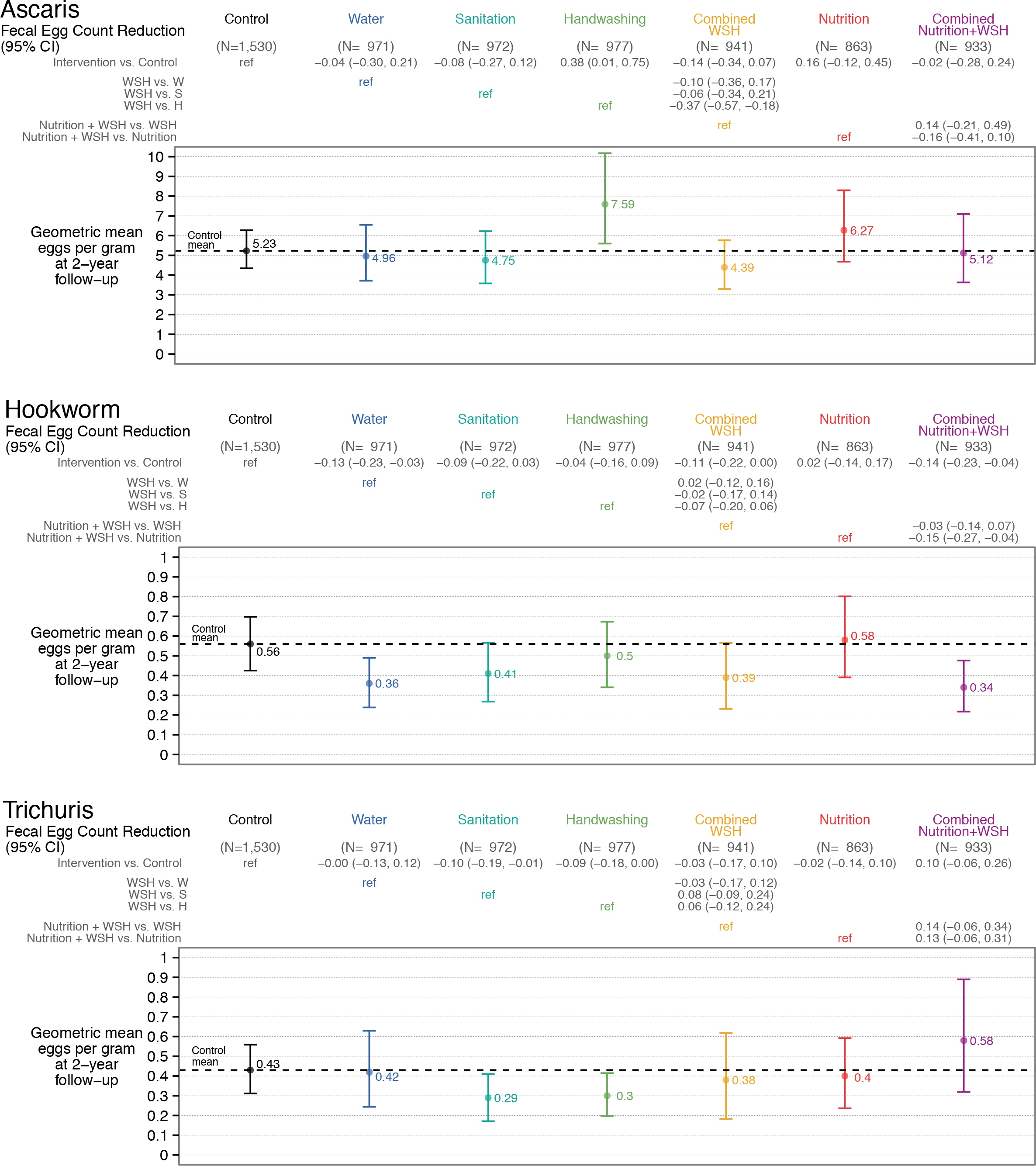
Geometric fecal egg count reduction (FECR) for *A. lumbricoides*, hookworm and *T. trichiura* infection in children aged 2-12 years measured with double-slide Kato-Katz 2.5 years after intervention initiation. Legend: FECR is defined as the ratio of mean egg count per gram minus one.

### Subgroup analyses

Subgroup analyses on index children, other children in the index household and children in other households in the compound yielded findings consistent with those using pooled data from all children. Average age at follow-up was 30 months (range: 22-38) for index children and 7 years (range: 3-12) for non-index children. Non-index children had higher infection prevalence (Table 2), consistent with previous evidence on these age groups [1]. The water intervention, which was implemented in the index household and showed a reduction in hookworm when using data from all enrolled children, substantially reduced hookworm among the older children living in the index household (PR=0.59 (0.39, 0.90)) but not among index children themselves who might have been consuming less water due to their younger age (PR=0.95 (0.52, 1.71)), nor among children in other households in the compound whose own households did not receive the water intervention (PR=0.75 (0.37, 1.51)). The handwashing intervention, which was implemented in the index household and did not achieve a reduction when using data from all children, also did not achieve a reduction among children living in the index household (Table 2). Similarly, the nutrition intervention, which was provided to index children only and did not achieve a reduction when using data from all enrolled children, also did not achieve a reduction among index children (Table 2). As expected, the reduction on hookworm and *T. trichiura* from the compound-level sanitation intervention was similar among all children in the compound (Table 2); however, most confidence intervals crossed the null, reflecting the small sample sizes of the subgroups. Point estimates suggested that the WSH and N+WSH interventions, which reduced hookworm among all children, achieved similar reductions in all three subgroups of children but, once again, most confidence intervals crossed the null due to small sample size (Table 2). Our full set of subgroup findings are reported elsewhere (https://osf.io/v2c8p/); we note that these analyses should be considered exploratory as they had limited statistical power.

**Table 2.** STH prevalence for index children, other children in index household and children in other households in compound

|  | All observations |  |  | Index children<sup>a</sup> |  |  | Other children in index household<sup>b</sup> |  |  | Children in non-index households<sup>c</sup> |  |  |
| --- | --- | --- | --- | --- | --- | --- | --- | --- | --- | --- | --- | --- |
|  | N | Prev | Prevalence ratio | N | Prev | Prevalence ratio | N | Prev | Prevalence ratio | N | Prev | Prevalence ratio |
| **Ascaris** |  |  |  |  |  |  |  |  |  |  |  |  |
| Control | 1530 | 36.8% |  | 823 | 31.3% |  | 496 | 44.4% |  | 211 | 40.3% |  |
| Water | 971 | 35.8% | 0.97 (0.86, 1.11) | 522 | 28.0% | 0.89 (0.74, 1.07) | 325 | 43.4% | 0.98 (0.82, 1.18) | 124 | 49.2% | 1.22 (0.91, 1.64) |
| Sanitation | 972 | 35.9% | 0.98 (0.88, 1.09) | 525 | 30.3% | 0.97 (0.83, 1.13) | 327 | 42.8% | 0.98 (0.84, 1.13) | 120 | 41.7% | 0.96 (0.73, 1.27) |
| Handwashing | 977 | 40.5% | 1.10 (0.98, 1.24) | 513 | 36.5% | 1.16 (0.97, 1.40) | 322 | 45.7% | 1.05 (0.89, 1.23) | 142 | 43.7% | 1.05 (0.80, 1.39) |
| WSH | 941 | 34.3% | 0.93 (0.83, 1.05) | 496 | 28.8% | 0.92 (0.78, 1.09) | 311 | 40.5% | 0.93 (0.78, 1.12) | 134 | 40.3% | 1.00 (0.74, 1.34) |
| Nutrition | 863 | 40.3% | 1.10 (0.98, 1.23) | 463 | 35.0% | 1.12 (0.96, 1.29) | 257 | 49.0% | 1.14 (0.95, 1.37) | 143 | 42.0% | 1.14 (0.84, 1.54) |
| Nutrition + WSH | 933 | 33.0% | 0.90 (0.80, 1.01) | 489 | 26.4% | 0.84 (0.72, 0.99) | 290 | 41.0% | 0.95 (0.80, 1.13) | 154 | 39.0% | 1.02 (0.75, 1.40) |
| **Hookworm** |  |  |  |  |  |  |  |  |  |  |  |  |
| Control | 1530 | 9.2% |  | 823 | 3.6% |  | 496 | 16.1% |  | 211 | 14.7% |  |
| Water | 971 | 6.4% | 0.69 (0.50, 0.95) | 522 | 3.4% | 0.95 (0.52, 1.71) | 325 | 9.5% | 0.59 (0.39, 0.90) | 124 | 10.5% | 0.75 (0.37, 1.51) |
| Sanitation | 972 | 7.0% | 0.76 (0.54, 1.06) | 525 | 2.7% | 0.73 (0.35, 1.51) | 327 | 12.5% | 0.78 (0.52, 1.17) | 120 | 10.8% | 0.78 (0.38, 1.58) |
| Handwashing | 977 | 8.3% | 0.90 (0.66, 1.22) | 513 | 3.5% | 0.96 (0.49, 1.90) | 322 | 13.4% | 0.84 (0.61, 1.16) | 142 | 14.1% | 0.95 (0.54, 1.69) |
| WSH | 941 | 6.6% | 0.71 (0.52, 0.99) | 496 | 3.0% | 0.83 (0.49, 1.40) | 311 | 11.3% | 0.70 (0.46, 1.08) | 134 | 9.0% | 0.63 (0.36, 1.09) |
| Nutrition | 863 | 9.5% | 1.03 (0.74, 1.43) | 463 | 5.0% | 1.36 (0.72, 2.59) | 257 | 14.0% | 0.90 (0.62, 1.30) | 143 | 16.1% | 1.37 (0.78, 2.42) |
| Nutrition + WSH | 933 | 6.2% | 0.67 (0.50, 0.91) | 489 | 3.1% | 0.84 (0.51, 1.38) | 290 | 9.7% | 0.62 (0.41, 0.94) | 154 | 9.7% | 0.58 (0.28, 1.20) |
| **Trichuris** |  |  |  |  |  |  |  |  |  |  |  |  |
| Control | 1530 | 7.5% |  | 823 | 5.2% |  | 496 | 9.5% |  | 211 | 11.8% |  |
| Water | 971 | 7.1% | 0.95 (0.68, 1.32) | 522 | 4.6% | 0.88 (0.54, 1.44) | 325 | 9.5% | 1.02 (0.63, 1.66) | 124 | 11.3% | 0.86 (0.42, 1.78) |
| Sanitation | 972 | 5.3% | 0.71 (0.52, 0.98) | 525 | 3.2% | 0.62 (0.38, 1.00) | 327 | 7.3% | 0.81 (0.49, 1.35) | 120 | 9.2% | 0.72 (0.41, 1.27) |
| Handwashing | 977 | 6.0% | 0.80 (0.59, 1.10) | 513 | 3.7% | 0.71 (0.41, 1.21) | 322 | 9.3% | 0.98 (0.61, 1.59) | 142 | 7.0% | 0.72 (0.34, 1.51) |
| WSH | 941 | 6.4% | 0.85 (0.59, 1.22) | 496 | 5.0% | 0.96 (0.63, 1.47) | 311 | 8.0% | 0.87 (0.54, 1.41) | 134 | 7.5% | 0.69 (0.32, 1.49) |
| Nutrition | 863 | 7.2% | 0.96 (0.69, 1.33) | 463 | 4.5% | 0.87 (0.55, 1.37) | 257 | 8.6% | 0.92 (0.57, 1.50) | 143 | 13.3% | 1.31 (0.61, 2.81) |
| Nutrition + WSH | 933 | 9.1% | 1.21 (0.89, 1.65) | 489 | 6.3% | 1.21 (0.79, 1.86) | 290 | 12.4% | 1.33 (0.88, 2.02) | 154 | 11.7% | 1.04 (0.52, 2.08) |
| **Any STH** |  |  |  |  |  |  |  |  |  |  |  |  |
| Control | 1530 | 43.4% |  | 823 | 35.2% |  | 496 | 54.2% |  | 211 | 49.8% |  |
| Water | 971 | 42.5% | 0.98 (0.87, 1.10) | 522 | 33.0% | 0.94 (0.79, 1.11) | 325 | 52.6% | 0.98 (0.85, 1.14) | 124 | 56.5% | 1.14 (0.89, 1.46) |
| Sanitation | 972 | 40.6% | 0.94 (0.84, 1.04) | 525 | 32.6% | 0.92 (0.79, 1.07) | 327 | 50.2% | 0.95 (0.82, 1.09) | 120 | 50.0% | 0.97 (0.75, 1.24) |
| Handwashing | 977 | 46.3% | 1.07 (0.96, 1.19) | 513 | 38.8% | 1.10 (0.93, 1.31) | 322 | 54.7% | 1.02 (0.89, 1.17) | 142 | 54.2% | 1.07 (0.85, 1.35) |
| WSH | 941 | 39.3% | 0.91 (0.81, 1.01) | 496 | 32.3% | 0.92 (0.79, 1.06) | 311 | 47.6% | 0.90 (0.76, 1.06) | 134 | 46.3% | 0.93 (0.70, 1.23) |
| Nutrition | 863 | 45.1% | 1.04 (0.93, 1.16) | 463 | 38.2% | 1.08 (0.95, 1.24) | 257 | 53.7% | 1.02 (0.86, 1.21) | 143 | 51.7% | 1.14 (0.88, 1.48) |
| Nutrition + WSH | 933 | 38.8% | 0.89 (0.81, 0.99) | 489 | 30.1% | 0.85 (0.74, 0.98) | 290 | 49.3% | 0.93 (0.80, 1.08) | 154 | 46.8% | 0.95 (0.74, 1.23) |

<sup>a</sup> Children born to enrolled pregnant women following enrolment. The nutrition intervention was delivered to index children only.
<sup>b</sup> Other children living in index child's household (index household). These include siblings of index children and children from other mothers in the
same household. The water and handwashing interventions were delivered to index households only.
<sup>c</sup> Children living in other households in index child's compound. The sanitation intervention was delivered to all households in the compound.

### Quality control

The kappa statistic for interrater agreement between two laboratory technicians was 1.00 for *A. lumbricoides* and 0.99 for hookworm and *T. trichiura*. The average kappa statistic for agreement between the laboratory technician that performed the original count and the experienced parasitologist was 0.92 for *A. lumbricoides*, 0.20 for hookworm, and 0.86 for *T. trichiura*. For hookworm, the kappa statistic decreased with the number of days since the slide had been prepared and the original count had been conducted at the field laboratory; the kappa statistic was 1.00 for samples where the experienced parasitologist counted the slides on the day of the original count, 0.33 for samples counted one day later, and 0.11 for samples counted 2-4 days later (Text S4).

## Discussion

### Effect of water treatment

In all intervention arms that included a water treatment component (the individual water treatment, WSH and N+WSH arms), we found a significant reduction in hookworm but not in *A. lumbricoides* or *T. trichiura*. These findings suggest waterborne hookworm transmission in our study population, and water treatment with chlorine combined with safe storage is effective in reducing this transmission. While infections of *A. lumbricoides* or *T. trichiura* are transmitted by ingesting embryonated ova, hookworm ova hatch in soil and larvae infect hosts by penetrating skin; however, one species, *A. duodenale*, can also be transmitted by ingesting larvae [1]. STH ova have been detected in drinking water in low-income countries [33], suggesting a potential reservoir. A systematic review identified three observational studies where water treatment by boiling and filtering was associated with reduced STH infection [6]. While chlorination is generally considered ineffective against STH ova [34], fragile hookworm ova and larvae could be more chlorine-susceptible than the hardier ova of *A. lumbricoides* or *T. trichiura*. The safe storage container with a narrow mouth and lid would also reduce STH contamination of stored drinking water by eliminating contact with hands, which are known reservoirs of ova and larvae [35]. An observational study found increased STH infection associated with unhygienic water storage [36]. Storage could also allow eggs to settle out of the water column before consumption [34]. However, while safe storage should similarly affect all three STH species, *A. lumbricoides* or *T. trichiura* were not reduced by our water intervention, potentially suggesting that the hookworm reduction is due to chlorine rather than safe storage; there are scarce data on the effectiveness of chlorine on hookworm. Alternatively, waterborne transmission could be more important for hookworm in this setting than for *A. lumbricoides* or *T. trichiura*.

### Effect of sanitation

Sanitation improvements isolating human feces from the environment would be expected to reduce the spread of ova into soil and reduce STH transmission by interrupting their life cycle. The WASH Benefits sanitation intervention with concrete-lined double-pit latrines, potties and scoops for feces management reduced *T. trichiura* and achieved a borderline reduction in hookworm but had no effect on*A. lumbricoides*. While the other two arms with a sanitation component (WSH, N+WSH) reduced hookworm to a similar degree as the single sanitation intervention, *T. trichiura* was not affected in these arms; the reduction in the sanitation arm for this species could thus be a chance finding and should be interpreted cautiously. Two previous sanitation trials in India found no impact on STH; however, both studies entailed community-level programs with relatively low adherence [10,11]. Clasen et al. (2014) reported 38% of households in intervention villages having a functional latrine vs. 10% in control villages [10]. Patil et al. (2014) found 41% of households in intervention villages vs. 22% in control villages had improved sanitation and 73-84% of adults in both groups reported daily open defecation [11]. WASH Benefits implemented a compound-level intervention with high adherence across arms. At follow-up, >95% of respondents in the sanitation, WSH and N+WSH arms had a latrine with a functional water seal, compared to <25% of controls [26]. In structured observations, field workers observed >90% of adults in sanitation arms using a hygienic latrine vs. 40% of controls [26]. Low adherence is therefore unlikely to explain the lack of impact on *A. lumbricoides* from sanitation or the lack of *T. trichiura* reduction in the WSH and N+WSH arms. However, it is possible that structured observations overestimated actual latrine use due to respondent reactivity [37]. Also, children continued open defecation despite sanitation access; only 37-54% of young children in sanitation arms were observed to defecate in a latrine or potty vs. 32% of controls [26]. Finally, WASH Benefits intervened on roughly 10% of compounds within a geographical area and did not implement community-level latrine coverage. Bangladesh has a high population density, and contamination with STH ova from surrounding non-study compounds could have entered intervention compounds on shoes/soles of compound residents or via surface runoff. Bangladeshi families also use soil from outside the compound to coat walls and courtyards. Community-level sanitation coverage may be more instrumental in improving child health than individual household sanitation in rural settings [38,39]; it is possible that community-level sanitation coverage is needed to impact STH infections.

The lack of sanitation impact on *A. lumbricoides* could also be due to its prolonged survival in soil, providing a persistent reservoir to sustain infection [13]. Hookworm ova degrade within hours in low-moisture environments [34], while hookworm larvae survive in soil for weeks [40] and *T. trichiura* ova for months [40]. In contrast, *A. lumbricoides* ova can survive in soil for several years in warm and saturated conditions [13]. A pilot assessment among study households found *A. lumbricoides* ova in 67% of courtyard soil samples; of these, 70% developed larvae when incubated (i.e., were viable) [41]. We also found high concentrations of fecal indicator bacteria and evidence of human and animal fecal markers in soil samples from study households, suggesting heavy fecal contamination in the ambient domestic environment [42,43]. We would expect that any reductions in fecal input into the environment would be reflected in a more immediate reduction in infections with hookworm and *T. trichiura* whose larvae/ova are shorter-lived in the environment than those of *A. lumbricoides*, which is consistent with our findings. Any protective effect from sanitation interventions against *A. lumbricoides* infections may not be apparent until existing ova in the environment from pre-intervention contamination are naturally inactivated.

### Effect of handwashing

Handwashing did not reduce STH infection except for a modest reduction in *T. trichiura* intensity. Previous hygiene programs have been shown to reduce STH infections [7–9].Two of the previous studies were conducted in schools [7,8], which may have fewer sources of fecal contamination than the domestic environment and therefore lower risk of re-contamination of hands following handwashing. It is also possible that these studies achieved better handwashing than WASH Benefits, potentially because our promotion primarily targeted caregivers rather than children, some of whom were too young to wash their own hands. A study in Bangladesh found *A. lumbricoides* ova in 51%, *T. trichiura* ova in 23% and hookworm larvae in 26% of fingernails [35], and nail clipping reduced parasite infections among Ethiopian children [9]. While our intervention promotion mentioned washing with soap under fingernails, the practices adopted by participants may not have been sufficient to remove ova/larvae from under nails. This could also explain why the *T. trichiura* intensity but not prevalence was reduced; if the intervention reduced but did not eliminate *T. trichiura* ova on hands, this could lead to a reduced worm burden without affecting prevalence.

### Effect of combined WSH interventions

We found no added benefit from combining WSH interventions. While we did not power the study to statistically detect differences between combined vs. individual interventions, the effect estimates suggest that the combined WSH and N+WSH packages achieved a similar magnitude of reduction in hookworm prevalence as the individual water and sanitation interventions. One possible explanation is that combined interventions might have lower user adherence as they require more complex behavior change [44]. However, adherence indicators were similar between individual and combined intervention arms in our study [26]. It is also possible that the primary barrier of sanitation (reducing spread of ova into water sources) and the secondary barrier of water treatment (reducing ingestion of ova/larvae) were operating on the same waterborne transmission pathway and there was thus no benefit from combining them. Nonetheless, combined WSH was the only intervention that achieved a small (albeit borderline non-significant) reduction in *A. lumbricoides*.

### Effect of nutrition

Lipid-based nutrient supplements, breastfeeding and complementary feeding promotion did not reduce STH prevalence/intensity alone or in combination with WSH, even when the analysis was restricted to index children directly receiving nutritional improvements. There was a borderline increase in *A. lumbricoides* prevalence in the nutrition arm. A recent study found increased hookworm in school children receiving micronutrient-fortified rice, raising concerns about fortification in settings with high (>15%) baseline prevalence [45]. Other studies found STH reductions from nutritional supplements [19]. WASH Benefits showed improved child growth and micronutrient status, and reduced anemia in the arms containing a nutrition component (nutrition, N+WSH), indicating that the intervention effectively improved nutritional status [22,46]. One reason for the lack of STH reduction despite improved nutrition could be the dual direction of possible biological associations between nutrition and STH infection. Breastfeeding and improved nutrition could decrease infection risk by improving immune response and repairing cells damaged by infection; conversely, it could increase risk by making nutrients available to helminths [19,20]. Chronic heavy STH infections can lead to malnutrition and growth faltering [40]. The hookworm reductions in the water and WSH arms were not reflected by improved growth in these arms. However, children in the WSH arm had a borderline reduction in the prevalence of anemia and iron deficiency as indicated by low ferritin [46], which would be consistent with the reduction in hookworm prevalence in this arm. Nutritional outcomes are likely interrelated with myriad causal effects and the impact of STH on growth should be further assessed.

### Findings in the context of MDA

This trial provides a novel investigation of water, sanitation, hygiene and nutrition interventions in a population with ongoing MDA. Our findings can inform policy dialogue about whether STH control policies, which currently emphasize MDA programs, could be strengthened by complementing these with water, sanitation and hygiene interventions [47]. Since 2008, the Bangladesh Ministry of Health has implemented biannual school-based deworming while pre-school-aged children are dewormed through the Expanded Program on Immunization [23]. The majority of infections we detected in enrolled children were low-intensity, suggesting that the deworming program successfully reduced the prevalence of heavy infections that drive the morbidity burden [1]. However, 43% of children in the control arm were infected with STH (mostly *A. lumbricoides*) despite several years of deworming, demonstrating ongoing transmission and suggesting that MDA alone is unlikely to break STH transmission in this setting.

Against this backdrop, we found a 30% relative reduction in hookworm prevalence from water treatment and combined WSH interventions, as well as a borderline reduction of similar size from sanitation improvements. While the reductions were comparable in magnitude to the reductions in child diarrhea and protozoan infections achieved by WASH Benefits [22,48] and other water treatment and hygiene trials in low-income countries [49,50], they are small compared to the typical cure rates from deworming [2,24], suggesting that water, sanitation and hygiene interventions alone would not sufficiently reduce STH morbidity in similar settings. However, while re-infection rates following deworming can be as high as 94% within 12 months of drug administration [4], the effects we report were observed 2.5 years after intervention initiation, suggesting sustained reductions in environmental transmission in a population receiving biannual MDA.

It is also possible that the effect of water, sanitation and hygiene on STH depends on background transmission intensity. In our study, WSH interventions had more pronounced effect on hookworm, which was relatively rare (9% control prevalence), than on *A. lumbricoides*, which was more common (37% control prevalence). This is consistent with a school-based trial in Kenya that found reduction in *A. lumbricoides* (9-14% prevalence in the study population) but not the more prevalent hookworm (28-29% prevalence) from a combined water, sanitation and hygiene intervention [16]. These findings suggest that in settings where deworming has been successfully implemented to reduce infection intensity and morbidity, water, sanitation and hygiene interventions can complement MDA programs in striving toward elimination by interrupting environmental transmission.

### Limitations

We measured STH infection using Kato-Katz, which has poor sensitivity when infection intensity is low. A systematic review demonstrated a sensitivity of 55% for *A. lumbricoides*, 53% for hookworm, 80% for *T. trichiura* for double-slide Kato-Katz for low-intensity infections [51]. As 95% of infections in our study were low-intensity, this could yield substantial false negatives in our outcome measurements. Recently developed sensitive nucleic acid-based diagnostics can detect infections that are missed by Kato-Katz [52,53]. We preserved an additional stool aliquot for validation analysis by quantitative polymerase chain reaction (qPCR). Preliminary analyses in a validation study using a subset of our specimens suggest that double-slide Kato-Katz had low to moderate sensitivity for all three STH while it had moderate specificity for *A. lumbricoides* and high specificity for *T. trichiura* and hookworm (Benjamin-Chung et al. 2018, *in prep*). Assuming non-differential misclassification by arm, imperfect sensitivity and specificity would bias our estimated intervention effects toward the null (Text S5). If the interventions reduced infection intensity, imperfect sensitivity could also lead to differential misclassification by arm, where a larger proportion of cases in the intervention arms would go undetected by Kato-Katz and intervention effects would therefore be biased away from the null.

Also, WASH Benefits was designed around its primary outcomes (length-for-age Z-score and diarrhea) so there was only sufficient statistical power to detect relatively large effects on hookworm and *T. trichiura* given their low prevalence. Post-hoc calculations suggested an MDE of 19% relative reduction for *A. lumbricoides*, 41% for hookworm, and 52% for *T. trichiura*. Future studies in low-prevalence settings should enroll sample sizes large enough to detect small effects and use sensitive diagnostics.

We conducted multiple comparisons, increasing the risk of chance findings; the *T. trichiura* reduction in the sanitation but not WSH and N+WSH arms could indicate random error. However, most observed reductions followed consistent patterns that are unlikely to be explained by chance. Hookworm prevalence and intensity showed internally consistent reductions of similar size in all arms with a water or sanitation component, while the only intervention that achieved a borderline reduction in *A. lumbricoides* was combined WSH - the most biologically plausible intervention to reduce environmental transmission.

Another limitation is that we assessed STH outcomes after 2.5 of intervention, which is a relatively short period of time to assess impact on *A. lumbricoides* given its long survival in soil [13]. This timeframe risks underestimating the long-term population benefit of reducing environmental soil contamination through improved sanitation. Longer-term follow-up of this population might provide a more accurate assessment of the long-term contribution of improved sanitation towards *A. lumbricoides* elimination.

Environmental conditions such as temperature, humidity and soil type affect the fate and transport of STH ova [34,40] and intervention effects are therefore likely to be setting-dependent. We controlled for month in our analysis to adjust for seasonality. Also, our geographically pair-matched randomization synchronized the timing of outcome measurement between arms, eliminating confounding from season as well as unmeasured tempo-spatial factors. However, our findings may not be generalizable to other settings with different climatic and geological conditions, or different levels of fecal contamination in the ambient environment. Similarly, our findings are relevant to other populations with MDA programs and relatively low intensity of STH infection. Future studies should investigate the effect of water, sanitation and hygiene improvements on STH infection and how these can augment MDA programs in high-intensity infection settings.

### Conclusions

In a setting with ongoing MDA and low-intensity infections, we found modest but sustained reductions in hookworm prevalence and intensity from water treatment, sanitation and combined WSH interventions. There was no STH reduction from handwashing and nutrition improvements. Intervention effects were more pronounced on hookworm than on *A. lumbricoides* and *T. trichiura;* this could be because of the short survival of hookworm in soil, precluding persistent environmental reservoirs of ova from pre-intervention contamination. Our findings highlight drinking water as an overlooked transmission route for hookworm and suggest that water treatment and sanitation interventions can augment MDA programs in striving towards breaking transmission.

## Acknowledgements

We greatly appreciate the families who participated in the study and the dedication of the icddr,b staff who delivered the interventions and collected the data and specimens. We thank Md. Alimojjaman for overseeing field activities and specimen collection, Shimul Das for overseeing specimen receipt, Md. Rabiul Karim, Shamsun Nahar Jothi, Habibur Rahman and A.S.M. Homaun Kabir Chowdhury for completing the Kato-Katz analyses, and Farida Nazib and Md. Abdullah Siddique at the icddr,b Parasitology Laboratory for providing guidance on microscopy.

## Supplemental Information

Text S1. Study protocol

Text S2. CONSORT checklist

Text S3. Intervention promotion and user adherence

Text S4. Quality assurance for Kato-Katz

Text S5. Bias in effect estimates with imperfect sensitivity and specificity under non-differential classification

Table S1: Enrolment characteristics of individuals with missing vs. observed outcomes

Table S2: Infection prevalence, all interventions vs. control

Table S3: Infection prevalence, combined vs. individual WSH interventions

Table S4: Infection prevalence, combined N+WSH vs. WSH and nutrition interventions

Table S5: Moderate/heavy infection prevalence, all interventions vs. control

Table S6: Moderate/heavy infection prevalence, combined vs. individual WSH interventions

Table S7: Moderate/heavy infection prevalence, combined N+WSH vs. WSH and nutrition interventions

Table S8: Fecal egg count reduction, all interventions vs. control

Table S9: Fecal egg count reduction, combined vs. individual WSH interventions

Table S10: Fecal egg count reduction, combined N+WSH vs. WSH and nutrition interventions

