## Supplemental Text for "Effects of water, sanitation, handwashing and nutritional interventions on soil-transmitted helminth infections in young children: a cluster-randomized controlled trial in rural Bangladesh"

**Supplemental Information**

Text S1. Study protocol
Text S2. CONSORT checklist
Text S3. Intervention promotion and user adherence
Text S4. Quality assurance for Kato-Katz
Text S5. Bias in effect estimates with imperfect sensitivity and specificity under non-differential classification

PROTOCOL
Biomedical Non-Exempt
Berkeley

Protocol # 2011-09-3652
Date Printed: 07/31/2018

PROTOCOL
Biomedical
Non-Exempt
Berkeley

Protocol # 2011-09-3652
Date Printed: 07/31/2018

---

**Protocol Title:** Measuring the benefits of sanitation, water quality, handwashing and nutrition interventions for improving health and development in rural Bangladesh

**Protocol Status:** APPROVED

**Date Submitted:** 10/26/2017

**Approval Period:** 10/30/2017-11/03/2018

**Important Note:** This Print View may not reflect all comments and contingencies for approval. Please check the comments section of the online protocol.
Questions that appear to not have been answered may not have been required for this submission. Please see the system application for more details.

#### * * * Amendment Application * * *

##### Amendment Application

1. **Summarize the amendment (or proposed changes) you wish to make to your study.**

We will change the Administrative Contact listed on the protocol from Melanie Gendell to Lucas Carlton.

2. **Explain the reason(s) for the proposed amendment(s).**

Melanie Gendell, the Program and Administrative Manager of the Colford-Hubbard Research Group, retired this year. We would like to add our new Program and Administrative Manager, Lucas Carlton, to this protocol as the Administrative Contact at UC Berkeley.

3. **Indicate how the change(s) impact the level of risk to subjects:**

| | |
|---|---|
| | Increase |
| Y | No Change |
| | Decrease |

4. **Describe any effects the change(s) will have regarding risk(s) to the subjects:**

n/a

5. **Will this amendment require the re-consent of any currently enrolled subjects?** N

**If YES, please explain.**

6. **Is this modification consistent with the scope of research activities as described in the proposal(s) for the grant(s) funding the research? (Check N/A if you have no external funding)** Y

---

| | |
|---|---|
| **Protocol Title:** | Measuring the benefits of sanitation, water quality, handwashing and nutrition interventions for improving health and development in rural Bangladesh |
| **Protocol Status:** | APPROVED |
| **Date Submitted:** | 10/26/2017 |
| **Approval Period:** | **10/30/2017-11/03/2018** |
| **Important Note:** | **This Print View may not reflect all comments and contingencies for approval. Please check the comments section of the online protocol. Questions that appear to not have been answered may not have been required for this submission. Please see the system application for more details.** |

**7. If this is an amendment or renewal application including changes to previously-approved consent/permission/assent documents, please include a version of the document(s) with tracked changes in the Attachments section under 'Other' in order to facilitate review. The list of sections that have been changed or modified will appear below:
Proceed to the appropriate section(s) and make your changes.**

---

---

**Protocol Title:** Measuring the benefits of sanitation, water quality, handwashing and nutrition interventions for improving health and development in rural Bangladesh

**Protocol Status:** APPROVED

**Date Submitted:** 10/26/2017

**Approval Period:** 10/30/2017-11/03/2018

**Important Note:** This Print View may not reflect all comments and contingencies for approval. Please check the comments section of the online protocol.
Questions that appear to not have been answered may not have been required for this submission. Please see the system application for more details.

#### * * * Personnel Information * * *

**Enter all UC Berkeley study personnel (if not previously entered) and relevant training information. Please read Personnel Titles and Responsibilities: Roles in eProtocol before completing this section.**

**Note: The Principal Investigator or Faculty Sponsor, Co-Principal Investigator, Student or Postdoctoral Investigator, Administrative Contact, and Other Contact can EDIT and SUBMIT. Other Personnel can only VIEW the protocol.**

##### Principal Investigator or Faculty Sponsor

**Name of Principal Investigator**
Jack COLFORD

**Degree (e.g., MS/PhD)**

**Title**
Professor

**Email**


**Phone**
+1 510 642-9370

**Fax**
+1 510 666-2551

**Department Name**
Pub Hlth-Epidemiology

**Mailing Address**
94720-7358

UCB status (select all that apply):

| | | | | | | | | | |
|---|---|---|---|---|---|---|---|---|---|
| X | Faculty | | Postdoc | | Grad | | Undergrad | | Other |

Faculty (with some exceptions), staff, and students engaged in human subjects research must complete either the biomedical or social-behavioral human research course through the online Collaborative Institutional Training Initiative (CITI), depending upon which is most germane to the research. ALL PIs on an NIH award are required to complete either CITI or NIH Training. See Training and Education for more information.

If applicable, please insert date (mm/dd/yy) of completion in appropriate box(es) below:

| CITI | NIH | Other Training (title & date completed) |
|---|---|---|

##### Administrative Contact

**Name of Administrative Contact**
Jade D. Benjamin-Chung

**Degree**
MPH

**Title**

**Email**

**Phone**

**Fax**

---

**Protocol Title:** Measuring the benefits of sanitation, water quality, handwashing and nutrition interventions for improving health and development in rural Bangladesh

**Protocol Status:** APPROVED

**Date Submitted:** 10/26/2017

**Approval Period:** 10/30/2017-11/03/2018

**Important Note:** This Print View may not reflect all comments and contingencies for approval. Please check the comments section of the online protocol. Questions that appear to not have been answered may not have been required for this submission. Please see the system application for more details.

| Department Name | Mailing Address |
|---|---|
| Pub Hlth-Epidemiology | |

UCB status (select all that apply):

| | | | | | | | | | |
|---|---|---|---|---|---|---|---|---|---|
| | Faculty | | Postdoc | X | Grad | | Undergrad | | Other |

#### Other Contact

| Name of Other Contact | Degree | Title |
|---|---|---|
| Lucas R CARLTON | | SRA III |
| **Email** | **Phone** | **Fax** |
| | +1 510 643-0238 | |
| **Department Name** | **Mailing Address** | |
| Pub Hlth-Epidemiology | 94720-7360 | |

UCB status (select all that apply):

| | | | | | | | | | | |
|---|---|---|---|---|---|---|---|---|---|---|
| | Faculty | | Postdoc | | Grad | | Undergrad | X | Other | Staff |

##### Other Personnel

| Name | Degree | Title | Department Name |
|---|---|---|---|
| Benjamin F. Arnold | | Epidemiologist | Pub Hlth-Epidemiology |
| Alan HUBBARD | | Associate Professor | Pub Hlth-Biostatistics |
| Lia C. FERNALD | | Associate Professor | Pub Hlth-Nutrition |
| Audrie Lin | PhD | Specialist | Pub Hlth-Epidemiology |
| Kara Nelson | | Professor | Civ Engr/CEE Environmental |
| Ayse Ercumen | PhD | Specialist | Pub Hlth-Epidemiology |

---

**Protocol Title:** Measuring the benefits of sanitation, water quality, handwashing and nutrition interventions for improving health and development in rural Bangladesh

**Protocol Status:** APPROVED

**Date Submitted:** 10/26/2017

**Approval Period:** **10/30/2017-11/03/2018**

**Important Note:** **This Print View may not reflect all comments and contingencies for approval. Please check the comments section of the online protocol.
Questions that appear to not have been answered may not have been required for this submission. Please see the system application for more details.**

---

---

| | |
|---|---|
| **Protocol Title:** | Measuring the benefits of sanitation, water quality, handwashing and nutrition interventions for improving health and development in rural Bangladesh |
| **Protocol Status:** | APPROVED |
| **Date Submitted:** | 10/26/2017 |
| **Approval Period:** | **10/30/2017-11/03/2018** |
| **Important Note:** | **This Print View may not reflect all comments and contingencies for approval. Please check the comments section of the online protocol. Questions that appear to not have been answered may not have been required for this submission. Please see the system application for more details.** |

#### * * * Vulnerable Subject Checklist * * *

##### Vulnerable Subject Checklist

| Yes | No | |
|---|---|---|
| Y | | Children/Minors |
| | N | Prisoners |
| Y | | Pregnant Women |
| | N | Fetuses |
| Y | | Neonates |
| Y | | Educationally Disadvantaged |
| Y | | Economically Disadvantaged |
| | N | Cognitively Impaired |
| | N | Other (i.e., any vulnerable subject population(s) not specified above) |

---

---

**Protocol Title:** Measuring the benefits of sanitation, water quality, handwashing and nutrition interventions for improving health and development in rural Bangladesh

**Protocol Status:** APPROVED

**Date Submitted:** 10/26/2017

**Approval Period:** 10/30/2017-11/03/2018

**Important Note:** This Print View may not reflect all comments and contingencies for approval. Please check the comments section of the online protocol.
Questions that appear to not have been answered may not have been required for this submission. Please see the system application for more details.

#### * * * Study Sites * * *

##### Study Sites

Select all study sites where data collection via subject interaction will take place:

International

X International Site(s) (specify country, region, and township or village)

Bangladesh

Local

X UC Berkeley

X UC Davis

UC Irvine

UC Los Angeles

UC Merced

UC Riverside

UC San Diego

UC San Francisco

UC Santa Barbara

UC Santa Cruz

Lawrence Berkeley National Laboratory

Alameda Unified School District (specify schools below)

Berkeley Unified School District (specify schools below)

Oakland Unified School District (specify schools below)

Other (Specify other Study Sites)

---

---

| | |
|---|---|
| **Protocol Title:** | Measuring the benefits of sanitation, water quality, handwashing and nutrition interventions for improving health and development in rural Bangladesh |
| **Protocol Status:** | APPROVED |
| **Date Submitted:** | 10/26/2017 |
| **Approval Period:** | **10/30/2017-11/03/2018** |
| **Important Note:** | **This Print View may not reflect all comments and contingencies for approval. Please check the comments section of the online protocol. Questions that appear to not have been answered may not have been required for this submission. Please see the system application for more details.** |

#### * * * General Checklist * * *

##### General Checklist

| Yes | No | |
|---|---|---|
| | N | Is the research receiving any federal funding (e.g., NIH, NSF, DOD, etc.)? |
| | N | Is another campus relying on UC Berkeley for IRB review by means of the UC System Memorandum of Understanding (MOU)? |
| Y | | Is another institution relying on UC Berkeley for IRB review by means of an Inter-institutional IRB Authorization Agreement? |
| | N | Will subjects be paid for participation? |
| | N | Is this protocol administratively supported by Campus Shared Services Team 9? |
| | N | Does this research fall under FDA regulations? |
| Y | | Any use of human blood, body fluids, tissues, or cells (including cell lines)* by drawing samples, accepting samples already drawn, receiving samples from any source, or in any other way? |
| | | If yes, Lab Location: ICDDRB Bangladesh |
| | | And Biological Use Authorization (BUA) #(s): |
| Y | | Will biological specimens be stored for future research projects? |
| | N | Will specimens be sent out of UCB as part of a research agreement? |
| | N | Will proprietary drug or device testing be done? |
| | N | Any use of embryonic stem cells? *NOTE: If research involves embryonic stem cells, see UCB Stem Cell Policy and Committee. |
| | N | Any use of medical devices or equipment cleared/approved for marketing? |
| | N | Any use of any experimental or investigational devices or equipment (i.e., not cleared/approved for marketing?) |
| Y | | Any use of commercially available drugs, reagents, or other chemicals administered to subjects (even if drugs themselves are not being studied)? |
| | N | Any use of investigational drugs, reagents, or chemicals (i.e., not cleared/approved for marketing)? |

---

---

**Protocol Title:** Measuring the benefits of sanitation, water quality, handwashing and nutrition interventions for improving health and development in rural Bangladesh

**Protocol Status:** APPROVED

**Date Submitted:** 10/26/2017

**Approval Period:** 10/30/2017-11/03/2018

**Important Note:** This Print View may not reflect all comments and contingencies for approval. Please check the comments section of the online protocol.
Questions that appear to not have been answered may not have been required for this submission. Please see the system application for more details.

#### * * * Funding * * *

##### Funding Checklist

If the research is not funded, check the "Not Funded" box below. If the research is funded, add the funding source to the appropriate table below.

NOTE: Only the Principal Investigator (PI) of the grant or subcontract can add his or her own SPO Funding information in this section. The PI of the grant must also be listed in the Personnel Information section of the protocol in one of the following roles: Principal Investigator or Faculty Sponsor, Student or Postdoctoral Investigator, Co-Principal Investigator, Administrative Contact, or Other Contact. Training Grants can be added by anyone in one of the aforementioned roles. For step-by-step instructions, see Add SPO Funding Quick Guide

Not Funded

SPO - Funding

| SPO ID | Sponsor | Sponsor Award ID | Prime Sponsor |
|---|---|---|---|

##### SPO - Funding

| | |
|---|---|
| SPO ID | 20130310 |
| Sponsor Award ID | |
| Sponsor | Bill & Melinda Gates Foundation |
| Prime Sponsor | |
| Funding Status | Pending |
| Principal Investigator | Colford Jr, John M |
| Co-Investigator (s) | |
| Admin Unit | SPH Administrative Services |
| Project Title | Measuring the Benefits of Sanitation, Water Quality, Handwashing and Nutrition Interventions for Improving Health and Development |
| Amount | $271,336 |
| Start | 9/17/2009 |
| End | 9/30/2015 |

---

| | |
|---|---|
| **Protocol Title:** | Measuring the benefits of sanitation, water quality, handwashing and nutrition interventions for improving health and development in rural Bangladesh |
| **Protocol Status:** | APPROVED |
| **Date Submitted:** | 10/26/2017 |
| **Approval Period:** | **10/30/2017-11/03/2018** |
| **Important Note:** | **This Print View may not reflect all comments and contingencies for approval. Please check the comments section of the online protocol. Questions that appear to not have been answered may not have been required for this submission. Please see the system application for more details.** |

| | |
|---|---|
| **Subcontracts** | Yes |

| | |
|---|---|
| **SPO ID** | 028064-002 |
| **Sponsor Award ID** | OPPGD759 |
| **Sponsor** | Bill & Melinda Gates Foundation |
| **Prime Sponsor** | |
| **Funding Status** | Active |
| **Principal Investigator** | Colford Jr, John M |
| **Co-Investigator (s)** | |
| **Admin Unit** | School of Public Health |
| **Project Title** | Measuring the Benefits of Sanitation, Water Quality and Handwashing Interventions for Improving Health and Development |
| **Amount** | $28,521,066 |
| **Start** | 9/17/2009 |
| **End** | 9/30/2015 |
| **Subcontracts** | No |

| | |
|---|---|
| **SPO ID** | 20131960 |
| **Sponsor Award ID** | |
| **Sponsor** | NIH National Institutes of Health - Miscellaneous |
| **Prime Sponsor** | |
| **Funding Status** | Pending |
| **Principal Investigator** | Colford Jr, John M |
| **Co-Investigator (s)** | |
| **Admin Unit** | School of Public Health |
| **Project Title** | Effects of Sanitation of Pathogen Transmission and Child Health in Bangladesh |
| **Amount** | $2,692,294 |
| **Start** | 9/1/2013 |
| **End** | 8/31/2017 |

---

| | |
|---|---|
| **Protocol Title:** | Measuring the benefits of sanitation, water quality, handwashing and nutrition interventions for improving health and development in rural Bangladesh |
| **Protocol Status:** | APPROVED |
| **Date Submitted:** | 10/26/2017 |
| **Approval Period:** | **10/30/2017-11/03/2018** |
| **Important Note:** | **This Print View may not reflect all comments and contingencies for approval. Please check the comments section of the online protocol. Questions that appear to not have been answered may not have been required for this submission. Please see the system application for more details.** |

| | |
|---|---|
| **Subcontracts** | Yes |

| | |
|---|---|
| **SPO ID** | 20141262 |
| **Sponsor Award ID** | |
| **Sponsor** | Stanford University |
| **Prime Sponsor** | World Bank |
| **Funding Status** | Funded |
| **Principal Investigator** | Colford Jr, John M |
| **Co-Investigator (s)** | |
| **Admin Unit** | Dean's Office, Public Health |
| **Project Title** | Assessment of Exposure Pathways to Fecal Contamination, Assocation with Diarrhea and Sanitation Coverage in Bangladesh |
| **Amount** | $40,744 |
| **Start** | 6/1/2013 |
| **End** | 4/30/2014 |
| **Subcontracts** | No |

| | |
|---|---|
| **SPO ID** | 20140633 |
| **Sponsor Award ID** | |
| **Sponsor** | NIH National Institute of Child Health & Human Development |
| **Prime Sponsor** | |
| **Funding Status** | Pending |
| **Principal Investigator** | Colford Jr, John M |
| **Co-Investigator (s)** | |
| **Admin Unit** | Dean's Office, Public Health |
| **Project Title** | Spillover Effects of Water, Sanitation, and Hygiene Interventions on Child Health |
| **Amount** | $368,089 |
| **Start** | 4/1/2014 |

---

| | |
|---|---|
| **Protocol Title:** | Measuring the benefits of sanitation, water quality, handwashing and nutrition interventions for improving health and development in rural Bangladesh |
| **Protocol Status:** | APPROVED |
| **Date Submitted:** | 10/26/2017 |
| **Approval Period:** | **10/30/2017-11/03/2018** |
| **Important Note:** | **This Print View may not reflect all comments and contingencies for approval. Please check the comments section of the online protocol. Questions that appear to not have been answered may not have been required for this submission. Please see the system application for more details.** |

| | |
|---|---|
| **End** | 3/31/2016 |
| **Subcontracts** | Yes |

| | |
|---|---|
| **SPO ID** | 20142940 |
| **Sponsor Award ID** | |
| **Sponsor** | UC Davis |
| **Prime Sponsor** | Thrasher Research Fund |
| **Funding Status** | Funded |
| **Principal Investigator** | John M Colford Jr |
| **Co-Investigator (s)** | |
| **Admin Unit** | SPH Divisional Rsrch and Cntrs |
| **Project Title** | Evaluating the impact of water, sanitation, hygiene and nutrition interventions on infant micronutrient status and anemia risk in the WASH Benefits study |
| **Amount** | $37,597 |
| **Start** | 7/1/2015 |
| **End** | 3/31/2017 |
| **Subcontracts** | No |

#### Funding - Other

---

---

**Protocol Title:** Measuring the benefits of sanitation, water quality, handwashing and nutrition interventions for improving health and development in rural Bangladesh

**Protocol Status:** APPROVED

**Date Submitted:** 10/26/2017

**Approval Period:** **10/30/2017-11/03/2018**

**Important Note:** **This Print View may not reflect all comments and contingencies for approval. Please check the comments section of the online protocol.**
**Questions that appear to not have been answered may not have been required for this submission. Please see the system application for more details.**

#### * * * Expedited Paragraphs * * *

**Request for Expedited Review**

**An expedited review procedure consists of a review of research involving human subjects by the IRB Chair, or by one or more experienced reviewers designated by the Chairperson from among the members of the committees.**

**In order to be eligible for expedited review, ALL aspects of the research must include activities that (1) present no more than minimal risk to human subjects, and (2) involve only procedures included in one or more of the specific categories listed below.**

**If requesting Expedited Review, select one or more of the applicable paragraph(s) below.**
**(DO NOT select any paragraph(s) if your protocol does not qualify for expedited review. Protocols that do not qualify for expedited review will be reviewed by the full (convened) Committee.)**

1. **Clinical studies of drugs and medical devices only when conditions (a) and (b) are met.**

   **a)** Research on drugs for which an investigational new drug application (21 CFR Part 312) is not required. (Note: Research on marketed drugs that significantly increases the risks or decreases the acceptability of the risks associated with the use of the product is not eligible for expedited review.)

   **b)** Research on medical devices for which

   **i)** an investigational device exemption application (21 CFR Part 812) is not required; or
   **ii)** the medical device is cleared/approved for marketing and the medical device is being used in accordance with its cleared/approved labeling.

X 2. **Collection of blood samples by finger stick, heel stick, ear stick, or venipuncture as follows:**

   **a)** From healthy, non-pregnant adults who weigh at least 110 pounds. For these subjects, the amounts drawn may not exceed 550 ml in an 8 week period and collection may not occur more frequently than 2 times per week; or

   **b)** From other adults and children, considering the age, weight, and health of the subjects, the collection procedure, the amount of blood to be collected, and the frequency with which it will be collected. For these subjects, the amount drawn may not exceed the lesser of 50 ml or 3 ml per kg in an 8 week period and collection may not occur more frequently than 2 times per week.

X 3. **Prospective collection of biological specimen for research purposes by non-invasive means.**

**Examples:**

**a)** hair and nail clippings in a non-disfiguring manner;
**b)** deciduous teeth at time of exfoliation or if routine patient care indicates a need for extraction;
**c)** permanent teeth if routine patient care indicates a need for extraction;
**d)** excreta and external secretions (including sweat);
**e)** uncannulated saliva collected either in an unstimulated fashion or stimulated by chewing gumbase or wax or by applying a dilute citric solution to the tongue;
**f)** placenta removed at delivery;
**g)** amniotic fluid obtained at the time of rupture of the membrane prior to or during labor;
**h)** supra- and subgingival dental plaque and calculus, provided the collection procedure is not more invasive than routine prophylactic scaling of the teeth and the process is accomplished in accordance with accepted prophylactic techniques;
**i)** mucosal and skin cells collected by buccal scraping or swab, skin swab, or mouth washings;
**j)** sputum collected after saline mist nebulization.

X 4. **Collection of data through non-invasive procedures (not involving general anesthesia or sedation) routinely employed in clinical practice, excluding procedures involving x rays or microwaves. Where medical devices are employed, they must be cleared/approved for marketing. (Studies intended to evaluate the safety and effectiveness of the medical device are not generally eligible for expedited review, including studies of cleared medical devices for new indications.)**

**Examples:**

**a)** physical sensors that are applied either to the surface of the body or at a distance and do not involve input of significant amounts of energy into the subject or an invasion of the subject's privacy;
**b)** weighing or testing sensory acuity;
**c)** magnetic resonance imaging;
**d)** electrocardiography, electroencephalography, thermography, detection of naturally occurring radioactivity, electroretinography, ultrasound, diagnostic infrared imaging, doppler blood flow, and echocardiography;
**e)** moderate exercise, muscular strength testing, body composition assessment, and flexibility testing where appropriate given the age, weight, and health of the individual.

**5. Research involving materials (data, documents, records, or specimens) that have been collected or will be collected solely for non-research purposes (such as medical treatment or diagnosis). (NOTE: Some research in this paragraph may be exempt from the HHS regulations for the protection of human subjects. 45 CFR 46.101(b)(4). This listing refers only to research that is not exempt.)**

**6. Collection of data from voice, video, digital, or image recordings made for research purposes.**

**7. Research on individual or group characteristics or behavior (including, but not limited to, research on perception, cognition, motivation, identity, language, communication, cultural beliefs or practices, and social behavior) or research employing survey, interview, oral history, focus group, program evaluation, human factors evaluation, or quality assurance methodologies. (NOTE: Some research in this category may be exempt from the HHS regulations for the protection of human subjects. 45 CFR 46.101(b)(2) and (b)(3). This listing refers only to research that is not exempt.)**

**8. Continuing review of research previously approved by the convened IRB as follows:**

**a)** Where (i) the research is permanently closed to the enrollment of new subjects; (ii) all subjects have completed all research-related interventions; and (iii) the research remains active only for long-term follow-up of subjects; or

**b)** Where no subjects have been enrolled and no additional risks have been identified; or

**c)** Where the remaining research activities are limited to data analysis.

**9. Continuing review of research, not conducted under an investigational new drug application or investigational device exemption where categories two (2) through eight (8) do not apply but the IRB has determined and documented at a convened meeting that the research involves no greater than minimal risk and no additional risks have been identified.**

---

---

**Protocol Title:** Measuring the benefits of sanitation, water quality, handwashing and nutrition interventions for improving health and development in rural Bangladesh

**Protocol Status:** APPROVED

**Date Submitted:** 10/26/2017

**Approval Period:** 10/30/2017-11/03/2018

**Important Note:** **This Print View may not reflect all comments and contingencies for approval. Please check the comments section of the online protocol. Questions that appear to not have been answered may not have been required for this submission. Please see the system application for more details.**

#### * * * Purpose, Background, Collaborative Research * * *

Old CPHS # (for Protocols approved before eProtocol)

**Study Title**

Measuring the benefits of sanitation, water quality, handwashing and nutrition interventions for improving health and development in rural Bangladesh

**Complete each section. When a question is not applicable, enter "N/A". Do not leave any sections blank.**

##### 1. Purpose

**Provide a brief explanation of the proposed research, including specific study hypothesis, objectives, and rationale.**

The goal of the WASH Benefits study is to generate rigorous evidence about the impacts of sanitation, water quality, and handwashing (WASH) and nutrition interventions on child health and development in the first years of life. The primary hypotheses of the study are:
H1: Water, sanitation, handwashing, nutrition and their combination improve child health and development.
H2: When delivered in combination, water, sanitation and handwashing interventions reduce child diarrhea more than when delivered individually.
H3: Combined Nutrient supplementation and WASH interventions improve child growth and development more than nutrient supplementation alone.

The study objectives are to:

1) Rigorously measure health benefits arising from low-cost WASH approaches including local promoters and subsidies for simple technologies (e.g. latrine improvements or potties for children, chlorine dispensers, and handwashing stations) and evaluate the degree to which, in resource-constrained settings, there is added health benefit to delivering multiple interventions concurrently (sanitation services, drinking water, and handwashing promotion)

Most of the burden of diarrheal disease is thought to be preventable with improvements in sanitation, water quality, and hygiene. However, in rural areas of low-income countries it is often prohibitively expensive to provide residents with networked sanitation and water treatment that provide microbiologically and chemically safe water and consistently separate feces from the environment. We have almost no evidence that allows direct comparison of the health benefits or cost-effectiveness of improvements in sanitation, water quality, and hygiene, nor on how the benefits of these interventions aggregate when provided in combination. Such evidence is critical for guiding the allocation of public and donor funds to achieve the maximum health impact given limited resources. In the absence of credible evidence, little change from the status quo can be expected, even if the impact of current practices is unknown. A rigorous evaluation of these interventions that documents changes in outcomes associated with long-run economic success, such as early childhood growth, could have at least two important influences on policy. First, such evidence could help to maximize the value of existing resources by shifting expenditures to the most cost-

---

**Protocol Title:** Measuring the benefits of sanitation, water quality, handwashing and nutrition interventions for improving health and development in rural Bangladesh

**Protocol Status:** APPROVED

**Date Submitted:** 10/26/2017

**Approval Period:** 10/30/2017-11/03/2018

**Important Note:** **This Print View may not reflect all comments and contingencies for approval. Please check the comments section of the online protocol. Questions that appear to not have been answered may not have been required for this submission. Please see the system application for more details.**

effective interventions. In addition, such evidence could help generate more resources for these sectors by resolving uncertainty regarding the efficacy of water, sanitation, and hygiene interventions and identifying simple technologies and approaches to behavior change that cost-effectively improve health and could be replicated at scale.
2) Measure the impact of lipid-based nutrient supplementation (LNS) alone and in combination with sanitation, water and hygiene interventions on child growth and development.
For children whose food intake is insufficient, LNS helps reduce gross energy shortfalls and provide essential micronutrients. The energy and micronutrients that LNS provides are likely to improve nutritional indicators, length-for-age and child development, particularly among children that are at highest risk for severe stunting. It is possible that improved nutrition alone can reduce the negative effects of infection on growth and development due to the improved ability of better-nourished children to fight off enteric infections and exhibit catch-up growth during the convalescent period.
A combined LNS+WASH intervention could have greater impacts on growth and development than LNS alone. The reasons for this are two-fold. First, the likely reciprocal relationship between enteric infection and malnutrition in young children suggests that the provision of joint interventions that interrupt both components of the “vicious cycle” may have effects that exceed interventions that interrupt just one component. Indeed, if there is a reciprocal relationship between enteric infection and malnutrition, the feedback loop toward decline could, in principal, be reversed and leveraged to enhance growth and development. There is a second plausible scenario that could result in larger combined effects: if the interventions are deployed together, the available energy for growth and development could be enhanced by improved utilization of the additional nutrition provided by LNS that would have not been available to the child without complementary infection control. Such improved utilization could be achieved by reductions in acute diarrheal disease and the chronic symptoms that characterize environmental enteropathy.
3) Measure the impact of nutritional supplements and household environmental interventions on environmental enteropathy biomarkers, and more clearly elucidate this potential pathway between environmental interventions and child growth and development.
If improvements in sanitation, water quality, and hand hygiene could reduce the severity of intestinal malabsorption from environmental enteropathy (EE) either by preventing its acquisition or by reversing the pathology, this would represent an important contribution to global public health. EE is an inflammatory disorder of the small intestine that results in reduced nutrient absorption and increased gut permeability (and thus increased immune system stimulation). The scientific literature to date suggests that EE is most likely caused by poor sanitation, water quality and hygiene in low income countries, but there are no studies that demonstrate a specific association between EE and environmental conditions separated from other exposures in a low income environment. The WASH Benefits study is uniquely positioned to gather randomized evidence about the impact of household environmental interventions on EE in young children. Furthermore, the study will also be positioned to gather rigorous evidence about the independent and combined impact of supplemental nutrition and WASH interventions on EE biomarkers.
4) Measure the impact of WASH and nutrition interventions on intestinal parasitic infection prevalence and intensity.
Observational studies suggest that environmental interventions can reduce parasite infection if sanitation and hygiene conditions improve in a large share of the population. Pre-school aged children spend much of their time in the immediate home environment. It is with this in mind that the WASH Benefits study focuses on household environmental interventions. There is a large body of evidence that documents household level clustering and within-household transmission of Cryptosporidium, Giardia, and E. histolytica, as well as soil-transmitted helminthes (Ascaris, Trichuris, and hookworm). The observed patterns suggest that intestinal parasites may be a useful marker of enteric pathogen transmission more broadly. The simultaneous measurement of parasitic infections, caregiver-reported diarrhea, and

environmental enteropathy biomarkers will allow us to explore this secondary hypothesis.

5) Measure the impact of interactions between the water, sanitation, hygiene and nutritional interventions and the mother and child's intestinal microbiome, immune function, internal biochemical environment and genetic disposition.

We expect that the interventions will have different effects in different children and that some of these differences will be mediated by differences in the mother and child's intestinal microbiome, immune function, internal biochemical environment and genetic disposition. We will collect maternal and child samples to permit future assessment of mother and child's intestinal microbiome, immune function, internal biochemical environment and genetic disposition, and so explore the importance of interactions between these characteristics and study outcomes. Measurements of immune function, internal biochemical environment including nutrition and genetic characteristics are especially dynamic areas of scientific research where important new insights are emerging each month. The capacity to explore these factors will provide the opportunity to understand the mechanism and generalizability of any impacts on child health and development that are observed within the study.
For example a child who acquires a specific population of microbiological organisms from her mother's intestinal microbiome may be more resistant to infection with intestinal pathogens and so less likely to benefit from sanitation interventions. As a second example, the haplotype HLA AW-31 is strongly associated with the development of tropical sprue, which is a disease of the small intestine that affects intestinal absorption (Menendez-Corrada R, Nettleship E, Santiago-Delfin E. 1986. HLA and Tropical Sprue. Lancet 328:1183-1185.). We would expect that interventions to impact this small intestinal pathology would be different in the presence of persons who genetically have HLA AW-31 compared to those who do not. This example illustrates the potential importance of genotype in the development of intestinal pathology, which could consequently modify the efficacy of the interventions. By exploring genetic characteristics, such as HLA AW-31 and those that will be described in the coming 20 years, we set ourselves up to optimally understand the mechanism of interaction between genetic characteristics and intervention efficacy.

As part of the assessment of lead, a particularly potent neurotoxin that can affect child intellectual development, we plan to leverage the study to assess the prevalence of lead exposure, identify the environmental source and pathway, and strive to understand the primary incentives that encourage environmental contamination with lead.

#### 2. Background

**Give relevant background (e.g., summarize previous/current related studies) on condition, procedure, product, etc. under investigation, including citations if applicable (attach bibliography in Attachments section).**

An estimated 2.2 million children under the age of 5 years die from diarrheal disease each year (WHO 2008). Children who survive multiple episodes of diarrhea and enteric infections commonly develop environmental enteropathy, an inflammatory disorder of the intestines that compromises nutrient absorption (Haghighi 1997). Repeated episodes of diarrhea and chronic environmental enteropathy in early childhood reduce growth and cognitive function, and impair school performance (Alderman 2006,

Checkley 2008, Lorntz 2006, Niehaus 2002, Petri 2008). This in turn can reduce income later in life (Boissiere 1985). Thus, repeated episodes of childhood diarrhea and enteric infection may exact a long-run toll, perpetuating a cycle of poverty and ill health.

Water, Sanitation, and Handwashing Interventions
Most of the burden of diarrheal disease is thought to be preventable with improvements in sanitation, water quality, and hygiene (Ezzati 2003). However, in rural areas of low-income countries it is often prohibitively expensive to provide residents with networked sanitation and water treatment that provide microbiologically and chemically safe water and consistently separate feces from the environment.

This has led to a movement towards alternative non-networked solutions including improved sanitation efforts, efforts to increase water treatment by households, and programs to increase handwashing with soap. Observational studies suggest that reducing open defecation is potentially important in reducing the transmission of both diarrhea and trachoma (Esrey 1996, Esrey 1991). There is evidence that water treatment, in particular with dilute chlorine solution, can reduce self-reported diarrhea (Arnold 2007, Clasen 2007). There is also strong evidence that handwashing with soap can dramatically reduce self-reported diarrhea as well as other diseases (Ejemot 2008, Rabie 2006). Some researchers however have called for evidence on more objective measures such as physical growth and cognition, rather than reports by family members (Schmidt 2009). We have almost no evidence that allows direct comparison of the health benefits or cost-effectiveness of improvements in sanitation, water quality, and hygiene, nor on how the benefits of these interventions aggregate when provided in combination.

Improved Sanitation
Improved sanitation effectively separates human excreta from human contact and the environment. The most common sanitation technologies in developing countries, particularly in rural areas, are various forms of private latrines. Observational studies suggest that households that receive improved sanitation experience 24% less diarrhea than households without sanitary facilities (Daniels 1990, Barreto 2007). However, there has never been a randomized controlled trial (RCT) to confirm these observational findings.

Full subsidies for appropriately designed latrines have resulted in high levels of coverage and regular use. For example, in the Gambia improved pit latrines were provided free of charge to 666 households in 32 villages. After 25 to 47 months each household was revisited; 77% of the provided latrines were still in use and 97% of latrines owners said they would make a new latrine when their current one was full (Simms 2005). In an evaluation of one of the Carter Center's subsidized latrine provision programs that included a random sample of 200 households across 50 villages in Niger, 86% of latrines were in regular use and 70% were clean after one year during unannounced visits (Diallo 2007). In southern India, a recent observational study found that despite high levels of latrine coverage (57%) following a community mobilization campaign, 40% of households with toilets continued to defecate in the open and there was no improvement in child diarrhea or growth – suggesting that health impacts are not guaranteed by high coverage and are likely context dependent (Arnold 2010).

The vast majority of gastrointestinal illness is caused by fecal-oral pathogen transmission through complex, environmentally mediated pathways including drinking water, ambient waters, hands, food, soil and vectors. Sanitation coverage is a primary barrier measure that aims to prevent fecal contamination from entering the environment and could plausibly break all of these pathways. However, it is uncertain whether increasing sanitation coverage effectively reduces environmental contamination and there are

---

**Protocol Title:** Measuring the benefits of sanitation, water quality, handwashing and nutrition interventions for improving health and development in rural Bangladesh

**Protocol Status:** APPROVED

**Date Submitted:** 10/26/2017

**Approval Period:** 10/30/2017-11/03/2018

**Important Note:** **This Print View may not reflect all comments and contingencies for approval. Please check the comments section of the online protocol. Questions that appear to not have been answered may not have been required for this submission. Please see the system application for more details.**

conflicting findings on the impact of sanitation improvements on child health.

Handwashing Promotion
There is significant evidence from randomized controlled trials that households receiving intense encouragement to regularly wash their hands with soap have less self-reported diarrhea and respiratory disease than households who continue their normal hand hygiene practices (Ejemot 2008, Rabie 2006). However, the existing evidence comes from trials which used very intensive interventions to encourage handwashing interventions that would not be practical to implement at scale. A key issue is the difficulty in knowing how to encourage greater take up of handwashing at reasonable cost.

One key barrier to handwashing is the difficulty and high use of water required in filling a basin, washing hands in the basin, and emptying the basin. In an observational evaluation in rural Bangladesh, households that had soap or water at their most convenient place to wash hands were twice as likely to wash their hands with soap after fecal contact than households that lacked these essential supplies (Luby 2009) [29]. Of course causality is not clear in the absence of a randomized controlled trial. A variety of simple low-cost handwashing stations have been developed which provide a place to wash hands and a source of flowing water. These can be as simple as plastic containers which one can tilt to create a stream of water by pulling rope with one's feet, or plastic containers plugged with sticks, which release a trickle of water when the stick is removed. A well-placed handwashing station provides a visual cue to spur handwashing and greatly increases the convenience of handwashing. There is considerable evidence across the social sciences that convenience is a key factor in promoting behavior change. In fact, one important lesson from the literature on behavioral change is that making something easy can be more effective at inducing change than education or promotional messaging (Kaplan 1986, Sallis 2008, Kremer 2009). Additionally, if they are placed outside of the latrines they are at least partially in public view, and there is evidence at least in a western context that people are much more likely to wash hands after using a toilet if they believe they may be observed (Pederson 1986, Ram et al 2010).

Water Treatment
There is little evidence that providing water supplies that meet the engineering definition of "improved" lead to health or social benefits for the population. In contrast, evidence from settings where diarrhea is a leading cause of death shows that improving the microbiological quality of drinking water markedly reduces reported diarrhea (Clasen 2006, Fewtrell 2005, Arnold 2007). Randomized controlled trials conducted using various household-based point-of-use water treatment technologies have demonstrated that households that consume regularly treated water report substantially less diarrhea than households using untreated water. However, self-reported diarrhea, which is the outcome measure for the majority of these studies, may be subject to measurement error potentially correlated with treatment. For this reason, long-term objective outcomes such as those planned for the proposed study (i.e. anthropometric measurements, cognition, and environmental enteropathy) may prove more convincing from a policy perspective and may be more likely to motivate action from policy makers (WHO & UNICEF Progress on drinking water & sanitation 2008).

Chlorination is the household water treatment solution of choice in many contexts given its safety and cost-effectiveness. It has been used in piped water systems around the world for almost a century. In many places where such infrastructure is absent or imperfectly maintained, dilute chlorine solution is marketed as a consumer good used in the home. Although chlorinating household drinking water reduces reported diarrhea by 20-40%, take-up has been low under the current social marketing model. Members of the

---

**Protocol Title:** Measuring the benefits of sanitation, water quality, handwashing and nutrition interventions for improving health and development in rural Bangladesh

**Protocol Status:** APPROVED

**Date Submitted:** 10/26/2017

**Approval Period:** **10/30/2017-11/03/2018**

**Important Note:** **This Print View may not reflect all comments and contingencies for approval. Please check the comments section of the online protocol. Questions that appear to not have been answered may not have been required for this submission. Please see the system application for more details.**

study team for this proposal have developed and piloted a chlorine dispenser in rural Kenya, which is a simple device that is installed at communal water sources to enable water treatment at the point of collection. When the dispenser is provided along with a local promoter to encourage its use, we find that take-up is on the order of 60-70% in communities with access to a dispenser, as compared to 5-10% in communities with access only to the traditional model of chlorine distribution. Moreover, in contrast to other strategies that we tested, chlorine use appears to be stable or even rising over time, likely because the dispenser technology makes water treatment cheap and easy for users, and harnesses positive peer effects: the public nature of the dispenser allows community members to implicitly and explicitly remind each other to treat their water.
In Bangladesh the study will use Aquatabs for chlorination. Aquatabs are effervescent water purification tablets that utilize sodium dichloroisocyanurate (NaDCC) as the chlorine donor. NaDCC was judged to be a safe and appropriate treatment for water by the World Health Organization (Clasen and Edmondson, 2006). In a previous field study in Geneva slum in Dhaka, use of Aquatabs consistently yielded an appropriate chlorine dose and dramatically improved drinking water quality (Clasen 2007). In pilot testing in rural Bangladesh Aquatabs provided with a safe water storage container were acceptable to the community with 78% of households having chlorine residual in store drinking water on unannounced follow-up visits.

Pilot Work (2010-12-2601)
The WASH Benefits pilot work has allowed the study teams to refine the sanitation, handwashing and water quality interventions and identify hardware and behavior change packages that result in high levels of uptake.

Based on the results of the pilot work, the following water, sanitation, and hygiene interventions will be implemented:

Intervention class
--------------------------------------------------------------
Sanitation:  Sanitation promotion, child potties, sani-scoop hoes to remove feces from household environments, latrine upgrades to a dual pit latrine

Handwashing:  Promotion of handwashing with soap or waterless hand sanitizers at critical times, handwashing stations, soapy water at handwashing locations

Water quality:  Chlorine tablets (Aquatabs) + safe storage vessels, water treatment promotion
--------------------------------------------------------------

Nutrition Intervention
There is abundant evidence that the prenatal period and the first two years of life are a critical window for intervention in growth and development: infection and poor nutrition during this window can negatively impact an individual’s long-term cognitive development and lifetime physiologic trajectory (Checkley 2003, Berkman 2002, Black 2008, Guerrant 1999, Niehaus 2002, Tarleton 2006, Bhutta 2008, Crimmins 2006, Grantham-McGregor 2007, Victora 2010). Nutritional interventions during the first years of life improve schooling and income in adolescents and adults up to 35 years later (Victora 2008, Hoddinott 2008). Yet, a systematic review of the impacts of complementary feeding and supplementation interventions reports that

---

**Protocol Title:** Measuring the benefits of sanitation, water quality, handwashing and nutrition interventions for improving health and development in rural Bangladesh

**Protocol Status:** APPROVED

**Date Submitted:** 10/26/2017

**Approval Period:** 10/30/2017-11/03/2018

**Important Note:** **This Print View may not reflect all comments and contingencies for approval. Please check the comments section of the online protocol. Questions that appear to not have been answered may not have been required for this submission. Please see the system application for more details.**

even the most successful of these interventions increase length-for-age Z-scores by 0.69 SDs, which is approximately 1/3 of the mean growth deficit for African and Southeast Asian populations (the mean intervention effect is 0.28 SDs) (Dewey 2008).

One hypothesis for why nutritional supplementation appears to be necessary but not sufficient to eliminate growth shortfalls is that chronic infection and colonization of the small intestine by fecal bacteria impedes nutrient absorption and creates low-level immune system stimulation, a condition called environmental enteropathy (Lunn 2000). Environmental enteropathy is characterized by damage to mucosa in the wall of small intestine that decreases its surface area for nutrient absorption and increases its permeability to antigenic molecules that stimulate immune system defenses (Lunn 2000, Campbell 2003). Biomarkers for intestinal permeability and immune system stimulation have been more strongly associated than acute diarrhea with growth shortfalls (Campbell 2003). The mucosal damage that characterizes environmental enteropathy is caused by the body's inflammatory response to the ingestion of fecal bacteria, and when people move to lower-bacteria environments the condition resolves (Haghighi 1997). Recently, nutritionists have hypothesized that reducing a child's fecal bacteria exposure during the first years of life through improved sanitation, handwashing or water treatment may improve gut function and subsequent growth (Humphrey 2009).

For children whose food intake is insufficient, lipid-based nutritional supplementation (LNS) helps reduce gross energy shortfalls and provide essential micronutrients. The energy and micronutrients that LNS provides are likely to improve nutritional indicators, length-for-age and child development (Adu-Afarwuah 2008, Adu-Afarwuah 2007, Dewey 2008, Walker 2007, Rosales 2009, Bryan 2004), particularly among children that are at highest risk for severe stunting (Phuka 2008, Phuka 2009). It is possible that improved nutrition alone can reduce the negative effects of infection on growth and development due to the improved ability of better-nourished children to fight off enteric infections and exhibit catch-up growth during the convalescence period (Guerrant 1992, Guerrant 2008).

The specific LNS we propose to use is a next generation supplement to Nutributter; members of our team (Drs. Dewey and Steward at UC Davis) have been involved in the development of the supplement and are currently deploying it in ongoing randomized, controlled trials in Bangladesh, Burkina Faso, Ghana and Malawi as part of the iLiNS project and related studies (iLiNS.org). Nutributter and related LNS interventions have demonstrated efficacy for improving child growth and development when provided daily after age six months (Adu-Afarwuah 2008, Adu-Afarwuah 2007). We propose a combined energy / micronutrient supplement because micronutrient supplementation alone is unlikely to have a large impact on linear growth (Ramakrishnan 1989). LNS is administered daily using 10 gram sachets that can be mixed into existing meals (e.g., porridge); a child eats two sachets per day. LNS is intended to supplement – and not replace – breastfeeding and locally available complementary foods, by providing 108 kcal/day and including a broad suite of essential fatty acids and micronutrients at dosages appropriate for children in this age group. It has an 18-month shelf life, does not spoil at high temperatures and costs as little as $0.10 per day. Its compliance has been over 88% in controlled trials (Adu- Afarwuah 2008), in part due to the ease of incorporating it into existing feeding routines. Breastfeeding is highly prevalent in both populations, and so we have focused on supplements that would not replace this essential source of nutrition (Black 2008).

Our collaborators at UC Davis have a series of ongoing randomized trials evaluating the impact of LNS supplementation provided to pregnant and lactating women and/or their infants in Ghana, Malawi, and Burkina Faso through the International Lipid Based Nutrient Supplementation Project (www.ilins.org). The

---

**Protocol Title:** Measuring the benefits of sanitation, water quality, handwashing and nutrition interventions for improving health and development in rural Bangladesh

**Protocol Status:** APPROVED

**Date Submitted:** 10/26/2017

**Approval Period:** **10/30/2017-11/03/2018**

**Important Note:** **This Print View may not reflect all comments and contingencies for approval. Please check the comments section of the online protocol. Questions that appear to not have been answered may not have been required for this submission. Please see the system application for more details.**

objectives of the project include the development of low-cost, acceptable LNS formulations using locally available ingredients and evaluation of the efficacy of reduced cost formulations of LNS for infants, young children and pregnant women. Acceptability trials have been conducted in all the three of the countries with positive results. Importantly, the iLiNS project has already demonstrated that LNS is acceptable among young children in similar cultures to the rural Bangladesh population.

Biomarkers for Environmental Enteropathy
Environmental enteropathy, an inflammatory disorder of the intestines that compromises nutrient absorption, is associated with child malnutrition and poor development (Haghighi 1997, Humphrey 2009, McKay 2010). Environmental enteropathy is one of the main hypothesized pathways for the impact of our interventions on growth and development. Measurement of environmental enteropathy symptoms will provide important information about the mechanism for intervention impacts (or lack of impact) in this study. Altered intestinal permeability is an indicator of environmental enteropathy, measured using a dual-sugar permeability test in which the lactulose:mannitol urinary excretion ratio is measured (Lunn 2000, Campbell 2003). The child is given a combination of the two sugars, lactulose and mannitol. Mannitol diffuses through a transcellular pathway and is used to assess the absorptive capacity and mucosal surface area of the enterocytes. Lactulose is typically minimally absorbed via the paracellular tight junctions and thus, it is used to assess epithelial integrity. A normal intestinal epithelium absorbs nearly all mannitol, but almost no lactulose. A damaged epithelium absorbs mannitol less efficiently and more lactulose. By measuring the lactulose: mannitol ratio in the urine passed over the subsequent 3-5 hours, the intestinal absorptive efficiency can be calculated and the severity of environmental enteropathy inferred.

Earlier studies demonstrated that environmental enteropathy as assessed by intestinal absorption is widespread in low income tropical countries where fecal contamination of water, food, and the environment are common in contrast to rarely being seen among normal residents of high income temperate countries (Haghighi 2003). Environmental enteropathy is acquired early in childhood. Stillborn children in tropical countries have normal intestinal small intestinal cellular structure (Haghighi 1979). During the first three months of life mannitol/lactulose absorption is normal in children in The Gambia compared to children in the UK, however after three months intestinal absorption among Gambian children progressively decreases during the first year of life (Lunn 1991). Recent studies in Bangladesh confirm intestinal malabsorption consistent with environmental enteropathy is present in children 3 – 24 months of age in the rural Dhamrai subdistrict; the degree of impairment in absorption increased in children between 3 and 12 months of age (Goto 2009a, Goto 2009b). The intestinal absorption and pathology of migrants who move from highly contaminated low income tropical countries to developed temperate countries normalizes within 3 to 5 years (Gerson 1971).

If improvements in sanitation, water quality, hand hygiene and nutrition could reduce the severity of intestinal malabsorption from environmental enteropathy either by preventing its acquisition or by reversing the pathology, this would represent an important contribution to global public health, and would be a useful outcome assessment for the larger planned intervention study. Nutritionists have recently argued that environmental enteropathy is most likely caused by poor sanitation and hygiene in low income countries (Humphrey 2009, McKay 2010). Yet, there are no studies that demonstrate a specific association between environmental enteropathy and poor sanitation separated from other exposures in a low income country environment, although one study from Rhodesia 30 years ago noted an association between intestinal absorption and socioeconomic status (Thomas 1976).

---

**Protocol Title:** Measuring the benefits of sanitation, water quality, handwashing and nutrition interventions for improving health and development in rural Bangladesh

**Protocol Status:** APPROVED

**Date Submitted:** 10/26/2017

**Approval Period:** **10/30/2017-11/03/2018**

**Important Note:** **This Print View may not reflect all comments and contingencies for approval. Please check the comments section of the online protocol. Questions that appear to not have been answered may not have been required for this submission. Please see the system application for more details.**

Pilot Environmental Enteropathy Work in Bangladesh (2010-11-2536)

In our Bangladesh environmental enteropathy pilot study, we selected 119 children from an existing cohort (SHEWA-B intervention assessment study) who lived in different levels of household environmental cleanliness based on sanitation, water quality and handwashing indicators. The children were between age 8 and 48 mo in May 2010 and lived in 83 different rural villages across Bangladesh.

The 66 children from households with improved household hygiene lived in homes with good sanitation (flush/septic/piped sewerage or a pit latrine with slab and water seal), good water quality (median E. coli < 10 CFU/100 ml in up to 8 samples collected over 24 mo), and favorable handwashing conditions (a dedicated location to wash hands stocked with soap and water). In contrast, the 53 children who lived in homes with poor household hygiene lacked adequate sanitation (open defecation, open pit latrines, slabs with broken water seals, toilets that flush to “somewhere else” or hanging toilets), had poor water quality (median E. coli ≥ 10 CFU/100 ml), and had unfavorable handwashing conditions (no dedicated location to wash hands, or a dedicated location that lacked either water or soap). The definitions of improved hygienic conditions were chosen to reflect indicators that we hope to improve through intervention in the WASH Benefits study.

Children in the two environments differed greatly in their growth: after statistical adjustment for potentially confounding differences, children in households with improved hygienic conditions had 0.54 SDs (95%CI 0.06, 1.01) higher HAZ than children in households with poor hygienic conditions (unadjusted difference = 0.91 SDs). Importantly, the children also differed in biomarkers for environmental enteropathy. After statistical adjustment for measures of socioeconomic status, children living in improved hygienic households had lactulose : mannitol (L:M) ratios that were –0.32 SDs lower than children living in poor hygienic conditions (95% CI –0.72, 0.08). Children in improved hygienic households also had lower Immunoglobulin G endotoxin core antibody (IgG EndoCAb) titers (–0.23 SDs, 95% CI: –0.63, 0.17) than children living in poor hygienic conditions. After adjusting for age and sex, the L:M ratio was also strongly associated with HAZ in the population: a 1-unit increase in the log L:M was associated with a –0.36 SDs reduction in HAZ (95% CI –0.64, –0.07).

These pilot results support our original rationale to conduct the main WASH Benefits study. However, because household environmental conditions in the pilot were not randomized, it remains possible that differences observed between the children in growth and EE biomarkers result from unmeasured or unquantifiable differences between groups that we cannot control for without an experiment. A randomized trial that delivers high impact household environmental interventions (i.e., interventions with good uptake and high efficacy at reducing pathogen transmission to young children) in large populations as we have in our Kenya and Bangladesh cohorts would provide more conclusive evidence.

Effect of the interventions on telomere length and allostatic load

Multiple in utero and early life exposures to biological and psychosocial stress may increase allostatic load (the cumulative biological damage from chronic stress) and increase susceptibility to disease later in life(Entringer et al., 2010; Shonkoff et al., 2009; Tomiyama et al., 2012). The attrition of telomeres, the repetitive DNA sequences protecting the tips of chromosomes, may serve as a biomarker of cumulative lifetime stress or play a causal role in the etiology of various diseases, or both(Entringer et al., 2011).

---

Telomere attrition may contribute to chromosomal instability, premature apoptosis, and organ damage(Armanios, 2013; Calado and Young, 2009). During the sensitive period of early postnatal life, cellular replication occurs at a rapid rate as the immune system, brain, and other systems develop(Zeichner et al., 1999). Since telomeres are a key determinant of tissue development and shorten at a dramatically faster rate in infancy compared to in adulthood, it is efficient to focus on early childhood factors that may accelerate telomere attrition(Frenck et al., 1998; Zeichner et al., 1999). Although telomere attrition within the context of various diseases in adult populations has been widely studied(Calado and Young, 2009; Lin et al., 2012), little is known about the pregnancy and early life risk factors associated with telomere attrition in infants from low-income countries. Complex pathways connect early life insults – micronutrient deficiencies, environmental enteropathy, and family violence – to adverse child health outcomes, and accumulating evidence implicates telomere attrition, allostatic load, inflammation, and growth factors as potentially important underlying mechanisms linking these environmental stressors and disease susceptibility. The trial design will enable us to a) measure the effect of the interventions on telomere attrition, b) examine the association between environmental enteropathy, telomere attrition, linear growth faltering, and poor cognitive development, and c) evaluate the impact of maternal psychological stress on child allostatic load, telomere length, growth trajectories, and cognitive development.

Micronutrient deficiencies may accelerate telomere attrition in children. Since the vast majority of cells are engaged in the DNA synthesis phase during early childhood development, the additive or synergistic effects of several micronutrient deficiencies could produce destructive effects on genome stability leading to negative health sequelae later in life(Fenech, 2005). Micronutrients maintain the genome by serving as cofactors for enzymes, participating in DNA synthesis and repair, and inhibiting oxidative stress-induced DNA damage(Bull and Fenech, 2008). In Bangladeshi children ages 24-48 months, 97% of children had inadequate folate intake(Arsenault et al., 2013). The thymidine-rich telomere repeat sequence, (TTAGGG)n, may be highly susceptible to folate-deficient conditions that favor the incorporation of elevated levels of uracil into the DNA rather than thymidine, which then leads to chromosome breaks(Blount et al., 1997). The impact of specific micronutrients on telomere length requires further study, and telomere attrition, in turn, may emerge as a sensitive marker of nutritional deficiency. This trial would be the first to examine the potential association of micronutrient deficiencies and telomere attrition and to measure the impact of nutrition interventions on telomere lengths in children.

The chronic infections and inflammation endemic in low-income countries with poor WASH may contribute to telomere attrition. When human adult subjects were experimentally exposed to a common cold virus, those with longer telomeres displayed more resistance to acute upper respiratory infection and clinical illness compared to those with shorter telomeres(Cohen et al., 2013). Celiac disease and environmental enteropathy share similar histologic features of intestinal inflammation, and the telomeres from small intestinal biopsies in individuals with celiac disease were shorter than the telomeres of healthy controls(Cottliar et al., 2003). An important question to explore within the context of the study is whether environmental enteropathy affects telomere attrition or whether telomere attrition exacerbates or increases susceptibility to environmental enteropathy. Furthermore, we will elucidate the potential associations between telomere lengths, linear growth, and cognitive development.

The association between psychological stress, telomere attrition, and growth faltering could be mediated through glucocorticoid and immune activation and oxidative stress. The hypothalamic-pituitary-adrenal (HPA) axis serves a vital role in the neuroendocrine systemic response to stress, and its contribution to telomere attrition has not yet been elucidated. Studies have demonstrated that chronic stress leads to glucocorticoid receptor resistance, a decreased sensitivity of immune cells to cortisol(Cohen et al., 2012;

Miller et al., 2002). Due to a lack of glucocorticoid regulation, a prolonged pro-inflammatory cytokine response ensues causing damage to multiple systems throughout the body. This dysregulation of the pro-inflammatory response is particularly detrimental when it occurs during the first two years of a child's life, the critical window of growth, because it negatively affects the growth hormone/insulin-like growth factor 1 (GH/IGF-1) endocrine axis. Growth hormone stimulates the secretion of IGF-1, an important regulator of cell proliferation, immunity, and inflammation(Deelen et al., 2014). During acute stress or infections, the body utilizes the pro-inflammatory cytokines to restrict growth and energy storage and instead, redirects energy to ensure survival(O'Connor et al., 2008). During the first two years of life, a period of rapid growth and development for a child, chronic stress induces a protracted pro-inflammatory response that diverts energy towards the immune system by dampening the anabolic activities of IGF-1 and instead promoting protein catabolism, thereby contributing to childhood stunting(Livingstone, 2013). Several child cohort studies have suggested associations between increased cortisol secretion, decreased IGF-1 levels, and greater risk of growth faltering(Cianfarani et al., 2002; Cianfarani et al., 1998; Idohou-Dossou et al., 2003; Kilic et al., 2004). The potential association between IGF-1 and telomeres has not yet been examined in children. Two potential applications of understanding this molecular pathway are 1) to develop biomarkers to evaluate the efficacy of interventions and 2) to generate possible targets for intervention to alter child growth trajectories.

Overall, this study will extend our understanding of how biological and social determinants, specifically micronutrient deficiencies, environmental enteropathy, and psychological stress, shape child health outcomes on a cellular level. This trial will be the first to evaluate the impact of water, sanitation, hygiene, and nutrition interventions on telomere length in infants living in Bangladesh. The insights from this research could build upon existing foundations to implement and assess holistic strategies to improve nutrition, decrease fecal contamination, reduce family stress, and ultimately cultivate a healthy environment to promote telomere elongation and advance child development.

Assessment of interactions

Since intervention assignment will be randomized the study groups will have similar population characteristics that will permit inferring that observed differences in outcome are attributable to the intervention. However, we expect that the interventions will have different effects in different children and that some of these differences will be mediated by differences in the mother and child's intestinal microbiome, immune function, internal biochemical environment and genetic disposition. For example a child who acquires a specific pattern of microbiological organisms from her mother's intestinal microbiome may be more resistant to infection with intestinal pathogens and so less likely to benefit from sanitation interventions. Immune function, internal biochemical environment including nutrition and genetic characteristics are especially dynamic areas of scientific research where important new insights are emerging each month. The capacity to explore these issues will provide the opportunity to understand the mechanism and generalizability of any impacts on child health and development that are observed within the study.

One exposure that has the potential to interact with child development is lead. Humans exposed to lead experience irreversible impairment of intellectual function (Bellinger et al., 1992). Two studies of residents living in rural Bangladesh remote from roads and industry report unexpectedly high blood lead levels including 14% of children in a rural area of Dinajpur District having blood lead levels >10 μg/dL (Mitra et al., 2009) (twice the current 5 µg/dL level used to identify US residents at high risk (Advisory Committee on Childhood Lead Poisoning Prevention, 2012)) and postpartum women in rural Matlab, where the median equivalent blood lead concentration of 6.0 µg/dL exceeded the high risk threshold (Bergkvist et al., 2010). A few studies have explored potential sources of lead in rural Bangladesh. Lead concentrations in soil used for agriculture in Mymensingh District, Bangladesh was twice as high as soil collected from adjacent plots used for non-agricultural domestic purposes (40.6 ppm versus 20.7 ppm) (Muhibbullah et al., 2005). Rice, the primary dietary staple in Bangladesh, collected from households in the Matlab study contained a median of 25 µg/kg of lead (Bergkvist et al., 2010).

Environmental Microbial Assessment

Fecal-oral pathogen transmission is a complex process. The complexity arises from a multitude of transmission pathways, a broad diversity of pathogens, the importance of environmental conditions, and interactions between the environment and human behavior. Water, sanitation and hygiene interventions present primary and secondary barriers that separate feces from the environment and should block enteric pathogen transmission. Measuring fecal contamination along environmentally mediated pathogen transmission pathways, including water, hands, soil, food and flies, will enable us to understand which of these pathways are successfully broken by our interventions and elucidate the factors behind their success or failure in improving child development outcomes. Detailed assessment of contamination along these pathways (including measurement of fecal indicator bacteria, microbial source tracking to differentiate between human and animal sources of contamination, and detection of common diarrheagenic pathogens such as pathogenic E. coli, Shigella and rotavirus using culture-based and molecular techniques) will allow a nuanced understanding of the impact of the interventions on disease transmission in young children in the rural Bangladeshi setting. In-depth information on environmental contamination will also allow us to explore the relationship between environmental enteropathy in children and fecal contamination in their living environment.

Spillovers of WASH Interventions
While there is a rich literature on the health effects of WASH interventions for children receiving such interventions, to our knowledge, no studies have measured the health effects of such interventions on children that are geographically proximate to WASH intervention recipients who did not receive the intervention themselves. Effects of interventions on those not receiving interventions are termed “spillovers,” and failing to account for spillovers in the same direction as the effect on the treated (“positive spillover”) will lead to underestimates of the efficacy and cost effectiveness of an intervention. As such, measurement of spillovers is important for the prioritization of interventions and allocation of public funds. Although many studies have applied mathematical models to spillovers of infectious diseases, very few studies have empirically measured spillovers for infectious diseases (Anderson & May 1992; Anderson & Medley 1985; Medley et al. 1993; Basáñez et al., 2012; Chan et al, 1994; Magalhães et al., 2011; Halloran et al., 2002; Bansal et al., 2006; O'Brien et al., 2007). Our proposed study will be one of the first to do so, and it will be the first to generate empirical spillover estimates for WASH interventions. This study will generate unique evidence to inform the estimation of the cost effectiveness of interventions and optimal resource allocation to maximize health benefits for rural populations in developing countries. It will concurrently advance the general methodology for spillover measurement in multiple disciplines.

Effect of the interventions on anemia
Anemia in preschool children results in poor physical growth, impaired cognitive development, reduced school achievement and, when severe, may result in increased mortality risk. The prevalence of preschool

---

**Protocol Title:** Measuring the benefits of sanitation, water quality, handwashing and nutrition interventions for improving health and development in rural Bangladesh

**Protocol Status:** APPROVED

**Date Submitted:** 10/26/2017

**Approval Period:** 10/30/2017-11/03/2018

**Important Note:** This Print View may not reflect all comments and contingencies for approval. Please check the comments section of the online protocol. Questions that appear to not have been answered may not have been required for this submission. Please see the system application for more details.

child anemia (hemoglobin concentration<110 g/L) has been estimated at 64% in Bangladesh. While it is recognized that anemia is a multifactoral disorder, the relative contribution to anemia from micronutrient deficiency, infection, and hemoglobinopathies among low-income populations has not been well characterized. It has been assumed that more than half of the burden of anemia can be attributed to iron deficiency, yet the interplay between undernutrition and infection may reveal a more complicated story. Although iron supplementation interventions have had modest success at reducing anemia risk, high rates remain even after supplementation. This may be due to a failure to address the clinical and sub-clinical infectious causes of anemia. Certain parasitic or other enteric infections such as diarrheal disease may also contribute to anemia due to blood loss or inflammation. It has been hypothesized that environmental enteropathy (EE) is one such condition in which repeated often subclinical enteric infection, thought to be due to poor water, sanitation, and hygiene, results in a chronic state of gut inflammation and nutrient malabsorption that may contribute to anemia (Prendergast et al, 2012). No interventions designed to both improve nutrition and reduce infection simultaneously have rigorously evaluated the impact on anemia. The trial's randomized, factorial design will enable us to experimentally measure the independent and combined effects of interventions to reduce infection and undernutrition in young children. The study's findings will contribute to the critical evidence gap regarding the causes of anemia and interventions to prevent it. The results of this study are likely to be broadly applicable to other rural, low income populations where food insecurity, poor access to safe water, and inadequate sanitation coexist and could identify new strategies to address this important and intractable problem.

##### 3. Collaborative Research

**a) If any non-UCB institutions or individuals are engaged in the research, explain their human research roles and what human subjects training they have/PI has planned to provide.**

The following collaborating institutions are are also engaged in the research, in that they will be involved in obtaining consent and collecting data data through contact with human subjects:

The International Centre for Diarrheal Disease Research, Bangladesh. This organization will employ the staff that will engage in the field work and have contact with human subjects. They will work with the raw data that includes personal identifying data. Mahbubur Rahman is a Public Health Specialist at ICDDR,B who is responsible for managing field logistics for this study.

Steve Luby from Stanford University will coordinate collaborators across institutions, provide input on intervention, protocol, and questionnaire development, and contribute to analysis and manuscript writing responsibilities. Mark Davis at Stanford will contribute to the immunological analysis in the environmental enteropathy subset.

Collaborators at UC Davis will support the development of the nutrition intervention, questions related to nutrition intervention, and uptake measures related to nutrition.

Collaborators at Emory University will provide input on measurement and will help to ensure that the Bangladesh protocol is aligned with the WASH Benefits Kenya protocol.

Michael Kremer from Innovations for Poverty Action will provide input on measurement and will help to ensure that the Bangladesh protocol is aligned with the WASH Benefits Kenya protocol.

---

**Protocol Title:** Measuring the benefits of sanitation, water quality, handwashing and nutrition interventions for improving health and development in rural Bangladesh

**Protocol Status:** APPROVED

**Date Submitted:** 10/26/2017

**Approval Period:** **10/30/2017-11/03/2018**

**Important Note:** **This Print View may not reflect all comments and contingencies for approval. Please check the comments section of the online protocol. Questions that appear to not have been answered may not have been required for this submission. Please see the system application for more details.**

Collaborators from Emory University and Innovations for Poverty Action are not directly engaged in the proposed research but will serve in an advisory role.

Wagner College. Dr. Mohammed Alauddin will contribute to the environmental enteropathy portion of the study. He is an Analytical Chemist, and he will measure one of the important biomarkers of EE, namely Lactoluse-Mannitol (LM) ratio in urine collected from children in the WASH Benefits study. Dr. Alauddin will employ highly sensitive technique of high performance liquid chromatography and mass spectrometry (LC-MS/MS) for the urinary analysis of lactoluse and mannitol in children. The analysis will be carried out in Dr. Alauddin's research laboratory at Wagner College backed by appropriate method validation and quality control check. In addition to the LM analysis in urine samples, Dr. Alauddin will collaborate with the UC Berkeley team and the ICDDRB team in Bangladesh in the data analysis, interpretation, report writing and successful completion of the project. He will have access to identifiable data.

VitMin Lab. Dr. Juergen Erhardt will conduct micronutrient and acute phase protein panel analyses of blood samples as part of the nutritional assessment. Dr. Erhardt will not have access to identifiable information. He is not involved in the study design.

**b) If any non-UCB institutions or individuals are collaborating in the research, complete the table below and attach any relevant IRB approvals in the Attachments section.**

##### Non-UCB institutions

| Institution Name | Individual Contact/ Affiliate of Institution | FWA # | Local IRB Review? (Y or N) | IRB Approval Date | IRB Approval Expiration Date |
|---|---|---|---|---|---|
| UC Davis | | | N | | |
| International Centre for Diarrheal Disease Research, Bangladesh | Mahbubur Rahman | 00001468 | Y | 04/09/2017 | 04/08/2018 |
| Emory University | | | N | | |
| Stanford University | Steve Luby | 00000935 | Y | 10/31/2016 | 10/31/2017 |
| Innovations for Poverty Action | Michael Kremer | | N | | |
| Wagner College | Mohammed Alauddin | None | N | | |
| VitMin Lab | Juergen Erhardt | | N | | |

---

#### 4. Qualifications of Study Personnel

**a) Explain expertise of Principal Investigator, Student/Postdoc Investigator, Faculty Sponsor (if applicable), any Co-Investigators or other key personnel listed in the application, and how it relates to their specific roles in the study team.**

John M. Colford, Jr., MD, PhD (Principal Investigator)
Dr. Colford obtained his M.D. from Johns Hopkins School of Medicine, and his Ph.D. in Epidemiology from the University of California, Berkeley (1996). He is a Professor of Epidemiology in the School of Public Health at UC Berkeley. Dr. Colford has extensive experience in and prior funding from NIH, CDC, and the USEPA for the conduct of randomized trials and other studies of drinking water and environmental health issues internationally and in the United States. Dr. Colford will ensure communication with the Gates Foundation, oversee collaboration between the University of California Berkeley and ICDDRB, oversee protocol approval process through the University of California Berkeley and assure that the project activities are aligned with the obligations to the Gates Foundation.

Benjamin Arnold, PhD (Key Personnel)
Dr. Arnold is an epidemiologist in Dr. Colford's research group. He has participated in numerous studies of water, sanitation and hygiene interventions throughout Asia, Africa and Latin America. He will contribute to the study design, development of survey instruments, data analysis and scientific publications.

Alan Hubbard, PhD (Key Personnel)
Dr. Hubbard is an Associate Professor of Biostatistics, School of Public Health, UC Berkeley. He will provide guidance for the study design and statistical analysis.

Lia Fernald, PhD (Key Personnel)
Dr. Fernald, Associate Professor of Public Health Nutrition, will lead the design and execution of the outcome measures concerned with child growth and development (a role similar to that which she currently has the on Gates/World Bank WSP Project).

Kara Nelson, PhD (Key Personnel)
Dr. Nelson is a Professor of Environmental Engineering. She will oversee the environmental microbial assessment and related data analysis activities.

Jade Benjamin Chung (Graduate Student Researcher)
Jade coordinated the pilot environmental enteropathy study in Bangladesh, and is now a doctoral student at UC Berkeley. Jade will assist with the environmental enteropathy assessment and nutrition intervention implementation for the main trial in Bangladesh.

Audrie Lin, PhD (Postdoctoral Researcher)
Ms. Lin will provide guidance on the environmental enteropathy assessment and related data analysis activities. She holds a PhD in Microbiology and has experience conducting field work in Bangladesh.

---

**Protocol Title:** Measuring the benefits of sanitation, water quality, handwashing and nutrition interventions for improving health and development in rural Bangladesh

**Protocol Status:** APPROVED

**Date Submitted:** 10/26/2017

**Approval Period:** 10/30/2017-11/03/2018

**Important Note:** **This Print View may not reflect all comments and contingencies for approval. Please check the comments section of the online protocol. Questions that appear to not have been answered may not have been required for this submission. Please see the system application for more details.**

Ayse Ercumen, PhD (Postdoctoral Researcher)
Dr. Ercumen will provide guidance to the environmental microbial assessment and related data analysis activities. She holds a PhD in Epidemiology and has experience conducting field work in Bangladesh.

Co-Investigators from Stanford University
Steve Luby, MD
Dr. Luby is a Professor of Infectious Diseases at Stanford University and is a Co-Principal Investigator. He will co-lead the study in Bangladesh. He will draft the protocol, coordinate collaborators, assign project implementation, analytic, report and manuscript writing responsibilities.

Amy Pickering, PhD (Postdoctoral Researcher)
Dr. Pickering will provide guidance to the environmental microbial assessment and related data analysis activities. She holds a PhD in Environmental Engineering and has experience conducting field work in Bangladesh.

Jessica Grembi, PhD Student
Ms. Grembi will lead the microbiome analysis.

Mark Davis, PhD
Dr. will support the immunological analysis in the environmental enteropathy subset.

Researchers from ICDDR,B
Leanne Unicomb, PhD, MMed Sci (Co-Investigator)
Dr. Unicomb (epidemiologist) is a Co-Principal Investigator.  She will supervise the Bangladesh based project implementation team, ensuring coordination and collaboration among research investigators, supporting project management, and drafts of the manuscript on the water treatment intervention and changing the water treatment indicators after intervention.

Tahmeed Ahmed, MBBS, PhD (Co-Investigator)
Dr. Ahmed is a pediatrician and nutritionist with broad experience conducting research on undernutrition among children in Bangladesh including collaborating with Dr. Dewey and colleagues at the University of California Davis on studies of Lipid based Nutrition Supplement in Bangladesh. Dr. Ahmed will oversee the nutrition intervention in Bangladesh, work with his team to ensure that measurements of child growth are valid, and assist in study design data interpretation and manuscript writing.

Mahbubur Rahman, MBBS
Mr. Rahman is Public Health Specialist at ICDDR,B who is responsible for managing field logistics for this study.

Abu Naser, MBBS
Mr. Naser is a Public Health Specialist at ICDDR,B who is responsible for the post-intervention assessment.

Co-investigators from UC Davis

Kathryn Dewey, PhD
Dr. Dewey is a Professor of Nutrition and Director of the Program in International and Community Nutrition

---

**Protocol Title:** Measuring the benefits of sanitation, water quality, handwashing and nutrition interventions for improving health and development in rural Bangladesh

**Protocol Status:** APPROVED

**Date Submitted:** 10/26/2017

**Approval Period:** **10/30/2017-11/03/2018**

**Important Note:** **This Print View may not reflect all comments and contingencies for approval. Please check the comments section of the online protocol. Questions that appear to not have been answered may not have been required for this submission. Please see the system application for more details.**

at UC Davis. Dr. Dewey's research area is maternal and infant nutrition, in both affluent and low income countries. She has conducted research in the U.S., Mexico, Costa Rica, Honduras, Guatemala, Peru, Indonesia, Thailand and Ghana. Her professional service includes extensive consultation for WHO, UNICEF, the Pan American Health Organization and the March of Dimes, and serving as President of the Society for International Nutrition Research (2000-02) and of the International Society for Research on Human Milk and Lactation (2006-2008). She currently serves as chair of the Technical Advisory Group for the Alive and Thrive Project and is a member of the Steering Committee for the WHO Multicentre Growth Reference Study and the NIH Technical Working Group on Iron and Malaria. Dr. Dewey will collaborate on the nutritional components of the study.

Christine Stewart, PhD MPH
Dr. Stewart is an Assistant Professor of Nutrition at UC Davis and her research focuses on maternal and child nutrition in low income countries. Dr. Stewart will oversee the nutrition intervention activities in Kenya, and will advise all co-investigators on the LNS intervention protocol, monitoring and nutritional outcome measurement in the study. She will also assist in study design, data interpretation and manuscript writing.

Kendra Byrd, M.S.
Kendra Byrd is a graduate student researcher who will be studying the association between inflammation, hemoglobinopathies and the iron regulatory protein hepcidin in the study data.

Charles Arnold, M.S.
Charles Arnold is a statistician who will be assisting with the analysis of the data on micronutrient status, infant feeding, and anemia.

Co-intestigators from Innovations for Poverty Action

Clair Null, PhD (IPA Afflilliate and Emory University)
Dr. Null is an Assistant Professor of Global Health at the Rollins School of Public Health, Emory University. Dr. Null is the Principal Investigator of the parallel study in Kenya. She received a PhD in Agricultural and Resource Economics at the University of California at Berkeley in 2009. She will assure that the activities in Bangladesh and Kenya are sufficiently
aligned so that the subsequent interventions will be sufficiently similar so that the data can be combined, and will oversee the translation of the Kenya experience to Bangladesh.

Michael Kremer, PhD (IPA Affiliate).
Dr. Kremer will lead the design and evaluation of the behavioral economics components of the study

Consultants
In addition to Co-investigators and collaborators at IPA, ICDDRB, and UC Davis, a number of other individuals are providing expert advise on various aspects of the research. These experts will not be engaged in human subjects activities related to the project. These experts include Tom Clasen (Senior Lecturer in the Department of Infectious and Tropical Diseases, London School of Hygiene & Tropical Medicine), Patricia K. Kariger (Ph.D.), Dr. Pavani Ram (Assistant Professor of Social and Preventative Medicine, University at Buffalo), Dr. Elli Leontsini (Associate Professor of International Health, Johns Hopkins University) and Dr.
Peter Winch (Professor of International Health, Johns Hopkins University).

Mohammed Alauddin, Wagner College
Dr. Alauddin graduated in Chemistry from the University of Dhaka in 1975, earned his Ph.D. in Analytical Chemistry from the University of Kentucky, USA in 1982. His research includes the application of analytical techniques in solving problems of environmental, biological and geological significance. Dr. Alauddin is actively engaged in projects dealing with women and child health issues in Bangladesh in collaboration with institutions in the USA, Australia and Bangladesh.

Juergen Erhardt, Hohenheim University
Juergen Erhardt received a PhD in human nutrition is an expert in laboratory assessments of Vitamin A and iron status.

b) **In case of international research, describe the expertise you have, or have access to, which prepares you to conduct research in this location and/or with this subject population, including specific qualifications (e.g., relevant coursework, background, experience, training). Also, explain your knowledge of local community attitudes and cultural norms, and cultural sensitivities necessary to carry out the research. See Human Subjects Research in an International Setting and <a href='http://cphs.berkeley.edu/international.pdf' target='_blank'>CPHS Guidelines on Research in an International Setting.**

The core team will be assisted by many persons at ICDDR,B (these persons are listed as key personnel on the IRB submission to ICDDR,B), in addition to international consultants with specific expertise and experience in nutrition, child development, sanitation, handwashing, and tropical enteropathy. All consultants will participate in the development of the intervention and training modules for their specific areas and develop field quality assurance protocols for their materials. They will not participate in activities involving contact with human subjects nor in the analysis of data that might include personal identifiers. The
consultants include Patricia Kariger, PhD (child development expert), Thomas Clasen, MSc, PhD, JD (water and sanitation expert), Pavani Ram, PhD (handwashing expert), Elli Leontsini, PhD (behavior change expert) and Peter Winch, PhD (behavior change expert).

---

#### * * * Subject Population * * *

##### 5. Subject Population

**a) Describe proposed subject population, stating age range, gender, race, ethnicity, language and literacy.**

The subject population will be young children and their mothers/guardians living in approximately 3-4 areas of Bangladesh where communities meet the following study criteria:

•Rural communities
•Drinking water
  o low levels of iron (<1mg/L on average) and arsenic (<50 mg/L on average) as documented in the collaborative assessments by the Government of Bangladesh and the British Geological Survey and internal testing using Hach iron kits
  o Sources known to be frequently contaminated with fecal indicator bacteria (including shallow tubewells)
•Low levels of fully hygienic latrines coverage as indicated by the Multiple Indicator Cluster Survey
•Levels of childhood stunting greater than or equal to 30%
•That the Government of Bangladesh, international non-governmental organizations working in Bangladesh and local government authorities report that no major water, sanitation, or focused nutrition programs are currently operating or planned in the area in the next 2 years.
  o (All participating communities will remain free to engage in intervention opportunities which they see as in their best interest. If water, sanitation, hygiene or nutrition promotion activities are initiated in the study community during the course of the trial, this will complicate the inferences we can draw from these areas, but we will not drop any such communities from the analysis, but, as noted in the analytic plan, we will retain the intention to treat analysis.)

  o Specifically avoiding hoar areas, hill tracks and coastal belts (i.e. significant flood risk areas)

Target children will be unborn children of pregnant women identified by report of their last menstrual period at enrollment. Target children will age to between 17 and 24 months over the course of the study. Older siblings or neighbors (age 18 – 27 months) of the target children will also be included in the study. The subject population will include both males and females, and no one will be excluded based on their race, ethnicity, language or literacy.

**b) State the maximum number of subjects planned for the study. This number should account for all subjects to be recruited, including those who may drop out or be found ineligible. Explain how number of subjects needed to answer the research question was determined.**

Figure 1 (attached) provides an overview of the WASH Benefits study design (The Kenya study is described in a separate protocol). The interventions will require about 3 months from the baseline survey to deliver. The follow-up rounds are planned for 12 and 24 months after intervention delivery.

In Bangladesh, we plan to enroll 90 clusters per intervention arm, a double-sized control arm, and 7 children per cluster. Because of the risk of pregnancy loss, we will enroll 8 pregnant women per cluster.

The control arm will include a maximum of 1,440 children and 1,440 mothers and the 6 intervention arms will include a maximum of 4,320 children and 4,320 mothers. The total number of children planned for the study is 5,040, but because of the uncertainty of pregnancy outcome our goal will be to enroll 5760 target children, 5760 older siblings/neighbors of the target children, and 5760 mothers of target children. We will also interview 540 promoters. We will collect additional samples from 1800 children aged approximately 18-27 months at baseline who live in the same compounds as target children. We will enroll 2,500 children neighboring children enrolled in the study as part of the spillover substudy. During the qualitative assessment of spillovers, we will ask 30 participants to mention the names of individuals who they are closely socially connected to. The maximum number of unique people that each person could mention is 9 people, and assuming no overlap in these individuals across interviews, the maximum number of people named is 270. The maximum number of subjects we will enroll in the whole study is 22,390 people (5760 target children + 5760 older siblings/neighbor children + 5760 mothers of target children + 540 promoters + 1800 children 18-27 months at baseline + 2500 children in the spillover study + 270 people in spillover qualitative assessments.)

HAZ Sample Size Calculations:
Using data from the SHEWA-B assessment conducted by the icddr,b Water and Sanitation Research Group with included a large sample of rural Bangladesh, the mean (SD) HAZ among the 1,413 children < 3 years old with height data is -1.825 (1.243). The village level intra-class correlation is 0.01. There are two hypotheses of interest: (H1) comparison of individual treatment arms vs. control, and (H3) comparison of the combined nutrition+WSH arm to the nutrition arm. We have slightly more power for comparison to control for due to the double sized control arm. We identified the minimum detectable effect (MDE) for the two comparisons with > 80% power. We have assumed a one sided α of 0.05. With the current design, we will have power to detect differences of +0.15 HAZ; this is approximately half the mean effect size observed in supplemental feeding intervention studies (Dewey 2008). We have targeted a smaller effect size because we expect the impacts from water, sanitation, and handwashing interventions to be smaller than from nutritional interventions.

Diarrhea Sample Size Calculations:
In the Bangladesh SHEWA-B dataset, described above, the 48 hour period prevalence of diarrhea is 12.5%. For all calculations, we have assumed that the diarrhea prevalence in the control group will be 12.0%. For comparisons of the combined WSH arm to any single treatment arm we assume that the single treatment arm prevalence is 8.0% (a 33% relative reduction from the control). There are two hypotheses of interest: (H1) comparison of individual treatment arms vs. control, and (H2) comparison of impacts in the combined WSH arm with the individual treatment arms. We identified the minimum detectable effect (MDE) for the two comparisons with > 80% power (see Table 1). We have assumed a one sided α of 0.05.

Environmental Enteropathy Subgroup Sample Size Calculations:
We will collect blood, serum, urine, and stool specimens from a subsample of 2,000 children in the study to measure biomarkers for environmental enteropathy. Environmental enteropathy is one of the key hypothesized mechanisms for intervention impact on child growth and development. The sample of 2,000 children will be distributed equally over four arms in the study – 500 children in each of the control arm, LNS alone arm, combined WSH arm, and the LNS+ combined WSH arm. Specimen collection will take place 3 months following baseline, at midline (1 year following baseline), and endline (2 years following

baseline). We expect that only 500 of the 2,000 children in the subsample will have been born at the time of our first EE assessment. We plan to collect specimens on the entire subsample at the 1-year and 2-year follow-up measurements.

We arrived at a sample size of 2,000 children (500 children per arm) using two outcome measurements we collected in our Bangladesh environmental enteropathy pilot study, the lactulose:mannitol (L/M) ratio and Endotoxin Core Antibody (EndoCAb). In the pilot, the 119 children ranged in age from 10 to 48 months and lived in 83 villages across rural Bangladesh. Using estimates of variability in the lactulose:mannitol (L/M) ratio and Endotoxin Core Antibody (EndoCAb), we estimate that this design will have greater than 80 percent power to detect differences between groups of –0.20 SDs in the L:M ratio and –0.25 SDs in IgG EndoCAb antibodies. These detectable differences are smaller than those observed between children in poor versus improved hygiene households in the pilot study: –0.42 SDs for L:M and –0.29 SDs for EndoCAb. In these calculations, we assumed a village-level intra-class correlation of 0.05 for L:M and 0.27 for EndoCAb, and child-level intra-class correlations (for repeated measures within children) that range between 0.5 and 0.9.

Intestinal Parasite Measurement in Target Children and their peers:
At baseline, we plan to collect stool specimens and blood spot samples from children 18 – 27 months who live in the same compound as the target children (7 children per cluster; 5,040 total). At the 2-year follow-up, we will collect stool specimens and blood spot samples from target children, from the same 18-27 month old children that provided a sample at baseline, and from an additional older child that is 5-12 years old at endline and lives in the same compound as the target child. The purpose of the stool collection is to measure the presence and intensity of intestinal parasite infections. For the stool samples, we propose to only measure protozoan parasites in this young age group because we expect the prevalence of these organisms to be reasonably high. The eventual goal of the blood spot collection will be to analyze the samples at some point in the future for intestinal helminths and protozoans using antigen-based assays.

We estimate that these samples will be sufficient to detect a relative reduction of 18% in infection prevalence. Our power calculations assume 50% prevalence in the control arm, a village intraclass correlation of 0.14, and 71% successful stool collection and analysis (10 / 14 samples per village), which is highly conservative.

Spillover Study Sample Size Calculations:
Since spillover effects are likely smaller than the direct effects of the intervention, we will need to measure outcomes in more children per arm in the spillover study than are enrolled in the main trial. We aim to detect a relative reduction of 6% in our primary outcomes. We calculated the prevalence and intraclass correlation coefficients (ICC) for diarrhea from a WASH Benefits pilot study and soil-transmitted helminths from a study of children in rural India. In the WASH Benefits pilot, the prevalence of Ascaris and Trichuris was 7.5% and 10.4%, respectively, in households with poor hygiene and 20.7% and 13.8%, respectively, in households with good hygiene. Our sample size calculations did not focus on respiratory illness due to their higher prevalence relative to diarrhea and helminth infection. The ICCs ranged from 0.023 to 0.153, and since these are somewhat larger than ICCs reported in the literature, we expect our sample size estimates are conservative. We assumed that we would measure 10 children per cluster. Assuming 80% power and a type I error of 0.05, we calculated the required sample size for each outcome of interest, adjusting for the ICC. Given these assumptions, the spillover study plan to enroll 2,000 children in 180 clusters (1,000 children and 90 clusters per arm).

**c) If any proposed subjects are children/minors, prisoners, pregnant women, those with physical or cognitive impairments, or others who are considered vulnerable to coercion or undue influence, state rationale for their involvement.**

The proposed subjects include very young children, pregnant women, and educationally and economically disadvantaged subjects. The goal of the WASH Benefits study is to generate rigorous evidence about the impacts of sanitation, water quality, handwashing, and nutrition interventions on child growth and development in the first years of life. There is abundant evidence aggregated over more than 325,000 children from around the world that the window for interventions to improve growth is in the first 1,000 days of life, including the 9 months before birth (Victora 2010). Meeting the study goals requires intervention in the middle of this development window. Enrolling pregnant women will ensure our ability to meet our sample size goals. The rural population that we are targeting in this study are very poor, and lack the water, sanitation, and hygiene infrastructure which we are assessing.

#### 6. Recruitment

**a) Explain how, where, when, and by whom prospective subjects will be identified/selected and approached for study participation. If researcher is subject's instructor, physician, or job supervisor, or if vulnerable subject groups will be recruited, explain what precautions will be taken to minimize potential coercion or undue influence to participate. See CPHS Guidelines on Recruitment for more information.**

The intervention trial will be implemented in 5 districts of rural Bangladesh. Eligible communities will be identified through area surveys conducted by ICDDRB research assistants. The fieldwork will be implemented by local field workers recruited and supervised by the Bangladesh-based scientific team. The trained fieldworkers will travel to the eligible communities and will ask community leaders for permission to conduct research within their community. If the community leaders agree, then the team will proceed with recruitment. A local village leader will accompany the field staff during their first visit to potential study households at the time of enrollment to ensure that subjects are fully informed of the implications of study participation and to avoid any potential mistrust in the community. Fieldworkers will approach randomly selected, eligible baris (a group of 3-20 households that share a common courtyard and are usually blood relatives) within a community. Only compounds that include a pregnant woman in the first or second trimester of her pregnancy with self reported low levels or no iron problems in their drinking water, who will stay in this household for the next 24 years are identified as eligible by the team for participation. Occasionally compounds may have multiple pregnant mothers and both eligible households are listed in this case. The prospect of participation in the study will be discussed with adults in the compound, including the pregnant mother/caregiver of the target infants. If a potential respondent is not able to read the consent form, the field worker will suggest that they invite a witness to help read the form with them so that they can be certain that they agree to participate in the procedures in the study. If a witness is present, both the respondent and witness will be asked to provide a thumb print or signature. After providing time for discussion among the compound residents and verbal interest to participate, a member of the field team records the GPS coordinates at the front door of the mother’s household. These coordinates will be compiled and analyzed using ArcGIS mapping software, to identify 8 closest mothers for a cluster. A buffer region of 1 km is then excluded around this cluster, before identifying the next cluster of 8 mothers. Field teamsreturn and seek formal informed consent from the head of the selected compounds and from the pregnant mother/guardian of target infants within each cluster during enrollment.

The intervention is a cluster randomized controlled trial. Each cluster will be a group of compounds that includes at least seven eligible children. The compounds within a single cluster will be located closely enough together so that a single hygiene promoter can reach each of the participating compounds by walking. If the compounds are too dispersed for a hygiene promoter to reach all of them on foot, then they will not be enrolled in the study. More than one cluster may be enrolled in a single village but clusters within the same village will need to be separated from each other by a minimum of 1km distance between the two closest households. Each of our water sanitation and hygiene interventions have both a hardware component (and our nutritional intervention requires ongoing provision of nutrient supplements) as well as a software, communication component. Neighboring clusters will not receive the hardware or supplies. Moreover, water sanitation, and hygiene interventions are generally plagued with low levels of uptake and regular use. Thus we do not anticipate spillover of the intervention into neighboring clusters. Since there will be at least 1 km distance, equivalent to 15-20 minutes walking, that separates the closest point of intersection between intervention areas we do not expect direct effects of the intervention to spill over into the closest cluster. In addition, each cluster will have its own community hygiene promoter, and so we will not be asking the same hygiene promoter to deliver different messages to different households. We will have GPS location information from each participating household and as part of the analysis evaluate whether effects of uptake for impact is affected by proximity to other interventions. Although we do not expect to see spillover effects, if we do will adjust for them in the analysis.

After 8 clusters have been enrolled in a particular geographical area, these 8 compound identifications will be sent to Dr. Ben Arnold at UC Berkeley, who will block randomize and assign each cluster to receive one of the 6 intervention or to the double sized control arm.

Field staff will enroll and randomize compounds into the intervention trial over a 10-month period. The enrollment will be stopped for two months after the first 80 clusters. The implementation team will follow and roll out the interventions. This time will allow the team to improve and learn from the experience so that the next rounds are more efficient.

Children born into existing study compounds, other than those born to pregnant mothers enrolled at baseline, will also receive the same intervention as the child in the compound who was originally enrolled. This will be essential to maintain a coherent and consistent intervention within each compound. Since LNS is a more targeted (and expensive) intervention, we plan to limit LNS provision to additional children born to the same mother as target children. For children who are born into existing study compounds, we will attempt to enroll them at midline and endline and if enrolled we will measure diarrhea, anthropometry (length, weight, head circumference) and child development outcomes (when possible: children would need to be > 4 months old to administer most tests). Due to cost and logistics, we do not plan to measure environmental enteropathy biomarkers or parasitic infections in the additional enrollees. We do not plan to include the additional newborn children in our primary analysis because the exposure to intervention for this subset will be shorter than for our index children, and their inclusion would increase the variability of intervention exposure length in our population (and would make defining what constitutes the “intervention” more difficult). However, children born into existing compounds may provide additional information about the impact of our interventions on very young children who are born into cleaner environments. For the spillover substudy, we will enroll children aged 0-5 years in neighboring compounds in the combined intervention and control arms at endline.

To assess the impact of WASH Benefits interventions on parasite infection among school-aged children, we will enroll one child aged 1-4 years besides the target child and one additional child aged 5-12 years in each study compound during endline. These children will be enrolled by a field team conducting the parasites assessment who visits the compound after the completion of the endline survey. A parent of each child will be asked to give parent permission for stool and blood collection. The parent will also be asked to give parent permission for the target child.

For the spillover substudy, we will enroll children aged 0-5 years in neighboring compounds in the combined intervention and control arms at endline. Compounds will be eligible to participate in the spillover substudy if they are located in close proximity to enrolled compounds in the combined intervention and control arms and if a child 0-59 months resides there at endline. Specifically, we will use GPS information to pre-define a perimeter around the study clusters in which compounds will be eligible to enroll in the spillover substudy. This perimeter will ensure sufficient distance remains between study clusters since our aim is to measure within-cluster spillovers rather than between-cluster spillovers. The diameter of the perimeter will be defined through piloting. We will use satellite imagery from Google Earth to identify potential compounds that will be eligible for the spillover study. These compounds will be located within 160 meters of compounds enrolled in the main study and within the perimeter described above. Because satellite images may be outdated, during endline data collection, field staff will ground truth the identified compounds upon arriving in the cluster using GPS devices. At that time, they will also determine which compounds within 160 meters of main study compounds have children 0-5 years. Compounds with children 0-5 years will be eligible for the spillover study and will be invited to participate. They will determine which study compounds have children under 0-5 years by asking neighboring compounds if any 0-5 year old children live there using the recruitment script in the attachments section.

**b) Describe any recruitment materials (e.g., letters, flyers, advertisements [note type of media/where posted], scripts for verbal recruitment, etc.) and letter of permission/cooperation from institutions, agencies or organizations where off-site subject recruitment will take place (e.g., another UC campus, clinic, school district). Attach these documents in Attachments section. Please see eProtocol Attachments Check List for Non-Exempt Applications for more information.**

Recruitment is integrated in to the consent process. Except for the spillover study and the parasites endline recruitment, there are no separate materials just for recruitment. Recruitment scripts for the spillover study and the parasite endline sample are attached in Section 22. Trained field staff will approach the eligible household and introduce themselves and describe the study and the participant involvement should they choose to enroll. The consent documents are attached.

**c) Will anyone who will be recruiting or enrolling human subjects for this research receive compensation for each subject enrolled into this protocol? If yes, please identify the individual(s) and the amount of payment (per subject and total).**

No

#### 7. Screening

**a) Provide criteria for subject inclusion and exclusion. If any inclusion/exclusion criteria are based on gender, race, or ethnicity, explain rationale for restrictions.**

---

**Protocol Title:** Measuring the benefits of sanitation, water quality, handwashing and nutrition interventions for improving health and development in rural Bangladesh

**Protocol Status:** APPROVED

**Date Submitted:** 10/26/2017

**Approval Period:** 10/30/2017-11/03/2018

**Important Note:** **This Print View may not reflect all comments and contingencies for approval. Please check the comments section of the online protocol. Questions that appear to not have been answered may not have been required for this submission. Please see the system application for more details.**

The study will be conducted in communities in 5 districts in rural Bangladesh. These communities must meet all of the following criteria:
•Have no on-going, externally-funded projects implementing or promoting water, sanitation and hygiene technologies or behaviors
•Have no on-going, externally-funded projects implementing or promoting nutritional supplementation or promotion of specific foods based on their micronutrient content
•Have water with an iron concentration of <3milligrams/L on average
•Have water with an arsenic concentration of <50 micrograms/L on average

Compounds (within eligible communities) will be eligible to participate if they include at least one pregnant woman currently living in the compound.  Within each enrolled compound, we will collect information from three types of children:

(1) Infants (target child) will be eligible to participate in the study if they are:
1. They were in utero at the baseline survey
2. Their parents/guardians are planning to stay in the study village for the next 24 months (if a mother is planning to give birth at her natal home and then return, she will still be a candidate for enrollment)

(2) Children < 36 months at baseline that are living in the compound of a target child will be eligible to participate in diarrhea measurement if:
1. They are 3 – 36 months old at the baseline survey
2. Their parents/guardians are planning to stay in the study village for the next 12 months

(3) In addition to the target child, up to two older siblings or older children from each enrolled bari that includes a target child will be eligible to participate in the intestinal parasite specimen measurement if:
1. They are between the ages of 18-27 months at baseline (parasite assessment in these children will be done at both baseline and endline)
2. They will be between the ages of 5-12 years at endline (parasite assessment in these children will only be done at endline)
3. Their parents/guardians are planning to stay in the study village for the next 12 months after baseline enrollment

There will be no exclusion criteria based on gender, race, or ethnicity.

Compounds will be eligible to participate in the spillover substudy if they are located in close proximity to enrolled compounds in the combined intervention and control arms and if a child 0-59 months resides there at endline. Specifically, we will use GPS information to pre-define a perimeter around the study clusters in which compounds will be eligible to enroll in the spillover substudy. This perimeter will ensure sufficient distance remains between study clusters since our aim is to measure within-cluster spillovers rather than between-cluster spillovers. The diameter of the perimeter will be defined through piloting. We will use satellite imagery from Google Earth to identify potential compounds that will be eligible for the spillover study. These compounds will be located within 160 meters of compounds enrolled in the main study and within the perimeter described above. Because satellite images may be outdated, field staff will ground truth the identified compounds upon arriving in the cluster using GPS devices. At that time, they will also determine which compounds within 160 meters of main study compounds have children 1-5 years. These compounds will be eligible for the spillover study.

**b) If prospective subjects will be screened via tests, interviews, etc., prior to entry into the "main" study, explain how, where, when, and by whom screening will be done. NOTE: Consent must be obtained for screening procedures as well as "main" study procedures. As appropriate, either: 1) create a separate "Screening Consent Form;" or 2) include screening information within the consent form for the main study.**

Field staff will assess child and household eligibility by asking caregivers the age of their children (or the approximate due date for pregnant women) when they visit the household to potentially enroll them.

**8. Compensation and Costs**

**a)**

Describe plan for compensation of subjects. If no compensation will be provided, this should be stated. If subjects will be compensated for their participation, explain in detail about the amount and methods/ terms of payment.

Include any provisions for partial payment if subject withdraws before study is complete.

When subjects are required to provide Social Security Number in order to be paid, this data must be collected separately from consent documentation. If applicable, describe security measures that will be used to protect subject confidentiality.

**If non-monetary compensation (e.g., course credit, services) will be offered, explain how**

No monetary compensation will be given to subjects for their participation. Participants in treatment arms will receive free sanitation hardware, handwashing hardware as part of the study, which they will retain after the completion of the study. Participants with target children in the household will additionally receive free nutritional supplements (LNS), detergent for making liquid soap and a regular supply of waterless hand sanitizer and water treatment supplies for the duration of their participation in the study.

**b) Discuss reasoning behind amount/method/terms of compensation, including appropriateness of compensation for the study population and avoiding undue influence to participate.**

In rural villages in Bangladesh, the standard wage is $1.50 per day. Thus, even modest compensation risks being coercive. ICDDRB's practice, consistent with the practice of other research organizations in Bangladesh, is to provide modest tangible benefits to participants in studies (for example the water treatment supplies included in some arms of this project) and then provide potential study participants with the clear option to participate or not.

---

**Protocol Title:** Measuring the benefits of sanitation, water quality, handwashing and nutrition interventions for improving health and development in rural Bangladesh

**Protocol Status:** APPROVED

**Date Submitted:** 10/26/2017

**Approval Period:** 10/30/2017-11/03/2018

**Important Note:** This Print View may not reflect all comments and contingencies for approval. Please check the comments section of the online protocol. Questions that appear to not have been answered may not have been required for this submission. Please see the system application for more details.

**c) Costs to Subjects. If applicable, describe any costs/charges which subjects or their insurance carriers will be expected to pay. (If there are no costs to subjects or their insurers, this should be stated.)**

Participation in the study will not result in any direct costs to subjects or their insurers, other than cases in which subjects elect to devote their own time to improving their water quality, sanitation or hygiene practices, as encouraged and facilitated by the interventions.

---

---

**Protocol Title:** Measuring the benefits of sanitation, water quality, handwashing and nutrition interventions for improving health and development in rural Bangladesh

**Protocol Status:** APPROVED

**Date Submitted:** 10/26/2017

**Approval Period:** 10/30/2017-11/03/2018

**Important Note:** This Print View may not reflect all comments and contingencies for approval. Please check the comments section of the online protocol. Questions that appear to not have been answered may not have been required for this submission. Please see the system application for more details.

#### * * * Study Procedures, Alternatives to Participation * * *

##### 9. Study Procedures

**a) Describe in chronological order of events how the research will be conducted, providing information about all study procedures (e.g., all interventions/interactions with subjects, data collection procedures etc.), including follow-up procedures. If any interviews, questionnaires, surveys, or focus groups will be conducted for the study, explain and attach one copy each of all study instruments (standard and/or non-standard) in the Attachments section. Please see eProtocol Attachments Check List for Non-Exempt Applications for more information. If the proposed research involves use of existing data/specimens, describe how data/specimens will be acquired.**

Please see Figure 1 (attached) for the overall chronology of the study. The interventions will require about 3 months from the baseline to implementation. The follow-up rounds are planned for 12 and 24 months after intervention delivery. The major activities involved in the study include:

1)Promoter Selection and Training
2)Baseline Assessment
3)Intervention Implementation
4)Midline Assessment
5)Endline Assessment

1) Promoter Selection and Training
The fieldwork will be implemented by local field workers recruited and supervised by the Bangladesh-based scientific team. For all study arms, local promoters will be nominated by community members participating in the study. Following the baseline survey icddr,b field staff will search for potential candidates by interviewing community members within each cluster. During interviews, they will describe promoter eligibility criteria and explain the role of the promoter in the community. Field staff will seek nominations from at least one member of each target household and at least two community leaders (includes school teachers, religious leaders, village doctors, NGO workers, and others) for a few candidates to work as a promoter in their community. Based on the nominations from the community, field staff will tally marks against each candidate on a checklist. Staff will identify the 3 nominated candidates with the highest scores and then they will visit them to inform the eligibility criteria, promoter roles and responsibilities, and benefits of being a promoter. Based on the discussion, if the candidate shows interest in being a promoter then field staff will proceed in collecting detailed socio-demographic information otherwise they will stop the interview and proceed to the next candidate. Following this, field staff with the support of other team members will analyze the data to identify eligible candidates for interviews. After randomization, the team will invite at least 3 preliminary eligible candidates from each intervention cluster for interview (both written and viva). Based on the performance of the interview one will be chosen as the promoter while other will be in the waiting list as a back-up promoter in case the selected promoter is unable to satisfactorily carry out his/her responsibilities or drop out due to any reasons.

All selected promoters will be invited for 2 -day basic training conducted by icddr,b staff which includes research and training team on interpersonal communication, introduction to behavior change communication strategies, basic adult learning theory, time management/planning and reporting producers. Upon successful completion of the initial training, and based on the trainer’s assessments of producers. Upon successful completion of the initial training, and based on the trainer's assessments of the two nominees from each village, one will be chosen as the promoter. The selected promoter of individual arm will subsequently attend a two-day training specific to the intervention they will be promoting and, if the cluster is randomized to the combined arm, an additional 4 days of training. For example, promoters in nutrition arms will address the importance of good nutrition for child development, the basics of a healthy diet for children in the target age range, the LNS product and nutrition specific health education modules that have been specifically developed for this project. Field staff along with the training team will assess the performance (knowledge, skill and attitude) of promoter in each quarter by using performance assessment tools and will do the grading according to their performance. Based on the assessment there will be refresher trainings at every quarter that will last for one to two days and present a review of general themes as well as any new behavior change communication strategies that have been developed. The refresher training will serve as a venue to share ideas, lessons learned and best practices among the promoters and their supervisors.

Shortly after the training workshops, community meetings will be conducted to introduce the intervention and to present the promoter and describe their role to the community and mothers participating in the study by a representative from icddr,b. The study representative will then accompany the promoter on his/her first 2-3 participant interactions (lasting ~1 hr each) and provide the promoter with feedback on his/her techniques. The promoter supervisors will each oversee ~12 promoters for the duration of the study. They will stay in touch with promoters through monthly meeting, monthly phone calls and site visits in each month. In order to ensure that study households are adequately supported, promoters will be asked to make frequent contact with study households in the first days and weeks after the intervention is launched, with interactions then tapering off after the first 6 months of the study to a long-term pattern of monthly visits by promoters to deliver LNS, Aquatabs, hand sanitizers and technical support on repairs as required in addition to provision of ongoing behavior change support, and check on uptake. Promoters will be compensated for their efforts at a rate that is commensurate with the government's pay to community health workers, with a strong emphasis on the prestige of being selected as a promoter (actualized in the form of a diploma from the training, a household visit kit, and study identification badge and potentially monthly top-up for promoters cell phone to facilitate communication between research staff and promoters).

2) Baseline Assessment (age -7 to 0 mo.)

After a pregnant mother has been enrolled in the study, trained ICDDRB staff will conduct a baseline assessment. Mothers will be asked standard questions about their (and spousal) education, activities, occupation, household assets, and current sanitation and hygiene practices. The baseline assessment in all participating households will include bar soap and detergent powder consumption measurements, latrine use (visual inspection), and numerous spot-check hygiene indicators (e.g. presence of animal or human feces in the household environment). The field team will collect these measurements at the three measurement rounds of the study (baseline, 1-year follow-up, 2-year follow-up). We will ask mothers to provide a 10 mL sample of blood for future genetic testing and testing of nutrition and other biochemical parameters, a urine sample for chemistry and a stool sample to permit future testing of the intestinal microbiome. For the blood sample we will separately freeze aliquots of sera for eventual biochemical analysis, and retain a clot for genetic testing. Although we are not currently funded to analyze these samples, we envision future interest in assessing the impact and controlling in the analysis for maternal nutritional deficiencies or other biochemical factors on child development, and so we will collect and archive this prenatal sample. We are requesting consent to hold these samples for up to 20 years because we want to take advantage of expected future advances in characterizing immunological and other parameters relevant to the pathophysiology of environmental enteropathy. We have not specified the tests, because we do not know which relevant tests will become available, though the field is making substantial advances each year, and so we expect that highly relevant tests will become available.

Fly Density
In the random subset of 720 households within each group we will assess environmental contamination at baseline by measuring fly density. Field workers will start by locating the nearest latrine, food preparation area (usually rural outdoor kitchens) and the garbage disposal site for the target household. To capture flies, they will use Revenge fly tapes from Roxide inc. We chose these passive sticky tapes because of their adhesion ability. They will cut out three 1.5 ft of these tapes and hang them parallel to each other near the sites. The field staff will set these traps between 9-10 am,ask the household not to disturb the tapes, and collect them after 24 hours. Trained field workers will count the number of flies in each trap and speciate them using a simple visual identification chart made from The Fauna of British India series (Aubertin and Smart 1940; Van Emden 1965; Nandi 2002).

Parasite Assessment (compound residents aged 18 to 27 months)
Since at enrollment the target children will not yet be born, and even those who are born are at low risk of parasitic infection, we will enroll older children age 18 to 27 months who live in the same compound as the target child as a proxy to assess the risk for parasitic infection in our target communities. These children will be the same age at baseline as the target will be at endline. Stool and blood spot samples will be collected from these children. Stool collection will require two visits to each household. On day 1, the field team conducting the survey will deliver to each caregiver a stool collection kit and instruct them how to collect stool from their children. Caregivers will be instructed to have their child defecate on a sheet of provided plastic, to use a provided plastic scoop (integrated into a storage container) to collect ~10 mL of fresh stool from the top of the pile. On day 2, field staff will return to the household to collect the stool sample. Field staff will aliquot fresh stool specimens for parasite microscopy at endline (see endline assessment section for details). We will maintain a cold chain of 4°C until the samples are transported to the field lab (<6 hours) where they will be stored at –20°C until shipment on dry ice to ICDDRB where they will be stored at final temperature of -80°C.

Paired with each stool sample we collect we will also collect a finger prick blood sample. One of the child's fingers will be cleaned using the disinfectant liquid and after drying completely prick to adapt to 0.25 mm using a spring-loaded disposable handset (BD Microtainer®). Six drops of blood (about 60 μl) will be collected using a filter disk. We will store the filter disk samples at 4°C. The filter disk will then be frozen and transported to the ICDDR,B lab, where they will be stored at –80°C. The eventual goal of the filter disk blood specimen collection will be to analyze them for intestinal helminthes, protozoans, and other pathogenic organisms using antigen-based assays (Luminex), but this protocol does not include the filter disk analysis activities. At endline, we will measure the study child’s hemoglobin concentration using Hemocue analyzers (Hemocue 301).  Results will be provided to the household in the field.

---

Subsample Environmental Enteropathy (EE) Assessment (6 months after baseline, midline at 15 months and endline at 27 months after baseline)
We will randomly select 80 clusters from the control group, 80 clusters from the combined water, sanitation, hygiene (WASH) group, 80 clusters from the nutrition group, and 54 clusters from the nutrition plus WASH group for markers of environmental enteropathy. This sample collection will be implemented over the course of 2 days per village at the household level. We plan to collect samples from 1500 infants at the first survey round, which will take place approximately 6 months after baseline (however, 1500 infants is an optimistic number as a significant fraction of mothers may be living in their parents' village during the first few months of their child's life or a fraction of infants will still be in utero and will thus be unavailable for sample collection). However, the entire sample of 1,500 will likely not be be present for the 1-year (midline) and 2-year (endline) assessments. Thus, at the midline and endline assessments, we will need to increase the sample size to 2,000 children. Each round of assessment will include the urine, serum, and stool collection described below. Furthermore, at the midline and endline assessments, we will collect blood, urine, saliva, hair, and stool specimens from a subsample of 2,000 children in the study to measure biomarkers for environmental enteropathy, including interleukin 6, interleukin 1-beta, F2-isoprostanes, and other biomarkers, allostatic load, and telomere length. Additionally, we will collect blood, saliva, hair, and urine from the 2,000 mothers of the children in the EE subsample to validate these newly discovered candidate environmental enteropathy markers and correlate them with the child results.

On the first day, consent, anthropometric measurements, blood pressure, heart rate, autonomic function, and skin conductance (sweat) measurements, provision of stool collection materials, and saliva, hair, and blood collection take place. On the second day we will collect urine and additional saliva samples from the study children and their mothers, pick up the stool from the study children, interview the mothers about their diets as well as their infant's diet during the past week and the previous 24 hours. On a non-consecutive day, a subsample of mothers (n=60) will be revisited to be interviewed about their diet as a quality control measure.

Saliva sampling:

Saliva and hair sampling, autonomic function testing:

On day 1, the field team will collect a total of 3 saliva specimens each from mothers and their children (before the blood draw, immediately following the blood draw, and at a later time point after the blood draw) using the previously approved Salimetrics swab. They will place the Salimetrics swab under the child's tongue for 2-3 minutes and store the swab in a tube. These Salimetrics samples will be used to measure salivary cortisol reactivity and cytokine response before, during, and after an acute minor stressor (the blood draw). A hair sample consisting of 3-4 strands of hair from each mother and child will also be taken with stainless steel scissors. Three to four strands of hair will be cut from three or four locations on the head. These hair samples will be used to measure hair cortisol and long term stress. To measure allostatic load, blood pressure measurements will be taken using a standard sphygmomanometer, heart rate and autonomic function will be measured using a portable electrocardiogram (Biopac ECG), and skin conductance (changes in sweat response, a test of the autonomic nervous system) will be measured using a galvanic skin response meter (MindWare GSR). The ECG and GSR will be attached before, during and after the acute minor stressor (the blood draw), to correlate autonomic function with cortisol activity. The mother will receive the blood pressure and heart rate results for herself and her child. The saliva and hair

---

**Protocol Title:** Measuring the benefits of sanitation, water quality, handwashing and nutrition interventions for improving health and development in rural Bangladesh

**Protocol Status:** APPROVED

**Date Submitted:** 10/26/2017

**Approval Period:** 10/30/2017-11/03/2018

**Important Note:** **This Print View may not reflect all comments and contingencies for approval. Please check the comments section of the online protocol. Questions that appear to not have been answered may not have been required for this submission. Please see the system application for more details.**

samples collected from day 1 will be used to measure cortisol reactivity, cortisol recovery, and cumulative cortisol levels using a commercial ELISA kit. To measure temporal variability of the EE biomarkers, on day 2 in the morning, the staff will collect an additional saliva sample from the child and the mother using the Oragene kit which consists of a soft sponge and tube. On day 2 in the morning, the staff will collect an additional saliva sample from the child and the mother using the Oragene kit which consists of a soft sponge and tube. Telomere length will be measured in saliva samples collected from the children and mothers on day 2. On day 2 in the afternoon, field workers will collect samples using the Oracal sampling workers will collect samples using the Oracol sampling tube. Specifically they will 1) Remove the sampler being careful to touch only the handle. 2) Holding the handle as a tooth brush. Insert it in child's mouth and the rub sponge against the gums for one to two minutes, ensuring that the sponge is completely wet/saturated. 3) Without touching the sponge, insert the sampler into the original Oracol tube, sponge down 4) Close the lid tightly 5) Put the Oracol sampler tube into a storage bag for transfer to a cooler. The Oracol device is used to collect crevicular fluid to study pathogen-specific antibodies. These Oracol samples will be assessed using new assays to measure pathogen specific IgA and IgG antibodies and compared to serological markers.

To measure telomere lengths, DNA will be extracted by using a commercial kit (QIAamp. We will use a validated published method to measure relative telomere lengths by quantitative PCR (Cawthon, 2002). Briefly, this method determines relative telomere lengths by measuring the factor by which each DNA sample differs from a reference DNA sample in its ratio of telomere repeat copy number (T) to single copy gene copy number (S) (Cawthon, 2002). The T/S ratio is proportional to the average telomere length. To convert the T/S ratio into base pairs, we will use a formula derived from the mean telomeric restriction fragment length from Southern blot analysis and the slope of the plot of mean telomeric restriction fragment length versus T/S (Entringer, 2011).

Urine specimen collection and analysis: From a subset of every fifth cluster (n=144), urine will be collected from the pregnant mother for micronutrient and iodine testing. These will be maintained at 4 degrees C at the field level and stored at -20 degree C at the field office.

Our field teams will collect urine samples from all eligible children and their mothers in a study cluster (up to 7 children) in one day per cluster. The field team will request the mother to collect a sample from her first urine of the day. Additionally, the field team will request that mothers not feed their children for at least one hour before they receive the lactulose-mannitol solution. The children will be weighed and measured using the same anthropometric procedures as described above. The lactulose-mannitol solution will be prepared at the ICDDR,B nutritional biochemistry lab using lactulose syrup and mannitol powder secured from international pharmaceutical suppliers. The lactulose-mannitol solution will be mixed with sterile water to produce a solution with a concentration of 250 mg of lactulose and 50 mg of mannitol per milliliter. The lactulose-mannitol assay requires the collection of an additional pre-LM urine sample to serve as a "control or baseline" urine for comparison with the post-LM urine. A pre-LM urine sample is a sample of urine (12 ml) that is collected during the 1-hour fasting period preceding the administration of lactulose-mannitol solution to the child. This additional pre-LM urine sample does not change the amount of time we are present in the household. For assay standardization and QA/QC purposes, we also plan to spike these pre-LM urine samples with fixed concentrations of lactulose, mannitol, or a known interfering compound during analysis.

Field workers will administer 2 ml of the solution per kilogram of body weight of the child. A urine collection bag equipped with a drainage tube will be attached to the infant immediately after dosing. Thirty minutes

---

**Protocol Title:** Measuring the benefits of sanitation, water quality, handwashing and nutrition interventions for improving health and development in rural Bangladesh

**Protocol Status:** APPROVED

**Date Submitted:** 10/26/2017

**Approval Period:** **10/30/2017-11/03/2018**

**Important Note:** **This Print View may not reflect all comments and contingencies for approval. Please check the comments section of the online protocol. Questions that appear to not have been answered may not have been required for this submission. Please see the system application for more details.**

after the infant consumes the sugar solution, mothers will be encouraged to breastfeed infants <6 months or offer water to their children >= 6 months to help their urination. Children over 6 months will be given purified drinking water 30 minutes after taking the sugar to help urination. Whenever the child urinates, the urine will be removed from the bag and placed in a container with 0.1% thimerosal (1 drop per 5 ml), a preservative. The total volume of urine collected after 5 hours will be noted, a 12ml well mixed sample will be stored at -80 degrees C, and the urine bag will be removed from the child.

Since the mannitol/lactulose concentration measurements necessitate the use of high performance liquid chromatography and mass spectrometry (LC-MS/MS), which is presently unavailable at icddr,b, the child urine samples will be shipped to the US and also analyzed in Bangladesh, where the LC-MS/MS labs are located. We plan to collaborate with Dr. Mohammad Alauddin and his team at Wagner College to analyze these urine samples. Dr. Alauddin has a mass spec machine at his lab in Wagner College that will need to be calibrated and standardized with the Pre-LM urine samples before he ships the mass spec machine to his lab in Dhaka, where the remaining urine samples will be analyzed. Oxidative stress and hypothalamic-pituitary-adrenal axis markers will also be measured in the urine samples from the mother and child: F2-isoprostanes will be analyzed using gas chromatography (GC)/negative ion chemical ionization (NICI) mass spectrometry (MS) (Morrow and Roberts 2002), 8-hydroxy-2'-deoxyguanosine (8-OHdG) will be measured in using high-performance liquid chromatography-tandem mass spectrometry (HPLC-MS/MS) (Weimann, Belling et al. 2002), and catecholamines (epinephrine, norepinephrine, and dopamine) will be measured using a commercial ELISA kit (Parks, Miller et al. 2009).

The urine samples will be analyzed for concentrations of mannitol and lactulose using high performance ion chromatography at the ICDDRB laboratory (Barboza 1999).

Venous blood specimen collection and analysis: Before the urine specimens are collected, trained phlebotomists will collect up to 7.7 ml of venous blood from each child (< 2.5% of total blood volume for infants > 2 kg). and 10 ml of venous blood from each mother. We plan to collect an additional 2.7 mL of blood for serum micronutrient biomarker analysis during endline and we believe that this additional blood volume are not physiologically significant since the children are around two years old during endline. Retinol binding protein, transferrin receptor, ferritin, hepcidin, folate and B12, c-reactive protein, and alpha-1 acid glycoprotein will be measured in these serum samples. The additional sample collection tube is required due to the fact that a number of these assays do not perform well in samples collected in trace element-free plasma collection tubes and to ensure that there is sufficient sample volume to meet the assay requirements. With this added tube, the total blood volume to be collected from children at endline will be 7.7 mL. Blood samples will be centrifuged within three hours of collection to separate the plasma and serum from the red blood cells. The plasma and serum will then be stored at –80°C. Commercially available ELISA kits will be used to measure total IgG, IgG endotoxin core antibodies (Coaset EndoCAb) and C-reactive protein, alpha-1 acid glycoprotein (AGP), interleukin 6 (IL-6), interleukin 1 (IL-1), tumor necrosis factor (TNF), insulin-like growth factor 1 (IGF-1), and other environmental enteropathy biomarkers. Maternal blood samples will be assayed for inflammation and stress biomarkers. Blood spots will be collected for luminex testing of antibodies to intestinal parasites. At least 50 microliters of each sample will be reserved for nutritional markers (iron, vitamin A, B12, folate) and the rest of the sample will be frozen to allow for the analysis of infection with enteric pathogens and environmental enteropathy biomarkers. Aliquots of the blood samples will be shipped to Mark Davis's lab at Stanford University (USA) for immunological analyses using CyTOF, a Time-of-Flight mass spectrometer to measure highly multi-parametric single cell data including cytokine panels and peripheral blood phenotyping. Aliquots of the blood will also be shipped to Hohenheim University (Germany) for micronutrient and acute phase protein panel analyses in Juergen Erhardt's lab.

Stool specimen collection: The field team will collect stool samples from all eligible children in the study cluster. Stool collection will require two visits to each household. On day 1, the field team will deliver to each caregiver a stool collection kit and instruct them how to collect stool from their children. Caregivers will be instructed to collect stool from their children on the following morning in the event that the child defecates before they report to a central location in the cluster for urine and serum sample collection. Caregivers will be instructed to have their child defecate in a plastic, non-absorbent diaper and to use a provided plastic scoop (integrated into a storage container) to collect ~10 mL of fresh stool from the top of the pile. On day 2, field staff will visit the households for the urine sample collections in each cluster (details above). Field staff will aliquot stool specimens (either collected by the caregiver in the early morning, or collected by field staff on Day 2) into 5 cryovials and maintain a cold chain of -20°C until the samples are transported to ICDDR,B (<6 hrs) where they will be stored at –80°C until they are analyzed using qPCR to identify Entamoeba histolytica, Giardia and Cryptosporidium. The remaining 4 aliquots will be retained for testing stool markers of environmental enteropathy and to assess the fecal microbiota and microbiome. At endline, the remaining fresh stool will be used to measure soil transmitted helminthes (as outlined in the endline parasite assessment).

Quality Assurance / Quality Control: We will include biological and technical replicates to ensure data validity. Aliquots from the same biological or environmental sample will be analyzed separately and compared. We will include negative controls daily for water testing. For example, the ELISA test can be performed twice on two separate days for the same sample. We will set aside an aliquot of each batch of L/M solution for further testing in case there are any batch inconsistencies. Multiple field research assistants can record anthropometric, blood pressure, and heart rate measurements for the same child to measure human error. Two percent of questionnaire assessments will be repeated by the supervisor within 7 days of data collection

A list of the modules/instruments that will be utilized at baseline include:

Module 2. Diarrhea and illness symptoms
Module 7. Handwashing assessment
Module 8. Sanitation assessment
Module 9. Child defecation and feces disposal assessment
Module 10. Water treatment, storage, and quality assessment
Module 14. Environmental enteropathy assessment (subset of study population)
Module 15. Intestinal parasites assessment (for older siblings of target children)
Module 18. Quantitative fly assessment (subset of households)
Module 20. Behavioral determinants
Module 22. Household food insecurity

3) Intervention Implementation

The interventions arms to be implemented include:

---

**Protocol Title:** Measuring the benefits of sanitation, water quality, handwashing and nutrition interventions for improving health and development in rural Bangladesh

**Protocol Status:** APPROVED

**Date Submitted:** 10/26/2017

**Approval Period:** 10/30/2017-11/03/2018

**Important Note:** This Print View may not reflect all comments and contingencies for approval. Please check the comments section of the online protocol.
Questions that appear to not have been answered may not have been required for this submission. Please see the system application for more details.

1) Improved water quality: chorine tablets (Aquatabs) + 5 liter safe storage vessels, water treatment promotion

2) Improved sanitation: sanitation promotion, child potties, sani-scoop hoes to remove feces from household environments, latrine upgrades to dual pit latrines

3) Improved handwashing: promotion of handwashing with soap or waterless hand sanitizers at critical times, handwashing stations, soapy water at handwashing locations. Specifically, for the kitchen a 16 liter bucket with tap fitting, stool, bowl and soapy water bottle will be provided. For the latrine, a 40 liter bucket with tap fitting, stool, bowl and soapy water bottle will be provided.

4) Combined water + sanitation + handwashing: this combined arm includes all interventions described above in #1, 2, & 3, with phased implementation

5)Nutrition supplementation: the nutrition supplement we will use provides a combination of energy and micronutrients delivered in 10 gram sachets (produced by Nutriset), to be mixed into existing meals (i.e. porridge) two times per day for target children age 6 mo. to 24 months in age. The supplement provides 108 kcal/day and includes a broad suite of essential fatty acids and micronutrients. The nutrition supplement is meant to supplement breastfeeding and locally available complementary foods. Additional messages about breast feeding and the consumption of micronutrient-rich complementary foods modeled on those recommended in the Guiding Principles for Complementary Feeding of the Breastfed Child [Dewey 2003] and the recent UNICEF Program Guide for Infant and Young Child Feeding Practices [Unicef 2011]

6) Nutrition supplementation + water + sanitation + handwashing: this arm will include all of the sub components described above, with phased implementation

The behavior change strategy and communication plan is attached in a separate document.

Two additional arms will be the control group.

Upon completion of the baseline data collection in 8 clusters, the research investigator overseeing site selection (Sania Ashraf) will compile the list of 8 clusters with their cluster IDs drawn from the database and listed in the order that the baseline data collection aws completed. An external research investigator (Swapon Biswas) will email the cluster IDs to a second offsite co-investigator (Ben Arnold). The offsite co-investigator will use prespecified statistical code to generate blocks of 8 assignments in random order specifying each of the 6 potential interventions (improved water quality; improved sanitation; improved handwashing; combined water, handwashing and sanitation; nutrition; nutrition plus improved hand washing; combined water, handwashing and sanitation) plus two assignments to the control group. He will return a password protected WASH B Bangladesh Treatment Assignment spreadsheet to Mahbubur Rahman, senior program manager for this study at icddr,b. It will be stored on a secure server that is backed up regularly. He and the four investigators in charge of delivering the interventions will be the only people who have access to the file to prevent unblinding the primary investigators and primary data analysts for the trial.

We will hold separate training sessions for community promoters who are delivering different interventions. Thus, when we convene a training session for community promoters who will be implementing the water

---

**Protocol Title:** Measuring the benefits of sanitation, water quality, handwashing and nutrition interventions for improving health and development in rural Bangladesh

**Protocol Status:** APPROVED

**Date Submitted:** 10/26/2017

**Approval Period:** 10/30/2017-11/03/2018

**Important Note:** **This Print View may not reflect all comments and contingencies for approval. Please check the comments section of the online protocol. Questions that appear to not have been answered may not have been required for this submission. Please see the system application for more details.**

quality intervention, there will be no training on handwashing promotion. This will reduce the risk of spillover of intervention from one intervention group to another. However, it also requires that enough clusters be enrolled and community promoters identified to be able to convene an intervention specific training.

In order to accumulate sufficient intervention-specific promoters for training there will be a period of up to 3 months between the baseline assessment and the intervention implementation. Field staff will return to the intervention compounds after the baseline assessment and randomization into intervention arms. ICDDRB staff and the community hygiene promoters will distribute handwashing stations, potties, sani scoops, provide Aquatabs and safe water storage containers, and/or LNS packets, depending on the study arm. The intervention team will work in collaboration with the Village Education Resources Center (VERC), who has considerable experience installing dual pit latrines in rural communities in Bangladesh, to install dual pit latrines in eligible compounds as determined from baseline data. Field staff will discuss the process of intervention implementation and messaging with study participants in each community as the interventions are being rolled out.

At the commencement of intervention roll out, community meetings will be held including community leaders to explain the study and the interventions. A trained, local female health promoter will deliver the behavior change messaging and will promote intervention use. The hygiene promoter will work with authority figures in the community to communicate messages depending on the assigned intervention, for example that child feces should be disposed in a latrine, that it is the occasional unseen contaminant in both water and on hands that needs to be protected against, and that key times to wash hands with soap include after defecation, after cleaning a child who has defecated, before preparing food and before eating or feeding their infants. Promoters will engage in a conversation with participants. They will observe household and compound conditions and personalize placement and use of enabling technologies, listen to people's concerns work with them to solve problems, respond to questions, encourage household members to use the enabling hardware and products in their presence to ensure understanding of use, and encourage and congratulate the adopters. Intervention promoters will also collect a subset of the indicators (LNS sachet consumption, hardware use indicators) on a monthly basis.

The nutrition intervention will be implemented in two study arms. During the first 6 months of life, promoters will encourage mothers to exclusively breast feed their children. When children turn 6 months of age and are starting to eat solid foods, community promoters will instruct mothers/guardians to continue breastfeeding along with offering solid food. Promoters will instruct mother/guardians to mix 1 sachet (10 mg) of the supplied nutrition supplement (LNS) and either feed it directly to the child or mix it with rice or other food fed to the child 2 times per day. The mother/guardian will be given a 1-month supply of the supplement at a time. In the event that the specified LNS is not immediately available at the study start-up, we will initiate the trial with Nutributter, an off-the-shelf product that is very similar in nutrient content to our research formulation of LNS (Appendix 8), and is available through a second factory in the United States (Edesia). The local health promoter will deliver the monthly supplies, will be responsible for delivering behavior change communication (BCC) messages encouraging continued breastfeeding, feeding of nutrient rich complementary foods, feeding frequency, and proper use of the supplement. The behavior change communication messages will follow the best practices for complementary feeding interventions specified by WHO and Unicef and utilize the recommendations and practices from the Alive and Thrive program in Bangladesh (Dewey 2003, Unicef 2011).

For all intervention arms, the hygiene promoter will visit the participating household frequently (i.e. 1 visit per week) early in the study; later in the study, the promoter visits will taper off to one visit per month.

If at any visit to an intervention household a community hygiene promoter identifies a serious illness or injury that she believes is related to the intervention (e.g. an injury associated with construction of a new pit latrine) then the hygiene promoter will inform her supervising field research assistant who will record the details and notify Dr. Md. Mahbubur Rahman.

4) In-Depth Environmental Assessment in Sanitation, Combined WASH and Control Arms

In order to assess the impact of sanitation improvements alone and in combination with water and hand hygiene interventions on fecal bacterial commination in the household environment, we will collect environmental samples from all households enrolled in the sanitation arm and the combined water, sanitation, and hygiene arm, and half of the households in the double-sized control arm. This environmental assessment will occur between 2 -10 months post intervention delivery, preceding the 1-year midline assessment. We will measure the level of fecal contamination along five transmission pathways, including water, hand, soil, food, and flies among 2160 households (720 households per arm). Each enrolled household will be visited twice, as discussed below (consent form in Appendix 1q).

On the first visit, we will collect the following samples for microbial analysis: hand rinse from target child, soil from the child's play area, stored drinking water, source drinking water, pond water, food to be served to the target child, and flies captured near the food preparation area. Field microbiologists will collect 250 mL of water samples in sterile Whirlpak bags from the household's tubewell as well as from drinking water storage containers, by asking participants to provide a glass of water that they would give to their child to drink. Index child hand samples will be collected by rinsing the hands of the index child, one at a time, in 200 mL of sterile water in a sterile Whirlpak bag (Pickering et al., 2010). In each study compound, ~100 grams of soil will be excavated using a disposable sterile scoop from approximately 20cm by 20cm area where the index child is currently playing or reported to most recently have played (Pickering et al., 2012). In Bangladesh food is usually prepared in the morning, then fed to children throughout the day. Any previously prepared food being stored in the household for consumption by the index child will be sampled by collecting ~25-50g directly from the storage pot using a sterile spoon, then placed into a sterile plastic bag. Flies will be captured using sticky tape or baited traps placed near the food preparation area at the beginning of the household visit.

All water, hand rinse, food, fly, and soil samples will be placed on ice and transported to the WASH Benefits field laboratory for quantitative analysis for E. coli and fecal coliforms within 8 hours by the Colilert most probable number method (IDEXX) (Eckner, 1998). Lab technicians will analyze water and hand rinse samples directly. They will analyze soil samples by first homogenizing the sample, then suspending and agitating a specified amount (5-20g) of the soil in sterile water for subsequent processing by the IDEXX method. Food samples will be processed by mixing an aliquot of ~25g with 100ml 0.1% peptone water (Islam et al., 2012). Flies (up to 5) will be removed from the traps with a sterile tweezer, placed in a sterile tube with 1ml sterile saline solution, crushed with a sterile pestle, then further diluted with saline solution for processing by IDEXX. Serial dilutions will be prepared as needed for pond, food, fly, and soil samples. One duplicate and one blank of sterile water will be analyzed for every 10th sample.

In addition to IDEXX analysis for E. coli and fecal coliforms, aliquots from all food and fly homogenized samples (mixed with sterile water) will be processed by membrane filtration and subsequently cultered for the presence of shigella and enterobacteriaceae in the field lab. For these same food and fly samples, IDEXX wells that are positive for E. coli post-incubation will be lanced and the contents removed and centrifuged to isolate E. coli cells. These cells will be frozen and transported to the icddr,b laboratory for DNA extraction and molecular analysis by multiplex PCR of the following pathogenic E.coli genes: eae, ial, bfp, ipaH, st, lt, aat, aaiC, stx1, stx2.

An aliquot of each soil sample will also be processed by Kato Katz microscopy for detection of Ascaris, hookworm, and Trichuris ova (Albonico et al., 2012)- the most common soil transmitted helminth infections in Bangladesh affecting children. In addition, aliquots from a subset of hand rinse, soil rinse, diluted food, and water samples (total of ~2000 samples) will be vacuum filtered and archived for subsequent DNA/RNA extraction, molecular fecal source tracking analysis (to differentiate fecal contamination of animal vs. human origin), and detection of two of the most common child diarrheal pathogens in Bangladesh: rotavirus and pathogenic E. coli. The archived filters (n~2000) will be transported to Stanford University for molecular analysis.

During the first household visit for in-depth environmental assessment, we will also administer a brief interview to the primary caregiver of the index child to assess caregiver reported diarrhea for the index child and other children < 60 months living in the compound. This will be followed by a second visit where we will administer the same questionnaire about children's health. The second visit is designed to fall within a plausible incubation period between child exposure to a diarrheal pathogen and illness.

During each of the two household visits, spot checks will be conducted to assess indicators of relevant behaviors (compliance with the interventions), including water treatment, latrine usage, child and animal feces management, hand hygiene, and food hygiene. These spot checks include: latrine features (slab, functional water seal), presence and quantity of human and animal feces in compound, presence and functionality of child feces management tools and potty, presence of soap and water at handwashing stations, presence of visible dirt on caregiver and child's hands, presence of chlorine residual in stored drinking water, drinking water storage container and extraction method, presence of clean cover over stored food, presence of animals in household and animal feces in compound, and presence of flies in food preparation area and near the latrine.

5. Longitudinal Environmental Assessment in Sanitation and Control Arms

To assess the long term impact of the sanitation intervention on microbiological contamination in the household environment, we will longitudinally follow 720 households in the sanitation and control arms (360 households per arm) through quarterly visits over two years, for a total of eight visits per household over the study period. The visits will start approximately 12 months after intervention implementation. In addition, we will collect a one-time soil sample for the detection of soil-transmitted helminth eggs from all households in the sanitation and two control arms (720 households/arm, 2160 households total). This sample will only be collected once – the timing will be synchronized with the collection of the stool samples from these households as part of the endline parasitic assessment.

Data Collection:
At each quarterly sampling round, we will collect samples from households' stored water, mother hands and children's hands for analysis of fecal indicator organisms (E. coli and fecal coliforms). First, concentrations of fecal indicator organisms in environmental samples and on hands have substantial

---

**Protocol Title:** Measuring the benefits of sanitation, water quality, handwashing and nutrition interventions for improving health and development in rural Bangladesh

**Protocol Status:** APPROVED

**Date Submitted:** 10/26/2017

**Approval Period:** 10/30/2017-11/03/2018

**Important Note:** **This Print View may not reflect all comments and contingencies for approval. Please check the comments section of the online protocol. Questions that appear to not have been answered may not have been required for this submission. Please see the system application for more details.**

temporal and spatial variability [Levy 2009, Pickering 2011, Ram 2011], and repeated samples have been recommended to adequately characterize contamination [Boehm 2002, Jensen 2004]. Fecal indicator detection frequency over repeated measurements has also been shown to better predict pathogen presence in tubewells than single samples [Ferguson 2012]. Additionally, we also anticipate temporal trends in fecal contamination of water and hands. Bangladesh has a dry season from November to April and a monsoon season from May to October, during which flooding is common. Seasons have a marked impact on environmental contamination in rural Bangladesh, with heavier contamination during the rainy season [Leber 2011, van Geen 2011]; wet conditions also lead to prolonged pathogen survival in the environment [Santamaria 2003]. Even within a given season we expect variation in the conditions that spread pathogens from feces into the environment; for example, the groundwater table is low during the early monsoon and equalizes with surface waters by the late monsoon [Knappett 2011]. Quarterly sampling will allow nuanced assessment of contamination during different seasons (i.e. early wet, late wet, early dry, late dry).

During two sampling rounds (quarterly visits 2 and 3), we will collect an additional aliquot of stored tubewell water, mothers' and children's hand rinse samples as well as a sample of soil from the household entrance and stored food served to young children from the same 720 households to measure selected pathogens and conduct microbial source tracking. We will analyze these samples with molecular methods for three of the most common diarrheagenic pathogens in Bangladesh (rotavirus, enterotoxigenic E. coli, Shigella) [Black 1981, Albert 1999] and for the protozoa that WASH Benefits measures in child stool (Cryptosporidium sp., Giardia sp. and Entamoeba histolytica). Additionally, food samples will be analyzed for Campylobacter jejuni, a probable contributor to diarrhea among Bangladeshi children under 12 months old [Taniuchi 2013], and toxins produced by Bacillus cereus, a bacteria common in rice and milk products that constitute the predominant complementary food [Haque 2005, Zhou 2008]. We will also run molecular assays for general, human, ruminant, and avian unique fecal markers in these samples to identify the source of the fecal contamination as originating from humans, cattle or poultry. These source tracking assays detect bacteria that are specific to fecal hosts, such as Bacteroidales; and the identification of host-specific genes by molecular methods allows their use for microbial source tracking [Stoeckel 2007]. Finally, we will analyze the food and soil samples collected during these two rounds for E. coli and fecal coliforms as well to aid with interpreting the pathogen data. This sampling will be done during the first year of the study period to capture the impact of the interventions during ongoing behavior change promotion and the two rounds of sampling (which will fall during the late dry and early wet seasons) will allow us to assess seasonal impacts.

The additional soil sample collected from all sanitation and control arm households at the time of the stool collection for parasitic assessment will be analyzed by microscopy for the soil-transmitted helminths that WASH Benefits measures in stool (Ascaris lumbricoides, Trichuris trichuria, Ancylostoma duodenale and Necator americanus).

Sample Processing:

Measurement of Fecal Indicator Organisms: Field microbiologists will collect 250 mL of water from household storage containers by asking participants to provide a glass of water that they would give to their child to drink. A composite soil sample will be collected from the entrance to the household and 50 g of soil will be excavated from a 30-cm by 30-cm area by scraping the soil surface until 50 g of sample has been obtained. A separate sample of 50 g will be collected from the tubewell area adjacent to the hand pump or to the tubewell platform if one exists. Soil will be scraped into a sterile centrifuge tube with a sterile disposable plastic scoop. Target child and mother hand samples will be collected by rinsing the hands, one at a time, in 200 mL of sterile water in a sterile Whirlpak bag. The field microbiologists will use a sterile collection container with scoop to collect 25 g of food prepared specifically for the target child, if possible, otherwise they will collect a rice-based food that the child has consumed recently or will consume. Only food samples that have been stored for three hours or longer outside of a refrigerator will be collected.

All samples will be placed on ice and transported to the ICDDR,B field laboratory for analysis for E. coli and fecal coliforms within 8 hours. Lab technicians will process aliquots of 100 mL from water and hand rinse samples following the standard IDEXX Colilert most probable number (MPN) method. Soil samples will first be homogenizedby vigorous shaking; 20 g of soil will then be mixed with 200 mL of sterile water and homogenized by mechanical agitation. Serial dilutions will be prepared as needed for pond and soil samples. Food samples will be processed by mechanically homogenizing 10 g with 100 mL of distilled water. After mixing, 10 mL of the solution will be diluted with 90 mL of distilled water and processed by IDEXX. A second aliquot of food will be dried overnight to determine the moisture content. Duplicates and sterile blanks will be run for every 10th sample.

Measurement of Selected Pathogens and Microbial Source Tracking: Aliquots of stored water, mother and child hand rinse samples, soil samples from the household entrance and stored food samples will be collected and pre-processed to concentrate organisms for molecular detection of enteric pathogens, and human and animal specific fecal markers during the two specified rounds of data collection. Aliquots from hand rinse samples (50-100 mL), stored water samples (100-500 mL) and homogenized soil samples (5g) and food samples (5g) will be designated for molecular work. Each water and hand rinse sample aliquot will be vacuum filtered through a 0.45 uM-pore sized filter (HA filter) in order to capture bacterial and viral DNA. Prior to filtration, 0.5 mL of 2.5 M $MgCl_2$ will be added to every 50 mL of sample filtered to facilitate the capture of virus particles on the filter. Filters will then be treated with 500 uL of RNA/DNA stabilizing agent (RNAlater, Qiagen), vacuum aspirated, then stored at -80°C until transport to UC Berkeley. An aliquot of 2g of homogenized soil samples will be measured out and placed in DNA free centrifuge tubes with 1mL of RNAlater, vortexed for 20 seconds, then placed at -80°C until transport to UC Berkeley. A second aliquot of 3g will be measured to determine the soil moisture content.

Archived filters and soil samples will be shipped to UC Berkeley at room temperature, then stored at -80°C until DNA extraction and molecular analysis. RNA and DNA will be extracted simultaneously from water and hand rinse filters using the MoBioPowerWater RNA isolation kit, and RNA and DNA will be extracted from soil samples using the MoBio RNA PowerSoil Total RNA isolation kit. Selected pathogens and fecal source markers will be detected using the PCR, multiplex PCR, and qPCR assays.

Food samples will be pre-processed by aliquoting the contents of the E. coli positive wells of IDEXX trays. The food microbiology lab at icddr,b will process these aliquots forthe presence of the pathogenic E.coli genes eae, ial, bfp, ipaH, st, lt, aat, aaiC, stx1, and stx2using PCR. The food microbiology lab will also analyse food samples directly for Bacillus cereus using culture-based methods.

Measurement of Soil-Transmitted Helminths: Soil samples will be cleaned and helminth eggs will be concentrated through settling, sieving, and floatation steps. Processed samples will be enumerated by direct microscopy in duplicate by two different lab technicians trained in parasitology. The number of Ascaris lumbricoides, Trichuris trichuria, Ancylostoma duodenale and Necator americanus eggs will be counted. Multiple microscopic slides will be prepared and read for each sample, if necessary. Egg counts will be multiplied by the total grams analyzed to determine the concentration of eggs per gram of soil. Samples will be processed and read within 6 hours of collection.

Monitoring Of Uptake:
At each quarterly visit, we will monitor the presence of functional latrine and feces disposal hardware, latrine use and feces disposal practices and presence and quantity of feces in the living environment using spot check observations. We propose collecting repeated measures because spot check observations are a noisy indicator of household behaviors because of temporal variation, and a longitudinal index based on repeated measurements can more finely distinguish meaningful behavioral patterns [Gorter 1998, Ruel 2002]. We also expect time trends in the uptake of the latrine intervention; uptake might initially be low due to unfamiliarity, increase with behavior change promotion and taper off as the novelty dissipates. Quarterly data will allow us to monitor these trends.

We will augment the spot checks with a novel, Passive Latrine Use Monitor (PLUM) that has been developed and validated by members of our team specifically for monitoring latrine usage. We will deploy 30 PLUM sensors in our study population of 720 households in rotating fashion (15 sensors per arm rotated in 24 waves per year in one-week periods) for the two-year study period. The PLUM sensors have been shown to be widely acceptable to households in rural India [Clasen 2012]. PLUM uses a passive infrared sensor and a door switch to discretely and anonymously measure latrine visits. The sensors will enable us to measure the number of latrine visits per household as well as a rich set of information about defecation practices (event frequency, timing, duration) impossible to collect without structured observation. The advantage of sensors over traditional structured observation is that they are less expensive and are less likely to cause measurement bias by reactivity, a demonstrated problem with structured observations of sanitation and hygiene practices [Ram 2010].

Monitoring of Child Health Outcomes:
At each visit, we will also administer a brief interview to the primary caregiver of the index child to assess caregiver reported diarrhea for the index child and other children < 60 months living in the compound.

6. Midline Assessment
Field teams will measure outcomes at 1 year following the initiation of intervention. Children will be between 8 and 15 months at the 1-year survey. This will be the first round in which the field team measures anthropometry. The team will collect information about how many weeks the children and their mothers stayed in another village to understand how many weeks they were out of interventions. The field teams will measure length, weight, and head circumference using standardized measurement techniques. Our anthropometric teams will have been trained and standardized in measurement techniques according to the FANTA and WHO guidelines (Cogill 2003, deOnis 2004). The child will be weighed using a calibrated scale and measure his/her length or height using a height board and head circumference using a tape measure. We will also measure maternal height and weight, and will conduct an assessment of short term maternal stress. The common modules that will be utilized at the midline assessment include:

Module 0. Tracking information
Module 1. Birthdate, age, and sex measurement
Module 2. Diarrhea and illness symptoms
Module 3. Deworming
Module 4. Anthropometry

Module 5. Vaccination history
Module 6. Child Food frequency questionnaire (24 hour and 7 day recall)
Module 7. Handwashing assessment
Module 8. Sanitation assessment
Module 9. Child defecation and feces disposal assessment
Module 10. Water treatment, storage, and quality assessment
Module 12. Home care environment
Module 13. Measures of spillover
Module 14. LNS measurement
Module 15. Environmental microbial assessment and Quantitative fly assessment (subset of households)
Module 16. Children's motor milestones using WHO validated tool and language development via Bangladesh adapted MacArthur Communicative Development Inventories at midline
Module 19. Maternal depression
Module 20. Environmental enteropathy assessment (subset of study population)

We will measure diarrhea morbidity using caregiver report with a 48-hour recall period. We will collect information on index children's deworming medications. The assessment of the sanitation interventions will be the spot checks of latrine structures to assess type, cleanliness, stated use, and state of repair as well as the presence of child feces or other feces that appears to be human in or near the compound. Measures of fly density and sentinal object (Child Toy) contamination will also be assessed.

Assessments of secondary child outcomes

Child:
In assessing the effects of the WASH and nutrition interventions on child health and wellbeing, we will measure some aspects of children's development that may be affected by the treatments. The links between nutrition and cognitive development are clear (Grantham-McGregor et al 2007, Walker et al 2007, Allen et al 2001, Sigman, 1995), but the pathways through which diarrhea and WASH interventions may affect child development are still speculative (Humphrey 2009, Walker et al 2011, Bowen et al 2012). A recent study demonstrated that intensive handwashing interventions for 7 months during the first 30 months of life predicted higher development scores across a range of domains (adaptive, personal-social, communication, cognitive, and motor) at 5-7 years of age (Bowen et al 2012). However, the precise mechanisms of how the treatments improved child development could not be determined. The present study provides the unique opportunity to rigorously examine the associations between WASH and nutrition interventions, child nutritional status, tropical enteropathy and child development outcomes. The findings have the potential of making novel contributions to the WASH, nutrition and child development fields.

We will measure children's motor and language development in all children ~8-15 months of age (that is, born during or after the baseline). Motor skills (sitting, walking, standing) will be assessed directly, using a WHO validated protocol (Wijnhoven et al 2004) adapted for use in Bangladesh. The motor milestone scale has been widely used throughout the world to detect nutritional effects on motor acquisition in Africa (Adu-Afarwuah 2007, Kariger et al 2005), Nepal (Siegel et al 2005) and Bangladesh (Hamadani et al 2013; Tofail et al 2006). Language skills (understanding and speaking words) will be assessed via parent report using the MacArthur Communicative Development Inventory (CDI) (Fenson et al 1994). The CDI is a well-established measure that has been used in more than 40 dialects to describe language development in

---

infants and young children as well as identify group differences in language development (Law &Roy, 2008). The measure provides a valid and reliable method for assessing language in large groups of very young children. Both measures were validated for use in Bangladesh, and have successfully discriminated development in populations with poorly nourished children (Hamadani et al 2010; Tofail 2006) .

Assessments for potential interactions

Child
We will collect a sample of blood for future genetic testing and testing of nutrition and other biochemical parameters, a urine sample for chemistry and a stool sample to permit future testing of the intestinal microbiome from each enrolled child. For the blood sample we will separately freeze aliquots of sera for eventual biochemical analysis, and retain a clot for genetic testing.

We will measure indicators that describe the intake of food for the infants by interviews conducted at the household level using a household survey methodology. The indicators and the instruments of the household survey will be adapted from WHO and UNICEF guidelines on "Indicators for assessing infant and young child feeding practices: Part 2 Measurement" (WHO, 2010). We will measure 24 hours recall and 7 days recall for the indicators of food frequency.

Mother
We will measure the mother’s height and weight and administer the parental stress index (Module 21).

We will collect data on the mother’s height and weight, and maternal depressive symptoms (Module 19) and maternal cognitive abilities to control for their influences on child growth and development. There is substantial evidence that maternal characteristics -- such as education, intelligence and depression -- are associated with infant undernutrition and poor developmental outcomes (Anoop et al 2004; Wachs et al 2009; Walker et al, 2007; 2011). The Centers for Epidemiological Studies-Depression Scale (CESD) (Radloff, 1977) is a brief, widely used measure of 20 statements that assess the likelihood of depressive symptomology. The ICCDR,B psychologists have adapted and used the CESD in various studies, and have noted relationships between higher scores (indicating depression risk), and stunting and lower developmental scores in young children (Black et al 2009; Nahar et al 2012). We will administer the adapted Bangladesh version of the CESD to all mothers of children 8-15 months of age.

Information on maternal education was collected at baseline, but to ensure we are adequately capturing the possible effects of maternal intelligence on child outcomes, we will measure cognitive functioning using three different measures: the Mini-Mental Status Exam, the Digit Span Task and the Verbal Fluency task.

The Mini-Mental Status Exam is a measure of cognitive functioning used in many parts of the world (Mitchell 2009) that has been adapted in Bangladesh for use with illiterate populations (known as BAMSE) (Kabir&Herlitz 2000) . It assesses orientation (knowledge of day, current prime minister), memory, simple calculation, capacity to carry out instructions, and summarization of a short (oral) story. The BAMSE has been used successfully in Bangladesh to detect the effects of early child health interventions on adolescent development (Barham&Calimeris 2008) and the association between malnutrition and cognitive development in older adults (Ferdous et al 2010). The BAMSE will be administered to all mothers of children 8-15 months of age.

---

**Protocol Title:** Measuring the benefits of sanitation, water quality, handwashing and nutrition interventions for improving health and development in rural Bangladesh

**Protocol Status:** APPROVED

**Date Submitted:** 10/26/2017

**Approval Period:** 10/30/2017-11/03/2018

**Important Note:** **This Print View may not reflect all comments and contingencies for approval. Please check the comments section of the online protocol. Questions that appear to not have been answered may not have been required for this submission. Please see the system application for more details.**

The other cognitive measures that we will use with mothers of children 8-15 months of age includes Digit Span and Verbal Fluency Tests. Both the tests are classic tests ofworking memory and short-term memory that has been used around the world. We will specifically use Digit Span (Backward) test that requires respondents to repeat back a string of digits (3-7 in length) in reverse order and has also been used in Bangladesh on primary school aged children (Baddely, 1992; Wechsler, 1994, 1997; Wasserman et al 2011, Huda et al 2001). We will also administer a Verbal Fluency task, which assesses cognitive processing speed by asking respondents to name as many animals as possible in 60 seconds. The test has been used in Jamaica (Baddely et al 1995), Tanzania (Jukes et al 2002) and nutritional trials in Bangladesh (Huda et al 1999; 2001). Both the tests were piloted at field level on 80 rural mothers for the current study with good test-retest reliability, r value for Digit Span and Verbal Fluency were 0.72 and 0.91 respectively and good correlation with socio-demographic variables. The Child Development Index module matches the age group. To control influences of maternal cognitive abilities on child growth and development we are measuring maternal intelligence using Backward Digit Span. Home care environment and maternal depression measurements will be collected.

Home Care Environment
It is well established that the home care environment has a large influence on child health and development (Bradley &Corwyn, 2005; Grantham-McGregor et al 2007; Walker et al 2007 and 2011). We will control for the influence of the home environment on child and health outcomes by collecting data using items adapted from the Home Measurement for Observation for the Environment (HOME) (Bradley et al 2001; Ertem et al 1997) and from the UNICEF Multi-Indicator Cluster Surveys (Kariger et al 2012). Items from these measures have been used to determine differences in child development in Bangladesh (Hamadani et al 2010)

Environmental microbial assessment
Measures of fly density, sentinel object (Child Toy) contamination, hand contamination and drinking water contamination will also be assessed within the subset of households that has been selected for environmental enteropathy measurements (80 clusters (500 households) from each of four arms-- the control group, the combined WASH interventions, the nutrition intervention and the combined WASH and nutrition interventions, for a total of 2000 households). In addition, we will measure hand contamination and drinking water contamination among a subset of up to 360 households per arm in the single intervention (water and hygiene) arms. The sample collection and analysis procedures are detailed below.

Drinking Water
In the 2000 households selected for environmental enteropathy measurements as well as in a subset of households in the water arm, we will ask the caregiver of the target child to give us a glass of water as if giving it to her child. We will collect 250 ml of this water in a sterile Whirlpak and record whether it came from a tubewell or a storage container. We will also ask the caregiver if the water has been treated in any way. All samples will be delivered on ice to the ICDDR,B Laboratory within 24 hours of retrieval for analysis. We will use standard membrane filtration methods to quantify the number of colony forming units (cfu) of E. coli. Sample aliquots of 100 ml will be filtered through 0.45 µm Millipore member filters and filtered samples will be plated on MI Agar.

Target Child Hand Rinse
In the 2000 households selected for environmental enteropathy measurements as well as in a subset of households in the hygiene arm, we will collect a hand rinse sample by rinsing the hands of the target child,

one at a time, in 200 mL of sterile water in a sterile Whirlpak bag. All samples will be delivered on ice to the ICDDR,B Laboratory within 24 hours of retrieval for analysis. We will use standard membrane filtration methods to quantify the number of colony forming units (cfu) of E. coli. Sample aliquots of 10 ml will be filtered through 0.45 μm Millipore member filters and filtered samples will be plated on MI Agar and incubated at 35°C.

Sentinel Toy
In the 2000 households selected for environmental enteropathy measurements, we will assess environmental contamination using a sentinel non-porous toy ball. The toy ball will be initially sterilized and stored in a sterile bag or aluminum foil until it is given to the selected households. After one day, a field assistant will return and ask the mother to locate the toy ball without touching it to avoid hand contamination. The field research assistant will use sterile gloves to retrieve the toy and place it in a sterile Whirlpak bag containing 200-250ml of recovery media (water with salts) or sterile water. The toy will be immersed and bathed in the recovery media for 15 seconds. The field research assistant will remove the toy, place the sealed bag on ice packs, and then wash the toy with soap and water before returning it to the household. All samples will be delivered on ice to the ICDDR,B Laboratory within 24 hours of retrieval for analysis. We will use standard membrane filtration methods to quantify the number of colony forming units (cfu) of thermotolerant fecal coliforms. Sample aliquots of 10 ml and 100 ml will be filtered through 0.45 μm Millipore member filters and filtered samples will be plated on MI Agar and incubated at 44.5°C modified fecal coliform (mFC). If necessary, 10-fold dilutions will be made and plated following the same protocol.

Fly Density
In the 1500 households selected for environmental enteropathy measurements, we will count and speciate flies caught in the latrine and food preparation areas of the compounds using the methods detailed under baseline assessment.

The assessment for the handwashing intervention will be the presence of soap/soapy water and water at the handwashing station, the per capita consumption of soap in the compound, and the assessment of visible dirt on mothers and children's hands. Assessment of hand contamination will be through the use of target child hand rinse samples to test for the presence of E. coli in the environmental enteropathy subset and in a subset of households in the single intervention arms, as discussed above.

The primary assessment of drinking water quality will be through the use of household drinking samples to test for the presence of E. coli in the environmental enteropathy subset and in a subset of households in the single intervention arms, as discussed above. We will also assess the presence of reportedly treated water through tests for chlorine residual, obvious evidence for unused Aquatabs, and the presence and accessibility of the provided storage container (e.g. not readily accessible where water is consumed).

The primary assessment of compliance with the LNS intervention will be through monitoring the remaining LNS sachets unused at the end of every month. In addition, caregiver knowledge, attitudes and practices related to LNS usage and complementary feeding practices will be assessed.

The assessments of each of the interventions will also include questions that address the major behavior

---

**Protocol Title:** Measuring the benefits of sanitation, water quality, handwashing and nutrition interventions for improving health and development in rural Bangladesh

**Protocol Status:** APPROVED

**Date Submitted:** 10/26/2017

**Approval Period:** 10/30/2017-11/03/2018

**Important Note:** This Print View may not reflect all comments and contingencies for approval. Please check the comments section of the online protocol. Questions that appear to not have been answered may not have been required for this submission. Please see the system application for more details.

change constructs for social cognitive theory, for new habit formation and for community mobilization.

Environmental Enteropathy biomarkers will also be measured, as described above (2. Baseline Assessment). We collected 5 ml of blood from each EE participant at midline.

7) Endline Assessment (21 – 27 month)
The final assessment will take place 2 years after the initiation of intervention. Children will be between 21 and 27 months in age. The common modules to be included are:

Module 0. Tracking information
Module 1. Birthdate, age, and sex measurement
Module 2. Diarrhea and illness symptoms
Module 3. Deworming
Module 4. Anthropometry
Module 5. Vaccination history
Module 6. Childfood frequency questionnaire (24 hour and 7 day recall)
Module 7. Handwashing assessment
Module 8. Sanitation assessment
Module 9. Child defecation and feces disposal assessment
Module 10. Water treatment, storage, and quality assessment
Module 12. Home care environment
Module 13. Measures for spillover
Module 14. LNS measurement
Module 15. Environmental microbial assessment and quantitative fly assessment (subset of households)
Module 16. Child development
Module 19.  Maternal depression
Module 20. Environmental enteropathy subsample
Module 21. Maternal intelligence

Trained field staff will repeat the anthropometric, child development, and diarrhea morbidity assessments. Spot checks and behavior change assessments will be performed as described above (see Midline Assessment). In addition to the anthropometric measurement of the index child the team will also measure length, weight, and head circumference for the older sibling next to closest in age of target child using standardized measurement techniques (procedure describe above in the midline assessment section).

Child developmental measures for endline will include the A-NOT-B and Tower Tests for measuring working memory, inhibition and executive function of the children. The A-NOT-B tasks require children to search for objects hidden in specified locations after short delays and reversals. The procedures are based on protocols described in Epsy, K. A., Kaufmann, P. M., McDiarmid, M. D., & Glisky, M. L. (Espy et al 1999). The Tower Test consists of building a tower with the child, and assesses how well children can inhibit responses and impulses and follow directions. The A-NOT-B and Tower Tests are direct tests of the child. In addition, we will use two parent report measures to gather information on development in various domains. These are (1) the MacArthur Communicative Development Inventory (CDI) (Fenson et al 1994; 2007), also administered at midline, that documents words children speak and understand; and (2) the extended and adapted version of the Ages and Stages Questionnaires (Bricker et al 1999; Fernald et al 2012), which will gather information on children's communication, gross motor and personal social skills. (

Boyce et al; Fernald et al 2012; Fenson et al 2007).

In addition, observations of child reactivity to a stressful event (the blood-draw, as described in the Environmental Enteropathy substudy above) will be made for the children enrolled in that substudy. Chronic exposure to stressors, such as overcrowding, poor quality housing, pollution, and family turmoil, is associated with poorer health and developmental outcomes across the lifespan (Shonkoff et al 2009). Early adversity may impact health and development by disrupting the child's ability to regulate stress response and recovery. Sustained physiological response to stress in animals has been linked with damage to the hippocampus, a brain structure important for learning and memory (National Scientific Council on the Developing Child, 2005/2014). For this measure, trained observers will use a simple coding scheme to rate child behavior before, during and after the blood draw (Blair et al 2008). Trained personnel will videotape each child's behavior, and code it using software and procedures recommended by C. Blair (personal communication, P. Kariger). All videotapes will be identified by a number only; no recordings will identify the respondents by name. These videotapes will be viewed only by trained personnel for coding of the child's response to the procedure. Caregivers will also be asked to complete a brief questionnaire on child temperament (subscales on fear reactions and soothability), derived from a measure developed by Ted Wachs (personal communication, P. Kariger), also used in the Gates funded MAL-ED project and in rural Bangladesh (Baker-Henningham et al 2009).

Information on maternal education was collected at baseline, but to ensure we are adequately capturing the possible effects of maternal intelligence on child outcomes, we will measure cognitive functioning using the Backward Digit Span Task (Baddelly, 1992; Wechsler, 1994, 1997). This short-term memory and information manipulation test requires respondents to repeat back a string of digits (3-7 in length) in reverse order. The test has also been used in Bangladesh on primary school aged children (Wasserman et al 2011, Huda et al 2001). The test was piloted at field level on 80 rural mothers for the current study with good test-retest reliability, r value 0.72, and scores showed good correlation with socio-demographic variables.

The assessment of the sanitation interventions will be the spot checks of latrine structures to assess type, cleanliness, stated use, and state of repair as well as the presence of child feces or other feces that appears to be human in or near the compound. We will also collect information about pit switching and pit emptying for the households who have received dual pit latrines. Measures of fly density, sentinel object (Child Toy) contamination, hand contamination and drinking water contamination will also be assessed.

We will collect a sample of blood and urine from each child for future testing of nutrition and other biochemical parameters and a stool sample to permit future testing of the intestinal microbiome from each enrolled child. Furthermore, after endline is completed, we will collect 10 additional monthly follow up stool samples from 60 children enrolled in the EE subset (20 children in the WASH arm, 20 children in the nutrition arm, and 20 children in the control arm). These additional stool samples will be used for future testing of the intestinal microbiome to assess how the interventions impact development and stability of the colonic microbial community. We will further assess how the microbial community interacts with immune functions such as intestinal inflammation and explore the microbiome as a critical link in the causal pathway between interventions and child growth and development. We will analyze the microbial community composition (16S rRNA sequencing), functional gene content (shotgun metagenomic sequencing), gene expression (metatranscriptomic profiling), and assess pathogen load(qPCR). In addition, these stool samples will be tested for the same intestinal inflammation biomarkers as described above for the EE subset (Midline Assessment ).

Environmental Enteropathy, allostatic load, and telomere length biomarkers will also be measured in a subset of participants, as described above (Midline Assessment). In addition, at endline, whole blood hemoglobin will be measured at the time of blood sample collection using a portable spectrophotometer (Hemocue 301). An aliquot of whole blood will be also be sent to the Thalassemia Center at Shishu Hospital to test for markers of inherited hemoglobin disorders (thalassemia and HbE). We will collect a total of 7.7 ml of blood from each EE participant at endline. The increase in blood volume at endline reflects the addition of the anemia measurement.

Furthermore, we have experienced differential refusals by treatment arm in the EE subsample during midline that may threaten the validity of the trial. We experience the highest number of refusals in the control arm. Since the control arm is the arm that we compare all the other arms against, this differential refusal rate is potentially a large problem. We hope to correct this issue by providing a small token of our appreciation, a plastic chair or equivalent item, to the control households at endline. After numerous discussions with the field team, the plastic chair was suggested as a token that did not interfere with our water, sanitation, handwashing, and nutrition interventions and would also be useful to the households in the EE subsample.

Throughout our midline enrollment, the majority of caregivers request blood grouping results. Thus, at endline, from the blood sample we are already planning to take, we will perform a blood grouping test on each mother and child enrolled in the EE subsample. The caregivers will be provided with the results.

Intestinal parasitic infections will also be measured at endline. We will collect stool and blood spot samples from 7 target children per cluster and up to two older children living in the same household compound. The two older children will include (1) the same child that was 18-27 months old at baseline and provided a baseline sample, and (2) an older 5-12 year old child that lives in the same compound as the target child. The procedures will be the same as described in the Baseline Assessment section. Additionally, at endline, an aliquot of each stool sample will also be processed by Kato Katz microscopy for detection of Ascaris, hookworm, and Trichuris ova (Albonico et al., 2012)- the most common soil transmitted helminth infections in Bangladesh affecting children. All members of study compounds will be offered deworming medicine at endline. Field workers will offer it to children because it is logistically too difficult to provide results of tests for worm infections to participants, so they provide it to them all regardless of results in case they are infected. This is a standard practice in research on soil-transmitted helminths. Furthermore, deworming is commonly distributed in mass drug administrations around the world (including Bangladesh) to all school age children, regardless of infection status. The remaining aliquots of stool will be frozen for future biomarker validation (including testing for parasite and environmental enteropathy markers).

Chlorine water testing will be conducted for households that report treating their water. Water samples will be collected from all households and after departure from the home, field staff will test on the samples from those that indicate using Aquatabs to treat their water. In addition, we will collect a sample from source of drinking water from all the households during endline to measure the arsenic (As) and manganese (Mn) concentration using EconoQuick kits.

Sustainability: We propose to add some questions in endline survey in different sections to assess whether participants will maintain their behaviour once we stop visiting their households and stop providing them supplies. The purpose of adding sustainability questions is to obtain a baseline for comparison to use after we survey again in 1-2 years. We have developed these questions through a process of discussion and revision with several of the WASH Benefits team members. Currently, we are in the process of finalizing the list of sustainability questions through a broader discussion. There are three domains of sustainability questions. The first domain includes questions regarding whether participants have the knowledge and skills to use and maintain latrines, safely store water, wash their hands, and provide nutrient-rich complementary foods to their children. The second domain includes questions regarding participants' self-efficacy to use and maintain latrines, safely store water, wash their hands, and provide nutrient-rich complementary food to their children. The third domain includes questions regarding some of the physical items and structures present at the time of the survey such as the types of water storage containers in the home and the condition of the latrines. We will also take a photograph of the latrine slab to use as a baseline.

For child health improvements to translate into economic gains, complementary investments in schooling and cognitive stimulation must be made (Cunha & Heckman, 2007). Such investments will only be undertaken if there is a belief (high subjective probability) that it will have a positive return. We plan to investigate the impact of the health interventions delivered by WASH-N on parental beliefs about the capability of their child. We will do this by comparing treated versus control siblings in experimental versus placebo clusters. We plan to follow these survey results with a longer term study of whether such expectations translate into higher schooling investments and whether the ability of parents to translate high expectations for their children into human capital (e.g. schooling) is affected by the level of physical capital (e.g. schools/teachers) in a given area. The data we plan to collect are crucial for identifying the effects of the program and understanding the causal linkages between health status in early childhood and later productivity.

Antibiotic resistance is a growing global public health challenge that could undo decades of progress in improving access to effective antibiotics and reducing the burden of infectious disease in the developing world (Laxminarayan et al., 2013). Oftentimes, antibiotics are used for nonspecific viral symptoms, hence reducing the burden of such infections may reduce the use of antibiotics which is directly linked to the development of resistance (WHO 2012; 2014). To test this hypothesis, we will add questions on antibiotic use and compare individuals in the treatments and control arms to see whether antibiotic use has diminished at the household level. We will also probe their general understanding of the appropriate use of antibiotics and how they are consumed. Such information is critical for halting the spread of antibiotic resistant pathogens.

Environmental microbial assessment
Measures of fly density, sentinel object (Child Toy) contamination, hand contamination, drinking water contamination and food contamination will also be assessed within the subset of households that has been selected for environmental enteropathy measurements (80 clusters (500 households) from each of four arms-- the control group, the combined WASH interventions, the nutrition intervention and the combined WASH and nutrition interventions, for a total of 2000 households). In addition, we will measure hand contamination, drinking water contamination and food contamination among a subset of up to 360 households per arm in the single intervention (water and hygiene) arms. The sample collection and analysis procedures are for the fly density and sentinel toy, hand and drinking water testing components have been described in the midline environmental microbial assessment section. The procedures for food testing are described below.

Field team will sample any previously prepared food being stored in the household for consumption by the

---

index child by collecting ~25-50g directly from the storage pot using a sterile spoon, then placed into a sterile plastic bag. Samples will be transported on ice to the field laboratory. 25g of sample will be homogenized with distilled water for 10x dilution. 1 mL of the homogenized slurry will be analyzed for E.coli using TBX media with the ISO 16649: Microbiology of food and animal feeding staff-Horizontal method for the enumeration of beta-glucuronidase-positive E. coli. Further decimal dilutions will be prepared as required.

Monitoring and Process Documentation

Valid evaluation of the study hypotheses requires a consistent intervention, but the WASH Benefits intervention is a large complex intervention spread over 5 districts. While an intervention of this scope and complexity involving this many individuals will inevitably have some deviation from optimal implementation as planned, meaningful interpretation of the results requires a rigorous assessment of how consistently the intervention activities were delivered. This assessment will include 3 components:
1)A monitoring system that tracks
a)procurement and distribution of commodities and technologies
b)recruitment, training and supervision of promoters
2)An unannounced fidelity assessment of the delivered intervention (Appendix 6)
a)The fidelity assessments will be conducted
i)In each of the initial 1216 intervention implementation clusters after 1, 2, 3 and 4 months of intervention
ii)In each of the second group of 1216 implementation clusters after 4 months of intervention
iii)In each of the third group of 1216 implementation clusters after 3 months of intervention
iv)In each of the fourth group of 1216 implementation clusters after 2 months of intervention
v)In each of the fifth group of 1216 implementation clusters after 1 month of intervention
vi)In each of these groups (i-v) subsets of BCC samples 50% which is 4 out of 8 households
vii)Assessments will continue in 24intervention clusters (4 blocks) per month (4from each of the 6 interventions with a subsets sample 50% )

Revised fidelity assessment strategy :
The revision of the fidelity assessment will be made to see the uptake status both for initial phases as well as new phases with a intention to have a similar number of samples from new phases and reducing samples from initial phases.

Stages:
Selection of 1 block from more than 6 months intervention implementation blocks from each phase (10 blocks phase = 1 stratum) and selection of 2 blocks from each phase (stratum) where intervention implementation less than or equal to 6 months.

Selection of 9 blocks (1 block from each phase) randomly after 6 assessment months and it will continue up to reaching 24 months by phase one.

Fidelity assessment will be ended in one point of time when the first phase will complete 24 months of intervention

b)The focus will be on the households that received the intervention and will address key elements of the intervention

i)Are each of the elements of hardware distributed to the appropriate home
ii)Are each of the elements of distributed hardware functional?
iii)Is there objective evidence of uptake
(1)Handwashing
(a) presence of a HW station
(b)Water present?
(c)Soap present?
(d)Soapy water bottle present?
(e)Soapy water present?
(f)Is all equipment in working order?
(2)Water
(a)Is the icddr,b provided water storage container present?
(b)Are Aquatabs present?
(c)Is drinking water in the storage container?
(d)Is the equipment in working order?
(e)Are the containers being used for storage of other liquids?
(f)Is there detectable residual chlorine
(3)Sanitation
(a)Is the potty available?
(b)Is the potty immediately available to the child?
(c)Is the potty in working order?
(d)Is the poop scoop available within 30 seconds?
(e)Is the poop scoop in working order?
(f)Does the latrine show signs of use?
(g)Is there an odor of feces in the latrine?
(h)Is the latrine in working order?
(4)Supply of LNS
(a)How many days ago was it delivered?
(b)How many sachets are present?
(c)How many sachets have been used?
iv)For each intervention, 1 – 3 key questions will be asked of persons who received the intervention to see if they received the principal message of the intervention
v)How many times has the community health worker visited
(1)in the preceding week?
(2)in the preceding month?
c)The data will be collected by smart phone, that is uploaded to ICDDRB server the day it is collected with reports produced every month and circulated to the management team.
d)Assessments are front loaded so that early problems can be addressed through refresher training, additional supervisory visits or other appropriate strategies.
3)A process documentation system that tracks operational problems and how they were addressed, and modification made to intervention and the reasons for these modifications.

A primary goal of the process documentation is to provide the CHP supervisors with the information necessary to ensure the intervention is implemented according to the prescribed strategy and to identify areas where additional communication and training would be helpful.  In addition an external consultant has been contracted to conduct process evaluation throughout the intervention.

---

**Protocol Title:** Measuring the benefits of sanitation, water quality, handwashing and nutrition interventions for improving health and development in rural Bangladesh

**Protocol Status:** APPROVED

**Date Submitted:** 10/26/2017

**Approval Period:** 10/30/2017-11/03/2018

**Important Note:** **This Print View may not reflect all comments and contingencies for approval. Please check the comments section of the online protocol. Questions that appear to not have been answered may not have been required for this submission. Please see the system application for more details.**

Structured Observation

Trained field workers will conduct structured observation in a subset of participating households to observe how the interventions are being used. We will randomly select 6 blocks from each of the nine Phases of study area. From each of the selected 54 blocks, we'll randomly select one household in each cluster. Therefore a total 432 households will be selected for structured observation during midline. If the uptake seems to be low during midline below the benchmark, we will consider conducting structured onservation during endline.

If the structured observation data suggest important difficulties in uptake and use of the interventions, then the project leadership will consider additional qualitative investigation (detailed below) or changes to the intervention.

Additional structured observations will be conducted in a total of 150 households (subset of the 2160 households enrolled in the in-depth environmental contamination assessment) in the sanitation alone arm (n=50), the combined sanitation, water, and hygiene arm (n=50), and the control arm (n=50) to characterize hygiene, water treatment, food hygiene, and sanitation behaviors in richer detail and with an emphasis on child exposure to feces or fecal contamination (consent form in Appendix 1r). Each structured observation will allow for observation of child defecation events and child feces disposal practices, child feeding events, latrine usage, food preparation, food storage, and handwashing behavior at critical times.

To complement the structured observations, we will obtain video surveillance of practices and activities in the same 150 households to capture key water, sanitation and hygiene practices and behaviors. Video surveillance will be conducted by local women trained to conduct video data collection; videographers will also record notes on behaviors during the observation. Behaviors includes personal hygiene, household water management (water usage and storage pattern), handwashing practices at critical times, object contact with children's hand and mouth, presence of feces within and surrounding the households, defecation practices of older children (3+ years) interactions with domestic and pet animals, dealing with and disposal of animal feces, contact with mother and family members. Observations will be made of how commonly children come in contact with soil in their own and neighboring compounds, play with children including those from neighboring compounds.

Qualitative Investigation

We will conduct up to 45 in-depth interviews (subset of the 2160 households enrolled in the in-depth environmental contamination assessment) in the sanitation alone arm (n=15), the combined sanitation, water, and hygiene arm (n=15), and the control arm (n=15) to explore individual beliefs and perception of certain practices that contribute to fecal contamination (consent from in Appendix 1s). We will focus on personal and household hygiene behaviors (handwashing, food preparation, water management), decision making processes, empowerment, defecation practices among children, women, sick and older people; animal rearing methods (open or corralled, within or outside the households) contributing to environmental fecal contamination. The in-depth interview guidelines are in Appendix 3 (Module 53).

We will conduct at least:

---

•30 interviews with primary and secondary caregivers of the children,
•5-10 interviews with the person accompany a child most of the time in absence of a caregiver (grandmothers, older siblings, aunts of the child).
•5-10 interviews with fathers of the children to explore their concern and role on child's health and development.

The principal and co-principal investigators will carefully review the intervention fidelity assessments and identify any areas of low uptake of interventions. Critical benchmarks for uptake based on unannounced visits are:

The principal and co-principal investigators will carefully review the intervention fidelity assessments and identify any areas of low uptake of interventions. Critical benchmarks for uptake based on unannounced visits are:

•Handwashing promotion households
o65% of households have at least one handwashing station with soap and water present
•Water quality intervention households
o65% of households with children 6 – 24 months of age have stored chlorinated drinking water
•Sanitation
o80% of households have a potty easily accessible to mother
o50% of households have a potty easily accessible to child 12 – 36 months of age
o80% of households have a sani-scoop easily accessible to mother
o80% of households have a latrine with a functional water seal
•Nutrition
o80% report hearing any messages on infant/child nutrition and or Sonamoni
o90% report at least one visit by the CHP in household to discuss infant and child nutrition
oWithin households with targeted children > 6 months of age, the stock of LNS sachets in 70% of households is consistent with daily use of two sachets per day

If any of the uptake measures are below the critical benchmarks, then a qualitative team will review the monitoring and process documentation in the low performing area, visit the site of the low uptake, meet with community hygiene promoters, supervisors and study subjects and troubleshoot the cause of the low uptake. Because these interventions have each been piloted and in the pilots achieved these benchmarks of uptake, we expect that uptake below the benchmark will indicate a problem where the intervention was not implemented as planned, and the investigation will identify needs to provide additional training or other support to achieve the planned intervention.

While unlikely, it is also possible that the community hygiene promoters will be implementing the intervention precisely as planned, but uptake is lower than expected. If uptake is below the benchmark in the setting where implementation followed the prescribed approach, the qualitative team will conduct more in-depth evaluation will be framed around the Integrated Behavioral Model for Water, Sanitation and Hygiene (IBM-WASH), based on the earlier Integrated Model for Hygiene, Point-of-use water treatment and Sanitation behaviors (IMHPS). The IBM-WASH was developed and refined based on the pilot phase of the WASH Benefits project, as well as the concurrent Cholera Behavior Change (CBC) study carried out in Mohammadpur, Dhaka. It also incorporates behavioural determinants from a number of previous models

---

used for WASH behaviour change interventions. This ecological (multi-level) model has 5 levels (Societal/structural, Community, Interpersonal/household, Individual and Behavioral/Habitual) and 3 dimensions (Contextual, Psychosocial and Product/Technology).

One investigation in one cluster will involve up to:
•5 interviews with implementation partners
•10 observations of household visits by CHPs
•10 household-level interviews with 3 individuals in each household: One person responsible for maintaining the product/hardware e.g. handwashing station, and two users of the product/hardware.
•10 household-level observations of the product, its condition and associated factors affecting its use.

We anticipate 10 investigations per year, for a total of up to 35x10 = 350 interviews and 10x10 = 100 observations of CHP visits and 10x10 =100 household-level observations.

The objective of these investigations is to rapidly identify problems that can be addressed throughout the intervention to ensure a consistent high quality intervention with regular uptake by study participants.

Ethnographic investigation
Two female researchers will stay for approximately two months in two communities at a time and collect data through observations during waking hours in a natural setting when the activities related to defecation, handling fecal matter or waste disposal occurs. The selected communities will not be enrolled in the WASH Benefits study, but will be near the WASH Benefits study area. The ethnographers will collect data through participant observation (unstructured), informal conversation, in-depth interviews (consent form in Appendix 1s) and focus group discussions (consent form in Appendix 1t) when necessary with community members, for a total of 15 in-depth interviews with adult males and females, with the primary and secondary caregivers of children to record beliefs and practices related to water, sanitation and hygiene. The researchers will also conduct 6 (depending on the community setting) focus groups with the community members. The focus group guidelines are in Appendix 3 (Module 54).

Lead Assessment
At baseline mothers have a blood sample collected using lead-free blood collection equipment, and are asked where most of the rice they eat comes from. We will randomly select 500 mothers for whom most of the rice they eat comes from their own fields and will analyze their blood lead levels in the nutritional biochemistry laboratory at icddr,b. Lead will be measured in whole blood by the Graphite Furnace Atomic Absorption Spectrometry (Shimadzu). For quality control, Standard Reference Material from the National Institute of Standards and Technology (SRM, NIST) is used for external quality control purpose. For internal quality of the assay, duplicates are run as well as recovery checks in every lot are performed to maintain the quality.
Qualitative researchers will visit 15 of the women with high blood lead levels and seek informed consent for participation in this component of the study (Appendix 1n). They will observe the women’s living environment and ask questions about various potential sources of lead exposure including pesticides, herbicides, fertilizer, industrial processes or wastes that use lead, cosmetics and canned food (Appendix 3, Module 50). Through these conversations they will generate a list of potential exposures. The anthropologists will visit four area shops that sell agrochemicals, seek the informed consent of shop proprietors (Appendix 1o) and ask questions to understand the range of chemicals available and their

typical selling pattern (Appendix 3, Module 51). They will use this information in discussion with the women and whomever in the household purchases and applies agrochemicals to develop a taxonomy that explicates how people in this area classify the various chemicals and so how questions can be framed to assess exposures. The anthropologists will also ask questions about the process of applying agrochemicals, including who mixes and applies them, how they are mixed and applied, where the chemicals are stored, and how the containers that stored the chemicals are used or disposed (Appendix 3, Module 51).
The project collaborators will review the findings from the in-depth interviews and use these to revise the multiple choice questions to explore potential exposures to lead in these communities (Appendix 3, Module 52). We will classify the 100 mothers with the highest blood lead levels as cases and the 100 mothers with the lowest blood lead levels as controls. Field research assistants will re-visit these mothers and seek their consent to participate in this component of the study (Appendix 1p). The field team will administer the questionnaire to the cases and controls.
During the household visit the field team will collect a sample of uncooked rice that was grown in the study participant's field and a sample of soil from the field. To collect the soil sample, the field worker will ask the respondent to identify the agricultural field where most of the rice that they consume is grown. They will identify the corner of the field that is the farthest from the household and designate the border of the field that is most closely aligned east to west access as the x-axis and the border that intersects it as the y-axis. They will consult a random number table that lists random integers between 1 and 9. They will use these 2 numbers as coordinates measured in meters to identify a point within the field to collect a soil sample. If the random number selected corresponds to a point that is not located within the field, the field team will select the next random number in the table. The field team will identify 3 coordinates within the field.
At each of the selected coordinates, field workers will collect soil to a depth of 25 cm (rooting depth) using a 2 cm diameter push corer. Field workers will extrude soil from the corer and placed in plastic, zip-lock bags. Soil and food samples will be shipped to Stanford, dried and ground. The samples will then be examined by X-ray fluorescence (XRF) spectroscopy to determine the concentrations of Pb. The XRF analysis will be conducted in the Environmental Measurements facility (http://em-1.stanford.edu/) at Stanford, which is shared analytical facility in the School of Earth Sciences.
We will analyze the questionnaire data, soil and food lead levels and assess which exposures are statistically associated with elevated blood lead levels.
The anthropology team will follow up on the exposures that are identified in the case-control study to determine how and why people come in contact with this source. If the analysis supports the hypothesis of agrochemical contamination as the primary pathway of lead exposure in this population, the field team will return to a subset of houses over the course of the season to collect a representative sample of agrochemicals to understand more thoroughly which chemicals are being applied and identify those that are most likely to contain lead for further testing.
Once the source of lead is confirmed, by confirmation of high levels of lead in the pathway statistically associated with high lead levels in the mothers, the Bangladesh anthropology team will return to the community to explore the patterns of lead containing product use and trade further. Where are the products purchased? Why are these specific products used, and where do these products come from? Who sells the products, and why do they sell these product and not others? We will trace back the source of production as far as possible and explore any existing regulations and the process of their enforcement.

Spillover substudy
The household with a child 0-59 months that is nearest to a compound enrolled in the combined intervention and control arms will be eligible for this study. We will administer a survey to measure caregiver reported diarrhea and respiratory illness of children under five years. We will define diarrhea as 3

or more loose or watery stools in 24 hours, or ≥1 stool with blood. Respiratory illness will be defined as a persistent cough or difficulty breathing in the 7 days before the interview. Symptoms will be reported by caregivers collected daily for the 7 days preceding the interview. We will collect a stool sample and analyze it for soil transmitted helminth (Ascaris, Trichuris, hookworm) ova in stool specimens among children 0-59 months. We focus on this age range because children in this range experience the greatest burden of diarrhea and respiratory diseases and are not covered by the national school-based deworming campaign. To measure helminth infection, field workers will deliver a stool collection kit, give instructions to the caregiver, and return the following day to collect the specimen. Field staff will collect specimens, preserve them, and detect helminth ova using the Kato-Katz technique.

Field workers will also visually inspect of mother and children's fingers, fingernails, and palms for dirt and assess the presence of a handwashing station and soapy water bottles in the kitchen/latrine. They will take a sample of drinking water typically provided to a child, and we will analyze the samples for E. Coli using DelAgua kits. Field workers will also count feces piles in courtyards. They will assess contamination of sentinel objects using the methods outlined above. They will assess fly density in the compound by recording the number of flies that land on sticky cards affixed to walls near where food is prepared for a 24 hour period.

To understand mechanisms of spillovers, we will also conduct in-depth interviews with up to 30 participants in the spillover substudy. We will ask about the places they learn about health and hygiene behaviors, how they interact with their neighbors, and other information that will help us ascertain potential mechanisms of spillovers.

**b) Explain who will conduct the procedures, where and when they will take place. Indicate frequency and duration of visits/sessions, as well as total time commitment for the study.**

Trained field staff will conduct all anthropometry, diarrhea morbidity, and child development assessments in the participant's home or at a nearby field office at the beginning of the study, one year following intervention implementation, and 2 years following intervention implementation. These assessments will take about 1 hour to complete, and we expect the baseline/enrollment visit to require 2 hours total.

Local health promoters will visit households several times weekly early in the study to encourage families to adopt the new behaviors, help with solving problems and answering questions. Visits will gradually taper off to at most monthly by the end of the two year period, though participants will be encouraged to contact the promoters if they encounter equipment breakage, or consumable product shortage. In the arms receiving the nutrition intervention, the promoters will deliver the supplement to target households, teach the mother/guardian about proper use of the supplement, and will deliver behavior change communications to encourage breastfeeding and proper feeding of complementary foods after 6 months of age. In arms receiving water, sanitation, and/or handwashing interventions, likewise, the promoters deliver technologies and supplies, look after repairs, problem solve, answer questions and doubts, encourage and congratulate the adopters.

Environmental Enteropathy assessment will be conducted by trained staff, including phlebotomists, at two household visits within participating communities. This assessment will take place at the beginning, middle and end of the study. The assessment will take about 10 hours and 15 minutes per participant and will occur during two days. On the first day, the field team visit each household to deliver the stool collection kit and provide instructions as well as conduct anthropometry, collect blood, saliva, and hair, and measure heart rate, blood pressure, autonomic function, and skin conductance. On the second day, they will collect stool, urine, saliva and administer the infant food frequency questionnaire. The first visit is expected to take 3 hours and 15 minutes and the second visit is expected to take 7 hours.

Stool specimen collection to test for intestinal parasites will be conducted by trained field staff during the final survey. Stool collection will require two visits to each household. The first visit will last about 20 minutes and will involve delivery of the stool collection kit and provision of instructions to the caregiver regarding stool collection. The following day, field staff will return to collect the stool specimen. This visit is expected to take 10 minutes.

The qualitative team will investigate barriers for uptake using unstructured and structured observations, in-depth interviews, and doer/non doer analysis as indicated by the situation. The objective of these investigations is to rapidly identify problems that can be addressed throughout the intervention to ensure a consistent high quality intervention with regular uptake by study participants.

**c) Identify any research procedures that are experimental/investigational. Experimental or investigational procedures are treatments or interventions that do not conform to commonly accepted clinical or research practice as may occur in medical, psychological, or educational settings. Note: if the study only involves standard research or clinical procedures, state "N/A."**

The element of this study that is not a standard public health interventions is the intensive data collection required to learn from the experience. Most of this data collection involves observation and collection of information that is not particularly culturally sensitive.

The LNS used in the study will be a slightly modified variant of Nutributter, which is a commercially-available supplemental feeding product sold by Nutriset. The specific LNS formulation that we will use has been developed by our nutrition colleagues at UC Davis and has been tested extensively in Bangladesh, Malawi, Ghana and Burkina Faso.

**d) If a placebo will be used, provide rationale and explain why active control is not appropriate.**

The control group will not receive a placebo.

**e) If any type of deception or incomplete disclosure will be used, explain what it will entail, why it is justified, and what the plans are to debrief subjects. See CPHS Guidelines on Deception and Incomplete Disclosure for more information. Any debriefing materials should be included in the Attachments section.**

N/A

**f) State if audio or video recording will occur and for what purpose (e.g. transcription, coding facial expressions).**

Audio, video taping and photography may be done during assessment visits to record interviews and observations. Video recording (with audio) for stress reactivity around the blood draw will be conducted as follows: A waterproof video camera will be placed on a tripod about 15-20 feet from the child-caregiver pair. Videotaping will begin 10 minutes prior to the blood draw, and continue until 25 minutes after the blood draw. At all times, the child will be seated on the lap of the caregiver. The camera will be focused on the child’s face. The camera will be swiveled as necessary to capture the child’s response. The videotapes will be coded at a later time for facial expressions and vocalizations. We will only code selected minutes before, during and after the blood draw. The video, audio records and the photos will be erased from cameras after transferring into the computer. The soft copies will be secured by the computer security system (password protection) until the completion of the study and only the investigator of this study will have the access to open it.  All media will be digitized and securely stored in password protected devices for up to 20 years. At that point or once the data is analyzed, the media files will be deleted from the devices. Some photographs or video sections may be used in public presentations and on project websites. Specific consent for these uses will be obtained from study participants. Identifiable information stored electronically on a removable medium or networked computer will be encrypted, as per CPHS policy.

#### 10. Alternatives to Participation

**Describe appropriate alternative resources, procedures, courses of treatment, if any, that are available to prospective subjects. If there are no appropriate alternatives to study participation, this should be stated. If the study does not involve treatment/intervention, enter "N/A" here.**

Prospective subjects are free to carry on with their current sanitation and hygiene practices, regardless of whether or not they chose to participate in the study. Those who choose not to participate can purchase materials (available outside the study) on their own to improve their child’s nutrition and/or their home environmental conditions.

---

---

| | |
|---|---|
| **Protocol Title:** | Measuring the benefits of sanitation, water quality, handwashing and nutrition interventions for improving health and development in rural Bangladesh |
| **Protocol Status:** | APPROVED |
| **Date Submitted:** | 10/26/2017 |
| **Approval Period:** | **10/30/2017-11/03/2018** |
| **Important Note:** | This Print View may not reflect all comments and contingencies for approval. Please check the comments section of the online protocol. Questions that appear to not have been answered may not have been required for this submission. Please see the system application for more details. |

#### * * * Radiation * * *

11. Ionizing Radiation (e.g. X-ray) and Non-ionizing Radiation (e.g. MRI)

a) Do you intend to use ionizing radioactive materials or ionizing radiation-producing devices in your research (e.g., injectable, oral, x-rays, etc.)? CAUTION: The UCB Radioactive Materials License does not permit human research using radioactive materials or radiation from such materials. N

If Yes, provide Radiation Use Authorization (RUA) number(s):

Note: The research may not proceed without an RUA. Please visit:

http://www.ehs.berkeley.edu/how-create-new-radiation-use-authorization-rua

b) Do you intend to use any non-ionizing radiation sources (laser or magnetic sources) in your research? N

If Yes, provide Laser Use Registration (LUR) number(s):

And/or Magnetic Inventory number(s):

Note: The research may not proceed without an LUR or Magnetic Inventory Number. Please visit:

http://ehs.berkeley.edu/laser-safety/how-do-i-register-my-new-laser

c) Describe the source of ionizing radiation or non-ionizing radiation.

---

---

**Protocol Title:** Measuring the benefits of sanitation, water quality, handwashing and nutrition interventions for improving health and development in rural Bangladesh

**Protocol Status:** APPROVED

**Date Submitted:** 10/26/2017

**Approval Period:** 10/30/2017-11/03/2018

**Important Note:** This Print View may not reflect all comments and contingencies for approval. Please check the comments section of the online protocol.
Questions that appear to not have been answered may not have been required for this submission. Please see the system application for more details.

#### * * * Medical Equipment, Investigational Devices * * *

##### 12. Medical Equipment

**If the research involves use of medical equipment, explain whether the equipment is approved for marketing and routinely employed in clinical practice.**

The following equipment will be used to collect blood and urine samples and for anthropometric measurements. These equipment are all available through major commercial retailers, so our understanding is that they are approved for marketing and routinely employed in clinical practice.

- ECLP Hypodermic needles
- Syringes
- Contact-activated lancets
- Trace metal needles for children
- Reusable tourniquets for children
- Urine specimen collection bags
- Head circumference measuring tape
- ITC Tenderfoot incision devices
- Powder-free nitrile gloves
- Seca 383, 413, 874 digital scales
- Salimetrics children's swabs
- Urine collection jar
- Pediatric urine collector
- Cotton swabs
- ShorrBoard infant/child measuring boards
- Sterile newborn U-bag, 24-hour style
- Prestige medical tourniquets
- 3m Transpore surgical tape

##### 13. Investigational Devices

**List in the table below all Investigational Devices to be used on subjects**

Investigational Devices

---

---

| | |
|---|---|
| **Protocol Title:** | Measuring the benefits of sanitation, water quality, handwashing and nutrition interventions for improving health and development in rural Bangladesh |
| **Protocol Status:** | APPROVED |
| **Date Submitted:** | 10/26/2017 |
| **Approval Period:** | **10/30/2017-11/03/2018** |
| **Important Note:** | **This Print View may not reflect all comments and contingencies for approval. Please check the comments section of the online protocol. Questions that appear to not have been answered may not have been required for this submission. Please see the system application for more details.** |

#### * * * Drugs, Reagents, or Chemicals * * *

##### 14. Drugs, Reagents, or Chemicals

**a) List in the table below all investigational drugs, reagents or chemicals to be administered to subjects during this study.**

**b) List in the table below all commercial drugs, reagents or chemicals to be administered to subjects during this study.**

| | |
|---|---|
| **Generic Drug Name and Synonyms** | Lactulose:mannitol solution |
| **Source of Drug** | International pharmaceutical supplier |
| **Manufacturer** | |
| **If not pre-formulated, where will the material be prepared and by whom?** | ICDDRB, by trained laboratory technicians. The solution will consist of 250 mg of lactulose and 50 mg of mannitol per milliliter of sterile water |
| **Indications for Use of Drug** | To measure intestinal absorption |
| **Dosage** | 2mL per kilogram body weight |
| **Route of Administration** | oral |
| N | Are these new or unapproved uses of these commercially available drugs, reagents, or chemicals? If Yes, you may be required to submit an Investigational New Drug Application (IND) to the FDA. See Off- Label Use of Marketed Drugs. |
| **IND# (if applicable)** | |

| | |
|---|---|
| **Generic Drug Name and Synonyms** | Aquatabs (chlorine tablets) |
| **Source of Drug** | in-country supplier |
| **Manufacturer** | Medentech |
| **If not pre-formulated, where will the material be prepared and by whom?** | |

---

| | |
|---|---|
| **Protocol Title:** | Measuring the benefits of sanitation, water quality, handwashing and nutrition interventions for improving health and development in rural Bangladesh |
| **Protocol Status:** | APPROVED |
| **Date Submitted:** | 10/26/2017 |
| **Approval Period:** | **10/30/2017-11/03/2018** |
| **Important Note:** | **This Print View may not reflect all comments and contingencies for approval. Please check the comments section of the online protocol. Questions that appear to not have been answered may not have been required for this submission. Please see the system application for more details.** |

| | |
|---|---|
| **Indications for Use of Drug** | water purification |
| **Dosage** | 5 mg per liter of water |
| **Route of Administration** | drinking water treatment |
| N | Are these new or unapproved uses of these commercially available drugs, reagents, or chemicals? If Yes, you may be required to submit an Investigational New Drug Application (IND) to the FDA. See Off- Label Use of Marketed Drugs. |
| **IND# (if applicable)** | |

---

---

**Protocol Title:** Measuring the benefits of sanitation, water quality, handwashing and nutrition interventions for improving health and development in rural Bangladesh

**Protocol Status:** APPROVED

**Date Submitted:** 10/26/2017

**Approval Period:** 10/30/2017-11/03/2018

**Important Note:** **This Print View may not reflect all comments and contingencies for approval. Please check the comments section of the online protocol. Questions that appear to not have been answered may not have been required for this submission. Please see the system application for more details.**

#### * * * Risks and Discomforts * * *

##### 15. Risks and Discomforts

**a) Describe all known risks and discomforts associated with study procedures, whether physical, psychological, economic or social (e.g., pain, stress, invasion of privacy, breach of confidentiality), noting the likelihood and degree of potential harm.**

There is minimal risk of physical, psychological, social, or legal injury from participation in this study.

No severe allergic or other reactions to Nutributter (a product with similar ingredients to the LNS used in this pilot study) were observed in similar studies in Ghana or Bangladesh (conducted by our UC Davis team members), and none are expected in this study. In the LNS arms, when the children are age 6 months we will ask about peanut allergies and will test children with a small sample of the LNS supplement. If a child is allergic, we will not give them LNS but we will retain them in the intention to treat analysis. Community health workers will be trained to tell mothers that they should be aware of any allergic reactions in their children after using the LNS supplement, and in the event of an allergic reaction, not to give the supplement.

The intervention hardware, improved latrines, child potties, handwashing stations, and aluminum water storage vessels are interventions that are widely promoted and used in a variety of contexts. The interventions involve a behavior change component that involves developing communication messages and training local health promoters to deliver these messages. This is a standard approach to public health promotion used throughout Bangladesh and other contexts.

Some aspects of the interventions and data collection activities might be uncomfortable for subjects to discuss, given cultural sensitivities surrounding the topic of defecation. Currently many young children defecate in the open and parents sometimes do not clean this up and sometimes clean it up with a hoe. When using a potty or sani-scoop, parents may be exposed to their children's feces in somewhat different ways than they previously were.

There is the slight risk of breach of confidentiality. Community members may see study staff entering other baris and homes of their neighbors (usually all family members within a bari), and may overhear interviews. We will make every effort to ensure that household surveys and structured interviews are conducted in privacy.

The measurements for environmental enteropathy involve administration naturally occurring sugars, and small children seem to enjoy the flavor. There is the risk that some children will not like the flavor and will be upset when it is administered. Collection of urine and stool may be uncomfortable to the parent or child. There is also the risk of short-term discomfort and pain during the collection of venous blood and blood spot samples.

**b) Discuss measures that will be taken to minimize risks and discomforts to subjects.**

We will discuss the objectives of the research with participants as part of delivering the intervention. All participants will be informed of potential discomforts during the consent process, and may decide not to participate at any time during the study. The study team will obtain informed consent from all participants. Individually identified information will be kept confidential.

Potential harms of the study include that people will give time to the study that would be better given to address other issues. We will address this risk by securing informed consent, and clarifying that study participants can drop out at any time, even in the middle of an interview or group discussion.

Discomfort during venous blood draw will be minimized by using trained phlebotomists to collect the specimens. Needles used to collect blood will be disposed of in a safe manner and will not be re-used. Field staff will be carefully trained to collect blood spot samples. We will explain the procedures to the parents and will be available to answer any questions they may have.

We will streamline the data collection procedures as much as possible in order to take as little of the subjects' time as possible. We will make every effort to put subjects at ease during discussions of sensitive topics such as defecation, by using culturally appropriate terminology or euphemisms as possible, and by reminding subjects at the outset that they are free to withdraw from the study activities at any point. It will be made clear during promotion activities that individuals are free to continue their current sanitation practices, and that no one should be coerced to adopt latrine usage. When parents are trained in use of the child's potty, they will be taught the importance of handwashing with soap afterwards.

If hypertension is identified for mothers and children of the EE subsample, we will refer them to the governmental hospitals for treatment. Similarly, if we identify anaemia among the children, we will also refer them to the nearby governmental hospitals.

To minimize the risk of breach of confidentiality, every effort will be made to conduct interviews in the privacy of the participant's home. Data collected during interviews and observations will be kept secure by study staff.

**c) If applicable, indicate if a particular study treatment or procedure may involve risks to the subject (or to the embryo or fetus, if the subject is or may become pregnant) that are currently unforeseeable.**

N/A

**d) If applicable, describe the Data Safety Monitoring Plan (DSMP). NIH may require a DSMP for some projects.**

An independent Data Monitoring Committee will be assembled in Bangladesh to monitor adverse events and safety and to advise research investigators. The Committee will include a multi-disciplinary team that will track adverse events in the nutrition arms of the study. This board will meet twice each year, once by phone and once in person.

---

**e) Explain how unanticipated negative outcomes/experiences or serious adverse events will be managed. (NOTE: This may apply in social-behavioral as well as biomedical research, e.g., undue stress or anxiety of subject, breach of confidentiality via loss of laptop computer with study data. Provisions should be made and described here if applicable.)**

The interventions used in the study have been used in many other settings and consistently found to be safe, but if hygiene promoters or study staff learn of a severe illness or injury that appears related to study activities, they will inform study supervisors who will inform Dr. Mahbubur Rahman at ICDDRB who will collect information on the event, advise appropriate clinical care for an affected participant, and will notify the Principal Investigator. The event will be documented on an adverse event form and will be submitted to the study investigators at ICDDRB and UC Berkeley. The ICDDRB investigator will report the event to both ICDDRB's Ethical Review Committee and the Data Monitoring Committee.

**f) Discuss plans for reporting unanticipated problems involving risks to subjects or others, or serious adverse events, to CPHS. (This applies to all types of research.) See Adverse Event and Unanticipated Problem Reporting.**

Adverse events will be reviewed by the UC Berkeley PI. The event will be reported to CPHS if the event 1) is unexpected; 2) is related or possibly related to study participation; AND 3) suggests that the research places subjects or others at a greater risk of harm than was previously known or recognized. If the event is determined by the PI to be reportable, an initial report will be submitted via email to the Director, Research Subject Projection at CPHS. The initial report will be submitted as soon as possible, but no later than 7 days after the PI learns of the event. The initial report will be followed by a formal written report within 14 days of learning of the incident. The formal report will be submitted to CPHS via eProtocol.

**g) Describe plans for provision of treatment for study-related injuries, and how costs of injury treatment will be covered. If the study involves more than minimal risk, indicate that the researchers are familiar with and will follow University of California policy in this regard, and will use recommended wording on any consent forms (see CPHS Informed Consent Guidelines).**

This study does not involve more than minimal risk. In the LNS arms, health promoters will recommend that caregivers stop using the LNS and notify one of the ICDDRB staff immediately should their child have any adverse reactions shortly after ingesting the supplement (such as vomiting, stomach pain, rash, breathing problems with wheezing). In the event of an adverse reaction, ICDDRB staff will assess the child’s condition and, if necessary, provide transport to the closest medical facility for treatment.

In the anthropometry and enteropathy assessment survey, children who are found to be acutely malnourished based on WHO/Unicef criteria (severely wasted [WHZ < -3] and/or bipedal edema) will be referred to the appropriate existing treatment programs. Children who are found to be infected with intestinal protozoan or helminth parasites will be referred to treatment at the closest health facility. Children with severe anemia (Hb<70 g/L) will be referred for treatment to the closest health facility. Families with children found to have β-thalassemia major will be informed at will be referred to the nearest facility with the capacity for counseling and treatment.

---

---

| | |
|---|---|
| **Protocol Title:** | Measuring the benefits of sanitation, water quality, handwashing and nutrition interventions for improving health and development in rural Bangladesh |
| **Protocol Status:** | APPROVED |
| **Date Submitted:** | 10/26/2017 |
| **Approval Period:** | **10/30/2017-11/03/2018** |
| **Important Note:** | **This Print View may not reflect all comments and contingencies for approval. Please check the comments section of the online protocol. Questions that appear to not have been answered may not have been required for this submission. Please see the system application for more details.** |

#### * * * Benefits, Confidentiality * * *

##### 16. Benefits

**Describe any potential benefits to the individual subject, group of subjects, and/or society. If subjects will not benefit directly from study procedures, this should be stated.**

**NOTE: Do not include compensation/payment of subjects in this section, as remuneration is not considered a "benefit" of participation in research.**

Participants will receive the results of all assessments and referrals to appropriate treatment will be made as necessary. Households in the intervention arms will additionally benefit from free sanitation, handwashing and water quality improvements, and nutrient supplements provided by the study. In the long term, the results of this study could benefit other children in Bangladesh and elsewhere by helping us understand the effects of providing nutrition supplements in combination with WASH interventions.

##### 17. Confidentiality and Privacy

NOTE: See CPHS Data Security Policy and Guidelines before completing this section.

**a) If reviewing or accessing Protected Health Information (PHI) from UC Berkeley's Tang Center, Optometry Clinic, Psychology Clinic, Intercollegiate Athletics, or Human Resources for activities preparatory to research, describe the process and confirm that the health information will not be removed from the "covered entity".**

N/A

**b) What identifiable participant data will you obtain? Note: Audio, photo, and video recordings are generally considered identifiable unless distinguishing features can be successfully masked.**

ICDDR,B field workers will collect participants’ names and dates of birth in order to be able to locate them for follow-up data collection. Such data will be collected for the mothers and the target children, as well as their older siblings and neighboring children included in the baseline assessment. We will also collect GPS coordinates of the location of each participating compound. Photographic images used for presentations and websites will not include personal information such as name or GPS coordinates. Field workers will also collect audio and video files as part of the qualitative assessments. Trained personnel will videotape children to code their behavior before, during, and after the blood draw as part of the child development measures.

**c) If obtaining existing data/specimens, will you have access to identifiers? Please see The Industry Alliance Office website for requirements when receiving existing data/specimens for research.**

UC Berkeley scientists will not have access to identifiers with the exception of images to be used in presentations. The ICDDR,B field workers, field managers, and Project Manager for the study will have access to personal identifiers. These names will only be used to guide survey enumerators to the respondents in follow-up survey rounds (no addresses or phone numbers are collected from household survey respondents). Researchers working with the data will only have numerical identification codes for each entry in the database.

**d) Explain how the confidentiality of subject information will be maintained. Include:**

**i. Who will have access to study records/specimens? If the study is subject to FDA regulations, include a statement that the FDA might inspect the records of the study.**

The ICDDR,B field workers, field managers, and Project Manager for the study will have access to personal identifiers. Lab scientists working with specimens will not have access to personal identifiers. Only trained personnel will have access to videotapes for processing and coding purposes.

**ii. How the records will be secured (e.g., password-protected computer, encrypted files, locked cabinet). Response should be consistent with CPHS Data Security Policy.**

ICDDRB takes a number of precautions to ensure the confidentiality of all information collected from subjects in the studies it conducts. The majority of data will be collected by PDA (personal digital assistant), which minimizes the risk of loss of confidentiality. GPS coordinates will be recorded digitally as well. The cognitive development and environmental enteropathy teams may use paper-based questionnaires. Interviewers will be trained to keep data confidential. Records of phone numbers collected from health promoters and households selected for structured interviews will be stored in a locked room at the research office in Bangladesh, to which only the project manager at ICDDRB will have access. Blood, stool and urine samples will be labeled with the same numerical ID code used on the household surveys. All videotapes will be identified by a number only; no recordings will identify the respondent by name. These videotapes will be viewed only by trained personnel for coding. For audio recordings taken during assessments, the digital output will be transferred to and stored on a secure computer, accessible only to the project manager. The records and videotapes containing identifiers will be stored in field offices in a locked cabinet. The blood, stool and urine samples will be transported to ICDDR,B and stored in a freezer. Access to the freezer room is restricted by a padlock. Samples will be analyzed within 24 months of collection. Identifiable information stored electronically on a removable medium or networked computer will be encrypted, as per CPHS policy.

Data collection instruments, photographs, videotapes, and audiotapes will be stored at ICDDRB after study completion. They will be stored securely under lock and key. The study PI and Project Manager will have access. After study completion, these materials will be digitized, and the hardcopy will be destroyed. Digitized materials will be securely stored for an indefinite period of time.

---

Identifiable information (including audio and video files) stored electronically on a removable medium or networked computer will be password protected and encrypted, as per CPHS policy. Digital copies of the data will not contain any identifying information. Hard copies of the survey forms (if any), including those with subject identifiers, will be stored in locked cabinets at ICDDRB.

**iii. How long study data will be retained.**

Digital copies of the data will be stored indefinitely after the conclusion of the study, for the purposes of additional analysis and informing future research project design. Forms (if any) and digital files containing identifying information will be retained indefinitely for use in a possible follow-up study to assess the long-term health and economic impacts of the interventions. Hard copies of forms which have been digitized will be destroyed at the conclusion of the project. Data resulting from research involving children will be stored for at least 7 years after the child reaches the age of 18 years.

We are interested in holding biological specimens that we collect for 20 years because we envision that these study subjects will be followed in subsequent studies for at least this long so the study population will be a group for whom there is active ongoing scientific interest. The issues that such samples allow us to explore-- how cellular level characteristics influence growth, development and ultimately adult function--are issues where the assays are actively developing in what they can measure. Every year brings substantial breakthroughs. We anticipate that these advances will continue over the next 20 years and permit important insights to be quickly realized without the cost and delay of conducting another decades long longitudinal study. The ICDDR,B WASH Benefits Principal Investigator will be the “gatekeeper” regarding future access and analysis of the stored samples.

**iv. When audio/video recordings will be transcribed and when they will be destroyed (if ever).**

Audiotapes and videotapes will be securely stored for an indefinite period of time. Identifiable information (including audio and video files) stored electronically on a removable medium or networked computer will be password protected and encrypted, as per CPHS policy.

**e) Identifiers should be removed from data/specimens as soon as possible following collection, except in cases where the identifiers are embedded (e.g., voices in audio or faces in video recordings). If data are coded in order to retain a link between the data and identifiable information, explain where the key to the code will be stored, how it will be protected, who will have access to it, and when it will be destroyed.**

Identifiers will be removed from the data at the time that the data is sent out for cleaning and analysis. Identifiers will not be destroyed, but will be stored indefinitely at ICDDRB, as we may return at some future date to evaluate long term intervention effectiveness. All personal identifiers other than the household and child identification codes will be removed from paper-based questionnaires prior to digitization. Researchers working with the data have only numerical identification codes for each entry in the database, while the paper forms with respondent names and corresponding study identification codes are stored in a locked room at the research office in Bangladesh. These names are used only to guide survey enumerators to the respondents in follow-up survey rounds (no addresses or phone numbers are collected from household survey respondents). The key to identifiers will be stored in field offices in a locked cabinet

---

**Protocol Title:** Measuring the benefits of sanitation, water quality, handwashing and nutrition interventions for improving health and development in rural Bangladesh

**Protocol Status:** APPROVED

**Date Submitted:** 10/26/2017

**Approval Period:** 10/30/2017-11/03/2018

**Important Note:** **This Print View may not reflect all comments and contingencies for approval. Please check the comments section of the online protocol. Questions that appear to not have been answered may not have been required for this submission. Please see the system application for more details.**

and/or password protected server. Only study personnel that require the key to complete the study will have access to the locked cabinet. Once the study is completed, the key to identifiers will be stored at ICDDRB in a locked cabinet and/or secure server.

**f) Describe how identifiable data will be transferred (e.g., courier, mail) or transmitted (e.g., file transfer software, file sharing, email). If transmitted via electronic networks, describe how you will secure the data while in transit (e.g., prior encryption). If not applicable, enter N/A.**

Information technology staff directly download the data to desktop computers that are connected to the icddr,b network. The database is password-protected on both PDAs and computers. Data are converted into specified format (STATA, SPSS) and send back to us for error checking and cleaning. Following data cleaning, electronic data are stored in data repository system (DRS) that is connected to the ICDDR,B server and is password-protected as well. Data is encrypted before it is transferred.

**g) Will subjects be asked to give permission for release of identifiable data (e.g., for future studies, publications, presentations, etc.), now or in the future? If so, explain here and include appropriate statements in the consent materials. See Media Records Release Form template for guidance.**

Participants will be asked for permission to use their images in for public presentations and/or on project websites. No other identifiable data will be included within or along with these photos (i.e. no names or other identifiable data). Appropriate statements have been included in consent materials. Special consent will be obtained to allow the use of this material in public presentations and/or public websites.

**h) Explain how subject privacy will be protected (e.g., conducting interviews in a discreet location).**

Field workers will ask to conduct interviews inside the household if possible in order to ensure the privacy of the respondent and the compound members. If non-compound members are present, the field worker will ask them to leave during the interview to ensure the privacy of the respondent. Compound members are often present, but for sensitive questions, field workers will politely ask them to leave so the interview can be conducted in private.

---

---

| | |
|---|---|
| **Protocol Title:** | Measuring the benefits of sanitation, water quality, handwashing and nutrition interventions for improving health and development in rural Bangladesh |
| **Protocol Status:** | APPROVED |
| **Date Submitted:** | 10/26/2017 |
| **Approval Period:** | **10/30/2017-11/03/2018** |
| **Important Note:** | **This Print View may not reflect all comments and contingencies for approval. Please check the comments section of the online protocol. Questions that appear to not have been answered may not have been required for this submission. Please see the system application for more details.** |

#### * * * Potential Financial Conflict of Interest * * *

##### 18. Potential Financial Conflict of Interest

Individuals who have independent roles in projects and who are responsible for the design, analysis, conduct, or reporting of the results of research performed (or to be performed) under a human subjects protocol must disclose whether or not they have a financial interest in or association with the sponsor or the company supplying materials, drugs, or devices for the project. This checklist pertains to the entire project team working under the protocol. Any individual who has a conflict must comply with University regulations and procedures for disclosure of financial conflict of interest.

**See Conflict of Interest Committee Website for more information.**

Please answer the following questions:

Does any member of the project team (defined as UCB or non-UCB personnel working under the protocol) with substantive responsibility for the design, conduct, or reporting of activities under the protocol, or any member of their immediate family (defined as spouse, dependent child or registered domestic partner) have any of the following relationships with the non-UC entity financing the research to be done under the protocol or the non-UC entity supplying materials, drugs or devices being tested under the protocol:

1. N **Positions of management (e.g., board member, scientific advisor, director, officer, partner, trustee, employee, consultant).**
2. N **Equity interest (e.g., stock, stock options, investment, or other ownership).**
3. N **Rights to a pending patent application or issued patent to any invention(s), or license rights or copyright for software that has a direct relationship to the project proposed.**

If the answer to any of the above is Yes, then each individual with any "Yes" response (s) must submit a Human Subjects Financial Conflict of Interest Form DIRECTLY to the Conflict of Interest (COI) Committee for a separate review.

NOTE: When review by the COI Committee is required, CPHS approval of protocols will be contingent upon the disclosure and resolution of all financial conflicts of interest, as determined by the COI Committee.

---

---

**Protocol Title:** Measuring the benefits of sanitation, water quality, handwashing and nutrition interventions for improving health and development in rural Bangladesh

**Protocol Status:** APPROVED

**Date Submitted:** 10/26/2017

**Approval Period:** 10/30/2017-11/03/2018

**Important Note:** This Print View may not reflect all comments and contingencies for approval. Please check the comments section of the online protocol.
Questions that appear to not have been answered may not have been required for this submission. Please see the system application for more details.

#### * * * Informed Consent * * *

##### 19. Informed Consent

Add the consent documents and/or waivers needed for this research using the table at the bottom of the page, including any translated versions. For any translated consent, include an affirmation of the translation's accuracy, indicating who is affirming the accuracy (PI, Co-PI, or Student Investigator), in the Consent/Waiver Description or in the Attachment section. Describe the consent process and provide justification for any waivers for each consent document, translation, and/or waiver. The various consent/waiver options are described below.

Note: DO NOT include child assent documents, parent permission documents or waivers here (these are addressed in the next section).

Altered and Unsigned Consent - A consent document that has omitted required information and does not include a place for a participant's signature. This means that CPHS is being asked to waive one or more elements of consent in addition to the requirement for documented consent.

Altered Consent Form - A consent form that has omitted required information. This means that the CPHS is asked to waive one or more required elements of informed consent. For example, if the purpose of the study will not be disclosed to participants in order to avoid bias, this option should be selected because disclosure of the "purpose" is a required element of informed consent. The form must include a signature line and date line for the individual to sign if he or she agrees to participate.

Consent Form - A standard consent document that embodies all of the required information (elements of informed consent) designed to help an individual make an informed decision about whether or not to participate in the research. The form must include a signature line and date line for the individual to sign if he or she agrees to participate. The Consent Form can also be presented as a "short form" document stating that the required elements of informed consent have been presented orally to the participant. When the short form method is used, a "summary" of the information that is presented to the participant must also be provided for CPHS approval and there must be an impartial witness to the oral presentation. The witness must sign the summary as well as the short form and the participant must sign the summary. The "short form" method may be used in circumstances where oral presentation of consent is preferable or necessary, e.g., subjects are illiterate in English or their native language.

Consent Waiver - No consent will be sought at all. This means that the CPHS is asked to waive the requirement for informed consent. This option is often appropriate for research that involves use of existing data or samples

---

**Protocol Title:** Measuring the benefits of sanitation, water quality, handwashing and nutrition interventions for improving health and development in rural Bangladesh

**Protocol Status:** APPROVED

**Date Submitted:** 10/26/2017

**Approval Period:** 10/30/2017-11/03/2018

**Important Note:** **This Print View may not reflect all comments and contingencies for approval. Please check the comments section of the online protocol. Questions that appear to not have been answered may not have been required for this submission. Please see the system application for more details.**

Unsigned Consent - A document that embodies all of the required information (elements of informed consent), but does not include a place for a participant to indicate with a signature that he or she agrees to take part in the research. This means that the CPHS is asked to waive the requirement for documented (signed) consent. For example, if consent will be obtained verbally or using a button on the web, this option should be selected.

•Informed Consent Guidelines, Templates and Sample Forms

•Informed Consent Policies and Procedures

•Consent Builder: Online Tool for Creating Consent Forms

**Informed Consent**

| Consent/Waiver Description | Consent Type | Consent Document |
|---|---|---|

**Informed Consent**

| | | | |
|---|---|---|---|
| Consent/Waiver Description (e.g. Consent for Group A, Waiver for Group B, Surrogate Consent for Group C) | Consent 1i | | |
| Consent Type | Consent Form | | |
| Attach Consent Document (in Word format) | X | Consent Document | 2011-09-3652_Colford_1i-Consent-Parasite-v21-English |

Explain how, where, when, and by whom informed consent will be obtained. If any vulnerable subject groups are involved, discuss relevant considerations. Note: If attaching multiple consent forms and consent process has already been described for another consent form, simply refer to the other form (e.g., consent process is the same as for Group A).

ICDDR,B field workers will approach eligible compounds and ask to speak with the pregnant woman/mother and head of compound. The study will be described, and participant involvement described. If the compound agrees to participate, the consent process will be completed. If a potential respondent is not able to read the consent form, field workers will suggest that they invite a witness to help read the form with them so that they can be certain that they agree to participate in the procedures in the study. If a witness is present, both the witness and respondent will be asked to provide a thumb print/signature.

| | | | |
|---|---|---|---|
| Consent/Waiver Description (e.g. Consent for Group A, Waiver for Group B, Surrogate Consent for Group C) | Consent 2i - Bengali | | |
| Consent Type | Consent Form | | |
| Attach Consent Document (in Word format) | X | Consent Document | 2011-09-3652_Colford_2i-Consent-Parasite-v21-Bengali |

---

Explain how, where, when, and by whom informed consent will be obtained. If any vulnerable subject groups are involved, discuss relevant considerations. Note: If attaching multiple consent forms and consent process has already been described for another consent form, simply refer to the other form (e.g., consent process is the same as for Group A).

ICDDR,B field workers will approach eligible compounds and ask to speak with the pregnant woman/mother and head of compound. The study will be described, and participant involvement described. If the compound agrees to participate, the consent process will be completed.

| | |
|---|---|
| Consent/Waiver Description (e.g. Consent for Group A, Waiver for Group B, Surrogate Consent for Group C) | Consent 1k1 |

| Consent Type | Consent Form | | |
|---|---|---|---|
| Attach Consent Document (in Word format) | X | Consent Document | 2011-09-3652_Colford_1k1-Consent-EnvAssess-v21-English |

Explain how, where, when, and by whom informed consent will be obtained. If any vulnerable subject groups are involved, discuss relevant considerations. Note: If attaching multiple consent forms and consent process has already been described for another consent form, simply refer to the other form (e.g., consent process is the same as for Group A).

ICDDR,B field workers will approach eligible compounds and ask to speak with the pregnant woman/mother and head of compound. The study will be described, and participant involvement described. If the compound agrees to participate, the consent process will be completed.

| | |
|---|---|
| Consent/Waiver Description (e.g. Consent for Group A, Waiver for Group B, Surrogate Consent for Group C) | Consent 2k1 - Bengali |

| Consent Type | Consent Form | | |
|---|---|---|---|
| Attach Consent Document (in Word format) | X | Consent Document | 2011-09-3652_Colford_2k1-Consent-EnvAssess-v19-Bengali |

Explain how, where, when, and by whom informed consent will be obtained. If any vulnerable subject groups are involved, discuss relevant considerations. Note: If attaching multiple consent forms and consent process has already been described for another consent form, simply refer to the other form (e.g., consent process is the same as for Group A).

ICDDR,B field workers will approach eligible compounds and ask to speak with the pregnant woman/mother and head of compound. The study will be described, and participant involvement described. If the compound agrees to participate, the consent process will be completed.

---

| | |
|---|---|
| **Protocol Title:** | Measuring the benefits of sanitation, water quality, handwashing and nutrition interventions for improving health and development in rural Bangladesh |
| **Protocol Status:** | APPROVED |
| **Date Submitted:** | 10/26/2017 |
| **Approval Period:** | **10/30/2017-11/03/2018** |
| **Important Note:** | **This Print View may not reflect all comments and contingencies for approval. Please check the comments section of the online protocol. Questions that appear to not have been answered may not have been required for this submission. Please see the system application for more details.** |

| | | | |
|---|---|---|---|
| Consent/Waiver Description (e.g. Consent for Group A, Waiver for Group B, Surrogate Consent for Group C) | Consent 1k3 | | |
| Consent Type | Consent Form | | |
| Attach Consent Document (in Word format) | X | Consent Document | 2011-09-3652_Colford_1k3-Consent-EnvAssess-hygiene-v21-English |

Explain how, where, when, and by whom informed consent will be obtained. If any vulnerable subject groups are involved, discuss relevant considerations. Note: If attaching multiple consent forms and consent process has already been described for another consent form, simply refer to the other form (e.g., consent process is the same as for Group A).

ICDDR,B field workers will approach eligible compounds and ask to speak with the pregnant woman/mother and head of compound. The study will be described, and participant involvement described. If the compound agrees to participate, the consent process will be completed.

| | | | |
|---|---|---|---|
| Consent/Waiver Description (e.g. Consent for Group A, Waiver for Group B, Surrogate Consent for Group C) | Consent 2k3 - Bengali | | |
| Consent Type | Consent Form | | |
| Attach Consent Document (in Word format) | X | Consent Document | 2011-09-3652_Colford_2k3-Consent-EnvAssess-hygiene-v19-Bengali |

Explain how, where, when, and by whom informed consent will be obtained. If any vulnerable subject groups are involved, discuss relevant considerations. Note: If attaching multiple consent forms and consent process has already been described for another consent form, simply refer to the other form (e.g., consent process is the same as for Group A).

ICDDR,B field workers will approach eligible compounds and ask to speak with the pregnant woman/mother and head of compound. The study will be described, and participant involvement described. If the compound agrees to participate, the consent process will be completed.

| | | | |
|---|---|---|---|
| Consent/Waiver Description (e.g. Consent for Group A, Waiver for Group B, Surrogate Consent for Group C) | Consent 1l | | |
| Consent Type | Consent Form | | |
| Attach Consent Document (in Word format) | X | Consent Document | 2011-09-3652_Colford_1l-Consent-5hr-SO-v18-English |

Explain how, where, when, and by whom informed consent will be obtained. If any vulnerable subject groups are involved, discuss relevant considerations. Note: If attaching multiple consent forms and consent process has already been described for another consent form, simply refer to the other form (e.g., consent process is the same as for Group A).

---

| | |
|---|---|
| **Protocol Title:** | Measuring the benefits of sanitation, water quality, handwashing and nutrition interventions for improving health and development in rural Bangladesh |
| **Protocol Status:** | APPROVED |
| **Date Submitted:** | 10/26/2017 |
| **Approval Period:** | **10/30/2017-11/03/2018** |
| **Important Note:** | **This Print View may not reflect all comments and contingencies for approval. Please check the comments section of the online protocol. Questions that appear to not have been answered may not have been required for this submission. Please see the system application for more details.** |

ICDDR,B field workers will approach eligible compounds and ask to speak with the pregnant woman/mother and head of compound. The study will be described, and participant involvement described. If the compound agrees to participate, the consent process will be completed.

| | | | |
|---|---|---|---|
| Consent/Waiver Description (e.g. Consent for Group A, Waiver for Group B, Surrogate Consent for Group C) | Consent 2l - Bengali | | |
| Consent Type | Consent Form | | |
| Attach Consent Document (in Word format) | X | Consent Document | 2011-09-3652_Colford_2l-Consent-5hr-SO-v18-Bengali |

Explain how, where, when, and by whom informed consent will be obtained. If any vulnerable subject groups are involved, discuss relevant considerations. Note: If attaching multiple consent forms and consent process has already been described for another consent form, simply refer to the other form (e.g., consent process is the same as for Group A).

ICDDR,B field workers will approach eligible compounds and ask to speak with the pregnant woman/mother and head of compound. The study will be described, and participant involvement described. If the compound agrees to participate, the consent process will be completed.

| | | | |
|---|---|---|---|
| Consent/Waiver Description (e.g. Consent for Group A, Waiver for Group B, Surrogate Consent for Group C) | Consent 1n | | |
| Consent Type | Consent Form | | |
| Attach Consent Document (in Word format) | X | Consent Document | 2011-09-3652_Colford_1n-Consent-Lead exposure-v17-English |

Explain how, where, when, and by whom informed consent will be obtained. If any vulnerable subject groups are involved, discuss relevant considerations. Note: If attaching multiple consent forms and consent process has already been described for another consent form, simply refer to the other form (e.g., consent process is the same as for Group A).

ICDDR,B field workers will approach eligible compounds and ask to speak with the pregnant woman/mother and head of compound. The study will be described, and participant involvement described. If the compound agrees to participate, the consent process will be completed.

| | |
|---|---|
| Consent/Waiver Description (e.g. Consent for Group A, Waiver for Group B, Surrogate Consent for Group C) | Consent 2n - Bengali |
| Consent Type | Consent Form |

---

**Protocol Title:** Measuring the benefits of sanitation, water quality, handwashing and nutrition interventions for improving health and development in rural Bangladesh

**Protocol Status:** APPROVED

**Date Submitted:** 10/26/2017

**Approval Period:** **10/30/2017-11/03/2018**

**Important Note:** **This Print View may not reflect all comments and contingencies for approval. Please check the comments section of the online protocol. Questions that appear to not have been answered may not have been required for this submission. Please see the system application for more details.**

| | | | |
|---|---|---|---|
| Attach Consent Document (in Word format) | X | Consent Document | 2011-09-3652_Colford_2n-Consent-Lead exposure-v17-Bengali |

Explain how, where, when, and by whom informed consent will be obtained. If any vulnerable subject groups are involved, discuss relevant considerations. Note: If attaching multiple consent forms and consent process has already been described for another consent form, simply refer to the other form (e.g., consent process is the same as for Group A).

ICDDR,B field workers will approach eligible compounds and ask to speak with the pregnant woman/mother and head of compound. The study will be described, and participant involvement described. If the compound agrees to participate, the consent process will be completed.

| | | | |
|---|---|---|---|
| Consent/Waiver Description (e.g. Consent for Group A, Waiver for Group B, Surrogate Consent for Group C) | Consent 1o | | |
| Consent Type | Consent Form | | |
| Attach Consent Document (in Word format) | X | Consent Document | 2011-09-3652_Colford_1o-Consent-Agrochemical-v17-English |

Explain how, where, when, and by whom informed consent will be obtained. If any vulnerable subject groups are involved, discuss relevant considerations. Note: If attaching multiple consent forms and consent process has already been described for another consent form, simply refer to the other form (e.g., consent process is the same as for Group A).

ICDDR,B field workers will approach eligible compounds and ask to speak with the pregnant woman/mother and head of compound. The study will be described, and participant involvement described. If the compound agrees to participate, the consent process will be completed.

| | | | |
|---|---|---|---|
| Consent/Waiver Description (e.g. Consent for Group A, Waiver for Group B, Surrogate Consent for Group C) | Consent 2o - Bengali | | |
| Consent Type | Consent Form | | |
| Attach Consent Document (in Word format) | X | Consent Document | 2011-09-3652_Colford_2o-Consent-Agrochemical-v17-Bengali |

Explain how, where, when, and by whom informed consent will be obtained. If any vulnerable subject groups are involved, discuss relevant considerations. Note: If attaching multiple consent forms and consent process has already been described for another consent form, simply refer to the other form (e.g., consent process is the same as for Group A).

ICDDR,B field workers will approach eligible compounds and ask to speak with the pregnant woman/mother and head of compound. The study will be described, and participant involvement described. If the compound agrees to participate, the consent process will be completed.

---

| | |
|---|---|
| **Protocol Title:** | Measuring the benefits of sanitation, water quality, handwashing and nutrition interventions for improving health and development in rural Bangladesh |
| **Protocol Status:** | APPROVED |
| **Date Submitted:** | 10/26/2017 |
| **Approval Period:** | **10/30/2017-11/03/2018** |
| **Important Note:** | **This Print View may not reflect all comments and contingencies for approval. Please check the comments section of the online protocol. Questions that appear to not have been answered may not have been required for this submission. Please see the system application for more details.** |

| | | | |
|---|---|---|---|
| Consent/Waiver Description (e.g. Consent for Group A, Waiver for Group B, Surrogate Consent for Group C) | Consent 1p | | |
| Consent Type | Consent Form | | |
| Attach Consent Document (in Word format) | X | Consent Document | 2011-09-3652_Colford_1p-Consent-Lead case control-v17-English |

Explain how, where, when, and by whom informed consent will be obtained. If any vulnerable subject groups are involved, discuss relevant considerations. Note: If attaching multiple consent forms and consent process has already been described for another consent form, simply refer to the other form (e.g., consent process is the same as for Group A).

ICDDR,B field workers will approach eligible compounds and ask to speak with the pregnant woman/mother and head of compound. The study will be described, and participant involvement described. If the compound agrees to participate, the consent process will be completed.

| | | | |
|---|---|---|---|
| Consent/Waiver Description (e.g. Consent for Group A, Waiver for Group B, Surrogate Consent for Group C) | Consent 2p - Bengali | | |
| Consent Type | Consent Form | | |
| Attach Consent Document (in Word format) | X | Consent Document | 2011-09-3652_Colford_2p-Consent-Lead case control-v17-Bengali |

Explain how, where, when, and by whom informed consent will be obtained. If any vulnerable subject groups are involved, discuss relevant considerations. Note: If attaching multiple consent forms and consent process has already been described for another consent form, simply refer to the other form (e.g., consent process is the same as for Group A).

ICDDR,B field workers will approach eligible compounds and ask to speak with the pregnant woman/mother and head of compound. The study will be described, and participant involvement described. If the compound agrees to participate, the consent process will be completed.

| | | | |
|---|---|---|---|
| Consent/Waiver Description (e.g. Consent for Group A, Waiver for Group B, Surrogate Consent for Group C) | Consent 1r | | |
| Consent Type | Consent Form | | |
| Attach Consent Document (in Word format) | X | Consent Document | 2011-09-3652_Colford_1r-Consent-Structured observation with video-v17-English |

---

**Protocol Title:** Measuring the benefits of sanitation, water quality, handwashing and nutrition interventions for improving health and development in rural Bangladesh

**Protocol Status:** APPROVED

**Date Submitted:** 10/26/2017

**Approval Period:** 10/30/2017-11/03/2018

**Important Note:** **This Print View may not reflect all comments and contingencies for approval. Please check the comments section of the online protocol. Questions that appear to not have been answered may not have been required for this submission. Please see the system application for more details.**

Explain how, where, when, and by whom informed consent will be obtained. If any vulnerable subject groups are involved, discuss relevant considerations. Note: If attaching multiple consent forms and consent process has already been described for another consent form, simply refer to the other form (e.g., consent process is the same as for Group A).

ICDDR,B field workers will approach eligible compounds and ask to speak with the pregnant woman/mother and head of compound. The study will be described, and participant involvement described. If the compound agrees to participate, the consent process will be completed.

| | | | |
|---|---|---|---|
| Consent/Waiver Description (e.g. Consent for Group A, Waiver for Group B, Surrogate Consent for Group C) | Consent 2r - Bengali | | |
| Consent Type | Consent Form | | |
| Attach Consent Document (in Word format) | X | Consent Document | 2011-09-3652_Colford_2r-Consent-Structured observation with video-v17-Bengali |

Explain how, where, when, and by whom informed consent will be obtained. If any vulnerable subject groups are involved, discuss relevant considerations. Note: If attaching multiple consent forms and consent process has already been described for another consent form, simply refer to the other form (e.g., consent process is the same as for Group A).

ICDDR,B field workers will approach eligible compounds and ask to speak with the pregnant woman/mother and head of compound. The study will be described, and participant involvement described. If the compound agrees to participate, the consent process will be completed.

| | | | |
|---|---|---|---|
| Consent/Waiver Description (e.g. Consent for Group A, Waiver for Group B, Surrogate Consent for Group C) | Consent 1s | | |
| Consent Type | Consent Form | | |
| Attach Consent Document (in Word format) | X | Consent Document | 2011-09-3652_Colford_1s-Consent-In depth interview environmental assessment-v17-English |

Explain how, where, when, and by whom informed consent will be obtained. If any vulnerable subject groups are involved, discuss relevant considerations. Note: If attaching multiple consent forms and consent process has already been described for another consent form, simply refer to the other form (e.g., consent process is the same as for Group A).

ICDDR,B field workers will approach eligible compounds and ask to speak with the pregnant woman/mother and head of compound. The study will be described, and participant involvement described. If the compound agrees to participate, the consent process will be completed.

---

**Protocol Title:** Measuring the benefits of sanitation, water quality, handwashing and nutrition interventions for improving health and development in rural Bangladesh

**Protocol Status:** APPROVED

**Date Submitted:** 10/26/2017

**Approval Period:** **10/30/2017-11/03/2018**

**Important Note:** **This Print View may not reflect all comments and contingencies for approval. Please check the comments section of the online protocol. Questions that appear to not have been answered may not have been required for this submission. Please see the system application for more details.**

| | | | |
|---|---|---|---|
| Consent/Waiver Description (e.g. Consent for Group A, Waiver for Group B, Surrogate Consent for Group C) | Consent 2s - Bengali | | |
| Consent Type | Consent Form | | |
| Attach Consent Document (in Word format) | X | Consent Document | 2011-09-3652_Colford_2s-Consent-In depth interview environmental assessment-v17-Bengali |

Explain how, where, when, and by whom informed consent will be obtained. If any vulnerable subject groups are involved, discuss relevant considerations. Note: If attaching multiple consent forms and consent process has already been described for another consent form, simply refer to the other form (e.g., consent process is the same as for Group A).

ICDDR,B field workers will approach eligible compounds and ask to speak with the pregnant woman/mother and head of compound. The study will be described, and participant involvement described. If the compound agrees to participate, the consent process will be completed.

| | | | |
|---|---|---|---|
| Consent/Waiver Description (e.g. Consent for Group A, Waiver for Group B, Surrogate Consent for Group C) | Consent 1t | | |
| Consent Type | Consent Form | | |
| Attach Consent Document (in Word format) | X | Consent Document | 2011-09-3652_Colford_1t-Consent-Focus-Group-Discussion-v17-English |

Explain how, where, when, and by whom informed consent will be obtained. If any vulnerable subject groups are involved, discuss relevant considerations. Note: If attaching multiple consent forms and consent process has already been described for another consent form, simply refer to the other form (e.g., consent process is the same as for Group A).

ICDDR,B field workers will approach eligible compounds and ask to speak with the pregnant woman/mother and head of compound. The study will be described, and participant involvement described. If the compound agrees to participate, the consent process will be completed.

| | | | |
|---|---|---|---|
| Consent/Waiver Description (e.g. Consent for Group A, Waiver for Group B, Surrogate Consent for Group C) | Consent 2t - Bengali | | |
| Consent Type | Consent Form | | |
| Attach Consent Document (in Word format) | X | Consent Document | 2011-09-3652_Colford_2t-Consent-Focus-Group-Discussion-v17-Bengali |

---

| | |
|---|---|
| **Protocol Title:** | Measuring the benefits of sanitation, water quality, handwashing and nutrition interventions for improving health and development in rural Bangladesh |
| **Protocol Status:** | APPROVED |
| **Date Submitted:** | 10/26/2017 |
| **Approval Period:** | **10/30/2017-11/03/2018** |
| **Important Note:** | **This Print View may not reflect all comments and contingencies for approval. Please check the comments section of the online protocol. Questions that appear to not have been answered may not have been required for this submission. Please see the system application for more details.** |

Explain how, where, when, and by whom informed consent will be obtained. If any vulnerable subject groups are involved, discuss relevant considerations. Note: If attaching multiple consent forms and consent process has already been described for another consent form, simply refer to the other form (e.g., consent process is the same as for Group A).

ICDDR,B field workers will approach eligible compounds and ask to speak with the pregnant woman/mother and head of compound. The study will be described, and participant involvement described. If the compound agrees to participate, the consent process will be completed.

| | | | |
|---|---|---|---|
| Consent/Waiver Description (e.g. Consent for Group A, Waiver for Group B, Surrogate Consent for Group C) | Consent 1u | | |
| Consent Type | Consent Form | | |
| Attach Consent Document (in Word format) | X | Consent Document | 2011-09-3652_Colford_1u-Consent-Parasites-v21-English |

Explain how, where, when, and by whom informed consent will be obtained. If any vulnerable subject groups are involved, discuss relevant considerations. Note: If attaching multiple consent forms and consent process has already been described for another consent form, simply refer to the other form (e.g., consent process is the same as for Group A).

ICDDR,B field workers will approach eligible compounds and ask to speak with the pregnant woman/mother and head of compound. The study will be described, and participant involvement described. If the compound agrees to participate, the consent process will be completed. If a potential respondent is not able to read the consent form, field workers will suggest that they invite a witness to help read the form with them so that they can be certain that they agree to participate in the procedures in the study. If a witness is present, both the witness and respondent will be asked to provide a thumb print/signature.

| | | | |
|---|---|---|---|
| Consent/Waiver Description (e.g. Consent for Group A, Waiver for Group B, Surrogate Consent for Group C) | Consent 2u-Bengali | | |
| Consent Type | Consent Form | | |
| Attach Consent Document (in Word format) | X | Consent Document | 2011-09-3652_Colford_2u-Consent-Longitudinal-EnvAssess-v21-Bengali |

Explain how, where, when, and by whom informed consent will be obtained. If any vulnerable subject groups are involved, discuss relevant considerations. Note: If attaching multiple consent forms and consent process has already been described for another consent form, simply refer to the other form (e.g., consent process is the same as for Group A).

ICDDR,B field workers will approach eligible compounds and ask to speak with the pregnant woman/mother and head of compound. The study will be described, and participant involvement described. If the compound agrees to participate, the consent process will be completed.

---

| | |
|---|---|
| **Protocol Title:** | Measuring the benefits of sanitation, water quality, handwashing and nutrition interventions for improving health and development in rural Bangladesh |
| **Protocol Status:** | APPROVED |
| **Date Submitted:** | 10/26/2017 |
| **Approval Period:** | **10/30/2017-11/03/2018** |
| **Important Note:** | **This Print View may not reflect all comments and contingencies for approval. Please check the comments section of the online protocol. Questions that appear to not have been answered may not have been required for this submission. Please see the system application for more details.** |

| | | | |
|---|---|---|---|
| Consent/Waiver Description (e.g. Consent for Group A, Waiver for Group B, Surrogate Consent for Group C) | Consent 1k2 | | |
| Consent Type | Consent Form | | |
| Attach Consent Document (in Word format) | X | Consent Document | 2011-09-3652_Colford_1k2-Consent-EnvAssess-water-v21-English |

Explain how, where, when, and by whom informed consent will be obtained. If any vulnerable subject groups are involved, discuss relevant considerations. Note: If attaching multiple consent forms and consent process has already been described for another consent form, simply refer to the other form (e.g., consent process is the same as for Group A).

ICDDR,B field workers will approach eligible compounds and ask to speak with the pregnant woman/mother and head of compound. The study will be described, and participant involvement described. If the compound agrees to participate, the consent process will be completed.

| | | | |
|---|---|---|---|
| Consent/Waiver Description (e.g. Consent for Group A, Waiver for Group B, Surrogate Consent for Group C) | Consent 2k2 - Bengali | | |
| Consent Type | Consent Form | | |
| Attach Consent Document (in Word format) | X | Consent Document | 2011-09-3652_Colford_2k2-Consent-EnvAssess-water-v19-Bengali |

Explain how, where, when, and by whom informed consent will be obtained. If any vulnerable subject groups are involved, discuss relevant considerations. Note: If attaching multiple consent forms and consent process has already been described for another consent form, simply refer to the other form (e.g., consent process is the same as for Group A).

ICDDR,B field workers will approach eligible compounds and ask to speak with the pregnant woman/mother and head of compound. The study will be described, and participant involvement described. If the compound agrees to participate, the consent process will be completed.

| | | | |
|---|---|---|---|
| Consent/Waiver Description (e.g. Consent for Group A, Waiver for Group B, Surrogate Consent for Group C) | Consent 1v | | |
| Consent Type | Consent Form | | |
| Attach Consent Document (in Word format) | X | Consent Document | 2011-09-3652_Colford_1v-Child cognitive development-v18-English |

---

| | |
|---|---|
| **Protocol Title:** | Measuring the benefits of sanitation, water quality, handwashing and nutrition interventions for improving health and development in rural Bangladesh |
| **Protocol Status:** | APPROVED |
| **Date Submitted:** | 10/26/2017 |
| **Approval Period:** | **10/30/2017-11/03/2018** |
| **Important Note:** | **This Print View may not reflect all comments and contingencies for approval. Please check the comments section of the online protocol. Questions that appear to not have been answered may not have been required for this submission. Please see the system application for more details.** |

Explain how, where, when, and by whom informed consent will be obtained. If any vulnerable subject groups are involved, discuss relevant considerations. Note: If attaching multiple consent forms and consent process has already been described for another consent form, simply refer to the other form (e.g., consent process is the same as for Group A).

ICDDR,B field workers will approach eligible compounds and ask to speak with the pregnant woman/mother and head of compound. The study will be described, and participant involvement described. If the compound agrees to participate, the consent process will be completed.

| | | | |
|---|---|---|---|
| Consent/Waiver Description (e.g. Consent for Group A, Waiver for Group B, Surrogate Consent for Group C) | Consent 2v-Bengali | | |
| Consent Type | Consent Form | | |
| Attach Consent Document (in Word format) | X | Consent Document | 2011-09-3652_Colford_2v-Child cognitive development-v18-Bengali |

Explain how, where, when, and by whom informed consent will be obtained. If any vulnerable subject groups are involved, discuss relevant considerations. Note: If attaching multiple consent forms and consent process has already been described for another consent form, simply refer to the other form (e.g., consent process is the same as for Group A).

ICDDR,B field workers will approach eligible compounds and ask to speak with the pregnant woman/mother and head of compound. The study will be described, and participant involvement described. If the compound agrees to participate, the consent process will be completed.

| | | | |
|---|---|---|---|
| Consent/Waiver Description (e.g. Consent for Group A, Waiver for Group B, Surrogate Consent for Group C) | Consent 1w | | |
| Consent Type | Consent Form | | |
| Attach Consent Document (in Word format) | X | Consent Document | 2011-09-3652_Colford_1w-Consent-Spillover compounds-v19-English |

Explain how, where, when, and by whom informed consent will be obtained. If any vulnerable subject groups are involved, discuss relevant considerations. Note: If attaching multiple consent forms and consent process has already been described for another consent form, simply refer to the other form (e.g., consent process is the same as for Group A).

ICDDR,B field workers will approach eligible compounds and ask to speak with the pregnant woman/mother and head of compound. The study will be described, and participant involvement described. If the compound agrees to participate, the consent process will be completed.

---

| | |
|---|---|
| **Protocol Title:** | Measuring the benefits of sanitation, water quality, handwashing and nutrition interventions for improving health and development in rural Bangladesh |
| **Protocol Status:** | APPROVED |
| **Date Submitted:** | 10/26/2017 |
| **Approval Period:** | **10/30/2017-11/03/2018** |
| **Important Note:** | **This Print View may not reflect all comments and contingencies for approval. Please check the comments section of the online protocol. Questions that appear to not have been answered may not have been required for this submission. Please see the system application for more details.** |

| | | | |
|---|---|---|---|
| Consent/Waiver Description (e.g. Consent for Group A, Waiver for Group B, Surrogate Consent for Group C) | Consent 2w - Bengali | | |
| Consent Type | Consent Form | | |
| Attach Consent Document (in Word format) | X | Consent Document | 2011-09-3652_Colford_2w-Consent-Spillover compounds-v19-Bengali |

Explain how, where, when, and by whom informed consent will be obtained. If any vulnerable subject groups are involved, discuss relevant considerations. Note: If attaching multiple consent forms and consent process has already been described for another consent form, simply refer to the other form (e.g., consent process is the same as for Group A).

ICDDR,B field workers will approach eligible compounds and ask to speak with the pregnant woman/mother and head of compound. The study will be described, and participant involvement described. If the compound agrees to participate, the consent process will be completed.

| | | | |
|---|---|---|---|
| Consent/Waiver Description (e.g. Consent for Group A, Waiver for Group B, Surrogate Consent for Group C) | Consent 1j | | |
| Consent Type | Consent Form | | |
| Attach Consent Document (in Word format) | X | Consent Document | 2011-09-3652_Colford_1j-Consent-EE-v21-English |

Explain how, where, when, and by whom informed consent will be obtained. If any vulnerable subject groups are involved, discuss relevant considerations. Note: If attaching multiple consent forms and consent process has already been described for another consent form, simply refer to the other form (e.g., consent process is the same as for Group A).

ICDDR,B field workers will approach eligible compounds and ask to speak with the pregnant woman/mother and head of compound. The study will be described, and participant involvement described. If the compound agrees to participate, the consent process will be completed. If a potential respondent is not able to read the consent form, field workers will suggest that they invite a witness to help read the form with them so that they can be certain that they agree to participate in the procedures in the study. If a witness is present, both the witness and respondent will be asked to provide a thumb print/signature.

| | | | |
|---|---|---|---|
| Consent/Waiver Description (e.g. Consent for Group A, Waiver for Group B, Surrogate Consent for Group C) | Consent 2j - Bengali | | |
| Consent Type | Consent Form | | |
| Attach Consent Document (in Word format) | X | Consent Document | 2011-09-3652_Colford_1j-Consent-EE-v21-English |

---

| | |
|---|---|
| **Protocol Title:** | Measuring the benefits of sanitation, water quality, handwashing and nutrition interventions for improving health and development in rural Bangladesh |
| **Protocol Status:** | APPROVED |
| **Date Submitted:** | 10/26/2017 |
| **Approval Period:** | **10/30/2017-11/03/2018** |
| **Important Note:** | **This Print View may not reflect all comments and contingencies for approval. Please check the comments section of the online protocol. Questions that appear to not have been answered may not have been required for this submission. Please see the system application for more details.** |

Explain how, where, when, and by whom informed consent will be obtained. If any vulnerable subject groups are involved, discuss relevant considerations. Note: If attaching multiple consent forms and consent process has already been described for another consent form, simply refer to the other form (e.g., consent process is the same as for Group A).

ICDDR,B field workers will approach eligible compounds and ask to speak with the pregnant woman/mother and head of compound. The study will be described, and participant involvement described. If the compound agrees to participate, the consent process will be completed.

Consent/Waiver Description (e.g. Consent for Group A, Waiver for Group B, Surrogate Consent for Group C) — Consent 1z

| Consent Type | Consent Form | | |
|---|---|---|---|
| Attach Consent Document (in Word format) | X | Consent Document | 2011-09-3652_Colford_1z-Consent-Microbiome-v20-English |

Explain how, where, when, and by whom informed consent will be obtained. If any vulnerable subject groups are involved, discuss relevant considerations. Note: If attaching multiple consent forms and consent process has already been described for another consent form, simply refer to the other form (e.g., consent process is the same as for Group A).

ICDDR,B field workers will approach eligible compounds and ask to speak with the pregnant woman/mother and head of compound. The study will be described, and participant involvement described. If the compound agrees to participate, the consent process will be completed. If a potential respondent is not able to read the consent form, field workers will suggest that they invite a witness to help read the form with them so that they can be certain that they agree to participate in the procedures in the study. If a witness is present, both the witness and respondent will be asked to provide a thumb print/signature.

Consent/Waiver Description (e.g. Consent for Group A, Waiver for Group B, Surrogate Consent for Group C) — Consent 2z - Bengali

| Consent Type | Consent Form | | |
|---|---|---|---|
| Attach Consent Document (in Word format) | X | Consent Document | 2011-09-3652_Colford_2z-Consent-Microbiome-v20-Bengali |

Explain how, where, when, and by whom informed consent will be obtained. If any vulnerable subject groups are involved, discuss relevant considerations. Note: If attaching multiple consent forms and consent process has already been described for another consent form, simply refer to the other form (e.g., consent process is the same as for Group A).

ICDDR,B field workers will approach eligible compounds and ask to speak with the pregnant woman/mother and head of compound. The study will be described, and participant involvement described. If the compound agrees to participate, the consent process will be completed.

---

| | |
|---|---|
| **Protocol Title:** | Measuring the benefits of sanitation, water quality, handwashing and nutrition interventions for improving health and development in rural Bangladesh |
| **Protocol Status:** | APPROVED |
| **Date Submitted:** | 10/26/2017 |
| **Approval Period:** | **10/30/2017-11/03/2018** |
| **Important Note:** | **This Print View may not reflect all comments and contingencies for approval. Please check the comments section of the online protocol. Questions that appear to not have been answered may not have been required for this submission. Please see the system application for more details.** |

| | | | |
|---|---|---|---|
| Consent/Waiver Description (e.g. Consent for Group A, Waiver for Group B, Surrogate Consent for Group C) | Consent 1aa | | |
| Consent Type | Consent Form | | |
| Attach Consent Document (in Word format) | X | Consent Document | 2011-09-3652_Colford_1aa-Consent-Spillover-qualitative-v21-English |

Explain how, where, when, and by whom informed consent will be obtained. If any vulnerable subject groups are involved, discuss relevant considerations. Note: If attaching multiple consent forms and consent process has already been described for another consent form, simply refer to the other form (e.g., consent process is the same as for Group A).

ICDDR,B field workers will approach eligible compounds and ask to speak with the pregnant woman/mother and head of compound. The study will be described, and participant involvement described. If the compound agrees to participate, the consent process will be completed.

| | | | |
|---|---|---|---|
| Consent/Waiver Description (e.g. Consent for Group A, Waiver for Group B, Surrogate Consent for Group C) | Consent 2aa - Bengali | | |
| Consent Type | Consent Form | | |
| Attach Consent Document (in Word format) | X | Consent Document | 2011-09-3652_Colford_2aa-Consent-Spillover-qualitative-Bengali |

Explain how, where, when, and by whom informed consent will be obtained. If any vulnerable subject groups are involved, discuss relevant considerations. Note: If attaching multiple consent forms and consent process has already been described for another consent form, simply refer to the other form (e.g., consent process is the same as for Group A).

ICDDR,B field workers will approach eligible compounds and ask to speak with the pregnant woman/mother and head of compound. The study will be described, and participant involvement described. If the compound agrees to participate, the consent process will be completed.

| | |
|---|---|
| Consent/Waiver Description (e.g. Consent for Group A, Waiver for Group B, Surrogate Consent for Group C) | Waiver for people named in spillover qualitative assessment |
| Consent Type | Consent Waiver |

For the CPHS to approve a waiver of one or more elements of informed consent, either criterion A or B must be met. Select the applicable criterion and provide justification in the box below.

Y **A.** (1) The research involves no more than minimal risk of harm to the subjects;

---

(2) The waiver or alteration will not adversely affect the rights and welfare of the subjects;

(3) The research could not practicably be carried out without the waiver or alteration; and

(4) Whenever appropriate, the subjects will be provided with pertinent information after participation.

In order to assess spillover effects via social networks, we would like to collect information about who WASH Benefits participants and their neighbors are closely socially connected to. In order to follow best practices used in the field of social network analysis, we plan to implement a name generator module in the qualitative assessment. This is a standard practice to understand diffusion of interventions through networks. If we do not ask for names but rather for other identifying information, it would be nearly impossible for us to determine whether WASH Benefits participants and their neighbors were closely socially connected. Thus, we feel that we would not be able to achieve our scientific objectives without asking for names. The information will be kept private and will not be shared with anyone outside our research team. We feel that there is minimal risk associated with this activity.

**B.** (1) The research or demonstration project is to be conducted by or subject to the approval of state or local officials and is designed to study, evaluate, or otherwise examine: (i) public benefit or service programs; (ii) procedures for obtaining benefits or service; (iii) possible changes in or alternatives to those programs or procedures; or (iv) possible changes in methods or levels of payment for benefits or services under those programs;and

(2) The research could not practicably be carried out without the waiver or alteration.

---

---

| | |
|---|---|
| **Protocol Title:** | Measuring the benefits of sanitation, water quality, handwashing and nutrition interventions for improving health and development in rural Bangladesh |
| **Protocol Status:** | APPROVED |
| **Date Submitted:** | 10/26/2017 |
| **Approval Period:** | **10/30/2017-11/03/2018** |
| **Important Note:** | **This Print View may not reflect all comments and contingencies for approval. Please check the comments section of the online protocol. Questions that appear to not have been answered may not have been required for this submission. Please see the system application for more details.** |

#### * * * Child Assent & Parent Permission * * *

##### 20. Child Assent and Parent/Guardian Permission

Add each child assent document, parent/guardian permission document, and/or waiver needed for this research using the table at the bottom of the page, including any translated versions. For any translated consent, include an affirmation of the translation's accuracy, indicating who is affirming the accuracy (PI, Co-PI, or Student Investigator), in the Consent/Waiver Description or in the Attachment section. Describe the consent process and provide justification for any waivers for each consent document, translation, and/or waiver. The various consent/waiver options are described below.

Altered and Unsigned Parent/Guardian Permission Form - A parent permission document that has omitted required information (elements) and does not include a place for a parent to indicate with a signature that he or she agrees to permit the child's participation. This means that CPHS is being asked to waive one or more elements of consent in addition to the requirement for documented consent.

Altered Parent/Guardian Permission Form - A permission form that has omitted required information (elements). This means that the CPHS is asked to waive one or more required elements of informed consent. However, the form must include signature and date lines for the parent(s)/guardian(s) to sign if the child is permitted to take part in the research.

Assent Document - A form or script of the information that will be conveyed to the child about the study. In general, researchers must obtain the affirmative agreement of children ages seven years and older for their participation. Assent forms should be written at a level understandable to the child. If the study includes a broad age range of children, more than one assent form may be needed (i.e., an assent form suitable for a 15 year old is not usually suitable for a 7 year old child).

Assent Waiver - No child assent will be sought at all. This means that CPHS is asked to waive the requirement for child assent. Among other circumstances, this option is appropriate when the capability of the child to understand the research is too limited or when the research holds out a prospect of direct benefit that is important to the health or well being of the child.

Parent/Guardian Permission Form - A document that embodies all of the required information (elements of informed consent) designed to help the parent/guardian of a child make an informed decision about whether or not to permit the child's participation in the research. The form must include signature and date lines for the parent(s)/guardian(s) to sign if the child is permitted to take part in the research.

Permission Waiver - No parent/guardian permission will be sought at all. This means that the CPHS is asked to waive the requirement for parent/guardian permission. This option, for example, is often appropriate for research designed to study conditions in children or a study population for which parental permission is not a reasonable requirement to protect the children (e.g., neglected or abused children).

Unsigned Parent/Guardian Permission - A parent permission document that embodies all of the required information (elements of informed consent), but does not include a place for a parent to indicate with a signature that he or she agrees to permit the child's participation. This means that the CPHS is asked to waive the requirement for documented (signed) consent.

**•Child Assent and Parent Permission Guidelines, Templates, and Sample Forms**

•Policies and Procedures on Child Assent and Parent Permission

**Documents and Waivers**

| Permission/Assent Description | Assent or Permission Type | Assent/Permission Document |
|---|---|---|

**Documents and Waivers**

| | |
|---|---|
| Permission/Assent Description (e.g. Assent for Group A, Permission for Group A, Waiver of Parent Permission for Group B, Assent for Group B etc) | Assent Waiver for EE, parasites, LNS, nutrition |
| Assent or Permission Type | Assent Waiver |

For CPHS to approve a waiver of child assent (e.g. no assent will be obtained from child/minor at all), either criterion A, B, C or D must be met. Please check the applicable criterion and provide justification in the box below.

Y **A.** The capability of some or all of the children is so limited that they cannot reasonably be consulted.

This children in this study are very young (age 0 - 30 months).  Mothers/guardians of these children will be asked to consent on behalf of these children.

**B.** The intervention or procedure involved in the research holds out the prospect of direct benefit to the health or well-being of the children and is available only in the context of the research.

**C.** (1) The research involves no more than minimal risk of harm to the subjects;

(2) The waiver or alteration will not adversely affect the rights and welfare of the subjects;

(3) The research could not practicably be carried out without the waiver or alteration; and

(4) Whenever appropriate, the subjects will be provided with pertinent information after participation.

---

**Protocol Title:** Measuring the benefits of sanitation, water quality, handwashing and nutrition interventions for improving health and development in rural Bangladesh

**Protocol Status:** APPROVED

**Date Submitted:** 10/26/2017

**Approval Period:** **10/30/2017-11/03/2018**

**Important Note:** **This Print View may not reflect all comments and contingencies for approval. Please check the comments section of the online protocol. Questions that appear to not have been answered may not have been required for this submission. Please see the system application for more details.**

**D.** (1) The research or demonstration project is to be conducted by or subject to the approval of state or local officials and is designed to study, evaluate, or otherwise examine: (i) public benefit or service programs; (ii) procedures for obtaining benefits or service; (iii) possible changes in or alternatives to those programs or procedures; or (iv) possible changes in methods or levels of payment for benefits or services under those programs; AND

(2) The research could not practicably be carried out without the waiver or alteration.

| | | | |
|---|---|---|---|
| Permission/Assent Description (e.g. Assent for Group A, Permission for Group A, Waiver of Parent Permission for Group B, Assent for Group B etc) | Assent for children 7-12 years in parasite assessment | | |
| Assent or Permission Type | Assent Document | | |
| Attach Assent or Permission documents (in Word format) | X | Assent/Permission Document | Appendix 1i-2 Parasite Assent Form |

Explain how, where, when, and by whom assent will be obtained from the minor/child.

After the parent has given consent or permission for their child to participate in the research, the field worker will ask to speak with the child. The field worker will verbally administer the assent form in Bengali (to be attached later once translation is completed). Assent will be obtained at the child's compound.

Describe any additional/appropriate measures that will be in place to protect this vulnerable population, if any.

We will also ask for informed consent from each child's parent.

---

---

**Protocol Title:** Measuring the benefits of sanitation, water quality, handwashing and nutrition interventions for improving health and development in rural Bangladesh

**Protocol Status:** APPROVED

**Date Submitted:** 10/26/2017

**Approval Period:** 10/30/2017-11/03/2018

**Important Note:** This Print View may not reflect all comments and contingencies for approval. Please check the comments section of the online protocol.
Questions that appear to not have been answered may not have been required for this submission. Please see the system application for more details.

#### * * * HIPAA * * *

##### 21. Health Insurance Portability and Accountability Act (HIPAA)

The HIPAA Privacy Rule establishes the right of an individual to authorize a covered entity, such as a health plan, health care clearinghouse or health care provider, to use and disclose his/her Protected Health Information (PHI) for research purposes. UC Berkeley's covered entities are the University Health Services (including its health care services on behalf of Intercollegiate Athletics) and the Optometry Clinic. The Privacy Rule defines the elements of individual information that comprise PHI and establishes the conditions under which PHI may be used or disclosed by covered entities for research purposes. It also includes provisions to allow an individual's PHI to be disclosed or used in research without their authorization (i.e., IRB waiver of authorization). For more information, see CPHS Guidelines HIPAA and Human Subjects Research.

a. Does the study involve use of Protected Health Information (PHI) from a "covered entity" outside of UC Berkeley (i.e. another organization or institution)? For more information, see HIPAA and Human Subjects Research.

N

If Yes, explain what arrangements have been made to comply with the HIPAA requirements of the entity from which the PHI will be obtained:

b. Does the study involve use of a "Limited Data Set" from a covered entity? For more information, see HIPAA and Human Subjects Research Please see The Industry Alliance Office website for limited data set requirements.

N

If Yes, patient authorization for use of the data set is not required; however, you must have a data use agreement in place with the data holder from which the data will be obtained as required by HIPAA. Contact the Industry Alliance Office for further information at (510) 642-5766.

c. Does the study involve use of Protected Health Information (PHI) from UC Berkeley's University Health Services (including its health care services on behalf of Intercollegiate Athletics) and/or the Optometry Clinic?

N

If Yes (and a limited data set will not be used), EITHER request/add a Waiver/Alteration of HIPAA Authorization below OR provide a HIPAA Authorization Form in the Attachments section of the

---

| | |
|---|---|
| **Protocol Title:** | Measuring the benefits of sanitation, water quality, handwashing and nutrition interventions for improving health and development in rural Bangladesh |
| **Protocol Status:** | APPROVED |
| **Date Submitted:** | 10/26/2017 |
| **Approval Period:** | **10/30/2017-11/03/2018** |
| **Important Note:** | **This Print View may not reflect all comments and contingencies for approval. Please check the comments section of the online protocol. Questions that appear to not have been answered may not have been required for this submission. Please see the system application for more details.** |

**protocol.**

HIPAA WAIVER/ALTERATION: For each waiver or alteration of the requirement for authorization from the patient for use of his or her PHI, provide justification in the table below.

Note: Use table below ONLY when requesting waiver/alteration of HIPAA authorization for use of PHI from UC Berkeley's Tang Health Center, the Human Resources Health Plan, Athletics and Recreational Sports, and/or the Optometry Clinic. For more information, see HIPAA and Human Subjects Research.

---

---

**Protocol Title:** Measuring the benefits of sanitation, water quality, handwashing and nutrition interventions for improving health and development in rural Bangladesh

**Protocol Status:** APPROVED

**Date Submitted:** 10/26/2017

**Approval Period:** 10/30/2017-11/03/2018

**Important Note:** **This Print View may not reflect all comments and contingencies for approval. Please check the comments section of the online protocol. Questions that appear to not have been answered may not have been required for this submission. Please see the system application for more details.**

#### * * * Attachments * * *

##### 22. Attachments

Add appropriate attachments (e.g., advertisements, data collection instruments, IRB approvals from collaborating institutions, etc.) in this section. Attachments MUST be in PDF or Word format. Please see eProtocol Attachments Check List for Non-Exempt Applications for more information.

| Document Type | Document Name | Attached Date | Submitted Date |
|---|---|---|---|
| CITI Certificate(s) | Jade CITI completion report_20111127 | 11/28/2011 | 01/12/2012 |
| Survey Instruments | Wash Benefits_Main Study Baseline_04_3 2012 | 03/06/2012 | 03/06/2012 |
| Other | Figure 1-Study Design | 03/06/2012 | 03/06/2012 |
| Other | Nutrition-Supplement-Analysis | 03/06/2012 | 03/06/2012 |
| Other | WASH Benefits Behavior Change Strategy Overview Feb 21 2012 | 03/16/2012 | 03/16/2012 |
| Other Institutions' IRB Approvals | Prot PR-11063_ERC approval_9 April 2012 | 04/13/2012 | 04/30/2012 |
| Notice of Intent to Rely Form | NOITR-597-0 | 05/30/2013 | 06/03/2013 |
| Survey Instruments | Wash Benefits Bangladesh Baseline Questionnaire_2013_03 | 07/08/2013 | 07/08/2013 |
| CITI Certificate(s) | Eugene CITI Completion Report 20130726 | 07/26/2013 | 07/28/2013 |
| CITI Certificate(s) | BenArnold-CITI | 07/28/2013 | 07/28/2013 |
| CITI Certificate(s) | CitiApprovalBiomedical_Hubbard2013 | 07/28/2013 | 07/28/2013 |
| CITI Certificate(s) | Colford CITI Completion Report | 07/28/2013 | 07/28/2013 |
| CITI Certificate(s) | Fernald CITI Group 1 Completion Report | 07/28/2013 | 07/28/2013 |
| CITI Certificate(s) | Audrie - Part 2 Citi | 07/28/2013 | 07/28/2013 |

---

**Protocol Title:** Measuring the benefits of sanitation, water quality, handwashing and nutrition interventions for improving health and development in rural Bangladesh

**Protocol Status:** APPROVED

**Date Submitted:** 10/26/2017

**Approval Period:** **10/30/2017-11/03/2018**

**Important Note:** **This Print View may not reflect all comments and contingencies for approval. Please check the comments section of the online protocol. Questions that appear to not have been answered may not have been required for this submission. Please see the system application for more details.**

| | | | |
|---|---|---|---|
| Other Institutions' IRB Approvals | Prot PR-11063_ERC certificate on yearly review_1 Sep 2013 | 09/27/2013 | 10/03/2013 |
| Survey Instruments | Wash Benefits Midline_Endline Instruments_131003 | 10/03/2013 | 10/03/2013 |
| CITI Certificate(s) | Ercumen_CITI Completion_2008 | 11/19/2013 | 11/19/2013 |
| CITI Certificate(s) | KNelson CITI Human Subjects completion report | 11/19/2013 | 11/19/2013 |
| Other Institutions' IRB Approvals | Protocol PR # 11063_ERC approval_addendum_17 NOV | 12/05/2013 | 12/05/2013 |
| Survey Instruments | wash benefits endline spillover compounds instrument_v2 | 02/25/2014 | 02/28/2014 |
| Other Institutions' IRB Approvals | Stanford WASH B Extension Dec 2013 ApprovalLetter-25863 | 03/26/2014 | 03/29/2014 |
| Recruitment Script(s) | WASH Benefits Bangladesh Spillover recruitment script | 04/08/2014 | 04/08/2014 |
| Inter-institutional Agreement | 2011-09-3652_IIA_Wagner | 07/21/2014 | 07/21/2014 |
| Other Institutions' IRB Approvals | Prot PR-11063_ERC approval addendum_10 June 2014 | 08/27/2014 | 08/29/2014 |
| Other Institutions' IRB Approvals | PR-11063_RRC Addendum Approval_13 April 2014 | 08/29/2014 | 08/29/2014 |
| Survey Instruments | Wash Benefits Midline & Endline Survey Questionnaires_no life experiences | 09/09/2014 | 09/12/2014 |
| Survey Instruments | EE Collection Form v7_12.11.2014 (Paper) | 11/12/2014 | 11/12/2014 |
| Recruitment Script(s) | WASH Benefits Bangladesh Parasites Recruitment script_v1 | 12/11/2014 | 12/11/2014 |

---

**Protocol Title:** Measuring the benefits of sanitation, water quality, handwashing and nutrition interventions for improving health and development in rural Bangladesh

**Protocol Status:** APPROVED

**Date Submitted:** 10/26/2017

**Approval Period:** **10/30/2017-11/03/2018**

**Important Note:** **This Print View may not reflect all comments and contingencies for approval. Please check the comments section of the online protocol.
Questions that appear to not have been answered may not have been required for this submission. Please see the system application for more details.**

| | | | |
|---|---|---|---|
| Survey Instruments | Spillover study qualitative instrument_v1 | 12/11/2014 | 12/11/2014 |
| Survey Instruments | Environmental Enteropathy Supplemental questionnaires | 08/31/2015 | 09/01/2015 |
| References | References | 09/23/2015 | 09/23/2015 |
| Other | 2011-09-3652_Colford_1j-Consent-EE-v21-English_Tracked Changes | 09/23/2015 | 09/23/2015 |
| Other Institutions' IRB Approvals | icddr,b PR-11063_Continuation Approval_24 Feb 2015_9 April 2015-8 April 2016 | 11/01/2015 | 11/01/2015 |
| Other Institutions' IRB Approvals | Stanford WASH Benefits IRB continuation approval Nov. 2015 | 12/18/2015 | 12/18/2015 |
| Other Institutions' IRB Approvals | PR-11063 ERC approval addendum 09Apr2016 | 03/29/2016 | 03/29/2016 |
| Other Institutions' IRB Approvals | PR#11063 (WASHB) Continuation approval upto April 8, 2018 | 10/19/2017 | 10/19/2017 |
| Other Institutions' IRB Approvals | Stanford IRB ApprovalLetter-25863 through Oct 2017 | 10/19/2017 | 10/19/2017 |

**Document Type** CITI Certificate(s)
**Document Name** Jade CITI completion report_20111127

**Document Type** Survey Instruments
**Document Name** Wash Benefits_Main Study Baseline_04_3 2012

**Document Type** Other
**Document Name** Figure 1-Study Design

---

| | |
|---|---|
| **Protocol Title:** | Measuring the benefits of sanitation, water quality, handwashing and nutrition interventions for improving health and development in rural Bangladesh |
| **Protocol Status:** | APPROVED |
| **Date Submitted:** | 10/26/2017 |
| **Approval Period:** | **10/30/2017-11/03/2018** |
| **Important Note:** | **This Print View may not reflect all comments and contingencies for approval. Please check the comments section of the online protocol. Questions that appear to not have been answered may not have been required for this submission. Please see the system application for more details.** |

| | |
|---|---|
| **Document Type** | Other |
| **Document Name** | Nutrition-Supplement-Analysis |
| **Document Type** | Other |
| **Document Name** | WASH Benefits Behavior Change Strategy Overview Feb 21 2012 |
| **Document Type** | Other Institutions' IRB Approvals |
| **Document Name** | Prot PR-11063_ERC approval_9 April 2012 |
| **Document Type** | Notice of Intent to Rely Form |
| **Document Name** | NOITR-597-0 |
| **Document Type** | Survey Instruments |
| **Document Name** | Wash Benefits Bangladesh Baseline Questionnaire_2013_03 |
| **Document Type** | CITI Certificate(s) |
| **Document Name** | Eugene CITI Completion Report 20130726 |
| **Document Type** | CITI Certificate(s) |
| **Document Name** | BenArnold-CITI |
| **Document Type** | CITI Certificate(s) |
| **Document Name** | CitiApprovalBiomedical_Hubbard2013 |
| **Document Type** | CITI Certificate(s) |
| **Document Name** | Colford CITI Completion Report |

---

**Protocol Title:** Measuring the benefits of sanitation, water quality, handwashing and nutrition interventions for improving health and development in rural Bangladesh

**Protocol Status:** APPROVED

**Date Submitted:** 10/26/2017

**Approval Period:** 10/30/2017-11/03/2018

**Important Note:** **This Print View may not reflect all comments and contingencies for approval. Please check the comments section of the online protocol. Questions that appear to not have been answered may not have been required for this submission. Please see the system application for more details.**

**Document Type** CITI Certificate(s)
**Document Name** Fernald CITI Group 1 Completion Report

**Document Type** CITI Certificate(s)
**Document Name** Audrie - Part 2 Citi

**Document Type** Other Institutions' IRB Approvals
**Document Name** Prot PR-11063_ERC certificate on yearly review_1 Sep 2013

**Document Type** Survey Instruments
**Document Name** Wash Benefits Midline_Endline Instruments_131003

**Document Type** CITI Certificate(s)
**Document Name** Ercumen_CITI Completion_2008

**Document Type** CITI Certificate(s)
**Document Name** KNelson CITI Human Subjects completion report

**Document Type** Other Institutions' IRB Approvals
**Document Name** Protocol PR # 11063_ERC approval_addendum_17 NOV

**Document Type** Survey Instruments
**Document Name** wash benefits endline spillover compounds instrument_v2

**Document Type** Other Institutions' IRB Approvals

---

| | |
|---|---|
| **Protocol Title:** | Measuring the benefits of sanitation, water quality, handwashing and nutrition interventions for improving health and development in rural Bangladesh |
| **Protocol Status:** | APPROVED |
| **Date Submitted:** | 10/26/2017 |
| **Approval Period:** | **10/30/2017-11/03/2018** |
| **Important Note:** | **This Print View may not reflect all comments and contingencies for approval. Please check the comments section of the online protocol. Questions that appear to not have been answered may not have been required for this submission. Please see the system application for more details.** |

| | |
|---|---|
| **Document Name** | Stanford WASH B Extension Dec 2013 ApprovalLetter-25863 |
| **Document Type** | Recruitment Script(s) |
| **Document Name** | WASH Benefits Bangladesh Spillover recruitment script |
| **Document Type** | Inter-institutional Agreement |
| **Document Name** | 2011-09-3652_IIA_Wagner |
| **Document Type** | Other Institutions' IRB Approvals |
| **Document Name** | Prot PR-11063_ERC approval addendum_10 June 2014 |
| **Document Type** | Other Institutions' IRB Approvals |
| **Document Name** | PR-11063_RRC Addendum Approval_13 April 2014 |
| **Document Type** | Survey Instruments |
| **Document Name** | Wash Benefits Midline & Endline Survey Questionnaires_no life experiences |
| **Document Type** | Survey Instruments |
| **Document Name** | EE Collection Form v7_12.11.2014 (Paper) |
| **Document Type** | Recruitment Script(s) |
| **Document Name** | WASH Benefits Bangladesh Parasites Recruitment script_v1 |
| **Document Type** | Survey Instruments |

---

| | |
|---|---|
| **Protocol Title:** | Measuring the benefits of sanitation, water quality, handwashing and nutrition interventions for improving health and development in rural Bangladesh |
| **Protocol Status:** | APPROVED |
| **Date Submitted:** | 10/26/2017 |
| **Approval Period:** | **10/30/2017-11/03/2018** |
| **Important Note:** | **This Print View may not reflect all comments and contingencies for approval. Please check the comments section of the online protocol. Questions that appear to not have been answered may not have been required for this submission. Please see the system application for more details.** |

| | |
|---|---|
| **Document Name** | Spillover study qualitative instrument_v1 |
| **Document Type** | Survey Instruments |
| **Document Name** | Environmental Enteropathy Supplemental questionnaires |
| **Document Type** | References |
| **Document Name** | References |
| **Document Type** | Other |
| **Document Name** | 2011-09-3652_Colford_1j-Consent-EE-v21-English_Tracked Changes |
| **Document Type** | Other Institutions' IRB Approvals |
| **Document Name** | icddr,b PR-11063_Continuation Approval_24 Feb 2015_9 April 2015-8 April 2016 |
| **Document Type** | Other Institutions' IRB Approvals |
| **Document Name** | Stanford WASH Benefits IRB continuation approval Nov. 2015 |
| **Document Type** | Other Institutions' IRB Approvals |
| **Document Name** | PR-11063 ERC approval addendum 09Apr2016 |
| **Document Type** | Other Institutions' IRB Approvals |
| **Document Name** | PR#11063 (WASHB) Continuation approval upto April 8, 2018 |
| **Document Type** | Other Institutions' IRB Approvals |

---

**Protocol Title:** Measuring the benefits of sanitation, water quality, handwashing and nutrition interventions for improving health and development in rural Bangladesh

**Protocol Status:** APPROVED

**Date Submitted:** 10/26/2017

**Approval Period:** **10/30/2017-11/03/2018**

**Important Note:** **This Print View may not reflect all comments and contingencies for approval. Please check the comments section of the online protocol. Questions that appear to not have been answered may not have been required for this submission. Please see the system application for more details.**

| **Document Name** | Stanford IRB ApprovalLetter-25863 through Oct 2017 |
| --- | --- |

---

---

| | |
|---|---|
| **Protocol Title:** | Measuring the benefits of sanitation, water quality, handwashing and nutrition interventions for improving health and development in rural Bangladesh |
| **Protocol Status:** | APPROVED |
| **Date Submitted:** | 10/26/2017 |
| **Approval Period:** | **10/30/2017-11/03/2018** |
| **Important Note:** | **This Print View may not reflect all comments and contingencies for approval. Please check the comments section of the online protocol. Questions that appear to not have been answered may not have been required for this submission. Please see the system application for more details.** |

#### * * * Assurance * * *

**Assurance**

**As Principal Investigator, I have ultimate responsibility for the performance of this study, the protection of the rights and welfare of the human subjects, and strict adherence by all co-investigators and research personnel to CPHS requirements, federal regulations, and state statutes for human subject's research.**

**I hereby assure the following:**

1. The information provided in this application is accurate to the best of my knowledge.

2. All experiments and procedures involving human subjects will be performed under my supervision or that of another qualified professional listed on this protocol.

3. This protocol covers the human subjects research activities described in the grant proposal(s) supporting this research and any such activities that are not covered have been/will be covered by a CPHS approved protocol.

4. The legally effective informed consent of all human subjects or their legally authorized representative will be obtained (unless waived) using only the current, approved consent form(s).

5. If any study subject experiences an unanticipated problem involving risks to subjects or others, and/or a serious adverse event, the CPHS will be informed promptly within no more than one week (7 calendar days), and receive a written report within no more than two weeks (14 calendar days), of recognition/ notification of the event.

6. No change in the design, conduct, or key personnel of this research will be implemented without prior CPHS review and approval, unless the changes are necessary to eliminate an apparent immediate hazard to subjects. Changes made to eliminate hazards to subjects will be reported to OPHS/CPHS via the AE/UP reporting process.

7. Applications for continuation review will be submitted in a timely manner prior to the expiration date to allow sufficient time for the renewal process. I understand that if approval expires, all research activity (including data analysis) must cease until I receive notice of re-approval by the CPHS.

---

**Protocol Title:** Measuring the benefits of sanitation, water quality, handwashing and nutrition interventions for improving health and development in rural Bangladesh

**Protocol Status:** APPROVED

**Date Submitted:** 10/26/2017

**Approval Period:** 10/30/2017-11/03/2018

**Important Note:** **This Print View may not reflect all comments and contingencies for approval. Please check the comments section of the online protocol.
Questions that appear to not have been answered may not have been required for this submission. Please see the system application for more details.**

8. Participants' complaints or requests for information about the study will be addressed appropriately.

9. I will promptly and completely comply with a CPHS decision to suspend or withdraw its approval for the project.

10. I will submit a study closure form at the conclusion of this project.

X **I have read and agree to the above assurances.**

---

---

**Protocol Title:** Measuring the benefits of sanitation, water quality, handwashing and nutrition interventions for improving health and development in rural Bangladesh

**Protocol Status:** APPROVED

**Date Submitted:** 10/26/2017

**Approval Period:** 10/30/2017-11/03/2018

**Important Note:** This Print View may not reflect all comments and contingencies for approval. Please check the comments section of the online protocol.
Questions that appear to not have been answered may not have been required for this submission. Please see the system application for more details.

#### * * * Event History * * *

##### Event History

| Date | Status | View Attachments | Letters |
| --- | --- | --- | --- |
| 10/30/2017 | AMENDMENT 17 FORM APPROVED | Y | Y |
| 10/30/2017 | AMENDMENT 17 FORM REVIEWER(S) ASSIGNED | | |
| 10/30/2017 | AMENDMENT 17 FORM SUBMITTED (CYCLE 1) | Y | |
| 10/27/2017 | AMENDMENT 17 FORM PANEL MANAGER REVIEW | | |
| 10/26/2017 | AMENDMENT 17 FORM SUBMITTED | Y | |
| 10/26/2017 | AMENDMENT 17 FORM CREATED | | |
| 10/24/2017 | CONTINUING REVIEW 4 FORM APPROVED | Y | Y |
| 10/19/2017 | CONTINUING REVIEW 4 FORM REVIEWER(S) ASSIGNED | | |
| 10/19/2017 | CONTINUING REVIEW 4 FORM SUBMITTED (CYCLE 1) | Y | |
| 10/05/2017 | CONTINUING REVIEW 4 FORM PANEL MANAGER REVIEW | | |
| 09/14/2017 | CONTINUING REVIEW 4 FORM SUBMITTED | Y | |
| 09/12/2017 | CONTINUING REVIEW 4 FORM CREATED | | |
| 11/02/2016 | CONTINUING REVIEW 3 FORM APPROVED | Y | Y |
| 10/28/2016 | CONTINUING REVIEW 3 FORM REVIEWER(S) ASSIGNED | | |

---

| | | | |
|---|---|---|---|
| 10/28/2016 | CONTINUING REVIEW 3 FORM SUBMITTED (CYCLE 1) | Y | |
| 10/11/2016 | CONTINUING REVIEW 3 FORM PANEL MANAGER REVIEW | | |
| 10/03/2016 | CONTINUING REVIEW 3 FORM SUBMITTED | Y | |
| 09/20/2016 | CONTINUING REVIEW 3 FORM CREATED | | |
| 04/11/2016 | AMENDMENT 16 FORM APPROVED | Y | Y |
| 04/08/2016 | AMENDMENT 16 FORM REVIEWER(S) ASSIGNED | | |
| 04/07/2016 | AMENDMENT 16 FORM SUBMITTED (CYCLE 1) | Y | |
| 04/07/2016 | AMENDMENT 16 FORM SUBMITTED (CYCLE 1) | Y | |
| 03/29/2016 | AMENDMENT 16 FORM PANEL MANAGER REVIEW | | |
| 03/29/2016 | AMENDMENT 16 FORM SUBMITTED | Y | |
| 03/29/2016 | AMENDMENT 16 FORM CREATED | | |
| 12/21/2015 | AMENDMENT 15 FORM APPROVED | Y | Y |
| 12/18/2015 | AMENDMENT 15 FORM REVIEWER(S) ASSIGNED | | |
| 12/18/2015 | AMENDMENT 15 FORM PANEL MANAGER REVIEW | | |
| 12/18/2015 | AMENDMENT 15 FORM SUBMITTED | Y | |
| 12/08/2015 | AMENDMENT 15 FORM CREATED | | |
| 11/06/2015 | AMENDMENT 14 FORM APPROVED | Y | Y |
| 11/03/2015 | AMENDMENT 14 FORM REVIEWER(S) ASSIGNED | | |
| 11/03/2015 | AMENDMENT 14 FORM PANEL MANAGER REVIEW | | |

---

**Protocol Title:** Measuring the benefits of sanitation, water quality, handwashing and nutrition interventions for improving health and development in rural Bangladesh

**Protocol Status:** APPROVED

**Date Submitted:** 10/26/2017

**Approval Period:** 10/30/2017-11/03/2018

**Important Note:** This Print View may not reflect all comments and contingencies for approval. Please check the comments section of the online protocol.
Questions that appear to not have been answered may not have been required for this submission. Please see the system application for more details.

| | | | |
|---|---|---|---|
| 11/01/2015 | AMENDMENT 14 FORM SUBMITTED | Y | |
| 09/28/2015 | AMENDMENT 14 FORM CREATED | | |
| 09/28/2015 | AMENDMENT 13 FORM APPROVED | Y | Y |
| 09/25/2015 | AMENDMENT 13 FORM REVIEWER(S) ASSIGNED | | |
| 09/23/2015 | AMENDMENT 13 FORM SUBMITTED (CYCLE 1) | Y | |
| 09/08/2015 | AMENDMENT 13 FORM PANEL MANAGER REVIEW | | |
| 09/01/2015 | AMENDMENT 13 FORM SUBMITTED | Y | |
| 08/25/2015 | AMENDMENT 13 FORM CREATED | | |
| 12/21/2014 | AMENDMENT 12 FORM APPROVED | Y | Y |
| 12/16/2014 | AMENDMENT 12 FORM REVIEWER(S) ASSIGNED | | |
| 12/15/2014 | AMENDMENT 12 FORM SUBMITTED (CYCLE 2) | Y | |
| 12/11/2014 | AMENDMENT 12 FORM SUBMITTED (CYCLE 1) | Y | |
| 11/17/2014 | AMENDMENT 12 FORM PANEL MANAGER REVIEW | | |
| 11/12/2014 | AMENDMENT 12 FORM SUBMITTED | Y | |
| 11/10/2014 | AMENDMENT 12 FORM CREATED | | |
| 10/16/2014 | AMENDMENT 11 FORM SUBMITTED (CYCLE 2) | Y | |
| 10/16/2014 | AMENDMENT 11 FORM WITHDRAWN | | |
| 10/16/2014 | AMENDMENT 11 FORM SUBMITTED (CYCLE 2) | Y | |
| 10/03/2014 | AMENDMENT 11 FORM TABLED | | |

---

**Protocol Title:** Measuring the benefits of sanitation, water quality, handwashing and nutrition interventions for improving health and development in rural Bangladesh

**Protocol Status:** APPROVED

**Date Submitted:** 10/26/2017

**Approval Period:** **10/30/2017-11/03/2018**

**Important Note:** **This Print View may not reflect all comments and contingencies for approval. Please check the comments section of the online protocol. Questions that appear to not have been answered may not have been required for this submission. Please see the system application for more details.**

| | | | |
|---|---|---|---|
| 09/26/2014 | AMENDMENT 11 FORM REVIEWER(S) ASSIGNED | | |
| 09/26/2014 | AMENDMENT 11 FORM SUBMITTED (CYCLE 1) | Y | |
| 09/23/2014 | AMENDMENT 11 FORM PANEL MANAGER REVIEW | | |
| 09/22/2014 | AMENDMENT 11 FORM SUBMITTED | Y | |
| 09/22/2014 | AMENDMENT 11 FORM CREATED | | |
| 09/17/2014 | AMENDMENT 10 FORM APPROVED | Y | Y |
| 09/15/2014 | AMENDMENT 10 FORM REVIEWER(S) ASSIGNED | | |
| 09/15/2014 | AMENDMENT 10 FORM SUBMITTED (CYCLE 3) | Y | |
| 09/12/2014 | AMENDMENT 10 FORM SUBMITTED (CYCLE 2) | Y | |
| 08/29/2014 | AMENDMENT 10 FORM SUBMITTED (CYCLE 1) | Y | |
| 08/21/2014 | AMENDMENT 10 FORM PANEL MANAGER REVIEW | | |
| 08/18/2014 | AMENDMENT 10 FORM SUBMITTED | Y | |
| 08/13/2014 | AMENDMENT 10 FORM CREATED | | |
| 07/21/2014 | AMENDMENT 9 FORM APPROVED | Y | Y |
| 07/21/2014 | AMENDMENT 9 FORM REVIEWER(S) ASSIGNED | | |
| 07/18/2014 | AMENDMENT 9 FORM PANEL MANAGER REVIEW | | |
| 07/16/2014 | AMENDMENT 9 FORM SUBMITTED | Y | |
| 07/15/2014 | AMENDMENT 9 FORM CREATED | | |
| 04/11/2014 | AMENDMENT 8 FORM APPROVED | Y | Y |

---

**Protocol Title:** Measuring the benefits of sanitation, water quality, handwashing and nutrition interventions for improving health and development in rural Bangladesh

**Protocol Status:** APPROVED

**Date Submitted:** 10/26/2017

**Approval Period:** 10/30/2017-11/03/2018

**Important Note:** This Print View may not reflect all comments and contingencies for approval. Please check the comments section of the online protocol.
Questions that appear to not have been answered may not have been required for this submission. Please see the system application for more details.

| | | | |
|---|---|---|---|
| 04/09/2014 | AMENDMENT 8 FORM REVIEWER(S) ASSIGNED | | |
| 04/08/2014 | AMENDMENT 8 FORM SUBMITTED (CYCLE 2) | Y | |
| 03/29/2014 | AMENDMENT 8 FORM SUBMITTED (CYCLE 1) | Y | |
| 03/03/2014 | AMENDMENT 8 FORM PANEL MANAGER REVIEW | | |
| 02/28/2014 | AMENDMENT 8 FORM SUBMITTED | Y | |
| 02/25/2014 | AMENDMENT 8 FORM CREATED | | |
| 01/29/2014 | AMENDMENT 7 FORM APPROVED | Y | Y |
| 01/24/2014 | AMENDMENT 7 FORM REVIEWER(S) ASSIGNED | | |
| 01/23/2014 | AMENDMENT 7 FORM SUBMITTED (CYCLE 3) | Y | |
| 01/23/2014 | AMENDMENT 7 FORM SUBMITTED (CYCLE 2) | Y | |
| 01/22/2014 | AMENDMENT 7 FORM SUBMITTED (CYCLE 1) | Y | |
| 01/21/2014 | AMENDMENT 7 FORM PANEL MANAGER REVIEW | | |
| 01/21/2014 | AMENDMENT 7 FORM SUBMITTED | Y | |
| 01/08/2014 | AMENDMENT 7 FORM CREATED | | |
| 12/12/2013 | AMENDMENT 6 FORM APPROVED | Y | Y |
| 12/06/2013 | AMENDMENT 6 FORM REVIEWER(S) ASSIGNED | | |
| 12/06/2013 | AMENDMENT 6 FORM SUBMITTED (CYCLE 1) | Y | |
| 12/05/2013 | AMENDMENT 6 FORM PANEL MANAGER REVIEW | | |
| 12/05/2013 | AMENDMENT 6 FORM SUBMITTED | Y | |

---

**Protocol Title:** Measuring the benefits of sanitation, water quality, handwashing and nutrition interventions for improving health and development in rural Bangladesh

**Protocol Status:** APPROVED

**Date Submitted:** 10/26/2017

**Approval Period:** 10/30/2017-11/03/2018

**Important Note:** This Print View may not reflect all comments and contingencies for approval. Please check the comments section of the online protocol.
Questions that appear to not have been answered may not have been required for this submission. Please see the system application for more details.

| | | | |
|---|---|---|---|
| 12/05/2013 | AMENDMENT 6 FORM CREATED | | |
| 12/04/2013 | AMENDMENT 5 FORM APPROVED | Y | Y |
| 12/04/2013 | AMENDMENT 5 FORM REVIEWER(S) ASSIGNED | | |
| 12/04/2013 | AMENDMENT 5 FORM PANEL MANAGER REVIEW | | |
| 11/26/2013 | AMENDMENT 5 FORM SUBMITTED | Y | |
| 11/26/2013 | AMENDMENT 5 FORM CREATED | | |
| 11/26/2013 | AMENDMENT 4 FORM APPROVED | Y | Y |
| 11/21/2013 | AMENDMENT 4 FORM REVIEWER(S) ASSIGNED | | |
| 11/20/2013 | DEVIATION 2 FORM APPROVED | Y | N |
| 11/20/2013 | DEVIATION 1 FORM APPROVED | Y | N |
| 11/19/2013 | AMENDMENT 4 FORM SUBMITTED (CYCLE 2) | Y | |
| 11/12/2013 | AMENDMENT 4 FORM SUBMITTED (CYCLE 1) | Y | |
| 11/12/2013 | AMENDMENT 4 FORM PANEL MANAGER REVIEW | | |
| 11/07/2013 | AMENDMENT 4 FORM SUBMITTED | Y | |
| 11/07/2013 | AMENDMENT 4 FORM CREATED | | |
| 10/29/2013 | CONTINUING REVIEW 2 FORM APPROVED | Y | Y |
| 10/28/2013 | DEVIATION 1 FORM REVIEWER(S) ASSIGNED | | |
| 10/28/2013 | DEVIATION 2 FORM REVIEWER(S) ASSIGNED | | |
| 10/25/2013 | CONTINUING REVIEW 2 FORM REVIEWER(S) ASSIGNED | | |

---

**Protocol Title:** Measuring the benefits of sanitation, water quality, handwashing and nutrition interventions for improving health and development in rural Bangladesh

**Protocol Status:** APPROVED

**Date Submitted:** 10/26/2017

**Approval Period:** 10/30/2017-11/03/2018

**Important Note:** This Print View may not reflect all comments and contingencies for approval. Please check the comments section of the online protocol.
Questions that appear to not have been answered may not have been required for this submission. Please see the system application for more details.

| | | | |
|---|---|---|---|
| 10/25/2013 | DEVIATION 2 FORM REVIEWER(S) ASSIGNED | | |
| 10/25/2013 | DEVIATION 2 FORM REVIEWER(S) ASSIGNED | | |
| 10/25/2013 | DEVIATION 2 FORM REVIEWER(S) ASSIGNED | | |
| 10/25/2013 | DEVIATION 1 FORM REVIEWER(S) ASSIGNED | | |
| 10/25/2013 | DEVIATION 1 FORM PANEL MANAGER REVIEW | | |
| 10/25/2013 | DEVIATION 2 FORM PANEL MANAGER REVIEW | | |
| 10/24/2013 | DEVIATION 2 FORM SUBMITTED | Y | |
| 10/24/2013 | DEVIATION 1 FORM SUBMITTED | Y | |
| 10/24/2013 | DEVIATION 2 FORM CREATED | | |
| 10/24/2013 | DEVIATION 1 FORM CREATED | | |
| 10/22/2013 | CONTINUING REVIEW 2 FORM SUBMITTED (CYCLE 1) | Y | |
| 10/08/2013 | CONTINUING REVIEW 2 FORM PANEL MANAGER REVIEW | | |
| 10/03/2013 | CONTINUING REVIEW 2 FORM SUBMITTED | Y | |
| 09/23/2013 | CONTINUING REVIEW 2 FORM CREATED | | |
| 07/30/2013 | AMENDMENT 3 FORM APPROVED | Y | Y |
| 07/29/2013 | AMENDMENT 3 FORM REVIEWER(S) ASSIGNED | | |
| 07/28/2013 | AMENDMENT 3 FORM SUBMITTED (CYCLE 4) | Y | |
| 07/25/2013 | AMENDMENT 3 FORM UNDO APPROVED | | |
| 07/24/2013 | AMENDMENT 3 FORM APPROVED | Y | Y |

---

**Protocol Title:** Measuring the benefits of sanitation, water quality, handwashing and nutrition interventions for improving health and development in rural Bangladesh

**Protocol Status:** APPROVED

**Date Submitted:** 10/26/2017

**Approval Period:** 10/30/2017-11/03/2018

**Important Note:** This Print View may not reflect all comments and contingencies for approval. Please check the comments section of the online protocol.
Questions that appear to not have been answered may not have been required for this submission. Please see the system application for more details.

| | | | |
|---|---|---|---|
| 07/24/2013 | AMENDMENT 3 FORM REVIEWER(S) ASSIGNED | | |
| 07/24/2013 | AMENDMENT 3 FORM SUBMITTED (CYCLE 3) | Y | |
| 07/16/2013 | AMENDMENT 3 FORM SUBMITTED (CYCLE 2) | Y | |
| 07/08/2013 | AMENDMENT 3 FORM SUBMITTED (CYCLE 1) | Y | |
| 06/12/2013 | AMENDMENT 3 FORM PANEL MANAGER REVIEW | | |
| 06/03/2013 | AMENDMENT 3 FORM SUBMITTED | Y | |
| 03/20/2013 | AMENDMENT 3 FORM CREATED | | |
| 11/16/2012 | AMENDMENT 2 FORM APPROVED | Y | Y |
| 11/09/2012 | AMENDMENT 2 FORM REVIEWER(S) ASSIGNED | | |
| 11/09/2012 | AMENDMENT 2 FORM SUBMITTED (CYCLE 1) | Y | |
| 11/08/2012 | AMENDMENT 2 FORM PANEL MANAGER REVIEW | | |
| 11/08/2012 | AMENDMENT 2 FORM SUBMITTED | Y | |
| 11/08/2012 | AMENDMENT 2 FORM CREATED | | |
| 10/24/2012 | CONTINUING REVIEW 1 FORM APPROVED | Y | Y |
| 10/18/2012 | CONTINUING REVIEW 1 FORM REVIEWER(S) ASSIGNED | | |
| 10/18/2012 | CONTINUING REVIEW 1 FORM SUBMITTED (CYCLE 2) | Y | |
| 10/17/2012 | CONTINUING REVIEW 1 FORM SUBMITTED (CYCLE 1) | Y | |
| 10/12/2012 | CONTINUING REVIEW 1 FORM PANEL MANAGER REVIEW | | |

---

**Protocol Title:** Measuring the benefits of sanitation, water quality, handwashing and nutrition interventions for improving health and development in rural Bangladesh

**Protocol Status:** APPROVED

**Date Submitted:** 10/26/2017

**Approval Period:** 10/30/2017-11/03/2018

**Important Note:** This Print View may not reflect all comments and contingencies for approval. Please check the comments section of the online protocol.
Questions that appear to not have been answered may not have been required for this submission. Please see the system application for more details.

| | | | |
|---|---|---|---|
| 10/11/2012 | CONTINUING REVIEW 1 FORM SUBMITTED | Y | |
| 10/03/2012 | CONTINUING REVIEW 1 FORM CREATED | | |
| 05/01/2012 | AMENDMENT 1 FORM APPROVED | Y | Y |
| 04/30/2012 | AMENDMENT 1 FORM SUBMITTED (CYCLE 4) | Y | |
| 04/16/2012 | AMENDMENT 1 FORM REVIEWER(S) ASSIGNED | | |
| 04/06/2012 | AMENDMENT 1 FORM CONDITIONAL APPROVAL | | |
| 04/01/2012 | AMENDMENT 1 FORM REVIEWER(S) ASSIGNED | | |
| 03/29/2012 | AMENDMENT 1 FORM REVIEWER(S) ASSIGNED | | |
| 03/26/2012 | AMENDMENT 1 FORM SUBMITTED (CYCLE 2) | Y | |
| 03/19/2012 | AMENDMENT 1 FORM REVIEWER(S) ASSIGNED | | |
| 03/16/2012 | AMENDMENT 1 FORM SUBMITTED (CYCLE 1) | Y | |
| 03/12/2012 | AMENDMENT 1 FORM PANEL MANAGER REVIEW | | |
| 03/12/2012 | AMENDMENT 1 FORM PANEL MANAGER REVIEW | | |
| 03/06/2012 | AMENDMENT 1 FORM SUBMITTED | Y | |
| 03/05/2012 | AMENDMENT 1 FORM CREATED | | |
| 01/17/2012 | NEW FORM APPROVED | Y | Y |
| 01/12/2012 | NEW FORM SUBMITTED (CYCLE 1) | Y | |
| 11/04/2011 | NEW FORM CONDITIONAL APPROVAL | | |
| 10/28/2011 | NEW FORM REVIEWER(S) ASSIGNED | | |
| 10/04/2011 | NEW FORM PANEL MANAGER REVIEW | | |

---

| | |
|---|---|
| **Protocol Title:** | Measuring the benefits of sanitation, water quality, handwashing and nutrition interventions for improving health and development in rural Bangladesh |
| **Protocol Status:** | APPROVED |
| **Date Submitted:** | 10/26/2017 |
| **Approval Period:** | **10/30/2017-11/03/2018** |
| **Important Note:** | **This Print View may not reflect all comments and contingencies for approval. Please check the comments section of the online protocol. Questions that appear to not have been answered may not have been required for this submission. Please see the system application for more details.** |

| | | |
|---|---|---|
| 10/04/2011 | NEW FORM PANEL ASSIGNED | |
| 10/03/2011 | NEW FORM SUBMITTED | Y |
| 09/29/2011 | NEW FORM CREATED | |

---------------------------------------------------------------------------------

| | |
|---|---|
| **Protocol Title:** | Measuring the benefits of sanitation, water quality, handwashing and nutrition interventions for improving health and development in rural Bangladesh |
| **Protocol Status:** | APPROVED |
| **Date Submitted:** | 10/26/2017 |
| **Approval Period:** | **10/30/2017-11/03/2018** |
| **Important Note:** | **This Print View may not reflect all comments and contingencies for approval. Please check the comments section of the online protocol. Questions that appear to not have been answered may not have been required for this submission. Please see the system application for more details.** |

**Disclaimer: The generated PDF may not duplicate the original format completely. We do not warrant the accuracy of the changed format.**

*** * * Attached Document * * ***

| Document Name | Created Date |
|---|---|
| Jade CITI completion report_20111127.pdf | 10/26/2017 |

### CITI Collaborative Institutional Training Initiative

#### Human Research Curriculum Completion Report
#### Printed on 11/27/2011

**Learner:** Jade Benjamin-Chung (username: jadebc)
**Institution:** University of California, Berkeley


**Group 1Biomedical Research Investigators and Key Personnel:**

**Stage 1. Basic Course Passed on 11/27/11** (Ref # 7074049)

| Required Modules | Date Completed | Score |
|---|---|---|
| Belmont Report and CITI Course Introduction | 11/25/11 | 3/3 (100%) |
| History and Ethical Principles | 11/25/11 | 6/6 (100%) |
| Basic Institutional Review Board (IRB) Regulations and Review Process | 11/25/11 | 5/5 (100%) |
| Informed Consent | 11/25/11 | 4/4 (100%) |
| Social and Behavioral Research for Biomedical Researchers | 11/25/11 | 3/4 (75%) |
| Records-Based Research | 11/25/11 | 2/2 (100%) |
| Genetic Research in Human Populations | 11/25/11 | 1/2 (50%) |
| Research With Protected Populations - Vulnerable Subjects: An Overview | 11/25/11 | 4/4 (100%) |
| Vulnerable Subjects - Research Involving Prisoners | 11/25/11 | 4/4 (100%) |
| Vulnerable Subjects - Research Involving Children | 11/25/11 | 3/3 (100%) |
| Vulnerable Subjects - Research Involving Pregnant Women, Human Fetuses, and Neonates | 11/27/11 | 3/3 (100%) |
| International Studies | 11/27/11 | 1/1 (100%) |
| Avoiding Group Harms: U.S. Research Perspectives | 11/27/11 | 3/3 (100%) |
| FDA-Regulated Research | 11/27/11 | 4/5 (80%) |
| Human Subjects Research at the VA | 11/27/11 | 2/3 (67%) |
| Research and HIPAA Privacy Protections | 11/27/11 | 5/5 (100%) |
| Vulnerable Subjects - Research Involving Workers/Employees | 11/27/11 | 4/4 (100%) |
| Hot Topics | 11/27/11 | no quiz |
| Conflicts of Interest in Research Involving Human Subjects | 11/27/11 | 3/5 (60%) |
| University of California, Berkeley | 11/27/11 | no quiz |
| Stem Cell Research Oversight (Part I) | 11/27/11 | 4/5 (80%) |

**For this Completion Report to be valid, the learner listed above must be affiliated with a CITI participating institution. Falsified information and unauthorized use of the CITI course site is unethical, and may be considered scientific misconduct by your institution.**

Paul Braunschweiger Ph.D.
Professor, University of Miami
Director Office of Research Education
CITI Course Coordinator

Return

PROTOCOL
Biomedical
Non-Exempt
Berkeley

Protocol # 2011-09-3652
Date Printed: 07/31/2018

-----------------------------------------------------------------------------------
---

**Protocol Title:** Measuring the benefits of sanitation, water quality, handwashing and nutrition interventions for improving health and development in rural Bangladesh

**Protocol Status:** APPROVED

**Date Submitted:** 10/26/2017

**Approval Period:** **10/30/2017-11/03/2018**

**Important Note:** **This Print View may not reflect all comments and contingencies for approval. Please check the comments section of the online protocol.**
**Questions that appear to not have been answered may not have been required for this submission. Please see the system application for more details.**

*** * * Attached Document * * ***

| Document Name | Created Date |
|---|---|
| Wash Benefits_Main Study Baseline_04_3 2012.pdf | 10/26/2017 |

ICDDR,B

### Wash Benefits

Baseline Survey

**asania**

**2/16/2012**

Catherine Wright 3/6/12 12:10 PM
**Deleted:** 9

Catherine Wright 3/6/12 12:10 PM
**Deleted:** 9

Catherine Wright 3/6/12 12:10 PM
**Deleted:** 9

Catherine Wright 3/6/12 12:10 PM
**Deleted:** 9

Catherine Wright 3/6/12 12:10 PM
**Deleted:** 10

Catherine Wright 3/6/12 12:10 PM
**Deleted:** 10

Catherine Wright 3/6/12 12:10 PM
**Deleted:** 11

Catherine Wright 3/6/12 12:10 PM
**Deleted:** 11

Catherine Wright 3/6/12 12:10 PM
**Deleted:** 12

Catherine Wright 3/6/12 12:10 PM
**Deleted:** 13
Catherine Wright 3/6/12 12:10 PM
**Deleted:** 19
Catherine Wright 3/6/12 12:10 PM
**Deleted:** 20
Catherine Wright 3/6/12 12:10 PM
**Deleted:** 23
Catherine Wright 3/6/12 12:10 PM
**Deleted:** 23
Catherine Wright 3/6/12 12:10 PM
**Deleted:** 29
Catherine Wright 3/6/12 12:10 PM
**Deleted:** 29
Catherine Wright 3/6/12 12:10 PM
**Deleted:** 35
Catherine Wright 3/6/12 12:10 PM
**Deleted:** 35
Catherine Wright 3/6/12 12:10 PM
**Deleted:** 35
Catherine Wright 3/6/12 12:10 PM
**Deleted:** 35
Catherine Wright 3/6/12 12:10 PM
**Deleted:** 36
Catherine Wright 3/6/12 12:10 PM
**Deleted:** 36
Catherine Wright 3/6/12 12:10 PM
**Deleted:** 36
Catherine Wright 3/6/12 12:10 PM
**Deleted:** 36
Catherine Wright 3/6/12 12:10 PM
**Deleted:** 37
Catherine Wright 3/6/12 12:10 PM
**Deleted:** 37
Catherine Wright 3/6/12 12:10 PM
**Deleted:** 39
Catherine Wright 3/6/12 12:10 PM
**Deleted:** 39
Catherine Wright 3/6/12 12:10 PM
**Deleted:** 40
Catherine Wright 3/6/12 12:10 PM
**Deleted:** 40
Catherine Wright 3/6/12 12:10 PM
**Deleted:** 40
Catherine Wright 3/6/12 12:10 PM
**Deleted:** 41
Catherine Wright 3/6/12 12:10 PM
**Deleted:** 41
Catherine Wright 3/6/12 12:10 PM
**Deleted:** 41
Catherine Wright 3/6/12 12:10 PM
**Deleted:** 42
Catherine Wright 3/6/12 12:10 PM
**Deleted:** 42
Catherine Wright 3/6/12 12:10 PM
**Deleted:** 43
Catherine Wright 3/6/12 12:10 PM
**Deleted:** 43
Catherine Wright 3/6/12 12:10 PM
**Deleted:** 43

LNS measurement .......................................................................................................................... 46

Catherine Wright 3/6/12 12:10 PM
**Deleted:** 43

This document includes the baseline survey and associated instructions. It specifies the population that needs to be included for the survey, indicators from the common modules in addition to country specific indicators. It guides the user through the survey process from the start to the end. Please note that the module numbers match the common module numbers but appear out of order in this document. The items include the common module variable number in the first column to faciliate bridging the survey question numbers to the common module question numbers. Modules 40 and 41 are additional modules or subsets of the common modules split to aid survey administration, only in the Bangladesh baseline. The sequence of the modules as it appears now is open for discussion and can be easily revised. There are placeholders for the modules that are not applicable for baseline.

**NOTES TO THE FIELD TEAM**

There are 4 groups of respondents of interest to us:

| POPULATION | DEFINITION | SAMPLES |
|---|---|---|
| 1. Target children | Children in utero at enrollment (primary study population) | N/A FOR BASELINE |
| 2. Mothers | Mothers of target children | BLOOD, STOOL, URINE |
| 3. Diarrhea cohort | Children < 36 months old at enrollment, living in study compounds | NO SAMPLES |
| 4. Parasite cohort | NOT FINALIZED<br>Children 18 – 27 months old at baseline (blood spot, stool)<br>~~School-aged children 6 – 12 years old at baseline (stool)?~~ | BLOOD, STOOL |

THE LISTING TEAM WILL GIVE THE FIELD RESEARCH OFFICERS A LIST OF HOUSEHOLDS, ASSOCIATED ADDRESSES AND RESPONDENT SPECIFICS TO HELP THE BASELINE TEAM LOCATE THE ELIGIBLE HOUSEHOLDS. THE HOUSEHOLDS WILL BE VISIBLY MARKED FOR IDENTIFICATION.

**Consent (All households)**

FOLLOW THESE STEPS CAREFULLY:

1. CONFIRM THE INFORMATION FROM THE LISTING WITH THE CURRENT ADDRESS/HOUSEHOLD.
2. VERIFY RESPONDENT ELIGIBILITY BY ASKING FOR PREGNANCY STATUS.
3. DETERMINE IF THERE ARE ANY OTHER ELIGIBLE RESPONDENTS IN THE COMPOUND.
4. IF RESPONDENT MEETS OUR CRITERIA, CAREFULLY RECORD **HOUSEHOLD ID** AT THE TOP OF THE CONSENT PAGE.
5. CONFIRM THAT THE ID EXACTLY MATCHES THE ID ASSIGNED TO THIS COMPOUND/HOUSEHOLD BY THE FIELD RESEARCH OFFICER.
6. AFTER WRITING IT CROSS IT OFF YOUR LIST.

READ THE CONSENT FORM VERBATIM TO THE RESPONDENT IN THE LANGUAGE OF THE WRITTEN CONSENT FORM AND THEN CONFIRM THAT SHE UNDERSTANDS.

ENTER THE HOUSEHOLD ID: |__|__|__|__|__|__|| |__|__|__|__|__|__|| |__|__|__|.

**Read**: Hello. My name is [name]. I am from Mohakhali Cholera Hospital (icddr,b) in Dhaka. We are here to talk to you today because your household has been selected for our study because [you are pregnant/you have a child 18-27 months old/you have a child 18-27 YEARS old]. READ REST OF CONSENT.

**GPS position:** Following the household's consent, stand in a position closest to the household's front door and record the GPS coordinates.

#### Wash Benefit Module 40

##### Geographical Identification

FILL THE FOLLOWING IDENTIFICATION INFORMATION FOR THE RESPONDENT VERY CAREFULLY.

| Module ID | WBM 40 | Q ID | |
|---|---|---|---|
| | 4001. | 00 | এফ.আর.এ কোড (FRA code): ÿÿ |
| C.101 | 4002. | 01 | তথ্য সংগ্রহের তারিখ [Date of data collection]: __/__/____ |
| C.18 | 4003. | | GPS point ID number [Auto-populate, if possible] |
| C.19 | 4004. | | GPS Latitude [Auto-populate, if possible] |
| C.20 | 4005. | | GPS Longitude[Auto-populate, if possible] |
| | 4006. | 04 | DISTRICT NAME: |
| | 4007. | 05 | UPAZILA NAME: |
| | 4008. | 06 | ইউনিয়নের নাম [Union name _____________ |
| | 4009. | 07 | খানার ঠিকানা Household Address: |
| | 4010. | 08 | খানা প্রধানের নাম [Name of Household head]_____________________________ |
| | 4011. | 09 | খানা প্রধানের বাবা/স্বামীর নাম [Father's/Husband's Name of HH head] _______________ |
| | 4012. | 10 | বাড়ির নাম [Bari Name]: ________________________________________ |
| | 4013. | 11 | গ্রামের নাম [Village Name]: ____________________________________ |
| C.01 | 4014. | 12 | ক্লাস্টার নাম্বার [Cluster No. [1-720] |
| C.02 | 4015. | 13 | বাড়ী নাম্বার [Bari No. [1-7] |
| C.03 | 4016. | 14 | **HH ID:** [1-7] |
| | 4017. | 15 | Unique HH ID: |
| C.04 | 4018. | 16 | মায়ের আইডি [Mother ID [1-7] **[PDA to store in MOTHER COHORT]** |
| C.05 | 4019. | 17 | টার্গেট শিশুর আইডি [Target child ID [1-7] **[PDA to store in Target Child Cohort]** |

Now look at the list provided to ensure the presence of the additional enrolled children in the bari

| | | Who is this child | নাম Name | লিঙ্গ Sex | জন্মতারিখ Birth date | Source of DOB (C.104) | নির্বাচন করার কারণ (একাধিক উত্তর হতে পারে) Reason for enrollment (select all that apply) **[PDA to store accordingly]** |
|---|---|---|---|---|---|---|---|
| **C.06.** | 4020. | 1 Target child<br>2 Another child of same mother<br>3 Child from the same bari | | 1 পুরুষ M<br>0 মহিলা F | দিন/মাস/বছর DD/MM/YYY | 1 = Confirmed DOB by valid vaccination/health card<br>2 = Mother/Relative remembers DOB<br>3 = Both 1 & 2<br>4 = Estimated DOB with 2 and event calendar | ☐1 ডায়রিয়া Diarrhea<br>☐2 পায়খানার নমুণা Stool specimen<br>☐3 রক্তের নমুণা Blood specimen |

asania 2/12/12 2:45 PM
**Comment:** FRO TO GIVE ASSISGNMENTS

asania 2/12/12 2:45 PM
**Comment:** 7 maximum

asania 2/16/12 4:30 PM
**Comment:** Composite of 12,13,14

asania 2/15/12 9:31 PM
**Comment:** As provided by the listing team

asania 2/15/12 9:33 PM
**Comment:** Change from CM

| | | | | | | | |
|---|---|---|---|---|---|---|---|
| **C.07.** | 4021. | 1 Target child<br>2 Another child of same mother<br>3 Child from the same bari | | 1 পুরুষ M<br>0 মহিলা F | দিন/মাস/ বছর<br>DD/MM/ YYY | 1 = Confirmed DOB by valid vaccination/health card<br>2 = Mother/Relative remembers DOB<br>3 = Both 1 & 2<br>4 = Estimated DOB with 2 and event calendar | ☐1 ডায়রিয়া Diarrhea<br>☐2 পায়খানার নমুণা Stool specimen<br>☐3 রক্তের নমুণা Blood specimen |
| **C.08.** | 4022. | 1 Target child<br>2 Another child of same mother<br>3 Child from the same bari | | 1 পুরুষ M<br>0 মহিলা F | দিন/মাস/ বছর<br>DD/MM/ YYY | 1 = Confirmed DOB by valid vaccination/health card<br>2 = Mother/Relative remembers DOB<br>3 = Both 1 & 2<br>4 = Estimated DOB with 2 and event calendar | ☐1 ডায়রিয়া Diarrhea<br>☐2 পায়খানার নমুণা Stool specimen<br>☐3 রক্তের নমুণা Blood specimen |
| **C.09.** | 4023. | 1 Target child<br>2 Another child of same mother<br>3 Child from the same bari | | 1 পুরুষ M<br>0 মহিলা F | দিন/মাস/ বছর<br>DD/MM/ YYY | 1 = Confirmed DOB by valid vaccination/health card<br>2 = Mother/Relative remembers DOB<br>3 = Both 1 & 2<br>4 = Estimated DOB with 2 and event calendar | ☐1 ডায়রিয়া Diarrhea<br>☐2 পায়খানার নমুণা Stool specimen<br>☐3 রক্তের নমুণা Blood specimen |
| **C.10.** | 4024. | 1 Target child<br>2 Another child of same mother<br>3 Child from the same bari | | 1 পুরুষ M<br>0 মহিলা F | দিন/মাস/ বছর<br>DD/MM/ YYY | 1 = Confirmed DOB by valid vaccination/health card<br>2 = Mother/Relative remembers DOB<br>3 = Both 1 & 2<br>4 = Estimated DOB with 2 and event calendar | ☐1 ডায়রিয়া Diarrhea<br>☐2 পায়খানার নমুণা Stool specimen<br>☐3 রক্তের নমুণা Blood specimen |

#### Wash Benefit Module 1

##### উত্তরদাতা সম্পর্কীত তথ্য RESPONDENT IDENTIFICATION

| | | |
|---|---|---|
| মডিউল আইডি Module ID | WBM 1 | |
| | 101. 1 | : What is your full name? প্রধান উত্তরদাতার নাম [Name of respondent: |
| | 102. | : \|_____\| প্রধান উত্তরদাতার পরিচয় [Status of main respondent]<br>1. প্রধান পরির্চযাকারী /মা [Mother of youngest child]<br>2. পরির্চযাকারী (পুরুষ) [Male caregiver]<br>3. পরির্চযাকারী (মহিলা) [Female caregiver] |
| | 103. | প্রধান উত্তরদাতার বয়স (বছরে) [Age of main respondent: *(in years)]* |
| | 104. | উত্তরদাতা কোন পূরণ করেছে? RESPONDENT MEETS WHICH ELIGIBILITY CRITERIA?<br>[1] গর্ভবতী PREGNANT<br>[2] ১৮-২৭ মাসের বাচ্চা আছে HAS CHILD 18-27 MONTHS OF AGE →SKIP to |
| I need to ask you some personal questions regarding your pregnancy because that is our project's eligibility criteria. I hope you do not mind and will cooperate with us. [see scripts of other studies] | | |
| C. 108 | 105. | **What was the date of your last menstrual period?** __/__/____<br>88 / 88 / 88 = No menstruation since their last pregnancy<br>99 = Don't know / not sure |
| C. 109 | 106. | \|__\| Record number of completed months<br>99 = Don't know / not sure |
| C.110 | 107. | \|___\|Source of Pregnancy Length<br>1 = Estimated by mother only<br>2 = Estimated by mother and a health practitioner, no ultrasound (last prenatal visit)<br>3 = Estimated by mother and health practitioner, using ultrasound (last prenatal visit) |
| | 108. | What will be the expected DOB of your child?<br>__/__/__ |
| | 109. | Actual DOB of target child (skip for baseline)<br>__/__/___ |
| At some point over the next several months, we may need to get in touch with you by phone to give you information about our project. Do you have a phone? If not, does someone in your compound have a phone? Maybe a neighbor? What is the number?<br><br>**Wash Benefit Module 0**<br>**TRACKING** | | |
| | 001 | Is there a phone available? 1 Yes 0 No (Skip to end) |
| C.11 | **002** | Primary phone number |
| C.12 | 003 | Does this phone number belong to you? 1 Yes 0 No |

<table>
<tr><td>C.13</td><td>004</td><td>IF NO, to whom does this number belong? (Specify relationship to respondent)<br>1 HUSBAND<br>2 IN LAWS<br>3 FATHER<br>4 MOTHER<br>5 FAMILY MEMBER<br>99 OTHER</td></tr>
<tr><td>C.14</td><td>005</td><td>Secondary phone number</td></tr>
<tr><td>C.15</td><td>006</td><td>Does this phone number belong to you? 1 Yes 0 No</td></tr>
<tr><td>C.16</td><td>007</td><td>IF NO, to whom does this number belong?<br>1 HUSBAND<br>2 IN LAWS<br>3 FATHER<br>4 MOTHER<br>5 FAMILY MEMBER<br>99 OTHER</td></tr>
<tr><td>~~C.17~~</td><td>~~008~~</td><td>~~Directions to the compound from a central location within the village~~</td></tr>
<tr><td></td><td>009</td><td>|_____| |_____| সবচেয়ে ছোট বাচ্চার মা কত ক্লাস পর্যন্ত পড়ালেখা শেষ করেছেন ? (জানি না = ***99***) Education of mother of the youngest child<br>*(Years of education completed, DK=99)*</td></tr>
<tr><td></td><td>011</td><td>|_____| |_____| সবচেয়ে ছোট বাচ্চার বাবার প্রধান পেশা [Main occupation of father of the youngest child]<br><br>Occupation Code:<br>1. কৃষক [Farmer/Cultivator]<br>2. ঘরে কাজ করে [Homemaker]<br>3. কৃষি শ্রমিক [Agri-labor]<br>4. শ্রমিক (কৃষি শ্রমিক ছাড়া) [Non-agri labor]<br>5. বেতনভুক্ত কর্মচারী (সরকারী/প্রাইভেট/এন.জি.ও) [Salaried job (Govt./Private/NGO)]<br>6. রাজমিস্ত্রী [Mason (*Rajmistri*)]<br>7. কাঠ মিস্ত্রী [Carpenter]<br>8. ভ্যান/রিকশা চালক [Van/Rickshaw puller]<br>9. জেলে [Fisherman]<br>10. নৌকা চালক/মাঝি [Boatman]<br>11. কর্মকার [Blacksmith]<br>12. স্বর্ণকার [Goldsmith]<br>13. কুমার/কুম্ভকার [Potter (soil smith)]<br>14. মুচি [Shoe polish /maker]<br>15. দোকানদার [Shopkeeper]<br>16. ফেরিওয়ালা [Vendor<br>21. কুটীর শিল্প [Cottage industry]<br>22. পোলট্রি/ ব্যবসার জন্য পশু লালন-পালনকারী [Poultry /livestock rearer]<br>23. বৈদ্যুতিক মিস্ত্রী [Electrician]<br>24. হোমিওপ্যাথি ডাক্তার [Homeopath]<br>25. আধ্যাতিক চিকিৎসক/ কবিরাজ/ ওঝাঁ [Spiritual healer/kabiraj/ Ojha]<br>26. পেশাদার ডাক্তার/উকিল [Professional practitioner (Doctor/lawyer)]<br>27. ইমাম/ ধর্মযাজক [Imam/priest]<br>28. অবসরপ্রাপ্ত চাকুরীজীবী [Retired service holder]<br>29. ছাত্র [Student]<br>30. বেকার [Unemployed]<br>31. অক্ষম [Disabled]<br>32. কাজের লোক [Domestic maid / servant]</td></tr>
</table>

asania 3/4/12 12:21 PM
**Comment:** cut

| | | | |
|---|---|---|---|
| | | (*Feriwala/howker*)]<br>17. ক্ষুদ্র ব্যবসায়ী (মূলধন <=১০০০০) [Petty trader, capital <=10000]<br>18. ব্যবসায়ী (মূলধন >১০০০০) [Business, capital >10000]<br>19. দর্জি [Tailor]<br>20. ড্রাইভার [Driver] | / servant]<br>33. জমিদার (শস্য উৎপাদন অথবা অন্য কোন কাজে কৃষকদের জমি বর্গা দেয়) [Landlord (Provide land for farmers for sharecropping or others)]<br>34. বিদেশে থাকে [Staying abroad]<br>35. মৃত/নিখোঁজ [Died/untraced]<br>99. জানিনা [Don't know]<br><br>77. অন্যান্য (বর্ণনা লিখুন) [Others (specify)] |
| | 010 | \|_____\| \|_____\| সবচেয়ে ছোট বাচ্চার বাবা কত ক্লাস পর্যন্ত পড়ালেখা শেষ করেছেন ? (জানি না = ***99***) Education of father of the youngest child *(Years of education completed, DK=99)* | |
| | 013 | \|_____\|How many children < 36 months do you have? | |
| | 012 | \|_____\| \|_____\| আপনার খানাতে/পরিবারে কত জন লোক প্রতিদিন একই হাড়িতে রান্না করে খান?How many people in this house eat from the same cooking pot every day (খানা বলতে একই হাড়িতে রান্না করে খায় এমন সদস্যদের বুঝানো হয়েছে) | |

asania 3/4/12 12:49 PM
**Deleted:** 012

asania 3/4/12 12:49 PM
**Deleted:** |_____| |_____| আপনার খানাতে/পরিবারে কত জন লোক প্রতিদিন একই হাড়িতে রান্না করে খান?How many people in this house eat from the same cooking pot every day (খানা বলতে একই হাড়িতে রান্না করে খায় এমন সদস্যদের বুঝানো হয়েছে)

| A. CENSUS OF HOUSEHOLDS ARE IN THIS COMPOUND (All households) | | | | | | |
|---|---|---|---|---|---|---|
| READ | Okay. Now I am going to ask you about each household separately. First, think about your own household.<br>MARK "99" IF RESPONDENT DOESN'T KNOW THE ANSWER TO ANY OF THE FOLLOWING QUESTIONS. | | | | | |
| | A.1 | A.2 | A.3 | A.4 | A.5 | A.6 |
| HH No. | What is the name of the household head? FOR REFERENCE ONLY | How many adults over 40 years live in this household? | How many young adults (19-40 years) live in this household? | How many school-aged children (4-18 years) live in this household? | How many young children (0-3 years) live in this household? | CALCULATE THE TOTAL NUMBER OF PEOPLE IN EACH HH AND CONFIRM |
| 1 | | | | | | |
| 2 | | | | | | |

#### Wash Benefit Module 6

##### CHILD FOOD FREQUENCY

asania 1/29/12 12:27 PM
**Comment:** Skip for baseline

Administer to:Target Children

Respondent:Child'sprimarycaregiver.

(The primary caregiver is the person that spends the most timewith the child.This is often the mother.)

**Introduction**

Now I would like to ask you some questions about feeding [NAME].FirstIneed to know if you will be able to tell me aboutfeeding[NAME]yesterdayandoverthelastweek.

| | | | |
|---|---|---|---|
| | | CHILD ID: | \|__\|__\|__\|__\|__\|__\| |

asania 2/27/12 5:50 PM
**Comment:** REMOVING SECTION FOR NOW.NO NEED TO PROGRAM NOW. BUT LEAVE SPACE FOR IT.

#### Wash Benefit Module 18

##### MATERNAL FFQ

**7-Day Food Frequency Questionnaire for Mothers**

Now I would like to ask you some questions regarding your nutritional status. In the past 7 days, that is since last [Tomorrow's Day], on how many days have you eaten the following foods:

| CM | WBM 18 | Foods | Number of days (0-7) |
|---|---|---|---|
| **C.1801.** | 1801. | Any kind of meat (cow/buffalo/sheep/goat/pig) | [__] |
| **C.1802.** | 1802. | Any kind of poultry meat | [__] |
| **C.1803.** | 1803. | Any kind of fresh fish | [__] |
| **C.1804.** | 1804. | Any kind of dried fish | [__] |
| **C.1805.** | 1805. | Snails, crabs, insects | [__] |
| **C.1806.** | 1806. | Eggs | [__] |
| **C.1807.** | 1807. | Milk | [__] |
| **C.1808.** | 1808. | Yogurt/Curd/Buttermilk | [__] |
| **C.1809.** | 1809. | Dal and other legumes | [__] |
| **C.1810.** | 1810. | Peanuts and other nuts | [__] |
| **C.1811.** | 1811. | Orange flesh sweet potato | [__] |
| **C.1812.** | 1812. | Pumpkins / orange or yellow squash | [__] |

| | | | |
|---|---|---|---|
| **C.1813.** | 1813. | Dark green leafy vegetables | [__] |
| **C.1814.** | 1814. | Red leafy vegetables | [__] |
| **C.1815.** | 1815. | Mango | [__] |
| **C.1816.** | 1816. | Guava | [__] |
| **C.1817.** | 1817. | Pineapple | [__] |
| **C.1818.** | 1818. | Indian Jujube | [__] |
| **C.1819.** | 1819. | Banana | [__] |
| **C.1820.** | 1820. | Cauliflower | [__] |
| **C.1821.** | 1821. | Tomato | [__] |

#### Wash Benefit Module 11

#### FOOD INSECURITY

**Household Food Insecurity and Access Scale**

Administer to: All study households during baseline

| | WBM 11 | **Occurrence Questions**<br>1 = Rarely (once or twice in the past four weeks)<br>2 = Sometimes (three to ten times in the past four weeks)<br>3 = Often (more than ten times in the past four weeks) | Response | If yes, how often did this happen? |
|---|---|---|---|---|
| **C.1101.** | 1101. | 1. In the past four weeks, did you worry that your household would not have enough food? | [0] No [skip to 1102]<br>[1] Yes | [1] Rarely<br>[2] Sometimes<br>[3] Often |
| **C.1102.** | 1102. | 2. In the past four weeks, were you or any household member not able to eat the kinds of foods you preferred because of a lack of resources? | [0] No [skip to 1103]<br>[1] Yes | [1] Rarely<br>[2] Sometimes<br>[3] Often |
| **C.1103.** | 1103. | 3. In the past four weeks, did you or any household member have to eat a limited variety of foods due to a lack of resources? | [0] No [skip to 1104]<br>[1] Yes | [1] Rarely<br>[2] Sometimes<br>[3] Often |
| **C.1104.** | 1104. | 4. In the past four weeks, did you or any household member have to eat some foods that you really did not want to eat because of a lack of resources to obtain other types of food? | [0] No [skip to 1105]<br>[1] Yes | [1] Rarely<br>[2] Sometimes<br>[3] Often |
| **C.1105.** | 1105. | 5. In the past four weeks, did you or any household member have to eat a smaller meal than you felt you needed because there was not enough food? | [0] No [skip to 1106]<br>[1] Yes | [1] Rarely<br>[2] Sometimes<br>[3] Often |
| **C.1106.** | 1106. | 6. In the past four weeks, did you or any household member have to eat fewer meals in a day because there was not enough food? | [0] No [skip to 1107]<br>[1] Yes | [1] Rarely<br>[2] Sometimes<br>[3] Often |
| **C.1107.** | 1107. | 7. In the past four weeks, was there ever no food to | [0] No [skip to 1108] | [1] Rarely |

| | | | | |
|---|---|---|---|---|
| | | eat of any kind in your household because of lack of resources to get food? | [1] Yes | [2] Sometimes<br>[3] Often |
| **C.1108.** | 1108. | 8. In the past four weeks, did you or any household member go to sleep at night hungry because there was not enough food? | [0] No [skip to 1109]<br>[1] Yes | [1] Rarely<br>[2] Sometimes<br>[3] Often |
| **C.1109.** | 1109. | 9. In the past four weeks, did you or any household member go a whole day and night without eating anything because there was not enough food? | [0] No [skip to 1110]<br>[1] Yes | [1] Rarely<br>[2] Sometimes<br>[3] Often |
| **C.1110.** | 1110. | 10. In the past four weeks, did you or any household member have to borrow rice because you did not have enough rice? | [0] No [skip to 1111]<br>[1] Yes | [1] Rarely<br>[2] Sometimes<br>[3] Often |
| **C.1111.** | 1111. | 11. Compared to other times, does your household face food deficiency during Kartik/chyatra (Monga)? | [0] No [skip to 1112]<br>[1] Yes | [1] No difference<br>[2] Reduced quality<br>[3] Reduced quantity |
| **C.1112.** | 1112. | 12. What would you think is the status of your household in terms of food availability? | | [1] Always deficit<br>[2]Deficit sometimes<br>[3] Adequate (Neither deficit nor surplus)<br>[4]Food surplus |

#### Wash Benefit Module 8

##### SANITATION (All households)

| | |
|---|---|
| SAY | Thank you so much for your participation so far. The next part of the survey is a bit sensitive. I would like to ask you some questions about the sanitation practices of people in your compound, including how you usually dispose of your children's feces. I would also like to make some observations. Are you comfortable with this? IF NOT, EXPLAIN THE IMPORTANCE OF THIS MODULE AND ENCOURAGE RESPONDENT TO PROCEED. |

OPEN DEFECATION (ADMINISTER TO ALL HOUSEHOLDS)

| | | | **Group A** | **B** | **C** | **D** | **E** |
|---|---|---|---|---|---|---|---|
| | WBM 8 | | **Men**<br>পুরুষ | **Women**<br>মহিলা | **Children**<br>**<3 years**<br><৩ বছরের বাচ্চা | **Children**<br>**3-7 years**<br>৩-৭ বছরের বাচ্চা | **Children**<br>**8-15**<br>৮-১৫ ছরের বাচ্চা |
| **C.801.** | 801. | এই খানার কোন সদস্যরা খোলা জায়গাতে পায়খানা করে কী? [Do [GROUP] in this household ever practice open defecation?<br>1 প্রতিদিন Daily<br>2 মাঝেমাঝে Occasionally<br>3 কখনই না Never **(Skip to 806)**<br>88. প্রযোজ্য নয় Not applicable (**806** নং প্রশ্নে যান) **(Skip to 806)**<br>99 জানি না (**806** নং প্রশ্নে যান) | ☐ | ☐ | ☐ | ☐ | ☐ |

| | | | | | | | |
|---|---|---|---|---|---|---|---|
| | | Don't Know **(Skip to 806)** | | | | | |
| **C.802.** | 802. | সাধারনত: একই জায়গাতে প্রতিবার যান কী? [Do [GROUP] go to more or less the same area every time?<br>1 হ্যাঁ Yes<br>2 না No (**805** নং প্রশ্নে যান) **(Skip to 805)**<br>99 জানি না Don't Know / Not Sure (**805** নং প্রশ্নে যান) **(Skip to 805)** | ☐ | ☐ | ☐ | ☐ | ☐ |
| **C.803.** | 803. | আপনার খানা থেকে উক্তস্থানে যেতে (শুধু যাওয়া) কতটুকু সময় লাগে (মিনিট) [How long does it take to walk (one way) from your house to the most commonly visited place? | I_____I MINUTES<br>99 জানি না Don't Know / Not Sure | | | | |
| **C.804.** | 804. | এই জায়গাটি ঐ গ্রামের মধ্যেই কী? [Is that place within the village?<br>1হ্যাঁ Yes<br>2 না No<br>99 জানি না Don't Know / Not Sure | I__I | | | | |

##### Optional Open Defecation Questions

| | | | Children < 3 years | Children 3 – 7 years | Children 8 – 15 years | Men | Women |
|---|---|---|---|---|---|---|---|
| **C.804a** | 805. | What are the main reasons that [GROUP] in your household practice open defecation?<br>☟DO NOT READ RESPONSES MARK ALL THAT APPLY | | | | | |
| | 1 | 1 No choice (nothing else is available) | ☐ | ☐ | ☐ | ☐ | ☐ |
| | 2 | 2 Cannot control where children defecate | | | ☐ | ☐ | ☐ |
| | 3 | 3 Privacy | ☐ | ☐ | ☐ | ☐ | ☐ |
| | 4 | 4 Habit / Routine | ☐ | ☐ | ☐ | ☐ | ☐ |
| | 5 | 5 Prefer to use the bush rather than a toilet | ☐ | ☐ | ☐ | ☐ | ☐ |
| | 6 | 6 Toilet not available at work / school | ☐ | ☐ | ☐ | ☐ | ☐ |
| | 7 | 7 Choose not to share toilets with in-laws / extended family (or cannot) | ☐ | ☐ | ☐ | ☐ | ☐ |
| | 8 | 8 Convenience | ☐ | ☐ | ☐ | ☐ | ☐ |
| | 9 | 9 Safety | ☐ | ☐ | ☐ | ☐ | ☐ |

| | | | | | | | |
|---|---|---|---|---|---|---|---|
| | 10 | 10 Comfort | ☐ | ☐ | ☐ | ☐ | ☐ |
| | 11 | 11 Sickness | ☐ | ☐ | ☐ | ☐ | ☐ |
| | 12 | 12 Latrine overflowed | ☐ | ☐ | ☐ | ☐ | ☐ |
| | 13 | 13 Latrine broken (superstructure and /or slab) | ☐ | ☐ | ☐ | ☐ | ☐ |
| | 14 | 14 Fear of latrine | ☐ | ☐ | ☐ | ☐ | ☐ |
| | 15 | 15 Don't know how to use the latrine | ☐ | ☐ | ☐ | ☐ | ☐ |
| | 16 | 16 Too young to use latrine | ☐ | ☐ | | | |
| | | | **Children < 3 years** | **Children 3 – 7 years** | **Children 8 – 15 years** | **Men** | **Women** |
| **C.X** | 806. | Do you know of other households in the community whose [GROUP] practice OD, even if they might have a toilet or latrine at their house?<br>1 Yes, Often<br>2 Yes, Sometimes<br>3 No, Never<br>99 Don't Know / Not Sure | ☐ | ☐ | ☐ | ☐ | ☐ |
| **C.X** | 807. | Do you see that children's stools are disposed in the yard / surrounding / community in your neighborhood / river?<br>1 Yes, Often<br>2 Yes, Sometimes<br>3 No, Never<br>99 Don't Know | | | | ☐ | ☐ |

#### Sanitation Facility

Administer to: All study households

| | | | |
|---|---|---|---|
| C.805. | 808. | Does your household have a toilet facility that is in use? Can I see it? | 1 Yes have toilet, can observe<br>2 Yes have toilet, refused observation **(Skip to 812)**<br>3 Yes have toilet, cannot observe **(Skip to 812)**<br>4 No toilet facility **(Skip to 812)** |
| C.806. | 809. | 👁 Observation: Note the type, condition and apparent use of the toilet:<br>1 Yes<br>2 No<br>88 Not Applicable / Could not observe / cannot tell | |
| **Exterior observation** | | | |
| 1 | 1 | পায়খানার চারপাশে ৩টি দেয়াল At least 3 walls around the toilet | *(1= হ্যাঁ [Yes], 0= না [No],* প্রযোজ্য নয় [not applicable] =888)] |
| 2 | 2 | Bamboo fences around the toilet | *(1= হ্যাঁ [Yes], 0= না [No],* প্রযোজ্য নয় [not applicable] =888)] |
| 3 | 3 | Door/curtain or walls that guarantee privacy around the toilet | *(1= হ্যাঁ [Yes], 0= না [No],* প্রযোজ্য নয় [not applicable] =888)] |
| 4 | 4 | Roof over toilet | *(1= হ্যাঁ [Yes], 0= না [No],* প্রযোজ্য নয় [not applicable] =888)] |
| 5 | 5 | Ventilation pipe | *(1= হ্যাঁ [Yes], 0= না [No],* প্রযোজ্য নয় [not applicable] =888)] |
| 6 | 6 | পায়খানায় যাবার রাস্তা দেখে বুঝা যাচ্ছে নিয়মিত ব্যবহার করা হয় (পরিষ্কার, জীর্ণ ইত্যাদি ) Path to the toilet suggests regular use (is clear, well-worn, without grass or any barriers etc.) | *(1= হ্যাঁ [Yes], 0= না [No],* প্রযোজ্য নয় [not applicable] =888)] |
| **Interior observation** | | | |
| 7 | 7 | Toilet has a slab | *(1= হ্যাঁ [Yes], 0= না [No],* প্রযোজ্য নয় [not applicable] =888)] |
| 8 | 8 | Raised footing around hole | *(1= হ্যাঁ [Yes], 0= না [No],* প্রযোজ্য নয় [not applicable] =888)] |
| 9 | 9 | Flush or poor Flush: | *(1= হ্যাঁ [Yes], 0= না [No],* প্রযোজ্য নয় [not applicable] =888)] |
| 9a | 9a | If Flush or Pour Flush:<br><br>Water seal condition: | [ 1 ] Functional water seal<br>[ 2 ] ওয়াটার সিল ভাঙ্গা Broken water seal<br>[ 3 ] ওয়াটার সিল নেই No water seal<br>[ 88 ] পায়খানাতে পানির সীল আছে/পর্যবেক্ষণ করা সম্ভব হয়নি N/A / could not observe / cannot tell |
| 9b | 9b | If Flush or Pour Flush: | [ 1 ] Piped sewer system<br>[ 2 ] Septic tank |

| | | | |
|---|---|---|---|
| | | Flushes to:<br><br>*(Ask / probe household members if necessary)* | [ 3 ] Pit latrine (off set)<br>[ 4 ] খাল, ড্রেন, নদী ইত্যাদির সাথে সংযোজন Somewhere else (canal, ditch, river, etc.)<br>[ 88 ] N/A |
| 10 | 10 | Main material of the floor (*select 1*) | [ 1 ] Mud<br>[ 2 ] Wood<br>[ 3 ] Cement<br>[ 4 ] Tile / brick<br>[ 5 ] Plastic<br>[ 88 ] N/A / could not observe / cannot tell |
| 11 | 11 | Bucket toilet | *(1= হ্যাঁ [Yes], 0= না [No], প্রযোজ্য নয় [*not applicable] =888)] |
| 12 | 12 | ঝুলন্ত পায়খানা Hanging toilet | [ 1 Yes ] [ 2 No ] [ 88 N/A] |
| 13 | 13 | Latrine appears to be in use (by your best judgment) | *(1= হ্যাঁ [Yes], 0= না [No], প্রযোজ্য নয় [*not applicable] =888)] |
| 14 | 14 | পায়খানা/টয়লেটে পায়খানার গন্ধ রয়েছে Odor of feces in the latrine/bathroom | *(1= হ্যাঁ [Yes], 0= না [No], প্রযোজ্য নয় [*not applicable] =888)] |
| 15 | 15 | পায়খানা/টয়লেটে প্রস্রাবের গন্ধ রয়েছে Odor of urine in the latrine/bathroom | *(1= হ্যাঁ [Yes], 0= না [No], প্রযোজ্য নয় [*not applicable] =888)] |
| 16 | 16 | স্লাব অথবা মেঝেতে পায়খানা দেখতে পাওয়া গেছে Stool is visible on the slab or floor | *(1= হ্যাঁ [Yes], 0= না [No], প্রযোজ্য নয় [*not applicable] =888)] |
| 17 | 17 | Drop hole is covered | *(1= হ্যাঁ [Yes], 0= না [No], প্রযোজ্য নয় [*not applicable] =888)] |
| 17 a | 17a | If yes, cover is correctly placed over the drop hole so that flies cannot enter/exit | *(1= হ্যাঁ [Yes], 0= না [No], প্রযোজ্য নয় [*not applicable] =888)] |
| 18 | 18 | মাছির উপস্থিতি রয়েছে Flies present | *(1= হ্যাঁ [Yes], 0= না [No], প্রযোজ্য নয় [*not applicable] =888)] |
| 19 | 19 | **General Characteristics**<br>Single or double pit latrine | [ 1 ] Single pit<br>[ 2 ] দুই পিট/গর্ত Double pit<br>[ 88 ] N/A / could not observe / cannot tell |
| 20 | 20 | কমপোষ্টিং টয়লেট Composting toilet | *(1= হ্যাঁ [Yes], 0= না [No], প্রযোজ্য নয় [*not applicable] =888)] |
| **C.807.** | 810. | 👁 Observation: Fullness of the pit – shine a light into the pit to see if solid waste is… | 1 শুকনা ময়লা/পায়খানা ৩ ফুটের চেয়ে বেশী দূরে (>৩ ফুট) Ver far from surface (>1 meter)<br>2 শুকনা ময়লা/পায়খানা ৩ ফুটের মধ্যে (<৩ ফুট) With 1 meter<br>3 পায়খানার খুব কাছাকাছি বা পায়খানা ভরে গেছে Ver close to surface or full |

| | | | |
|---|---|---|---|
| | | | 88 পায়খানাতে পানির সীল আছে/পর্যবেক্ষণ করা সম্ভব হয়<br>Water seal latrine / non direct pit latrine / could not observe |
| C. 808 | 811. | 👁 Observation: What materials for anal cleansing are present inside or immediately outside the latrine? | *(1= হ্যাঁ [Yes], 0= না [No],* প্রযোজ্য নয় *[*not applicable] =888)]<br>1.Leaves/grass<br>2.Twigs / sticks<br>3. Rag or cloth<br>4. Stones<br>5. Hygienic (toilet) paper<br>6. Water container / vessel<br>7. Water tap<br>8. Soap<br>9. Ash or soil for cleansing<br>10. Newspaper |
| C. 809 | 812. | আপনার খানার কে কে এই পায়খানা/টয়লেটটি পায়খানার জন্য ব্যবহার করে? Please tell me about who in your family uses the latrine for defecation. | |
| 1 | 1 | < **3** বাচ্চা Children <3 | 1. সব সময় [Always]<br>2. প্রায়ই [Usually]<br>3. মাঝেমাঝে [Sometime]<br>4. কখনই না [Never]<br>888. প্রযোজ্য নয় [ N/A] |
| 2 | 2 | ৩-৭ বাচ্চা Children 3-7 | 1. সব সময় [Always]<br>2. প্রায়ই [Usually]<br>3. মাঝেমাঝে [Sometime]<br>4. কখনই না [Never]<br>888. প্রযোজ্য নয় [ N/A] |
| 3 | 3 | ৮-১৫ বাচ্চা Children 8-15 | 1. সব সময় [Always]<br>2. প্রায়ই [Usually]<br>3. মাঝেমাঝে [Sometime]<br>4. কখনই না [Never]<br>888. প্রযোজ্য নয় [ N/A] |
| 4 | 4 | পুরুষ **Men** | 1. সব সময় [Always]<br>2. প্রায়ই [Usually]<br>3. মাঝেমাঝে [Sometime]<br>4. কখনই না [Never]<br>888. প্রযোজ্য নয় [ N/A] |
| 5 | 5 | মহিলা **Women** | 1. সব সময় [Always]<br>2. প্রায়ই [Usually]<br>3. মাঝেমাঝে [Sometime]<br>4. কখনই না [Never]<br>888. প্রযোজ্য নয় [ N/A] |
| C. 810 | 813. | অন্য কোন খানা কি এই পায়খানা ব্যবহার করে? Do you share this toilet with other households? | *1=* হ্যাঁ *[Yes]*<br>*0=* না *[No].......skip to 815* |
| C.811 | 814. | কতগুলো খানা মিলে এই পায়খানাটি ব্যবহার করে ? How many HHs use it? | (Don't know=99) |

| | | | |
|---|---|---|---|
| C.811 | 814. | কতগুলো খানা মিলে এই পায়খানাটি ব্যবহার করে ? How many HHs use it? | (Don't know=99)<br>I____I I____I |
| C.812 | 815. | How many people including children in your household use this toilet? | I____I I____I |
| C. 813 | 816. | পায়খানার মালিকানা ? (প্রশ্ন করুন) [***Ask***: Who owns the toilet facility?] | শুধুমাত্র ঐ খানার জন্য [Only for the household] 1<br>কয়েকঘর মিলে/ অংশীদার [Shared] 2<br>অন্য কেউ [Someone else] 3<br>পাবলিক [Public] 4<br>প্রযোজ্য নয় [Not applicable].................................................888 |
| C. 814 | 817. | এই স্থানে এই পায়খানাটি কত বছর ধরে আছে? (জানি না=999) [How long have you had the present latrine in this place? [In years] (Don't [illegible] | A. I_____I I_____I Years<br>B. [illegible] |

### Wash Benefit Module 9

##### Child defecation and feces disposal practice

| | | | |
|---|---|---|---|
| **Child defecation and feces disposal** | | | |
| Administer to: All study households ( সকল খানায় হবে) | | | |
| C.912 | 901. | I_____I I_____I কত বছর বয়স থেকে একটি শিশু সাধারনতঃ টয়লেট/পায়খানা ব্যবহার করা শুরু করে (মাসে)? [At what age (in months) does a child start using a latrine, if at all?] | |
| C.902 | 902. | I_____I আপনার (<৩ বছর) শিশু সর্বশেষ কখন পায়খানা করেছে? (একাধিক শিশু থাকলে সবচেয়ে বড় বাচ্চার ক্ষেত্রে জিজ্ঞেস করুন) [When was the last time your youngest child/infant (<3 years) defecated?] | 1. আজ [Today]<br>2. গতকাল [Yesterday]<br>3. ২ দিন বা তার আগে [Before 2 or more days back]<br>4. কখনই না/ মনে করতে পারছি না [Never/can't remember]....... নং প্রশ্নে চলে যান (skips to 906)<br>5. বলতে রাজি না [Refused] .........**906** নং প্রশ্নে চলে যান (skips to 906)<br>88. প্রযোজ্য নয় [Not applicable]......... **908** নং প্রশ্নে চলে যান (skips 908) |
| **C.903.** | 903. | I_____I আপনার শিশু সর্বশেষ কোথায় পায়খানা করেছে? [Where did the child defecate the last time?]<br>**Note:** উত্তর পড়ে শোনাবেন না, তাকে নির্দিষ্ট করে বলতে বলুন । এবিষয়ে যতক্ষণ পর্যন্ত কিছু বলার থাকে ততক্ষণ পর্যন্ত তাকে বলতে উৎসাহিত করুন [Don't read the answer, encourage by asking if there is anything else until he/she mentions there in nothing else and check all mentioned.] | 1. পটি (উঠানে) [Potty (in the courtyard)]<br>2. পটি (ঘরের ভিতরে) [Potty (inside the house)]<br>3. উঠানে (পটি ছাড়া) [Courtyard (without potty)]<br>4. ঘরের ভিতরে (পটি ছাড়া) [Inside the house (without potty)]<br>5. উঠানের বাহিরে খোলা জায়গাতে [Open space outside the front ya<br>6. ঝোপ-ঝাড়ে/জঙ্গলে Bush/jungle<br>7. পায়খানায়/টয়লেটে [In toilet] ......**906** নং প্রশ্নে চলে যান (Skip to 9<br>8. কাথা/ন্যাপি/ডায়পার Katha/nappy/diaper<br>77. অন্যান্য (বর্ণনা লিখুন) [Other (Please describe)]....................<br>99. জানি না [Don't Know]............. **906** নং প্রশ্নে চলে যান (Skip |

| | | | |
|---|---|---|---|
| | | | 906) |
| **C.904.** | 904. | \|_____\| **†mB cvqLvbv wK Kiv n‡qwQj?**<br>[What was done to the feces?] | **হ্যাঁ [Yes].....1, না [No]......0**<br>1. যেখানে পায়খানা করেছিল সেখানেই ফেলে রাখা হয়েছিল [It is left there]**906** নং প্রশ্নে চলে যান (Skip to 906)<br>2. \|_____\| টয়লেট/পায়খানার ভিতরে ফেলা/ধোয়া হয়েছিল [Put/rinsed into toilet or latrine]<br>3. \|_____\| ড্রেনে/নর্দমার ভিতরে ফেলা/ধোয়া হয়েছিল [Put/rinsed drain or ditch]<br>4. \|_____\|ঝোপ-ঝাড়ে/জঙ্গলে ফেলা হয়েছিল [Thrown into Bush/jungle<br>5. \|_____\| ময়লা আবর্জনার মধ্যে ফেলা হয়েছিল [Thrown into garbage]<br>6. \|_____\| নিদির্ষ্ট গর্তে ফেলা হয়েছিল [Thrown into a specific pit child's feces]<br>7. \|_____\| মাটির নীচে পুতে ফেলা হয়েছিল [Buried]<br>77. \|_____\| অন্যান্য (বর্ণনা লিখুন) [Other (specify)]....<br>99. \|_____\| জানি না [DK]**906** নং প্রশ্নে চলে যান (Skip to 906) |
| **C.905.** | 905. | How did you handle the feces? ☝ Do Not Read Responses. Mark All that Apply (1 = Yes) | ☐ 1 Hands only (bare hands)<br>☐ 2 Hands and cloth / paper / leaves<br>☐ 3 Scrap material to scoop feces<br>☐ 4 Potty<br>☐ 5 Local agricultural hoe/instrument<br>☐ 6 Sani-scoop<br>☐ 7 Did nothing<br>☐ 77 Others (specify)___________<br>☐ 99 Don't know / not sure |
| | 906. | আপনার ৩ বছরের ছোট বাচ্চার পায়খানা কে কে ফেলে? (যতজন ফেলে সবাইকে কোড করুন)<br>[Who disposes your under 3 child's feces? | **হ্যাঁ [Yes].....1, না [No]......0**<br>1. \|_____\| মা [Mother]<br>2. \|_____\| বাবা [Father]<br>3. \|_____\| বোন [Sister]<br>4. \|_____\| ভাই [Brother]<br>5. \|_____\| খালা/মামী/চাচী/ফুফু/দাদী [Aunt/grandmother]<br>6. \|_____\| খালু/মামা/চাচা/ফুফা/দাদা [Uncle/grantfather]<br>7. \|_____\| কেউ পায়খানা ফেলে না [Nobody dispose]<br>77. \|_____\| অন্যান্য (নির্দিষ্ট করে লিখুন) Others (Specify)<br>......................... |
| | 907. | ৩ বছরের ছোট বাচ্চার পায়খানা ফেলার স্থানটি রান্না ঘর থেকে কতটুকু দূরে? [How far away is this disposal site to the kitchen?] | ১. ১০ কদমের কম [Less than 10 steps]<br>২. ১০ কদমের বেশী [More than 10 steps] |
| **C.906.** | 908. | \|_____\| আপনার খানায় ৩-৭ বছরের অন্য কোন শিশু আছে কি? [Is there any other child | 1 হ্যাঁ [Yes]<br>0. না [No].......... **913** নং প্রশ্নে চলে যান (skips to 913) |

| | | | |
|---|---|---|---|
| | | between the age of 3-7 years?] | |
| **C.907.** | 909. | আপনার খানায় ৩-৭ বছরের যে শিশুটি আছে তার বয়স কত (মাসে)? (একাধিক শিশু থাকলে সবচেয়ে বড় বাচ্চার ক্ষেত্রে জিজ্ঞেস করুন) | .......................... মাস |
| **C.908.** | 910. | \|_____\| **আপনার শিশু (৩-৭ বছরের) সর্বশেষ কোথায় পায়খানা করেছে? (একাধিক শিশু থাকলে সবচেয়ে বড় বাচ্চার ক্ষেত্রে জিজ্ঞেস করুন)** [Where did your oldest child (aged 3-7) defecate the last time?] | 1. পটি (উঠানে) [Potty (in the courtyard)]<br>2. পটি (ঘরের ভিতরে) [Potty (inside the house)]<br>3. উঠানে (পটি ছাড়া) [Courtyard (without potty)]<br>4. ঘরের ভিতরে (পটি ছাড়া) [Inside the house (without potty)]<br>5. উঠানের বাহিরে খোলা জায়গাতে [Open space outside the fro yard<br>6. ঝোপ-ঝাড়ে/জঙ্গলে Bush/jungle<br>7. পায়খানায়/টয়লেটে [In toilet] ...**913** নং প্রশ্নে চলে যান (Skip to 913)<br>8. কাথা/ন্যাপি/ডায়পার Katha/nappy/diaper<br>77. অন্যান্য (বর্ণনা লিখুন) [Other (Please describe)]....................<br>99. জানি না [Don't Know]............. **913** নং প্রশ্নে চলে যান (Skip 913) |
| **C.909.** | 911. | \|_____\|সেই পায়খানা কি করা হয়েছিল? [What is done to the feces?] | **হ্যাঁ [Yes].....1, না [No]......0**<br>1. যেখানে পায়খানা করেছিল সেখানেই ফেলে রাখা হয়েছিল [It is left there]<br>2. \|_____\| টয়লেট/পায়খানার ভিতরে ফেলা/ধোয়া হয়েছিল [Put/rinsed into toilet or latrine]<br>3. \|_____\| ড্রেনে/নর্দমার ভিতরে ফেলা/ধোয়া হয়েছিল [Put/rinsed into drain or ditch]<br>4. \|_____\|ঝোপ-ঝাড়ে/জঙ্গলে ফেলা হয়েছিল [Thrown into Bush/jungle<br>5. \|_____\| ময়লা আবর্জনার মধ্যে ফেলা হয়েছিল [Thrown into garbage]<br>6. \|_____\| নির্দিষ্ট গর্তে ফেলা হয়েছিল [Thrown into a specifi pit for child's feces]<br>7. \|_____\| মাটির নীচে পুতে ফেলা হয়েছিল [Buried]<br>77. \|_____\| অন্যান্য (বর্ণনা লিখুন) [Other (specify)]...........<br>99. \|_____\| জানি না [DK]**913** নং প্রশ্নে চলে যান (Skip to 913 |
| **C.910.** | 912. | How did you handle the feces? ☝ <u>Do Not Read Responses. Mark All that Apply</u> (1 = Yes) | ☐ 1 Hands only (bare hands)<br>☐ 2 Hands and cloth / paper / leaves<br>☐ 3 Scrap material to scoop feces<br>☐ 4 Potty<br>☐ 5 Local agricultural hoe/instrument<br>☐ 6 Sani-scoop<br>☐ 7 Did nothing |

| | | | | | |
|---|---|---|---|---|---|
| | | | ☐ | 77 | Others (specify)___________ |
| | | | ☐ | 99 | Don't know / not sure |

| | | | |
|---|---|---|---|
| **Child Potty Use**<br>**( শিশুর পটি ব্যবহার সর্ম্পকীত তথ্য)** | | | |
| Administer to: All study households ( সকল খানায় হবে) | | | |
| C.815 | 913. | Does your household have a potty that children use for defecation? আপনার খানাতে কি শিশুদের পায়খানা করার পটি আছে? | 1 Yes হ্যাঁ<br>2 No না **(Skip to 919)** (919 নং প্রশ্নে যান)<br>99 Don't know / not sure জানি না **(Skip to 919)** (919 নং প্রশ্নে যান) |
| C.816 | 914. | In the last week, how often did your child use the potty? গত এক সপ্তাহের মধ্যে, শিশুটি কতবার পটি ব্যবহার করছে?<br>[ *If multiple children use the potty, ask about the youngest child* ][যদি একাধিক শিশু ব্যবহার করে থাকে তবে সবচেয়ে ছোট শিশুর জন্য প্রশ্ন করুন] | 1 Every time প্রতিবার<br>2 More than half of all defecation events, but n every time যতবার পায়খানা করেছে তার ৫০ ভাগ বা তার বেশী কিন্তু প্রতিবার নয়<br>3 যতবার পায়খানা করেছে তার ৫০ ভাগের কম কিন্তু মাঝেমাঝে Less than half of all defecation events<br>4 আগে করেছি কিন্তু এখন আর করি না Used to use but no longer use it **(Skip to 919)** (919 নং প্রশ্নে যান)<br>5 কখনই না Never **(Skip to 919)** (919 নং প্রশ্নে যান)<br>99 জানি না Don't know / not sure **(Skip to 919)** (9 নং প্রশ্নে যান) |
| C.817 | 915. | What is the age of the child (or children) who is using the potty? যে শিশুটি (বা শিশুরা) পটি ব্যবহার করছে তার বয়স কত? | <u>Mark All that Apply.</u> Yes = 1 হ্যাঁ = 1, না = 0<br>1 ☐ < 1 year ১ বছরের কম<br>2 ☐ 1- < 3 years ১-<৩ বছর<br>3 ☐ 3- < 5 years ৩-<৫ বছর<br>4 ☐ >= 5 years >=৫ বছর |
| C.818 | 916. | Could I please see the potty?<br>👁 Observation: *Record how long it takes to produce the potty.*<br>আমি কি পটিটি দেখতে পারি<br>👁 আমি কি পটিটি দেখতে পারি (পটিটি আনতে কতটুকু সময় লেগেছে তা সেকেন্ডে গণনা করুন) | ☐☐:☐☐MINUTES : SECONDS (99:99 if could not measure) মিনিট : সেকেন্ড (পটি দেখাতে না পারলে 999 কোড করুন) |
| C.819 | 917. | 👁 Observation: Potty condition.<br>পটিটির অবস্থা কি তা পর্যবেক্ষণ করুন (একাধিক উত্তর হতে পারে) | <u>Mark All that Apply.</u> Yes = 1 হ্যাঁ = 1, না = 0<br>1 Easily accessible when needed by the child প্রয়োজনের |

| | | | | |
|---|---|---|---|---|
| | | | | লিড ছাড়া অন্য কিছু দিয়ে ঢাকা ছিল<br>6.Dry শুকনা<br>7.Broken so that it is unusableভেঙ্গে গেছে তাই ইহা ব্যবহার করা সম্ভব না<br>8.Covered in dust / signs of non-use পটি ধুলায় ঢাকা/ ব্যবহার না করার চিহ্ন রয়েছে<br>9.Cannot produce a potty পটি দেখাতে পারে নাই |
| C.820 | 918. | ☐Where do you usually dispose of feces from the potty? এই পটি থেকে আপনি পায়খানা কোথায় ফেলেন? | | 1 Latrine পায়খানায়/টয়লেটে<br>2 Open Pit / separate pit for child or animal fecesখোলা গর্তে / শিশুর বা পশুপাখির পায়খানা ফেলার গর্তে<br>3 Bury it / Covered Pit মাটির নীচে পুতে ফেলা হয়ে ঢাকা গর্তে<br>4 Undefined open site near the compound (including open garbage disposal sites / dumps) উঠানের কাছে খোলা জায়গা<br>5 Bush / forest / fieldঝোপ-ঝাড়ে/জঙ্গলে/মাঠে<br>6 Nearby water (pond, canal, river) পানির উৎসের কাছে (পুকুর/সেচ নালা/ নদী)<br>77 Other (specify) অন্যান্য (লিখুন)____________ |
| **Sani Scoop Use** (সেনিস্কুপ ব্যবহার) | | | | |
| Administer to: All households ( সকল খানায় হবে) | | | | |
| C.821 | 919. | ☐Does your household have a dedicated tool [sani scoop] to clean up feces around your household?<br>আপনার খানার চারপাশের পায়খানা পরিষ্কার করার জন্য আপনার খানায় কোন নির্দিষ্ট কিছু আছে কি? | | 1 Yes হ্যাঁ<br>2 No না (1001 নং প্রশ্নে যান) **(Skip to 1001)**<br>99 Don't know / not sure জানি না (1001 নং প্রশ্নে যান) **(Skip to 1001)** |
| C.822 | 920. | How often do you use the [sani scoop]? আপনি কখন কখন সেনিস্কুপ ব্যবহার করেন? | | 1 Multiple times per day প্রতিদিন কয়েকবার<br>2 Once per day প্রতিদিন একবার<br>3 A few times each week প্রতি সপ্তাহে কয়েকবার<br>4 Less than once per week সপ্তাহে একবারের কম<br>5 Used to use it, but no longer use it প্রথমদিকে ব্যবহার করেছে কিন্তু এখন আর করে না (1001 নং প্রশ্নে যান) **(Skip to 1001)**<br>6 Never কখনই না (1001 নং প্রশ্নে যান) **(Skip to 1001)** |
| C.823 | 921. | What do you use the [sani scoop] for? ☝<u>Do Not Read Responses.</u><br>আপনি কি কি কাজে সেনিস্কুপ ব্যবহার করেন? (একাধিক উত্তর হতে পারে) ☝ উত্তর পড়ে শোনাবেন না | | <u>Mark All that Apply.</u> Yes = 1 হ্যাঁ = 1, না = 0<br>1 ☐ Clean up animal feces জীবজন্তুর পায়খানা ফেলার কাজে<br>2 ☐ Clean up child feces বাচ্চার পায়খানা ফেলার কাজে<br>3 ☐ Clean up garbage ময়লা আবর্জনা ফেলার কাজে<br>4 ☐ Take the scoop to the field (for work) কৃষিকাজে/মাঠে |

| | | | | | |
|---|---|---|---|---|---|
| | | | 5 | ☐ | Digging / gardening গর্ত তৈরীর কাজে |
| | | | 77 | ☐ | Other (specify) অন্যান্য (বর্ণনা লিখুন)______ |
| C.824 | 922. | Could I please see the [sani scoop]? 👁 Observation: Sani scoop condition.<br>আমি কি সেনিস্কুপটি দেখতে পারি? (একাধিক উত্তর হতে পারে)<br>👁 পর্যবেক্ষণ: সেনিস্কুপটির অবস্থা পর্যবেক্ষণ করুন | Mark All that Apply. Yes = 1 হ্যাঁ = 1, না = 0 | | |
| | | | 1 | ☐ | Visible signs of feces on the sani scoop<br>সেনিস্কুপে পায়খানার চিহ্ন আছে |
| | | | 2 | ☐ | Dry শুকনা |
| | | | 3 | ☐ | Broken and needs repair<br>ভেঙ্গে গেছে এবং মেরামত করা দরকার |
| | | | 4 | ☐ | Easily accessible when needed by an adult<br>প্রয়োজনের সময় বড়রা যেন পায় এমন কাছাকাছি জায়গাতে রা |
| | | | 5 | ☐ | Signs that the sani scoop is not used<br>সেনিস্কুপ ব্যবহার না করার চিহ্ন রয়েছে |
| | | | 6 | ☐ | Cannot produce a sani scoop<br>সেনিস্কুপ দেখাতে পারে নাই |

C.825 923.

I'd like to ask a few more questions about the types of feces you pick up with the [sani scoop] and how you dispose of the different types of feces.

[Ask about each type of feces separately.].

সেনিস্কুপ দিয়ে আপনি যেসব পায়খানা ফেলেন এবং কিভাবে ফেলেন সেসম্পর্কে জানার জন্য এখন আমি আপনাকে আরো কিছু প্রশ্ন করতে চাই।

(প্রতিটি পায়খানার ধরনের কথা উল্লেখ করে কোড করুন)

(শিশু/পশুপাখির পায়খানা ফেলার কাজে ব্যবহার না করলে প্রযোজ্য নয় ঘরটিতে টিক দিন)

Where do you dispose the feces?
পায়খানা কোথায় ফেলা হয়েছিল?

✋ Do Not Read Responses. উত্তর পড়ে শোনাবেন না

Mark All that Apply (একাধিক উত্তর হতে পারে) 1 = হ্যাঁ Yes

| | A | B | C | D | E | F |
|---|---|---|---|---|---|---|
| Feces পায়খানা | Use Sani Scoop for disposal?পায়খানা ফেলার জন্য সেনিস্কুপ ব্যবহার করে? | Latrine পায়খানায় | Open Pit খোলা গর্ত | Bush /Farm ঝোপে ঝাড়ে/খোলা জায়গায় | Surface Water পানিতে | Dig Hole and cove গর্ত করা হয় এবং ঢেে |
| 1 Child শিশুর | ☐ | | | | | |
| 2 Cow গোবর | ☐ | | | | | |
| 3 Poultry | | | | | | |

| | | | | |
|---|---|---|---|---|
| | | | / pigeons হাঁস-মুরগীর | |
| | | | 4 Goat ছাগলের | ☐ |
| | | | 5 Pig শুকরের | ☐ |
| | | | 6 Dog or cat কুকুর বা বিড়ালের | ☐ |
| | | | 6 Not applicable প্রযোজ্য নয় | ☐ |

#### Wash Benefit Module 10

##### Water Access

Thank you very much. Now, I would like to ask you some questions about the water you and your family drink at home.

| C.10 WATER TREATMENT, STORAGE, AND QUALITY (All households) | | | | |
|---|---|---|---|---|
| SAY | Thank you very much. Now, I would like to ask you some questions about the water you and your family drink at home. | | | |
| C.1001 | 1001. | How do you store drinking water?<br><br>👁 ASK THE Q ESTION AND OBSERVE IF POSSIBLE | [1] IN PLASTIC OR METAL CONTAINE<br>[2] IN CLAY POTS<br>[3] ROOF TANK OR CISTERN<br>[4] DO NOT STORE WATER | |
| C.1002 | 1002. | Is there a child 0-36 months in this household who drinks water? | [1] YES<br>[2] NO | |
| C.1003 | 1003. | **IF YES** → ASK: If your child 0-3 years wanted a drink of water right now, could you show me how you wo give it to him/her?<br>**IF NO** → ASK: If you wanted a drink of water right now, could you show me how you would get it?<br><br>ATTEMPT TO FOLLOW RESPONDENT WHEN SHE GOES TO RETRIEVE THE WATER. IF WATER IS STORED IN ANOTHER ROOM, ASK HER IF IT IS OKAY TO ENTER THAT ROOM. CIRCLE "99" IF YOU CANNOT OBSERVE. | | |
| | | 👁WHAT DID RESPONDENT DO BEFORE TAKING THE WATER? | YES | NO |
| 2 | 2 | RINSED GLASS/CONTAINER WITH DRINKING WATER BEFORE FILLING | [1] | [2] |
| 3 | 3 | WASHED HANDS WI H WATER BEFORE DRINKING WATER WAS OBTAINED | [1] | [2] |

| | | | | |
|---|---|---|---|---|
| 4 | 4 | WASHED HANDS WITH SOAP BEFORE DRINKING WATER OBTAINED | [1] | [2] |
| | | 👁FROM WHERE DID THE RESPONDENT TAKE THE WATER? | YES | NO |
| 5 | 5 | BROUGHT DIRECTLY FROM WATER SOURCE | [1] | [2] |
| 6 | 6 | BROUGHT IRECTLY FROM STORAGE CONTAINER | [1] | [2] |
| 7 | 7 | BROUGHT DIRECTLY FROM WATER FILTER | [1] | [2] |
| | 8 | STORED WATER WAS COVERED | [1] | [2] |
| | | 👁HOW DID SHE GET THE WATER INTO THE CUP? | YES | NO |
| 9 | 9 | HANDS TOUCHED / CONTACTED THE DRINKING WATER | [1] | [2] |
| 10 | 10 | CONTAINER/GLASS DIPPED INTO WATER CONTAINER | [1] | [2] |
| 11 | 11 | LADLE USED TO OBTAIN WATE | [1] | [2] |
| 12 | 12 | WATER POURED FROM CONTAINER | [1] | [2] |
| 13 | 13 | WATER POURED FROM TAP | [1] | [2] |

| | | | | |
|---|---|---|---|---|
| C.1004 | 1004. | How long ago did you or somebody in your home collect this water? ( 99 = Don't know; 88 = No Water) | A.\|__\|__\|HOURS<br>B.\|__\|__\|DAYS | |
| C.1005 | 1005. | Have you done anything to make this water less cloudy or safer to drink? | [1] YES<br>[2] NO<br>[99] DON'T KNOW/NOT SURE | <br>→ **SKIP to 1007**<br>→ **SKIP to 1007** |
| C.1006 | 1006. | What method(s) did you use?<br>DO NOT READ RESPONSES. PROBE UNTIL RESPONDENT IS FINISHED. CIRCLE ALL THAT APPLY. | | |
| 1 | 1 | Aquatabs | [1] Yes [0] No | |
| 2 | 2 | Waterguard / bottled chlorine | [1] Yes [0] No | |
| 3 | 3 | Boil | [1] Yes [0] No | |
| 4 | 4 | Strain it through clothor other material | [1] Yes [0] No | |
| 5 | 5 | Water filter [ceramic, sand, composite] | [1] Yes [0] No | |
| 6 | 6 | Solar disinfection (SODIS) | [1] Yes [0] No | |
| 7 | 7 | Let it stand and settle | [1] Yes [0] No | |
| 8 | 8 | Biosand filter | [1] Yes [0] No | |
| 9 | 9 | Coagulant (alum) | [1] Yes [0] No | |

| | | | | |
|---|---|---|---|---|
| 10 | 10 | PUR (flocculant + disinfectant) | [1] Yes [0] No | |
| C.1007 | 1007. | Do you ever treat your drinking water or do anything to make it less cloudy? | [1] YES<br>[0] NO | → **SKIP TO 1010** |
| C.1008 | 1008. | When was the last time you treated your water or did anything to make it less cloudy?<br><br>DO NOT READ RESPONSES | [1] TODAY<br>[2] YESTERDAY<br>[3] WITHIN THE PAST WEEK<br>[4] WITHIN THE PAST 2 WEEKS<br>[5] WITHIN THE PAST MONTH<br>[6] WITHIN THE PAST YEAR<br>[99] DON'T KNOW/NOT SURE | |
| C.1009 | 1009. | What are <u>all the ways</u> you treat your drinking water?<br>DO NOT READ RESPONSES. PROBE UNTIL RESPONDENT IS FINISHED. CIRCLE ALL THAT APPLY. | YES | |
| 1 | 1 | Aquatabs | [1] Yes [0] No | |
| 2 | 2 | Waterguard / bottled chlorine | [1] Yes [0] No | |
| 3 | 3 | Boil | [1] Yes [0] No | |
| 4 | 4 | Strain it through clothor other material | [1] Yes [0] No | |
| 5 | 5 | Water filter [ceramic, sand, composite] | [1] Yes [0] No | |
| 6 | 6 | Solar disinfection (SODIS) | [1] Yes [0] No | |
| 7 | 7 | Let it stand and settle | [1] Yes [0] No | |
| 8 | 8 | Biosand filter | [1] Yes [0] No | |
| 9 | 9 | Coagulant (alum) | [1] Yes [0] No | |
| 10 | 10 | PUR (flocculant + disinfectant) | [1] Yes [0] No | |

| | | | |
|---|---|---|---|
| **DRINKING WATER SOURCES (ALL HOUSEHOLDS)** | | | |
| ASK THE FOLLOWING QUESTIONS ABOUT THE RESPONDENT'S PRIMARY AND SECONDARY WATER SOURCES. | | | |
| C.1016 | 1010. | What type of water source does your household collect most of the water that you use from? | 1 Shallow tubewell<br>2 Deep tubewell<br>3 Piped into home<br>4 Piped into yard<br>5 Borewell in yard<br>6 Public tap<br>7 Public borewell<br>8 Shallow well with concrete reinforcement<br>9 Shallow well (no concrete)<br>10 River, lake, stream<br>11 Other: _____________ |
| | [illegible] | Can you tell me how long it takes you | \| \|Hours |

| | | | |
|---|---|---|---|
| | | to walk one-way to this PRIMARY water source from your home? | |__|Min |
| C.1018 | 1012. | Can you please tell me whether you use water from this source for… | |
| 1 | 1 | Drinking | [1] Yes [0] No |
| 2 | 2 | Cooking/ washing dishes | [1] Yes [0] No |
| 3 | 3 | Laundry | [1] Yes [0] No |
| 4 | 4 | Bathing/hand washing | [1] Yes [0] No |
| 5 | 5 | Irrigate crop land or garden | [1] Yes [0] No |
| 6 | 6 | Water for livestock | [1] Yes [0] No |
| C.1019 | 1013. | Does your household pay any money to use this source? | 1 Yes 0 No |
| C.1020 | 1014. | Does your household currently use a secondary water source? If so, what type is it? | 1 Shallow tubewell<br>2 Deep tubewell<br>3 Piped into home<br>4 Piped into yard<br>5 Borewell in yard<br>6 Public tap<br>7 Public borewell<br>8 Shallow well with concrete reinforcement<br>9 Shallow well (no concrete)<br>10 River, lake, stream<br>11 Other: _____________ |
| C1021 | 1015. | Can you tell me how long it takes you to walk one-way to this SECONDARY water source from your home? | |__|Hours<br>|__|Min |
| C1022 | 1016. | Can you please tell me whether you use water from this source for… | |
| 1 | 1 | Drinking | [1] Yes [0] No |
| 2 | 2 | Cooking/ washing dishes | [1] Yes [0] No |
| 3 | 3 | Laundry | [1] Yes [0] No |
| 4 | 4 | Bathing/hand washing | [1] Yes [0] No |
| 5 | 5 | Irrigate crop land or garden | [1] Yes [0] No |

| | | | |
|---|---|---|---|
| 6 | 6 | Water for livestock | [1] Yes [0] No |
| C1023 | 1017. | Does your household pay any money to use this source? | [1] Yes [0] No |
| C1024 | 1018. | Does your household collect rainwater? | [1] Yes [0] No |
| C1025 | 1019. | Is your household currently collecting rainwater? | [1] Yes [0] No |
| C1026 | 1020. | Can you please tell me whether you use rainwater for | |
| 1 | 1 | Drinking | [1] Yes [0] No |
| 2 | 2 | Cooking/ washing dishes | [1] Yes [0] No |
| 3 | 3 | Laundry | [1] Yes [0] No |
| 4 | 4 | Bathing/hand washing | [1] Yes [0] No |
| 5 | 5 | Irrigate crop land or garden | [1] Yes [0] No |
| 6 | 6 | Water for livestock | [1] Yes [0] No |
| **C.1027.** | 1021. | Thinking about ALL the water that your household obtains from ALL sources, approximately how many 20-litre containers of water would you say that your household uses IN TOTAL EACH DAY?<br><br>***Encourage respondent to estimate. "Don't know"=999. Use decimals as needed (e.g. one half=0.5) | ☐10-liter containers per day<br><br>☐20-liter containers per day |
| **C.1010.** | 1022. | ☐How frequently do you treat your water using [Aquatabs]? | Do Not Read Responses.<br>1 Every time they collect water<br>2 Sometimes / occasionally<br>3 Treated water in the beginning [of the program] but not any more<br>4 Never treated water with [Aquatabs] **(Ski to 1026)**<br>99 Don't know / not sure |
| **C.1011.** | 1023. | ☐Is the drinking water stored in your household today treated with [Aquatabs]? | 1 Yes, all of it<br>2 Yes, some of it<br>3 Not treated |

| | | | |
|---|---|---|---|
| | | | 4 No water in the house<br>99 Don't know |
| C.1012. | 1024. | ☐N Ask to see the treated water. Observe: Is the water covered? | 1 Yes<br>2 No<br>3 No water stored in the house<br>88 Not applicable / refused |
| C.1013. | 1025. | Approximately how long ago did you treat the water with [Aquatabs]? | HH:MM ☐☐:☐☐ ( 9 9 : 9 9 Don't know ) |

**Residual Chlorine Test**

Administer to: All study households

*Ask to collect a water sample from the source identified in C.1003. Explain that some (but not all) of the samples may be tested to see if there is any chlorine in the water.*

| | | | |
|---|---|---|---|
| C.101 | 1026. | ☐ May I collect a small sample of your drinking water? | 1 Yes<br>2 No / refused<br>3 No drinking water available to test |
| C.101 | 1027. | Collect a small water sample from stored drinking water for target children.<br>Test for free residual chlorinein a discrete location after you leave the household.<br>Only test samples collect from households who report to have treated their water with some form of chlorine (C.1009 = 1, 2, or 11).If the household did not report treating the water with chlorine, discard the sample without testing it and record 8 88 in the field below. | 8 88 Did not report that the water was treate with chlorine<br>9 9 9 Could not test<br>Level of Free Residual Chlorine: ☐☐.☐mg / L |

| | | | | |
|---|---|---|---|---|
| | | **HOUSEHOLD MICROBIOLOGICAL SAMPLE (ADMINISTER TO RANDOM SUBSAMPLE HOUSEHOLD)** | | |
| C.1501 | 1028. | May I collect a small sample of your drinking water to test for bacteria in our office? | [1] YES, SAMPLE COLLECTED<br>[2] NO/REFUSED | → **SKIP TO C.1033** |
| | 1029. | | HH Water Sample Time Log (can be automated i a bar-code scanner) | |

| | DD/MM/YY | HH:MM |
|---|---|---|

| | | | | | |
|---|---|---|---|---|---|
| | | | A | □□/□□/□□<br>HH sample collected | □□:□□ |
| | | | B | □□/□□/□□<br>HH sample arrived at lab | □□:□□ |
| | | | C | □□/□□/□□<br>HH sample analyzed | □□:□□ |

| | | | | |
|---|---|---|---|---|
| c.1503 | 1030. | HH sample, volume of water filtered (in milliliters) | □□□ ml | |
| C.1504 | 1031. | Household sample lab result, *E. coli* (5555 = Too numerous to count, 9999 = Not analyzed) | □□□□ (CFU / 100 ml) | |
| C.1505 | 1032. | HH sample qualifier | 1. Below detection limit<br>2. Above detection limit<br>3. Sample could not be analyzed | |
| | | **** WRITE THE CLUSTER/HH ID <u>VERY CLEARLY</u> ON THE WHIRLPACK BEFORE PROCEEDING**** | | |
| | | PAIRED SOURCE WATER SAMPLE FOR HH WATER SAMPLE<br>IF RESPONDENT REPORTED COLLECTING WATER FROM A <u>SHARED SOURCE</u> → **SKIP TO 701** | | |
| C.1506 | 1033. | IF RESPONDENT REPORTED COLLECTING WATER FROM A <u>PRIVATE WATER SOURCE</u> AND THAT WATER SOURCE IS <u>WITHIN THE COMPOUND</u> (INCLUDING RAINWATER), ASK :<br><br>May I collect a small sample of your source water to test for bacteria in our office?<br><br>RECORD WHETHER SAMPLE WAS COLLECTED: | [1] YES, SAMPLE COLLECTED<br>[2] NO, REFUSED | → **SKIP TO 701** |
| C.1507 | 1034. | Water source ID<br>If Source is a shared source, list the source ID.<br>If Source is located in the compound, only list the cluster/HH ID, and record "99" in the Source ID.<br>If the shared source located is not on the preprinted list, enter "00" | A □□□ / □□ Cluster / HH ID<br>B □□ Source ID (if a shared source) | |

| | | | |
|---|---|---|---|
| | 1035. | Source sample collected (DD/MM/YY, HH:MM) | □□/□□/□□ □□:□□ |
| C.1508 | 1036. | 👁 Record the source water type | 1 Shallow tubewell<br>2 Deep tubewell<br>3 Piped into home<br>4 Piped into yard<br>5 Borewell in yard<br>6 Public tap<br>7 Public borewell<br>8 Shallow well with concrete reinforcement<br>9 Shallow well (no concrete)<br>10 River, lake, stream<br>11 Other: ______________ |
| | | ** **WRITE THE SOURCE ID <u>VERY CLEARLY</u> ON THE WHIRLPACK BEFORE PROCEEDING*** | |

#### Wash Benefit Module 7

##### HANDWASHING

| | WBM 7 | | |
|---|---|---|---|
| C.700 | 700. | RECORD WHETHER THE RESPONDENT HAS WASHED HER HANDS AT ANY TIME BEFORE THIS QUESTION | [1] OBSERVED RESPONDENT WASHING HANDS<br>[2] DID NOT OBSERVE RESPONDENT WASHING HANDS |
| C.701 | 701. | READ: Thank you. Now, I would like to do a quick inspection of your hands. I hope you don't mind. Can you please show me your hands?<br><br>BOTH HANDS SHOULD BE SHOWN (NOT JUST ONE HAND). USE <u>APPEARANCE CODES</u> BELOW TO RECORD THE DESCRIPTION THAT BEST DESCRIBES THE DEFINITIONS THE LEVEL OF CLEANLINESS.<br><u>APPEARANCE CODES:</u><br>[1] VISIBLE DIRT (DIRT/MUD/SOIL/ASH OR ANY OTHER MATERIAL IS VISIBLE)<br>[2] UNCLEAN APPEARANCE (NO DIRT IS VISIBLE ON THIS PART OF THE HAND BUT, IN GENERAL, THIS PART OF THE HAND APPEARS UNCLEAN)<br>[3] CLEAN (OBSERVED PART OF | 👁 MOTHER<br><br>**Left Hand**<br>A\|__\|FINGERNAILS<br>B\|__\| PALMS<br>C\|__\|FINGER PADS<br><br>**Right Hand**<br>D\|__\|FINGERNAILS<br>E\|__\| PALMS<br>F\|__\|FINGER PADS |

| | | | |
|---|---|---|---|
| C.701 | 701. | READ: Thank you. Now, I would like to do a quick inspection of your hands. I hope you don't mind. Can you please show me your hands?<br><br>BOTH HANDS SHOULD BE SHOWN (NOT JUST ONE HAND). USE APPEARANCE CODES BELOW TO RECORD THE DESCRIPTION THAT BEST DESCRIBES THE DEFINITIONS THE LEVEL OF CLEANLINESS.<br>APPEARANCE CODES:<br>[1] VISIBLE DIRT (DIRT/MUD/SOIL/ASH OR ANY OTHER MATERIAL IS VISIBLE)<br>[2] UNCLEAN APPEARANCE (NO DIRT IS VISIBLE ON THIS PART OF THE HAND BUT, IN GENERAL, THIS PART OF THE HAND APPEARS UNCLEAN)<br>[3] CLEAN (OBSERVED PART OF THE HAND IS CLEAN AS WOULD APPEAR AFTER SOMEONE WASHES HANDS OR TAKES A BATH)<br>N/A; OBSERVATION NOT POSSIBLE/REFUSED | 👁 MOTHER<br><br>**Left Hand**<br>A\|__\|FINGERNAILS<br>B\|__\| PALMS<br>C\|__\|FINGER PADS<br><br>**Right Hand**<br>D\|__\|FINGERNAILS<br>E\|__\| PALMS<br>F\|__\|FINGER PADS |
| **C.701** | 702. | ASK: Please show me [NAME]'s hands.<br><br>ASK TO SEE THE HANDS OF THE RESPONDENT'S OLDEST CHILD 0-36 MONTHS (SEE C.1) IF RESPONDENT DOESN'T HAVE A CHILD 0-36 MONTHS → **SKIP TO NEXT PAGE**.<br><br>APPEARANCE CODES:<br>[4] VISIBLE DIRT (DIRT/MUD/SOIL/ASH OR ANY OTHER MATERIAL IS VISIBLE)<br>[5] UNCLEAN APPEARANCE (NO DIRT IS VISIBLE ON THIS PART OF THE HAND BUT, IN GENERAL, THIS PART OF THE HAND APPEARS UNCLEAN)<br>[6] CLEAN (OBSERVED PART OF THE HAND IS CLEAN AS WOULD APPEAR AFTER SOMEONE WASHES HANDS OR TAKES A BATH)<br>[88] N/A; OBSERVATION NOT POSSIBLE/REFUSED/NO CHILD 0-36 MONTHS | 👁 OLDEST CHILD 0-36 MONTHS<br><br>**Left Hand**<br>A\|__\|FINGERNAILS<br>B\|__\| PALMS<br>C\|__\|FINGER PADS<br><br>**Right Hand**<br>D\|__\|FINGERNAILS<br>E\|__\| PALMS<br>F\|__\|FINGER PADS |

| | | | |
|---|---|---|---|
| C.702c | 704. | পর্যবেক্ষনঃ প্রাথমিক/ **প্রধান** হাত ধোয়ার স্থানটিতে কোন কোন উপাদান আছে? (আপনি যদি দেখে থাকেন তাহলে "1"কোড করুন আর না দেখলে"0" কোড করুন) [Observation: Which of the following are present at the primary hand washing station? (If you observe the listed item, write "1" for "yes" in the box below. If you do not observe the listed item, write "0" for "no" in the box below.)]<br>*1= হ্যাঁ [Yes], 0= না [No]* | MATERIALS PRESENT<br><br>[1] **পানি** WATER<br>[2] **গোসল/হাত ধোয়ার সাবান (লাক্স, লাইফবয়)** BAR SOAP (Body/han<br>[3] **অন্য যে কোন ধরনের সাবান (হুইল)** BAR SOAP (other)<br>[4] **গুড়া পাউডার** POWDERED SOAP<br>[5] **সাবান-পানি** SOAPY WATER<br>[6] **তরল সাবান** LIQUID SOAP<br>[7] **ছাই** Ash<br>[8] **মাটি /বালু** Mud/Sand<br>[9] **আইসিডিডিআর, বি-র দেওয়া ট্যাপসহ ড্রাম** ICDDR,B drum with t<br>[10] হাত ধোয়ার স্থানটি ভিজা ছিল/ব্যবহারের চিহ্ন ছিল Moisture below handwas station<br>[11] (**বালতি** Bucket<br>[12] **বেসিন, জগ** Basin/Jug<br>[13] **কিছুই নাই** NONE OF THE ABOVE<br>**[**777] **অন্যান্য** OTHER (SPECIFY): _____________ |
| | 705. | **প্রধান হাত ধোয়ার স্থানটি সাবান জাতীয় যেকোন ধরনের উপাদান (যা হাত ধোয়ার কাজে ব্যবহার করা হয়/ সাবান পানি ও পানি একসাথে উপস্থিত আছে কী?**<br>[Soap/soapy water and water present together in PHWS] | 1. হ্যাঁ [Yes]<br>0. না [No] |
| C.702c | 706. | প্রাথমিক/ **প্রধান** হাত ধোয়ার স্থানটি রান্নাঘর থেকে কত কদম দূরে? [Observation: The primary hand washing station is away from the kitchen---Count in steps, allow in continuous numbers] | \|____\| |
| C.702d | 707. | প্রাথমিক/ **প্রধান** হাত ধোয়ার স্থানটি পায়খানা থেকে কত কদম দূরে? [Observation: The primary hand washing station is away from the toilet---Count in steps, allow in continuous numbers] | \|____\| |
| C.703 | 708. | উত্তরদাতাকে প্রশ্ন করুনঃ আপনার হাতধোয়ার জন্য প্রাথমিক/ **প্রধান** স্থান ছাড়া অন্যকোন জায়গা আছে কি? [Ask the respondent: "Is there anywhere else you wash your hands?" (Record code in box)]: | 1. হ্যাঁ [Yes]<br>0. না [No] **714** নং প্রশ্নে চলে যান (skips to 714) |
| C.703a | 709. | পর্যবেক্ষনঃ দ্বিতীয় প্রধান হাত ধোয়ার স্থানটি কোথায় তা রেকর্ড করুন [Observation: Record the location of the secondary hand washing station]. | 1. ঘরের ভিতরে/ কাছে (≤৬ কদম) In/near main house (≤6 feet to entrance)<br>2. পায়খানার ভিতরে/ কাছে (≤৬ কদম) In/near latrine (≤6 feet to entra<br>3. রান্নার স্থানে ভিতরে/ কাছে (≤৬ কদম) In/near cooking area (≤6 feet entrance)<br>4. ঘর, পায়খানা, রান্নার স্থানে থেকে >৬ কদম >6 feet away from main hc latrine <u>and</u> cooking area |

| | | | |
|---|---|---|---|
| C.703c | 710. | পর্যবেক্ষনঃ দ্বিতীয় প্রধান হাত ধোয়ার স্থানটিতে কোন কোন উপাদান আছে? (আপনি যদি দেখে থাকেন তাহলে "1"কোড করুন আর না দেখলে"0" কোড করুন) [Observation: Which of the following are present at the secondary hand washing station? (If you observe the listed item, write "1" for "yes" in the box below. If you do not observe the listed item, write "0" for "no" in the box below.) ] | *1= হ্যাঁ [Yes], 0= না [No]*<br>[1] পানি WATER<br>[2] গোসল/হাত ধোয়ার সাবান (লাক্স, লাইফবয়) BAR SOAP (Body/han<br>[3] অন্য যে কোন ধরনের সাবান (হুইল) BAR SOAP (other)<br>[4] গুড়া পাউডার POWDERED SOAP<br>[5] সাবান-পানি SOAPY WATER<br>[6] তরল সাবান LIQUID SOAP<br>[7] ছাই Ash<br>[8] মাটি /বালু Mud/Sand<br>[9] আইসিডিডিআর, বি-র দেওয়া ট্যাপসহ ড্রাম ICDDR,B drum with t<br>[10] হাত ধোয়ার স্থানটি ভিজা ছিল/ব্যবহারের চিহ্ন ছিল Moisture below handwas station<br>[11] (বালতি Bucket<br>[12] বেসিন, জগ Basin/Jug<br>[13] কিছুই নাই NONE OF THE ABOVE<br>অন্যান্য OTHER (SPECIFY): ______________ |
| | 711. | **দ্বিতীয় প্রধান হাত ধোয়ার স্থানটিতে সাবান জাতীয় যেকোন ধরনের উপাদান (যা হাত ধোয়ার কাজে ব্যবহার করা হয়/সাবান পানি ও পানি একসাথে উপস্থিত আছে কী?** [Soap/soapy water and water present together in SHWS] | 1. হ্যাঁ [Yes]<br>0. না [No] |
| C.703c | 712. | দ্বিতীয় প্রধান হাত ধোয়ার স্থানটি রান্নাঘর থেকে কত কদম দূরে? [Observation:T he secondary hand washing station is away from the kitchen---Count in steps, allow in continuous numbers]] | \|____\| |
| C.703d | 713. | দ্বিতীয় প্রধান হাত ধোয়ার স্থানটি পায়খানা থেকে কত কদম দূরে? [Observation: The secondary hand washing station is away from the toilet---Count in steps, allow in continuous numbers] | \|____\| |
| INDICATOR 4: HANDWASHING DEMONSTRATION, SOAP PRESENCE | | | |
| Thank you. Please show me where you most often wash your hands after defecation. (Note: this may be the same place you already observed. Go to the place identified by the respondent and record location). | | | |
| C.704a | 714. | 👁OBSERVE AND RECORDLOCATION OF HANDWASHING (CIRCLE ONE) | 1. ঘরের ভিতরে/ কাছে (≤৬ কদম) In/near main house (≤6 feet t entrance)<br>2. পায়খানার ভিতরে/ কাছে (≤৬ কদম) In/near latrine (≤6 feet to entrance)<br>3. রান্নার স্থানে ভিতরে/ কাছে (≤৬ কদম) In/near cooking area (≤6 to entrance)<br>4. ঘর, পায়খানা, রান্নার স্থানে থেকে >৬ কদম >6 feet away from m house, latrine <u>and</u> cooking area<br>**5.** নির্দিষ্ট কোন জায়গায় নাই |
| **PREPARE YOUR STOP WATCH FOR TIMING.** | | | |
| C.705 | 715. | Now, can you demonstrate how you | [1] DEMONSTRATED |

| | | | |
|---|---|---|---|
| C.705 | 715. | Now, can you demonstrate how you normally clean your hands <u>after defecation</u>? Try in the same manner as you would if I wasn't here.<br>TIME HOW LONG IT TAKES THE RESPONDENT TO BRING MATERIALS TO THAT PLACE (START AT THE MOMENT YOU FINISH ASKING THE QUESTION). WHEN RESPONDENT STARTS CLEANING HER HANDS, TIME DURATION OF RUBBING HANDS TOGETHER. | [1] DEMONSTRATED<br><br>[2] COULD NOT DEMONSTRATE<br>IF NOT → **RECORD REASON:**<br>**______________________________________AND <u>SKIP TO 721</u>** |
| C.705b | 716. | 👁OBSERVE AND RECORD TIME TO PRODUCE (Bangladesh: <u>SOAP)</u> FOR CLEANING HANDS<br>MARK "00:00" IF SOAP IS ALREADY PRESENT AT HANDWASHING PLACE. (Bangladesh ONLY) MARK "88:88" IF SOAP IS N T USED | \|____:____\| mm:ss |
| C.705c | 717. | 👁OBSERVE AND RECORD LENGTH OF TIME RESPONDENT SPENDS <u>RUBBING HER HANDS</u> TOGETHER: | \|____:____\| mm:ss |
| C.705d | 718. | 👁OBSERVE AND RECORD <u>ALL MATERIALS</u> USED FOR HAND CLEANSING (CIRCLE ALL THAT APPLY)<br>1 Yes 0 No | 1. WATER<br>2. BAR SOAP (Body/hand BAR)<br>3. BAR SOAP (other)<br>4. POWDERED SOAP<br>5. SOAPY WATER<br>6. CLOTH OR LEAVES<br>7. BASIN/PITCHER<br>8. ASH<br>9. MUD<br>88.OTHER (SPECIFY): __________________________ |
| C.705e | 719. | 👁OBSERVE AND RECORD WHETHER BOTH HAN S WERE CLEANED | [1] YES, BOTH HANDS CLEANED<br>[2] NO, ONLY ONE HAND CLEANED |
| C.705f | 720. | 👁OBSERVE AND RECORD HOW RESPONDENT DRIES HANDS 1 Yes 0 No | 1.DRIES BY WIPING HER HANDS ON HER CLOTHES 2.DRIES BY WIPING HANDS ON ANOTHER 3.CLOTH DRIES BY WIPING HER HANDS ON 4.MATERIAL (NOT CLOTH)<br>5. DRIES BY SHAKING HER HANDS IN THE AIR<br>6.NOT DRIED<br>88.OTHER, SPECIFY_______________________ |
| C.705g | 721. | IF RESPONDENT <u>COULD NOT DEMONSTRATE</u>, OR <u>DID NOT USE SOAP</u>, ASK: | |

| | | | | | |
|---|---|---|---|---|---|
| C.705g | 721. | IF RESPONDENT COULD NOT DEMONSTRATE, OR DID NOT USE SOAP, ASK:<br>Do you have soap in your house that you use for handwashing? Can you bring it to me?<br><br>👁OBSERVE AND RECORD TIME TO PRODUCE SOAP FOR WASHING HANDS<br>MARK "88:88" IF ALREADY TIMED IN C.704a<br>MARK "99:99" IF OBSERVATION NOT POSSIBLE OR SOAP NOT AVAILABLE | \|____:____\| mm:ss | | |
| C. 706a | 722. | 👁OBSERVE: WAS THERE ANY OBVIOUS REACTIVITY DURING THE COURSE OF HIS DEMONSTRATION? | [1] YES<br>[0] NO → skip to 723 | | |
| b | 1 | Longer time spent cleaning/rubbing hands | [1] YES<br>[0] NO | | |
| c | 2 | Using soap where they otherwise would not | [1] YES<br>[0] NO | | |
| d | 3 | Other, (BRIEFLY explain) ________________________ | [1] YES<br>[0] NO | | |
| | | | ___________ | | |
| C. 707 | 723. | ASK: "Please tell me about all of the times you wash your hands with soap."<br>CIRCLE "1" IN COLUMN A IF CRITICAL TIME IS MENTIONED WITHOUT PROMPTING.<br><br>AFTER THE RESPONDENT FINISHES NAMING ALL THE TIMES ASK "IS THERE ANY OTHER TIME YOU WASH YOUR HANDS WITH SOAP?" AND STOP WHEN THE RESPONDENT SAYS THERE IS NO OTHER TIME.<br><br>FOR EACH TIME MENTIONED WITHOUT PROMPTING ASK COLUMN B.<br><br>FOR QUESTIONS C.707A-G WHERE [1] IS NOT CIRCLED IN COLUMN A, PROMPT BY ASKING:<br>Do you usually wash your hands with soap and water [CRITICAL TIME]? IF THE RESPONDENT SAYS "YES" THEN IMMEDIATELY ASK HOW OFTEN AND MARK APPROPRIATE ANSWER | A.<br>MENTIONED WITH NO PROMPTING | B.<br>Only for times mentioned with NO PROMPTING, ask: How often do you wash your hands with soap and water [insert event]?<br><br>[1] ALWAYS<br>[2] SOMETIMES<br>[3] RARELY | C.<br>REPLIED "A BEING PROM<br><br>[1] ALWAYS<br>[2] SOMETIME<br>[3] RARELY<br>[4] NO |

| | | |
|---|---|---|
| d | 7 | BEFORE FEEDING A CHILD |
| e | 8 | AFTER CLEANING A CHILD'S ANUS |
| f | 9 | AFTER DISPOSING OF CHILDREN'S FECES |
| g | 10 | AFTER TDEFECATION |
| h | 11 | AFTER RETURNING FROM OUTSIDE THE COMPOUND |
| m | 12 | AFTER TOUCHING A SICK PERSON |
| o | 13 | AFTER HANDLING LIVESTOCK |
| | 14 | AFTER HANDLING COW-DUNG (OR ANY ANIMAL FECES) |
| j | 15 | OTHER (SPECIFY): ________________________________________ |
| i | 16 | RESPONDENT NEVER WASHES HANDS WITH SOAP |

#### Wash Benefit Module 19

#### PARENTAL STRESS

#### Wash Benefit Module 17

#### MATERNAL INTELLIGENCE

#### Wash Benefit Module 12

##### HOME CARE ENVIRONMENT

asania 1/30/12 5:09 PM
Comment: N/A FOR BASELINE

#### Wash Benefit Module 13

##### MEASURES FOR SPILLOVER

Administer to: All study households

| | | | |
|---|---|---|---|
| **C.1301.** | 1301. | What is the name of your nearest health facility?<br>Enter "99" if Don't know / not sure | |
| **C.1301.a** | 1302. | In the past year, (since this time last year) approximately how many times did you visit this health facility for any reason (including with your children)? | \|__\|__\| TIMES |
| **C.1301.b** | 1303. | Approximately how long does it take to walk there (in minutes, one-way, from where we are now)?<br><br>Enter "999" if Don't know / not sure | \|__\|__\|__\| MINUTES |
| **C.1302.** | 1304. | What is the name of the ECD center [preschool] where your children go (or will go when they are old enough)? | |
| **C.1303.** | 1305. | What is the name of the primary school where your children go (or will go when they are old enough)? | |
| **C.1304.** | 1306. | What is the name of the market that you most often go to when you buy or sell things? | |
| **C.1304.a** | 1307. | How often do you go to that market? | 0 Never<br>1 Once per week or more<br>2 Once every 2 weeks<br>3 Once every 3 weeks<br>4 Once every 4 weeks (every month)<br>5 Less than once every month<br>99 Don't know / not sure |
| **C.1304.b** | 1308. | Approximately how long does it take to walk to the market (in minutes, one-way, from where we are now)?<br><br>Enter "999" if Don't know / not sure | \|__\|__\|__\| MINUTES |
| **C.1305.** | 1309. | What is the name of the church/mosque/temple (if any) that you attend? | |
| **C.1305.a** | 1310. | How often do you go to church? | 0 Never<br>1 Once per week or more<br>2 Once every 2 weeks<br>3 Once every 3 weeks<br>4 Once every 4 weeks (every month)<br>5 Less than once every month |

| | | | |
|---|---|---|---|
| | | | 99 Don’t know / not sure |
| **C.1305.b** | 1311. | Approximately how long does it take to walk to the church (in minutes, one-way, from where we are now)?<br><br>Enter “999” if Don’t know / not sure | |__|__|__| MINUTES |
| **C.1306.** | 1312. | Confirm that you have recorded a GPS location linked to this household | [ 1 ] Confirmed, GPS point recorded<br>[ 99 ] Could not record |

#### Wash Benefit Module 41

##### Household Assets

asania 2/9/12 10:32 AM
**Comment:** MOVE TO THE END OF THE INTERVIEW WITH THE HOUSEHOLD BECAUSE THIS DOESN’T INVOLVE THE RESPONDENT

<table>
<tr><td colspan="3">Thank you very much for your time. Now I would talk about <u>the house that you live in</u>. First, I would like to observe the material your house is made of. Can I take a look at your house?<br><br>OBSERVE MAIN MATERIAL OF THE HOUSE:</td></tr>
<tr><td></td><td>4101.</td><td>ছাদ তৈরীতে কি কি উপাদান ব্যবহার করা হয়েছ ? (পরির্দশন করে যাচাই করুন) [Main material of the roof]<br>কাঁচা (বাঁশ/খড়) [Kaccha (bamboo / thatch)] 1<br>টিন [Tin] 2<br>সিমেন্ট/ কংক্রিট/ টালি [Cement / concrete / tiled]…3</td></tr>
<tr><td></td><td>4102.</td><td>দেয়াল তৈরীতে কি কি উপাদান ব্যবহার করা হয়েছ ? (পরির্দশন করে যাচাই করুন) [Main material of the walls]<br>পাট/বাঁশ/মাটি (কাঁচা) [Jute / bamboo / mud (kaccha)] 1<br>কাঠ [Wood] .......................................... 2<br>ইট/সিমেন্ট [Brick / cement] ..................... 3<br>টিন [Tin]................................................ 4</td></tr>
<tr><td></td><td>4103.</td><td>মেঝে তৈরীতে কি কি উপাদান ব্যবহার করা হয়েছ ? (পরির্দশন করে যাচাই করুন) [Main material of the floor]<br>মাটি/বাঁশ (কাঁচা) [Earth / bamboo (kaccha)] ......... 1<br>কাঠ [Wood] 2<br>ইট/ সিমেন্ট [Cement / concrete] .......................... 3</td></tr>
<tr><td colspan="3">Thank you. Now I would like to ask you some questions about your household assets.</td></tr>
<tr><td></td><td>4104.</td><td>খানার ঘরটিতে কতগুলো কক্ষ আছে? (রান্নাঘর ও বাথরুম বাদে) [How many rooms the households have (exclude bathroom and Kitchen)?].</td></tr>
<tr><td></td><td>4105.</td><td><u>হ্যাঁ (Yes)....1, না (No)....0, জানি না (DK).....999</u><br>a. |_____|বিদ্যুৎ [Electricity]<br>b. |_____||_____|আলমারী/ওয়ারড্রব (সংখ্যা) [Number of Almirah or wardrobe]<br>c. |_____||_____|টেবিল (সংখ্যা) [Number of tables]<br>d. |_____||_____|চেয়ার/বেঞ্চ (সংখ্যা) [Number of chair or bench]<br>e. |_____||_____|হাত ঘড়ি/দেয়াল ঘড়ি (সংখ্যা) [Number of watch or clock]<br>f. |_____||_____|খাট (সংখ্যা) [Number of khat]<br>g. |_____||_____|চৌকি (সংখ্যা) [Number of chouki ]<br>h. |_____|রেডিও [A radio that is working]<br>i. |_____|টেলিভিশন (সাদা/কালো) [A B/W television that is working]<br>j. |_____|টেলিভিশন (রঙিন) [A color television that is working]</td></tr>
</table>

k. _____|রেফ্রিজারেটর [Refrigerator]
l. |_____|সাইকেল (খেলনা সাইকেল নয়) A bicycle (used for commercial purposes not toy for children)
m. |_____|মটর সাইকেল [A motorcycle]
n. |_____|সেলাই মেশিন [A sewing machine]
o. |_____||_____|মোবাইল ফোন (সংখ্যা) [Number of Mobile phones]
p. |_____|ল্যান্ড ফোন [A land phone]

4106. আপনার খানায় রান্নার জন্য প্রধানত কি ধরনের জ্বালানী ব্যবহার করা হয় ? [What type of fuel does your household mainly use for cooking?]

কাঠ [Wood] ........ 01
শস্যের অবশিষ্টাংশ/ঘাস [Crop residue / grass ] ........ 02
শুকনা গোবর [Dung cakes ] ........ 03
কয়লা [Coal / coke / lignite ] ........ 04
কাঠ কয়লা [Charcoal ] ........ 05
কেরোসিন [Kerosene ] ........ 06
বিদ্যুৎ [Electricity ] ........ 07
তরল গ্যাস/প্রাকৃতিক গ্যাস [Liquid gas / gas ] ........ 08
বায়ো-গ্যাস [Bio-gas] ........ 09
অন্যান্য [Other] ........ 77
(বর্ণণা লিখুন) [*Specify other]* ______________________
জানি না [Don't know ] ........ 99

4107. |_____|আপনার খানার মোট মাসিক আয় কত? (সকল আয়ের উৎসের যোগফল যেমনঃ বেতন, ঘর ভাড়া, কৃষিখাত ইত্যাদি) For the purpose of our research, would you please tell us your total monthly household income? (Please sum up your income from all sources like wage, rent, agriculture etc.)

1. 4000 টাকা পর্যন্ত [UP TO 4000 Taka]
2. 4001-6000 টাকা [4001-6000 Taka]
3. 6001- 8000 টাকা [6001- 8000 Taka]
4. 8001- 10000টাকা [8001- 10000 Taka]
5. 10001- 12000 টাকা [10001- 12000 Taka]
6. 12001- 15000 টাকা [12001- 15000 Taka]
7. 15001-20000 টাকা [15001-20000 Taka]
8. 20001- 25000 টাকা [20001- 25000 Taka]
9. 25001- 30000 টাকা [25001- 30000 Taka]
10. 30000 টাকার উপরে [Above 30000 Taka]

4108. |____| বসতবাড়ি মালিকানা [What is the ownership status of the house where your household is currently living?]

নিজের বাড়ি [Self-owned] ........ 1..... **4110** নং প্রশ্নে চলে যান (skips to **4110**)
ভাড়া বাড়ি [Rental] ........ 2
সরকারী জমি [Govt. land] ........ 3..... **4110** নং প্রশ্নে চলে যান (skips to **4110**)
কোন জমিদারের/জোতদারের বাড়িতে থাকে [Owned by a landlord] ........ 4..... **4110** নং প্রশ্নে চলে যান (skips to **4110**)
বাড়ি ভাড়া না দিয়ে অন্যের বাড়িতে থাকে [Living in someone's house without giving rent] .......5........ **4110** নং প্রশ্নে চলে যান (skips to **4110**)

asania 2/15/12 10:00 PM
**Comment:** Fix skip notes

4109. যদি **704** - এর উত্তর ***2*** হয় (ভাড়া বাড়ি), তবে বসতবাড়িটির আয়তন কি **200** বর্গফুটের কম? [If 704 answer is 2

| | | | | |
|---|---|---|---|---|
| | | (rental) then is the area less than 200 sq. ft.?] | | |
| | 4110. | \|____\|\|____\|\|____\|\|____\|\|____\| উত্তর হ্যাঁ হলে, খানার সদস্যদের বসতবাড়িতে মোট কতটুকু জমি আছে (ডেসিমেল)? (How much homestead land (decimal) does your household own?]<br><br>বলতে রাজি হয়নি [Refused]…...............................8 (skip to 709)<br>জানি না [Don't know]…..................…...............99 | | |
| | 4111. | আপনার মতে সামাজিক প্রেক্ষাপটে আপনার খানাটির অবস্থান কি রকম [How would you describe your economic status?]<br>ধনী [Dhoni].................................................1<br>উচ্চ মধ্যবিত্ত [Uchho modho bitto] ................2<br>মধ্যবিত্ত [Modho Bitto]..................................3<br>দরিদ্র [Doridro or Nimno motho bitto] ..........4<br>হতদরিদ্র [Hotodoridro] ................................5 | | |
| | 4112. | I'm going to read you a list of animals. Please tell me how many of each stays <u>in your compound</u>, and how many belong to <u>your household</u>. IF NONE, ENTER "00", IF UNKNOWN, ENTER "99" | COMPOUND | HOUSEHOLD |
| | 1 | CATTLE | \|__\|__\| | \|__\|__\| |
| | 2 | GOATS | \|__\|__\| | \|__\|__\| |
| | 3 | CHICKENS | \|__\|__\| | \|__\|__\| |
| | 4113. | How often do the cattleroam free in the compound?<br>READ RESPONSES.<br>[1] ALWAYS ROAM FREE<br>[2] SOMETIMES ROAM FREE<br>[3] NEVER ROAM FREE<br>4. N/A; ONLY HAVE CHICKENS | | |
| | 4114. | Do chickens or other poultry ever go inside <u>your main house</u>?<br>READ RESPONSES.<br>[1] ALWAYS<br>[2] SOMETIMES<br>[3] NEVER<br>4. N/A; NO CHICKENS | | |

**Wash Benefit Module 42**

**Feces Observed in and Around the Compound**

Administer to: All study households

👁 Observation: For the following:

*Record the number of piles of human feces you observe in eacharea (up to 10 piles)*

55 Too numerous to count (more than 10 piles)

99 Cannot tell / could not observe

👁Within the courtyard

| | | | |
|---|---|---|---|
| **C.826.** | 4201. | **Human** feces <u>within the **courtyard**</u> that could be considered open defecation | ☐☐ |

| | | | |
|---|---|---|---|
| C.827. | 4202. | **Animal** feces present within the **courtyard**<br>(mark all that apply) | ☐1 Poultry (chicken, duck, pigeon)<br>☐2 Cow / Buffalo<br>☐3 Goat / Sheep<br>☐4 Pig<br>☐5 Dog or Cat<br>☐8 Other<br>☐9 NO FECES OBSERVED<br>☐99 COULD NOT OBSERVE |
| In the area where the target child spends the most time<br>*Probe and identify where the target child spends the most time. At baseline, the target child will not be present, so the search should be defined as the area where the pregnant mother spends the most time.* | | | |
| C.828. | 4203. | **Human** fecesin the **area where the target child spends the most time** that could be considered open defecation | ☐☐ |
| C.829. | 4204. | **Animal** feces in the **area where the target child spends the most time**<br>(mark all that apply) | ☐1 Poultry (chicken, duck, pigeon)<br>☐2 Cow / Buffalo<br>☐3 Goat / Sheep<br>☐4 Pig<br>☐5 Dog or Cat<br>☐8 Other<br>☐9 NO FECES OBSERVED<br>☐99 COULD NOT OBSERVE |
| Behind the study house | | | |
| C.830. | 4205. | **Human** fecesbehindthe **house** that could be considered open defecation | ☐☐ |
| C.831. | 4206. | **Animal** feces **behind the house**<br>(mark all that apply) | ☐1 Poultry (chicken, duck, pigeon)<br>☐2 Cow / Buffalo<br>☐3 Goat / Sheep<br>☐4 Pig<br>☐5 Dog or Cat<br>☐8 Other<br>☐9 NO FECES OBSERVED<br>☐99 COULD NOT OBSERVE |

#### Wash Benefit Module 5

##### VACCINATION HISTORY

asania 2/15/12 9:21 PM
**Comment:** Skip for baseline

PUT AT END. EASY TO COLLECT AND COPY FROM VACCINATION CARD. PUT BEFORE HH OBSERVATION.

Removed body because not needed now.

Administer to: Target children
Respondent: Child's primary caregiver.
Thank you for your time. I would now like to ask you for your child's (index) vaccination card. If she has one verbally confirm each vaccination as you enter it.

| | | | |
|---|---|---|---|
| | 501. | CHILD ID: | \|__\|__\|__\|__\|__\|__\|__\| |
| | 502. | CHILD NAME | ______________________ |

#### Wash Benefit Module 5

##### VACCINATION HISTORY

Administer to: Target children
Respondent: Child's primary caregiver.
Thank you for your time. I would now like to ask you for your child's (index) vaccination card. If she has one verbally confirm each vaccination as you enter it.

| | | | |
|---|---|---|---|
| | 501. | CHILD ID: | \|__\|__\|__\|__\|__\|__\| |
| | 502. | CHILD NAME | ______________________ |
| C.501 | 503. | Do you have a card where [NAME'S] vaccinations are written down? [*If YES*: May I see it please?] (আপনার শিশুর (নাম বলুন) কোন টিকা কার্ড আছে কী? হ্যাঁ হলে বলুন: আমি কী টিকা কার্ডটি দেখতে পারি?) | 1 Yes, seen (হ্যাঁ, দেখাতে পেরেছে) **(Skip to V122)** (**V122** নং প্রশ্নে চলে যান)<br>2 Yes, not seen (হ্যাঁ, দেখাতে পারে নাই)<br>3 No card (কোন টিকা কার্ড নাই) |

asania 2/15/12 9:25 PM
**Comment:** Pda will open this from the tracking section

mi_sheuli 2/15/12 9:25 PM
**Comment:** CHILD id NOT TO BE FILLED TWICE. WILL DISCUSS WITH MAHMUD BHAI.

#### Wash Benefit Module 3

##### DEWORMING

Administer to:

Pregnant mothers **[PDA to open MOTHERS COHORT]**

Children < 36 months at enrollment **[PDA to open DIARRHEA COHORT]**

~~18-27mo children that provide stool specimens~~**[~~PDA to open~~ PARASITE ~~COHORT~~]**

Respondent: Child's primary caregiver or pregnant mother.
Now I would like to ask you about any medications you might have taken for intestinal worms.

| | | | |
|---|---|---|---|
| C.301 | 301. | In the last <u>six months</u>, has [NAME] received a pill or drug for intestinal worms? | [1] YES<br>[2] NO → **Skip to 305**<br>[99] DK/NOT SURE → **Skip to 305** |
| C.302 | 302. | Where did [NAME] receive the drug for intestinal worms? | [1] AT HOME/IN VILLAGE<br>[2] AT A CLINIC OR HEALTH FACILITY<br>[3] AT A SCHOOL<br>[99] DK/NOT SURE |
| C.303 | 303. | Did [NAME] receive the drug as part of a larger campaign? | [1] YES<br>[2] NO<br>[99] DK/NOT SURE |
| C.304 | 304. | Approximately how long ago did [NAME] receive the drug?<br><br>MARK "99" FOR MONTHS IF DK/NOT SURE | \|__\|MONTHS (0-6)<br>\|__\|__\|WEEKS |
| C.305 | 305. | Has the [NAME] eaten any dirt or soil?<br>*Ask for each recall period:* | |
| | 1 | 1. Today | ☐ 1 Yes ☐ 2 No ☐ 99 Don't know |
| | 2 | 2. Yesterday | ☐ 1 Yes ☐ 2 No ☐ 99 Don't know |

| | | | | | | |
|---|---|---|---|---|---|---|
| **Wash Benefit Module 2**<br><br>**DIARRHEA AND ILLNESS SYMPTOMS (Children < 36 months)** | | | | | | |
| Administer to: Children < 36 months living in a study compound at baseline<br>This may include the child in the household or in other households in the bari.<br>PDA TO OPEN CHILD ID FROM **DIARRHEA COHORT**<br>IF study compound DOES NOT HAVE A CHILD 0-36 MONTHS → **SKIP TO C.305** | | | | | | |
| Respondent: Child's primary caregiver | | | | | | |
| | WBM 2 | Thank you. Now I am going to ask you some questions about the health of your [child/children]. Please answer each question as accurately as you can. If you don't know the answer to a question, say "I don't know." We will begin with [NAME OF CHILD 1]. | | | | |
| | | | **A** | **B** | **C** | **D** |
| | | **Did [NAME] have [SYMPTOM] :** | **Today**<br>**1=Yes**<br>**0=No**<br>**999= Don't know** | **Yesterday**<br>**1=Yes**<br>**0=No**<br>**999= Don't know** | **Day before Yesterday**<br>**1=Yes**<br>**0=No**<br>**999= Don't know** | **In the last 7 days** (since this day last week)<br>**1=Yes**<br>**0=No**<br>**999= Don't know** |
| **C.201** | 201. | Fever | | | | |
| **C.202** | 202. | Diarrhea | | | | |
| **C.203** | 203. | 3 or more bowel movements in 24 hours | | | | |
| **C.204** | 204. | Number of bowl movements each day | | | | |
| **C.205** | 205. | Watery or soft stool (unformed) | | | | |
| **C.206** | 206. | Blood in the stool | | | | |
| **C.207** | 207. | Skin rash (anywhere on the body) | | | | |
| **C.208** | 208. | Constant cough | | | | |
| **C.209** | 209. | Congestion / runny nose | | | | |
| **C.210** | 210. | Panting / wheezing / difficulty breathing | | | | |
| **C.211** | 211. | Bruising, scrapes or cuts | | | | |
| **C.212** | 212. | Toothache / teething | | | | |
| **C.213** | 213. | *If answered Yes to C.202* | | | | |

#### Wash Benefit Module 4

#### Anthropometry

#### Wash Benefit Module 16

#### Child Development

#### Wash Benefit Module 14

#### LNS measurement

mi_sheuli 2/15/12 1:56 PM
**Comment:** Skip for baseline

PROTOCOL
Biomedical
Non-Exempt
Berkeley

Protocol # 2011-09-3652
Date Printed: 07/31/2018

---------------------------------------------------------------------------------
---

| | |
|---|---|
| **Protocol Title:** | Measuring the benefits of sanitation, water quality, handwashing and nutrition interventions for improving health and development in rural Bangladesh |
| **Protocol Status:** | APPROVED |
| **Date Submitted:** | 10/26/2017 |
| **Approval Period:** | **10/30/2017-11/03/2018** |
| **Important Note:** | **This Print View may not reflect all comments and contingencies for approval. Please check the comments section of the online protocol. Questions that appear to not have been answered may not have been required for this submission. Please see the system application for more details.** |

*** * * Attached Document * * ***

| Document Name | Created Date |
|---|---|
| Figure 1-Study Design.pdf | 10/26/2017 |

**Figure 1. WASH Benefits study design overview**

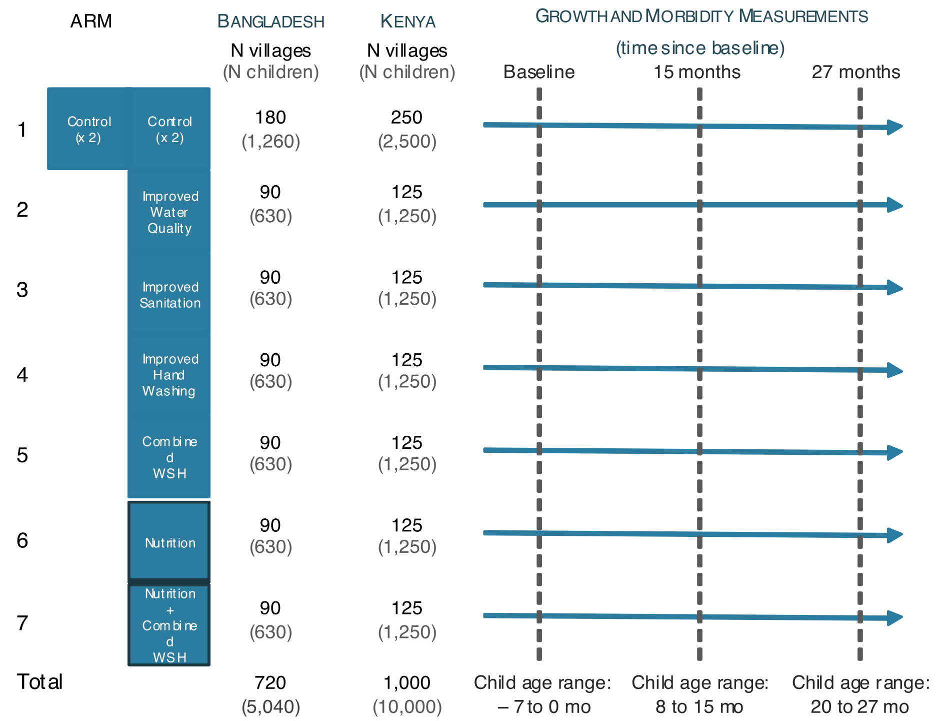

PROTOCOL
Biomedical
Non-Exempt
Berkeley

Protocol # 2011-09-3652
Date Printed: 07/31/2018

-----------------------------------------------------------------------------------
---

**Protocol Title:** Measuring the benefits of sanitation, water quality, handwashing and nutrition interventions for improving health and development in rural Bangladesh

**Protocol Status:** APPROVED

**Date Submitted:** 10/26/2017

**Approval Period:** **10/30/2017-11/03/2018**

**Important Note:** **This Print View may not reflect all comments and contingencies for approval. Please check the comments section of the online protocol.**
**Questions that appear to not have been answered may not have been required for this submission. Please see the system application for more details.**

*** * * Attached Document * * ***

| Document Name | Created Date |
|---|---|
| Nutrition-Supplement-Analysis.pdf | 10/26/2017 |

APPENDIX 4

NUTRIENT CONTENT COMPARISON OF THE PROPOSED LNS TO MMP AND NUTRIBUTTER

| Nutrient | Unit | WHO/FAO RNIs for children 1-3 y* | MMP† | | Nutributter‡ | | Proposed LNS‡ | | |
|---|---|---|---|---|---|---|---|---|---|
| | | | Content | % RNI | Content | % RNI | Content | % RNI | Chemical form |
| Dose | g | | | | 20 | | 20 | | |
| Energy | kcal | | | | 108 | | 118 | | |
| Fat | g | | | | 7 | | 9.6 | | |
| Linoleic acid | g | | | | 1.29 | | 4.46 | | |
| Alpha-linolenic acid | g | | | | 0.29 | | 0.58 | | |
| Ratio of LA to ALA | | | | | 4.4 | | 7.7 | | |
| Protein | g | | | | 2.6 | | 2.6 | | |
| ***Vitamins*** | | | | | | | | | |
| Vitamin A | µg | 400 | 400 | 100% | 400 | 100% | 400 | 100% | Retyinyl acetate |
| Vitamin D | µg | 5 | 5 | 100% | NA | | 5 | 100% | Cholecalciferol (D3) |
| Vitamin E | mg | 5 | 5 | 100% | NA | | 6 | 120% | DL-alpha-tocopherol acetate |
| Vitamin K | µg | 15 | NA | | NA | | 30 | 200% | Phylloquinone 5% |
| Vitamin C | mg | 30 | 30 | 100% | 30 | 100% | 30 | 100% | L-ascorbic acid |
| Biotin | µg | 8 | NA | | NA | | NA | | |
| Folic acid | µg | 150 | 150 | 100% | 80 | 53% | 150 | 100% | Pteroyl monoglutamic acid |
| Thiamine (B1) | mg | 0.5 | 0.5 | 100% | 0.3 | 60% | 0.5 | 100% | Thiamin hydrochloride |
| Riboflavin (B2) | mg | 0.5 | 0.5 | 100% | 0.4 | 80% | 0.5 | 100% | Riboflavin |
| Niacin | mg | 6 | 6 | 100% | 4 | 67% | 6 | 100% | Niacinamide |
| Pantothenic acid (B5) | mg | 2 | NA | | 1.8 | 90% | 2 | 100% | Calcium pantothenate |
| Vitamin B6 | mg | 0.5 | 0.5 | 100% | 0.3 | 60% | 0.5 | 100% | Pyridoxine hydrochloride |
| Vitamin B12 | µg | 0.9 | 0.9 | 100% | 0.5 | 56% | 0.9 | 100% | Cyanocobalamin (0.1%) |
| ***Minerals*** | | | | | | | | | |
| Calcium§ | mg | 500 | NA | | 100 | 20% | 280 | 56% | Tri-calcium phosphate |

| Nutrient | Unit | WHO/FAO RNIs for children 1-3 y* | MMP† | | Nutributter‡ | | Proposed LNS‡ | | |
|---|---|---|---|---|---|---|---|---|---|
| | | | Content | % RNI | Content | % RNI | Content | % RNI | Chemical form |
| Copper¶ | mg | 0.34 | 0.56 | 165% | 0.2 | 59% | 0.34 | 100% | Encapsulated copper sulfate |
| Iodine | µg | 90 | 90 | 100% | 90 | 100% | 90 | 100% | Potassium iodate |
| Iron** | mg | 5.8 | 10.0 | 172% | 9 | 155% | 6 | 103% | Encapsulated ferrous sulfate |
| Magnesium§ | mg | 60 | NA | | 16 | 27% | 40 | 67% | Magnesium citrate |
| Manganese | mg | 1.2 | NA | | 0.08 | 7% | 1.2 | 100% | Manganeze sulfate |
| Phosphorous§ | mg | 460 | NA | | 82 | 18% | 190 | 41% | Tri-calcium phosphate & Di-potassium phosphate |
| Potassium | mg | | NA | | 152 | | 200 | | Di-potassium phosphate & potassium chloride |
| Selenium | µg | 17 | 17 | 100% | 10 | 59% | 20 | 118% | Sodium selenite 1.5% |
| Zinc** | mg | 4.1 | 4.1 | 100% | 4 | 98% | 8 | 195% | Zinc sulfate |

*RNI=Recommended Nutrient Intake; MMP=Multiple Micronutrient Powder; LNS=Lipid-based nutrient supplement
† Formulation for UNICEF/WHO/WFP Multiple micronutrient powder
‡ In malaria endemic areas, it is recommended that the supplement be split into two 10 g servings in one day to reduce the iron consumed in a single bolus dose
§ The calcium, phosphorus, and magnesium content of Nutributter and the proposed LNS do not meet the RNI for technical reasons
¶ The IOM RDA level for copper for infants 1-3 y is shown here. MNPs use the FAO/IAEA/WHO 1996 recommended intake value for copper
** The RNI for iron and zinc is that assumed under a diet of moderate bioavailability.

----------------------------------------------------------------------------------
---

| | |
|---|---|
| **Protocol Title:** | Measuring the benefits of sanitation, water quality, handwashing and nutrition interventions for improving health and development in rural Bangladesh |
| **Protocol Status:** | APPROVED |
| **Date Submitted:** | 10/26/2017 |
| **Approval Period:** | **10/30/2017-11/03/2018** |
| **Important Note:** | **This Print View may not reflect all comments and contingencies for approval. Please check the comments section of the online protocol. Questions that appear to not have been answered may not have been required for this submission. Please see the system application for more details.** |

*** * * Attached Document * * ***

| Document Name | Created Date |
|---|---|
| WASH Benefits Behavior Change Strategy Overview Feb 21 2012.pdf | 10/26/2017 |

icddr,b
KNOWLEDGE FOR GLOBAL LIFESAVING SOLUTIONS

**A randomized controlled trial of the benefits of water sanitation hygiene plus nutrition interventions on child growth (WASHplus BENEFITS main trial)**

#### Behavior Change Strategy and Communication Plan:

- **For 6 different interventions to promote Home Water Treatment, Sanitation, Handwashing and Nutrition behaviors, separately or in combination, in rural Bangladesh**

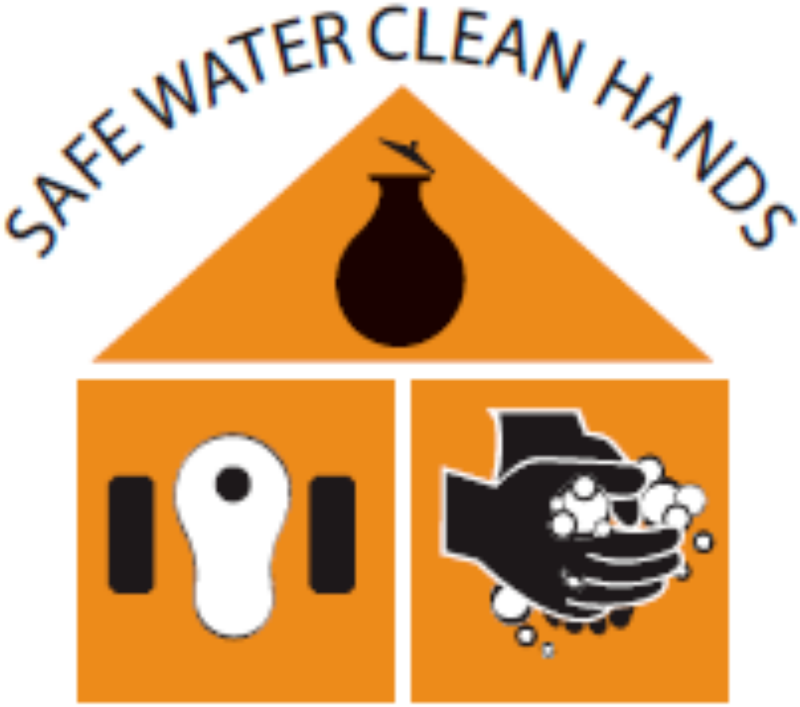

### SAFE ENVIRONMENT STRONG KIDS

Funded by:

- Bill and Melinda Gates Foundation

---

Version of: February 21, 2012

#### Overview of WASHplus Benefits Behavior Change Strategy

| **“Safe Water Clean Hands Safe Environment Strong Kids!”** | | | |
|---|---|---|---|
| **Target groups** | **Behavioral objectives** | **Key factors of behavior change to be influenced by the intervention** | **Intervention Activities** |
| All household members and caregivers of children born into the trial to 2 years old<br><br>Handwashi ng station household volunteers<br><br>Community health promoters (CHPs)<br><br>Pregnant women | To **handwash with soap** after defecation and during food preparation<br><br>To replenish handwashing supplies and remind family to handwash<br><br>To counsel, model and advocate for handwashing in the households of trial children and in the extended community<br><br>Establish hand washing routines before giving birth | Stable access to two low cost, enabling technologies:<br>-handwashing stations<br>-soapy water dispensers where/when needed<br>Fixed location in context of ultiple water sources<br><br>Skills on how to handwash and prepare soapy water<br><br>Convenience<br>Ease of practice and use<br>Self-efficacy<br><br>Family and community support in maintaining the technologies and replenishing the supplies<br><br>Health protection from diarrheal illness<br><br>Invoking feelings of disgust<br><br>Counseling to remind, point out the benefits, negotiate & problem-solve, as needed<br><br>Responding to aspirations of nurture, happiness, safety, modernity, scientific soundness & esthetics<br><br>Positive social norms<br><br>Cues and habit formation<br><br>Addressing humoral explanations of cold | **Community Health Promoters (CHPs)** conduct periodic:<br>-home counseling visits<br>-extended family meetings, and<br>-follow ups with handwashing station volunteers<br><br>To:<br>-Provide the technologies<br>-Teach the skills<br>- Arrange for social support<br>- Communicate:<br>-the benefits of use and practice<br>-the change in social norms<br>Aided by flipcharts (initially), cue cards, & other channels and media<br>-Congratulate and encourage<br>Aided by tokens of appreciation (stickers, soap, certificates for caretakers, etc)<br>-Problem-solve, as needed<br>-Inspire<br><br>**Handwashing station family volunteers** conduct daily monitoring of handwashing supplies, encourage and problem-solve as needed<br><br>ICDDRB trains and supports CHPs on:<br>-technical competence in handwashing<br>-Counseling and communication skills<br>-Supplying enabling technologies, communication aids, and tokens of appreciation<br>-Supplying enabling technologies to CHPs themselves |

| "Safe Water Clean Hands Safe Environment Strong Kids!" | | | |
|---|---|---|---|
| **Target groups** | **Behavioral objectives** | **Key factors of behavior change to be influenced by the intervention** | **Intervention Activities** |
| Children born into the trial to 2 years old, and all of their household members<br><br>Drinking water guarding household volunteers<br><br>Community health promoters (CHPs)<br><br>Pregnant women | To **drink chlorinated water**, treated at the point of use up to 24 hours earlier<br><br>To replenish and re-treat the household drinking water every 24 hours or less<br>To remind family to give baby only chlorinated water<br><br>To counsel, model and advocate for water chlorination in the households of trial children<br><br>Establish water chlorination routines | Stable access to two low cost, enabling technologies:<br>-chlorination tablets<br>--water storage containers for treated drinking water<br><br>Self-efficacy through<br>-Skills on how to treat and handle the water correctly<br><br>-Key knowledge of<br>-the lasting effect of chlorine compared to other water treatments<br>-the inconsequential effects of chlorine smell to children and pregnant women<br>-regular users noticing smell less with time<br>-Distinction from medicine<br><br>Interpretation of smell as a sign of water safety rather than a sign of alarm<br><br>Convenience<br>Ease of practice and use<br>No maintenance<br><br>Family support (males + elders)<br><br>Health protection from diarrheal illness<br><br>Invoking feelings of disgust<br><br>Counseling to point out the benefits, negotiate & problem-solve, as needed<br><br>Responding to aspirations of nurture, happiness, safety, modernity, and scientific soundness | **Community Health Promoters (CHPs)** conduct periodic:<br>-home counseling visits<br>-follow ups with drinking water guarding family volunteers, and<br>-extended family meetings, if needed<br><br>To:<br>-Provide the technologies<br>-Teach the skills<br>-Arrange for social support<br>-Communicate:<br>-the key knowledge on chlorine<br>-the benefits of use and practice<br>-the change in social norms<br>Aided by flipcharts (initially), cue cards, & other channels and media<br>-Congratulate and encourage<br>Aided by tokens of appreciation (stickers, certificates for caretakers, etc)<br>-Problem-solve, as needed<br>-Inspire<br><br>**Drinking water guarding family volunteers** conduct daily monitoring of drinking water, encourage and problem-solve as needed<br><br>ICDDRB trains and supports CHPs on:<br>-technical competence in water chlorination<br>-Counseling and communication skills<br>-Supplying enabling technologies, communication aids, and tokens of appreciation<br>-Supplying enabling technologies to CHPs themselves |

| "Safe Water Clean Hands Safe Environment Strong Kids!" | | | |
|---|---|---|---|
| **Target groups** | **Behavioral objectives** | **Key factors of behavior change to be influenced by the intervention** | **Intervention Activities** |
| | before giving birth | Social norm creation for POU<br><br>Cues and habit formation | |
| Extended family of children born into the trial to 2 years old (bari level)<br><br>Caregivers of children born into the trial<br><br>Floor, yard, and latrine keeper household volunteers<br><br>Community Health Promoters (CHPs) | To **use dual pit improved latrines**<br><br>To **potty train** their young children, beginning at x months of age; to empty the potty in latrine or safe pit<br><br>To collect child and animal feces from the floor and yard **using sani-scoop** at least twice a day<br><br>To counsel, model and advocate for sanitation in the households of trial children and in the extended community | Availability of three low cost, enabling technologies to all eligible:<br>-dual pit improved latrines<br>-child potties<br>-sani-scoops<br><br>Self-efficacy through Knowledge and skills on how to potty train, use saniscoop, build an improved dual pit latrine, or convert a current one, and switch and empty pit when necessary<br><br>Self-efficacy of children<br><br>Comfort and convenience<br>Ease of practice and use<br><br>Family and community support in maintaining the technologies<br><br>Shared responsibility<br><br>Cleanliness and safety of the home and the environment<br>Favorable environment (clean latrine)<br><br>(for latrine)<br>Purdah and privacy, esp. for women<br><br>Invoking feelings of shame and disgust<br><br>Perceived health threats<br><br>Counseling to remind, point | **Community Health Promoters (CHPs)** conduct periodic:<br>-home counseling visits<br>-extended family meetings, and<br>-follow ups with floor/yard/latrine keeper volunteers<br>Coordinate with community partner for latrine building (VERC)<br><br>To:<br>-Provide the technologies<br>-Teach the skills<br>- Arrange for social support<br>- Communicate:<br>-the benefits of use and practice<br>-the change in social norms<br>Aided by flipcharts (initially), cue cards, & other channels and media<br>-Congratulate and encourage<br>Aided by tokens of appreciation (stickers, certificates for caretakers, etc)<br>-Problem-solve, as needed<br>-Inspire<br><br>**Floor, yard and latrine keeper volunteers** conduct daily monitoring, encourage and problem-solve as needed<br><br>ICDDRB in collaboration with VERC trains and supports CHPs on:<br>-technical competence in sanitation<br>-Counseling and communication skills<br>-Supplying enabling technologies, communication aids, and tokens of appreciation |

| "Safe Water Clean Hands Safe Environment Strong Kids!" | | | |
|---|---|---|---|
| **Target groups** | **Behavioral objectives** | **Key factors of behavior change to be influenced by the intervention** | **Intervention Activities** |
| Pregnant women | Prepare for sanitation routines before giving birth | out the benefits, negotiate & problem-solve, as needed<br><br>Aspirations of nurture, happiness, safety, modernity, esthetics, & social status<br><br>Positive social norms<br><br>Cues and habit formation | |
| Mothers of children born into the trial<br><br>Caregivers of children born into the trial<br><br>Community Health Promoters | -To initiate breastfeeding within ½ hour and continue **exclusive breastfeeding** until 180 days<br><br>Continue to breast feed Practice active, **complementary feeding** during baby's 6-24 months of life<br><br>Supplement baby's food with **LNS** twice a day<br><br>**Handwash with soap** before baby food preparation and feeding<br><br>To counsel, model and advocate for | Exclusive breastfeeding skills and tips (self-efficacy)<br><br>Support for early and exclusive breastfeeding from birth attendant and family<br><br>Active feeding skills<br><br>Key knowledge that: - complementary food is whatever you have at home (no need for special food for your baby)<br>-Use of LNS / continued feeding/breastfeeding during illness is necessary<br><br>(for LNS)<br>Convenience<br>Ease of use<br><br>Familial support for LNS in absence of mother (caregivers)<br><br>Counseling to remind, point out the benefits, negotiate & | **Community Health Promoters (CHPs)** conduct:<br>-home counseling visits of pregnant mothers<br>-breastfeeding initiation visit at birth<br>-periodic home counseling visits<br><br>To:<br>-Teach the skills<br>-Arrange for social support<br>- Provide LNS<br>-Communicate:<br>-the key knowledge on breastfeeding, complementary feeding and LNS<br>-the benefits of use and practice<br>-the change in social norms<br>Aided by flipcharts (initially), cue cards, & other channels and media<br>-Congratulate and encourage<br>Aided by tokens of appreciation (stickers, certificates, etc)<br>-Problem-solve, as needed<br>-Inspire<br><br>ICDDRB in collaboration with A&T trains and supports CHPs on:<br>-technical competence in breastfeeding and complementary feeding<br>-Counseling and communication skills<br>-Supplying LNS, communication aids, and |

| “Safe Water Clean Hands Safe Environment Strong Kids!” | | | |
|---|---|---|---|
| **Target groups** | **Behavioral objectives** | **Key factors of behavior change to be influenced by the intervention** | **Intervention Activities** |
| (CHPs)<br><br><br><br><br><br><br>Pregnant women | healthy nutrition in the households of trial children<br><br>Prepare for exclusive breastfeeding before giving birth | problem-solve, as needed<br><br>Responding to aspirations of nurture, happiness, and safety<br><br>Positive social norms<br><br>Cues and habit formation | tokens of appreciation<br>-Supplying LNS to CHPs themselves (with eligible children in the age window) |

PROTOCOL
Biomedical
Non-Exempt
Berkeley

---------------------------------------------------------------------------------

| | |
|---|---|
| **Protocol Title:** | Measuring the benefits of sanitation, water quality, handwashing and nutrition interventions for improving health and development in rural Bangladesh |
| **Protocol Status:** | APPROVED |
| **Date Submitted:** | 10/26/2017 |
| **Approval Period:** | **10/30/2017-11/03/2018** |
| **Important Note:** | **This Print View may not reflect all comments and contingencies for approval. Please check the comments section of the online protocol. Questions that appear to not have been answered may not have been required for this submission. Please see the system application for more details.** |

*** * * Attached Document * * ***

| Document Name | Created Date |
|---|---|
| Prot PR-11063_ERC approval_9 April 2012.pdf | 10/26/2017 |

icddr,b

### Memorandum

9 April 2012

To: Dr Stephen P Luby
Principal Investigator of research protocol # PR-11063
Centre for Communicable Diseases (CCD)

From: Professor AKM Nurul Anwar
Chairman
Ethical Review Committee (ERC)

Sub: Approval of research protocol # PR-11063

Thank you for your memos dated 29 February 2012 and 27 March 2012 attaching the modified version of your research protocol # PR-11063 entitled **"A randomized controlled trial of the benefits of water sanitation hygiene plus nutrition interventions on child growth"** addressing the issues raised by the full committee in its December meeting held on 28 December 2011 to the satisfaction of the Committee. Accordingly, the Committee approved the research protocol. You will be required to observe the following terms and conditions in implementing the research protocol:

1. The research protocol is approved for 12-month period from the date of approval of the protocol. Annual approval for further continuation, if needed, shall be obtained before expiration of the approval.
2. The ERC approval shall automatically be revoked after one year if the protocol is not started. After one year, you shall have to seek approval for revalidation of the protocol by the RRC & ERC before starting.
3. You should notify the IRB Secretariat of the start date of the protocol for updating in the integrated navision system. The protocol start date will not be updated in the navision system until receiving information from you. Therefore you will not be able to operate budget code and continue spending funds under the research protocol.
4. As Principal Investigator, the ultimate responsibility for scientific and ethical conduct including the protection of the rights and welfare of study participants vest upon you. You shall also be responsible for ensuring competence, integrity and ethical conduct of other investigators and staff directly involved in this research protocol.
5. You shall conduct the study in accordance with the ERC-approved protocol and shall fully comply with any subsequent determinations by the ERC.
6. You shall obtain prior approval from the Research Review Committee and the ERC for any modification in the approved research protocol and/or approved

GPO Box 128, Dhaka 1000, Bangladesh.
**For Packages :** 68, Shaheed Tajuddin Ahmed Sarani, Mohakhali, Dhaka 1212, Bangladesh.

consent form(s), except in case of emergency to safeguard/eliminate apparent immediate hazards to study participants. Such changes must immediately be reported to the ERC Chairman.

7. You shall recruit/enrol participants for this study strictly adhering to the criteria mentioned in the research protocol.

8. You shall obtain legally effective informed consent (i.e. consent should be free from coercion or undue influence) from the selected study participants or their legally responsible representative, as approved in the protocol, using the approved consent form prior to their enrollment in this study. Before obtaining consent, all prospective study participants must be adequately informed about the purpose(s) of the study, its methods and procedures, and also what would be done if they agree and also if they do not agree to participate in the study.

   They must be informed that their participation in the study is voluntary and that they can withdraw their participation any time without any prejudice. Signed consent forms should be preserved for a period of at least five years following official termination of the study.

9. You shall promptly report the occurrence of any Adverse Event or Serious Adverse Event or unanticipated problems of potential risk to study participants or others to the ERC in writing within 24 hours of such occurrences.

10. Any significant new findings, developing during the course of this study that might affect the risks and benefits and thus influence either participation in the study or continuation of participation should be reported in writing to the participants and the ERC.

11. Data and/or samples should be collected and interviews should be conducted, as specified in the ERC-approved protocol, and confidentiality must be maintained. Data/samples must be protected by reasonable security, safeguarding against risks such as their loss or unauthorized access, destructions, used by others, and modification or disclosure of data. Data/samples should not be disclosed, made available to or use for purposes other than those specified in the protocol, and shall be preserved for a period, as specified under Centre's policies/practices.

12. You shall promptly and fully comply with the decision of the ERC to suspend or withdraw its approval for the research protocol.

13. You shall report progress of research to the ERC for continuing review of the implementation of the research protocol as stipulated in the ERC Guidelines. Relevant excerpt of ERC Guidelines and '*Annual/Completion Report for Research Protocol involving Human Subjects*' are attached for your information and guidance.

14. The RRC should be immediately notified if the protocol is discontinued before the expected date of completion.

I wish you success in running the above-mentioned study.

Copy: Coordination Manager, RAS

PROTOCOL
Biomedical
Non-Exempt
Berkeley

---------------------------------------------------------------------------------
---

**Protocol Title:** Measuring the benefits of sanitation, water quality, handwashing and nutrition interventions for improving health and development in rural Bangladesh

**Protocol Status:** APPROVED

**Date Submitted:** 10/26/2017

**Approval Period:** 10/30/2017-11/03/2018

**Important Note:** This Print View may not reflect all comments and contingencies for approval. Please check the comments section of the online protocol.
Questions that appear to not have been answered may not have been required for this submission. Please see the system application for more details.

*** * * Attached Document * * ***

| Document Name | Created Date |
|---|---|
| NOITR-597-0.pdf | 10/26/2017 |

#### IRB Reliance #597 : Study Information

| | |
|---|---|
| Reliance #: | 597 |

Title of Study

WASH Benefits Main Study Bangladesh

| | |
|---|---|
| Principal Investigator: | Jack Colford |
| Primary Awardee Institution: | UC Berkeley |
| Degree: | MD, PhD |
| Title: | Professor |
| Department: | - |
| E-mail address: | |
| Phone: | - |
| Name of Reviewing Sponsor: | Bill and Melinda Gates Foundation |
| UC Location providing IRB Review: | UC Berkeley |

Reviewing Award Information

No records found.

| | |
|---|---|
| Type of Study: | Pending IRB |
| Review Type: | Pending IRB |

Provide a brief lay (max 500 words) synopsis or abstract of the entire study.

The goal of the WASH Benefits study is to generate rigorous evidence about the impacts of sanitation, water quality and handwashing interventions on child health and development in the first years of life.The core scientific objective of WASH Benefits is to evaluate the spectrum of benefits obtained through improved sanitation, water quality, hygiene promotion, and nutrition interventions. More specifically, there are three scientific goals that the study addresses:•    Rigorously measure health benefits arising from low-cost approaches including local promoters and simple technologies (e.g. latrine improvements or potties for children, chlorine based tablets, and handwashing stations).•    Evaluate the degree to which, in resource-constrained settings, there is added health benefit to delivering multiple interventions concurrently (sanitation facilities, drinking water, and handwashing promotion).•    Measure the impact of lipid-based nutrient supplementation alone and in combination with sanitation, water and hygiene interventions on child growth and development. />The current study is designed as two, highly comparable randomized trials in Bangladesh and Kenya. The study has two main phases. The first phase was a 2-year pilot that will continue through 2011. The pilot has allowed the study teams to refine the sanitation, handwashing and water quality interventions and identify hardware and behavior change packages that result in high levels of uptake.

Reviewing Research Coordinators

Research Coordinator #1
First Name: Jack
Last Name: Colford
Degree: MD, PhD
Title: Professor
Type: review

© 2013 UC Regents

**IRB Reliance #597 : Reliance Information**

#### Assigned Relying Principal Invesigator 1

| | |
|---|---|
| Name: | Christine Stewart |
| Degree: | PhD |
| Title: | Assistant Professor |
| E-mail Address: | |
| Campus: | UC Davis |

Will the Relying PI/LI recruit and consent subjects? no

a. Type (e.g., adults, children) and number of subjects

N/A

b. How, when, and by whom potential subjects will be approached

N/A

c. The informed consent process

N/A

Relying PIs/LIs scope of work and activities.

The relying PIs at UC Davis will be involved in the design of the study, creation of data collection forms, analysis of data, and interpretation/publication of results. The focus of the work of the UC Davis team relates to the development, implementation, and evaluation of the nutrition components of the trial.

Award Information

No records found.

Relying Research Coordinators

No records found.

Relying Key Study Personnel

Key Study Personnel #1
First Name: Kathryn
Last Name: Dewey
Title: Professor
Degree: PhD


#### Assurances

+ I certify that the information provided in this application is complete and correct.

+ I certify that I will follow the IRB-approved Protocol.

+ I will comply all applicable federal and state laws regarding the protection of human subjects in research.

+ I will make sure that the personnel performing this study are qualified and adhere to the provisions of this IRB-approved protocol.

+ I will not modify the protocol or any attached materials without first obtaining review and approval from the Reviewing IRB.

+ I will accept responsibility for the conduct of this study at this site, the ethical performance of the project, and the protection of the rights and welfare of the human subjects who are directly involved at this site.

**[X] PI has asserted the above assurances to the Reviewing IRB.**

© 2013 UC Regents

PROTOCOL
Biomedical
Non-Exempt
Berkeley

Protocol # 2011-09-3652
Date Printed: 07/31/2018

-----------------------------------------------------------------------------------------

**Protocol Title:** Measuring the benefits of sanitation, water quality, handwashing and nutrition interventions for improving health and development in rural Bangladesh

**Protocol Status:** APPROVED

**Date Submitted:** 10/26/2017

**Approval Period:** 10/30/2017-11/03/2018

**Important Note:** This Print View may not reflect all comments and contingencies for approval. Please check the comments section of the online protocol.
Questions that appear to not have been answered may not have been required for this submission. Please see the system application for more details.

#### * * * Attached Document * * *

| Document Name | Created Date |
| --- | --- |
| Wash Benefits Bangladesh Baseline Questionnaire_2013_03.pdf | 10/26/2017 |

ICDDR,B

### Wash Benefits

Baseline Survey

asania

2/16/2012

This document includes the baseline survey and associated instructions. It specifies the population that needs to be included for the survey, indicators from the common modules in addition to country specific indicators. It guides the user through the survey process from the start to the end. Please note that the module numbers match the common module numbers but appear out of order in this document. The items include the common module variable number in the first column to faciliate bridging the survey question numbers to the common module question numbers. Modules 40 and 41 are additional modules or subsets of the common modules split to aid survey administration, only in the Bangladesh baseline. The sequence of the modules as it appears now is open for discussion and can be easily revised. There are placeholders for the modules that are not applicable for baseline.

**NOTES TO THE FIELD TEAM**

**চার ভাগে উত্তর আসবে** There is 4 groups of respondents of interest to us:

| POPULATION | DEFINITION | SAMPLES |
|---|---|---|
| 1. নিদ্দিষ্ট শিশু Target children | গবেষণায় আওতাধীন শিশু Children in utero at enrollment (primary study population) | বেইজ লাইনে প্রজোয্য নয় N/A FOR BASELINE |
| 2. মা/প্রধান পরিচর্যাকারী Mothers | নিদ্দিষ্ট শিশু র মা/প্রধান পরিচর্যাকারী Mothers of target children | রক্ত,মল,প্রস্রাব BLOOD, STOOL, URINE |
| 3. ডায়রিয়া কোহট/অংশ Diarrhea cohort | ৩৬ মাস বয়সের নীচের শিশু যে এই খানা/বাড়ী/কম্পাউন্ডের মধ্যে বাস করে Children < 36 months old at enrollment, living in study compounds | নমূনা লাগবে না NO SAMPLES |
| 4. পরজীবি কোহট/অংশ Parasite cohort | <u>NOT FINALIZED</u> বেইজ লাইনে ১৮-২৭ মাস বয়সে শিশুর রক্ত,মলের নমূনা সংগ্রহ করতে হবে যে এই খানা/বাড়ী/কম্পাউন্ডের মধ্যে বাস করে Children 18 – 27 months old at baseline (blood spot, stool) ~~School-aged children 6 – 12 years old at baseline (stool)?~~ | রক্ত,মল BLOOD, STOOL |

তালিকা/লিস্টিং দল খানার নিদ্দিষ্ট সদস্য এবং তাদের ঠিকানাসহ তালিকা রিসার্চ অফিসারকে প্রদান করবে। পরবর্তীতে এই তালিকা বেইজ লাইন দলকে খানার নির্ধারিত সদস্যাকে পেতে সহায়তা করবে । বেইজ লাইন দলকে এই তালিকায় খানার নির্ধারিত সদস্যাকে চিহ্নিত করেও দিতে হবে। THE LISTING TEAM WILL GIVE THE FIELD RESEARCH OFFICERS A LIST OF HOUSEHOLDS, ASSOCIATED ADDRESSES AND RESPONDENT SPECIFICS TO HELP THE BASELINE TEAM LOCATE THE ELIGIBLE HOUSEHOLDS. THE HOUSEHOLDS WILL BE VISIBLY MARKED FOR IDENTIFICATION.

খানার সম্মতি**<u>Consent (All households)</u>**

নিলিখিত বিষয়গুলো জরুরী FOLLOW THESE STEPS CAREFULLY:

1. নিদ্দিষ্ট সদস্য/খানার বর্তমান ঠিকানার সঠিক বর্তমান ঠিকানার তথ্য CONFIRM THE INFORMATION FROM THE LISTING WITH THE CURRENT ADDRESS/HOUSEHOLD.
2. নির্ধারিত সদস্যার গর্ভধারনের সঠিক তথ্য যাচাই VERIFY RESPONDENT ELIGIBILITY BY ASKING FOR PREGNANCY STATUS.
3. নির্ধারিত গর্ভবতী ছাড়াও এই বাড়ীতে অন্য গর্ভবতী আছে কিনা নিদ্দিষ্ট তা করুন DETERMINE IF THERE ARE ANY OTHER ELIGIBLE RESPONDENTS IN THE COMPOUND.
4. যদি উত্তরদাতা আমাদের সমস্ড় শর্তাদি পূরণ করতে পারে তাহলে খানা আইডি খানার সম্মতি পত্রের নিদ্দিষ্ট অংশে রেকর্ড করুন IF RESPONDENT MEETS OUR CRITERIA, CAREFULLY RECORD **HOUSEHOLD ID** AT THE TOP OF THE CONSENT PAGE.
5. তালিকায় দেয়া আইডি এবং নির্ধারিত কাজের জন্য বাড়ী/কম্পাউন্ড /খানার আইডি মিলিয়ে দেখুন এবং নিদ্দিষ্ট করুন CONFIRM THAT THE ID <u>EXACTLY MATCHES THE ID ASSIGNED TO THIS COMPOUND/HOUSEHOLD BY THE FIELD RESEARCH OFFICER</u>.
6. আইডি লেখার পর তালিকায় দেয়া আইডির সাথে পুনরায় মিলিয়ে দেখুন AFTER WRITING IT CROSS IT OFF YOUR LIST.

লিখিত সম্মতিপত্র পড়ার সময়ে সম্মতিপত্রটি মৌখিকভাবে উত্তরদাতাকে /বোঝাতে হবে এবং তার সম্মতিতে খানার আইডি বসাতে হবে READ THE CONSENT FORM VERBATIM TO THE RESPONDENT IN THE LANGUAGE OF THE WRITTEN CONSENT FORM AND THEN CONFIRM THAT SHE UNDERSTANDS.

খানার আইডি নং ENTER THE HOUSEHOLD ID: |__|__|__|__|__||__|__|__|__|__||__|__|__|.

সম্মতি পত্র পড়ুন **Read**: ওহে। আমার নাম ----{নাম বলুন}|আমি ঢাকা মহাখলী কলেরা হাসপাতাল(আইসিডিডিআরবি) থেকে এসেছি। আজ আমরা আপনার খানায় গবেষণা বিষয়ে কথা বলতে এসেছি যেহেতু আপনার খানাটি এই ষ্টাডীর জন্য নির্বাচিত হয়েছে ( যেহেতু আপনি গর্ভবতী/আপনার ১৮-২৭ মাসের একটি শিশু আছে|Hello. My name is [name]. I am from Mohakhali Cholera Hospital (icddr,b) in Dhaka. We are here to talk to you today because your household has been selected for our study because [you are pregnant/you have a child 18-27 months old]. READ REST OF CONSENT.

জিপিএস অবস্থান **GPS position:** খানার সম্মতি অনুযায়ী খানার দরজার কাছে দাড়ান এবং জিপিএস-এ রেকর্ড করুন Following the household's consent, stand in a position closest to the household's front door and record the GPS coordinates.

##### Wash Benefit Module 40 Geographical Identification

নিচে উত্তরদাতার র্নিদিষ্টকরন তথ্য গুরুত্বের সাথে পূরন করুন FILL THE FOLLOWING IDENTIFICATION INFORMATION FOR THE RESPONDENT VERY CAREFULLY.

| Module ID | WBM 40 | Q ID | |
|---|---|---|---|
| | 4001. | 00 | এফ.আর.এ কোড (FRA code): ?? |
| C.101 | 4002. | 01 | তথ্য সংগ্রহের তারিখ [Date of data collection]: ??/??/???? |
| C.18 | 4003. | | জিপি এস রিডিং পয়েন্ট আই ডি নাম্বার (GPS point ID number) [Auto-populate, if possible] |
| C.19 | 4004. | | জিপি এস ল্যাটিচিউড রিডিং (GPS Latitude [Auto-populate, if possible]) |
| C.20 | 4005. | | জিপি এস লংগিচিউিড রডিং রডিং (GPS Longitude[Auto-populate, if possible]) |
| | 4006. | 04 | জেলার নাম(DISTRICT NAME): |
| | 4007. | 05 | উপজেলার নাম (UPAZILA NAME): |
| | 4008. | 06 | ইউনিয়নের নাম [Union name _____________ |
| | 4009. | 07 | খানার ঠিকানা Household Address: |
| | 4010. | 08 | খানা প্রধানের নাম [Name of Household head]_____________________________ |
| | 4011. | 09 | খানা প্রধানের বাবা/স্বামীর নাম [Father's/Husband's Name of HH head] ________________ |
| | 4012. | 10 | বাড়ির নাম [Bari Name]: _________________________________________ |
| | 4013. | 11 | গ্রামের নাম [Village Name]: ______________________________________ |
| C.01 | 4014. | 12 | ক্লাস্টার নাম্বার [Cluster No. [1-720] |
| C.02 | 4015. | 13 | বাড়ী নাম্বার [Bari No. [1-7] |
| C.03 | 4016. | 14 | **HH ID:** [1-7] |
| | 4017. | 15 | Unique HH ID: |
| C.04 | 4018. | 16 | মায়ের আইডি [Mother ID [1-7] **[PDA to store in MOTHER COHORT]** |
| C.05 | 4019. | 17 | টার্গেট শিশুর আইডি [১-৭][Target child ID [1-7] **[PDA to store in Target Child Cohort]** |
| | 4019.a | | **\|_____\| এই বাড়ীতে ৩৬ মাসের নীচে কতজন শিশু আছে?** How many children do you have < 36 months in this bari? |

| | | | | | | | |
|---|---|---|---|---|---|---|---|
| | শিশুর প্রদত্ত তালিকা দেখুন এবং শিশুর উপস্থিতি উল্লেখ করুন (Now look at the list provided to ensure the presence of the additional enrolled children in the bari) | | | | | | |
| | | শিশুর পরিচয় (Who is this child?) | নাম (Name) | লিঙ্গ (Sex) | জন্ম তারিখ(Birth date) | জন্ম তারিখের উৎস(Source of DOB )(C.104) | নির্বাচন করার কারণ (একাধিক উত্তর হতে পারে) Reason for enrollment (select all that apply) **[PDA to store accordingly]** |
| **C.06.** | 4020. | 1. র্নিদিষ্ট শিশু (Target child)<br>2 .একই মায়ের অন্য শিশু(Another child of same mother)<br>3. একই বাড়ীর | | 1 পুরুষ M<br>0 মহিলা F | দিন/মাস/বছর DD/MM/YY | 1 = কার্ড দ্বারা নিশ্চিত হয়েছিল (Confirmed DOB by valid vaccination/health card) | ☐1 ডায়রিয়া Diarrhea<br>☐2 পায়খানার নমুণা Stool specimen<br>☐3 রক্তের নমুণা Blood specimen |

| | | | | | | | |
|---|---|---|---|---|---|---|---|
| | | অন্য খানার শিশু (Child from the same bari)<br>4. একই খানার অন্য মায়ের শিশু(Another child of another mother in the same HH) | | | | 2 = মা/আত্নীয় দ্বারা নিশ্চিত হয়েছিল (Mother/Relative remembers DOB)<br>3 = উভয় (Both 1 & 2)<br>4 = ঘটনা দ্বারা নিশ্চিত হয়েছিল (Estimated DOB with 2 and event calendar) | |
| **C.07.** | 4021. | 1. র্নিদিষ্ট শিশু (Target child)<br>2 .একই মায়ের অন্য শিশু(Another child of same mother)<br>3. একই বাড়ীর অন্য খানার শিশু (Child from the same bari)<br>4. একই খানার অন্য মায়ের শিশু(Another child of another mother in the same HH) | | 1 পুরুষ M<br>0 মহিলা F | দিন/মাস/বছর DD/MM/YYY | 1 = কার্ড দ্বারা নিশ্চিত হয়েছিল (Confirmed DOB by valid vaccination/health card)<br>2 = মা/আত্নীয় দ্বারা নিশ্চিত হয়েছিল (Mother/Relative remembers DOB)<br>3 = উভয় (Both 1 & 2)<br>4 = ঘটনা দ্বারা নিশ্চিত হয়েছিল (Estimated DOB with 2 and event calendar) | ☐1 ডায়রিয়া Diarrhea<br>☐2 পায়খানার নমুণা Stool specimen<br>☐3 রক্তের নমুণা Blood specimen |
| **C.08.** | 4022. | 1. র্নিদিষ্ট শিশু (Target child)<br>2 .একই মায়ের অন্য শিশু(Another child of same mother)<br>3. একই বাড়ীর অন্য খানার শিশু (Child from the same bari)<br>4. একই খানার অন্য মায়ের শিশু(Another child of another | | 1 পুরুষ M<br>0 মহিলা F | দিন/মাস/বছর DD/MM/YYY | 1 = কার্ড দ্বারা নিশ্চিত হয়েছিল (Confirmed DOB by valid vaccination/health card)<br>2 = মা/আত্নীয় দ্বারা নিশ্চিত হয়েছিল (Mother/Relative remembers DOB)<br>3 = উভয় | ☐1 ডায়রিয়া Diarrhea<br>☐2 পায়খানার নমুণা Stool specimen<br>☐3 রক্তের নমুণা Blood specimen |

| | | | | | | | |
|---|---|---|---|---|---|---|---|
| | | mother in the same HH) | | | | (Both 1 & 2)<br>4 = ঘটনা দ্বারা নিশ্চিত হয়েছিল (Estimated DOB with 2 and event calendar) | |
| C.09. | 4023. | 1. র্নিদিষ্ট শিশু (Target child)<br>2 .একই মায়ের অন্য শিশু(Another child of same mother)<br>3. একই বাড়ীর অন্য খানার শিশু (Child from the same bari)<br>4. একই খানার অন্য মায়ের শিশু(Another child of another mother in the same HH) | | 1 পুরুষ M<br>0 মহিলা F | দিন/মাস/বছর DD/MM/YYY | 1 = কার্ড দ্বারা নিশ্চিত হয়েছিল (Confirmed DOB by valid vaccination/health card)<br>2 = মা/আত্বীয় দ্বারা নিশ্চিত হয়েছিল (Mother/Relative remembers DOB)<br>3 = উভয় (Both 1 & 2)<br>4 = ঘটনা দ্বারা নিশ্চিত হয়েছিল (Estimated DOB with 2 and event calendar) | ☐1 ডায়রিয়া Diarrhea<br>☐2 পায়খানার নমুণা Stool specimen<br>☐3 রক্তের নমুণা Blood specimen |
| C.10. | 4024. | 1. র্নিদিষ্ট শিশু (Target child)<br>2 .একই মায়ের অন্য শিশু(Another child of same mother)<br>3. একই বাড়ীর অন্য খানার শিশু (Child from the same bari)<br>4. একই খানার অন্য মায়ের শিশু(Another child of another mother in the same HH) | | 1 পুরুষ M<br>0 মহিলা F | দিন/মাস/বছর DD/MM/YYY | 1 = কার্ড দ্বারা নিশ্চিত হয়েছিল (Confirmed DOB by valid vaccination/health card)<br>2 = মা/আত্বীয় দ্বারা নিশ্চিত হয়েছিল (Mother/Relative remembers DOB)<br>3 = উভয় (Both 1 & 2)<br>4 = ঘটনা দ্বারা নিশ্চিত হয়েছিল (Estimated DOB with 2 and event calendar) | ☐1 ডায়রিয়া Diarrhea<br>☐2 পায়খানার নমুণা Stool specimen<br>☐3 রক্তের নমুণা Blood specimen |

**Wash Benefit Module 1 RESPONDENT IDENTIFICATION**

| | | |
|---|---|---|
| মডিউল আইডি<br>Module ID | WBM 1 | |
| | 101. | : What is your full name? প্রধান উত্তরদাতার নাম [Name of respondent: |
| | 102. | : **I_____I** প্রধান উত্তরদাতার পরিচয় [Status of main respondent]<br>1. প্রধান পরির্চযাকারী /মা [Mother of youngest child]<br>2. পরির্চযাকারী (পুরুষ) [Male caregiver]<br>3. পরির্চযাকারী (মহিলা) [Female caregiver] |
| | 103. | প্রধান উত্তরদাতার বয়স (বছরে) [Age of main respondent: *(in years)]* |
| | 104. | উত্তরদাতা কোন শর্তটা পূরণ করেছে? RESPONDENT MEETS WHICH ELIGIBILITY CRITERIA?<br>[1] গর্ভবতী PREGNANT<br>[2] ১৮-২৭ মাসের বাচ্চা আছে HAS CHILD 18-27 MONTHS OF AGE |
| আমি আপনাকে আপনার গর্ভবতী হওয়া বিষয়ে কিছু ব্যক্তিগত প্রশ্ন এই গবেষনার জন্য করতে চাই ।আমি আশা করি এ বিষয়ে আপনি কিছু মনে করবেন না এবং আমাদিগকে সহায়তা করবেন । I need to ask you some personal questions regarding your pregnancy because that is our project's eligibility criteria. I hope you do not mind and will cooperate with us. [see scripts of other studies] | | |
| C. 108 | 105. | উত্তরদাতার সর্বশেষ মাসিকের তারিখ (**What was the date of your last menstrual period?)**▯▯/▯▯/▯▯▯▯<br>88 / 88 / 88 = সর্বশেষ গর্ভধারণের পর থেকে মাসিক না হলে (No menstruation since their last pregnancy)<br>99 = জানি না (Don't know / not sure) |
| C. 109 | 106. | \|__\| গর্ভধারণের পূর্নাঙ্গ মাস গণনা করুন( Record number of completed months)<br>99 = জানি না (Don't know / not sure) |
| C.110 | 107. | \|___\|গর্ভধারণের মসয়কালের উৎস(Source of Pregnancy Length)<br>1 = শুধুমাত্র মায়ের নির্ণয় (Estimated by mother only)<br>2 = মায়ের এবং স্বাস্থ্য সেবাদানকারীর নির্ণয় ,আলট্রাসাউন্ড ছাড়া(Estimated by mother and a health practitioner, no ultrasound (last prenatal visit)<br>3 = মা, স্বাস্থ্য সেবাদানকারী এবং আলট্রাসাউন্ড দ্বারা নির্ণয় Estimated by mother and health practitioner, using ultrasound (last prenatal visit) |
| | 108. | ▯▯/▯▯/▯▯ কখন আপনার শিশু জন্মাবে? (What will be the expected DOB of your child?)<br>__/__/__ |
| | 109. | নির্দিষ্ট শিশুর প্রকৃত বয়স (বেইজ লাইনে লাগবে না)Actual DOB of target child (skip for baseline)<br>__/__/___ |
| | 110. | Do you have any paln to go to your mother's house for delivary porpoise?<br>1. Yesহ্যাঁ<br>0. No না (Skip to module 0) |
| | 111. | আপনার মায়ের বাড়ী যে গ্রামে সেই গ্রামের নাম কি, যেখানে আপনি বাচ্চার জন্মগ্রহনের সময় যাবেন? What is the name of your mother's (maternal) village, where you might go to give birth?<br>0. Same as this একই গ্রাম<br>1. গ্রামের নাম লিখুন _______________ |
| আগামী কয়েক মাস আমাদের এই প্রকল্পের কিছু বিষয়ে আপনাকে কিছু তথ্য দেওয়ার জন্য ফোনে যোগাযোগ রাখতে চাই । আপনার কি কোন ফোন আছে? যদি না থাকে তাহলে আপনার বাড়ীতে কারো বা প্রতিবেশীর ফোন আছে কি? তার নাম্বার কত? At some point over the next several months, we may need to get in touch with you by phone to give you information about our project. Do you have a phone? If not, does someone in your compound have a phone? Maybe a neighbor? What is the number? | | |

**Wash Benefit Module 0 TRACKING**

| | | |
|---|---|---|
| | 001 | উত্তরদাতার/ উত্তরদাতার খানার কোন সদস্যের কী কোন ফোন নাম্বার আছে?(Is there a phone available?) 1 Yes<br>0 No (Skip to 005) |
| C.11 | **002** | -----------প্রাথমিক ফোন নাম্বার (Primary phone number) |
| C.12 | 003 | এই ফোনে কি আপন াকে সরাসরি পাওয়া যাবে? (Does this phone number belong to you?) 1 Yes(Skip to 005) 0 No |
| C.13 | 004 | যদি আপন াকে পাওয়া না যায় তাহলে এই নাম্বারটি কার? IF NO, to whom does this number belong? (Specify |

| | | |
|---|---|---|
| | | relationship to respondent)<br>1 . স্বামী (HUSBAND)<br>2. শ্বশুর IN LAWS<br>3. পিতা(FATHER)<br>4.মা(MOTHER)<br>5. পরিবারের সদস্য(FAMILY MEMBER)<br>99. অন্যান্য (নাম ও উত্তরদাতার সাথে সম্পর্ক লিখুন) (OTHER) |
| | 004.a | দ্বিতীয় কোন ফোন নাম্বার আছে কী?(Is there a phone available?)<br>1 Yes 0 No (Skip to 008) |
| C.14 | 005 | -----------দ্বিতীয় ফোন নাম্বার Secondary phone number |
| C.15 | 006 | এই ফোনে কি আপন াকে সরাসরি পাওয়া যাবে? Does this phone number belong to you? 1 Yes(Skip to 008) 0 No |
| C.16 | 007 | যদি আপন াকে পাওয়া না যায় তাহলে এই নাম্বারটি কার? IF NO, to whom does this number belong?<br>1. স্বামী (HUSBAND)<br>2. শ্বশুর IN LAWS<br>3. পিতা(FATHER)<br>4 .মা(MOTHER)<br>5 .পরিবারের সদস্য(FAMILY MEMBER)<br>99 .অন্যান্য (নাম ও উত্তরদাতার সাথে সম্পর্ক লিখুন) (OTHER) |
| C.17 | 008 | এই খানাটির আশেপাশে কোন উল্লেখযোগ্য কিছু আছে কি? (Location/landmark) |
| | 009 | \|_____\| \|_____\| সবচেয়ে ছোট বাচ্চার মা কত ক্লাস পর্যন্ত পড়ালেখা শেষ করেছেন ? (জানি না = ***99***) Education of mother of the youngest child *(Years of education completed, DK=99)* |
| | 010 | \|_____\| \|_____\| সবচেয়ে ছোট বাচ্চার বাবা কত ক্লাস পর্যন্ত পড়ালেখা শেষ করেছেন ? (জানি না = ***99***) Education of father of the youngest child *(Years of education completed, DK=99)* |
| | 011 | \|_____\| \|_____\| সবচেয়ে ছোট বাচ্চার বাবার প্রধান পেশা [Main occupation of father of the youngest child]<br><br>পেশা কোড Occupation Code:<br><br>1. কৃষক [Farmer/Cultivator]<br>2. ঘরে কাজ করে [Homemaker]<br>3. কৃষি শ্রমিক [Agri-labor]<br>4. শ্রমিক (কৃষি শ্রমিক ছাড়া) [Non-agri labor]<br>5. বেতনভুক্ত কর্মচারী (সরকারী/প্রাইভেট/এন.জি.ও) [Salaried job (Govt./Private/NGO)]<br>6. রাজমিস্ত্রী [Mason (*Rajmistri*)]<br>7. কাঠ মিস্ত্রী [Carpenter]<br>8. ভ্যান/রিকশা চালক [Van/Rickshaw puller]<br>9. জেলে [Fisherman]<br>10. নৌকা চালক/মাঝি [Boatman]<br>11. কর্মকার [Blacksmith]<br>12. স্বর্ণকার [Goldsmith]<br>13. কুমার/কুম্ভকার [Potter (soil smith)]<br>14. মুচি [Shoe polish /maker]<br>15. দোকানদার [Shopkeeper]<br>16. ফেরিওয়ালা [Vendor (*Feriwala/howker*)]<br>17. ক্ষুদ্র ব্যবসায়ী (মূলধন <=১০০০০) [Petty trader, capital <=10000]<br>21. কুটীর শিল্প [Cottage industry]<br>22. পোলট্রি/ ব্যবসার জন্য পশু লালন-পালনকারী [Poultry /livestock rearer]<br>23. বৈদ্যুতিক মিস্ত্রী [Electrician]<br>24. হোমিওপ্যাথি ডাক্তার [Homeopath]<br>25. আধ্যাতিক চিকিৎসক/ কবিরাজ/ ওঝাঁ [Spiritual healer/kabiraj/ Ojha]<br>26. পেশাদার ডাক্তার/উকিল [Professional practitioner (Doctor/lawyer)]<br>27. ইমাম/ ধর্মযাজক [Imam/priest]<br>28. অবসরপ্রাপ্ত চাকুরীজীবী [Retired service holder]<br>29. ছাত্র [Student]<br>30. বেকার [Unemployed]<br>31. অক্ষম [Disabled]<br>32. কাজের লোক [Domestic maid / servant]<br>33. জমিদার (শস্য উৎপাদন অথবা অন্য কোন কাজে কৃষকদের জমি বর্গা দেয়) [Landlord (Provide land for farmers for sharecropping |

| | | | |
|---|---|---|---|
| | | 18. ব্যবসায়ী (মূলধন >১০০০০) [Business, capital >10000]<br>19. দর্জি [Tailor]<br>20. ড্রাইভার [Driver] | or others)]<br>34. বিদেশে থাকে [Staying abroad]<br>35. মৃত/নিখোঁজ [Died/untraced]<br>99. জানিনা [Don't know]<br><br>77. অন্যান্য (বর্ণনা লিখুন) [Others (specify)] |
| | 012 | \|_____\| \|_____\| আপনার খানাতে/পরিবারে কত জন লোক প্রতিদিন একই হাড়িতে রান্না করে খান?How many people in this house eat from the same cooking pot every day (খানা বলতে একই হাড়িতে রান্না করে খায় এমন সদস্যদের বুঝানো হয়েছে) | |
| | 013 | \|_____\| আপনার খানাতে ৩৬ মাসের নীচে কতজন শিশু আছে? How many children do you have < 36 months do you have? | |
| | 014 | \|____\| এই বাড়ীতে ৩৬ মাসের নীচে কতজন শিশু আছে? How many children do you have < 36 months in this bari? | |
| | 014 | \|_____\| এই বাড়ীতে কতগুলো খানা আছে? How many HHs do you have in this bari? | |

A. এই বাড়ীর খানাগুলোর সেনসাস (সকল খানার) CENSUS OF HOUSEHOLDS ARE IN THIS COMPOUND (All households)

| | | | | | | |
|---|---|---|---|---|---|---|
| পড়ে শোনান READ | ঠিক আছে। আমি এখন আপনাকে সকল খানার সম্পর্কে আলাদাভাবে কিছু প্রশ্ন করব। প্রথমে আপনার খানা সম্পর্কে বলুন। Okay. Now I am going to ask you about each household separately. First, think about your own household.<br>MARK "99" IF RESPONDENT DOESN'T KNOW THE ANSWER TO ANY OF THE FOLLOWING QUESTIONS. | | | | | |
| | A.1 | A.2 | A.3 | A.4 | A.5 | A.6 |
| খানার নাম্বার HH No. | খানা প্রধানের নাম কি? What is the name of the household head? FOR REFERENCE ONLY | এই খানাতে ৪০ বছরের উপরের বয়সের কতজন সদস্য আছে? How many adults over 40 years live in this household? | এই খানাতে (>১৮-৪০) বছরের বয়সের কতজন সদস্য আছে? How many young adults (>18-40 years) live in this household? | এই খানাতে (>৩-১৮) বছরের বয়সের কতজন সদস্য আছে?How many school-aged children (>3-18 years) live in this household? | এই খানাতে (০-৩) বছরের বয়সের কতজন সদস্য আছে?How many young children (0-3 years) live in this household? | প্রতিটি খানায় কতজন সদস্য আছে তা গননা করুন এবং নিশ্চিত হোন CALCULATE THE TOTAL NUMBER OF PEOPLE IN EACH HH AND CONFIRM |
| 1 | | | | | | |
| 2 | | | | | | |

**Wash Benefit Module 18 MATERNAL FFQ**

৭-দিনের খাদ্য গ্রহণের তালিকা বিষয়ে মায়ের জন্য প্রশ্নপত্র **7-Day Food Frequency Questionnaire for Mothers**

আমি এখন আপনাকে আপনার পুষ্টিসংক্রান্ত বিষয়ে কিছু প্রশ্ন করতে চাই। গত ৭দিনের মধ্যে কতদিন আপনি নিম্নলিখিত খাবারগুলো খেয়েছেন তা বলুন। Now I would like to ask you some questions regarding your nutritional status. In the past 7 days, that is since last [Tomorrow's Day], on how many days have you eaten the following foods:

| **CM** | **WBM 18** | **Foods**(খাদ্য) | দিনের সংখ্যা (০-৭)**Number of days (0-7)** |
|---|---|---|---|
| **C.1801.** | 1801. | যে কোন মাংস( গরু/ভেড়া/ছাগল/মহিষ/শুকর)<br>Any kind of meat (cow/buffalo/sheep/goat/pig) | [__] |
| **C.1802.** | 1802. | যে কোন মাংস(মুরগী/হাস/পাখি)<br>Any kind of poultry meat | [__] |
| **C.1803.** | 1803. | যে কোন মাছ<br>Any kind of fresh fish | [__] |

| | | | |
|---|---|---|---|
| **C.1804.** | 1804. | যে কোন শুকনা মাছ<br>Any kind of dried fish | [__] |
| **C.1805.** | 1805. | চিংড়ি, কাকড়া<br>Shrimp/ parwn, crabs, | [__] |
| **C.1806.** | 1806. | ডিম Eggs | [__] |
| **C.1807.** | 1807. | দুধ Milk | [__] |
| **C.1808.** | 1808. | পনির/দই/মাঠা<br>Yogurt/Curd/Buttermilk | [__] |
| **C.1809.** | 1809. | ডাল এবং ডাল জাতীয় Dal and other legumes | [__] |
| **C.1810.** | 1810. | চীনা বাদাম বা অন্যান্য যে কোন বাদাম<br>Peanuts and other nuts | [__] |
| **C.1811.** | 1811. | কমলা রংয়ের আলু যার ভিতরের রং কমলা<br>Orange flesh sweet potato | [__] |
| **C.1812.** | 1812. | গাজর, মিষ্টি কুমড়া, লাল রংয়ের মিষ্টি মরিচ, কুমড়া যার ভিতরের রং হলুদ বা কমলা? Carrot, pumpkins, red papper, orange or yellow squash | [__] |
| **C.1813.** | 1813. | গাঢ়-সবুজ রং-এর শাক জাতীয় Dark green leafy vegetables (cabbage will not be included) | [__] |
| **C.1814.** | 1814. | লাল রং-এর শাক জাতীয়( যেমন: লাল শাক)<br>Red leafy vegetables | [__] |
| **C.1815.** | 1815. | আম Mango | [__] |
| **C.1816.** | 1816. | পেয়ারা Guava | [__] |
| **C.1817.** | 1817. | আনারস Pineapple | [__] |
| **C.1818.** | 1818. | বড়ই Indian Jujube | [__] |
| **C.1819.** | 1819. | কলা Banana | [__] |
| **C.1820.** | 1820. | ফুলকপি Cauliflower | [__] |
| **C.1821.** | 1821. | টমেটো Tomato | [__] |
| | 1822. | সারা বছর ধরে আপনি/ আপনার খানা সদস্যরা যে পরিমান চাল খান তার কতটুকু আপনারা নিজের জমিতে চাষ করেন? How much of the rice that you eat throughout the year comes from land that your family cultivates? | 1) একটুও না None<br>2) সামান্য, অর্ধেকেরও কম Some, but less than half 3) প্রায় অর্ধেক About half 4) অর্ধেকেরও বেশী More than half<br>5) সবটুকু বা প্রায় সবটুকু All or nearly all |
| | 1823. | কে কে বা কারা এই জমি চাষ করেন? (একাধিক উত্তর হতে পারে) | 1. [__]নিজের জমি নিজে চাষ করেন<br>২. [__]নিজের জমি অন্যে চাষ করেন/ বর্গা দিয়েছেন<br>৩. [__]অন্যের জমি নিজে চাষ করেন/বর্গা |

| | | | |
|---|---|---|---|
| | | | নিয়েছেন/বন্ধক নিয়েছেন<br>৪. [__]নিজের জমি আগে চাষ করতাম এখন বন্ধক দিয়েছি |

**Wash Benefit Module 11 FOOD INSECURITY**

খানার খাদ্য অনিশ্চিয়তা এবং অধিকার সম্পর্কিত **Household Food Insecurity and Access Scale**

বেইজ লাইনে সকল খানার জন্য প্রযোজ্য Administer to: All study households during baseline

| | WBM 11 | ঘটনা **Occurrence Questions**<br>1 = কদাচিৎ (বিগত ৪ সপ্তাহে একবার বা দুইবার ঘটেছে) Rarely (once or twice in the past four weeks)<br>2 = মাঝে মাঝে (বিগত ৪ সপ্তাহে তিনবার -দশবার ঘটেছে) Sometimes (three to ten times in the past four weeks)<br>3 = প্রায়ই (বিগত ৪ সপ্তাহে দশবারের বেশী ঘটেছে) Often (more than ten times in the past four weeks) | উত্তর Response | উত্তর যদি হ্যাঁ হয় তাহলে কতবার এমন ঘটেছে ?If yes, how often did this happen? |
|---|---|---|---|---|
| **C.1101.** | 1101. | 1. . বিগত ৪ সপ্তাহে আপনাদের খানায় অভাবের কারণে যথেষ্ট খাবার ছিল না, এরকম অবস্থায় আপনার কোন দুশ্চিন্তা হয়েছিল কি? In the past four weeks, did you worry that your household would not have enough food? | [0] না (পরবর্তী 1102 নং প্রশ্নে যান) No [skip to 1102]<br>[1] হ্যাঁ Yes | [1] কদাচিৎ Rarely<br>[2] মাঝে মাঝে Sometimes<br>[3] প্রায়ই Often |
| **C.1102.** | 1102. | 2. আপনারা সাধারণত যে ধরণের খাবার খেয়ে থাকেন, গত ৪ সপ্তাহে অভাবের কারণে আপনি বা আপনাদের খানার কোনো সদস্য কি সে ধরণের খাবার খেতে পারেননি? In the past four weeks, were you or any household member not able to eat the kinds of foods you preferred because of a lack of resources? | [0] না (পরবর্তী 1103 নং প্রশ্নে যান) No [skip to 1103]<br>[1] হ্যাঁ | [1] কদাচিৎ Rarely<br>[2] মাঝে মাঝে Sometimes<br>[3] প্রায়ই Often |
| **C.1103.** | 1103. | 3. গত ৪ সপ্তাহে আপনি বা আপনাদের খানার কোনো সদস্যকে কি অভাবের কারণে সীমিত রকমের (ভিন্নতা) খাবার খেতে হয়েছে, অর্থাৎ আগে যত পদ খেতেন গত ৪ সপ্তাহে অভাবের কারণে তার চেয়ে কম পদ খেতে হয়েছে? In the past four weeks, did you or any household member have to eat a limited variety of foods due to a lack of resources? | [0] না (পরবর্তী 1104 নং প্রশ্নে যান) No [skip to 1104]<br>[1] হ্যাঁ | [1] কদাচিৎ Rarely<br>[2] মাঝে মাঝে Sometimes<br>[3] প্রায়ই Often |
| **C.1104.** | 1104. | 4. গত ৪ সপ্তাহে আপনাকে বা আপনার খানার কোনো সদস্যকে, যে খাবার সাধারনত আপনারা খান না, অভাবের কারনে তা খেয়ে থাকতে হয়েছে কি? In the past four weeks, did you or any household member have to eat some foods that you really did not want to eat because of a lack of resources to obtain other types of food? | [0] না (পরবর্তী 1105 নং প্রশ্নে যান) No [skip to 1105]<br>[1] হ্যাঁ | [1] কদাচিৎ Rarely<br>[2] মাঝে মাঝে Sometimes<br>[3] প্রায়ই Often |
| **C.1105.** | 1105. | 5. গত ৪ সপ্তাহে আপনাকে বা আপনার খানার কোনো সদস্যকে কি সাধারনত প্রতিবেলা যে পরিমান খান, অভাবের কারনে তার থেকে কম খেয়ে থাকতে হয়েছে? In the past four weeks, did you or any household member have to eat a smaller meal than you felt you needed because there was not enough food? | [0] না (পরবর্তী 1106 নং প্রশ্নে যান) No [skip to 1106]<br>[1] হ্যাঁ | [1] কদাচিৎ Rarely<br>[2] মাঝে মাঝে Sometimes<br>[3] প্রায়ই Often |
| **C.1106.** | 1106. | 6. গত ৪ সপ্তাহে আপনাকে বা খানার কোনো সদস্যকে, অভাবের কারনে কোনো বেলা না খেয়ে থাকতে হয়েছে কি? In the past four weeks, did you or any household member have to eat fewer meals in a day because there was not enough food? | [0] না (পরবর্তী 1107 নং প্রশ্নে যান) No [skip to 1107]<br>[1] হ্যাঁ | [1] কদাচিৎ Rarely<br>[2] মাঝে মাঝে Sometimes<br>[3] প্রায়ই Often |
| **C.1107.** | 1107. | 7. গত ৪ সপ্তাহে এমনকি হয়েছে যে আপনার খানায় অভাবের কারনে খাওয়ার জন্য কোন খাবার ছিল না? In the past four weeks, was there ever no food to eat of any kind in your household because of lack of resources to get food? | [0] না (পরবর্তী 1108 নং প্রশ্নে যান) No [skip to 1108]<br>[1] হ্যাঁ | 1] কদাচিৎ Rarely<br>[2] মাঝে মাঝে Sometimes<br>[3] প্রায়ই Often |
| **C.1108.** | 1108. | 8. গত ৪ সপ্তাহে আপনাকে বা আপনার খানার অন্য কোনো সদস্যকে কি ঘরে যথেষ্ট খাবার না থাকার কারনে ক্ষুধার্ত অবস্থায় রাতে ঘুমাতে হয়েছে? In the past four weeks, did you or any household member go to sleep at night hungry because there was not enough food? | [0] না (পরবর্তী 1109 নং প্রশ্নে যান) No [skip to 1109]<br>[1] হ্যাঁ | 1] কদাচিৎ Rarely<br>[2] মাঝে মাঝে |

| | | | | |
|---|---|---|---|---|
| | | | | Sometimes<br>[3] প্রায়ই Often |
| **C.1109.** | 1109. | 9. গত ৪ সপ্তাহে আপনাকে বা আপনার খানার অন্য কোনো সদস্যকে কি যথেষ্ট খাবার না থাকার কারনে সারাদিন এবং সারারাত না খেয়ে থাকতে হয়েছে? In the past four weeks, did you or any household member go a whole day and night without eating anything because there was not enough food? | [0] না (পরবর্তী 1110 নং প্রশ্নে যান) No [skip to 1110]<br>[1] হ্যাঁ | 1] কদাচিৎ Rarely<br>[2] মাঝে মাঝে Sometimes<br>[3] প্রায়ই Often |
| **C.1110.** | 1110. | 10. গত ৪ সপ্তাহে আপনার খানায় যথেষ্ট চাল না থাকার কারনে কি আপনাকে বা আপনার খানার অন্য কোনো সদস্য কে চাল ধার করে এনে খেতে হয়েছে? In the past four weeks, did you or any household member have to borrow rice because you did not have enough rice? | [0] না (পরবর্তী 1111 নং প্রশ্নে যান) No [skip to 1111]<br>[1] হ্যাঁ | 1] কদাচিৎ Rarely<br>[2] মাঝে মাঝে Sometimes<br>[3] প্রায়ই Often |
| **C.1111.** | 1111. | 11. বছরের অন্যান্য সময়ের তুলনায় আপনার খানা কি কার্তিক/চৈত্র মাসে ( মঙ্গাকালীন সময়) খাদ্যের অভাবে থাকে?<br>Compared to other times, does your household face food deficiency during Kartik/chyatra (Monga)? | [0] না (পরবর্তী 1112 নং প্রশ্নে যান) No [skip to 1112]<br>[1] হ্যাঁ | [1] কোন পার্থক্য নাই<br>No difference<br>[2] গুনগত মান কমে<br>Reduced quality<br>[3] পরিমানগত মান কমে<br>Reduced quantity |
| **C.1112.** | 1112. | 12. আপনার খানার আয় ও খাদ্যের খরচ হিসেব করে আপনি আপনাদের অবস্থাকে কি বলবেন? What would you think is the status of your household in terms of food availability? | | [1] সবসময় ঘাটতি<br>Always deficit<br>[2] মাঝে মাঝে ঘাটতি<br>Deficit sometimes<br>[3] ঘাটতিও না উদ্বৃত্তও না<br>Adequate (Neither deficit nor surplus)<br>[4] খাদ্য উদ্বৃত্ত থাকে<br>Food surplus |

**Wash Benefit Module 7 HANDWASHING**

| | WBM 7 | | |
|---|---|---|---|
| C.700 | 700. | এই প্রশ্নের আগে উত্তরদাতা যে কোন সময় তার হাত ধুয়েছিল কিনা তা পর্যবেক্ষণ করুন।RECORD WHETHER THE RESPONDENT HAS WASHED HER HANDS AT ANY TIME BEFORE THIS QUESTION | [1] উত্তরদাতা হাত ধুয়েছিল OBSERVED RESPONDENT WASHING HANDS<br>[2] উত্তরদাতা হাতধোয় নাই DID NOT OBSERVE RESPONDENT WASHING HANDS |
| C.701 | 701. | আপনাকে ধন্যবাদ । এখন আমি আপনার হাতগুলো দেখতে চাই। আশা করি আপনি কিছু মনে করবেন না। আপনি কি আমাকে দেখাবেন? READ: Thank you. Now, I would like to do a quick inspection of your hands. I hope you don't mind. Can you please show me your hands?<br><br>দুই হাতই দেখতে হবে(প্রথমে এক হাত)। এরপর নিচের উল্লেখিত হাতের অবস্থার কোড দেখে রেকর্ড করতে হবে। BOTH HANDS SHOULD BE SHOWN (NOT JUST ONE HAND). USE <u>APPEARANCE CODES</u> BELOW TO RECORD THE DESCRIPTION | 👁 MOTHER<br><br>**বাম হাত Left Hand**<br>A\|__\| হাতের নখ FINGERNAILS<br>B\|__\| হাতের তালু PALMS<br>C\|__\| আঙ্গুলের সম্মুখভাগ FINGER PADS<br><br>**ডান হাত Right Hand**<br>D\|__\| হাতের নখ FINGERNAILS<br>E\|__\| হাতের তালু PALMS<br>F\|__\| আঙ্গুলের সম্মুখভাগ FINGER PADS |

| | | | |
|---|---|---|---|
| | | THAT BEST DESCRIBES THE DEFINITIONS THE LEVEL OF CLEANLINESS.<br>APPEARANCE CODES:<br>[1] ময়লা স্পষ্টভাবে দেখা যাচ্ছিল VISIBLE DIRT (DIRT/MUD/SOIL/ASH OR ANY OTHER MATERIAL IS VISIBLE)<br>[2] ময়লা স্পষ্টভাবে দেখা নাগেলেও অপরিচ্ছন্নভাব ছিল UNCLEAN APPEARANCE (NO DIRT IS VISIBLE ON THIS PART OF THE HAND BUT, IN GENERAL, THIS PART OF THE HAND APPEARS UNCLEAN)<br>[3] পরিষ্কার ছিল CLEAN (OBSERVED PART OF THE HAND IS CLEAN AS WOULD APPEAR AFTER SOMEONE WASHES HANDS OR TAKES A BATH)<br>৮৮. পর্যবেক্ষণ করা সম্ভব হয়নি/প্রত্যাখান N/A; OBSERVATION NOT POSSIBLE/REFUSED | |
| **C.701** | 702. | ASK: দয়া করে আমাকে(শিশুর নাম ধরে) হাতগুলো দেখাও । Please show me [NAME]'s hands.<br><br>উত্তরদাতার সবচেয়ে বড় বাচ্চার (০-৩৬ মাস) হাত দেখুন । এই বয়সের বাচ্চা না থাকলে স্কিপ কর‍ুন। TO SEE THE HANDS OF THE RESPONDENT'S OLDEST CHILD 0-36 MONTHS (SEE C.1) IF RESPONDENT DOESN'T HAVE A CHILD 0-36 MONTHS → **SKIP TO NEXT PAGE**.<br><br>APPEARANCE CODES:<br>[1] ময়লা স্পষ্টভাবে দেখা যাচ্ছিল(**ময়লা/কাঁদা/মাটি/ছাই /এছাড়া অন্য কিছু**)VISIBLE DIRT (DIRT/MUD/SOIL/ASH OR ANY OTHER MATERIAL IS VISIBLE)<br>[2] ময়লা স্পষ্টভাবে দেখা না গেলেও অপরিচ্ছন্নভাব ছিল(হাতে **ময়লা দেখা না গেলেও** অপরিচ্ছন্ন ছিল)UNCLEAN APPEARANCE (NO | সবচেয়ে বড় বাচ্চার ০-৩৬ মাস OLDEST CHILD 0-36 MONTHS<br><br>**বাম হাত Left Hand**<br>A\|__\| হাতের নখ FINGERNAILS<br>B\|__\| হাতের তালু PALMS<br>C\|__\| আঙ্গুলের সম্মুখভাগ FINGER PADS<br><br>**ডান হাত Right Hand**<br>D\|__\| হাতের নখ FINGERNAILS<br>E\|__\| হাতের তালু PALMS<br>F\|__\| আঙ্গুলের সম্মুখভাগ FINGER PADS |

| | | | |
|---|---|---|---|
| | | DIRT IS VISIBLE ON THIS PART OF THE HAND BUT, IN GENERAL, THIS PART OF THE HAND APPEARS UNCLEAN)<br>[3] পরিষ্কার ছিল(হাতধোয়ার পর/গোসলের পর হাত দেখলে পরিষ্কার হবে) CLEAN (OBSERVED PART OF THE HAND IS CLEAN AS WOULD APPEAR AFTER SOMEONE WASHES HANDS OR TAKES A BATH)<br>[88] পর্যবেক্ষণ করা সম্ভব হয়নি/প্রত্যাখান/০-৩৬ মাসের বাচ্চা নাই N/A; OBSERVATION NOT POSSIBLE/REFUSED/ NO CHILD 0-36 MONTHS | |
| C.702a | 703. | পর্যবেক্ষনঃ প্রাথমিক/ **প্রধান** হাত ধোয়ার স্থানটি কোথায় তা রেকর্ড করুন<br>**[Observation: Record the location where the primary hand washing station is located?]** | 1. ঘরের ভিতরে/ কাছে (≤৬ কদম) In/near main house (≤6 feet to entrance)<br>2. পায়খানার ভিতরে/ কাছে (≤৬ কদম) In/near latrine (≤6 feet to entrance)<br>3. রান্নার স্থানের ভিতরে/ কাছে (≤৬ কদম) In/near cooking area (≤6 feet to entrance)<br>4. ঘর, পায়খানা, রান্নার স্থানে থেকে >৬ কদম >6 feet away from main house, latrine and cooking area<br>**5.** নির্দিষ্ট কোন জায়গায় নাই No specific place 714 নং প্রশ্নে চলে যান (skips to 714) |
| C.702c | 704. | পর্যবেক্ষনঃ প্রাথমিক/ **প্রধান** হাত ধোয়ার স্থানটিতে কোন কোন উপাদান আছে? (আপনি যদি দেখে থাকেন তাহলে "1" কোড করুন আর না দেখলে "0" কোড করুন) [Observation: Which of the following are present at the primary hand washing station? (If you observe the listed item, write "1" for "yes" in the box below. If you do not observe the listed item, write "0" for "no" in the box below.)]<br>*1= হ্যাঁ [Yes], 0= না [No]* | MATERIALS PRESENT<br>[1] **পানি** WATER<br>[2] **গোসল/হাত ধোয়ার সাবান (লাক্স, লাইফবয়)** BAR SOAP (Body/hand Bar)<br>[3] **অন্য যে কোন ধরনের সাবান (হুইল)** BAR SOAP (other)<br>[4] **গুড়া পাউডার** POWDERED SOAP<br>[5] **সাবান-পানি** SOAPY WATER<br>[6] **তরল সাবান** LIQUID SOAP<br>[7] **ছাই** Ash<br>[8] **মাটি /বালু** Mud/Sand<br>[9] **আইসিডিডিআর, বি-র দেওয়া ট্যাপসহ বালতি** ICDDR,B drum with tap<br>[10] হাত ধোয়ার স্থানটি ভিজা ছিল/ব্যবহারের চিহ্ন ছিল Moisture below handwashing station<br>[11] **বালতি** Bucket<br>[12] **বেসিন, জগ** Basin/Jug<br>[13] **কিছুই নাই** NONE OF THE ABOVE<br>**[777]** **অন্যান্য** OTHER (SPECIFY): _____________ |

| | | | |
|---|---|---|---|
| | 705. | **প্রধান হাত ধোয়ার স্থানটি সাবান জাতীয় যেকোন ধরনের উপাদান (যা হাত ধোয়ার কাজে ব্যবহার করা হয়/ সাবান পানি ও পানি একসাথে উপস্থিত আছে কী?** [Soap/soapy water and water present together in PHWS] | 1. হ্যাঁ [Yes]<br>0. না [No] |
| C.702c | 706. | প্রাথমিক/ **প্রধান** হাত ধোয়ার স্থানটি রান্নাঘর থেকে কত কদম দূরে? [Observation: The primary hand washing station is away from the kitchen---Count in steps, allow in continuous numbers] | \|____\| |
| C.702d | 707. | প্রাথমিক/ **প্রধান** হাত ধোয়ার স্থানটি পায়খানা থেকে কত কদম দূরে? [Observation: The primary hand washing station is away from the toilet---Count in steps, allow in continuous numbers] | \|____\| |
| C.703 | 708. | উত্তরদাতাকে প্রশ্ন করুনঃ আপনার হাতধোয়ার জন্য প্রাথমিক/ **প্রধান** স্থান ছাড়া অন্যকোন জায়গা আছে কি? [Ask the respondent: "Is there anywhere else you wash your hands?" (Record code in box)]: | 1. হ্যাঁ [Yes]<br>0. না [No] **714** নং প্রশ্নে চলে যান (skips to 714) |
| C.703a | 709. | পর্যবেক্ষনঃ দ্বিতীয় প্রধান হাত ধোয়ার স্থানটি কোথায় তা রেকর্ড কর্"ন [Observation: Record the location of the secondary hand washing station]. | 1. ঘরের ভিতরে/ কাছে (≤৬ কদম) In/near main house (≤6 feet to entrance)<br>2. পায়খানার ভিতরে/ কাছে (≤৬ কদম) In/near latrine (≤6 feet to entrance)<br>3. রান্নার স্থানে ভিতরে/ কাছে (≤৬ কদম) In/near cooking area (≤6 feet to entrance)<br>4. ঘর, পায়খানা, রান্নার স্থানে থেকে >৬ কদম >6 feet away from main house, latrine <u>and</u> cooking area |
| C.703c | 710. | পর্যবেক্ষনঃ দ্বিতীয় প্রধান হাত ধোয়ার স্থানটিতে কোন কোন উপাদান আছে? (আপনি যদি দেখে থাকেন তাহলে "1" কোড কর্"ন আর না দেখলে "0" কোড কর্"ন) [Observation: Which of the following are present at the secondary hand washing station? (If you observe the listed item, write "1" for "yes" in the box below. If you do not observe the listed item, write "0" for "no" in the box below.) ] | *<u>1= হ্যাঁ [Yes], 0= না [No]</u>*<br>[1] **পানি** WATER<br>[2] **গোসল/হাত ধোয়ার সাবান (লাক্স, লাইফবয়)** BAR SOAP (Body/hand Bar)<br>[3] **অন্য যে কোন ধরনের সাবান (হুইল)** BAR SOAP (other)<br>[4] **গুড়া পাউডার** POWDERED SOAP<br>[5] **সাবান-পানি** SOAPY WATER<br>[6] **তরল সাবান** LIQUID SOAP<br>[7] **ছাই** Ash<br>[8] **মাটি /বালু** Mud/Sand<br>[9] **আইসিডিডিআর, বি-র দেওয়া ট্যাপসহ বালতি** ICDDR,B drum with tap<br>[10] হাত ধোয়ার স্থানটি ভিজা ছিল/ব্যবহারের চিহ্ন ছিল Moisture below handwashing station<br>[11] **(বালতি** Bucket<br>[12] **বেসিন, জগ** Basin/Jug<br>[13] **কিছুই নাই** NONE OF THE ABOVE<br>**৭৭৭. অন্যান্য** OTHER (SPECIFY): _____________ |

| | | | |
|---|---|---|---|
| | 711. | **দ্বিতীয় প্রধান হাত ধোয়ার স্থানটিতে সাবান জাতীয় যেকোন ধরনের উপাদান (যা হাত ধোয়ার কাজে ব্যবহার করা হয়/সাবান পানি ও পানি একসাথে উপস্থিত আছে কী?** [Soap/soapy water and water present together in SHWS] | 1. হ্যাঁ [Yes]<br>0. না [No] |
| C.703c | 712. | দ্বিতীয় প্রধান হাত ধোয়ার স্থানটি রান্নাঘর থেকে কত কদম দূরে? [Observation:The secondary hand washing station is away from the kitchen---Count in steps, allow in continuous numbers]] | \|____\| |
| C.703d | 713. | দ্বিতীয় প্রধান হাত ধোয়ার স্থানটি পায়খানা থেকে কত কদম দূরে? [Observation: The secondary hand washing station is away from the toilet---Count in steps, allow in continuous numbers] | \|____\| |
| INDICATOR 4: HANDWASHING সাবানের উপস্থিতিতে হাতধোয়া দেখানো DEMONSTRATION, SOAP PRESENCE | | | |
| আপনাকে ধন্যবাদ। আপনি দয়া করে আমাকে পায়খানা থেকে আসার পর যেখানে বেশীরভাগ সময় হাত ধোন সেই স্থানটি দেখান(নোট ;পূর্বে পর্যবেক্ষীত স্থানও হতে পারে। এরপর উত্তরদাতা কর্তৃক চিহ্নিত স্থানে যেতে হবে এবং রেকর্ড করতে হবে।Thank you. Please show me where you most often wash your hands after defecation. (Note: this may be the same place you already observed. Go to the place identified by the respondent and record location). | | | |
| C.704 a | 714. | 👁 হাতধোয়ার স্থান পর্যবেক্ষণ করুন(একটিকে গোলাকার করুন)। OBSERVE AND RECORDLOCATION OF HANDWASHING (CIRCLE ONE) | 1. ঘরের ভিতরে/ কাছে (≤৬ কদম) In/near main house (≤6 feet to entrance)<br>2. পায়খানার ভিতরে/ কাছে (≤৬ কদম) In/near latrine (≤6 feet to entrance)<br>3. রান্নার স্থানে ভিতরে/ কাছে (≤৬ কদম) In/near cooking area (≤6 feet to entrance)<br>4. ঘর, পায়খানা, রান্নার স্থানে থেকে >৬ কদম >6 feet away from main house, latrine <u>and</u> cooking area<br>**5.** নির্দিষ্ট কোন জায়গায় নাই |
| **সময় গননার জন্য ষ্টপ ওয়াচ প্রস্তুতকরন PREPARE YOUR STOP WATCH FOR TIMING.** | | | |
| C.705 | 715. | পায়খানা থেকে আসার পর যেভাবে আপনি আপনার হাতধোন একইভাবে আমাকে এখন তা করে দেখান। Now, can you demonstrate how you normally clean your hands <u>after defecation</u>? Try in the same manner as you would if I wasn't here.<br>TIME HOW LONG IT TAKES THE RESPONDENT TO BRING MATERIALS TO THAT PLACE (START AT THE MOMENT YOU FINISH ASKING THE QUESTION). WHEN RESPONDENT STARTS CLEANING HER HANDS, TIME DURATION OF RUBBING HANDS TOGETHER. | [1] দেখানো হয়েছে DEMONSTRATED<br><br>[2] দেখানো হয়নি COULD NOT DEMONSTRATE<br>IF NOT → **RECORD REASON:**<br>**__________________________________AND <u>SKIP TO 721</u>** |
| C.705 b | 716. | 👁পর্যবেক্ষণ করুন এবং হাতধোয়ার সাবান আনতে কতটুকু সময় লেগেছে তা রেকর্ড করুন। যদি হাতধোয়ার স্থানে | \|____:____\| মি.mm. সে.ss |

| | | | |
|---|---|---|---|
| | | সাবান থাকে তাহলে ০০:০০ বসান। যদি সাবান ব্যবহার না করে থাকে তাহলে ৮৮:৮৮ বসান । OBSERVE AND RECORD TIME TO PRODUCE (Bangladesh: SOAP) FOR CLEANING HANDS MARK "00:00" IF SOAP IS ALREADY PRESENT AT HANDWASHING PLACE. (Bangladesh ONLY) MARK "88:88" IF SOAP IS NT USED | |
| C.705 c | 717. | ় পর্যবেক্ষণ এবং হাতঘষার সময় রেকর্ড করুন \| উত্তরদাতা যখন হাত পরিস্কারের জন্য দুহাত ঘষবে সেই সময় গনণা করতে হবে । OBSERVE AND RECORD LENGTH OF TIME RESPONDENT SPENDS RUBBING HER HANDS TOGETHER: | \|____:____\| মি.mm. সে.ss |
| C.705 d | 718. | ় পর্যবেক্ষণ এবং হাত ধোয়ার সকল উপাদান রেকর্ড করুন OBSERVE AND RECORD ALL MATERIALS USED FOR HAND CLEANSING (CIRCLE ALL THAT APPLY) 1 হ্যাঁ Yes 0 না No | 1. **পানি** WATER<br>2. **বার সাবান** BAR SOAP (Body/hand BAR)<br>3. **বার সাবান** (অন্যান্য)BAR SOAP (other)<br>4. **গুড়া পাউডার** POWDERED SOAP<br>5. **সাবান-পানি** SOAPY WATER<br>6. **কাপড়/পাতা** CLOTH OR LEAVES<br>7. **বেসিন/কলস**BASIN/PITCHER<br>8. **ছাই** ASH<br>9. মাটি/বালু MUD<br>88. **অন্যান্য উলে- খ করুন** OTHER (SPECIFY):<br>______________________________ |
| C.705 e | 719. | ় পর্যবেক্ষণ এবং ২ হাত ধোয়া রেকর্ড করুন \|OBSERVE AND RECORD WHETHER BOTH HANS WERE CLEANED | [1] হ্যাঁ, ২-হাত ধুয়েছে YES, BOTH HANDS CLEANED<br>[2] না ,১-হাত ধুয়েছে NO, ONLY ONE HAND CLEANED |
| C.705f | 720. | ় পর্যবেক্ষণ এবং হাত শুকানো রেকর্ড করুন OBSERVE AND RECORD HOW RESPONDENT DRIES HANDS<br>1 Yes 0 No | 1. পরিধেয় কাপড় DRIES BY WIPING HER HANDS ON HER CLOTHES<br>2.এক হাত অন্য হাতের সাথে ঘষে DRIES BY WIPING HER HANDS ON ANOTHER<br>3. পরিধেয় ছাড়া অন্য পরিষ্কার কাপড় Clean CLOTH DRIES BY WIPING HER HANDS ON<br>৪. পরিধেয় ছাড়া অন্য অপরিষ্কার কাপড় Unclean CLOTH DRIES BY WIPING HER HANDS ON<br>5. কাপড় ছাড়া অন্য জিনিস যেমন: টিস্যু, কাগজ ইত্যাদি MATERIAL (NOT CLOTH)<br>5. বাতাসে/হাত ঝাঁকিয়ে বাতাসে শুকানো হয়েছে DRIES BY SHAKING HER HANDS IN THE AIR |

| | | | |
|---|---|---|---|
| | | | 6. হাত শুকানো হয় নাই NOT DRIED<br>88. অন্যান্য উলে- খ কর ্ন OTHER,<br>SPECIFY_________________________ |
| C.705 g | 721. | **যদি উত্তরদাতা হাতধোয়া না দেখান অথবা সাবান ব্যবহার না করে তখন তাকে তার খানায় হাতধোয়ার জন্য সাবান আছে কিনা জানতে হবে এবং তা আনতে বলতে হবে?**IF RESPONDENT COULD NOT DEMONSTRATE, OR DID NOT USE SOAP, ASK:<br>Do you have soap in your house that you use for handwashing? Can you bring it to me?<br><br>পর্যবেক্ষণ কর ্ন এবং হাতধোয়ার সময় রেকর্ড কর ্ন। যদি 716 তে উত্তর/সময় উলে- খ থাকে তাহলে ৮৮:৮৮ বসান। যদি খানাতেসাবান না থাকে তাহলে ৯৯:৯৯ বসান । OBSERVE AND RECORD TIME TO PRODUCE SOAP FOR WASHING HANDS MARK “88:88” IF ALREADY TIMED IN C.704a MARK “99:99” IF OBSERVATION NOT POSSIBLE OR SOAP NOT AVAILABLE | \|____:____\| মি.mm. সে.ss |
| C. 706a | 722. | পর্যবেক্ষণ কর ্ন : **উত্তরদাতার হাতধোয়া দেখানোর সময়ে এ বিষয়ে কোন প্রতিক্রিয়া দেখতে পেয়েছিলেন কী?**OBSERVE: WAS THERE ANY OBVIOUS REACTIVITY DURING THE COURSE OF HIS DEMONSTRATION? | [1] *হ্যাঁ YES*<br>[0] *না (৭২৩ নং এ চলে যাবে)* NO → skip to 723 |
| b | 1 | অধিককক্ষণ যাবৎ হাত ধুয়েছে কিনা<br>Longer time spent cleaning/rubbing hands | [1] *হ্যাঁ* YES<br>[0] *না* NO |
| c | 2 | সাবান দিয়ে হাত ধুয়েছে কিনা অন্যথায়<br>Using soap where they otherwise would not | [1] *হ্যাঁ* YES<br>[0] *না* NO |
| d | 3 | **অন্যান্য (লিখুন)** Other, (BRIEFLY explain)<br>__________________________<br>_ | [1] *হ্যাঁ* YES<br>[0] *না* NO |

| | | | | | |
|---|---|---|---|---|---|
| | 723. | প্রশ্ন করুন ; আপনি কখন কখন সাবান দিয়ে হাত ধোন । ASK: "Please tell me about all of the times you wash your hands with soap."<br><br>A কলা ম (লিড ছাড়া)-এ যখন উত্তরদাতা যে কাজ করবে তা উলে- খ করতে হবে । ১নং গোলাকার হবে যদি নির্দিষ্ট বিষয়ে উত্তর পাওয়া যায়। CIRCLE "1" IN COLUMN A IF CRITICAL TIME IS MENTIONED WITHOUT PROMPTING.<br><br>উত্তদাতার সকল কাজ উলে- খ করার পর অন্য আর কোন কাজে সাবান দিয়ে হাত ধোন কিনা এ ব্যপারে জিজ্ঞাসা করুন? AFTER THE RESPONDENT FINISHES NAMING ALL THE TIMES ASK "IS THERE ANY OTHER TIME YOU WASH YOUR HANDS WITH SOAP?" AND STOP WHEN THE RESPONDENT SAYS THERE IS NO OTHER TIME.<br><br>B কলাম-এ লিড দেওয়া ছাড়া প্রতিবার ব্যবহার করে কিনা তা উলে- খ করুন । EACH TIME MENTIONED WITHOUT PROMPTING ASK COLUMN B.<br><br>C.707A-G নং প্রশ্নের উত্তর যদি A কলা ম [1] না হয় তখন সাবান ও পানি দিয়ে আপনি আপনার হাতধোন কিনা তা জিজ্ঞাসা করুন। যদি উত্তদাতা হ্যাঁ বলে তখন C কলা ম-এ হাতধোয়ার সংখ্যা/বার উলে- খ করুন । FOR QUESTIONS C.707A-G WHERE [1] IS NOT CIRCLED IN COLUMN A, PROMPT BY ASKING:<br>Do you usually wash your hands with soap and water [CRITICAL TIME]? IF THE RESPONDENT SAYS "YES" THEN IMMEDIATELY ASK HOW OFTEN AND MARK APPROPRIATE ANSWER CHOICE IN COLUMN C. | A.<br>লিড দেওয়া ছাড়া হাতধোয়া উলে- খ করুন<br>MENTIONED WITH NO PR OMPTING | B.<br>লিড দেওয়া ছাড়া শুধুমাত্র হাতধোয়ার সংখ্যা/বার উলে- খ করুন । সাবান ও পানি দিয়ে কত বার আপনার হাতধোন তা বলুন । Only for times mentioned with NO PROMPTING, ask: How often do you wash your hands with soap and water [insert event]?<br><br>[1] সবসময়/প্রতিবার ALWAYS<br>[2] মাঝে মাঝে/ অর্ধেকের কম বার SOMETIMES<br>[3] খুব কম সময়/ দু-এক বার RARELY | C.<br>লিড দেওয়ার পর শুধুমাত্র হাতধোয়ার সংখ্যা/বার উলে- খ করুন । সাবান ও পানি দিয়ে কত বার আপনার হাতধোন তা বলুন । REPLIED "AFTER BEING PROMPTED<br><br>[1] সবসময়/প্রতিবার ALWAYS<br>[2] মাঝে মাঝে/ অর্ধেকের কম বার SOMETIMES<br>[3] খুব কম সময়/ দু-এক বার RARELY<br>[4] না NO<br>[88] প্রযোজ্য নয় Not applicable |
| a | 1 | খাবার তৈরী করার আগে BEFORE PREPARING FOOD | | | |
| | 2 | মাংস/মাছ কাটার পর AFTER HANDLING MEAT/FISH | | | |
| | 3 | ফল/সব্জি কাটার আগে BEFORE CUTTING FRUITS AND VEGETABLES | | | |
| | 4 | ভর্তা/ম্যাসড খাবার তৈরী করার আগে BEFORE MAKING MASHED | | | |

| | | |
|---|---|---|
| | | FOOD (BHORTA) |
| b | 5 | খাবারের পূর্বে BEFORE EATING |
| c | 6 | খাবারের পরে AFTER EATING |
| d | 7 | শিশুকে খাওয়ানোর আগে BEFORE FEEDING A CHILD |
| e | 8 | বাচ্চাকে সৌচানোর পর AFTER CLEANING A CHILD'S ANUS |
| f | 9 | বাচ্চার পায়খানা ফেলার পর AFTER DISPOSING OF CHILDREN'S FECES |
| g | 10 | পায়খানার পর AFTER TDEFECATION |
| h | 11 | কম্পাউন্ড/বাড়ীর বাহির থেকে ফেরার পর AFTER RETURNING FROM OUTSIDE THE COMPOUND |
| m | 12 | রুগ্ন ব্যক্তিকে ধরার পর AFTER TOUCHING A SICK PERSON |
| o | 13 | পশু-পাখি ধরার পর AFTER HANDLING LIVESTOCK |
| | 14 | গোবর ধরার পর AFTER HANDLING COW-DUNG (OR ANY ANIMAL FECES) |
| j | 15 | **অন্যান্য (উলে- খ করুন)** OTHER (SPECIFY): ______________________________ _______________ |
| i | 16 | **উত্তরদাতা কখনই সাবান দিয়ে হাতধোয় নি** RESPONDENT NEVER WASHES HANDS WITH SOAP Skip to module 8 |
| | 17 | **উঠান ঝাড়ু দেওয়া/ময়লা পরিষ্কারের পর** After weaping courtyard |
| | 18 | **রান্নার পর** After cooking |
| | 19 | **থালা-বাসন ধোয়ার পর** After washing dishes |

**Wash Benefit Module 8 SANITATION (All households)**

| | |
|---|---|
| SAY | এই গবেষণায় আপনার অংশ গ্রহনের জন্য আপনাকে ধন্যবাদ। ষ্টাডির এই অংশটি কিছুটা স্পর্শকাতর । আমি আপনাকে আপনার/এই বড়ীর পয়ঃ ব্যবস্থাপনা/পয়ঃ নিষ্কাশন ও তার অনুশীলন এবং কিভাবে তা করেন এ ব্যপারে জানতে চাই \| আমি কিছুটা পর্যবেক্ষণ করতে চাই। আপনার স্বাচ্ছন্দবোধ হবে কি ? যদি না হয় তাহলে উত্তরদাতাকে ইহার গুরুত্ব বোঝাতে চেষ্টা করুন।Thank you so much for your participation so far. The next part of the survey is a bit sensitive. I would like to ask you some questions about the sanitation practices of people in your compound, including how you usually dispose of your children's feces. I would also like to make some observations. Are you comfortable with this? IF NOT, EXPLAIN THE IMPORTANCE OF THIS MODULE AND ENCOURAGE RESPONDENT TO PROCEED. |

খোলা-পায়খানা(সকল খানার ) OPEN DEFECATION (ADMINISTER TO ALL HOUSEHOLDS)

| | | | **Group A** | **B** | **C** | **D** | **E** |
|---|---|---|---|---|---|---|---|
| | WBM 8 | | **Men** পুরুষ | **Women** মহিলা | **Children <3 years** <৩ বছরের বাচ্চা | **Children 3-<8 years** ৩-<৮ বছরের বাচ্চা | **Children 8-15** ৮-১৫ ছরের বাচ্চা |
| **C.801.** | 801. | **এই খানার কোন সদস্যরা খোলা জায়গাতে পায়খানা করে কী?** [Do [GROUP] in this household ever | ☐ | ☐ | ☐ | ☐ | ☐ |

| | | | | | | | |
|---|---|---|---|---|---|---|---|
| | | practice open defecation?<br>1 প্রতিদিন Daily<br>2 মাঝেমাঝে Occasionally<br>3 কখনই না Never **(Skip to 806)**<br>88. প্রযোজ্য নয় Not applicable (**806** নং প্রশ্নে যান) **(Skip to 806)**<br>99 জানি না (**806** নং প্রশ্নে যান) Don't Know **(Skip to 806)** | | | | | |
| **C.802.** | 802. | সাধারনত: একই জায়গাতে প্রতিবার যান কী? [Do [GROUP] go to more or less the same area every time?<br>1 হ্যাঁ Yes<br>2 না No (**805** নং প্রশ্নে যান) **(Skip to 805)**<br>99 জানি না Don't Know / Not Sure (**805** নং প্রশ্নে যান) **(Skip to 805)** | ☐ | ☐ | ☐ | ☐ | ☐ |
| **C.803.** | 803. | আপনার খানা থেকে উক্তস্থানে যেতে (শুধু যাওয়া) কতটুকু সময় লাগে (মিনিট) [How long does it take to walk (one way) from your house to the most commonly visited place? | \|______\| মিনিট MINUTES<br>99 জানি না Don't Know / Not Sure | | | | |
| **C.804.** | 804. | এই জায়গাটি ঐ গ্রামের মধ্যেই কী? [Is that place within the village?<br>1হ্যাঁ Yes<br>2 না No<br>99 জানি না Don't Know / Not Sure | \|__\| | | | | |

খোলা পায়খানা বিষয়ে প্রশ্নাবলী **Open Defecation Questions**

| | | | **Men** পুরুষ | **Women** মহিলা | **Children <3 years** <৩ বছরের বাচ্চা | **Children 3-<8 years** ৩-<৮ বছরের বাচ্চা | **Children 8-16** ৮-১৫ ছরের বাচ্চা |
|---|---|---|---|---|---|---|---|
| **C.804a** | 805. | আপনার খানার লোকজনের খোলা জায়গায় পায়খানা করার প্রধান কারন কি কি? উত্তর শুনানো যাবে না।<br>What are the main reasons that [GROUP] in your household practice open defecation?<br>☟DO NOT READ RESPONSES MARK ALL THAT APPLY | | | | | |
| | 1 | 1 কোনও কিছুই নাই /পায়খানা নাই No choice (nothing else is available) | ☐ | ☐ | ☐ | ☐ | ☐ |
| | 2 | 2 কোথায় শিশু পায়খানা করবে তা নিয়ন্ত্রন করতে পারে না Cannot control where young children defecate | | | ☐ | ☐ | ☐ |
| | 3 | 3 গোপনীয়তা Privacy | ☐ | ☐ | ☐ | ☐ | ☐ |
| | 4 | 4 অভ্যাস/নিয়মিত কাজ Habit / Routine | ☐ | ☐ | ☐ | ☐ | ☐ |
| | 5 | 5 পায়খানা থেকে বাঁশঝাড় পছন্দ Prefer to use the bush rather than a toilet | ☐ | ☐ | ☐ | ☐ | ☐ |

| | | | | | | | |
|---|---|---|---|---|---|---|---|
| | 6 | 6 কাজে/স্কুলে পায়খানা/টয়লেট সহজে **পাওয়া না গেলে** Toilet not available at work / school | ☐ | ☐ | ☐ | ☐ | ☐ |
| | 7 | ভাগের পায়খানা/টয়লেট পছন্দ **না হলে** Choose not to share toilets with in-laws / extended family (or cannot) | ☐ | ☐ | ☐ | ☐ | ☐ |
| | 8 | 8 সুবিধাজনক Convenience | ☐ | ☐ | ☐ | ☐ | ☐ |
| | 9 | 9 নিরাপদ Safety | ☐ | ☐ | ☐ | ☐ | ☐ |
| | 10 | 10 আরামদায়ক Comfort | ☐ | ☐ | ☐ | ☐ | ☐ |
| | 11 | 11 অসুস্থতা/ পাতলা পায়খানা Sickness/diarrhoea | ☐ | ☐ | ☐ | ☐ | ☐ |
| | 12 | 12 পায়খানা/টয়লেট ভরে গেছে Latrine overflowed | ☐ | ☐ | ☐ | ☐ | ☐ |
| | 13 | 13 পায়খানা/টয়লেট ভেঙে গেছে(উপরের অংশ/স্লাব) Latrine broken (superstructure and /or slab) | ☐ | ☐ | ☐ | ☐ | ☐ |
| | 14 | 14 পায়খানা/টয়লেটে যেতে ভয় পায় Fear of latrine | ☐ | ☐ | ☐ | ☐ | ☐ |
| | 15 | 15 পায়খানা/টয়লেট কিভাবে ব্যবহার করতে হয় তা জানে না Don't know how to use the latrine | ☐ | ☐ | ☐ | ☐ | ☐ |
| | 16 | 16 **বাচ্চা** খুব ছোট হওয়ায় পায়খানা/টয়লেট ব্যবহার করে না Too young to use latrine | ☐ | ☐ | | | |
| | 17 | বয়স্ক (খুবই বৃদ্ধ) Old age (too old) | | | | | |
| | | | <৩ বছরের বাচ্চা **Children < 3 years** | ৩-<৮ বছরের বাচ্চা **Children 3 – <8 years** | ৮-১৫ বছরের বাচ্চা **Children 8 – 15 years** | পুরুষ **Men** | মহিলা **Women** |
| **C.X** | 806. | **পায়খানা থাকা সত্ত্বেও এই বাড়ীর অন্য কোন পরিবার খোলা জায়গায় পায়খানা করে কিনা তা আপনি জানেন কী?** Do you know of other households in the bari/compound whose [GROUP] practice OD, even if they might have a toilet or latrine at their house?<br>1 **হ্যাঁ, প্রায়ই** Yes, Often<br>2 **হ্যাঁ, মাঝে মাঝে** Yes, Sometimes<br>3 **না, কখনই না** No, Never **(Skip to 808)**<br>88. **প্রযোজ্য নয়** Not applicable (**808** নং প্রশ্নে যান) **(Skip to 808)**<br>99 **জানি না/নিশ্চিত না** Don't Know / Not Sure | ☐ | ☐ | ☐ | ☐ | ☐ |
| **C.X** | 807. | **আপনি শিশুদের পায়খানা উঠানে/উঠানের চার-পাশে/আশেপাশে /নদীতে ফেলে দিতে দেখেছেন কী?** Do you see that children's stools are disposed in the yard / surrounding / community in your neighborhood / | | | | ☐ | ☐ |

| | | |
|---|---|---|
| | | river?<br>1 হ্যাঁ, প্রায়ই Yes, Often<br>2 হ্যাঁ, মাঝে মাঝে Yes, Sometimes<br>3 না, কখনই না No, Never<br>99 জানি না/নিশ্চিত না Don't Know / Not Sure |

| পয়ঃ ব্যবস্থাপনা সুবিধাদি **Sanitation Facility** | | | |
|---|---|---|---|
| Administer to: All study households | | | |
| **C.805.** | 808. | আপনার খানায় কোন টয়লেট/পায়খানা আছে কি?<br>আমি কি তা দেখতে পারি?<br>Does your household have a toilet facility that is in use? Can I see it? | 1 হ্যাঁ আছে,পর্যবেক্ষন করতে পারি Yes have toilet, can observe<br>2 হ্যাঁ আছে,পর্যবেক্ষন করতে দেওয়া হয়নি/প্রত্যাখান (৮১২ নং এ যান ) Yes have toilet, refused observation **(Skip to 812)**<br>3 হ্যাঁ আছে,পর্যবেক্ষন করা যায়নি (৮১২ নং এ যান ) Yes have toilet, cannot observe **(Skip to 812)**<br>4 কোন পায়খানা নাই No toilet facility **(Skip to MODULE 9)** |
| **C.806.** | 809. | 👁 পর্যবেক্ষন; ধরণ,অবস্থা এবং পুনঃব্যবহার অবস্থা Observation: Note the type, condition and apparent use of the toilet:<br>1 হ্যাঁ Yes<br>2 না No<br>88 প্রযোজ্য নয়/ পর্যবেক্ষন করতে পারেনি/বলতে পারেনি Not Applicable / Could not observe / cannot tell | |
| বর্হিরাংশ পর্যবেক্ষণ **Exterior observation** | | | |
| 1 | 1 | পায়খানার চারপাশে ৩টি ইটের/ টিনের/মাটির দেয়াল At least 3 walls around the toilet | *(1= হ্যাঁ [Yes], 0= না [No],* প্রযোজ্য নয় *[*not applicable] =888)] |
| 2 | 2 | পায়খানার চারপাশে ৩টি বাঁশের বেড়ার দেয়াল Bamboo fences around the toilet | *(1= হ্যাঁ [Yes], 0= না [No],* প্রযোজ্য নয় *[*not applicable] =888)] |
| 3 | 3 | পায়খানার চারপাশে দরজা/পর্দা দিয়ে ঘেরা Door/curtain or walls that guarantee privacy around the toilet | *(1= হ্যাঁ [Yes], 0= না [No],* প্রযোজ্য নয় *[*not applicable] =888)] |
| 4 | 4 | পায়খানার ছাদ Roof over toilet | *(1= হ্যাঁ [Yes], 0= না [No],* প্রযোজ্য নয় *[*not applicable] =888)] |
| 5 | 5 | বাতাস বের হওয়ার পথ Ventilation pipe | *(1= হ্যাঁ [Yes], 0= না [No],* প্রযোজ্য নয় *[*not applicable] =888)] |
| 6 | 6 | পায়খানায় যাবার রাস্তা দেখে বুঝা যাচ্ছে নিয়মিত ব্যবহার করা হয় (পরিষ্কার, জীর্ণ ইত্যাদি ) Path to the toilet suggests regular use (is clear, well-worn, without grass or any barriers etc.) | *(1= হ্যাঁ [Yes], 0= না [No],* প্রযোজ্য নয় *[*not applicable] =888)] |
| ভিতরের অংশ পর্যবেক্ষণ **Interior observation** | | | |
| 7 | 7 | টয়লেটে স্-াব আছে Toilet has a slab | *(1= হ্যাঁ [Yes], 0= না [No],* প্রযোজ্য নয় *[*not applicable] =888)] |
| 8 | 8 | গর্তের পাশে পাদানী আছে Raised footing around hole | *(1= হ্যাঁ [Yes], 0= না [No],* প্রযোজ্য নয় *[*not applicable] =888)] |
| 9 | 9 | পানি ঢেলে ফ্লাস করা যায় Flush or poor Flush: | *(1= হ্যাঁ [Yes], 0= না [No],* প্রযোজ্য নয় *[*not applicable] =888)] [If 0/888 skip to 12] |

| | | | |
|---|---|---|---|
| 9a | 9a | যদি পানি ঢেলে ফ্লাস করা যায় If Flush or Pour Flush:<br><br>পানির সীলের অবস্থা Water seal condition: | [ 1 ] ওয়াটার সিল ভাল আছে/কাজ করে Functional water seal<br>[ 2 ] ওয়াটার সিল ভাঙ্গা Broken water seal<br>[ 3 ] ওয়াটার সিল নেই No water seal<br>[ 88 ] পায়খানাতে পানির সীল আছে/পর্যবেক্ষণ করা সম্ভব হয়নি N/A / could not observe / cannot tell |
| 9b | 9b | যদি পানি ঢেলে ফ্লাস করা যায় If Flush or Pour Flush:<br><br>পানি ঢাললে/দিলে Flushes to:<br><br>*(খানার সদস্যকে জিজ্ঞাসা/যাচাই করুন Ask / probe household members if necessary)* | ১. টয়লেটিতে পয়ঃনিষ্কাশন পাইপের সাথে সংযোগ করে দেয়া [Piped sewer system]<br>২. টয়লেটিতে সেপটিক ট্যাংক বসানো আছে [Septic tank]<br>৩. সেপটিক ট্যাংক নাই কিন্তু ফ্লাস করে বা পানি ঢেলে পায়খানা দুরে বা পিটের মধ্যে সরিয়ে দেয়া যায় [Flush to pit latrine ]<br>৪. ফ্লাশ-টয়লেট অথবা পানি ঢেলে ফ্লাস করা টয়লেট যা কোন খাল, ড্রেন, নদী ইত্যাদির সাথে সংযোজন করার ফলে অস্বাস্থ্যকর অবস্থার সৃষ্টি করে থাকে [Flush or pour flush toilet connected to somewhere else (canal, ditch, river, etc.)] |
| | 9c | পায়খানার অবস্থা কেমনCondition of the latrine | 1. স্বাস্থ্যসম্মত পায়খানা Hygienic latrine<br>2. অস্বাস্থ্যকর পায়খানা Non hygienic latrine |
| 10 | 10 | মেঝের প্রধান উপাদান Main material of the floor (*select 1*) | [ 1 ] মাটি Mud<br>[ 2 ] কাঠ Wood<br>[ 3 ] সিমেন্ট Cement<br>[ 4 ] টাইলস/ইটTile / brick<br>[ 5 ] প্লাষ্টিক Plastic<br>[ 88 ] প্রযোজ্য নয়/পর্যবেক্ষণ সম্ভব হয় নি N/A / could not observe / cannot tell |
| 11 | 11 | Bucket toilet | *(1= হ্যাঁ [Yes], 0= না [No],* **প্রযোজ্য নয়** *[*not applicable] =888)] |
| 12 | 12 | ঝুলন্ত পায়খানা Hanging toilet | [ 1 Yes ] [ 2 No ] [ 88 N/A] |
| 13 | 13 | ব্যবহৃত হচ্ছে বুঝা যায় এমন Latrine appears to be in use (by your best judgment) | *(1= হ্যাঁ [Yes], 0= না [No],* **প্রযোজ্য নয়** *[*not applicable] =888)] |
| 14 | 14 | **পায়খানা/টয়লেটে পায়খানার গন্ধ রয়েছে** Odor of feces in the latrine/bathroom | *(1= হ্যাঁ [Yes], 0= না [No],* **প্রযোজ্য নয়** *[*not applicable] =888)] |
| 15 | 15 | **পায়খানা/টয়লেটে প্রস্রাবের গন্ধ রয়েছে** Odor of urine in the latrine/bathroom | *(1= হ্যাঁ [Yes], 0= না [No],* **প্রযোজ্য নয়** *[*not applicable] =888)] |
| 16 | 16 | স্লাব অথবা মেঝেতে **পায়খানা দেখতে পাওয়া গেছে** Stool is visible on the slab or floor | *(1= হ্যাঁ [Yes], 0= না [No],* **প্রযোজ্য নয়** *[*not applicable] =888)] |
| 17 | 17 | ড্রপ হোল(মল-মূত্র প্রবেশ গর্ত) ঢাকা Drop hole is covered | *(1= হ্যাঁ [Yes], 0= না [No],* **প্রযোজ্য নয়** *[*not applicable] =888)][If 0/888 skip to 18] |
| 17 a | 17a | যদি ড্রপ হোল (মল-মূত্র প্রবেশ গর্ত)ঢাকা থাকে তাহলে মাছি আসা/যাওয়া করতে পারে কি If yes, cover is correctly placed over the drop hole so that flies cannot enter/exit | *(1= হ্যাঁ [Yes], 0= না [No],* **প্রযোজ্য নয়** *[*not applicable] =888)] |
| 18 | 18 | **মাছির উপস্থিতি রয়েছে** Flies present | *(1= হ্যাঁ [Yes], 0= না [No],* **প্রযোজ্য নয়** *[*not applicable] =888)] |
| 19 | 19 | **সাধারন বৈশিষ্ট General Characteristics**<br>একটি অথবা দুই পিট/গর্ত টয়লেট Single or double pit | [ 1 ] একটি পিট/গত Single pit<br>[ 2 ] দুই পিট/গর্ত Double pit |

| | | | |
|---|---|---|---|
| | | latrine | [ 88 ] প্রযোজ্য নয়/পর্যবেক্ষণ সম্ভব হয় নি /বলতে পারে নিN/A / could not observe / cannot tell |
| 20 | 20 | কমপোষ্টিং টয়লেট Composting toilet | *(1= হ্যাঁ [Yes], 0= না [No],* প্রযোজ্য নয় *[*not applicable] =888)] |
| **C.807.** | 810. | 👁 **পর্যবেক্ষন কর ন** Observation: টয়লেট-এর গর্ত মলে ভরেছে কিনা-আলোর সাহায্যে পর্যবেক্ষণ করুন Fullness of the pit – shine a light into the pit to see if solid waste is… | 1 **শুকনা ময়লা/পায়খানা ৩ ফুটের চেয়ে বেশী দূরে (>৩ ফুট)** Very far from surface (>1 meter)<br>2 **শুকনা ময়লা/পায়খানা ৩ ফুটের মধ্যে (<৩ ফুট)** Within 1 meter<br>3 **পায়খানার খুব কাছাকাছি বা পায়খানা ভরে গেছে** Very close to surface or full<br>88 **পায়খানাতে পানির সীল আছে/পর্যবেক্ষণ করা সম্ভব হয়নি** Water seal latrine / non direct pit latrine / could not observe |
| C. 808 | 811. | 👁 **পর্যবেক্ষন কর ন** Observation: : সৌচ কাজের জন্য টয়লেট-এর ভেতরে এবং কাছে **কী কী উপাদান রয়েছে** What materials for anal cleansing are present inside or immediately outside the latrine? | *(1= হ্যাঁ [Yes], 0= না [No],* প্রযোজ্য নয় *[*not applicable] =888)]<br>1.*পাতা/ঘাস* Leaves/grass<br>2.*কাঠি* Twigs / sticks<br>3. *কাপড়* Rag or cloth<br>4. *পাথর* Stones<br>5. *স্বাস্থ সম্মত কাগজ*Hygienic (toilet) paper<br>6. *পানির পাত্র/বদনা/মগ* Water container / vessel<br>7. *পানির ট্যাপ* Water tap<br>8. *সাবান* Soap<br>9. *ছাই অথবা মাটি* Ash or soil for cleansing<br>10. *খবরের কাগজ* Newspaper<br>11. *কিছুই নাই* Nothing |
| C. 809 | 812. | **আপনার খানার কে কে এই পায়খানা/টয়লেটটি পায়খানার জন্য ব্যবহার করে?** Please tell me about who in your family uses the latrine for defecation. | |
| 1 | 1 | < **3** বাচ্চা Children <3 | 1. সব সময় [Always]<br>2. প্রায়ই [Usually]<br>3. মাঝেমাঝে [Sometime]<br>4. কখনই না [Never]<br>888. প্রযোজ্য নয় [ N/A] |
| 2 | 2 | ৩-<৮ বাচ্চা Children 3-<8 | 1. সব সময় [Always]<br>2. প্রায়ই [Usually]<br>3. মাঝেমাঝে [Sometime]<br>4. কখনই না [Never]<br>888. প্রযোজ্য নয় [ N/A] |
| 3 | 3 | ৮-১৫ বাচ্চা Children 8-15 | 1. সব সময় [Always]<br>2. প্রায়ই [Usually]<br>3. মাঝেমাঝে [Sometime]<br>4. কখনই না [Never]<br>888. প্রযোজ্য নয় [ N/A] |
| 4 | 4 | পুরুষ Men | 1. সব সময় [Always]<br>2. প্রায়ই [Usually]<br>3. মাঝেমাঝে [Sometime]<br>4. কখনই না [Never]<br>888. প্রযোজ্য নয় [ N/A] |
| 5 | 5 | মহিলা Women | 1. সব সময় [Always]<br>2. প্রায়ই [Usually] |

| | | | |
|---|---|---|---|
| | | | 3. মাঝেমাঝে [Sometime]<br>4. কখনই না [Never]<br>888. প্রযোজ্য নয় [ N/A] |
| C. 810 | 813. | অন্য কোন খানা কি এই পায়খানা ব্যবহার করে? Do you share this toilet with other households? | *1= হ্যাঁ [Yes]*<br>*0= না [No].......skip to 815* |
| C.811 | 814. | কতগুলো খানা মিলে এই পায়খানাটি ব্যবহার করে ? How many HHs use it? | (Don't know=99)<br>\|____\|\|____\| |
| C.812 | 815. | এইখানার শিশুসহ আপনারা কতজন এই পায়খানাটি ব্যবহার করেন? How many people including children in your household use this toilet? | \|____\|\|____\| |
| C. 813 | 816. | পায়খানার মালিকানা ? (প্রশ্ন করুন) [***Ask***: Who owns the toilet facility?] | শুধুমাত্র ঐ খানার জন্য [Only for the household] 1<br>কয়েকঘর মিলে/ অংশীদার [Shared] 2<br>অন্য কেউ [Someone else] 3<br>পাবলিক [Public] 4<br>প্রযোজ্য নয় [Not applicable]................................................888 |
| C. 814 | 817. | এই স্থানে এই পায়খানাটি কত বছর ধরে আছে? (জানি না=999) [How long have you had the present latrine in this place? [In years] (Don't know = 99)] | A. \|_____\|\|_____\| বৎসর Years<br>B. \|_____\|\|_____\| মাস Months |

ওয়াস বেনিফিট মডিউল-৯ **Wash Benefit Module 9**

শিশুর পায়খানা ব্যবস্থাপনা এবং মল নিষ্কাশন অভ্যাস **Child defecation and feces disposal practice**

| | | | |
|---|---|---|---|
| শিশুর পায়খানা ব্যবস্থাপনা এবং মল নিষ্কাশন অভ্যাস **Child defecation and feces disposal** | | | |
| Administer to: All study households ( সকল খানায় হবে) | | | |
| C.912 | 901. | \|_____\|\|_____\| কত বছর বয়স থেকে একটি শিশু সাধারনতঃ টয়লেট/পায়খানা ব্যবহার করা শুরু করে (মাসে)? [At what age (in months) does a child start using a latrine, if at all?] | |
| C.902 | 902. | \|_____\| আপনার (<৩ বছর) শিশু সর্বশেষ কখন পায়খানা করেছে? (একাধিক শিশু থাকলে সবচেয়ে বড় বাচ্চার ক্ষেত্রে জিজ্ঞেস করুন) [When was the last time your youngest child/infant (<3 years) defecated?] | 1. আজ [Today]<br>2. গতকাল [Yesterday]<br>3. ২ দিন বা তার আগে [Before 2 or more days back]<br>4. কখনই না/ মনে করতে পারছি না [Never/can't remember]....... **906** নং প্রশ্নে চলে যান (skips to 906)<br>5. বলতে রাজি না [Refused] ..........**906** নং প্রশ্নে চলে যান (skips to 906)<br>88. প্রযোজ্য নয় [Not applicable]......... **908** নং প্রশ্নে চলে যান (skips to 908) |
| **C.903.** | 903. | \|_____\| আপনার শিশু সর্বশেষ কোথায় পায়খানা করেছে? [Where did the child defecate the last time?]<br>**Note:** উত্তর পড়ে শোনাবেন না, তাকে নির্দিষ্ট করে বলতে বলুন । এ বিষয়ে যতক্ষণ পর্যন্ত কিছু বলার থাকে ততক্ষণ পর্যন্ত তাকে বলতে উৎসাহিত করুন [Don't read the answer, encourage by asking if there is anything else until he/she mentions there in nothing else and check all mentioned.] | 1. পটি (উঠানে) [Potty (in the courtyard)]<br>2. পটি (ঘরের ভিতরে) [Potty (inside the house)]<br>3. উঠানে (পটি ছাড়া) [Courtyard (without potty)]<br>4. ঘরের ভিতরে (পটি ছাড়া) [Inside the house (without potty)]<br>5. উঠানের বাহিরে খোলা জায়গাতে [Open space outside the front yard<br>6. ঝোপ-ঝাড়ে/জঙ্গলে Bush/jungle<br>7. পায়খানায়/টয়লেটে [In toilet] ......**906** নং প্রশ্নে চলে যান (Skip to 906)<br>8. কাথা/ন্যাপি/ডায়পার Katha/nappy/diaper<br>77. অন্যান্য (বর্ণনা লিখুন) [Other (Please describe)]..................... |

| | | | |
|---|---|---|---|
| | | | 99. জানি না [Don't Know]............. **906** নং প্রশ্নে চলে যান (Skip to 906) |
| **C.904.** | 904. | \|_____\| সেই পায়খানা কি করা হয়েছিল? [What was done to the feces?] | **হ্যাঁ [Yes].....1, না [No]......0**<br>1. যেখানে পায়খানা করেছিল সেখানেই ফেলে রাখা হয়েছিল [It is left there]**905.a** নং প্রশ্নে চলে যান (Skip to 906)<br>2. \|_____\| টয়লেট/পায়খানার ভিতরে ফেলা/ধোয়া হয়েছিল [Put/rinsed into toilet or latrine]<br>3. \|_____\| ড্রেনে/নর্দমার ভিতরে ফেলা/ধোয়া হয়েছিল [Put/rinsed into drain or ditch]<br>4. \|_____\|ঝোপ-ঝাড়ে/জঙ্গলে ফেলা হয়েছিল [Thrown into Bush/jungle<br>5. \|_____\| ময়লা আবর্জনার মধ্যে ফেলা হয়েছিল [Thrown into garbage]<br>6. \|_____\| নির্দিষ্ট গর্তে ফেলা হয়েছিল [Thrown into a specific pit for child's feces]<br>7. \|_____\| মাটির নীচে পুতে ফেলা হয়েছিল [Buried]<br>77. \|_____\| অন্যান্য (বর্ণনা লিখুন) [Other (specify)]....<br>99. \|_____\| জানি না [DK]**906** নং প্রশ্নে চলে যান (Skip to 906) |
| **C.905.** | 905. | আপনি কিভাবে পায়খানা পরিষ্কার করেন? উত্তর পড়ে শুনানো যাবে না \|হ্যাঁ হলে ১ বসান ।did you handle the feces? ☟Do Not Read Responses. Mark All that Apply (1 = Yes) | 1. \|_____\|নগ্ন/খালি হাতে Hands only (bare hands)<br>2. \|_____\|হাতে এবং কাপড়/পাতা/কাগজ Hands and cloth / paper / leaves<br>3. \|_____\| পায়খানা ফেলার জন্য কোন উপাদান ব্যবহার করে Scrap material to scoop fece<br>4. \|_____\|পটি Potty<br>5. \|_____\|স্থানীয় কৃষিকাজ-এর হাতিয়ার Local agricultural hoe/instrument<br>6. \|_____\|সেনি স্কুপ Sani-scoop<br>7. \|_____\|কোন কিছুই করা হয় না Did nothing<br>77 \|_____\|অন্যান্য (উলে- খ কর ন)Others (specify)____________<br>99 \|_____\|জানি না Don't know / not sure |
| C.905.b | 905.a | পায়খানা করার পর কতক্ষণ এই পায়খানা এখানে পরে ছিল? How long after defecation did you dispose of the feces? | \|____\|Hour ঘন্টা<br>\|____\|Minute মিনিট |
| **C.906.** | 906. | আপনার ৩ বছরের ছোট বাচ্চার পায়খানা কে কে ফেলে? (যতজন ফেলে সবাইকে কোড কর ন) [Who disposes your under 3 child's feces? | **হ্যাঁ [Yes].....1, না [No]......0**<br>1. \|_____\| মা [Mother]<br>2. \|_____\| বাবা [Father]<br>3. \|_____\| বোন [Sister]<br>4. \|_____\| ভাই [Brother]<br>5. \|_____\| খালা/মামী/চাচী/ফুফু/দাদী [Aunt/grandmother]<br>6. \|_____\| খালু/মামা/চাচা/ফুফা/দাদা [Uncle/grantfather]<br>7. \|_____\| কেউ পায়খানা ফেলে না [Nobody dispose]<br>77. \|_____\| অন্যান্য (নির্দিষ্ট করে লিখুন) Others (Specify) ........................<br>888. \|_____\|প্রযোজ্য নয় [Not applicable] |
| | 907. | ৩ বছরের ছোট বাচ্চার পায়খানা ফেলার স্থানটি রান্না ঘর থেকে কতটুকু দূরে? [How far away is this disposal site to the kitchen?] | \|_____\| কদম দূরে? [steps]<br>**888.** প্রযোজ্য নয় [Not applicable] |

| | | | |
|---|---|---|---|
| **C.907.** | 908. | \|_____\| **আপনার খানায় ৩-<৮ বছরের অন্য কোন শিশু আছে কি?** [Is there any other child between the age of 3-<8 years?] | 1 হ্যাঁ [Yes]<br>0. না [No].......... **913** নং প্রশ্নে চলে যান (skips to 913) |
| **C.908.** | 909. | আপনার খানায় ৩-<৮ বছরের যে শিশুটি আছে তার বয়স কত (মাসে)? (একাধিক শিশু থাকলে সবচেয়ে বড় বাচ্চার ক্ষেত্রে জিজ্ঞেস করুন) | ........................... মাস |
| **C.909.** | 910. | \|_____\| **আপনার শিশু (৩-৮ বছরের) সর্বশেষ কোথায় পায়খানা করেছে? (একাধিক শিশু থাকলে সবচেয়ে বড় বাচ্চার ক্ষেত্রে জিজ্ঞেস করুন)** [Where did your oldest child (aged 3-7) defecate the last time?] | 1. পটি (উঠানে) [Potty (in the courtyard)]<br>2. পটি (ঘরের ভিতরে) [Potty (inside the house)]<br>3. উঠানে (পটি ছাড়া) [Courtyard (without potty)]<br>4. ঘরের ভিতরে (পটি ছাড়া) [Inside the house (without potty)]<br>5. উঠানের বাহিরে খোলা জায়গাতে [Open space outside the front yard<br>6. ঝোপ-ঝাড়ে/জঙ্গলে Bush/jungle<br>7. পায়খানায়/টয়লেটে [In toilet] ...**913** নং প্রশ্নে চলে যান (Skip to 913)<br>8. কাথা/ন্যাপি/ডায়পার Katha/nappy/diaper<br>77. অন্যান্য (বর্ণনা লিখুন) [Other (Please describe)]....................<br>99. জানি না [Don't Know]............. **913** নং প্রশ্নে চলে যান (Skip to 913) |
| **C.910.** | 911. | \|_____\|**সেই পায়খানা কি করা হয়েছিল?** [What is done to the feces?] | **হ্যাঁ [Yes].....1, না [No]......0**<br>1. যেখানে পায়খানা করেছিল সেখানেই ফেলে রাখা হয়েছিল [It is left there]**912.a** নং প্রশ্নে চলে যান (Skip to 912.a)<br>2. \|_____\| টয়লেট/পায়খানার ভিতরে ফেলা/ধোয়া হয়েছিল [Put/rinsed into toilet or latrine]<br>3. \|_____\| ড্রেনে/নর্দমার ভিতরে ফেলা/ধোয়া হয়েছিল [Put/rinsed into drain or ditch]<br>4. \|_____\|ঝোপ-ঝাড়ে/জঙ্গলে ফেলা হয়েছিল [Thrown into Bush/jungle<br>5. \|_____\| ময়লা আবর্জনার মধ্যে ফেলা হয়েছিল [Thrown into garbage]<br>6. \|_____\| নির্দিষ্ট গর্তে ফেলা হয়েছিল [Thrown into a specific pit for child's feces]<br>7. \|_____\| মাটির নীচে পুতে ফেলা হয়েছিল [Buried]<br>77. \|_____\| অন্যান্য (বর্ণনা লিখুন) [Other (specify)]................<br>99. \|_____\| জানি না [DK]**913** নং প্রশ্নে চলে যান (Skip to 913) |
| **C.910.** | 912. | **আপনি কিভাবে পায়খানা হ্যান্ডলিং (ব্যবস্থাপনা )করেন?** How did you handle the feces? ☞Do Not Read Responses. Mark All that Apply (1 = Yes) | ☐ 1 নগ্ন/খালি হাতে Hands only (bare hands)<br>☐ 2 হাতে এবং কাপড়/পাতা/কাগজ Hands and cloth / paper / leaves<br>☐ 3 ফেলানো সামগ্রী মাটিতে পুঁতে Scrap material to scoop feces<br>☐ 4 পটি Potty<br>☐ 5 স্থানীয় কৃষিকাজ-এর হাতিয়ার Local agricultural hoe/instrument<br>☐ 6 সেনি স্কুপ Sani-scoop<br>☐ 7 কোন কিছুই করা হয় না Did nothing<br>☐ 77 অন্যান্য (উল্লেখ করুন) Others (specify)____________ |

| | | | |
|---|---|---|---|
| | | | ☐ 99 জানি না Don't know / not sure |
| C.910.b | 912.a | পায়খানা করার পর কতক্ষণ এই পায়খানা এখানে পরে ছিল? How long after defecation did you dispose of the feces? | \|____\|Hour ঘন্টা<br>\|____\|Minute মিনিট |

| | | | |
|---|---|---|---|
| **Child Potty Use**<br>**( শিশুর পটি ব্যবহার সর্ম্পকীত তথ্য)** | | | |
| Administer to: All study households ( সকল খানায় হবে) | | | |
| C.815 | 913. | Does your household have a potty that children use for defecation? আপনার খানাতে কি শিশুদের পায়খানা করার পটি আছে? | 1 Yes হ্যাঁ<br>2 No না **(Skip to 919)** (919 নং প্রশ্নে যান)<br>99 Don't know / not sure জানি না **(Skip to 919)** (919 নং প্রশ্নে যান) |
| C.816 | 914. | In the last week, how often did your child use the potty? গত এক সপ্তাহের মধ্যে, শিশুটি কতবার পটি ব্যবহার করছে?<br>[ *If multiple children use the potty, ask about the youngest child* ][যদি একাধিক শিশু ব্যবহার করে থাকে তবে যে সবচেয়ে বেশী ব্যবহার করে সেই শিশুর জন্য প্রশ্ন করুন] | 1 Every time প্রতিবার<br>2 More than half of all defecation events, but not every time যতবার পায়খানা করেছে তার ৫০ ভাগ বা তার বেশী কিন্তু প্রতিবার নয়<br>3 যতবার পায়খানা করেছে তার ৫০ ভাগের কম কিন্তু মাঝেমাঝে Less than half of all defecation events<br>4 আগে করেছি কিন্তু এখন আর করি না Used to use it, but no longer use it **(Skip to 919)** (919 নং প্রশ্নে যান)<br>5 কখনই না Never **(Skip to 919)** (919 নং প্রশ্নে যান)<br>99 জানি না Don't know / not sure **(Skip to 919)** (919 নং প্রশ্নে যান) |
| C.817 | 915. | What is the age of the child (or children) who is using the potty? যে শিশুটি (বা শিশুরা) পটি ব্যবহার করছে তার বয়স কত? | <u>Mark All that Apply.</u> Yes = 1 হ্যাঁ = 1, না = 0<br>1 ☐ < 1 year ১ বছরের কম<br>2 ☐ 1- < 3 years ১-<৩ বছর<br>3 ☐ 3- < 5 years ৩-<৫ বছর<br>4 ☐ >= 5 years >=৫ বছর |
| C.818 | 916. | Ask the respondent to identify the place where the child typically uses the potty. Then ask her to retrieve the potty<br>Could I please see the potty?<br>👁 Observation: *Record how long it takes to produce the potty.*<br>আমি কি পটিটি দেখতে পারি<br>👁 আমি কি পটিটি দেখতে পারি (পটিটি আনতে কতটুকু সময় লেগেছে তা সেকেন্ডে গণনা করুন) | ☐☐:☐☐MINUTES : SECONDS (99:99 if could not measure) মিনিট : সেকেন্ড (পটি দেখাতে না পারলে 999 কোড করুন) |
| C.819 | 917. | 👁 Observation: Potty condition.<br>পটিটির অবস্থা কি তা পর্যবেক্ষণ করুন (একাধিক উত্তর হতে পারে) | <u>Mark All that Apply.</u> Yes = 1 হ্যাঁ = 1, না = 0<br>1.Easily accessible when needed by the child প্রয়োজনের সময় বাচ্চা যেন |

| | | | |
|---|---|---|---|
| | | | পায় এমন কাছাকাছি জায়গাতে রয়েছে<br>2.Easily accessible when needed by the mother প্রয়োজনের সময় মা যেন পায় এমন কাছাকাছি জায়গাতে রয়েছে<br>3.Visible signs of feces inside / on the potty / removable pot ভিতরে/ পটিতে পায়খানার চিহ্ন আছে<br>4.Potty was covered with the lid পটি লিড দিয়ে ঢাকা ছিল<br>5. Potty was covered with anything other than the lid পটি লিড ছাড়া অন্য কিছু দিয়ে ঢাকা ছিল<br>6.Dry শুকনা<br>7.Broken/or missing parts(tray) so that it is unusable ভেঙ্গে গেছে তাই ইহা ব্যবহার করা সম্ভব না<br>8.Covered in dust / signs of non-use পটি ধুলায় ঢাকা/ ব্যবহার না করার চিহ্ন রয়েছে<br>9.Cannot produce a potty পটি দেখাতে পারে নাই |
| C.820 | 918. | ☐Where do you usually dispose of feces from the potty? সাধারনতঃ এই পটি থেকে আপনি পায়খানা কোথায় ফেলেন? | 1 Latrine পায়খানায়/টয়লেটে<br>2 Open Pit / separate pit for child or animal feces খোলা গর্তে / শিশুর বা পশুপাখির পায়খানা ফেলার গর্তে<br>3 Bury it / Covered Pit মাটির নীচে পুতে ফেলা হয়েছিল/ ঢাকা গর্তে<br>4 Undefined open site near the compound (including open garbage disposal sites / dumps) উঠানের কাছে খোলা জায়গা<br>5 Bush / forest / field ঝোপ-ঝাড়ে/জঙ্গলে/মাঠে<br>6 Nearby water (pond, canal, river) পানির উৎসের কাছে (পুকুর/সেচ নালা/ নদী)<br>77 Other (specify) অন্যান্য (লিখুন)_______________ |

**Sani Scoop Use**
**(সেনিস্কুপ ব্যবহার)**

Administer to: All households ( সকল খানায় হবে)

| | | | |
|---|---|---|---|
| C.821 | 919. | ☐Does your household have a dedicated tool [sani scoop] to clean up feces around your household? আপনার খানার চারপাশের পায়খানা পরিষ্কার করার জন্য আপনার খানায় কোন নির্দিষ্ট কিছু আছে কি? | 1 Yes হ্যাঁ<br>2 No না (1001 নং প্রশ্নে যান) **(Skip to 1001)**<br>99 Don't know / not sure জানি না (1001 নং প্রশ্নে যান) **(Skip to 1001)** |
| C.822 | 920. | How often do you use the [sani scoop]? আপনি কখন কখন পায়খানা পরিষ্কার করার জন্য কোদাল/সেনিস্কুপ ব্যবহার করেন? | 1 Multiple times per day প্রতিদিন কয়েকবার<br>2 Once per day প্রতিদিন একবার<br>3 A few times each week প্রতি সপ্তাহে কয়েকবার<br>4 Less than once per week সপ্তাহে একবারের কম<br>5 Used to use it, but no longer use it প্রথমদিকে ব্যবহার করেছে কিন্তু এখন আর করে না (1001 নং প্রশ্নে যান) **(Skip to 1001)**<br>6 Never কখনই না (1001 নং প্রশ্নে যান) **(Skip to 1001)** |
| C.823 | 921. | What do you use the [sani scoop] for? ☟Do Not Read Responses. আপনি কি কি কাজে কোদাল/সেনিস্কুপ ব্যবহার করেন? (একাধিক উত্তর হতে পারে) ☟ উত্তর পড়ে শোনাবেন না | Mark All that Apply. Yes = 1 হ্যাঁ = 1, না = 0<br>1 ☐ Clean up animal feces জীবজন্তুর পায়খানা ফেলার কাজে<br>2 ☐ Clean up child feces বাচ্চার পায়খানা ফেলার কাজে<br>3 ☐ Clean up garbage ময়লা আবর্জনা ফেলার কাজে<br>4 ☐ Take the scoop to the field (for work) কৃষিকাজে/মাঠে ইহা নিয়ে যাই |

| | | | |
|---|---|---|---|
| | | | 5 ☐ Digging / gardening গর্ত তৈরীর কাজে<br>77 ☐ Other (specify) অন্যান্য (বর্ণনা লিখুন)_______ |
| C.824 | 922. | Could I please see the [sani scoop]?<br>👁 Observation: Sani scoop condition.<br>আমি কি কোদাল/সেনিস্কুপটি দেখতে পারি? (একাধিক উত্তর হতে পারে)<br>👁 পর্যবেক্ষণ: সেনিস্কুপটির অবস্থা পর্যবেক্ষণ করুন | Mark All that Apply. Yes = 1 হ্যাঁ = 1, না = 0<br>1 ☐ Visible signs of feces on the sani scoop কোদাল/সেনিস্কুপে পায়খানার চিহ্ন আছে<br>2 ☐ Dry শুকনা<br>3 ☐ Broken and needs repair ভেঙ্গে গেছে এবং মেরামত করা দরকার<br>4 ☐ Easily accessible when needed by an adult প্রয়োজনের সময় বড়রা যেন পায় এমন কাছাকাছি জায়গাতে রয়েছে<br>5 ☐ Signs that the sani scoop is not used কোদাল/সেনিস্কুপ ব্যবহার না করার চিহ্ন রয়েছে<br>6 ☐ Cannot produce a sani scoop কোদাল/সেনিস্কুপ দেখাতে পারে নাই |
| C.825 | 923. | I'd like to ask a few more questions about the types of feces you pick up with the [sani scoop] and how you dispose of the different types of feces.<br>[Ask about each type of feces separately.].<br>কোদাল/সেনিস্কুপ দিয়ে আপনি যেসব পায়খানা ফেলেন এবং কিভাবে ফেলেন সেসম্পর্কে জানার জন্য এখন আমি আপনাকে আরো কিছু প্রশ্ন করতে চাই।<br>(প্রতিটি পায়খানার ধরনের কথা উল্লেখ করে কোড করুন)<br>(শিশু/পশুপাখির পায়খানা ফেলার কাজে ব্যবহার না করলে প্রযোজ্য নয় ঘরটিতে টিক দিন) | Where do you dispose the feces?<br>পায়খানা কোথায় ফেলা হয়েছিল?<br>✋ Do Not Read Responses. উত্তর পড়ে শোনাবেন না<br>Mark All that Apply (একাধিক উত্তর হতে পারে) 1 = হ্যাঁ Yes, না = 0,<br>88 Not applicable প্রযোজ্য নয় |

| | A | B | C | D | E | F |
|---|---|---|---|---|---|---|
| Feces পায়খানা | Use Sani Scoop for disposal? পায়খানা ফেলার জন্য কোদাল/সেনিস্কুপ ব্যবহার করে? | Latrine পায়খানায় | Open Pit খোলা গর্তে | Bush/ Farm ঝোপে ঝাড়ে/খোলা জায়গায় | Surface Water পানিতে | Dig Hole and cover গর্ত করা হয় এবং ঢেকে রাখা হয় |
| 1 Child (<8 years) শিশুর (<৮ বছর) | ☐ | | | | | |
| 2 Cow গোবর | ☐ | | | | | |
| 3 Poultry / | ☐ | | | | | |

| | | | | |
|---|---|---|---|---|
| | | | pigeons হাঁস-মুরগীর | |
| | | | 4 Goat ছাগলের | ☐ |
| | | | 5 Pig শুকরের | ☐ |
| | | | 6 Dog or cat কুকুর বা বিড়ালের | ☐ |

**Wash Benefit Module 10 Water Access**

| C.10 WATER TREATMENT, STORAGE, AND QUALITY (All households) | | | | | |
|---|---|---|---|---|---|
| SAY | Thank you very much. Now, I would like to ask you some questions about the water you and your family drink at home. | | | | |
| C.1001 | 1001. | আপনি কিভাবে খাবার পানি সংরক্ষন করেন ? [How do you store drinking water?<br><br>👁 **প্রশ্ন করুন এবং সম্ভব হলে পর্যবেক্ষন করুন** ASK THE QESTION AND OBSERVE IF POSSIBLE | [1] প্লাষ্টিক অথবা ধাতব পাত্রে IN PLASTIC OR METAL CONTAINERS<br>[2] মাটির পাত্রে IN CLAY POTS<br>[3] ছাদে পানির ট্যাংক ROOF TANK OR CISTERN<br>[4] পানি সংরক্ষণ করা হয় না DO NOT STORE WATER | | |
| C.1002 | 1002. | এই খানায় ০-<৩৬ মাসের কোন শিশু আছে কি যে পানি পান করে ?Is there a child 0-<36 months in this household who drinks water? | [1] হ্যাঁ YES<br>[2] না NO<br>[88] Not applicable প্রযোজ্য নয় | | |
| C.1003 | 1003. | **IF YES → প্রশ্ন করুন এবং পর্যবেক্ষন করুনঃ যদি আপনার শিশু (<৩ বছর) এখন পানি খেতে চাইত তাহলে আপনি কিভাবে তাকে পানি দিতেন তা আমাকে দেখান** ASK: If <u>your child 0-3 years</u> wanted a drink of water right now, could you show me how you would give it to him/her?<br>**IF NO →** ASK: **যদি আপনি এখন পানি খেতে চাইতেন তাহলে আপনি কিভাবে পানি নিতেন তা আমাকে দেখান** If <u>you</u> wanted a drink of water right now, could you show me how you would get it?<br><br>উত্তরদাতার পানি দেওয়ার প্রক্রিয়াকে পর্যবেক্ষণ/অনুসরণ করুন । যদি অন্য ঘরে পানি সংরক্ষিত এবং সেখানে প্রবেশাধিকার থাকে তাহলে প্রক্রিয়াকে পর্যবেক্ষণ/অনুসরণ করুন । অন্যথায় ৯৯ কে গোলাকার করতে হবে। ATTEMPT TO FOLLOW RESPONDENT WHEN SHE GOES TO RETRIEVE THE WATER. IF WATER IS STORED IN ANOTHER ROOM, ASK HER IF IT IS OKAY TO ENTER THAT ROOM. CIRCLE "99" IF YOU CANNOT OBSERVE. | | | |
| | | 👁 উত্তরদাতা পানি নেওয়ার আগে কি করেছিল ? WHAT DID RESPONDENT DO <u>BEFORE TAKING THE WATER</u>? | **হ্যাঁ** YES | **না** NO | **জানি না** D/K |
| 2 | 2 | **গ-াস বা পাত্রটিতে পানি ঢালার পূর্বে গ-াস/পাত্রটি খাবার পানি দিয়ে ধুয়েছিল** RINSED GLASS/CONTAINER WITH DRINKING WATER BEFORE FILLING | [1] | [2] | [99] |
| 3 | 3 | **পানি ঢালার পূর্বে হাত ধুয়েছিল** WASHED HANDS WITH WATER BEFORE DRINKING WATER WAS OBTAINED | [1] | [2] | [99] |
| 4 | 4 | **পানি ঢালার পূর্বে সাবান দিয়ে হাত ধুয়েছিল** WASHED HANDS WITH SOAP BEFORE DRINKING WATER OBTAINED | [1] | [2] | [99] |
| | | 👁 উত্তরদাতা কোথা থেকে পানি সংগ্রহ করেছিল? <u>FROM WHERE</u> DID THE RESPONDENT TAKE THE WATER? | **হ্যাঁ** YES | **না** NO | **জানি না** D/K |
| 5 | 5 | সরাসরি পানির উৎস থেকে BROUGHT DIRECTLY FROM WATER SOURCE | [1] | [2] | [99] |

| | | | | | |
|---|---|---|---|---|---|
| 6 | 6 | সরাসরি খোলা পানির পাত্র থেকে BROUGHT IRECTLY FROM uncovered STORAGE CONTAINER | [1] | [2] | [99] |
| 7 | 7 | সরাসরি পানির ফ্লিটার থেকে BROUGHT DIRECTLY FROM WATER FILTER | [1] | [2] | [99] |
| | 8 | সরাসরি ঢেকে রাখা পানির পাত্র থেকে STORED WATER WAS COVERED | [1] | [2] | [99] |
| | | 👁 উত্তরদাতা কিভাবে গ-াসে/কাপে/পাত্রে পানি দিয়েছিল? <u>HOW DID SHE GET THE WATER</u> INTO THE CUP? | হ্যাঁ YES | না NO | জানি না D/K |
| 9 | 9 | গ-াস বা পাত্রটিতে ঢালা পানির ভিতরে তার হাত বা হাতের আঙ্গুল লেগেছিল কি? HANDS TOUCHED / CONTACTED THE DRINKING WATER | [1] | [2] | [9] |
| 10 | 10 | গ-াস বা পাত্রটিতে পানি ভরার সময় তা পানির পাত্রের ভিতর ডুবিয়ে পানি ভরা হয়েছিল CONTAINER/GLASS DIPPED INTO WATER CONTAINER | [1] | [2] | [99] |
| 11 | 11 | গ-াসে পানি ভরার জন্য লম্বা হাতলযুক্ত কোন চামচ/মগ ব্যবহার করা হয়েছিল LADLE USED TO OBTAIN WATE | [1] | [2] | [99] |
| 12 | 12 | পানির পাত্র থেকে সরাসরি পানি ঢেলেছিল WATER POURED FROM CONTAINER | [1] | [2] | [99] |
| 13 | 13 | সরাসরি টিউবওয়েল/ পানির উৎস থেকে এনেছে WATER POURED FROM TAP | [1] | [2] | [99] |

| | | | | |
|---|---|---|---|---|
| C.1004 | 1004. | আপনি/এ খানার অন্যকেউ কতক্ষণ আগে এই সংরক্ষণকৃত পানি সংগ্রহ করেছেন? How long ago did you or somebody in your home collect this water? ( 99 = Don't know জানি না; 87 = No Water কোন পানি নাই, 88 Not applicable প্রযোজ্য নয় ) If 88, skip to 1007 | A.\|__\|__\| ঘন্টা HOURS<br>B.\|__\|__\| দিন DAYS | |
| C.1005 | 1005. | এই খাবার পানি নিরাপদ করার জন্য কোন কিছু করেছেন কি? Have you done anything to make this water less cloudy or safer to drink? | [1] হ্যাঁYES<br>[2] না NO<br>[99] জানি না DON'T KNOW/NOT SURE | <br>→ **SKIP to 1007**<br>→ **SKIP to 1007** |
| C.1006 | 1006. | কোন পদ্ধতি কি আপনি ব্যবহার করেন? What method(s) did you use? উত্তর পড়ে শুনাবেন না \|উত্তরদাতাকে উত্তর বলতে সাহায্য করা যাবে তার উত্তর দেয়া শেষ হলে।DO NOT READ RESPONSES. PROBE UNTIL RESPONDENT IS FINISHED. CIRCLE ALL THAT APPLY. | | |
| 1 | 1 | আকুয়্যাট্যাবস_Aquatabs | [1] হ্যাঁYes [0] না No | |
| 2 | 2 | ক্লোরিন (তরল) মিশিয়ে নেই (ওয়াটার গার্ড নয়) Waterguard / bottled chlorine | [[1] হ্যাঁYes [0] না No | |
| 3 | 3 | পানি ফুটিয়ে নেই Boil | [1] হ্যাঁYes [0] না No | |
| 4 | 4 | কাপড়ে ছেকে নেই Strain it through clothor other material | [1] হ্যাঁYes [0] না No | |
| 5 | 5 | পানির ফিল্টার ব্যবহার করে (সিরামিক, বালু ইত্যাদি) Water filter [ceramic, sand, composite] | [1] হ্যাঁ Yes [0] না No | |
| 6 | 6 | সূর্যের আলোতে জীবানুমুক্তকরণ Solar disinfection (SODIS) | [1] হ্যাঁ Yes [0] না No | |
| 7 | 7 | থিতিয়ে নেই Let it stand and settle | [1] হ্যাঁ Yes [0] না No | |
| 8 | 8 | স্যান্ড ফিল্টার এর মাধ্যমে বিশুদ্ধ করে নেই Biosand filter | [1] হ্যাঁ Yes [0] না No | |

| | | | | |
|---|---|---|---|---|
| 9 | 9 | ফিটকিরি যোগ করে Coagulant (alum) | [1] হ্যাঁ Yes [0] না No | |
| 10 | 10 | পার PUR (flocculant + disinfectant) | [1] হ্যাঁ Yes [0] না No | |
| C.1007 | 1007. | আপনি কখনও আপনার খাবার পানি পরিশোধন করার জন্য কিছু করেছিলেন কি? Do you ever treat your drinking water or do anything to make it less cloudy? | [1] হ্যা YES<br>[0] না NO | → **1010 নং এ চলে যান SKIP TO 1010** |
| C.1008 | 1008. | **সর্বশেষ কখন আপনি** আপনার খাবার পানি পরিশোধন করেছিলেন ? When was the last time you treated your water or did anything to make it less cloudy?<br><br>উত্তর পড়ে শুনানো যাবে না DO NOT READ RESPONSES | [1] আজ TODAY<br>[2] গতকালYESTERDAY<br>[3] গত সপ্তাহের মধ্যে WITHIN THE PAST WEEK<br>[4] গত দুই সপ্তাহের মধ্যে WITHIN THE PAST 2 WEEKS<br>[5] গত মাসের মধ্যে WITHIN THE PAST MONTH<br>[6] গত বছরের মধ্যে WITHIN THE PAST YEAR<br>[99] জানি না/নিশ্চিত না DON'T KNOW/NOT SURE | |
| C.1009 | 1009. | কোন কোন উপায়ে আপনি পানি পরিশোধন করে থাকেন ? What are all the ways you treat your drinking water?<br>উত্তর পড়ে শুনানো যাবে না । উত্তর দেয়া শেষ হলে উত্তর পেতে সাহায্য করা যেতে পারে ।DO NOT READ RESPONSES. PROBE UNTIL RESPONDENT IS FINISHED. CIRCLE ALL THAT APPLY. | | |
| 1 | 1 | আকুয়াট্যাবস_Aquatabs | [1] হ্যাঁYes [0] না No | |
| 2 | 2 | ক্লোরিন (তরল) মিশিয়ে নেই (ওয়াটার গার্ড নয়) Waterguard / bottled chlorine | [[1] হ্যাঁYes [0] না No | |
| 3 | 3 | পানি ফুটিয়ে নেই Boil | [1] হ্যাঁYes [0] না No | |
| 4 | 4 | কাপড়ে ছেকে নেই Strain it through clothor other material | [1] হ্যাঁYes [0] না No | |
| 5 | 5 | পানির ফিল্টার ব্যবহার করে (সিরামিক, বালু ইত্যাদি) Water filter [ceramic, sand, composite] | [1] হ্যাঁYes [0] না No | |
| 6 | 6 | সূর্যের আলোতে জীবানুমুক্তকরণ Solar disinfection (SODIS) | [1] হ্যাঁYes [0] না No | |
| 7 | 7 | থিতিয়ে নেই Let it stand and settle | [1] হ্যাঁYes [0] না No | |
| 8 | 8 | স্যান্ড ফিল্টার এর মাধ্যমে বিশুদ্ধ করে নেই Biosand filter | [1] হ্যাঁYes [0] না No | |
| 9 | 9 | ফিটকিরি যোগ করে Coagulant (alum) | [1] হ্যাঁYes [0] না No | |
| 10 | 10 | PUR (flocculant + disinfectant) | [1] হ্যাঁYes [0] না No | |

| পানের পানির উৎস **DRINKING WATER SOURCES (ALL HOUSEHOLDS)** | | | |
|---|---|---|---|
| উত্তর দাতাকে তার **পানির প্রধান এবং দ্বিতীয় উৎস বিষয়ে প্রশ্ন কর**ুন ASK THE FOLLOWING QUESTIONS ABOUT THE RESPONDENT'S PRIMARY AND SECONDARY WATER SOURCES. | | | |
| C.1016 | 1010. | আপনার খানায় ব্যবহৃত পানির প্রধান উৎস কি ? What type of water source does your household collect most of the water that you use from? | 1 অগভীর টিউবওয়েল Shallow tubewell<br>2 গভীর টিউবওয়েল Deep tubewell<br>3 ঘরের ভিতর ট্যাপ বা পাইপের পানি Piped into home |

| | | | |
|---|---|---|---|
| | | | 4 উঠানে ট্যাপ বা পাইপের পানি Piped into yard<br>5 গভীর পাতকুয়া (উঠানে)Borewell in yard<br>6 পাবলিক ট্যাপ Public tap<br>7 পাবলিক গভীর পাতকুয়া Public borewell<br>8 অগভীর পাকা পাতকুয়া Shallow well with concrete reinforcement<br>9 অগভীর কাঁচা পাতকুয়া Shallow well (no concrete)<br>10 নদী/বাধ/লেক/পুকুর/সেচ নালা থেকে সরাসরি সংগৃহীত পানি River, lake, stream<br>11 অন্যান্য (লিখুন) Other: _____________ |
| C.1017 | 1011. | আপনি আমাকে বলবেন আপনার বাড়ী থেকে প্রধান পানির উৎসে একবার যেতে কতক্ষণ সময় লাগে? Can you tell me how long it takes you to walk one-way to this PRIMARY water source from your home? | \|__\|ঘন্টা Hours<br>\|__\|মিনিট Min |
| C.1018 | 1012. | আপনি আমাকে বলবেন কোন কোন কাজের জন্য এই উৎসের পানি ব্যবহার করেন Can you please tell me whether you use water from this source for… | |
| 1 | 1 | পানি পান Drinking | [1] হ্যাঁYes [0] না No |
| 2 | 2 | রান্না-বান্না/বাসন ধোয়া Cooking/ washing dishes | 1] হ্যাঁYes [0] না No |
| 3 | 3 | কাপড় ধোয়া Laundry | 1] হ্যাঁYes [0] না No |
| 4 | 4 | গোসল/হাতধোয়া Bathing/hand washing | 1] হ্যাঁYes [0] না No |
| 5 | 5 | কৃষিকাজ/বাগান Irrigate crop land or garden | 1] হ্যাঁYes [0] না No |
| 6 | 6 | পশু-পালন Water for livestock | 1] হ্যাঁYes [0] না No |
| C.1019 | 1013. | আপনার খানাকে এই পানির উৎস ব্যবহারের জন্য কি কোন প্রকার টাকা দিতে হয়? your household pay any money to use this source? | ] হ্যাঁYes [0] না No |
| C.1020 | 1014. | আপনার খানায় পানি ব্যবহারের জন্য কি ২য় কোন উৎস ব্যবহার করতে হয়? যদি হয় তাহলে কি ধরনের ? Does your household currently use a secondary water source? If so, what type is it? | 1. অগভীর টিউবওয়েল Shallow tubewell<br>2. গভীর টিউবওয়েল Deep tubewell<br>3. ঘরের ভিতর ট্যাপ বা পাইপের পানি Piped into home<br>4. উঠানে ট্যাপ বা পাইপের পানি Piped into yard<br>5. গভীর পাতকুয়া (উঠানে)Borewell in yard<br>6. পাবলিক ট্যাপ Public tap<br>7. পাবলিক গভীর পাতকুয়া Public borewell<br>8. অগভীর পাকা পাতকুয়া Shallow well with concrete reinforcement<br>9. অগভীর কাঁচা পাতকুয়া Shallow well (no concrete)<br>10. নদী/বাধ/লেক/পুকুর/সেচ নালা থেকে সরাসরি সংগৃহীত পানি River, lake, stream<br>11. অন্যান্য (লিখুন) Other: _____________<br>88] ২য় কোন পানির উৎস ব্যবহার করতে হয় না Don't use any secondary water source (Skip to 1017.a) |
| C1021 | 1015. | আপনি আমাকে বলবেন আপনার বাড়ী থেকে ২য় পানির উৎসে একবার যেতে কতক্ষণ সময় লাগে? Can you | \|__\|ঘন্টা Hours<br>\|__\|মিনিট Min |

| | | | |
|---|---|---|---|
| | | tell me how long it takes you to walk one-way to this SECONDARY water source from your home? | |
| C1022 | 1016. | আপনি আমাকে বলবেন কোন কোন কাজের জন্য এই উৎসের পানি ব্যবহার করেন Can you please tell me whether you use water from this source for… | |
| 1 | 1 | পানি পান Drinking | [1] হ্যাঁYes [0] না No |
| 2 | 2 | রান্না-বান্না/বাসন ধোয়া Cooking/ washing dishes | 1] হ্যাঁYes [0] না No |
| 3 | 3 | কাপড় ধোয়া Laundry | 1] হ্যাঁYes [0] না No |
| 4 | 4 | গোসল/হাতধোয়া Bathing/hand washing | 1] হ্যাঁYes [0] না No |
| 5 | 5 | কৃষিকাজ/বাগান Irrigate crop land or garden | 1] হ্যাঁYes [0] না No |
| 6 | 6 | পশু-পালন Water for livestock | 1] হ্যাঁYes [0] না No |
| C1023 | 1017. | আপনার খানাকে এই পানির উৎস ব্যবহারের জন্য কি কোন প্রকার টাকা দিতে হয়? your household pay any money to use this source? | 1] হ্যাঁYes [0] না No |
| | 1017.a | আপনার খানার খাবার পানির প্রধান উৎস কি? What is your main drinking water source? | 1. অগভীর টিউবওয়েল Shallow tubewell<br>2. গভীর টিউবওয়েল Deep tubewell<br>3. ঘরের ভিতর ট্যাপ বা পাইপের পানি Piped into home<br>4. উঠানে ট্যাপ বা পাইপের পানি Piped into yard<br>5. গভীর পাতকুয়া (উঠানে)Borewell in yard<br>6. পাবলিক ট্যাপ Public tap<br>7. পাবলিক গভীর পাতকুয়া Public borewell<br>8. অগভীর পাকা পাতকুয়া Shallow well with concrete reinforcement<br>9. অগভীর কাঁচা পাতকুয়া Shallow well (no concrete)<br>10. নদী/বাধ/লেক/পুকুর/সেচ নালা থেকে সরাসরি সংগৃহীত পানি River, lake, stream<br>11. অন্যান্য (লিখুন) Other: _____________<br>88] Not applicable |
| C1024 | 1018. | আপনার খানা কি কখনও বৃষ্টির পানি সংগ্রহ করত ? Does your household collect rainwater? | 1] হ্যাঁYes [0] না No |
| C1025 | 1019. | আপনার খানা কি বর্তমানে বৃষ্টির পানি সংগ্রহ করে ? Is your household currently collecting rainwater? | 1] হ্যাঁYes [0] না No if no, skip to 1301 |
| C1026 | 1020. | আপনি আমাকে বলবেন কি আপনার খানা কি কি কাজে বৃষ্টির পানি ব্যবহার করে ? Can you please tell me whether you use rainwater for | |
| 1 | 1 | পানি পান Drinking | [1] হ্যাঁYes [0] না No |
| 2 | 2 | রান্না-বান্না/বাসন ধোয়া Cooking/ washing dishes | 1] হ্যাঁYes [0] না No |

| | | | |
|---|---|---|---|
| 3 | 3 | কাপড় ধোয়া Laundry | 1] হ্যাঁYes [0] না No |
| 4 | 4 | গোসল/হাতধোয়া Bathing/hand washing | 1] হ্যাঁYes [0] না No |
| 5 | 5 | কৃষিকাজ/বাগান Irrigate crop land or garden | 1] হ্যাঁYes [0] না No |
| 6 | 6 | পশু-পালন Water for livestock | 1] হ্যাঁYes [0] না No |
| **Skip to 1301** | | | |
| ~~C.1027.~~ | ~~1021.~~ | ~~আপনি ভেবে বলবেন কি আপনার পরিবার মোট সকল উৎস থেকে দৈনিক আনুমানিক ২০ লিটার পাত্রের কতটি পাত্র পানি পায়? ***উত্তরদাতাকে উত্তর দিতে অনুপ্রেরনা দেন*** জানা না থাকলে ৯৯৯ বসান(দূরত্ব আধা কিমি হলে ০.৫ লিখতে হবে)। Thinking about ALL the water that your household obtains from ALL sources, approximately how many 20-litre containers of water would you say that your household uses IN TOTAL EACH DAY?~~<br>~~***Encourage respondent to estimate. "Don't know"=999. Use decimals as needed (e.g. one half=0.5)~~ | ~~☐ দৈনিক ১০ লিটার পাত্র 10-liter containers per day~~<br>~~☐ দৈনিক ২০ লিটার পাত্র 20-liter containers per day~~ |
| C.1010. | 1022. | আপনার খানায় কত ঘন ঘন খাবার পানিতে আকুয়াট্যাবস মেশানো হয়?<br>How frequently do you treat your water using [Aquatabs]? | Do Not Read Responses.<br>1 যতবার পানি সংগ্রহ করা হয়/সবসময় Every time they collect water<br>2 মাঝেমাঝে Sometimes / occasionally<br>3 প্রথমদিকে দিয়েছিলাম কিন্তু এখন আর দেই না Treated water in the beginning [of the program] but not any more<br>4 আকুয়াট্যাবস দিয়ে কখনও পানি পরিশোধন করি নাই Never treated water with [Aquatabs] **(Skip to 1026)**<br>99 জানি না Don't know / not sure |
| C.1011. | 1023. | আপনার খানায় সংরক্ষনকৃত খাবার পানি কি কুয়াট্যাবস দ্বারা পরিশোধন করা হয়েছে?<br>Is the drinking water stored in your household today treated with [Aquatabs]? | 1 হ্যাঁ, সবটুকু পানি Yes, all of it<br>2 হ্যাঁ, কিছু পানি Yes, some of it<br>3 না Not treated<br>4 খানায় কোন পানি নাই No water in the house<br>99 জানি না Don't know |
| C.1012. | 1024. | পর্যবেক্ষন করুন: আকুয়াট্যাবস দ্বারা পরিশোধিত পানির পাত্র দেখাতে বলুন: পানির পাত্র ঢাকা ছিল কী? Ask to see the treated water. Observe: Is the water covered? | 1 হ্যাঁ Yes<br>2 না No<br>3 খানায় পাত্রে কোন পানি নাই No water stored in the house<br>88 প্রযোজ্য নয়/প্রত্যাখ্যান Not applicable / refused |
| C.1013. | 1025. | পানি পরিশোধন করার জন্য কত সময় আগে আকুয়াট্যাবস ব্যবহার করা হয়েছিল?<br>Approximately how long ago did you treat the | ঘিন্টা HH: মনিট MM ☐☐:☐☐ ( 9 9 : 9 9 Don't know ) |

| | | |
|---|---|---|
| | | water with [Aquatabs]? |

ক্লোরিন অবশেষ পরীক্ষা **Residual Chlorine Test**

সকল খানার জন্য প্রযোজ্য Administer to: All study households

*Ask to collect a water sample from the source identified in C.1003. Explain that some (but not all) of the samples may be tested to see if there is any chlorine in the water.*

| | | | |
|---|---|---|---|
| **C.1014.** | 1026. | ক্লোরিণের জন্য আমি আপনার খাবার পানি পরীক্ষা করতে পারি কী?<br>May I collect a small sample of your drinking water? | 1 হ্যাঁ Yes<br>2 না/প্রত্যাখান No / refused<br>3 খানায় কোন পানি নাই No drinking water available to test |
| **C.1015.** | 1027. | ক্লোরিন অবশেষ পরীক্ষার জন্য সংরক্ষনকৃত পানি যা নির্দিষ্ট বাচ্চা খায় তার নমুণা সংগ্রহ করুন \| ভিন্ন ভিন্ন অবস্থানেই খানার সংরক্ষিত খাবার পানির ক্লারিন অবশেষ পরীক্ষা করুন । শুধুমাত্র সেই খানার পানি পরীক্ষা করতে হবে যারা বলবে সংরক্ষিত খাবার পানিতে ক্লারিন মিশিনো হয়েছে(C.1009 = 1, 2, or 11)। যদি এই খানায় সংরক্ষিত খাবার পানিতে ক্লারিন না মেশানো হয়, তাহলে পানি পরীক্ষা থেকে বিরত থাকুন এবং ৮৮ কোড করুন।<br>Collect a small water sample from stored drinking water for target children.<br>Test for free residual chlorinein a discrete location after you leave the household.<br>Only test samples collect from households who report to have treated their water with some form of chlorine (C.1009 = 1, 2, or 11).If the household did not report treating the water with chlorine, discard the sample without testing it and record 8 88 in the field below. | ৮৮৮ ক্লোরিন মেশানো হয়েছে কিনা এ বিষয়ে রিপোর্ট দিতে পারে নি<br>8 88 Did not report that the water was treated with chlorine<br>**999** পরীক্ষা করা হয় নাই<br>9 9 9 Could not test<br>ক্লোরিন অবশেষের পরিমান Level of Free Residual Chlorine: ☐☐.☐ মি.গ্রাম/লিটার mg / L |

| | | | | |
|---|---|---|---|---|
| | | খানায় অনুজীবি সংক্রান্ড় নমূনা (খানা দৈবচয়িতভাবে নির্বাচিত হবে)**HOUSEHOLD MICROBIOLOGICAL SAMPLE (ADMINISTER TO RANDOM SUBSAMPLE HOUSEHOLD)** | | |
| C.1501 | 1028. | ব্যাকটেরিয়া আছে কিনা জানার জন্য আমি আপনার খাবার পানির কিছু পরিমান সংগ্রহ করতে পারি কী?<br>May I collect a small sample of your drinking water to test for bacteria in our office? | [1] হ্যাঁ YES, SAMPLE COLLECTED<br>[2] না NO/REFUSED | → **SKIP TO C.1033** |
| C.1502 | 1029. | খানার নমুনা সংগ্রহীত হয়েছে HH sample collected (DD/MM/YY, HH:MM) | খানার নমুনা পানির রেকর্ড HH Water Sample Time Log (can be automated in a bar-code scanner) | |

| | DD/MM/YY | HH:MM |
|---|---|---|
| A | ☐☐/☐☐/☐☐<br>খানার পানির নমুনা সংগ্রহের তারিখ<br>HH sample collected | ☐☐:☐☐<br>সময় |

| | | | | |
|---|---|---|---|---|
| | | | B □□/□□/□□ খানার পানির নমুনা ল্যাবে পৌছানোর তারিখ HH sample arrived at lab | □□:□□ সময় |
| | | | C □□/□□/□□ খানার পানির নমুনা এনালাইসিসের তারিখ HH sample analyzed | □□:□□ সময় |
| c.1503 | 1030. | খানার পানির নমুনা, ফিল্টারকৃত পানির পরিমান (মিলি)<br>HH sample, volume of water filtered (in milliliters) | □□□ মিলি ml | |
| C.1504 | 1031. | খানার পানির নমুনার ল্যাব রেজাল্ট, ই.কোলাই (*E. coli)*<br>(5555 = অসংখ্য, 9999 = এনালাইসিস করা হয় নাই<br>Household sample lab result, *E. coli* (5555 = Too numerous to count, 9999 = Not analyzed) | □□□□ (CFU / 100 ml) | |
| C.1505 | 1032. | খানার নমুনার অবস্থা<br>HH sample qualifier | 1. ডিটেকশন লেভেলের নীচে Below detection limit<br>2. ডিটেকশন লেভেলের উপরে Above detection limit<br>3. নমুনা এনালাইসিস করা হয় নাই Sample could not be analyzed | |
| | | প্রস্তুতির আগে ক্লাষ্টার/খানা আই ডি বোতলে/র্হুয়েলপ্যাকে স্পষ্ট অক্ষরে লিখুন** **WRITE THE CLUSTER/HH ID VERY CLEARLY ON THE WHIRLPACK BEFORE PROCEEDING**** | | |
| | | খানার সমন্বিত পানির উৎসের নমূনা PAIRED SOURCE WATER SAMPLE FOR HH WATER SAMPLE<br>I যদি উত্তরদাতা বলেন যে ভাগের/সেয়াড পানির উৎস থেকে পানি সংগ্রহ করা হয়েছে তাহলে **701** নং প্রশ্নে চলে যান ।IF RESPONDENT REPORTED COLLECTING WATER FROM A SHARED SOURCE → **SKIP TO 701** | | |
| C.1506 | 1033. | যদি উত্তরদাতা বলেন যে নিজস্ব পানির উৎস থেকে পানি সংগ্রহ করা হয়েছে এবং পানির উৎসটি বাড়ীর মধ্যেই অবস্থিত(রেইন ওয়াটারসহ)I F RESPONDENT REPORTED COLLECTING WATER FROM A PRIVATE WATER SOURCE AND THAT WATER SOURCE IS WITHIN THE COMPOUND (INCLUDING RAINWATER), ASK :<br><br>প্রশ্ন কর্হুন;আমি কী আমাদের অফিসে আপনার উৎসের পানি পরীক্ষা করার জন্য কিছু পানি সংগ্রহ করতে পারি ? May I collect a small sample of your source water to test for bacteria in our office?<br><br>পানি সংরক্ষিত হলেই সংগ্রহের রেকর্ড কর্হুন। RECORD WHETHER SAMPLE WAS COLLECTED: | [1] হ্যাঁ ,নমূনা নেয়া হয়েছে YES, SAMPLE COLLECTED<br>[2] না,প্রত্যাখাত হয়েছে NO, REFUSED | → **SKIP TO 701** |
| C.1507 | 1034. | উৎসের পানির আই ডি Water source ID<br>যদি উৎস ভাগের/সেয়াড হয় তখন উৎস আইডির তালিকা দেখুন। If Source is a shared source, list the source ID.<br>যদি উৎস বাড়ীর /কম্পাউন্ড মধ্যে হয় তখন ক্লাস্টার/খানা আইডি হবে উৎস আইডি এবং ৯৯ কোড রেকর্ড করতে হবে। If Source is located in the compound, only list the cluster/HH ID, and record "99" in the Source ID.<br>যদি উৎস ভাগের/সেয়াড হয় এবং ইহা যদি তালিকাতে না থাকে তখন ০০ রেকর্ড করতে হবে। If the shared source located is not on the preprinted list, enter "00" | A □□□ / □□ ক্লাস্টার/খানা আইডি Cluster / HH ID<br>B □□উৎস আইডি (যদি উৎস একই/সেয়াড হয় )Source ID (if a shared source) | |

| | | | |
|---|---|---|---|
| | 1035. | উৎস-এর সংরক্ষিত পানির নমুনা সংরক্ষণকাল (দিন/মাস/বছর.ঘন্টা:মিনিট)Source sample collected (DD/MM/YY, HH:MM) | ☐☐/☐☐/☐☐ ☐☐:☐☐ |
| C.1508 | 1036. | 👁 উৎস-এর পানির ধরণ রেকর্ড কর ন Record the source water type | 12. অগভীর টিউবওয়েল Shallow tubewell<br>13. গভীর টিউবওয়েল Deep tubewell<br>14. ঘরের ভিতর ট্যাপ বা পাইপের পানি Piped into home<br>15. উঠানে ট্যাপ বা পাইপের পানি Piped into yard<br>16. গভীর পাতকূয়া (উঠানে)Borewell in yard<br>17. পাবলিক ট্যাপ Public tap<br>18. পাবলিক গভীর পাতকূয়া Public borewell<br>19. অগভীর পাকা পাতকূয়া Shallow well with concrete reinforcement<br>20. অগভীর কাঁচা পাতকূয়া Shallow well (no concrete)<br>21. নদী/বাধ/লেক/পুকুর/সেচ নালা থেকে সরাসরি সংগৃহীত পানি River, lake, stream<br>22. অন্যান্য (লিখুন) Other: _____________ |
| | | প্রস্তুতির আগে ক্লাষ্টার/খানা আই ডি বোতলে/র য়েলপ্যাকে স্পষ্ট অক্ষরে লিখুন ** WRITE THE SOURCE ID VERY CLEARLY ON THE WHIRLPACK BEFORE PROCEEDING** | |

Wash Benefit Module 12 HOME CARE ENVIRONMENT

Administer to: All study households

If target child is not born at baseline, administer this module if the mother has another child < 36 months

**Learning Stimulation**

## C.1201.

☐☐☐☐☐☐

Record the child ID number of the child about whom these questions are asked

*If the index child is not born at baseline, administer this module if an older sibling is < 36 months*

## C.1202.

☐☐How many children's books or picture books do you have for (name)?

00 None

99 Don't know / refused

## C.1203.

I am interested in learning about the things that (name) plays with when he/she is at home.

Does he/she play with:

1 Yes

2 No

88 Don't know / not sure

| | | |
|---|---|---|
| A | ☐ | Homemade toys (such as dolls, cars, or other toys made at home)? |
| B | ☐ | Toys from a shop or manufactured toys? |
| C | ☐ | Household objects (such as bowls or pots) or objects found outside (such as sticks, rocks, animal shells or leaves)? |

Sometimes adults taking care of children have to leave the house to go shopping, wash clothes, or for other reasons and have to leave young childre

On how many days in the past week was (name):

0 None
99 Don't know / not sure

| | Days | |
|---|---|---|
| A | ☐ | Left alone for more than an hour? |
| B | ☐ | Left in the care of another child, that is, someone less than 10 years old, for more than an hour? |

**C.1204.**
In the past 3 days, did you or any household member over 15 years of age engage in any of the following activities with (name):
If YES, ask:
Who engaged in this activity with (name)?
Circle all that apply.

| | | A | B | C | D |
|---|---|---|---|---|---|
| | | **Mother** | **Father** | **Other** | **Nobody** |
| 1 | Read books to or looked at picture books with (name)? | Y | Y | Y | Y |
| 2 | Told stories to (name)? | Y | Y | Y | Y |
| 3 | Sang songs to (name) or with (name), including lullabies? | Y | Y | Y | Y |
| 4 | Took (name) outside the home, compound, yard or enclosure? | Y | Y | Y | Y |
| 5 | Played with (name)? | Y | Y | Y | Y |
| 6 | Named, counted, or drew things to or with (name)? | Y | Y | Y | Y |

**Spanking**

**C.1205.**
☐ Sometimes kids mind pretty well and sometimes they don't. About how many times have you spanked your child in the past week?
1 None
2 1 or 2 times
3 3–7 times
4 8 or more times
99 Don't know / not sure / refused

**Teaching**

**C.1206.**
☐ Some parents spend time teaching their children new skills while other parents believe children learn best on their own. Which of the followin
attitude?
1 Parent always teaches
2 Parent usually teaches
3 Usually learn on own
4 Always learn on own

99 Don't know / not sure

**Other**

## C.1207.

☐ Children seem to demand attention when their parents are busy, doing housework, for example. How often do you talk to your child while yo

1 Always talking
2 Often talking
3 Sometimes
4 Rarely talk
5 Never talk
99 Don't know / not sure

## C.1208.

☐ Does your child have a regular feeding schedule?

1 Yes
2 No

## C.1209.

☐ Does your child see his/her father or father figure on a daily basis?

1 Yes
2 No

**Observation: Parental Responsiveness**

1 Yes
2 No

| | | |
|---|---|---|
| **C.1210.** | ☐ | (Mother/Guardian) Spontaneously spoke to child twice or more (excluding scolding) |
| **C.1211.** | ☐ | (Mother/Guardian) Responded verbally to child's speech or verbal bids for attention |
| **C.1212.** | ☐ | (Mother/Guardian) Provided toys or interesting activities for child |
| **C.1213.** | ☐ | (Mother/Guardian) Caressed, kissed, or hugged child at least once |
| **C.1214.** | ☐ | (Mother/Guardian) Kept child in view/could see child/looked at (him/her) often |

**Observation: Other**

1 Yes
2 No

| | | |
|---|---|---|
| **C.1215.** | ☐ | (Mother/Guardian) Interfered with child's actions or restricted child from exploring more than 3 times |

| | | |
|---|---|---|
| **C.1216.** | ☐ | Child’s play environment is safe (no potentially dangerous health or structural hazards within a toddler’s or infant’s range) |
| **C.1217.** | ☐ | Reading material (newspapers, magazines, etc.) is present and visible |
| **C.1218.** | ☐ | Child and child’s clothing appear clean |
| **C.1219.** | ☐ | (Mother/Guardian) Slapped or spanked child at least once |
| **C.1220.** | ☐ | There is evidence that older children are handling the child inappropriately (i.e., handling roughly, hitting, etc.) |

**ওয়াস বেনিফিটস কমন মডিউল-১২**

Home Care Environment Version Number 3.0 (2011-09-07)

**সকল খানায় করতে হবে**

**যদি বেইজলাইনের সময় টার্গেট শিশুটি জন্মগ্রহন না করে থাকে তবে ঐ মায়ের <৩৬ মাসের অন্য শিশু থাকলে তার জন্য এই মডিউলটি পূরণ করুন**

| **IDENTIFICATION** | |
|---|---|
| **0.1. CLUSTER ID:** | ☐☐☐☐ |
| **0.2. HOUSEHOLD ID:** | ☐☐☐ |

**শেখার অনুপ্রেরণা**

**C.1221.**
☐☐☐☐☐☐

**যে শিশুর জন্য এই প্রশ্নগুলো করা হবে তার আইডি নাম্বার লিখুন**

**যদি বেইজলাইনের সময় টার্গেট শিশুটি জন্মগ্রহন না করে থাকে তবে ঐ মায়ের <৩৬ মাসের অন্য শিশু থাকলে তার জন্য এই মডিউলটি পূরণ করুন**

**C.1222.**
☐☐শিশুটির [ নাম বলুন] কতগুলো ছোটদের বই/ছবির বই আছে?

0 একটিও না

99 জানি না/প্রত্যাখ্যান

**C.1223.**
শিশুটি যখন বাড়িতে থাকে তখন সে যেসব জিনিস দিয়ে খেলে তা জানার ব্যাপারে আমি আগ্রহী। সে কি নিম্ন লিখিত জিনিস দিয়ে খেলে?

1 হ্যাঁ

2 না

88 জানি না/নিশ্চিত না

| | | |
|---|---|---|
| A | ☐ | ঘরে তৈরী খেলনা, যেমন: পুতুল, গাড়ি অথবা ঘরে তৈরী অন্য খেলনা |
| B | ☐ | দোকান থেকে কেনা অথবা দোকানে তৈরী খেলনা |

| | | |
|---|---|---|
| C | ☐ | ঘরের কোন জিনিস [যেমন: বাটি বা পাত্র] অথবা বাহিরের কোন জিনিস [যেমন: লাঠি, পাথর, ঝিনুক বা পাতা |

## C.1224.

বাচ্চার পরিচর্যাকারীকে অনেক সময় বাচ্চাকে বাড়িতে রেখে কেনাকাটা করার জন্য বা অন্য কাজের জন্য বাহিরে যেতে হয়

গত সপ্তাহে শিশুটির [নাম বলুন] ক্ষেত্রে নিম্নলিখিত ঘটনাগুলো কতবার ঘটেছে?

0 একটিও না

99 জানি না/প্রত্যাখ্যান

| | দিন | |
|---|---|---|
| A | ☐ | এক ঘন্টার চেয়ে বেশী সময়ের জন্য একা রেখে গিয়েছিলেন |
| B | ☐ | এক ঘন্টার চেয়ে বেশী সময়ের জন্য ১০ বছরের ছোট অন্যকোন বাচ্চার কাছে রেখে গিয়েছিলেন |

## C.1225.

গত ৩ দিনে আপনার খানার ১৫ বছরের উপরের বয়সের কোন সদস্য শিশুটির [নাম বলুন] সাথে নিম্নলিখিত কোন কার্যক্রমের সাথে জরিত ছিল কী?

হ্যাঁ হলে, প্রশ্ন করুন: শিশুটির [নাম বলুন] সাথে এইসব কাজে জরিত ছিল?

| | | A | B | C | D |
|---|---|---|---|---|---|
| | | মা<br>হ্যাঁ= 1<br>না= 0 | বাবা<br>হ্যাঁ= 1<br>না= 0 | অন্যকেউ<br>হ্যাঁ= 1<br>না= 0 | কেউ না<br>হ্যাঁ= 1<br>না= 0 |
| 1 | (শিশুটির [নাম বলুন] সাথে বই পড়েছে বা ছবির বই দেখেছে কী?) | Y | Y | Y | Y |
| 2 | (শিশুটিকে [নাম বলুন] গল্প শুনিয়েছে কী?) | Y | Y | Y | Y |
| 3 | (শিশুটিকে [নাম বলুন] ঘুমপাড়ানি গান সহ অন্য গান শুনিয়েছিল বা শিশুটির সাথে গান গেয়েছিল কী?) | Y | Y | Y | Y |
| 4 | (শিশুটিকে [নাম বলুন] ঘরের বাহিরে, উঠানে বা বাড়ীর কাছাকাছি কোন জায়গাতে নিয়ে গিয়েছিল কী?) | Y | Y | Y | Y |
| 5 | (শিশুটির [নাম বলুন] সাথে খেলা করেছিল কী?) | Y | Y | Y | Y |
| 6 | (শিশুটিকে/শিশুটির [নাম বলুন] সাথে নাম বলা, গনণা করা বা ছবি একেঁছিল কী?) | Y | Y | Y | Y |

**আঘাত করা**

## C.1226.

☐ মাঝেমাঝে শিশুরা ভাল থাকে এবং মাঝেমাঝে বিরক্ত করে। গত সপ্তাহে আপনি আপনার শিশুকে কতবার আঘাত করেছেন? [শারিরীক আঘাত])

1 একবারও না

2 ১-২ বার

3 ৩-৭ বার

4 ৮ বার বা তারচেয়ে বেশী বার

99 জানি না/নিশ্চিত না/প্রত্যাখ্যান

শেখানো

## C.1227.

☐ কিছু বাবা/মা তাদের শিশুদের নতুন নতুন জিনিস শিখান, আবার অনেক বাবা/মা মনে করেন শিশুরা একা একাই ভাল শিখতে পারে। আপনার মতে নিম্নের কোনটি সবচেয়ে ভাল--)

1 বাবা/মা সব সময় শিখাবে

2 বাবা/মা সাধারনত শিখাবে

3 শিশু সাধারনত একা শিখবে

4 শিশু সবসময় একা শিখবে

99 জানি না/নিশ্চিত না

**Other**

## C.1228.

☐ বাবা/মা যখন ঘরের কাজ বা এমন অন্যান্য কাজে ব্যস্ত থাকে তখন শিশুরা তাদের মনোযোগ দাবী করে। আপনি যখন কাজ করেন তখন আপনার শিশুর সাথে কতবার কথা বলেন?

1 সব সময় কথা বলি

2 বেশিরভাগ সময় কথা বলি

3 মাঝেমাঝে কথা বলি

4 খুব কম কথা বলি

5 কখনই কথা বলি না

99 জানি না/নিশ্চিত না

## C.1229.

☐ আপনার শিশুর খাওয়ার কোন নির্দিষ্ট সময় তালিকা আছে কী?

1 হ্যাঁ

0. না

## C.1230.

☐ আপনার শিশু তার বাবাকে প্রতিদিন দেখে কী?

1 হ্যাঁ

0. না

**পর্যবেক্ষন ঃ বাবা/মার বাচ্চার প্রতি আগ্রহ**

1 হ্যাঁ

0. না

| | | |
|---|---|---|
| C.1231. | ☐ | মা/উত্তরদাতা স্বতস্ফূর্তভাবে বাচ্চার সাথে ২ বার বা তারচেয়ে বেশী বার কথা বলেছিল |
| C.1232. | ☐ | মা/উত্তরদাতার সাথে বাচ্চা কথা বলার পর মা কথা বলেছে |
| C.1233. | ☐ | মা/ উত্তরদাতা বাচ্চাকে খেলনা দিয়েছে বা বাচ্চার সাথে আনন্দদায়ক কোন আচরন করেছিল |
| C.1234. | ☐ | মা/ উত্তরদাতা বাচ্চাকে অন্তত একবার আদর, চুমু দিয়েছিল বা জরিয়ে ধরেছিল |
| C.1235. | ☐ | মা/ উত্তরদাতা বাচ্চাকে চোখে চোখে রেখেছিল বা তার দিকে প্রায়ই তাকাচ্ছিল |

পর্যবেক্ষন ঃ অন্যান্য

1 হ্যাঁ

0. না

| | | |
|---|---|---|
| C.1236. | ☐ | মা/অভিভাবক ৩ বারের বেশী বাচ্চার বিভিন্ন আচরনে বাধা দিচ্ছিল |
| C.1237. | ☐ | বাচ্চার খেলার পরিবেশ নিরাপদ, শিশুর নাগাল সীমার মধ্যে কোন বিপদজনক কিছু নেই |
| C.1238. | ☐ | পড়ার উপকরণ যেমন: খবরের কাগজ, ম্যাগাজিন ইত্যাদি আছে এবং দেখা যাচ্ছিল |
| C.1239. | ☐ | বাচ্চা এবং বাচ্চার কাপড় পরিষ্কার ছিল |
| C.1240. | ☐ | মা/অভিভাবক কমপক্ষে একবার বাচ্চাকে চড় মেরেছিল বা আঘাত করেছিল |
| C.1241. | ☐ | শিশুটির সাথে বড় বাচ্চারা (<১২ বছর) সঠিক আচরন করছিল না [খারাপভাবে টানাটানি করছিল, আঘাত করছিল] তার প্রমান ছিল |

Wash Benefit Module 13 MEASURES FOR SPILLOVER

**সকল খানার জন্য প্রযোজ্য** Administer to: All study households

| | | | |
|---|---|---|---|
| **C.1301.** | 1301. | **আপনার সবচেয়ে কাছের হেলথ ফ্যাসিলিটি (স্বাস্থ্যসেবা পাওয়ার সুযোগ)-এর নাম কি? যদি না জানে/নিশ্চিত না হয় তবে ৯৯ বসান। খানার কেউ ডাক্তার হলে বা ডাক্তার খানায় এসে চিকিৎসা করলে ৫৫ লিখুন।** What is the name of your nearest health facility?<br>Enter "99" if Don't know / not sure | |
| **C.1301.a** | 1302. | **গত ১ বছরে আপনি/আপনার শিশু মোটামুটি কতবার যে কোন অসুখে এই সুযোগ গ্রহন করেছিলেন?** In the past year, (since this time last year) approximately how many times did you visit this health facility for any reason (including with your children)? | \|__\|__\| বার TIMES |
| **C.1301.b** | 1303. | **আপনি সেখানে কিভাবে যান?** What is your primary mode of travel to the health facility? | **১. পায়ে হেঁটে**<br>**2.** **রিকশা/ভ্যান/নৌকা**<br>**3.** **ব্যাটারি চালিত অটোরিকশা** |

| | | | |
|---|---|---|---|
| | | | 4. ট্যাম্পু/লেগুনা/ সিএনজি/ বাস<br>5. প্রযোজ্য নয় |
| **C.1301.c** | **1303.a** | **আনুমানিক সেখানে যেতে কতক্ষণ সময় লাগে(আপনার খানা থেকে শুধু যেতে কত মিনিট সময় লাগে) ? যদি না জানে/নিশ্চিত না হয় তবে ৯৯ বসান।** Approximately how long does it take to travel there (in minutes, one-way, from where we are now)?<br><br>Enter "999" if Don't know / not sure | \|__\|__\|__\| মিনিটে MINUTES |
| **C.1302.** | 1304. | **আপনার কাছের প্রাক- প্রাইমারী স্কুলের -এর নাম কি ?** যেখানে আপনার শিশু যেতে পারে/বড় হলেও যেতে পারবে । **যদি না জানে/নিশ্চিত না হয় তবে ৯৯, যদি না থাকে তবে ৮৮ বসান।**What is the name of the ECD center [preschool] where your children go (or will go when they are old enough)? | |
| **C.1303.** | 1305. | **আপনার কাছের প্রাইমারী স্কুলের -এর নাম কি ?** যেখানে আপনার শিশু যেতে পারে/বড় হলেও যেতে পারবে । What is the name of the primary school where your children go (or will go when they are old enough)? | |
| **C.1304.** | 1306. | **আপনার কাছের বাজার -এর নাম কি** যেখানে আপনি সবসময়ই কেনা-বেচা করতে যান? । What is the name of the market that you most often go to when you buy or sell things? | |
| **C.1304.a** | 1307. | **কতবার আপনি** সেখানে যান? How often do you go to that market? | 0 **কখনই না** Never<br>1 **সপ্তাহে একবার অথবা তার অধিক** Once per week or more<br>2 **প্রতি ২ সপ্তাহে একবার** Once every 2 weeks<br>3 **প্রতি ৩ সপ্তাহে একবার** Once every 3 weeks<br>4 **প্রতি ৪ সপ্তাহে একবার(প্রতি মাসে)** Once every 4 weeks (every month)<br>5 **প্রতি মাসে একবারের কম** Less than once every month<br>99 **জানি না/নিশ্চিত না** Don't know / not sure |
| | 1308. | **আপনি সেখানে কিভাবে যান?** What is your primary mode of travel to the health facility? | 1. পায়ে হেঁটে<br>2. রিকশা/ভ্যান/নৌকা<br>3. ব্যাটারি চালিত অটোরিকশা<br>4. ট্যাম্পু/লেগুনা/ সিএনজি/ বাস<br>5. প্রযোজ্য নয় |
| **C.1304.b** | **1308.a** | **আনুমানিক সেখানে যেতে কতক্ষণ সময় লাগে(আপনার খানা থেকে শুধু যেতে কত মিনিট সময় লাগে) ? যদি না জানে/নিশ্চিত না হয় তবে ৯৯ বসান।** Approximately how long does it take to travel there (in minutes, one-way, from where we are now)?<br><br>Enter "999" if Don't know / not sure | \|__\|__\|__\| মিনিটে MINUTES |
| **C.1305.** | 1309. | **আপনার কাছের মসজিদ/মন্দির/গীর্জা/প্যাগোডা -এর নাম কি?** যেখানে আপনি প্রার্থনা করতে যান । What is the name of the church/mosque/temple (if any) that you attend? | |
| **C.1305.a** | 1310. | **কতবার আপনি** সেখানে যান? How often do you go to church? | ০. **কখনই না** Never<br>1 **সপ্তাহে একবার অথবা তার অধিক** Once per week or more<br>2 **প্রতি ২ সপ্তাহে একবার** Once every 2 weeks<br>3 **প্রতি ৩ সপ্তাহে একবার** Once every 3 weeks<br>4 **প্রতি ৪ সপ্তাহে একবার(প্রতি মাসে)** Once every 4 weeks (every month)<br>5 **প্রতি মাসে একবারের কম** Less than once every month |

| | | | |
|---|---|---|---|
| | | | 99 জানি না/নিশ্চিত না Don't know / not sure |
| | 1311. | আপনি সেখানে কিভাবে যান? What is your primary mode of travel to the health facility? | 1. পায়ে হেঁটে<br>2. রিকশা/ভ্যান/নৌকা<br>3. ব্যাটারি চালিত অটোরিকশা<br>4. ট্যাম্পু/লেগুনা/ সিএনজি/ বাস<br>5. প্রযোজ্য নয় |
| **C.1305.b** | **1312.a** | আনুমানিক সেখানে যেতে কতক্ষণ সময় লাগে(আপনার খানা থেকে শুধু যেতে কত মিনিট সময় লাগে) ? যদি না জানে/নিশ্চিত না হয় তবে ৯৯ বসান। Approximately how long does it take to travel there (in minutes, one-way, from where we are now)?<br><br>Enter "999" if Don't know / not sure | \|__\|__\|__\| মিনিটে MINUTES |
| **C.1306.** | 1312. | খানাটির জিপিএস অবস্থান রেকর্ড নি শ্চিত করুন। Confirm that you have recorded a GPS location linked to this household | [ 1 ] খানাটির জিপিএস অবস্থান রেকর্ড নি শ্চিত হয়েছে Confirmed, GPS point recorded<br>[ 99 ] রেকর্ড নি শ্চিত হয়নি Could not record |

**Wash Benefit Module 41 Household Assets**

| | | |
|---|---|---|
| আপনার সময়ের জন্য আপনাকে ধন্যবাদ।আমি আপনার খানা সম্বন্ধে জানতে চাই যেখানে আপনি বসবাস করেন। প্রথমেই আমি আপনার ঘর কী দিয়ে তৈরী সেটা দেখতে চাই । তাহলে আমি কি ইহা দেখতে পারি? Thank you very much for your time. Now I would talk about <u>the house that you live in</u>. First, I would like to observe the material your house is made of. Can I take a look at your house?<br><br>OBSERVE MAIN MATERIAL OF THE HOUSE: | | |
| | 4101. | ছাদ তৈরীতে কি কি উপাদান ব্যবহার করা হয়েছে ? (পরির্দশন করে যাচাই করুন) [Main material of the roof]<br>কাঁচা (বাঁশ/খড়) [Kaccha (bamboo / thatch)] 1<br>টিন [Tin] 2<br>সিমেন্ট/ কংক্রিট/ টালি [Cement / concrete / tiled]…3 |
| | 4102. | দেয়াল তৈরীতে কি কি উপাদান ব্যবহার করা হয়েছে ? (পরির্দশন করে যাচাই করুন) [Main material of the walls]<br>পাট/বাঁশ/মাটি (কাঁচা) [Jute / bamboo / mud (*kaccha*)] .........1<br>কাঠ [Wood] .................................................... 2<br>ইট/সিমেন্ট [Brick / cement] ............................... 3<br>টিন [Tin]................................................ 4 |
| | 4103. | মেঝে তৈরীতে কি কি উপাদান ব্যবহার করা হয়েছে ? (পরির্দশন করে যাচাই করুন) [*Main material of the floor]*<br>মাটি/বাঁশ (কাঁচা) [Earth / bamboo (kaccha)] .......................1<br>কাঠ [Wood] 2<br>ইট/ সিমেন্ট [Cement / concrete] .......................................3 |
| Thank you. Now I would like to ask you some questions about your household assets. | | |
| | 4104. | খানার ঘরটিতে কতগুলো কক্ষ আছে? (রান্নাঘর ও বাথরুম বাদে) [How many rooms the households have (exclude bathroom and Kitchen)?]. |
| | 4105. | আপনার খানায় নিম্নলিখিত জিনিসগুলো আছে কী? [How many does the households have<br><u>হ্যাঁ (Yes)....1, না (No)....0, জানি না (DK).....999</u><br>a. \|_____\|বিদ্যুৎ [Electricity]<br>b. \|_____\|\|_____\|আলমারী/ওয়ারড্রব (সংখ্যা) [Number of Almirah or wardrobe]<br>c. \|_____\|\|_____\|টেবিল (সংখ্যা) [Number of tables]<br>d. \|_____\|\|_____\|চেয়ার/বেঞ্চ (সংখ্যা) [Number of chair or bench]<br>e. \|_____\|\|_____\|হাত ঘড়ি/দেয়াল ঘড়ি (সংখ্যা) [Number of watch or clock]<br>f. \|_____\|\|_____\|খাট (সংখ্যা) [Number of khat]<br>g. \|_____\|\|_____\|চৌকি (সংখ্যা) [Number of chouki ]<br>h. \|_____\|রেডিও [A radio that is working]<br>i. \|_____\|টেলিভিশন (সাদা/কালো) [A B/W television that is working]<br>j. \|_____\|টেলিভিশন (রঙিন) [A color television that is working] |

| | | |
|---|---|---|
| | | k. _____\|রেফ্রিজারেটর [Refrigerator]<br>l. \|_____\|সাইকেল (খেলনা সাইকেল নয়) A bicycle (used for commercial purposes not toy for children)<br>m. \|_____\|মটর সাইকেল [A motorcycle]<br>n. \|_____\|সেলাই মেশিন [A sewing machine]<br>o. \|_____\|\|_____\|মোবাইল ফোন (সংখ্যা) [Number of Mobile phones]<br>p. \|_____\|ল্যান্ড ফোন [A land phone] |
| | 4106. | আপনার খানায় রান্নার জন্য প্রধানত কি ধরনের জ্বালানী ব্যবহার করা হয় ? [What type of fuel does your household mainly use for cooking?]<br>কাঠ [Wood] ........ 01<br>শস্যের অবশিষ্টাংশ/ঘাস [Crop residue / grass ] ........ 02<br>শুকনা গোবর [Dung cakes ] ........ 03<br>কয়লা [Coal / coke / lignite ] ........ 04<br>কাঠ কয়লা [Charcoal ] ........ 05<br>কেরোসিন [Kerosene ] ........ 06<br>বিদ্যুৎ [Electricity ] ........ 07<br>তরল গ্যাস/প্রাকৃতিক গ্যাস [Liquid gas / gas ] ........ 08<br>বায়ো-গ্যাস [Bio-gas] ........ 09<br>অন্যান্য [Other] ........ 77<br>(বর্নণা লিখুন) [*Specify other]* ____________________<br>জানি না [Don't know ] ........ 99 |
| | 4107. | \|_____\|আপনার খানার মোট মাসিক আয় কত? (সকল আয়ের উৎসের যোগফল যেমনঃ বেতন, ঘর ভাড়া, কৃষিখাত ইত্যাদি) For the purpose of our research, would you please tell us your total monthly household income? (Please sum up your income from all sources like wage, rent, agriculture etc.)<br>1. 4000 টাকা পর্যন্ত [UP TO 4000 Taka]<br>2. 4001-6000 টাকা [4001-6000 Taka]<br>3. 6001- 8000 টাকা [6001- 8000 Taka]<br>4. 8001- 10000টাকা [8001- 10000 Taka]<br>5. 10001- 12000 টাকা [10001- 12000 Taka]s<br>6. 12001- 15000 টাকা [12001- 15000 Taka]<br>7. 15001-20000 টাকা [15001-20000 Taka]<br>8. 20001- 25000 টাকা [20001- 25000 Taka]<br>9. 25001- 30000 টাকা [25001- 30000 Taka]<br>10. 30000 টাকার উপরে [Above 30000 Taka] |
| | 4108. | \|____\| বসতবাড়ি মালিকানা [What is the ownership status of the house where your household is currently living?]<br>নিজের বাড়ি [Self-owned] ........ 1<br>ভাড়া বাড়ি [Rental] ........ 2<br>সরকারী জমি [Govt. land] ........ 3<br>কোন জমিদারের/জোতদারের বাড়িতে থাকে [Owned by a landlord] ........ 4<br>বাড়ি ভাড়া না দিয়ে অন্যের বাড়িতে থাকে [Living in someone's house without giving rent] .......5 |
| | 4109. | খানার সদস্যদের বসতবাড়িতে কোন জমি আছে কি?<br>1. হ্যাঁ<br>0. না (skip to 4111) |
| | 4110. | \|____\|\|____\|\|____\|\|____\|\|____\| উত্তর হ্যাঁ হলে, খানার সদস্যদের বসতবাড়িতে মোট কতটুকু জমি আছে (ডেসিমেল)? (How much homestead land (decimal) does your household own?]<br>বলতে রাজি হয়নি [Refused]........ 88<br>জানি না [Don't know]........ 99 |
| | 4111. | খানার সদস্যদের বসতবাড়ি ছাড়া অন্য কোন জমি আছে কি?<br>1. হ্যাঁ<br>০. না (skip to 4113) |
| | 4112. | \|____\|\|____\|\|____\|\|____\|\|____\| উত্তর হ্যাঁ হলে, খানার সদস্যদের বসতবাড়িতে মোট কতটুকু জমি আছে (ডেসিমেল)? (How much homestead land (decimal) does your household own?] |

| | | | | |
|---|---|---|---|---|
| | | বলতে রাজি হয়নি [Refused]……………………………88<br>জানি না [Don't know]………………….…............99 | | |
| | 4113. | আপনার মতে সামাজিক প্রেক্ষাপটে আপনার খানাটির অবস্থান কি রকম [How would you describe your economic status?]<br>ধনী [Dhoni]............................................................1<br>উচ্চ মধ্যবিত্ত [Uchho modho bitto] ............................2<br>মধ্যবিত্ত [Modho Bitto] ............................................3<br>দরিদ্র [Doridro or Nimno motho bitto] ....................4<br>হতদরিদ্র [Hotodoridro]............................................5 | | |
| | 4114. | আমি পশুর নামের তালিকা আপনাকে পড়ে শুনাতে চাচ্ছি ।এ গুলো কোনটি আপনার কম্পাউন্ড/বাড়ীতে কতগুলো আছে এবং নিজের কয়টা আছে তা আমাকে বলুন । যদি না থাকে তাহলে ০০ বসান । আর যদি অজানা থাকে তাহলে ৯৯ বসান I'm going to read you a list of animals. Please tell me how many of each stays <u>in your compound</u>, and how many belong to <u>your household</u>. IF NONE, ENTER "00", IF UNKNOWN, ENTER "99" | কম্পাউন্ড/বাড়ী COMPOUND | খানা HOUSEHOLD |
| | 1 | গরু CATTLE | \|__\|__\| | \|__\|__\| |
| | 2 | ছাগল GOATS | \|__\|__\| | \|__\|__\| |
| | 3 | মুরগী/হাঁস CHICKENS | \|__\|__\| | \|__\|__\| |
| | 4115. | কতবার গরু/ছাগল তাদের ঘরের বাহিরে বাড়ীতে/উঠানে ছেড়ে দেন? How often do the cattleroam free in the compound?<br>উত্তরগুলো পড়ুন READ RESPONSES.<br>[1] সবসময় ছেড়ে দেন ALWAYS ROAM FREE<br>[2] মাঝে মাঝে ছেড়ে দেন SOMETIMES ROAM FREE<br>[3] কখনও ছেড়ে দেয়া হয় না NEVER ROAM FREE<br>4. প্রযোজ্য নয়; গরু/ছাগল ছাড়া শুধুমাত্র মুরগী/হাঁস থাকলে N/A; ONLY HAVE CHICKENS | | |
| | 4116. | মুরগী অথবা অন্যান্য পাখি জাতীয় কোন কিছু আপনার প্রধান <u>থাকার</u> ঘরে প্রবেশ করে কি? Do chickens or other poultry ever go inside <u>your main house</u>?<br>READ RESPONSES.<br>[1] সবসময় ALWAYS<br>[2] মাঝে মাঝে SOMETIMES<br>[3] কখনও না NEVER<br>4. প্রযোজ্য নয়; মুরগী/হাঁস না থাকলে N/A; NO CHICKENS | | |

**Wash Benefit Module 42 Feces Observed in and Around the Compound**

সকল খানার জন্য প্রযোজ্য Administer to: All study households

পর্যবেক্ষণ Observation: For the following:

প্রত্যেক জায়গায় মানুষের মল/পায়খানার স্তুপের সংখ্যা পর্যবেক্ষণ করুন (১০টি পর্যন্ত) *Record the number of piles of human feces you observe in each area (up to 10 piles)*

55 অধিক সংখ্যক (স্তুপের সংখ্যা ১০টির উপরে) Too numerous to count (more than 10 piles)

99 বলা যাচ্ছে না/ পর্যবেক্ষণ করা সম্ভব হয়নি Cannot tell / could not observe

উঠানে Within the courtyard

| | | | |
|---|---|---|---|
| **C.826.** | 4201. | উঠানে মানুষের মল/পায়খানা পড়ে আছে যা উম্মুক্ত পায়খানা হিসেবে বিবেচিত হবে **Human** feces <u>within the **courtyard**</u> that could be considered open defecation<br><br>55 অধিক সংখ্যক (স্তুপের সংখ্যা ১০টির উপরে) Too numerous to count (more than 10 piles) | ☐☐ |

| | | | |
|---|---|---|---|
| | | 99 বলা যাচ্ছে না/ **পর্যবেক্ষণ** করা সম্ভব হয়নি Cannot tell / could not observe | |
| **C.827.** | 4202. | **উঠানে পশুর/পাখির মল/পায়খানা পড়ে আছে (নিদ্দিষ্টভাবে উলে- খ করুন)** **Animal** feces present <u>within the **courtyard**</u> (mark all that apply)<br><br>55 অধিক সংখ্যক **(স্তুপের সংখ্যা ১০টির উপরে)** Too numerous to count (more than 10 piles)<br>99 বলা যাচ্ছে না/ **পর্যবেক্ষণ** করা সম্ভব হয়নি Cannot tell / could not observe | ☐1 **পাখি বিশেষ(মুরগী/হাঁস/কবুতর** )Poultry (chicken, duck, pigeon)<br>☐2 **গরু/মহিষ** Cow / Buffalo<br>☐3 **ছাগল/ভেড়া** Goat / Sheep<br>☐4**শুকর** Pig<br>☐5 **কুকুর অথবা বিড়াল** Dog or Cat<br>☐8 **অন্যান্য** Other<br>☐9 **কোনও মল/পায়খানা পাওয়া যায়নি** NO FECES OBSERVED<br>☐99 **পর্যবেক্ষণ** করা সম্ভব হয়নি COULD NOT OBSERVE |
| | 4202.a | **অন্যকোন খানা এই উঠান ব্যবহার করে কী?** Do any other household share this <u>**courtyard**</u> | **১. হ্যাঁ**<br>**০. না** |
| নির্দিষ্ট শিশু বেশীরভাগ সময় যেখানে সময় কাটায় তা নিরুপন ও **যাচাই** করুন । বেইজ লাইনে নির্দিষ্ট শিশু থাকবে না সেজন্য গর্ভবতী মা যেখানে বেশীরভাগ সময় কাটায় তা নিরুপন করুন<br>In the area where the target child spends the most time<br>*Probe and identify where the target child spends the most time. At baseline, the target child will not be present, so the search should be defined as the area where the pregnant mother spends the most time.* | | | |
| **C.828.** | 4203. | **মানুষের মল/পায়খানা পড়ে আছে যেখানে** নির্দিষ্ট শিশু (বেইজলাইনে গর্ভবতী মা) বেশীরভাগ সময় কাটায়। **ইহা উন্মুক্ত পায়খানা হিসেবে বিবেচিত হবে** **Human** feces<u>in the **area where the target child spends the most time**</u> that could be considered open defecation<br><br>55 অধিক সংখ্যক **(স্তুপের সংখ্যা ১০টির উপরে)** Too numerous to count (more than 10 piles)<br>99 বলা যাচ্ছে না/ **পর্যবেক্ষণ** করা সম্ভব হয়নি Cannot tell / could not observe | ☐☐ |
| **C.829.** | 4204. | **উঠানে পশুর/পাখির মল/পায়খানা পড়ে আছে যেখানে** নির্দিষ্ট শিশু (বেইজলাইনে গর্ভবতী মা)বেশীরভাগ সময় সময় কাটায় **আছে । (নিদ্দিষ্টভাবে উলে- খ করুন)** **Animal** feces in the <u>**area where the target child spends the most time**</u><br>(mark all that apply)<br><br>55 অধিক সংখ্যক **(স্তুপের সংখ্যা ১০টির উপরে)** Too numerous to count (more than 10 piles)<br>99 বলা যাচ্ছে না/ **পর্যবেক্ষণ** করা সম্ভব হয়নি Cannot tell / could not observe | ☐1 **পাখি বিশেষ(মুরগী/হাঁস/কবুতর** )Poultry (chicken, duck, pigeon)<br>☐2 **গরু/মহিষ** Cow / Buffalo<br>☐3 **ছাগল/ভেড়া** Goat / Sheep<br>☐4**শুকর** Pig<br>☐5 **কুকুর অথবা বিড়াল** Dog or Cat<br>☐8 **অন্যান্য** Other<br>☐9 **কোনও মল/পায়খানা পাওয়া যায়নি** NO FECES OBSERVED<br>☐99 **পর্যবেক্ষণ** করা সম্ভব হয়নি COULD NOT OBSERVE |
| **নিদ্দিষ্ট খানার পিছনে** Behind the study house | | | |

| | | | |
|---|---|---|---|
| **C.830.** | 4205. | **খানার পিছনে মানুষের মল/পায়খানা পড়ে আছে/থাকলে। ইহা উন্মুক্ত পায়খানা হিসেবে বিবেচিত হবে Human** feces<u>behindthe **house**</u> that could be considered open defecation<br><br>55 অধিক সংখ্যক **(স্তুপের সংখ্যা ১০টির উপরে)** Too numerous to count (more than 10 piles)<br>99 বলা যাচ্ছে না/ **পর্যবেক্ষণ** করা সম্ভব হয়নি Cannot tell / could not observe | ☐☐ |
| **C.831.** | 4206. | **খানার পিছনে পশুর মল/পায়খানা পড়ে আছে/থাকলে। (নির্দ্দিষ্টভাবে উলে- খ কর ন) Animal** feces **<u>behind the house</u>**<br>(mark all that apply)<br><br>55 অধিক সংখ্যক **(স্তুপের সংখ্যা ১০টির উপরে)** Too numerous to count (more than 10 piles)<br>99 বলা যাচ্ছে না/ **পর্যবেক্ষণ** করা সম্ভব হয়নি Cannot tell / could not observe | ☐1 **পাখি বিশেষ(মুরগী/হাঁস/কবুতর** )Poultry (chicken, duck, pigeon)<br>☐2 **গর /মহিষ** Cow / Buffalo<br>☐3 **ছাগল/ভেড়া** Goat / Sheep<br>☐4**শুকর** Pig<br>☐5 **কুকুর অথবা বিড়াল** Dog or Cat<br>☐8 **অন্যান্য** Other<br>☐9 **কোনও মল/পায়খানা পাওয়া যায়নি** NO FECES OBSERVED<br>☐99 **পর্যবেক্ষণ** করা সম্ভব হয়নি COULD NOT OBSERVE |

**Wash Benefit Module 2 DIARRHEA AND ILLNESS SYMPTOMS (Children < 36 months)**

৩৬ মাসের নীচের যে সকল শিশু খানায়/বাড়ীতে বাস করে (বেইজ লাইলের জন্য) । Administer to: Children < 36 months living in a study compound at baseline
This may include the child in the household or in other households in the bari.
PDA TO OPEN CHILD ID FROM **DIARRHEA COHORT**
যদি উত্তরদাতার 0-36 মাসের শিশু না থাকে তাহলে এখানেই শেষ করুন নং প্রশ্ন যেতে হবে | IF study compound DOES NOT HAVE A CHILD 0-36 MONTHS → **SKIP TO END**

**উত্তরদাতা শিশুর প্রধান পরিচর্যাকারী** Respondent: Child's primary caregiver

| | | | | | | |
|---|---|---|---|---|---|---|
| | WBM 2 | **আপনাকে ধন্যবাদ । আমি এখন** আপনাকে আপনার শিশুর স্বাস্থ্য বিষয়ে কিছু প্রশ্ন জিজ্ঞাসা করব। দয়া করে আপনি **আমাকে ঠিকমতো প্রত্যক প্রশ্নের জবাব দিবেন। যদি উত্তর না জানা থাকে তখন বলবেন জানি না। শিশুটির নাম ধরে শুর‍ু করতে হবে।** Thank you. Now I am going to ask you some questions about the health of your [child/children]. Please answer each question as accurately as you can. If you don't know the answer to a question, say "I don't know." We will begin with [NAME OF CHILD 1]. | | | | |
| | | | **A** | **B** | **C** | **D** |
| | | **শিশুটির নাম বলুন এবং তার লক্ষণ গুলো ছিল কী?Did [NAME] have [SYMPTOM] :** | আজ **Today** ১=হ্যাঁ YES ০=না NO ৯৯৯=জানি না DK | গতকাল **Yesterday** ১=হ্যাঁ YES ০=না NO ৯৯৯=জানি না DK | গত পরশুদিন **Day before Yesterday** ১=হ্যাঁ YES ০=না NO ৯৯৯=জানি না DK | গত ৭দিনে In **the last 7 days** (since this day last week) ১=হ্যাঁ YES ০=না NO ৯৯৯=জানি না না DK |
| **C.201** | 201. | জ্বর Fever | | | | |
| **C.202** | 202. | **পাতলা পায়খানা** Diarrhea | | | | |
| **C.203** | 203. | ২৪ ঘন্টায় ৩ বার বা তার বেশী পায়খানা করেছে কিনা? 3 or more bowel movements in 24 hours | | | | |
| **C.204** | 204. | দৈনিক পায়খানায় সংখ্যা Number of bowl movements each day | | | | |
| **C.205** | 205. | পানির মত বা নরম পায়খানা? Watery or soft stool (unformed) | | | | |
| **C.206** | 206. | পায়খানায় রক্ত Blood in the stool | | | | |
| **C.207** | 207. | শরীর বা মাথায় চুলকানি, রেশ?Skin rash (anywhere on the body) | | | | |
| **C.208** | 208. | দীর্ঘ কাশি Constant cough | | | | |
| **C.209** | 209. | নাক বন্ধ/নাক দিয়ে পানি পড়া?Congestion / runny nose | | | | |
| **C.210** | 210. | শ্বাস নেওয়ার সময় শোঁ শোঁ করে আওয়াজ বা শ্বাসকষ্ট? (যে কোন ধরনের শ্বাসকষ্ট, তবে নাক বন্ধ থাকার কারণে শ্বাসকষ্ট হলে তা বাদ দিতে হবে) Panting / wheezing / difficulty breathing | | | | |
| **C.211** | 211. | দাগ, থেঁৎলে যাওয়া বা কাটা Bruising, scrapes or cuts | | | | |
| **C.212** | 212. | দাঁতের ব্যাথাToothache / teething | | | | |

**Wash Benefit Module 4**

Administer to: Children at midline and endline

| IDENTIFICATION | |
|---|---|
| **0.1. CLUSTER ID:** | ☐☐☐☐ |
| **0.2. HOUSEHOLD ID:** | ☐☐☐ |
| **0.3. CHILD ID** | ☐☐☐☐☐☐ |

| | | |
|---|---|---|
| **C.401**<br>**FRA ID** | ## | ☐☐ |
| **C.402**<br>**Name of FRA** | Full Name | |
| **C.403**<br>**Is mother wearing heavy clothing during weight measurement?** | 1 = Light clothing<br>2 = Light clothing plus sweater<br>3 = Heavy clothing | ☐ |
| **C.404**<br>**Weight of Mother**<br>**Measurement #1** | Weight<br>(kg) | ☐☐.☐ |
| **C.405**<br>**Weight of Mother**<br>**Measurement #2** | Weight<br>(kg) | ☐☐.☐ |
| **C.406**<br>**Weight of Mother**<br>**Measurement #3**<br>(If difference between<br>measures 1 & 2 is ≥ 0.1 kg) | Weight<br>(kg) | ☐☐.☐ |

| | | |
|---|---|---|
| **C.407**<br>**Is child wearing clothing during weight measurement?** | 0 = No Clothes<br>1 = Only Shirt<br>2 = Only Pants<br>3 = Both Shirt & Pants | ☐ |
| **C.408**<br>**Weight of Mother + Child**<br>**Measurement #1** | Weight<br>(kg) | ☐☐.☐ |
| **C.409**<br>**Weight of Mother + Child**<br>**Measurement #2** | Weight<br>(kg) | ☐☐.☐ |
| **C.410**<br>**Weight of Mother + Child**<br>**Measurement #3**<br>(If difference between<br>measures 1 & 2 is ≥ 0.1 kg) | Weight<br>(kg) | ☐☐.☐ |
| **C.411 – C.413**<br>reserved for child weight measurement without mother (follow-up visits) | | |
| **C.414**<br>**Length of Child**<br>**Measurement #1** | Length<br>(cm) | ☐☐☐.☐ |
| **C.415**<br>**Length of Child**<br>**Measurement #2** | Length<br>(cm) | ☐☐☐.☐ |
| **C.416**<br>**Length of Child**<br>**Measurement #3**<br>(If difference between<br>measures 1 & 2 is ≥ 0.5 cm) | Length<br>(cm) | ☐☐☐.☐ |

| | | |
|---|---|---|
| **C.417**<br>**Length Measurement Method** | Child was:<br><br>1 = lying (recumbent)<br>2 = standing | □ |
| **C.418**<br>**Head Circumference**<br>**Measurement #1** | Circumference<br>(cm) | □□.□ |
| **C.419**<br>**Head Circumference**<br>**Measurement #2** | Circumference<br>(cm) | □□.□ |
| **C.420**<br>**Head Circumference**<br>**Measurement #3**<br>(If difference between<br>measures 1 & 2 is ≥ 0.5 cm) | Circumference<br>(cm) | □□.□ |
| **C.421**<br>**Does the child have swollen feet (bi-pedal edema)?** | 1 = Yes (**>> Referral**)<br>2 = No | □ |

ওয়াস বেনিফিটস কমন মডিউল-৪

এনথ্রোপোমেট্রিক পরিমাপক

বেইজলাইনের সময় যে সকল খানায় <৩ মাসের শিশু আছে

| **IDENTIFICATION** | |
|---|---|
| **0.1.** ক্লাস্টার আইডি: | □□□□ |
| **0.2.** খানা আইডি: | □□□ |
| **0.3.** শিশুর আইডিঃ | □□□□□□ |

| | | |
|---|---|---|
| **C.401**<br>**এফ.আর.এ আইডি** | ## | □□ |
| **C.402**<br>**এফ.আর.এ-র নাম** | **পুরো নাম** | |
| **C.403**<br>ওজন নেওয়ার সময় বাচ্চার মা ভারী কাপড় পড়েছিল কী? | 1 = হালকা কাপড়<br>2 = সোয়েটারসহ হালকা কাপড়<br>3 = ভারী কাপড় | □ |
| **C.404**<br>মায়ের ওজন<br>পরিমাপ#1 | ওজন(কেজি)<br>0 =ওজন নেওয়া সম্ভব হয় নি | □□.□ |
| **C.405**<br>মায়ের ওজন<br>পরিমাপ#2 | ওজন(কেজি)<br>0 =ওজন নেওয়া সম্ভব হয় নি | □□.□ |
| **C.406**<br>মায়ের ওজন<br>পরিমাপ#3<br>(যদি পরিমাপ#1 এবং পরিমাপ#2 এর মধ্যে ≥ **0.1** কেজিপার্থক্য থাকে) | ওজন(কেজি)<br>0 =ওজন নেওয়া সম্ভব হয় নি | □□.□ |
| **C.407**<br>ওজন নেওয়ার সময় বাচ্চা কোন কাপড় পড়েছিল কী? | 0 = কোন কাপড় নেই<br>1 = শুধুমাত্র শার্ট<br>2 = শুধুমাত্র প্যান্ট<br>3 = শার্ট এবং প্যান্ট উভয় | □ |
| **C.408**<br>মা+বাচ্চার ওজন<br>পরিমাপ#1 | ওজন(কেজি)<br>0 =ওজন নেওয়া সম্ভব হয় নি | □□.□ |

| | | |
|---|---|---|
| **C.409**<br>মা+বাচ্চার ওজন<br>পরিমাপ#2 | ওজন(কেজি)<br><br>0 =ওজন নেওয়া সম্ভব হয় নি | □□.□ |
| **C.410**<br>মা+বাচ্চার ওজন<br>পরিমাপ#3<br>(যদি পরিমাপ#1 এবং পরিমাপ#2 এর মধ্যে ≥ **0.1** কেজিপার্থক্য থাকে) | ওজন(কেজি)<br><br>0 =ওজন নেওয়া সম্ভব হয় নি | □□.□ |
| **C.411 – C.413**<br>reserved for child weight measurement without mother (follow-up visits) | | |
| **C.414**<br>বাচ্চার উচ্চতা<br>পরিমাপ#1 | উচ্চতা(সেমি)<br><br>0=উচ্চতা নেওয়া সম্ভব হয়নি | □□□.□ |
| **C.415**<br>বাচ্চার উচ্চতা<br>পরিমাপ#2 | উচ্চতা(সেমি)<br><br>0=উচ্চতা নেওয়া সম্ভব হয়নি | □□□.□ |
| **C.416**<br>বাচ্চার উচ্চতা<br>পরিমাপ#3<br>(যদি পরিমাপ#1 এবং পরিমাপ#2 এর মধ্যে ≥ **0.5** সেমি.পার্থক্য থাকে) | উচ্চতা(সেমি)<br><br>0=উচ্চতা নেওয়া সম্ভব হয়নি | □□□.□ |
| **C.417**<br>উচ্চতা পরিমাপক পদ্ধতি | 1 = শুয়ে মাপা হয়েছে<br>2 = দাড়িঁয়ে মাপা হয়েছে | □ |
| **C.418**<br>মাথার পরিধি<br>পরিমাপ#1 | পরিধি(সেমি)<br><br>0 =মাথার পরিধি নেওয়া সম্ভব হয়নি | □□.□ |

| | | |
|---|---|---|
| **C.419**<br>মাথার পরিধি<br>পরিমাপ#2 | পরিধি(সেমি)<br><br>0 =মাথার পরিধি নেওয়া সম্ভব হয়নি | ☐☐.☐ |
| **C.420**<br>মাথার পরিধি<br>পরিমাপ#3<br>(যদি পরিমাপ#1 এবং পরিমাপ#2 এর মধ্যে ≥ **0.5** সেমি.পার্থক্য থাকে) | পরিধি(সেমি)<br><br>0 =মাথার পরিধি নেওয়া সম্ভব হয়নি | ☐☐.☐ |
| **C.421**<br>**Does the child have swollen feet (bi-pedal edema)?** | 1 = Yes (**>> Referral**)<br>2 = No | ☐ |

PROTOCOL
Biomedical
Non-Exempt
Berkeley

Protocol # 2011-09-3652
Date Printed: 07/31/2018

---------------------------------------------------------------------------------------

**Protocol Title:** Measuring the benefits of sanitation, water quality, handwashing and nutrition interventions for improving health and development in rural Bangladesh

**Protocol Status:** APPROVED

**Date Submitted:** 10/26/2017

**Approval Period:** **10/30/2017-11/03/2018**

**Important Note:** **This Print View may not reflect all comments and contingencies for approval. Please check the comments section of the online protocol. Questions that appear to not have been answered may not have been required for this submission. Please see the system application for more details.**

*** * * Attached Document * * ***

| Document Name | Created Date |
| --- | --- |
| Eugene CITI Completion Report 20130726.pdf | 10/26/2017 |

### CITI Collaborative Institutional Training Initiative

#### Human Research Curriculum Completion Report
#### Printed on 7/26/2013

**Learner:** Eugene Konagaya (username: ekonagaya)
**Institution:** University of California, Berkeley
**Contact Information** 832 Redondo Ave. Apt.B
832 Redondo Ave. Apt.B
Long Beach, CA 90804 United States
Department: Public Health


**Group 1 Biomedical Research Investigators and Key Personnel:**

**Stage 1. Basic Course Passed on 07/26/13** (Ref # 10876380)

| Required Modules | Date Completed | Score |
|---|---|---|
| Belmont Report and CITI Course Introduction | 07/26/13 | 3/3 (100%) |
| History and Ethical Principles | 07/26/13 | 6/6 (100%) |
| Basic Institutional Review Board (IRB) Regulations and Review Process | 07/26/13 | 5/5 (100%) |
| Informed Consent | 07/26/13 | 4/4 (100%) |
| Social and Behavioral Research (SBR) for Biomedical Researchers | 07/26/13 | 4/4 (100%) |
| Records-Based Research | 07/26/13 | 2/2 (100%) |
| Genetic Research in Human Populations | 07/26/13 | 2/2 (100%) |
| Research With Protected Populations - Vulnerable Subjects: An Overview | 07/26/13 | 4/4 (100%) |
| Vulnerable Subjects - Research Involving Prisoners | 07/26/13 | 4/4 (100%) |
| Vulnerable Subjects - Research Involving Children | 07/26/13 | 3/3 (100%) |
| Vulnerable Subjects - Research Involving Pregnant Women, Human Fetuses, and Neonates | 07/26/13 | 3/3 (100%) |
| International Studies | 07/26/13 | 3/3 (100%) |
| FDA-Regulated Research | 07/26/13 | 5/5 (100%) |
| Research and HIPAA Privacy Protections | 07/26/13 | 5/5 (100%) |
| Vulnerable Subjects - Research Involving Workers/Employees | 07/26/13 | 4/4 (100%) |
| Hot Topics | 07/26/13 | no quiz |
| Conflicts of Interest in Research Involving Human Subjects | 07/26/13 | 5/5 (100%) |
| Avoiding Group Harms - U.S. Research Perspectives | 07/26/13 | 3/3 (100%) |
| Unanticipated Problems and Reporting Requirements in | 07/26/13 | 6/6 (100%) |

| | | |
|---|---|---|
| Biomedical Research | | |
| University of California, Berkeley | 07/26/13 | no quiz |
| Stem Cell Research Oversight (Part I) | 07/26/13 | 4/5 (80%) |

**For this Completion Report to be valid, the learner listed above must be affiliated with a CITI participating institution. Falsified information and unauthorized use of the CITI course site is unethical, and may be considered scientific misconduct by your institution.**

Paul Braunschweiger Ph.D.
Professor, University of Miami
Director Office of Research Education
CITI Course Coordinator

Return

PROTOCOL
Biomedical
Non-Exempt
Berkeley

-----------------------------------------------------------------------------------

**Protocol Title:** Measuring the benefits of sanitation, water quality, handwashing and nutrition interventions for improving health and development in rural Bangladesh

**Protocol Status:** APPROVED

**Date Submitted:** 10/26/2017

**Approval Period:** **10/30/2017-11/03/2018**

**Important Note:** **This Print View may not reflect all comments and contingencies for approval. Please check the comments section of the online protocol. Questions that appear to not have been answered may not have been required for this submission. Please see the system application for more details.**

*** * * Attached Document * * ***

| Document Name | Created Date |
|---|---|
| BenArnold-CITI.pdf | 10/26/2017 |

### CITI Collaborative Institutional Training Initiative

Print This Report

**Monday, October 5, 2009**

**CITI Course Completion Record # 212399**
**for BENJAMIN ARNOLD**

To whom it may concern:

On *8/9/2006*, *BENJAMIN ARNOLD* (username=benarnold) completed all *CITI Program* requirements for the Collaborative Institutional Training Initiative.

**Learner Institution:** *University of California, Berkeley*

**Learner Group:** *Group 1Biomedical Research Investigators and Key Personnel*

**Learner Group Description:**

| **The Required Modules for *Group 1Biomedical Research Investigators and Key Personnel* are:** | **Date completed** |
|---|---|
| Introduction | |
| History and Ethical Principles | 07/06/06 |
| Basic Institutional Review Board (IRB) Regulations and Review Process | 07/06/06 |
| Informed Consent | 07/06/06 |
| Social and Behavioral Research for Biomedical Researchers | 08/08/06 |
| Records-Based Research | 08/09/06 |
| Genetic Research in Human Populations | 08/09/06 |
| Research With Protected Populations - Vulnerable Subjects: An Overview | 08/09/06 |
| Vulnerable Subjects - Research with Prisoners | 08/09/06 |
| Vulnerable Subjects - Research Involving Minors | 08/09/06 |
| Vulnerable Subjects - Research Involving Pregnant Women and Fetuses in Utero | 08/09/06 |
| Group Harms: Research With Culturally or Medically Vulnerable Groups | 08/09/06 |

| | |
|---|---|
| FDA-Regulated Research | 08/09/06 |
| International Research | |
| Human Subjects Research at the VA | 08/09/06 |
| HIPAA and Human Subjects Research | 08/09/06 |
| Workers as Research Subjects-A Vulnerable Population | 08/09/06 |
| Hot Topics | |
| Conflicts of Interest in Research Involving Human Subjects | 08/09/06 |
| University of California, Berkeley | |

**For this Completion Report to be valid, the learner listed above must be affiliated with a CITI participating institution. Falsified information and unauthorized use of the CITI course site is unethical, and may be considered scientific misconduct by your institution.**

Paul Braunschweiger Ph.D.
Professor, University of Miami
Director Office of Research Education
CITI Course Coordinator

**CR # 212399**

PROTOCOL
Biomedical
Non-Exempt
Berkeley

Protocol # 2011-09-3652
Date Printed: 07/31/2018

-----------------------------------------------------------------------------------

| | |
|---|---|
| **Protocol Title:** | Measuring the benefits of sanitation, water quality, handwashing and nutrition interventions for improving health and development in rural Bangladesh |
| **Protocol Status:** | APPROVED |
| **Date Submitted:** | 10/26/2017 |
| **Approval Period:** | **10/30/2017-11/03/2018** |
| **Important Note:** | **This Print View may not reflect all comments and contingencies for approval. Please check the comments section of the online protocol. Questions that appear to not have been answered may not have been required for this submission. Please see the system application for more details.** |

*** * * Attached Document * * ***

| Document Name | Created Date |
|---|---|
| CitiApprovalBiomedical_Hubbard2013.pdf | 10/26/2017 |

### CITI Collaborative Institutional Training Initiative

#### Human Research Curriculum Completion Report
#### Printed on 4/12/2013

**Learner:** Alan Hubbard (username: ahubb40)
**Institution:** University of California, Berkeley
**Contact Information** School of Public Health, Div. of Biostatistics
101 Haviland Hall, MC 7358
University of California
Berkeley, CA 94720 USA


**Group 1 Biomedical Research Investigators and Key Personnel:**

**Stage 1. Basic Course Passed on 04/12/13** (Ref # 6200015)

| Required Modules | Date Completed | Score |
|---|---|---|
| Belmont Report and CITI Course Introduction | 04/10/13 | 2/3 (67%) |
| History and Ethical Principles | 04/10/13 | 6/6 (100%) |
| Basic Institutional Review Board (IRB) Regulations and Review Process | 04/11/13 | 5/5 (100%) |
| Informed Consent | 04/11/13 | 4/4 (100%) |
| Social and Behavioral Research for Biomedical Researchers | 04/11/13 | 3/4 (75%) |
| Records-Based Research | 04/11/13 | 2/2 (100%) |
| Genetic Research in Human Populations | 04/11/13 | 2/2 (100%) |
| Research With Protected Populations - Vulnerable Subjects: An Overview | 04/11/13 | 4/4 (100%) |
| Vulnerable Subjects - Research Involving Prisoners | 04/11/13 | 4/4 (100%) |
| Vulnerable Subjects - Research Involving Children | 04/11/13 | 3/3 (100%) |
| Vulnerable Subjects - Research Involving Pregnant Women, Human Fetuses, and Neonates | 04/11/13 | 3/3 (100%) |
| International Studies | 04/11/13 | 3/3 (100%) |
| FDA-Regulated Research | 04/11/13 | 5/5 (100%) |
| Human Subjects Research at the VA | 04/11/13 | 2/3 (67%) |
| Research and HIPAA Privacy Protections | 04/11/13 | 3/5 (60%) |
| Vulnerable Subjects - Research Involving Workers/Employees | 04/12/13 | 4/4 (100%) |
| Hot Topics | 04/12/13 | no quiz |
| Conflicts of Interest in Research Involving Human Subjects | 04/12/13 | 3/5 (60%) |
| Avoiding Group Harms: U.S. Research Perspectives | 04/12/13 | 3/3 (100%) |
| Unanticipated Problems and Reporting Requirements in Biomedical Research | 04/12/13 | 3/6 (50%) |

| | | |
|---|---|---|
| University of California, Berkeley | 04/12/13 | no quiz |
| Stem Cell Research Oversight (Part I) | 04/12/13 | 4/5 (80%) |

**For this Completion Report to be valid, the learner listed above must be affiliated with a CITI participating institution. Falsified information and unauthorized use of the CITI course site is unethical, and may be considered scientific misconduct by your institution.**

Paul Braunschweiger Ph.D.
Professor, University of Miami
Director Office of Research Education
CITI Course Coordinator

Return

PROTOCOL
Biomedical
Non-Exempt
Berkeley

Protocol # 2011-09-3652
Date Printed: 07/31/2018

---------------------------------------------------------------------------------
---

**Protocol Title:** Measuring the benefits of sanitation, water quality, handwashing and nutrition interventions for improving health and development in rural Bangladesh

**Protocol Status:** APPROVED

**Date Submitted:** 10/26/2017

**Approval Period:** **10/30/2017-11/03/2018**

**Important Note:** **This Print View may not reflect all comments and contingencies for approval. Please check the comments section of the online protocol.
Questions that appear to not have been answered may not have been required for this submission. Please see the system application for more details.**

*** * * Attached Document * * ***

| Document Name | Created Date |
|---|---|
| Colford CITI Completion Report.pdf | 10/26/2017 |

### CITI Collaborative Institutional Training Initiative

#### Human Research Curriculum Completion Report
#### Printed on 7/27/2013

**Learner:** John Colford (username:)
**Institution:** University of California, Berkeley
**Contact Information** 101 Haviland MC 7358
Division of Epidemiology
2625 Middlefield PMB 860
Berkeley, California 94720-7358 United States
Department: Public Health


**Group 1 Biomedical Research Investigators and Key Personnel:**

**Stage 1. Basic Course Passed on 07/27/13** (Ref # 10875285)

| Required Modules | Date Completed | Score |
|---|---|---|
| Belmont Report and CITI Course Introduction | 07/26/13 | 3/3 (100%) |
| History and Ethical Principles | 07/26/13 | 5/6 (83%) |
| Basic Institutional Review Board (IRB) Regulations and Review Process | 07/27/13 | 5/5 (100%) |
| Informed Consent | 07/27/13 | 4/4 (100%) |
| Social and Behavioral Research (SBR) for Biomedical Researchers | 07/27/13 | 4/4 (100%) |
| Records-Based Research | 07/27/13 | 2/2 (100%) |
| Genetic Research in Human Populations | 07/27/13 | 2/2 (100%) |
| Research With Protected Populations - Vulnerable Subjects: An Overview | 07/27/13 | 4/4 (100%) |
| Vulnerable Subjects - Research Involving Prisoners | 07/27/13 | 4/4 (100%) |
| Vulnerable Subjects - Research Involving Children | 07/27/13 | 3/3 (100%) |
| Vulnerable Subjects - Research Involving Pregnant Women, Human Fetuses, and Neonates | 07/27/13 | 3/3 (100%) |
| International Studies | 07/27/13 | 3/3 (100%) |
| FDA-Regulated Research | 07/27/13 | 5/5 (100%) |
| Research and HIPAA Privacy Protections | 07/27/13 | 3/5 (60%) |
| Vulnerable Subjects - Research Involving Workers/Employees | 07/27/13 | 4/4 (100%) |
| Hot Topics | 07/27/13 | no quiz |
| Conflicts of Interest in Research Involving Human Subjects | 07/27/13 | 4/5 (80%) |
| Avoiding Group Harms - U.S. Research Perspectives | 07/27/13 | 3/3 (100%) |
| Unanticipated Problems and Reporting Requirements in Biomedical Research | 07/27/13 | 4/6 (67%) |

| | | |
|---|---|---|
| University of California, Berkeley | 07/27/13 | no quiz |
| Stem Cell Research Oversight (Part I) | 07/27/13 | 5/5 (100%) |

**For this Completion Report to be valid, the learner listed above must be affiliated with a CITI participating institution. Falsified information and unauthorized use of the CITI course site is unethical, and may be considered scientific misconduct by your institution.**

Paul Braunschweiger Ph.D.
Professor, University of Miami
Director Office of Research Education
CITI Course Coordinator

Return

PROTOCOL
Biomedical
Non-Exempt
Berkeley

---------------------------------------------------------------------------------
---

| | |
|---|---|
| **Protocol Title:** | Measuring the benefits of sanitation, water quality, handwashing and nutrition interventions for improving health and development in rural Bangladesh |
| **Protocol Status:** | APPROVED |
| **Date Submitted:** | 10/26/2017 |
| **Approval Period:** | **10/30/2017-11/03/2018** |
| **Important Note:** | This Print View may not reflect all comments and contingencies for approval. Please check the comments section of the online protocol. Questions that appear to not have been answered may not have been required for this submission. Please see the system application for more details. |

*** * * Attached Document * * ***

| Document Name | Created Date |
|---|---|
| Fernald CITI Group 1 Completion Report.pdf | 10/26/2017 |

### CITI Collaborative Institutional Training Initiative

#### Human Research Curriculum Completion Report
#### Printed on 7/28/2013

**Learner:** Lia Fernald (username: liafernald)
**Institution:** University of California, Berkeley


**Group 1 Biomedical Research Investigators and Key Personnel:**

**Stage 1. Basic Course Passed on 07/28/13** (Ref # 10881428)

| Required Modules | Date Completed | Score |
|---|---|---|
| Belmont Report and CITI Course Introduction | 07/27/13 | 3/3 (100%) |
| History and Ethical Principles | 07/27/13 | 5/6 (83%) |
| Basic Institutional Review Board (IRB) Regulations and Review Process | 07/27/13 | 5/5 (100%) |
| Informed Consent | 07/27/13 | 3/4 (75%) |
| Social and Behavioral Research (SBR) for Biomedical Researchers | 07/27/13 | 4/4 (100%) |
| Records-Based Research | 07/27/13 | 2/2 (100%) |
| Genetic Research in Human Populations | 07/27/13 | 2/2 (100%) |
| Research With Protected Populations - Vulnerable Subjects: An Overview | 07/27/13 | 4/4 (100%) |
| Vulnerable Subjects - Research Involving Prisoners | 07/27/13 | 4/4 (100%) |
| Vulnerable Subjects - Research Involving Children | 07/28/13 | 3/3 (100%) |
| Vulnerable Subjects - Research Involving Pregnant Women, Human Fetuses, and Neonates | 07/28/13 | 3/3 (100%) |
| International Studies | 07/28/13 | 3/3 (100%) |
| FDA-Regulated Research | 07/28/13 | 5/5 (100%) |
| Research and HIPAA Privacy Protections | 07/28/13 | 2/5 (40%) |
| Vulnerable Subjects - Research Involving Workers/Employees | 07/28/13 | 4/4 (100%) |
| Hot Topics | 07/28/13 | no quiz |
| Conflicts of Interest in Research Involving Human Subjects | 07/28/13 | 4/5 (80%) |
| Avoiding Group Harms - U.S. Research Perspectives | 07/28/13 | 3/3 (100%) |
| Unanticipated Problems and Reporting Requirements in Biomedical Research | 07/28/13 | 4/6 (67%) |
| University of California, Berkeley | 07/28/13 | no quiz |
| Stem Cell Research Oversight (Part I) | 07/28/13 | 3/5 (60%) |

**For this Completion Report to be valid, the learner listed above must be**

**affiliated with a CITI participating institution. Falsified information and unauthorized use of the CITI course site is unethical, and may be considered scientific misconduct by your institution.**

Paul Braunschweiger Ph.D.
Professor, University of Miami
Director Office of Research Education
CITI Course Coordinator

Return

PROTOCOL
Biomedical
Non-Exempt
Berkeley

Protocol # 2011-09-3652
Date Printed: 07/31/2018

-------------------------------------------------------------------------------------

| | |
|---|---|
| **Protocol Title:** | Measuring the benefits of sanitation, water quality, handwashing and nutrition interventions for improving health and development in rural Bangladesh |
| **Protocol Status:** | APPROVED |
| **Date Submitted:** | 10/26/2017 |
| **Approval Period:** | **10/30/2017-11/03/2018** |
| **Important Note:** | **This Print View may not reflect all comments and contingencies for approval. Please check the comments section of the online protocol. Questions that appear to not have been answered may not have been required for this submission. Please see the system application for more details.** |

*** * * Attached Document * * ***

| Document Name | Created Date |
|---|---|
| Audrie - Part 2 Citi.pdf | 10/26/2017 |

### CITI Collaborative Institutional Training Initiative

#### Human Research Curriculum Completion Report
#### Printed on 7/26/2013

**Learner:** Audrie Lin (username: audrielin)
**Institution:** University of California, Berkeley


**Group 1 Biomedical Research Investigators and Key Personnel:**

**Stage 1. Basic Course Passed on 07/26/13** (Ref # 10872675)

| Required Modules | Date Completed | Score |
|---|---|---|
| Belmont Report and CITI Course Introduction | 07/26/13 | 3/3 (100%) |
| History and Ethical Principles | 07/26/13 | 6/6 (100%) |
| Basic Institutional Review Board (IRB) Regulations and Review Process | 07/26/13 | 5/5 (100%) |
| Informed Consent | 07/26/13 | 4/4 (100%) |
| Social and Behavioral Research (SBR) for Biomedical Researchers | 07/26/13 | 2/4 (50%) |
| Records-Based Research | 07/26/13 | 2/2 (100%) |
| Genetic Research in Human Populations | 07/26/13 | 2/2 (100%) |
| Research With Protected Populations - Vulnerable Subjects: An Overview | 07/26/13 | 4/4 (100%) |
| Vulnerable Subjects - Research Involving Prisoners | 07/26/13 | 3/4 (75%) |
| Vulnerable Subjects - Research Involving Children | 07/26/13 | 3/3 (100%) |
| Vulnerable Subjects - Research Involving Pregnant Women, Human Fetuses, and Neonates | 07/26/13 | 3/3 (100%) |
| International Studies | 07/26/13 | 2/3 (67%) |
| FDA-Regulated Research | 07/26/13 | 5/5 (100%) |
| Research and HIPAA Privacy Protections | 07/26/13 | 5/5 (100%) |
| Vulnerable Subjects - Research Involving Workers/Employees | 07/26/13 | 4/4 (100%) |
| Hot Topics | 07/26/13 | no quiz |
| Conflicts of Interest in Research Involving Human Subjects | 07/26/13 | 5/5 (100%) |
| Avoiding Group Harms - U.S. Research Perspectives | 07/26/13 | 3/3 (100%) |

| | | |
|---|---|---|
| Unanticipated Problems and Reporting Requirements in Biomedical Research | 07/26/13 | 5/6 (83%) |
| University of California, Berkeley | 07/26/13 | no quiz |
| Stem Cell Research Oversight (Part I) | 07/26/13 | 5/5 (100%) |

**For this Completion Report to be valid, the learner listed above must be affiliated with a CITI participating institution. Falsified information and unauthorized use of the CITI course site is unethical, and may be considered scientific misconduct by your institution.**

Paul Braunschweiger Ph.D.
Professor, University of Miami
Director Office of Research Education
CITI Course Coordinator

Return

PROTOCOL
Biomedical
Non-Exempt
Berkeley

Protocol # 2011-09-3652
Date Printed: 07/31/2018

-----------------------------------------------------------------------------------

**Protocol Title:** Measuring the benefits of sanitation, water quality, handwashing and nutrition interventions for improving health and development in rural Bangladesh

**Protocol Status:** APPROVED

**Date Submitted:** 10/26/2017

**Approval Period:** **10/30/2017-11/03/2018**

**Important Note:** **This Print View may not reflect all comments and contingencies for approval. Please check the comments section of the online protocol. Questions that appear to not have been answered may not have been required for this submission. Please see the system application for more details.**

*** * * Attached Document * * ***

| Document Name | Created Date |
| --- | --- |
| Prot PR-11063_ERC certificate on yearly review_1 Sep 2013.PDF | 10/26/2017 |

icddr,b

### Approval for continuation of research activity

This is to certify that icddr,b research protocol # PR-11063 titled "***A randomized controlled trial of the benefits of water sanitation hygiene plus nutrition intervention on child growth***": PI- Dr Leanne Unicomb of the Centre for Communicable Diseases (CCD) had been approved by Ethical Review Committee (ERC) on 9 April 2012.

The ERC undertakes annual/periodic review of all ERC-approved protocols for reappraisal. The ERC approval for implementation of any research protocol is not, however, affected unless any unanticipated problems involving risks to the study participants or any serious or continuing noncompliance of the ERC Guidelines are detected in the implementation of the study, during the review period.

The review undertaken as of 9 April 2013 to oversee the implementation of the above protocol reveals no Adverse Event (AE) or Serious Adverse Event (SAE) or unanticipated problems involving risks to the study participants or any serious or continuing noncompliance of the ERC Guidelines. Therefore, the ERC is pleased to **approve** the protocol for continuation of its activity for next one year starting **from 8 April 2013 to 9 April 2014**.

The continuing review application must be submitted to the IRB Secretariat for this study to continue beyond 9 April 2014. All necessary materials for continuing review must be reviewed with sufficient time for review and issuing continued approval before the expiration date. Failure to initiate a continuing review application in a timely fashion may result in discontinuation of study activities until approval can be renewed. Performing study activities, including data analysis, beyond the expiration date results in noncompliance of federal regulations.

Other terms and conditions for implementation of your research protocol, as contained in our memo dated 9 April 2012 according initial approval of the research protocol shall, however, remain unchanged.

Mamun

Professor Kazi Zulfiquer Mamun
***MBBS, M.Trop. Med, PhD.***
Chairperson
Ethical Review Committee of icddr,b
1 September 2013

GPO Box 128, Dhaka 1000, Bangladesh.
**For Packages :** 68, Shaheed Tajuddin Ahmed Sarani, Mohakhali, Dhaka 1212, Bangladesh.

PROTOCOL
Biomedical
Non-Exempt
Berkeley

Protocol # 2011-09-3652
Date Printed: 07/31/2018

---------------------------------------------------------------------------------

**Protocol Title:** Measuring the benefits of sanitation, water quality, handwashing and nutrition interventions for improving health and development in rural Bangladesh

**Protocol Status:** APPROVED

**Date Submitted:** 10/26/2017

**Approval Period:** 10/30/2017-11/03/2018

**Important Note:** **This Print View may not reflect all comments and contingencies for approval. Please check the comments section of the online protocol. Questions that appear to not have been answered may not have been required for this submission. Please see the system application for more details.**

*** * * Attached Document * * ***

| Document Name | Created Date |
| --- | --- |
| Wash Benefits Midline_Endline Instruments_131003.pdf | 10/26/2017 |

ICDDR,B

Wash Benefits

---

Midline and Endline Instruments

6/19/2013

**NOTES TO THE FIELD TEAM**

চার ভাগে উত্তর আসবে There is4 groups of respondents of interest to us:

| POPULATION | DEFINITION |
|---|---|
| 1. নিদ্দিষ্ট শিশু Target children | গবেষণায় আওতাধীন শিশু Children in utero at enrollment (primary study population) |
| 2. মা/প্রধান পরিচর্যাকারী Mothers | নিদ্দিষ্ট শিশুর মা/প্রধান পরিচর্যাকারী Mothers of target children |
| 3. ডায়রিয়া কোহট/অংশ Diarrhea cohort | ৩৬ মাস বয়সের নীচের শিশু যে এই খানা/ বাড়ী/কম্পাউন্ডের মধ্যে বাস করে Children < 36 months old at enrollment, living in study compounds |
| 4. পরজীবি কোহট/অংশ Parasite cohort | বেইজ লাইনে ১৮-২৭ মাস বয়সে শিশুর রক্ত,মলেরনমূনা সংগ্রহ করতে হবে যে এই খানা/বাড়ী/কম্পাউন্ডের মধ্যে বাস করেChildren 18 – 27 months old at baseline (blood spot, stool) |

**ভৌগলিক/অবস্থান পরিচিতি Geographical Identification**

নিচে উত্তরদাতার নির্দিষ্টকরন তথ্য গুরম্নত্বের সাথে পূরন করম্নন FILL THE FOLLOWING IDENTIFICATION INFORMATION FOR THE RESPONDENT VERY CAREFULLY.

| Module ID | WBM 40 | |
|---|---|---|
| | 4001. | এফ.আর.এ কোড (FRA code): ÿÿ |
| C.101 | 4002. | তথ্য সংগ্রহের তারিখ [Date of data collection]: __/__/____ |
| | dataid | \|__\|__\|__\|__\|__\|__\| |
| | 4003. | |
| | 4004. | |
| | 4005. | |
| | 4006. | জেলার নাম(DISTRICT NAME): |
| | 4007. | উপজেলার নাম (UPAZILA NAME): |
| | 4008. | ইউনিয়নের নাম [Union name _____________ |
| | 4009. | খানার ঠিকানা Household Address: |
| | 4010. | খানা প্রধানের নাম [Name of Household head]______________________________ |
| | 4011. | খানা প্রধানের বাবা/স্বামীর নাম [Father's/Husband's Name of HH head] ________________ |
| | 4012. | বাড়ির নাম [Bari Name]: __________________________________________ |
| | 4013. | গ্রামের নাম [Village Name]: _______________________________________ |
| | 4014. | |
| | 4015. | বাড়ী নাম্বার [Bari No. [1-8] |
| | 4016. | **HH ID:** [1-7] |
| | 4017. | |
| | 4018. | |
| | 4019. | |
| | 4019.a | **\|_____\|** এই বাড়ীতে ৩৬ মাসের নীচে কতজন শিশু আছে? How many children do you have < 36 months in this bari? |

| শিশুর প্রদত্ত তালিকা দেখুন এবং শিশুর উপস্থিতি উলেস্নখ করম্নন (Now look at the list provided to ensure the presence of the additional enrolled children in the bari) | | | | | | | |
|---|---|---|---|---|---|---|---|
| | | শিশুর পরিচয় (Who is this child?) | নাম (Name) | লিঙ্গ (Sex) | জন্ম তারিখ(Birth date) | জন্ম তারিখেরউৎস(Source of DOB )(C.104) | নির্বাচন করার কারণ (একাধিক উত্তর হতে পারে) Reason for enrollment (select all that apply) [**PDA** |

| | | | | | | | |
|---|---|---|---|---|---|---|---|
| | | | | | | | to store accordingly] |
| C.06. | 4020. | 1. নির্দিষ্ট শিশু (Target child)<br>2 .একই মায়ের অন্য শিশু(Another child of same mother)<br>3. একই বাড়ীর অন্য খানার শিশু (Child from the same bari)<br>4. একই খানার অন্য মায়ের শিশু(Another child of another mother in the same HH) | | 1 পুরুষ M<br>0 মহিলা F | দিন/মাস/বছর DD/MM/YY | 1 = কার্ড দ্বারা নিশ্চিত হয়েছিল (Confirmed DOB by valid vaccination/health card)<br>2 = মা/আত্মীয় দ্বারা নিশ্চিতহয়েছিল (Mother/Relative remembers DOB)<br>3 = উভয় (Both 1 & 2)<br>4 = ঘটনা দ্বারা নিশ্চিতহয়েছিল (Estimated DOB with 2 and event calendar) | ☐1 ডায়রিয়া Diarrhea<br>☐2 পায়খানার নমুণা Stool specimen<br>☐3 রক্তের নমুণা Blood specimen |
| C.07. | 4021. | 1. নির্দিষ্ট শিশু (Target child)<br>2 .একই মায়ের অন্য শিশু(Another child of same mother)<br>3. একই বাড়ীর অন্য খানার শিশু (Child from the same bari)<br>4. একই খানার অন্য মায়ের শিশু(Another child of another mother in the same HH) | | 1 পুরুষ M<br>0 মহিলা F | দিন/মাস/বছর DD/MM/YYY | 1 = কার্ড দ্বারা নিশ্চিত হয়েছিল (Confirmed DOB by valid vaccination/health card)<br>2 = মা/আত্মীয় দ্বারা নিশ্চিত হয়েছিল (Mother/Relative remembers DOB)<br>3 = উভয় (Both 1 & 2)<br>4 = ঘটনা দ্বারা নিশ্চিতহয়েছিল (Estimated DOB with 2 and event calendar) | ☐1 ডায়রিয়া Diarrhea<br>☐2 পায়খানার নমুণা Stool specimen<br>☐3 রক্তের নমুণা Blood specimen |
| C.08. | 4022. | 1. নির্দিষ্ট শিশু (Target child)<br>2 .একই মায়ের অন্য শিশু(Another child of same mother)<br>3. একই বাড়ীর অন্য খানার শিশু | | 1 পুরুষ M<br>0 মহিলা F | দিন/মাস/বছর DD/MM/YYY | 1 = কার্ড দ্বারা নিশ্চিত হয়েছিল (Confirmed DOB by valid vaccination/health card)<br>2 = মা/আত্মীয় দ্বারা নিশ্চিত হয়েছিল (Mother/Relative remembers DOB)<br>3 = উভয় (Both 1 | ☐1 ডায়রিয়া Diarrhea<br>☐2 পায়খানার নমুণা Stool specimen<br>☐3 রক্তের নমুণা Blood specimen |

| | | | | | | | |
|---|---|---|---|---|---|---|---|
| | | (Child from the same bari)<br>4. একই খানার অন্য মায়ের শিশু(Another child of another mother in the same HH) | | | | & 2)<br>4 = ঘটনা দ্বারা নিশ্চিত হয়েছিল (Estimated DOB with 2 and event calendar) | |
| **C.09.** | 4023. | 1. নির্দিষ্ট শিশু (Target child)<br>2 .একই মায়ের অন্য শিশু(Another child of same mother)<br>3. একই বাড়ীর অন্য খানার শিশু (Child from the same bari)<br>4. একই খানার অন্য মায়ের শিশু(Another child of another mother in the same HH) | | 1 পুরুষ M<br>0 মহিলা F | দিন/মাস/বছর DD/MM/YYY | 1 = কার্ড দ্বারা নিশ্চিত হয়েছিল (Confirmed DOB by valid vaccination/health card)<br>2 = মা/আত্মীয় দ্বারা নিশ্চিত হয়েছিল (Mother/Relative remembers DOB)<br>3 = উভয় (Both 1 & 2)<br>4 = ঘটনা দ্বারা নিশ্চিতহয়েছিল (Estimated DOB with 2 and event calendar) | ☐1 ডায়রিয়া Diarrhea<br>☐2 পায়খানার নমুণা Stool specimen<br>☐3 রক্তের নমুণা Blood specimen |
| **C.10.** | 4024. | 1. নির্দিষ্ট শিশু (Target child)<br>2 .একই মায়ের অন্য শিশু(Another child of same mother)<br>3. একই বাড়ীর অন্য খানার শিশু (Child from the same bari)<br>4. একই খানার অন্য মায়ের শিশু(Another child of another mother in the same HH) | | 1 পুরুষ M<br>0 মহিলা F | দিন/মাস/বছর DD/MM/YYY | 1 = কার্ড দ্বারা নিশ্চিত হয়েছিল (Confirmed DOB by valid vaccination/health card)<br>2 = মা/আত্মীয় দ্বারা নিশ্চিতহয়েছিল (Mother/Relative remembers DOB)<br>3 = উভয় (Both 1 & 2)<br>4 = ঘটনা দ্বারা নিশ্চিতহয়েছিল (Estimated DOB with 2 and event calendar) | ☐1 ডায়রিয়া Diarrhea<br>☐2 পায়খানার নমুণা Stool specimen<br>☐3 রক্তের নমুণা Blood specimen |

| | | |
|---|---|---|
| | 001 | উত্তরদাতার/ উত্তরদাতার খানার কোন সদস্যের কী কোন ফোন নাম্বার আছে?(Is there a phone available?) 1 Yes 0 No (Skip to 005) |
| C.11 | **002** | -----------প্রাথমিক ফোন নাম্বার (Primary phone number) |
| C.12 | 003 | এই ফোনে কি আপন াকে সরাসরি পাওয়া যাবে? (Does this phone number belong to you?) 1 Yes(Skip to 005) 0 No |
| C.13 | 004 | যদি আপন াকে পাওয়া না যায় তাহলে এই নাম্বারটি কার? IF NO, to whom does this number belong? (Specify relationship to respondent)<br>1 . স্বামী (HUSBAND)<br>2. শ্বশুর IN LAWS<br>3.পিতা(FATHER)<br>4.মা(MOTHER)<br>5.পরিবারের সদস্য(FAMILY MEMBER)<br>99.অন্যান্য (নাম ও উত্তরদাতার সাথে সম্পর্ক লিখুন)(OTHER) |
| | 004.a | দ্বিতীয় কোন ফোন নাম্বার আছে কী?(Is there a phone available?)<br>1 Yes 0 No (Skip to 008) |
| C.14 | 005 | -----------দ্বিতীয় ফোন নাম্বার Secondary phone number |
| C.15 | 006 | এই ফোনে কি আপন াকে সরাসরি পাওয়া যাবে? Does this phone number belong to you? 1 Yes(Skip to 008) 0 No |
| C.16 | 007 | যদি আপন াকে পাওয়া না যায় তাহলে এই নাম্বারটি কার? IF NO, to whom does this number belong?<br>1. স্বামী (HUSBAND)<br>2. শ্বশুর IN LAWS<br>3. পিতা(FATHER)<br>4 .মা(MOTHER)<br>5 .পরিবারের সদস্য(FAMILY MEMBER)<br>99 .অন্যান্য(নাম ও উত্তরদাতার সাথে সম্পর্ক লিখুন) (OTHER) |
| C.17 | 008 | এই খানাটির আশেপাশে কোন উলেস্নখযোগ্য কিছু আছে কি? (Location/landmark) |
| | 009 | **\|_____\| \|_____\|** সবচেয়ে ছোট বাচ্চার মা কত ক্লাস পর্যন্ত পড়ালেখা শেষ করেছেন ? (জানি না = ***99***) Education of mother of the youngest child *(Years of education completed, DK=99)* |
| | 010 | **\|_____\| \|_____\|** সবচেয়ে ছোট বাচ্চার বাবা কত ক্লাস পর্যন্ত পড়ালেখা শেষ করেছেন ? (জানি না = ***99***) Education of father of the youngest child *(Years of education completed, DK=99)* |
| | 011 | **\|_____\| \|_____\|** সবচেয়ে ছোট বাচ্চার বাবার প্রধান পেশা [Main occupation of father of the youngest child]<br>পেশা কোড Occupation Code:<br>1. কৃষক [Farmer/Cultivator]<br>2. ঘরে কাজ করে [Homemaker]<br>3. কৃষি শ্রমিক [Agri-labor]<br>4. শ্রমিক (কৃষি শ্রমিক ছাড়া) [Non-agri labor]<br>5. বেতনভুক্ত কর্মচারী (সরকারী/প্রাইভেট/এন.জি.ও) [Salaried job (Govt./Private/NGO)]<br>6. রাজমিস্ত্রী [Mason (*Rajmistri*)]<br>7. কাঠ মিস্ত্রী [Carpenter]<br>8. ভ্যান/রিকশা চালক [Van/Rickshaw puller]<br>9. জেলে [Fisherman]<br>10. নৌকা চালক/মাঝি [Boatman]<br>11. কর্মকার [Blacksmith]<br>12. স্বর্ণকার [Goldsmith]<br>13. কুমার/কুম্ভকার [Potter (soil smith)]<br>21. কুটীর শিল্প [Cottage industry]<br>22. পোলট্রি/ ব্যবসার জন্য পশু লালন-পালনকারী [Poultry /livestock rearer]<br>23. বৈদ্যুতিক মিস্ত্রী [Electrician]<br>24. হোমিওপ্যাথি ডাক্তার [Homeopath]<br>25. আধ্যাতিক চিকিৎসক/ কবিরাজ/ ওঝাঁ [Spiritual healer/kabiraj/ Ojha]<br>26. পেশাদার ডাক্তার/উকিল [Professional practitioner (Doctor/lawyer)]<br>27. ইমাম/ ধর্মযাজক [Imam/priest]<br>28. অবসরপ্রাপ্ত চাকুরীজীবী [Retired service holder]<br>29. ছাত্র [Student]<br>30. বেকার [Unemployed]<br>31. অক্ষম [Disabled]<br>32. কাজের লোক [Domestic maid / |

| | | |
|---|---|---|
| | | 14. মুচি [Shoe polish /maker]<br>15. দোকানদার [Shopkeeper]<br>16. ফেরিওয়ালা [Vendor (*Feriwala/howker*)]<br>17. ক্ষুদ্র ব্যবসায়ী (মূলধন <=১০০০০) [Petty trader, capital <=10000]<br>18. ব্যবসায়ী(মূলধন >১০০০০) [Business, capital >10000]<br>19. দর্জি [Tailor]<br>20. ড্রাইভার [Driver]<br>servant]<br>33. জমিদার (শস্য উৎপাদন অথবা অন্য কোন কাজে কৃষকদের জমি বর্গা দেয়) [Landlord (Provide land for farmers for sharecropping or others)]<br>34. বিদেশে থাকে [Staying abroad]<br>35. মৃত/নিখোঁজ [Died/untraced]<br>99. জানিনা [Don’t know]<br><br>77. অন্যান্য (বর্ণনা লিখুন) [Others (specify) |
| | 012 | **\|_____\| \|_____\| আপনার খানাতে/পরিবারে কত জন লোক প্রতিদিন একই হাড়িতে রান্না করে খান?** How many people in this house eat from the same cooking pot every day (খানা বলতে একই হাড়িতে রান্না করে খায় এমন সদস্যদের বুঝানো হয়েছে) |
| | 013 | **\|_____\| আপনার খানাতে ৩৬ মাসের নীচে কতজন শিশু আছে?** How many children do you have < 36 months do you have? |
| | 014 | **\|____\| এই বাড়ীতে ৩৬ মাসের নীচে কতজন শিশু আছে?** How many children do you have < 36 months in this bari? |
| | 014 | **\|_____\|এই বাড়ীতে কতগুলো খানা আছে?** How many HHs do you have in this bari? |

| A. এই বাড়ীর খানাগুলোর সেনসাস (সকল খানার) CENSUS OF HOUSEHOLDS ARE IN THIS COMPOUND (All households) | | | | | | |
|---|---|---|---|---|---|---|
| পড়ে শোনান READ | ঠিক আছে। আমি এখন আপনাকে সকল খানার সম্পর্কে আলাদাভাবে কিছু প্রশ্ন করব। প্রথমে আপনার খানা সম্পর্কে বলুন। Okay. Now I am going to ask you about each household separately. First, think about your own household. MARK “99” IF RESPONDENT DOESN’T KNOW THE ANSWER TO ANY OF THE FOLLOWING QUESTIONS. | | | | | |
| | A.1 | A.2 | A.3 | A.4 | A.5 | A.6 |
| খানার নাম্বার HH No. | খানা প্রধানের নাম কি? What is the name of the household head? FOR REFERENCE ONLY | এই খানাতে ৪০ বছরের উপরের বয়সের কতজন সদস্য আছে? How many adults over 40 years live in this household? | এই খানাতে (>১৮-৪০) বছরের বয়সের কতজন সদস্য আছে? How many young adults (>18-40 years) live in this household? | এই খানাতে (>৩-১৮) বছরের বয়সের কতজন সদস্য আছে?How many school-aged children (>3-18 years) live in this household? | এই খানাতে (০-৩) বছরের বয়সের কতজন সদস্য আছে?How many young children (0-3 years) live in this household? | প্রতিটি খানায় কতজন সদস্য আছে তা গণণা করুন এবং নিশ্চিত হোন CALCULATE THE TOTAL NUMBER OF PEOPLE IN EACH HH AND CONFIRM |
| 1 | | | | | | |
| 2 | | | | | | |

**Wash Benefit Module 1RESPONDENT IDENTIFICATION**

| মডিউল আইডিModule ID | WBM 1 | |
|---|---|---|
| | 101. | : What is your full name? প্রধান উত্তরদাতার নাম [Name of respondent: |
| | 102. | : **\|_____\| প্রধান উত্তরদাতার পরিচয়**[Status of main respondent]<br><br>1. প্রধান পরির্চযাকারী /মা [Mother of youngest child]<br>2. পরির্চযাকারী (পুরুষ) [Male caregiver]<br>3. পরির্চযাকারী (মহিলা) [Female caregiver] |
| | 103. | প্রধান উত্তরদাতার বয়স (বছরে)[Age of main respondent: *(in years)]* |
| | 104. | উত্তরদাতা কোন শর্তটা পুরণ করেছে?RESPONDENT MEETS WHICH ELIGIBILITY CRITERIA?<br>[1] গর্ভবতীPREGNANT<br><br>[2] ১৮-২৭ মাসের বাচ্চা আছেHAS CHILD 18-27 MONTHS OF AGE →SKIP to |

| | | |
|---|---|---|
| আমি আপনাকে আপনার গর্ভবতী হওয়া বিষয়ে কিছু ব্যক্তিগত প্রশ্ন এই গবেষনার জন্য করতে চাই।আমি আশা করি এ বিষয়ে আপনি কিছু মনে করবেন না এবং আমাদিগকে সহায়তা করবেন। I need to ask you some personal questions regarding your pregnancy because that is our project's eligibility criteria. I hope you do not mind and will cooperate with us. [see scripts of other studies] | | |
| C. 108 | 105. | উত্তরদাতার সর্বশেষ মাসিকের তারিখ (**What was the date of your last menstrual period?)**__/__/____<br>88 / 88 / 88 = সর্বশেষ গর্ভধারণের পর থেকে মাসিক না হলে (No menstruation since their last pregnancy)<br>99 = জানি না (Don't know / not sure) |
| C. 109 | 106. | \|__\| গর্ভধারণের পূর্নাঙ্গ মাস গণনা করুন( Record number of completed months)<br>99 = জানি না (Don't know / not sure) |
| C.110 | 107. | \|___\|গর্ভধারণের মসয়কালের উৎস(Source of Pregnancy Length)<br>1 = শুধুমাত্র মায়ের নির্ণয় (Estimated by mother only)<br>2 = মায়ের এবং স্বাস্থ্য সেবাদানকারীর নির্ণয় ,আলট্রাসাউন্ড ছাড়া(Estimated by mother and a health practitioner, no ultrasound (last prenatal visit)<br>3 = মা, স্বাস্থ্য সেবাদানকারী এবং আলট্রাসাউন্ড দ্বারা নির্ণয়Estimated by mother and health practitioner, using ultrasound (last prenatal visit) |
| | 108. | __/__/__ কখন আপনার শিশু জন্মাবে? (What will be the expected DOB of your child?)<br>__/__/__ |
| | 109. | নির্দিষ্ট শিশুর প্রকৃত বয়স (বেইজ লাইনে লাগবে না)Actual DOB of target child (skip for baseline)<br>__/__/___ |
| | 110. | Do you have any paln to go to your mother's house for delivery porpoise?<br>1. Yesহ্যাঁ<br>0. No না (Skip to module 0) |
| | 111. | আপনার মায়ের বাড়ী যে গ্রামে সেই গ্রামের নাম কি, যেখানে আপনি বাচ্চার জন্মগ্রহনের সময় যাবেন?<br>What is the name of your mother's (maternal) village, where you might go to give birth?<br>0. Same as this একই গ্রাম<br>1. গ্রামের নাম লিখুন______________ |
| আগামী কয়েক মাস আমাদের এই প্রকল্পের কিছু বিষয়ে আপনাকেকিছু তথ্য দেওয়ার জন্যফোনেযোগাযোগ রাখতে চাই । আপনার কি কোন ফোন আছে? যদি না থাকেতাহলে আপনার বাড়ীতে কারো বাপ্রতিবেশীর ফোন আছে কি? তার নাম্বার কত? At some point over the next several months, we may need to get in touch with you by phone to give you information about our project. Do you have a phone? If not, does someone in your compound have a phone? Maybe a neighbor? What is the number? | | |

**Wash Benefit Module 3 DEWORMING**

**Wash Benefit Module 2DIARRHEA AND ILLNESS SYMPTOMS (Children < 36 months)**

৩৬ মাসের নীচের যে সকল শিশু খানায়/বাড়ীতে বাস করে (বেইজ লাইলের জন্য) । Administer to: Children < 36 months living in a study compound at baseline. This may include the child in the household or in other households in the bari.

PDA TO OPEN CHILD ID FROM **DIARRHEA COHORT**

যদিউত্তরদাতার 0-36 মাসেরশিশু না থাকেতাহলে এখানেই শেষ করম্ননননংপ্রশ্ন যেতে হবে | IF study compound DOES NOT HAVE A CHILD 0-36 MONTHS → **SKIP TO END**

| উত্তরদাতা শিশুর প্রধান পরিচর্যাকারী Respondent: Child's primary caregiver | | | | | | |
|---|---|---|---|---|---|---|
| | WBM 2 | **আপনাকে ধন্যবাদ । আমি এখন** আপনাকে আপনার শিশুর স্বাস্থ্য বিষয়ে কিছু প্রশ্ন জিজ্ঞাসা করব। দয়া করে আপনি **আমাকে ঠিকমতো প্রত্যক প্রশ্নের জবাব দিবেন। যদি উত্তর না জানা থাকে তখন বলবেন জানি না। শিশুটির নাম ধরে শুরু করতে হবে।** Thank you. Now I am going to ask you some questions about the health of your [child/children]. Please answer each question as accurately as you can. If you don't know the answer to a question, say "I don't know." We will begin with [NAME OF CHILD 1]. | | | | |
| | | | **A** | **B** | **C** | **D** |
| | | **শিশুটির নাম বলুন এবং তার লক্ষণ গুলো ছিল কী?Did [NAME] have [SYMPTOM] :** | আজ **Today** ১=হ্যাঁ YES ০=না NO ৯৯৯=জানি না DK | গতকাল **Yesterday** ১=হ্যাঁ YES ০=না NO ৯৯৯=জানি না DK | গত পরশুদিন **Day before Yesterday** ১=হ্যাঁ YES ০=না NO ৯৯৯=জানি না DK | গত৭দিনে In **the last 7 days** (since this day last week) ১=হ্যাঁ YES ০=না NO ৯৯৯=জানি না না DK |
| **C.201** | 201. | জ্বর Fever | | | | |
| **C.202** | 202. | **পাতলা পায়খানা** Diarrhea | | | | |
| **C.203** | 203. | ২৪ ঘনটায় ৩ বার বা তার বেশী পায়খানা করেছে কিনা? 3 or more bowel movements in 24 hours | | | | |
| **C.204** | 204. | দৈনিক পায়খানায় সংখ্যা Number of bowl movements each day | | | | |
| **C.205** | 205. | পানির মত বা নরম পায়খানা? Watery or soft stool (unformed) | | | | |
| **C.206** | 206. | পায়খানায় রক্ত Blood in the stool | | | | |
| **C.207** | 207. | শরীর বা মাথায় চুলকানি, রেশ?Skin rash (anywhere on the body) | | | | |
| **C.208** | 208. | দীর্ঘ কাশি Constant cough | | | | |
| **C.209** | 209. | নাক বন্ধ/নাক দিয়ে পানি পড়া?Congestion / runny nose | | | | |
| **C.210** | 210. | শ্বাস নেওয়ার সময় শোঁ শোঁ করে আওয়াজ বা শ্বাসকষ্ট? (যে কোন ধরনের শ্বাসকষ্ট, তবে নাক বন্ধ থাকার কারণে শ্বাসকষ্ট হলে তা বাদ দিতে হবে) Panting / wheezing / difficulty breathing | | | | |
| **C.211** | 211. | দাগ, থেঁৎলে যাওয়া বা কাটা Bruising, scrapes or cuts | | | | |
| **C.212** | 212. | দাঁতের ব্যাথাToothache / teething | | | | |

#### Wash Benefits Module 4 Anthropometry

| | | |
|---|---|---|
| **C.401**<br>**FRA ID** | ## | ☐☐ |
| **C.402**<br>**Name of FRA** | Full Name | |
| **C.403**<br>**Is mother wearing heavy clothing during weight measurement?** | 1 = Light clothing<br>2 = Light clothing plus sweater<br>3 = Heavy clothing | ☐ |
| **C.404**<br>**Weight of Mother**<br>**Measurement #1** | Weight<br>(kg) | ☐☐.☐ |
| **C.405**<br>**Weight of Mother**<br>**Measurement #2** | Weight<br>(kg) | ☐☐.☐ |
| **C.406**<br>**Weight of Mother**<br>**Measurement #3**<br>(If difference between measures 1 & 2 is ≥ 0.1 kg) | Weight<br>(kg) | ☐☐.☐ |
| **C.407**<br>**Is child wearing clothing during weight measurement?** | 0 = No Clothes<br>1 = Only Shirt<br>2 = Only Pants<br>3 = Both Shirt & Pants | ☐ |
| **C.408**<br>**Weight of Mother + Child**<br>**Measurement #1** | Weight<br>(kg) | ☐☐.☐ |
| **C.409**<br>**Weight of Mother + Child**<br>**Measurement #2** | Weight<br>(kg) | ☐☐.☐ |

| | | |
|---|---|---|
| **C.410**<br>**Weight of Mother + Child**<br>**Measurement #3**<br>(If difference between measures 1 & 2 is ≥ 0.1 kg) | Weight<br>(kg) | □□.□ |
| **C.411 – C.413**<br>reserved for child weight measurement without mother (follow-up visits) | | |
| **C.414**<br>**Length of Child**<br>**Measurement #1** | Length<br>(cm) | □□□.□ |
| **C.415**<br>**Length of Child**<br>**Measurement #2** | Length<br>(cm) | □□□.□ |
| **C.416**<br>**Length of Child**<br>**Measurement #3**<br>(If difference between measures 1 & 2 is ≥ 0.5 cm) | Length<br>(cm) | □□□.□ |
| **C.417**<br>**Length Measurement Method** | Child was:<br><br>1 = lying (recumbent)<br>2 = standing | □ |
| **C.418**<br>**Head Circumference**<br>**Measurement #1** | Circumference<br>(cm) | □□.□ |
| **C.419**<br>**Head Circumference**<br>**Measurement #2** | Circumference<br>(cm) | □□.□ |
| **C.420**<br>**Head Circumference**<br>**Measurement #3**<br>(If difference between measures 1 & 2 is ≥ 0.5 cm) | Circumference<br>(cm) | □□.□ |

| | | |
|---|---|---|
| **C.421**<br>**Does the child have swollen feet (bi-pedal edema)?** | 1 = Yes (**>> Referral**)<br>2 = No | □ |

ওয়াস বেনিফিটস কমন মডিউল-৪

এনথ্রোপোমেট্রিক পরিমাপক

বেইজলাইনের সময় যে সকল খানায় <৩ মাসের শিশু আছে

| | | |
|---|---|---|
| **C.401**<br>এফ.আর.এ আইডি | ## | □□ |
| **C.402**<br>এফ.আর.এ-র নাম | পুরো নাম | |
| **C.403**<br>ওজন নেওয়ার সময় বাচ্চার মা ভারী কাপড় পড়েছিল কী? | 1 = হালকা কাপড়<br>2 = সোয়েটারসহ হালকা কাপড়<br>3 = ভারী কাপড় | □ |
| **C.404**<br>মায়ের ওজন<br>পরিমাপ#1 | ওজন(কেজি)<br>0 =ওজন নেওয়া সম্ভব হয় নি | □□.□ |
| **C.405**<br>মায়ের ওজন<br>পরিমাপ#2 | ওজন(কেজি)<br>0 =ওজন নেওয়া সম্ভব হয় নি | □□.□ |
| **C.406**<br>মায়ের ওজন<br>পরিমাপ#3<br>(যদি পরিমাপ#1 এবং পরিমাপ#2 এর মধ্যে ≥ **0.1** কেজিপার্থক্য থাকে) | ওজন(কেজি)<br>0 =ওজন নেওয়া সম্ভব হয় নি | □□.□ |
| **C.407**<br>ওজন নেওয়ার সময় বাচ্চা কোন কাপড় পড়েছিল কী? | 0 = কোন কাপড় নেই<br>1 = শুধুমাত্র শার্ট<br>2 = শুধুমাত্র প্যান্ট<br>3 = শার্ট এবং প্যান্ট উভয় | □ |

| | | |
|---|---|---|
| **C.408**<br>মা+বাচ্চার ওজন<br>পরিমাপ#1 | ওজন(কেজি)<br><br>0 =ওজন নেওয়া সম্ভব হয় নি | ☐☐.☐ |
| **C.409**<br>মা+বাচ্চার ওজন<br>পরিমাপ#2 | ওজন(কেজি)<br><br>0 =ওজন নেওয়া সম্ভব হয় নি | ☐☐.☐ |
| **C.410**<br>মা+বাচ্চার ওজন<br>পরিমাপ#3<br>(যদি পরিমাপ#1 এবং পরিমাপ#2 এর মধ্যে ≥ **0.1** কেজিপার্থক্য থাকে) | ওজন(কেজি)<br><br>0 =ওজন নেওয়া সম্ভব হয় নি | ☐☐.☐ |
| **C.411 – C.413**<br>reserved for child weight measurement without mother (follow-up visits) | | |
| **C.414**<br>বাচ্চার উচ্চতা<br>পরিমাপ#1 | উচ্চতা(সেমি)<br><br>0=উচ্চতা নেওয়া সম্ভব হয়নি | ☐☐☐.☐ |
| **C.415**<br>বাচ্চার উচ্চতা<br>পরিমাপ#2 | উচ্চতা(সেমি)<br><br>0=উচ্চতা নেওয়া সম্ভব হয়নি | ☐☐☐.☐ |
| **C.416**<br>বাচ্চার উচ্চতা<br>পরিমাপ#3<br>(যদি পরিমাপ#1 এবং পরিমাপ#2 এর মধ্যে ≥ **0.5** সেমি.পার্থক্য থাকে) | উচ্চতা(সেমি)<br><br>0=উচ্চতা নেওয়া সম্ভব হয়নি | ☐☐☐.☐ |
| **C.417**<br>উচ্চতা পরিমাপক পদ্ধতি | 1 = শুয়ে মাপা হয়েছে<br>2 = দাড়িঁয়ে মাপা হয়েছে | ☐ |
| **C.418**<br>মাথার পরিধি<br>পরিমাপ#1 | পরিধি(সেমি)<br><br>0 =মাথার পরিধি নেওয়া সম্ভব হয়নি | ☐☐.☐ |

| | | |
|---|---|---|
| **C.419**<br>মাথার পরিধি<br>পরিমাপ#2 | পরিধি(সেমি)<br><br>0 =মাথার পরিধি নেওয়া সম্ভব হয়নি | ☐☐.☐ |
| **C.420**<br>মাথার পরিধি<br>পরিমাপ#3<br>(যদি পরিমাপ#1 এবং পরিমাপ#2 এর মধ্যে ≥ **0.5** সেমি.পার্থক্য থাকে) | পরিধি(সেমি)<br><br>0 =মাথার পরিধি নেওয়া সম্ভব হয়নি | ☐☐.☐ |
| **C.421**<br>**Does the child have swollen feet (bi-pedal edema)?** | 1 = Yes (**>> Referral**)<br>2 = No | ☐ |

**Wash Benefit Module 5 Vaccination**

**Vaccination** (টিকাদান)

V101. ☐ Do you have a card where [NAME'S] vaccinations are written down? [*If YES*: May I see it please?] (আপনার শিশুর (নাম বলুন) কোন টিকা কার্ড আছে কী? হ্যাঁ হলে বলুন: আমি কী টিকা কার্ডটি দেখতে পারি?)

1 Yes, seen (হ্যাঁ, দেখাতে পেরেছে) **(Skip to V122)** (**V122** নং প্রশ্নে চলে যান)

2 Yes, not seen (হ্যাঁ, দেখাতে পারে নাই)

3 No card (কোন টিকা কার্ড নাই)

V102. ☐ Did [NAME] ever receive any vaccinations to prevent him/her from getting diseases, including vaccinations received in a national immunization campaign? (আপনার শিশু [নাম বলুন] রোগ থেকে রৰা পাওয়ার জন্য কখনও কোন টিকা বা জাতীয় টিকা দিবসে কোন টিকা নিয়েছিল কী?)

1 Yes (হ্যাঁ)

2 No (না) (Skip toN201)

99 Don't know / not sure (জানি না/নিশ্চিত না) (Skip toN201)

V103. *For children without a vaccination card* (যে শিশুর কোন টিকা কার্ড নেই)

Please tell me if [NAME] received any of the following vaccinations: (আপনার শিশু [নাম বলুন] নিম্নলিখিত কোন টিকা নিয়ে থাকলে আমাকে তা বলুন)

| | | |
|---|---|---|
| 1 | BCG vaccination against tuberculosis that is an injection in the arm or shoulder that usually causes a scar? (যৰা প্রতিরোধের জন্য বিসিজি-র টিকা দেওয়ার কারণে বাহু বা কাধে ৰত চিহ্ন আছে কী?) | ☐ 1. Yes (হ্যাঁ)<br>☐ 0. No (না)<br>☐ 88. N/A (প্রযোজ্য নয়)<br>☐ 99. Don't Know (জানি না) |
| 2 | Polio vaccine, that is, (pink) drops in the mouth? (মুখে কোন পোলিও টিকার [ গোলাপী রং] ফোঁটা দেওয়া হয়েছিল কী?) | ☐ 1. Yes (হ্যাঁ)<br>☐ 0. No (না)১২১.৪ নং প্রশ্নে যান (skip to 121.4)<br>☐ 88. N/A (প্রযোজ্য নয়) ১২১.৪নং প্রশ্নে যান (skip to 121.4)☐ 99. Don't Know (জানি না)১২১.৪ নং প্রশ্নে যান (skip to 121.4) |

| | | |
|---|---|---|
| 3 | How many times was the polio vaccine received? (কতবার পোলিও টিকা দেওয়া হয়েছিল?) | ☐ 9 = DK (জানি না) |
| 4 | A Penta (DPT + Hib vaccination + Hepatitis-B vaccination ) vaccination, that is, an injection given in the thigh or buttocks, sometimes at the same time as the polio drops? (উরম্ন বা পাছাতে কোন ডিপিটি-র টিকা দেওয়া হয়েছিল কী, যা অনেক সময় পোলিও টিকার সাথে দেওয়া হয়?) | ☐ 1. Yes (হ্যাঁ)<br>☐ 0. No (না) ১২১.৬ নং প্রশ্নে যান (skip to 121.6)<br>☐ 88. N/A (প্রযোজ্য নয়) ১২১.৬ নং প্রশ্নে যান (skip to121.6)<br>☐ 99. Don't Know (জানি না) ১২১.৬ নং প্রশ্নে যান (skip to121.6) |
| 5 | How many times was the Penta vaccine received? (কতবার ডিপিটি-র টিকা দেওয়া হয়েছিল?) | ☐ 9 = DK(জানি না) |
| 6. | A MR (measles-rubella) injection, that is, a shot in the arm at the age of 9 months or older to prevent him/her from getting measles and rubella? (হাম প্রতিরোধের জন্য ৯ মাস বয়সে হামের টিকা বাহুতে দেওয়া হয়েছে কী?) | ☐ 1. Yes (হ্যাঁ)<br>☐ 0. No (না)<br>☐ 88. N/A (প্রযোজ্য নয়)<br>☐ 99. Don't Know (জানি না) |
| 7 | A measles injection, that is, a shot in the arm at the age of 15 months or older to prevent him/her from getting measles? (হাম প্রতিরোধের জন্য ১৫ মাস বয়সে হামের টিকা বাহুতে দেওয়া হয়েছে কী?) | ☐ 1. Yes (হ্যাঁ)<br>☐ 0. No (না)<br>☐ 88. N/A (প্রযোজ্য নয়)<br>☐ 99. Don't Know (জানি না) |
| 11 | Within the last six months, has [NAME] received a vitamin A dose (like this / any of these)?<br>SHOW COMMON TYPES OF AMPULES/CAPSULES/SYRUPS (গত ছয় মাসের মধ্যে শিশুটি (নাম বলুন) এই রকম/অনেকটা এই রকমের কোন 'ভিটামিন এ' ক্যাপসুল খেয়েছিল কী? (ভিটামিন এ ক্যাপসুল দেখান) | ☐ 1. Yes (হ্যাঁ)<br>☐ 0. No (না)<br>☐ 88. N/A (প্রযোজ্য নয়)<br>☐ 99. Don't Know (জানি না) |

V104. *For children with a vaccination card:* (যে শিশুর টিকা কার্ড আছে)

Copy the vaccination date for each vaccine from the card. (টিকা কার্ড থেকে প্রতিটি টিকা প্রদানের তারিখ লিখুন)

Record "**88**" in the DAY column if card shows that a vaccination was given, but no date is was recorded. (যদি টিকা কার্ডে উলেস্নখ থাকে টিকা দিয়েছে কিন্তু কোন তারিখ না থাকলে দিনের কলামে [**DAY column**]"**88**"রেকর্ড করম্নন)

Record "**99**" in the DAY column if vaccination not given. (টিকা না দিলে দিনের কলামে [**DAY column**]"**99**"রেকর্ড করম্নন )

| | | DAY | MONTH | YEAR |
|---|---|---|---|---|
| 1 | BCG | I__I__I | I__I__I | I__I__I |
| 2 | POLIO 0 | I__I__I | I__I__I | I__I__I |
| 3 | POLIO 1 | I__I__I | I__I__I | I__I__I |
| 4 | POLIO 2 | I__I__I | I__I__I | I__I__I |
| 5 | POLIO 3 | I__I__I | I__I__I | I__I__I |
| 6 | Penta 1 | I__I__I | I__I__I | I__I__I |
| 7 | Penta 2 | I__I__I | I__I__I | I__I__I |
| 8 | Penta 3 | I__I__I | I__I__I | I__I__I |
| 9 | MR | I__I__I | I__I__I | I__I__I |
| 10 | MEASLES | I__I__I | I__I__I | I__I__I |

| | | | | |
|---|---|---|---|---|
| 11 | VITAMIN A (with measles vaccine) | I__I__I | I__I__I | I__I__I |

**Wash Benefit Module 6 CHILD FOOD FREQUENCY**

Administer to:Target Children(নির্দ্দিষ্ট শিশুর জন্য প্রযোজ্য)

Respondent: Child'sprimarycaregiver.( উত্তরদাতা শিশুর প্রধান পরিচর্চাকারী)

প্রধান পরিচর্চাকারী তিনিই হবেন যিনি সবচেয়ে বেশীসময় শিশুর জন্য ব্যয় করেন। সাধারনতঃ মা-ই প্রধান পরিচর্চাকারী।(The primary caregiver is the person that spends the most timewith the child.This is often the mother.)

**ভূমিকা**

আমি এখন আপনার শিশুর (নাম) খাদ্যগ্রহণ সংক্রান্ত বিষয়ে কিছু প্রশ্ন করতে চাই। প্রথমে আমার জানা দরকার যে গতকাল এবং গত ৭ দিনের মধ্যে শিশুর (নাম) খাদ্যগ্রহণ সম্পর্কে বলতে পারবেন।

**Introduction**Now I would like to ask you some questions about feeding [NAME]. First I need to know if you will be able to tell me about feeding [NAME] yesterday and over the last week.

**C.601.**

আপনি কি জানেন শিশু (নাম) গতকাল কি কি খেয়েছে?

Do you know what [NAME] consumed yesterday?

☐ 1 হ্যাঁ (৬০৩ নং প্রশ্নে যান) Yes [skip C.603]

☐ 2 না No

**C.602.**

যদি না হয় তবে এমন কেউ কি আছে যে জানে শিশু কি খেয়েছিল এবং যিনি আমাদের প্রশ্নসমূহের উত্তর দিয়ে সহায়তা করবে?

*If No,* Is there someone else who knows what the child ate, who can sit with us today and help answer questions?

☐ 1 হ্যাঁ Yes

☐ 2 নাNo

**C.603.**

শিশুর সাথে উত্তরদাতার সম্পর্ক

Respondent relationship to the participating child

☐ 1 মা Mother

☐ 2 বাবা Father

☐ 3 বোন Sister

☐ 77 অন্যান্য (নির্দ্দিষ্ট করুন) ___________________

Other (specify) _________________

শিশু যে খাবার খেয়েছে যেমন বুকের দুধ, তরল খাবার এবং অন্যান্য খাবার

**Breastfeeding, Liquids, and Foods Eaten by the Child**

**C.604.**

জন্মের কতক্ষণ পর শিশুকে (নাম) বুকের দুধ দিয়েছিলেন? **(উত্তর গুলো পড়ে শোনান)**

How long after the birth did you first put [name] to the breast?

- ☐ 0 প্রথম আধা ঘন্টার মধ্যে within the first half hour
- ☐ 1 প্রথম এক ঘন্টার মধ্যে Within the first hour
- ☐ 2 এক ঘন্টার বেশী কিন্তু ২৪ ঘন্টার মধ্যে More than one hour but less than 24 hours
- ☐ 3 ২৪ ঘন্টার বেশী More than 24 hours
- ☐ 88 কখনো না/প্রযোজ্য নয় (৬০৮ নং প্রশ্নে যান) Never breastfed / Not Applicable (**Skip to C.608**)
- ☐ 99 জানি না/ মনে নেই Don't know / not sure

**C.604a**

জন্মের পরপরই শিশুকে (নাম) শাল দুধ দিয়ে কি বুকের দুধ খাওয়ানো শুরু করেছিলেন?

When [NAME] was born, did you start breastfeeding him/her immediately with the very first breast milk (colostrum)?

- ☐ 1 হ্যাঁYes
- ☐ 2 না No

**C.605.**

শিশুটি কি এখনো বুকের দুধ পান করছে, নাকি সে সম্পূর্নভাবে বারতি খাবার খাচ্ছে?( যদি শিশু বুকের দুধ খায় তাহলে ১ কোড করম্নন)

Is the baby still breastfeeding, or is he/she completely weaned?

- ☐ 1 এখনো বুকের দুধ পান করে ( ৬০৭ নং প্রশ্নে যান) **Still**, breastfeeding (**Skip to C.607)**
- ☐ 2 বাড়তি খাবার খাচ্ছে. কোন বুকের দুধ পাচ্ছে না (weaned, not receiving any breast milk)

**C.606.**

শিশুটির (নাম) যখন সর্বশেষ বুকের দুধ পান করেছিল,তখন তার বয়স কত ছিল ?

How old was [NAME] the last time he/she was breastfed?

৯৯= জানি না / নিশ্চিত না 99 Don't know / not sure

☐☐ দিন

☐☐ মাস

☐☐

**C.607.**

এখন আমাকে বলুন শিশুটি (নাম) গতকাল কতবার বুকের দুধ পান করেছে ?

Now I would like you to tell me how many times [NAME] breastfed yesterday.

☐☐বার TIMES

**C. 607a**

বুকের দুধ ছাড়া শিশুকে (নাম)সর্বপ্রথম কি খাওয়ানো হয়েছে ?

What was the very first thing that [Name] consumed apart from breast milk?

☐১মধু Honey

☐২ চিনির পানি Sugary water

☐৩ তাল মিছরি Tal misree ( Crystalline sugar cube from palm sugar)

☐৪ শুধু পানি   Plain water

☐৫ কিছুই না ( ৬০৭ সি নং প্রশ্নে যান)   Nothing (**Skip to C.60 7c)**

☐৭৭   অন্যান্য (নির্দ্দিষ্ট করুন)   _________________ others (specify) _________________

**C. 607b**

শিশুটি কোন বয়সে এই খাবার খেয়েছিল ?

At what age did [Name] consume this item?

☐☐**দিন**[যদি মা দিন বলতে পারে ]DAYS [if mother mentioned days]অথবা   OR

☐☐মাস [যদি মা মাস বলতে পারে ]   MONTHS [if mother mentioned months]

**C. 607c**

গত সপ্তাহে কি কোন দিন শিশুর (নাম) ক্ষুধামন্দা হয়েছিল?

In the past week, were there any days that [NAME] lost his/her appetite?

| 1. উত্তর **Response** | 2. কতদিন **How many days?** |
|---|---|
| ☐ ১ হ্যাঁ Yes<br><br>☐ ২ না (৬০৮ নং প্রশ্নে যান ) No [skip to 608]<br><br>☐ ৯৯ জানি না (৬০৮ নং প্রশ্নে যান )<br><br>99 Don't Know [skip to 608] | ☐দিন Days |

এখন গতকাল সারাদিন আর সারারাতে (শেষ ২৪ ঘন্টা, সাক্ষাৎকারের সময় থেকে) আপনার শিশু [নাম] কি কি তরল খাবার খেয়েছে আমি সে সম্পর্কে কিছু জানতে চাইবো। আপনার শিশু যে তরল খাবার খেয়েছে, আমি সে সম্পর্কে জানতে চাই, তা অন্য যে কোন খাবারের সাথে মিশিয়ে হোক না কেন ।

তালিকা থেকে প্রশ্নগুলো পড়ুন এবং সঠিক ঘরে টিক চিহ্ন দিন

Now I would like to ask you about liquids that [NAME] may have had yesterday during the day or at night (last 24hours, starting from the time interview). I am interested in whether your child had the item even if it was combined with other foods.

*For each item on the list, read the question below and tick the appropriate box.*

**C.608.**

কোন পানীয় পান করেছে কিনা? (তালিকা থেকে প্রত্যেকটি প্রশ্ন পড়ে শোনান )

Did [NAME] drink/have any [ITEM FROM LIST]?   *Read question 14 times, once for each item*

| খাবার এর নাম **Food Item** | এই পানীয় টা খেয়েছে/ পান করেছে কিনা?<br><br>**Drink / take / eat this item?** | গতকাল দিনে বা রাতে কত বার (নাম) এই খাবারটি খেয়েছে/ পান করেছে ? (তালিকা থেকে)<br><br>**How many times yesterday during the day or night did [NAME] consume [ITEM FROM LIST]?** |
|---|---|---|
| 1. পানি Water? | ☐ ১ হ্যাঁ Yes | ☐☐বার Times |

| | | | |
|---|---|---|---|
| | | ☐ ২ না No<br>☐ ৯৯ জানি না<br>99 Don't Know | |
| 2. | চিনির পানি Sugar water? | ☐ ১ হ্যাঁ Yes<br>☐ ২ না No<br>☐ ৯৯ জানি না<br>99 Don't Know | ☐☐বার Times |
| 3. | প্রাণিজ দুধ, কৌটার দুধ, গুড়া দুধ?<br>Milk, including any animal fresh milk, milk in tin or box, or powdered milk? | ☐ ১ হ্যাঁ Yes<br>☐ ২ না No<br>☐ ৯৯ জানি না<br>99 Don't Know | ☐☐বার Times |
| 4. | শিশু খাদ্য যেমন ন্যান, ল্যাকটোজেন, বায়োমিল অথবা মাই বয়? Infant formula such as NAN or Lactogen or Biomeal or My Boy? | ☐ ১ হ্যাঁ Yes<br>☐ ২ না No<br>☐ ৯৯ জানি না<br>99 Don't Know | ☐☐বার Times |
| 5. | দুধ চা? Tea made with milk? | ☐ ১ হ্যাঁ Yes<br>☐ ২ না No<br>☐ ৯৯ জানি না<br>99 Don't Know | ☐☐বার Times |
| 6. | রং চা? Tea made without milk? | ☐ ১ হ্যাঁ Yes<br>☐ ২ না No<br>☐ ৯৯ জানি না<br>99 Don't Know | ☐☐বার Times |
| 7. | দই? Yogurt? | ☐ ১ হ্যাঁ Yes<br>☐ ২ না No<br>☐ ৯৯ জানি না<br>99 Don't Know | ☐☐বার Times |
| 8. | দুধ বা পানি দিয়ে ফুটানো শস্য জাতীয় খাবার যেমন সুজি, চালের গুড়া ইত্যাদি ?<br>Thin Porridge or Barley? | ☐ ১ হ্যাঁ Yes<br>☐ ২ না No<br>☐ ৯৯ জানি না<br>99 Don't Know | ☐☐বার Times |
| 9. | ফলের রস? Fruit juice | ☐ ১ হ্যাঁ Yes<br>☐ ২ না No<br>☐ ৯৯ জানি না<br>99 Don't Know | ☐☐বার Times |
| 10. | পানির মত পাতলা স্যুপ [কোন শক্ত উপাদান ছাড়া]<br>Clear broth [soup without solid ingredient] | ☐ ১ হ্যাঁ Yes<br>☐ ২ না No<br>☐ ৯৯ জানি না<br>99 Don't Know | ☐☐বার Times |
| 11. | ডাবের পানি (অন্য কিছুর সাথে না মিশিয়ে)<br>Coconut water (plain) | ☐ ১ হ্যাঁ Yes<br>☐ ২ না No<br>☐ ৯৯ জানি না<br>99 Don't Know | ☐☐বার Times |
| 12. | হরলিকসবা অন্য কোন সম্পুরক পানিয় Horlicks (or any other | ☐ ১ হ্যাঁ Yes | ☐☐বার Times |

| | |
|---|---|
| supplementary drinks) | ☐ ২ না No<br>☐ ৯৯ জানি না<br>99 Don't Know |

এর পর গতকাল সারাদিন আর রাতে (শেষ ২৪ ঘনটা, সাক্ষাৎকারের সময় থেকে) আপনার শিশু [নাম]কি কি খেয়েছে আমি সে সম্পর্কে কিছু প্রশ্ন জিঙ্গাসা করবো। আপনার শিশু যা কিছু খেয়েছে আমি সে সম্পর্কে সব কিছুই জানতে চাই, তা বাসায় বা অন্য যে কোন জায়গায় হোক না কেন ।

নির্দেশনার জন্য আলাদা পৃষ্ঠা ব্যবহার করুন,যা মা কেমনে করতে সাহায্য করবে। মাকে বলতে দিন শিশুটি কি খেয়েছিল। শিশু টি যা খেয়েছিল , প্রতিটি খাবার ( অথবা উপাদান ) এ গোল চিহ্ন দিন এবং হ্যাঁ তে টিক চিহ্ন দিন

Next I would like to ask you some questions about the foods that [NAME] ate yesterday during the day or at night (last 24hours, starting from the time interview). I would like to know everything that [NAME] ate, whether at home or someplace else.

*Use the separate page of instructions, with questions to help the mother remember. Do not read the list below. Let the mother tell you what the child ate. Circle each food (or ingredient) that the child ate, and tick "[√] 1 Yes" for that food group.*

<u>নিম্নের তালিকাটি পড়ে শুনাবেন না</u>

**C.609.**

| | | |
|---|---|---|
| | শিশু গতকাল যে খাবার/খাদ্য তৈরীর উপাদান খেয়েছে(ঝোল অথবা সিদ্ধ যা ই হোক না কেন)<br>Foods/ingredients in recipes (may be in a sauce or porridge) eaten by the child yesterday | |
| 1 | খিচুড়ী Khichuri<br>( খিচুড়ী কি কি দিয়ে রান্না হয়েছে জেনে নিয়ে ফুড গ্রপ অনুযায়ী নীচে কোড করুন)<br>(tick 'yes' and ask for the ingredients and code below according to food group) | ☐ ১ হ্যাঁ Yes<br>☐২ না No |
| 2 | পানিতে বা দুধে সিদ্ধ করা শস্য জাতীয় খাবার যেমন: সুজি,ভাত, রুটি, নুডুলস, অন্যান্য শস্য জাতীয় দানাদার খাদ্য<br>Porridge Rice Bread/ Roti Noodles Other foods made by grain | ☐ ১ হ্যাঁ Yes<br>☐২ না No |
| 3 | মিষ্টি কুমড়া, গাজর, অন্যান্য হলুদ সবজী<br>Pumpkin Carrots Other yellow vegetable | ☐ ১ হ্যাঁ Yes<br>☐২ না No |
| 4 | আলু সাদা মিষ্টি আলু<br>Potato White sweet potato | ☐ ১ হ্যাঁ Yes<br>☐২ নার No |
| 5 | মিষ্টি কুমড়া শাক, সরিষা শাক, মটর শুটি শাক, পুই শাক, অন্যান্য গাঢ় সবুজ শাক<br>Pumpkin Mustard Bean Pigeon pea/ Motorshuti Other dark green<br>leaves leaves leaves leaves<br>leaves | ☐ ১ হ্যাঁ Yes<br>☐২ না No |
| 6 | পাকা আম , পাকা পেঁপেঁ<br>Ripe mango Ripe papaya | ☐ ১ হ্যাঁ Yes<br>☐২ না No |
| 7 | কলা, আনারস, পেয়ারা, আপেল, আঙ্গুর, কমলা অন্যান্য ফল<br>Banana Pineapple Guava Apple Grape Orange Other fruit | ☐ ১ হ্যাঁ Yes<br>☐২ না No |
| 8 | টমেটো, পিঁয়াজ , মাশরুম, ঢেড়স, তাজা সীম/মটর শুটি অন্যান্য সবজি<br>Tomato Onion Mushroom Lady's finger Fresh bean / Motorshuti other Vegetable | ☐ ১ হ্যাঁ Yes<br>☐২ না No |
| 9 | কলিজা গিলা<br>Liver Kidney Heart | ☐ ১ হ্যাঁ Yes<br>☐২ না No |
| 10 | যে কোন মাংস ,পশু পাখিসহ<br>Any type of meat / flesh, including from birds and animals | ☐ ১ হ্যাঁ Yes |

| | | | |
|---|---|---|---|
| | | ☐২ না | No |
| 11 | যে কোন ধরণের ডিম<br>Any type of egg | ☐ ১ হ্যাঁ<br>☐২ না | Yes<br>No |
| 12 | তাজা মাছ, শুটকি মাছ, অন্যান্য মাছ / সামুদ্রিক খাবার<br>Fresh fish Dried fish Other fish / seafood | ☐ ১ হ্যাঁ<br>☐২ না | Yes<br>No |
| 13 | সীম,ডাল,সয়া, চীনাবাদাম, কেশর, ভারীচীনাবাদাম অন্যান্য যে কোন ডাল বা বাদাম জাতীয় খাদ্য<br>Beans Peas / Soya Groundnut Cashew Pounded Any other<br>Lentils<br>groundnut legume or nut | ☐ ১ হ্যাঁ<br>☐২ না | Yes<br>No |
| 14 | পনির দই অন্যান্য দুধ জাতীয় খাদ্য<br>Cheese Yogurt Other milk products | ☐ ১ হ্যাঁ<br>☐২ না | Yes<br>No |
| 15 | উদ্ভিজ্জ তেল (ডালডা) পশুর চর্বি এক ধরনের মাখন<br>Vegetable oil Animal fat Margarine | ☐ ১ হ্যাঁ<br>☐২ না | Yes<br>No |
| 16 | চকলেট মিষ্টি/ ক্যান্ডি পিঠা মিষ্টি বিস্কুট<br>Chocolate Sweets / candies Cake Cookies / sweet biscuits | ☐ ১ হ্যাঁ<br>☐২ না | Yes<br>No |
| 17 | স্বাদ বৃদ্ধি কারক রসুন মসলা<br>Seasonings Garlic Spices | ☐ ১ হ্যাঁ<br>☐২ না | Yes<br>No |
| 18 | চিংড়ি, কাকড়া<br>Prawns Crab | ☐ ১ হ্যাঁ<br>☐২ না | Yes<br>No |
| 19 | যদি খাদ্য তালিকায় না থাকে তাহলে নিচে খাবারের নাম লিখুন।<br>*If not on list above, write food(s) here and at bottom* | ☐ ১ হ্যাঁ<br>☐২ না | Yes<br>No |

**C.610.**

আপনি গতকাল সারাদিন আর রাতে (শেষ ২৪ ঘন্টা, সাক্ষাৎকারের সময় থেকে) আপনার শিশু [নাম] কি কি খেয়েছে তা বললেন [পিছনের পৃষ্ঠার গোল করা খাবারে নাম গুলো পড়ুন] এছাড়া আপনার শিশু [নাম] নাস্তাসহ আর কি কিছু খেয়েছে ?

You mentioned that [NAME] ate [read back circled foods on previous page] yesterday during the day or at night (last 24hours, starting from the time interview).

Did [NAME] have any other food at all, including snacks?

☐১ হ্যাঁ Yes

☐ ২ না No

যদি হ্যা হয়, তাহলে পূর্বের পৃষ্ঠার প্রশ্ন গুলো আবার করুন এবং গোল চিহ্ন দিন । মনে করা শেষ হলে, যদি সেই দলের কোন খাবার বা খাদ্য উপাদানে গোল করা না হয় তাহলে না তে টিক চিহ্ন দিন।

যদি C.609এবং C.610 এর সব উত্তর না হয় তাহলে C.611 নং প্রশ্নে যান ।

*If "yes", use the same probing questions and circle on the list on previous page.* ***At the end of the recall tick "[√] 2 No" if no food or ingredient is circled for that group.***

If C.609 and C.610 are all 'No' skip to C.611

**C. 610a**

গতকাল সারাদিন আর সারারাতে (শেষ ২৪ ঘন টা, সাক্ষাৎকারের সময় থেকে) আপনার শিশু [নাম] কতবার শক্ত, আধা শক্ত আথবা নরম খাবার খেয়েছে তরল খাবার বাদে?

নোট: সকল ধরনের দুধ বা পানি দিয়ে ফুটানো শস্য জাতীয় খাবার ।

How many times did [NAME] eat solid, semi-solid, or soft foods other than liquids yesterday during the day or night (last 24hours, starting from the time interview)? *Note: include all forms of porridge, including thin porridge.*

☐☐বার Times

আমি এখন আপনাকে আপনার শিশু [নাম] গত ৭দিনে [সাক্ষ্যাৎকারের দিন থেকে] কি কি খাবার খেয়েছে সে বিষয়ে কিছু প্রশ্ন করতে চাই।প্রতিটি খাবারএর কথা আমি জানতে চাইবো, গত ৭দিনের মধ্যে কতদিন আপনার শিশু নিম্নলিখিত খাবার গুলো খেয়েছেন তা বলুন।

Now I would like to ask you some questions about foods [NAME] ate in the last 7 days, since last [INTERVIEW DAY]. For each food I ask about, please tell me how many days in the last 7 days you think the child ate that food.

যদি শিশুটি (নাম) খাবার খেয়ে থাকে, এমনকি এটি যদি অন্যান্য খাবারের সংঙ্গে সংযুক্ত থাকে, উদাহরণস্বরূপ,যদি শিশুটি (নাম) মুরগির মাংসের ঝোল খায় যা পিয়াজ, টমেটো, মুরগির মাংস দিয়ে তৈরী তাহলে আমি যখন আপনাকে প্রশ্ন করবো মাংস সম্পর্কে তখন আপনি "হ্যাঁ" বলবেন আবার যখন আমি আপনাকে প্রশ্ন করবো সবজি সম্পর্কে তখন আপনি আবার "হ্যাঁ" বলবেন যদি আপনার শিশু (নাম) দুটো খাবার ইখেয়ে থাকে, কিন্তু যদি মুরগির মাংস আর সবজি কোনটাই না খেয়ে থাকে তাহলে "হ্যাঁ" বলবেন না কারণ সে এগুলা খায় নি।

I would like to know if [NAME] had the food, even if it was combined with other foods. For example, if [NAME] ate a sauce or relish made with chicken, onions, and tomatoes, you should say "yes" when I ask about meat, and again "yes" when I ask about vegetables. However, if [NAME] only had the broth, not the chicken or vegetables, do not say "yes" because they did not eat it.

তালিকার প্রতিটি খাবারের জন্য, প্রশ্নটি পরুন এবং উওর দাতা যে কয়দিন বলে সেই নাম্বার টি খালি ঘরে লিখুন (০-৭)

*For each item on the list, read the question below and fill in the number of days the respondent says (0-7).*

**C.611.**

গত সাত দিনে আপনার শিশু [নাম] কয়দিন এই খাবার খেয়েছে [ তালিকার খাবার]? **(তালিকা থেকে প্রত্যেকটি প্রশ্ন পড়ে শোনান )**

How many days in the last 7 days did [NAME] have [ITEM FROM LIST]?

| | খাবারগুলো (উপাদান অনুযায়ী ভাগ করা হয়েছে) যা শিশু গত সাত দিনে খেয়েছেFoods (in groups) eaten by the child in the last seven days | শিশুটি যে কয় দিন খাবার খেয়েছে =(০-৭)<br>Number of days food was eaten by child (0-7)<br>খেয়েছে,কিন্তু কয়দিন খেয়েছে তা জানেনা = ৬৬<br>Eaten, don't know how many days = **66**<br>খেয়েছে কি খায়নিজানেনা = ৯৯<br>Don't know if eaten or not = **99** |
|---|---|---|
| 1 | খিচুড়ী ( খিচুড়ী কি কি দিয়ে রান্না হয়েছে জেনে নিয়ে ফুড গ্রুপ অনুযায়ী নীচে কোড করুন), ভাত, রুটি?<br>Khichuri (tick 'yes' and ask for the ingredients and code below according to food group), rice, bread? | I__I__I |
| 2 | মিষ্টি কুমড়া, গাজর ?<br>Pumpkin, carrots? | I__I__I |
| 3 | আলু, মিষ্টি আলু অথবা অন্যান্য যে কোন সাদা রং এর আলু জাতীয় খাদ্য?<br>Potato, sweet potatoes or any other white colored root or tuber? | I__I__I |
| 4 | গাঢ় সবুজ শাক যেমনমিষ্টি কুমড়া শাক, সরিষা শাক দিয়ে তৈরী ঝোল বা কোন মজাদার খাবার?<br>Any sauce or relish made with dark green leaves such as pumpkin leaves or mustard leaves? | I__I__I |
| 5 | পাকা আম বা পাকা পেঁপে?<br>Ripe mango or ripe papaya,? | I__I__I |
| 6 | অন্যান্য যে কোন ফল যেমন পেয়ারা ,আপেল,আঙ্গুর, কলা কমলা ? | I__I__I |

| | | |
|---|---|---|
| | Any other fruit such as, guava, apple, grape or banana, orange | |
| 7 | অন্যান্য যে কোন সবজি যেমন টমেটো,পিঁয়াজ,মাশরুম,সীম,মটরশুটি ?<br>Any other vegetable such as tomato, onions, mushroom, beans/ green peas or any other? | I__I__I |
| 8 | যে কোন মাংস ,পাখি বা প্রাণীর মাংস সহ?<br>Any type of meat, including from birds or from animals? | I__I__I |
| 9 | যে কোন ধরণের ডিম ?<br>Any type of egg? | I__I__I |
| 10 | যে কোন ধরণের শুটকি মাছ অথবা তাজামাছ?<br>Any type of dried fish or fresh fish? | I__I__I |
| 11 | সীম, মটর,ডাল,চীনাবাদাম বা অন্যান্য বাদাম?<br>Any dishes made with beans, peas, lentils, groundnut, or other nuts,? | I__I__I |
| 12 | যে কোন দুধ, পনির, দই বা অন্যান্য দুধের তৈরী খাদ্য?<br>Any milk, cheese, yogurt, or foods/drinks made with milk? | I__I__I |
| 13 | উদ্ভিজ্জ তেল, পশুর চর্বি , ঘি, মাখন বা অন্য যে কোন খাবার যা এই গুলো দিয়ে তৈরী ?<br>Vegetable oil, fat from animals, ghee, butter or any foods made with these? | I__I__I |
| 14 | মিষ্টি খাবার যেমন: চকলেট, মিষ্টি/ ক্যান্ডি, পিঠা অথবা মিষ্টি বিস্কুট<br>Sweet foods such as chocolate, sweets/candies, cake or cookies/sweet biscuits, mishti? | I__I__I |

আমি এখন আপনার কাছে আপনার শিশুর [নাম] শিশু খাদ্য এবং কিছু বিশেষ খাবার যা শিশুরা মাঝে মাঝে খায় সে বিষয়ে কিছু প্রশ্ন করতে চাই ।যদি আপনি এই খাবার সম্পর্কে আগেই বলে থাকেন, তবে আনুগ্রহ করে আবার বলুন যাতে আমি এই বিশেষ খাবার গুলো নির্ভূল ভাবে লিখতে পারি ।

Now I would like to ask you about infant formula and about some special foods that are sometimes given to infants and small children. Even if you already told me about the food, please tell me again so I can be sure to write down these special foods.

**C.612.**

গত সাত দিনের মধ্যে কয়দিন আপনার শিশু [নাম] এই খাবার খেয়েছে [সাক্ষাৎকারের দিন থেকে] [তালিকার খাবার]?

On how many days in the last 7 days, since last [INTERVIEW DAY], did [NAME] have any [ITEM FROM LIST]?

| | | |
|---|---|---|
| | শিশু খাদ্য এবং বিশেষ খাবার যা শিশু গত সাত দিনে খেয়েছে<br>Infant formula and special foods eaten by the child in the last seven days | শিশুটি যে কয় দিন খাবার খেয়েছে =(০-৭)<br>Number of days food was eaten by child (0-7)<br>খেয়েছে,কিন্তু কয়দিন খেয়েছে তা জানেনা = ৬৬<br>Eaten, don't know how many days = **66**<br>খেয়েছে কি খায়নিজানেনা = ৯৯<br>Don't know if eaten or not = **99** |
| 1 | শিশু খাদ্য যেমন ল্যাকটোজেন অথবা নান অথবা বায়োমিল,মাইবয় অন্যান্য?<br>Infant formula such as Lactogen or NAN or Biomeal, My Boy, or others? | I__I__I |
| 1.1 | অন্যান্য হলে ,নির্দিষ্ট করুন? If others, What type?<br>_______________________________________________ | |
| 2 | পানিতে বা দুধে সিদ্ধ করা শস্য জাতীয় খাবার যেমন: সুজি অথবা অন্যান্য খাবার যা ভূট্টার তৈরী, যেগুলো দোকানে কিনতে পাওয়া যায়?<br>Porridge or other food made with corn/ vutta, of the type bought in stores? | I__I__I |

| | | |
|---|---|---|
| 3 | অথবা শিশুদের খাদ্য শস্য যেমন সেরেলাক?<br>Other baby cereal such as Cerelac or other? | I__I__I |
| 3.1 | অন্যান্য হলে ,নির্দিষ্ট করুন? If others, What type?<br>_______________________________________________ | |
| 4 | পুষ্টিকণা, মনিমিক্স মিশ্রিত খাবার [গুড়া বা মাইক্রোনিউট্রিয়েন্ড দানা যা বাজারে পাওয়া যায়]?<br>Foods to which you added pushtikona/ monimix/ other [a powder or micronutrient sprinkles available in the market]? | I__I__I |
| 4.1 | অন্যান্য হলে ,নির্দিষ্ট করুন? If other, what type?<br>_____________________________________________________ | |
| 5 | সোনামনি যা আমাদের থেকে পেয়েছেন?<br>Shonamoni you received from us? | I__I__I<br>যদি ওয়াশ-বেনিফিট এ অন্তর্ভূক্ত হওয়ার পর পর হয় তাহলে "**00**" কোড করুন ।<br>যদি শিশুটির বয়স ৬ মাসের বেশী হয় এবং সে ওয়াশ-বেনিফিট থেকে কোন LNS না পেয়ে থাকে তাহলে "**88**" কোড করুন ।<br>[Fill in "**00**" at enrollment.<br>Fill in "**88**" if infant does not receive LNS from WASH Benefits] |
| 6 | অন্য যে কোন (এল এন এস) বা সম্পূরক পুষ্টি/পুষ্টি প্যাকেট?<br>Any other [Lipid-based nutrient supplement (LNS)]? | I__I__I |
| 6.1 | যদি হ্যা হয়, দেখাতে বলুন এবং নাম লিখুন: ________________________________<br>If Yes, ask to see and write name/type: | |

এখন আমি আপনার কাছে ভিটামিন/মিনারেল এর বড়ি অথবা ড্রপস সর্ম্পকে জানতে চাইবো।

Now I would like to ask you about vitamin/mineral pills or drops.

**C.613.**
গত সাত দিনে কয়দিন আপনার শিশু [নাম] ভিটামিন/মিনারেল এর বড়ি অথবা ড্রপস নিয়েছে?

On how many days did [NAME] have any vitamin/mineral pills or drops in the last 7 days?

(০ - ৭)=**দিন** [০০ হলে C.615 নং প্রশ্নে যান]

(0 – 7) = DAYS[*if* 00 then skip to C.615]

৬৬ = খেয়েছে,কিন্তু কয়দিন খেয়েছে তা জানেনা

66 Child had, but number of days not known

৯৯ =খেয়েছে কি খায়নিজানেনা

99 Don't know if child had or not

☐☐

**C.614.**
যদি শিশু ভিটামিন/মিনারেল এর বড়ি অথবা ড্রপস নিয়ে থাকে : কি ধরনের?

If baby was given vitamin / mineral drops or pills: What type?

_____________________________________________________

উত্তর দাতাকে মোড়ক দেখাতে বলুন এবং উপরের লাইনে নাম লিখুন।

*Ask the respondent to show the package and write the name on the line above.*

👁 **পর্যবেক্ষণ:** বড়ি অথবা ড্রপস এর নামের , তথ্যের উৎস কি?Observation:
What is the source of the information on the pill or drop name?

- ☐ 1 তথ্য পাওয়া যায় নি Information not available
- ☐ 2 তথ্য সংগ্রহকারী মোড়ক দেখেছে Data collector saw package
- ☐ 3 উওর দাতা মনেকরে নাম বলতে পেরেছে Respondent remembered and told name

**C.615.**
শিশু কি কখনো ময়লা বা মাটি খিয়েছে?Has the child eaten any dirt or soil?

মনে করার জন্য প্রতিটি সময় আলাদা ভাবে জানতে চান: *Ask for each recall period:*

| | | | | |
|---|---|---|---|---|
| 1 | আজ Today | ☐ 1 Yes | ☐ 2 No | ☐ 99 Don't know |
| 2 | গতকাল Yesterday | ☐ 1 Yes | ☐ 2 No | ☐ 99 Don't know |
| 3 | গত কালের আগের দিন Day before yesterday | ☐ 1 Yes | ☐ 2 No | ☐ 99 Don't know |
| 4 | গত সাত দিন (আজকের দিন থেকে গত এক সপ্তাহ?) In the past 7 days (since this day last week?) | ☐ 1 Yes | ☐ 2 No | ☐ 99 Don't know |

#### Wash Benefit Module 7 HANDWASHING

| | WBM 7 | | |
|---|---|---|---|
| C.700 | 70 | এই প্রশ্নের আগে উত্তররদাতা যে কোন সময় তার হাত ধুয়েছিল কিনা তা পর্যবেক্ষণ করুন।RECORD WHETHER THE RESPONDENT HAS WASHED HER HANDS AT ANY TIME BEFORE THIS QUESTION | [1] উত্তররদাতা হাত ধুয়েছিলOBSERVED RESPONDENT WASHING HANDS<br>[2] উত্তররদাতা হাতধোয় নাইDID NOT OBSERVE RESPONDENT WASHING HANDS |
| C.701 | 70 | আপনাকে ধন্যবাদ । এখন আমি আপনার হাতগুলো দেখতে চাই। আশা করি আপনি কিছু মনে করবেন না। আপনি কি আমাকে দেখাবেন? READ: Thank you. Now, I would like to do a quick inspection of your hands. I hope you don't mind. Can you please show me your hands?<br><br>দুই হাতই দেখতে হবে(প্রথমে এক হাত)। এরপর নিচের উলেস্নখিত হাতের অবস্থার কোড দেখে রেকর্ড করতে হবে। BOTH HANDS SHOULD BE SHOWN (NOT JUST ONE HAND). USE APPEARANCE CODES BELOW TO RECORD THE DESCRIPTION THAT BEST DESCRIBES THE DEFINITIONS THE LEVEL OF CLEANLINESS.<br>APPEARANCE CODES:<br>[1] ময়লা স্পষ্টভাবে দেখা যাচ্ছিল VISIBLE DIRT (DIRT/MUD/SOIL/ASH OR ANY OTHER MATERIAL IS VISIBLE)<br>[2] ময়লা স্পষ্টভাবে দেখা নাগেলেও অপরিচ্ছন্নভাব ছিলUNCLEAN | 👁 MOTHER<br><br>**বাম হাত Left Hand**<br>A\|__\| হাতের নখ FINGERNAILS<br>B\|__\| হাতের তালু PALMS<br>C\|__\| আঙ্গুলের সম্মুখভাগ FINGER PADS<br><br>**ডান হাত Right Hand**<br>D\|__\| হাতের নখ FINGERNAILS<br>E\|__\| হাতের তালু PALMS<br>F\|__\| আঙ্গুলের সম্মুখভাগ FINGER PADS |

| | | | |
|---|---|---|---|
| | | APPEARANCE (NO DIRT IS VISIBLE ON THIS PART OF THE HAND BUT, IN GENERAL, THIS PART OF THE HAND APPEARS UNCLEAN)<br>[3] পরিষ্কার ছিল CLEAN (OBSERVED PART OF THE HAND IS CLEAN AS WOULD APPEAR AFTER SOMEONE WASHES HANDS OR TAKES A BATH)<br>৮৮. পর্যবেক্ষণ করা সম্ভব হয়নি/প্রত্যাখান N/A; OBSERVATION NOT POSSIBLE/REFUSED | |
| **C.701** | 70 | ASK: দয়া করে আমাকে(শিশুর নাম ধরে) হাতগুলো দেখাও । Please show me [NAME]'s hands.<br><br>উত্তরদাতার সবচেয়ে বড় বাচ্চার (০-৩৬ মাস) হাত দেখুন । এই বয়সের বাচ্চা না থাকলে স্কিপ করু। TO SEE THE HANDS OF THE RESPONDENT'S OLDEST CHILD 0-36 MONTHS (SEE C.1) IF RESPONDENT DOESN'T HAVE A CHILD 0-36 MONTHS → **SKIP TO NEXT PAGE**.<br><br>APPEARANCE CODES:<br>[1] ময়লা স্পষ্টভাবে দেখা যাচ্ছিল(**ময়লা/কাঁদা/মাটি/ছাই/এছাড়া অন্য কিছু**)VISIBLE DIRT (DIRT/MUD/SOIL/ASH OR ANY OTHER MATERIAL IS VISIBLE)<br>[2] ময়লা স্পষ্টভাবে দেখা না গেলেও অপরিচ্ছন্নভাব ছিল(হাতে **ময়লা দেখা না গেলেও**অপরিচ্ছন্ন ছিল)UNCLEAN APPEARANCE (NO DIRT IS VISIBLE ON THIS PART OF THE HAND BUT, IN | 👁সবচেয়ে বড় বাচ্চার ০-৩৬ মাস OLDEST CHILD 0-36 MONTHS<br><br>**বাম হাত Left Hand**<br>A\|__\| হাতের নখ FINGERNAILS<br>B\|__\| হাতের তালু PALMS<br>C\|__\| আঙ্গুলের সম্মুখভাগ FINGER PADS<br><br>**ডান হাত Right Hand**<br>D\|__\| হাতের নখ FINGERNAILS<br>E\|__\| হাতের তালু PALMS<br>F\|__\| আঙ্গুলের সম্মুখভাগ FINGER PADS |

| | | | |
|---|---|---|---|
| C.702a | 7 | পর্যবেক্ষনঃ প্রাথমিক/**প্রধান** হাত ধোয়ার স্থানটি কোথায় তা রেকর্ড | 1. ঘরের ভিতরে/ কাছে (≤৬ কদম) In/near main house (≤6 feet to entrance)<br>2. পায়খানার ভিতরে/ কাছে(≤৬ কদম) In/near latrine (≤6 feet to entrance)<br>3. রান্নার স্থানের ভিতরে/ কাছে (≤৬ কদম) In/near cooking area (≤6 feet to entrance) |

| | | |
|---|---|---|
| | | [3] পরিষ্কার ছিল(হাতধোয়ার পর/গোসলের পর হাত দেখলে পরিষ্কার হবে) CLEAN (OBSERVED PART OF THE HAND IS CLEAN AS WOULD APPEAR AFTER SOMEONE WASHES HANDS OR TAKES A BATH)<br>[88] পর্যবেক্ষণ করা সম্ভব হয়নি/প্রত্যাখান/০-৩৬ মাসেরবাচ্চা নাই N/A; OBSERVATION NOT POSSIBLE/REFUSED/NO CHILD 0-36 MONTHS |
| | 70 | |

| | | | |
|---|---|---|---|
| | | করুন **[Observation: Record the location where the primary hand washing station is located?]** | 4. ঘর, পায়খানা, রান্নার স্থানে থেকে >৬ কদম>6 feet away from main house, latrine <u>and</u> cooking area<br>**5.** নির্দিষ্ট কোন জায়গায় নাই No specific place 714 নং প্রশ্নে চলে যান (skips to 714) |
| C.702c | 7 | পর্যবেক্ষনঃ প্রাথমিক/ **প্রধান** হাত ধোয়ার স্থানটিতে কোন কোন উপাদান আছে? (আপনি যদি দেখে থাকেন তাহলে "1" কোড করুন আর না দেখলে "0" কোড করুন) [Observation: Which of the following are present at the primary hand washing station? (If you observe the listed item, write "1" for "yes" in the box below. If you do not observe the listed item, write "0" for "no" in the box below.)] *<u>1= হ্যাঁ[Yes], 0=না [No]</u>* | MATERIALS PRESENT<br>[1]............................................পানি WATER<br>[2]............................................গোসল/হাত ধোয়ার সাবান (লাক্স, লাইফবয়) BAR SOAP (Body/hand Bar)<br>[3]............................................অন্য যে কোন ধরনের সাবান (হুইল) BAR SOAP (other)<br>[4]............................................গুড়া পাউডার POWDERED SOAP<br>[5] সাবান-পানি SOAPY WATER<br>[6] তরল সাবান LIQUID SOAP<br>[7] ছাই Ash<br>[8] মাটি /বালু Mud/Sand<br>[9] আইসিডিডিআর, বি-র দেওয়া ট্যাপসহ বালতি ICDDR,B drum with tap<br>[10] হাত ধোয়ার স্থানটি ভিজা ছিল/ব্যবহারের চিহ্ন ছিল Moisture below handwashing station<br>[11] বালতি Bucket<br>[12] বেসিন, জগ Basin/Jug<br>[13] কিছুই নাই NONE OF THE ABOVE<br>[777] অন্যান্য OTHER (SPECIFY): ______________ |
| | 7 | **প্রধানহাত ধোয়ার স্থানটি সাবান জাতীয় যেকোন ধরনের উপাদান (যা হাত ধোয়ার কাজে ব্যবহার করা হয়/ সাবান পানি ও পানি একসাথে উপস্থিত আছে কী?** [Soap/soapy water and water present together in PHWS] | 1. হ্যাঁ [Yes]<br>0. ............................................না [No] |
| C.702c | 7 | প্রাথমিক/ **প্রধান** হাত ধোয়ার স্থানটি রান্নাঘর থেকে কত কদম দূরে? [Observation: The primary hand washing station is away from the kitchen---Count in steps, allow in continuous numbers] | \|____\| |

| | | | |
|---|---|---|---|
| C.702d | 7 | প্রাথমিক/ **প্রধান** হাত ধোয়ার স্থানটি পায়খানা থেকে কত কদম দূরে? [Observation: The primary hand washing station is away from the toilet---Count in steps, allow in continuous numbers] | \|____\| |
| C.703 | 7 | উত্তরদাতাকে প্রশ্ন করম্ননঃ আপনার হাতধোয়ার জন্য প্রাথমিক/ **প্রধান** স্থান ছাড়া অন্যকোন জায়গা আছে কি? [Ask the respondent: "Is there anywhere else you wash your hands?" (Record code in box)]: | 1. হ্যাঁ [Yes]<br>0. না [No] **714**নং প্রশ্নে চলে যান (skips to 714) |
| C.703a | 7 | পর্যবেক্ষনঃ দ্বিতীয় প্রধান হাত ধোয়ার স্থানটি কোথায় তা রেকর্ড করুন [Observation: Record the location of the secondary hand washing station]. | 1. ঘরের ভিতরে/ কাছে (≤৬ কদম) In/near main house (≤6 feet to entrance)<br>2. পায়খানার ভিতরে/ কাছে(≤৬ কদম) In/near latrine (≤6 feet to entrance)<br>3. রান্নার স্থানে ভিতরে/ কাছে (≤৬ কদম) In/near cooking area (≤6 feet to entrance)<br>4. ঘর, পায়খানা, রান্নার স্থানে থেকে >৬ কদম>6 feet away from main house, latrine <u>and</u> cooking area |
| C.703c | 7 | পর্যবেক্ষনঃ দ্বিতীয় প্রধান হাত ধোয়ার স্থানটিতে কোন কোন উপাদান আছে? (আপনি যদি দেখে থাকেন তাহলে "1" কোড করুন আর না দেখলে "0" কোড করুন) [Observation: Which of the following are present at the secondary hand washing station? (If you observe the listed item, write "1" for "yes" in the box below. If you do not observe the listed item, write "0" for "no" in the box below.) ] | *<u>1= হ্যাঁ[Yes], 0=না [No]</u>*<br>[1]........................................................................................................................পানি WATER<br>[2]........................................................................................................................গোসল/হাত ধোয়ার সাবান (লাক্স, লাইফবয়) BAR SOAP (Body/hand Bar)<br>[3]........................................................................................................................অন্য যে কোন ধরনের সাবান (হুইল) BAR SOAP (other)<br>[4]........................................................................................................................গুড়া পাউডার POWDERED SOAP<br>[5] সাবান-পানি SOAPY WATER<br>[6] তরল সাবান LIQUID SOAP<br>[7] ছাই Ash<br>[8] মাটি /বালু Mud/Sand<br>[9] আইসিডিডিআর, বি-র দেওয়া ট্যাপসহ বালতি ICDDR,B drum with tap<br>[10] হাত ধোয়ার স্থানটি ভিজা ছিল/ব্যবহারের চিহ্ন ছিল Moisture below handwashing station<br>[11] (বালতি Bucket<br>[12] বেসিন, জগ Basin/Jug<br>[13] কিছুই নাই NONE OF THE ABOVE<br>৭৭৭. অন্যান্য OTHER (SPECIFY): ______________ |
| | 7 | **দ্বিতীয় প্রধানহাত ধোয়ার স্থানটিতে সাবান জাতীয় যেকোন ধরনের উপাদান (যা** | 1. হ্যাঁ [Yes]<br>0........................................................................................................................না [No] |

| | | | |
|---|---|---|---|
| | | **হাত ধোয়ার কাজে ব্যবহার করা হয়/সাবান পানি ও পানি একসাথে উপস্থিত আছে কী?** [Soap/soapy water and water present together in SHWS] | |
| C.703c | 7 | দ্বিতীয় প্রধান হাত ধোয়ার স্থানটি রান্নাঘর থেকে কত কদম দূরে? [Observation:T he secondary hand washing station is away from the kitchen---Count in steps, allow in continuous numbers]] | \|____\| |
| C.703d | 7 | দ্বিতীয় প্রধান হাত ধোয়ার স্থানটি পায়খানা থেকে কত কদম দূরে? [Observation: The secondary hand washing station is away from the toilet---Count in steps, allow in continuous numbers] | \|____\| |
| INDICATOR 4: HANDWASHING সাবানেরউপস্থিতিতে হাতধোয়া দেখানো DEMONSTRATION, SOAP PRESENCE | | | |
| আপনাকে ধন্যবাদ। আপনি দয়া করে আমাকে পায়খানা থেকে আসার পর যেখানে বেশীরভাগ সময় হাত ধোন সেই স্থানটি দেখান(নোট ;পূর্বে পর্যবেক্ষীত স্থানও হতে পারে। এরপর উত্তরদাতা কর্তক চিহ্নিত স্থানে যেতে হবে এবং রেকর্ড করতে হবে।Thank you. Please show me where you most often wash your hands after defecation. (Note: this may be the same place you already observed. Go to the place identified by the respondent and record location). | | | |
| C.704a | 7 | 👁 হাতধোয়ার স্থান পর্যবেক্ষণ করুন(একটিকে গোলাকার করুন)। OBSERVE AND RECORDLOCATION OF HANDWASHING (CIRCLE ONE) | 1. ঘরের ভিতরে/ কাছে (≤৬ কদম) In/near main house (≤6 feet to entrance)<br>2. পায়খানার ভিতরে/ কাছে(≤৬ কদম) In/near latrine (≤6 feet to entrance)<br>3. রান্নার স্থানে ভিতরে/ কাছে (≤৬ কদম) In/near cooking area (≤6 feet to entrance)<br>4. ঘর, পায়খানা, রান্নার স্থানে থেকে >৬ কদম>6 feet away from main house, latrine and cooking area<br>5. নির্দিষ্ট কোন জায়গায় নাই |
| **সময় গনণার জন্য ষ্টপ ওয়াচ প্রস্তুতকরন PREPARE YOUR STOP WATCH FOR TIMING.** | | | |
| C.705 | 7 | পায়খানা থেকে আসার পর যেভাবে আপনি আপনার হাতধোন একইভাবে আমাকে এখন তা করে দেখান। Now, can you demonstrate how you normally clean your hands after defecation? Try in the same manner as you would if I wasn't here. TIME HOW LONG | [1] দেখানো হয়েছে DEMONSTRATED<br><br>[2] দেখানো হয়নি COULD NOT DEMONSTRATE<br>IF NOT → **RECORD REASON: ________________________________________AND SKIP TO 721** |

| | | | |
|---|---|---|---|
| | | IT TAKES THE RESPONDENT TO BRING MATERIALS TO THAT PLACE (START AT THE MOMENT YOU FINISH ASKING THE QUESTION). WHEN RESPONDENT STARTS CLEANING HER HANDS, TIME DURATION OF RUBBING HANDS TOGETHER. | |
| C.705b | 7 | 👁 পর্যবেক্ষণ করন্নন এবং হাতধোয়ার সাবান আনতে কতটুকু সময় লেগেছে তা রেকর্ড করন্নন। যদি হাতধোয়ার স্থানে সাবান থাকে তাহলে ০০:০০ বসান। যদি সাবান ব্যবহার না করে থাকে তাহলে ৮৮:৮৮ বসান । OBSERVE AND RECORD TIME TO PRODUCE (Bangladesh: <u>SOAP)</u> FOR CLEANING HANDS MARK "00:00" IF SOAP IS ALREADY PRESENT AT HANDWASHING PLACE. (Bangladesh ONLY) MARK "88:88" IF SOAP IS N T USED | \|____:____\| মি.mm. সে.ss |
| C.705c | 7 | 👁 পর্যবেক্ষণ এবং হাতঘষার সময় রেকর্ড করুন \| উত্তরদাতা যখন হাত পরিস্ক ারের জন্য দুহাত ঘষবে সেই সময় গনণা করতে হবে । OBSERVE AND RECORD LENGTH OF TIME RESPONDENT SPENDS <u>RUBBING HER HANDS</u> TOGETHER: | \|____:____\| মি.mm. সে.ss |
| C.705d | 7 | 👁 পর্যবেক্ষণ এবং হাত ধোয়ার সকল উপাদান রেকর্ড করুন OBSERVE AND RECORD <u>ALL MATERIALS</u> USED FOR HAND CLEANSING | 1. পানি WATER<br>2. **বার সাবান** BAR SOAP (Body/hand BAR)<br>3. **বার সাবান** (অন্যান্য)BAR SOAP (other)<br>4. **গুড়া পাউডার** POWDERED SOAP<br>5. **সাবান-পানি** SOAPY WATER |

| | | | |
|---|---|---|---|
| | | (CIRCLE ALL THAT APPLY) 1 হ্যাঁ Yes 0 না No | 6. কাপড়/পাতা CLOTH OR LEAVES<br>7. বেসিন/কলসBASIN/PITCHER<br>8. ছাই ASH<br>9. মাটি/বালু MUD<br>88. অন্যান্য উল্লেখ করুন OTHER (SPECIFY): ___________________________ |
| C.705e | 7 | 👁 পর্যবেক্ষণ এবং ২ হাত ধোয়া রেকর্ড করুন \|OBSERVE AND RECORD WHETHER BOTH HAN S WERE CLEANED | [1] হ্যাঁ, ২-হাত ধুয়েছে YES, BOTH HANDS CLEANED<br>[2] না ,১-হাত ধুয়েছে NO, ONLY ONE HAND CLEANED |
| C.705f | 7 | 👁 পর্যবেক্ষণ এবং হাত শুকানোরেকর্ড করুনOBSERVE AND RECORD HOW RESPONDENT DRIES HANDS<br>1 Yes 0 No | 1. পরিধেয় কাপড় DRIES BY WIPING HER HANDS ON HER CLOTHES<br>2.এক হাত অন্য হাতের সাথে ঘষে DRIES BY WIPING HER HANDS ON ANOTHER<br>3. পরিধেয় ছাড়া অন্য পরিষ্কার কাপড় Clean CLOTH DRIES BY WIPING HER HANDS ON<br>৪. পরিধেয় ছাড়া অন্য অপরিষ্কার কাপড় Unclean CLOTH DRIES BY WIPING HER HANDS ON<br>5. কাপড় ছাড়া অন্য জিনিস যেমন: টিস্যু, কাগজ ইত্যাদিMATERIAL (NOT CLOTH)<br>5. বাতাসে/হাত ঝাঁকিয়ে বাতাসে শুকানো হয়েছে DRIES BY SHAKING HER HANDS IN THE AIR<br>6. হাত শুকানো হয় নাই NOT DRIED<br>88. অন্যান্য উল্লেখ করুন OTHER, SPECIFY________________________ |
| C.705g | 7 | যদি উত্তরদাতা হাতধোয়া না দেখান অথবা সাবান ব্যবহার না করে তখন তাকে তার খানায় হাতধোয়ার জন্য সাবান আছে কিনা জানতে হবে এবং তা আনতে বলতে হবে?IF RESPONDENT COULD NOT DEMONSTRATE, OR DID NOT USE SOAP, ASK: Do you have soap in your house that you use for handwashing? Can you bring it to me?<br>👁 পর্যবেক্ষণ করুন এবং হাতধোয়ার সময় রেকর্ড করুন। যদি 716 তে উত্তর/সময় উল্লেখ থাকে তাহলে ৮৮:৮৮ বসান। যদি খানাতেসাবান না থাকে তাহলে ৯৯:৯৯ বসান । OBSERVE AND | \|____:____\| মি.mm. সে.ss |

| | | |
|---|---|---|
| | | RECORD TIME TO PRODUCE SOAP FOR WASHING HANDS<br>MARK “88:88” IF ALREADY TIMED IN C.704a<br>MARK “99:99” IF OBSERVATION NOT POSSIBLE OR SOAP NOT AVAILABLE |

| | | | |
|---|---|---|---|
| C. 706a | 722. | 👁 পর্যবেক্ষণ করুন : **উত্তরদাতার হাতধোয়া দেখানোর সময়ে এ বিষয়ে কোন প্রতিক্রিয়া দেখতে পেয়েছিলেন কী?**OBSERVE: WAS THERE ANY OBVIOUS REACTIVITY DURING THE COURSE OF HIS DEMONSTRATION? | [1] *হ্যাঁ YES*<br>[0] *না (৭২৩ নং এ চলে যাবে)* NO → skip to 723 |
| b | 1 | অধিকক্ষণ যাবৎ হাত ধুয়েছে কিনা<br>Longer time spent cleaning/rubbing hands | [1] *হ্যাঁ* YES<br>[0] *না* NO |
| c | 2 | সাবান দিয়ে হাত ধুয়েছে কিনা অন্যথায় | [1] *হ্যাঁ* YES |

| | | | |
|---|---|---|---|
| | | Using soap where they otherwise would not | [0] না NO |
| d | 3 | অন্যান্য (লিখুন) Other, (BRIEFLY explain) ___________________________ | [1] হ্যাঁ YES<br>[0] না NO |

| | | | | | |
|---|---|---|---|---|---|
| | 72[illegible] | প্রশ্ন করুন ; আপনি কখন কখন সাবানদিয়ে হাত ধোন । ASK: "Please tell me about all of the times you wash your hands with soap."<br><br>A কলা ম (লিড ছাড়া)-এ যখন উত্তরদাতা যে কাজ করবে তাউল্লেখ করতে হবে । ১নং গোলাকার হবে যদি নির্দিষ্ট বিষয়ে উত্তর পাওয়া যায়। CIRCLE "1" IN COLUMN A IF CRITICAL TIME IS MENTIONED WITHOUT PROMPTING.<br><br>উত্তরদাতার সকল কাজ উল্লেখ করার পর অন্য আর কোন কাজে সাবানদিয়ে হাত ধোন কিনা এ ব্যপারে জিজ্ঞাসা করুন? AFTER THE RESPONDENT FINISHES NAMING ALL THE TIMES ASK "IS THERE ANY OTHER TIME YOU WASH YOUR HANDS WITH SOAP?" AND STOP WHEN THE RESPONDENT SAYS THERE IS NO OTHER TIME.<br><br>B কলাম-এ লিড দেওয়া ছাড়া প্রতিবার ব্যবহার করে কিনা তা উল্লেখ করুন । EACH TIME MENTIONED WITHOUT PROMPTING ASK COLUMN B.<br><br>C.707A-G নং প্রশ্নের উত্তর যদি A কলা ম [1] না হয় তখন সাবান ও পানি দিয়ে আপনি আপনার হাতধোন কিনা তা জিজ্ঞাসা করুন। যদি উত্তরদাতা হ্যাঁ বলে তখন C কলা ম-এ হাতধোয়ার সংখ্যা/বার উল্লেখ করুন । FOR QUESTIONS C.707A-G WHERE [1] IS NOT CIRCLED IN COLUMN A, PROMPT BY ASKING:<br>Do you usually wash your hands with soap and water [CRITICAL TIME]? IF THE RESPONDENT SAYS "YES" THEN IMMEDIATELY ASK HOW OFTEN AND MARK APPROPRIATE ANSWER CHOICE IN COLUMN C. | A.<br>লিড দেওয়া ছাড়া হাতধোয়া উল্লেখ করুন<br>MENTIONED WITH NO PROMPTING | B.<br>লিড দেওয়া ছাড়া শুধুমাত্র হাতধোয়ার সংখ্যা/বার উল্লেখ করুন । সাবান ও পানি দিয়ে কত বার আপনার হাতধোন তা বলুন । Only for times mentioned with NO PROMPTING, ask: How often do you wash your hands with soap and water [insert event]?<br><br>[1] সবসময়/প্রতিবার ALWAYS<br>[2] মাঝে মাঝে/ অর্ধেকের কম বার SOMETIMES<br>[3] খুব কম সময়/ দু-এক বার RARELY | C.<br>লিড দেওয়ার পর শুধুমাত্র হাতধোয়ার সংখ্যা/বার উল্লেখ করুন । সাবান ও পানি দিয়ে কত বার আপনার হাতধোন তা বলুন ।<br>REPLIED "AFTER BEING PROMPTED<br><br>[1] সবসময়/প্রতিবার ALWAYS<br>[2] মাঝে মাঝে/ অর্ধেকের কম বার SOMETIMES<br>[3] খুব কম সময়/ দু-এক বার RARELY<br>[4] না NO<br>[88] প্রযোজ্য নয় Not applicable |
| a | 1 | খাবার তৈরী করার আগে BEFORE PREPARING FOOD | | | |
| | 2 | মাংস/মাছ কাটার পর AFTER HANDLING MEAT/FISH | | | |
| | 3 | ফল/সব্জি কাটার আগেBEFORE CUTTING FRUITS AND VEGETABLES | | | |
| | 4 | ভর্তা/ম্যাসড খাবার তৈরী করার আগে BEFORE MAKING MASHED FOOD (BHORTA) | | | |
| b | 5 | খাবারের পূর্বেBEFORE EATING | | | |
| c | 6 | খাবারের পরে AFTER EATING | | | |
| d | 7 | শিশুকে খাওয়ানোর আগে BEFORE FEEDING A CHILD | | | |
| e | 8 | বাচ্চাকে সৌচানোর পরAFTER CLEANING A CHILD'S ANUS | | | |
| f | 9 | বাচ্চার পায়খানা ফেলার পরAFTER DISPOSING OF CHILDREN'S FECES | | | |
| g | 10 | পায়খানার পরAFTER TDEFECATION | | | |
| h | 11 | কম্পাউন্ড/বাড়ীর বাহির থেকে ফেরার পর AFTER RETURNING FROM OUTSIDE THE COMPOUND | | | |
| m | 12 | রুগ্ন ব্যক্তিকে ধরার পরAFTER TOUCHING A SICK PERSON | | | |
| o | 13 | পশু-পাখি ধরার পরAFTER HANDLING LIVESTOCK | | | |
| | 14 | গোবর ধরার পর AFTER HANDLING COW-DUNG (OR ANY ANIMAL FECES) | | | |

| | | |
|---|---|---|
| j | 15 | অন্যান্য (উল্লেখ করুন) OTHER (SPECIFY): ______________________________ |
| i | 16 | উত্তরদাতা কখনই সাবান দিয়ে হাতধোয় নি RESPONDENT NEVER WASHES HANDS WITH SOAP Skip to module 8 |
| | 17 | উঠান ঝাড়ু দেওয়া/ময়লা পরিষ্কারের পর After weaping courtyard |
| | 18 | রান্নার পর After cooking |
| | 19 | থালা-বাসন ধোয়ার পর After washing dishes |

**Wash Benefit Module 8SANITATION (All households)**

| | |
|---|---|
| SAY | এই গবেষণায় আপনার অংশ গ্রহনের জন্য আপনাকে ধন্যবাদ। ষ্টাডির এই অংশটি কিছুটা স্পর্শকাতর । আমি আপনাকে আপনার/এই বড়ীর পয়ঃ ব্যবস্থাপনা/পয়ঃ নিষ্কাশন ও তার অনুশীলন এবং কিভাবে তা করেনএ ব্যপারে জানতে চাই\| আমি কিছুটা পর্যবেক্ষণ করতে চাই। আপনার স্বাচ্ছন্দবোধ হবে কি ?যদি না হয় তাহলে উত্তরদাতাকে ইহার গুরুত্ব বোঝাতে চেষ্টা করুন।Thank you so much for your participation so far. The next part of the survey is a bit sensitive. I would like to ask you some questions about the sanitation practices of people in your compound, including how you usually dispose of your children's feces. I would also like to make some observations. Are you comfortable with this? IF NOT, EXPLAIN THE IMPORTANCE OF THIS MODULE AND ENCOURAGE RESPONDENT TO PROCEED. |

খোলা-পায়খানা(সকল খানার )OPEN DEFECATION (ADMINISTER TO ALL HOUSEHOLDS)

| | | | | | | | |
|---|---|---|---|---|---|---|---|
| | | | **Group A** | **B** | **C** | **D** | **E** |
| | WBM 8 | | **Men** পুরুষ | **Women** মহিলা | **Children <3 years** <৩ বছরের বাচ্চা | **Children 3-<8 years** ৩-<৮ বছরের বাচ্চা | **Children** ৮-১৫ ছরের বাচ্চা |
| | | এই খানার কোন সদস্যরা খোলা জায়গাতে পায়খানা করে কী? [Do [GROUP] in this household ever practice open defecation?<br>1 প্রতিদিন Daily<br>2 মাঝেমাঝে Occasionally<br>3 কখনই না Never **(Skip to 806)**<br>88. প্রযোজ্য নয় Not applicable (**806**নং প্রশ্নে যান)**(Skip to 806)**<br>99 জানি না (**806**নং | ☐ | ☐ | ☐ | ☐ | ☐ |

| | | | | | | | |
|---|---|---|---|---|---|---|---|
| | | প্রশ্নে যান) Don't Know **(Skip to 806)** | | | | | |
| | | সাধারনতঃ একই জায়গাতে প্রতিবার যান কী? [Do [GROUP] go to more or less the same area every time?<br><br>1হ্যাঁ Yes<br><br>2না No (**805**নং প্রশ্নে যান)**(Skip to 805)**<br><br>99জানি না Don't Know / Not Sure (**805** নং প্রশ্নে যান)**(Skip to 805)** | ☐ | ☐ | ☐ | ☐ | ☐ |
| | | আপনার খানা থেকে উক্তস্থানে যেতে (শুধু যাওয়া) কতটুকু সময় লাগে (মিনিট) [How long does it take to walk (one way) from your house to the most commonly visited place? | I______I মিনিট MINUTES<br><br>99 জানি না Don't Know / Not Sure | | | | |
| | | এই জায়গাটি ঐ গ্রামের মধ্যেই কী? [Is that place within the village?<br><br>1হ্যাঁ Yes<br><br>2না No<br><br>99জানি না Don't Know / Not Sure | I__I | | | | |

Open Defecation Questions

| | | | **Men**<br>পুরম্নষ | **Women**<br>মহিলা | **Children**<br>**<3 years**<br><৩ বছরের বাচ্চা | **Children**<br>**3-<8 years**<br>৩-<৮ বছরের বাচ্চা | **Children**<br>৮-১৫ ছরের বাচ্চা |
|---|---|---|---|---|---|---|---|
| **C.804 a** | | আপনার খানার লোকজনের খোলাজায়গায় পায়খানা করার | | | | | |

| | | | | | | | |
|---|---|---|---|---|---|---|---|
| | | প্রধান কারন কি কি? উত্তর শুনানো যাবে না।What are the main reasons that [GROUP] in your household practice open defecation?<br>DO NOT READ RESPONSES<br>MARK ALL THAT APPLY | | | | | |
| | 1 | 1 কোনও কিছুই নাই /পায়খানা নাইNo choice (nothing else is available) | ☐ | ☐ | ☐ | ☐ | ☐ |
| | 2 | 2কোথায়শিশু পায়খানা করবে তা নিয়ন্ত্রন করতে পারে না Cannot control where young children defecate | | | ☐ | ☐ | ☐ |
| | 3 | 3 গোপনীয়তাPrivacy | ☐ | ☐ | ☐ | ☐ | ☐ |
| | 4 | 4 অভ্যাস/নিয়মিত কাজHabit / Routine | ☐ | ☐ | ☐ | ☐ | ☐ |
| | 5 | 5 পায়খানা থেকে বাঁশঝাড় পছন্দ Prefer to use the bush rather than a toilet | ☐ | ☐ | ☐ | ☐ | ☐ |
| | 6 | 6 কাজে/স্কুলে পায়খানা/টয়লেট সহজে পাওয়া না গেলে Toilet not available at work / school | ☐ | ☐ | ☐ | ☐ | ☐ |
| | 7 | ভাগের পায়খানা/টয়লেট পছন্দ না হলে Choose not to share toilets with in-laws / extended | ☐ | ☐ | ☐ | ☐ | ☐ |

| | | | | | | | |
|---|---|---|---|---|---|---|---|
| | | family (or cannot) | | | | | |
| | 8 | 8 সুবিধাজনক Convenience | ☐ | ☐ | ☐ | ☐ | ☐ |
| | 9 | 9 নিরাপদ Safety | ☐ | ☐ | ☐ | ☐ | ☐ |
| | 10 | 10 আরামদায়কComfort | ☐ | ☐ | ☐ | ☐ | ☐ |
| | 11 | 11 অসুস্থতা/ পাতলা পায়খানাSickness/diarrhoea | ☐ | ☐ | ☐ | ☐ | ☐ |
| | 12 | 12 পায়খানা/টয়লেট ভরে গেছে Latrine overflowed | ☐ | ☐ | ☐ | ☐ | ☐ |
| | 13 | 13 পায়খানা/টয়লেট ভেঙে গেছে(উপরের অংশ/সস্নাব) Latrine broken<br>(superstructure and /or slab) | ☐ | ☐ | ☐ | ☐ | ☐ |
| | 14 | 14 পায়খানা/টয়লেটে যেতে ভয় পায় Fear of latrine | ☐ | ☐ | ☐ | ☐ | ☐ |
| | 15 | 15 পায়খানা/টয়লেট কিভাবে ব্যবহার করতে হয় তা জানে না Don’t know how to use the latrine | ☐ | ☐ | ☐ | ☐ | ☐ |
| | 16 | 16 **বাচ্চা**খুব ছোট হওয়ায় পায়খানা/টয়লেট ব্যবহার করে না Too young to use latrine | ☐ | ☐ | | | |
| | 17 | বয়স্ক (খুবই বৃদ্ধ)Old age (too old) | | | | | |
| | | | **<৩ বছরের** বাচ্চা**Children < 3** | **৩-<৮ বছরের** বাচ্চা**Childr** | **৮-১৫ বছরের** বাচ্চা**Childr** | **পুরম্নষ Men** | **মহিলা Wome** |

| | | | | | | | |
|---|---|---|---|---|---|---|---|
| | | | years | en<br><br>3 – <8 years | en<br><br>8 – 15 years | | n |
| **C.X** | | পায়খানা থাকা সত্ত্বেও এই বাড়ীর অন্য কোনপরিবারখোলা জায়গায় পায়খানা করে কিনা তা আপনি জানেন কী? Do you know of other households in the bari/compound whose [GROUP] practice OD, even if they might have a toilet or latrine at their house?<br><br>1 হ্যাঁ, প্রায়ই Yes, Often<br><br>2হ্যাঁ, মাঝে মাঝে Yes, Sometimes<br><br>3না, কখনই না No, Never**(Skip to 808)**<br><br>88. প্রযোজ্য নয় Not applicable (**808** নং প্রশ্নে যান)**(Skip to 808)**<br><br>99 জানি না/নিশ্চিত না Don't Know / Not Sure | ☐ | ☐ | ☐ | ☐ | ☐ |
| **C.X** | | আপনি শিশুদের পায়খানা উঠানে/উঠানের চার-পাশে/আশেপাশে /নদীতে ফেলে দিতে দেখেছেন কী? Do you see that children's stools are disposed in the yard / surrounding / | | | | ☐ | ☐ |

| | | |
|---|---|---|
| | | community in your neighborhood / river?<br>1 হ্যাঁ, প্রায়ই Yes, Often<br>2 হ্যাঁ, মাঝে মাঝে Yes, Sometimes<br>3 না, কখনই না No, Never<br>99 জানি না/নিশ্চিত না Don't Know / Not Sure |

| | | | |
|---|---|---|---|
| পয়ঃ ব্যবস্থাপনা সুবিধাদি Sanitation Facility | | | |
| Administer to: All study households | | | |
| C | | আপনার খানায় কোন টয়লেট/পায়খানা আছে কি?<br>আমি কি তা দেখতে পারি?<br>Does your household have a toilet facility that is in use? Can I see it? | 1 হ্যাঁ আছে,পর্যবেক্ষন করতে পারি Yes have toilet, can observe<br>2 হ্যাঁ আছে,পর্যবেক্ষন করতে দেওয়া হয়নি/প্রত্যাখান (৮১২ নং এ যান ) Yes have toilet, refused observation **(Skip to 812)**<br>3 হ্যাঁ আছে,পর্যবেক্ষন করা যায়নি(৮১২ নং এ যান ) Yes have toilet, cannot observe **(Skip to 812)**<br>4 কোন পায়খানা নাই No toilet facility **(Skip to MODULE 9)** |
| | | 👁 পর্যবেক্ষন; ধরণ,অবস্থা এবং পুনঃব্যবহার অবস্থা Observation: Note the type, condition and apparent use of the toilet:<br>1 হ্যাঁ Yes<br>2 না No<br>88 প্রযোজ্য নয়/ পর্যবেক্ষন করতে | |

| | | | |
|---|---|---|---|
| | | পারেনি/বলতে পারেনি<br>Not Applicable / Could not observe / cannot tell | |
| বর্হিরাংশ পর্যবেক্ষণ **Exterior observation** | | | |
| 1 | 1 | পায়খানার চারপাশে ৩টি ইটের/ টিনের/মাটির দেয়াল At least 3 walls around the toilet | *(1=হ্যাঁ[Yes], 0= না [No],* প্রযোজ্য নয় *[*not applicable] =888)] |
| 2 | 2 | পায়খানার চারপাশে ৩টি বাঁশের বেড়ার দেয়ালBamboo fences around the toilet | *(1=হ্যাঁ[Yes], 0= না [No],* প্রযোজ্য নয় *[*not applicable] =888)] |
| 3 | 3 | পায়খানার চারপাশে দরজা/পর্দা দিয়ে ঘেরাDoor/curtain or walls that guarantee privacy around the toilet | *(1=হ্যাঁ[Yes], 0= না [No],* প্রযোজ্য নয় *[*not applicable] =888)] |
| 4 | 4 | পায়খানার ছাদ Roof over toilet | *(1=হ্যাঁ[Yes], 0= না [No],* প্রযোজ্য নয় *[*not applicable] =888)] |
| 5 | 5 | বাতাস বের হওয়ার পথ Ventilation pipe | *(1=হ্যাঁ[Yes], 0= না [No],* প্রযোজ্য নয় *[*not applicable] =888)] |
| 6 | 6 | পায়খানায় যাবার রাস্তা দেখে বুঝা যাচ্ছে নিয়মিত ব্যবহার করা হয় (পরিষ্কার, জীর্ণ ইত্যাদি )Path to the toilet suggests regular use (is clear, well-worn, without grass or any barriers etc.) | *(1=হ্যাঁ[Yes], 0= না [No],* প্রযোজ্য নয় *[*not applicable] =888)] |
| ভিতরের অংশ পর্যবেক্ষণ **Interior observation** | | | |
| 7 | 7 | টয়লেটে স্লাব আছেToilet has a slab | *(1=হ্যাঁ[Yes], 0= না [No],* প্রযোজ্য নয় *[*not applicable] =888)] |
| 8 | 8 | গর্তের পাশে পাদানী আছেRaised footing around hole | *(1=হ্যাঁ[Yes], 0= না [No],* প্রযোজ্য নয় *[*not applicable] =888)] |
| 9 | 9 | পানি ঢেলে ফ্লাস করা যায় Flush or poor Flush: | *(1=হ্যাঁ[Yes], 0= না [No],* প্রযোজ্য নয় *[*not applicable] =888)] [If 0/888 skip to 12] |
| 9a | 9a | যদি পানি ঢেলে ফ্লাস করা যায় If Flush or Pour Flush:<br><br>পানির সীলের অবস্থা Water seal condition: | [ 1 ] ওয়াটার সিল ভাল আছে/কাজ করে Functional water seal<br>[ 2 ] ওয়াটার সিল ভাঙ্গা Broken water seal<br>[ 3 ] ওয়াটার সিল নেই No water seal<br>[ 88 ] পায়খানাতে পানির সীল আছে/পর্যবেৰণ করা |

| | | | |
|---|---|---|---|
| | | | সম্ভব হয়নিN/A / could not observe / cannot tell |
| 9b | 9b | যদি পানি ঢেলে ফ্লাস করা যায়If Flush or Pour Flush:<br><br>পানি ঢাললে/দিলে Flushes to:<br><br>(খানার সদস্যকে জিজ্ঞাসা/যাচাই করুন *Ask / probe household members if necessary)* | টয়লেটিতে পয়ঃনিষ্কাশন পাইপের সাথে সংযোগ করে দেয়া [Piped sewer system]<br><br>টয়লেটিতে সেপটিক ট্যাংক বসানো আছে [Septic tank]<br><br>সেপটিক ট্যাংক নাই কিন্তু ফ্লাস করে বা পানি ঢেলে পায়খানা দুরে বা পিটের মধ্যে সরিয়ে দেয়া যায় [Flush to pit latrine ]<br><br>ফ্লাশ-টয়েলেট অথবা পানি ঢেলে ফ্লাস করা টয়লেট যা কোন খাল, ড্রেন, নদী ইত্যাদির সাথে সংযোজন করার ফলে অস্বাস্থ্যকর অবস্থার সৃষ্টি করে থাকে[Flush or pour flush toilet connected to somewhere else (canal, ditch, river, etc.)] |
| | 9c | পায়খানার অবস্থা কেমনCondition of the latrine | 1. স্বাস্থ্যসম্মত পায়খানাHygienic latrine<br><br>2. অস্বাস্থ্যকর পায়খানাNon hygienic latrine |
| 10 | 10 | মেঝের প্রধান উপাদান Main material of the floor (*select 1*) | [ 1 ] মাটি Mud<br><br>[ 2 ] কাঠ Wood<br><br>[ 3 ] সিমেন্ট Cement<br><br>[ 4 ] টাইলস/ইটTile / brick<br><br>[ 5 ] প্লাষ্টিক Plastic<br><br>[ 88 ] প্রযোজ্য নয়/পর্যবেক্ষণ সম্ভব হয় নি N/A / could not observe / cannot tell |
| 11 | 11 | Bucket toilet | *(1=হ্যাঁ[Yes], 0= না [No],* প্রযোজ্য নয় *[*not applicable] =888)] |
| 12 | 12 | ঝুলন্ত পায়খানা Hanging toilet | [ 1 Yes ] [ 2 No ] [ 88 N/A] |
| 13 | 13 | ব্যবহৃত হচ্ছে বুঝা যায় এমন Latrine appears to be in use (by your best judgment) | *(1=হ্যাঁ[Yes], 0= না [No],* প্রযোজ্য নয় *[*not applicable] =888)] |
| 14 | 14 | পায়খানা/টয়লেটে পায়খানার গন্ধ রয়েছেOdor of feces in the latrine/bathroom | *(1=হ্যাঁ[Yes], 0= না [No],* প্রযোজ্য নয় *[*not applicable] =888)] |
| 15 | 15 | পায়খানা/টয়লেটে প্রস্রাবের গন্ধ রয়েছেOdor of urine in the latrine/bathroom | *(1=হ্যাঁ[Yes], 0= না [No],* প্রযোজ্য নয় *[*not applicable] =888)] |

| | | | |
|---|---|---|---|
| 16 | 16 | স্লাব অথবা মেঝেতে **পায়খানা দেখতে পাওয়া গেছে** Stool is visible on the slab or floor | *(1=হ্যাঁ[Yes], 0= না [No],* **প্রযোজ্য নয়** *[*not applicable] =888)] |
| 17 | 17 | ড্রপ হোল(মল-মূত্র প্রবেশ গর্ত) ঢাকা Drop hole is covered | *(1=হ্যাঁ[Yes], 0= না [No],* **প্রযোজ্য নয়** *[*not applicable] =888)][If 0/888 skip to 18] |
| 17a | 17a | যদি ড্রপ হোল (মল-মূত্র প্রবেশ গর্ত)ঢাকা থাকে তাহলে মাছি আসা/যাওয়া করতে পারে কি If yes, cover is correctly placed over the drop hole so that flies cannot enter/exit | *(1=হ্যাঁ[Yes], 0= না [No],* **প্রযোজ্য নয়** *[*not applicable] =888)] |
| 18 | 18 | **মাছির উপস্থিতি রয়েছে** Flies present | *(1=হ্যাঁ[Yes], 0= না [No],* **প্রযোজ্য নয়** *[*not applicable] =888)] |
| 19 | 19 | **সাধারন বৈশিষ্ট General Characteristics**<br><br>একটি অথবা দুই পিট/গর্ত টয়লেট Single or double pit latrine | [ 1 ] একটি পিট/গত Single pit<br><br>[ 2 ] দুই পিট/গর্ত Double pit<br><br>[ 88 ] প্রযোজ্য নয়/পর্যবেক্ষণ সম্ভব হয় নি /বলতে পারে নিN/A / could not observe / cannot tell |
| 20 | 20 | কমপোষ্টিং টয়লেট Composting toilet | *(1=হ্যাঁ[Yes], 0= না [No],* **প্রযোজ্য নয়** *[*not applicable] =888)] |
| | | 👁 **পর্যবেক্ষন করুন** Observation: টয়লেট-এর গর্ত মলে ভরেছে কিনা-আলোর সাহায্যে পর্যবেৰণ করুনFullness of the pit – shine a light into the pit to see if solid waste is… | 1 **শুকনা ময়লা/পায়খানা ৩ ফুটের চেয়ে বেশী দূরে (>৩ ফুট)** Very far from surface (>1 meter)<br><br>2 **শুকনা ময়লা/পায়খানা ৩ ফুটের মধ্যে (<৩ ফুট)** Within 1 meter<br><br>3 **পায়খানার খুব কাছাকাছি বা পায়খানা ভরে গেছে** Very close to surface or full<br><br>88 **পায়খানাতে পানির সীল আছে/পর্যবেৰণ করা সম্ভব হয়নি**Water seal latrine / non direct pit latrine / could not observe |
| C. 808 | | 👁 **পর্যবেক্ষন করুন** Observation: : সৌচ কাজের জন্য টয়লেট-এর ভেতরে এবং কাছে **কী কী উপাদান রয়েছে** What materials for anal cleansing are present inside or immediately outside the latrine? | *(1=হ্যাঁ[Yes], 0= না [No],* **প্রযোজ্য নয়** *[*not applicable] =888)]<br><br>1.*পাতা/ঘাস* Leaves/grass<br><br>2.*কাঠি* Twigs / sticks |

| | | | |
|---|---|---|---|
| | | | 3. *কাপড়* Rag or cloth<br>4. *পাথর* Stones<br>5. *স্বাস্থ সম্মত কাগজ*Hygienic (toilet) paper<br>6. *পানির পাত্র/বদনা/মগ*Water container / vessel<br>7. *পানির ট্যাপ* Water tap<br>8. *সাবান* Soap<br>9. *ছাই অথবা মাটি* Ash or soil for cleansing<br>10. *খবরের কাগজ* Newspaper<br>11. *কিছুই নাই* Nothing |
| C. 809 | | আপনার খানার কে কে এই পায়খানা/টয়লেটটি পায়খানার জন্য ব্যবহার করে? Please tell me about who in your family uses the latrine for defecation. | |
| 1 | 1 | <**3**বাচ্চা Children <3 | 1. সব সময় [Always]<br>2. প্রায়ই [Usually]<br>3. মাঝেমাঝে [Sometime]<br>4. কখনই না [Never]<br>888. প্রযোজ্য নয় [ N/A] |
| 2 | 2 | ৩-<৮বাচ্চা Children 3-<8 | 1. সব সময় [Always]<br>2. প্রায়ই [Usually]<br>3. মাঝেমাঝে [Sometime]<br>4. কখনই না [Never]<br>888. প্রযোজ্য নয় [ N/A] |
| 3 | 3 | ৮-১৫বাচ্চা Children 8-15 | 1. সব সময় [Always]<br>2. প্রায়ই [Usually] |

| | | | |
|---|---|---|---|
| | | | 3. মাঝেমাঝে [Sometime]<br>4. কখনই না [Never]<br>888. প্রযোজ্য নয় [ N/A] |
| 4 | 4 | পুরম্নষMen | 1. সব সময় [Always]<br>2. প্রায়ই [Usually]<br>3. মাঝেমাঝে [Sometime]<br>4. কখনই না [Never]<br>888. প্রযোজ্য নয় [ N/A] |
| 5 | 5 | মহিলাWomen | 1. সব সময় [Always]<br>2. প্রায়ই [Usually]<br>3. মাঝেমাঝে [Sometime]<br>4. কখনই না [Never]<br>888. প্রযোজ্য নয় [ N/A] |
| C. 810 | | অন্য কোন খানা কি এই পায়খানা ব্যবহার করে?Do you share this toilet with other households? | *1=হ্যাঁ[Yes]*<br>*0=না[No].......skip to 815* |
| C.811 | | কতগুলো খানা মিলে এই পায়খানাটি ব্যবহার করে ? How many HHs use it? | (Don't know=99)<br>I____I I____I |
| C.812 | | এইখানার শিশুসহ আপনারা কতজন এই পায়খানাটি ব্যবহার করেন? How many people including children in your household use this toilet? | I____I I____I |
| C. 813 | | পায়খানার মালিকানা ? (প্রশ্ন করুন) [***Ask***: Who owns the toilet facility?] | শুধুমাত্র ঐ খানার জন্য [Only for the household] 1<br>কয়েকঘর মিলে/ অংশীদার [Shared] 2<br>অন্য কেউ [Someone else] 3<br>পাবলিক [Public]4<br>প্রযোজ্য নয় [Not applicable]...............................................8 |

| | | | |
|---|---|---|---|
| | | | 88 |
| C. 814 | | এই স্থানে এই পায়খানাটি কত বছর ধরে আছে? (জানি না=999) [How long have you had the present latrine in this place? [In years] (Don't know = 99)] | \|_____\|\|_____\| বৎসর Years<br>\|_____\|\|_____\| মাস Months |

**Wash Benefit Module 9 Child defecation and feces disposal practice**

| **Child defecation and feces disposal** | | | |
|---|---|---|---|
| Administer to: All study households | | | |
| C.912 | | \|_____\|\|_____\| [At what age (in months) does a child start using a latrine, if at all?] | |
| C.902 | | \|_____\| [When was the last time your youngest child/infant (<3 years) defecated?] | [Today]<br>[Yesterday]<br>[Before 2 or more days back]<br>[Never/can't remember]....... (skips to 906)<br>[Refused] ………. (skips to 906)<br>88. [Not applicable]......... (skips to 908) |
| | | \|_____\| [Where did the child defecate the last time?]<br>**<u>Note:</u>** [Don't read the answer, encourage by asking if there is anything else until he/she mentions there in nothing else and check all mentioned.] | [Potty (in the courtyard)]<br>[Potty (inside the house)]<br>[Courtyard (without potty)]<br>[Inside the house (without potty)]<br>[Open space outside the front yard<br>Bush/jungle<br>[In toilet] ...... (Skip to 906)<br>Katha/nappy/diaper<br>77. [Other (Please describe)]....................<br>99. [Don't Know].............(Skip to 906) |
| | | \|_____\| What was done to the feces?] | **<u>[Yes].....1, [No]......0</u>** |

<table>
<tr><td></td><td></td><td></td><td>1. [It is left there] (Skip to 906)<br>2. |_____| [Put/rinsed into toilet or latrine]<br>3. |_____| [Put/rinsed into drain or ditch]<br>4. |_____| [Thrown into Bush/jungle<br>5. |_____| [Thrown into garbage]<br>6. |_____| [Thrown into a specific pit for child’s feces]<br>7. |_____| [Buried]<br>77. |_____| [Other (specify)]....<br>99. |_____| [DK] (Skip to 906)</td></tr>
<tr><td></td><td></td><td>|did you handle the feces? <u>Do Not Read Responses. Mark All that Apply</u> (1 = Yes)</td><td>|_____| Hands only (bare hands)<br>|_____| Hands and cloth / paper / leaves<br>|_____| Scrap material to scoop fece<br>|_____| Potty<br>|_____| Local agricultural hoe/instrument<br>|_____| Sani-scoop<br>|_____| Did nothing<br>77 |_____|Others (specify)___________<br>99 |_____|Don’t know / not sure</td></tr>
<tr><td>C.905.b</td><td>905.a</td><td>How long after defecation did you dispose of the feces?</td><td>|____|Hour<br>|____|Minute</td></tr>
<tr><td></td><td></td><td>[Who disposes your under 3 child’s feces?</td><td><u>[Yes].....1, [No]......0</u><br>|_____| [Mother]<br>|_____| [Father]<br>|_____| [Sister]<br>|_____| [Brother]<br>|_____| [Aunt/grandmother]</td></tr>
</table>

| | | | |
|---|---|---|---|
| | | | \|_____\| [Uncle/grantfather]<br><br>\|_____\| **[**Nobody dispose**]**<br><br>77. \|_____\| Others (Specify) ........................<br><br>888. \|_____\| [Not applicable] |
| | | **[**How far away is this disposal site to the kitchen?] | \|_____\|[steps]<br><br>**888.** [Not applicable] |
| | | \|_____\| [Is there any other child between the age of 3-<8 years?] | 1 [Yes]<br><br>0. [No].......... (skips to 913) |
| | | আপনার খানায় ৩-<৮ বছরের যে শিশুটি আছে তার বয়স কত (মাসে)? (একাধিক শিশু থাকলে সবচেয়ে বড় বাচ্চার ক্ষেত্রে জিজ্ঞেস করুন) | ........................... বছর<br><br>........................... মাস |
| | | \|_____\| [Where did your oldest child (aged 3-7) defecate the last time?] | [Potty (in the courtyard)]<br><br>[Potty (inside the house)]<br><br>[Courtyard (without potty)]<br><br>[Inside the house (without potty)]<br><br>[Open space outside the front yard<br><br>Bush/jungle<br><br>[In toilet] ... (Skip to 913)<br><br>Katha/nappy/diaper<br><br>77. [Other (Please describe)]....................<br><br>99. [Don’t Know].............. (Skip to 913) |
| | | \|_____\| [What is done to the feces?] | **হ্যাঁ [Yes].....1, না [No]......0**<br><br>1. **যেখানে পায়খানা করেছিল সেখানেই ফেলে রাখা হয়েছিল** [ওঃ রং ষবভঃ ঃযবৎব]**৯১২.ধ নং প্রশ্নে চলে যান** (Skip to 912.a)<br><br>2. \|______\| **টয়লেট/পায়খানার ভিতরে ফেলা/ধোয়া হয়েছিল** [Put/rinsed into toilet or latrine]<br><br>3. \|______\|**ড্রেনে/নর্দমার ভিতরে ফেলা/ধোয়া হয়েছিল** [Put/rinsed into drain or ditch] |

| | | | |
|---|---|---|---|
| | | | 4. \|_____\|ঝোপ-ঝাড়ে/জঙ্গলে ফেলা হয়েছিল [Thrown into Bush/jungle<br><br>5. \|_____\|ময়লা আবর্জনার মধ্যে ফেলা হয়েছিল [Thrown into garbage]<br><br>6. \|_____\|নিদির্ষ্ট গর্তে ফেলা হয়েছিল [Thrown into a specific pit for child's feces]<br><br>7. \|_____\|মাটির নীচে পুতে ফেলা হয়েছিল [Buried]<br><br>77. \|_____\|অন্যান্য (বর্ণনা লিখুন) [Other (specify)]................<br><br>99. \|_____\| জানি না [DK]৯১৩ নং প্রশ্নে চলে যান (Skip to 913) |
| | | আপনি কিভাবে পায়খানা হ্যান্ডলিং (ব্যবস্থাপনা )করেন?How did you handle the feces? ☝ Do Not Read Responses. Mark All that Apply (1 = Yes) | ☐ 1 নগ্ন/খালি হাতে Hands only (bare hands)<br>☐ 2 হাতে এবং কাপড়/পাতা/কাগজ Hands and cloth / paper / leaves<br>☐ 3 ফেলানো সামগ্রী মাটিতে পুঁতে Scrap material to scoop feces<br>☐ 4 পটি Potty<br>☐ 5 স্থানীয় কৃষিকাজ-এর হাতিয়ার Local agricultural hoe/instrument<br>☐ 6 সেনি স্কুপ Sani-scoop<br>☐ 7 কোন কিছুই করা হয় না Did nothing<br>☐ 77 অন্যান্য (উলেস্নখ করুন) Others (specify)____________<br>☐ 99 জানি না Don't know / not sure |
| C.910.b | 912.a | পায়খানা করার পর কতক্ষণ এই পায়খানা এখানে পরে ছিল? How long after | \|_____\|Hour ঘনটা<br><br>\|_____\|Minute মিনিট |

| | | |
|---|---|---|
| | | defecation did you dispose of the feces? |

| | | | |
|---|---|---|---|
| Child Potty Use<br><br>( শিশুর পটি ব্যবহার সম্পর্কীত তথ্য) | | | |
| Administer to: All study households ( সকল খানায় হবে) | | | |
| C.815 | | Does your household have a potty that children use for defecation? আপনার খানাতে কি শিশুদের পায়খানা করার পটি আছে? | 1 Yes হ্যাঁ<br><br>2 No না**(Skip to 919)** (919নং প্রশ্নে যান)<br><br>99 Don't know / not sure জানি না **(Skip to 919)** (919নং প্রশ্নে যান) |
| C.816 | | In the last week, how often did your child use the potty? গত এক সপ্তাহের মধ্যে, শিশুটি কতবার পটি ব্যবহার করছে?<br><br>[ *If multiple children use the potty, ask about the youngest child* ][যদি একাধিক শিশু ব্যবহার করে থাকে তবে যে সবচেয়ে বেশী ব্যবহার করে সেই শিশুর জন্য প্রশ্ন করুন] | 1 Every time প্রতিবার<br><br>2 More than half of all defecation events, but not every time যতবার পায়খানা করেছে তার ৫০ ভাগ বা তার বেশী কিন্তু প্রতিবার নয়<br><br>3 যতবার পায়খানা করেছে তার ৫০ ভাগের কম কিন্তু মাঝেমাঝে Less than half of all defecation events<br><br>4 আগে করেছি কিন্তু এখন আর করি না Used to use it, but no longer use it **(Skip to 919)** (919নং প্রশ্নে যান)<br><br>5 কখনই না Never **(Skip to 919)** (919নং প্রশ্নে যান)<br><br>99 জানি না Don't know / not sure **(Skip to 919)** (919নং প্রশ্নে যান) |
| C.817 | | What is the age of the child (or children) who is using the potty? যে শিশুটি (বা শিশুরা) পটি ব্যবহার করছে তার বয়স কত? | 1 ☐ < 1 year ১ বছরের কম<br>2 ☐ 1- < 3 years ১-<৩ বছর<br>3 ☐ 3- < 5 years ৩-<৫ বছর<br>4 ☐ >= 5 years>=৫ বছর<br><br><u>Mark All that Apply.</u> Yes = 1 হ্যাঁ = 1, না = 0 |

| | | | |
|---|---|---|---|
| C.818 | | Ask the respondent to identify the place where the child typically uses the potty. Then ask her to retrieve the potty<br><br>Could I please see the potty?<br><br>👁 Observation: *Record how long it takes to produce the potty.*<br><br>আমি কি পটিটি দেখতে পারি<br><br>👁আমি কি পটিটি দেখতে পারি (পটিটি আনতে কতটুকু সময় লেগেছে তা সেকেন্ডে গণনা করুন) | ☐☐:☐☐MINUTES : SECONDS (99:99 if could not measure) মিনিট : সেকেন্ড(পটি দেখাতে না পারলে 999 কোড করুন) |
| C.819 | | 👁 Observation: Potty condition.<br><br>পটিটির অবস্থা কি তা পর্যবেক্ষণ করুন (একাধিক উত্তর হতে পারে) | Mark All that Apply. Yes = 1 হ্যাঁ = 1, না = 0<br><br>1.Easily accessible when needed by the child প্রয়োজনের সময় বাচ্চা যেন পায় এমন কাছাকাছি জায়গাে রয়েছে<br><br>2.Easily accessible when needed by the mother প্রয়োজনের সময় মা যেন পায় এমন কাছাকাছি জায়গাতে র<br><br>3.Visible signs of feces inside / on the potty / removable pot ভিতরে/ পটিতে পায়খানার চিহ্ন আছে<br><br>4.Potty was covered with the lid পটি লিডদিয়ে ঢাক<br><br>5. Potty was covered with anything other than th পটি লিডছাড়াঅন্য কিছু দিয়ে ঢাকা ছিল<br><br>6.Dry শুকনা<br><br>7.Broken/or missing parts(tray) so that it is unusableভেঙ্গে গেছে তাই ইহা ব্যবহার করা সম্ভব না<br><br>8.Covered in dust / signs of non-use পটি ধুলায় ঢা ব্যবহার না করার চিহ্ন রয়েছে<br><br>9.Cannot produce a potty পটি দেখাতে পারে নাই |
| C.820 | | ☐Where do you usually dispose of feces from the potty? সাধারনতঃ এই পটি থেকে আপনি পায়খানা কোথায় ফেলেন? | 1 Latrine পায়খানায়/টয়লেটে<br><br>2 Open Pit / separate pit for child or animal fecesখোলা গর্তে/ শিশুর বা পশুপাখির |

| | | | |
|---|---|---|---|
| | | | পায়খানা ফেলার গর্তে<br><br>3 Bury it / Covered Pit মাটির নীচে পুতে ফেলা হয়েছিল/ ঢাকা গর্তে<br><br>4 Undefined open site near the compound (including open garbage disposal sites / dumps) উঠানের কাছে খোলা জায়গা<br><br>5 Bush / forest / fieldঝোপ-ঝাড়ে/জঙ্গলে/মাঠে<br><br>6 Nearby water (pond, canal, river) পানির উৎসের কাছে (পুকুর/সেচ নালা/নদী)<br><br>77 Other (specify) অন্যান্য (লিখুন)_______________ |
| Sani Scoop Use<br><br>**(সেনিস্কুপ ব্যবহার)** | | | |
| Administer to: All households ( সকল খানায় হবে) | | | |
| C.821 | | ☐Does your household have a dedicated tool [sani scoop] to clean up feces around your household?<br><br>আপনার খানার চারপাশের পায়খানা পরিষ্কার করার জন্য আপনার খানায় কোন নির্দিষ্ট কিছু আছে কি? | 1 Yes হ্যাঁ<br><br>2 No না (1001নং প্রশ্নে যান)**(Skip to 1001)**<br><br>99 Don't know / not sure জানি না (1001 নং প্রশ্নে যান)**(Skip to 1001)** |
| C.822 | | How often do you use the [sani scoop]? আপনি কখন কখন পায়খানা পরিষ্কার করার জন্য কোদাল/সেনিস্কুপ ব্যবহার করেন? | 1 Multiple times per day প্রতিদিন কয়েকবার<br><br>2 Once per day প্রতিদিন একবার<br><br>3 A few times each week প্রতি সপ্তাহে কয়েকবার<br><br>4 Less than once per week সপ্তাহে একবারের কম<br><br>5 Used to use it, but no longer use it প্রথমদিকে ব্যবহার করেছে কিন্তু এখন আর করে না(1001 নং প্রশ্নে যান)**(Skip to 1001)**<br><br>6 Never কখনই না(1001 নং প্রশ্নে যান)**(Skip to 1001)** |

| | | | |
|---|---|---|---|
| C.823 | | What do you use the [sani scoop] for? ☟Do Not Read Responses.<br><br>আপনি কি কি কাজেকোদাল/সেনিস্কুপ ব্যবহার করেন? (একাধিক উত্তর হতে পারে)☟ উত্তর পড়ে শোনাবেন না | Mark All that Apply. Yes = 1 হ্যাঁ = 1, না = 0 |

| | | |
|---|---|---|
| 1 | ☐ | Clean up animal feces জীবজন্তুর পায়খানা |
| 2 | ☐ | Clean up child feces বাচ্চার পায়খানা ফেলা |
| 3 | ☐ | Clean up garbage ময়লা আবর্জনা ফেলার ক |
| 4 | ☐ | Take the scoop to the field (for work) কৃ |
| 5 | ☐ | Digging / gardening গর্ত তৈরীর কাজে |
| 77 | ☐ | Other (specify) অন্যান্য (বর্ণনা লিখুন)_____ |

| | | | |
|---|---|---|---|
| C.824 | | Could I please see the [sani scoop]? 👁 Observation: Sani scoop condition.<br><br>আমি কি কোদাল/সেনিস্কুপটি দেখতে পারি? (একাধিক উত্তর হতে পারে)<br><br>👁পর্যবেৰণ:সেনিস্কুপটির অবস্থা পর্যবেৰণ করম্নন | Mark All that Apply. Yes = 1 হ্যাঁ = 1, না = 0 |

| | | |
|---|---|---|
| 1 | ☐ | Visible signs of feces on the sani scoop<br>কোদাল/সেনিস্কুপে পায়খানার চিহ্ন আছে |
| 2 | ☐ | Dry শুকনা |
| 3 | ☐ | Broken and needs repair<br>ভেঙ্গে গেছে এবং মেরামত করা দরকার |
| 4 | ☐ | Easily accessible when needed by an ad<br>প্রয়োজনের সময় বড়রা যেন পায় এমন কাছাকাছি রয়েছে |
| 5 | ☐ | Signs that the sani scoop is not used<br>কোদাল/সেনিস্কুপ ব্যবহার না করার চিহ্ন রয়েছে |
| 6 | ☐ | Cannot produce a sani scoop<br>কোদাল/সেনিস্কুপ দেখাতে পারে নাই |

| | | | |
|---|---|---|---|
| C.825 | | I'd like to ask a few more questions about the types of feces you pick up with the [sani scoop] and how you dispose of the different types of feces.<br><br>[Ask about each type of feces separately.].<br><br>কোদাল/সেনিস্কুপ দিয়ে আপনি যেসব | Where do you dispose the feces?<br><br>পায়খানা কোথায়ফেলা হয়েছিল?<br><br>☟ Do Not Read Responses. উত্তর পড়ে শোনাবেন<br><br>Mark All that Apply (একাধিক উত্তর হতে পারে) 1<br><br>88 Not applicable প্রযোজ্য নয় |

| | A | B | C | D | E | F |
|---|---|---|---|---|---|---|

| | | | Feces পায়খানা | Use Sani Scoop for disposal? পায়খানা ফেলার জন্য কোদাল/সেনিস্কুপ ব্যবহার করে? | Latrine পায়খানায় | Open Pit খোলা গর্তে | Bush/Farm ঝোপে ঝাড়ে/খোলা জায়গায় | Surface Water পানিতে | Dig Hole and [illegible] গর্ত করা হয় এ[illegible] |
|---|---|---|---|---|---|---|---|---|---|
| | | পায়খানা ফেলেন এবং কিভাবে ফেলেন সেসম্পর্কে জানার জন্য এখন আমি আপনাকে আরো কিছু প্রশ্ন করতে চাই।<br>(প্রতিটি পায়খানার ধরনের কথা উলেস্নখ করে কোড করম্নন)<br>(শিশু/পশুপাখির পায়খানা ফেলার কাজে ব্যবহার না করলে প্রযোজ্য নয় ঘরটিতে টিক দিন) | | | | | | | |
| | | | 1 Child (<8 years) শিশুর (<৮ বছর) | ☐ | | | | | |
| | | | 2 Cow গোবর | ☐ | | | | | |
| | | | 3 Poultry / pigeons হাঁস-মুরগীর | ☐ | | | | | |
| | | | 4 Goat ছাগলের | ☐ | | | | | |
| | | | 5 Pig শুকরের | ☐ | | | | | |
| | | | 6 Dog | ☐ | | | | | |

| | | | | |
|---|---|---|---|---|
| | | | or cat<br>কুকুর বা বিড়ালের | |
| | | | 88 Not applicable | ☐ |

**Wash Benefit Module 10 Water Access**

আপনাকে ধন্যবাদ। আমি আপনাকে এবং আপনার খানার সদস্যদের বাড়ীতে খাবার পানি বিষয়ে কিছু প্রশ্ন করতে চাই।

Thank you very much. Now, I would like to ask you some questions about the water you and your family drink at home.

| WATER TREATMENT, STORAGE, AND QUALITY (All households) | | | |
|---|---|---|---|
| SAY | Thank you very much. Now, I would like to ask you some questions about the water you and your family drink at home. | | |
| C.1001 | 1001. | আপনি কিভাবে খাবার পানি সংরক্ষন করেন ? [How do you store drinking water?<br>👁প্রশ্ন করুন এবং সম্ভব হলে পর্যবেক্ষন করুন ASK THE QESTION AND OBSERVE IF POSSIBLE | [1] প্লাষ্টিক অথবা ধাতব পাত্রে IN PLASTIC OR METAL CONTAINERS<br>[2] মাটির পাত্রে IN CLAY POTS<br>[3] ছাদে পানির ট্যাংক ROOF TANK OR CISTERN<br>[4] পানি সংরক্ষণ করা হয় নাDO NOT STORE WATER |
| C.1002 | 1002. | এই খানায় ০-<৩৬ মাসের কোন শিশু আছে কি যে পানি পান করে ?Is there a child 0-<36 months in this household who drinks water? | [1] হ্যাঁ YES<br>[2] না NO<br>[88] Not applicable প্রযোজ্য নয় |
| C.1003 | 1003. | **IF YES→প্রশ্ন করুন এবং পর্যবেক্ষন করুনঃ যদি আপনার শিশু (<৩ বছর) এখন পানি খেতে চাইত তাহলে আপনি কিভাবে তাকে পানি দিতেন তা আমাকে দেখান**ASK: If <u>your child 0-3 years</u> wanted a drink of water right now, could you show me how you would give it to him/her?<br>**IF NO→** ASK: **যদি আপনি এখন পানি খেতে চাইতেন তাহলে আপনি কিভাবে পানি নিতেন তা আমাকে দেখান**If <u>you</u> wanted a drink of water right now, could you show me how you would get it?<br>উত্তরদাতার পানি দেওয়ার প্রক্রিয়াকে পর্যবেক্ষণ/অনুসরণ করুন । যদি অন্য ঘরে পানি সংরক্ষিত এবং সেখানে প্রবেশাধিকার থাকে তাহলে প্রক্রিয়াকে পর্যবেক্ষণ/অনুসরণ করুন । অন্যথায় ৯৯ কে গোলাকার করতে হবে। ATTEMPT TO FOLLOW RESPONDENT WHEN SHE GOES TO RETRIEVE THE WATER. IF WATER IS STORED IN ANOTHER ROOM, ASK HER IF IT IS OKAY TO ENTER THAT ROOM. CIRCLE "99" IF YOU CANNOT OBSERVE. | |

| | | | | | |
|---|---|---|---|---|---|
| | | 👁 উত্তরদাতা পানি নেওয়ার আগে কি করেছিল ?WHAT DID RESPONDENT DO <u>BEFORE TAKING THE WATER</u>? | হ্যাঁ YES | না NO | জানি নাD/K |
| 2 | 2 | গ্লাস বা পাত্রটিতে পানি ঢালার পূর্বে গ্লাস/পাত্রটি খাবার পানি দিয়ে ধুয়েছিলRINSED GLASS/CONTAINER WITH DRINKING WATER BEFORE FILLING | [1] | [2] | [99] |
| 3 | 3 | পানি ঢালার পূর্বে হাত ধুয়েছিলWASHED HANDS WITH WATER BEFORE DRINKING WATER WAS OBTAINED | [1] | [2] | [99] |
| 4 | 4 | পানি ঢালার পূর্বে সাবান দিয়ে হাত ধুয়েছিলWASHED HANDS WITH SOAP BEFORE DRINKING WATER OBTAINED | [1] | [2] | [99] |
| | | 👁 উত্তরদাতা কোথা থেকে পানিসংগ্রহ করেছিল?<u>FROM WHERE</u> DID THE RESPONDENT TAKE THE WATER? | হ্যাঁ YES | না NO | জানি না D/K |

| | | | | | |
|---|---|---|---|---|---|
| 5 | 5 | সরাসরি পানির উৎস থেকে BROUGHT DIRECTLY FROM WATER SOURCE | [1] | [2] | [99] |
| 6 | 6 | সরাসরি খোলা পানির পাত্র থেকে BROUGHT IRECTLY FROM uncoveredSTORAGE CONTAINER | [1] | [2] | [99] |
| 7 | 7 | সরাসরি পানির ফ্লিটার থেকে BROUGHT DIRECTLY FROM WATER FILTER | [1] | [2] | [99] |
| | 8 | সরাসরি ঢেকে রাখা পানির পাত্র থেকে STORED WATER WAS COVERED | [1] | [2] | [99] |
| | | 👁 উত্তরদাতা কিভাবে গ্লাসে/কাপে/পাত্রে পানি দিয়েছিল? HOW DID SHE GET THE WATER INTO THE CUP? | হ্যাঁ YES | না NO | জানি না D/K |
| 9 | 9 | গ্লাস বা পাত্রটিতে ঢালা পানির ভিতরে তার হাত বা হাতের আঙ্গুল লেগেছিল কি?HANDS TOUCHED / CONTACTED THE DRINKING WATER | [1] | [2] | [9 ] |
| 10 | 10 | গ্লাস বা পাত্রটিতে পানি ভরার সময় তা পানির পাত্রের ভিতর ডুবিয়ে পানি ভরা হয়েছিলCONTAINER/GLASS DIPPED INTO WATER CONTAINER | [1] | [2] | [99] |
| 11 | 11 | গ্লাসে পানি ভরার জন্য লম্বা হাতলযুক্ত কোন চামচ/মগ ব্যবহার করা হয়েছিলLADLE USED TO OBTAIN WATE | [1] | [2] | [99] |
| 12 | 12 | পানির পাত্র থেকে সরাসরি পানি ঢেলেছিলWATER POURED FROM CONTAINER | [1] | [2] | [99] |
| 13 | 13 | সরাসরি টিউবওয়েল/ পানির উৎস থেকে এনেছেWATER POURED FROM TAP | [1] | [2] | [99] |

| | | | | |
|---|---|---|---|---|
| C.1004 | 1004. | আপনি/এ খানার অন্যকেউ কতক্ষণ আগে এই সংরক্ষণকৃত পানি সংগ্রহকরেছেন? How long ago did you or somebody in your home collect this water? ( 99 = Don't knowজানি না; 87 = No Water কোন পানি নাই, 88 Not applicableপ্রযোজ্য নয় ) If 88, skip to 1007 | A.\|__\|__\| ঘন্টা HOURS<br>B.\|__\|__\| দিনDAYS | |
| C.1005 | 1005. | এই খাবার পানিনিরাপদ করার জন্য কোন কিছু করেছেন কি?Have you done anything to make this water less cloudy or safer to drink? | [1] হ্যাঁYES<br>[2] না NO<br>[99] জানি না DON'T KNOW/NOT SURE | → **SKIP to 1007**<br>→ **SKIP to 1007** |
| C.1006 | 1006. | কোন পদ্ধতি কি আপনি ব্যবহার করেন? What method(s) did you use?<br>উত্তর পড়ে শুনাবেন না।উত্তরদাতাকে উত্তর বলতে সাহায্য করা যাবে তার উত্তর দেয়া শেষ হলে।DO NOT READ RESPONSES. PROBE UNTIL RESPONDENT IS FINISHED. CIRCLE ALL THAT APPLY. | | |
| 1 | 1 | আকুয়াট্যাবসAquatabs | [1] হ্যাঁYes [0] না No | |
| 2 | 2 | ক্লোরিন (তরল) মিশিয়ে নেই (ওয়াটার গার্ড নয়)Waterguard / bottled chlorine | [[1] হ্যাঁYes [0] না No | |
| 3 | 3 | পানি ফুটিয়ে নেইBoil | [1] হ্যাঁYes [0] না No | |
| 4 | 4 | কাপড়ে ছেকে নেইStrain it through clothor other material | [1] হ্যাঁYes [0] না No | |
| 5 | 5 | পানির ফিল্টার ব্যবহার করে (সিরামিক, বালু ইত্যাদি) Water filter [ceramic, sand, composite] | [1] হ্যাঁYes [0] না No | |

| | | | | |
|---|---|---|---|---|
| 6 | 6 | সূর্যের আলোতে জীবানুমুক্তকরণSolar disinfection (SODIS) | [1] হ্যাঁYes [0] না No | |
| 7 | 7 | থিতিয়ে নেই Let it stand and settle | [1] হ্যাঁYes [0] না No | |
| 8 | 8 | স্যান্ড ফিল্টার এর মাধ্যমে বিশুদ্ধ করে নেইBiosand filter | [1] হ্যাঁYes [0] না No | |
| 9 | 9 | ফিটকিরি যোগ করে Coagulant (alum) | [1] হ্যাঁYes [0] না No | |
| 10 | 10 | পার PUR (flocculant + disinfectant) | [1] হ্যাঁYes [0] না No | |
| C.1007 | 1007. | আপনি কখনও আপনার খাবার পানি পরিশোধন করার জন্য কিছু করেছিলেন কি?Do you ever treat your drinking water or do anything to make it less cloudy? | [1] হ্যা YES<br>[0] নাNO | → 1010 নং এ চলে যান SKIP TO 1010 |
| C.1008 | 1008. | সর্বশেষ কখন আপনিআপনার খাবার পানি পরিশোধন করেছিলেন ? When was the last time you treated your water or did anything to make it less cloudy?<br><br>উত্তর পড়ে শুনানো যাবে না DO NOT READ RESPONSES | [1] আজ TODAY<br>[2] গতকালYESTERDAY<br>[3] গত সপ্তাহের মধ্যে WITHIN THE PAST WEEK<br>[4] গত দুই সপ্তাহের মধ্যে WITHIN THE PAST 2 WEEKS<br>[5] গত মাসের মধ্যে WITHIN THE PAST MONTH<br>[6] গত বছরের মধ্যে WITHIN THE PAST YEAR<br>[99] জানি না/নিশ্চিত না DON'T KNOW/NOT SURE | |
| C.1009 | 1009. | কোন কোন উপায়ে আপনি পানি পরিশোধন করে থাকেন ? What are <u>all the ways</u> you treat your drinking water?<br>উত্তর পড়ে শুনানো যাবে না । উত্তর দেয়া শেষ হলে উত্তর পেতে সাহায্য করা যেতে পারে ।DO NOT READ RESPONSES. PROBE UNTIL RESPONDENT IS FINISHED. CIRCLE ALL THAT APPLY. | | |
| 1 | 1 | আকুয়াট্যাবসAquatabs | [1] হ্যাঁYes [0] না No | |
| 2 | 2 | ক্লোরিন (তরল) মিশিয়ে নেই (ওয়াটার গার্ড নয়)Waterguard / bottled chlorine | [[1] হ্যাঁYes [0] না No | |
| 3 | 3 | পানি ফুটিয়ে নেইBoil | [1] হ্যাঁYes [0] না No | |
| 4 | 4 | কাপড়ে ছেকে নেইStrain it through clothor other material | [1] হ্যাঁYes [0] না No | |
| 5 | 5 | পানির ফিল্টার ব্যবহার করে (সিরামিক, বালু ইত্যাদি) Water filter [ceramic, sand, composite] | [1] হ্যাঁYes [0] না No | |
| 6 | 6 | সূর্যের আলোতে জীবানুমুক্তকরণSolar disinfection (SODIS) | [1] হ্যাঁYes [0] না No | |
| 7 | 7 | থিতিয়ে নেই Let it stand and settle | [1] হ্যাঁYes [0] না No | |
| 8 | 8 | স্যান্ড ফিল্টার এর মাধ্যমে বিশুদ্ধ করে নেইBiosand filter | [1] হ্যাঁYes [0] না No | |
| 9 | 9 | ফিটকিরি যোগ করে Coagulant (alum) | [1] হ্যাঁYes [0] না No | |
| 10 | 10 | PUR (flocculant + disinfectant) | [1] হ্যাঁYes [0] না No | |

| পানের পানির উৎস **DRINKING WATER SOURCES (ALL HOUSEHOLDS)** | | | |
|---|---|---|---|
| উত্তরদাতাকে তার **পানির প্রধান এবং দ্বিতীয় উৎস বিষয়ে প্রশ্ন করুন** ASK THE FOLLOWING QUESTIONS ABOUT THE RESPONDENT'S PRIMARY AND SECONDARY WATER SOURCES. | | | |
| C.1016 | 1010. | আপনার খানায়ব্যবহৃত পানির প্রধান উৎস কি ?What type of water source does your household collect most of the water that you use from? | 1 অগভীর টিউবওয়েল Shallow tubewell<br>2 গভীর টিউবওয়েল Deep tubewell<br>3 ঘরের ভিতর ট্যাপ বা পাইপের পানি Piped into home<br>4 উঠানে ট্যাপ বা পাইপের পানি Piped into yard<br>5 গভীর পাতকূয়া (উঠানে)Borewell in yard<br>6 পাবলিক ট্যাপ Public tap<br>7 পাবলিক গভীর পাতকূয়া Public borewell<br>8 অগভীর পাকা পাতকূয়া Shallow well with concrete reinforcement<br>9 অগভীর কাঁচা পাতকূয়া Shallow well (no concrete)<br>10 নদী/বাধ/লেক/পুকুর/সেচ নালাথেকে সরাসরি সংগৃহীত পানি River, lake, stream<br>11 অন্যান্য (লিখুন) Other: _____________ |
| C.1017 | 1011. | আপনি আমাকে বলবেন আপনার বাড়ী থেকে প্রধান পানির উৎসে একবার যেতে কতক্ষণ সময় লাগে? Can you tell me how long it takes you to walk one-way to this PRIMARY water source from your home? | \|__\|ঘন্টা Hours<br>\|__\|মিনিট Min |
| C.1018 | 1012. | আপনি আমাকে বলবেন কোন কোন কাজের জন্য এই উৎসের পানি ব্যবহার করেন Can you please tell me whether you use water from this source for… | |
| 1 | 1 | পানি পান Drinking | [1] হ্যাঁ Yes [0] না No |
| 2 | 2 | রান্না-বান্না/বাসন ধোয়া Cooking/ washing dishes | 1] হ্যাঁ Yes [0] না No |
| 3 | 3 | কাপড় ধোয়া Laundry | 1] হ্যাঁ Yes [0] না No |
| 4 | 4 | গোসল/হাতধোয়া Bathing/hand washing | 1] হ্যাঁ Yes [0] না No |
| 5 | 5 | কৃষিকাজ/বাগান Irrigate crop land or garden | 1] হ্যাঁ Yes [0] না No |
| 6 | 6 | পশু-পালন Water for livestock | 1] হ্যাঁ Yes [0] না No |
| C.1019 | 1013. | আপনার খানাকে এই পানির উৎস ব্যবহারের জন্য কি কোন প্রকার টাকা দিতে হয়? your household pay any money to use this source? | [1] হ্যাঁ Yes [0] না No |
| C.1020 | 1014. | আপনার খানায় পানি ব্যবহারের জন্য কি ২য় কোন উৎস ব্যবহার করতে হয়? যদি হয় তাহলে কি ধরনের ? Does your household currently | 1. অগভীর টিউবওয়েল Shallow tubewell<br>2. গভীর টিউবওয়েল Deep tubewell<br>3. ঘরের ভিতর ট্যাপ বা পাইপের পানি Piped into home<br>4. উঠানে ট্যাপ বা পাইপের পানি Piped into yard<br>5. গভীর পাতকূয়া (উঠানে)Borewell in yard |

| | | | |
|---|---|---|---|
| | | source? If so, what type is it? | 7. পাবলিক গভীর পাতকূয়া Public borewell<br>8. অগভীর পাকা পাতকূয়া Shallow well with concrete reinforcement<br>9. অগভীর কাঁচা পাতকূয়া Shallow well (no concrete)<br>10. নদী/বাধ/লেক/পুকুর/সেচ নালাথেকে সরাসরি সংগৃহীত পানি River, lake, stream<br>11. অন্যান্য (লিখুন) Other: _____________<br>88] ২য় কোন পানির উৎস ব্যবহার করতে হয় না Don't use any secondary water source (Skip to 1017.a) |
| C1021 | 1015. | আপনি আমাকে বলবেন আপনার বাড়ী থেকে ২য় পানির উৎসে একবার যেতে কতক্ষণ সময় লাগে? Can you tell me how long it takes you to walk one-way to this SECONDARY water source from your home? | \|__\|ঘনটা Hours<br>\|__\|মিনিট Min |
| C1022 | 1016. | আপনি আমাকে বলবেন কোন কোন কাজের জন্য এই উৎসের পানি ব্যবহার করেন Can you please tell me whether you use water from this source for… | |
| 1 | 1 | পানি পান Drinking | [1] হ্যাঁ Yes [0] না No |
| 2 | 2 | রান্না-বান্না/বাসন ধোয়া Cooking/ washing dishes | 1] হ্যাঁ Yes [0] না No |
| 3 | 3 | কাপড় ধোয়া Laundry | 1] হ্যাঁ Yes [0] না No |
| 4 | 4 | গোসল/হাতধোয়া Bathing/hand washing | 1] হ্যাঁ Yes [0] না No |
| 5 | 5 | কৃষিকাজ/বাগান Irrigate crop land or garden | 1] হ্যাঁ Yes [0] না No |
| 6 | 6 | পশু-পালন Water for livestock | 1] হ্যাঁ Yes [0] না No |
| C1023 | 1017. | আপনার খানাকে এই পানির উৎস ব্যবহারের জন্য কি কোন প্রকার টাকা দিতে হয়? your household pay any money to use this source? | 1] হ্যাঁ Yes [0] না No |
| | 1017.a | আপনার খানার খাবার পানির প্রধান উৎস কি? What is your main drinking water source? | 1. অগভীর টিউবওয়েল Shallow tubewell<br>2. গভীর টিউবওয়েল Deep tubewell<br>3. ঘরের ভিতর ট্যাপ বা পাইপের পানি Piped into home<br>4. উঠানে ট্যাপ বা পাইপের পানি Piped into yard<br>5. গভীর পাতকূয়া (উঠানে)Borewell in yard<br>6. পাবলিক ট্যাপ Public tap<br>7. পাবলিক গভীর পাতকূয়া Public borewell<br>8. অগভীর পাকা পাতকূয়া Shallow well with concrete reinforcement<br>9. অগভীর কাঁচা পাতকূয়া Shallow well (no concrete)<br>10. নদী/বাধ/লেক/পুকুর/সেচ নালাথেকে সরাসরি সংগৃহীত পানি River, lake, |

| | | | |
|---|---|---|---|
| | | | stream<br>11. অন্যান্য (লিখুন) Other: ____________<br>88] Not applicable |
| C1024 | 1018. | আপনার খানা কি কখনও বৃষ্টির পানি সংগ্রহ করত ? Does your household collect rainwater? | 1] হ্যাঁ Yes [0] না No |
| C1025 | 1019. | আপনার খানা কি বর্তমানে বৃষ্টির পানি সংগ্রহ করে ? Is your household currently collecting rainwater? | 1] হ্যাঁ Yes [0] না No if no, skip to 1301 |
| C1026 | 1020. | আপনি আমাকে বলবেন কি আপনার খানা কি কি কাজে বৃষ্টির পানি ব্যবহার করে ? Can you please tell me whether you use rainwater for | |
| 1 | 1 | পানি পান Drinking | [1] হ্যাঁ Yes [0] না No |
| 2 | 2 | রান্না-বান্না/বাসন ধোয়া Cooking/ washing dishes | 1] হ্যাঁ Yes [0] না No |
| 3 | 3 | কাপড় ধোয়া Laundry | 1] হ্যাঁ Yes [0] না No |
| 4 | 4 | গোসল/হাতধোয়া Bathing/hand washing | 1] হ্যাঁ Yes [0] না No |
| 5 | 5 | কৃষিকাজ/বাগান Irrigate crop land or garden | 1] হ্যাঁ Yes [0] না No |
| 6 | 6 | পশু-পালন Water for livestock | 1] হ্যাঁ Yes [0] না No |
| **Skip to 1301** | | | |
| ~~C.1027.~~ | ~~1021.~~ | ~~আপনি ভেবে বলবেন কি আপনার পরিবার মোট সকল উৎস থেকে দৈনিক আনুমানিক ২০-লিটার পাত্রের কতটি পাত্র পানি পায়? ***উত্তরদাতাকে উত্তর দিতে অনুপ্রেরনা দেন*** জানা না থাকলে ৯৯৯ বসান(দূরত্ব আধা কিমি হলে ০.৫ লিখতে হবে)। Thinking about ALL the water that your household obtains from ALL sources, approximately how many 20-litre containers of water would you say that your household uses IN TOTAL EACH DAY?~~<br><br>~~***Encourage respondent to estimate. "Don't know"=999. Use decimals as needed (e.g. one half=0.5)~~<br><br>~~☐দৈনিক ১০-লটার পাত্র 10-liter containers per day~~<br><br>~~☐দৈনিক ২০-লিটার পাত্র 20-liter containers per day~~ | |
| **C.1010.** | 1022. | আপনার খানায় কত ঘন ঘন খাবার পানিতে আকুয়াট্যাবস মেশানো হয়? How frequently do you treat your water using [Aquatabs]? | Do Not Read Responses.<br>1 যতবার পানি সংগ্রহ করা হয়/সবসময়Every time they collect water<br>2 মাঝেমাঝেSometimes / occasionally<br>3 প্রথমদিকে দিয়েছিলাম কিন্তু এখন আর দেই নাTreated water in the beginning [of the program] but not any more<br>4 আকুয়াট্যাবস দিয়ে কখনও পানি পরিশোধন করি নাইNever treated water with [Aquatabs] **(Skip to 1026)**<br>99 জানি না Don't know / not sure |

| | | | |
|---|---|---|---|
| C.10 | 1023. | আপনার খানায় সংরক্ষনকৃত খাবার পানি কি কুয়াট্যাবস দ্বারা পরিশোধন করা হয়েছে?<br>Is the drinking water stored in your household today treated with [Aquatabs]? | 1 হ্যাঁ, সবটুকু পানিYes, all of it<br>2 হ্যাঁ, কিছু পানি Yes, some of it<br>3 নাNot treated<br>4 খানায় কোন পানি নাই No water in the house<br>99 জানি না Don't know |
| C.10 | 1024. | ◎ পর্যবেৰন করম্নন: আকুয়াট্যাবস দ্বারা পরিশোধিত পানির পাত্র দেখাতে বলুন: পানির পাত্র ঢাকা ছিল কী? Ask to see the treated water. Observe: Is the water covered? | 1 হ্যাঁYes<br>2 নাNo<br>3 খানায় পাত্রে কোন পানি নাই No water stored in the house<br>88 প্রযোজ্য নয়/প্রত্যাখ্যান Not applicable / refused |
| C.10 | 1025. | পানি পরিশোধন করার জন্য কত সময় আগেআকুয়াট্যাবস ব্যবহার করা হয়েছিল?<br>Approximately how long ago did you treat the water with [Aquatabs]? | ঘিনটা HH: মনিট MM ☐☐:☐☐ ( 9 9 : 9 9 Don't know ) |

ক্লোরিন অবশেষ পরীক্ষা **Residual Chlorine Test**

সকল খানার জন্য প্রযোজ্য Administer to: All study households

*Ask to collect a water sample from the source identified in C.1003. Explain that some (but not all) of the samples may be tested to see if there is any chlorine in the water.*

| | | | |
|---|---|---|---|
| C.10 | 1026. | ক্লোরিণের জন্য আমি আপনার খাবার পানি পরীৰা করতে পারি কী?<br>May I collect a small sample of your drinking water? | 1 হ্যাঁ Yes<br>2 না/প্রত্যাখান No / refused<br>3 খানায় কোন পানি নাই No drinking water available to test |
| C.10 | 1027. | ক্লোরিন অবশেষ পরীক্ষার জন্য সংরক্ষনকৃত পানি যা নিদ্দিষ্ট বাচচা খায় তার নমুণা সংগ্রহ করম্নন। ভিন্ন ভিন্ন অবস্থানেই খানার সংরক্ষিত খাবার পানির ক্লারিন অবশেষ পরীক্ষা করুন । শুধুমাত্র সেই খানার পানি পরীক্ষা করতে হবে যারা বলবে সংরক্ষিত খাবার পানিতে ক্লারিন মিশিনো হয়েছে(C.1009 = 1, 2, or 11)। যদি এই খানায় সংরক্ষিত খাবার পানিতে ক্লারিন না মেশানো হয়, তাহলে পানি পরীক্ষা থেকে বিরত থাকুন এবং ৮৮ কোড করুন।<br>Collect a small water sample from stored drinking water for target children.<br>Test for free residual chlorinein a discrete | ৮৮৮ ক্লোরিন মেশানো হয়েছে কিনা এ বিষয়ে রিপোর্ট দিতে পারে নি<br>8 88 Did not report that the water was treated with chlorine<br>**999**পরীক্ষা করা হয় নাই<br>9 9 9 Could not test<br>ক্লোরিন অবশেষের পরিমান Level of Free Residual Chlorine: ☐☐.☐ মি.গ্রাম/লিটারmg / L |

| | | | | |
|---|---|---|---|---|
| | | location after you leave the household.<br>Only test samples collect from households who report to have treated their water with some form of chlorine (C.1009 = 1, 2, or 11).If the household did not report treating the water with chlorine, discard the sample without testing it and record 8 88 in the field below. | | |
| খানায় অনুজীবি সংক্রান্ত নমূনা (খানা দৈবচয়িতভাবে নির্বাচিত হবে)**HOUSEHOLD MICROBIOLOGICAL SAMPLE (ADMINISTER TO RANDOM SUBSAMPLE HOUSEHOLD)** | | | | |
| C.1501 | 1028. | ব্যাকটেরিয়া আছে কিনা জানার জন্য আমি আপনার খাবার পানির কিছু পরিমান সংগ্রহ করতে পারি কী?<br>May I collect a small sample of your drinking water to test for bacteria in our office? | [1] হ্যাঁ YES, SAMPLE COLLECTED<br>[2] না NO/REFUSED | <br>→ **SKIP TO C.1033** |
| C.1502 | 1029. | খানার নমুনা সংগ্রহীত হয়েছে<br>HH sample collected (DD/MM/YY, HH:MM) | খানার নমূনা পানির রেকর্ড HH Water Sample Time Log (can be automated in a bar-code scanner) | |

| | DD/MM/YY | HH:MM |
|---|---|---|
| A | ☐☐/☐☐/☐☐<br>খানার পানির নমুনা সংগ্রহের তারিখ HH sample collected | ☐☐:☐☐<br>সময় |
| B | ☐☐/☐☐/☐☐<br>খানার পানির নমুনা ল্যাবে পৌছানোর তারিখ HH sample arrived at lab | ☐☐:☐☐<br>সময় |
| C | ☐☐/☐☐/☐☐<br>খানার পানির নমুনা এনালাইসিসের তারিখ HH sample analyzed | ☐☐:☐☐<br>সময় |

| | | | |
|---|---|---|---|
| c.1503 | 1030. | খানার পানির নমুনা, ফিল্টারকৃত পানির পরিমান (মিলি)<br>HH sample, volume of water filtered (in milliliters) | ☐☐☐মিলি ml |
| C.1504 | 1031. | খানার পানির নমুনার ল্যাব রেজাল্ট, ই.কোলাই (*E. coli)*<br>(5555 = অসংখ্য, 9999 = এনালাইসিস করা হয় নাই<br>Household sample lab result, *E. coli* (5555 = Too numerous to count, 9999 | ☐☐☐☐ (CFU / 100 ml) |

| | | | | |
|---|---|---|---|---|
| | | = Not analyzed) | | |
| C.1505 | 1032. | খানার নমুনার অবস্থা<br>HH sample qualifier | 1. ডিটেকশন লেভেলের নীচে Below detection limit<br>2. ডিটেকশন লেভেলের উপরে Above detection limit<br>3. নমুনা এনালাইসিস করা হয় নাই Sample could not be analyzed | |
| | | প্রস্তুতির আগে ক্লাষ্টার/খানা আই ডি বোতলে/হুয়েলপ্যাকে স্পষ্ট অক্ষরে লিখুন** **WRITE THE CLUSTER/HH ID VERY CLEARLY ON THE WHIRLPACK BEFORE PROCEEDING**** | | |
| | | খানার সমন্বিত পানির উৎসের নমুনা PAIRED SOURCE WATER SAMPLE FOR HH WATER SAMPLE<br>। যদি উত্তরদাতা বলেন যে ভাগের/সেয়াড পানির উৎস থেকে পানি সংগ্রহ করা হয়েছে তাহলে **701 নং** প্রশ্নে চলে যান ।IF RESPONDENT REPORTED COLLECTING WATER FROM A SHARED SOURCE→ **SKIP TO 701** | | |
| C.1506 | 1033. | যদি উত্তরদাতা বলেন যে নিজস্ব পানির উৎস থেকে পানি সংগ্রহ করা হয়েছে এবং পানির উৎসটি বাড়ীর মধ্যেই অবস্থিত(রেইন ওয়াটারসহ)। F RESPONDENT REPORTED COLLECTING WATER FROM A PRIVATE WATER SOURCE AND THAT WATER SOURCE IS WITHIN THE COMPOUND (INCLUDING RAINWATER), ASK :<br><br>প্রশ্ন করুন;আমি কী আমাদের অফিসে আপনার উৎসের পানি পরীক্ষা করার জন্য াকছু পানি সংগ্রহ করতে পারি ? May I collect a small sample of your source water to test for bacteria in our office?<br><br>পানি সংরক্ষিত হলেই সংগ্রহের রেকর্ড করুন। RECORD WHETHER SAMPLE WAS COLLECTED: | [1] হ্যাঁ ,নমূনা নেয়া হয়েছে YES, SAMPLE COLLECTED<br>[2] না,প্রত্যাখাত হয়েছে NO, REFUSED | → **SKIP TO 701** |
| C.1507 | 1034. | উৎসের পানির আই ডি Water source ID<br>যদি উৎস ভাগের/সেয়াড হয় তখন উৎস আইডির তালিকা দেখুন। If Source is a shared source, list the source ID.<br>যদি উৎস বাড়ীর /কম্পাউন্ড মধ্যে হয় তখন ক্লাস্টার/খানা আইডি হবে উৎস আইডি এবং ৯৯ কোড রেকর্ড করতে হবে। If Source is located in the compound, only list the cluster/HH ID, and record "99" in the Source ID.<br>যদি উৎস ভাগের/সেয়াড হয় এবং ইহা যদি তালিকাতে না থাকে তখন ০০ রেকর্ড করতে হবে। If the shared source located is not on the preprinted list, enter "00" | A ☐☐☐ / ☐☐ক্লাস্টার/খানা আইডি Cluster / HH ID<br>B ☐☐উৎস আইডি (যদি উৎস একই/সেয়াড হয় )Source ID (if a shared source) | |
| | 1035. | উৎস-এর সংরক্ষিত পানির নমূনা সংরক্ষণকাল (দিন/মাস/বছর.ঘন টা:মিনিট)Source sample collected | ☐☐/☐☐/☐☐☐☐:☐☐ | |

| | | | |
|---|---|---|---|
| | | (DD/MM/YY, HH:MM) | |
| C.1508 | 1036. | উৎস-এর পানির ধরণ রেকর্ড করুন Record the source water type | 12. অগভীর টিউবওয়েল Shallow tubewell<br>13. গভীর টিউবওয়েল Deep tubewell<br>14. ঘরের ভিতর ট্যাপ বা পাইপের পানি Piped into home<br>15. উঠানে ট্যাপ বা পাইপের পানি Piped into yard<br>16. গভীর পাতকূয়া (উঠানে)Borewell in yard<br>17. পাবলিক ট্যাপ Public tap<br>18. পাবলিক গভীর পাতকূয়া Public borewell<br>19. অগভীর পাকা পাতকূয়া Shallow well with concrete reinforcement<br>20. অগভীর কাঁচা পাতকূয়া Shallow well (no concrete)<br>21. নদী/বাধ/লেক/পুকুর/সেচ নালাথেকে সরাসরি সংগৃহীত পানি River, lake, stream<br>22. অন্যান্য (লিখুন) Other: _____________ |
| | | প্রস্তুতির আগে ক্লাষ্টার/খানা আই ডি বোতলে/রুয়েলপ্যাকে স্পষ্ট অক্ষরে লিখুন **** WRITE THE SOURCE ID VERY CLEARLY ON THE WHIRLPACK BEFORE PROCEEDING**** | |

**Wash Benefit Module 12 Measures of the Home Environment**

| **Interview items** | |
|---|---|
| **Learning stimulation** | |
| How many children's books or picture books do you have for (*name*)? | None<br>[___ ___] |
| I am interested in learning about the things that (*name*) plays with when he/she is at home.<br><br>Does he/she play with:<br><br>[A] homemade toys (such as dolls, cars, or other toys made at home)?<br><br>[B] toys from a shop or manufactured toys?<br><br>[C] household objects (such as bowls or pots) or objects found outside (such as sticks, rocks, animal shells or leaves)?<br><br>*If the respondent says "YES" to the categories above, then probe to learn specifically what the child plays with to ascertain the response* | Y N DK<br><br>Homemade toys 1 2 8<br><br>Toys from a shop 1 2 8<br><br>Household objects or outside objects 1 2 8 |

| | |
|---|---|
| Sometimes adults taking care of children have to leave the house to go shopping, wash clothes, or for other reasons and have to leave young children.<br><br>On how many days in the past week was (*name*):<br><br>[A] Left alone for more than an hour?<br><br>[B] Left in the care of another child, that is, someone less than 10 years old, for more than an hour?<br><br>*If 'none' enter' 0'. If 'don't know' enter'8'* | Number of days left alone for more than an hour __<br><br>Number of days left with other child for more than an hour __ |

| | Mother | Father | Other | No one |
|---|---|---|---|---|
| In the past 3 days, did you or any household member over 15 years of age engage in any of the following activities with (*name*):<br><br>*If yes, ask:*<br>who engaged in this activity with (*name*)?<br><br>*Circle all that apply. We need to be sure that all circled are tallied, so that a cumulative (1-3 for each item below) can be computed.* | | | | |
| [A] Read books to or looked at picture books with (*name*)? | Y | Y | Y | Y |
| [B] Told stories to (*name*)? | Y | Y | Y | Y |
| [C] Sang songs to (*name*) or with (*name*), including lullabies? | Y | Y | Y | Y |
| [D] Took (*name*) outside the home, compound, yard or enclosure? | Y | Y | Y | Y |
| [E] Played with (*name*)? | Y | Y | Y | Y |
| [F] Named, counted, or drew things to or with (*name*)? | Y | Y | Y | Y |

**Wash Benefit Module 13 MEASURES FOR SPILLOVER**

সকল খানার জন্য প্রযোজ্য Administer to: All study households

| | | | |
|---|---|---|---|
| **C.1301.** | 1301. | আপনার সবচেয়ে কাছের হেলথ ফ্যাসিলিটি (স্বাস্থ্যসেবা পাওয়ার সুযোগ)-এর নাম কি?যদি না জানে/নিশ্চিত না হয় তবে ৯৯ বসান। খানার কেউ ডাক্তার হলে বা ডাক্তার খানায় এসে চিকিৎসা করলে ৫৫ লিখুন।What is the name of your nearest health facility?<br>Enter "99" if Don't know / not sure | |
| **C.1301.a** | 1302. | গত ১ বছরে আপনি/আপনার শিশু মোটামুটি কতবার যে কোন অসুখে এই সুযোগ গ্রহন করেছিলেন? In the past year, (since this time last year) approximately how many times did you visit this health facility for any reason (including with your children)? | \|__\|__\|বার TIMES |

| | | | |
|---|---|---|---|
| C.1301.b | 1303. | আপনি সেখানে কিভাবে যান? What is your primary mode of travel to the health facility? | ১. পায়ে হেঁটে<br>2. রিকশা/ভ্যান/নৌকা<br>3. ব্যাটারি চালিত অটোরিকশা<br>4. ট্যাম্পু/লেগুনা/ সিএনজি/ বাস<br>5. প্রযোজ্য নয় |
| C.1301.c | 1303.a | আনুমানিক সেখানে যেতে কতক্ষণ সময় লাগে(আপনার খানা থেকে শুধু যেতে কত মিনিট সময় লাগে) ? যদি না জানে/নিশ্চিত না হয় তবে ৯৯ বসান। Approximately how long does it take to travel there (in minutes, one-way, from where we are now)?<br><br>Enter "999" if Don't know / not sure | \|__\|__\|__\| মিনিটে MINUTES |
| C.1302. | 1304. | আপনার কাছের প্রাক- প্রাইমারী স্কুলের -এর নাম কি ? যেখানে আপনার শিশু যেতে পারে/বড় হলেও যেতে পারবে । যদি না জানে/নিশ্চিত না হয় তবে ৯৯, যদি না থাকে তবে ৮৮ বসান।What is the name of the ECD center [preschool] where your children go (or will go when they are old enough)? | |
| C.1303. | 1305. | আপনার কাছের প্রাইমারী স্কুলের -এর নাম কি ? যেখানে আপনার শিশু যেতে পারে/বড় হলেও যেতে পারবে । What is the name of the primary school where your children go (or will go when they are old enough)? | |
| C.1304. | 1306. | আপনার কাছের বাজার -এর নাম কিযেখানে আপনি সবসময়ই কেনা-বেচা করতে যান? । What is the name of the market that you most often go to when you buy or sell things? | |
| C.1304.a | 1307. | কতবার আপনি সেখানে যান? How often do you go to that market? | 0 কখনই না Never<br>1 সপ্তাহে একবার অথবা তার অধিক Once per week or more<br>2 প্রতি ২ সপ্তাহে একবার Once every 2 weeks<br>3 প্রতি ৩ সপ্তাহে একবার Once every 3 weeks<br>4 প্রতি ৪ সপ্তাহে একবার(প্রতি মাসে) Once every 4 weeks (every month)<br>5 প্রতি মাসে একবারের কম Less than once every month<br>99 জানি না/নিশ্চিত না Don't know / not sure |
| | 1308. | আপনি সেখানে কিভাবে যান? What is your primary mode of travel to the health facility? | 1. পায়ে হেঁটে<br>2. রিকশা/ভ্যান/নৌকা<br>3. ব্যাটারি চালিত অটোরিকশা<br>4. ট্যাম্পু/লেগুনা/ সিএনজি/ বাস<br>5. প্রযোজ্য নয় |
| C.1304.b | 1308.a | আনুমানিক সেখানে যেতে কতক্ষণ সময় লাগে(আপনার খানা থেকে শুধু যেতে কত মিনিট সময় লাগে) ? যদি না জানে/নিশ্চিত না হয় তবে ৯৯ বসান। Approximately how long does it take to travel there (in minutes, one-way, from where we are now)?<br><br>Enter "999" if Don't know / not sure | \|__\|__\|__\| মিনিটে MINUTES |
| C.1305. | 1309. | আপনার কাছের মসজিদ/মন্দির/গীর্জা/প্যাগোডা -এর নাম কি? যেখানে আপনি প্রার্থনা করতে যান । What is the name of the church/mosque/temple (if any) that you attend? | |
| C.1305.a | 1310. | কতবার আপনি সেখানে যান? How often do you go to church? | ০. কখনই না Never<br>1 সপ্তাহে একবার অথবা তার অধিক Once per week or more<br>2 প্রতি ২ সপ্তাহে একবার Once every 2 weeks<br>3 প্রতি ৩ সপ্তাহে একবার Once every 3 weeks |

| | | | |
|---|---|---|---|
| | | | 4 প্রতি ৪ সপ্তাহে একবার(প্রতি মাসে) Once every 4 weeks (every month)<br>5 প্রতি মাসে একবারের কম Less than once every month<br>99 জানি না/নিশ্চিত না Don't know / not sure |
| | 1311. | আপনি সেখানে কিভাবে যান? What is your primary mode of travel to the health facility? | 1. পায়ে হেঁটে<br>2. রিকশা/ভ্যান/নৌকা<br>3. ব্যাটারি চালিত অটোরিকশা<br>4. ট্যাম্পু/লেগুনা/ সিএনজি/ বাস<br>5. প্রযোজ্য নয় |
| **C.1305.b** | **1312.a** | আনুমানিক সেখানে যেতে কতক্ষণ সময় লাগে(আপনার খানা থেকে শুধু যেতে কত মিনিট সময় লাগে) ? যদি না জানে/নিশ্চিত না হয় তবে ৯৯ বসান। Approximately how long does it take to travel there (in minutes, one-way, from where we are now)?<br><br>Enter "999" if Don't know / not sure | \|__\|__\|__\| মিনিটে MINUTES |
| **C.1306.** | 1312. | খানাটির জিপিএস অবস্থান রেকর্ড নিশ্চিত করুন। Confirm that you have recorded a GPS location linked to this household | [ 1 ] খানাটির জিপিএস অবস্থান রেকর্ড নিশ্চিত হয়েছে Confirmed, GPS point recorded<br>[ 99 ] রেকর্ড নিশ্চিত হয়নি Could not record |

**Wash Benefit Module 14LNS UPTAKE**

| **Adherence to Sonamoni** | | | |
|---|---|---|---|
| **N.1401** | Is this a nutrition intervention arm? | 1. Yes<br>0. No | If No, skip to 1801 |
| **N.1402** | Did they receive LNS yet? | 1. Yes<br>2. No,child is below 6 months<br>3. No, as they were from home for last one month<br>4. Others, specify__________ | Stop the interview here. |
| **N.1403** | If yes, please show me the items you received? Record your observation.<br><br>DO NOT READ THE RESPONSES. MULTIPLE RESPONSES ALLOWED. | Items (Yes=1, No=0)<br>1. Sonamoni<br>2. Plastic container for Sonamoni | |
| **N.1404** | How many days ago were the Sonamoni sachets distributed?<br>(FRAs ask household and see the records ) | I__II__I days | |
| **N.1405** | How many unused Sonamoni sachets you had at the time of last distribution?<br>(FRAs calculate and put the number) | I__II__I Sachets | |
| **N.1406** | How many Sonamoni sachets did you receive during last distribution? (FRAs ask household & see the records) | I__I__I Sachets | |
| **N.1407** | How many Sonamoni sachets are unused at present?<br>(FRAs count and put the number) | I__I__I Sachets | |

| | | | |
|---|---|---|---|
| **N.1408** | During the past week, how many days did you feed Sonamoni to your child?” | \|__\| days | If 0 then skip to 510 |
| **N.1409** | How much supplement per day did you feed most of the time during the last seven days?<br><br>DO NOT READ THE RESPONSES. SINGLE RESPONSE ALLOWED | \|___\|. \|__ \| sachets | |
| **N.1410** | Did anybody else ever share Sonamoni with the target child? | 1. Yes<br>0. No | If no, skip to 512 |
| **N.1411** | If yes, who?<br><br>DO NOT READ THE RESPONSES. MULTIPLE RESPONSES ALLOWED. | Relationship (Yes=1, No=0)<br>1. Sibling<br>2. Other child/children<br>3. Adult relative(s)<br>4. Other adult(s) | |
| **N.1412** | If yes, how did you primarily feed the Sonamoni to your Child during last seven days?<br><br>DO NOT READ THE RESPONSES. SINGLE RESPONSES ALLOWED. | Supplementation (Yes=1, No=0)<br>1. Eaten alone<br>2. Mixed with rice<br>3. Mixed with other food | |
| **N.1413** | What did you use to feed the child last time you fed Sonamoni? | 1. Spoon<br>2. Mother's fingers<br>3. Directly from sachet<br>4. Child fed him/herself by hand | If 2 or 4 then ask N.514 or skip to N.516 |
| **N.1414** | Did you wash your hands last time you fed Sonamoni to your child?<br>Or<br>Did your child washed hands last time s/he ate Sonamoni by hands? | 1. Yes<br>0. No | If no, skip to N.516 |
| **N.1415** | If yes, what did you use to wash your hands last time you fed Sonamoni to your child?<br>DO NOT READ THE RESPONSES. MULTIPLE RESPONSES ALLOWED. | Used (Yes==1, No=0)<br>1. Bar soap<br>2. Detergent<br>3. Liquid soap<br>4. Soapy water<br>5. Ash<br>6. Mud<br>7. Only water | |
| **N.1416** | If no to N.511 then why Sonamoni was not eaten?<br><br>DO NOT READ THE RESPONSES. MULTIPLE RESPONSES ALLOWED. | Reasons (Yes=1, No=0)<br>1. Child didn’t like<br>2. Not sure about the product<br>3. Peer pressure not to take<br>4. Child was sick<br>5. Away from home<br>6. Gave supplement to another person<br>77. Others (Specify) | |
| **N.1417** | Were any of the Sonamoni sachets damaged or | 1. Yes | |

| | | |
|---|---|---|
| | opened prior or during distribution? | 0. No |
| **N.1418** | What shortcomings or problems are there in your opinion with Sonamoni?<br><br>DO NOT READ THE RESPONSES. MULTIPLE RESPONSES ALLOWED. | Shortcoming/problems (Yes=1, No=0)<br>1. Irregular supply<br>2. Taste is not good<br>3. Child vomit after swallow<br>4. Color of the food changes when mixed with Sonamoni<br>5. Limited supply<br>6. Bad smell<br>7. Causes indigestion<br>8. Loose stool<br>9. Allergic Reaction (skin rash)<br>10. Child doesn't like to eat Sonamoni<br>11. Difficult to take out of the sachet<br>12. No Problem<br>13. Increased appetite<br>14. Child won't eat other food because they are eating Sonamoni<br>77. Others, ( Specify ) |
| **N.1419** | What is that you liked about the supplement?<br>DO NOT READ THE RESPONSES. MULTIPLE RESPONSES ALLOWED. | Benefits (Yes=1, No=0)<br>1. Increased appetite<br>2. Weight gain<br>3. Increased energy<br>4. Easy to feed<br>5. Child liked it<br>6. Child remains playful<br>7. Good taste<br>8. Good smell<br>9. Don't have to feed other foods because Sonamoni meets the child's needs<br>77. Other, Specify ___________ |

#### **Wash Benefit Module 15** Environmental Sampling (Water, Hands, Sentinel Toys and Fly Density)

DAY 1

SECTION 1. DELIVER TOY BALL AND FLY TAPE

PROMPT: Please deliver the toy ball to the target child or respondent.

1.1 Who did you give the ball to?

1 = Target child

2 = Respondent

3 = Other caregiver

4 = Ball not delivered

77 = Other (specify)

1.2 (if question 1.1 is not 4) Record the time that the toy ball was delivered (24H format, HH:MM).

1.3 (if question 1.1 is 4) Why was the toy ball not delivered?

1 = Respondent/caregiver refused

2 = Did not have a toy ball to deliver

77 = Other (specify)

PROMPT: Can you please show me the area where you prepare food?

PROMPT: Hang fly tape as close as possible to the food preparation area If there is more than one food prep area, hang the strips near where food was most recently prepared. Do NOT hang the strips over or near to a cooking fire. Ask the respondents to leave the tape undisturbed.

1.4 Record the time that the fly paper was hung at the food preperation area (24H format, HH:MM)

PROMPT: Can you please show me your primary latrine area?

PROMPT: Hang fly tape as close as possible to the primary latrine area. If there is more than one primary latrine area, hang the strips near the latrine that the respondent used most recently. Ask the respondents to leave the tape undisturbed.

1.5 Record the time that the fly paper was hung at the primary latrine area (24H format, HH:MM)

SECTION 2. HAND RINSE SAMPLING

| | |
|---|---|
| 2.1 | Record whether the respondent has washed heror target child's hands at any time before this question since you arrived at the household. Select all that apply.<br><br>[1] Observed respondent washing hands<br><br>[2] Did not observe handwashing directly but respondent had wet hands<br><br>[3] Did not observe respondent washing hands or with wet hands<br><br>[4] Observed respondent washing/wiping target child's hands<br><br>[5] Did not observe handwashing directly but target child had wet hands<br><br>[6] Did not observe respondent washing/wiping target child's hands or child with wet hands |

<table>
<tr><td rowspan="3">.<br><br>2.2</td><td rowspan="3">READ: Thank you. Now, I would like to do a quick inspection of your hands. I hope you don't mind. Can you please show me your hands?<br><br>BOTH HANDS SHOULD BE SHOWN (NOT JUST ONE HAND). USE APPEARANCE CODES BELOW TO RECORD THE DESCRIPTION THAT BEST DESCRIBES THE LEVEL OF CLEANLINESS.<br><br>ASK: Please show me [TARGET CHILD NAME]'s hands. DO NOT TOUCH HANDS.</td><td colspan="2">MOTHER/CAREGIVER</td></tr>
<tr><td>Left Hand<br><br>A|__|FINGERNAILS<br><br>B|__| PALMS<br><br>C|__|FINGER PADS</td><td>Right Hand<br><br>D|__|FINGERNAILS<br><br>E|__| PALMS<br><br>F|__|FINGER PADS</td></tr>
<tr><td colspan="2">TARGET CHILD</td></tr>
</table>

| | | | |
|---|---|---|---|
| | | Left Hand<br><br>G\|__\|FINGERNAILS<br><br>H\|__\| PALMS<br><br>I \|__\|FINGER PADS | Right Hand<br><br>J \|__\|FINGERNAILS<br><br>K \|__\| PALMS<br><br>L \|__\|FINGER PADS |

APPEARANCE CODES:

VISIBLE DIRT (DIRT/MUD/SOIL/ASH OR ANY OTHER MATERIAL IS VISIBLE)

UNCLEAN APPEARANCE (NO DIRT IS VISIBLE ON THIS PART OF THE HAND BUT, IN GENERAL, THIS PART OF THE HAND APPEARS UNCLEAN)

CLEAN (OBSERVED PART OF THE HAND IS CLEAN AS WOULD APPEAR AFTER SOMEONE WASHES HANDS OR TAKES A BATH)

[88] N/A

[99] OBSERVATION NOT POSSIBLE/REFUSED/TARGET CHILD NOT AVAILABLE

PROMPT: Collect a hand rinse sample from the target child by rinsing both hands in same Whirlpak bag.

2.3 Please enter the 4-digit unique numerical ID.

PROMPT: Please label the whirlpak with the following label: H.[PID].[DAY].[MONTH]

2.4 Has a child hand rinse sample been collected successfully? Select all that apply.

1 = Yes, both hands rinsed

2 = No, only one hand rinsed

3 = No, some sample was spilled

4 = No, the inside of the sample bag was contaminated

5 = No, sample not collected

2.5 (if 2.4 is not 5) Record time that sample was collected (24H format, HH:MM).

SECTION 3. WATER SAMPLING

If (target child's name) wanted a drink of water right now, could you show me how you would give it to him/her? [If target child it too young to drink water, ask: If your child < 3 years wanted a drink of water right now, could you show me how you would give it to him/her?] [If the mother has no children < 3 years, ask: If you wanted a drink of water right now, could you show me how you would get it?]

Ask the question and observe.

| | | | | |
|---|---|---|---|---|
| 3.1 | IS WATER FOR DRINKING CURRENTLY AVAILABLE? | YES<br>NO → SKIP TO SECTION 4 | | |
| | WHAT DID RESPONDENT DO BEFORE TAKING THE WATER? | YES | NO | D/K |
| 3.2 | Rinsed glass / cup with drinking water before filling | [1] | [2] | [99] |
| 3.3 | Washed hands with water before drinking water was obtained | [1] | [2] | [99] |
| 3.4 | Washed hands with soap before drinking water obtained | [1] | [2] | [99] |
| | FROM WHERE DID THE RESPONDENT TAKE THE WATER? | YES | NO | D/K |
| 3.5 | Brought directly from the water source | [1] | [2] | [99] |
| 3.6 | Brought directly from water stored in a container | [1] | [2] | [99] |
| 3.7 | Brought directly from water filter | [1] | [2] | [99] |
| 3.8 | Stored water was covered or in a narrow mouth vessel | [1] | [2] | [99] |
| | HOW DID SHE GET THE WATER INTO THE GLASS/CUP? | YES | NO | D/K |
| 3.9 | Hands touched / contacted the drinking water | [1] | [2] | [99] |
| 3.10 | Glass/cup dipped into water container | [1] | [2] | [99] |
| 3.11 | Ladle used to obtain water | [1] | [2] | [99] |

| | | | | |
|---|---|---|---|---|
| 3.12 | Water poured from container | [1] | [2] | [99] |
| 3.13 | Water poured from tap / handpump | [1] | [2] | [99] |

3.14 (ask/observe) What is the source of the drinking water?

1 = Tubewell

2 = Unprotected spring

3 = Protected spring

4 = Unprotected dug well

5 = Protected dug well

6 = Rainwater collection

7 = Cart with small tank / drum

8 = Tanker truck

9 = Surface water (river, dam, lake, pond, stream, canal, irrigation channel)

10 = Piped water into dwelling

11 = Piped water into yard / plot

77 = Other (specify)

99 = DK

3.15 (obs) (if 3.6 is 1) What is the type of the container?

1= Kolshi[B] /Jerrycan[K]

2 = Jug

3 = Topaz (provided by icddrb) [B]

4 = Bucket

5 = Other wide mouth container

6 = Other narrow mouth container

99 = Could not observe

3.16 (if 3.6 is 1) How long ago did you or somebody in your home collect this water?  (99 = Don't know)

A ☐☐ HOURS

B ☐☐ DAYS

3.17 (if 3.6 is 1) Have you done anything to make this water less cloudy or safer to drink?

1 = Yes

2 = No → Skip to 3.20

99 = DK/Not sure

3.18 (if 3.17 is 1) How was this water treated? ✋Do Not Read Responses. Mark all that apply.

| | | |
|---|---|---|
| 1 | [1] Yes [2] No | Aquatabs[B] / Chlorine dispenser [K] (field code differs by country) |
| 2 | [1] Yes [2] No | Waterguard / bottled chlorine |
| 3 | [1] Yes [2] No | Boil |
| 4 | [1] Yes [2] No | Strain it through cloth or other material |
| 5 | [1] Yes [2] No | Water filter [ceramic, sand, composite] |
| 6 | [1] Yes [2] No | Solar disinfection (SODIS) |
| 7 | [1] Yes [2] No | Let it stand and settle |
| 8 | [1] Yes [2] No | Biosand filter |
| 9 | [1] Yes [2] No | Lifestraw Family Filter (VestergaardFrandsen distributed) [K] |
| 10 | [1] Yes [2] No | Coagulant (alum) |
| 11 | [1] Yes [2] No | PUR (flocculant + disinfectant) |
| 12 | [1] Yes [2] No | Other (specify) |

3.19 (if 3.17 is 1) Approximately how long ago did you treat the water?

HH:MM ☐☐:☐☐ ( 99:99 Don't know )

PROMPT: (if 3.18 is 1=Y or 2=Y) Please collect a drinking water sample by pouring the water from the glass provided by the participant into a Whirlpak with sodium thiosulphate.

PROMPT: (if 3.18 is 1=Nand 2=N) Please collect a drinking water sample by pouring the water from the glass provided by the participant into a regular Whirlpak.

3.20 Please enter the 4-digit unique numerical ID.

PROMPT: Please label the whirlpak with the following label: W.[PID].[DAY].[MONTH]

3.21 Has a drinking water sample been collected for microbiological testing?

1 = Yes

2 = No

3.22 (if 3.21 is 1) Record time that sample was collected (24H format, HH:MM)

PROMPT: (if 3.18 is 1=Y or 2=Y) Please fill a plastic tube with stored water for chlorine testing.

3.23 (if 3.18 is 1=Y or 2=Y) Has a sample from the storage container been collected for chlorine testing?

1 = Yes

2 = No

3.24 (if 3.23 is 1) Record time that sample was collected (24H format, HH:MM)

Beginning of Tubewell Section [B]

3.25 (obs) (if 3.14 is 1) Are there any materials attached to the tubewell mouth?

1 = No materials are covering the tubewell mouth

2 = Cloth

3 = Plastic

4 =Bamboo

77 = Other

3.26 (obs) (if 3.14 is 1) Did the respondent prime the tubewell prior to retrieving the glass of water?

1 = Yes

2 = No

3.27 (ask) (if 3.26 is 1) What kind of water was used to prime the tubewell?

1= Stored water from same tubewell

2 = Water from another tubewell

3 = Pond water

77 = Other

3.28 (ask) (if 3.14 is 1) What is the depth of the tubewell? (ft) (99 = Don't know)

3.29 (obs) (if 3.14 is 1) Does the tubewell have a platform?

1 = Yes

2 = No

99 = Could not observe

3.30 (obs) (if. 3.29 is 1) Is the platform intact?

1 = Yes

2 = No

99 = Could not observe

3.31 (obs) (if 3.14 is 1) Is faulty drainage allowing ponding within 2 meters of the tubewell?

1 = Yes

2 = No

99 = Could not observe

3.32 (obs) (if 3.14 is 1) Is there a latrine within 10 meters of the tubewell?

1 = Yes

2 = No

99 = Could not observe

3.33 (obs) (if 3.32 is 1) How many steps is the closest latrine <10 m from the tubewell?

3.34 (obs) (if 3.32 is 1) How many latrines are present within 10 meters of the tubewell?

3.35 (obs) (if 3.14 is 1) Is there a pond within 10 meters of the tubewell?

1 = Yes

2 = No

99 = Could not observe

End of Tubewell Section [B]

SECTION 4. RESIDUAL CHLORINE MEASUREMENT

PROMPT: (if 3.23 is 1) Measure the free chlorine residual in the stored water sample.Please leave the compound before conducting the chlorine test.

4.1 (record) (if 3.23 is 1) Free chlorine: _____._____ mg/L

4.2 (record) (if 3.23 is 1) Record time that chlorine was measured (24H format, HH:MM)

DAY 2

SECTION 1. SENTINEL TOY SAMPLING

1.1(ask) Did you receive a toy ball from our team yesterday?

1 = Yes

2 = No → Skip to Section 2

PROMPT: Could you please show me where the ball that was delivered to your household yesterday is currently located?

1.2 (ask) May I now rinse the toy ball that was delivered to your household yesterday?

1 = Yes

2 = Ball was lost → Skip to Section 2

3 = Ball was given away to another household → Skip to Section 2

4 = Refused → Skip to Section 2

77 = Could not retrieve the ball for other reason (specify) → Skip to Section 2

1.3 (obs) Where is the ball located?

1 = Outside: on dirt

2 = Outside: on concrete/cement/wood

3 = Outside: in container

4 = Outside: in another home

5 = Inside: on dirt floor

6 = Inside: on concrete/cement/wood floor

7 = Inside: on surface other than ground, not in container (e.g. on bed, on table, etc.)

8 = Inside: in storage container/cabinet

9 = In child's hands

77 = Other (specify)

1.4.a (ask)In your opinion, how much did (target child's name) play with the toy ball over the past 24 hours? (read each choice)

1 = Several times (4 or more times)

2 = Few times (2-3 times)

3 = Only once since he/she got the ball

4 = Never

99 = DK/Not sure

1.4.b In your opinion, how much time total did (target child's name) spend playing with the toy ball over the past 24 hours? (enter hours)

1.5.a (ask) In your opinion, how much did any of the other children in the household or bari play with the toy ball in the past 24 hours? (read each choice)

1 = Several times (4 or more times)

2 = Few times (2-3 times)

3 = Only once since he/she got the ball

4 = Never

99 = DK/Not sure

1.5.b In your opinion, how much time total did other children in the household or bari spend playing with the toy ball over the past 24 hours? (enter hours)

1.6 (ask) Did children from other baris play with the toy ball in the past 24 hours? (read each choice)

1 = They played with it within this bari

2 = They played with it outside this bari

3 = No

99 = DK/Not sure

1.7 (ask) From what you saw, did the children play with the ball mostly inside the home (indoors), mostly outside the home (outdoors) or equal amount inside and outside the home?

1 = Mostly inside the home

2 = Mostly outside the home

3 = Equal amounts inside and outside the home

99 = DK/Not sure

1.8 (ask) How did the child play with the ball? Read each choice.

1 = With hands

2 = With feet

3 = With hands and feet

99 = DK/Not sure

1.9 (ask) Did you do anything to clean the ball?

1 = Yes

2 = No

1.10 (ask) (if 1.9 is 1) What did you do to clean the ball? Select all that apply. Read each choice.

1 = Washed with water only

2 = Washed with water and soap

3 = Wiped with towel

4 = Wiped on clothes

77 = Other (specify)

1.11 (ask) (if 1.9 is 1) How long ago did you clean the ball? (Enter hours, 99=DK)

1.12 (obs) Have children played with the ball since you arrived at the household? (select all that apply)

1 = Target child has played with ball

2 = Other children in the household have played with ball

3 = No children have played with ball

1.13 (obs) Toy appearance.

1 = Unused

2 = Used, clean appearance

3 = Used, unclean appearance

4 = Used, visibly dirty

5 =  Used, visibly wet

1.14 (obs) Toy condition.

1 = Good condition

2 = Bulging (from being stepped on, sat on, heat applied, etc)

3 = Deflated (ball has lost air)

4 = Burst/Torn (ball has visible hole)

PROMPT: Collect a toy rinse sample by rinsing the toy in the Whirlpak bag.

1.15 Please enter the 4-digit unique numerical ID.

PROMPT: Please label the whirlpak with the following label: T.[PID].[DAY].[MONTH]

1.16 Has a toy rinse sample been collected successfully?

1 = Yes, toy rinsed successfully

2 = No, some sample was spilled

3 = No, the inside of the sample bag was contaminated

4 = No, sample not collected

1.17 (if 1.16 is not 4) Record time that sample was collected (24H format, HH:MM).

SECTION 2. FLY DENSITY

2.1 Observe fly activity in the food preparation area. Estimate the total number of flies seen.

1 = None

2 = Low (1-5)

3 = Moderate (6-10)

4 = High (>10)

2.2 Was the fly tape hung under a roof (protected from rain)?

1 = Yes

2 = No

2.3 Is the food prep area inside or outside?

1 = Walls and roof

2 = Walls but no roof

3 = Roof but no walls

4 = No roof and no walls

2.4 (record) How many steps are the strips hung from the food preparation area?

2.5 (obs) Was the fly tape in the food preparation area tampered with or did it fall down?

2.6 (record) What is the total number of flies counted on all strips?

2.7(record) (if 2.6 is not 0) Write the number of each species observed:

Musca domestica______

Lesser house fly______

Blow/bottle fly______

Flesh fly/sarcophaga______

Other______

Cannot distinguish______

2.8 Record the time you performed the fly density observation at the food preparation area (24H format, HH:MM)

2.9 Observe fly activity in the latrine area. Estimate the total number of flies seen.

1 = None

2 = Low (1-5)

3 = Moderate (6-10)

4 = High (>10)

2.10 Was the fly tape hung under a roof (protected from rain)?

1 = Yes

2 = No

2.11 (record) How many steps are the strips hung from the latrine area?

2.12 (obs) Was the fly tape in the latrine area tampered with or did it fall down?

2.13 (record) What is the total number of flies counted on all strips?

2.14(record) (if 2.13 is not 0) Write the number of each species observed:

Musca domestica______

Lesser house fly______

Blow/bottle fly ______

Flesh fly/sarcophaga ______

Other ______

Cannot distinguish______

2.15 Record the time you performed the fly density observation at the latrine area (24H format, HH:MM)

#### MacArthur Adapted Communicative Development Inventory

##### Words and Gestures (MWG)

**Score : YES=1, NO=0**

| **Very easy** | **Un**<br><br>**(Yes=1, no=0)** | **Say**<br><br>**(Yes=1, no=0)** |
|---|---|---|
| 1.Come here | | |
| 2.Look at me | | |
| 3. Wake up | | |
| 4. Go to mom or dad | | |
| 5. Rattle | | |
| 6. Water | | |
| 7. Milk | | |
| 8.Glass | | |
| 9. Pillow | | |
| 10. Moon | | |
| 11.Mommy | | |
| 12. Daddy | | |
| 13. Grand father or mother | | |
| 14. Baby | | |
| 15. Peack-a-boo | | |
| 16. Clap hand | | |
| 17. To eat | | |
| 18. To bite | | |

| |
|---|
| 19. To sit |
| 20. Bye bye |
| Total very easy |

| **Easy** | **Un**<br><br>**(Yes=1, no=0)** | **Say**<br><br>**(Yes=1, no=0)** |
|---|---|---|
| 21. Are you hurt | | |
| 22. Give it to mom | | |
| 23. Love/hug me | | |
| 24. Want to pass urine | | |
| 25.Don't touch | | |
| 26.Meow | | |
| 27. Chicken | | |
| 28. Car | | |
| 29. Ball | | |
| 30.Rice | | |
| 31. Biscuit | | |
| 32. Puff.rice | | |
| 33. Pant | | |
| 34. Mirror | | |
| 35. Its dirty | | |

| |
|---|
| 36. Brush teeth |
| 37.To bang |
| 38. To play |
| 39. To walk |
| 40. To dance |
| Total Easy |

| **Moderate** | **Un**<br>**(Yes=1, no=0)** | **Say**<br>**(Yes=1, no=0)** |
|---|---|---|
| 41. Snake | | |
| 42. Khichuri | | |
| 43. Meat | | |
| 44. Jackfruit | | |
| 45. Noodles | | |
| 46. Eye | | |
| 47. Nose | | |
| 48. Beard | | |
| 49. Kitchen | | |
| 50. ে Table | | |
| 51. Cowshed | | |
| 52. Jug | | |
| 53. Pitcher | | |
| 54. Scissor | | |

| |
|---|
| 55. Box |
| 56. Shop |
| 57. Tree |
| 58. School room |
| 59. To open |
| 60. To write |
| Total Moderate |

| Difficult words | **Un**<br><br>**(Yes=1, no=0)** | **Say**<br><br>**(Yes=1, no=0)** |
|---|---|---|
| 61. Be careful | | |
| 62. Play doll | | |
| 63. All | | |
| 64. Tell ryhmes | | |
| 65. To get | | |
| 66. Dreid rice | | |
| 67. Hua hu | | |
| 68. Tiger | | |
| 69. Shoulder | | |
| 70. Little bit later | | |
| 71. Hide & seek | | |
| 72. Top (Toy) | | |

| |
|---|
| 73.Rain |
| 74. Almira/ showcase |
| 75. Van |
| 76. Khunti |
| 77. That |
| 78. To draw |
| 79. Radio |
| 80. Beside |
| Total score |

| **Very difficult words** | **Un**<br><br>**(Yes=1, no=0)** | **Say**<br><br>**(Yes=1, no=0)** |
|---|---|---|
| 81. Jacket | | |
| 82. Why |  |  |
| 83. Women | | |
| 84. Field | | |
| 85. Inside | | |
| 86. Person | | |
| 87. Brow | | |
| 88. Garden | | |
| 89. Block | | |
| 90. How |  |  |
| 91. Lace | | |

| |
|---|
| 92. Clean |
| 93. Chi chi |
| 94. Noon |
| 95. Her, His |
| 96. Buffelo |
| 97. Small pitcer |
| 98. Train |
| 99. Carrot |
| 100. Same |
| Total very difficult |
| **Total Scores** |

Motor milestone survey form

Household ID)

Will be filled according to ChildID 6-24 months

Note-1: FRAs will take this measurement with the help of FRO.

Note 2: Please fill up as many separate questionnaires as many 6-24 months children are in a household.

**Section 1: Motor milestone development**

The data collector will observe some of the activities of the child listed here and may need to ask the mother/caregiver for assistance. Answer any questions the mother/caregiver has. Observe each item and note in the column next. Clarify items as follows

**No (inability)**, the child tried but failed to perform the test item because it surpassed his or her developmental level

**Yes**, the child performed the test item according to the specific criteria

**Refused**, the child was calm and alert but just refused to cooperate

**Unable to test**, the child could not be tested on this milestone because his or her emotional state (drowsiness, fussiness or crying) was interfering with testing, the child was sick or child's caregiver was distraught

**Date of achievement**, the date that CHP entered in her record form. CHP records the date as caregiver reported and cross checked during her visit to the household.

| **Test item** | **(A) Examiner Report** | **(B) Date of Achievement** | | |
|---|---|---|---|---|
| | 0= No (inability)<br>1= Yes<br>222= Unable to test<br>666= Refused | *Only enter dates for milestones achieved for the first time from the CHP's record form.* | | |
| | Skip column B in baseline and go to next question. | D | M | Y |
| M 101. Sitting without support -10 secs | 0 1 222 666 | __ __ | __ __ | __ __ |
| M 102. Hands-and-knees crawling -3 | 0 1 222 666 | | | |

| | | | | |
|---|---|---|---|---|
| steps | | __ __ | __ __ | __ __ |
| M 103. Standing with assistance -10 secs | 0 1 222 666 | __ __ | __ __ | __ __ |
| M 104. Walking with assistance -5 steps | 0 1 222 666 | __ __ | __ __ | __ __ |
| M 105. Standing alone assistance | 0 1 222 666 | __ __ | __ __ | __ __ |
| M 106. Walking alone crawling- 5 steps | 0 1 222 666 | __ __ | __ __ | __ __ |
| M 107. Squats without support | 0 1 222 666 | __ __ | __ __ | __ __ |
| M 108. Balances on right foot with support- 2 secs | 0 1 222 666 | __ __ | __ __ | __ __ |
| M 109. Balances on left foot with support- 2 secs | 0 1 222 666 | __ __ | __ __ | __ __ |
| M 110. Jumps forward - 4 inches | 0 1 222 666 | __ __ | __ __ | __ __ |
| M 111. Balances on right foot for 2 seconds alone | 0 1 222 666 | __ __ | __ __ | __ __ |
| M 112. Balances on left foot for 2 seconds alone | 0 1 222 666 | __ __ | __ __ | __ __ |

**Wash Benefit Module 19 :Maternal Depression**

#### Center for Epidemiologic Studies Depression Scale (CES-D), NIMH

Below is a list of the ways you might have felt or behaved. Please tell me how often you have felt this way during the past week.

| | During the Past Week | | | |
|---|---|---|---|---|
| | Rarely or none of the time (less than 1 day ) | Some or a little of the time (1-2 days) | Occasionally or a moderate amount of time (3-4 days) | Most or all of the time (5-7 days) |

| | | | | |
|---|---|---|---|---|
| 1. I was bothered by things that usually don't bother me. | ☐ | ☐ | ☐ | ☐ |
| 2. I did not feel like eating; my appetite was poor. | ☐ | ☐ | ☐ | ☐ |
| 3. I felt that I could not shake off the blues even with help from my family or friends. | ☐ | ☐ | ☐ | ☐ |
| 4. I felt I was just as good as other people. | ☐ | ☐ | ☐ | ☐ |
| 5. I had trouble keeping my mind on what I was doing. | ☐ | ☐ | ☐ | ☐ |
| 6. I felt depressed. | ☐ | ☐ | ☐ | ☐ |
| 7. I felt that everything I did was an effort. | ☐ | ☐ | ☐ | ☐ |
| 8. I felt hopeful about the future. | ☐ | ☐ | ☐ | ☐ |
| 9. I thought my life had been a failure. | ☐ | ☐ | ☐ | ☐ |
| 10. I felt fearful. | ☐ | ☐ | ☐ | ☐ |
| 11. My sleep was restless. | ☐ | ☐ | ☐ | ☐ |
| 12. I was happy. | ☐ | ☐ | ☐ | ☐ |
| 13. I talked less than usual. | ☐ | ☐ | ☐ | ☐ |
| 14. I felt lonely. | ☐ | ☐ | ☐ | ☐ |
| 15. People were unfriendly. | ☐ | ☐ | ☐ | ☐ |
| 16. I enjoyed life. | ☐ | ☐ | ☐ | ☐ |
| 17. I had crying spells. | ☐ | ☐ | ☐ | ☐ |
| 18. I felt sad. | ☐ | ☐ | ☐ | ☐ |
| 19. I felt that people dislike me. | ☐ | ☐ | ☐ | ☐ |
| 20. I could not get "going." | ☐ | ☐ | ☐ | ☐ |

SCORING: zero for answers in the first column, 1 for answers in the second column, 2 for answers in the third column, 3 for answers in the fourth column. The scoring of positive items is reversed. Possible range of scores is zero to 60, with the higher scores indicating the presence of more symptomatology.

**Wash Benefit Module 41 Household Assets**

<table>
<tr><td colspan="3">আপনার সময়ের জন্য আপনাকে ধন্যবাদ।আমি আপনার খানা সম্মন্ধে জানতে চাই যেখানে আপনি বসবাস করেন। প্রথমেই আমি আপনার ঘর কী দিয়ে তৈরী সেটা দেখতে চাই । তাহলে আমি কি ইহা দেখতে পারি? Thank you very much for your time. Now I would talk about <u>the house that you live in</u>. First, I would like to observe the material your house is made of. Can I take a look at your house?<br><br>OBSERVE MAIN MATERIAL OF THE HOUSE:</td></tr>
<tr><td></td><td>4101.</td><td>ছাদ তৈরীতে কি কি উপাদান ব্যবহার করা হয়েছে ? (পরির্দশন করে যাচাই করুন) [Main material of the roof]<br>কাঁচা (বাঁশ/খড়) [Kaccha (bamboo / thatch)] 1<br>টিন [Tin] 2<br>সিমেন্ট/ কংক্রিট/ টালি [Cement / concrete / tiled]…3</td></tr>
<tr><td></td><td>4102.</td><td>দেয়াল তৈরীতে কি কি উপাদান ব্যবহার করা হয়েছে ? (পরির্দশন করে যাচাই করুন) [Main material of the walls]<br>পাট/বাঁশ/মাটি (কাঁচা) [Jute / bamboo / mud (kaccha)] .... 1<br>কাঠ [Wood] ................................................................ 2<br>ইট/সিমেন্ট [Brick / cement] ............................ 3<br>টিন [Tin]................................................ 4</td></tr>
<tr><td></td><td>4103.</td><td>মেঝে তৈরীতে কি কি উপাদান ব্যবহার করা হয়েছে ? (পরির্দশন করে যাচাই করুন) [Main material of the floor]<br>মাটি/বাঁশ (কাঁচা) [Earth / bamboo (kaccha)] ...................1<br>কাঠ [Wood] 2<br>ইট/ সিমেন্ট [Cement / concrete] ..................................3</td></tr>
<tr><td colspan="3">Thank you. Now I would like to ask you some questions about your household assets.</td></tr>
<tr><td></td><td>4104.</td><td>খানার ঘরটিতে কতগুলো কক্ষ আছে? (রান্নাঘর ও বাথরুম বাদে) [How many rooms the households have (exclude bathroom and Kitchen)?].</td></tr>
<tr><td></td><td>4105.</td><td>আপনার খানায় নিম্নলিখিত জিনিসগুলো আছে কী? [How many does the households have <u>হ্যাঁ (Yes)....1, না(No)....0,জানি না (DK).....999</u><br>a. |_____|বিদ্যুৎ [Electricity]<br>b. |_____||_____|আলমারী/ওয়ারড্রব (সংখ্যা) [Number of Almirah or wardrobe]<br>c. |_____||_____|টেবিল (সংখ্যা) [Number of tables]<br>d. |_____||_____|চেয়ার/বেঞ্চ(সংখ্যা) [Number of chair or bench]<br>e. |_____||_____|হাত ঘড়ি/দেয়াল ঘড়ি(সংখ্যা) [Number of watch or clock]<br>f. |_____||_____|খাট(সংখ্যা) [Number of khat]<br>g. |_____||_____|চৌকি (সংখ্যা) [Number of chouki ]<br>h. |_____|রেডিও [A radio that is working]<br>i. |_____|টেলিভিশন (সাদা/কালো) [A B/W television that is working]<br>j. |_____|টেলিভিশন (রঙিন) [A color television that is working]<br>k. _____|রেফ্রিজারেটর [Refrigerator]<br>l. |_____|সাইকেল (খেলনা সাইকেল নয়) A bicycle (used for commercial purposes not toy for children)<br>m. |_____|মটর সাইকেল[A motorcycle]<br>n. |_____|সেলাই মেশিন [A sewing machine]<br>o. |_____||_____|মোবাইল ফোন(সংখ্যা) [Number of Mobile phones]<br>p. |_____|ল্যান্ড ফোন[A land phone]</td></tr>
<tr><td></td><td>4106.</td><td>আপনার খানায় রান্নার জন্য প্রধানত কি ধরনের জ্বালানী ব্যবহার করা হয় ?[What type of fuel does your household mainly use for cooking?]<br>কাঠ [Wood] ........................................................................ 01<br>শস্যের অবশিষ্টাংশ/ঘাস [Crop residue / grass ]................................ 02<br>শুকনা গোবর [Dung cakes ] ............. 03<br>কয়লা [Coal / coke / lignite ]........................................................ 04<br>কাঠ কয়লা [Charcoal ]................................................................ 05<br>কেরোসিন [Kerosene ] ............. 06</td></tr>
</table>

<table>
<tr><td></td><td></td><td colspan="3">বিদ্যুৎ [Electricity ] ........................................................................ 07<br>তরল গ্যাস/প্রাকৃতিক গ্যাস [Liquid gas / gas ] .................................. 08<br>বায়ো-গ্যাস [Bio-gas] ..................................................................... 09<br>অন্যান্য [Other] ............................................................................... 77<br>(বর্নণা লিখুন) [Specify other] ______________________<br>জানি না [Don't know ] .................................................................. 99</td></tr>
<tr><td></td><td>4107.</td><td colspan="3">|_____|আপনার খানার মোট মাসিক আয় কত? (সকল আয়ের উৎসের যোগফল যেমনঃ বেতন, ঘর ভাড়া, কৃষিখাত ইত্যাদি) For the purpose of our research, would you please tell us your total monthly household income? (Please sum up your income from all sources like wage, rent, agriculture etc.)<br><br>1. 4000 টাকা পর্যন্ত [UP TO 4000 Taka]<br>2. 4001-6000 টাকা [4001-6000 Taka]<br>3. 6001- 8000 টাকা [6001- 8000 Taka]<br>4. 8001- 10000টাকা [8001- 10000 Taka]<br>5. 10001- 12000 টাকা [10001- 12000 Taka]s<br>6. 12001- 15000 টাকা [12001- 15000 Taka]<br>7. 15001-20000 টাকা [15001-20000 Taka]<br>8. 20001- 25000 টাকা [20001- 25000 Taka]<br>9. 25001- 30000 টাকা [25001- 30000 Taka]<br>10. 30000 টাকার উপরে [Above 30000 Taka]</td></tr>
<tr><td></td><td>4108.</td><td colspan="3">|____| বসতবাড়ি মালিকানা [What is the ownership status of the house where your household is currently living?]<br>নিজের বাড়ি [Self-owned] ............................................................................ 1<br>ভাড়া বাড়ি [Rental] ........................................................................................ 2<br>সরকারী জমি [Govt. land] ................................................................................ 3.<br>কোন জমিদারের/জোতদারের বাড়িতে থাকে [Owned by a landlord].................... 4<br>বাড়ি ভাড়া না দিয়ে অন্যের বাড়িতে থাকে [Living in someone's house without giving rent] .......5</td></tr>
<tr><td></td><td>4109.</td><td colspan="3">খানার সদস্যদের বসতবাড়িতেকোন জমি আছে কি?<br>1. হ্যাঁ<br>0. না(skip to 4111)</td></tr>
<tr><td></td><td>4110.</td><td colspan="3">|____||____||____||____||____| উত্তর হ্যাঁ হলে, খানার সদস্যদের বসতবাড়িতে মোট কতটুকু জমি আছে (ডেসিমেল)? (How much homestead land (decimal) does your household own?]<br><br>বলতে রাজি হয়নি [Refused]…..................................88<br>জানি না [Don't know]….................…...............99</td></tr>
<tr><td></td><td>4111.</td><td colspan="3">খানার সদস্যদের বসতবাড়ি ছাড়া অন্য কোন জমি আছে কি?<br>1. হ্যাঁ<br>০. না (skip to 4113)</td></tr>
<tr><td></td><td>4112.</td><td colspan="3">|____||____||____||____||____| উত্তর হ্যাঁ হলে, খানার সদস্যদের বসতবাড়িতে মোট কতটুকু জমি আছে (ডেসিমেল)? (How much homestead land (decimal) does your household own?]<br>বলতে রাজি হয়নি [Refused]…................................88<br>জানি না [Don't know]….................…...............99</td></tr>
<tr><td></td><td>4113.</td><td colspan="3">আপনার মতে সামাজিক প্রেক্ষাপটে আপনার খানাটির অবস্থান কি রকম [How would you describe your economic status?]<br>ধনী [Dhoni] ........................................................... 1<br>উচ্চ মধ্যবিত্ত [Uchho modho bitto].......................... 2<br>মধ্যবিত্ত [Modho Bitto] ........................................... 3<br>দরিদ্র [Doridro or Nimno motho bitto]................... 4<br>হতদরিদ্র [Hotodoridro]........................................... 5</td></tr>
<tr><td></td><td>4114.</td><td>আমি পশুর নামের তালিকা আপনাকে পড়ে শুনাতে চাচ্ছি ।এ গুলো কোনটি আপনার কম্পাউন্ড/াবাড়ীতে কতগুলো আছে এবং নিজের কয়টা আছে তা আমাকে বলুন । যদি না থাকে তাহলে ০০ বসান । আর যদি অজানা থাকে তাহলে ৯৯ বসান I'm going to read you a list of animals. Please tell me</td><td>কম্পাউন্ড/াবাড়ী<br>COMPOUND</td><td>খানা HOUSEHOLD</td></tr>
</table>

| | | | | |
|---|---|---|---|---|
| | | how many of each stays in your compound, and how many belong to your household. IF NONE, ENTER "00", IF UNKNOWN, ENTER "99" | | |
| | 1 | গরু CATTLE | \|__\|__\| | \|__\|__\| |
| | 2 | ছাগল GOATS | \|__\|__\| | \|__\|__\| |
| | 3 | মুরগী/হাঁস CHICKENS | \|__\|__\| | \|__\|__\| |
| | 4115. | **কতবার গরম্ন**/ছাগল তাদের ঘরের বাহিরে বাড়ীতে/উঠানে ছেড়ে দেন? How often do the cattleroam free in the compound?<br>উত্তরগুলো পড়ুন READ RESPONSES.<br>[1] সবসময় ছেড়ে দেন ALWAYS ROAM FREE<br>[2] মাঝে মাঝে ছেড়ে দেন SOMETIMES ROAM FREE<br>[3] কখনও ছেড়ে দেয়া হয় না NEVER ROAM FREE<br>4. প্রযোজ্য নয়; **গরম্ন**/ছাগল ছাড়া শুধুমাত্র মুরগী/হাঁসথাকলে N/A; ONLY HAVE CHICKENS | | |
| | 4116. | মুরগী অথবা অন্যান্য পাখি জাতীয় কোন কিছু আপনার প্রধান থাকার ঘরে প্রবেশ করে কি? Do chickens or other poultry ever go inside your main house?<br>READ RESPONSES.<br>[1] সবসময় ALWAYS<br>[2] মাঝে মাঝে SOMETIMES<br>[3] কখনও না NEVER<br>4. প্রযোজ্য নয়; মুরগী/হাঁসনাথাকলে N/A; NO CHICKENS | | |

| | | | |
|---|---|---|---|
| **Wash Benefit Module42Feces Observed in and Around the Compound** | | | |
| **সকল খানার জন্য প্রযোজ্য** Administer to: All study households | | | |
| **পর্যবেক্ষণ** Observation: For the following:<br>**প্রত্যেক জায়গায় মানুষের মল/পায়খানার স্তুপের সংখ্যা পর্যবেক্ষণ করুন (১০টি পর্যন্ত)** *Record the number of piles of human feces you observe in eacharea (up to 10 piles)*<br>55অধিক সংখ্যক (**স্তুপের সংখ্যা ১০টির উপরে**) Too numerous to count (more than 10 piles)<br>99 বলা যাচ্ছে না/**পর্যবেক্ষণ**করা সম্ভব হয়নি Cannot tell / could not observe | | | |
| **উঠানে** Within the courtyard | | | |
| **C.826.** | 4201. | **উঠানে মানুষের মল/পায়খানা পড়ে আছে যা উন্মুক্ত পায়খানা হিসেবে বিবেচিত হবে Human** feces<u>within the</u> **<u>courtyard</u>** that could be considered open defecation<br>55 অধিক সংখ্যক (**স্তুপের সংখ্যা ১০টির উপরে**) Too numerous to count (more than 10 piles)<br>99 বলা যাচ্ছে না/**পর্যবেক্ষণ**করা সম্ভব হয়নি Cannot tell / could not observe | ☐☐ |
| **C.827.** | 4202. | **উঠানে পশুর/পাখির মল/পায়খানা পড়ে আছে(নির্দিষ্টভাবে উল্লেখ করুন) Animal** feces present <u>within the</u> **<u>courtyard</u>**(mark all that apply) | ☐1 **পাখি বিশেষ(মুরগী/হাঁস/কবুতর)**Poultry (chicken, duck, pigeon)<br>☐2 **গরু/মহিষ** Cow / Buffalo |

| | | | |
|---|---|---|---|
| | | 55 অধিক সংখ্যক (স্তুপের সংখ্যা ১০টির উপরে) Too numerous to count (more than 10 piles)<br>99 বলা যাচ্ছে না/পর্যবেক্ষণকরা সম্ভব হয়নি Cannot tell / could not observe | ☐3 ছাগল/ভেড়া Goat / Sheep<br>☐4শুকর Pig<br>☐5 কুকুর অথবা বিড়াল Dog or Cat<br>☐8 অন্যান্য Other<br>☐9 কোনও মল/পায়খানা পাওয়া যায়নি NO FECES OBSERVED<br>☐99 পর্যবেক্ষণকরা সম্ভব হয়নি COULD NOT OBSERVE |
| | 4202.a | অন্যকোন খানা এই উঠান ব্যবহার করে কী?Do any other household share this **courtyard** | ১. হ্যাঁ<br>০. না |
| ৈনির্দিষ্টশিশু বেশীরভাগ সময় যেখানে সময় কাটায় তা নিরন্নপন ও যাচাইকরন্নন। বেইজ লাইনে নির্দিষ্টশিশু থাকবে না সেজন্য গর্ভবতী মা যেখানে বেশীরভাগ সময় কাটায় তা নিরন্নপনকরন্নন In the area where the target child spends the most time<br>*Probe and identify where the target child spends the most time. At baseline, the target child will not be present, so the search should be defined as the area where the pregnant mother spends the most time.* | | | |
| **C.828.** | 4203. | মানুষের মল/পায়খানা পড়ে আছে যেখানে নির্দিষ্টশিশু (বেইজলাইনে গর্ভবতী মা) বেশীরভাগ সময় কাটায়। ইহা উন্মুক্ত পায়খানা হিসেবে বিবেচিত হবে **Human** feces**in the area where the target child spends the most time** that could be considered open defecation<br>55 অধিক সংখ্যক (স্তুপের সংখ্যা ১০টির উপরে) Too numerous to count (more than 10 piles)<br>99 বলা যাচ্ছে না/পর্যবেক্ষণকরা সম্ভব হয়নি Cannot tell / could not observe | ☐☐ |
| **C.829.** | 4204. | পশুর/পাখির মল/পায়খানা পড়ে আছে যেখানে নির্দিষ্টশিশু (বেইজলাইনে গর্ভবতী মা)বেশীরভাগ সময় সময় কাটায় আছে ।(নিদ্দিষ্টভাবে উল্লেখ করুন) **Animal** feces in the **area where the target child spends the most time** (mark all that apply)<br>55 অধিক সংখ্যক (স্তুপের সংখ্যা ১০টির উপরে) Too numerous to count (more than 10 piles)<br>99 বলা যাচ্ছে না/পর্যবেক্ষণকরা সম্ভব হয়নি Cannot tell / could not observe | ☐1 পাখি বিশেষ(মুরগী/হাঁস/কবুতর )Poultry (chicken, duck, pigeon)<br>☐2 গরু/মহিষ Cow / Buffalo<br>☐3 ছাগল/ভেড়া Goat / Sheep<br>☐4শুকর Pig<br>☐5 কুকুর অথবা বিড়াল Dog or Cat<br>☐8 অন্যান্য Other<br>☐9 কোনও মল/পায়খানা পাওয়া যায়নি NO FECES OBSERVED<br>☐99 পর্যবেক্ষণকরা সম্ভব হয়নি |

| | | | |
|---|---|---|---|
| | | | COULD NOT OBSERVE |
| 👁 নিদ্দিষ্ট খানার পিছনে Behind the study house | | | |
| **C.830.** | 4205. | খানার পিছনে মানুষের মল/পায়খানা পড়ে আছে/থাকলে। ইহা উম্মুক্ত পায়খানা হিসেবে বিবেচিত হবে **Human** feces<u>behindthe</u> **<u>house</u>** that could be considered open defecation<br><br>55 অধিক সংখ্যক (স্তুপের সংখ্যা ১০টির উপরে) Too numerous to count (more than 10 piles)<br>99 বলা যাচ্ছে না/পর্যবেক্ষণকরা সম্ভব হয়নি Cannot tell / could not observe | ☐☐ |
| **C.831.** | 4206. | খানার পিছনে পশুর মল/পায়খানা পড়ে আছে/থাকলে। (নিদ্দিষ্টভাবে উল্লেখ করুন) **Animal** feces **<u>behind the house</u>** (mark all that apply)<br><br>55 অধিক সংখ্যক (স্তুপের সংখ্যা ১০টির উপরে) Too numerous to count (more than 10 piles)<br>99 বলা যাচ্ছে না/পর্যবেক্ষণকরা সম্ভব হয়নি Cannot tell / could not observe | ☐1 পাখি বিশেষ(মুরগী/হাঁস/কবুতর)Poultry (chicken, duck, pigeon)<br>☐2 গরু/মহিষ Cow / Buffalo<br>☐3 ছাগল/ভেড়া Goat / Sheep<br>☐4শুকর Pig<br>☐5 কুকুর অথবা বিড়াল Dog or Cat<br>☐8 অন্যান্য Other<br>☐9 কোনও মল/পায়খানা পাওয়া যায়নি NO FECES OBSERVED<br>☐99 পর্যবেক্ষণকরা সম্ভব হয়নি COULD NOT OBSERVE |

#### WASH Benefits Module 225 HOURS STRUCTURED OBSERVATIONS

HHID: ____________________

*Before you begin the observation ask about the people that are currently present in the household. This is so you know how to code each person that is present during the observation.* [PDA will save each entry as Mother, Father according to the codes in col 1.]

Target Household Person Index:

| **1.** Person | **2.** Sex<br>1.M<br>0.F | **3.** Enter the age (in years) | 4.Does the child:<br>**1.** Crawl<br>**2.** Walk<br>**0.** None of the above |
|---|---|---|---|
| **1.** Pregnant woman/Mother/ primary care giver<br>**2.** Father/ male care giver<br>**3.** Other adult(If 1-3, Skip col. 3)<br>**4.** Index child<br>**5.** Other Child | | | |

*Next, before you begin the observation take a look around the compound and locate possible hand washing stations.* <u>[PDA will save each entry as HWS 1, HWS 2 etc.]</u>

Hand washing station index:

| Hand washing station | Location | Hand cleansing agents present (LOOK AT HW) |
|---|---|---|
| HWS1<br>HWS2<br>HWS3 | 1…In/near main house (≤6 ft to entrance)<br><br>2…In/near latrine (≤6 ft to entrance)<br><br>3…In/near cooking area (≤6 ft to entrance)<br><br>4… >6 ft away from main house, latrine <u>and</u> cooking area | <u>[allow multiple options]</u><br><br>**[14]** ........ WATER (present directly from pump/tubewell/water source)<br>**[15]** ........ WATER (present in a container for handwashing purpose)<br>**[16]** ........ BAR SOAP (Body/hand Bar)<br>**[17]** ........ BAR SOAP (other)<br>**[18]** ........ POWDERED SOAP<br>**[19]** SOAPY WATER<br>**[20]** LIQUID SOAP<br>**[21]** Ash<br>**[22]** Mud/Sand<br>**[23]** ICDDR,B provided drum with tap<br>**[24]** Soapy water bottle<br>**[25]** (B) ICDDR,B provided smaller bucket<br>**[26]** (B) Other bucket<br>[27] (B) Basin<br>**[28]** Jug<br>**[29]** None of the above |

Determine an appropriate location so that child defecation, feeding events and hand washing event can be detected. You are allowed to move during this observation. Always keep the primary care giver in the line of vision. There might be more than one event going on at the same time. Focus on one event at a time. Record details about each observation according to the guide. We will focus on the target household for this observation. If you have notes or comments about related events please record them in paper with the HH ID, during these 5 hours.

Start time: --------/--------hh/mm (24 hour format)

End time: --------/---------hh/mm (24 hour format)

| **0.** Who are you observing? [PDA: open HH person list] | **1.** Which event are you observing? | 2. PDA to auto-record time at each observation | 3. Open sections within each event |
|---|---|---|---|
|  | **1)** Toilet use / defecation | --------/---------hh/mm | 1 Defecation/ Feces |

| | | | |
|---|---|---|---|
| **1.** Open target index list<br>**2.** Other household<br>- Adult, M<br>- Adult, F<br>- < 5 Child, M<br>- < 5 Child, F | | | Handling/ Disposal<br><br>2 Hand washing |
| | **2)** Other contact with feces | | 1 Defecation/ Feces Handling/ Disposal<br><br>3 Hand washing |
| | **3)** Food handling<br>**1.** Mashing food (bhorta)<br>**2.** Handling raw meat/fish<br>**3.** Cutting fruits<br>**4.** Cutting vegetables to be cooked<br>**5.** Cutting VEGETABLE FOR salad | | 1 Hand washing |
| | **4)** Eating | | 1. Other LNS event<br><br>2 Hand washing |
| | **5)** Eating (Index child) | | 1 Hand washing<br><br>2 Infant feeding |
| | **6)** Washing hands | | 1 Hand washing |
| | **7)** Water event | | 1 Water |

Defecation/ Feces Handling/ Disposal Record end time [allow option]

| **1.** Whose feces is it? | 2. Location of fecal contact | 3. Person handling feces: | 4. Feces disposed using (allow multiple options): | 5. Disposal site: | **6.** Hardware cleaning |
|---|---|---|---|---|---|
| **1.** [From person list]<br>**2.** Cow dung<br>**3.** Goat feces<br>**4.** Chicken feces<br>**5.** Other animal feces (if not known) | 1… Inside toilet (skip col. 7)<br><br>2… In the courtyard<br><br>3… Inside the household<br><br>4… Outside the household but not in the courtyard (beside/behind)<br><br>5… Potty<br><br>6… | Open list<br><br>99. No one (skip to end) | **1.** Hands only<br>**2.** Cloth/paper/leaves<br>**3.** Scrap material to scoop feces<br>**4.** Potty [do col.6, or else skip]<br>**5.** Local agricultural hoe<br>**6.** Sani scoop [do col.6, or else skip]<br>**7.** Did nothing<br>**8.** Other<br><br>**99.** Could not observe | **1.** Improved Latrine<br>**2.** Unimproved latrine<br>**3.** Open Pit / separate pit for child or animal feces<br>**4.** Bury it / Covered Pit<br>**5.** Undefined open site near the compound<br>**6.** Garbage disposal sites / dumps)<br>**7.** Bush / | **1.** Cleaned at tube well<br>**2.** Cleaned in the latrine<br>**3.** Cleaned at nearby water source (pond)<br>**4.** Cleaned in the courtyard using self poured water<br>**5.** Did not clean during this event |

| | | | | | |
|---|---|---|---|---|---|
| | Bush/Jungle/field<br>1. Other | | | forest / field<br>8. Nearby water (pond, canal, river)<br>99. Could not observe | observation |

Hand washing Record end time [allow option]

| 1. Were both hands cleansed? | 2. When were the hands cleansed? | 2. Hand cleansing materials: | 3. How were the hands washed? [multiple options] | 4. How were hands dried? | 5. Hand washing station index number | 6.Location if not at indexed HWS |
|---|---|---|---|---|---|---|
| **1....** Yes<br>**2....** No [skip to end]<br>99.Could not observe [skip to end] | (mostly applicable for food preparation times)<br>**1.** Before<br>**2.** After | **1....** Water only<br>**2....** Bar soap<br>**3....** Powdered Soap and water<br>**4....** Soapy water and rinse water<br>**5....** Soapy water only<br>**6....** Sanitizer<br>**7....** Other, Specify<br>9.... Could not observe | 1 Y 0 N<br>**1.** palms<br>**2.** back of hands<br>**3.** between fingers of hands<br>**4.** under fingernails | **1....** Not Dried<br>**2....** Air dried<br>**3....** Towel/Cloth<br>**4....** Clothing<br>99.Could not observe | [PDA open HW list to choose from] skip col 6<br>(put 99 if at a place not indexed) | *Record first option that fits*<br>1...In/near main house (≤6 ft to entrance)<br>2...In/near latrine (≤6 ft to entrance)<br>3...In/near cooking area (≤6 ft to entrance)<br>4... >6 ft away from main house, latrine <u>and</u> cooking area |

Water Event

| 1. What are you observing? | Where did respondent take the water from? | What type of storage container is it? | Treatment/treated | Handling<br>[multiple response] |
|---|---|---|---|---|

| | | | | |
|---|---|---|---|---|
| | | [multiple options] | | |
| **1.** Water collection<br>**2.** Water treatment<br>**3.** Water handling<br>4. Water storage<br>**5.** Drinking event | **1.** BROUGHT DIRECTLY FROM WATER SOURCE<br>**2.** BROUGHT DIRECTLY FROM STORAGE CONTAINER<br>**3.** BROUGHT DIRECTLY FROM WATER FILTER<br>**4.** STORED WATER WAS COVERED | **1.** 5L icddrb Kolshi<br>**2.** <5l container<br>**3.** >5L container<br>**4.** Hari (wide container)<br>**5.** Jug<br>**6.** Glass | **1.** Aquatabs<br>**2.** Boiling<br>**3.** Straining<br>**4.** Did not treat<br>**5.** Could not determine | **1.** rinsed glass/container with drinking water before filling<br>**2.** washed hands with water (no soap) before drinking water was obtained<br>**3.** washed hands with soap before drinking water obtained<br>**4.** hands touched / contacted the drinking water during retrieval<br>**5.** container/glass dipped into water container<br>**6.** ladle used to obtain water<br>**7.** water poured from container<br>**8.** water poured from tap |

Infant Feeding/Eating Record end time [allow option]

| 1. Type of food (allow multiple options) | 2. Person feeding (allow multiple options) | 3. How was LNS served: | 4. *How was LNS fed to child:* | 5. Consumption: |
|---|---|---|---|---|
| 1. Breastfed<br>2. LNS (fill Col 3,4,5 else skip)<br>3. Water/sugar water/honey water<br>4. Milk<br>5. Formula<br>6. Boiled rice<br>7. Khichuri (rice, lentil, potato, tomato, brinjal, beans)<br>8. Semolina/Porridge /Grains<br>9. Lentil<br>10. Eggs<br>11. Juice<br>12. Other liquids (ex. soup)<br>13. leafy vegetable<br>14. orange or yellow fruits/vegetables<br>15. other fruits/veg<br>16. Cheese/yogurt<br>17. Meat/fish/eggs<br>18. Packaged food (biscuits, chips, candy)<br>99. Other foods/drinks not observed | 1. Child (self)<br>2. Primary Caregiver, F<br>3. Primary caregiver, M<br>4. Secondary Caregiver F<br>5. Secondary caregiver, M<br>6. <6 year Child, F<br>7. <6 year Child, M<br>8. ≥7-15 year Child, F<br>9. ≥7-15 year Child, M<br>10. Other Adult, F<br>16. Other Adult, M | 1. Mixed into food with a spoon<br>2. Mixed into food by hand.<br>3. Fed directly from the packet | 1. By hand<br>2. By spoon<br>3. Direct from packet | 1. Consumed full packet<br>2. Partial and thrown away<br>3. Partial and eaten later |

Other LNS event

| 1. Did they consume LNS? | 2. Who consumed or took the LNS? | 3.How much was consumed? |
|---|---|---|
| 1.... Yes<br>2.... No [skip to end]<br>99.Could not observe [skip to end] | | |
| | 1. [From person list] | 1. Just a taste<br>2. Partial packet consumed<br>3. Whole packet consumed<br>4. Whole packet taken, consumption not observed |

Additional questions at the end of observation time: 1 Y 0 N

1. Did you feed LNS to your child today before I arrived?
2. Are you planning to feed LNS to your child later this afternoon or this evening?

WASH Benefits Module 50

**Lead exposure in-depth interview guide**

Note: Ask questions to the mother and the person in the household with the most knowledge regarding the exposure

#### Questions for mother

1. **How long have you lived in the current location?**
2. **How much of the rice that you eat comes from the fields that your household farm?**
3. **During which season do you purchase rice that is grown from somewhere else?**
4. **How much of the vegetables that you eat come from the fields your household farm?**
5. **Which vegetables do you purchase in the local market?**
6. **During which season do you purchase these vegetables?**
7. **How much of the fruits that you eat come from your household farm?**
8. **Which fruits do you purchase in the local market?**
9. **During which season do you purchase these fruits?**
10. **Do you or anyone in your household work with paint, solder, industrial waste or batteries?**
    a) **If yes, how often do you work with it?**
    b) **Are any of the batteries the type used in cars, trucks or motorcycle that contain acid?**
    c) **How are these batteries handled?**
11. **Do you or anyone in your household work in manufacturing or recycling goods? If yes, explain. Detail any contact with metals, solvents, chemicals, exhaust.**
12. **How close is the nearest road that motor vehicles use regularly to your home?**
13. **What foods do your purchase that come in a metal can? How often do you consume each of these foods?**
14. **How often do you wear kohl (makeup)? From where do you obtain the makeup? Probe details on whether it is a manufactured product or made. If it is a manufactured product, record the brand name of the product.**
15. **Does your house have tin roofs or tin walls? How long have you had them?**
16. **What material is your cooking pot made from?**
17. **What materials are your dinner plates made from? Are they painted?**
18. **What material is your water pot made from?**

#### Questions for person most knowledgeable regarding household farming

19. **How many harvests of rice does your land produce each year?**
20. **What other crops do you grow on your land (including small gardens)?**
21. **How often did you apply fertilizer on your land in the last year?**
22. **How do you decide how much fertilizer to use?**
23. **Did you use the same type of fertilizer for each application? If there were different types, probe and explicate the differences, e.g. different types of fertilizers for different times of year or for different crop?**
24. **What are the names of the types of fertilizers that you used? If there is more than one name, describe the roles of the different types.**
25. **How do you decide which fertilizer to use?**
26. **How do you decide how much fertilizer to use?**
27. **What packaging does the fertilizer come in?**
28. **Do you have to mix or otherwise prepare the fertilizer? If yes,**
    a) **Who is involved? (e.g. other family members?)**
    b) **Where is it mixed?**

c) Please describe the process.

29. What do you do with the leftover fertilizer packaging? Probe: Any re-use the packaging/container?
30. Who applies the fertilizer?
31. Describe the process of applying the fertilizer.
32. What do you do with leftover fertilizer?

33. How often did you apply pesticide on your land in the last year?
34. Did you use the same type of pesticide for each application? If there were different types, probe and explicate the differences, e.g. different types of pesticides for different times of year? for different crops? for different pests?
35. What are the names of the types of pesticides that you used? If there is more than one name, describe the roles of the different types.
36. Did you use pesticides before you saw any pests or do you always wait until you see pests? Why do you follow this practice?
37. How do you decide which pesticide to use?
38. How do you decide how much pesticide to use?
39. What packaging does the pesticide come in?
40. Do you have to mix or otherwise prepare the pesticide? If yes,
    a) Who is involved? (e.g. other family members?)
    b) Where is it mixed?
    c) Please describe the process.
41. What do you do with the leftover pesticide packaging? Probe: Any re-use the packaging/container?
42. Who applies the pesticide?
43. Describe the process of applying the pesticide.
44. What do you do with leftover pesticide?

45. How often did you apply herbicide on your land in the last year?
46. If he/she applied herbicide, why did you apply it?
47. Did you use the same type of herbicide for each application? If there were different types, probe and explicate the differences, e.g. different types of herbicides for different times of year? for different crops? for different pests?
48. What are the names of the types of herbicides that you used? If there is more than one name, describe the roles of the different types.
49. Did you use herbicides before you saw any weeds or do you always wait until you see weeds? Why do you follow this practice?
50. How do you decide which herbicide to use?
51. How do you decide how much herbicide to use?
52. What packaging does the herbicide come in?
53. Do you have to mix or otherwise prepare the herbicide? If yes,
    a) Who is involved? (e.g. other family members?)
    b) Where is it mixed?
    c) Please describe the process.
54. What do you do with the leftover herbicide packaging? Probe: Any re-use the packaging/container?
55. Who applies the herbicide?
56. Describe the process of applying the herbicide?
57. What do you do with leftover herbicide?

58. How often did you apply fungicide on your crops in the last year?
59. If he/she applied fungicide, why did you apply it?

60. **Did you use the same type of fungicide for each application? If there were different types, probe and explicate the differences, e.g. different types of fungicides for different times of year? for different crops? for different pests?**
61. **What are the names of the types of fungicides that you used? If there is more than one name, describe the roles of the different types.**
62. **Did you use fungicides before you saw any fungus or do you always wait until you see fungus? Why do you follow this practice?**
63. **How do you decide which fungicide to use?**
64. **How do you decide how much fungicide to use?**
65. **What packaging does the fungicide come in?**
66. **Do you have to mix or otherwise prepare the fungicide? If yes,**
    a) **Who is involved? (e.g. other family members?)**
    b) **Where is it mixed?**
    c) **Please describe the process.**
67. **What do you do with the leftover fungicide packaging? Probe: Any re-use the packaging/container?**
68. **Who applies the fungicide?**
69. **Describe the process of applying the fungicide?**
70. **What do you do with leftover fungicide?**

71. **Did you apply any other chemicals to crops/lands in the last year that we have not already discussed? If yes, explicate what else he/she used, how often, its name, and why it was used.**

**ওয়াশ বেনিফিট মডিউল ৫০ (Module 50)**

**সীসার প্রভাব সম্পর্কিত নিবিড় সাক্ষাৎকারের গাইডলাইন**

[ নোট: মা এবং পরিবারের সবচেয়ে তথ্য সমৃদ্ধ ব্যাক্তিকে এক্সপোজার বিষয়ক প্রশ্নগুলো করুন ]

**মায়ের জন্য প্রশ্নাবলী**

১. বর্তমান জায়গায় আপনি কতদিন ধরে বসবাস করছেন?

২. আপনারা যে ভাত খান তার কি পরিমাণ আপনাদের গৃহস্থালী খামার থেকে আসে?

৩. আপনি সাধারণত কোন ঋতু/সময়ে চাল কেনেন যা অন্য কোন জায়গায় উৎপাদন করা হয়?

৪. আপনার খাবারের কি পরিমাণ সবজি আপনার নিজের খামার/গৃহস্থালী থেকে আসে?

৫. স্থানীয় বাজার থেকে আপনি কি ধরণের সবজি কেনেন?

৬. এই সব সবজি আপনি কোন ঋতু/সময়ে কেনেন?

৭. আপনার খাবারের কি পরিমাণ ফলমূল আপনার নিজের খামার/গৃহস্থালী থেকে আসে?

৮. স্থানীয় বাজার থেকে আপনি কি কি ফলমূল কেনেন?

৯. কোন ঋতুতে/সময়ে আপনি এইসব ফল কেনেন?

১০. আপনি অথবা আপনার পরিবারের কেউ কি রং, ঝালাই, শিল্পবর্জ্য অথবা ব্যাটারির কাজ করে?

ক) যদি হ্যাঁ হয়, তাহলে কত ঘনঘন এগুলো করেন?

খ) এখানে কি কোন এসিডযুক্ত ব্যাটারী আছে যা গাড়ি,ট্রাক অথবা মোটর সাইকেলে ব্যবহৃত হয়।

গ) সে সব ব্যাটারিগুলো কিভাবে নাড়াচাড়া করেন?

১১. আপনি অথবা আপনার পরিবারের কেউ কি পণ্য প্রক্রিয়াজাতকরণ এবং পূর্ণ প্রক্রিয়াজাতকরণের কাজের সাথে যুক্ত আছে? যদি থাকে দয়া করে বর্ণনা করুন। ধাতু,দ্রাবক/তরল পদার্থ, রাসায়নিক, ধোয়া নির্গম নল প্রভৃতির সাথে যে কোন ধরণের স্পর্শ বিস্তারিতভাবে বর্ণনা করুন।

১২. নিয়মিত মোটরযান চলাচলকারী রাস্তা আপনার বাড়ী থেকে কত কাছে?

১৩. ধাতব পাত্রে প্রক্রিয়াজাত কি কি ধরণের খাবার আপনি কেনেন? আপনি কত ঘনঘন এই সব খাবার খান?

১৪. আপনি সাধারণত কখন কখন সুরমা (প্রসাধন) ব্যাবহার করেন? কোথা থেকে আপনি এই প্রসাধন নেন? এটা কোন প্রক্রিয়াজাতকৃত অথবা তৈরীকৃত পণ্য কিনা তা বিস্তারিত জানতে চেস্টা করুন। এটা যদি প্রক্রিয়াজাত পণ্য হয় তাহলে ব্রান্ড নাম সংগ্রহ করুন ।

১৫. আপনার ঘরে কি কোন টিনের ছাদ বা টিনের বেড়া আছে? কত দিন ধরে আছে?

১৬. আপনার রান্নার হাড়ি পাতিল কি দিয়ে তৈরী?

১৭. আপনার খাবার থালা বাসন কি দিয়ে তৈরী/কিসের তৈরী? এগুলো কি রং করা?

১৮. আপনার পানির পাত্র, কি দিয়ে তৈরী/ কিসের তৈরী?

**গৃহস্থালী খামার সম্পর্কে সবচেয়ে তথ্যসমৃদ্ধ ব্যাক্তির জন্য প্রশ্ন**

১৯. প্রতি বছর আপনার জমি থেকে কত ধরণের ধান সংগ্রহ/তোলা হয়?

২০. আপনার জমিতে অন্য আর কি কি ধরণের ফসল আপনি চাষ করেন (ছোট বাগান সহ)?

২১. গতবছর আপনার জমিতে আপনি কত ঘনঘন রাসায়নিক সার ব্যবহার করেছেন?

২২. কি পরিমান রাসায়নিক সার ব্যবহার করতে হবে সেটা আপনি কিভাবে ঠিক করেন?

২৩. প্রতিক্ষেত্রে আপনি কি একই ধরণের রাসায়নিক সার ব্যবহার করেন? যদি ভিন্ন ধরণের হয় তাহলে পার্থক্যগুলো বিশদভাবে জানার চেস্টা করুন। যেমন, বিভিন্ন ফসলের জন্য বছরের বিভিন্ন সময়ে বিভিন্ন ধরণের রাসায়নিক সার?

২৪. যে যে ধরণের রাসায়নিক সার আপনি ব্যবহার করেছেন, সেগুলোর নাম কি? যদি একাধিক ধরণের থাকে তাহলে তাদের প্রত্যেকটির ভূমিকা বর্ণনা করুন।

২৫. কোন ধরণের রাসায়নিক সার ব্যবহার করা হবে সেটা আপনি কিভাবে নির্ধারণ করেন?

২৬. কি পরিমাণ রাসায়নিক সার ব্যবহার করা হবে সেটা আপনি কিভাবে নির্ধারণ করেন?

২৭. কি ধরণের মোড়কে রাসায়নিক সারগুলো থাকে?

২৮. অপনাকে কি কখনো রাসায়নিক সার মিশিয়ে বা অন্য কোন উপায়ে প্রস্তুত করতে হয়? যদি হয়,

ক) কে সেটা করে/কারা কারা যুক্ত থাকে (যেমন, পরিবারের অন্যান্য সদস্য?)

খ) কোথায় সেগুলো মেশানো হয়?

গ) দয়া করে সার মেশানোর পদ্ধতিটি বর্ণনা করুন।

২৯. সারের পরিত্যাক্ত মোড়কগুলো দিয়ে আপনি কি করেন? প্রোব: মোড়ক/ধারণ পাত্রের কোন প্রকার পুনঃব্যবহার?

৩০. কে এই রাসায়নিক সার প্রয়োগ করে?

৩১. এই রাসায়নিক সার প্রয়োগের পদ্ধতি বর্ণনা করুন।

৩২. পরিত্যাক্ত রাসায়নিক সার দিয়ে আপনি কি করেন?

৩৩. গতবছর কতবার/কতদিন পর পর আপনি জমিতে কীটনাশক ব্যবহার করেছেন?

৩৪. প্রতিক্ষেত্রে আপনি কি একই ধরণের কীটনাশক ব্যবহার করছেন? যদি ভিন্ন ধরণের হয় তাহলে পার্থক্যগুলো বিশদভাবে জানার চেস্টা করুন করুন। যেমন, বছরের বিভিন্ন সময়ে বিভিন্ন ধরণের কীটনাশক, বিভিন্ন ফসলের জন্য, বিভিন্ন পোকামাকড়ের জন্য।

৩৫. আপনি যে যে ধরণের কীটনাশক ব্যবহার করেছেন সেগুলোর নাম কি? যদি একাধিক ধরণের থাকে তাহলে তাদের প্রত্যেকটির ভূমিকা বর্ণনা করুন।

৩৬. পোকা মাকড় দেখার আগে কি আপনি কীটনাশক ব্যবহার করেছেন অথবা আপনি সবসময়ই পোকা-মাকড় দেখা পর্যন্ত অপেক্ষা করেন? আপনি কেন এই রীতি মেনে চলেন?

৩৭. কোন ধরণের কীটনাশক ব্যবহার করা হবে এটা আপনি কিভাবে নির্ধারণ করেন?

৩৮. কি পরিমাণ কীটনাশক ব্যবহার করা হবে এটা আপনি কিভাবে নির্ধারণ করেন?

৩৯. কি ধরণের মোড়কে কীটনাশকগুলো থাকে?

৪০. আপনাকে কি কখনও কীটনাশক মিশিয়ে বা অন্য কোন উপায়ে প্রস্তুত করতে হয়? যদি হয়,

ক) কে সেটা করে/কারা কারা এ কাজের সাথে যুক্ত থাকে (যেমন, পরিবারের অন্যান্য সদস্য?)

খ) কোথায় এ মিশ্রণের কাজটি করা হয়?

গ) দয়া করে পদ্ধতিটি বর্ণনা করুন।

৪১. পরিত্যাক্ত কীটনাশকের মোড়কগুলো দিয়ে আপনি কি করেন? প্রোব: মোড়ক/ধারণ পাত্রের কোন প্রকার পুনঃব্যবহার?

৪২. কে এই কীটনাশক প্রয়োগ করেন?

৪৩. কীটনাশক প্রয়োগের পদ্ধতি বর্ণনা করুন।

৪৪. অবশিষ্ট কীটনাশক দিয়ে আপনি কি করেন?

৪৫. গত বছরে কত ঘনঘন আপনি জমিতে আগাছা নাশক ব্যাবহার করেছেন?

৪৬. সে যদি আগাছানাশক ব্যাবহার করে তাহলে কেন এটা করেছে?

৪৭. আপনি কি প্রতিক্ষেত্রেই একই ধরণের আগাছানাশক প্রয়োগ করেছেন যদি ভিন্ন ধরণের হয় তাহলে পার্থক্যগুলো বিশদভাবে জানার চেস্টা করুন করুন। যেমন: বছরে বিভিন্ন সময়ে, বিভিন্ন ধরণের আগাছানাশক বিভিন্ন ফসলের জন্য? বিভিন্ন পোকামাকড়ের জন্য?

৪৮. আপনি যেসব আগাছানাশক ব্যবহার করেছেন সেগুলোর নাম কি? যদি একাধিক ধরণের থাকে তাহলে তাদের প্রত্যেকটির ভূমিকা বর্ণনা করুন।

৪৯. কোন আগাছা দেখার আগেই কি আপনি আগাছানাশক ব্যবহার করেছেন? অথবা আগাছা দেখা পর্যন্ত অপেক্ষা করেন? আপনি কেন এই রীতি অনুসরণ করেন?

৫০. কোন ধরণের আগাছানাশক ব্যবহার করা হবে এটা আপনি কিভাবে নির্ধারণ করেন?

৫১. কি পরিমাণ আগাছানাশক ব্যবহার করা হবে এটা আপনি কিভাবে নির্ধারণ করেন?

৫২. কি ধরণের মোড়কে এই আগাছানাশকগুলো থাকে?

৫৩. আপনাকে কি কখনও আগাছানাশক মিশিয়ে বা অন্য কোন উপায়ে প্রস্তুত করতে হয়? যদি হ্যাঁ হয়,

ক) কে সেটা করে/ কারা কারা এ কাজের সাথে যুক্ত থাকে (যেমন, পরিবারের অন্যান্য সদস্য?)

খ) কোথায় মেশানো হয়?

গ) দয়া করে পদ্ধতিটি বর্ণনা করুন।

৫৪. পরিত্যাক্ত আগাছানাশকের মোড়কগুলো দিয়ে আপনি কি করেন? মোড়ক/ধারণ পাত্রের পুনঃব্যবহার প্রোব করুন ।

৫৫. কে এই আগাছানাশক প্রয়োগ করেন?

৫৬. আগাছানাশক প্রয়োগ পদ্ধতি বর্ণনা করুন।

৫৭. অবশিষ্ট আগাছানাশক দিয়ে আপনি কি করেন?

৫৮. গতবছর কতবার/কতদিন পর পর আপনার জমিতে ছত্রাকনাশক ব্যবহার করেছেন?

৫৯. সে যদি ছত্রাকনাশক ব্যবহার করে থাকে তাহলে কেন করেছে?

৬০. প্রতিক্ষেত্রে কি আপনি একই ধরণের ছত্রাকনাশক ব্যবহার করছেন? যদি ভিন্ন ধরণের হয় তাহলে পার্থক্যগুলো বিশদভাবে জানার চেস্টা করুন করুন। প্রোব: বছরে বিভিন্ন সময়ে, বিভিন্ন ধরণের ছত্রাকনাশক বিভিন্ন ফসলের জন্য? বিভিন্ন পোকামাকড়ের জন্য?

৬১. আপনি যেসব ছত্রাকনাশক ব্যবহার করেছেন সেগুলোর নাম কি? যদি একাধিক ধরণের থাকে তাহলে তাদের প্রত্যেকটির ভূমিকা বর্ণনা করুন।

৬২. কোন ছত্রাক দেখার আগেই কি আপনি ছত্রাকনাশক ব্যবহার করেছেন? অথবা ছত্রাক দেখা পর্যন্ত অপেক্ষা করেন? আপনি কেন এই রীতি অনুসরণ করেন?

৬৩. কোন ধরণের ছত্রাকনাশক ব্যবহার করা হবে এটা আপনি কিভাবে নির্ধারণ করেন?

৬৪. কি পরিমাণ ছত্রাকনাশক ব্যবহার করা হবে এটা আপনি কিভাবে নির্ধারণ করেন?

৬৫. কি ধরণের মোড়কে এই ছত্রাকনাশকগুলো থাকে?

৬৬. আপনাকে কি কখনও ছত্রাকনাশক মিশিয়ে বা অন্য কোন উপায়ে প্রস্তুত করতে হয়? যদি হ্যাঁ হয়,

ক) কে সেটা করে/কারা কারা এ কাজের সাথে যুক্ত থাকে (যেমন, পরিবারের অন্যান্য সদস্য?)

খ) কোথায় মেশানো হয়?

গ) দয়া করে পদ্ধতিটি বর্ণনা করুন

৬৭. পরিত্যাক্ত ছত্রাকনাশকের মোড়কগুলো দিয়ে আপনি কি করেন? মোড়ক/ধারণ পাত্রের পুনঃব্যবহার প্রোব করুন।

৬৮. কে এই ছত্রাকনাশক প্রয়োগ করেন?

৬৯. ছত্রাকনাশকের প্রয়োগ পদ্ধতি বর্ণনা করুন।

৭০. অবশিষ্ট ছত্রাকনাশক দিয়ে আপনি কি করেন?

৭১. আপনি কি গত বছরে জমিতে/ফসলে অন্য কোন রাসায়নিক প্রয়োগ করেছেন যা আমরা ইতিমধ্যে আলোচনা করিনি। যদি হ্যাঁ হয়, তাহলে সে কি ব্যবহার করেছে, কতবার, এর নাম এবং কেন এটা ব্যবহার করা হয়েছে বিস্তারিত বর্ণনা করুন।

**Agrochemical salesman in-depth interview guide**

1. **How long have you worked in the current location?**
2. **How many types of fertilizer do you sell?**
3. **What names do you and farmers use to refer to the different types of fertilizers?**
4. **Why do farmers purchase different types of fertilizers?**
5. **Where do you secure each type of these fertilizers?**
6. **Where are each type of these fertilizers manufactured?**
7. **Which fertilizer is your biggest seller? Why?**

8. **How many types of pesticide do you sell?**
9. **What names do you and farmers use to refer to the different types of pesticides?**
10. **Why do farmers purchase different types of pesticides?**
11. **Where do you secure each type of these pesticides?**
12. **Where are each type of these pesticides manufactured?**
13. **Which pesticide is your biggest seller? Why?**

14. **How many types of herbicide do you sell?**
15. **What names do you and farmers use to refer to the different types of herbicides?**
16. **Why do farmers purchase different types of herbicides?**
17. **Where do you secure each type of these herbicides?**
18. **Where are each type of these herbicides manufactured?**
19. **Which herbicide is your biggest seller? Why?**

20. **How many types of fungicide do you sell?**
21. **What names do you and farmers use to refer to the different types of fungicides?**
22. **Why do farmers purchase different types of fungicides?**
23. **Where do you secure each type of these fungicides?**
24. **Where are each type of these fungicides manufactured?**
25. **Which fungicide is your biggest seller? Why?**

**ওয়াশ বেনিফিট মডিউল ৫১ (Module 51)**

কৃষি-রাসায়নিক বিক্রেতার জন্য নিবিড় সাক্ষাৎকার গাইডলাইন

১. আপনি কতদিন ধরে এই এলাকায় কাজ করছেন?
২. আপনি কত ধরনের সার বিক্রি করেন?
৩. বিভিন্ন ধরনের সারকে বোঝাতে আপনি এবং অন্যান্য কৃষকরা কি কি নাম ব্যবহার করে থাকেন?
৪. কেন কৃষকরা বিভিন্ন ধরনের সার ক্রয় করে?
৫. এই সারগুলোর প্রত্যেকটিকে আপনারা কোথায় থেকে সংগ্রহ করেন?
৬. এই সারগুলো প্রত্যেকটি কোথায় কোথায় উৎপাদিত/প্রক্রিয়াজাতকরণ হয়?
৭. কোন সারটি সবচেয়ে বেশী বিক্রি হয়? কেন?
৮. কত ধরনের কীটনাশক আপনি বিক্রি করেন?
৯. বিভিন্ন ধরনের কীটনাশক বোঝাতে আপনি এবং অন্যান্য কৃষকরা কি কি নাম ব্যবহার করেন?
১০. কেন কৃষকরা বিভিন্ন ধরনের কীটনাশক ক্রয় করে?
১১. এই কীটনাশকগুলোর প্রত্যেকটিকে আপনারা কোথায় থেকে সংগ্রহ করেন?
১২. এই কীটনাশকগুলোর প্রত্যেকটি কোথায় কোথায় উৎপাদিত/প্রক্রিয়াজাতকরণ হয়?
১৩. কোন কীটনাশকটি সবচেয়ে বেশী বিক্রি হয়? কেন?

১৪. কত ধরনের আগাছানাশক আপনি বিক্রি করেন?

১৫. বিভিন্ন ধরনের আগাছানাশক বোঝাতে আপনি এবং অন্যান্য কৃষকরা কি কি নাম ব্যবহার করেন?
১৬. কেন কৃষকরা বিভিন্ন ধরনের আগাছানাশক ক্রয় করে?
১৭. এই আগাছানাশকগুলোর প্রত্যেকটিকে আপনারা কোথায় থেকে সংগ্রহ করেন?
১৮. এই আগাছানাশকগুলোর প্রত্যেকটি কোথায় কোথায় উৎপাদিত/প্রক্রিয়াজাতকরণ হয়?
১৯. কোন আগাছানাশকটি সবচেয়ে বেশী বিক্রি হয়? কেন?

২০. কত ধরনের ছত্রাকনাশক আপনি বিক্রি করেন?
২১. বিভিন্ন ধরনের ছত্রাকনাশক বোঝাতে আপনি এবং অন্যান্য কৃষকরা কি কি নাম ব্যবহার করেন?
২২. কেন কৃষকরা বিভিন্ন ধরনের ছত্রাকনাশক ক্রয় করে?
২৩. এই ছত্রাকনাশকগুলোর প্রত্যেকটিকে আপনারা কোথায় থেকে সংগ্রহ করেন?
২৪. এই ছত্রাকনাশকগুলোর প্রত্যেকটি কোথায় কোথায় উৎপাদিত/প্রক্রিয়াজাতকরণ হয়?
২৫. কোন ছত্রাকনাশকটি সবচেয়ে বেশী বিক্রি হয়? কেন?

WASH Benefits Module 52

**সীসার প্রভাব সম্পর্কীত প্রশ্নপত্র (Lead exposure case-control questionnaire)**

Note: **সীসার প্রভাব সম্পর্কে ভাল ধারনা আছে এমন মা অথবা পরিবারের অন্য সদস্যদের জন্য প্রশ্নাবলী**
(Ask questions to the mother and the person in the household with the most knowledge regarding the exposure)

**Questions for mother** (মায়ের জন্য প্রশ্নসমূহ)

1. **বর্তমান এই খানাতে আপনি কতদিন যাবত বসবাস করছেন? (How long have you lived in the current location?)**
   ________

2. **আপনি/ আপনার খানা সদস্যরা যে পরিমান চাল খান তার কতটুকু আপনার/আপনাদের পরিবারের চাষের জমি থেকে আসে? (How much of the rice that you eat comes from the fields that your household farm?)**
   1) একটুও না (None)
   2) সামান্য, অর্ধেকেরও কম (Some but less than half)
   3) প্রায় অর্ধেক(About half)
   4) অর্ধেকেরও বেশী, কিন্তু সবটুকু না(More than half, but not all)
   5) সবটুকু (All)

3. **আপনি/ আপনার খানা সদস্যরা যে পরিমান শাকসব্জি খান তার কতটুকু আপনার/আপনাদের পরিবারের চাষের জমি থেকে আসে? (How much of the vegetables that you eat come from the fields your household farm?)**
   1) একটুও না (None)
   2) সামান্য, অর্ধেকেরও কম (Some but less than half)
   3) প্রায় অর্ধেক(About half)
   4) অর্ধেকেরও বেশী, কিন্তু সবটুকু না(More than half, but not all)
   5) সবটুকু (All)

4. **আপনি/ আপনার খানা সদস্যরা যে পরিমান ফলমূল খান তার কতটুকু আপনার/আপনাদের পরিবারের বাগান থেকে আসে? (How much of the fruits that you eat come from your household farm?)**
   1) একটুও না (None)
   2) সামান্য, অর্ধেকেরও কম (Some but less than half)
   3) প্রায় অর্ধেক(About half)
   4) অর্ধেকেরও বেশী, কিন্তু সবটুকু না (More than half, but not all)
   5) সবটুকু (All)

5. আপনি কি নিচের উপাদানগুলোর যেকোনটি নিয়ে বছরে অন্ততঃ কয়েকবার কাজ করেন? (প্রত্যেকটি অপশনের ক্ষেত্রে আলাদাভাবে উত্তর দিন। হ্যাঁ=1, না=0)[Do you work at least a few times per year with any of these materials? (answer each separately Yes = 1, No = 0)]
   1) রং (Paint) ___
   2) ঝালাই (Solder) ___
   3) শিল্প বর্জ্য (Industrial waste) ___
   4) ট্রাক, গাড়ি বা মোটর সাইকেলের ব্যাটারি (Truck, car or motorcycle batteries) ___
   5) নির্মান শিল্পের উপাদান (Manufacturing metals) ___
   6) দ্রাবক (Solvents) ___
   7) রাসায়নিক পদার্থসমূহ (Chemicals) ___
   8) নির্গত ধোঁয়া (Exhaust fumes) ______
   9) পুর্নব্যবহারযোগ্য পণ্য (Recycling goods) _____

6. আপনার পরিবারের কোন সদস্য কি নিচের উপাদানগুলোর যেকোনটি নিয়ে বছরে অন্ততঃ কয়েকবার কাজ করে? (প্রত্যেকটি অপশনের ক্ষেত্রে আলাদাভাবে উত্তর দিন। হ্যাঁ=1, না=0) [Does anyone in your household work at least a few times per year with any of these materials? (answer each separately Yes = 1, No = 0)]
   1) রং (Paint) ___
   2) ঝালাই (Solder) ___
   3) শিল্প বর্জ্য (Industrial waste) ___
   4) ট্রাক, গাড়ি বা মোটর সাইকেলের ব্যাটারি (Truck, car or motorcycle batteries) ___
   5) নির্মান শিল্পের উপাদান (Manufacturing metals) ___
   6) দ্রাবক (Solvents) ___
   7) রাসায়নিক পদার্থসমূহ (Chemicals) ___
   8) নির্গত ধোঁয়া (Exhaust fumes) ______
   9) পুর্নব্যবহারযোগ্য পণ্য (Recycling goods) _____

7. নিয়মিতভাবে মটরযান চলাচলের রাস্তাটি আপনার বাড়ি থেকে কতটা কাছাকাছি অবস্থিত? (How close is the nearest road that motor vehicles use regularly to your home?)
   1) 50 মিটারের কম (< 50 meters)
   2) 51 থেকে 200 মিটারের মধ্যে (51 – 200 meters)
   3) 201 থেকে 500 মিটারের মধ্যে (201 – 500 meters)
   4) 501 থেকে 1,000 মিটারের মধ্যে (501 – 1,000 meters)
   5) 1,000 মিটারের বেশি (> 1,000 meters)

8. কতদিন পর পর আপনি ধাতবপাত্রে প্রক্রিয়াজাত করা খাবার কিনে থাকেন? (How often do you consume food that comes from a metal can?)
   1) কখনোই না (Never)
   2) বছরে কয়েক বার (A few times per year)
   3) প্রায় প্রতি মাসে এক বার (About once per month)
   4) প্রায় প্রতি সপ্তাহে এক বার (About once per week)
   5) অধিকাংশ দিনই (Most days)

9. আপনি কি প্রায়ই দোকান থেকে কেনা মেকআপ ব্যবহার করেন? (How often do you wear kohl (makeup) that you purchase from a store?)
   1) কখনোই না (Never)
   2) বছরে কয়েক বার (A few times per year)

3) প্রায় প্রতি মাসে এক বার (About once per month)
4) প্রায় প্রতি সপ্তাহে এক বার (About once per week)
5) অধিকাংশ দিনই (Most days)

10. আপনার ঘরের ছাদ বা দেয়াল কি টিনের তৈরি? (Does you house have tin roofs or tin walls?)
    1) হ্যাঁ (Yes)
    2) না (No)
11. উত্তর হ্যাঁ হলে, কতদিন থেকে আপনার ঘরের ছাদ বা দেয়াল টিনের তৈরি? (If yes, how long have you had them?) _______
12. আপনার হাঁড়ি-পাতিল কোন উপাদান দিয়ে তৈরি? (প্রত্যেকটি অপশনের ক্ষেত্রে আলাদাভাবে উত্তর দিন । হ্যাঁ=1, না=0) [What materials are your cooking pot made from? (answer each separately Yes = 1, No = 0)]
    1) অ্যালুমিনিয়াম (Aluminum)
    2) রংবিহীন কাদামাটি/ সিরামিক (Unpainted clay / ceramic)
    3) রং করা কাদামাটি/ সিরামিক (Painted clay / ceramic)
    4) ঢালাই লোহা (Cast iron)
13. আপনার খাবারের থালা, খাবার পরিবেশনের ডিশ এবং হাঁড়ি-পাতিল কিসের তৈরি? (প্রত্যেকটি অপশনের ক্ষেত্রে আলাদাভাবে উত্তর দিন । হ্যাঁ=1, না=0) [What materials are your plates, serving dishes and utensils made from? (answer each separately Yes = 1, No = 0)]
    1) অ্যালুমিনিয়াম (Aluminum)
    2) রংবিহীন কাদামাটি/ সিরামিক (Unpainted clay / ceramic)
    3) রং করা কাদামাটি/ সিরামিক (Painted clay / ceramic)
    4) ঢালাই লোহা (Cast iron)
    5) প্লাস্টিক (plastic)

পারিবারিক চাষাবাদ সম্বন্ধে জ্ঞানবুদ্ধিসম্পন্ন ব্যক্তির জন্য প্রশ্নাবলীঃ (Questions for person most knowledgeable regarding household farming)

14. আপনার জমিতে প্রতিবছর কয় ধরনের ধান উৎপাদন করা হয়? (How many harvests of rice does your land produce each year? ___
15. আপনার জমিতে ধান ছাড়া আর কি কি শস্য উৎপাদন করে থাকেন (ছোট বাগানসহ) ? (প্রত্যেকটি শস্যের ক্ষেত্রে আলাদাভাবে উত্তর দিন । হ্যাঁ=1, না=0) [What other crops do you grow on your land (including small gardens)? (answer each separately Yes = 1, No = 0)]
    1) মসুর ডাল (Lentils)
    2) ছোলা [Chick peas (chola)]
    3) বেগুন (Eggplant)
    4) টমেটো (Tomato)
    5) শসা (Cucumbers)
    6) ঢেঁড়স (Lady fingers)
    7) লেটুস (Lettuce)
    8) পেঁয়াজ (Onions)
    9) গাজর (Carrots)
    10) আলু (Potatoes)
    11) আম (Mango)
    12) কমলা (Komla)
    13) লিঁচু (Lychee)
    14) (Jackfruit)

15) (Banana)
88. Others: Specify___________________________

সার সংক্রান্ত প্রশ্নাবলী (Fertilizer Questions)

16. **গত বছর আপনার জমিতে কতবার সার প্রয়োগ করেছেন? (How many times did you apply fertilizer on your land in the last year?) ____**
17. **আমি আপনাকে প্রচলিত সারের একটি তালিকা পড়ে শোনাব। দয়া করে আপনি আমাকে বলবেন, এসব সারের মধ্যে কোন কোন সার গত বছর আপনি কতবার আপনার জমিতে ধান উৎপাদনের জন্য প্রয়োগ/ব্যবহার করেছেন? (I will read a list of available fertilizers. Please tell me how many times you applied each of these types of fertilizer on the land you used for** growing rice **in the last year.)**
    1) টাইপ A [Type A (Type names will be identified in the qualitative study)]
    2) টাইপ B (Type B)
    3) টাইপ C (Type C)
    4) টাইপ D (Type D)
    5) টাইপ E (Type E)
    6) অন্যান্য (Other) _______
18. **আমি আপনাকে পুনরায় প্রচলিত সারের তালিকাটি পড়ে শোনাব। দয়া করে আপনি আমাকে বলবেন, এসব সারের মধ্যে কোন কোন সার গত বছর আপনি কতবার আপনার জমিতে অন্যান্য শস্য (ধান ছাড়া) উৎপাদনের জন্য প্রয়োগ/ব্যবহার করেছেন? (I will again read the list of available fertilizers. Please tell me how many times you applied each of these types of fertilizer on the land you used for** growing other crops **in the last year.)**
    1) টাইপ A[Type A (Type names will be identified in the qualitative study)]
    2) টাইপ B(Type B)
    3) টাইপ C(Type C)
    4) টাইপ D(Type D)
    5) টাইপ E(Type E)
    6) অন্যান্য (Other) _______
19. **সারগুলো জমিতে প্রয়োগ করার পূর্বে কে এগুলো প্রস্তুত করেন/নাড়াচাড়া করেন? (প্রত্যেকটি অপশনের ক্ষেত্রে আলাদাভাবে উত্তর দিন । হ্যাঁ=1, না=0) [Who handled the fertilizer while** preparing **to apply it? (answer each separately Yes = 1, No = 0)]**
    1) সবচেয়ে ছোট শিশুর বাবা (Father of the youngest child in the household)
    2) সবচেয়ে ছোট শিশুর মা (Mother of the youngest child in the household)
    3) সবচেয়ে ছোট শিশুর সহোদর (Sibling of the youngest child in the household)
    4) সবচেয়ে ছোট শিশুর দাদা-দাদী (Grandparent of the youngest child in the household)
    5) সবচেয়ে ছোট শিশুর চাচাতো ভাই-বোন (Cousin of the youngest child in the household)
    6) সবচেয়ে ছোট শিশুর চাচা/মামা (Uncle of the youngest child in the household)
    7) সবচেয়ে ছোট শিশুর চাচী/মামী (Aunt of the youngest child in the household)
    8) অন্যান্য (Other)_________________

20. **সারগুলো প্রয়োগ করার/ছিটানোর জন্য কোথায় প্রস্তুত করা হয়েছিল? (Where was the fertilizer prepared for application)**
    1) বাড়িতে (In the house)
    2) বাড়ির কাছে (Near the house)
    3) জমিতে (In the field)
    4) অন্যান্য (Other)_________________________

21. **জমিতে ব্যবহারের পর অবশিষ্ট সারগুলো আপনি কি করেছিলেন? (What did you do with the leftover fertilizer?)**
    1) অবশিষ্ট ছিল না (Did not have any)

2) বাড়িতে সংরৰন করেছিলাম (Stored it in the house)
3) কাউকে দান করে দিয়েছিলাম বা বিক্রি করে দিয়েছিলাম (Gave it away or sold it)
4) অন্যান্য (Other)

22. জমিতে ব্যবহারের পর সারের অবশিষ্ট প্যাকেটগুলো আপনি কি করেছিলেন? (প্রত্যেকটি অপশনের ৰেত্রে আলাদাভাবে উত্তর দিন । হ্যাঁ=1, না=0) [What did you do with the leftover fertilzer packaging? (answer each separately Yes = 1, No = 0]
1) বাড়িতে সংরৰন করেছিলাম (Stored it at home)
2) খাদ্য মজুদ রাখার কাজে ব্যবহার করেছিলাম (Used it to store food)
3) পানি সংরৰনের কাজে ব্যবহার করেছিলাম (Used it to store water)
4) গৃহস্থালী সামগ্রী সংরৰনের কাজে ব্যবহার করেছিলাম (Used it to store household goods)
5) শিশুদের খেলতে দিয়েছিলাম (Let the children play with it)
6) ফেলে দিয়েছিলাম (Discarded it)

23. কে সারগুলো জমিতে দিয়েছিল? (প্রত্যেকটি অপশনের ৰেত্রে আলাদাভাবে উত্তর দিন । হ্যাঁ=1, না=0) [Who applied the fertilizer to the land? (answer each separately Yes = 1, No = 0]
1) সবচেয়ে ছোট শিশুর বাবা (Father of the youngest child in the household)
2) সবচেয়ে ছোট শিশুর মা (Mother of the youngest child in the household)
3) সবচেয়ে ছোট শিশুর সহোদর (Sibling of the youngest child in the household)
4) সবচেয়ে ছোট শিশুর দাদা-দাদী (Grandparent of the youngest child in the household)
5) সবচেয়ে ছোট শিশুর চাচাতো ভাই-বোন (Cousin of the youngest child in the household)
6) সবচেয়ে ছোট শিশুর চাচা/মামা (Uncle of the youngest child in the household)
7) সবচেয়ে ছোট শিশুর চাচী/মামী (Aunt of the youngest child in the household)
8) অন্যান্য (Other)_________________

#### কীটনাশক বিষয়ক প্রশ্নাবলী (Pesticide Questions)

24. গত বছর আপনার জমিতে কতবার কীটনাশক প্রয়োগ করেছেন? (How many times did you apply pesticide on your land in the last year?) ____
25. আমি আপনাকে প্রচলিত কীটনাশকের একটি তালিকা পড়ে শোনাব। দয়া করে আপনি আমাকে বলবেন, এসব কীটনাশকের মধ্যে কোন কোন কীটনাশক গত বছর আপনি কতবার আপনার জমিতে ধান উৎপাদনের জন্য প্রয়োগ/ব্যবহার করেছেন? (I will read a list of available pesticides. Please tell me how many times you applied each of these types of pesticide on the land you used for growing rice in the last year.
1) টাইপ A[Type A (Type names will be identified in the qualitative study)]
2) টাইপ B(Type B)
3) টাইপ C(Type C)
4) টাইপ D(Type D)
5) টাইপ E(Type E)
6) অন্যান্য (Other) ________

i. আমি আপনাকে পুনরায় প্রচলিত কীটনাশকের তালিকাটি পড়ে শোনাব। দয়া করে আপনি আমাকে বলবেন, এসব কীটনাশকের মধ্যে কোন কোন কীটনাশক গত বছর আপনি কতবার আপনার জমিতে অন্যান্য শস্য (ধান ছাড়া) উৎপাদনের জন্য প্রয়োগ/ব্যবহার করেছেন? (I will again read the list of available pesticides. Please tell me how many times you applied each of these types of pesticide on the land you used for growing other crops in the last year.)
1) টাইপ A[Type A (Type names will be identified in the qualitative study)]
2) টাইপ B(Type B)
3) টাইপ C(Type C)

4) টাইপ D(Type D)
5) টাইপ E(Type E)
6) অন্যান্য (Other) ________

ii. কীটনাশকগুলো জমিতে প্রয়োগ করার পূর্বে কে এগুলো প্রস্তুত করেন/নাড়াচাড়া করেন? (প্রত্যেকটি অপশনের ক্ষেত্রে আলাদাভাবে উত্তর দিন । হ্যাঁ=1, না=0) (Who handled the pesticide while preparing to apply it? (answer each separately Yes = 1, No = 0)

1) সবচেয়ে ছোট শিশুর বাবা (Father of the youngest child in the household)
2) সবচেয়ে ছোট শিশুর মা (Mother of the youngest child in the household)
3) সবচেয়ে ছোট শিশুর সহোদর (Sibling of the youngest child in the household)
4) সবচেয়ে ছোট শিশুর দাদা-দাদী (Grandparent of the youngest child in the household)
5) সবচেয়ে ছোট শিশুর চাচাতো ভাই-বোন (Cousin of the youngest child in the household)
6) সবচেয়ে ছোট শিশুর চাচা/মামা (Uncle of the youngest child in the household)
7) সবচেয়ে ছোট শিশুর চাচী/মামী (Aunt of the youngest child in the household)
8) অন্যান্য (Other)_________________

36) কীটনাশকগুলো প্রয়োগ করার/ছিটানোর জন্য কোথায় প্রস্তুত করা হয়েছিল? (Where was the pesticide prepared for application)

1) বাড়িতে (In the house)
2) বাড়ির কাছে (Near the house)
3) জমিতে (In the field)
4) অন্যান্য (Other)_________________________

37) জমিতে ব্যবহারের পর অবশিষ্ট কীটনাশকগুলো আপনি কি করেছিলেন? (What did you do with the leftover pesticide?

1) অবশিষ্ট ছিল না (Did not have any)
2) বাড়িতে সংরক্ষন করেছিলাম (Stored it in the house)
3) কাউকে দান করে দিয়েছিলাম বা বিক্রি করে দিয়েছিলাম (Gave it away or sold it)
4) অন্যান্য (Other)

38) জমিতে ব্যবহারের পর কীটনাশকের অবশিষ্ট প্যাকেটগুলো আপনি কি করেছিলেন? (প্রত্যেকটি অপশনের ক্ষেত্রে আলাদাভাবে উত্তর দিন । হ্যাঁ=1, না=0) What did you do with the leftover pesticide packaging? (answer each separately Yes = 1, No = 0)

1) বাড়িতে সংরক্ষন করেছিলাম (Stored it at home)
2) খাদ্য মজুদ রাখার কাজে ব্যবহার করেছিলাম (Used it to store food)
3) পানি সংরক্ষনের কাজে ব্যবহার করেছিলাম (Used it to store water)
4) গৃহস্থালী সামগ্রী সংরক্ষনের কাজে ব্যবহার করেছিলাম (Used it to store household goods)
5) শিশুদের খেলতে দিয়েছিলাম (Let the children play with it)
6) ফেলে দিয়েছিলাম (Discarded it)

39) কে কীটনাশকগুলো জমিতে দিয়েছিল? (প্রত্যেকটি অপশনের ক্ষেত্রে আলাদাভাবে উত্তর দিন । হ্যাঁ=1, না=0) (Who applied the pesticide to the land? (answer each separately Yes = 1, No = 0)

1) সবচেয়ে ছোট শিশুর বাবা (Father of the youngest child in the household)
2) সবচেয়ে ছোট শিশুর মা (Mother of the youngest child in the household)
3) সবচেয়ে ছোট শিশুর সহোদর (Sibling of the youngest child in the household)
4) সবচেয়ে ছোট শিশুর দাদা-দাদী (Grandparent of the youngest child in the household)
5) সবচেয়ে ছোট শিশুর চাচাতো ভাই-বোন (Cousin of the youngest child in the household)
6) সবচেয়ে ছোট শিশুর চাচা/মামা (Uncle of the youngest child in the household)
7) সবচেয়ে ছোট শিশুর চাচী/মামী (Aunt of the youngest child in the household)

8) অন্যান্য (Other)________________

আগাছানাশক সংক্রান্ত প্রশ্নাবলী (Herbicide Questions)

40) গত বছর আপনার জমিতে কতবার আগাছানাশক প্রয়োগ করেছেন? (How many times did you apply herbicide on your land in the last year?) ____

41) আমি আপনাকে প্রচলিত আগাছানাশকে র একটি তালিকা পড়ে শোনাব। দয়া করে আপনি আমাকে বলবেন, এসব আগাছানাশকে র মধ্যে কোন কোন আগাছানাশক গত বছর আপনি কতবার আপনার জমিতে ধান উৎপাদনের জন্য প্রয়োগ/ব্যবহার করেছেন? [I will read a list of available herbicides. Please tell me how many times you applied each of these types of herbicide on the land you used for growing rice in the last year.]
   1) টাইপ A[Type A (Type names will be identified in the qualitative study)]
   2) টাইপ B(Type B)
   3) টাইপ C(Type C)
   4) টাইপ D(Type D)
   5) টাইপ E(Type E)
   6) অন্যান্য (Other) _______

42) আমি আপনাকে পুনরায় প্রচলিত আগাছানাশকে র তালিকাটি পড়ে শোনাব। দয়া করে আপনি আমাকে বলবেন, এসব আগাছানাশকে র মধ্যে কোন কোন আগাছানাশক গত বছর আপনি কতবার আপনার জমিতে অন্যান্য শস্য (ধান ছাড়া) উৎপাদনের জন্য প্রয়োগ/ব্যবহার করেছেন? (I will again read the list of available herbicides. Please tell me how many times you applied each of these types of herbicide on the land you used for growing other crops in the last year.)
   1) টাইপ A[Type A (Type names will be identified in the qualitative study)]
   2) টাইপ B(Type B)
   3) টাইপ C(Type C)
   4) টাইপ D(Type D)
   5) টাইপ E(Type E)
   6) অন্যান্য (Other) _______

43) আগাছানাশক গুলো জমিতে প্রয়োগ করার পূর্বে কে এগুলো প্রস্তুত করেন/নাড়াচাড়া করেন? (প্রত্যেকটি অপশনের ক্ষেত্রে আলাদাভাবে উত্তর দিন । হ্যাঁ=1, না=0) [Who handled the herbicide while preparing to apply it? (answer each separately Yes = 1, No = 0)]
   1) সবচেয়ে ছোট শিশুর বাবা (Father of the youngest child in the household)
   2) সবচেয়ে ছোট শিশুর মা (Mother of the youngest child in the household)
   3) সবচেয়ে ছোট শিশুর সহোদর (Sibling of the youngest child in the household)
   4) সবচেয়ে ছোট শিশুর দাদা-দাদী (Grandparent of the youngest child in the household)
   5) সবচেয়ে ছোট শিশুর চাচাতো ভাই-বোন (Cousin of the youngest child in the household)
   6) সবচেয়ে ছোট শিশুর চাচা/মামা (Uncle of the youngest child in the household)
   7) সবচেয়ে ছোট শিশুর চাচী/মামী (Aunt of the youngest child in the household)
   8) অন্যান্য (Other)________________

44) আগাছানাশক গুলো প্রয়োগ করার/ছিটানোর জন্য কোথায় প্রস্তুত করা হয়েছিল? [Where was the herbicide prepared for application]
   1) বাড়িতে (In the house)
   2) বাড়ির কাছে (Near the house)

3) জমিতে (In the field)
4) অন্যান্য (Other)_________________________

45) জমিতে ব্যবহারের পর অবশিষ্ট আগাছানাশক গুলো আপনি কি করেছিলেন? [What did you do with the leftover herbicide? ]
1) অবশিষ্ট ছিল না (Did not have any)
2) বাড়িতে সংরক্ষন করেছিলাম (Stored it in the house)
3) কাউকে দান করে দিয়েছিলাম বা বিক্রি করে দিয়েছিলাম (Gave it away or sold it)
4) অন্যান্য (Other)

46) জমিতে ব্যবহারের পর আগাছানাশকে র অবশিষ্ট প্যাকেটগুলো আপনি কি করেছিলেন? (প্রত্যেকটি অপশনের ক্ষেত্রে আলাদাভাবে উত্তর দিন । হ্যাঁ=1, না=0) [What did you do with the leftover herbicide packaging? (answer each separately Yes = 1, No = 0)]
1) বাড়িতে সংরক্ষন করেছিলাম (Stored it at home)
2) খাদ্য মজুদ রাখার কাজে ব্যবহার করেছিলাম (Used it to store food)
3) পানি সংরক্ষনের কাজে ব্যবহার করেছিলাম (Used it to store water)
4) গৃহস্থালী সামগ্রী সংরক্ষনের কাজে ব্যবহার করেছিলাম (Used it to store household goods)
5) শিশুদের খেলতে দিয়েছিলাম (Let the children play with it)
6) ফেলে দিয়েছিলাম (Discarded it)

47) কে আগাছানাশক গুলো জমিতে দিয়েছিল? (প্রত্যেকটি অপশনের ক্ষেত্রে আলাদাভাবে উত্তর দিন । হ্যাঁ=1, না=0) [Who applied the herbicide to the land? (answer each separately Yes = 1, No = 0)]
1) সবচেয়ে ছোট শিশুর বাবা (Father of the youngest child in the household)
2) সবচেয়ে ছোট শিশুর মা (Mother of the youngest child in the household)
3) সবচেয়ে ছোট শিশুর সহোদর (Sibling of the youngest child in the household)
4) সবচেয়ে ছোট শিশুর দাদা-দাদী (Grandparent of the youngest child in the household)
5) সবচেয়ে ছোট শিশুর চাচাতো ভাই-বোন (Cousin of the youngest child in the household)
6) সবচেয়ে ছোট শিশুর চাচা/মামা (Uncle of the youngest child in the household)
7) সবচেয়ে ছোট শিশুর চাচী/মামী (Aunt of the youngest child in the household)
8) অন্যান্য (Other)_________________

ছত্রাকনাশক ঔষধ সংক্রান্ত প্রশ্নাবলী (Fungicide Questions)

48) গত বছর কতবার আপনি আপনার জমিতে ছত্রাকনাশক ঔষধ প্রয়োগ করেছেন? (How many times did you apply fungicide on your land in the last year?) ____
49) আমি আপনাকে প্রচলিত ছত্রাকনাশক ঔষধের একটি তালিকা পড়ে শোনাব। দয়া করে আপনি আমাকে বলবেন, এসব (ছত্রাকনাশক) ঔষধের মধ্যে কোন কোন ধরনের ঔষধ গত বছর আপনি কতবার আপনার জমিতে ধান উৎপাদনের জন্য প্রয়োগ/ব্যবহার করেছেন? (I will read a list of available fungicides. Please tell me how many times you applied each of these types of fungicide on the land you used for growing rice in the last year.)
1) টাইপ A [Type A (Type names will be identified in the qualitative study)]
2) টাইপ B (Type B)
3) টাইপ C (Type C)
4) টাইপ D (Type D)
5) টাইপ E (Type E)

6) অন্যান্য (Other) _______

50) আমি আপনাকে পুনরায় প্রচলিত ছত্রাকনাশক ঔষধের তালিকাটি পড়ে শোনাব। দয়া করে আপনি আমাকে বলবেন, এসব (ছত্রাকনাশক) ঔষধের মধ্যে কোন ধরনের ঔষধ গত বছর আপনি কতবার আপনার জমিতে অন্যান্য শস্য (ধান ছাড়া) উৎপাদনের জন্য প্রয়োগ/ব্যবহার করেছেন? I will again read the list of available fungicides. Please tell me how many times you applied each of these types of fungicide on the land you used for growing other crops in the last year.

1) টাইপ A[Type A (Type names will be identified in the qualitative study)]
2) টাইপ B(Type B)
3) টাইপ C(Type C)
4) টাইপ D(Type D)
5) টাইপ E(Type E)
6) অন্যান্য (Other) _______

51) ছত্রাকনাশক ঔষধগুলো জমিতে প্রয়োগ করার পূর্বে কে এগুলো প্রস্তুত করেন/নাড়াচাড়া করেন? (প্রত্যেকটি অপশনের ক্ষেত্রে আলাদাভাবে উত্তর দিন । হ্যাঁ=1, না=0) [Who handled the fungicide while preparing to apply it? (answer each separately Yes = 1, No = 0)]

1) সবচেয়ে ছোট শিশুর বাবা (Father of the youngest child in the household)
2) সবচেয়ে ছোট শিশুর মা (Mother of the youngest child in the household)
3) সবচেয়ে ছোট শিশুর সহোদর (Sibling of the youngest child in the household)
4) সবচেয়ে ছোট শিশুর দাদা-দাদী (Grandparent of the youngest child in the household)
5) সবচেয়ে ছোট শিশুর চাচাতো ভাই-বোন (Cousin of the youngest child in the household)
6) সবচেয়ে ছোট শিশুর চাচা/মামা (Uncle of the youngest child in the household)
7) সবচেয়ে ছোট শিশুর চাচী/মামী (Aunt of the youngest child in the household)
8) অন্যান্য (Other)________________

52) ছত্রাকনাশক ঔষধগুলো প্রয়োগ করার/ছিটানোর জন্য কোথায় প্রস্তুত করা হয়েছিল? [Where was the fungicide prepared for application]

1) বাড়িতে (In the house)
2) বাড়ির কাছে (Near the house)
3) জমিতে (In the field)
4) অন্যান্য (Other)________________________

53) জমিতে ব্যবহারের পর অবশিষ্ট ছত্রাকনাশক ঔষধগুলো আপনি কি করেছিলেন? [What did you do with the leftover fungicide? ]

1) অবশিষ্ট ছিল না (Did not have any)
2) বাড়িতে সংরক্ষন করেছিলাম (Stored it in the house)
3) কাউকে দান করে দিয়েছিলাম বা বিক্রি করে দিয়েছিলাম (Gave it away or sold it)
4) অন্যান্য (Other)

54) জমিতে ব্যবহারের পর ছত্রাকনাশক ঔষধের অবশিষ্ট প্যাকেটগুলো আপনি কি করেছিলেন? (প্রত্যেকটি অপশনের ক্ষেত্রে আলাদাভাবে উত্তর দিন । হ্যাঁ=1, না=0) [What did you do with the leftover fungicide packaging? (answer each separately Yes = 1, No = 0)]

1) বাড়িতে সংরক্ষন করেছিলাম (Stored it at home)
2) খাদ্য মজুদ রাখার কাজে ব্যবহার করেছিলাম (Used it to store food)
3) পানি সংরক্ষনের কাজে ব্যবহার করেছিলাম (Used it to store water)
4) গৃহস্থালী সামগ্রী সংরক্ষনের কাজে ব্যবহার করেছিলাম (Used it to store household goods)
5) শিশুদের খেলতে দিয়েছিলাম (Let the children play with it)

6) ফেলে দিয়েছিলাম (Discarded it)

55) কে ছত্রাকনাশক ঔষধগুলো জমিতে দিয়েছিল? (প্রত্যেকটি অপশনের ক্ষেত্রে আলাদাভাবে উত্তর দিন । হ্যাঁ=1, না=0) [Who applied the fungicide to the land? (answer each separately Yes = 1, No = 0)]
1) সবচেয়ে ছোট শিশুর বাবা (Father of the youngest child in the household)
2) সবচেয়ে ছোট শিশুর মা (Mother of the youngest child in the household)
3) সবচেয়ে ছোট শিশুর সহোদর (Sibling of the youngest child in the household)
4) সবচেয়ে ছোট শিশুর দাদা-দাদী (Grandparent of the youngest child in the household)
5) সবচেয়ে ছোট শিশুর চাচাতো ভাই-বোন (Cousin of the youngest child in the household)
6) সবচেয়ে ছোট শিশুর চাচা/মামা (Uncle of the youngest child in the household)
7) সবচেয়ে ছোট শিশুর চাচী/মামী (Aunt of the youngest child in the household)
8) অন্যান্য (Other)_________________

#### WASH Benefits Module 53
#### In-Depth Interview Guidelines

**In-depth environmental assessment (assessment of exposure pathways to fecal contamination, association with diarrhea and sanitation coverage in Bangladesh)**

**Drinking water sources, handling and storage practices**

- Can you tell me about how you collect and store drinking water?
- What is your primary/main source of drinking water? (probe- type, tube well, pond and others, their conditions, secondary and other sources, seasonal impacts. Emphasize tube well and pond water). Observe the source and describe objectively.
- Who from your household is responsible to collect drinking water? (probe- primary and secondary collectors, when and why?)
- How is the collected water stored and for how long? Note the common storage containers. Give me an example of typical collection and storage of water and usage by the household members.
- How can these water sources or stored water be contaminated by human and/or animal feces? What are the possible pathways that could contaminate the water by humanfeces including children feces and/or animal feces(draw a map of assumed paths of fecal flow)?
- How do young children in your household get water for drinking?
- What, if any, are the difficulties associated with collecting and storing safe water?

**Availability, treating, scarcityand consumption of water**

- Can you tell me if your household ever experiences drinking water scarcity? What do you do in those times?
- Do members of your household consume water differently? How?
- Water treating or purifying behaviors:Do you do anything to your drinking water to make it safer to drink? What are the common water treatment methods in your household?
- Tell me about typical consumption of water by each of your household members including children by age group?
- Tell me yesterday's practices and consumption (note date, temperature, season etc.)

**Contamination of hands and handwashing practices**

- How frequently and when do you wash your hands? probe- key times (before having different types of food, after using toilet and cleaning child's anus). Illustrate a typical day's practices (may be yesterday or today).

- Do you think either/both of your hands could be contaminated by human feces? Probe- handwashing during these key times(draw a map from which activities fecal contamination may occur to hands)
- What agent do you use to wash your hands? Probe- why and when with what?
- How often do you clean your young children's hands? When and how?
- How often do your young children wash their own hands? When and how?
- Do you think your children's hands could be contaminated by human feces?
- Do you find handwashing with soap difficult or easy? Why?

**Contamination of soil**

- Does your household do anything to keep your courtyard clean? Probe – what is done, how often?
- How could courtyard soil be contaminated by human feces? Probe- open defecation by children and adults, during day/night or during rain. Locate spots and describe.
- Where do young children in your compound defecate? When and why? Who disposes of those feces, how and where? Is there any variation in these practices? What are the differences and why? Observe feces disposal sites. Probe- each sites.
- Is there any child potty available in households in this household?
- Tell me about how often your children come into contact with soil and feces?
  - Child and animal feces
  - Type of animal, children from neighboring households
- How frequently do your children (by age group) spend time in courtyard or verandah and for how long? Who else accompanies them? Do children play alone or in groups? Observe- floor materials (concrete or mud), is there any child potty in the household available?
- How much time do your children spend on thecourtyard ground and why? Who watches over children's behavior when they play in courtyard? Probe- absence and present of primary and secondary caregivers.
- What do the caregivers do when a child puts something from courtyard ground/soil into their mouth?
- What do the caregivers think when a child puts something from soil into their mouth?

**Contamination of food**

- Please tell me about food preparation and preservation processes followed by your household? What is your role in preparing and storing daily food? Probe- main food and snacks, occasional/seasonal foods.
- How do you store those foods and for how long? Probe- when and why?
- Who is responsible for maintaining food storage and hygiene in your household? Probe- social and economic power structure in the household (mother/father-in-laws, husbands).
- How do prepared and stored food gets into contact with hands? What do you do to keep your food safe? Probe- hand hygiene before preparing food, heating and reheating before serving food to child, child consumption of food and feeding methods. Who feeds, when and how?

**Free roaming domestic animals**

- How many domestic animals do you have? Make a count. Identify the majority.
- Who takes care of animals and animal sheds?
- When and why are animal feces visible in compound/inside household and for how long?
- How can a child come into contact with animals?
- Is there any chance that a child can put animal feces from the courtyard ground into their mouth? Probe- which animals, when and where?
- What do parents do if a child puts animal feces in their mouth? Please describe.

**Contamination via flies**

- What do you think about the presence of different flies in your house and surrounding environment? Ask specifically about type (size, color, other characteristics).

- Why are these flies found around/inside the household? Where do they come from? When is the fly density high and why?
- Is there a chance that these flies can land on food? How frequently? Probe- time and season.
- Do you feel that flies pose any health risks to your household? Why or why not?

**Availability and type of toilet facility**

- Please tell me about your household’s sanitation practices. Where do the household members usually defecate? Probe- practice open defecation, individual or shared latrine.
- How many latrines do you have? Probe- type and other facilities available for each toilet.Map the compound marking toilets and distances from drinking water source, pond or other water sources and from living room, kitchen etc.
- Who is responsible for cleaning and maintaining the toilet? Probe- man, women, specific roles, frequency.
- What are the hardware available in this household to dispose of feces from courtyard or inside the house? Probe- child and animal feces, adult human feces, feral animal feces.
- What happens during different seasons of the year? Probe- when latrine pit is full, feces float over pan, during rainy season, when latrine is broken.
- Where do the feces from the latrinein your household end up? In ditches, ponds, drains, small rivers, other water bodies.
- What do you find difficult about managing sanitation in your household? Please describe.
- What is easy about managing sanitation in your household? Please describe.

#### WASH Benefits Module 53: In-Depth Interview Guidelines in Bengali for Environmental Assessment

**পরিবেশগত নিবিড় মূল্যায়ন (বাংলাদেশে ডায়রিয়া এবং স্যানিটেশন সুবিধার অন্তর্ভূক্তি, মলদুষণের গমনাগমনের মূল্যায়ন)**

**পানীয় জলের উৎসসমূহ, পরিচালনা এবং সংরৰণ চর্চাসমূহ**

- আপনি কি বলতে পারেন কিভাবে পানি সংগ্রহ ও সংরৰণ করেন?
- আপনার প্রাথমিক/প্রধান পান করার পানির উৎস কী কী? (জিজ্ঞাসা করম্নন- ধরণ, টিউবওয়েল, পুকুর বা অন্যান্য, সেগুলির ভৌত অবস্থা, দ্বিতীয় ও অন্যান্য উৎসসমূহ, মৌসুমী প্রভাবসমূহ। টিউবওয়েল ও পুকুরের পানির উপর গুরম্নত্ব বেশী দিন)। এই উৎসগুলি পর্যবেৰণ করম্নন এবং নৈর্ব্যক্তিকভাবে বর্ণনা করম্নন।আপনার গৃহস্থালীতে পান করার পানি সংগ্রহ করা কার দায়িত্ব? (জিজ্ঞাসা করম্নন- প্রাথমিক ও দ্বিতীয় সংগ্রহকারী, কখন ও কেন?)
- সংগৃহীত পানি কিভাবে সংরৰণ করা হয় এবং কত সময়ের জন্য? সাধারণভাবে ব্যবহার করা হয় এমন সংরৰণ পাত্রগুলি লিপিবদ্ধ করম্নন। আপনার পরিবারের সদস্যগণ কর্তৃক সাধারণ প্রাত্যহিক একটি দিনে পানি সংগ্রহ, সংরৰণ ও ব্যবহারের উদাহরণ দিন।
- এই পানির উৎসগুলি বা সংরৰণকৃত পানি কিভাবে মানুষ বা পশু-পাখির মল দ্বারা দুষিত হতে পারে? এই সম্ভাব্য দুষণের গমনাগমন পথগুলি কী কী যা প্রাপ্ত বয়স্ক মানুষ, শিশু বা পশু-পাখির মলের দ্বারা দুষিত হয়?
- আপনার পরিবারের শিশুরা কিভাবে খাবার পানি পায়?
- যদি থেকে থাকে তাহলে নিরাপদ পানি সংগ্রহ এবং সংরৰণের সাথে সংশিস্নষ্ট অসুবিধাগুলি কী কী?

**সহজপ্রাপ্যতা, পানি নিরাপদকরণ, পানি স্বল্পতা এবং পানির ব্যবহার**

- আমাকে কি বলতে পারেন আপনার পরিবার কখনো পানির স্বল্পতার মুখোমুখি হয়েছেন কি না? ঐ সময়গুলিতে আপনি কী করেছেন?
- আপনার পরিবারের সদস্যদের পানির ব্যবহারের পরিমাণে ভিন্নতা রয়েছে কি? কিভাবে ও কেন?
- পানি নিরাপদ বা বিশুদ্ধকরণ আচরণঃ আপনি কি আপনার পানি পানের জন্য নিরাপদ করতে কোন কিছু করেন? আপনার গৃহস্থালীতে পানি বিশুদ্ধকরণের সাধারণ পদ্ধতিগুলি কী কী?
- শিশুদের বয়সভেদসহ আপনার পরিবারের সকল সদস্যদের সাধারণভাবে পানি ব্যবহার সম্পর্কে বলুন।
- গতকালকের পানি ব্যবহার ও এ সংক্রান্ত চর্চা সম্পর্কে বলুন (তারিখ, ঐদিনের তাপমাত্রা, তখনকার সিজন ইত্যাদি লিপিবদ্ধ করম্নন)

**হাতে জীবানুর সংক্রমণ এবং হাত ধোয়ার চর্চা**

- কত ঘনঘন এবং কিভাবে আপনি হাত ধোন? জিজ্ঞাসা করম্নন- প্রধান প্রধান সময়গুলি (বিভিন্ন ধরণের খাবার খাওয়ার আগে, টয়লেট ব্যবহারের পর এবং বাচ্চাকে শৌচানোর পর)। কোন একটি সাধারণ দিনের হাত ধোয়ার চর্চাগুলি বর্ণনা করম্নন (হতে পারে গতকাল বা আজ)।

- আপনি কি মনে করেন যে কোন একটি বা উভয় হাতই মানুষের মল দ্বারা দুষিত হতে পারে? জিজ্ঞাসা করুন- ঐ প্রধান প্রধান সময়গুলিতে হাত ধোয়া (একটি চিত্র অংকন করুন যেসব কর্মকান্ড থেকে হাতে মলের সংক্রমন ঘটতে পারে)।
- হাত ধোয়ার জন্য আপনি কী কী ব্যবহার করেন? জিজ্ঞাসা করুন- কেন এবং কখন কখন কী কী দিয়ে?
- কতবার আপনি আপনার শিশুর হাত পরিস্কার করেন/ধোন? কখন কখন এবং কিভাবে?
- কতবার আপনার শিশুরা তাদের নিজেদের হাত ধোয়? কখন কখন এবং কিভাবে?
- আপনি কি মনে করেন আপনার শিশুর হাত মানুষের মল দ্বারা দুষিত হতে পারে?
- সাবান দিয়ে হাত ধোয়া আপনার কাছে সহজ না কঠিন মনে হয়? কেন?

**মাটি দূষণ**

- আপনার উঠানটিকে পরিস্কার রাখার জন্য আপনার পরিবার কী কী করে? জিজ্ঞাসা করুন- কী কী করা হয়, কিভাবে এবং কত সময় পর পর?
- উঠান কিভাবে মানুষের মল দ্বারা দুষিত হতে পারে? জিজ্ঞাসা করুন- শিশু এবং বয়স্কগণ, দিনে বা রাতে বা বৃষ্টির সময় খোলা জায়গায় মলত্যাগ করা। স্থানগুলি চিহ্নিত করুন এবং বর্ণনা করুন।
- আপনার বাড়ীতে ছোট ছোট শিশুরা কোথায় পায়খানা করে? কখন কখন এবং কোথায়? ছোট শিশুদের মল কে কোথায় এবং কিভাবে অপসারণ করে? এই চর্চাগুলিতে কি কোন পার্থক্য হয়? পার্থক্যগুলি কী কী এবং কেন হয়? যেখানে যেখানে মল ফেলা হয় সেই স্থানগুলি পর্যবেক্ষণ করুন এবং প্রত্যেক স্থানগুলি নিয়ে আরও প্রশ্ন জিজ্ঞাসা করুন।
- এই পরিবারে কি কোন শিশুদের মলত্যাগ করার জন্য পটি রয়েছে?
- আমাকে বলুন কত বার আপনার শিশুটি মাটি এবং মলের সংস্পর্শে আসে?
  - শিশুদের এবং পশু-পাখির মল
  - মলের ধরণ, পার্শ্ববর্তী বাড়ীর শিশুদের মল
- কত সময় পর পর আপনার শিশুরা (বয়সদল ভেদে) উঠান বা বারান্দায় সময় কাটায় এবং কত সময় ধরে? তাদেরকে কে সঙ্গ দেয়? শিশুরা কি একা না অনেকে মিলে খেলাধুলা করে? পর্যবেক্ষণ করুন- ঘরের বা বারান্দার মেঝের ধরণ (পাকা বা মাটি), ঘরে শিশুদের মলত্যাগ করার জন্য কোন ধরণের পটি আছে কি না।
- আপনার শিশু কত সময় উঠানে সময় কাটায় এবং কেন? কে তাদের আচার-আচরণ লক্ষ্য করে যখন তারা উঠানের মাটিতে খেলাধুলা করে? জিজ্ঞাসা করুন- মায়ের বা অন্য যত্নগ্রহণকারীর উপস্থিতি এবং অনুপস্থিতিতে।
- যখন শিশুরা উঠানের মাটি থেকে তাদের মুখে কোন কিছু দেয় তখন মায়েরা কী করেন?
- বাচ্চারা যখন মাটি থেকে কোন কিছু মুখে দেয় তখন মায়েরা কী ভাবেন বা মনে করেন?

**খাদ্য দূষণ**

- দয়া করে আমাকে আপনাদের পরিবারের খাবার তৈরী এবং সংরক্ষণ প্রক্রিয়া সম্পর্কে বলুন। প্রতিদিনকার খাবার তৈরী এবং রক্ষণাবেক্ষণে আপনার ভূমিকা কী? জিজ্ঞাসা করুন- প্রধান প্রধান খাবার, নাস্তা, মৌসুমি খাবার।
- এই খাবারগুলি কিভাবে ও কতক্ষণের জন্য সংরক্ষণ করেন? জিজ্ঞাসা করুন- কখন কখন ও কেন?
- আপনার পরিবারে খাবার সংরক্ষণ ও তা স্বাস্থ্যসম্মত রাখার দায়িত্ব কার? জিজ্ঞাসা করুন- পরিবারের সামাজিক ও অর্থনৈতিক ক্ষমতা (শ্বশুর-শ্বাশুড়ী, স্বামী)
- তৈরীকৃত ও সংরক্ষিত খাবারগুলি কিভাবে হাতের সংস্পর্শে আসে? আপনার পরিবারের জন্য খাবার নিরাপদ রাখার জন্য কী কী করেন? জিজ্ঞাসা করুন- খাবার তৈরীর পূর্বে হাত ধোয়া, বাচ্চাকে খাওয়ানোর আগে বার বার খাবার গরম করা, শিশুদের খাবারের পরিমাণ এবং খাওয়ানোর পদ্ধতি। কে খাওয়ায়, কখন কখন ও কেন?

**উন্মুক্তভাবে থাকা গৃহপালিত পশু-প্রাণীসমূহ**

- আপনার কতগুলি গৃহপালিত পশু রয়েছে? সংখ্যা গণনা করুন, সবচেয়ে বেশী কোনগুলি তা চিহ্নিত করুন।
- পশু-পাখী ও খোয়ার বা গোয়ালঘর কে দেখাশোনা করে
- কখন কখন এবং কেন পশু-পাখীর মল আপনার বাড়ীর উঠানে ও আশে পাশে দেখা যায় এবং কত সময় ধরে?
- একটি বাচ্চা শিশু কিভাবে পশু-পাখীর সংস্পর্শে আসতে পারে?
- এমন কি কোন সম্ভাবনা আছে যে শিশুরা উঠানের মাটি থেকে পশু-পাখির কোন মল মুখে দিতে পারে?
- বাচ্চারা যদি পশু-পাখির মল মুখে দেয় তবে মা-বাবারা কী করেন? বর্ণনা করুন।

**মশা-মাছির মাধ্যমে দূষণ/সংক্রমণ**

- আপনার বাড়ীর ভেতর এবং আশে পাশে বিভিন্ন ধরণের মাছির উপস্থিতি সম্পর্কে আপনার কী মনে হয়? সুনির্দিষ্টভাবে ধরণগুলি সম্পর্কে জিজ্ঞাসা করুন (আকৃতি, রং, অন্যান্য বৈশিষ্ট্যসমূহ)
- এই ধরণের মশা-মাছিগুলি আপনার ঘরের ভেতর বা আশে পাশে কেন পাওয়া যায়? এগুলি কোথা থেকে আসে? কখন কখন মশা-মাছির উপদ্রব বেশী হয় ও কেন?

- এমনি কি কোন সুযোগ আছে যে মশা-মাছি খাবারের উপর বসে? কত ঘন ঘন এমন হয়? জিজ্ঞাসা করম্নন- সময় ও ঋতু।
- আপনি কি মনে করেন যে মাছিরা আপনার পরিবারের জন্য কোন স্বাস্থ্য ঝুকি বয়ে আনে? কেন অথবা কেন নয়?

**টয়লেট সুবিধাদির ধরণ ও সহজলভ্যতা**

- আপনার পরিবারের স্বাস্থ্যসম্মত চর্চা সম্পর্কে বলুন। তারা কোথায় সচরাচর পায়খানা/মলত্যাগ করেন? জিজ্ঞাসা করম্নন- খোলা জায়গায়, ব্যক্তিগত/নিজস্ব বা অন্যের ল্যাট্রিনে।
- আপনার কয়টি ল্যাট্রিন আছে? জিজ্ঞাস করম্নন- প্রতিটি ল্যাট্রিনের ধরন ও অন্যান্য সুবিধাসমূহ। এই বাড়ীটির একটি চিত্র অংকন করম্নন যেখানে ল্যাট্রিন/টয়লেট সুবিধাদি এবং খাবার পানির উৎস, পুকুর বা অন্যান্য উৎস, থাকার ঘর ও রান্না ঘর থেকে তার দূরত্ব চিহ্নিত করম্নন।
- টয়লেট ব্যবস্থাপনা ও পরিস্কার রাখার দায়িত্ব কার? জিজ্ঞাসা করম্নন- পুরম্নষ, নারী, নির্দিষ্ট ভূমিকা, কত সময় পর পর?
- উঠান থেকে অপসারণ করার জন্য এই পরিবারের কোন কোন যন্ত্র-পাতি রয়েছে। জিজ্ঞসা করম্নন- শিশু ও পশুর মল, বয়স্ক দের ও অপোষা প্রাণীদের।
- বছরের বিভিন্ন মৌসুমে কী হয়? জিজ্ঞাস করম্নন- ল্যাট্রিন যখন ভরে যায়, প্যানের উপর মল ভাসতে থাকে, বা ল্যাট্রিন যখন ভেঙ্গে যায়।
- ল্যাট্রিন থেকে মল কোথায় গিয়ে শেষ পৌছে? নালায়, পুকুরে, ডোবায়, ছোট নদী, গাঙ্গে, বা অন্যান্য কোন পানিতে।
- আপনার পরিবারের সদস্যদের মলত্যাগ ও পরিবেশ মলমুক্ত রাখার জন্য কী কী অসুবিধার মুখোমুখি হন বর্ণনা করম্নন।
- এ ধরণের স্যানিটেশন কাজে কোন বিষয়গুলি সহজ বলে মনে হয় ও কেন? বর্ণনা করম্নন।

#### WASH Benefits Module 54
#### Focus Group Discussion Guidelines

**In-depth environmental assessment (assessment of exposure pathways to fecal contamination, association with diarrhea and sanitation coverage in Bangladesh)**

**Drinking water sources, handling and storage practices**

- What are the primary/main sources of drinking water in this area? Probe- type, tube well, pond and others, their conditions, secondary and other sources, seasonal impacts. Emphasize tube-well and pond water. Observe the source and describe objectively.
- What are the common drinking water collection and storage methods? Rank these methods and probe all methods for clarification and understanding.
- Who from the households is usually responsible to collect and store drinking water? (probe- primary and secondary collectors, when and why?
- How can these water sources or stored water be contaminated by human and/or animal feces? What are the possible pathways that could contaminate the water by human feces including children feces and/or animal feces?(draw a map of assumed paths of fecal flow, check with the maps drawn from in-depth interviewsforupdate/modification)

**Availability, treating, scarcityand consumption of water**

- When is there drinking water scarcity in this area?
- What do the household members do at that time?
- How much difference is there in collecting, storing and consuming water?
- What are the common or popular water treating or purifying methods in this area? Describe to me the most popular or common method.
- Are there certain times of the year when treating or purifying water is more or less common?
- Tell me about typical consumption of water by household members. Probe- men, women, children by age group?

**Contamination of hands and handwashing practices**

- How frequently and when do people wash their hands? Probe- three key times (before having different types of food, after using toilet and cleaning child's anus). Describe a typical person's handwashing practices (could be a caregiver/homemaker).
- Do people think that either/both of their hands could be contaminated by human feces? Probe- handwashing during these key times. (draw a map from which activities fecal contamination may occur to hands)
- Do people wash the left or right hands more often? When and why?
- What agentsare available in this area that people use to wash their hands? Probe- all agents, rank on availability and frequency of use, why and when with what?
- How frequently do young children have their hands washed by their caregivers?
- How often do young children wash their own hands?
- Do people think that young children's hands could be contaminated by human and/or animal feces? Who from the household can encourage or influence handwashing? Probe- Grandfather, father-in-law, husband, school going children?

**Contamination of courtyard, soil of surrounding household environment**

- How could courtyard soil become contaminated by human feces? Probe- open defecation by children and adults, elderly, sick, during day/night or during rain, locatespots and describe. Ask for an example that they can explain. Draw a map with sources of feces and contamination.
- Where do young children defecate? When and why? Who disposes of those feces, how and where?Probe- common, fixed, temporary sites.
- Is there any variation in feces disposal practices? What are the differences and why? Observe feces disposal sites. Probe- each site.
- Are there child potties available in households in this community?
- In which place do young children spend most of their time other than lap of adults? Describe the places
- How frequently dochildren (by age group) spend time in courtyard or verandah and for how long?
- Who else accompanies them? Do children play alone or in groups in household or courtyard?
- Tell me about how often children from this community come into contact with soil and feces.
  - Courtyard or veranda or household floor soil
  - Child and animal feces
  - Type of animals, children from neighboring households
- How much time do children spend on the courtyard ground and why? by age group
  - <3 years
  - 3-6 years
  - 6+ years
- Who watches over children's behavior when they play in courtyard? Probe- absence and present of primary and secondary caregivers.
- What are the materials that young children can put into their mouth when playing or resting on the ground? List those materials, rank and describe.
- What do the caregivers do when a child puts something from courtyard ground/soil into their mouth?
- What do the caregivers think when a child puts something from soil into their mouth?

**Contamination of Food**

- Tell me about the food preparation methods are used in this area?
- What are the common food storing methods in this area? Rank and identify the main method and explain.
- Who plays the most important role in preparing and storing daily food? Who designates these people and why? Probe- main food and snacks, occasional/seasonal foods.
- Who from the household is responsible for maintaining food storage and hygiene in your area? Probe- social and economic power structure in the household (mother/father-in-laws, husbands).
- Who decides food preparation methods and why?

- What are the ways hands contact prepared and stored food? Whose hands get frequent contact? When, how and why? Explain at least one way.
- When is food for young children prepared during the day? How long is food stored?
- What do people do to keep their food safe? Probe- hand hygiene before preparing food, heating and reheating before serving food to child, child consumption of food and feeding methods. Who feeds, when and how?

**Free roaming domestic animals**

- How many domestic animals doesa typical household have in your area? Make a count. Identify the majority. Make at least 3 categories. Probe to cover all categories.
- Who takes care of animals and animal sheds and disposes of animal feces?
- When and why are animal feces visible in compound/inside household and for how long?
- How can a child come into contact with animals? Which animals do they like? Do parents allow children to touch animals?
- When are the chances that a child can put animal feces from the courtyard ground into their mouth? Probe- which animals, when and where?
- What do parents do if a child puts animal feces in their mouth? Why or why not?
- Are animals allowed in the house? In food preparation areas?

**Contamination via flies**

- What do people think of flies visible around their household environment?
- What are the types of flies you see around your household environment?
- Which one is most dangerous and why? Describe the most dangerous (size, color and other characteristics).
- What are the reasons for presence of different flies in house, courtyard and surrounding environment? Ask specifically type (size, color, other characteristics).
- Why are these flies found around/inside the household? Where do they come from? When is the fly density high and why?
- Is there a chance that these flies can land on food? How frequently? Probe- time and season.

**Availability and type of toilet facility**

- Where do the people in this area usually defecate? Probe- practice open defecation, individual or shared latrine.
- How many latrines does a household usually own? Probe- type and other facilities available for each toilet.Map the compound marking toilets and distances from drinking water source, pond or other water sources and from living room, kitchen etc.
- Who is responsible for cleaning and maintaining the toilet? Probe- man, women, specific roles, frequency.
- What are the hardware available in this community to dispose of feces from courtyard or inside the house? Probe- child and animal feces, adult human feces, feral animal feces?
- How does sanitation change during the different seasons of the year? Probe- when latrine pit is full, feces floats over pan, during rainy season, when latrine is broken.
- Where do the feces from the latrines end up? In ditches, ponds, drains, small rivers, other water bodies?
- What is easy about managing sanitation in your household? Please describe.

---

#### Module 54 – Focus Group Discussion Guideliens in Bengali for environmental assessment

---

**পরিবেশগত নিবিড় মূল্যায়ন (বাংলাদেশে ডায়রিয়া এবং স্যানিটেশন সুবিধার অন্তর্ভূক্তি, মলদুষণের গমনাগমনের মূল্যায়ন)**

**পানীয় জলের উৎসসমূহ, পরিচালনা এবং সংরৰণ চর্চাসমূহ**

- এই এলাকার খাবার পানির প্রধান প্রধান উৎসগুলি কী কী? জিজ্ঞাসা করুন - ধরণ, টিউবওয়েল, পুকুর বা অন্যান্য, সেগুলির ভৌত অবস্থা, দ্বিতীয় ও অন্যান্য উৎসসমূহ, মৌসুমী প্রভাবসমূহ। টিউবওয়েল ও পুকুরের পানির উপর গুরুত্ব বেশী দিন)। এই উৎসগুলি পর্যবেক্ষণ করুন এবং নৈর্ব্যক্তিকভাবে বর্ণনা করুন।
- গৃহস্থালীতে সাধারণত খাবার পানি সংগ্রহ করা কার দায়িত্ব? (জিজ্ঞাসা করুন- প্রাথমিক ও দ্বিতীয় সংগ্রহকারী, কখন ও কেন?)
- পানি সংগ্রহ ও রক্ষণাবেক্ষণের সাধারণ পদ্ধতিগুলি কী কী? এই পদ্ধতিগুলি পর্যায়ক্রমে লিখুন এবং পরিষ্কারভাবে বোঝার জন্য আরও প্রয়োজনীয় প্রশ্ন করুন।
- আপনার গৃহস্থালীতে পান করার পানি সংগ্রহ করা কার দায়িত্ব? (জিজ্ঞাসা করুন- প্রাথমিক ও দ্বিতীয় সংগ্রহকারী, কখন ও কেন?)
- এই পানির উৎসগুলি বা সংরক্ষণকৃত পানি কিভাবে মানুষ বা পশু-পাখির মল দ্বারা দুষিত হতে পারে? এই সম্ভাব্য দুষণের গমনাগমন পথগুলি কী কী যা প্রাপ্ত বয়স্ক মানুষ, শিশু বা পশু-পাখির মলের দ্বারা দুষিত হয়? (অনুমানকৃত বা সম্ভাব্য এই মল দূষণের একটি গমনাগমন চিত্র অংকন করুন, নিবিড় সাক্ষাৎকার থেকে প্রাপ্ত চিত্রের সাথে চেক করুন, অতঃপর পরিবর্তন করুন)

##### সহজপ্রাপ্যতা, পানি নিরাপদকরণ, পানি স্বল্পতা এবং পানির ব্যবহার

- এই এলাকায় কখন কখন পানির স্বল্পতা দেখা দেয়?
- এই সময় এখানকার মানুষজন কী করেন?
- এই সময়ে অন্যান্য সময়ের সাথে পানি সংগ্রহ, সংরক্ষণ এবং ব্যবহারে কী ধরণের পার্থক্য দেখা যায়?
- আপনার গৃহস্থালীতে পানি বিশুদ্ধকরণের সাধারণ পদ্ধতিগুলি কী কী? সাধারণ বা জনপ্রিয় পদ্ধতিগুলি বর্ণনা করুন।
- কোন নির্দিষ্ট সময় কি রয়েছে যখন পানি নিরাপদকরণ বা বিশুদ্ধকরণ কম বা বেশী হয়ে থাকে?
- শিশুদের বয়সভেদসহ আপনার পরিবারের সকল সদস্যদের সাধারণভাবে পানি ব্যবহার সম্পর্কে বলুন।

##### হাতে জীবানুর সংক্রমণ এবং হাত ধোয়ার চর্চা

- কত ঘনঘন এবং কিভাবে আপনি হাত ধোন? জিজ্ঞাসা করুন- প্রধান প্রধান সময়গুলি (বিভিন্ন ধরণের খাবার খাওয়ার আগে, টয়লেট ব্যবহারের পর এবং বাচ্চাকে শৌচানোর পর)। কোন একটি সাধারণ দিনের হাত ধোয়ার চর্চাগুলি বর্ণনা করুন (হতে পারে গতকাল বা আজ)।
- মানুষ কি মনে করে যে তাদের কোন একটি বা উভয় হাতই মানুষের মল দ্বারা দুষিত হতে পারে? জিজ্ঞাসা করুন- ঐ প্রধান প্রধান সময়গুলিতে হাত ধোয়া (একটি চিত্র অংকন করুন যেসব কর্মকান্ড থেকে হাতে মলের সংক্রমন ঘটতে পারে)।
- মানুষেরা কি বাম বা ডান হাত বেশী বেশী ধোয়? কখন কখন এবং কেন?
- হাত ধোয়ার জন্য এখানে কোন কোন উপাদানগুলি সহজে পাওয়া যায়? জিজ্ঞাসা করুন- সব ধরণের উপাদান, প্রাপ্তি ভেদে ও ঘন ঘন ব্যবহারে উপরি পর্যায়ে সাজান? কেন এবং কখন কখন?
- কতবার আপনি আপনার শিশুর হাত পরিষ্কার করেন/ধোন? কখন কখন এবং কিভাবে?
- কতবার আপনার শিশুরা তাদের নিজেদের হাত ধোয়? কখন কখন এবং কিভাবে?
- আপনি কি মনে করেন আপনার শিশুর হাত মানুষের মল দ্বারা দুষিত হতে পারে? পরিবারের কোন ব্যক্তি হাত ধোয়াকে উৎসাহিত বা প্রভাবিত করতে পারেন? জিজ্ঞসা করুন- দাদা, শ্বশুর, স্বামী, স্কুলে যায় এমন বাচ্চারা।

### উঠানের ও ঘরের আশেপাশের মাটি দুষণ

- বাড়ীর উঠান কিভাবে মানুষের মল দ্বারা দুষিত হতে পারে? জিজ্ঞাসা করুন- শিশু এবং বয়স্কগণ, অসুস্থ্য ব্যক্তি, দিনে বা রাতে বা বৃষ্টির সময় খোলা জায়গায় মলত্যাগ করা। স্থানগুলি চিহ্নিত করুন এবং বর্ণনা করুন। তারা ব্যক্ত করতে পারে এমন একটি উদাহরণ দিতে বলুন। মলের উৎস ও তার সংক্রমনের একটি চিত্র অংকন করুন।
- আপনার বাড়ীতে ছোট ছোট শিশুরা কোথায় পায়খানা করে? কখন কখন এবং কোথায়? জিজ্ঞাসা করুন- সাধারণ, নির্দিষ্ট ও অস্থায়ী স্থানগুলি সম্পর্কে।
- ছোট শিশুদের মল কে কোথায় এবং কিভাবে অপসারণ করে? এই চর্চাগুলিতে কি কোন পার্থক্য হয়? পার্থক্যগুলি কী কী এবং কেন হয়? যেখানে যেখানে মল ফেলা হয় সেই স্থানগুলি পর্যবেক্ষণ করুন এবং প্রত্যেক স্থানগুলি নিয়ে আরও প্রশ্ন জিজ্ঞাসা করুন।
- এই পরিবারে কি কোন শিশুদের মলত্যাগ করার জন্য পটি রয়েছে?
- ছোট ছোট শিশুরা তাদের মায়ের/বয়স্কদের কোল ছাড়া আর কোথায় কোথায় অধিকাংশ সময় কাটায়? স্থানগুলির বর্ণনা করুন
- শিশুরা কত সময় উঠানে সময় কাটায় বা বারান্দায় এবং কেন?
- তাদেরকে কে সঙ্গ দেয়? শিশুরা কি একা না অনেকে মিলে খেলাধুলা করে?
- বলুন কিভাবে আপনাদের বাচ্চারা মাটি ও মলের সংস্পর্শে আসে।
  - উঠান বা বারান্দা বা ঘরের ভেতরের মাটি
  - বাচ্চাদের এবং পশু-পাখির মল
  - পশু-পাখির ধরন, প্রতিবেশীদের বাচ্চাদের
- কত সময় বাচ্চারা উঠানের মাটিতে সময় কাটায় এবং কেন? দলভেদে
  - <৩ বছর

- <৩ -৬ বছর
- ৬+বছর

- ক তাদের আচার-আচরণ লৰ্য করে যখন তারা উঠানের মাটিতে খেলাধুলা করে? জিজ্ঞাসা করম্নন- মায়ের বা অন্য যত্নগ্রহণকারীর উপস্থিতি এবং অনুপস্থিতিতে।
- খেলাধুলা বা সময় কাটানোর বেলায় বাচ্চারা কোন কোন জিনিস মুখে দিতে পারে? জিনিসগুলির নাম লিখুন, উপরি-পর্যায়ক্রমে সাজান ও বর্ণনা করম্নন।
- যখন শিশুরা উঠানের মাটি থেকে তাদের মুখে কোন কিছু দেয় তখন মায়েরা কী করেন?
- বাচ্চারা যখন মাটি থেকে কোন কিছু মুখে দেয় তখন মায়েরা কী ভাবেন বা মনে করেন?

**খাদ্য দূষণ**

- দয়া করে আমাকে এই এলাকার খাবার তৈরী এবং সংরৰণ প্রক্রিয়া সম্পর্কে বলুন।
- এই এলাকার সাধারণ খাবার সংরৰণ পদ্ধতিগুলি কী কী? প্রধান পদ্ধতিগুলি উপরি-পর্যায়ক্রমে সাজান এবং ব্যাখ্যা করম্নন।
- প্রতিদিনকার খাবার তৈরী ও সংরৰনে কে সবচেয়ে বেশী ভূমিকা পালন করে? কে তাদেরকে সেই দায়িত্ব দেয় ও কেন? জিজ্ঞাসা করম্নন- প্রধান প্রধান খাবার, নাস্ত্মা, মৌসুমি খাবার।
- এই অঞ্চলে সাধারণত খাবার সংরৰণ ও তা স্বাস্থ্যসম্মত রাখার দায়িত্ব কার? জিজ্ঞাসা করম্নন- পরিবারের সামাজিক ও অর্থনৈতিক ৰমতা (শ্বশুর-শ্বাশুড়ী, স্বামী)
- খাবার তৈরীর পদ্ধতি সম্পর্কে কে সিদ্ধান্ত্ম নেয় এবং কেন?
- তৈরীকৃত ও সংরৰিত খাবারগুলি কিভাবে হাতের সংস্পর্শে আসে? কার হাতে বার বার ধরা হয়?
- খাবার নিরাপদ রাখার জন্য কী কী করা হয়? জিজ্ঞাসা করম্নন- খাবার তৈরীর পূর্বে হাত ধোয়া, বাচ্চাকে খাওয়ানোর আগে বার বার খাবার গরম করা, শিশুদের খাবারের পরিমাণ এবং খাওয়ানোর পদ্ধতি। কে খাওয়ায়, কখন কখন ও কেন?
- বাচ্চাদের জন্য খাবার দিনের কোন সময়ে তৈরী করা হয়? কতৰণ সংরৰণ করা হয়?

**উন্মুক্তভাবে থাকা গৃহপালিত পশু-প্রাণীসমূহ**

- সাধারণত আপনাদের এলাকায় একটি সাধারণ গৃহস্থালীতে কতগুলি গৃহপালিত পশু-পাখি থাকে? গণনা করম্নন। সবচেয়ে বেশী কোনগুলি তা চিহ্নিত করম্নন। কমপৰে ৩ টি শ্রেণী করম্নন। সবগুলি শ্রেণী সম্পর্কে জিজ্ঞাসা করম্নন।
- পশু-পাখী ও খোয়ার বা গোয়ালঘর কে দেখাশোনা করে
- কখন কখন এবং কেন পশু-পাখীর মল আপনার বাড়ীর উঠানে ও আশে পাশে দেখা যায় এবং কত সময় ধরে?
- একটি বাচ্চা শিশু কিভাবে পশু-পাখীর সংস্পর্শে আসতে পারে? কোন প্রাণী তারা পছন্দ করে? মা-বাবারা কি বাচ্চাদেরকে পশু-পাখির সংস্পর্শে আসতে দেয়?
- এমন কি কোন সম্ভাবনা আছে যে শিশুরা উঠানের মাটি থেকে পশু-পাখির কোন মল মুখে দিতে পারে? কোন প্রাণীর, কখন ও কোথায়?
- বাচ্চারা যদি পশু-পাখির মল মুখে দেয় তবে মা-বাবারা কী করেন? কেন বা কেন নয়?
- বাড়ীতে কি পশুপাখি আসতে দেয়া হয়? রান্না ঘরে বা যেখানে খাবার তৈরী হয়?

**মশা-মাছির মাধ্যমে দূষণ/সংক্রমণ**

- আপনার বাড়ীর ভেতর এবং আশে পাশে বিভিন্ন ধরণের মাছির উপস্থিতি সম্পর্কে আপনার কী মনে হয়? সুনির্দিষ্টভাবে ধরণগুলি সম্পর্কে জিজ্ঞাসা করম্নন (আকৃতি, রং, অন্যান্য বৈশিষ্ট্যসমূহ)
- কোন ধরণের মাছি বাড়ীর চারপাশে দেখা যায়? কোনগুলি বেশী বিপদজনক ও কেন? বেশী বিপদজনকগুলি বর্ণনা করম্নন (আকৃতি, রং, অন্যান্য বৈশিষ্ট্যসমূহ)
- এই ধরণের মশা-মাছিগুলি আপনার ঘরের ভেতর বা আশে পাশে কেন পাওয়া যায়? এগুলি কোথা থেকে আসে? কখন কখন মশা-মাছির উপদ্রব বেশী হয় ও কেন?
- এমনি কি কোন সুযোগ আছে যে মশা-মাছি খাবারের উপর বসে? কত ঘন ঘন এমন হয়? জিজ্ঞাসা করম্নন- সময় ও ঋতু।
- আপনি কি মনে করেন যে মাছিরা আপনার পরিবারের জন্য কোন স্বাস্থ্য ঝুকি বয়ে আনে? কেন অথবা কেন নয়?

**টয়লেট সুবিধাদির ধরণ ও সহজলভ্যতা**

- এ এলাকার মানুষজন সচরাচর কোথায় সচরাচর পায়খানা/মলত্যাগ করেন? জিজ্ঞাসা করম্নন- খোলা জায়গায়, ব্যক্তিগত/নিজস্ব বা অন্যের ল্যাট্রিনে।
- একটি পরিবারের কয়টি ল্যাট্রিন আছে? জিজ্ঞাস করম্নন- প্রতিটি ল্যাট্রিনের ধরন ও অন্যান্য সুবিধাসমূহ। এই বাড়ীটির একটি চিত্র অংকন করম্নন যেখানে ল্যাট্রিন/টয়লেট সুবিধাদি এবং খাবার পানির উৎস, পুকুর বা অন্যান্য উৎস, থাকার ঘর ও রান্না ঘর থেকে তার দূরত্ব চিহ্নিত করম্নন।
- টয়লেট ব্যবস্থাপনা ও পরিস্কার রাখার দায়িত্ব কার? জিজ্ঞাসা করম্নন- পুরম্নষ, নারী, নির্দিষ্ট ভূমিকা, কত সময় পর পর?
- উঠান থেকে অপসারণ করার জন্য এই পরিবারের কোন কোন যন্ত্র-পাতি রয়েছে। জিজ্ঞসা করম্নন- শিশু ও পশুর মল, বয়স্ক দের ও অপোষা প্রাণীদের।
- বছরের বিভিন্ন মৌসুমে কী হয়? জিজ্ঞাস করম্নন- ল্যাট্রিন যখন ভরে যায়, প্যানের উপর মল ভাসতে থাকে, বা ল্যাট্রিন যখন ভেঙ্গে যায়।
- ল্যাট্রিন থেকে মল কোথায় গিয়ে শেষ পৌছে? নালায়, পুকুরে, ডোবায়, ছোট নদী, গাঙ্গে, বা অন্যান্য কোন পানিতে।
- পরিবারের সদস্যরা মলত্যাগ ও পরিবেশ মলমুক্ত রাখার জন্য কী কী অসুবিধার মুখোমুখি হন বর্ণনা করম্নন।

#### Household Latrine Survey

Note: Ask these Questions to any adult family member
(এই প্রশ্নগুলো ল্যাট্রিন সম্পর্কে তথ্য দিতে পারবে খানার এমন কোন প্রাপ্ত বয়স্ক সদস্যকে জিজ্ঞাস করতে হবে)

---

**Section-1: Household unique identifier**

1.1 বাড়ী নং (Bari ID): .. .. .. .. .. .. ... .. .. .. .. .. .. .. .. .. .. .. ... .. .. .. .. .. .. .. .. .. .. .. .. .. .. . .. [ ][ ][ ]

1.2 খানা নং**(**Household ID): ........................................................................................................ [ ][ ][ ][ ][ ]

**(Please follow the specific code sheet)**

1.3 ইউনিয়নের নাম (Union name): ........................................................................................ [ ][ ][ ]

1.4 **ঠিকানা** (Address): খানা প্রধানের নাম: ..................................................................................

(Name of household head)

খানা প্রধানের বাবার/স্বামীর নাম: ...........................................................................................

(Father's/ husband's name)

সাক্ষাতকার প্রদানকারীর নাম: ...............................................................................................

(Name of respondent)

সাক্ষাতকার প্রদানকারীর সাথে খানা প্রধানের সম্পর্ক:

(Relation with HH head)

1.বাবা (father) 2.মা (mother) 3.বড়ভাই (elder brother) 4. দাদা ((grand-father))

5. দাদী (grand-mother) 6. চাচা (uncle) 7. চাচী (unty) 8. মামা (maternal uncle)

77. অন্যান্য:

বাড়ীর নাম: ....................................................................................................................... [ ][ ][ ]

(Bari Name)

গ্রাম: .................................................................................................................................

(Village)

বাড়ীর অবস্থান (বিস্তারিত লিখুন): ..........................................................................................

Location (specify)

1.5 সাক্ষাতকার গ্রহণকারীর নাম এবং কোড (Interviewer name & code): ................................................ [ ][ ][ ][ ]

[ ][ ][ ][ ][ ][ ]

1.6 তথ্য সংগ্রহের তারিখ (Date of data collection/observation/spot check): / /

**Section 2: Respondent and household demographics**

| খানার সদস্যদের নাম [**List of household members (name)]**<br>**(খানার সবচেয়ে ছোট বাচ্চাকে দিয়ে শুরু করতে হবে)** | বয়স [Age] | লিঙ্গ [Sex]<br>1. Male<br>2. Female | শিক্ষা [Education]<br>99-জানিনা 88-**প্রযোজ্য নয়** | পেশা [Occupations] | ল্যাট্রিন ব্যবহার (Latrine user) |
|---|---|---|---|---|---|
| **1.** | | | | | |
| **2.** | | | | | |
| **3.** | | | | | |
| **4.** | | | | | |
| **5.** | | | | | |
| **6.** | | | | | |
| **7.** | | | | | |
| **8.** | | | | | |
| **9.** | | | | | |
| **10.** | | | | | |
| **11.** | | | | | |
| **12.** | | | | | |
| **13.** | | | | | |
| **14.** | | | | | |
| **15.** | | | | | |

Occupation Code:

**36.** কৃষক [Farmer/Cultivator]
**37.** ঘরে কাজ করে [Homemaker]
**38.** কৃষি শ্রমিক [Agri-labor]
**39.** শ্রমিক (কৃষি শ্রমিক ছাড়া) [Non-agri labor]
**40.** বেতনভুক্ত কর্মচারী (সরকারী/প্রাইভেট/এন.জি.ও) [Salaried job (Govt./Private/NGO)]
**41.** রাজমিস্ত্রী [Mason (*Rajmistri*)]
**42.** কাঠ মিস্ত্রী [Carpenter]
**43.** ভ্যান/রিকশা চালক [Van/Rickshaw puller]
**44.** জেলে [Fisherman]
**45.** নৌকা চালক/মাঝি [Boatman]
**46.** কর্মকার [Blacksmith]
**47.** স্বর্ণকার [Goldsmith]
**48.** কুমার/কুম্ভকার [Potter (soil smith)]
**49.** মুচি [Shoe polish /maker]
**50.** দোকানদার [Shopkeeper]
**51.** ফেরিওয়ালা [Vendor (*Feriwala/howker*)]
**52.** ক্ষুদ্র ব্যবসায়ী (মূলধন <=১০০০০) [Petty trader, capital <=10000]
**53.** ব্যবসায়ী(মূলধন >১০০০০) [Business, capital >10000]
**54.** দর্জি [Tailor]
**55.** ড্রাইভার [Driver]
**56.** কুটীর শিল্প [Cottage industry]
**57.** পোলট্রি/ ব্যবসার জন্য পশু লালন-পালনকারী [Poultry /livestock re
**58.** বৈদ্যুতিক মিস্ত্রী [Electrician]
**59.** হোমিওপ্যাথি ডাক্তার [Homeopath]
**60.** আধ্যাতিক চিকিৎসক/ কবিরাজ/ ওঝাঁ [Spiritual healer/kabiraj/ Ojh
**61.** পেশাদার ডাক্তার/উকিল [Professional practitioner (Doctor/lawye
**62.** ইমাম/ ধর্মযাজক [Imam/priest]
**63.** অবসরপ্রাপ্ত চাকুরীজীবী [Retired service holder]
**64.** ছাত্র [Student]
**65.** বেকার [Unemployed]
**66.** অক্ষম [Disabled]
**67.** কাজের লোক [Domestic maid / servant]
**68.** জমিদার (শস্য উৎপাদন অথবা অন্য কোন কাজে কৃষকদের জমি বর্গা দে
[Landlord (Provide land for farmers for sharecropping or o
**69.** বিদেশে থাকে [Staying abroad]
**70.** মৃত/নিখোঁজ [Died/untraced]
99. জানিনা [Don't know]

88. প্রযোজ্য নয় [Not applicable]

77. অন্যান্য (বর্ণনা লিখুন) [Others (specify)]

..................................................................

**Self-reported data**

**Section-3: Access to latrine**

1. আপনার/আপনার পরিবারের সদস্যদের কি ল্যাট্রিন ব্যবহারের সুযোগ আছে [Do you/family members have access to a latrine]?

   1. হ্যাঁ [Yes] 2. না [No]

   (যদি উত্তর হ্যাঁ হয় তাহলে প্রশ্ন-২ এ যান অথবা 'না' হয় তাহলে প্রশ্ন-২ এ যান তারপর সেকশন-৫ এ যান) [if answer is 'no', then ask ques-2 and skip to section-5]

2. আপনার/আপনার পরিবারের সদস্যরা কোথায় পায়খানা করেন [Where do you/family members defecate]?

   1. উঠানে [yard] 2.উঠানের বাহিরে খোলা জায়গাতে [Open space outside the front yard]

   3. ঝোপ-ঝাড়ে/জঙ্গলে [Bush/jungle] 4.ল্যাট্রিন /টয়লেটে [In toilet]

   77. অন্যান্য (বর্ণনা লিখুন) [Other (Please describe)]................................

3. ল্যাট্রিন কি আপনার [Do you own the latrine (that you use)]?

   1. হ্যাঁ [Yes] 2.না [No] (যদি উত্তর হ্যাঁ হয় তাহলে প্রশ্ন-৪ এ যান অথবা না হয় তাহলে **প্রশ্ন-৬** এ যান )[if answer is 'No' skip to **quest-6**]

4. ল্যাট্রিনের মালিকানার ধরন (Type of ownership)?

1. একক মালিকানা (self own) 2. যৌথ মালিকানা (jointly own)

(যদি উত্তর একক মালিকানা হয় তাহলে **সেকশন-৪** এ যান) [if self own, go to **section-4**]

5. যদি ল্যাট্রিনটি যৌথমালিকানায় হয় তাহলে অংশীদার কে?

1. আত্মীয় (একই বাড়ীর মধ্যে) [relative] 2. প্রতিবেশী [neighbor] 77.**অন্যান্য** [Other]................................

6. আপনি/আপনার পরিবারের সদস্যরা কি অন্য খানা/বাড়ীর ল্যাট্রিন ব্যবহার করেন [Do you/family members use another household’s latrine]?

1. হ্যাঁ [Yes] 2. না [No]

(যদি উত্তর হ্যাঁ হয় তাহলে ৭ নং প্রশ্ন জিজ্ঞাসা করুন এবং সেকশন-৫ এ যান, যতি উত্তর না হয় তাহলে সেকশন-৪ এ যান)

(If answer is ‘yes’, ask question-7 and skip to section-5, if answer is ‘no’, skip to section-4)

7. কোন খানার ল্যাট্রিন আপনি/আপনার পরিবারের সদস্যরা শেয়ার করেন (বাচ্চারা সহ) (আত্মীয় না প্রতিবেশীর ল্যাট্রিন তা জানতে হবে)
[Which household do you share this latrine (including children)]?

1. আত্মীয় (একই বাড়ীর মধ্যে) [relative] 2. প্রতিবেশী [neighbor] 77.**অন্যান্য** [Other]................................

**Section-4: Duration of latrine use and pit emptying**

8. আপনি/আপনার পরিবারের সদস্যরা কত মাস ধরে এই ল্যাট্রিন ব্যবহার করছেন [How long have you been using this latrine]?
(দিন/বছর যেতে তা মাসে লিখতে হবে)

☐☐ MM

9. এখন যে ল্যাট্রিন ব্যবহার করছেন তার কোন পিট/গর্ত কি কখনও খালি করা হয়েছে ? [Did you ever empty a pit of existing latrine]

1. হ্যাঁ [Yes] 2. না [No]

( যদি উত্তর হ্যাঁ হয় তাহলে প্রশ্ন- ১০ এ যান অথবা না হয় তাহলে প্রশ্ন-১৫ তে চলে যান) [if ‘no’, skip to question-15]

10. কত বার এই ল্যাট্রিনের পিট খালি করা হয়েছে? [How many times did you empty pit of existing latrine] ☐☐

11. সর্ব শেষ কবে এই ল্যাট্রিনের পিট খালি করা হয়েছিল? [When did you last empty the pit of existing latrine] MM ☐☐

**(যদি যৌথ মালিকানায় হয় তাহলে প্রশ্নঃ ১২ করতে হবে)** [if answer is Jointly own then ask to **quest-12**]

12. সর্ব শেষ কার খরচে এই ল্যাট্রিনের পিট খালি করা হয়েছিল? [Who bear the cost for last empty the pit]

1. নিজের 2.প্রতিবেশী 3.আত্মীয়

13. কি ভাবে পিট খালি করা হয়েছিল [How did you empty]?

1. নিজে/পরিবারের সদস্য [By yourself]    2. সুইপার ভাড়া করে [By hiring sweeper]    77. অন্যান্য [Other].. .. .. .. .. .. ..

14. যদি সুইপার ভাড়া করা হয় তাহলে সর্ব শেষ পিট খালি করতে কত টাকা খরচ হয়েছিল [If, emptied hiring a sweeper, how much does it cost for empty the pit last time]?    টাকা [ ☐☐☐☐

15. গত এক বছরে আপনার/আপনাদের ল্যাট্রিন কি মেরামত করা হয়েছে [Did you ever repair latrine structure]?

1. হ্যাঁ [Yes]    2. না [No]

(যদি উত্তর হ্যাঁ হয় তাহলে প্রশ্ন-১৬ অথবা না হয়, তাহলে সেকশন-৫ এ চলে যান) [if answer is 'no', skip to secktio-5]

16. গত এক বছরে আপনার/আপনাদের ল্যাট্রিন কতবার মেরামত করা হয়েছে [How many times did you repair the structure (within last one year)?    ☐☐

17. কোন অংশ সব চেয়ে বেশী মেরামত করেছেন এবং কত বার [Which part did you repair most]?

1. দেয়াল/ বেড়া [Walls/fences]    ☐☐

2. দরজা [Door]    ☐☐

3. ছাদ [Roof]    ☐☐

77. অন্যান্য [Other].. … … … … … … … … … … … … … … … … … … … … … … …

18. সর্ব শেষ কোন অংশ মেরামত করেছেন [Which part did you repair last]?

1. দেয়াল/ বেড়া [Walls/fences]
2. দরজা [Door]
3. ছাদ [Roof]
77. অন্যান্য [Other-describe] . . . . . . . . . . . . . . . . . . . . . . . . . . . . . . . . . . . . . . . .

19. কে মেরামত করেছিল [Who repaired it]?

1. নিজে/পরিবারের সদস্য [By yourself]    2. শ্রমিক ভাড়া করে [hire daily labor]

77. অন্যান [other].............................

20. সর্ব শেষ মেরামতে কত খরচ হয়েছিল [How much did it cost to repair (last time repaired)]?

☐☐☐☐ টাকা [Taka]

**Section-5: consent on latrine upgrade**

21. যদি আমরা গবেষণার প্রয়োজনে নতুন ল্যাট্রিন স্থাপন করে দিতে চাই, আপনি কি তাতে রাজী হবেন [If we want to set up a new latrine for research purpose, will you agree]?

1. হ্যাঁ [Yes] 2. না [No]

[যদি উত্তর হ্যাঁ হয় তাহলে প্রশ্ন-২৩ এ যান অথবা না হলে প্রশ্ন-২২ এ যান, যদি ল্যাট্রিন না থাকে তহলে প্রশ্ন ২১ ও ২২ দুটোই প্রযোজ্য হবে] [if answer is 'No' skip to **quest-22**]

22. যদি উত্তর (২১ নং প্রশ্নের) না হয় তাহলে কারন বর্ণনা করুন (বিস্তারিত)ঃ … … … … … … … … … … … … … … … … … … … … … … … … …

… … … … … … … … … … … … … … … … … … …… … … … … … … … … … … … … … … … … … …… … … … … … … … … … … … … … … … … … …

… … … … … … … … … … … … … … … … … … …… … … … … … … … … … … … … … … … … … …… … … … … … … … … … … … … … … … … … …

23. যদি আমরা আপনার ল্যাট্রিন গবেষণার প্রয়োজনে সংস্কার করতে চাই, আপনি কি তাতে রাজী হবেন [If we want to upgrade your existing latrine for research purpose, will you agree]?

1. হ্যাঁ [Yes] 2. না [No] যদি উত্তর হ্যাঁ হয় তাহলে প্রশ্ন-২৫ এ যান,উত্তর না হলে প্রশ্ন-২৪ এ যান] [if answer is 'No' skip to **quest-24**]

24. যদি উত্তর (২৩ নং প্রশ্নের) না হয় তাহলে কারন বর্ণনা করুন (বিস্তারিত)ঃ … … … … … … … … … … … … …… … … … … … … … … … … … … … … … … …

…… … … … … … … … … … … … … … … … … … …… … … … … … … … … … … … … … … … … …

25. ল্যাট্রিন স্থাপনের জন্য আপনার কতটুকু জায়গা আছে (আত্মীয়-স্বজন ও প্রতিবেশী সহ) (নিজের মালিকানায় নাও হতে পারে কিন্তু আত্মীয়/প্রতিবেশীর জায়গায় পায়খানা স্থাপনের অনুমতি আছে) [How much land do you have access to install a latrine]? [Including neighbor, relative and kin]

(ডেসিমেল)[decimal] ☐☐

26. ল্যাট্রিন স্থাপনের জন্য আপনার নিজের মালিকানাধীন কতটুকু জায়গা আছে [How much land do you own to install a latrine]? ☐☐

(ডেসিমেল) [decimal]

(যদি ল্যাট্রিন স্থাপনের জন্য নিজের জায়গা থাকে তাহলে প্রশ্ন-৩১ এ যেতে হবে, জায়গা না থাকলে প্রশ্ন-২৭ এ যেতে হবে) [if HH own land for latrine installation then skip to quest-31]

27. আপনার আত্মীয়-স্বজন এবং প্রতিবেশীরা কি (তাদের জায়গায়) ল্যাট্রিন স্থাপনের অনুমতি দেবেন

[Would your neighbors and relatives (kin group) permit you to install a latrine in their land]

1. হ্যাঁ [Yes] 2. না [No]

[যদি উত্তর হ্যাঁ হয় তাহলে প্রশ্ন-৩১ এ যান. উত্তর না হলে প্রশ্ন ২৮ এ যান] [if answer is 'No' skip to **quest-28**]

28. যদি উত্তর (২৭ নং প্রশ্নের) না হয় তাহলে কারন বর্ণনা করুন (বিস্তারিত) [Describe if answer is no to quest-27]:

… … … … … … … … … … … … … … … … … … …… … … … … … … … … … … … … … … … … … …… … … … … … … … … … … … … … … … … … …

… … … … … … … … … … … … … … … … … …… … … … … … … … … … … … … … … … …… … … … … … … … … … … … … … … …

29. আপনার আত্মীয়-স্বজন এবং প্রতিবেশীরা কি (তাদের জায়গায়) স্থাপিত ল্যাট্রিন কম পক্ষে দুই (২) বছর ব্যবহারের অনুমতি দেবেন
[Would your neighbors and relatives (kin group) permit you to use that latrine at least 2 year]?

1. হ্যাঁ [Yes] 2. না [No]

[যদি উত্তর হ্যাঁ হয় তাহলে প্রশ্ন-৩১ এ যান, না হলে প্রশ্ন ৩০ যান] [if answer is 'No' skip to **quest-30**]

30. যদি উত্তর (২৯ নং প্রশ্নের) না হয় তাহলে কারন বর্ণনা করুন (বিস্মারিত [Describe if answer is no to quest-29]… … … … … … … …

… … … … … … … … … … … … … … … … … …… … … … … … … … … … … … … … … … …… … … … … … … … … … … … … … … …

… … … … … … … … … … … … … … … … … …… … … … … … … … … … … … … … … … …… … … … … … … … … … … … … … … …

31. যে জায়গায় নতুন ল্যাট্রিন /পিট স্থানপন করা হবে তার নিকট বর্তী পানির উৎসগুলো বিস্মারিত বর্ননা করুন (পনির উৎসের নাম ও দূরত্ব):

[nearest water sources and distance from proposed duel pit latrine place]

| পানির উৎসের নাম<br><br>(Source of water) | দূরত্ব (প্রস্মাবিত নতুন ল্যাট্যিন/পিট স্থাপনের জায়গা)<br><br>(Distance) | গভীরতা (টিউব-ওয়েল হলে)<br><br>(Depth-if tub-well) |
|---|---|---|

**Section-6: Observation of household latrines (by FRA or data collector)**

32. খানায় কতগুলো ল্যাট্রিন ব্যবহৃত হচ্ছে (নিয়মিত/অনিয়মিত) [How many latrines is in-use? (regular/ irregular)] ☐☐

33. খানায় যে ল্যাট্রিনগুলো ব্যবহৃত হচ্ছে (নিয়মিত/অনিয়মিত) তার ধরন [Type of latrines is in-use in HH? (regular/ irregular)]

ল্যাট্রিন-১ [Latrine: 1] ধরন[Type:] . . . . . . . . . . . . . . . . . . . . . . . . . . . . . . . . . . . . . . . . . .

ল্যাট্রিন-২ [Latrine: 2] ধরন[Type:] . . . . . . . . . . . . . . . . . . . . . . . . . . . . . . . . . . . . . . . . . .

স্বাস্থ্য সম্মত-টয়েলেট (Improved sanitation facilities)-

ফ্লাশ-টয়লেট অথবা পানি ঢেলে ফ্লাস করা টয়লেট [Flush or pour flush toilet flushed to]:

টয়লেটটিতে পয়ঃনিষ্কাশন পাইপের সাথে সংযোগ করে দেয়া [Piped sewer system]................................01

টয়লেটটিতে সেপটিক ট্যাংক বসানো আছে [Septic tank]......................................................................02

সেপটিক ট্যাংক নাই কিন্তু ফ্লাস করে বা পানি ঢেলে পায়খানা দুরে পিটের মধ্যে সরিয়ে দেয়া যায় ..........................03

[Flush to pit latrine (Off set)]

পিট-টয়লেট (স্লাব এবং ওয়াটার সিল আছে) [Pit latrine with slab & water seal] ...............................................04

পিট-টয়লেট (স্লাব আছে কিন্তু ওয়াটার সিল নেই তবে ঢাকনা দেয়ার ব্যবস্থা আছে) .............................................................05

[Pit latrine with slab & no water seal but with a lid]

পিট-টয়লেট (স্স্লাব এবং ফ্ল্যাপ আছে কিন্তু ওয়াটার সিল নেই)
[Pit latrine with slab and flap, no water seal]..................................................45

**বায়ু চলাচল উপযোগী উন্নত ল্যাট্রিন** [Ventilated Improved Pit (VIP) latrine]………………………46

কমপোষ্টিং টয়লেট **(পায়খানা এবং প্রস্রাব করার জন্য আলাদা আলাদা ঘর এবং সাথে আলাদা কোথাও পানির ব্যাবস্থা আছে)**

[Composting toilet, *(Composting toilet ensure separation of urine, water and excreta)*] .................06

**(পায়খানা এবং প্রস্রাব করার জন্য আলাদা আলাদা ঘর এবং সাথে আলাদা কোথাও পানির ব্যাবস্থা আছে)**

ফ্লাশ-টয়েলেট অথবা পানি ঢেলে ফ্লাস করা টয়লেট যা কোন খাল, ড্রেন, নদী ইত্যাদির সাথে সংযোজন

করার ফলে অস্বাস্থ্যকর অবস্থার সৃষ্টি করে থাকে.........................................................................................07

[Flush or pour flush toilet connected to somewhere else (canal, ditch, river, etc.)]

পিট/গর্ত পায়খানা, স্লাব নেই এবং যেখান থেকে মশা/মাছি যাওয়া আসা করতে পারে এবং দূর্গন্ধ ছড়ায়.......................................08

[Pit latrine without slab/open pit]

পিট/গর্ত পায়খানা স্লাব আছে তবে,ওয়াটার সিল নেই অথবা ওয়াটার সিল ভাঙ্গা এবং কোন ঢাকনাও নেই ...................................09

[Pit latrine with slab & no water seal/broken water seal and no lid]

ঝুলন্ত পায়খানা [Hanging toilet/latrine] ....................................................................................................10

<u>খোলা পায়খানা/ টয়লেট (**Open defecation**):</u>

কোন পায়খানা নেই/জঙ্গলে/ঝোপে ঝাড়ে/ খোলা জায়গায় [No facility/bush/field]....................................................11

**অন্যান্য (নির্দিষ্ট করে লিখুন)** [Others: Specify].......................................................................................77

৩৪. ল্যাট্রিনের উপরি কাঠামোর বিস্তারিত বর্ননা [ structure of latrine]

ল্যাট্রিন-১ [Latrine: 1]

1.দেয়াল/ বেড়া [Walls/fences] . . . . . . . . . . . . . . . . . . . . . . . . . . . . . . . . . . . . . . . . . . . . . . . . . . . . . .

2.দরজা [Door] . . . . . . . . . . . . . . . . . . . . . . . . . . . . . . . . . . . . . . . . . . . . . . . . . . . . . . . . . . . . . .

3.ছাদ [Roof] . . . . . . . . . . . . . . . . . . . . . . . . . . . . . . . . . . . . . . . . . . . . . . . . . . . . . . . . . . . . . . .

ল্যাট্রিন-২ [Latrine: 2]

1.দেয়াল/ বেড়া [Walls/fences] . . . . . . . . . . . . . . . . . . . . . . . . . . . . . . . . . . . . . . . . . . . . . . . . . . . . . .

2.দরজা [Door] . . . . . . . . . . . . . . . . . . . . . . . . . . . . . . . . . . . . . . . . . . . . . . . . . . . . . . . . . . . . .

3.ছাদ [Roof] . . . . . . . . . . . . . . . . . . . . . . . . . . . . . . . . . . . . . . . . . . . . . . . . . . . . . . . . . . . . .

35. নিয়মিত খাবার পানির উৎস (রেসপনডেন্ট চিহ্নিত করবেন) [Source of regular drinking water (as defined by the respondents)]?

1. **অগভীর টিউবওয়েল (২৫০ ফিটের কম)**[Shallow tube well] (<250 feet)

2. **গভীর টিউবওয়েল (২৫০ ফিটের বেশী)** [Deep tube well]. (250+ feet)

3. **নদী/বাধ/লেক/পুকুর/সেচ নালাথেকে জীবানুমুক্তকরন পানি**

[Pathogen treatment plant (Pond Sand Filter)]: River/dam/lake/ponds/stream/canal/irrigation channel.]

4. **নদী/বাধ/লেক/পুকুর/সেচ নালাথেকে সরাসরি সংগৃহীত পানি**

[Directly from River/dam/lake/ponds/stream/canal/irrigation channel]

77. অন্যান্য [Other] (বর্ণনা করুন): . . . . . . . . . . . . . . . . . . . . . . . .

36. ল্যাট্রিন (সবচেয়ে বেশী ব্যবহৃত হয় যে ল্যাট্রিনটা) ও খাবার পানির উৎসের মধ্যে দূরত্ব (আনুমানিক) [Distance (approximately) between latrine (mostly in use) and drinking water source (tube-well)]?

**(নিজের/আত্মীয়/প্রতিবেশীর মালিকানার খাবার পানির উৎসের দূরুত্ব)**

<৩০ ক[illegible] [St]eps] (৩০ কদমের বেশী হলে মাপার দরকার নাই) ☐☐   > ৩০ এর বেশী (>more than 30 [ste]ps] ☐

37. বর্তমান ল্যাট্রিনে কয়টি পিট আছে (বর্তমানে ব্যবহৃত হচ্ছে বা খানার সদস্যরা যে ল্যাট্রিন বেশী ব্যবহার করে) [Number of pit in existing latrine? (currently in use, mostly used by the household)] ☐☐

38. পিটে কতটি রিং ব্যবহৃত হয়েছে (প্রশ্ন করতে হবে) [Number of rings used in the pit? (need to ask)] ☐☐

39. রিং এর বর্তমান অবস্থা কি (যে রিং গুলো দেখা যায়) [What is the current condition of the rings]?

| | | |
|---|---|---|
| পিট-১ [Pit-1]: | 1. ভাল [Functional] | 2. ভাঙ্গা [Broken] |
| পিট-২ [Pit-2]: | 1. ভাল [Functional] | 2. ভাঙ্গা [Broken] |

40. যদি রিং ভাঙ্গা হয়, তাহলে তার বিস্তারিত বর্ণনা করুন [If ring is broken, describe the situation]?

পিট-১ [Pit-1]: . . . . . . . . . . . . . . . . . . . . . . . . . . . . . . . . . . . . . . . . . . . . . . . . . . . . . . . . . . . . . . . .

পিট-২ [Pit-2]: . . . . . . . . . . . . . . . . . . . . . . . . . . . . . . . . . . . . . . . . . . . . . . . . . . . . . . . . . . . . . . .

41. পিট থেকে কি পায়খানা বাইরে বের হচ্ছে [Does feces spill out from the pit]?

1. হ্যাঁ [Yes] 2. না [No]

(যদি উত্তর হ্যাঁ হয় তাহলে প্রশ্ন-৪২ এ যান অথবা 'না' হলে ৪৫ নং প্রশ্নে চলে যান) [if answer is 'yes' then describe (question-42), otherwise skip to question-45]

42. পায়খানা বের হয়ে কোথায় পরছে [Where the spilled feces get to]? … … … … … … … … … … … … … … … … … … … … … … … … …
… … … … … … … … … … … … … … … … … … …… … … … … … … … … … … … … … … … … … … …… … … … … … … … … … … … … … … … … … …

… … … … … … … … … … … … … … … … … … …… … … … … … … … … … … … … … … … … … … …… … … … … … … … … … … … … … … … … … …

43. পিটের চার পাশে পায়খানা দেখা যাচ্ছে কি [Is there feces visible around the pit]?

1. হ্যাঁ [Yes] 2. না [No]

44. পায়খানা কি ভাবে পিট এর বাইরে বের হচ্ছে তা বর্ণনা করুন [Describe how does feces come out from the pit]? . … … … … … …
… … … … … … … … … … … … … … … … … … …… … … … … … … … … … … … … … … … … … … …… … … … … … … … … … … … … … … … … … …

… … … … … … … … … … … … … … … … … … …… … … … … … … … … … … … … … … … … … … …… … … … … … … … … … … … … … … … … … …

45. ল্যাট্রিনের স্লাবের [Floor] বর্তমান অবস্থা কি [What is the current condition of latrine slab]?

1. ভাল [Functional] 2. ভাঙ্গা [Broken]

46. যদি স্লাব [Floor] ভাঙ্গা হয় বা খারাপ হয় তাহলে তার বর্ণনা করুন [If slab is damaged or broken describe it]? … … … … … … …

… … … … … … … … … … … … … … … … … … …… … … … … … … … … … … … … … … … … … … …… … … … … … … … … … … … … … … … … … …

… … … … … … … … … … … … … … … … … … …… … … … … … … … … … … … … … … … … … … …… … … … … … … … … … … … … … … … … … …

47. স্লাবে [Floor] কি পায়খানা দেখা যাচ্ছে [Are there any visible feces on the slab]?

1. হ্যাঁ [Yes] 2. না [No]

(যদি উত্তর হ্যাঁ হয় তাহলে প্রশ্ন-৪৮ এ যান অথবা 'না' হয় তাহলে ৪৯ নং প্রশ্নে চলে যান) [if answer is 'no', skip to the question-48]

48. যদি হ্যাঁ হয়, তার বর্ণনা করুন (কোন অংশে দেখা যাচ্ছে, পরিমান) [If yes, describe it? (which part it appears, quantity… … … …

… … … … … … … … … … … … … … … … … … …… … … … … … … … … … … … … … … … … … … …… … … … … … … … … … … … … … … … … … …

… … … … … … … … … … … … … … … … … … …… … … … … … … … … … … … … … … … … … … …… … … … … … … … … … … … … … … … … … …

49. প্যানে পায়খানা দেখা যাচ্ছে কি [Is there any feces appear on the pan]?

1. হ্যাঁ [Yes] 2.না [No] 3. নাই [No pan]

( যদি উত্তর হ্যাঁ হয় তাহলে প্রশ্ন- ৫০ এ যান অথবা 'না'হলে ৫১ নং প্রশ্নে চলে যান) [if answer is 'no', skip to the question-50]

50. যদি হ্যাঁ হয়, তার বর্ণনা করুন [If yes, describe it]? . . . . . . . . . . . . . . . . . . . . . . . . . . . . . . . . . . . . . . . . . . . . . . . . . . . . . . . . . . . . . . . . . . . . . . . . . . . . . . . . . . . . . . . . . . . . . . . . . . . . . . . . . . . . . . . . . . . . . . . . . . . . . . . . . . . . . . . . . . . . . . . . . . . . . . . . . . . . . . . . . . . . . . . . . . . . . . . . . . . . . . . . . . . . . . . . . . . . .

51. ল্যাট্রিনে কি ওয়াটার সিল/সাইফুন দেখা যাচ্ছে [Is there is any water seal/siphon appear in the latrine]?

1. হ্যাঁ [Yes]    2. না [No]    3. নাই [No water seal]

(যদি দেখা সম্ভব হয় দেখতে হবে, প্যান পায়খানায় পরিপূর্ন থাকলে দেখা নাও যেতে পারে। পরিপূর্ণ থাকলে ল্যাট্রিনে পানি ঢেলে দেখতে হবে। যদি উত্তর হ্যাঁ হয় তাহলে পরবর্তী প্রশ্নে চলে যান) [if it is possible to watch, if latrine is overflow it may not be visible. If the answer is ‘yes’ proceed]

52. ওয়াটার সিল/সাইফুন এর বর্তমান অবস্থা কি [What is the current condition of the water seal]?

1. ভাল [Functional]    2.ভাঙ্গা [Broken]

53. যদি ভাঙ্গা হয়, তা বর্ণনা করুন (এর মধ্যে পানি নেই, এর মধ্য দিয়ে পিটের পায়খানা সরাসরি দেখা যাচ্ছে, গর্ত দেখা যাচ্ছে) [If broken, describe it (no water in it, pit content appears through the water seal, big hollow appears etc.)]? . . . . . . . . . . . . . . . . . . . . . . . . . . . . . . . . . . . . . . . . . . . . . . . . . . . . . . . . . . . . . . . . . . . . . . . . . . . . . . . . . . . . . . . . . . . . . . . . . . . . . . . . . . . . . . . . . . . . . . . . . . . . . . . . . . . . . . . . . . . . . . . . . . . . . . . . . . .

**Interviewer instruction sheet**

১. যদি কোন খানায় ল্যাট্রিন না থাকে তাকে নতুন ল্যাট্রিনের জন্য বিবেচিত হবে (**নতুন ল্যাট্রিন হচ্ছে উপরিকাঠাম সহ সমস্ৰ উপাদান নতুন** )।

২. যদি ল্যাট্রিনের উপরি কাঠামোর শুধু ছাদ/দরজা/বেড়া না থাকে তাহলে তা মেড়ামত করে দেয়া জন্য বিবেচিত হবে।

৩. যদি ল্যাট্রিনের উপরি কাঠামো নস্ট হয়ে যায় (ব্যবহারের অনুপযগি, একই সাথে উপর দিয়ে পানি পরে, দরজা ভাঙ্গা, বেড়া ভাঙ্গা) তাহলে নতুন উপরি কাঠামো দেয়া হবে জন্য বিবেচিত হবে।

৪. কোন ল্যাট্রিনের একটা পিট/গর্ত থাকলে তা ডুয়েল পিট ল্যাট্রিনে রূপান্ৰর করে দেয়া জন্য বিবেচিত হবে (যদি প্রথম পিট/বর্তমান পিট ভাল থাকে)।

৫. কোন ল্যট্রিনের বর্তমান পিট (রিং ও স্লাব ভাঙ্গা/ রিং ভাঙ্গা এবং পায়খান বাইরে পরছে) তাহলে তাকে নতুন ল্যাট্রিন দেয়া জন্য বিবেচিত হবে।

৬. যদি কোন ল্যাট্রিনে ৩ টার কম (১ বা ২ টা) রিং থাকে তাহলে নতুন ল্যাট্রিনের জন্য বিবেচিত হবে।

৭. সব ল্যাট্রিনের সাইফুন দেয়ার জন্য বিবেচনা করা হবে।

PROTOCOL
Biomedical
Non-Exempt
Berkeley

-----------------------------------------------------------------------------------
---

**Protocol Title:** Measuring the benefits of sanitation, water quality, handwashing and nutrition interventions for improving health and development in rural Bangladesh

**Protocol Status:** APPROVED

**Date Submitted:** 10/26/2017

**Approval Period:** **10/30/2017-11/03/2018**

**Important Note:** **This Print View may not reflect all comments and contingencies for approval. Please check the comments section of the online protocol.**
**Questions that appear to not have been answered may not have been required for this submission. Please see the system application for more details.**

*** * * Attached Document * * ***

| Document Name | Created Date |
| --- | --- |
| Ercumen_CITI Completion_2008.pdf | 10/26/2017 |

**CITI Collaborative Institutional Training Initiative**

#### Human Research Curriculum Completion Report
#### Printed on Friday, March 7, 2008

**Learner:** Ayse Ercumen (username: aercumen)
**Institution:** University of California, Berkeley
**Contact Information** 2299 Piedmont Avenue, #485
Berkeley, CA 94720 USA
Department: School of Public Health


**Group 1:** Biomedical Research Investigators and Key Personnel

**Stage 1. Basic Course Passed on 03/07/08** (Ref # 1661285)

| Required Modules | Date Completed | Score |
|---|---|---|
| Introduction | 03/07/08 | no quiz |
| History and Ethical Principles | 03/07/08 | 5/5 (100%) |
| Basic Institutional Review Board (IRB) Regulations and Review Process | 03/07/08 | 5/5 (100%) |
| Informed Consent | 03/07/08 | 4/4 (100%) |
| Social and Behavioral Research for Biomedical Researchers | 03/07/08 | 4/4 (100%) |
| Records-Based Research | 03/07/08 | 2/2 (100%) |
| Genetic Research in Human Populations | 03/07/08 | 2/2 (100%) |
| Research With Protected Populations - Vulnerable Subjects: An Overview | 03/07/08 | 4/4 (100%) |
| Vulnerable Subjects - Research with Prisoners | 03/07/08 | 4/4 (100%) |
| Vulnerable Subjects - Research Involving Minors | 03/07/08 | 3/3 (100%) |
| Vulnerable Subjects - Research Involving Pregnant Women and Fetuses in Utero | 03/07/08 | 3/3 (100%) |
| International Research | 03/07/08 | no quiz |
| Group Harms: Research With Culturally or Medically Vulnerable Groups | 03/07/08 | 3/3 (100%) |
| FDA-Regulated Research | 03/07/08 | 5/5 (100%) |
| Human Subjects Research at the VA | 03/07/08 | 3/3 (100%) |
| HIPAA and Human Subjects Research | 03/07/08 | 1/2 (50%) |
| Workers as Research Subjects-A Vulnerable Population | 03/07/08 | 4/4 (100%) |
| Hot Topics | 03/07/08 | no quiz |
| Conflicts of Interest in Research Involving Human Subjects | 03/07/08 | 2/2 (100%) |
| University of California, Berkeley | 03/07/08 | no quiz |

**For this Completion Report to be valid, the learner listed above must be affiliated with a CITI participating institution. Falsified information and unauthorized use of the CITI course site is unethical, and may be considered scientific misconduct by your institution.**

Paul Braunschweiger Ph.D.
Professor, University of Miami
Director Office of Research Education
CITI Course Coordinator

Return

PROTOCOL
Biomedical
Non-Exempt
Berkeley

-----------------------------------------------------------------------------------
---

**Protocol Title:** Measuring the benefits of sanitation, water quality, handwashing and nutrition interventions for improving health and development in rural Bangladesh

**Protocol Status:** APPROVED

**Date Submitted:** 10/26/2017

**Approval Period:** **10/30/2017-11/03/2018**

**Important Note:** **This Print View may not reflect all comments and contingencies for approval. Please check the comments section of the online protocol. Questions that appear to not have been answered may not have been required for this submission. Please see the system application for more details.**

*** * * Attached Document * * ***

| Document Name | Created Date |
|---|---|
| KNelson CITI Human Subjects completion report.pdf | 10/26/2017 |

### COLLABORATIVE INSTITUTIONAL TRAINING INITIATIVE (CITI)

#### HUMAN RESEARCH CURRICULUM COMPLETION REPORT

**Printed on 11/19/2013**

| | |
|---|---|
| **LEARNER** | Kara Nelson (ID: 3875445)<br>CEE Dept MS 1710<br>University of California<br>Berkeley<br>CA 94720-1710<br>USA |
| **DEPARTMENT** | Civil and Environmental Engineering |
| **PHONE** | 5106435023 |
| **EMAIL** | |
| **INSTITUTION** | University of California, Berkeley |
| **EXPIRATION DATE** | 11/19/2015 |

**GROUP 1 BIOMEDICAL RESEARCH INVESTIGATORS AND KEY PERSONNEL**

| | |
|---|---|
| **COURSE/STAGE**: | Basic Course/1 |
| **PASSED ON**: | 11/19/2013 |
| **REFERENCE ID**: | 11787325 |

| REQUIRED MODULES | DATE COMPLETED | SCORE |
|---|---|---|
| Belmont Report and CITI Course Introduction | 11/19/13 | 3/3 (100%) |
| History and Ethical Principles | 11/19/13 | 6/6 (100%) |
| Basic Institutional Review Board (IRB) Regulations and Review Process | 11/19/13 | 4/5 (80%) |
| Informed Consent | 11/19/13 | 3/4 (75%) |
| Social and Behavioral Research (SBR) for Biomedical Researchers | 11/19/13 | 3/4 (75%) |
| Records-Based Research | 11/19/13 | 1/2 (50%) |
| Genetic Research in Human Populations | 11/19/13 | 2/2 (100%) |
| Research With Protected Populations - Vulnerable Subjects: An Overview | 11/19/13 | 4/4 (100%) |
| Vulnerable Subjects - Research Involving Prisoners | 11/19/13 | 4/4 (100%) |
| Vulnerable Subjects - Research Involving Children | 11/19/13 | 3/3 (100%) |
| Vulnerable Subjects - Research Involving Pregnant Women, Human Fetuses, and Neonates | 11/19/13 | 3/3 (100%) |
| International Studies | 11/19/13 | 3/3 (100%) |
| FDA-Regulated Research | 11/19/13 | 4/5 (80%) |
| Research and HIPAA Privacy Protections | 11/19/13 | 5/5 (100%) |
| Vulnerable Subjects - Research Involving Workers/Employees | 11/19/13 | 4/4 (100%) |
| Hot Topics | 11/19/13 | No Quiz |
| Conflicts of Interest in Research Involving Human Subjects | 11/19/13 | 4/5 (80%) |
| Avoiding Group Harms - U.S. Research Perspectives | 11/19/13 | 3/3 (100%) |
| Unanticipated Problems and Reporting Requirements in Biomedical Research | 11/19/13 | 4/6 (67%) |
| University of California, Berkeley | 11/19/13 | No Quiz |
| Stem Cell Research Oversight (Part I) | 11/19/13 | 3/5 (60%) |

**For this Completion Report to be valid, the learner listed above must be affiliated with a CITI Program participating institution or be a paid Independent Learner. Falsified information and unauthorized use of the CITI Progam course site is unethical, and may be considered research misconduct by your institution.**

Paul Braunschweiger Ph.D.
Professor, University of Miami
Director Office of Research Education
CITI Program Course Coordinator

CITI Collaborative Institutional Training Initiative at the University of Miami

PROTOCOL
Biomedical
Non-Exempt
Berkeley

Protocol # 2011-09-3652
Date Printed: 07/31/2018

-----------------------------------------------------------------------------------
---

**Protocol Title:** Measuring the benefits of sanitation, water quality, handwashing and nutrition interventions for improving health and development in rural Bangladesh

**Protocol Status:** APPROVED

**Date Submitted:** 10/26/2017

**Approval Period:** **10/30/2017-11/03/2018**

**Important Note:** This Print View may not reflect all comments and contingencies for approval. Please check the comments section of the online protocol.
Questions that appear to not have been answered may not have been required for this submission. Please see the system application for more details.

*** * * Attached Document * * ***

| Document Name | Created Date |
| --- | --- |
| Protocol PR # 11063_ERC approval_addendum_17 NOV.pdf | 10/26/2017 |

icddr,b

### Memorandum

17 November 2013

To: Dr Leanne Unicomb
Principal Investigator of research protocol # PR-11063
Centre for Communicable Diseases (CCD)

From: Professor Kazi Zulfiquer Mamun
Chairperson
Ethical Review Committee (ERC)

Sub: Approval of an addendum to research protocol # PR-11063

Thank you for your memo dated 10 November 2013 attaching the modified version of your addendum proposal to previously approved research protocol # PR-11063 titled "**A randomized controlled trial of the benefits of water sanitation hygiene plus nutrition interventions on child growth** through expedited review process. I have the pleasure to inform you that the addendum proposal to the above research protocol is approved through expedited review mechanism.

Other terms and conditions for implementation of your research protocol, as contained in our memo dated 9 April 2012 according approval of the research protocol shall, however, remain unchanged.

Thank you.

Cc: Director, CCD

GPO Box 128, Dhaka 1000, Bangladesh.
**For Packages :** 68, Shaheed Tajuddin Ahmed Sarani, Mohakhali, Dhaka 1212, Bangladesh.

PROTOCOL
Biomedical
Non-Exempt
Berkeley

Protocol # 2011-09-3652
Date Printed: 07/31/2018

-------------------------------------------------------------------------------------

**Protocol Title:** Measuring the benefits of sanitation, water quality, handwashing and nutrition interventions for improving health and development in rural Bangladesh

**Protocol Status:** APPROVED

**Date Submitted:** 10/26/2017

**Approval Period:** **10/30/2017-11/03/2018**

**Important Note:** **This Print View may not reflect all comments and contingencies for approval. Please check the comments section of the online protocol. Questions that appear to not have been answered may not have been required for this submission. Please see the system application for more details.**

*** * * Attached Document * * ***

| Document Name | Created Date |
|---|---|
| wash benefits endline spillover compounds instrument_v2.pdf | 10/26/2017 |

ICDDR,B

### Wash Benefits

Endline Instruments for Compounds Enrolled in Spillover Substudy

1/30/2014

**This questionnaire will be administered to the compounds with children 0-<60 months closest to compounds enrolled in the combined intervention arm and control arm.**

খানার সম্মতি**Consent (All households)**

নিম্নলিখিত বিষয়গুলো জরুরী FOLLOW THESE STEPS CAREFULLY:

1. এই বাড়িতে পাঁচ বছরের নিচে শিশু আছে এবং এই বাড়িতে ওয়াশ বেনিফিটের অন্তর্ভূক্ত বাড়ির সবচেয়ে কাছের বাড়ি, এই বিষয় নিশ্চিত হয়ে এই বাড়ির অন্তর্ভূক্ততার যোগ্যতা যাচাই করুন।VERIFY ELIGIBILITY BY CONFIRMING THAT THE COMPOUND IS THE CLOSEST COMPOUND WITH A CHILD UNDER FIVE YEARS TO A COMPOUND ENROLLED IN WASH BENEFITS.
2. যদি উত্তরদাতা আমাদের সমস্ত শর্তাদি পূরণ করতে পারে তাহলে খানা আইডি খানার সম্মতি পত্রের নিদ্দিষ্ট অংশে রেকর্ড করুন IF RESPONDENT MEETS OUR CRITERIA, CAREFULLY RECORD **HOUSEHOLD ID** AT THE TOP OF THE CONSENT PAGE.
3. তালিকায় দেয়া আইডি এবং নির্ধারিত কাজের জন্য বাড়ী/কম্পাউন্ড /খানার আইডি মিলিয়ে দেখুন এবং নিদ্দিষ্ট করুন CONFIRM THAT THE ID EXACTLY MATCHES THE ID ASSIGNED TO THIS COMPOUND/HOUSEHOLD BY THE FIELD RESEARCH OFFICER.

লিখিত সম্মতিপত্র পড়ার সময়ে সম্মতিপত্রটি মৌখিকভাবে উত্তরদাতাকে /বোঝাতে হবে এবং তার সম্মতিতে খানার আইডি বসাতে হবে READ THE CONSENT FORM VERBATIM TO THE RESPONDENT IN THE LANGUAGE OF THE WRITTEN CONSENT FORM AND THEN CONFIRM THAT SHE UNDERSTANDS.

খানার আইডি নং ENTER THE HOUSEHOLD ID: |__|__|__|__|__||__|__|__|__|__||__|__|__|.

খানার আইডি নং ENTER THE HOUSEHOLD ID OF THE COMPOUND ENROLLED IN THE MAIN STUDY WHICH THIS COMPOUND IS ADJACENT TO: |__|__|__|__|__||__|__|__|__|__||__|__|__|.

সম্মতি পত্র পড়ুন **Read**: ওহে। আমার নাম ----{নাম বলুন}|আমি ঢাকা মহাখলী কলেরা হাসপাতাল(আইসিডিডিআরবি) থেকে এসেছি। আজ আমরা আপনার খানায় গবেষণা বিষয়ে কথা বলতে এসেছি যেহেতু আপনার খানাটি এই ষ্টাডীর জন্য নির্বাচিত হয়েছে ( যেহেতু আপনার বাড়িতে পাঁচ বছরের নিচে শিশু আছে |Hello. My name is [name]. I am from Mohakhali Cholera Hospital (icddr,b) in Dhaka. We are here to talk to you today because your household has been selected for our study because there are children under five years who live in your household. READ REST OF CONSENT.

##### ভৌগলিক/অবস্থান পরিচিতি Geographical Identification

নিচে উত্তরদাতার নিঁদিষ্টকরন তথ্য গুরত্ত্বের সাথে পূরন করুন FILL THE FOLLOWING IDENTIFICATION INFORMATION FOR THE RESPONDENT VERY CAREFULLY.

| Module ID | WBM 40 | |
|---|---|---|
| | 4001. | এফ.আর.এ কোড (FRA code): ÿÿ |
| C.101 | 4002. | তথ্য সংগ্রহের তারিখ [Date of data collection]: __/__/____ |
| | dataid | \|__\|__\|__\|__\|__\| |
| | 4003. | |
| | 4004. | |
| | 4005. | |
| | 4006. | জেলার নাম(DISTRICT NAME): |
| | 4007. | উপজেলার নাম (UPAZILA NAME): |
| | 4008. | ইউনিয়নের নাম [Union name ____________ |
| | 4009. | খানার ঠিকানা Household Address: |
| | 4010. | খানা প্রধানের নাম [Name of Household head]__________________________ |
| | 4011. | খানা প্রধানের বাবা/স্বামীর নাম [Father's/Husband's Name of HH head] |

| | | |
|---|---|---|
| | | _______________ |
| | 4012. | বাড়ির নাম [Bari Name]: ________________________________________ |
| | 4013. | গ্রামের নাম [Village Name]: _____________________________________ |
| | 4014. | |
| | 4015. | বাড়ী নাম্বার [Bari No. [1-8] |
| | 4016. | **HH ID:** [1-7] |
| | 4017. | |
| | 4018. | |
| | 4019. | |
| | 4019.a | **\|_____\| এই বাড়ীতে ৩৬ মাসের নীচে কতজন শিশু আছে?** How many children do you have < 60 months in this bari? |

**Contact information**

| | | |
|---|---|---|
| | 001 | উত্তরদাতার/ উত্তরদাতার খানার কোন সদস্যের কী কোন ফোন নাম্বার আছে?(Is there a phone available?) 1 Yes 0 No (Skip to 005) |
| C.11 | **002** | -----------প্রাথমিক ফোন নাম্বার (Primary phone number) |
| C.12 | 003 | এই ফোনে কি আপন াকে সরাসরি পাওয়া যাবে? (Does this phone number belong to you?) 1 Yes(Skip to 005) 0 No |
| C.13 | 004 | যদি আপন াকে পাওয়া না যায় তাহলে এই নাম্বারটি কার? IF NO, to whom does this number belong? (Specify relationship to respondent)<br>1 . স্বামী (HUSBAND)<br>2. শ্বশুর IN LAWS<br>3.পিতা(FATHER)<br>4.মা(MOTHER)<br>5.পরিবারের সদস্য(FAMILY MEMBER)<br>99.অন্যান্য (নাম ও উত্তরদাতার সাথে সম্পর্ক লিখুন)(OTHER) |
| | 004.a | দ্বিতীয় কোন ফোন নাম্বার আছে কী?(Is there a phone available?)<br>1 Yes 0 No (Skip to 008) |
| C.14 | 005 | -----------দ্বিতীয় ফোন নাম্বার Secondary phone number |
| C.15 | 006 | এই ফোনে কি আপন াকে সরাসরি পাওয়া যাবে? Does this phone number belong to you? 1 Yes(Skip to 008) 0 No |
| C.16 | 007 | যদি আপন াকে পাওয়া না যায় তাহলে এই নাম্বারটি কার? IF NO, to whom does this number belong?<br>1. স্বামী (HUSBAND)<br>2. শ্বশুর IN LAWS<br>3. পিতা(FATHER)<br>4 .মা(MOTHER)<br>5 .পরিবারের সদস্য(FAMILY MEMBER)<br>99 .অন্যান্য(নাম ও উত্তরদাতার সাথে সম্পর্ক লিখুন) (OTHER) |
| C.17 | 008 | এই খানাটির আশেপাশে কোন উলেস্নখযোগ্য কিছু আছে কি? (Location/landmark) |
| | 009 | **\|_____\| \|_____\| সবচেয়ে ছোট বাচ্চার মা কত ক্লাস পর্যন্ত পড়ালেখা শেষ করেছেন ? (জানি না = *99*)** Education of mother of the youngest child *(Years of education completed, DK=99)* |
| | 010 | **\|_____\| \|_____\| সবচেয়ে ছোট বাচ্চার বাবা কত ক্লাস পর্যন্ত পড়ালেখা শেষ করেছেন ? (জানি না** |

| | | |
|---|---|---|
| | | = ***99***) Education of father of the youngest child *(Years of education completed, DK=99)* |
| | 011 | **I_____I I_____I** সবচেয়ে ছোট বাচ্চার বাবার প্রধান পেশা [Main occupation of father of the youngest child]<br>পেশা কোড Occupation Code:<br>1. কৃষক [Farmer/Cultivator]<br>2. ঘরে কাজ করে [Homemaker]<br>3. কৃষি শ্রমিক [Agri-labor]<br>4. শ্রমিক (কৃষি শ্রমিক ছাড়া) [Non-agri labor]<br>5. বেতনভুক্ত কর্মচারী (সরকারী/প্রাইভেট/এন.জি.ও) [Salaried job (Govt./Private/NGO)]<br>6. রাজমিস্ত্রী [Mason (*Rajmistri*)]<br>7. কাঠ মিস্ত্রী [Carpenter]<br>8. ভ্যান/রিকশা চালক [Van/Rickshaw puller]<br>9. জেলে [Fisherman]<br>10. নৌকা চালক/মাঝি [Boatman]<br>11. কর্মকার [Blacksmith]<br>12. স্বর্ণকার [Goldsmith]<br>13. কুমার/কুম্ভকার [Potter (soil smith)]<br>14. মুচি [Shoe polish /maker]<br>15. দোকানদার [Shopkeeper]<br>16. ফেরিওয়ালা [Vendor (*Feriwala/howker*)]<br>17. ক্ষুদ্র ব্যবসায়ী (মূলধন <=১০০০০) [Petty trader, capital <=10000]<br>18. ব্যবসায়ী(মূলধন >১০০০০) [Business, capital >10000]<br>19. দর্জি [Tailor]<br>20. ড্রাইভার [Driver]<br>21. কুটীর শিল্প [Cottage industry]<br>22. পোলট্রি/ ব্যবসার জন্য পশু লালন-পালনকারী [Poultry /livestock rearer]<br>23. বৈদ্যুতিক মিস্ত্রী [Electrician]<br>24. হোমিওপ্যাথি ডাক্তার [Homeopath]<br>25. আধ্যাতিক চিকিৎসক/ কবিরাজ/ ওঝাঁ [Spiritual healer/kabiraj/ Ojha]<br>26. পেশাদার ডাক্তার/উকিল [Professional practitioner (Doctor/lawyer)]<br>27. ইমাম/ ধর্মযাজক [Imam/priest]<br>28. অবসরপ্রাপ্ত চাকুরীজীবী [Retired service holder]<br>29. ছাত্র [Student]<br>30. বেকার [Unemployed]<br>31. অক্ষম [Disabled]<br>32. কাজের লোক [Domestic maid / servant]<br>33. জমিদার (শস্য উৎপাদন অথবা অন্য কোন কাজে কৃষকদের জমি বর্গা দেয়) [Landlord (Provide land for farmers for sharecropping or others)]<br>34. বিদেশে থাকে [Staying abroad]<br>35. মৃত/নিখোঁজ [Died/untraced]<br>99. জানিনা [Don't know]<br>77. অন্যান্য (বর্ণনা লিখুন) [Others (specify)] |
| | 012 | **I_____I I_____I** আপনার খানাতে/পরিবারে কত জন লোক প্রতিদিন একই হাড়িতে রান্না করে খান?How many people in this house eat from the same cooking pot every day (খানা বলতে একই হাড়িতে রান্না করে খায় এমন সদস্যদের বুঝানো হয়েছে) |
| | 013 | **I_____I** আপনার খানাতে ৩৬ মাসের নীচে কতজন শিশু আছে? How many children do you have < 60 months do you have? |
| | 014 | **I____I** এই বাড়ীতে ৩৬ মাসের নীচে কতজন শিশু আছে? How many children do you have < 60 months in this bari? |
| | 014 | **I_____I**এই বাড়ীতে কতগুলো খানা আছে? How many HHs do you have in this bari? |

| A. এই বাড়ীর খানাগুলোর সেনসাস (সকল খানার) CENSUS OF HOUSEHOLDS ARE IN THIS COMPOUND (All households) | | | | | | | |
|---|---|---|---|---|---|---|---|
| পড়ে শোনান READ | ঠিক আছে। আমি এখন আপনাকে সকল খানার সম্পর্কে আলাদাভাবে কিছু প্রশ্ন করব। প্রথমে আপনার খানা সম্পর্কে বলুন। Okay. Now I am going to ask you about each household separately. First, think about your own household.<br>MARK "99" IF RESPONDENT DOESN'T KNOW THE ANSWER TO ANY OF THE FOLLOWING QUESTIONS. Record the GPS location of each household. | | | | | | |
| | A.1 | A.2 | A.3 | A.4 | A.5 | A.6 | A.7 |
| খানার নাম্বার HH No. | খানা প্রধানের নাম কি? What is the name of the household head? | এই খানাতে ৪০ বছরের উপরের বয়সের কতজন সদস্য আছে? How many adults | এই খানাতে (>১৮-৪০) বছরের বয়সের কতজন সদস্য আছে? How many young | এই খানাতে (>৩-১৮) বছরের বয়সের কতজন সদস্য আছে?How many | এই খানাতে (০-৩) বছরের বয়সের কতজন সদস্য আছে?How many young | প্রতিটি খানায় কতজন সদস্য আছে তা গণণা করুন এবং নিশ্চিত হোন CALCULATE THE TOTAL | GPS coordinates |

| | FOR REFERENCE ONLY | over 40 years live in this household? | adults (>18-40 years) live in this household? | school-aged children (>3-18 years) live in this household? | children (0-3 years) live in this household? | NUMBER OF PEOPLE IN EACH HH AND CONFIRM |
|---|---|---|---|---|---|---|
| 1 | | | | | | |
| 2 | | | | | | |

Information about children under five years. Autopopulate the number of rows in the following table based on the response to 4019.a

| | | HH. no. | নাম (Name) | লিঙ্গ (Sex) | জন্ম তারিখ(Birth date) | জন্ম তারিখেরউৎস(Source of DOB )(C.104) |
|---|---|---|---|---|---|---|
| **C.06.** | 4020 | | | 1 পুরম্নষ M<br>0 মহিলা F | দিন/মাস/বছর<br>DD/MM/YY | 1 = কার্ড দ্বারা নিশ্চিত হয়েছিল (Confirmed DOB by valid vaccination/health card)<br>2 = মা/আত্মীয় দ্বারা নিশ্চিতহয়েছিল (Mother/Relative remembers DOB)<br>3 = উভয় (Both 1 & 2)<br>4 = ঘটনা দ্বারা নিশ্চিতহয়েছিল (Estimated DOB with 2 and event calendar) |
| **C.07.** | 4021 | | | 1 পুরম্নষ M<br>0 মহিলা F | দিন/মাস/বছর<br>DD/MM/YYY | 1 = কার্ড দ্বারা নিশ্চিত হয়েছিল (Confirmed DOB by valid vaccination/health card)<br>2 = মা/আত্মীয় দ্বারা নিশ্চিত হয়েছিল (Mother/Relative remembers DOB)<br>3 = উভয় (Both 1 & 2)<br>4 = ঘটনা দ্বারা নিশ্চিতহয়েছিল (Estimated DOB with 2 and event calendar) |
| **C.08.** | 4022 | | | 1 পুরম্নষ M<br>0 মহিলা F | দিন/মাস/বছর<br>DD/MM/YYY | 1 = কার্ড দ্বারা নিশ্চিত হয়েছিল (Confirmed DOB by valid vaccination/health card)<br>2 = মা/আত্মীয় দ্বারা নিশ্চিত হয়েছিল (Mother/Relative remembers DOB)<br>3 = উভয় (Both 1 & 2)<br>4 = ঘটনা দ্বারা নিশ্চিত হয়েছিল (Estimated DOB with 2 and event calendar) |
| **C.09.** | 4023 | | | 1 পুরম্নষ M<br>0 মহিলা F | দিন/মাস/বছর<br>DD/MM/YYY | 1 = কার্ড দ্বারা নিশ্চিত হয়েছিল (Confirmed DOB by valid vaccination/health card)<br>2 = মা/আত্মীয় দ্বারা নিশ্চিত হয়েছিল (Mother/Relative remembers DOB)<br>3 = উভয় (Both 1 & 2)<br>4 = ঘটনা দ্বারা নিশ্চিতহয়েছিল (Estimated DOB with 2 and event calendar) |

| | | | | | | |
|---|---|---|---|---|---|---|
| **C.10.** | 4024 | | | 1 পুরুষ M<br>0 মহিলা F | দিন/মাস/বছর<br>DD/MM/YYY | 1 = কার্ড দ্বারা নিশ্চিত হয়েছিল (Confirmed DOB by valid vaccination/health card)<br>2 = মা/আত্মীয় দ্বারা নিশ্চিতহয়েছিল (Mother/Relative remembers DOB)<br>3 = উভয় (Both 1 & 2)<br>4 = ঘটনা দ্বারা নিশ্চিতহয়েছিল (Estimated DOB with 2 and event calendar) |

**Wash Benefit Module 1RESPONDENT IDENTIFICATION**

| | | |
|---|---|---|
| মডিউল আইডিModule ID | WBM 1 | |
| | 101. | : What is your full name? প্রধান উত্তরদাতার নাম [Name of respondent: |
| | 102. | : \|_____\| প্রধান উত্তরদাতার পরিচয়[Status of main respondent]<br><br>1. প্রধান পরির্চযাকারী /মা [Mother of child closest to 2 years old]<br>2. পরির্চযাকারী (পুরুষ) [Male caregiver]<br>3. পরির্চযাকারী (মহিলা) [Female caregiver] |
| | 103. | প্রধান উত্তরদাতার বয়স (বছরে)[Age of main respondent: *(in years)]* |

**Wash Benefit Module 3 DEWORMING**

**(**Administer to: Children < 60 months. This may include the child in the household or in other households in the bari.)

**C.301.**

☐ Within the last six months, has [NAME] received a pill or drug for intestinal worms?

1 Yes
2 No **(Done)**
99 Don't know / not sure **(Done)**

**C.302.**

☐ Where did [NAME] receive the drug for intestinal worms?

1 At home / in the village
2 At a hospital or health facility
3 At a school
99 Don't know / not sure

**C.303.**

☐ Did [NAME] receive the drug as part of a large campaign?

1 Yes

2 No

99 Don't know / not sure

**C.304.**

Approximately how long ago did [NAME] receive the drug?

*If more recent than 1 month, record weeks*

99 Don't know / not sure

A ☐ MONTHS (0 – 6)

B ☐☐ WEEKS

**C.305.**

Has the [NAME] intentionally eaten any dirt or soil?

*Ask for each recall period:*

| | | |
|---|---|---|
| 1 | Today | ☐ 1 Yes ☐ 2 No ☐ 99 Don't know |
| 2 | Yesterday | ☐ 1 Yes ☐ 2 No ☐ 99 Don't know |
| 3 | Day before yesterday | ☐ 1 Yes ☐ 2 No ☐ 99 Don't know |
| 4 | In the past 7 days (since this day last week?) | ☐ 1 Yes ☐ 2 No ☐ 99 Don't know |

| |
|---|
| Administer to:<br>গর্ভবতী মা Pregnant mothers **[PDA to open MOTHERS COHORT]**<br>৩৬ মাসের নীচের শিশু Children < 36 months at enrollment **[PDA to open DIARRHEA C(** |

**Wash Benefit Module 2DIARRHEA AND ILLNESS SYMPTOMS (Children < 60 months)**

৩৬ মাসের নীচের যে সকল শিশু খানায়/বাড়ীতে বাস করে (বেইজ লাইলের জন্য) । Administer to: Children < 60 mo household or in other households in the bari.

PDA TO OPEN CHILD ID FROM **DIARRHEA COHORT**

| উত্তরদাতা শিশুর প্রধান পরিচর্যাকারী Respondent: Child's primary caregiver | | | | |
|---|---|---|---|---|
| | WBM 2 | আপনাকে ধন্যবাদ । আমি এখন আপনাকে আপনার শিশুর স্বাস্থ্য বিষয়ে কিছু প্রশ্ন জিজ্ঞাসা করব। দয়া ক জবাব দিবেন। যদি উত্তর না জানা থাকে তখন বলবেন জানি না। শিশুটির নাম ধরে শুরু ক ask you some questions about the health of your [child/children]. Please answer you don't know the answer to a question, say "I don't know." We will begin with | | |
| | | | **A** | **B** |
| | | **শিশুটির নাম বলুন এবং তার লক্ষণ গুলো ছিল কী?Did [NAME] have [SYMPTOM] :** | আজ<br>**Today**<br>১=হ্যাঁ YES<br>০=না NO<br>৯৯৯=জানি না DK | গতকাল<br>**Yesterday**<br>১=হ্যাঁ YES<br>০=না NO<br>৯৯৯=জানি না DK |
| **C.201** | 201. | জ্বর Fever | | |
| **C.202** | 202. | পাতলা পায়খানা Diarrhea | | |
| **C.203** | 203. | ২৪ ঘন্টায় ৩ বার বা তার বেশী পায়খানা করেছে কিনা? 3 or more bowel movements in 24 hours | | |
| **C.204** | 204. | দৈনিক পায়খানায় সংখ্যা Number of bowl movements each day | | |
| **C.205** | 205. | পানির মত বা নরম পায়খানা? Watery or soft stool (unformed) | | |
| **C.206** | 206. | পায়খানায় রক্ত Blood in the stool | | |
| **C.207** | 207. | শরীর বা মাথায় চুলকানি, রেশ?Skin rash (anywhere on the body) | | |
| **C.208** | 208. | দীর্ঘ কাশি Constant cough | | |
| **C.209** | 209. | নাক বন্ধ/নাক দিয়ে পানি পড়া?Congestion / runny nose | | |
| **C.210** | 210. | শ্বাস নেওয়ার সময় শোঁ শোঁ করে আওয়াজ বা শ্বাসকষ্ট? (যে কোন ধরনের শ্বাসকষ্ট, তবে নাক বন্ধ থাকার কারণে শ্বাসকষ্ট হলে তা বাদ দিতে হবে) Panting / wheezing / difficulty breathing 9 | | |
| **C.211** | 211. | দাগ, থেঁৎলে যাওয়া বা কাটা Bruising, scrapes or cuts | | |
| **C.212** | 212. | দাঁতের ব্যাথাToothache / teething | | |

**C.601.**

শিশু কি কখনো ময়লা বা মাটি খিয়েছে?Has the child eaten any dirt or soil?

মনে করার জন্য প্রতিটি সময় আলাদা ভাবে জানতে চান: *Ask for each recall period:*

| | | | | |
|---|---|---|---|---|
| 1 | আজ Today | ☐ 1 Yes | ☐ 2 No | ☐ 99 Don't know |
| 2 | গতকাল Yesterday | ☐ 1 Yes | ☐ 2 No | ☐ 99 Don't know |
| 3 | গত কালের আগের দিন Day before yesterday | ☐ 1 Yes | ☐ 2 No | ☐ 99 Don't know |
| 4 | গত সাত দিন (আজকের দিন থেকে গত এক সপ্তাহ?) In the past 7 days (since this day last week?) | ☐ 1 Yes | ☐ 2 No | ☐ 99 Don't know |

**Wash Benefit Module 7 HANDWASHING**

| | WBM 7 | | |
|---|---|---|---|
| C.700 | 70 | এই প্রশ্নের আগে উত্তরদাতা যে কোন সময় তার হাত ধুয়েছিল কিনা তা পর্যবেক্ষণ করুন।RECORD WHETHER THE RESPONDENT HAS WASHED HER HANDS AT ANY TIME BEFORE THIS QUESTION | [1] উত্তরদাতা হাত ধুয়েছিলOBSERVED RESPONDENT WASHING HANDS<br>[2] উত্তরদাতা হাতধোয় নাইDID NOT OBSERVE RESPONDENT WASHING HANDS |
| C.701 | 70 | আপনাকে ধন্যবাদ । এখন আমি আপনার হাতগুলো দেখতে চাই। আশা করি আপনি কিছু মনে করবেন না। আপনি কি আমাকে দেখাবেন? READ: Thank you. Now, I would like to do a quick inspection of your hands. I hope you don't mind. Can you please show me your hands?<br><br>দুই হাতই দেখতে হবে(প্রথমে এক হাত)। এরপর নিচের উলেস্নখিত হাতের অবস্থার কোড দেখে রেকর্ড করতে হবে। BOTH HANDS SHOULD BE SHOWN (NOT JUST ONE HAND). USE <u>APPEARANCE CODES</u> BELOW TO RECORD THE DESCRIPTION THAT BEST DESCRIBES THE DEFINITIONS THE LEVEL OF CLEANLINESS.<br><u>APPEARANCE CODES:</u><br>[1] ময়লা স্পষ্টভাবে দেখা যাচ্ছিল VISIBLE DIRT (DIRT/MUD/SOIL/ASH OR ANY OTHER MATERIAL IS VISIBLE)<br>[2] ময়লা স্পষ্টভাবে দেখা নাগেলেও অপরিচ্ছন্নভাব ছিলUNCLEAN APPEARANCE (NO DIRT IS VISIBLE ON THIS PART OF THE HAND BUT, IN GENERAL, THIS PART OF THE HAND APPEARS | 👁 MOTHER<br><br>**বাম হাত Left Hand**<br>A\|__\| হাতের নখ FINGERNAILS<br>B\|__\| হাতের তালু PALMS<br>C\|__\| আঙ্গুলের সম্মুখভাগ FINGER PADS<br><br>**ডান হাত Right Hand**<br>D\|__\| হাতের নখ FINGERNAILS<br>E\|__\| হাতের তালু PALMS<br>F\|__\| আঙ্গুলের সম্মুখভাগ FINGER PADS |

| | | | |
|---|---|---|---|
| | | UNCLEAN)<br>[3] পরিষ্কার ছিল CLEAN (OBSERVED PART OF THE HAND IS CLEAN AS WOULD APPEAR AFTER SOMEONE WASHES HANDS OR TAKES A BATH)<br>৮৮. পর্যবেক্ষণ করা সম্ভব হয়নি/প্রত্যাখান N/A; OBSERVATION NOT POSSIBLE/REFUSED | |
| **C.701** | 70 | ASK: দয়া করে আমাকে(শিশুর নাম ধরে) হাতগুলো দেখাও । Please show me [NAME]'s hands.<br><br>উত্তরদাতার সবচেয়ে বড় বাচ্চার (০-৩৬ মাস) হাত দেখুন । এই বয়সের বাচ্চা না থাকলে স্কিপ করু। TO SEE THE HANDS OF THE RESPONDENT'S OLDEST CHILD 0-36 MONTHS (SEE C.1) IF RESPONDENT DOESN'T HAVE A CHILD 0-36 MONTHS → **SKIP TO NEXT PAGE**.<br><br>APPEARANCE CODES:<br>[1] ময়লা স্পষ্টভাবে দেখা যাচ্ছিল(**ময়লা/কাঁদা/মাটি/ছাই/ এছাড়া অন্য কিছু**)VISIBLE DIRT (DIRT/MUD/SOIL/ASH OR ANY OTHER MATERIAL IS VISIBLE)<br>[2] ময়লা স্পষ্টভাবে দেখা না গেলেও অপরিচ্ছন্নভাব ছিল(হাতে **ময়লা দেখা না গেলেও**অপরিচ্ছন্ন ছিল)UNCLEAN APPEARANCE (NO DIRT IS VISIBLE ON THIS PART OF THE HAND BUT, IN GENERAL, THIS PART OF THE HAND APPEARS UNCLEAN)<br>[3] পরিষ্কার ছিল(হাতধোয়ার পর/গোসলের পর হাত দেখলে | 👁সবচেয়ে বড় বাচ্চার ০-৩৬ মাস OLDEST CHILD 0-36 MONTHS (If no children 0-36 months, child 36-60 months)<br><br>**বাম হাত Left Hand**<br>A\|__\| হাতের নখ FINGERNAILS<br>B\|__\| হাতের তালু PALMS<br>C\|__\| আঙ্গুলের সম্মুখভাগ FINGER PADS<br><br>**ডান হাত Right Hand**<br>D\|__\| হাতের নখ FINGERNAILS<br>E\|__\| হাতের তালু PALMS<br>F\|__\| আঙ্গুলের সম্মুখভাগ FINGER PADS |

| | | | |
|---|---|---|---|
| C.702a | 7 | পর্যবেক্ষনঃ প্রাথমিক/**প্রধান** হাত ধোয়ার স্থানটি কোথায় তা রেকর্ড | 1. ঘরের ভিতরে/ কাছে (≤৬ কদম) In/near main house (≤6 feet to entrance)<br>2. পায়খানার ভিতরে/ কাছে(≤৬ কদম) In/near latrine (≤6 feet to entrance)<br>3. রান্নার স্থানের ভিতরে/ কাছে (≤৬ কদম) In/near cooking area (≤6 feet to entrance) |

| | | |
|---|---|---|
| | | APPEAR AFTER SOMEONE WASHES HANDS OR TAKES A BATH)<br>[88] পর্যবেক্ষণ করা সম্ভব হয়নি/প্রত্যাখান/০-৩৬ মাসেরবাচ্চা নাই N/A; OBSERVATION NOT POSSIBLE/REFUSED/NO CHILD 0-36 MONTHS |
| | 70 | |

| | | | |
|---|---|---|---|
| | | করম্নন<br>**[Observation: Record the location where the primary hand washing station is located?]** | 4. ঘর, পায়খানা, রান্নার স্থানে থেকে >৬ কদম>6 feet away from main house, latrine and cooking area<br>**5.** নির্দিষ্ট কোন জায়গায় নাই No specific place 714 নং প্রশ্নে চলে যান (skips to 714) |
| C.702c | 7 | পর্যবেক্ষনঃ প্রাথমিক/ **প্রধান** হাত ধোয়ার স্থানটিতে কোন কোন উপাদান আছে? (আপনি যদি দেখে থাকেন তাহলে "1" কোড করম্নন আর না দেখলে "0" কোড করম্নন)<br>[Observation: Which of the following are present at the primary hand washing station? (If you observe the listed item, write "1" for "yes" in the box below. If you do not observe the listed item, write "0" for "no" in the box below.)]<br>*1= হ্যাঁ[Yes], 0=না [No]* | MATERIALS PRESENT<br>[1] ............................................ পানি WATER<br>[2] ............................................ গোসল/হাত ধোয়ার সাবান (লাক্স, লাইফবয়) BAR SOAP (Body/hand Bar)<br>[3] ............................................ অন্য যে কোন ধরনের সাবান (হুইল) BAR SOAP (other)<br>[4] ............................................ গুড়া পাউডার POWDERED SOAP<br>[5] সাবান-পানি SOAPY WATER<br>[6] তরল সাবান LIQUID SOAP<br>[7] ছাই Ash<br>[8] মাটি /বালু Mud/Sand<br>[9] আইসিডিডিআর, বি-র দেওয়া ট্যাপসহ বালতি ICDDR,B drum with tap<br>[10] হাত ধোয়ার স্থানটি ভিজা ছিল/ব্যবহারের চিহ্ন ছিল Moisture below handwashing station<br>[11] বালতি Bucket<br>[12] বেসিন, জগ Basin/Jug<br>[13] কিছুই নাই NONE OF THE ABOVE<br>[777] অন্যান্য OTHER (SPECIFY): _____________ |
| | 7 | **প্রধানহাত ধোয়ার স্থানটি সাবান জাতীয় যেকোন ধরনের উপাদান (যা হাত ধোয়ার কাজে ব্যবহার করা হয়/ সাবান পানি ও পানি একসাথে উপস্থিত আছে কী?**<br>[Soap/soapy water and water present together in PHWS] | 1. হ্যাঁ [Yes]<br>0. ............................................ না [No] |
| C.702c | 7 | প্রাথমিক/ **প্রধান** হাত ধোয়ার স্থানটি রান্নাঘর থেকে কত কদম দূরে?<br>[Observation: The primary hand | \|____\| |

| | | | |
|---|---|---|---|
| | | washing station is away from the kitchen---Count in steps, allow in continuous numbers] | |
| C.702d | 7 | প্রাথমিক/ প্রধান হাত ধোয়ার স্থানটি পায়খানা থেকে কত কদম দূরে? [Observation: The primary hand washing station is away from the toilet---Count in steps, allow in continuous numbers] | \|____\| |
| C.703 | 7 | উত্তরদাতাকে প্রশ্ন করুননঃ আপনার হাতধোয়ার জন্য প্রাথমিক/ প্রধান স্থান ছাড়া অন্যকোন জায়গা আছে কি? [Ask the respondent: "Is there anywhere else you wash your hands?" (Record code in box)]: | 1. হ্যাঁ [Yes]<br>0. না [No] **714**নং প্রশ্নে চলে যান (skips to 714) |
| C.703a | 7 | পর্যবেক্ষনঃ দ্বিতীয় প্রধান হাত ধোয়ার স্থানটি কোথায় তা রেকর্ড করুন [Observation: Record the location of the secondary hand washing station]. | 1. ঘরের ভিতরে/ কাছে (≤৬ কদম) In/near main house (≤6 feet to entrance)<br>2. পায়খানার ভিতরে/ কাছে(≤৬ কদম) In/near latrine (≤6 feet to entrance)<br>3. রান্নার স্থানে ভিতরে/ কাছে (≤৬ কদম) In/near cooking area (≤6 feet to entrance)<br>4. ঘর, পায়খানা, রান্নার স্থানে থেকে >৬ কদম>6 feet away from main house, latrine <u>and</u> cooking area |
| C.703c | 7 | পর্যবেক্ষনঃ দ্বিতীয় প্রধান হাত ধোয়ার স্থানটিতে কোন কোন উপাদান আছে? (আপনি যদি দেখে থাকেন তাহলে "1" কোড করুন আর না দেখলে "0" কোড করুন) [Observation: Which of the following are present at the secondary hand washing station? (If you observe the listed item, write "1" for "yes" in the box below. If | *<u>1= হ্যাঁ[Yes], 0=না [No]</u>*<br>[1]......................................................পানি WATER<br>[2] ......................................................গোসল/হাত ধোয়ার সাবান (লাক্স, লাইফবয়) BAR SOAP (Body/hand Bar)<br>[3] ......................................................অন্য যে কোন ধরনের সাবান (হুইল) BAR SOAP (other)<br>[4] ......................................................গুড়া পাউডার POWDERED SOAP<br>[5] সাবান-পানি SOAPY WATER<br>[6] তরল সাবান LIQUID SOAP<br>[7] ছাই Ash<br>[8] মাটি /বালু Mud/Sand<br>[9] আইসিডিডিআর, বি-র দেওয়া ট্যাপসহ বালতি ICDDR,B drum with tap<br>[10] হাত ধোয়ার স্থানটি ভিজা ছিল/ব্যবহারের চিহ্ন ছিল Moisture below handwashing station<br>[11] (বালতি Bucket |

| | | | |
|---|---|---|---|
| | | you do not observe the listed item, write “0” for “no” in the box below.) ] | [12] বেসিন, জগ Basin/Jug<br>[13] কিছুই নাই NONE OF THE ABOVE<br>৭৭৭. অন্যান্য OTHER (SPECIFY): _____________ |
| | 7 | দ্বিতীয় প্রধানহাত ধোয়ার স্থানটিতে সাবান জাতীয় যেকোন ধরনের উপাদান (যা হাত ধোয়ার কাজে ব্যবহার করা হয়/সাবান পানি ও পানি একসাথে উপস্থিত আছে কী? [Soap/soapy water and water present together in SHWS] | 1. হ্যাঁ [Yes]<br>0. ........................................................................................................না [No] |
| C.703c | 7 | দ্বিতীয় প্রধান হাত ধোয়ার স্থানটি রান্নাঘর থেকে কত কদম দূরে? [Observation:T he secondary hand washing station is away from the kitchen---Count in steps, allow in continuous numbers]] | \|____\| |
| C.703d | 7 | দ্বিতীয় প্রধান হাত ধোয়ার স্থানটি পায়খানা থেকে কত কদম দূরে? [Observation: The secondary hand washing station is away from the toilet---Count in steps, allow in continuous numbers] | \|____\| |
| INDICATOR 4: HANDWASHING সাবানেরউপস্থিতিতে হাতধোয়া দেখানো DEMONSTRATION, SOAP PRESENCE | | | |
| আপনাকে ধন্যবাদ। আপনি দয়া করে আমাকে পায়খানা থেকে আসার পর যেখানে বেশীরভাগ সময় হাত ধোন সেই স্থানটি দেখান(নোট ;পূর্বে পর্যবেক্ষীত স্থানও হতে পারে। এরপর উত্তরদাতা কর্তক চিহ্নিত স্থানে যেতে হবে এবং রেকর্ড করতে হবে।Thank you. Please show me where you most often wash your hands after defecation. (Note: this may be the same place you already observed. Go to the place identified by the respondent and record location). | | | |
| C.704a | 7 | 👁 হাতধোয়ার স্থান পর্যবেক্ষণ করুনন(একটিকে গোলাকার করুনন)। OBSERVE AND RECORDLOCATION OF HANDWASHING (CIRCLE ONE) | 1. ঘরের ভিতরে/ কাছে (≤৬ কদম) In/near main house (≤6 feet to entrance)<br>2. পায়খানার ভিতরে/ কাছে(≤৬ কদম) In/near latrine (≤6 feet to entrance)<br>3. রান্নার স্থানে ভিতরে/ কাছে (≤৬ কদম) In/near cooking area (≤6 feet to entrance)<br>4. ঘর, পায়খানা, রান্নার স্থানে থেকে >৬ কদম>6 feet away from main house, latrine and cooking area<br>5. নির্দিষ্ট কোন জায়গায় নাই |

| সময় গনণার জন্য ষ্টপ ওয়াচ প্রস্তুতকরন **PREPARE YOUR STOP WATCH FOR TIMING.** | | | |
|---|---|---|---|
| C.705 | 7 | পায়খানা থেকে আসার পর যেভাবে আপনি আপনার হাতধোন একইভাবে আমাকে এখন তা করে দেখান। Now, can you demonstrate how you normally clean your hands after defecation? Try in the same manner as you would if I wasn't here. TIME HOW LONG IT TAKES THE RESPONDENT TO BRING MATERIALS TO THAT PLACE (START AT THE MOMENT YOU FINISH ASKING THE QUESTION). WHEN RESPONDENT STARTS CLEANING HER HANDS, TIME DURATION OF RUBBING HANDS TOGETHER. | [1] দেখানো হয়েছে DEMONSTRATED<br><br>[2] দেখানো হয়নি COULD NOT DEMONSTRATE<br>IF NOT → **RECORD REASON: ________________________________________AND SKIP TO 721** |
| C.705b | 7 | 👁পর্যবেক্ষণ করম্নন এবং হাতধোয়ার সাবান আনতে কতটুকু সময় লেগেছে তা রেকর্ড করম্নন। যদি হাতধোয়ার স্থানে সাবান থাকে তাহলে ০০:০০ বসান। যদি সাবান ব্যবহার না করে থাকে তাহলে ৮৮:৮৮ বসান । OBSERVE AND RECORD TIME TO PRODUCE (Bangladesh: SOAP) FOR CLEANING HANDS MARK "00:00" IF SOAP IS ALREADY PRESENT AT HANDWASHING PLACE. (Bangladesh ONLY) MARK "88:88" IF SOAP IS N T USED | \|____:____\| মি.mm. সে.ss |

| | | | |
|---|---|---|---|
| C.705c | 7 | 👁 পর্যবেক্ষণ এবং হাতঘষার সময় রেকর্ড করুন \| উত্তরদাতা যখন হাত পরিস্কারের জন্য দুহাত ঘষবে সেই সময় গণণা করতে হবে ।<br>OBSERVE AND RECORD LENGTH OF TIME RESPONDENT SPENDS RUBBING HER HANDS TOGETHER: | \|____:____\| মি.mm. সে.ss |
| C.705d | 7 | 👁 পর্যবেক্ষণ এবং হাত ধোয়ার সকল উপাদান রেকর্ড করুন<br>OBSERVE AND RECORD ALL MATERIALS USED FOR HAND CLEANSING (CIRCLE ALL THAT APPLY) 1 হ্যাঁ Yes 0 না No | 1. পানি WATER<br>2. বার সাবান BAR SOAP (Body/hand BAR)<br>3. বার সাবান (অন্যান্য)BAR SOAP (other)<br>4. গুড়া পাউডার POWDERED SOAP<br>5. সাবান-পানি SOAPY WATER<br>6. কাপড়/পাতা CLOTH OR LEAVES<br>7. বেসিন/কলসBASIN/PITCHER<br>8. ছাই ASH<br>9. মাটি/বালু MUD<br>88. অন্যান্য উল্লেখ করুন OTHER (SPECIFY): ____________________________ |
| C.705e | 7 | 👁 পর্যবেক্ষণ এবং ২ হাত ধোয়া রেকর্ড করুন \|OBSERVE AND RECORD WHETHER BOTH HAN S WERE CLEANED | [1] হ্যাঁ, ২-হাত ধুয়েছে YES, BOTH HANDS CLEANED<br>[2] না ,১-হাত ধুয়েছে NO, ONLY ONE HAND CLEANED |
| C.705f | 7 | 👁 পর্যবেক্ষণ এবং হাত শুকানোরেকর্ড করুনOBSERVE AND RECORD HOW RESPONDENT DRIES HANDS<br>1 Yes 0 No | 1. পরিধেয় কাপড় DRIES BY WIPING HER HANDS ON HER CLOTHES<br>2.এক হাত অন্য হাতের সাথে ঘষে DRIES BY WIPING HER HANDS ON ANOTHER<br>3. পরিধেয় ছাড়া অন্য পরিষ্কার কাপড় Clean CLOTH DRIES BY WIPING HER HANDS ON<br>৪. পরিধেয় ছাড়া অন্য অপরিষ্কার কাপড় Unclean CLOTH DRIES BY WIPING HER HANDS ON<br>5. কাপড় ছাড়া অন্য জিনিস যেমন: টিস্যু, কাগজ ইত্যাদিMATERIAL (NOT CLOTH)<br>5. বাতাসে/হাত ঝাঁকিয়ে বাতাসে শুকানো হয়েছে DRIES BY SHAKING HER HANDS IN THE AIR<br>6. হাত শুকানো হয় নাই NOT DRIED<br>88. অন্যান্য উল্লেখ করুন OTHER, SPECIFY________________________ |
| C.705g | 7 | যদি উত্তরদাতা হাতধোয়া না দেখান অথবা সাবান ব্যবহার না করে তখন তাকে তার খানায় | \|____:____\| মি.mm. সে.ss |

| | | |
|---|---|---|
| | | হাতধোয়ার জন্য সাবান আছে কিনা জানতে হবে এবং তা আনতে বলতে হবে?IF RESPONDENT COULD NOT DEMONSTRATE, OR DID NOT USE SOAP, ASK: Do you have soap in your house that you use for handwashing? Can you bring it to me?<br><br>পর্যবেক্ষণ করুন এবং হাতধোয়ার সময় রেকর্ড করুন। যদি 716 তে উত্তর/সময় উল্লেখ থাকে তাহলে ৮৮:৮৮ বসান। যদি খানাতেসাবান না থাকে তাহলে ৯৯:৯৯ বসান । OBSERVE AND RECORD TIME TO PRODUCE SOAP FOR WASHING HANDS MARK "88:88" IF ALREADY TIMED IN C.704a MARK "99:99" IF OBSERVATION NOT POSSIBLE OR SOAP NOT AVAILABLE |

| | | | |
|---|---|---|---|
| C. 706a | 722. | পর্যবেক্ষণ করুন : উত্তরদাতার হাতধোয়া দেখানোর সময়ে এ বিষয়ে কোন প্রতিক্রিয়া দেখতে পেয়েছিলেন কী?OBSERVE: WAS THERE ANY OBVIOUS REACTIVITY DURING THE | [1] *হ্যাঁ YES*<br>[0] *না (৭২৩ নং এ চলে যাবে)* NO → skip to 723 |

| | | | |
|---|---|---|---|
| | | COURSE OF HIS DEMONSTRATION? | |
| b | 1 | অধিকক্ষণ যাবৎ হাত ধুয়েছে কিনা<br>Longer time spent cleaning/rubbing hands | [1] হ্যাঁ YES<br>[0] না NO |
| c | 2 | সাবান দিয়ে হাত ধুয়েছে কিনা অন্যথায়<br>Using soap where they otherwise would not | [1] হ্যাঁ YES<br>[0] না NO |
| d | 3 | অন্যান্য (লিখুন) Other, (BRIEFLY explain) | [1] হ্যাঁ YES<br>[0] না NO |

| | | | | | |
|---|---|---|---|---|---|
| | 72[illegible] | প্রশ্ন করুন ; আপনি কখন কখন সাবানদিয়ে হাত ধোন । ASK: "Please tell me about all of the times you wash your hands with soap."<br><br>A কলা ম (লিড ছাড়া)-এ যখন উত্তরদাতা যে কাজ করবে তাউল্লেখ করতে হবে । ১নং গোলাকার হবে যদি নির্দিষ্ট বিষয়ে উত্তর পাওয়া যায়। CIRCLE "1" IN COLUMN A IF CRITICAL TIME IS MENTIONED WITHOUT PROMPTING.<br><br>উত্তরদাতার সকল কাজ উল্লেখ করার পর অন্য আর কোন কাজে সাবানদিয়ে হাত ধোন কিনা এ ব্যপারে জিজ্ঞাসা করুন? AFTER THE RESPONDENT FINISHES NAMING ALL THE TIMES ASK "IS THERE ANY OTHER TIME YOU WASH YOUR HANDS WITH SOAP?" AND STOP WHEN THE RESPONDENT SAYS THERE IS NO OTHER TIME.<br><br>B কলাম-এ লিড দেওয়া ছাড়া প্রতিবার ব্যবহার করে কিনা তা উল্লেখ করুন । EACH TIME MENTIONED WITHOUT PROMPTING ASK COLUMN B.<br><br>C.707A-G নং প্রশ্নের উত্তর যদি A কলা ম [1] না হয় তখন সাবান ও পানি দিয়ে আপনি আপনার হাতধোন কিনা তা জিজ্ঞাসা করুন। যদি উত্তরদাতা হ্যাঁ বলে তখন C কলা ম-এ হাতধোয়ার সংখ্যা/বার উল্লেখ করুন । FOR QUESTIONS C.707A-G WHERE [1] IS NOT CIRCLED IN COLUMN A, PROMPT BY ASKING:<br>Do you usually wash your hands with soap and water [CRITICAL TIME]? IF THE RESPONDENT SAYS "YES" THEN IMMEDIATELY ASK HOW OFTEN AND MARK APPROPRIATE ANSWER CHOICE IN COLUMN C. | A.<br>লিড দেওয়া ছাড়া হাতধোয়া উল্লেখ করুন<br>MENTIONED WITH NO PR OMPTING | B.<br>লিড দেওয়া ছাড়া শুধুমাত্র হাতধোয়ার সংখ্যা/বার উল্লেখ করুন । সাবান ও পানি দিয়ে কত বার আপনার হাতধোন তা বলুন । Only for times mentioned with NO PROMPTING, ask: How often do you wash your hands with soap and water [insert event]?<br><br>[1] সবসময়/প্রতিবার ALWAYS<br>[2] মাঝে মাঝে/ অর্ধেকের কম বার SOMETIMES<br>[3] খুব কম সময়/ দু-এক বার RARELY | C.<br>লিড দেওয়ার পর শুধুমাত্র হাতধোয়ার সংখ্যা/বার উল্লেখ করুন । সাবান ও পানি দিয়ে কত বার আপনার হাতধোন তা বলুন । REPLIED "AFTER BEING PROMPTED<br><br>[1] সবসময়/প্রতিবার ALWAYS<br>[2] মাঝে মাঝে/ অর্ধেকের কম বার SOMETIMES<br>[3] খুব কম সময়/ দু-এক বার RARELY<br>[4] না NO<br>[88] প্রযোজ্য নয় Not applicable |
| a | 1 | খাবার তৈরী করার আগে BEFORE PREPARING FOOD | | | |
| | 2 | মাংস/মাছ কাটার পর AFTER HANDLING MEAT/FISH | | | |
| | 3 | ফল/সব্জি কাটার আগেBEFORE CUTTING FRUITS AND VEGETABLES | | | |
| | 4 | ভর্তা/ম্যাসড খাবার তৈরী করার আগে BEFORE MAKING MASHED FOOD (BHORTA) | | | |
| b | 5 | খাবারের পূর্বেBEFORE EATING | | | |
| c | 6 | খাবারের পরে AFTER EATING | | | |
| d | 7 | শিশুকে খাওয়ানোর আগে BEFORE FEEDING A CHILD | | | |
| e | 8 | বাচ্চাকে সৌচানোর পরAFTER CLEANING A CHILD'S ANUS | | | |
| f | 9 | বাচ্চার পায়খানা ফেলার পরAFTER DISPOSING OF CHILDREN'S FECES | | | |

| | | |
|---|---|---|
| g | 10 | পায়খানার পরAFTER TDEFECATION |
| h | 11 | কম্পাউন্ড/বাড়ীর বাহির থেকে ফেরার পর AFTER RETURNING FROM OUTSIDE THE COMPOUND |
| m | 12 | রুগ্ন ব্যক্তিকে ধরার পরAFTER TOUCHING A SICK PERSON |
| o | 13 | পশু-পাখি ধরার পরAFTER HANDLING LIVESTOCK |
| | 14 | গোবর ধরার পর AFTER HANDLING COW-DUNG (OR ANY ANIMAL FECES) |
| j | 15 | অন্যান্য (উল্লেখ করুন) OTHER (SPECIFY): ___________________________ |
| i | 16 | উত্তরদাতা কখনই সাবান দিয়ে হাতধোয় নি RESPONDENT NEVER WASHES HANDS WITH SOAP Skip to module 8 |
| | 17 | উঠান ঝাড়ু দেওয়া/ময়লা পরিষ্কারের পর After weaping courtyard |
| | 18 | রান্নার পর After cooking |
| | 19 | থালা-বাসন ধোয়ার পর After washing dishes |

**Wash Benefit Module 8SANITATION (All households)**

| | |
|---|---|
| SAY | এই গবেষণায় আপনার অংশ গ্রহনের জন্য আপনাকে ধন্যবাদ। ষ্টাডির এই অংশটি কিছটা স্পর্শকাতর । আমি আপনাকে আপনার/এই বড়ীর পয়ঃ ব্যবস্থাপনা/পয়ঃ নিষ্কাশন ও তার অনুশীলন এবং কিভাবে তা করেনএ ব্যপারে জানতে চাই\| আমি কিছুটা পর্যবেক্ষণ করতে চাই। আপনার স্বাচ্ছন্দবোঃ হবে কি ?যদি না হয় তাহলে উত্তরদাতাকে ইহার গুরুত্ব বোঝাতে চেষ্টা করুন।Thank you so much for your participation so far. The next part of the survey is a bit sensitive. I would like to ask you some questions about the sanitation practices of people in your compound, including how you usually dispose of your children’s feces. I would also like to make some observations. Are you comfortable with this? IF NOT, EXPLAIN THE IMPORTANCE OF THIS MODULE AND ENCOURAGE RESPONDENT TO PROCEED. |

খোলা-পায়খানা(সকল খানার )OPEN DEFECATION (ADMINISTER TO ALL HOUSEHOLDS)

| | | | Group A | B | C | D | E |
|---|---|---|---|---|---|---|---|
| | WBM 8 | | Men পুরুষ | Women মহিলা | Children <3 years <৩ বছরের বাচ্চা | Children 3-<8 years ৩-<৮ বছরের বাচ্চা | Children ৮-১৫ ছরের বাচ্চা |
| | | এই খানার কোন সদস্যরা খোলা জায়গাতে পায়খানা করে কী? [Do [GROUP] in this household ever practice open defecation? 1 প্রতিদিন Daily | ☐ | ☐ | ☐ | ☐ | ☐ |

| | | | | | | | |
|---|---|---|---|---|---|---|---|
| | | 2 মাঝেমাঝে Occasionally<br><br>3 কখনই না Never **(Skip to 806)**<br><br>88. প্রযোজ্য নয় Not applicable (**806**নং প্রশ্নে যান)**(Skip to 806)**<br><br>99 জানি না (**806**নং প্রশ্নে যান) Don't Know **(Skip to 806)** | | | | | |
| | | সাধারনত: একই জায়গাতে প্রতিবার যান কী? [Do [GROUP] go to more or less the same area every time?<br><br>1হ্যাঁ Yes<br><br>2না No (**805**নং প্রশ্নে যান)**(Skip to 805)**<br><br>99জানি না Don't Know / Not Sure (**805** নং প্রশ্নে যান)**(Skip to 805)** | ☐ | ☐ | ☐ | ☐ | ☐ |
| | | আপনার খানা থেকে উক্তস্থানে যেতে (শুধু যাওয়া) কতটুকু সময় লাগে (মিনিট) [How long does it take to walk (one way) from your house to the most commonly visited place? | I_____I মিনিট MINUTES<br><br>99 জানি না Don't Know / Not Sure | | | | |
| | | এই জায়গাটি ঐ গ্রামের মধ্যেই কী? [Is that place within the village?<br><br>1হ্যাঁ Yes<br><br>2না No<br><br>99জানি না Don't Know / Not Sure | I__I | | | | |

Open Defecation Questions

| | | | Men<br>পুরম্নষ | Women<br>মহিলা | Children<br><3 years<br><৩ বছরের বাচ্চা | Children<br>3-<8 years<br>৩-<৮ বছরের বাচ্চা | Children<br>৮-১৫ ছরের বাচ্চা |
|---|---|---|---|---|---|---|---|
| C.804 a | | আপনার খানার লোকজনের খোলাজায়গায় পায়খানা করার প্রধান কারন কি কি? উত্তর শুনানো যাবে না ।What are the main reasons that [GROUP] in your household practice open defecation?<br>✋DO NOT READ RESPONSES<br>MARK ALL THAT APPLY | | | | | |
| | 1 | 1 কোনও কিছুই নাই /পায়খানা নাইNo choice (nothing else is available) | ☐ | ☐ | ☐ | ☐ | ☐ |
| | 2 | 2কোথায়শিশু পায়খানা করবে তা নিয়ন্ত্রন করতে পারে না Cannot control where young children defecate | | | ☐ | ☐ | ☐ |
| | 3 | 3 গোপনীয়তাPrivacy | ☐ | ☐ | ☐ | ☐ | ☐ |

| | | | | | | | |
|---|---|---|---|---|---|---|---|
| | 4 | 4 অভ্যাস/নিয়মিত কাজHabit / Routine | ☐ | ☐ | ☐ | ☐ | ☐ |
| | 5 | 5 পায়খানা থেকে বাঁশঝাড় পছন্দ Prefer to use the bush rather than a toilet | ☐ | ☐ | ☐ | ☐ | ☐ |
| | 6 | 6 কাজে/স্কুলে পায়খানা/টয়লেট সহজে পাওয়া না গেলে Toilet not available at work / school | ☐ | ☐ | ☐ | ☐ | ☐ |
| | 7 | ভাগের পায়খানা/টয়লেট পছন্দ না হলে Choose not to share toilets with in-laws / extended family (or cannot) | ☐ | ☐ | ☐ | ☐ | ☐ |
| | 8 | 8 সুবিধাজনক Convenience | ☐ | ☐ | ☐ | ☐ | ☐ |
| | 9 | 9 নিরাপদ Safety | ☐ | ☐ | ☐ | ☐ | ☐ |
| | 10 | 10 আরামদায়কComfort | ☐ | ☐ | ☐ | ☐ | ☐ |
| | 11 | 11 অসুস্থতা/ পাতলা পায়খানাSickness/diarrhoea | ☐ | ☐ | ☐ | ☐ | ☐ |
| | 12 | 12 পায়খানা/টয়লেট ভরে গেছে Latrine overflowed | ☐ | ☐ | ☐ | ☐ | ☐ |
| | 13 | 13 পায়খানা/টয়লেট ভেঙে গেছে(উপরের অংশ/সম্বাব) Latrine | ☐ | ☐ | ☐ | ☐ | ☐ |

| | | | | | | | |
|---|---|---|---|---|---|---|---|
| | | broken<br><br>(superstructure and /or slab) | | | | | |
| | 14 | 14 পায়খানা/টয়লেটে যেতে ভয় পায় Fear of latrine | ☐ | ☐ | ☐ | ☐ | ☐ |
| | 15 | 15 পায়খানা/টয়লেট কিভাবে ব্যবহার করতে হয় তা জানে না Don't know how to use the latrine | ☐ | ☐ | ☐ | ☐ | ☐ |
| | 16 | 16 বাচ্চাখুব ছোট হওয়ায় পায়খানা/টয়লেট ব্যবহার করে না Too young to use latrine | ☐ | ☐ | | | |
| | 17 | বয়স্ক (খুবই বৃদ্ধ)Old age (too old) | | | | | |
| | | | **<৩ বছরের বাচ্চাChildren < 3 years** | **৩-<৮ বছরের বাচ্চাChildren<br><br>3 – <8 years** | **৮-১৫ বছরের বাচ্চাChildren<br><br>8 – 15 years** | **পুরুষ Men** | **মহিলা Women** |
| **C.X** | | **পায়খানা থাকা সত্ত্বেও এই বাড়ীর অন্য কোনপরিবারখোলা জায়গায় পায়খানা করে কিনা তা আপনি জানেন কী?** Do you know of other households in the bari/compound whose [GROUP] practice OD, even if they might have a toilet or latrine at their house?<br><br>1 হ্যাঁ, প্রায়ই | ☐ | ☐ | ☐ | ☐ | ☐ |

| | | | | | | | |
|---|---|---|---|---|---|---|---|
| | | Yes, Often<br><br>2হ্যাঁ, মাঝে মাঝে Yes, Sometimes<br><br>3না, কখনই না No, Never**(Skip to 808)**<br><br>88. প্রযোজ্য নয় Not applicable (**808** নং প্রশ্নে যান)**(Skip to 808)**<br><br>99 জানি না/নিশ্চিত না Don’t Know / Not Sure | | | | | |
| **C.X** | | আপনি শিশুদের পায়খানা উঠানে/উঠানের চার-পাশে/আশেপাশে /নদীতে ফেলে দিতে দেখেছেন কী? Do you see that children’s stools are disposed in the yard / surrounding / community in your neighborhood / river?<br><br>1 হ্যাঁ, প্রায়ই Yes, Often<br><br>2 হ্যাঁ, মাঝে মাঝে Yes, Sometimes<br><br>3 না, কখনই না No, Never<br><br>99 জানি না/নিশ্চিত না Don’t | | | | ☐ | ☐ |

| | | |
|---|---|---|
| | | Know / Not Sure |

| | | | |
|---|---|---|---|
| পয়ঃ ব্যবস্থাপনা সুবিধাদি Sanitation Facility | | | |
| Administer to: All study households | | | |
| C | | আপনার খানায় কোন টয়লেট/পায়খানা আছে কি?<br>আমি কি তা দেখতে পারি?<br>Does your household have a toilet facility that is in use? Can I see it? | 1 হ্যাঁ আছে,পর্যবেক্ষন করতে পারি Yes have toilet, can ob<br>2 হ্যাঁ আছে,পর্যবেক্ষন করতে দেওয়া হয়নি/প্রত্যাখান (৮১২ নং ) Yes have toilet, refused observation **(Skip to 812)**<br>3 হ্যাঁ আছে,পর্যবেক্ষন করা যায়নি(৮১২ নং এ যান ) Yes hav toilet, cannot observe **(Skip to 812)**<br>4 কোন পায়খানা নাই No toilet facility **(Skip to MODULE** |
| | | 👁 পর্যবেক্ষন; ধরণ,অবস্থা এবং পুনঃব্যবহার অবস্থা Observation: Note the type, condition and apparent use of the toilet:<br>1 হ্যাঁ Yes<br>2 না No<br>88 প্রযোজ্য নয়/ পর্যবেক্ষন করতে পারেনি/বলতে পারেনি Not Applicable / Could not observe / cannot tell | |
| বর্হিরাংশ পর্যবেক্ষণ **Exterior observation** | | | |
| 1 | 1 | পায়খানার চারপাশে ৩টি ইটের/ টিনের/মাটির দেয়াল At least 3 walls around the toilet | *(1=হ্যাঁ[Yes], 0= না [No],* প্রযোজ্য নয় *[*not applicable] =888 |
| 2 | 2 | পায়খানার চারপাশে ৩টি বাঁশের বেড়ার দেয়ালBamboo fences around the | *(1=হ্যাঁ[Yes], 0= না [No],* প্রযোজ্য নয় *[*not applicable] =888 |

| | | | |
|---|---|---|---|
| | | toilet | |
| 3 | 3 | পায়খানার চারপাশে দরজা/পর্দা দিয়ে ঘেরাDoor/curtain or walls that guarantee privacy around the toilet | *(1=হ্যাঁ[Yes], 0= না [No],* প্রযোজ্য নয় *[*not applicable] =888 |
| 4 | 4 | পায়খানার ছাদ Roof over toilet | *(1=হ্যাঁ[Yes], 0= না [No],* প্রযোজ্য নয় *[*not applicable] =888 |
| 5 | 5 | বাতাস বের হওয়ার পথ Ventilation pipe | *(1=হ্যাঁ[Yes], 0= না [No],* প্রযোজ্য নয় *[*not applicable] =888 |
| 6 | 6 | পায়খানায় যাবার রাস্ৰা দেখে বুঝা যাচ্ছে নিয়মিত ব্যবহার করা হয় (পরিষ্কার, জীর্ণ ইত্যাদি )Path to the toilet suggests regular use (is clear, well-worn, without grass or any barriers etc.) | *(1=হ্যাঁ[Yes], 0= না [No],* প্রযোজ্য নয় *[*not applicable] =888 |
| ভিতরের অংশ পর্যবেক্ষণ **Interior observation** | | | |
| 7 | 7 | টয়লেটে স্লাব আছেToilet has a slab | *(1=হ্যাঁ[Yes], 0= না [No],* প্রযোজ্য নয় *[*not applicable] =888 |
| 8 | 8 | গর্তের পাশে পাদানী আছেRaised footing around hole | *(1=হ্যাঁ[Yes], 0= না [No],* প্রযোজ্য নয় *[*not applicable] =888 |
| 9 | 9 | পানি ঢেলে ফ্লাস করা যায় Flush or poor Flush: | *(1=হ্যাঁ[Yes], 0= না [No],* প্রযোজ্য নয় *[*not applicable] =888 0/888 skip to 12] |
| 9a | 9a | যদি পানি ঢেলে ফ্লাস করা যায় If Flush or Pour Flush:<br><br>পানির সীলের অবস্থা Water seal condition: | [ 1 ] ওয়াটার সিল ভাল আছে/কাজ করে Functional water seal<br>[ 2 ] ওয়াটার সিল ভাঙ্গা Broken water seal<br>[ 3 ] ওয়াটার সিল নেই No water seal<br>[ 88 ] পায়খানাতে পানির সীল আছে/পর্যবেৰণ করা সম্ভব হয়া could not observe / cannot tell |
| 9b | 9b | যদি পানি ঢেলে ফ্লাস করা যায়If Flush or Pour Flush:<br><br>পানি ঢাললে/দিলে Flushes to:<br><br>*(খানার সদস্যকে জিজ্ঞাসা/যাচাই করুন Ask / probe household members if* | টয়লেটিতে পয়ঃনিষ্কাশন পাইপের সাথে সংযোগ করে দেয়া [Piped sew<br>system]<br>টয়লেটিতে সেপটিক ট্যাংক বসানো আছে [Septic tank]<br>সেপটিক ট্যাংক নাই কিন্ৰ ফ্লাস করে বা পানি ঢেলে পায়খানা দুরে বা পিটের সরিয়ে দেয়া যায় [Flush to pit latrine ]<br>ফ্লাশ-টয়েলেট অথবা পানি ঢেলে ফ্লাস করা টয়লেট যা কোন খাল, ড্রেন, নদী সাথে সংযোজন করার ফলে অস্বাস্থ্যকর অবস্থার সৃষ্টি করে থাকে[Flush or flush toilet connected to somewhere else (canal, dit |

| | | | |
|---|---|---|---|
| | | *necessary)* | river, etc.)] |
| | 9c | পায়খানার অবস্থা কেমনCondition of the latrine | 1. স্বাস্থ্যসম্মত পায়খানাHygienic latrine<br><br>2. অস্বাস্থ্যকর পায়খানাNon hygienic latrine |
| 10 | 10 | মেঝের প্রধান উপাদান Main material of the floor (*select 1*) | [ 1 ] মাটি Mud<br><br>[ 2 ] কাঠ Wood<br><br>[ 3 ] সিমেন্ট Cement<br><br>[ 4 ] টাইলস/ইটTile / brick<br><br>[ 5 ] প্লাষ্টিক Plastic<br><br>[ 88 ] প্রযোজ্য নয়/পর্যবেক্ষণ সম্ভব হয় নি N/A / could not obser cannot tell |
| 11 | 11 | Bucket toilet | *(1=হ্যাঁ[Yes], 0= না [No],* **প্রযোজ্য নয়** *[*not applicable] =888 |
| 12 | 12 | ঝুলন্ত পায়খানা Hanging toilet | [ 1 Yes ] [ 2 No ] [ 88 N/A] |
| 13 | 13 | ব্যবহৃত হচ্ছে বুঝা যায় এমন Latrine appears to be in use (by your best judgment) | *(1=হ্যাঁ[Yes], 0= না [No],* **প্রযোজ্য নয়** *[*not applicable] =888 |
| 14 | 14 | **পায়খানা/টয়লেটে পায়খানার গন্ধ রয়েছে**Odor of feces in the latrine/bathroom | *(1=হ্যাঁ[Yes], 0= না [No],* **প্রযোজ্য নয়** *[*not applicable] =888 |
| 15 | 15 | **পায়খানা/টয়লেটে প্রস্রাবের গন্ধ রয়েছে**Odor of urine in the latrine/bathroom | *(1=হ্যাঁ[Yes], 0= না [No],* **প্রযোজ্য নয়** *[*not applicable] =888 |
| 16 | 16 | স্লাব অথবা মেঝেতে **পায়খানা দেখতে পাওয়া গেছে** Stool is visible on the slab or floor | *(1=হ্যাঁ[Yes], 0= না [No],* **প্রযোজ্য নয়** *[*not applicable] =888 |
| 17 | 17 | ড্রপ হোল(মল-মূত্র প্রবেশ গর্ত) ঢাকা Drop hole is covered | *(1=হ্যাঁ[Yes], 0= না [No],* **প্রযোজ্য নয়** *[*not applicable] =888 0/888 skip to 18] |
| 17a | 17a | যদি ড্রপ হোল (মল-মূত্র প্রবেশ গর্ত)ঢাকা থাকে তাহলে মাছি আসা/যাওয়া করতে পারে কি If yes, cover is correctly placed over the drop hole so that flies cannot enter/exit | *(1=হ্যাঁ[Yes], 0= না [No],* **প্রযোজ্য নয়** *[*not applicable] =888 |

| | | | |
|---|---|---|---|
| 18 | 18 | **মাছির উপস্থিতি রয়েছে** Flies present | *(1=হ্যাঁ[Yes], 0= না [No],* **প্রযোজ্য নয়** *[*not applicable] =888 |
| 19 | 19 | **সাধারন বৈশিষ্ট General Characteristics**<br><br>একটি অথবা দুই পিট/গর্ত টয়লেট Single or double pit latrine | [ 1 ] একটি পিট/গত Single pit<br><br>[ 2 ] দুই পিট/গর্ত Double pit<br><br>[ 88 ] প্রযোজ্য নয়/পর্যবেক্ষণ সম্ভব হয় নি /বলতে পারে নিN/A / could observe / cannot tell |
| 20 | 20 | কমপোষ্টিং টয়লেট Composting toilet | *(1=হ্যাঁ[Yes], 0= না [No],* **প্রযোজ্য নয়** *[*not applicable] =888 |
| | | **পর্যবেক্ষন করুন** Observation: টয়লেট-এর গর্ত মলে ভরেছে কিনা-আলোর সাহায্যে পর্যবেৰণ করম্ননFullness of the pit – shine a light into the pit to see if solid waste is… | 1 **শুকনা ময়লা/পায়খানা ৩ ফুটের চেয়ে বেশী দূরে (>৩** Very far from surface (>1 meter)<br><br>2 **শুকনা ময়লা/পায়খানা ৩ ফুটের মধ্যে (<৩ ফু** Within 1 meter<br><br>3 **পায়খানার খুব কাছাকাছি বা পায়খানা ভরে** Very close to surface or full<br><br>88 **পায়খানাতে পানির সীল আছে/পর্যবেৰণ করা হয়নি**Water seal latrine / non direct pit latrine / could observe |
| C. 808 | | **পর্যবেক্ষন করুন** Observation: : সৌচ কাজের জন্য টয়লেট-এর ভেতরে এবং কাছে **কী কী উপাদান রয়েছে** What materials for anal cleansing are present inside or immediately outside the latrine? | *(1=হ্যাঁ[Yes], 0= না [No],* **প্রযোজ্য নয়** *[*not applicable] =888<br><br>1.*পাতা/ঘাস* Leaves/grass<br><br>2.*কাঠি* Twigs / sticks<br><br>3. *কাপড়* Rag or cloth<br><br>4. *পাথর* Stones<br><br>5. *স্বাস্থ সম্মত কাগজ*Hygienic (toilet) paper<br><br>6. *পানির পাত্র/বদনা/মগ*Water container / vessel<br><br>7. *পানির ট্যাপ* Water tap<br><br>8. *সাবান* Soap<br><br>9. *ছাই অথবা মাটি* Ash or soil for cleansing<br><br>10. *খবরের কাগজ* Newspaper |

| | | | |
|---|---|---|---|
| | | | 11. কিছুই নাই Nothing |
| C. 809 | | আপনার খানার কে কে এই পায়খানা/টয়লেটটি পায়খানার জন্য ব্যবহার করে? Please tell me about who in your family uses the latrine for defecation. | |
| 1 | 1 | <**3**বাচ্চা Children <3 | 1. সব সময় [Always]<br>2. প্রায়ই [Usually]<br>3. মাঝেমাঝে [Sometime]<br>4. কখনই না [Never]<br>888. প্রযোজ্য নয় [ N/A] |
| 2 | 2 | ৩-<৮বাচ্চা Children 3-<8 | 1. সব সময় [Always]<br>2. প্রায়ই [Usually]<br>3. মাঝেমাঝে [Sometime]<br>4. কখনই না [Never]<br>888. প্রযোজ্য নয় [ N/A] |
| 3 | 3 | ৮-১৫বাচ্চা Children 8-15 | 1. সব সময় [Always]<br>2. প্রায়ই [Usually]<br>3. মাঝেমাঝে [Sometime]<br>4. কখনই না [Never]<br>888. প্রযোজ্য নয় [ N/A] |
| 4 | 4 | পুরুষMen | 1. সব সময় [Always]<br>2. প্রায়ই [Usually]<br>3. মাঝেমাঝে [Sometime]<br>4. কখনই না [Never] |

| | | | |
|---|---|---|---|
| | | | 888. প্রযোজ্য নয় [ N/A] |
| 5 | 5 | মহিলাWomen | 1. সব সময় [Always]<br><br>2. প্রায়ই [Usually]<br><br>3. মাঝেমাঝে [Sometime]<br><br>4. কখনই না [Never]<br><br>888. প্রযোজ্য নয় [ N/A] |
| C. 810 | | অন্য কোন খানা কি এই পায়খানা ব্যবহার করে?Do you share this toilet with other households? | *1=হ্যাঁ[Yes]*<br><br>*0=না[No].......skip to 815* |
| C.811 | | কতগুলো খানা মিলে এই পায়খানাটি ব্যবহার করে ? How many HHs use it? | (Don't know=99)<br><br>I____I I____I |
| C.812 | | এইখানার শিশুসহ আপনারা কতজন এই পায়খানাটি ব্যবহার করেন? How many people including children in your household use this toilet? | I____I I____I |
| C. 813 | | পায়খানার মালিকানা ? (প্রশ্ন করুন) [***Ask***: Who owns the toilet facility?] | শুধুমাত্র ঐ খানার জন্য [Only for the household] 1<br><br>কয়েকঘর মিলে/ অংশীদার [Shared] 2<br><br>অন্য কেউ [Someone else] 3<br><br>পাবলিক [Public]4<br><br>প্রযোজ্য নয় [Not applicable]...............................................888 |
| C. 814 | | এই স্থানে এই পায়খানাটি কত বছর ধরে আছে? (জানি না=999) [How long have you had the present latrine in this place? [In years] (Don't know = 99)] | \|_____\|\|_____\| বৎসর Years<br><br>\|_____\|\|_____\| মাস Months |

**Wash Benefit Module 9 Child defecation and feces disposal practice**

| | | | |
|---|---|---|---|
| wkïi cvqLvbv e¨e¯’vcbv Ges gj wb®‹vkb Af¨vm **Child defecation and feces disposal** | | | |
| Administer to: All study households ( **mKj Lvbvq n‡e**) | | | |
| C.912 | | \|_____\|\|_____\|**KZ eQi eqm †_‡K GKwU wkï mvavibZt Uq‡jU/cvqLvbv e¨envi Kiv ïi" K‡i (gv‡m)?** [At what age (in months) does a child start using a latrine, if at all?] | |
| C.902 | | \|_____\| **Avcbvi (<3 eQi) wkï me©†kl KLb cvqLvbv K‡i‡Q? (GKvwaK wkï _vK‡j me‡P‡q eo ev"Pvi †¶‡Î wR‡Ám Ki"b)** [When was the last time your youngest child/infant (<3 years) defecated?] | **AvR** [Today]<br><br>**MZKvj** [Yesterday]<br><br>**2 w`b ev Zvi Av‡M** [Before 2 or more days back]<br><br>**KLbB bv/ g‡b Ki‡Z cviwQ bv** [Never/can't remember]..... **bs cÖ‡kœ P‡j hvb** (skips to 906)<br><br>**ej‡Z ivwR bv** [Refused] ……….**906bs cÖ‡kœ P‡j hvb** (ski[illegible] 906)<br><br>88. **cÖ‡hvR¨ bq** [Not applicable]......... **908bs cÖ‡kœ P‡j** (skips to 908) |
| | | \|_____\| **Avcbvi wkï me©†kl ‡Kv_vq cvqLvbv K‡i‡Q?** [Where did the child defecate the last time?]<br><br><u>Note:</u>**DËi c‡o †kvbv‡eb bv, Zv‡K wbw`©ó K‡i ej‡Z ejyb \| Gwel‡q hZ¶Y ch©šÍ wKQy ejvi _v‡K ZZ¶Y ch©šÍ Zv‡K ej‡Z DrmvwnZ Ki"b** [Don't read the answer, encourage by asking if there is anything else until he/she mentions there in nothing else and check all mentioned.] | **cwU (DVv‡b)**[Potty (in the courtyard)]<br><br>**cwU (N‡ii wfZ‡i)**[Potty (inside the house)]<br><br>**DVv‡b (cwU Qvov)**[Courtyard (without potty)]<br><br>**N‡ii wfZ‡i (cwU Qvov)** [Inside the house (without potty)<br><br>**DVv‡bi evwn‡i †Lvjv RvqMv‡Z** [Open space outside the yard<br><br>**‡Svc-Sv‡o/R½‡j** Bush/jungle<br><br>**cvqLvbvq/Uq‡j‡U** [In toilet] ......**906bs cÖ‡kœ P‡j hvb** ([illegible] 906)<br><br>**Kv_v/b¨vwc/Wvqcvi** Katha/nappy/diaper<br><br>77. **Ab¨vb¨ (eY©bv wjLyb)** [Other (Please |

| | | | |
|---|---|---|---|
| | | | describe)]....................<br><br>99. **Rvwb bv** [Don't Know].............**906bs cÖ‡kœ P‡j h**[illegible] to 906) |
| | | \|_____\| **†mB cvqLvbv wK Kiv n‡qwQj?**[What was done to the feces?] | **n¨uv [Yes].....1, bv [No]......0**<br><br>1. **†hLv‡b cvqLvbv K‡iwQj †mLv‡bB †d‡j ivLv n‡qwQj** [[illegible] there]**905.abs cÖ‡kœ P‡j hvb** (Skip to 906)<br><br>2. \|_____\| **Uq‡jU/cvqLvbvi wfZ‡i †djv/†avqv n‡qwQj** [Put/rinsed into toilet or latrine]<br><br>3. \|_____\|**‡Wª‡b/b`©gvi wfZ‡i †djv/†avqv n‡qwQj** [Put/rinsed into drain or ditch]<br><br>4. \|_____\|**‡Svc-Sv‡o/R½‡j †djv n‡qwQj** [Thrown into Bush/jungle<br><br>5. \|_____\|**gqjv AveR©bvi g‡a¨ †djv n‡qwQj** [Thrown in[illegible] garbage]<br><br>6. \|_____\|**wbw`ó© M‡Z© †djv n‡qwQj** [Thrown into a pit for child's feces]<br><br>7. \|_____\|**gvwUi bx‡P cy‡Z †djv n‡qwQj** [Buried]<br><br>77. \|_____\|**Ab¨vb¨ (eY©bv wjLyb)** [Other (specify)]....<br><br>99. \|_____\| **Rvwb bv** [DK]**906 bs cÖ‡kœ P‡j hvb** (Skip t[illegible] |
| | | **Avcwb wKfv‡e cvqLvbv cwi®‹vi K‡ib?DËi c‡o ïbv‡bv hv‡e bv\|n¨vu n‡j 1 emvb \|**did you handle the feces? ☝ <u>Do Not Read Responses. Mark All that Apply</u> (1 = Yes) | \|_____\|**bMœ/Lvwj nv‡Z** Hands only (bare hands)<br><br>\|_____\|**nv‡Z Ges Kvco/cvZv/KvMR** Hands and cloth / p[illegible] leaves<br><br>\|_____\| **cvqLvbv †djvi Rb¨ †Kvb Dcv`vb e¨envi K‡i** Scra[illegible] material to scoop fece<br><br>\|_____\|**cwU** Potty<br><br>\|_____\|**¯'vbxq K…wlKvR-Gi nvwZqvi** Local agricultural hoe/instrument<br><br>\|_____\|**‡mwb ¯‹zc** Sani-scoop<br><br>\|_____\|**‡Kvb wKQyB Kiv nq bv** Did nothing |

<table>
<tr><td></td><td></td><td></td><td>77 |_____|Ab¨vb¨ (D‡j-L Ki“b)Others (specify)_________<br><br>99 |_____|Rvwb bvDon't know / not sure</td></tr>
<tr><td>C.905.b</td><td>905.a</td><td>cvqLvbv Kivi ci KZÿY GB cvqLvbv GLv‡b c‡i wQj? How long after defecation did you dispose of the feces?</td><td>|____|Hour N›Uv<br><br>|____|Minute wgwbU</td></tr>
<tr><td></td><td></td><td>Avcbvi 3 eQ‡ii †QvU ev”Pvi cvqLvbv †K ‡K †d‡j? (hZRb †d‡j mevB‡K †KvW Ki“b) [Who disposes your under 3 child's feces?</td><td><u>n¨uv [Yes].....1, bv [No]......0</u><br><br>|_____| gv [Mother]<br><br>|_____| evev [Father]<br><br>|_____| ‡evb [Sister]<br><br>|_____| fvB [Brother]<br><br>|_____| Lvjv/gvgx/PvPx/dydz/`v`x [Aunt/grandmother<br><br>|_____| Lvjy/gvgv/PvPv/dzdv/`v`v [Uncle/grantfather]<br><br>|_____| ‡KD cvqLvbv †d‡j bv [Nobody dispose]<br><br>77. |_____| Ab¨vb¨ (wbw`©ó K‡i wjLyb) Others (Speci<br>........................<br><br>888. |_____|cÖ‡hvR¨ bq [Not applicable]</td></tr>
<tr><td></td><td></td><td>3 eQ‡ii †QvU ev”Pvi cvqLvbv ‡djvi ¯’vbwU ivbœv Ni †_‡K KZUzKz `~‡i?</b> <b>[</b>How far away is this disposal site to the kitchen?]</td><td>|_____| <b>K`g`~‡i?[steps]<br><br>888. cÖ‡hvR¨ bq [Not applicable]</td></tr>
<tr><td></td><td></td><td>|_____| Avcbvi Lvbvq 3-<8 eQ‡ii Ab¨ †Kvb wkï Av‡Q wK? [Is there any other child between the age of 3-<8 years?]</td><td>1 n¨uv [Yes]<br><br>0. bv [No]..........913bs cÖ‡kœ P‡j hvb (skips to 913)</td></tr>
<tr><td></td><td></td><td>Avcbvi Lvbvq 3-<8 eQ‡ii ‡h wkïwU Av‡Q Zvi eqm KZ (gv‡m)? (GKvwaK wkï _vK‡j me‡P‡q eo ev”Pvi †¶‡Î wR‡Ám Ki“b)</td><td>.......................... eQi<br><br>.......................... gvm</td></tr>
<tr><td></td><td></td><td>|_____| Avcbvi wkï (3-8 eQ‡ii) me©†kl ‡Kv_vq cvqLvbv K‡i‡Q? (GKvwaK wkï _vK‡j me‡P‡q eo ev”Pvi †¶‡Î wR‡Ám</td><td>cwU (DVv‡b) [Potty (in the courtyard)]<br><br>cwU (N‡ii wfZ‡i) [Potty (inside the house)]</td></tr>
</table>

| | | | |
|---|---|---|---|
| | | **Ki"b)** [Where did your oldest child (aged 3-7) defecate the last time?] | **DVv‡b (cwU Qvov)** [Courtyard (without potty)]<br><br>**N‡ii wfZ‡i (cwU Qvov)** [Inside the house (without potty)<br><br>**DVv‡bi evwn‡i †Lvjv RvqMv‡Z** [Open space outside the yard<br><br>**‡Svc-Sv‡o/R½‡j** Bush/jungle<br><br>**cvqLvbvq/Uq‡j‡U** [In toilet] ...**913bs cÖ‡kœ P‡j hvb** (Ski 913)<br><br>**Kv_v/b¨vwc/Wvqcvi** Katha/nappy/diaper<br><br>77. **Ab¨vb¨ (eY©bv wjLyb)** [Other (Please describe)]....................<br><br>99. **Rvwb bv** [Don't Know].............**913bs cÖ‡kœ P‡j h** to 913) |
| | | \|_____\|**†mB cvqLvbv wK Kiv n‡qwQj?** [What is done to the feces?] | <u>**n¨uv [Yes].....1, bv [No]......0**</u><br><br>1. **†hLv‡b cvqLvbv K‡iwQj †mLv‡bB †d‡j ivLv n‡qwQj** [I there]**912.a bs cÖ‡kœ P‡j hvb** (Skip to 912.a)<br><br>2. \|_____\| **Uq‡jU/cvqLvbvi wfZ‡i †djv/†avqv n‡qwQj** [Put/rinsed into toilet or latrine]<br><br>3. \|_____\|**‡Wª‡b/b`©gvi wfZ‡i †djv/†avqv n‡qwQj** [Put/rinsed into drain or ditch]<br><br>4. \|\_\_\_\_\_\|**‡Svc-Sv‡o/R½‡j †djv n‡qwQj** [Thrown into Bush/jungle<br><br>5. \|\_\_\_\_\_\|**gqjv AveR©bvi g‡a¨ †djv n‡qwQj** [Thrown in garbage]<br><br>6. \|\_\_\_\_\_\|**wbw`ó© M‡Z© †djv n‡qwQj** [Thrown into a pit for child's feces]<br><br>7. \|_____\|**gvwUi bx‡P cy‡Z †djv n‡qwQj** [Buried]<br><br>77. \|_____\|**Ab¨vb¨ (eY©bv wjLyb)** [Other (specify)].......<br><br>99. \|_____\| **Rvwb bv** [DK]**913 bs cÖ‡kœ P‡j hvb** (Skip to |
| | | **Avcwb wKfv‡e cvqLvbv n¨vÛwjs (e¨e¯'vcbv )K‡ib?**How did you handle | ☐ 1 **bMœ/Lvwj nv‡Z** Hands only |

| | | | | | |
|---|---|---|---|---|---|
| | | the feces? Do Not Read Responses. Mark All that Apply (1 = Yes) | | | (bare hands) |
| | | | ☐ | 2 | **nv‡Z Ges Kvco/cvZv/KvMR** Hands and cloth / paper / leaves |
| | | | ☐ | 3 | **†djv‡bv mvgMÖx gvwU‡Z cyu‡Z** Scrap material to scoop feces |
| | | | ☐ | 4 | **cwU** Potty |
| | | | ☐ | 5 | **¯’vbxq K…wlKvR-Gi nvwZqvi** Local agricultural hoe/instrument |
| | | | ☐ | 6 | **†mwb ¯‹zc**Sani-scoop |
| | | | ☐ | 7 | **‡Kvb wKQyB Kiv nq bv** Did nothing |
| | | | ☐ | 77 | Ab¨vb¨ (D‡jøL Ki“b) Others (specify)___________ |
| | | | ☐ | 99 | **Rvwb bv**Don't know / not sure |
| C.910.b | 912.a | **cvqLvbv Kivi ci KZÿY GB cvqLvbv GLv‡b c‡i wQj?** How long after defecation did you dispose of the feces? | \|____\|Hour **N›Uv**<br>\|____\|Minute **wgwbU** | | |

| | | | |
|---|---|---|---|
| Child Potty Use<br><br>**( শিশুর পটি ব্যবহার সর্ম্পকীত তথ্য)** | | | |
| Administer to: All study households ( সকল খানায় হবে) | | | |
| C.815 | | Does your household have a potty that children use for defecation? আপনার খানাতে কি শিশুদের পায়খানা করার পটি আছে? | 1 Yes হ্যাঁ<br>2 No না**(Skip to 919)** (919নং প্রশ্নে যান)<br>99 Don't know / not sure জানি না **(Skip t[illegible]** (919নং প্রশ্নে যান) |
| C.816 | | In the last week, how often did your child use the potty? গত এক সপ্তাহের মধ্যে, শিশুটি কতবার পটি ব্যবহার করছে?<br><br>[ *If multiple children use the potty,* | 1 Every time প্রতিবার<br>2 More than half of all defecation event not every time যতবার পায়খানা করেছে তার ৫০ ভাগ বা তা |

| | | | |
|---|---|---|---|
| | | *ask about the youngest child* ][যদি একাধিক শিশু ব্যবহার করে থাকে তবে যে সবচেয়ে বেশী ব্যবহার করে সেই শিশুর জন্য প্রশ্ন করুন] | কিন্তু প্রতিবার নয়<br>3 যতবার পায়খানা করেছে তার ৫০ ভাগের কম মাঝেমাঝে Less than half of all defecation events<br>4 আগে করেছি কিন্তু এখন আর করি না use it, but no longer use it **(Skip to 919)** (919নং প্রশ্নে য<br>5 কখনই না Never **(Skip to 919)** (919নং<br>99 জানি না Don't know / not sure **(Skip to** (919নং প্রশ্নে যান) |
| C.817 | | What is the age of the child (or children) who is using the potty? যে শিশুটি (বা শিশুরা) পটি ব্যবহার করছে তার বয়স কত? | Mark All that<br>Yes = 1 হ্যাঁ = 0<br>1 ☐ < 1 year ১ বছরের কম<br>2 ☐ 1- < 3 years ১-<৩ বছর<br>3 ☐ 3- < 5 years ৩-<৫ বছর<br>4 ☐ >= 5 years>=৫ বছর |

Sani Scoop Use

**(সেনিস্কুপ ব্যবহার)**

Administer to: All households ( সকল খানায় হবে)

| | | | |
|---|---|---|---|
| C.821 | | ☐Does your household have a dedicated tool [sani scoop] to clean up feces around your household?<br>আপনার খানার চারপাশের পায়খানা পরিষ্কার করার জন্য আপনার খানায় কোন নির্দিষ্ট কিছু আছে কি? | 1 Yes হ্যাঁ<br>2 No না (1001নং প্রশ্নে যান)**(Skip to 1001)**<br>99 Don't know / not sure জানি না (1001 নং প্রশ্নে যান)**(Skip to 1001)** |
| C.822 | | How often do you use the [sani scoop]? আপনি কখন কখন পায়খানা পরিষ্কার করার জন্য কোদাল/সেনিস্কুপ ব্যবহার করেন? | 1 Multiple times per day প্রতিদিন কয়েকবার<br>2 Once per day প্রতিদিন একবার<br>3 A few times each week প্রতি সপ্তাহে কয়েকবার<br>4 Less than once per week সপ্তাহে একবারের ক |

| | | | |
|---|---|---|---|
| | | | 5 Used to use it, but no longer use it প্রথমদিবে করেছে কিন্তু এখন আর করে না(1001 নং প্রশ্নে যান)(Skip to 1001)<br>6 Never কখনই না(1001 নং প্রশ্নে যান)(Skip to 1 |
| C.823 | | What do you use the [sani scoop] for? ☝Do Not Read Responses.<br>আপনি কি কি কাজেকোদাল/সেনিস্কুপ ব্যবহার করেন? (একাধিক উত্তর হতে পারে)☝ উত্তর পড়ে শোনাবেন না | Mark All that Apply. Yes = 1 হ্যাঁ = 1, না = 0<br>1 ☐ Clean up animal feces জীবজন্তুর পায়খানা ফেল<br>2 ☐ Clean up child feces বাচ্চার পায়খানা ফেলার ব<br>3 ☐ Clean up garbage ময়লা আবর্জনা ফেলার কাজে<br>4 ☐ Take the scoop to the field (for work) কৃষিকা<br>5 ☐ Digging / gardening গর্ত তৈরীর কাজে<br>77 ☐ Other (specify) অন্যান্য (বর্ণনা লিখুন)_______ |

**Wash Benefit Module 10 Water Access**

আপনাকে ধন্যবাদ। আমি আপনাকে এবং আপনার খানার সদস্যদের বাড়ীতে খাবার পানি বিষয়ে কিছু প্রশ্ন করতে চাই।

Thank you very much. Now, I would like to ask you some questions about the water you and your family drink at home.

| WATER TREATMENT, STORAGE, AND QUALITY (All households) | | | |
|---|---|---|---|
| SAY | Thank you very much. Now, I would like to ask you some questions about the water you and your family drink at home. | | |
| C.1001 | 1001. | আপনি কিভাবে খাবার পানি সংরক্ষন করেন ? [How do you store drinking water?<br>👁প্রশ্ন করুন এবং সম্ভব হলে পর্যবেক্ষন করুন ASK THE QESTION AND OBSERVE IF POSSIBLE | [1] প্লাষ্টিক অথবা ধাতব পাত্রে IN PLASTIC OR METAL CONTAINERS<br>[2] মাটির পাত্রে IN CLAY POTS<br>[3] ছাদে পানির ট্যাংক ROOF TANK OR CISTERN<br>[4] পানি সংরক্ষণ করা হয় নাDO NOT STORE WATER |
| C.1002 | 1002. | এই খানায় ০-<৩৬ মাসের কোন শিশু আছে কি যে পানি পান করে ?Is there a child 0-<36 months in this household who drinks water? (if there is no child 0-<36 months, ask this question for a child 36-<60 months) | [1] হ্যাঁ YES<br>[2] না NO<br>[88] Not applicable প্রযোজ্য নয় |
| C.1003 | 1003. | **IF YES→**প্রশ্ন করুন এবং পর্যবেক্ষন করুনঃ যদি আপনার শিশু (<৩ বছর) এখন পানি খেতে চাইত তাহলে আপনি কিভাবে তাকে পানি দিতেন তা আমাকে দেখানASK: If your child 0-36 months wanted a drink of water right now, could you show me how you would give it to him/her? (If no child 0-36 months lives in the compound, ask this question for a child 36-<60 months)<br>**IF NO→** ASK: যদি আপনি এখন পানি খেতে চাইতেন তাহলে আপনি কিভাবে পানি নিতেন তা আমাকে দেখানIf you wanted a drink of water right now, could you show me how you would get it?<br>উত্তরদাতার পানি দেওয়ার প্রক্রিয়াকে পর্যবেক্ষণ/অনুসরণ করুন । যদি অন্য ঘরে পানি সংরক্ষিত এবং সেখানে প্রবেশাধিকার থাকে তাহলে প্রক্রিয়াকে পর্যবেক্ষণ/অনুসরণ করুন । অন্যথায় ৯৯ কে গোলাকার করতে হবে। ATTEMPT TO FOLLOW RESPONDENT WHEN SHE GOES TO RETRIEVE THE WATER. IF WATER IS STORED IN ANOTHER ROOM, ASK HER IF IT IS OKAY TO ENTER THAT ROOM. CIRCLE “99” IF YOU CANNOT OBSERVE. | |

| | | | | | |
|---|---|---|---|---|---|
| | | 👁 উত্তরদাতা পানি নেওয়ার আগে কি করেছিল ?WHAT DID RESPONDENT DO BEFORE TAKING THE WATER? | হ্যাঁ YES | না NO | জানি নাD/K |
| 2 | 2 | গ্লাস বা পাত্রটিতে পানি ঢালার পূর্বে গ্লাস/পাত্রটি খাবার পানি দিয়ে ধুয়েছিলRINSED GLASS/CONTAINER WITH DRINKING WATER BEFORE FILLING | [1] | [2] | [99] |
| 3 | 3 | পানি ঢালার পূর্বে হাত ধুয়েছিলWASHED HANDS WITH WATER BEFORE DRINKING WATER WAS OBTAINED | [1] | [2] | [99] |
| 4 | 4 | পানি ঢালার পূর্বে সাবান দিয়ে হাত ধুয়েছিলWASHED HANDS WITH SOAP BEFORE DRINKING WATER OBTAINED | [1] | [2] | [99] |
| | | 👁 উত্তরদাতা কোথা থেকে পানিসংগ্রহ করেছিল?FROM WHERE DID THE RESPONDENT TAKE THE WATER? | হ্যাঁ YES | না NO | জানি না D/K |
| 5 | 5 | সরাসরি পানির উৎস থেকে BROUGHT DIRECTLY FROM WATER SOURCE | [1] | [2] | [99] |
| 6 | 6 | সরাসরি খোলা পানির পাত্র থেকে BROUGHT IRECTLY FROM uncoveredSTORAGE CONTAINER | [1] | [2] | [99] |
| 7 | 7 | সরাসরি পানির ফ্লিটার থেকে BROUGHT DIRECTLY FROM WATER FILTER | [1] | [2] | [99] |
| | 8 | সরাসরি ঢেকে রাখা পানির পাত্র থেকে STORED WATER WAS COVERED | [1] | [2] | [99] |
| | | 👁 উত্তরদাতা কিভাবে গ্লাসে/কাপে/পাত্রে পানি দিয়েছিল? HOW DID SHE GET THE WATER INTO THE CUP? | হ্যাঁ YES | না NO | জানি না D/K |
| 9 | 9 | গ্লাস বা পাত্রটিতে ঢালা পানির ভিতরে তার হাত বা হাতের আঙ্গুল লেগেছিল কি?HANDS TOUCHED / CONTACTED THE DRINKING WATER | [1] | [2] | [9 ] |
| 10 | 10 | গ্লাস বা পাত্রটিতে পানি ভরার সময় তা পানির পাত্রের ভিতর ডুবিয়ে পানি ভরা হয়েছিলCONTAINER/GLASS DIPPED INTO WATER CONTAINER | [1] | [2] | [99] |
| 11 | 11 | গ্লাসে পানি ভরার জন্য লম্বা হাতলযুক্ত কোন চামচ/মগ ব্যবহার করা হয়েছিলLADLE USED TO OBTAIN WATE | [1] | [2] | [99] |
| 12 | 12 | পানির পাত্র থেকে সরাসরি পানি ঢেলেছিলWATER POURED FROM CONTAINER | [1] | [2] | [99] |
| 13 | 13 | সরাসরি টিউবওয়েল/ পানির উৎস থেকে এনেছেWATER POURED FROM TAP | [1] | [2] | [99] |

| | | | | |
|---|---|---|---|---|
| C.1004 | 1004. | আপনি/এ খানার অন্যকেউ কতক্ষণ আগে এই সংরক্ষণকৃত পানি সংগ্রহকরেছেন? How long ago did you or somebody in your home collect this water? ( 99 = Don't knowজানি না; 87 = No Water কোন পানি নাই, 88 Not applicableপ্রযোজ্য নয়<br>) If 88, skip to 1007 | A.\|__\|__\| ঘন্টা HOURS<br>B.\|__\|__\| দিনDAYS | |
| C.1005 | 1005. | এই খাবার পানিনিরাপদ করার জন্য কোন কিছু করেছেন কি?Have you done anything to make this water less cloudy or safer to drink? | [1] হ্যাঁYES<br>[2] না NO<br>[99] জানি না DON'T KNOW/NOT SURE | <br>→ SKIP to 1007<br>→ SKIP to 1007 |
| C.1006 | 1006. | কোন পদ্বতি কি আপনি ব্যবহার করেন? What method(s) did you | | |

| | | | | |
|---|---|---|---|---|
| | | use?<br>উত্তর পড়ে শুনাবেন না \|উত্তরদাতাকে উত্তর বলতে সাহায্য করা যাবে তার উত্তর দেয়া শেষ হলে ।DO NOT READ RESPONSES. PROBE UNTIL RESPONDENT IS FINISHED. CIRCLE ALL THAT APPLY. | | |
| 1 | 1 | আকুয়াট্যাবসAquatabs | [1] হ্যাঁ Yes [0] না No | |
| 2 | 2 | ক্লোরিন (তরল) মিশিয়ে নেই (ওয়াটার গার্ড নয়)Waterguard / bottled chlorine | [[1] হ্যাঁ Yes [0] না No | |
| 3 | 3 | পানি ফুটিয়ে নেইBoil | [1] হ্যাঁ Yes [0] না No | |
| 4 | 4 | কাপড়ে ছেকে নেইStrain it through clothor other material | [1] হ্যাঁ Yes [0] না No | |
| 5 | 5 | পানির ফিল্টার ব্যবহার করে (সিরামিক, বালু ইত্যাদি) Water filter [ceramic, sand, composite] | [1] হ্যাঁ Yes [0] না No | |
| 6 | 6 | সূর্যের আলোতে জীবানুমুক্তকরণSolar disinfection (SODIS) | [1] হ্যাঁ Yes [0] না No | |
| 7 | 7 | থিতিয়ে নেই Let it stand and settle | [1] হ্যাঁ Yes [0] না No | |
| 8 | 8 | স্যান্ড ফিল্টার এর মাধ্যমে বিশুদ্ধ করে নেইBiosand filter | [1] হ্যাঁ Yes [0] না No | |
| 9 | 9 | ফিটকিরি যোগ করে Coagulant (alum) | [1] হ্যাঁ Yes [0] না No | |
| 10 | 10 | পার PUR (flocculant + disinfectant) | [1] হ্যাঁ Yes [0] না No | |
| C.1007 | 1007. | আপনি কখনও আপনার খাবার পানি পরিশোধন করার জন্য কিছু করেছিলেন কি?Do you ever treat your drinking water or do anything to make it less cloudy? | [1] হ্যা YES<br>[0] নাNO | → 1010 নং এ চলে<br>SKIP TO 1010 |
| C.1008 | 1008. | সর্বশেষ কখন আপনিআপনার খাবার পানি পরিশোধন করেছিলেন ? When was the last time you treated your water or did anything to make it less cloudy?<br><br>উত্তর পড়ে শুনানো যাবে না DO NOT READ RESPONSES | [1] আজ TODAY<br>[2] গতকালYESTERDAY<br>[3] গত সপ্তাহের মধ্যে WITHIN THE PAST WEEK<br>[4] গত দুই সপ্তাহের মধ্যে WITHIN THE PAST 2 WEEKS<br>[5] গত মাসের মধ্যে WITHIN THE PAST MONTH<br>[6] গত বছরের মধ্যে WITHIN THE PAST YEAR<br>[99] জানি না/নিশ্চিত না DON'T KNOW/NOT SURE | |
| C.1009 | 1009. | কোন কোন উপায়ে আপনি পানি পরিশোধন করে থাকেন ? What are all the ways you treat your drinking water?<br>উত্তর পড়ে শুনানো যাবে না । উত্তর দেয়া শেষ হলে উত্তর পেতে সাহায্য করা যেতে পারে ।DO NOT READ RESPONSES. PROBE UNTIL RESPONDENT IS FINISHED. CIRCLE ALL THAT APPLY. | | |
| 1 | 1 | আকুয়াট্যাবসAquatabs | [1] হ্যাঁ Yes [0] না No | |
| 2 | 2 | ক্লোরিন (তরল) মিশিয়ে নেই (ওয়াটার গার্ড নয়)Waterguard / bottled | [[1] হ্যাঁ Yes [0] না No | |

| | | | |
|---|---|---|---|
| | | chlorine | |
| 3 | 3 | পানি ফুটিয়ে নেইBoil | [1] হ্যাঁYes [0] না No |
| 4 | 4 | কাপড়ে ছেকে নেইStrain it through clothor other material | [1] হ্যাঁYes [0] না No |
| 5 | 5 | পানির ফিল্টার ব্যবহার করে (সিরামিক, বালু ইত্যাদি) Water filter [ceramic, sand, composite] | [1] হ্যাঁYes [0] না No |
| 6 | 6 | সূর্যের আলোতে জীবানুমুক্তকরণSolar disinfection (SODIS) | [1] হ্যাঁYes [0] না No |
| 7 | 7 | থিতিয়ে নেই Let it stand and settle | [1] হ্যাঁYes [0] না No |
| 8 | 8 | স্যান্ড ফিল্টার এর মাধ্যমে বিশুদ্ধ করে নেইBiosand filter | [1] হ্যাঁYes [0] না No |
| 9 | 9 | ফিটকিরি যোগ করে Coagulant (alum) | [1] হ্যাঁYes [0] না No |
| 10 | 10 | PUR (flocculant + disinfectant) | [1] হ্যাঁYes [0] না No |

| পানের পানির উৎস **DRINKING WATER SOURCES (ALL HOUSEHOLDS)** | | | |
|---|---|---|---|
| উত্তরদাতাকে তার পানির প্রধান এবং দ্বিতীয় উৎস বিষয়ে প্রশ্ন করুন ASK THE FOLLOWING QUESTIONS ABOUT THE RESPONDENT'S PRIMARY AND SECOND | | | |
| C.1016 | 1010. | আপনার খানায়ব্যবহৃত পানির প্রধান উৎস কি ?What type of water source does your household collect most of the water that you use from? | 1 অগভীর টিউবওয়েল Shallow tubewell<br>2 গভীর টিউবওয়েল Deep tubewell<br>3 ঘরের ভিতর ট্যাপ বা পাইপের পানি Piped into home<br>4 উঠানে ট্যাপ বা পাইপের পানি Piped into yard<br>5 গভীর পাতকুয়া (উঠানে)Borewell in yard<br>6 পাবলিক ট্যাপ Public tap<br>7 পাবলিক গভীর পাতকূয়া Public borewell<br>8 অগভীর পাকা পাতকূয়া Shallow well with concrete reinforcement<br>9 অগভীর কাঁচা পাতকূয়া Shallow well (no concrete)<br>10 নদী/বাধ/লেক/পুকুর/সেচ নালাথেকে সরাসরি সংগৃহীত পানি River, lake, stream<br>11 অন্যান্য (লিখুন) Other: _____________ |
| C.1017 | 1011. | আপনি আমাকে বলবেন আপনার বাড়ী থেকে প্রধান পানির উৎসে একবার যেতে কতক্ষণ সময় লাগে? Can you tell me how long it takes you to walk one-way to this PRIMARY water source from your home? | \|__\|ঘনটা Hours<br>\|__\|মিনিট Min |
| C.1018 | 1012. | আপনি আমাকে বলবেন কোন কোন কাজের জন্য এই উৎসের পানি ব্যবহার করেন Can you please tell me whether you use water from this source for… | |
| 1 | 1 | পানি পান Drinking | [1] হ্যাঁYes [0] না No |
| 2 | 2 | রান্না-বান্না/বাসন ধোয়া Cooking/ washing dishes | 1] হ্যাঁYes [0] না No |

| | | | |
|---|---|---|---|
| 3 | 3 | কাপড় ধোয়া Laundry | 1] হ্যাঁ Yes [0] না No |
| 4 | 4 | গোসল/হাতধোয়া Bathing/hand washing | 1] হ্যাঁ Yes [0] না No |
| 5 | 5 | কৃষিকাজ/বাগান Irrigate crop land or garden | 1] হ্যাঁ Yes [0] না No |
| 6 | 6 | পশু-পালন Water for livestock | 1] হ্যাঁ Yes [0] না No |
| C.1019 | 1013. | আপনার খানাকে এই পানির উৎস ব্যবহারের জন্য কি কোন প্রকার টাকা দিতে হয়? your household pay any money to use this source? | [1] হ্যাঁ Yes [0] না No |
| C.1020 | 1014. | আপনার খানায় পানি ব্যবহারের জন্য কি ২য় কোন উৎস ব্যবহার করতে হয়? যদি হয় তাহলে কি ধরনের ? Does your household currently use a secondary water source? If so, what type is it? | 1. অগভীর টিউবওয়েল Shallow tubewell<br>2. গভীর টিউবওয়েল Deep tubewell<br>3. ঘরের ভিতর ট্যাপ বা পাইপের পানি Piped into home<br>4. উঠানে ট্যাপ বা পাইপের পানি Piped into yard<br>5. গভীর পাতকূয়া (উঠানে)Borewell in yard<br>6. পাবলিক ট্যাপ Public tap<br>7. পাবলিক গভীর পাতকূয়া Public borewell<br>8. অগভীর পাকা পাতকূয়া Shallow well with concrete reinforcement<br>9. অগভীর কাঁচা পাতকূয়া Shallow well (no concrete)<br>10. নদী/বাধ/লেক/পুকুর/সেচ নালাথেকে সরাসরি সংগৃহীত পানি River, lake, stream<br>11. অন্যান্য (লিখুন) Other: _____________<br>88] ২য় কোন পানির উৎস ব্যবহার করতে হয় না Don't use any secondary wate |
| C1021 | 1015. | আপনি আমাকে বলবেন আপনার বাড়ী থেকে ২য় পানির উৎসে একবার যেতে কতক্ষণ সময় লাগে? Can you tell me how long it takes you to walk one-way to this SECONDARY water source from your home? | \|__\|ঘন্টা Hours<br>\|__\|মিনিট Min |
| C1022 | 1016. | আপনি আমাকে বলবেন কোন কোন কাজের জন্য এই উৎসের পানি ব্যবহার করেন Can you please tell me whether you use water from this source for… | |
| 1 | 1 | পানি পান Drinking | [1] হ্যাঁ Yes [0] না No |

| | | | |
|---|---|---|---|
| 2 | 2 | রান্না-বান্না/বাসন ধোয়া Cooking/ washing dishes | 1] হ্যাঁ Yes [0] না No |
| 3 | 3 | কাপড় ধোয়া Laundry | 1] হ্যাঁ Yes [0] না No |
| 4 | 4 | গোসল/হাতধোয়া Bathing/hand washing | 1] হ্যাঁ Yes [0] না No |
| 5 | 5 | কৃষিকাজ/বাগান Irrigate crop land or garden | 1] হ্যাঁ Yes [0] না No |
| 6 | 6 | পশু-পালন Water for livestock | 1] হ্যাঁ Yes [0] না No |
| C1023 | 1017. | আপনার খানাকে এই পানির উৎস ব্যবহারের জন্য কি কোন প্রকার টাকা দিতে হয়? your household pay any money to use this source? | 1] হ্যাঁ Yes [0] না No |
| | 1017.a | আপনার খানার খাবার পানির প্রধান উৎস কি? What is your main drinking water source? | 1. অগভীর টিউবওয়েল Shallow tubewell<br>2. গভীর টিউবওয়েল Deep tubewell<br>3. ঘরের ভিতর ট্যাপ বা পাইপের পানি Piped into home<br>4. উঠানে ট্যাপ বা পাইপের পানি Piped into yard<br>5. গভীর পাতকূয়া (উঠানে)Borewell in yard<br>6. পাবলিক ট্যাপ Public tap<br>7. পাবলিক গভীর পাতকূয়া Public borewell<br>8. অগভীর পাকা পাতকূয়া Shallow well with concrete reinforcement<br>9. অগভীর কাঁচা পাতকূয়া Shallow well (no concrete)<br>10. নদী/বাধ/লেক/পুকুর/সেচ নালাথেকে সরাসরি সংগৃহীত পানি River, lake, stream<br>11. অন্যান্য (লিখুন) Other: _____________<br>88] Not applicable |
| C1024 | 1018. | আপনার খানা কি কখনও বৃষ্টির পানি সংগ্রহ করত ? Does your household collect rainwater? | 1] হ্যাঁ Yes [0] না No |
| C1025 | 1019. | আপনার খানা কি বর্তমানে বৃষ্টির পানি সংগ্রহ করে ? Is your household currently collecting rainwater? | 1] হ্যাঁ Yes [0] না No if no, skip to 1301 |
| C1026 | 1020. | আপনি আমাকে বলবেন কি আপনার খানা কি কি কাজে বৃষ্টির পানি ব্যবহার করে ? Can you please tell me whether you use rainwater for | |
| 1 | 1 | পানি পান Drinking | [1] হ্যাঁ Yes [0] না No |

| | | | |
|---|---|---|---|
| 2 | 2 | রান্না-বান্না/বাসন ধোয়া Cooking/ washing dishes | 1] হ্যাঁ Yes [0] না No |
| 3 | 3 | কাপড় ধোয়া Laundry | 1] হ্যাঁ Yes [0] না No |
| 4 | 4 | গোসল/হাতধোয়া Bathing/hand washing | 1] হ্যাঁ Yes [0] না No |
| 5 | 5 | কৃষিকাজ/বাগান Irrigate crop land or garden | 1] হ্যাঁ Yes [0] না No |
| 6 | 6 | পশু-পালন Water for livestock | 1] হ্যাঁ Yes [0] না No |
| | | | **Skip to 1301** |
| **C.1010.** | 1021. | আপনার খানায় কত ঘন ঘন খাবার পানিতে আকুয়াট্যাবস মেশানো হয়? How frequently do you treat your water using [Aquatabs]? | Do Not Read Responses.<br>1 যতবার পানি সংগ্রহ করা হয়/সবসময়Every time they collect water<br>2 মাঝেমাঝেSometimes / occasionally<br>3 প্রথমদিকে দিয়েছিলাম কিন্তু এখন আর দেই নাTreated water in the beginn[illegible] more<br>4 আকুয়াট্যাবস দিয়ে কখনও পানি পরিশোধন করি নাইNever treated water [illegible]<br>99 জানি না Don't know / not sure |
| **C.101[illegible]** | 1022. | আপনার খানায় সংরক্ষনকৃত খাবার পানি কি কুয়াট্যাবস দ্বারা পরিশোধন করা হয়েছে? Is the drinking water stored in your household today treated with [Aquatabs]? | 1 হ্যাঁ, সবটুকু পানিYes, all of it<br>2 হ্যাঁ, কিছু পানি Yes, some of it<br>3 নাNot treated<br>4 খানায় কোন পানি নাই No water in the house<br>99 জানি না Don't know |
| **C.101[illegible]** | 1023. | পর্যবেৰন করুন: আকুয়াট্যাবস দ্বারা পরিশোধিত পানির পাত্র দেখাতে বলুন: পানির পাত্র ঢাকা ছিল কী? Ask to see the treated water. Observe: Is the water covered? | 1 হ্যাঁYes<br>2 নাNo<br>3 খানায় পাত্রে কোন পানি নাই No water stored in the house<br>88 প্রযোজ্য নয়/প্রত্যাখ্যান Not applicable / refused |
| **C.101[illegible]** | 1024. | পানি পরিশোধন করার জন্য কত সময় আগেআকুয়াট্যাবস ব্যবহার করা হয়েছিল? Approximately how long ago did you treat the water with [Aquatabs]? | ঘিন টা HH: মনিট MM ☐☐:☐☐ ( 9 9 : 9 9 Don't know ) |
| খানায় অনুজীবি সংক্রান্ত নমূনা (খানা দৈবচয়িতভাবে নির্বাচিত হবে)**HOUSEHOLD MICROBIOLOGICAL SAMPLE (ADMINISTER TO ALL HOUSEHOLDS)** | | | |

| | | | | |
|---|---|---|---|---|
| C.1501 | 1025. | ব্যাকটেরিয়া আছে কিনা জানার জন্য আমি আপনার খাবার পানির কিছু পরিমান সংগ্রহ করতে পারি কী?<br>May I collect a small sample of your drinking water to test for bacteria in our office? | [1] হ্যাঁ YES, SAMPLE COLLECTED<br>[2] না NO/REFUSED | <br>→ |
| C.1502 | 1026. | খানার নমুনা সংগ্রহীত হয়েছে HH sample collected (DD/MM/YY, HH:MM) | খানার নমুনা পানির রেকর্ড HH Water Sample Time Log (can be automated in a bar-[illegible]<br><br>\| \| DD/MM/YY \| HH:MM \|<br>\| A \| □□/□□/□□ খানার পানির নমুনা সংগ্রহের তারিখ HH sample collected \| □□:□□ সময় \|<br>\| B \| □□/□□/□□ খানার পানির নমুনা ল্যাবে পৌছানোর তারিখ HH sample arrived at lab \| □□:□□ সময় \|<br>\| C \| □□/□□/□□ খানার পানির নমুনা এনালাইসিসের তারিখ HH sample analyzed \| □□:□□ সময় \| | |
| c.1503 | 1027. | খানার পানির নমুনা, ফিল্টারকৃত পানির পরিমান (মিলি)<br>HH sample, volume of water filtered (in milliliters) | □□□মিলি ml | |
| C.1504 | 1028. | খানার পানির নমুনার ল্যাব রেজাল্ট, ই.কোলাই (*E. coli)*<br>(5555 = অসংখ্য, 9999 = এনালাইসিস করা হয় নাই<br>Household sample lab result, *E. coli* (5555 = Too numerous to count, 9999 = Not analyzed) | □□□□ (CFU / 100 ml) | |
| C.1505 | 1029. | খানার নমুনার অবস্থা<br>HH sample qualifier | 1. ডিটেকশন লেভেলের নীচে Below detection limit<br>2. ডিটেকশন লেভেলের উপরে Above detection limit<br>3. নমুনা এনালাইসিস করা হয় নাই Sample could not be analyzed | |
| | | প্রস্তুতির আগে ক্লাষ্টার/খানা আই ডি বোতলে/হুয়েলপ্যাকে স্পষ্ট অক্ষরে লিখুন** **WRITE THE CLUSTER/HH ID <u>VERY CLEARLY</u> ON T[illegible] PROCEEDING**** | | |
| | | খানার সমন্বিত পানির উৎসের নমুনা PAIRED SOURCE WATER SAMPLE FOR HH WATER SAMPLE<br>। যদি উত্তরদাতা বলেন যে ভাগের/সেয়াড পানির উৎস থেকে পানি সংগ্রহ করা হয়েছে তাহলে **701** নং প্রশ্নে চলে যান।IF RESPONDENT REPORTED C[illegible] SHARED SOURCE→ **SKIP TO 701** | | |

**Wash Benefit Module 13 MEASURES FOR SPILLOVER**
সকল খানার জন্য প্রযোজ্য Administer to: All study households

| | | | |
|---|---|---|---|
| **C.1301.** | 1301. | আপনার সবচেয়ে কাছের হেলথ ফ্যাসিলিটি (স্বাস্থ্যসেবা পাওয়ার সুযোগ)-এর নাম কি?যদি না জানে/নিশ্চিত না হয় তবে ৯৯ বসান। খানার কেউ ডাক্তার হলে বা ডাক্তার খানায় এসে চিকিৎসা করলে ৫৫ লিখুন।What is the name of your nearest health facility?<br>Enter "99" if Don't know / not sure | |
| **C.1301.a** | 1302. | গত ১ বছরে আপনি/আপনার শিশু মোটামুটি কতবার যে কোন অসুখে এই সুযোগ গ্রহন করেছিলেন? <u>In the past year</u>, (since this time last year) approximately how many times did you visit this health facility for any reason (including with your children)? | \|__\|__\|বার TIMES |
| **C.1301.b** | 1303. | আপনি সেখানে কিভাবে যান? What is your primary mode of travel to the health facility? | ১. পায়ে হেঁটে<br>**2.** রিকশা/ভ্যান/নৌকা<br>**3.** ব্যাটারি চালিত অটোরিকশা<br>**4.** ট্যাম্পু/লেগুনা/ সিএনজি/ বাস<br>**5.** প্রযোজ্য নয় |
| **C.1301.c** | **1303.a** | আনুমানিক সেখানে যেতে কতক্ষণ সময় লাগে(আপনার খানা থেকে শুধু যেতে কত মিনিট সময় লাগে) ? যদি না জানে/নিশ্চিত না হয় তবে ৯৯ বসান। Approximately how long does it take to travel there (in minutes, one-way, from where we are now)?<br><br>Enter "999" if Don't know / not sure | \|__\|__\|__\| মিনিটে MINUTES |
| **C.1302.** | 1304. | আপনার কাছের প্রাক- প্রাইমারী স্কুলের -এর নাম কি ? যেখানে আপনার শিশু যেতে পারে/বড় হলেও যেতে পারবে । যদি না জানে/নিশ্চিত না হয় তবে ৯৯, যদি না থাকে তবে ৮৮ বসান।What is the name of the ECD center [preschool] where your children go (or will go when they are old enough)? | |
| **C.1303.** | 1305. | আপনার কাছের প্রাইমারী স্কুলের -এর নাম কি ? যেখানে আপনার শিশু যেতে পারে/বড় হলেও যেতে পারবে । What is the name of the primary school where your children go (or will go when they are old enough)? | |
| **C.1304.** | 1306. | আপনার কাছের বাজার -এর নাম কিযেখানে আপনি সবসময়ই কেনা-বেচা করতে যান? । What is the name of the market that you most often go to when you buy or sell things? | |
| **C.1304.a** | 1307. | কতবার আপনি সেখানে যান? How often do you go to that market? | 0 কখনই না Never<br>1 সপ্তাহে একবার অথবা তার অধিক Once per week or more<br>2 প্রতি ২ সপ্তাহে একবার Once every 2 weeks<br>3 প্রতি ৩ সপ্তাহে একবার Once every 3 weeks<br>4 প্রতি ৪ সপ্তাহে একবার(প্রতি মাসে) Once every 4 weeks (every month)<br>5 প্রতি মাসে একবারের কম Less than once every month<br>99 জানি না/নিশ্চিত না Don't know / not sure |
| | 1308. | আপনি সেখানে কিভাবে যান? What is your primary mode of travel to the health facility? | **1.** পায়ে হেঁটে<br>**2.** রিকশা/ভ্যান/নৌকা |

| | | | |
|---|---|---|---|
| | | | 3. ব্যাটারি চালিত অটোরিকশা<br>4. ট্যাম্পু/লেগুনা/ সিএনজি/ বাস<br>5. প্রযোজ্য নয় |
| C.1304.b | 1308.a | আনুমানিক সেখানে যেতে কতক্ষণ সময় লাগে(আপনার খানা থেকে শুধু যেতে কত মিনিট সময় লাগে) ? যদি না জানে/নিশ্চিত না হয় তবে ৯৯ বসান। Approximately how long does it take to travel there (in minutes, one-way, from where we are now)?<br><br>Enter "999" if Don't know / not sure | \|__\|__\|__\| মিনিটে MINUTES |
| C.1305. | 1309. | আপনার কাছের মসজিদ/মন্দির/গীর্জা/প্যাগোডা -এর নাম কি? যেখানে আপনি প্রার্থনা করতে যান । What is the name of the church/mosque/temple (if any) that you attend? | |
| C.1305.a | 1310. | কতবার আপনি সেখানে যান? How often do you go to church? | ০. কখনই না Never<br>1 সপ্তাহে একবার অথবা তার অধিক Once per week or more<br>2 প্রতি ২ সপ্তাহে একবার Once every 2 weeks<br>3 প্রতি ৩ সপ্তাহে একবার Once every 3 weeks<br>4 প্রতি ৪ সপ্তাহে একবার(প্রতি মাসে) Once every 4 weeks (every month)<br>5 প্রতি মাসে একবারের কম Less than once every month<br>99 জানি না/নিশ্চিত না Don't know / not sure |
| | 1311. | আপনি সেখানে কিভাবে যান? What is your primary mode of travel to the health facility? | 1. পায়ে হেঁটে<br>2. রিকশা/ভ্যান/নৌকা<br>3. ব্যাটারি চালিত অটোরিকশা<br>4. ট্যাম্পু/লেগুনা/ সিএনজি/ বাস<br>5. প্রযোজ্য নয় |
| C.1305.b | 1312.a | আনুমানিক সেখানে যেতে কতক্ষণ সময় লাগে(আপনার খানা থেকে শুধু যেতে কত মিনিট সময় লাগে) ? যদি না জানে/নিশ্চিত না হয় তবে ৯৯ বসান। Approximately how long does it take to travel there (in minutes, one-way, from where we are now)?<br><br>Enter "999" if Don't know / not sure | \|__\|__\|__\| মিনিটে MINUTES |

**Wash Benefit Module 15 Environmental Sampling (Water, Hands, Sentinel Toys and Fly Density). Administer to all households.**

DAY 1

SECTION 1. DELIVER TOY BALL AND FLY TAPE

PROMPT: Please deliver the toy ball to the target child or respondent.

1.1 Who did you give the ball to?

1 = Target child

2 = Respondent

3 = Other caregiver

4 = Ball not delivered

77 = Other (specify)

1.2 (if question 1.1 is not 4) Record the time that the toy ball was delivered (24H format, HH:MM).

1.3 (if question 1.1 is 4) Why was the toy ball not delivered?

1 = Respondent/caregiver refused

2 = Did not have a toy ball to deliver

77 = Other (specify)

PROMPT: Can you please show me the area where you prepare food?

PROMPT: Hang fly tape as close as possible to the food preparation area If there is more than one food prep area, hang the strips near where food was most recently prepared. Do NOT hang the strips over or near to a cooking fire. Ask the respondents to leave the tape undisturbed.

1.4 Record the time that the fly paper was hung at the food preperation area (24H format, HH:MM)

PROMPT: Can you please show me your primary latrine area?

PROMPT: Hang fly tape as close as possible to the primary latrine area. If there is more than one primary latrine area, hang the strips near the latrine that the respondent used most recently. Ask the respondents to leave the tape undisturbed.

1.5 Record the time that the fly paper was hung at the primary latrine area (24H format, HH:MM)

SECTION 3. WATER SAMPLING

If (target child's name) wanted a drink of water right now, could you show me how you would give it to him/her? [If target child it too young to drink water, ask: If your child < 3 years wanted a drink of water right now, could you show me how you would give it to him/her?] [If the mother has no children < 3 years, ask: If you wanted a drink of water right now, could you show me how you would get it?]

Ask the question and observe.

| | | | | |
|---|---|---|---|---|
| 3.1 | IS WATER FOR DRINKING CURRENTLY AVAILABLE? | YES<br>NO → SKIP TO SECTION 4 | | |
| | WHAT DID RESPONDENT DO BEFORE TAKING THE WATER? | YES | NO | D/K |
| 3.2 | Rinsed glass / cup with drinking water before filling | [1] | [2] | [99] |
| 3.3 | Washed hands with water before drinking water was obtained | [1] | [2] | [99] |
| 3.4 | Washed hands with soap before drinking water obtained | [1] | [2] | [99] |
| | FROM WHERE DID THE RESPONDENT TAKE THE WATER? | YES | NO | D/K |
| 3.5 | Brought directly from the water source | [1] | [2] | [99] |
| 3.6 | Brought directly from water stored in a container | [1] | [2] | [99] |
| 3.7 | Brought directly from water filter | [1] | [2] | [99] |
| 3.8 | Stored water was covered or in a narrow mouth vessel | [1] | [2] | [99] |
| | HOW DID SHE GET THE WATER INTO THE GLASS/CUP? | YES | NO | D/K |
| 3.9 | Hands touched / contacted the drinking water | [1] | [2] | [99] |
| 3.10 | Glass/cup dipped into water container | [1] | [2] | [99] |
| 3.11 | Ladle used to obtain water | [1] | [2] | [99] |
| 3.12 | Water poured from container | [1] | [2] | [99] |

| | | | | |
|---|---|---|---|---|
| 3.13 | Water poured from tap / handpump | [1] | [2] | [99] |

3.14 (ask/observe) What is the source of the drinking water?

1 = Tubewell

2 = Unprotected spring

3 = Protected spring

4 = Unprotected dug well

5 = Protected dug well

6 = Rainwater collection

7 = Cart with small tank / drum

8 = Tanker truck

9 = Surface water (river, dam, lake, pond, stream, canal, irrigation channel)

10 = Piped water into dwelling

11 = Piped water into yard / plot

77 = Other (specify)

99 = DK

3.15 (obs) (if 3.6 is 1) What is the type of the container?

1= Kolshi[B] /Jerrycan[K]

2 = Jug

3 = Topaz (provided by icddrb) [B]

4 = Bucket

5 = Other wide mouth container

6 = Other narrow mouth container

99 = Could not observe

3.16 (if 3.6 is 1) How long ago did you or somebody in your home collect this water? (99 = Don't know)

A ☐☐ HOURS

B ☐☐ DAYS

3.17 (if 3.6 is 1) Have you done anything to make this water less cloudy or safer to drink?

1 = Yes

2 = No → Skip to 3.20

99 = DK/Not sure

3.18 (if 3.17 is 1) How was this water treated? ✋Do Not Read Responses. Mark all that apply.

| | | |
|---|---|---|
| 1 | [1] Yes [2] No | Aquatabs[B] / Chlorine dispenser [K] (field code differs by country) |
| 2 | [1] Yes [2] No | Waterguard / bottled chlorine |
| 3 | [1] Yes [2] No | Boil |
| 4 | [1] Yes [2] No | Strain it through cloth or other material |
| 5 | [1] Yes [2] No | Water filter [ceramic, sand, composite] |
| 6 | [1] Yes [2] No | Solar disinfection (SODIS) |
| 7 | [1] Yes [2] No | Let it stand and settle |
| 8 | [1] Yes [2] No | Biosand filter |
| 9 | [1] Yes [2] No | Lifestraw Family Filter (VestergaardFrandsen distributed) [K] |
| 10 | [1] Yes [2] No | Coagulant (alum) |
| 11 | [1] Yes [2] No | PUR (flocculant + disinfectant) |
| 12 | [1] Yes [2] No | Other (specify) |

3.19 (if 3.17 is 1) Approximately how long ago did you treat the water?

HH:MM ☐☐:☐☐ ( 99:99 Don't know )

PROMPT: (if 3.18 is 1=Y or 2=Y) Please collect a drinking water sample by pouring the water from the glass provided by the participant into a Whirlpak with sodium thiosulphate.

PROMPT: (if 3.18 is 1=Nand 2=N) Please collect a drinking water sample by pouring the water from the glass provided by the participant into a regular Whirlpak.

3.20 Please enter the 4-digit unique numerical ID.

PROMPT: Please label the whirlpak with the following label: W.[PID].[DAY].[MONTH]

3.21 Has a drinking water sample been collected for microbiological testing?

1 = Yes

2 = No

3.22 (if 3.21 is 1) Record time that sample was collected (24H format, HH:MM)

PROMPT: (if 3.18 is 1=Y or 2=Y) Please fill a plastic tube with stored water for chlorine testing.

3.23 (if 3.18 is 1=Y or 2=Y) Has a sample from the storage container been collected for chlorine testing?

1 = Yes

2 = No

3.24 (if 3.23 is 1) Record time that sample was collected (24H format, HH:MM)

DAY 2

SECTION 1. SENTINEL TOY SAMPLING

1.1(ask) Did you receive a toy ball from our team yesterday?

1 = Yes

2 = No → Skip to Section 2

PROMPT: Could you please show me where the ball that was delivered to your household yesterday is currently located?

1.2 (ask) May I now rinse the toy ball that was delivered to your household yesterday?

1 = Yes

2 = Ball was lost → Skip to Section 2

3 = Ball was given away to another household → Skip to Section 2

4 = Refused → Skip to Section 2

77 = Could not retrieve the ball for other reason (specify) → Skip to Section 2

1.3 (obs) Where is the ball located?

1 = Outside: on dirt

2 = Outside: on concrete/cement/wood

3 = Outside: in container

4 = Outside: in another home

5 = Inside: on dirt floor

6 = Inside: on concrete/cement/wood floor

7 = Inside: on surface other than ground, not in container (e.g. on bed, on table, etc.)

8 = Inside: in storage container/cabinet

9 = In child's hands

77 = Other (specify)

1.4.a (ask)In your opinion, how much did (target child's name) play with the toy ball over the past 24 hours? (read each choice)

1 = Several times (4 or more times)

2 = Few times (2-3 times)

3 = Only once since he/she got the ball

4 = Never

99 = DK/Not sure

1.4.b In your opinion, how much time total did (target child's name) spend playing with the toy ball over the past 24 hours? (enter hours)

1.5.a (ask) In your opinion, how much did any of the other children in the household or bari play with the toy ball in the past 24 hours? (read each choice)

1 = Several times (4 or more times)

2 = Few times (2-3 times)

3 = Only once since he/she got the ball

4 = Never

99 = DK/Not sure

1.5.b In your opinion, how much time total did other children in the household or bari spend playing with the toy ball over the past 24 hours? (enter hours)

1.6 (ask) Did children from other baris play with the toy ball in the past 24 hours? (read each choice)

1 = They played with it within this bari

2 = They played with it outside this bari

3 = No

99 = DK/Not sure

1.7 (ask) From what you saw, did the children play with the ball mostly inside the home (indoors), mostly outside the home (outdoors) or equal amount inside and outside the home?

1 = Mostly inside the home

2 = Mostly outside the home

3 = Equal amounts inside and outside the home

99 = DK/Not sure

1.8 (ask) How did the child play with the ball? Read each choice.

1 = With hands

2 = With feet

3 = With hands and feet

99 = DK/Not sure

1.9 (ask) Did you do anything to clean the ball?

1 = Yes

2 = No

1.10 (ask) (if 1.9 is 1) What did you do to clean the ball? Select all that apply. Read each choice.

1 = Washed with water only

2 = Washed with water and soap

3 = Wiped with towel

4 = Wiped on clothes

77 = Other (specify)

1.11 (ask) (if 1.9 is 1) How long ago did you clean the ball? (Enter hours, 99=DK)

1.12 (obs) Have children played with the ball since you arrived at the household? (select all that apply)

1 = Target child has played with ball

2 = Other children in the household have played with ball

3 = No children have played with ball

1.13 (obs) Toy appearance.

1 = Unused

2 = Used, clean appearance

3 = Used, unclean appearance

4 = Used, visibly dirty

5 = Used, visibly wet

1.14 (obs) Toy condition.

1 = Good condition

2 = Bulging (from being stepped on, sat on, heat applied, etc)

3 = Deflated (ball has lost air)

4 = Burst/Torn (ball has visible hole)

PROMPT: Collect a toy rinse sample by rinsing the toy in the Whirlpak bag.

1.15 Please enter the 4-digit unique numerical ID.

PROMPT: Please label the whirlpak with the following label: T.[PID].[DAY].[MONTH]

1.16 Has a toy rinse sample been collected successfully?

1 = Yes, toy rinsed successfully

2 = No, some sample was spilled

3 = No, the inside of the sample bag was contaminated

4 = No, sample not collected

1.17 (if 1.16 is not 4) Record time that sample was collected (24H format, HH:MM).

SECTION 2. FLY DENSITY

2.1 Observe fly activity in the food preparation area. Estimate the total number of flies seen.

1 = None

2 = Low (1-5)

3 = Moderate (6-10)

4 = High (>10)

2.2 Was the fly tape hung under a roof (protected from rain)?

1 = Yes

2 = No

2.3 Is the food prep area inside or outside?

1 = Walls and roof

2 = Walls but no roof

3 = Roof but no walls

4 = No roof and no walls

2.4 (record) How many steps are the strips hung from the food preparation area?

2.5 (obs) Was the fly tape in the food preparation area tampered with or did it fall down?

2.6 (record) What is the total number of flies counted on all strips?

2.7(record) (if 2.6 is not 0) Write the number of each species observed:

Musca domestica______

Lesser house fly______

Blow/bottle fly______

Flesh fly/sarcophaga______

Other______

Cannot distinguish______

2.8 Record the time you performed the fly density observation at the food preparation area (24H format, HH:MM)

2.9 Observe fly activity in the latrine area. Estimate the total number of flies seen.

1 = None

2 = Low (1-5)

3 = Moderate (6-10)

4 = High (>10)

2.10 Was the fly tape hung under a roof (protected from rain)?

1 = Yes

2 = No

2.11 (record) How many steps are the strips hung from the latrine area?

2.12 (obs) Was the fly tape in the latrine area tampered with or did it fall down?

2.13 (record) What is the total number of flies counted on all strips?

2.14(record) (if 2.13 is not 0) Write the number of each species observed:

Musca domestica______

Lesser house fly______

Blow/bottle fly ______

Flesh fly/sarcophaga ______

Other ______

Cannot distinguish______

2.15 Record the time you performed the fly density observation at the latrine area (24H format, HH:MM)

**Wash Benefit Module 41 Household Assets**

<table>
<tr><td colspan="3">আপনার সময়ের জন্য আপনাকে ধন্যবাদ।আমি আপনার খানা সম্মন্ধে জানতে চাই যেখানে আপনি বসবাস করেন। প্রথমেই আমি আপনার ঘর কী দিয়ে তৈরী সেটা দেখতে চাই । তাহলে আমি কি ইহা দেখতে পারি? Thank you very much for your time. Now I would talk about <u>the house that you live in</u>. First, I would like to observe the material your house is made of. Can I take a look at your house?<br><br>OBSERVE MAIN MATERIAL OF THE HOUSE:</td></tr>
<tr><td></td><td>4101.</td><td>ছাদ তৈরীতে কি কি উপাদান ব্যবহার করা হয়েছে ? (পরির্দশন করে যাচাই করুন) [Main material of the roof]<br>কাঁচা (বাঁশ/খড়) [Kaccha (bamboo / thatch)] 1<br>টিন [Tin] 2<br>সিমেন্ট/ কংক্রিট/ টালি [Cement / concrete / tiled]…3</td></tr>
<tr><td></td><td>4102.</td><td>দেয়াল তৈরীতে কি কি উপাদান ব্যবহার করা হয়েছে ? (পরির্দশন করে যাচাই করুন) [Main material of the walls]<br>পাট/বাঁশ/মাটি (কাঁচা) [Jute / bamboo / mud (kaccha)] .... 1<br>কাঠ [Wood] .................................................................. 2<br>ইট/সিমেন্ট [Brick / cement] ............................ 3<br>টিন [Tin]................................................. 4</td></tr>
<tr><td></td><td>4103.</td><td>মেঝে তৈরীতে কি কি উপাদান ব্যবহার করা হয়েছে ? (পরির্দশন করে যাচাই করুন) [Main material of the floor]<br>মাটি/বাঁশ (কাঁচা) [Earth / bamboo (kaccha)] ..................1<br>কাঠ [Wood] 2<br>ইট/ সিমেন্ট [Cement / concrete] ..................................3</td></tr>
<tr><td colspan="3">Thank you. Now I would like to ask you some questions about your household assets.</td></tr>
<tr><td></td><td>4104.</td><td>খানার ঘরটিতে কতগুলো কক্ষ আছে? (রান্নাঘর ও বাথরুম বাদে) [How many rooms the households have (exclude bathroom and Kitchen)?].</td></tr>
<tr><td></td><td>4105.</td><td>আপনার খানায় নিম্নলিখিত জিনিসগুলো আছে কী? [How many does the households have <u>হ্যাঁ (Yes)....1, না(No)....0,জানি না (DK).....999</u><br>a.|_____|বিদ্যুৎ [Electricity]<br>b. |_____||_____|আলমারী/ওয়ারড্রব (সংখ্যা) [Number of Almirah or wardrobe]<br>c. |_____||_____|টেবিল (সংখ্যা) [Number of tables]<br>d. |_____||_____|চেয়ার/বেঞ্চ(সংখ্যা) [Number of chair or bench]<br>e. |_____||_____|হাত ঘড়ি/দেয়াল ঘড়ি(সংখ্যা) [Number of watch or clock]<br>f. |_____||_____|খাট(সংখ্যা) [Number of khat]<br>g. |_____||_____|চৌকি (সংখ্যা) [Number of chouki ]<br>h. |_____|রেডিও [A radio that is working]<br>i. |_____|টেলিভিশন (সাদা/কালো) [A B/W television that is working]<br>j. |_____|টেলিভিশন (রঙিন) [A color television that is working]<br>k. _____|রেফ্রিজারেটর [Refrigerator]<br>l. |_____|সাইকেল (খেলনা সাইকেল নয়) A bicycle (used for commercial purposes not toy for children)<br>m. |_____|মটর সাইকেল[A motorcycle]<br>n. |_____|সেলাই মেশিন [A sewing machine]<br>o. |_____||_____|মোবাইল ফোন(সংখ্যা) [Number of Mobile phones]<br>p. |_____|ল্যান্ড ফোন[A land phone]</td></tr>
<tr><td></td><td>4106.</td><td>আপনার খানায় রান্নার জন্য প্রধানত কি ধরনের জ্বালানী ব্যবহার করা হয় ?[What type of fuel does your household mainly use for cooking?]<br>কাঠ [Wood] ...................................................................... 01<br>শস্যের অবশিষ্টাংশ/ঘাস [Crop residue / grass ]................................. 02<br>শুকনা গোবর [Dung cakes ] ............. 03<br>কয়লা [Coal / coke / lignite ]........................................................ 04<br>কাঠ কয়লা [Charcoal ]................................................................. 05<br>কেরোসিন [Kerosene ] ............. 06<br>বিদ্যুৎ [Electricity ] ....................................................................... 07</td></tr>
</table>

| | | | | |
|---|---|---|---|---|
| | | তরল গ্যাস/প্রাকৃতিক গ্যাস [Liquid gas / gas ] .................................. 08<br>বায়ো-গ্যাস [Bio-gas] .................................................................... 09<br>অন্যান্য [Other] ............................................................................ 77<br>(বর্নণা লিখুন) [*Specify other]* ____________________<br>জানি না [Don't know ] ................................................................. 99 | | |
| | 4107. | \|_____\|আপনার খানার মোট মাসিক আয় কত? (সকল আয়ের উৎসের যোগফল যেমনঃ বেতন, ঘর ভাড়া, কৃষিখাত ইত্যাদি) For the purpose of our research, would you please tell us your total monthly household income? (Please sum up your income from all sources like wage, rent, agriculture etc.)<br>1. 4000 টাকা পর্যন্ত [UP TO 4000 Taka]<br>2. 4001-6000 টাকা [4001-6000 Taka]<br>3. 6001- 8000 টাকা [6001- 8000 Taka]<br>4. 8001- 10000টাকা [8001- 10000 Taka]<br>5. 10001- 12000 টাকা [10001- 12000 Taka]s<br>6. 12001- 15000 টাকা [12001- 15000 Taka]<br>7. 15001-20000 টাকা [15001-20000 Taka]<br>8. 20001- 25000 টাকা [20001- 25000 Taka]<br>9. 25001- 30000 টাকা [25001- 30000 Taka]<br>10. 30000 টাকার উপরে [Above 30000 Taka] | | |
| | 4108. | \|____\| বসতবাড়ি মালিকানা [What is the ownership status of the house where your household is currently living?]<br>নিজের বাড়ি [Self-owned] .............................................................. 1<br>ভাড়া বাড়ি [Rental] .......................................................................... 2<br>সরকারী জমি [Govt. land] ................................................................. 3.<br>কোন জমিদারের/জোতদারের বাড়িতে থাকে [Owned by a landlord].................... 4<br>বাড়ি ভাড়া না দিয়ে অন্যের বাড়িতে থাকে [Living in someone's house without giving rent] .......5 | | |
| | 4109. | খানার সদস্যদের বসতবাড়িতেকোন জমি আছে কি?<br>1. হ্যাঁ<br>0. না(skip to 4111) | | |
| | 4110. | \|____\|\|____\|\|____\|\|____\|\|____\| উত্তর হ্যাঁ হলে, খানার সদস্যদের বসতবাড়িতে মোট কতটুকু জমি আছে (ডেসিমেল)? (How much homestead land (decimal) does your household own?]<br>বলতে রাজি হয়নি [Refused]…..................................88<br>জানি না [Don't know]…................…..............99 | | |
| | 4111. | খানার সদস্যদের বসতবাড়ি ছাড়া অন্য কোন জমি আছে কি?<br>1. হ্যাঁ<br>০. না (skip to 4113) | | |
| | 4112. | \|____\|\|____\|\|____\|\|____\|\|____\| উত্তর হ্যাঁ হলে, খানার সদস্যদের বসতবাড়িতে মোট কতটুকু জমি আছে (ডেসিমেল)? (How much homestead land (decimal) does your household own?]<br>বলতে রাজি হয়নি [Refused]…..................................88<br>জানি না [Don't know]…................…..............99 | | |
| | 4113. | আপনার মতে সামাজিক প্রেক্ষাপটে আপনার খানাটির অবস্থান কি রকম [How would you describe your economic status?]<br>ধনী [Dhoni] .......................................................... 1<br>উচ্চ মধ্যবিত্ত [Uchho modho bitto] ......................... 2<br>মধ্যবিত্ত [Modho Bitto] .......................................... 3<br>দরিদ্র [Doridro or Nimno motho bitto] ................. 4<br>হতদরিদ্র [Hotodoridro] ......................................... 5 | | |
| | 4114. | আমি পশুর নামের তালিকা আপনাকে পড়ে শুনাতে চাচ্ছি ।এ গুলো কোনটি আপনার কম্পাউন্ড/াবাড়ীতে কতগুলো আছে এবং নিজের কয়টা আছে তা আমাকে বলুন । যদি না থাকে তাহলে ০০ বসান । আর যদি অজানা থাকে তাহলে ৯৯ বসান I'm going to read you a list of animals. Please tell me how many of each stays in your | কম্পাউন্ড/াবাড়ী COMPOUND | খানা HOUSEHOLD |

| | | | | |
|---|---|---|---|---|
| | | compound, and how many belong to your household. IF NONE, ENTER "00", IF UNKNOWN, ENTER "99" | | |
| | 1 | গরু CATTLE | \|__\|__\| | \|__\|__\| |
| | 2 | ছাগল GOATS | \|__\|__\| | \|__\|__\| |
| | 3 | মুরগী/হাঁস CHICKENS | \|__\|__\| | \|__\|__\| |
| | 4115. | কতবার গরু/ছাগল তাদের ঘরের বাহিরে বাড়ীতে/উঠানে ছেড়ে দেন? How often do the cattleroam free in the compound?<br>উত্তরগুলো পড়ুন READ RESPONSES.<br>[1] সবসময় ছেড়ে দেন ALWAYS ROAM FREE<br>[2] মাঝে মাঝে ছেড়ে দেন SOMETIMES ROAM FREE<br>[3] কখনও ছেড়ে দেয়া হয় না NEVER ROAM FREE<br>4. প্রযোজ্য নয়; গরু/ছাগল ছাড়া শুধুমাত্র মুরগী/হাঁসথাকলে N/A; ONLY HAVE CHICKENS | | |
| | 4116. | মুরগী অথবা অন্যান্য পাখি জাতীয় কোন কিছু আপনার প্রধান থাকার ঘরে প্রবেশ করে কি? Do chickens or other poultry ever go inside your main house?<br>READ RESPONSES.<br>[1] সবসময় ALWAYS<br>[2] মাঝে মাঝে SOMETIMES<br>[3] কখনও না NEVER<br>4. প্রযোজ্য নয়; মুরগী/হাঁসনাথাকলে N/A; NO CHICKENS | | |

**Wash Benefit Module42Feces Observed in and Around the Compound**

সকল খানার জন্য প্রযোজ্য Administer to: All study households

পর্যবেক্ষণ Observation: For the following:

প্রত্যেক জায়গায় মানুষের মল/পায়খানার স্তুপের সংখ্যা পর্যবেক্ষণ করুন (১০টি পর্যন্ত) *Record the n piles of human feces you observe in eacharea (up to 10 piles)*

55অধিক সংখ্যক (স্তুপের সংখ্যা ১০টির উপরে) Too numerous to count (more than 10 piles)

99 বলা যাচ্ছে না/পর্যবেক্ষণকরা সম্ভব হয়নি Cannot tell / could not observe

উঠানে Within the courtyard

| | | | |
|---|---|---|---|
| **C.826.** | 4201. | উঠানে মানুষের মল/পায়খানা পড়ে আছে যা উম্মুক্ত পায়খানা হিসেবে বিবেচিত হবে **Human** feces<u>within the</u> **<u>courtyard</u>** that could be considered open defecation<br><br>55 অধিক সংখ্যক (স্তুপের সংখ্যা ১০টির উপরে) Too numerous to count (more than 10 piles)<br>99 বলা যাচ্ছে না/পর্যবেক্ষণকরা সম্ভব হয়নি Cannot tell / could not observe | ☐☐ |
| **C.827.** | 4202. | উঠানে পশুর/পাখির মল/পায়খানা পড়ে আছে(নির্দিষ্টভাবে উল্লেখ করুন) **Animal** feces present <u>within the</u> **<u>courtyard</u>**(mark all that apply) | ☐1 পাখি বিশেষ(মুরগী/হাঁস/কবুতর )Poultry (chicken, duck, pigeon)<br>☐2 গরু/মহিষ Cow / Buffalo<br>☐3 ছাগল/ভেড়া Goat / Sheep |

| | | | |
|---|---|---|---|
| | | 55 অধিক সংখ্যক (স্তুপের সংখ্যা ১০টির উপরে) Too numerous to count (more than 10 piles)<br>99 বলা যাচ্ছে না/পর্যবেক্ষণকরা সম্ভব হয়নি Cannot tell / could not observe | ☐4শুকর Pig<br>☐5 কুকুর অথবা বিড়াল Dog or Cat<br>☐8 অন্যান্য Other<br>☐9 কোনও মল/পায়খানা পাওয়া যায়নি NO FECES OBSERVED<br>☐99 পর্যবেক্ষণকরা সম্ভব হয়নি COULD NOT OBSERVE |
| | 4202.a | অন্যকোন খানা এই উঠান ব্যবহার করে কী?Do any other household share this **courtyard** | ১. হ্যাঁ<br>০. না |
| ◉নির্দিষ্টশিশু বেশীরভাগ সময় যেখানে সময় কাটায় তা নিরম্নপন ও যাচাইকরম্নন। বেইজ লাইনে নির্দিষ্টশিশু থাকবে না সেজন্য গর্ভবতী মা যেখানে বেশীরভাগ সময় কাটায় তা নিরম্নপনকরম্নন In the area where the target child spends the most time<br>*Probe and identify where the target child spends the most time. At baseline, the target child will not be present, so the search should be defined as the area where the pregnant mother spends the most time.* | | | |
| **C.828.** | 4203. | মানুষের মল/পায়খানা পড়ে আছে যেখানে নির্দিষ্টশিশু (বেইজলাইনে গর্ভবতী মা) বেশীরভাগ সময় কাটায়। ইহা উম্মুক্ত পায়খানা হিসেবে বিবেচিত হবে **Human** feces in the **area where the target child spends the most time** that could be considered open defecation<br>55 অধিক সংখ্যক (স্তুপের সংখ্যা ১০টির উপরে) Too numerous to count (more than 10 piles)<br>99 বলা যাচ্ছে না/পর্যবেক্ষণকরা সম্ভব হয়নি Cannot tell / could not observe | ☐☐ |
| **C.829.** | 4204. | পশুর/পাখির মল/পায়খানা পড়ে আছে যেখানে নির্দিষ্টশিশু (বেইজলাইনে গর্ভবতী মা)বেশীরভাগ সময় সময় কাটায় আছে । (নিদ্দিষ্টভাবে উল্লেখ করুন) **Animal** feces in the **area where the target child spends the most time** (mark all that apply)<br>55 অধিক সংখ্যক (স্তুপের সংখ্যা ১০টির উপরে) Too numerous to count (more than 10 piles)<br>99 বলা যাচ্ছে না/পর্যবেক্ষণকরা সম্ভব হয়নি Cannot tell / could not observe | ☐1 পাখি বিশেষ(মুরগী/হাঁস/কবুতর )Poultry (chicken, duck, pigeon)<br>☐2 গরু/মহিষ Cow / Buffalo<br>☐3 ছাগল/ভেড়া Goat / Sheep<br>☐4শুকর Pig<br>☐5 কুকুর অথবা বিড়াল Dog or Cat<br>☐8 অন্যান্য Other<br>☐9 কোনও মল/পায়খানা পাওয়া যায়নি NO FECES OBSERVED<br>☐99 পর্যবেক্ষণকরা সম্ভব হয়নি COULD NOT OBSERVE |
| ◉ নিদ্দিষ্ট খানার পিছনে Behind the study house | | | |

| | | | |
|---|---|---|---|
| **C.830.** | 4205. | **খানার পিছনে মানুষের মল/পায়খানা পড়ে আছে/থাকলে। ইহা উম্মুক্ত পায়খানা হিসেবে বিবেচিত হবে Human** feces<u>behindthe</u> **<u>house</u>** that could be considered open defecation<br><br>55 অধিক সংখ্যক (**স্তুপের সংখ্যা ১০টির উপরে**) Too numerous to count (more than 10 piles)<br>99 বলা যাচ্ছে না/**পর্যবেক্ষণ**করা সম্ভব হয়নি Cannot tell / could not observe | ☐☐ |
| **C.831.** | 4206. | **খানার পিছনে পশুর মল/পায়খানা পড়ে আছে/থাকলে। (নিদ্দিষ্টভাবে উল্লেখ করুন) Animal** feces **<u>behind the house</u>** (mark all that apply)<br><br>55 অধিক সংখ্যক (**স্তুপের সংখ্যা ১০টির উপরে**) Too numerous to count (more than 10 piles)<br>99 বলা যাচ্ছে না/**পর্যবেক্ষণ**করা সম্ভব হয়নি Cannot tell / could not observe | ☐1 পাখি বিশেষ(মুরগী/হাঁস/কবুতর)Poultry (chicken, duck, pigeon)<br>☐2 গরু/মহিষ Cow / Buffalo<br>☐3 ছাগল/ভেড়া Goat / Sheep<br>☐4শুকর Pig<br>☐5 কুকুর অথবা বিড়াল Dog or Cat<br>☐8 অন্যান্য Other<br>☐9 কোনও মল/পায়খানা পাওয়া যায়নি NO FECES OBSERVED<br>☐99 **পর্যবেক্ষণ**করা সম্ভব হয়নি COULD NOT OBSERVE |

---------------------------------------------------------------------------------

**Protocol Title:** Measuring the benefits of sanitation, water quality, handwashing and nutrition interventions for improving health and development in rural Bangladesh

**Protocol Status:** APPROVED

**Date Submitted:** 10/26/2017

**Approval Period:** **10/30/2017-11/03/2018**

**Important Note:** **This Print View may not reflect all comments and contingencies for approval. Please check the comments section of the online protocol. Questions that appear to not have been answered may not have been required for this submission. Please see the system application for more details.**

*** * * Attached Document * * ***

| Document Name | Created Date |
|---|---|
| Stanford WASH B Extension Dec 2013 ApprovalLetter-25863.pdf | 10/26/2017 |

### STANFORD UNIVERSITY

*Stanford, CA 94305 [Mail Code 5579]*

Ronald L. Ariagno, M.D. (650) 724-7541

CHAIR, PANEL ON MEDICAL HUMAN SUBJECTS (650) 725-8013

#### Certification of Human Subjects Approvals

**Date:** December 17, 2013

**To:** Stephen P Luby, M.D., Medicine - Med/Infectious Diseases
Bianca Monique Velasquez, Amy Janel Pickering, Rossana Gonzalez-Ayala

**From:** Ronald L. Ariagno, M.D., Administrative Panel on Human Subjects in Medical Research

**Protocol** Randomized Controlled Trial of the Benefits of Water Sanitation Hygiene Plus Nutrition Interventions on Child Growth.

**Protocol ID:** 25863 **IRB Number:** 351 (Panel: 4)

The IRB approved human subjects involvement in your research project on 12/17/2013. **'Prior to subject recruitment and enrollment, if this is: a Cancer-related study, you must obtain Cancer Center Scientific Review Committee (SRC) approval;** a CTRU study, you must obtain CTRU approval; a VA study, you must obtain VA R and D Committee approval; and if a contract is involved, it must be signed.'

The expiration date of this approval is 12/17/2014 at Midnight. If this project is to continue beyond that date, you must submit an updated protocol in advance for the IRB's re-approval. If this protocol is used in conjunction with any other human use it must be re-approved. Proposed changes to approved research must be reviewed and approved prospectively by the IRB. No changes may be initiated without prior approval by the IRB, except where necessary to eliminate apparent immediate hazards to subjects. (Any such exceptions must be reported to the IRB within 10 working days.) Unanticipated problems involving risks to participants or others and other events or information, as defined and listed in the Report Form, must be submitted promptly to the IRB. (See Events and Information that Require Prompt Reporting to the IRB at http://humansubjects.stanford.edu.)

All continuing projects and activities must be reviewed and re-approved on or before Midnight of the expiration date. The approval period will be less than one year if so determined by the IRB. It is your responsibility to resubmit the project to the IRB for continuing review and to report the completion of the protocol to the IRB within 30 days.

Please remember that all data, including all signed consent form documents, must be retained for a minimum of three years past the completion of this research. Additional requirements may be imposed by your funding agency, your department, or other entities. (See Policy on Retention of and Access to Research Data at http://stanford.edu/dept/DoR/rph/2-10.html.)

This institution is in compliance with requirements for protection of human subjects, including 45 CFR 46, 21 CFR 50 and 56, and 38 CFR 16.

Includes: Add Longitudinal study

Waiver of Individual Authorization under 45 CFR 164.512(i)(2)(ii)(A),(B),(C), pursuant to information provided in the HIPAA section of the protocol application.

Ronald L. Ariagno, M.D., Chair

**Approval Period:** 12/17/2013 THROUGH 12/17/2014

**Review Type:** REGULAR - CONTINUING REVIEW

**Funding:** University of California Berkeley, Prime Bill and Melinda Gates Foundation , SPO: 108704

**Assurance Number:** FWA00000935 (SU)

LELAND STANFORD JUNIOR UNIVERSITY
DIE LUFT DER FREIHEIT WEHT
1891

### STANFORD UNIVERSITY

*Stanford, CA 94305 [Mail Code 5579]*

Ronald L. Ariagno, M.D. (650) 724-7541
CHAIR, PANEL ON MEDICAL HUMAN SUBJECTS (650) 725-8013

#### Certification of Human Subjects Approvals

**Date:** December 17, 2013

**To:** Stephen P Luby, M.D., Medicine - Med/Infectious Diseases
Bianca Monique Velasquez, Amy Janel Pickering, Rossana Gonzalez-Ayala

**From:** Ronald L. Ariagno, M.D., Administrative Panel on Human Subjects in Medical Research

**Protocol** Randomized Controlled Trial of the Benefits of Water Sanitation Hygiene Plus Nutrition Interventions on Child Growth.

**Protocol ID:** 25863 **IRB Number:** 351 (Panel: 4)

The IRB approved human subjects involvement in your research project on 12/17/2013. **'Prior to subject recruitment and enrollment, if this is: a Cancer-related study, you must obtain Cancer Center Scientific Review Committee (SRC) approval;** a CTRU study, you must obtain CTRU approval; a VA study, you must obtain VA R and D Committee approval; and if a contract is involved, it must be signed.'

The expiration date of this approval is 12/17/2014 at Midnight. If this project is to continue beyond that date, you must submit an updated protocol in advance for the IRB's re-approval. If this protocol is used in conjunction with any other human use it must be re-approved. Proposed changes to approved research must be reviewed and approved prospectively by the IRB. No changes may be initiated without prior approval by the IRB, except where necessary to eliminate apparent immediate hazards to subjects. (Any such exceptions must be reported to the IRB within 10 working days.) Unanticipated problems involving risks to participants or others and other events or information, as defined and listed in the Report Form, must be submitted promptly to the IRB. (See Events and Information that Require Prompt Reporting to the IRB at http://humansubjects.stanford.edu.)

All continuing projects and activities must be reviewed and re-approved on or before Midnight of the expiration date. The approval period will be less than one year if so determined by the IRB. It is your responsibility to resubmit the project to the IRB for continuing review and to report the completion of the protocol to the IRB within 30 days.

Please remember that all data, including all signed consent form documents, must be retained for a minimum of three years past the completion of this research. Additional requirements may be imposed by your funding agency, your department, or other entities. (See Policy on Retention of and Access to Research Data at http://stanford.edu/dept/DoR/rph/2-10.html.)

This institution is in compliance with requirements for protection of human subjects, including 45 CFR 46, 21 CFR 50 and 56, and 38 CFR 16.

Includes: Add Longitudinal study

Waiver of Individual Authorization under 45 CFR 164.512(i)(2)(ii)(A),(B),(C), pursuant to information provided in the HIPAA section of the protocol application.

Ronald L. Ariagno, M.D., Chair

**Approval Period:** 12/17/2013 THROUGH 12/17/2014

**Review Type:** REGULAR - CONTINUING REVIEW

**Funding:** World Bank , SPO: 112788

**Assurance Number:** FWA00000935 (SU)

### STANFORD UNIVERSITY

*Stanford, CA 94305 [Mail Code 5579]*

Ronald L. Ariagno, M.D. (650) 724-7541

CHAIR, PANEL ON MEDICAL HUMAN SUBJECTS (650) 725-8013

#### Certification of Human Subjects Approvals

**Date:** December 17, 2013

**To:** Stephen P Luby, M.D., Medicine - Med/Infectious Diseases
Bianca Monique Velasquez, Amy Janel Pickering, Rossana Gonzalez-Ayala

**From:** Ronald L. Ariagno, M.D., Administrative Panel on Human Subjects in Medical Research

**Protocol** Randomized Controlled Trial of the Benefits of Water Sanitation Hygiene Plus Nutrition Interventions on Child Growth.

**Protocol ID:** 25863 **IRB Number:** 351 (Panel: 4)

The IRB approved human subjects involvement in your research project on 12/17/2013. **'Prior to subject recruitment and enrollment, if this is: a Cancer-related study, you must obtain Cancer Center Scientific Review Committee (SRC) approval;** a CTRU study, you must obtain CTRU approval; a VA study, you must obtain VA R and D Committee approval; and if a contract is involved, it must be signed.'

The expiration date of this approval is 12/17/2014 at Midnight. If this project is to continue beyond that date, you must submit an updated protocol in advance for the IRB's re-approval. If this protocol is used in conjunction with any other human use it must be re-approved. Proposed changes to approved research must be reviewed and approved prospectively by the IRB. No changes may be initiated without prior approval by the IRB, except where necessary to eliminate apparent immediate hazards to subjects. (Any such exceptions must be reported to the IRB within 10 working days.) Unanticipated problems involving risks to participants or others and other events or information, as defined and listed in the Report Form, must be submitted promptly to the IRB. (See Events and Information that Require Prompt Reporting to the IRB at http://humansubjects.stanford.edu.)

All continuing projects and activities must be reviewed and re-approved on or before Midnight of the expiration date. The approval period will be less than one year if so determined by the IRB. It is your responsibility to resubmit the project to the IRB for continuing review and to report the completion of the protocol to the IRB within 30 days.

Please remember that all data, including all signed consent form documents, must be retained for a minimum of three years past the completion of this research. Additional requirements may be imposed by your funding agency, your department, or other entities. (See Policy on Retention of and Access to Research Data at http://stanford.edu/dept/DoR/rph/2-10.html.)

This institution is in compliance with requirements for protection of human subjects, including 45 CFR 46, 21 CFR 50 and 56, and 38 CFR 16.

Includes: Add Longitudinal study

Waiver of Individual Authorization under 45 CFR 164.512(i)(2)(ii)(A),(B),(C), pursuant to information provided in the HIPAA section of the protocol application.

Ronald L. Ariagno, M.D., Chair

**Approval Period:** 12/17/2013 THROUGH 12/17/2014

**Review Type:** REGULAR - CONTINUING REVIEW

**Funding:** Woods Institute for the Environment , SPO: 112476

**Assurance Number:** FWA00000935 (SU)

LELAND STANFORD JUNIOR UNIVERSITY
DIE LUFT DER FREIHEIT WEHT
1891

PROTOCOL
Biomedical
Non-Exempt
Berkeley

Protocol # 2011-09-3652
Date Printed: 07/31/2018

---------------------------------------------------------------------------------
---

| | |
|---|---|
| **Protocol Title:** | Measuring the benefits of sanitation, water quality, handwashing and nutrition interventions for improving health and development in rural Bangladesh |
| **Protocol Status:** | APPROVED |
| **Date Submitted:** | 10/26/2017 |
| **Approval Period:** | **10/30/2017-11/03/2018** |
| **Important Note:** | **This Print View may not reflect all comments and contingencies for approval. Please check the comments section of the online protocol. Questions that appear to not have been answered may not have been required for this submission. Please see the system application for more details.** |

*** * * Attached Document * * ***

| Document Name | Created Date |
|---|---|
| 2011-09-3652_Colford_1i-Consent-Parasite-v21-English.pdf | 10/26/2017 |

**Appendix 1i: English consent for parasitic assessment of children (target child, children aged 18-27 months at baseline or 5-12 years at endline) in the same compound**

Protocol Title: Effect of hand washing, water treatment, sanitation and nutritional supplement interventions on child health and development in rural Bangladesh

Principal Investigator's Name: Dr. Leanne Unicomb

**Purpose of the research**

Hello/Assalamualaikum/Namaste. My name is ________ and I work with the ICDDR,B (Cholera Hospital) in Dhaka. We are conducting research on diarrheal diseases and through this particular study we want to learn about the health impact of diarrheal diseases in children. We are interested in learning if the exposure of a child to diarrheal disease has long term effects. We are conducting this research with scientists at the University of California, Berkeley in the United States.

**Why are we inviting you to participate in the study?**

We are interested in enrolling your child because s/he is between the age of 18-27 months. Diarrhea is common among children in this age group. We are interested to know the risk of parasites for children in this bari. Therefore, we hope you will allow your child to participate into this study upon learning more details of participation.

**What is expected from the participants of the research study?**

Participation in this study will only span a few hours. To achieve the aim of the project we will collect a blood and a stool sample from your child. Both the samples will help us understand whether your child has been exposed to parasites diarrhea by conducting laboratory tests.

If you agree to participate, a field research person will visit your household twice for this purpose to collect a stool sample and a blood prick from your child. On the day before the collection a field member will deliver a stool collection kit and instruct you how to collect stool from your child. You will be instructed to collect your child's stool on the following morning, if your child defecates before their arrival, by having your child defecate on a sheet of provided plastic and use a plastic scoop to collect a small amount of fresh stool from the top of the pile into a container. The field person will collect this container when they come to collect the other specimens. We will examine whether your child has any worm infections, but this analysis will be done in Dhaka, and we will not be able to share the results with you. We will offer all members of your compound deworming medication regardless of the stool sample result.

The blood sample will be collected through a finger prick. Your child will experience a momentary pinch and approximately seven drops of blood will be collected by our trained field staff. One drop of blood will be used to test your child for anemia, a condition of lower than normal red blood cells. The field member can provide you with the results of this test during their visit.

In addition, if you are enrolled in the sanitation or control arms of the study, we will collect one soil sample from your household entrance to look for worm eggs.

**Risks & Benefits**

Although we will try to protect your identity there is some possibility that confidentiality could be compromised. The blood drops will be collected by a trained professional. Your child may feel some momentary pain during the blood collection. There is no direct benefit for participating in this study, but your child's participation will help us to gain knowledge on diarrheal disease in children.

**Confidentiality**

All data and specimens collected will be kept confidential as allowed by the law of this country.

Confidentiality of the data and test results will be strictly maintained. We will use the information only for the purpose of the study, and we will not use your name in sharing and publishing the results of this study.

**Future use of information**

The blood and stool samples may be stored until the end of the study, so they can be analyzed in the lab. If you agree, the information, along with some of the blood and stool collected will be stored for a long time after the study ends. This is because new laboratory techniques will become available in the future to help us better understand how diarrheal diseases affect children's health. The information collected from this study may be shared with other researchers if needed, but we will strictly maintain your confidentiality.

**Right not to participate and withdraw**

Taking part in the study is completely voluntary. You have the right to decline to allow your child to participate or to withdraw your child at any point in this study without penalty or loss of benefits to which you are otherwise entitled. You may choose not to answer any or all of the questions that will be asked. You can drop out of this study at any time, even in the middle of an interview. You have the right to refuse to take part in this study, which will not affect your family's treatment at the Cholera Hospital in the future.

**Principle of compensation**

You need not pay us to take part in this study, and similarly we will not pay you money for attending in the study.

**Persons to contact:**

If you have any questions, you can ask me any time. If you have additional questions about the survey, you may contact:

Masud Parvez, Center of Communicable Diseases, ICDDR,B, Mohakhali, Dhaka 1212.

If you have questions about your rights as a participant of a research study, or if you think some harm has been done to you because of the research, you may contact or meet IRB Secretariat, M. A. Salam Khan:

M. A. Salam Khan, IRB Secretariat, phone: 9886498 or PABX 8860523-32 ext. 3206

If you agree to participate in our study and to allow your child to participate in our study, please indicate that by putting your signature or your left thumb impression at the specified space below

I consent to storing my child's blood and stool samples long term [ ]

Thank you for your cooperation

____________________________________ ____________________

Signature or left thumb impression of Guardian Date

____________________________________ ____________________

Signature or left thumb impression of the witness Date

____________________________________ ____________________

Signature of the PI or his/her representative Date

PROTOCOL
Biomedical
Non-Exempt
Berkeley

Protocol # 2011-09-3652
Date Printed: 07/31/2018

---------------------------------------------------------------------------------

| | |
|---|---|
| **Protocol Title:** | Measuring the benefits of sanitation, water quality, handwashing and nutrition interventions for improving health and development in rural Bangladesh |
| **Protocol Status:** | APPROVED |
| **Date Submitted:** | 10/26/2017 |
| **Approval Period:** | **10/30/2017-11/03/2018** |
| **Important Note:** | **This Print View may not reflect all comments and contingencies for approval. Please check the comments section of the online protocol. Questions that appear to not have been answered may not have been required for this submission. Please see the system application for more details.** |

*** * * Attached Document * * ***

| Document Name | Created Date |
|---|---|
| 2011-09-3652_Colford_2i-Consent-Parasite-v21-Bengali.pdf | 10/26/2017 |

icddr,b
KNOWLEDGE FOR GLOBAL LIFESAVING SOLUTIONS

**Appendix 2i: Bengali consent for parasitic assessment of children (target child, children aged 18-27 months at baseline or 5-12 years at endline) in the same compound**

Protocol Title: Effect of hand washing, water treatment, sanitation and nutritional supplement interventions on child health and development in rural Bangladesh

cavb M‡elK: W: wj‡qb BDwbK¤

M‡elYvi D‡Ïk¨

AvmmvjvgAvjvBKg/ bg¯vi| Avgvi bvg---------------- | Avwg XvKvi AvBwmwWwWAvi, we (K‡jiv nvmcvZvj) †Z KvR Kwi| Avgiv Wvqwiqv †ivM wb‡q M‡elYv KiwQ Ges GB M‡elYvi gva¨‡g Avgiv wkk‡`i ¯v‡¯¨i Dci Wvqwiqv †iv‡Mi cfve m¤ú‡K Rvb‡Z Pvw"Q| wkï‡`i Wvqwiqv ‡iv‡Mi SwKi †Kvb `xN‡gqvw` cfve i‡q‡Q wKbv, Zv Rvb‡Z Avgiv AvMnx|

Avgiv †Kb Avcbv‡K GB M‡elYvq AskMn‡Y AvgšY Rvbvw"Q?

Avgiv GB Lvbv‡K M‡elYvq AšÍf³ Ki‡Z Pvw"Q KviY GLv‡b Kgc‡¶ GKwU 18-27 gv‡mi ev"Pv Av‡Q| GB eqmx ev"Pv‡`i g‡a¨ Wvqwiqv †ekx ‡`Lv hvq| Avgiv GB evoxi ev"Pv‡`i ciRxexi SwK m¤ú‡K Rvb‡Z AvMnx| Avgiv Avkv KiwQ AskMnY m¤ú‡K we¯ÍvwiZ Rvbvi ci Avcwb Avcbvi wkï‡K GB M‡elYvq AskMnb Ki‡Z w`‡Z ivRx n‡eb|

M‡elYvq AskMnYKvixi Kv‡Q cZ¨vkv Kx?

GB M‡elYvq AskMnY Ki‡j K‡qK N›Uv mgq e¨q n‡e| GB M‡elYvi D‡Ïk¨ mdj Ki‡Z n‡j wkïi kixi †_‡K K‡qK †dvUv i³ Ges wKQ cvqLvbvi bgbv msMn Ki‡Z n‡e| bgbv `wU cix¶vMv‡i cix¶vi gva¨‡g Avgiv eS‡Z m¶g ne †h, Avcbvi wkïi Wvqwiqv nevi SwK †Kgb|

Avcwb AskMnY Ki‡Z ivRx n‡j, GKRb gvV M‡elK Avcbvi wkkmn (Uv‡MU) GB DVv‡bi m‡ev"P Avi I `BRb wkïi cvqLvbv Ges i‡³i bgbv msMn Ki‡Z `Bevi Avcbvi Lvbvq Avm‡e| bgbv msMn Kivi Av‡Mi w`b GKRb gvVKgx Avcbv‡K cvqLvbv msMn Kivi mvgMx w`‡q hv‡e Ges wKfv‡e Avcbvi wkïi cvqLvbv msMn Ki‡Z n‡e Zv †`wL‡q ‡`‡e| c‡ii w`b mKv‡j hw` gvVKgx ‡cŠQv‡bvi c‡e wkï cvqLvbv K‡i, Z‡e Avcwb Avcbvi wkïi cvqLvbv msMn Ki‡eb| cvqLvbv msMn Kivi Rb¨ Avcbv‡K GKwU cøwóKwkU †`qv n‡e, †hLv‡b wkï cvqLvbv Ki‡e Ges Avcwb GKwU cøwóK (‡Qwb) ¯‹c e¨envi K‡i Dci †_‡K Aí GKU m`¨ (GBgvÎ Kiv) cvqLvbv †evZ‡j fi‡eb| Avgv‡`i gvVKgx hLb Ab¨vb¨ bgbv msMn Kivi Rb¨ Avm‡e, ZLb GB †evZjwU Avcbvi KvQ †_‡K wb‡q †b‡e| Avgiv cixÿvi gva¨‡g ‡`L‡ev ‡h Avcbvi wkïi kix‡i ‡Kvb Kwgi msµvgb Av‡Q wKbv, wKš GB cixÿwU XvKvq Kiv n‡e Ges Gi djvdj Avcbv‡K Rvbv‡bv n‡e bv| Avgiv Avcbvi wkïi cvqLvbv cixÿvi djvdj hvB‡nvK bv †Kb Avcbvi DVv‡bi mKj m`m¨‡K Kwgi Jla w`‡ev|

Avcbvi wkïi kixi †_‡K mPv‡jv m‡Pi gva¨‡g i³ msMn Kiv n‡e| Avgv‡`i cwk¶Ycvß gvVKgxi gva¨‡g K‡qK †dvUv i³ msMn Kiv n‡e| Avcbvi wkïi i³ msMn Kivi gn‡Z GKU †LvPv jvM‡e|i‡³i GK‡dvUv w`‡q Avcbvi wkïi G¨wbwgqv cixÿv Kiv n‡e| G¨wbwgqv n‡jv i‡³i g‡a¨ ‡jwnZ KwYKv mvfwe‡Ki Zjbvq Kg _vKv|gvV M‡elK hLb GB cixÿvi Rb¨ Avcbvi Lvbvq Avm‡e, ZLb ‡m GB cixÿvi djvdj Avcbv‡K w`‡q w`‡e|

GQvovI, hw` Avcwb m¨wb‡Ukb ev K‡›Uvj M‡elYvq ZwjKvf³ n‡q _v‡Kb Z‡e Avgiv Kwgi wWg Av‡Q wKbv Zv †`Lvi Rb¨ Avcbv Lvbvi c‡ek¯j †_‡K gwUi bgbv msMn Kie|

SwK Ges mweav

hw`I Avgiv Avcbvi cwiPq †Mvcb ivLvi †Póv Kie Ze Zv cKwkZ nIqvi m¤vebv Av‡Q| cwk¶Ycvß Ges †ckv`vi †jv‡Ki gva¨‡gi³ msMn Kiv n‡e| Avcbvi wkï i³ msMn Kivi mgq mvgwqK mg‡qi Rb¨ wKQUv e¨_v †c‡Z cv‡i| Avcwb M‡elYvq AskMn‡Yi Rb¨ mivmwi †Kvb ai‡Yi mnvqZvI cv‡eb bv| wKš Avcbvi wkïi AskMnY, wkï‡`i Wvqwiqv †ivM m¤ú‡K Ávbjv‡f Avgv‡`i‡K mnvqZv Ki‡e|

†MvcbxqZv

mKj Z_¨ Ges msMnxZ bgbv GB †`‡ki AvBb Abhvqx †MvcbxqZvi mv‡_ ivLv n‡e| Z_¨ Ges bgbvi djvd‡ji †MvcbxqZv K‡Vvi fv‡e cvjb Kiv n‡e|

Avgiv ïa M‡elYvi c‡qvR‡b GB Z_¨ e¨envi Kie Ges Avcbvi bvg A_ev cwiP‡qi †Kvb mÎ GB M‡elYv msµvšÍ †Kvb wi‡cvU/ cKvkbvq D‡jøL Kiv n‡e bv|

fwel¨‡Z Z‡_¨i e¨envi

GKB mg‡q j¨v‡e cix¶v Kivi Rb¨ GB i³ Ges cvqLvbvi bgbv M‡elYvi †kl mgq chšÍ msi¶Y Kiv n‡Z cv‡i| hw` Avcwb iwR _v‡Kb, Z‡e GB i³ Ges cvqLvbvi bgbvi cvkvcvwk msMnKZ Z_¨I M‡elYvi †kl mgq chšÍ msi¶Y Kiv n‡Z cv‡i| Gi d‡j fwel¨‡Z bZb j¨ve‡iUix c×wZ wkï‡`i ¯v‡¯¨i Dci Wvqwiqv †iv‡Mi cfve fvjfv‡e eS‡Z mvnvh¨ Ki‡e| GB M‡elYvq msMnxZ Z_¨ c‡qvR‡b Ab¨vb¨ M‡el‡Ki mv‡_ wewbgq n‡Z cv‡i, wKš Z‡_¨i †MvcYxqZv K‡Vvifv‡e cvjb Kiv n‡e|
‡¯”Qv AskMnY
GB M‡elYvq AskMnY m¤cYfv‡e Avcbvi B”Qvi Ici wbfi K‡i| Avcwb GB M‡elYvq Avcbvi wkï‡K AskMnY bvI Ki‡Z w`‡Z cv‡ib| Avcbvi wjwLZ AbgwZ †`qvi cieZx‡ZI Avcwb B”Qv Ki‡j ‡h †Kvb mg‡q GgbwK bgbv msM‡ni gvSLv‡bI Avcbvi AbgwZ cZ¨vL¨vb Ki‡Z cv‡ib| GB M‡elYvq AskMnY bv Kivi AwaKvi Avcbvi Av‡Q Ges GRb¨ fwel¨‡Z Avcbvi cwiev‡ii m`m¨‡`i AvBwmwWwWAvi,we-i ¯v¯¨‡mev MnY‡Yi Dci †Kvb cfve co‡e bv|
¶wZciY
GB M‡elYvq AskMn‡Yi Rb¨ Avcbvi mivmwi †Kvb ai‡Yi A_%bwZK LiP/ e¨q n‡e bv Ges GKBfv‡e Avcwb GB M‡elYvq AskMn‡Yi Rb¨ mivmwi †Kvb ai‡Yi A_%bwZK mnvqZvI cv‡eb bv|
‡hvMv‡hvM
Avcbvi †Kv‡bv ck _vK‡j Avgv‡K wRÁvmv Ki‡Z cv‡ib| hw` M‡elYv m¤úwKZ AwZwi³ †Kvb ck _v‡K Zvn‡j Avcwb gvm` cvi‡fR (wmwmwW, AvBwmwWwWAvi,we, gnvLwj, XvKv-1212) Gi mv‡_ 8860523-32 (G·- 120) †Uwj‡dvb b¤‡i mivmwi †hvMv‡hvM Ki‡Z cv‡ib|
GB M‡elYvq Avcbvi AwaKvi-msµvšÍ †Kvb ck _vK‡j A_ev M‡elYvi Kvi‡Y Avcbvi †Kvb ¶wZ n‡Z cv‡i e‡j hw` g‡b K‡ib, Zvn‡j Avcwb †Uwj‡dvb b¤‡i A_ev mivmwi AvBAviwe †m‡µUwi‡qU, Gg G mvjvg Lvb Gi mv‡_ †hvMv‡hvM Ki‡Z cv‡ib|
Gg G mvjvg Lvb, AvBAviwe †m‡µUwi‡qU, 9886498 A_ev wcGwеG· 8860523-32 (G·- 3206)

Avcwb hw` GB M‡elYvq AskMnY Ki‡Z AvMnx nb Zvn‡j wb‡gi wbawiZ ¯v‡b ¯v¶i A_ev evg e×v½jxi Qvc w`b|

Awg `xN mgq a‡i Avgvi wkïi i³ Ges cvqLvbv msi¶‡Yi AbgwZ w`jvg.........[ ]

Avcbvi mn‡hwMZvi Rb¨ ab¨ev`|

.......................................... ................................
Awffve‡Ki ¯v¶i/evg e×v½wji Qvc ZwiL

.......................................... ................................
mv¶xi ¯v¶i/evg e×v½wji Qvc ZwiL

.......................................... ................................
wc AvB/ M‡el‡Ki cwZwbwai ¯v¶i ZwiL

PROTOCOL
Biomedical
Non-Exempt
Berkeley

Protocol # 2011-09-3652
Date Printed: 07/31/2018

-----------------------------------------------------------------------------------
---

**Protocol Title:** Measuring the benefits of sanitation, water quality, handwashing and nutrition interventions for improving health and development in rural Bangladesh

**Protocol Status:** APPROVED

**Date Submitted:** 10/26/2017

**Approval Period:** **10/30/2017-11/03/2018**

**Important Note:** **This Print View may not reflect all comments and contingencies for approval. Please check the comments section of the online protocol.
Questions that appear to not have been answered may not have been required for this submission. Please see the system application for more details.**

*** * * Attached Document * * ***

| Document Name | Created Date |
|---|---|
| 2011-09-3652_Colford_1k1-Consent-EnvAssess-v21-English.pdf | 10/26/2017 |

#### Appendix 1k-1: English consent form for environmental assessment (drinking water, hand rinse, sentinel toy, food sampling and fly density measurements)

Protocol Title: Effect of hand washing, water treatment, sanitation and nutritional supplement interventions on child health and development in rural Bangladesh

Principal Investigator's name: Dr. Stephen P. Luby

##### Purpose of the research

Hello (Assalamualaikum/Namaste). My name is ________ and I work with the ICDDR,B (Cholera Hospital) in Dhaka. We are doing research on diarrheal diseases in children. Through this research we want to learn how certain conditions in sanitation, water quality and hygiene practices impact health. We are conducting this research in collaboration with scientists at University of California, Berkeley in the United States.

**Why are we inviting you to participate in the study?**
We are seeking households that have a child less than 3 years old and have already taken part in other activities from this study. We would like to tell you more detail about this study and hope you and your child will volunteer to participate.

**What is expected from the participants of the research study?**
If you decide to take part in the study, during two visits we will conduct some specific activities. In the first visit, which we would like to do today, we will ask you questions about your home and compound, and some practices in your home. We will also give your children one toy ball and set up fly traps in the latrine and kitchen areas of your compound. We will also wash your young child's hands in clean water and keep the water used to wash the hands. We will also take a very small amount of stored food that young children in your household will eat. Finally, we will ask you to give us a glass of water that you would give to your children to drink. We will take a small amount of the water that you give us.

We will return tomorrow for a second visit, wash the balls in water, keep the water used to wash the toy. Your child may choose to play with the ball or not; this is up to him or her. During this second visit, we will count the flies caught at the fly trap in the latrine and kitchen areas of your compound. We will also ask you a few short questions. Each visit will be less than 1 hour.

**Risks & Benefits**
There are no major risks involved in this study. Your children will receive 2 toy balls as a gift. You will not directly benefit from participating in this study but this participation will enable us to gain knowledge regarding fecal contamination in association with diarrhea.

**Privacy, anonymity and confidentiality**
We will do everything we can keep what you tell us confidential as allowed by the law of this country. We will not use your real name when we write out the data. All the information we collect will be kept locked. We will use the information only for the purpose of the study. We will not use your name when we share and publish the results of this study. We expect the steps we take will keep all of your information confidential, but it is possible that because of mistakes or unforeseen events, it could become compromised.

**Future use of information**
If it is necessary, we may share the information we collected from this study with other researchers. If this is done, we will not use your real name and we will maintain your confidentiality. We will store the information we collect from you for a long time after the end of the study.

**Right not to participate and withdraw**

Taking part in the study is completely voluntary. You may choose not to answer any or all of the questions that will ask. You can drop out of this study at any time, even in the middle of an interview. You have the right to refuse to take part in this study, which will not affect your family's treatment at the Cholera Hospital in the future.

**Principle of compensation**
You do not need to pay us to take part in this study, and we will not pay you money for taking part in the study.

**Persons to contact:**

If you have any questions, you can ask me any time. If you have additional questions about the survey, you may contact: Masud Parvez: phone: 01817541872, Center of Communicable Diseases, ICDDR,B, Mohakhali, Dhaka 1212.

If you have questions about your rights as a participant of a research study, or if you think some harm has been done to you because of this study, you may contact or meet him personally at following address:

M. A. Salam Khan, IRB Secretariat, phone: 9886498 or PABX 8860523-32 ext. 3206

If you agree to participate in our study and for your children to participate in our study, please indicate that by putting your signature or your left thumb impression at the specified space below

Thank you for your cooperation

__________________________________ __________________

Signature or left thumb impression of Participant Date

__________________________________ __________________

Signature or left thumb impression of the witness Date

__________________________________ __________________

Signature of the PI or his/her representative Date

PROTOCOL
Biomedical
Non-Exempt
Berkeley

Protocol # 2011-09-3652
Date Printed: 07/31/2018

-----------------------------------------------------------------------------------
---

**Protocol Title:** Measuring the benefits of sanitation, water quality, handwashing and nutrition interventions for improving health and development in rural Bangladesh

**Protocol Status:** APPROVED

**Date Submitted:** 10/26/2017

**Approval Period:** **10/30/2017-11/03/2018**

**Important Note:** **This Print View may not reflect all comments and contingencies for approval. Please check the comments section of the online protocol.**
**Questions that appear to not have been answered may not have been required for this submission. Please see the system application for more details.**

*** * * Attached Document * * ***

| Document Name | Created Date |
| --- | --- |
| 2011-09-3652_Colford_2k1-Consent-EnvAssess-v19-Bengali.pdf | 10/26/2017 |

Q.T/ S.G/ Cl. ID/ Co. ID/ B. ID/ HH ID
Household ID: □/□/ □□/ □□/□□/□□

icddr,b
KNOWLEDGE FOR GLOBAL LIFESAVING SOLUTIONS

### Appendix 2k-1: Bengali consent for environmental assessment (drinking water, hand rinse, sentinel toy and fly density measurements)

পরিবেশগত প্রভাব মূল্যায়নের জন্য সম্মতিপত্র (খাবার পানি, হাত ধোয়া, খেলনা এবং মাছির ঘনত্ব পরিমাপ)

গবেষণার শিরোনাম:বাংলাদেশের গ্রামীণ এলাকায় হাত ধোয়া, পানি বিশুদ্ধকরণ, স্যানিটেশন ওপুষ্টিকর সম্পূরক খাবার প্রদান এবং শিশুর স্বাস্থ্য ও বৃদ্ধির উপর তার প্রভাব।

**প্রধান গবেষক**: ডঃ লিয়েন ইউনিকম্ব

**গবেষণার উদ্দেশ্য:**

আসলামুআলাইকুম/আদাব। আমার নাম ----------------------------। আমরা ঢাকা (মহাখালী) কলেরা হাসপাতালে কাজ করি।আমরা শিশুদের ডায়রিয়াজনিত রোগ-বালাই নিয়ে গবেষণা করছি। কতিপয় স্যানিটেশন ব্যবস্থা, পানির গুনাগুন এবং স্বাস্থ্যসম্মত অভ্যাসসমূহ স্বাস্থ্যের উপর কিভাবে প্রভাব ফেলে এই গবেষণার মাধ্যমে আমরা তা বোঝার চেষ্টা করছি।

আমরা কেন আপনাকে এই গবেষণায় অংশগ্রহনে আমন্ত্রণ জানাচ্ছি?

আমরা আপনাকে/আপনার খানাকে এই গবেষণায় অংশগ্রহণের জন্য আমন্ত্রন জানাচ্ছি কারণ আপনার ৩ বছরের কম বয়সের একটি শিশু আছে এবং আপনি/আপনার খানা ইতোমধ্যেই আমাদের গবেষণার অন্যান্য কর্মকান্ডে অংশগ্রহন করেছেন। আমরা আপনাকে/আপনাদেরকে এই গবেষণার বিষয়ে আরো বিস্তারিতভাবে জানাতে চাই। আমরা আশা করছি আপনি এবং আপনার শিশু স্বেচ্ছায় এই গবেষণায় অংশগ্রহন করবেন।

গবেষণায় অংশগ্রহণকারীর কাছে প্রত্যাশা কী?

যদি আপনি/আপনার খানা এই গবেষণায় অংশগ্রহণে সম্মত থাকেন, তাহলে ২ বার আমরা আপনার খানা পরিদর্শন করব এবং এসময় নির্দিষ্ট কিছু কর্মকান্ড পরিচালনা করব। প্রথম পরিদর্শনের অংশ হিসাবে আজকে আমরা আপনাকে আপনার বাড়ি ও বাড়ির চারপাশের পরিবেশ, আপনার শিশুর স্বাস্থ্য এবং আপনার বাড়িতে প্রচলিত অভ্যাস সম্পর্কে কিছু প্রশ্ন জিজ্ঞাসা করব। আমরা আপনার শিশুকে ১টা খেলনার বল দিব এবং আপনার বাড়ির পায়খানা ও রান্নাঘরের আশেপাশে মাছি ধরার ফাঁদ বাধঁব/সেট করব। আমরা আপনার সবচেয়ে ছোট শিশুর হাতগুলো পরিস্কার পানি দিয়ে ধোব/ধৌত করব এবং হাতধোয়া পানিটুকুও সংরক্ষন করব। আপনি আপনার শিশুকে খাওয়ার জন্য যে পানি দেন, সেই পানি থেকে আমাদেরকে একগস্নাস পানি দিতে অনুরোধ করব। আপনার দেওয়া এই পানি থেকে আমরা নমুনা হিসাবে সামান্য পানি সংগ্রহ করব।

দ্বিতীয়বার পরিদর্শনের জন্য আমরা আগামীকাল আবার আপনার বাড়িতে আসব। আমরা পানি দিয়ে আপনার শিশুকে দেওয়া খেলনা বলগুলো ধোব/ধৌত করব এবং বল পানিটুকু সংরক্ষন করব। আপনার শিশু খেলনা বলগুলো দিয়ে খেলতেও পারে আবার নাও খেলতে পারে। এটা সম্পূর্ন তার ইচ্ছার উপর নির্ভর করে। দ্বিতীয় পরিদর্শনের সময় আমরা আপনার বাড়ির পায়খানা ও রান্নাঘরের

আশেপাশে বেধেঁ রাখা ফাঁদে/ ফ্লাই ট্রেপে আটকা পড়া মাছির সংখ্যা গননা করব। আমরা আপনাকে কিছু প্রশ্নও জিজ্ঞাসা করব। প্রত্যেকবার পরিদর্শনকালে আমরা আপনার খানাতে প্রায় ১ ঘন্টা সময় কাটাব।

ঝূঁকি এবং সুবিধাসমূহ:

এই গবেষণায় অংশগ্রহনে আপনার কোন ঝুঁকি নেই। এই গবেষণায় অংশগ্রহনের জন্য আপনি কোন ধরণের আর্থিক সহায়তাও পাবেন না। কিন্তু উপহার হিসাবে আপনার শিশু ১টা খেলনার বল পাবে।

নিজস্বতা, গোপনীয়তা ও বিশ্বাসযোগ্যতা:

এই দেশের আইন অনুযায়ী আপনার/ আপনাদের দেওয়া সমস্ত তথ্যগোপনীয়তার সাথে রক্ষণাবেক্ষণ করা হবে। আমরা আমাদের গবেষণার কোন লেখায়/ফলাফলে আপনার প্রকৃত নাম ব্যবহার করব না। আমাদের সংগৃহীত সকল তথ্য তালাবদ্ধ থাকবে। আমরা কেবলমাত্র আমাদের গবেষণার কাজে এসব তথ্য ব্যবহার করব। আমরা এই গবেষণার ফলাফল বিনিময় ও প্রকাশনার ক্ষেত্রে আপনার নাম ব্যবহার করব না। আমরা আপনার দেওয়া তথ্যসমূহ গোপন রাখার জন্য সম্ভাব্য সবধরনের চেষ্টা করব। তা সত্ত্বেও ভুলবশতঃ অনাকাঙ্খিত কোন ঘটনা ঘটলে সেজন্য মার্জনা প্রার্থনা করছি।

ভবিষ্যতে তথ্যের ব্যবহার:

এই গবেষণায় সংগৃহীত তথ্য প্রয়োজনে আমরা অন্যান্য গবেষকের সাথে বিনিময় করতে পারি। এবং কারো সাথে তথ্য বিনিময় করা হলে আমরা সেখানে আপনার প্রকৃত নাম ব্যবহার করব না এবং আপনার দেওয়া তথ্যের গোপণীয়তা ও নিজস্বতা কঠোরভাবে রক্ষা করব।

গবেষণায় অংশগ্রহন না করা বা নাম প্রত্যাহারের অধিকারঃ

এই গবেষণায় অংশগ্রহন করা সম্পূর্নভাবে আপনার ইচ্ছার উপর নির্ভর করছে। এই গবেষণায় আপনাকে যেসব প্রশ্ন জিজ্ঞাসা করা হবে তার যেকোনটির উত্তর আপনি চাইলে নাও দিতে পারেন। এই গবেষণায় অংশগ্রহনে সম্মত হবার পরেও গবেষণা চলাকালীন যেকোন সময় আপনি আপনার নাম প্রত্যাহার করে নিতে পারেন। এই গবেষণায় অংশগ্রহণ না করার অধিকার আপনার আছে এবং এ জন্য ভবিষ্যতে আপনার পরিবারের সদস্যদের কলেরা হাসপাতালে স্বাস্থ্যসেবা গ্রহণের উপর কোন প্রভাব পড়বে না।

ক্ষতিপূরণ :

এই গবেষণায় অংশগ্রহণের জন্য আপনার কোন ধরনের অর্থনৈতিক খরচ/ ব্যয় বহন করতে হবে না এবং একইভাবে আপনাকেও আমরা কোন ধরনের অর্থ প্রদান করব না।

যোগাযোগ:

আপনার কোনো প্রশ্ন থাকলে যেকোন সময় আপনি তা আমাকে জিজ্ঞাসা করতে পারেন। যদি গবেষণা সম্পর্কিত অতিরিক্ত কোন প্রশ্ন থাকে তাহলে আপনি জনাব মাসুদ পারভেজ (প্রোগ্রাম অন ইনফেক্সাস ডিজিজেজ এন্ড ভ্যাকসিন সাইন্সেস, আইসিডিডিআর,বিঃ মহাখালি, ঢাকা-১২১২) এর সাথে ০১৮৭৫৪১৮৭২ মোবাইল নম্বরে সরাসরি যোগাযোগ করতে পারেন। এই গবেষণায় আপনার অধিকার-সংক্রান্ত কোন প্রশ্ন থাকলে অথবা গবেষণার কারণে আপনার অর্থনৈতিক কোন ক্ষতি সাধিত হয়েছে বলে মনে করেন, তাহলে আপনি ৯৮৮৬৪৯৮ অথবা ৮৮৬০৫২৩-৩২ এক্স- ৩২০৬ টেলিফোন নম্বরে অথবা সরাসরি এম এ সালাম খান, আইসিডিডিআরবি সেক্রেটারিয়েট এর সাথে যোগাযোগ করতে পারেন।

আপনি যদি এই গবেষণায় অংশগ্রহণ করতে আগ্রহী হন তাহলে নিম্নের নির্ধারিত স্থানে স্বাক্ষর অথবা বাম বৃদ্ধাঙ্গুলীর ছাপ দিন।

আপনার সহযোগিতার জন্য ধন্যবাদ।

______________________________ ______________________
উত্তরদাতার স্বাক্ষর/বাম বৃদ্ধাঙ্গুলির ছাপ তারিখ

______________________________ ______________________
স্বাক্ষীর স্বাক্ষর/বাম বৃদ্ধাঙ্গুলির ছাপ তারিখ

______________________________ ______________________
পি আই/ গবেষকের প্রতিনিধির স্বাক্ষর তারিখ

PROTOCOL
Biomedical
Non-Exempt
Berkeley

Protocol # 2011-09-3652
Date Printed: 07/31/2018

-----------------------------------------------------------------------------------

| | |
|---|---|
| **Protocol Title:** | Measuring the benefits of sanitation, water quality, handwashing and nutrition interventions for improving health and development in rural Bangladesh |
| **Protocol Status:** | APPROVED |
| **Date Submitted:** | 10/26/2017 |
| **Approval Period:** | **10/30/2017-11/03/2018** |
| **Important Note:** | **This Print View may not reflect all comments and contingencies for approval. Please check the comments section of the online protocol. Questions that appear to not have been answered may not have been required for this submission. Please see the system application for more details.** |

*** * * Attached Document * * ***

| Document Name | Created Date |
|---|---|
| 2011-09-3652_Colford_1k3-Consent-EnvAssess-hygiene-v21-English.pdf | 10/26/2017 |

**Appendix 1k-3: English consent form for environmental assessment (hand rinse and food measurements in hygiene arm)**

Protocol Title: Effect of hand washing, water treatment, sanitation and nutritional supplement interventions on child health and development in rural Bangladesh

Principal Investigator's name: Dr. Stephen P. Luby

#### Purpose of the research

Hello (Assalamualaikum/Namaste). My name is ________ and I work with the ICDDR,B (Cholera Hospital) in Dhaka. We are doing research on diarrheal diseases in children. Through this research we want to learn how certain conditions in sanitation, water quality and hygiene practices impact health. We are conducting this research in collaboration with scientists at University of California, Berkeley in the United States.

**Why are we inviting you to participate in the study?**
We are seeking households that have a child less than 3 years old and have already taken part in other activities from this study. We would like to tell you more detail about this study and hope you and your child will volunteer to participate.

**What is expected from the participants of the research study?**
If you decide to take part in the study, we will conduct some specific activities. During our visit, which we would like to do today, we will ask you questions about your home and compound, and some practices in your home. We will also take a very small amount of stored food that young children in your household will eat. We will also wash your young child's hands in clean water and keep the water used to wash the hands. The visit will be less than 1 hour.

**Risks & Benefits**
There are minimal risks involved in this study. There is a slight risk of breach of confidentiality, but we will do our best to minimize this risk. You will not directly benefit from participating in this study but this participation will enable us to gain knowledge regarding fecal contamination in association with diarrhea.

**Privacy, anonymity and confidentiality**

All data and specimens collected will be kept confidential to the greatest extent possible. . Researchers at ICDDR,B will have access to some of your personal information that could identify you, such as your name and your phone number. They will remove the identifiable information before sharing it with researchers at partner institutions, such as UC Berkeley. Your research records will be kept in a locked cabinet, and computer-based data, will be stored in an encrypted format on a password-protected server.

Other persons who may have access to your information include research groups that oversee the safety of the study including institutes such as the University of California Berkeley and the Bill and Melinda Gates Foundation, USA. None of the researchers outside of ICDDR,B will be able to identify you or your child from information you provide. We will use the information only for the purpose of the study. We will not use your name when we share and publish the results of this study. We expect the steps we take will keep all of your information confidential, but it is possible that because of mistakes or unforeseen events, it could become compromised.

**Future use of information**
If it is necessary, we may share the information we collected from this study with other researchers. If this is done, we will not use your real name and we will maintain your confidentiality. We will store the information we collect from you for a long time after the end of the study.

**Right not to participate and withdraw**
Taking part in the study is completely voluntary. You have the right to decline to allow your child to participate or to withdraw your child at any point in this study without penalty or loss of benefits to which you are otherwise entitled. You may choose not to answer any or all of the questions that will ask. You can drop out of this study at any time, even in the middle of an interview. You have the right to refuse to take part in this study, which will not affect your family's treatment at the Cholera Hospital in the future.

**Principle of compensation**
You do not need to pay us to take part in this study, and we will not pay you money for taking part in the study.

**Persons to contact:**

If you have any questions, you can ask me any time. If you have additional questions about the survey, you may contact: Masud Parvez: phone: 01817541872,Center of Communicable Diseases, ICDDR,B, Mohakhali, Dhaka 1212.

If you have questions about your rights as a participant of a research study, or if you think some harm has been done to you because of this study, you may contact or meet him personally at following address:

M. A. Salam Khan, IRB Secretariat, phone: 9886498 or PABX 8860523-32 ext. 3206

If you agree to participate in our study and for your children to participate in our study, please indicate that by putting your signature or your left thumb impression at the specified space below

Thank you for your cooperation

____________________________________ ____________________

Signature or left thumb impression of Participant Date

____________________________________ ____________________

Signature or left thumb impression of the witness Date

____________________________________ ____________________

Signature of the PI or his/her representative Date

PROTOCOL
Biomedical
Non-Exempt
Berkeley

Protocol # 2011-09-3652
Date Printed: 07/31/2018

-----------------------------------------------------------------------------------
---

**Protocol Title:** Measuring the benefits of sanitation, water quality, handwashing and nutrition interventions for improving health and development in rural Bangladesh

**Protocol Status:** APPROVED

**Date Submitted:** 10/26/2017

**Approval Period:** **10/30/2017-11/03/2018**

**Important Note:** **This Print View may not reflect all comments and contingencies for approval. Please check the comments section of the online protocol.**
**Questions that appear to not have been answered may not have been required for this submission. Please see the system application for more details.**

*** * * Attached Document * * ***

| Document Name | Created Date |
| --- | --- |
| 2011-09-3652_Colford_2k3-Consent-EnvAssess-hygiene-v19-Bengali.pdf | 10/26/2017 |

#### Appendix 2k-3: English consent form for environmental assessment (hand rinse measurements in hygiene arm)

পরিবেশগত প্রভাব মূল্যায়নের জন্য সম্মতিপত্র (হাত ধোয়া পরিমাপ)

গবেষণার শিরোনাম: বাংলাদেশের গ্রামীণ এলাকায় হাত ধোয়া, পানি বিশুদ্ধকরণ, স্যানিটেশন ও পুষ্টিকর সম্পূরক খাবার প্রদান এবং শিশুর স্বাস্থ্য ও বৃদ্ধির উপর তার প্রভাব।

**প্রধান গবেষক:** ডঃ স্টিফেন পি. লুবি

**গবেষণার উদ্দেশ্য:**

আসলামুআলাইকুম/আদাব। আমার নাম ----------------------------। আমরা ঢাকা (মহাখালী) কলেরা হাসপাতালে কাজ করি। আমরা শিশুদের ডায়রিয়াজনিত রোগ-বালাই নিয়ে গবেষণা করছি। কতিপয় স্যানিটেশন ব্যবস্থা, পানির গুনাগুন এবং স্বাস্থ্যসম্মত অভ্যাসসমূহ স্বাস্থ্যের উপর কিভাবে প্রভাব ফেলে এই গবেষণার মাধ্যমে আমরা তা বোঝার চেষ্টা করছি।

**আমরা কেন আপনাকে এই গবেষণায় অংশগ্রহনে আমন্ত্রণ জানাচ্ছি?**

আমরা আপনাকে/আপনার খানাকে এই গবেষণায় অংশগ্রহণের জন্য আমন্ত্রন জানাচ্ছি কারণ আপনার ৩ বছরের কম বয়সের একটি শিশু আছে এবং আপনি/আপনার খানা ইতোমধ্যেই আমাদের গবেষণার অন্যান্য কর্মকান্ডে অংশগ্রহন করেছেন। আমরা আপনাকে/আপনাদেরকে এই গবেষণার বিষয়ে আরো বিস্তারিতভাবে জানাতে চাই। আমরা আশা করছি আপনি এবং আপনার শিশু স্বেচ্ছায় এই গবেষণায় অংশগ্রহন করবেন।

**গবেষণায় অংশগ্রহণকারীর কাছে প্রত্যাশা কী?**

যদি আপনি/আপনার খানা এই গবেষণায় অংশগ্রহণে সম্মত থাকেন, তাহলে আমরা আপনার খানাতে নির্দিষ্ট কিছু কর্মকান্ড পরিচালনা করব। আজকে পরিদর্শনকালে আমরা আপনাকে আপনার বাড়ি ও বাড়ির চারপাশের পরিবেশ, এবং আপনার বাড়িতে প্রচলিত অভ্যাস সম্পর্কে কিছু প্রশ্ন জিজ্ঞাসা করব। আমরা পরিষ্কার পানি দিয়ে আপনার ছোট শিশুর হাতগুলো ধোয়াব এবং হাতধোয়া সেই পানিটা নমুনা হিসাবে সংগ্রহ করব। পরিদর্শনকালে আমরা আপনার খানাতে প্রায় ১ ঘন্টা সময় কাটাব।

**ঝুঁকি এবং সুবিধাসমূহ:**

এই গবেষণায় আপনার এবং আপনার শিশুর ঝুঁকির সম্ভাবনা নুন্যতম। আপনার পরিচয়সংক্রান্ত তথ্যসমূহ গোপন না থাকার সামান্য সম্ভাবনা রয়েছে কিন্তু আমরা আপনার পরিচয়সংক্রান্ত তথ্যসমূহ গোপন রাখার জন্য সম্ভাব্য সবধরনের চেষ্টা করব। কিন্তু আপনার শিশুর অংশগ্রহণ, শিশুদের ডায়রিয়া রোগ সম্পর্কে জ্ঞানলাভে আমাদেরকে সহায়তা করবে।

**নিজস্বতা, গোপনীয়তা ও বিশ্বাসযোগ্যতা:**

আমরা আপনার/ আপনাদের দেওয়া সমস্ত তথ্যের গোপনীয়তা বজায় রাখার জন্য সর্বোচ্চ চেষ্টা করব। আপনাদের দেওয়া কিছু তথ্য আইসিডিডিআর,বি র গবেষকদের কাছে থাকবে যার ফলে আপনাকে সনাক্ত করা সম্ভব, যেমন আপনার নাম এবং ফোন নম্বর। তারা আপনাদের দেওয়া তথ্য থেকে আপনার সনাক্তকরণ মুলক তথ্য সমুহ বাদ দিয়ে অন্যান্য গবেষকদের সাথে শেয়ার করবে (যেমন আমেরিকার ক্যালিফোর্নিয়া বার্কলে বিশ্ববিদ্যালয়ের গবেষক), যার ফলে কোন গবেষকই আপনাকে চিহ্নিত করতে পারবে না। গবেষণার সব রেকর্ড তালাবদ্ধ করে রাখা হবে ও কমপিউটারে তথ্যাবলী পাসওআরড দিয়ে সংরক্ষন করা হবে ।

অন্যান্য ব্যক্তি অথবা গবেষণার সাথে সংশ্লিষ্ট দল আপনার তথ্য দেখতে পারবে যদি গবেষণার কারণে প্রয়োজন মনে করে। এরা হলেন- আমেরিকার ক্যালিফোর্নিয়া বার্কলে বিশ্ববিদ্যালয়ের গবেষক ও গবেষণার স্পন্সর (বিল ও মেলিন্ডা গেটস ফাউন্ডেশন)। আইসিডিডিআর,বি র গবেষকদের ছাড়া কেউ আপনাকে এবং আপনার শিশুকে সনাক্ত করতে পারবে না । আমরা শুধুমাত্র গবেষণার কাজে এই তথ্য ব্যবহার করব এবং আপনার নাম অথবা পরিচয়ের কোন সূত্র এই গবেষনা-সংক্রান্ত কোন রিপোর্টে/ প্রকাশনায়

উল্লেখ করা হবে না। আপনার কাছ থেকে প্রাপ্ত সকল তথ্য ও নমুনার সর্বোচ্চ গোপনীয়তা রক্ষার জন্য আমরা সর্বাত্মক চেষ্টা করব, তবুও ভুলবশতঃ অনাকাঙ্খিত কোন ঘটনার কারনে গোপনীয়তা রক্ষা নিশ্চিত করতে না পারলে সেজন্য মার্জনা প্রার্থনা করছি।

#### ভবিষ্যতে তথ্যের ব্যবহার:

এই গবেষণায় সংগৃহীত তথ্য প্রয়োজনে আমরা অন্যান্য গবেষকের সাথে বিনিময় করতে পারি। এবং কারো সাথে তথ্য বিনিময় করা হলে আমরা সেখানে আপনার প্রকৃত নাম ব্যবহার করব না এবং আপনার দেওয়া তথ্যের গোপণীয়তা কঠোরভাবে রক্ষা করব।

#### গবেষণায় অংশগ্রহন না করা বা নাম প্রত্যাহারের অধিকারঃ

এ গবেষণায় আপনার শিশুর অংশগ্রহণ সেচ্ছামূলক এবং আপনার শিশুর অংশগ্রহণ করা বা না করা সম্পূর্ণভাবে আপনার ইচ্ছার উপর নির্ভর করছে। এই গবেষণায় আপনাকে যেসব প্রশ্ন জিজ্ঞাসা করা হবে তার যেকোনটির উত্তর আপনি চাইলে নাও দিতে পারেন। এই গবেষণায় অংশগ্রহনে সম্মত হবার পরেও গবেষণা চলাকালীন যেকোন সময় আপনি আপনার নাম প্রত্যাহার করে নিতে পারেন। এই গবেষণায় অংশগ্রহণ না করার অধিকার আপনার আছে এবং এ জন্য ভবিষ্যতে আপনার পরিবারের সদস্যদের কলেরা হাসপাতালে স্বাস্থ্যসেবা গ্রহণের উপর কোন প্রভাব পড়বে না।

## ক্ষতিপূরণ :

এই গবেষণায় অংশগ্রহণের জন্য আপনার কোন ধরনের অর্থনৈতিক খরচ/ ব্যয় বহন করতে হবে না এবং একইভাবে আপনাকেও আমরা কোন ধরনের অর্থ প্রদান করব না।

## যোগাযোগ:

আপনার কোনো প্রশ্ন থাকলে যেকোন সময় আপনি তা আমাকে জিজ্ঞাসা করতে পারেন। যদি গবেষণা সম্পর্কিত অতিরিক্ত কোন প্রশ্ন থাকে তাহলে আপনি জনাব মাসুদ পারভেজ (প্রোগ্রাম অন ইনফেক্সাস ডিজিজেজ এন্ড ভ্যাকসিন সাইন্সেস, আইসিডিডিআর,বি: মহাখালি, ঢাকা-১২১২) এর সাথে ০১৮১৭৫৪১৮৭২ মোবাইল নম্বরে সরাসরি যোগাযোগ করতে পারেন। এই গবেষণায় আপনার অধিকার-সংক্রান্ত কোন প্রশ্ন থাকলে অথবা গবেষণার কারণে আপনার অর্থনৈতিক কোন ক্ষতি সাধিত হয়েছে বলে মনে করেন, তাহলে আপনি ৯৮৮৬৪৯৮ অথবা ৮৮৬০৫২৩-৩২ এক্স- ৩২০৬ টেলিফোন নম্বরে অথবা সরাসরি এম এ সালাম খান, আইসিডিডিআরবি সেক্রেটারিয়েট এর সাথে যোগাযোগ করতে পারেন।

আপনি যদি এই গবেষণায় অংশগ্রহণ করতে আগ্রহী হন তাহলে নিম্নের নির্ধারিত স্থানে স্বাক্ষর অথবা বাম বৃদ্ধাঙ্গুলীর ছাপ দিন।

আপনার সহযোগিতার জন্য ধন্যবাদ।

________________________ ____________________

উত্তরদাতার স্বাক্ষর/বাম বৃদ্ধাঙ্গুলির ছাপ তারিখ

________________________ ____________________

স্বাক্ষীর স্বাক্ষর/বাম বৃদ্ধাঙ্গুলির ছাপ তারিখ

________________________ ____________________

পি আই/ গবেষকের প্রতিনিধির স্বাক্ষর তারিখ

PROTOCOL
Biomedical
Non-Exempt
Berkeley

Protocol # 2011-09-3652
Date Printed: 07/31/2018

---------------------------------------------------------------------------------
---

**Protocol Title:** Measuring the benefits of sanitation, water quality, handwashing and nutrition interventions for improving health and development in rural Bangladesh

**Protocol Status:** APPROVED

**Date Submitted:** 10/26/2017

**Approval Period:** **10/30/2017-11/03/2018**

**Important Note:** **This Print View may not reflect all comments and contingencies for approval. Please check the comments section of the online protocol. Questions that appear to not have been answered may not have been required for this submission. Please see the system application for more details.**

*** * * Attached Document * * ***

| Document Name | Created Date |
| --- | --- |
| 2011-09-3652_Colford_1I-Consent-5hr-SO-v18-English.pdf | 10/26/2017 |

**Appendix 1L: English consent form for 5-hour structured observation**

Protocol Title: Effect of hand washing, water treatment, sanitation and nutritional supplement interventions on child health and development in rural Bangladesh

Principal Investigator's name: Dr. Stephen Luby

**Purpose of the research**

Hello (Assalamualaikum/Nomoshkar). My name is ________ and I work with the ICDDR,B (Cholera Hospital) in Dhaka. We are interested in conducting research on diarrheal diseases in children to learn how it affects their physical and mental development. Through this research we want to learn about the direct health benefits of some simple interventions to improve the health of children under three.

**Why are we inviting you to participate in the study?**

Because you are expecting to deliver a baby in the next 8 months/you have a child under 3 years, and we are interested in evaluating child development early in life, we would like to invite you to participate in this study.

**What is expected from the participants of the research study?**

If you agree to take part, I will find a place to sit in your home or courtyard so that I will not be in your way. I will observe hygiene related activities in your home and take some notes. I may look around the courtyard to observe some of these practices and may ask you some questions in the end. I will be here for a total of about five hours.

**Risks & Benefits**

There is no cost to you for participation in this research. You will not receive any compensation for being in the study. We will only collect information related to health and hygiene. The process of having someone in your home for a longer time may be uncomfortable to you. However, we do not expect any harm to come to you or your family because of being observed.

**Privacy, anonymity and confidentiality**

We will secure all information collected from you in a locked cabinet, and none other than designated staff of this research study will have access to that information. We would, however, like to inform you that disclosure of your information is subject to the laws of Bangladesh. Your name and identity will not be used in reporting and presenting study findings.

**Right not to participate and withdraw**

You are free to decide whether or not to take part in the study. If you decide against participating in the study, and also if you withdraw your consent at any time after participation, you and your family will continue to receive the same benefits that you have otherwise been eligible for.

**Persons to contact:**

If you have any question, you can ask me any time. If you have additional questions about the survey, you may contact:

Abu Naser, Center of Communicable Diseases, ICDDR,B, Mohakhali, Dhaka 1212. Phone: 8860523-32 # 120

If you have questions about your rights as a participant of a research study, or if you think some harm has been done to you because of the survey, you may contact or meet him personally at following address:

M. A. Salam Khan, IRB Secretariat, phone: 9886498 or PABX 8860523-32 ext. 3206

If you agree to our proposal of enrolling your household in our study, please indicate that by putting your signature or your left thumb impression at the specified space below

Thank you for your cooperation

_________________________ ________________ ___/______/________
Name of study representative Signature of Investigator Date:

_____________________ __________________________________ ___/____/__
Name of study participant Signature/thumb impression of participant Date:

PROTOCOL
Biomedical
Non-Exempt
Berkeley

Protocol # 2011-09-3652
Date Printed: 07/31/2018

---------------------------------------------------------------------------------
---

| | |
|---|---|
| **Protocol Title:** | Measuring the benefits of sanitation, water quality, handwashing and nutrition interventions for improving health and development in rural Bangladesh |
| **Protocol Status:** | APPROVED |
| **Date Submitted:** | 10/26/2017 |
| **Approval Period:** | **10/30/2017-11/03/2018** |
| **Important Note:** | **This Print View may not reflect all comments and contingencies for approval. Please check the comments section of the online protocol. Questions that appear to not have been answered may not have been required for this submission. Please see the system application for more details.** |

*** * * Attached Document * * ***

| Document Name | Created Date |
|---|---|
| 2011-09-3652_Colford_2I-Consent-5hr-SO-v18-Bengali.pdf | 10/26/2017 |

| Q.T/ S.G/ Cl. ID/ Co. ID/ B. ID/ HH ID |
|---|
| Household ID: ☐/☐/ ☐☐/ ☐☐/☐☐/☐☐ |

### Appendix 2L: Bengali consent for 5-hour structured observation

**স্বেচ্ছা সম্মতিপত্রঃ পাঁচ ঘনটা ব্যাপি পর্যবেক্ষণ**

আন্তর্জাতিক উদরাময় গবেষণা কেনদ্র বাংলাদেশ (আই সি ডি ডি আর, বি )

গবেষণার শিরোনাম:

বাংলাদেশের গ্রামীণ এলাকায় হাত ধোয়া, পানি বিশুদ্ধকরণ, স্যানিটেশন এবং সম্পূরক খাবার প্রদান এবং শিশুর স্বাস্থ্য ও বৃদ্ধির উপর তার প্রভাব

প্রধান গবেষকের নাম: ড. স্টিফেন পি লুবি

গবেষনার উদ্দেশ্য:

আস্‌সালামুআলাইকুম। আমার নাম ( ___________ ) এবং আমি আইসিডিডিআর, বি (কলেরা হাসপাতাল) তে কাজ করি। আমরা শিশুদের ডায়রিয়া রোগ এবং ইহা কিভাবে তাদের শারীরিক ও মানসিক বৃদ্ধিকে প্রভাবিত করে তা নিয়ে গবেষণা করতে আগ্রহী। আমরা এই গবেষণার মাধ্যমে তিন বছরের ছোট বাচ্চাদের স্বাস্থ্য উন্নয়নের মাধ্যমে স্বাস্থ্যগত সুবিধা সম্পর্কে জানতে চাচ্ছি।

আমরা কেন আপনাকে এই গবেষনায় অংশগ্রহনে আমন্ত্রণ জানাচ্ছি?

আমরা আপনাকে এই গবেষণায় অংশগ্রহণের জন্য আমন্ত্রন জানাচ্ছি কারণ আপনি পরবর্তী ৮ মাসের মধ্যে একটি শিশুর জন্ম দেবেন বা আপনার বাড়ীর ৩ বছরের কম বয়সের শিশু আছে এবং আমরা শিশুদের জীবনের শুরু থেকেই তাদের বৃদ্ধি পর্যবেক্ষণ করতে আগ্রহী।

গবেষনায় অংশগ্রহণকারীর কাছে প্রত্যাশা কী ?

আপনি যদি এই গবেষণায় অংশগ্রহণ করতে চান, তাহলে আমি আপনার ঘরে বা উঠানে বা আঙিনায় এমন একটি জায়গায় বসব যেখান থেকে আপনি এবং আপনার খানার অন্যান্য সদস্যগণ কখন কোথায় কি করছেন তা দেখতে পারি কিন্তু তাতে যেন আপনাদের দৈনন্দিন কাজে কোন সমস্যার সৃষ্টি না হয়। আমি আপনার বাড়ির দৈনন্দিন কাজর্কম র্পযবেক্ষণ করার জন্য আপনার ঘরের ভিতর এবং বাহিরের চারপাশ একটু ঘুরে দেখব এবং পর্যবেক্ষন শেষে আপনাকে কিছু প্রশ্ন করব। এছাড়া আমি কিছু তথ্য লিখে রাখব। প্রায় পাঁচ ঘণ্টা ধরে পর্যবেক্ষণের জন্য আপনার অনুমতি সাপেক্ষে আমি আপনার খানাতে থাকব।

ঝুঁকি এবং সুবিধা:

এই গবেষণায় অংশগ্রহণে আপনার কোনো ঝুঁকি নেই। এই গবেষণায় অংশগ্রহণের জন্য আপনি তাৎক্ষণিক কোন সুবিধা পাবেন না। আমরা শুধুমাত্র খানাতে দৈনন্দিন স্বাস্থ্যগত আচরণ সম্পর্কিত তথ্য সংগ্রহ করব। তবে গবেষণার কাজে পাঁচ ঘনটা সময়ের জন্য আপনার খানাতে/ বাড়িতে থাকাটা হয়তোবা আপনাদের জন্য কিছুটা অসস্বিদায়ক হতে পারে। তবে পর্যবেক্ষণের মাধ্যমে উত্তর সংগ্রহে আপনার বা আপনার পরিবারের জন্য কোন অসুবিধা হবে না বলে আশা করছি। তবে আমরা আশা করছি এই গবেষণায় প্রাপ্ত ফলাফল বাংলাদেশের জনসাধারণের পানি, পয়ঃব্যবস্থা এবং স্বাস্থ্যসম্মত অবস্থা বুঝতে সহায়তা করবে এবং এই অবস্থার উন্নতি সাধনের লক্ষ্যে কাজ করার সুযোগ সৃষ্টি হবে।

খরচ এবং ক্ষতিপূরণ :

এই গবেষণায় অংশগ্রহণের জন্য আপনার সারসরি কোন ধরনের অর্থনৈতিক খরচ হবে না এবং আপনি গবেষণায় অংশগ্রহনের জন্য সরাসরি কোন ধরণের অর্থনৈতিক বা অন্য কোন সহায়তাও পাবেন না।

নিজস্বতা, গোপনীয়তা ও বিশ্বাসযোগ্যতা :

এই দেশের আইন অনুযায়ী আমরা আপনার/ আপনাদের দেওয়া সমস্ত তথ্যের গোপনীয়তা বজায় রাখার জন্য সর্বোচ্চ চেষ্টা করব। সংগৃহীত তথ্য এবং এর ফলাফল গোপনীয়তার সাথে রক্ষণাবেক্ষণ করা হবে। আমরা শুধুমাত্র গবেষণার কাজে এই তথ্য ব্যবহার করব এবং আপনার নাম অথবা পরিচয়ের কোন সূত্র এই গবেষনা-সংক্রান্ত কোন রিপোর্ট/ প্রকাশনায় উল্লেখ করা হবে না।

স্বেচ্ছা অংশগ্রহণ :

এই গবেষণায় অংশগ্রহণ সম্পূর্ণভাবে আপনার ইচ্ছার ওপর র্নিভর করে। আপনার লিখিত অনুমতি দেয়ার পরবর্তীতেও আপনি ইচ্ছা করলে যে কোন সময়ে আপনার অনুমতি প্রত্যাখ্যান করতে পারেন। এই গবেষণায় অংশগ্রহণ না করার জন্য অথবা অংশগ্রণের পরবর্তীতে কোন এক পর্যায়ে অংশগ্রহণে অসম্মতি প্রকাশ করার জন্য আপনার পরিবার কোন ধরনের স্বাভাবিক বা প্রাপ্য সুযোগ থেকে কখনই বঞ্চিত হবেন না।

যোগাযোগ:

আপনার কোনো প্রশ্ন থাকলে আমাকে জিজ্ঞাসা করতে পারেন। যদি গবেষণা সম্পর্কিত অতিরিক্ত কোন প্রশ্ন থাকে তাহলে আপনি **ডাঃ আবু নাছের**,(পিআইডিভিএস, আইসিডিডিআর,বিঃ মহাখালি, ঢাকা-১২১২) এর সাথে ৮৮১৯৪১৯-২০ (এক্স- ১২০) টেলিফোন নম্বরে সরাসরি যোগাযোগ করতে পারেন। এই গবেষণায় আপনার অধিকার-সংক্রান্ত কোন প্রশ্ন থাকলে অথবা গবেষনার কারণে আপনার কোন ক্ষতি হতে পারে বলে যদি মনে করেন, তাহলে আপনি ৯৮৮৬৪৯৮ অথবা ৮৮৬০৫২৩ (এক্স- ৩২০৬) টেলিফোন নম্বরে অথবা সরাসরি এম এ সালাম খান, আইসিডিডিআরবি সেক্রেটারিয়েট এর সাথে যোগাযোগ করতে পারেন।

আপনি যদি এই গবেষণায় অংশগ্রহণ করতে আগ্রহী হন তাহলে নিম্নের নির্ধারিত স্থানে স্বাক্ষর অথবা বাম বৃদ্ধাঙ্গুলীর ছাপ দিন।

আপনার সহযোগিতার জন্য ধন্যবাদ।

............................................ ..................................

উত্তরদাতার স্বাক্ষর/বাম বৃদ্ধাঙ্গুলির ছাপ তারিখ

............................................ ..................................

সাক্ষীর স্বাক্ষর/বাম বৃদ্ধাঙ্গুলির ছাপ তারিখ

............................................ ..................................

পি আই/ গবেষকের প্রতিনিধির স্বাক্ষর তারিখ

PROTOCOL
Biomedical
Non-Exempt
Berkeley

Protocol # 2011-09-3652
Date Printed: 07/31/2018

-----------------------------------------------------------------------------------
---

**Protocol Title:** Measuring the benefits of sanitation, water quality, handwashing and nutrition interventions for improving health and development in rural Bangladesh

**Protocol Status:** APPROVED

**Date Submitted:** 10/26/2017

**Approval Period:** **10/30/2017-11/03/2018**

**Important Note:** **This Print View may not reflect all comments and contingencies for approval. Please check the comments section of the online protocol. Questions that appear to not have been answered may not have been required for this submission. Please see the system application for more details.**

*** * * Attached Document * * ***

| Document Name | Created Date |
|---|---|
| 2011-09-3652_Colford_1n-Consent-Lead exposure-v17-English.pdf | 10/26/2017 |

**Appendix 1n: English consent form for household in-depth lead exposure interview**

Protocol Title: Effect of hand washing, water treatment, sanitation and nutritional supplement interventions on child health and development in rural Bangladesh

Principal Investigator's name: Dr. Leanne Unicomb

**Purpose of the research**

Hello (Assalamualaikum/Nomoshkar). My name is ________ and I work with the ICDDR,B (Cholera Hospital) in Dhaka. We are interested in conducting research on exposures that people in this area have to lead. Sometimes lead exposure can cause illness so we want to understand them better.

**Why are we inviting you to participate in the study?**

Because a sample of blood collected from you had an elevated level of lead. We are interested in exploring how you and your family might be exposed.

**What is expected from the participants of the research study?**

If your household decides to enroll in we will ask you a series of questions. We will ask also questions of the person who does most of the farming. We will want to look at your home and the fields your family works in. As it is very difficult to write all the conversation in a note pad, so, if you agree, we would like to record this sessions in a tape recorder. We would also like to take some notes from this session. This session will last about an hour. We would like to return to ask additional questions sometime in the future. Your decision to participate in this part of the study is optional and will not affect your household participation in WASH Benefits.

**Risks & Benefits**

We do not forsee any risk to the study. If we identify a source of lead, we will make recommendations on how to limit your family's exposure.

**Privacy, anonymity and confidentiality**

All data collected will be kept confidential as allowed by the law of this country. Other persons who may have access to your test results include research groups that oversee the safety of the study including institutes such as the University of California Berkeley and the Bill and Melinda Gates Foundation, USA. Because of the potential need to release information to these parties, we cannot guarantee absolute confidentiality. However, we will ensure every effort that we can to preserve your confidentiality. There is no expiration date to this permission. This information will be coded and it is unlikely that anyone will be able to trace it to you and your child, protecting your privacy.

**Future use of information**

The information collected from this study may be shared with other researchers if needed, but we will strictly maintain your confidentiality and privacy.

**Right not to participate and withdraw**

Taking part in the study is completely voluntary. "If, at any point during the study, you decide you do not wish to participate, we will withdraw your permission and we will destroy your soil, rice samples and data recorded from you." You have the right to refuse participation in this study, which will not affect your family's treatment at the Cholera Hospital in the future.

**Principle of compensation**

You need not pay us to take part in this study, and similarly we will not pay you money for attending in the study.

**Persons to contact**:

If you have any question, you can ask me any time. If you have additional questions about the survey, you may contact:

Dr. Mahbubur Rahman, Centre of Communicable Diseases, ICDDR,B, Mohakhali, Dhaka 1212. Phone: 8860523-32 # 120

If you have questions about your rights as a participant of a research study, or if you think some harm has been done to you because of the survey, you may contact or meet him personally at following address:

M. A. Salam Khan, IRB Secretariat, phone: 9886498 or PABX 8860523-32 ext. 3206
If you agree to our proposal of enrolling your household in our study, please indicate that by putting your signature or your left thumb impression at the specified space below
Thank you for your cooperation

Consent of the mother for sample collection: YES_______ NO ______

_____________________________________ _____________________
Signature or left thumb impression of Participant Date

_____________________________________ ___________________
Signature of the PI or his/her representative Date

PROTOCOL
Biomedical
Non-Exempt
Berkeley

----------------------------------------------------------------------------------

| | |
|---|---|
| **Protocol Title:** | Measuring the benefits of sanitation, water quality, handwashing and nutrition interventions for improving health and development in rural Bangladesh |
| **Protocol Status:** | APPROVED |
| **Date Submitted:** | 10/26/2017 |
| **Approval Period:** | **10/30/2017-11/03/2018** |
| **Important Note:** | **This Print View may not reflect all comments and contingencies for approval. Please check the comments section of the online protocol. Questions that appear to not have been answered may not have been required for this submission. Please see the system application for more details.** |

*** * * Attached Document * * ***

| Document Name | Created Date |
|---|---|
| 2011-09-3652_Colford_2n-Consent-Lead exposure-v17-Bengali.pdf | 10/26/2017 |

icddr,b
KNOWLEDGE FOR GLOBAL LIFESAVING SOLUTIONS

### Appendix 2n: Bengali consent for for household in-depth lead exposure interview

**এপেন্ডিক্স ১এনঃ সীসার সংস্পর্শে আসা খানার নিবিড় সাক্ষাৎকারের জন্য সম্মতিপত্র**

গবেষণার শিরোনাম: বাংলাদেশের গ্রামীণ এলাকায় হাত ধোয়া, পানি বিশুদ্ধকরণ, স্যানিটেশন ও পুষ্টিকর সম্পূরক খাবার প্রদান এবং শিশুর স্বাস্থ্য ও বৃদ্ধির উপর তার প্রভাব।

**প্রধান গবেষকঃ** ডঃ লিয়েন ইউনিকম্ব

**গবেষণার উদ্দেশ্যঃ**

আসসালামু আলাইকুম/নমস্কার। আমার নাম----------- এবং আমি ঢাকায় আইসিডিডিআর,বি তে (কলেরা হাসপাতাল) কাজ করি। এই এলাকার মানুষজনের সীসার সংস্পর্শে আসার বিষয়টি নিয়ে আমরা গবেষণা করতে ইচ্ছুক। সীসার সংস্পর্শতা মাঝে মাঝে অসুস্থ্যতার কারণ হতে পারে, তাই আমরা এই বিষয়টি ভালভাবে বুঝতে চাই।

**কেন আপনাকে আমরা এই গবেষণায় অংশগ্রহণের জন্য আমন্ত্রন জানাচ্ছি?**

কারণ আপনার কাছ থেকে সংগৃহীত রক্তে উচ্চ মাত্রার সীসার উপস্থিতি রয়েছে। আমরা খুঁজে বের করতে ইচ্ছুক কিভাবে আপনি আক্রান্ত হয়েছেন এবং আপনার পরিবার এটা দ্বারা আক্রান্ত হতে পারে।

**গবেষণায় অংশগ্রহণকারীর কাছে প্রত্যাশা**

আপনার পরিবার যদি এই গবেষণায় অংশগ্রহণের সিদ্ধান্ত নেন তবে আমরা ধারাবাহিকভাবে কিছু প্রশ্ন করব। যারা কৃষিকাজে বেশিরভাগ সময়ে নিয়োজিত থাকেন আমরা তাদেরকে আরো প্রশ্ন করব। আমরা আপনার বাড়ী ও কৃষি জমি দেখতে চাইব যেখানে আপনার পরিবারের সদস্যরা কাজ করেন। যেহেতু আপনার সাথে কথোপকথনের সম্পূর্ণ বিবরণটি খাতায় লেখা কষ্টকর, তাই আপনি যদি রাজী থাকেন, আমরা আপনার সাথে কথোপকথনের বিষয়টি টেপ রেকর্ড করতে চাই। আমরা একই সাথে কিছু বিষয় খাতায়ও লিখে রাখব। এর জন্য এক ঘনটার মত সময় লাগবে। আরও কিছু প্রশ্ন করার জন্য আমরা ভবিষ্যতে কোন এক সময় আবারও আসতে পারি। গবেষণার এই পর্বে অংশগ্রহণে আপনার সিদ্ধান্ত সম্পূর্ণরূপে ঐচ্ছিক এবং এটা ওয়াশ বেনিফিট কার্যক্রমে আপনার পরিবারের অংশগ্রহণে কোন প্রভাব ফেলবে না।

**ঝুঁকি এবং সুবিধা**

এই গবেষণায় আমরা কোন ঝুঁকির সম্ভাবনা দেখছি না। আমরা যদি সীসার উৎস চিহ্নিত করতে পারি, কিভাবে আপনার পরিবারে সীসা দ্বারা আক্রান্ত হওয়ার মাত্রা রোধ করা যায়, সে বিষয়ে আমরা পরামর্শ দেব ।

**গোপনীয়তা, নাম হীনতা ও বিশ্বস্ততা**

আপনার কাছ থেকে প্রাপ্ত সকল তথ্য ও নমুনা এই দেশের আইন অনুযায়ী গোপন রাখা হবে। কোন গবেষকই আপনাকে চিহ্নিত করতে পারবে না। গবেষণা দল যারা এই গবেষণার নিরাপত্তার তত্ত্বাবধানে রয়েছেন এবং আমেরিকার ক্যালিফোর্নিয়া বার্কলে বিশ্ববিদ্যালয় এবং বিল ও মেলিন্ডা গেটস ফাউন্ডেশন আপনার দেওয়া তথ্য এবং এই অডিও রেকর্ডিং ব্যবহার করতে পারবেন। যেহেতু এই প্রতিষ্ঠানসমূহ ও ব্যক্তিবর্গকে যে কোন সময় তথ্য দিতে হতে পারে তাই পুরোপুরি গোপনীয়তা রক্ষার বিষয়টি আমরা নিশ্চিত করতে পারছি না। তবুও সর্বোচ্চ গোপনীয়তা রক্ষার জন্য আমরা সর্বাত্মক চেষ্টা করব। এদের এই তথ্য ব্যবহারে অনুমতির জন্য কোন নির্ধারিত সময়সীমা নেই। আপনার গোপনীয়তা রক্ষার্থে এই তথ্যগুলোকে কোড করা হবে যার ফলে কারো পক্ষে আপনাকে সনাক্ত করা প্রায় অসম্ভব।

**ভবিষ্যতে তথ্যের ব্যবহার**

এই গবেষণা হতে প্রাপ্ত তথ্য প্রয়োজনে অন্য গবেষকদের সাথে আদান-প্রদান করতে হতে পারে। তবে আমরা খুব সতর্কতার সাথে আপনার বিশ্বস্ততা ও গোপনীয়তা রক্ষা করব।

**গবেষণায় অংশগ্রহণ না করার ও নিজেকে প্রত্যাহারের অধিকার**

এই গবেষণায় আপনার অংশগ্রহণ সম্পূর্ণরূপে স্বেচ্ছাসেবামূলক। যদি গবেষণা চলাকালে যেকোন সময়েও আপনি উক্ত গবেষণায় অংশগ্রহন না করার সিদ্ধান্ত গ্রহন করেন, আমরা আপনার অংশগ্রহণের সম্মতি প্রত্যাহার করেদিব এবং আমরা আপনার কাছ থেকে নেওয়া চাল, মাটির নমুনা এবং আপনার দেওয়া সকল তথ্য নষ্ট করে ফেলব। এই গবেষণা থেকে নিজেকে প্রত্যাহার করার অধিকার আপনার আছে, এজন্য কলেরা হাসপাতালে আপনার পরিবারের চিকিৎসাক্ষেত্রে কোন সমস্যা হবে না।

**ক্ষতিপূরণ নীতি**

এই গবেষণায় অংশগ্রহণের জন্য আপনাকে কোন টাকাপয়সা দিতে হবে না একই ভাবে আমাদের পক্ষ থেকেও আপনাকে কোন টাকা পয়সা দেয়া হবেনা।

**যোগাযোগের ব্যক্তিবর্গ**

আপনার যদি কোন প্রশ্ন থাকে, তবে তা যে কোন সময়ে আমাকে জিজ্ঞাসা করতে পারেন। এই গবেষণা/ জরিপ বিষয়ে অপনার যদি অতিরিক্ত কোন প্রশ্ন থাকে তবে তা জানার জন্য যোগাযোগ করতে পারেন। যোগাযোগের ঠিকানাঃ

ডাঃ মাহাবুবুর রহমান, সেন্টার ফর কমিউনিকেবল ডিজিজেস, আইসিডিডিআর,বি, মহাখালী, ঢাকা-১২১২। ফোন: ৮৮৬০৫২৩-৩২ # ১২৩।

একজন গবেষণা অংশগ্রহণকারী হিসেবে আপনার অধিকার সম্পর্কে যদি জানতে চান অথবা যদি আপনি মনে করেন যে আপনার কাছ থেকে তথ্য সংগ্রহের কারণে আপনার কোন ক্ষতি হয়েছে তাবে আপনি নিম্নোক্ত ঠিকানায় যোগাযোগ অথবা সরাসরি দেখা করতে পারেন। যোগাযোগের ঠিকানাঃ

এম, এ সালাম খান, আই আর বি সেক্রেটারিয়েট, ফোনঃ ৯৮৮৬৪৯৮ অথবা PABX, ৮৮৬০৫২৩-৩২, এক্স. ৩২০৬।

আপনি যদি আমাদের প্রস্তাব অনুযায়ী আপনার খানাকে এই গবেষণায় যুক্ত করতে সম্মত হন তবে নীচে উলেস্নখিত স্থানে আপনার স্বাক্ষর অথবা বাম হাতের বৃদ্ধাঙ্গুলীর ছাপ দিন। আপনার সহযোগিতার জন্য ধন্যবাদ।

নমুনা সংগ্রহের জন্য মা'র সম্মতি: হ্যাঁ ---------না--------

---------------------------------------- ----------------------

অংশগ্রহণকারীর স্বাক্ষর/ বাম হাতের বৃদ্ধাঙ্গুলীর ছাপ তারিখ

---------------------------------------- ----------------------

প্রধান গবেষক অথবা তার প্রতিনিধির স্বাক্ষর তারিখ

PROTOCOL
Biomedical
Non-Exempt
Berkeley

Protocol # 2011-09-3652
Date Printed: 07/31/2018

-----------------------------------------------------------------------------------

**Protocol Title:** Measuring the benefits of sanitation, water quality, handwashing and nutrition interventions for improving health and development in rural Bangladesh

**Protocol Status:** APPROVED

**Date Submitted:** 10/26/2017

**Approval Period:** **10/30/2017-11/03/2018**

**Important Note:** **This Print View may not reflect all comments and contingencies for approval. Please check the comments section of the online protocol. Questions that appear to not have been answered may not have been required for this submission. Please see the system application for more details.**

*** * * Attached Document * * ***

| Document Name | Created Date |
|---|---|
| 2011-09-3652_Colford_1o-Consent-Agrochemical-v17-English.pdf | 10/26/2017 |

**Appendix 1o: English consent form for agrochemical in-depth interview**

Protocol Title: Effect of hand washing, water treatment, sanitation and nutritional supplement interventions on child health and development in rural Bangladesh

Principal Investigator's name: Dr. Leanne Unicomb

**Purpose of the research**

Hello (Assalamualaikum/Nomoshkar). My name is ________ and I work with the ICDDR,B (Cholera Hospital) in Dhaka. We are interested in conducting research on exposures that people in this area have to lead. Sometimes lead exposure can cause illness so we want to understand them better.

**Why are we inviting you to participate in the study?**

Sometimes chemicals used in agriculture contain lead. We want to understand what chemicals the farmers in this area purchase.

**What is expected from the participants of the research study?**

If you decide to enroll in we will ask you a series of questions. We will want to look at your shop. We will also collect a small sample of soil and some rice that is cultivated from your land. We may want to come and talk again. As it is very difficult to write all the conversation in a note pad, so, if you agree, we would like to record this sessions in a tape recorder. We would also like to take some notes from this session. This session will last about an hour. We would like to return to ask additional questions sometime in the future.

**Risks & Benefits**

This study presents minimal risk to you. Although we will try to protect your identity there is some possibility that confidentiality could be compromised.You will not directlybenefit for participating in this study, but your participation will help us to gain knowledge on exposures that people in this area have to lead. If we identify a source of lead, we will inform the government and this may affect which chemicals are allowed to be sold.

**Privacy and confidentiality**

All data and specimens collected will be kept confidential as allowed by the law of this country. None of these researchers will be able to identify you. Other persons who may have access to you're your responses and the audio recodings include research groups that oversee the safety of the study including institutes such as the University of California Berkeley and the Bill and Melinda Gates Foundation, USA. Because of the potential need to release information to these parties, we cannot guarantee absolute confidentiality. However, we will ensure every effort that we can to preserve your confidentiality. There is no expiration date to this permission. This information will be coded and it is unlikely that anyone will be able to trace it to you protecting your privacy.

**Future use of information**

The information collected from this study may be shared with other researchers if needed, but we will strictly maintain your confidentiality and privacy.

**Right not to participate and withdraw**

Taking part in the study is completely voluntary. If you wish, you can withdraw your permission to participate at any time during the study and we will erase/destroy your soil and rice samples or any data recorded from you. You have the right to refuse participation in this study, which will not affect your family's treatment at the Cholera Hospital in the future.

**Principle of compensation**

You need not pay us to take part in this study, and similarly we will not pay you money for attending in the study.

**Persons to contact**:

If you have any question, you can ask me any time. If you have additional questions about the survey, you may contact:

Dr. Mahbubur Rahman, Centre of Communicable Diseases, ICDDR,B, Mohakhali, Dhaka 1212. Phone: 8860523-32 # 120

If you have questions about your rights as a participant of a research study, or if you think some harm has been done to you because of the survey, you may contact or meet him personally at following address:

M. A. Salam Khan, IRB Secretariat, phone: 9886498 or PABX 8860523-32 ext. 3206
If you agree to our proposal of enrolling your household in our study, please indicate that by putting your signature or your left thumb impression at the specified space below
Thank you for your cooperation

Consent of the mother for sample collection: YES_______ NO ______

_______________________________________ ____________________
Signature or left thumb impression of Participant Date

_______________________________________ ____________________
Signature of the PI or his/her representative Date

---------------------------------------------------------------------------------------------

**Protocol Title:** Measuring the benefits of sanitation, water quality, handwashing and nutrition interventions for improving health and development in rural Bangladesh

**Protocol Status:** APPROVED

**Date Submitted:** 10/26/2017

**Approval Period:** **10/30/2017-11/03/2018**

**Important Note:** **This Print View may not reflect all comments and contingencies for approval. Please check the comments section of the online protocol.
Questions that appear to not have been answered may not have been required for this submission. Please see the system application for more details.**

*** * * Attached Document * * ***

| Document Name | Created Date |
|---|---|
| 2011-09-3652_Colford_2o-Consent-Agrochemical-v17-Bengali.pdf | 10/26/2017 |

Q.T/ S.G/ Cl. ID/ Co. ID/ B. ID/ HH ID

Household ID: □/□/ □□/ □□/□□/□□

icddr,b KNOWLEDGE FOR GLOBAL LIFESAVING SOLUTIONS

### Appendix 2o: Bengali consent for agrochemical in-depth interview

**এপেন্ডিক্স ১ও: কৃষিক্ষেত্রে ব্যবহৃত রাসায়নিক বিষয়ে (এগ্রোকেমিক্যাল) নিবিড় সাক্ষাৎকারের জন্য সম্মতিপত্র**

গবেষণার শিরোনাম: বাংলাদেশের গ্রামীণ এলাকায় হাত ধোয়া, পানি বিশুদ্ধকরণ, স্যানিটেশন ও পুষ্টিকর সম্পূরক খাবার প্রদান এবং শিশুর স্বাস্থ্য ও বৃদ্ধির উপর তার প্রভাব।

**প্রধান গবেষকঃ** ড: লিয়েন ইউনিকম্ব

**গবেষণার উদ্দেশ্য**

আসসালামু আলাইকুম/নমস্কার। আমার নাম----------- এবং আমি ঢাকায় আইসিডিডিআর,বি তে (কলেরা হাসপাতাল) কাজ করি। এই এলাকার মানুষজনের সীসার সংস্পর্শে আসার বিষয়টি নিয়ে আমরা গবেষণা করতে ইচ্ছুক। সীসার সংস্পর্শতা মাঝে মাঝে অসুস্থ্যতার কারণ হতে পারে, তাই আমরা এই বিষয়টি ভালভাবে বুঝতে চাই।

**কেন আপনাকে আমরা এই গবেষণায় অংশগ্রহণের জন্য আমন্ত্রন জানাচ্ছি?**

কৃষিকাজে ব্যবহৃত রাসায়নিকগুলোর মধ্যে কখনও কখনও সীসা থাকে। আমরা জানতে চাই, এই এলাকায় কৃষকেরা কী ধরণের রাসায়নিক ক্রয় করে থাকে।

**এই গবেষণায় অংশগ্রহণকারীর কাছে কি প্রত্যাশা করা হচ্ছে?**

আপনি যদি এই গবেষণায় অংশগ্রহণের সিদ্ধান্ড় নেন তবে আমরা আপনাকে কিছু ধারাবাহিক প্রশ্ন করব। আমরা আপনার দোকান দেখতে চাইব। আপনার যে জমিতে ধান চাষ করেন সেখান থেকে সামান্য মাটি এবং আপনার জমিতে উৎপাদিত কিছু ধান/চাল আমরা নমুনা হিসাবে সংগ্রহ করব । আমরা পরবর্তীতে আবারও কথা বলার জন্য আসতে পারি। যেহেতু আপনার সাথে কথোপকথনের সম্পূর্ণ বিবরণটি খাতায় লেখা কষ্টকর, তাই যদি আপনি রাজী থাকেন, আমরা আপনার সাথে কথোপকথনের এই বিষয়টি টেপ রেকর্ড করতে চাই। আমরা একই সাথে কিছু বিষয় খাতায়ও লিখে রাখব। এর জন্য এক ঘন্টা সময় লাগবে। আরও কিছু প্রশ্ন করার জন্য আমরা ভবিষ্যতে কোন এক সময় আবারও আসতে পারি।

**ঝুঁকি এবং সুবিধা**

এই গবেষণায় আপনার ঝুঁকির সম্ভাবনা নুন্যতম। আমরা আপনার পরিচয়সংক্রান্ত তথ্যসমূহ গোপন রাখার জন্য সম্ভাব্য সবধরনের চেষ্টা করব। তা সত্ত্বেও ভুলবশতঃ অনাকাঙ্খিত কোন ঘটনা ঘটলে সেজন্য মার্জনা প্রার্থনা করছি।এই গবেষণায় অংশগ্রহনের জন্য আপনি সরাসরিভাবে কোন ধরনের লাভবান হবেন না, তবে আপনার অংশগ্রহন সীসার সংস্পর্শতার কারনে এই এলাকার জনগনের কী ধরনের অসুস্থতার সম্মুখীন হয় সেসম্পর্কে আমাদেরকে জানতে সহায়তা করবে। যদি আমরা সীসার কোন উৎস চিহ্নিত করতে পারি তবে আমরা সরকারকে অবগত করব এবং এর জন্য কিছু কিছু রাসায়নিক বিক্রির উপর বাধ্যবাধকতা আসতে পারে।

**গোপনীয়তা ও বিশ্বস্ততা**

আপনার কাছ থেকে প্রাপ্ত সকল তথ্য ও নমুনা এই দেশের আইন অনুযায়ী গোপন রাখা হবে। কোন গবেষকই আপনাকে চিহ্নিত করতে পারবে না। গবেষণা দল যারা এই গবেষণার নিরাপত্তার তত্ত্বাবধানে রয়েছেন এবং আমেরিকার ক্যালিফোর্নিয়া বার্কলে বিশ্ববিদ্যালয় এবং বিল ও মেলিন্ডা গেটস ফাউন্ডেশন আপনার দেওয়া তথ্য এবং এই অডিও রেকর্ডিং ব্যবহার করতে পারবেন। যেহেতু এই প্রতিষ্ঠানসমূহ ও ব্যক্তিবর্গকে যে কোন সময় তথ্য দিতে হতে পারে তাই পুরোপুরি গোপনীয়তা রক্ষার বিষয়টি আমরা নিশ্চিত করতে পারছি না। তবুও সর্বোচ্চ গোপনীয়তা রক্ষার জন্য আমরা সর্বাত্মক চেষ্টা করব। এদের এই তথ্য ব্যবহারে অনুমতির জন্য কোন নির্ধারিত সময়সীমা নেই। আপনার গোপনীয়তা রক্ষার্থে এই তথ্যগুলোকে কোড করা হবে যার ফলে কারো পক্ষে আপনাকে সনাক্ত করা প্রায় অসম্ভব।

**ভবিষ্যতে তথ্যের ব্যাবহার**

এই গবেষণা হতে প্রাপ্ত তথ্য প্রয়োজনে অন্য গবেষকদের সাথে আদান-প্রদান করতে হতে পারে। তবে আমরা খুব সতর্কতার সাথে আপনার বিশ্বস্ততা ও গোপনীয়তা রক্ষা করব।

**গবেষণায় অংশগ্রহণে অস্বীকৃতি ও নিজেকে বাদ দেওয়ার অধিকার**

এই গবেষণায় আপনার অংশগ্রহণ সম্পূর্ণরূপে স্বেচ্ছাসেবামূলক। আপনি আপনার অংশগ্রহণ প্রত্যাহার করতে পারবেন এবং আপনি যদি ইচ্ছে করেন তবে আমরা আপনার কাছ থেকে ধারণকৃত তথ্যও মুছে/নষ্ট করে ফেলব। এমনকি গবেষণায় অংশগ্রহনে সম্মতি দেয়ার পরবর্তীতেও আপনি ইচ্ছা করলে গবেষণা চলাকালীন যে কোন সময়ে আপনার অনুমতি প্রত্যাখ্যান করতে পারেন।এই

গবেষণা থেকে নিজেকে প্রত্যাহার করার অধিকার আপনার আছে, এজন্য কলেরা হাসপাতালে আপনার পরিবারের চিকিৎসাক্ষেত্রে কোন সমস্যা হবে না।

**ক্ষতিপূরণ নীতি**

এই গবেষণায় অংশগ্রহণের জন্য আপনাকে কোন টাকাপয়সা দিতে হবে না একই ভাবে আমাদের পক্ষ থেকেও আপনাকে কোন টাকা পয়সা দেয়া হবে না।

**যোগাযোগের ব্যক্তিবর্গ**

আপনার যদি কোন প্রশ্ন থাকে, তবে তা যে কোন সময়ে আমাকে জিজ্ঞাসা করতে পারেন। এই গবেষণা/ জরিপ বিষয়ে অপনার যদি অতিরিক্ত কোন প্রশ্ন থাকে তবে তা জানার জন্য যোগাযোগ করতে পারেন। যোগাযোগের ঠিকানাঃ

ডাঃ মাহাবুবুর রহমান, সেন্টার ফর কমিউনিকেবল ডিজিজেস, আইসিডিডিআর,বি, মহাখালী, ঢাকা-১২১২। ফোন: ৮৮৬০৫২৩-৩২ # ১২৩।

একজন গবেষণা অংশগ্রহণকারী হিসেবে আপনার অধিকার সম্পর্কে যদি জানতে চান অথবা যদি আপনি মনে করেন যে আপনার কাছ থেকে তথ্য সংগ্রহের কারণে আপনার কোন ক্ষতি হয়েছে তাবে আপনি নিম্নোক্ত ঠিকানায় যোগাযোগ অথবা সরাসরি দেখা করতে পারেন।যোগাযোগের ঠিকানাঃ

এম, এ সালাম খান, আই আর বি সেক্রেটারিয়েট, ফোনঃ ৯৮৮৬৪৯৮ অথবা PABX, ৮৮৬০৫২৩-৩২, এক্স. ৩২০৬।

আপনি যদি আমাদের প্রস্তাব অনুযায়ী আপনার খানাকে এই গবেষণায় যুক্ত করতে সম্মত হন তবে নীচে উল্লেখিত স্থানে আপনার স্বাক্ষর অথবা বাম হাতের বৃদ্ধাঙ্গুলীর ছাপ দিন। আপনার সহযোগিতার জন্য ধন্যবাদ।

নমুনা সংগ্রহের জন্য মা'র সম্মতি: হ্যাঁ ---------না--------

---------------------------------------- ----------------------

অংশগ্রহণকারীর স্বাক্ষর/ বাম হাতের বৃদ্ধাঙ্গুলীর ছাপ তারিখ

--------------------------------------- ----------------------

প্রধান গবেষক অথবা তার প্রতিনিধির স্বাক্ষর তারিখ

--------------------------------------------------------------------------------
---

**Protocol Title:** Measuring the benefits of sanitation, water quality, handwashing and nutrition interventions for improving health and development in rural Bangladesh

**Protocol Status:** APPROVED

**Date Submitted:** 10/26/2017

**Approval Period:** **10/30/2017-11/03/2018**

**Important Note:** **This Print View may not reflect all comments and contingencies for approval. Please check the comments section of the online protocol.
Questions that appear to not have been answered may not have been required for this submission. Please see the system application for more details.**

*** * * Attached Document * * ***

| Document Name | Created Date |
|---|---|
| 2011-09-3652_Colford_1p-Consent-Lead case control-v17-English.pdf | 10/26/2017 |

**Appendix 1p: English consent form for lead case-control study**

Protocol Title: Effect of hand washing, water treatment, sanitation and nutritional supplement interventions on child health and development in rural Bangladesh

Principal Investigator's name: Dr. Leanne Unicomb

**Purpose of the research**

Hello (Assalamualaikum/Nomoshkar). My name is ________ and I work with the ICDDR,B (Cholera Hospital) in Dhaka. We are interested in conducting research on exposures that people in this area have to lead. Sometimes lead exposure can cause illness so we want to understand them better.

**Why are we inviting you to participate in the study?**

Some chemicals and metals can affect health and child development. We are interested in exploring how you and your family might be exposed to chemicals or metals in the environment.

**What is expected from the participants of the research study?**

If your household decides to enroll in we will ask you a series of questions. We will ask also questions of the person who does most of the farming. We will want to look at your home and the fields your family works in. We would like to collect a sample of soil from the field and a sample of uncooked rice that you cooked in the field. This session will last about an hour. We would like to return to ask additional questions sometime in the future. Your decision to participate in this part of the study is optional and will not affect your household participation in WASH Benefits.

**Risks & Benefits**

This study presents minimal risk to you. Although we will try to protect your identity there is some possibility that confidentiality could be compromised.You will not directly benefit for participating in this study, but your participation will help us to gain knowledge on lead exposures and associated risk factors for children and residents in this area. If our study identifies a dangerous exposure in your environment, we will make recommendations on how to limit your family's exposure.

**Privacy and confidentiality**

All data and specimens collected will be kept confidential as allowed by the law of this country. The soil and rice samples and the test results will be coded without your name or personal information, and stored separately for analysis by the researchers. None of these researchers will be able to identify you or your family as the sample donors. Other persons who may have access to your test results include research groups that oversee the safety of the study including institutes such as the University of California Berkeley and the Bill and Melinda Gates

Foundation, USA. Because of the potential need to release information to these parties, we cannot guarantee absolute confidentiality. However, we will ensure every effort that we can to preserve your confidentiality. There is no expiration date to this permission. This information will be coded and it is unlikely that anyone will be able to trace it to you and your child, protecting your privacy.

**Future use of information**

The information collected from this study may be shared with other researchers if needed, but we will strictly maintain your confidentiality and privacy.

**Right not to participate and withdraw**

Taking part in the study is completely voluntary. If, at any point during the study, you decide you do not wish to participate, we will withdraw your permission and we will destroy your soil, rice samples and data recorded from you.You have the right to refuse participation in this study, which will not affect your family's treatment at the Cholera Hospital in the future.

**Principle of compensation**

You need not pay us to take part in this study, and similarly we will not pay you money for attending in the study.

**Persons to contact**:

If you have any question, you can ask me any time. If you have additional questions about the survey, you may contact:

Dr. Mahbubur Rahman, Centre of Communicable Diseases, ICDDR,B, Mohakhali, Dhaka 1212. Phone: 8860523-32 # 120

If you have questions about your rights as a participant of a research study, or if you think some harm has been done to you because of the survey, you may contact or meet him personally at following address:

M. A. Salam Khan, IRB Secretariat, phone: 9886498 or PABX 8860523-32 ext. 3206
If you agree to our proposal of enrolling your household in our study, please indicate that by putting your signature or your left thumb impression at the specified space below
Thank you for your cooperation

Consent of the mother for sample collection: YES_______ NO ______

_______________________________________ _____________________

Signature or left thumb impression of Participant Date

Signature of the PI or his/her representative    Date

PROTOCOL
Biomedical
Non-Exempt
Berkeley

Protocol # 2011-09-3652
Date Printed: 07/31/2018

-----------------------------------------------------------------------------------

| | |
|---|---|
| **Protocol Title:** | Measuring the benefits of sanitation, water quality, handwashing and nutrition interventions for improving health and development in rural Bangladesh |
| **Protocol Status:** | APPROVED |
| **Date Submitted:** | 10/26/2017 |
| **Approval Period:** | **10/30/2017-11/03/2018** |
| **Important Note:** | **This Print View may not reflect all comments and contingencies for approval. Please check the comments section of the online protocol. Questions that appear to not have been answered may not have been required for this submission. Please see the system application for more details.** |

*** * * Attached Document * * ***

| Document Name | Created Date |
|---|---|
| 2011-09-3652_Colford_2p-Consent-Lead case control-v17-Bengali.pdf | 10/26/2017 |

#### Appendix 2p: Bengali consent for lead case-control study

গবেষণার শিরোনাম: বাংলাদেশের গ্রামীণ এলাকায় হাত ধোয়া, পানি বিশুদ্ধকরণ, স্যানিটেশন ও পুষ্টিকর সম্পূরক খাবার প্রদান এবং শিশুর স্বাস্থ্য ও বৃদ্ধির উপর তার প্রভাব।

**প্রধান গবেষকঃ** ড: লিয়েন ইউনিকম্ব

**গবেষণার উদ্দেশ্য**

আসসালামু আলাইকুম/নমস্কার। আমার নাম----------- এবং আমি ঢাকায় আইসিডিডিআর,বি তে (কলেরা হাসপাতাল) কাজ করি। এই এলাকার মানুষজনের সীসার সংস্পর্শে আসার বিষয়টি নিয়ে আমরা গবেষণা করতে ইচ্ছুক। সীসার সংস্পর্শতা মাঝে মাঝে অসুস্থ্যতার কারণ হতে পারে, তাই আমরা এই বিষয়টি ভালভাবে বুঝতে চাই।

**কেন আপনাকে আমরা এই গবেষণায় অংশগ্রহণের জন্য আমন্ত্রন জানাচ্ছি?**

কৃষিকাজে ব্যবহৃত রাসায়নিকগুলোর মধ্যে কখনও কখনও সীসা থাকে। আমরা জানতে চাই, এই এলাকায় কৃষকেরা কী ধরণের রাসায়নিক ক্রয় করে থাকে।

**এই গবেষণায় অংশগ্রহণকারীর কাছে কি প্রত্যাশা করা হচ্ছে?**

আপনি যদি এই গবেষণায় অংশগ্রহণের সিদ্ধান্ত নেন তবে আমরা আপনাকে কিছু ধারাবাহিক প্রশ্ন করব। আমরা আপনার দোকান দেখতে চাইব। আমরা পরবর্তীতে আবারও কথা বলার জন্য আসতে পারি। যেহেতু আপনার সাথে কথোপকথনের সম্পূর্ণ বিবরণটি খাতায় লেখা কষ্টকর, তাই যদি আপনি রাজী থাকেন, আমরা আপনার সাথে কথোপকথনের এই বিষয়টি টেপ রেকর্ড করতে চাই। আমরা একই সাথে কিছু বিষয় খাতায়ও লিখে রাখব। এর জন্য এক ঘনটা সময় লাগবে। আরও কিছু প্রশ্ন করার জন্য আমরা ভবিষ্যতে কোন এক সময় আবারও আসতে পারি।

**ঝুঁকি এবং সুবিধা**

এই গবেষণায় আপনার ঝুঁকির সম্ভাবনা নুন্যতম। আমরা আপনার পরিচয়সংক্রান্ত তথ্যসমূহ গোপন রাখার জন্য সম্ভাব্য সবধরনের চেষ্টা করব। তা সত্ত্বেও ভুলবশতঃ অনাকাঙ্খিত কোনঢ় ঘটনা ঘটলে সেজন্য মার্জনা প্রার্থনা করছি। এই গবেষণায় অংশগ্রহন করলে আপনি সরাসরিভাবে কোন ধরনের লাভবান হবেন না, তবে আপনার অংশগ্রহন সীসার সংস্পর্শতার কারনে এই এলাকার জনগনের কী ধরনের অসুস্থতার সম্মুখীন হয় সেসম্পর্কে আমাদেরকে জানতে সহায়তা করবে। যদি আমরা সীসার কোন উৎস চিহ্নিত করতে পারি তবে আমরা সরকারকে অবগত করব এবং এর জন্য কিছু কিছু রাসায়নিক বিক্রির উপর বাধ্যবাধকতা আসতে পারে।

**গোপনীয়তা ও বিশ্বস্ততা**

আপনার কাছ থেকে প্রাপ্ত সকল তথ্য ও নমুনা এই দেশের আইন অনুযায়ী গোপন রাখা হবে। কোন গবেষকই আপনাকে চিহ্নিত করতে পারবে না। গবেষণা দল যারা এই গবেষণার নিরাপত্তার তত্ত্বাবধানে রয়েছেন এবং আমেরিকার ক্যালিফোর্নিয়া বার্কলে বিশ্ববিদ্যালয় এবং বিল ও মেলিন্ডা গেটস ফাউন্ডেশন আপনার দেওয়া তথ্য এবং এই অডিও রেকর্ডিং ব্যবহার করতে পারবেন। যেহেতু এই প্রতিষ্ঠানসমূহ ও ব্যক্তিবর্গকে যে কোন সময় তথ্য দিতে হতে পারে তাই পুরোপুরি গোপনীয়তা রক্ষার বিষয়টি আমরা নিশ্চিত করতে পারছি না। তবুও সর্বোচ্চ গোপনীয়তা রক্ষার জন্য আমরা সর্বাত্মক চেষ্টা করব। এদের এই তথ্য ব্যবহারে অনুমতির জন্য কোন নির্ধারিত সময়সীমা নেই। আপনার গোপনীয়তা রক্ষার্থে এই তথ্যগুলোকে কোড করা হবে যার ফলে কারো পক্ষে আপনাকে সনাক্ত করা প্রায় অসম্ভব।

**ভবিষ্যতে তথ্যের ব্যাবহার**

এই গবেষণা হতে প্রাপ্ত তথ্য প্রয়োজনে অন্য গবেষকদের সাথে আদান-প্রদান করতে হতে পারে। তবে আমরা খুব সতর্কতার সাথে আপনার বিশ্বস্ততা ও গোপনীয়তা রক্ষা করব।

**গবেষণায় অংশগ্রহণে অস্বীকৃতি ও নিজেকে বাদ দেওয়ার অধিকার**

এই গবেষণায় আপনার অংশগ্রহণ সম্পূর্ণরূপে স্বেচ্ছাসেবামূলক। এই গবেষণা চলাকালীন যে কোন সময়ে আপনি চাইলে অংশগ্রহণ নাও করতে পারেন,আমরা আপনার অনুমতি প্রত্যাহার করব এবং আপনার কাছ থেকে নেওয়া মাটি, চালের নমুণা এবং তথ্য নষ্ট করে ফেলব।এমনকি গবেষণায় অংশগ্রহনে সম্মতি দেয়ার পরবর্তীতেও আপনি ইচ্ছা করলে গবেষণা চলাকালীন যে কোন সময়ে

আপনার অনুমতি প্রত্যাখ্যান করতে পারেন। এই গবেষণা থেকে নিজেকে প্রত্যাহার করার অধিকার আপনার আছে, এজন্য কলেরা হাসপাতালে আপনার পরিবারের চিকিৎসাক্ষেত্রে কোন সমস্যা হবে না।

**ক্ষতিপূরণ নীতি**

এই গবেষণায় অংশগ্রহণের জন্য আপনাকে কোন টাকাপয়সা দিতে হবে না একই ভাবে আমাদের পক্ষ থেকেও আপনাকে কোন টাকা পয়সা দেয়া হবে না।

**যোগাযোগের ব্যক্তিবর্গ**

আপনার যদি কোন প্রশ্ন থাকে, তবে তা যে কোন সময়ে আমাকে জিজ্ঞাসা করতে পারেন। এই গবেষণা/ জরিপ বিষয়ে অপনার যদি অতিরিক্ত কোন প্রশ্ন থাকে তবে তা জানার জন্য যোগাযোগ করতে পারেন। যোগাযোগের ঠিকানাঃ

ডাঃ মাহাবুবুর রহমান, সেন টার ফর কমিউনিকেবল ডিজিজেস, আইসিডিডিআর,বি, মহাখালী, ঢাকা-১২১২। ফোন: ৮৮৬০৫২৩-৩২ # ১২৩।

একজন গবেষণা অংশগ্রহণকারী হিসেবে আপনার অধিকার সম্পর্কে যদি জানতে চান অথবা যদি আপনি মনে করেন যে আপনার কাছ থেকে তথ্য সংগ্রহের কারণে আপনার কোন ক্ষতি হয়েছে তাবে আপনি নিম্নোক্ত ঠিকানায় যোগাযোগ অথবা সরাসরি দেখা করতে পারেন।যোগাযোগের ঠিকানাঃ

এম, এ সালাম খান, আই আর বি সেক্রেটারিয়েট, ফোনঃ ৯৮৮৬৪৯৮ অথবা PABX, ৮৮৬০৫২৩-৩২, এক্স. ৩২০৬।

আপনি যদি আমাদের প্রস্তাব অনুযায়ী আপনার খানাকে এই গবেষণায় যুক্ত করতে সম্মত হন তবে নীচে উল্লেখিত স্থানে আপনার স্বাক্ষর অথবা বাম হাতের বৃদ্ধাঙ্গুলীর ছাপ দিন। আপনার সহযোগিতার জন্য ধন্যবাদ।

নমুনা সংগ্রহের জন্য মা'র সম্মতি: হ্যাঁ ---------না--------

---------------------------------------- ----------------------

অংশগ্রহণকারীর স্বাক্ষর/ বাম হাতের বৃদ্ধাঙ্গুলীর ছাপ তারিখ

---------------------------------------- ----------------------

প্রধান গবেষক অথবা তার প্রতিনিধির স্বাক্ষর তারিখ

PROTOCOL
Biomedical
Non-Exempt
Berkeley

Protocol # 2011-09-3652
Date Printed: 07/31/2018

-------------------------------------------------------------------------------------

| | |
|---|---|
| **Protocol Title:** | Measuring the benefits of sanitation, water quality, handwashing and nutrition interventions for improving health and development in rural Bangladesh |
| **Protocol Status:** | APPROVED |
| **Date Submitted:** | 10/26/2017 |
| **Approval Period:** | **10/30/2017-11/03/2018** |
| **Important Note:** | **This Print View may not reflect all comments and contingencies for approval. Please check the comments section of the online protocol. Questions that appear to not have been answered may not have been required for this submission. Please see the system application for more details.** |

*** * * Attached Document * * ***

| Document Name | Created Date |
|---|---|
| 2011-09-3652_Colford_1r-Consent-Structured observation with video-v17-English.pdf | 10/26/2017 |

**Appendix 1r: English consent form for structured observations with video recording**

Protocol Title: Effect of hand washing, water treatment, sanitation and nutritional supplement interventions on child health and development in rural Bangladesh

Principal Investigator's name: Dr. Stephen P. Luby

#### Purpose of the research

Hello (Assalamualaikum/Namaste). My name is ________ and I work with the ICDDR,B (Cholera Hospital) in Dhaka. We are doing research on diarrheal diseases in children. Through this research we want to learn how certain conditions in sanitation, water quality and hygiene practices impact health.

**Why are we inviting you to participate in the study?**

We are seeking households that are already taking part in other activities from this study. We would like to tell you more detail about this study and hope you and your child will volunteer to participate. You are living in a rural social and natural environment from where we would like to learn something more. Additionally, there was pregnant women and a young child in your household and we are interested to talk about their health.

**What is expected from the participants of the research study?**

If you agree to take part, I will find a place to sit in your home or courtyard so that I will not be in your way. I will observe activities in your home and the daily activities of your child and take some notes. I may look around the courtyard and may ask you some questions in the end. I will also take a video of you and your child during your daily activities. I will be here for a total of about five hours.

**Risks & Benefits**

There are no major risks involved in this study. There is no monetary compensation for taking part in this study.

**Privacy, anonymity and confidentiality**

We will do everything we can keep what you tell us confidential as allowed by the law of this country. We will not use your real name when we write out the data. All the information we collect will be kept locked. We will use the information only for the purpose of the study. We will not use your name when we share and publish the results of this study. We expect the steps we take will keep all of your information confidential, but it is possible that because of mistakes or unforeseen events, it could become compromised. We may want to use some of the photographs, audio, or video recordings of you in public presentations related to the research. There is a Media Records Release Form attached that outlines several possible uses and asks for your specific consent to use these items in each way. If you agree to allow these items to be used after this research study is over, please read, initial, and sign the Media Records Release Form in addition to this consent form. We will not use any photographs, recordings, or other identifiable information about you in any future presentation without your consent. The recorded footage will be stored securely until data analysis is complete, and will then be erased.

**Future use of information**

If it is necessary, we may share the information we collected from this study with other researchers. If this is done, we will not use your real name and we will maintain your confidentiality.

**Right not to participate and withdraw**

Taking part in the study is completely voluntary. You may choose not to answer any or all of the questions that will ask. You can drop out of this study at any time, even in the middle of an interview. You have the right to refuse to take part in this study, which will not affect your family's treatment at the Cholera Hospital in the future.

**Principle of compensation**
You do not need to pay us to take part in this study, and we will not pay you money for taking part in the study.

**Persons to contact:**

If you have any questions, you can ask me any time. If you have additional questions about the survey, you may contact: Muhammad Faruqe Hussain: phone: 01711437326 from the Water Sanitation and Hygiene Research Group, Centre for Communicable Diseases (CCD), icddr,b Mohakhali, Dhaka 1212.

If you have questions about your rights as a participant of a research study, or if you think some harm has been done to you because of this study, you may contact or meet him personally at following address:

M. A. Salam Khan, IRB Secretariat, phone: 9886498 or PABX 8860523-32 ext. 3206

If you agree to our proposal of enrolling your household in our study, please indicate that by putting your signature or your left thumb impression at the specified space below

Thank you for your cooperation

_________________________________________ _____________________

Signature or left thumb impression of Participant Date

_________________________________________ _____________________

Signature or left thumb impression of the witness Date

_________________________________________ _____________________

Signature of the PI or his/her representative Date

**Separate consent to photography and video taping**

During the study we would also like to take photographs and videos of you, your child, your family members and activities including your natural environment. These photographs and videos may be used, if necessary, in presentations or reports at scientific meetings or to the general public. Even if you do not permit us to take photographs or video tape you might be considered to take part in this study. We will not use any photographs, recordings, or other identifiable information about you in any future presentation without your consent

Please sign next to "YES" or "NO" if you agree with possibly having photos taken and shared. In any use of these records, your name will not be identified.

1. The records can be studied by the research team for use in the research project.

   Photo __________ Audio __________ Video __________

2. The records can be used for scientific publications.

   Photo __________ Audio __________ Video __________

3. The records can be shown at meetings of scientists interested in the study of ___________.

   Photo __________ Audio __________ Video __________

4. The records can be shown in classrooms to students.

   Photo __________ Audio __________ Video __________

5. The records can be shown in public presentations to non-scientific groups.

   Photo __________ Audio __________ Video __________

PROTOCOL
Biomedical
Non-Exempt
Berkeley

Protocol # 2011-09-3652
Date Printed: 07/31/2018

--------------------------------------------------------------------------------

| | |
|---|---|
| **Protocol Title:** | Measuring the benefits of sanitation, water quality, handwashing and nutrition interventions for improving health and development in rural Bangladesh |
| **Protocol Status:** | APPROVED |
| **Date Submitted:** | 10/26/2017 |
| **Approval Period:** | **10/30/2017-11/03/2018** |
| **Important Note:** | **This Print View may not reflect all comments and contingencies for approval. Please check the comments section of the online protocol. Questions that appear to not have been answered may not have been required for this submission. Please see the system application for more details.** |

*** * * Attached Document * * ***

| Document Name | Created Date |
|---|---|
| 2011-09-3652_Colford_2r-Consent-Structured observation with video-v17-Bengali.pdf | 10/26/2017 |

#### Appendix 2r: Bangla consent form for structured observations with video recording

**স্বেচ্ছা সম্মতিপত্র**

আন্তর্জাতিক উদরাময় গবেষণা কেন্দ্র, বাংলাদেশ (আই সি ডি ডি আর, বি )

মহাখালী, ঢাকা।

**গবেষণার শিরোনামঃ**

বাংলাদেশের গ্রামীণ এলাকায় হাত ধোয়া, পানি বিশুদ্ধকরণ, স্যানিটেশন ও পুষ্টিকর সম্পূরক খাবার প্রদান এবং শিশুর স্বাস্থ্য ও বৃদ্ধির উপর তার প্রভাব।

**প্রধান গবেষক:**

ডঃ স্টিফেন পি. লুবি

**গবেষণার উদ্দেশ্যঃ**

আস্সালামুআলাইকুম/নমস্কার। আমার নাম --------------------------------। আমি ঢাকার মহাখালীতে অবস্থিত আইসিডিডিআর,বি (কলেরা হাসপাতাল) এ চাকরী করি। শিশু-স্বাস্থ্য ও ডায়রিয়া বিষয়ক আমরা একটি গবেষণা কাজ করছি। এই গবেষণার মাধ্যমে আমরা জানতে চাই নির্দিষ্ট কিছু স্বাস্থ্যসম্মত বিষয়, পানির গুনাগুণ এবং স্বাস্থ্যসম্মত অভ্যাসসমূহ স্বাস্থ্যের উপর কিভাবে প্রভাব ফেলে ।

**আমরা আপনাকে কেন এই গবেষণায় অংশগ্রহণের জন্য আমন্ত্রণ জানাচ্ছি?**

আমরা কিছু গৃহস্থালী/খানা দেখতে চাচ্ছি যারা এই গবেষণার অন্যান্য কার্যক্রমে ইতিমধ্যেই অংশগ্রহণ করছে। আমরা এই গবেষণা সম্পর্কে আপনাদেরকে আরও বিস্তারিত কিছু জানাতে চাই এবং আশা করি যে আপনি এবং আপনার শিশু এই গবেষণায় স্বেচ্ছায় অংশগ্রহণ করবেন। আপনারা একটি গ্রামীণ সামাজিক ও প্রাকৃতিক পরিবেশে বাস করছেন যেখান থেকে আমরা আরও কিছু শিখতে চাই। তাছাড়া এই খানায় একজন গর্ভবতী মহিলা ছিলেন এবং ছোট শিশুরাও রয়েছে যাদের স্বাস্থ্য নিয়ে আমরা আলাপ করবো।

**এই গবেষণায় অংশগ্রহণকারীদের কাছ থেকে প্রত্যাশা কি?**

যদি আপনি এই গবেষণায় অংশগ্রহণে রাজী থাকেন তাহলে আমি আপনার ঘর বা উঠানের কাছাকাছি এমন একটি স্থানে বসবো কিন্তু আপনার চলার পথের বাঁধা হবো না। আমি আপনার বাড়ী এবং আপনার শিশুর প্রতিদিনকার কর্মকান্ড সমূহ পর্যবেক্ষণ করবো ও তা লিখে রাখবো। এছাড়াও আমি আপনার উঠানের চারিদিকে ঘুরে দেখতে পারি এবং পরিশেষে কিছু প্রশ্নও জিজ্ঞাসা করতে পারি। আমি আপনার বাড়ীর প্রাকৃতিক পরিবেশ, আপনারা এবং আপনার শিশুর দৈনন্দিন কর্মকান্ড চলাকালীন কিছু ছবি ও ভিডিও চিত্রও নেবো। আমি আপনার বাড়ীতে মোট ৫ ঘন্টার মত থাকবো।

**ঝুঁকি এবং সুবিধাসমূহ**

এই গবেষণায় অংশগ্রহনে আপনার কোন ঝুঁকি নেই। এই গবেষণায় অংশগ্রহনের জন্য আপনি কোন ধরণের আর্থিক সহায়তাও পাবেন না। কিন্তু আপনার শিশুর অংশগ্রহণ, শিশুদের ডায়রিয়া রোগ সম্পর্কে জ্ঞানলাভে আমাদেরকে সহায়তা করবে।

## একান্ৰতা, গোপনীয়তা, এবং বিশ্বাসযোগ্যতা

এই দেশের আইন অনুযায়ী আপনি আমাদেরকে যা বলবেন তা গোপনীয় রাখার জন্য সব কিছুই করবো। আমরা আমাদের গবেষণার কোন লেখায়/ফলাফলে আপনার প্রকৃত নাম ব্যবহার করব না। আমাদের সংগৃহীত সকল তথ্য তালাবদ্ধ থাকবে। আমরা কেবলমাত্র আমাদের গবেষণার কাজে এসব তথ্য ব্যবহার করব। আমরা যখন এই গবেষণার ফলাফল প্রকাশ করবো এবং অন্যকেরকে জানাবো তখন আপনার নাম ব্যবহার করবো না। আমরা আশা করি আমরা যে সকল পদৰেপ গ্রহণ করবো তা আপনার প্রদত্ত তথ্যকে গোপন রাখতে সাহায্য করবে। আমরা এই গবেষণার ফলাফল বিনিময় ও প্রকাশনার ৰেত্রে আপনার নাম ব্যবহার করব না। আমরা আপনার দেওয়া তথ্যসমূহ গোপন রাখার জন্য সম্ভাব্য সবধরনের চেষ্টা করব। আমরা এই গবেষণা সম্পর্কিত আপনার কিছু ছবি, অডিও এবং ভিডিও জনসাধারণের সামনে উপস্থাপন করতে চাই। এখানে একটা মিডিয়া রেকর্ড রিলিজ ফরম্ সংযুক্ত আছে যেটাতে এর অনেক সম্ভাব্য ব্যবহার এবং প্রত্যেকভাবে জিনিসপত্র ব্যবহার করার জন্য আপনার সুনির্দিষ্ট অনুমতি প্রার্থণা করা হবে। গবেষণা শেষ হওয়ার পরে এই জিনিসপত্র ব্যবহার করার অনুমতি দিতে সম্মত হলে দয়া করে পড়ুন এবং মিডিয়া রেকর্ড রিলিজ ফরম্ সহ অনুমতি পত্রে সই করম্নন। আপনার অনুমতি ছাড়া আমরা আপনার কোন ছবি, ভিডিও অথবা অন্যান্য সনাক্তযোগ্য তথ্য ভবিষ্যতে ব্যবহার করবো না। তা সত্ত্বেও ভুলবশতঃ অনাকাঙ্খিত কোন ঘটনা ঘটলে সেজন্য মার্জনা প্রার্থনা করছি।

#### ভবিষ্যতে তথ্যের ব্যবহার

এই গবেষণায় সংগৃহীত তথ্য প্রয়োজনে আমরা অন্যান্য গবেষকের সাথে বিনিময় করতে পারি। এবং কারো সাথে তথ্য বিনিময় করা হলে আমরা সেখানে আপনার প্রকৃত নাম ব্যবহার করব না এবং আপনার দেওয়া তথ্যের গোপণীয়তা ও নিজস্বতা কঠোরভাবে রক্ষা করব।

#### অংশগ্রহণ না করার এবং অংশগ্রহণ বাদ দেওয়ার অধিকার

এই গবেষণায় আপনার অংশগ্রহণ সম্পূর্ণভাবে স্বেচ্ছামূলক। আপনাকে জিজ্ঞাসা করা সকল বা যে কোন প্রশ্নের উত্তর আপনি যদি দিতে না চান তবে কোন অসুবিধা নেই অথবা আপনি আপনার পছন্দ অনুযায়ী প্রশ্নের উত্তরও দিতে পারেন। এই গবেষণা থেকে যে কোন সময় আপনার অংশগ্রহণ বাদ দিতে পারেন, এমনকি কোন সাৰাৎকার চলাকালীন যেকোন সময়কালেও। এই গবেষণায় অংশগ্রহণ না করার অধিকার আপনার আছে এবং এ জন্য ভবিষ্যতে আপনার পরিবারের সদস্যদের কলেরা হাসপাতালে স্বাস্থ্যসেবা গ্রহণের উপর কোন প্রভাব পড়বে না।

## ৰতিপূরণ

এই গবেষণায় অংশগ্রহণের জন্য আপনার কোন ধরনের অর্থনৈতিক খরচ/ ব্যয় বহন করতে হবে না এবং একইভাবে আপনাকেও আমরা কোন ধরনের অর্থ প্রদান করব না।

## যোগাযোগঃ

আপনার যদি কোন প্রশ্ন থাকে আপনি যে কোন সময় আমাকে তা জিজ্ঞাসা করতে পারেন। এছাড়াও যদি আপনার এই জরিপ বা গবেষণা সম্পর্কে আরও কোন প্রশ্ন থাকে তবে আপনি ওয়াটার, স্যানিটেশন এন্ড হাইজিন রিসার্চ গ্রম্নপ, সেন্টার ফর কমিউনিকেবল ডিজিজেজ, আইসিডিডিআ,বি, মহাখালী, ঢাকা -১২১২ এর রিসার্চ ইনভেস্টিগেটর জনাব মুহাম্মদ ফারম্নক হুসাইন এর মোবাইল নম্বর- ০১৭১১৪৩৭৩২৬ এ যোগাযোগ করতে পারেন।

যদি গবেষণায় অংশগ্রহণের অধিকার সম্পর্কে আপনার কোন প্রশ্ন থাকে অথবা এই গবেষণার কারণে আপনার কোনরূপ ৰতি সাধিত হয়েছে তবে নিম্নলিখিত ব্যক্তির সাথে আপনি যোগাযোগ করতে বা ব্যক্তিগতভাবে দেখা করতে পারেনঃ

জনাব এম এ সালাম খান, আইআরবি সেক্রেটারিয়েট, ফোন-৯৮৮৬৪৯৮ অথবা পিএবিএক্স ৮৮৬০৫২৩-৩২, এক্স. ৩২০৬.

যদি আপনি আমাদের প্রস্ৰাবের সাথে একমত পোষণ করেন (রাজী থাকেন) ও এই গবেষণায় অংশগ্রহণ করতে আগ্রহী হন তাহলে এই গবেষণায় আপনার গৃহস্থালী (খানা) কে অন্তর্ভূক্ত করা হবে। দয়া করে নিম্নের ফাকা স্থানে আপনি আপনার স্বাৰর বা বাম বৃদ্ধাঙ্গুলির ছাপ দিয়ে আপনার সম্মতি প্রদান করম্নন।

আপনার সহযোগিতার জন্য আপনাকের অনেক ধন্যবাদ।

--------------------------------- -------------

অংশগ্রহণকারীর স্বাৰর বা বাম হাতের বৃদ্ধাঙ্গুলির ছাপ তারিখ

--------------------------------- -------------

একজন স্বাৰীর স্বাৰর বা বাম হাতের বৃদ্ধাঙ্গুলির ছাপ তারিখ

--------------------------------- -------------

প্রিন্সিপাল ইনভেস্টিগেটর বা তার প্রতিনিধির স্বাৰর তারিখ

**ছবি তোলা ও ভিডিও করার জন্য আলাদা সম্মতি**

আপনি যদি অনুমতি দেন তাহলে গবেষণা চলাকালীন সময়ে আমরা আপনার ও আপনার শিশুর, পরিবারের অন্যান্য সদস্যদের, তাদের কার্যক্রম এবং আপনার খানার আশেপাশের প্রাকৃতিক পরিবেশের স্থির ও চলামান কিছু ছবি (ভিডিও) তুলবো। এই ছবি বা ভিডিও, যদি প্রয়োজন হয়, তবে কোন প্রেজেনেটশনে বা লেখালেখিতে (রিপোর্টে) ব্যবহার করা হতে পারে অথবা বৈজ্ঞানিক আলোচনা বা মিটিং ও সাধারন মানুষের কাছে উপস্থাপন করা হবে। ছবি তোলার বা ভিডিও করার অনুমতি না দিলেও আপনি এই গবেষণায় অংশগ্রহন করতে পারবেন। আপনার কোন ছবি, ধারনকৃত ভিডিওচিত্র অথবা আপনার পরিচিতির সাথে সর্ম্পকিত কোন তথ্য আপনার সম্মতি ছাড়া ভবিষ্যতে আমাদের কোন প্রেজেনেটশনে বা লেখালেখিতে (রিপোর্টে) ব্যবহার করব না।

দয়া করে সম্ভাব্য ছবি তোলা ও ধারনকৃত ভিডিও চিত্র নেয়া এবং তা গবেষণার স্বার্থে সংশিস্নষ্টদের সাথে বিনিময়ের ৰেত্রে আপনার অনুমতি আছে কিনা তা নিম্নের হ্যাঁ বা না এর উপর স্বাৰর করে অনুমতি প্রদান করম্নন। এসব ছবি ও ধারনকৃত ভিডিও চিত্র ব্যবহারের ৰেত্রে আপনার নাম প্রকাশ করা হবে না।

১। এসব ধারনকৃত ছবি ও ভিডিও চিত্র ভবিষ্যতে গবেষনার কাজে ব্যবহার করা হতে পারে।

ছবি-----------------অডিও---------------------ভিডিও------------------

২। এসব ধারনকৃত ছবি ও ভিডিও চিত্র সায়েনিটফিক প্রকাশনার  কাজে ব্যবহার করা হতে পারে।

ছবি-----------------অডিও----------------------ভিডিও------------------------------

৩। এসব ধারনকৃত ছবি ও ভিডিও চিত্র বিভিন্ন বৈজ্ঞানিক সভায় দেখানো হতে পারে।

ছবি-----------------অডিও----------------------ভিডিও--------------

৪। এসব ধারনকৃত ছবি ও ভিডিও চিত্র শ্রেনীকক্ষে ছাত্র-ছাত্রীদেরকে দেখানো হতে পারে।

ছবি-----------------অডিও----------------------ভিডিও--------------

৫। এসব ধারনকৃত ছবি ও ভিডিও চিত্র জনসাধারনের জন্য বিভিন্ন সভায় দেখানো হতে পারে।

ছবি-----------------অডিও----------------------ভিডিও--------------

PROTOCOL
Biomedical
Non-Exempt
Berkeley

Protocol # 2011-09-3652
Date Printed: 07/31/2018

---------------------------------------------------------------------------------------------

**Protocol Title:** Measuring the benefits of sanitation, water quality, handwashing and nutrition interventions for improving health and development in rural Bangladesh

**Protocol Status:** APPROVED

**Date Submitted:** 10/26/2017

**Approval Period:** **10/30/2017-11/03/2018**

**Important Note:** **This Print View may not reflect all comments and contingencies for approval. Please check the comments section of the online protocol. Questions that appear to not have been answered may not have been required for this submission. Please see the system application for more details.**

*** * * Attached Document * * ***

| Document Name | Created Date |
|---|---|
| 2011-09-3652_Colford_1s-Consent-In depth interview environmental assessment-v17-English.pdf | 10/26/2017 |

**Appendix 1s: English consent form for in depth interview in relationship to environmental assessment**

Protocol Title: Effect of hand washing, water treatment, sanitation and nutritional supplement interventions on child health and development in rural Bangladesh

Principal Investigator's name: Dr. Stephen P. Luby

#### Purpose of the research

Hello (Assalamualaikum/Namaste). My name is ________ and I work with the ICDDR,B (Cholera Hospital) in Dhaka located at Mohakhali. We are doing research on child health and diarrheal diseases. Through this research we want to learn how certain conditions in sanitation, water quality and hygiene practices impact health.

**Why are we inviting you to participate in the study?**

We are seeking households that are already taking part in other activities from this study. We would like to tell you more detail about this study and hope you and your child will volunteer to participate. You are living in a rural social and natural environment from where we would like to learn something more. Additionally, there is either a pregnant woman or a young child in your household and we are interested to talk about their health.

**What is expected from the participants of the research study?**

With your permission we would like to enroll your compound and household in this study. If you agree to take part in the study, I will request you to allocate me 60 to 90 minutes to talk about your water, sanitation and hygiene related facilities and activities in your compound and household . I will also ask about your child's health and development. With your permission we will audio and video record our conversation.

**Risks & Benefits**

This study presents minimal risk to you and your child. Although we will try to protect your identity there is some possibility that confidentiality could be compromised. There is no added benefit to you or your immediate family from participating in this interview or assessment.

**Privacy and confidentiality**

We will do everything we can to keep what you tell us confidential as allowed by the law of this country. We will not use your real name when we write out the data. All the information we collect will be kept locked. We will use the information only for the purpose of the study. We will not use your name when we share and publish the results of this study. We expect the steps we take will keep all of your information confidential, but it is possible that because of mistakes or unforeseen events, it could become compromised. The recorded data from the tape recorder will not be publicly shared, nor will participants be identifiable in any published works. The recorded data will be stored securely until data analysis is complete, and will then be erased.

**Future use of information**

If it is necessary, we may share the information we collected from this study with other researchers. If this is done, we will not use your real name and we will maintain your confidentiality.

**Right not to participate and withdraw**

Taking part in the study is completely voluntary. You may choose not to answer any or all of the questions that will ask. You can drop out of this study at any time, even in the middle of an interview. You have the right to refuse to take part in this study, which will not affect your family's treatment at the Cholera Hospital in the future.

**Principle of compensation**
You do not need to pay us to take part in this study, and we will not pay you money for taking part in the study.

**Persons to contact:**

If you have any questions, you can ask me any time. If you have additional questions about the survey, you may contact: Muhammad Faruqe Hussain: phone: 01711437326 from the Water Sanitation and Hygiene Research Group, Centre for Communicable Diseases (CCD), icddr,b , Mohakhali, Dhaka 1212.

If you have questions about your rights as a participant of a research study, or if you think some harm has been done to you because of this study, you may contact or meet him personally at following address:

M. A. Salam Khan, IRB Secretariat, phone: 9886498 or PABX 8860523-32 ext. 3206

If you agree to our proposal of enrolling your household in our study, please indicate that by putting your signature or your left thumb impression at the specified space below

Thank you for your cooperation

______________________________________ ____________________

Signature or left thumb impression of Participant Date

______________________________________ ____________________

Signature or left thumb impression of the witness Date

______________________________________ ____________________

Signature of the PI or his/her representative Date

---------------------------------------------------------------------------------
---

**Protocol Title:** Measuring the benefits of sanitation, water quality, handwashing and nutrition interventions for improving health and development in rural Bangladesh

**Protocol Status:** APPROVED

**Date Submitted:** 10/26/2017

**Approval Period:** **10/30/2017-11/03/2018**

**Important Note:** This Print View may not reflect all comments and contingencies for approval. Please check the comments section of the online protocol.
Questions that appear to not have been answered may not have been required for this submission. Please see the system application for more details.

*** * * Attached Document * * ***

| Document Name | Created Date |
| --- | --- |
| 2011-09-3652_Colford_2s-Consent-In depth interview environmental assessment-v17-Bengali.pdf | 10/26/2017 |

#### Appendix 2s: Bangla consent form for in depth interview in relationship to environmental assessment

<u>**স্বেচ্ছা সম্মতিপত্র**</u>
আন্তর্জাতিক উদরাময় গবেষণা কেন্দ্র, বাংলাদেশ (আই সি ডি ডি আর, বি )
মহাখালী, ঢাকা।

**গবেষণার শিরোনামঃ**
বাংলাদেশের গ্রামীণ এলাকায় হাত ধোয়া, পানি বিশুদ্ধকরণ, স্যানিটেশন ও পুষ্টিকর সম্পূরক খাবার প্রদান এবং শিশুর স্বাস্থ্য ও বৃদ্ধির উপর তার প্রভাব।

**প্রধান গবেষক:**
ডঃ স্টিফেন পি. লুবি

**গবেষণার উদ্দেশ্যঃ**
আস্সালামুআলাইকুম/নমস্কার। আমার নাম -------------------------------। আমি ঢাকার মহাখালীতে অবস্থিত আইসিডিডিআর,বি (কলেরা হাসপাতাল) এ চাকরী করি। শিশু-স্বাস্থ্য ও ডায়রিয়া বিষয়ক আমরা একটি গবেষণা কাজ করছি। এই গবেষণার মাধ্যমে আমরা জানতে চাই নির্দিষ্ট কিছু স্বাস্থ্যসম্মত বিষয়, পানির গুনাগুণ এবং স্বাস্থ্যসম্মত অভ্যাসসমূহ স্বাস্থ্যের উপর কিভাবে প্রভাব ফেলে।

**আমরা আপনাকে কেন এই গবেষণায় অংশগ্রহণের জন্য আমন্ত্রণ জানাচ্ছি?**
আমরা কিছু গৃহস্থালী/খানা দেখতে চাচ্ছি যারা এই গবেষণার অন্যান্য কার্যক্রমে ইতিমধ্যেই অংশগ্রহণ করছে। আমরা এই গবেষণা সম্পর্কে আপনাদেরকে আরও বিস্তারিত কিছু জানাতে চাই এবং আশা করি যে আপনি এবং আপনার শিশু এই গবেষণায় স্বেচ্ছায় অংশগ্রহণ করবেন। আপনারা একটি গ্রামীণ সামাজিক ও প্রাকৃতিক পরিবেশে বাস করছেন যেখান থেকে আমরা আরও কিছু শিখতে চাই। তাছাড়া এই খানায় একজন গর্ভবতী মহিলা আছেন অথবা ছোট শিশু রয়েছে যাদের স্বাস্থ্য নিয়ে আমরা আলাপ করবো।

**এই গবেষণায় অংশগ্রহণকারীদের কাছ থেকে কি প্রত্যাশা করা হচ্ছে?**
আপনার অনুমতিক্রমে আমরা এই বাড়ী বা খানাটিকে এই গবেষণায় অন্তর্ভূক্ত করব। যদি আপনি এই গবেষণায় অংশগ্রহণে রাজী থাকেন তাহলে আমাকে আপনার খানার পানি, স্যানিটেশন এবং স্বাস্থ্যসম্মত আচরণ সংক্রান্ত ব্যবস্থা ও কর্মকান্ড নিয়ে কথা বলার জন্য ৬০ মিনিট-১.৫ঘন্টা সময় দেওয়ার জন্য অনুরোধ জানাবো। আমি আপনার শিশুর স্বাস্থ্য ও তার বিকাশ সম্পর্কেও কিছু কথা বলবো। আপনি অনুমতি দিলে আমাদের উভয়ের কথাবার্তা আমাদের কাজের সুবিধার্থে অডিও ও ভিডিও টেপ রেকর্ড করবো।

**ঝুঁকি এবং সুবিধাসমূহ**
এই গবেষণায় অংশগ্রহনে আপনার এবং আপনার শিশুর ঝুঁকির সম্ভাবনা নুন্যতম। আমরা আপনার পরিচয় সংক্রান্ত তথ্যসমূহ গোপন রাখার জন্য সম্ভাব্য সবধরনের চেষ্টা করব। তা সত্ত্বেও ভুলবশতঃ অনাকাঙ্খিত কোন ঘটনা ঘটলে সেজন্য মার্জনা প্রার্থনা করছি।এই গবেষণায় অংশগ্রহনের জন্য আপনি অথবা আপনার পরিবার অতিরিক্ত কোন ধরণের সহায়তা পাবেন না।

**গোপনীয়তা এবং বিশ্বাসযোগ্যতা**
এই দেশের আইন অনুযায়ী আপনার/ আপনাদের দেওয়া সমস্ত তথ্য গোপনীয়তার সাথে রক্ষণাবেক্ষণ করা হবে। আমরা আমাদের গবেষণার কোন লেখায়/ফলাফলে আপনার প্রকৃত নাম ব্যবহার করব না। আমাদের সংগৃহীত সকল তথ্য তালাবদ্ধ থাকবে। আমরা কেবলমাত্র আমাদের গবেষণার কাজে এসব তথ্য ব্যবহার করব। আমরা এই গবেষণার ফলাফল বিনিময় ও প্রকাশনার ক্ষেত্রে আপনার নাম ব্যবহার করব না। আমরা আপনার দেওয়া তথ্যসমূহ গোপন রাখার জন্য সম্ভাব্য সবধরনের চেষ্টা করব। তা সত্ত্বেও ভুলবশতঃ অনাকাঙ্খিত কোন ঘটনা ঘটলে সেজন্য মার্জনা প্রার্থনা করছি। আপনার রেকর্ডকৃত তথ্য জনগনের কাছে প্রকাশ করা হবে না অথবা কোন প্রকাশনায় আপনার পরিচয় প্রকাশ করা হবে না। গবেষণার ফলাফল প্রকাশের পূর্ব পর্যন্ত আপনার রেকর্ডকৃত তথ্য নিরাপদে সংরক্ষন করা হবে এবং গবেষণার ফলাফল প্রকাশের পর তা মুছে ফেলা হবে।

**ভবিষ্যতে তথ্যের ব্যবহার**

এই গবেষণায় সংগৃহীত তথ্য প্রয়োজনে আমরা অন্যান্য গবেষকের সাথে বিনিময় করতে পারি। এবং কারো সাথে তথ্য বিনিময় করা হলে আমরা সেখানে আপনার প্রকৃত নাম ব্যবহার করব না এবং আপনার দেওয়া তথ্যের গোপণীয়তা ও নিজস্বতা কঠোরভাবে রক্ষা করব।

**অংশগ্রহণ না করার এবং অংশগ্রহণ বাদ দেওয়ার অধিকার**

এই গবেষণায় অংশগ্রহন করা সম্পূর্নভাবে আপনার ইচ্ছার উপর নির্ভর করছে। এই গবেষণায় আপনাকে যেসব প্রশড়ব জিজ্ঞাসা করা হবে তার যেকোনটির উত্তর আপনি চাইলে নাও দিতে পারেন। এই গবেষণায় অংশগ্রহনে সম্মত হবার পরেও গবেষণা চলাকালীন যেকোন সময় আপনি আপনার নাম প্রত্যাহার করে নিতে পারেন। এই গবেষণায় অংশগ্রহণ না করার অধিকার আপনার আছে এবং এ জন্য ভবিষ্যতে আপনার পরিবারের সদস্যদের কলেরা হাসপাতালে স্বাস্থ্যসেবা গ্রহণের উপর কোন প্রভাব পড়বে না।

**ৰতিপূরণ**

এই গবেষণায় অংশগ্রহণের জন্য আপনার কোন ধরনের অর্থনৈতিক খরচ/ ব্যয় বহন করতে হবে না এবং একইভাবে আপনাকেও আমরা কোন ধরনের অর্থ প্রদান করব না।

**যোগাযোগঃ**

আপনার যদি কোন প্রশ্ন থাকে আপনি যে কোন সময় আমাকে তা জিজ্ঞাসা করতে পারেন। এছাড়াও যদি আপনার এই জরিপ বা গবেষণা সম্পর্কে আরও কোন প্রশ্ন থাকে তবে আপনি ওয়াটার, স্যানিটেশন এন্ড হাইজিন রিসার্চ গ্রম্নপ, সেন টার ফর কমিউনিকেবল ডিজিজেজ, আইসিডিডিআ,বি, মহাখালী, ঢাকা - ১২১২ এর রিসার্চ ইনভেস্টিগেটর জনাব মুহাম্মদ ফারম্নক হুসাইন এর মোবাইল নম্বর- ০১৭১১৪৩৭৩২৬ এ যোগাযোগ করতে পারেন।

যদি গবেষণায় অংশগ্রহণের অধিকার সম্পর্কে আপনার কোন প্রশ্ন থাকে অথবা এই গবেষণার কারণে আপনার কোনরূপ ৰতি সাধিত হয়েছে তবে নিম্নলিখিত ব্যক্তির সাথে আপনি যোগাযোগ করতে বা ব্যক্তিগতভাবে দেখা করতে পারেনঃ

জনাব এম এ সালাম খান, আইআরবি সেক্রেটারিয়েট, ফোন-৯৮৮৬৪৯৮ অথবা পিএবিএক্স ৮৮৬০৫২৩-৩২, এক্স. ৩২০৬.

যদি আপনি আমাদের প্রস্ৰাবের সাথে একমত পোষণ করেন (রাজী থাকেন) ও এই গবেষণায় অংশগ্রহণ করতে আগ্রহী হন তাহলে এই গবেষণায় আপনার গৃহস্থালী (খানা) কে অন্তর্ভূক্ত করা হবে। দয়া করে নিম্নের ফাকা স্থানে আপনি আপনার স্বাৰর বা বাম বৃদ্ধাঙ্গুলির ছাপ দিয়ে আপনার সম্মতি প্রদান করম্নন।

আপনার সহযোগিতার জন্য আপনাকের অনেক ধন্যবাদ।

________________________________ ____________

অংশগ্রহণকারীর স্বাৰর বা বাম হাতের বৃদ্ধাঙ্গুলির ছাপ তারিখ

________________________________ ____________

একজন স্বাৰীর স্বাৰর বা বাম হাতের বৃদ্ধাঙ্গুলির ছাপ তারিখ

________________________________ ____________

প্রিন্সিপাল ইনভেস্টিগেটর বা তার প্রতিনিধির স্বাৰর তারিখ

PROTOCOL
Biomedical
Non-Exempt
Berkeley

Protocol # 2011-09-3652
Date Printed: 07/31/2018

---------------------------------------------------------------------------------------

| | |
|---|---|
| **Protocol Title:** | Measuring the benefits of sanitation, water quality, handwashing and nutrition interventions for improving health and development in rural Bangladesh |
| **Protocol Status:** | APPROVED |
| **Date Submitted:** | 10/26/2017 |
| **Approval Period:** | **10/30/2017-11/03/2018** |
| **Important Note:** | **This Print View may not reflect all comments and contingencies for approval. Please check the comments section of the online protocol. Questions that appear to not have been answered may not have been required for this submission. Please see the system application for more details.** |

*** * * Attached Document * * ***

| Document Name | Created Date |
|---|---|
| 2011-09-3652_Colford_1t-Consent-Focus-Group-Discussion-v17-English.pdf | 10/26/2017 |

#### Appendix 1t: English consent form for Focus Group Discussion (FGD)

Protocol Title: Effect of hand washing, water treatment, sanitation and nutritional supplement interventions on child health and development in rural Bangladesh

Principal Investigator's name: Dr. Stephen P. Luby

##### Purpose of the research

Hello (Assalamualaikum/Namaste). My name is ________ and I work with the ICDDR,B (Cholera Hospital) in Dhaka. We are doing research on diarrheal diseases in children. Through this research we want to learn how certain conditions in sanitation, water quality and hygiene practices impact health.

**Why are we inviting you to participate in the study?**
We are seeking some participants who have been living within our study area and can provide us some valuable information about the local water, sanitation and hygiene practices. We know that you are undertaking a great responsibilities for your children, all other family members including maintaining the household environment. Your are also living in a rural social and natural environment from where we would like to learn something more. We are interested to know some of rural water, sanitation and hygiene behaviour, particularly about our family members during every day life that may help us to understand how environmental contaminations may occur including fecal matters and draw a pathway map.

**What is expected from the participants of the research study?**
If you agree to take part in the discussion, I will let you know a common place and request you to present and meet with some other participants invited for discussion. I will talk about some water sanitation and hygiene activities and practices of household members in their daily activities including your child. We will take notes of our discussion and conversations. To make sure we captured well the important information and discussion points we will tape record. I would ask your permission to tape record the discussion session. We would request you to provide us 1 hour of your valuable time. One of our colleagues will assist us taking notes and tape record our conversation.

**Risks & Benefits**
This study presents minimal risk to you and your child. Although we will try to protect your identity there is some possibility that confidentiality could be compromised. You will not directly benefit from participating in this study but this participation will enable us to gain knowledge regarding fecal contamination in association with diarrhea.

**Privacy and confidentiality**
We will do everything we can keep what you tell us confidential as allowed by the law of this country. We will not use your real name when we write out the data. All the information we collect will be kept locked. We will use the information only for the purpose of the study. We will not use any of your name when we share and publish the results of this study. We expect the steps we take will keep all of your information confidential, but it is possible that because of mistakes or unforeseen events, it could become compromised. The tape recordings will not be publicly shared, nor will participants be identifiable in any published works. The recordings will be stored securely until data analysis is complete, and will then be erased.

**Future use of information**
If it is necessary, we may share the information we collected from this study with other researchers. If this is done, we will not use your real name and we will maintain your confidentiality.

**Right not to participate and withdraw**
Taking part in the study is completely voluntary. You may choose not to answer any or all of the questions that will ask. You can drop out of this study at any time, even in the middle of the discussion. You have the right to refuse to take part in this study, which will not affect your family's treatment at the Cholera Hospital in the future.

**Principle of compensation**
You do not need to pay us to take part in this study, and we will not pay you money for taking part in the study unless you have some local transportation cost to attend in the discussion and some refreshments.

**Persons to contact:**

If you have any questions, you can ask me any time even during the discussion. But if you ask any question now or before starting the discussion it would be helpful for us. If you have additional questions about the study or discussion, you may contact: Muhammad Faruqe Hussain: phone: 01711437326 from the Water Sanitation and Hygiene Research Group, Centre for Communicable Diseases (CCD), icddr,b, Mohakhali, Dhaka 1212.

If you have questions about your rights as a participant of a research study, or if you think some harm has been done to you because of this study, you may contact or meet him personally at following address:

M. A. Salam Khan, IRB Secretariat, phone: 9886498 or PABX 8860523-32 ext. 3206

If you agree to our proposal of participation in the Focus Group Discussion, please indicate that by putting your signature or your left thumb impression at the specified space below.

Thank you for your cooperation

______________________________________ ____________________

Signature or left thumb impression of Participant Date

______________________________________ ____________________

Signature or left thumb impression of the witness Date

________________________________________ ____________________

Signature of the PI or his/her representative Date

Name of the moderator:

Name of the note taker:

________________________________________ ____________________

Signature or left thumb impression of the witness Date

________________________________________ ____________________

Signature of the PI or his/her representative Date

PROTOCOL
Biomedical
Non-Exempt
Berkeley

Protocol # 2011-09-3652
Date Printed: 07/31/2018

---------------------------------------------------------------------------------------------

**Protocol Title:** Measuring the benefits of sanitation, water quality, handwashing and nutrition interventions for improving health and development in rural Bangladesh

**Protocol Status:** APPROVED

**Date Submitted:** 10/26/2017

**Approval Period:** **10/30/2017-11/03/2018**

**Important Note:** **This Print View may not reflect all comments and contingencies for approval. Please check the comments section of the online protocol. Questions that appear to not have been answered may not have been required for this submission. Please see the system application for more details.**

*** * * Attached Document * * ***

| Document Name | Created Date |
| --- | --- |
| 2011-09-3652_Colford_2t-Consent-Focus-Group-Discussion-v17-Bengali.pdf | 10/26/2017 |

Q.T/ S.G/ Cl. ID/ Co. ID/ B. ID/ HH ID
Household ID: □/□/ □□/ □□/□□/□□

#### Appendix 2t: Bengali consent form for Focus Group Discussion (FGD)

স্বেচ্ছা সম্মতিপত্র
আন্তর্জাতিক উদরাময় গবেষণা কেন্দ্র, বাংলাদেশ (আই সি ডি ডি আর, বি )

মহাখালী, ঢাকা।

**গবেষণার শিরোনামঃ**

বাংলাদেশের গ্রামীণ এলাকায় হাত ধোয়া, পানি বিশুদ্ধকরণ, স্যানিটেশন ও পুষ্টিকর সম্পূরক খাবার প্রদান এবং শিশুর স্বাস্থ্য ও বৃদ্ধির উপর তার প্রভাব।

**প্রধান গবেষক:**

ড: স্টিফেন পি. লুবি

**গবেষণার উদ্দেশ্যঃ**

আস্সালামুআলাইকুম/নমস্কার। আমার নাম -------------------------------। আমি ঢাকার মহাখালীতে অবস্থিত আইসিডিডিআর,বি (কলেরা হাসপাতাল) এ চাকরী করি। শিশু-স্বাস্থ্য ও ডায়রিয়া বিষয়ক আমরা একটি গবেষণা কাজ করছি। এই গবেষণার মাধ্যমে আমরা জানতে চাই নির্দিষ্ট কিছু স্বাস্থ্যসম্মত বিষয়, পানির গুনাগুণ এবং স্বাস্থ্যসম্মত অভ্যাসসমূহ স্বাস্থ্যের উপর কিভাবে প্রভাব ফেলে ।

**আমরা আপনাকে কেন এই গবেষণায় অংশগ্রহণের জন্য আমন্ত্রণ জানাচ্ছি?**

আমরা কিছু সংখ্যক অংশগ্রহণকারী খুজছি যারা আমাদের গবেষণাধীন এলাকায় বাস করছেন এবং স্থানীয় পানি, স্বাস্থ্যসম্ভবত অভ্যাসসমূহের চর্চা সম্পর্কে মূল্যবান তথ্য দিতে পারেন। আমরা জানি যে আপনারা শিশুদের লালন-পালনসহ পরিবারের অন্যান্য সদস্যদের দেখাশোনা ও গৃহস্থালীর পরিবেশ রক্ষণাবেক্ষণের গুরু দায়িত্ব গ্রহণ করেছেন। তাছাড়া আপনারা একটি প্রাকৃতিক ও সামাজিক পরিবেশে বাস করছেন যেখান থেকে আমরা আরও অনেক বেশী কিছু জানতে চাই। আমরা গ্রামীণ আপনাদের পানি, স্বাস্থ্যসম্মত অভ্যাস ও চর্চা বিশেষ করে আপনাদের নিজেদের পরিবারের সদস্যদের প্রাত্যহিক জীবন-যাপন সংক্রান্ত বিষয়ে জানতে চাই যা আমাদেরকে পরিবেশের বিভিন্ন দূষণ এমনকি মল সংক্রান্ত দুষণ ঘটতে পারে তা এবং কোথায় থেকে কোথায় কিভাবে যায় তার একটি চিত্র অংকন করতে চাই।

**এই গবেষণায় অংশগ্রহণকারীদের কাছ থেকে প্রত্যাশা কি?**

যদি আপনি এই আলোচনায় অংশগ্রহণে রাজী থাকেন তাহলে আমি আপনাকে একটি স্থানে উপস্থিত হতে এবং এই আলোচনায় আমন্ত্রিত আরও কয়েকজন অংশগ্রহণকারীর সাথে সাক্ষাতের জন্য অনুরোধ করবো। আমি আপনাদের ও আপনাদের এলাকার মানুষের ও শিশুদের দৈনন্দিন জীবন-যাপনে পানি, স্বাস্থ্যসম্মত অভ্যাস ও চর্চা বা কর্মকান্ডসমূহ সম্পর্কে আলোচনা করবো। আমরা আমাদের আলোচনা ও কথোপকথনগুলি লিখে নিবো। আপনাদের সাথে আমাদের এই গুরুত্বপূর্ণ আলোচনার বিষয়বস্তুসমূহ নিশ্চিতভাবে বোঝার জন্য আমরা কথাগুলি রেকর্ড করবো। আমি প্রত্যাশা করবো আমাদের এই আলোচনা টেপ রেকর্ড করতে আপনারা আমাদেরকে অনুমতি দিবেন। আমরা এই আলোচনার জন্য এক ঘন্টা সময় প্রদানের জন্য আপনাদেরকে অনুরোধ করবো। আমাদের একজন সহকর্মী উপস্থিত থেকে আমাদের আলোচনা লেখা ও টেপ রেকর্ড করতে সাহায্য করবেন।

**ঝুঁকি এবং সুবিধাসমূহ**

এই গবেষণায় অংশগ্রহনে আপনার কোন ঝুঁকি নেই। আমরা আপনার পরিচয়সংক্রান্ত তথ্যসমূহ গোপন রাখার জন্য সম্ভাব্য সবধরনের চেষ্টা করব। তা সত্ত্বেও ভুলবশতঃ অনাকাঙ্খিত কোন ঘটনা ঘটলে সেজন্য মার্জনা প্রার্থনা করছি।

এই গবেষণায় অংশগ্রহনের জন্য আপনি সরাসরি কোন ধরণের সহায়তা পাবেন না। কিন্তু আপনার অংশগ্রহণ, ডায়রিয়া রোগ সম্পর্কে জ্ঞানলাভে আমাদেরকে সহায়তা করবে।

## গোপনীয়তা এবং বিশ্বাসযোগ্যতা

এই দেশের আইন অনুযায়ী আপনি আমাদেরকে যা বলবেন তা গোপনীয় রাখার জন্য সব কিছুই করবো। আমরা আমাদের গবেষণার কোন লেখায়/ফলাফলে আপনার প্রকৃত নাম ব্যবহার করব না। আমাদের সংগৃহীত সকল তথ্য তালাবদ্ধ থাকবে। আমরা কেবলমাত্র আমাদের গবেষণার কাজে এসব তথ্য ব্যবহার করব। আমরা যখন এই গবেষণার ফলাফল প্রকাশ করবো এবং অন্যকেরকে জানাবো তখন আপনার নাম ব্যবহার করবো না। আমরা আশা করি আমরা যে সকল পদক্ষেপ গ্রহণ করবো তা আপনার প্রদত্ত তথ্যকে গোপন রাখতে সাহায্য করবে।

আমরা এই গবেষণার ফলাফল বিনিময় ও প্রকাশনার ক্ষেত্রে আপনার নাম ব্যবহার করব না। আমরা আপনার দেওয়া তথ্যসমূহ গোপন রাখার জন্য সম্ভাব্য সবধরনের চেষ্টা করব। তা সত্ত্বেও ভুলবশতঃ অনাকাঙ্খিত কোন ঘটনা ঘটলে সেজন্য মার্জনা প্রার্থনা করছি। টেপ রেকর্ডগুলি কারো কাছে বা অংশগ্রহণকারীগণের কোনরূপ পরিচয় কোন কাজে প্রকাশ করা হবে না। এই রেকর্ডিংগুলি তথ্য বিশ্লেষণ হওয়া পর্যন্ত নিরাপত্তার সাথে সংরক্ষিত থাকবে এবং তারপর তা মুছে ফেলা/নস্ট করে ফেলা হবে।

#### ভবিষ্যতে তথ্যের ব্যবহার

এই গবেষণায় সংগৃহীত তথ্য প্রয়োজনে আমরা অন্যান্য গবেষকের সাথে বিনিময় করতে পারি। এবং কারো সাথে তথ্য বিনিময় করা হলে আমরা সেখানে আপনার প্রকৃত নাম ব্যবহার করব না এবং আপনার দেওয়া তথ্যের গোপণীয়তা ও নিজস্বতা কঠোরভাবে রক্ষা করব।

#### অংশগ্রহণ না করার এবং অংশগ্রহণ বাদ দেওয়ার অধিকার

এই গবেষণায় আপনার অংশগ্রহণ সম্পূর্ণভাবে স্বেচ্ছামূলক। আপনাকে জিজ্ঞাসা করা সকল বা যে কোন প্রশ্নের উত্তর আপনি যদি দিতে না চান তবে কোন অসুবিধা নেই অথবা আপনি আপনার পছন্দ অনুযায়ী প্রশ্নের উত্তরও দিতে পারেন। এই গবেষণা থেকে যে কোন সময় আপনার অংশগ্রহণ বাদ দিতে পারেন, এমনকি আলোচনা চলাকালীন যেকোন সময়কালেও। এই গবেষণায় অংশগ্রহণ না করার অধিকার আপনার আছে এবং এ জন্য ভবিষ্যতে আপনার পরিবারের সদস্যদের কলেরা হাসপাতালে স্বাস্থ্যসেবা গ্রহণের উপর কোন প্রভাব পড়বে না।

**ক্ষতিপূরণ**

এই গবেষণায় অংশগ্রহণের জন্য আপনার কোন ধরনের অর্থনৈতিক খরচ/ ব্যয় বহন করতে হবে না এবং একইভাবে আপনাকেও আমরা কোন ধরনের অর্থ প্রদান করব না যদি না অংশগ্রহণের জন্য এই আলোচনার স্থানে আসতে এবং কিছুটা আপ্যায়নের জন্য কোন খরচ হয়।

**যোগাযোগঃ**

আপনার যদি কোন প্রশ্ন থাকে আপনি যে কোন সময় আমাকে তা জিজ্ঞাসা করতে পারেন এমনকি আলোচনা চলাকালীন সময়েও। তবে আপনি যদি এখনই বা আলোচনা শুরু হওয়ার পূর্বেই প্রশ্নগুলি জিজ্ঞাসা করেন তাহলে আমাদের জন্য সুবিধা হয়। এছাড়াও যদি আপনার এই গবেষণা বা আলোচনা সম্পর্কে আরও কোন প্রশ্ন থাকে তবে আপনি ওয়াটার, স্যানিটেশন এন্ড হাইজিন রিসার্চ গ্রুপ, সেন্টার ফর কমিউনিকেবল ডিজিজেজ, আইসিডিডিআ,বি, মহাখালী, ঢাকা -১২১২ এর রিসার্চ ইনভেস্টিগেটর জনাব মুহাম্মদ ফারুক হুসাইন এর মোবাইল নম্বর- ০১৭১১৪৩৭৩২৬ এ যোগাযোগ করতে পারেন।

যদি গবেষণায় অংশগ্রহণের অধিকার সম্পর্কে আপনার কোন প্রশ্ন থাকে অথবা এই গবেষণার কারণে আপনার কোনরূপ ক্ষতি সাধিত হয়েছে তবে নিম্নলিখিত ব্যক্তির সাথে আপনি যোগাযোগ করতে বা ব্যক্তিগতভাবে দেখা করতে পারেনঃ
জনাব এম এ সালাম খান, আইআরবি সেক্রেটারিয়েট, ফোন-৯৮৮৬৪৯৮ অথবা পিএবিএক্স ৮৮৬০৫২৩-৩২, এক্স. ৩২০৬.
যদি আপনি দলীয় আলোচনায় অংশগ্রহণে আমাদের প্রস্তাবের সাথে একমত পোষণ করেন (রাজী থাকেন) তাহলে দয়া করে নিম্নের ফাকা স্থানে আপনি আপনার স্বাক্ষর বা বাম বৃদ্ধাঙ্গুলির ছাপ দিয়ে আপনার সম্মতি প্রদান করুন।
এই আলোচনায় অংশগ্রহণ করে সহযোগিতার জন্য আপনাকে অনেক ধন্যবাদ।

| | |
|---|---|
| ........................................... | ................................. |
| উত্তরদাতার স্বাক্ষর/বাম বৃদ্ধাঙ্গুলির ছাপ | তারিখ |
| ........................................... | ................................. |
| স্বাক্ষীর স্বাক্ষর/বাম বৃদ্ধাঙ্গুলির ছাপ | তারিখ |
| ........................................... | ................................. |
| পি আই/ গবেষকের প্রতিনিধির স্বাক্ষর | তারিখ |

মডারেটরের নামঃ

নোটগ্রহণকারীর নামঃ

| | |
|---|---|
| --- | --- |
| একজন স্বাক্ষীর স্বাক্ষর বা বাম হাতের বৃদ্ধাঙ্গুলির ছাপ | তারিখ |
| --- | --- |
| প্রিন্সিপাল ইনভেস্টিগেটর বা তার প্রতিনিধির স্বাক্ষর | তারিখ |

PROTOCOL
Biomedical
Non-Exempt
Berkeley

Protocol # 2011-09-3652
Date Printed: 07/31/2018

-----------------------------------------------------------------------------------

| | |
|---|---|
| **Protocol Title:** | Measuring the benefits of sanitation, water quality, handwashing and nutrition interventions for improving health and development in rural Bangladesh |
| **Protocol Status:** | APPROVED |
| **Date Submitted:** | 10/26/2017 |
| **Approval Period:** | **10/30/2017-11/03/2018** |
| **Important Note:** | **This Print View may not reflect all comments and contingencies for approval. Please check the comments section of the online protocol. Questions that appear to not have been answered may not have been required for this submission. Please see the system application for more details.** |

*** * * Attached Document * * ***

| Document Name | Created Date |
|---|---|
| 2011-09-3652_Colford_1u-Consent-Parasites-v21-English.pdf | 10/26/2017 |

**Appendix 1u: English consent form for longitudinal environmental assessment**
Protocol Title: Effect of hand washing, water treatment, sanitation and nutritional supplement interventions on child health and development in rural Bangladesh

Principal Investigator's name: Dr. Leanne Unicomb

#### Purpose of the research

Hello (Assalamualaikum/Namaste). My name is ________ and I work with the ICDDR,B (Cholera Hospital) in Dhaka. We are conducting a research on diarrheal diseases in children to learn how certain conditions in sanitation, water quality and hygiene practices impact health. This research is being conducted in collaboration with scientists at the University of California Berkeley in the United States.

**Why are we inviting you to participate in the study?**
We are seeking households that are already taking part in other activities from this existing study. We would like to tell you more detail about this study and hope you and your child will volunteer to participate in this study.

**What is expected from the participants of the research study?**
If you decide to take part in the study, we will visit your household now and we will continue to visit you once every three months over the next two years (for a total of eight visits). At each visit, we will collect some samples from your compound environment, including a water sample from your stored drinking water, and during two of these visits we will also collect some soil from the household entrance and a small amount of food that you feed to your young children. At each visit, we will also rinse your hands and your child's hands in clean water and collect the rinsed water. We will take all of these samples back to our field lab to analyse them for bacterial contamination. These samples will help us understand what makes your children become sick by conducting laboratory tests. We will also take a look at your compound with your permission to observe your water, latrine and handwashing facilities. We will also ask you about your children's health. Each visit will be less than 1 hour. Additionally, after 3 months from now, we will install a sensor device in your latrine, which does not capture any video or audio information. We will leave this device in your latrine for one week and then remove it. The sensors capture the timing of when someone enters or exits the latrine. We will do this in your household twice within 2 years.

**Risks & Benefits**
This study presents minimal risk to you and your child. There is a slight risk of breach of confidentiality, but we will do our best to minimize this risk. You will not directly benefit from participating in this study but this participation will enable us to gain knowledge regarding fecal contamination in association with diarrhea.

**Confidentiality**

All data and specimens collected will be kept confidential to the greatest extent possible. Researchers at ICDDR,B will have access to some of your personal information that could identify you, such as your name and your phone number. They will remove the identifiable information before sharing it with researchers at partner institutions, such as UC Berkeley. Your research records will be kept in a locked cabinet, and computer-based data, will be stored in an encrypted format on a password-protected server.

Other persons who may have access to your information include research groups that oversee the safety of the study including institutes such as the University of California Berkeley and the Bill and Melinda Gates Foundation, USA. None of the researchers outside of ICDDR,B will be able to identify you or your child from information you provide. We will use the information only for the purpose of the study. We will not use your name when we share and publish the results of this study. We expect the steps we take will keep all of your information confidential, but it is possible that because of mistakes or unforeseen events, it could become compromised.

**Future use of information**
If it is necessary, we may share the information we collected from this study with other researchers. If this is done, we will not use your real name and we will maintain your confidentiality. We will store the information we collect from you for a long time after the end of the study. Data resulting from research involving children will be stored for at least 7 years after the child reaches the age of 18 years.

**Right not to participate and withdraw**
Taking part in the study is completely voluntary. You have the right to decline to allow your child to participate or to withdraw your child at any point in this study without penalty or loss of benefits to which you are otherwise entitled. You may choose not to answer any or all of the questions that will ask. You can drop out of this study at any time, even in the middle of an interview. You have the right to refuse to take part in this study, which will not affect your family's treatment at the Cholera Hospital in the future.

**Principle of compensation**
You do not need to pay us to take part in this study, and we will not pay you money for taking part in the study.

**Persons to contact:**

If you have any questions, you can ask me any time. If you have additional questions about the survey, you may contact: Mahfuza Islamphone:+880-2-8819419-20, Ext: 118 , Center of Communicable Diseases, ICDDR,B, Mohakhali, Dhaka 1212.

If you have questions about your rights as a participant of a research study, or if you think some harm has been done to you because of this study, you may contact or meet IRB Secretariat, M.A. Salam Khan:

M. A. Salam Khan, IRB Secretariat, phone: 9886498 or PABX 8860523-32 ext. 3206

If you agree to participate in our study and to allow your child to participate in our study, please indicate that by putting your signature or your left thumb impression at the specified space below

Thank you for your cooperation

______________________________________ ____________________

Signature or left thumb impression of Participant Date

______________________________________ ____________________

Signature or left thumb impression of the witness Date

______________________________________ ___________________

Signature of the PI or his/her representative Date

PROTOCOL
Biomedical
Non-Exempt
Berkeley

Protocol # 2011-09-3652
Date Printed: 07/31/2018

-----------------------------------------------------------------------------------

| | |
|---|---|
| **Protocol Title:** | Measuring the benefits of sanitation, water quality, handwashing and nutrition interventions for improving health and development in rural Bangladesh |
| **Protocol Status:** | APPROVED |
| **Date Submitted:** | 10/26/2017 |
| **Approval Period:** | **10/30/2017-11/03/2018** |
| **Important Note:** | **This Print View may not reflect all comments and contingencies for approval. Please check the comments section of the online protocol. Questions that appear to not have been answered may not have been required for this submission. Please see the system application for more details.** |

*** * * Attached Document * * ***

| Document Name | Created Date |
|---|---|
| 2011-09-3652_Colford_2u-Consent-Longitudinal-EnvAssess-v21-Bengali.pdf | 10/26/2017 |

**Appendix 2u:** Bengali consent form for longitudinal environmental assessment

cwi‡ekMZ cfve gj¨vq‡bi Rb¨ m¤§wZcÎ

M‡elYvi wk‡ivbvg:evsjv‡`‡ki MvgxY GjvKvq nvZ †avqv, cwb weï×KiY, m¨wb‡Ukb I cwóKi m¤úiK Lvevi c`vb Ges wkïi ¯v¯¨ I ewׄi Dci Zvi cfve|

cavb M‡elK:W: wj‡qb BDwbK¤

M‡elYvi D‡Ïk¨:

AvmjvgAvjvBKg/Av`ve| Avgvi bvg ----------------------------| Avgiv XvKv (gnvLvjx) K‡jiv nvmcvZv‡j KvR Kwi|Avgiv wkï‡`i WvqwiqvRwbZ †ivM-evjvB wb‡q M‡elYv KiwQ Ges Rvb‡Z †Póv KiwQ m¨wb‡Ukb e¨e¯v ¯v‡¯¨i Dci wKfv‡e cfve ‡d‡j |

Avgiv †Kb Avcbv‡K GB M‡elYvq AskMn‡b AvgšY Rvbw"Q?

Avgiv Avcbv‡K/Avcbvi Lvbv‡K GB M‡elYvq AskMn‡Yi Rb¨ Avgšb Rvbw"Q KviY Avcwb/Avcbvi Lvbv BwZg‡a¨B Avgv‡`i M‡elYvi Ab¨vb¨ KgKv‡Û AskMnb K‡i‡Qb| Avgiv Avcbv‡K/Avcbv‡`i‡K GB M‡elYvi wel‡q Av‡iv we¯ÍwiZfv‡e Rvbv‡Z PvB Ges Avgiv Avkv KiwQ Avcwb Ges Avcbvi wkï ‡¯"Qvq GB M‡elYvq AskMnb Ki‡eb| GQvov GB Lvbvq GKRb MfeZx gv Av‡Q A_ev eZgv‡b GLv‡b †QvU wkï i‡q‡Q hv‡`i ¯v¯¨ wb‡q Avgiv K_v ej‡ev|

M‡elYvq AskMnYKvixi Kv‡Q cZ¨vkv Kx?

hw` Avcwb/Avcbvi Lvbv GB M‡elYvq AskMn‡Y m¤§Z _v‡Kb, Zvn‡j GLb Avgviv Avcbvi Lvbv cwi`kbKie Ges cwZ wZb gvm ci ci Avcbvi Lvbv cw`kb Kie hv cieZx 2 eQi a‡i Pj‡e ( ‡gvU 8 evi cw`kb Kiv n‡e) c‡Z¨Kevi cwi`kbKv‡j Avgiv Avcbvi/Avcbv‡`i msi¶bKZ cwbi cvÎ n‡Z Lvevi cwb bgbv wnmv‡e msMn Kie|Avcbv‡K Ges Avcbvi wkïi nvZ‡avqve Ges †mB nvZ‡avqv cwb bgbv wnmv‡e msMn Kie| `BwU cwi`kbKv‡j Avcbv Lvbvi c‡ek ¯v‡bi gwUi bgbv Ges Avcbvi †QvU wkï‡K †h Lvevi Lv Iqvb †mLvb †_‡K mvgvb¨ cwigvb Lvevi msMn Kie| Gme msMnKZ bgbv Avgiv cix¶v Kivi/we‡køl‡bi Rb¨ M‡elYvMv‡i wb‡q hve| wK Kvi‡Y wkïiv Am¯ nq Gme bgbv Avgv‡`i‡K Zv Rvb‡Z mnvqZv Ki‡e| Avgiv Avcbvi AbgwZ mv‡c‡¶ Avcbvi/Avcbv‡`i cwb, cvqLvbv Ges nvZ‡avqvi m‡hvM-mweaw` ch‡e¶Y Kie| Avgiv Avcbvi wkïi ¯v¯¨ m¤ú‡K wKQ ckl wRÁvmv Kie| c‡Z¨Kevi cwi`kbKv‡j Avgiv Avcbvi Lvbv‡Z cvq 1 N›Uv mgq KvUve|GQvov AvgivcvqwZbgvm ci Avcbvi Lvbvi cvqLvbv‡Z GKwU ms‡e`bkxj hb ¯vcb Kie hv wfwWI ev AwWI Z_¨ aviY Ki‡e bv| Avgiv GB hb GK mvßvn ci Avcbvi cvqLvbv †_‡K mwi‡q †be Ges aviYKZ Z_¨ ewZj Kie | GB KvRwU Avgiv Avcbvi Lvbv‡Z AvMvgx 2 eQ‡ii g‡a¨ †gvU 2 evi Kie|

SwK Ges mweavmgn:

GB M‡elYvq Avcbvi Ges Avcbvi wkïi SwKi m¤vebv bb¨Zg| Avcbvi cwiPqmsµvšÍ Z_¨mgn †Mvcb bv _vKvi mvgvb¨ m¤vebv i‡q‡Q wKš Avgiv Avcbvi cwiPqmsµvšÍ Z_¨mgn †Mvcb ivLvi Rb¨ m¤ve¨ meai‡bi †Póv Kie| Zv m‡ËI fjekZt AbvKw•LZ †Kvb NUbv NU‡j ‡mRb¨ gvRbv cv_bv KiwQ|GB M‡elYvq AskMn‡bi Rb¨ Avcwb mivmwi †Kvb ai‡Yi mnvqZv cv‡eb bv|wKš Avcbvi wkïi AskMnY, wkï‡`i Wvqwiqv †ivM m¤ú‡K Ávbjv‡f Avgv‡`i‡K mnvqZv Ki‡e|

wekvm‡hvM¨Zv:

Avgiv Avcbvi/ Avcbv‡`i †`Iqv mg¯Í Z‡_¨i †MvcbxqZv eRvq ivLvi Rb¨ m‡ev"P †Póv Kie| Avcbv‡`i †`Iqv wKQZ_¨ AvBwmwWwWAvi,we i M‡elK‡`i Kv‡Q _vK‡e hvi d‡j Avcbv‡K mbv³ Kiv m¤e, ‡hgb Avcbvi bvg Ges ‡dvb b¤i | Zviv

Avcbv‡`i †`Iqv Z_¨ ‡_‡K Avcbvi mbv³KiY gjK Z_¨ mgn ev` w`†q Ab¨vb¨ M‡elK†`i mv‡\_ ‡kqvi Ki‡e (‡hgb Av‡gwiKvi K¨wj‡dwbqv evK‡j wekwe`¨vj‡qi M‡elK), hvi d‡j‡Kvb M‡elKB Avcbv‡K wPwýZ Ki‡Z cvi‡e bv | M‡elYvi me †iKW Zvjvex× K‡i ivLv n‡e I Kgwc¤úDUv‡i Z_¨vejx cvmIAviW w`†q msi¶b Kiv n‡e|

Ab¨vb¨ e¨w³ A_ev M‡elYvi mv‡_ mswkøó `j Avcbvi Z\_¨ ‡`L‡Z cvi‡e hw` M‡elYvi Kvi‡Y c‡qvRb g‡b K‡i| Giv n‡jb- Av‡gwiKvi K¨wj‡dwbqv evK‡j wekwe`¨vj‡qi M‡elK I M‡elYvi ¯úYi (wej I ‡gwjÛv ‡MUm dvD‡Ûkb) | AvBwmwWwWAvi,we i M‡elK†`i Qvov †KD Avcbv‡K Ges Avcbvi wkï‡K mbv³ Ki‡Z cvi‡e bv | Avgiv ïagvÎ M‡elYvi Kv‡R GB Z_¨ e¨envi Kie Ges Avcbvi bvg A_ev cwiP‡qi †Kvb mÎ GB M‡elbv-msµvšÍ †Kvb wi‡cvU/ cKvkbvq D‡jøLKiv n‡e bv| Avcbvi KvQ ‡_‡K cvß mKj Z_¨ I bgbvi m‡ev"P ‡MvcbxqZv i¶vi Rb¨ Avgiv mevZ¥K ‡Póv Kie, ZeI fjekZt AbvKvw•LZ †Kvb NUbvi Kvi‡b ‡MvcbxqZv i¶v wbwðZ Ki‡Z bv cvi‡j ‡mRb¨ gvRbv cv_bv KiwQ|

fwel¨‡Z Z‡_¨i e¨envi:

GB M‡elYvq msMnxZ Z_¨ c‡qvR‡b Avgiv Ab¨vb¨ M‡el‡Ki mv‡_ wewbgq Ki‡Z cwi|Kv‡iv mv‡_ Z_¨ wewbgq Kiv n‡j Avgiv †mLv‡b Avcbvi cKZ bvg e¨envi Kie bv Ges Avcbvi †`Iqv Z‡_¨i †MvcYxqZv I wbR¯Zv K‡Vvi fv‡e i¶v Kie|

M‡elYvq AskMnb bv Kiv ev bvg cZ¨vnv‡ii AwaKvit

G M‡elYvq Avcbvi wkïi AskMnY ‡m"QvgjK Ges Avcbvi wkïi AskMnY Kiv ev bv Kiv m¯úYfv‡e Avcbvi B"Qvi Dci wbfi Ki‡Q| GB M‡elYvq Avcbv‡K †hme ck wRÁvmv Kiv n‡e Zvi †h‡KvbwUi DËi Avcwb PvB‡j bvI w`‡Z cv‡ib| GB M‡elYvq AskMn‡b m¤§Z nevi c‡iI M‡elYv PjvKvjxb †h‡Kvb mgq Avcwb Avcbvi bvg cZ¨vnvi K‡i wb‡Z cv‡ib| GB M‡elYvq AskMnY bv Kivi AwaKvi Avcbvi Av‡Q Ges G Rb¨ fwel¨‡Z Avcbvi cwiev‡ii m`m¨‡`i K‡jiv nvmcvZv‡j ¯v¯¨‡mev Mn‡Yi Dci †Kvb cfve co‡e bv|

¶wZciY :

GB M‡elYvq AskMn‡Yi Rb¨ Avcbvi †Kvb ai‡bi A_%bwZK LiP/ e¨q enb Ki‡Z n‡e bv Ges GKBfv‡e Avcbv‡KI Avgiv †Kvb ai‡bi A_ c`vb Kie bv|

‡hvMv‡hvM:

Avcbvi †Kv‡bv ck _vK‡j Avgv‡K wRÁvmv Ki‡Z cv‡ib| hw` M‡elYv m¤úwKZ AwZwi³ †Kvb ck \_v‡K Zvn‡j Avcwb miKvi gvm` cvi‡fR (‡m›Uvi di KwgDwbK¨vej wWwR‡Rm, AvBwmwWwWAvi,we: gnvLwj, XvKv-1212) Gi mv‡_ 01817541872 †Uwj‡dvb b¤‡i mivmwi †hvMv‡hvM Ki‡Z cv‡ib|

GB M‡elYvq Avcbvi AwaKvi-msµvšÍ †Kvb ck _vK‡j A_ev M‡elYvi Kvi‡Y Avcbvi A_%bwZK †Kvb ¶wZ mwaZ n‡q‡Q e‡j g‡b K‡ib, Zvn‡j Avcwb †Uwj‡dvb b¤‡i A_ev mivmwi AvBAviwe †m‡µUwi‡qU Gg G mvjvg Lvb Gi mv‡_ †hvMv‡hvM Ki‡Z cv‡ib|

Gg G mvjvg Lvb, AvBAviwe †m‡µUwi‡qU, 9886498 A_ev wcGweG· 8860523-32 (G·- 3206)

Avcwb hw` GB M‡elYvq AskMnY Ki‡Z AvMnx nb Zvn‡j wb‡gi wbawiZ ¯v‡b ¯v¶i A\_ev evg e×v½jxi Qvc w`b|

Avcbvi mn‡hwMZvi Rb¨ ab¨ev`|

____________________________ _____________________

DËi`vZvi ¯v¶i/evg e×v½wji Qvc ZwiL

____________________________ ______________________

¯v¶xi ¯v¶i/evg e×v½wji Qvc ZwiL

____________________________ ______________________

wc AvB/ M‡el‡Ki cwZwbwai ¯v¶i ZwiL

PROTOCOL
Biomedical
Non-Exempt
Berkeley

-----------------------------------------------------------------------------------

| | |
|---|---|
| **Protocol Title:** | Measuring the benefits of sanitation, water quality, handwashing and nutrition interventions for improving health and development in rural Bangladesh |
| **Protocol Status:** | APPROVED |
| **Date Submitted:** | 10/26/2017 |
| **Approval Period:** | **10/30/2017-11/03/2018** |
| **Important Note:** | **This Print View may not reflect all comments and contingencies for approval. Please check the comments section of the online protocol. Questions that appear to not have been answered may not have been required for this submission. Please see the system application for more details.** |

*** * * Attached Document * * ***

| Document Name | Created Date |
|---|---|
| 2011-09-3652_Colford_1k2-Consent-EnvAssess-water-v21-English.pdf | 10/26/2017 |

**Appendix 1k-2: English consent form for environmental assessment (drinking water and food measurements in water arm)**

Protocol Title: Effect of hand washing, water treatment, sanitation and nutritional supplement interventions on child health and development in rural Bangladesh

Principal Investigator's name: Dr. Stephen P. Luby

#### Purpose of the research

Hello (Assalamualaikum/Namaste). My name is ________ and I work with the ICDDR,B (Cholera Hospital) in Dhaka. We are doing research on diarrheal diseases in children. Through this research we want to learn how certain conditions in sanitation, water quality and hygiene practices impact health. We are conducting this research in collaboration with scientists at University of California, Berkeley in the United States.

**Why are we inviting you to participate in the study?**
We are seeking households that have a child less than 3 years old and have already taken part in other activities from this study. We would like to tell you more detail about this study and hope you and your child will volunteer to participate.

**What is expected from the participants of the research study?**
If you decide to take part in the study, we will conduct some specific activities. During our visit, which we would like to do today, we will ask you questions about your home and compound, and some practices in your home. We will also take a very small amount of stored food that young children in your household will eat. Finally, we will ask you to give us a glass of water that you would give to your children to drink. We will take a small amount of the water that you give us. The visit will be less than 1 hour.

**Risks & Benefits**
There are minimal risks involved in this study. There is a slight risk of breach of confidentiality, but we will do our best to minimize this risk. You will not directly benefit from participating in this study but this participation will enable us to gain knowledge regarding fecal contamination in association with diarrhea.

**Privacy, anonymity and confidentiality**

All data and specimens collected will be kept confidential to the greatest extent possible.. Researchers at ICDDR,B will have access to some of your personal information that could identify you, such as your name and your phone number. They will remove the identifiable information before sharing it with researchers at partner institutions, such as UC Berkeley. Your research records will be kept in a locked cabinet, and computer-based data, will be stored in an encrypted format on a password-protected server.

Other persons who may have access to your information include research groups that oversee the safety of the study including institutes such as the University of California Berkeley and the Bill and Melinda Gates Foundation, USA. None of the researchers outside of ICDDR,B will be able to identify you or your child from information you provide. We will use the information only for the purpose of the study. We will not use your name when we share and publish the results of this study. We expect the steps we take will keep all of your information confidential, but it is possible that because of mistakes or unforeseen events, it could become compromised.

**Future use of information**
If it is necessary, we may share the information we collected from this study with other researchers. If this is done, we will not use your real name and we will maintain your confidentiality. We will store the information we collect from you for a long time after the end of the study.

**Right not to participate and withdraw**
Taking part in the study is completely voluntary. You have the right to decline to allow your child to participate or to withdraw your child at any point in this study without penalty or loss of benefits to which you are otherwise entitled. You may choose not to answer any or all of the questions that will ask. You can drop out of this study at any time, even in the middle of an interview. You have the right to refuse to take part in this study, which will not affect your family's treatment at the Cholera Hospital in the future.

**Principle of compensation**
You do not need to pay us to take part in this study, and we will not pay you money for taking part in the study.

**Persons to contact:**

If you have any questions, you can ask me any time. If you have additional questions about the survey, you may contact: Masud Parvez: phone: 01817541872,Center of Communicable Diseases, ICDDR,B, Mohakhali, Dhaka 1212.

If you have questions about your rights as a participant of a research study, or if you think some harm has been done to you because of this study, you may contact or meet him personally at following address:

M. A. Salam Khan, IRB Secretariat, phone: 9886498 or PABX 8860523-32 ext. 3206

If you agree to participate in our study and for your children to participate in our study, please indicate that by putting your signature or your left thumb impression at the specified space below

Thank you for your cooperation

______________________________________ ____________________

Signature or left thumb impression of Participant Date

______________________________________ ____________________

Signature or left thumb impression of the witness Date

______________________________________ ____________________

Signature of the PI or his/her representative Date

PROTOCOL
Biomedical
Non-Exempt
Berkeley

Protocol # 2011-09-3652
Date Printed: 07/31/2018

---------------------------------------------------------------------------------------

| | |
|---|---|
| **Protocol Title:** | Measuring the benefits of sanitation, water quality, handwashing and nutrition interventions for improving health and development in rural Bangladesh |
| **Protocol Status:** | APPROVED |
| **Date Submitted:** | 10/26/2017 |
| **Approval Period:** | **10/30/2017-11/03/2018** |
| **Important Note:** | **This Print View may not reflect all comments and contingencies for approval. Please check the comments section of the online protocol. Questions that appear to not have been answered may not have been required for this submission. Please see the system application for more details.** |

*** * * Attached Document * * ***

| Document Name | Created Date |
|---|---|
| 2011-09-3652_Colford_2k2-Consent-EnvAssess-water-v19-Bengali.pdf | 10/26/2017 |

#### Appendix 2k-2: English consent form for environmental assessment (drinking water measurements in water arm)

পরিবেশগত প্রভাব মূল্যায়নের জন্য সম্মতিপত্র (খাবার পানি পরিমাপ)

গবেষণার শিরোনাম: বাংলাদেশের গ্রামীণ এলাকায় হাত ধোয়া, পানি বিশুদ্ধকরণ, স্যানিটেশন ওপুষ্টিকর সম্পূরক খাবার প্রদান এবং শিশুর স্বাস্থ্য ও বৃদ্ধির উপর তার প্রভাব।

**প্রধান গবেষক**: ডঃ স্টিফেন পি. লুবি

### গবেষণার উদ্দেশ্যঃ

আসলামুআলাইকুম/আদাব। আমার নাম ----------------------------। আমরা ঢাকা (মহাখালী) কলেরা হাসপাতালে কাজ করি।আমরা শিশুদের ডায়রিয়াজনিত রোগ-বালাই নিয়ে গবেষণা করছি। কতিপয় স্যানিটেশন ব্যবস্থা, পানির গুনাগুন এবং স্বাস্থ্যসম্মত অভ্যাসসমূহ স্বাস্থ্যের উপর কিভাবে প্রভাব ফেলে এই গবেষণার মাধ্যমে আমরা তা বোঝার চেষ্টা করছি।

##### আমরা কেন আপনাকে এই গবেষণায় অংশগ্রহনে আমন্ত্রণ জানাচ্ছি?

আমরা আপনাকে/আপনার খানাকে এই গবেষণায় অংশগ্রহণের জন্য আমন্ত্রন জানাচ্ছি কারণ আপনার ৩ বছরের কম বয়সের একটি শিশু আছে এবং আপনি/আপনার খানা ইতোমধ্যেই আমাদের গবেষণার অন্যান্য কর্মকান্ডে অংশগ্রহন করেছেন। আমরা আপনাকে/আপনাদেরকে এই গবেষণার বিষয়ে আরো বিস্তারিতভাবে জানাতে চাই। আমরা আশা করছি আপনি এবং আপনার শিশু স্বেচ্ছায় এই গবেষণায় অংশগ্রহন করবেন।

##### গবেষণায় অংশগ্রহণকারীর কাছে প্রত্যাশা কী?

যদি আপনি/আপনার খানা এই গবেষণায় অংশগ্রহণে সম্মত থাকেন, তাহলে আমরা আপনার খানাতে নির্দিষ্ট কিছু কর্মকান্ড পরিচালনা করব। আজকে পরিদর্শনকালে আমরা আপনাকে আপনার বাড়ি ও বাড়ির চারপাশের পরিবেশ, এবং আপনার বাড়িতে প্রচলিত অভ্যাস সম্পর্কে কিছু প্রশ্ন জিজ্ঞাসা করব। আপনি আপনার শিশুকে খাওয়ার জন্য যে পানি দেন, সেই পানি থেকে আমাদেরকে এক গ্লাস পানি দিতে অনুরোধ করব। আপনার দেওয়া এই পানি থেকে আমরা নমুনা হিসাবে সামান্য পানি সংগ্রহ করব। পরিদর্শনকালে আমরা আপনার খানাতে প্রায় ১ ঘনটা সময় কাটাব।

### ঝুঁকি এবং সুবিধাসমূহঃ

এই গবেষণায় আপনার এবং আপনার শিশুর ঝুঁকির সম্ভাবনা নুন্যতম। আপনার পরিচয়সংক্রান্ত তথ্যসমূহ গোপন না থাকার সামান্য সম্ভাবনা রয়েছে কিন্তু আমরা আপনার পরিচয়সংক্রান্ত তথ্যসমূহ গোপন রাখার জন্য সম্ভাব্য সবধরনের চেষ্টা করব। কিন্তু আপনার শিশুর অংশগ্রহণ, শিশুদের ডায়রিয়া রোগ সম্পর্কে জ্ঞানলাভে আমাদেরকে সহায়তা করবে।

### নিজস্বতা, গোপনীয়তা ও বিশ্বাসযোগ্যতাঃ

আমরা আপনার/ আপনাদের দেওয়া সমস্ত তথ্যের গোপনীয়তা বজায় রাখার জন্য সর্বোচ্চ চেষ্টা করব। আপনাদের দেওয়া কিছু তথ্য আইসিডিডিআর,বি র গবেষকদের কাছে থাকবে যার ফলে আপনাকে সনাক্ত করা সম্ভব, যেমন আপনার নাম এবং ফোন নম্বর। তারা আপনাদের দেওয়া তথ্য থেকে আপনার সনাক্তকরণ মুলক তথ্য সমুহ বাদ দিয়ে অন্যান্য গবেষকদের সাথে শেয়ার করবে (যেমন আমেরিকার ক্যালিফোর্নিয়া বার্কলে বিশ্ববিদ্যালয়ের গবেষক), যার ফলে কোন গবেষকই আপনাকে চিহ্নিত করতে পারবে না। গবেষণার সব রেকর্ড তালাবদ্ধ করে রাখা হবে ও কমপিউটারে তথ্যাবলী পাসওআরড দিয়ে সংরক্ষন করা হবে ।

অন্যান্য ব্যক্তি অথবা গবেষণার সাথে সংশ্লিষ্ট দল আপনার তথ্য দেখতে পারবে যদি গবেষণার কারণে প্রয়োজন মনে করে। এরা হলেন- আমেরিকার ক্যালিফোর্নিয়া বার্কলে বিশ্ববিদ্যালয়ের গবেষক ও গবেষণার স্পন্সর (বিল ও মেলিন্ডা গেটস ফাউন্ডেশন) । আইসিডিডিআর,বি র গবেষকদের ছাড়া কেউ আপনাকে এবং আপনার শিশুকে সনাক্ত করতে পারবে না । আমরা শুধুমাত্র গবেষণার কাজে এই তথ্য ব্যবহার করব এবং আপনার নাম অথবা পরিচয়ের কোন সূত্র এই গবেষনা-সংক্রান্ত কোন রিপোর্টে/ প্রকাশনায়

উল্লেখ করা হবে না। আপনার কাছ থেকে প্রাপ্ত সকল তথ্য ও নমুনার সর্বোচ্চ গোপনীয়তা রক্ষার জন্য আমরা সর্বাত্মক চেষ্টা করব, তবুও ভুলবশতঃ অনাকাঙ্খিত কোন ঘটনার কারনে গোপনীয়তা রক্ষা নিশ্চিত করতে না পারলে সেজন্য মার্জনা প্রার্থনা করছি।

#### ভবিষ্যতে তথ্যের ব্যবহার:

এই গবেষণায় সংগৃহীত তথ্য প্রয়োজনে আমরা অন্যান্য গবেষকের সাথে বিনিময় করতে পারি। এবং কারো সাথে তথ্য বিনিময় করা হলে আমরা সেখানে আপনার প্রকৃত নাম ব্যবহার করব না এবং আপনার দেওয়া তথ্যের গোপণীয়তা ও নিজস্বতা কঠোরভাবে রক্ষা করব।

#### গবেষণায় অংশগ্রহন না করা বা নাম প্রত্যাহারের অধিকারঃ

এ গবেষণায় আপনার শিশুর অংশগ্রহণ সেচ্ছামূলক এবং আপনার শিশুর অংশগ্রহণ করা বা না করা সম্পূর্ণভাবে আপনার ইচ্ছার উপর নির্ভর করছে। এই গবেষণায় আপনাকে যেসব প্রশ্ন জিজ্ঞাসা করা হবে তার যেকোনটির উত্তর আপনি চাইলে নাও দিতে পারেন। এই গবেষণায় অংশগ্রহনে সম্মত হবার পরেও গবেষণা চলাকালীন যেকোন সময় আপনি আপনার নাম প্রত্যাহার করে নিতে পারেন। এই গবেষণায় অংশগ্রহণ না করার অধিকার আপনার আছে এবং এ জন্য ভবিষ্যতে আপনার পরিবারের সদস্যদের কলেরা হাসপাতালে স্বাস্থ্যসেবা গ্রহণের উপর কোন প্রভাব পড়বে না।

## ক্ষতিপূরণ :

এই গবেষণায় অংশগ্রহণের জন্য আপনার কোন ধরনের অর্থনৈতিক খরচ/ ব্যয় বহন করতে হবে না এবং একইভাবে আপনাকেও আমরা কোন ধরনের অর্থ প্রদান করব না।

## যোগাযোগ:

আপনার কোনো প্রশ্ন থাকলে যেকোন সময় আপনি তা আমাকে জিজ্ঞাসা করতে পারেন। যদি গবেষণা সম্পর্কিত অতিরিক্ত কোন প্রশ্ন থাকে তাহলে আপনি জনাব মাসুদ পারভেজ (প্রোগ্রাম অন ইনফেক্সাস ডিজিজেজ এন্ড ভ্যাকসিন সাইন্সেস, আইসিডিডিআর,বি: মহাখালি, ঢাকা-১২১২) এর সাথে ০১৮১৭৫৪১৮৭২ মোবাইল নম্বরে সরাসরি যোগাযোগ করতে পারেন। এই গবেষণায় আপনার অধিকার-সংক্রান্ত কোন প্রশ্ন থাকলে অথবা গবেষণার কারণে আপনার অর্থনৈতিক কোন ক্ষতি সাধিত হয়েছে বলে মনে করেন, তাহলে আপনি ৯৮৮৬৪৯৮ অথবা ৮৮৬০৫২৩-৩২ এক্স- ৩২০৬ টেলিফোন নম্বরে অথবা সরাসরি এম এ সালাম খান, আইসিডিডিআরবি সেক্রেটারিয়েট এর সাথে যোগাযোগ করতে পারেন।

আপনি যদি এই গবেষণায় অংশগ্রহণ করতে আগ্রহী হন তাহলে নিম্নের নির্ধারিত স্থানে স্বাক্ষর অথবা বাম বৃদ্ধাঙ্গুলীর ছাপ দিন।

আপনার সহযোগিতার জন্য ধন্যবাদ।

________________________ __________________

উত্তরদাতার স্বাক্ষর/বাম বৃদ্ধাঙ্গুলির ছাপ তারিখ

________________________ __________________

স্বাক্ষীর স্বাক্ষর/বাম বৃদ্ধাঙ্গুলির ছাপ তারিখ

________________________ __________________

পি আই/ গবেষকের প্রতিনিধির স্বাক্ষর তারিখ

PROTOCOL
Biomedical
Non-Exempt
Berkeley

Protocol # 2011-09-3652
Date Printed: 07/31/2018

-----------------------------------------------------------------------------------

| | |
|---|---|
| **Protocol Title:** | Measuring the benefits of sanitation, water quality, handwashing and nutrition interventions for improving health and development in rural Bangladesh |
| **Protocol Status:** | APPROVED |
| **Date Submitted:** | 10/26/2017 |
| **Approval Period:** | **10/30/2017-11/03/2018** |
| **Important Note:** | **This Print View may not reflect all comments and contingencies for approval. Please check the comments section of the online protocol. Questions that appear to not have been answered may not have been required for this submission. Please see the system application for more details.** |

*** * * Attached Document * * ***

| Document Name | Created Date |
|---|---|
| 2011-09-3652_Colford_1v-Child cognitive development-v18-English.pdf | 10/26/2017 |

**Appendix 1v: English consent form assessment of child's cognitive development**
Protocol Title: Effect of hand washing, water treatment, sanitation and nutritional supplement interventions on child cognitive development in rural Bangladesh

Principal Investigator's name: Dr. Leanne Unicomb

Protocol Title: Effect of hand washing, water treatment, sanitation and nutritional supplement interventions on child health and development in rural Bangladesh

**Purpose of the research**

Hello/Assalamualaikum/Namaste. My name is ________ and I work with the ICDDR,B (Cholera Hospital) in Dhaka. We are conducting research on diarrheal diseases and through this particular study we want to learn about the health impact of diarrheal diseases in children. We are interested in learning if the exposure of a child to diarrheal disease has long term effects.

**Why are we inviting you to participate in this study?**

We are interested in enrolling your household because you have a child between the age of 8-15 months, and your household has already participated in other parts of this study. Diarrhea is common among children in this age group. We are interested to know how diarrhea affects child development outcomes, such as motor and language skills. Therefore, we hope you will allow your child to participate into this study upon learning more details of participation.

**What is expected from the participants of the research study?**

If you decide to participate, and let your child participate, all of our activities here today will take about one hour. We will ask you some general things about your household, such as things your child likes to play with. We will also ask you to complete three measures that are like games, where we ask you to do some things like remember numbers and words and repeat a story. We will ask you some questions about which words your child can uderstand and speak. We would also like to see what motor skills your child can do, such as sitting, standing and walking. For this motor skill activity, we will try to get your child to perform these various behaviors. This will take about 10-15 minutes to complete. Finally, we will ask you some questions about your well-being.

**Risks & Benefits**

There are no major risks involved in this study. Some of the interview questions may make you uncomfortable, but you are free to decline to answer any questions or stop the interview at any time.

Your child may get tired during the physical motor activity, although it is very brief. To put your child at ease, you and/or a caregiver may sit with them during any of the activities, and you are free to stop the assessments at any time.

There is no anticipated direct benefit to you or your child for participating in this study. However, it is hoped that the information gained from the study will help improve our knowledge of how diarrhea affects young children.

**Privacy, anonymity and confidentiality**

We will do everything we can keep what you tell us confidential as allowed by the law of this country. We will not use your real name or your child's name when we write out the data. All the information we collect will be kept locked. We will use the information only for the purpose of the study. We will not use your name or your child's name when we share and publish the results of this study. We expect the steps we take will keep all of your information confidential, but it is possible that because of mistakes or unforeseen events, it could become compromised.

**Future use of information**
If it is necessary, we may share the information we collected from this study with other researchers. If this is done, we will not use your real name and we will maintain your confidentiality.

**Right not to participate and withdraw**
Taking part in the study is completely voluntary. You may choose not to answer any or all of the questions that will ask, and you can refuse for your child to participate in the motor skills activity. You can drop out of this study at any time, even in the middle of an interview. You have the right to refuse to take part in this study, which will not affect your family's treatment at the Cholera Hospital in the future.

**Principle of compensation**
You do not need to pay us to take part in this study, and we will not pay you money for taking part in the study.

**Persons to contact:**

If you have any question, you can ask me any time. If you have additional questions about these activities, you may contact:

Dr. Fahmida Tofail, Scientist, Child Development Unit (CDU), Dhaka Hospital, ICDDRB, 68 Shaheed Tajuddin Ahmed Sarani, Mohakhali, Dhaka-1212, Bangladesh Phone no. +880 2 9840 523-32; Ext. 2350 Cell: +880 1715 700370

If you have questions about your rights as a participant of a research study, or if you think some harm has been done to you because of this study, you may contact or meet him personally at following address:

M. A. Salam Khan, IRB Secretariat, phone: 9886498 or PABX 8860523-32 ext. 3206

If you agree to our proposal of enrolling your household in our study, please indicate that by putting your signature or your left thumb impression at the specified space below

Thank you for your cooperation

_______________________________________ ____________________

Signature or left thumb impression of Participant Date

_______________________________________ ____________________

Signature or left thumb impression of the witness Date

__________________________________________ ____________________

Signature of the PI or his/her representative Date

---------------------------------------------------------------------------------
---

**Protocol Title:** Measuring the benefits of sanitation, water quality, handwashing and nutrition interventions for improving health and development in rural Bangladesh

**Protocol Status:** APPROVED

**Date Submitted:** 10/26/2017

**Approval Period:** **10/30/2017-11/03/2018**

**Important Note:** **This Print View may not reflect all comments and contingencies for approval. Please check the comments section of the online protocol.**
**Questions that appear to not have been answered may not have been required for this submission. Please see the system application for more details.**

*** * * Attached Document * * ***

| Document Name | Created Date |
|---|---|
| 2011-09-3652_Colford_2v-Child cognitive development-v18-Bengali.pdf | 10/26/2017 |

#### Appendix 2v: Bangla consent form assessment of child's cognitive development

গবেষণার শিরোনাম:বাংলাদেশের গ্রামীণ এলাকায় হাত ধোয়া, পানি বিশুদ্ধকরণ, স্যানিটেশন ওপুষ্টিকর সম্পূরক খাবার প্রদান এবং শিশুর মানসিকবৃদ্ধির উপর তার প্রভাব।

**প্রধান গবেষক**: ড: লিয়েন ইউনিকম্ব

**গবেষণার উদ্দেশ্য:**

আস্‌সালামুআলাইকুম/ নমস্কার। আমার নাম--------------------------------------------------- আমি আইসিডিডিআর,বি (কলেরা হাসপাতাল) ঢাকায় কাজ করি। আমরা ডাইরিয়া নিয়ে একটি গবেষণা করছি এবং আমরা জানতে চাই শিশু স্বাস্থের উপর ডাইরিয়ার কি প্রভাব। আমরা আরও জানতে চাই ডাইরিয়া হলে অদুর ভবিষ্যতে শিশুর কোন সমস্য হয় কিনা।

**কেন আপনাকে এই গবেষণার জন্য অংশগ্রহণ করতে বলা হচ্ছে:**

আমরা আপনার খানাকে এই গবেষণায় অন্তর্ভুক্ত করতে চাই কারন আপনার খানাতে একজন ৮-১৫ মাস বয়সী শিশু রয়েছে এবং আপনি ইতিমধ্যে গবেষণার প্রধান অংশে অন্তর্ভুক্ত হয়েছেন। এই বয়সের বাচ্চাদের জন্য ডাইরিয়া একটি প্রধান রোগ। আমরা জানতে চাই বাচ্চাদের ভাষাগত এবং অংগসঞ্চালন দক্ষতা বিকাশে ডাইরিয়ার কোন বিরূপ প্রভাব আছে কিনা। আমরা আশা করছি আপনি আপনার শিশুকে এই গবেষণায় অংশগ্রহনের অনুমতি দিবেন।

**এই গবেষণায় অংশগ্রহণে আপনার করনীয়:**

আপনি যদি আপনার শিশুকে অংশগ্রহন করার অনুমতি দেন তাহলে গবেষণা সংক্রান্ত সকল কার্যাবলি সম্পন্ন করতে আমাদের আজ ১ ঘন্টা সময় প্রয়োজন হবে। আমরা আপনার খানা সংক্রান্ত কিছু প্রশ্ন করব যেমন অপনার শিশু কি নিয়ে খেলতে চায়। আপনি আপনার শিশুকে কিছু সংখ্যা, অক্ষর এবং একটি গল্প বলবেন যা আপনার শিশু পূনরাবৃত্তি করবে এবং এটি একধরণের খেলার মত হবে। আমরা আপনার শিশু কি ধরণের শব্দ বুঝতে এবং বলতে পারে তা জানতে চাইব। আমরা আরও জানতে চাই আপনার শিশু বসতে, দাড়াতে এবং হাটতে পারে কিনা। সেজন্য আমরা আপনার শিশুকে বসতে, দাড়াতে এবং হাটতে বলব। এর জন্য ১০-১৫ মিনিট সময় লাগবে। অবশেষে আমরা আপনার শিশুর মানসিক এবং শারীরিক সুস্থতাবোধ সম্পর্কে জানতে চাইব।

**ঝুঁকি এবং সুবিধাদি:**

**এই গবেষণায় বড় কোন ঝুঁকি নাই। কিছু প্রশ্নের উত্তর দিতে আপনি স্বাচ্ছন্দ বোধ নাও করতে পারেন। কিন্তু আপনি চাইলে এইসকল প্রশ্নের উত্তর নাওদিতে পারেন।**

**আপনার শিশু বসা, দাড়ানো এবং হাটার সময় সামান্য ক্লান্ত বোধ করতে পারে। আপনি এবং আপনার শিশু যেকোন সময় বসতে বা আরাম করতে পারবেন অথবা অমাকে থামাতে পারবেন।**

**এই গবেষণায় অংশ গ্রহন করলে আপনি ও আপনার শিশু কোন সুবিধা পাবেননা। কিন্তু গবেষনা লব্ধ ফলাফল শিশুর ভাষাগত এবং অংগসঞ্চালন দক্ষতা বিকাশে ডাইরিয়ার প্রভাব সম্পর্কে জ্ঞানার্জনে সহায়তা করবে।**

**নিজস্বতা, গোপনীয়তা ও বিশ্বাসযোগ্যতা:**

এই দেশের আইন অনুযায়ী আপনার দেওয়া সমস্ত তথ্যগোপনীয়তার সাথে রক্ষণাবেক্ষণ করা হবে। আমরা আমাদের গবেষণার কোন লেখায়/ফলাফলে আপনার প্রকৃত নাম ব্যবহার করব না। আমাদের সংগৃহীত সকল তথ্য তালাবদ্ধ থাকবে। আমরা কেবলমাত্র

আমাদের গবেষণার কাজে এসব তথ্য ব্যবহার করব। আমরা এই গবেষণার ফলাফল বিনিময় ও প্রকাশনার ৰেত্রে আপনার নাম ব্যবহার করব না। আমরা আপনার দেওয়া তথ্যসমূহ গোপন রাখার জন্য সম্ভাব্য সবধরনের চেষ্টা করব। তা সত্ত্বেও ভুলবশতঃ অনাকাঙ্খিত কোন ঘটনা ঘটলে সেজন্য মার্জনা প্রার্থনা করছি।

#### ভবিষ্যতে তথ্যের ব্যবহারঃ

এই গবেষণায় সংগৃহীত তথ্য প্রয়োজনে আমরা অন্যান্য গবেষকের সাথে বিনিময় করতে পারি। এবং কারো সাথে তথ্য বিনিময় করা হলে আমরা সেখানে আপনার প্রকৃত নাম ব্যবহার করব না এবং আপনার দেওয়া তথ্যের গোপণীয়তা ও নিজস্বতা কঠোরভাবে রক্ষা করব।

#### গবেষণায় অংশগ্রহন না করা বা নাম প্রত্যাহারের অধিকারঃ

এই গবেষণায় অংশগ্রহন করা সম্পূর্নভাবে আপনার ইচ্ছার উপর নির্ভর করছে। এই গবেষণায় আপনাকে যেসব প্রশ্ন জিজ্ঞাসা করা হবে তার যেকোনটির উত্তর আপনি চাইলে নাও দিতে পারেন। এই গবেষণায় অংশগ্রহনে সম্মত হবার পরেও গবেষণা চলাকালীন যেকোন সময় আপনি আপনার নাম প্রত্যাহার করে নিতে পারেন। এই গবেষণায় অংশগ্রহণ না করার অধিকার আপনার আছে এবং এ জন্য ভবিষ্যতে আপনার পরিবারের সদস্যদের কলেরা হাসপাতালে স্বাস্থ্যসেবা গ্রহণের উপর কোন প্রভাব পড়বে না।

## ক্ষতিপূরণ :

এই গবেষণায় অংশগ্রহণের জন্য আপনার কোন ধরনের অর্থনৈতিক খরচ/ ব্যয় বহন করতে হবে না এবং একইভাবে আপনাকেও আমরা কোন ধরনের অর্থ প্রদান করব না।

## যোগাযোগঃ

আপনার কোনো প্রশ্ন থাকলে যেকোন সময় আপনি তা আমাকে জিজ্ঞাসা করতে পারেন। যদি গবেষণা সম্পর্কিত অতিরিক্ত কোন প্রশ্ন থাকে তাহলে আপনি ডাঃ ফাহমিদা তোফায়েল (চাইল্ড ডেভেলপমেন্ট ইউনিট, আইসিডিডিআর,বিঃ মহাখালি, ঢাকা-১২১২) এর সাথে ০১৭১৫৭০০৩৭০ মোবাইল নম্বরে সরাসরি যোগাযোগ করতে পারেন।

এই গবেষণায় আপনার অধিকার-সংক্রান্ত কোন প্রশ্ন থাকলে অথবা গবেষণার কারণে আপনার অর্থনৈতিক কোন ক্ষতি সাধিত হয়েছে বলে মনে করেন, তাহলে আপনি ৯৮৮৬৪৯৮ অথবা ৮৮৬০৫২৩-৩২ এক্স- ৩২০৬ টেলিফোন নম্বরে অথবা সরাসরি এম এ সালাম খান, আইসিডিডিআরবি সেক্রেটারিয়েট এর সাথে যোগাযোগ করতে পারেন।

আপনি যদি এই গবেষণায় অংশগ্রহণ করতে আগ্রহী হন তাহলে নিম্নের নির্ধারিত স্থানে স্বাক্ষর অথবা বাম বৃদ্ধাঙ্গুলীর ছাপ দিন।

আপনার সহযোগিতার জন্য ধন্যবাদ।

______________________ ______________________

উত্তরদাতার স্বাক্ষর/বাম বৃদ্ধাঙ্গুলির ছাপ তারিখ

______________________ ______________________

স্বাক্ষীর স্বাক্ষর/বাম বৃদ্ধাঙ্গুলির ছাপ তারিখ

______________________ ______________________

পি আই/ গবেষকের প্রতিনিধির স্বাক্ষর তারিখ

PROTOCOL
Biomedical
Non-Exempt
Berkeley

-----------------------------------------------------------------------------------

**Protocol Title:** Measuring the benefits of sanitation, water quality, handwashing and nutrition interventions for improving health and development in rural Bangladesh

**Protocol Status:** APPROVED

**Date Submitted:** 10/26/2017

**Approval Period:** **10/30/2017-11/03/2018**

**Important Note:** **This Print View may not reflect all comments and contingencies for approval. Please check the comments section of the online protocol. Questions that appear to not have been answered may not have been required for this submission. Please see the system application for more details.**

*** * * Attached Document * * ***

| Document Name | Created Date |
|---|---|
| 2011-09-3652_Colford_1w-Consent-Spillover compounds-v19-English.pdf | 10/26/2017 |

**Appendix 1w: English consent form for neighboring compounds in spillover study**

**International Centre for Diarrhoeal Disease Research, Bangladesh (ICDDR,B)**

Protocol Title: Effect of hand washing, water treatment, sanitation and nutritional supplement interventions on child health and development in rural Bangladesh

Principal Investigator's name: Dr. Stephen Luby

**Purpose of the research**

Hello (Assalamualaikum/Nomoshkar). My name is ________ and I work with the ICDDR,B (Cholera Hospital) in Dhaka. We are interested in conducting research on diarrheal diseases in children to learn how it affects their physical and mental development. Through this research we want to learn about the health benefits of some simple interventions to improve the sanitation, water quality, hygiene practices or nutritional status for children under five. We want to understand whether children who live in the same communities as children receiving these interventions benefit from them as well.

**Why are we inviting you to participate in the study?**

Because you live near to recipients of water, sanitation, and hygiene interventions, we are interested in evaluating your child's health, we would like to invite you to participate in this study.

**What is expected from the participants of the research study?**

If you agree to participate, a field research person will visit your household twice for this purpose. Participation in this study will only span a few hours. In the first visit, which we would like to do today, we will ask you questions about your home and compound, and some practices in your home. We will ask you questions about sociodemographic information, your water, sanitation, and hygiene practices and the health of your children under five years old. With your permission, they will inspect your sanitation facilities as well. We will also ask mothers or caregivers about whether your child has been sick.

We will also give your children one toy ball and set up fly traps in the latrine and kitchen areas of your compound. We will ask you to give us a glass of water that you would give to your children to drink. We will take a small amount of the water that you give us.

We will return tomorrow for a second visit, wash the balls in water, keep the water used to wash the toy. Your child may choose to play with the ball or not; this is up to him or her. During this second visit, we will count the flies caught at the fly trap in the latrine and kitchen areas of your compound. We will also ask you a few short questions. Each visit will be less than 1 hour.

We will also collect a stool sample from your children under five years old. We will examine whether your child has any worm infections, but this analysis will be done in Dhaka, and we will not be able to share the results with you. We will offer your child deworming medication regardless of the stool sample result. Today, a field member will deliver a stool collection kit and instruct you how to collect stool from your child. We will ask you to collect your child's stool before the field worker returns tomorrow by having your child defecate on a sheet of provided plastic and use a plastic scoop to collect a small amount of fresh stool from the top of the pile into a container. The field person will collect this container when they come to collect the other specimens.

**Risks & Benefits**

There are minimal risks involved in this study. There is a slight risk of breach of confidentiality. Your child's participation will help us to gain knowledge on diarrheal disease in children.

**Privacy, anonymity and confidentiality**

All data and specimens collected will be kept confidential to the greatest extent possible.. Researchers at ICDDR,B will have access to some of your personal information that could identify you, such as your name. They will remove the identifiable information before sharing it with researchers at partner institutions, such as UC Berkeley. Your research records will be kept in a locked cabinet, and computer-based data, will be stored in an encrypted format on a password-protected server.

Other persons who may have access to your information include research groups that oversee the safety of the study including institutes such as the University of California Berkeley and the Bill and Melinda Gates Foundation, USA. None of the researchers outside of ICDDR,B will be able to identify you or your child from information you provide. We will use the information only for the purpose of the study. We will not use your name when we share and publish the results of this study. We expect the steps we take will keep all of your information confidential, but it is possible that because of mistakes or unforeseen events, it could become compromised.

**Future use of information**

The information collected from this study may be shared with other researchers if needed, but we will strictly maintain your confidentiality and privacy. In the future, we may wish to perform additional tests on the blood and stool specimens that already collected. The samples will be stored at ICDDR,B. No further consent will be taken from you. If you change your mind about having your specimen saved for further testing, you may contact us and the sample will be discarded and not saved.

**Right not to participate and withdraw**

Taking part in the study is completely voluntary. You have the right to decline to allow your child to participate or to withdraw your child at any point in this study without penalty or loss of benefits to which you are otherwise entitled. The members of the households may choose not to answer any or all of the questions asked. You can drop out of this study at any time, even in the middle of an interview or sample collection. If you wish, we will withdraw your permission and we will destroy your samples and those of your child and remove data from the database. You have the right to refuse participation in this study, which will not affect your family's treatment at the Cholera Hospital in the future.

**Principle of compensation**

You need not pay us to take part in this study, and similarly we will not pay you money for attending in the study.

**Persons to contact:**

If you have any question, you can ask me any time. If you have additional questions about the survey, you may contact:

Abu Naser, Center of Communicable Diseases, ICDDR,B, Mohakhali, Dhaka 1212. Phone: 8860523-32 # 120

If you have questions about your rights as a participant of a research study, or if you think some harm has been done to you because of the survey, you may contact or meet him personally at following address:

M. A. Salam Khan, IRB Secretariat, phone: 9886498 or PABX 8860523-32 ext. 3206

If you agree to our proposal of enrolling your household in our study, please indicate that by putting your signature or your left thumb impression at the specified space below

Thank you for your cooperation

Consent to enroll into the study: YES______ NO______

Consent of the mother for sample collection: YES_______ NO ______

________________________________________ _____________________

Signature or left thumb impression of Participant Date

________________________________________ _____________________

Signature or left thumb impression of the witness Date

________________________________________ ____________________

Signature of the PI or his/her representative Date

PROTOCOL
Biomedical
Non-Exempt
Berkeley

---------------------------------------------------------------------------------
---

**Protocol Title:** Measuring the benefits of sanitation, water quality, handwashing and nutrition interventions for improving health and development in rural Bangladesh

**Protocol Status:** APPROVED

**Date Submitted:** 10/26/2017

**Approval Period:** **10/30/2017-11/03/2018**

**Important Note:** **This Print View may not reflect all comments and contingencies for approval. Please check the comments section of the online protocol.
Questions that appear to not have been answered may not have been required for this submission. Please see the system application for more details.**

*** * * Attached Document * * ***

| Document Name | Created Date |
|---|---|
| 2011-09-3652_Colford_2w-Consent-Spillover compounds-v19-Bengali.pdf | 10/26/2017 |

#### Appendix 2w: Bengali consent form for neighboring compounds in spillover study

### আন্ৰর্জাতিক উদারাময় গবেষণা কেন্দ্র বাংলাদেশ (আই সি ডি ডি আর, বি )

**গবেষণার শিরোনাম:** বাংলাদেশের গ্রামীণ এলাকায় হাত ধোয়া, পানি বিশুদ্ধকরণ, স্যানিটেশন এবং সম্পূরক খাবার প্রদান এবং শিশুর স্বাস্থ্য ও বৃদ্ধির উপর তার প্রভাব

**প্রধান গবেষকের নাম:** ড: লিয়েন ইউনিকম্ব

**গবেষণার উদ্দেশ্য:**

আস্‌সালামুআলাইকুম। আমার নাম ( ___________ ) এবং আমি আইসিডিডিআর, বি (কলেরা হাসপাতাল) তে কাজ করি। আমরা শিশুদের ডায়রিয়া রোগ এবং ইহা কিভাবে তাদের শারীরিক ও মানসিক বৃদ্ধিকে প্রভাবিত করে তা নিয়ে গবেষণা করতে আগ্রহী। আমরা এই গবেষণার মাধ্যমে কিছু সাধারন ইনটারভেনশন যেমন- পায়খানার উন্নয়ন, পানির গুনগতমান এবং স্বাস্থ্যসম্মত অভ্যাস বা পাঁচ বছরের ছোট বাচ্চাদের পুষ্টিগত অবস্থা ইত্যাদির উন্নয়নের মাধ্যমে স্বাস্থ্যগত সুবিধা সম্পর্কে জানতে চাচ্ছি। একই সমাজের/এলাকার শিশুরা যারা এই ইনটাভেনশনগুলো পেয়েছে এবং যারা পায় নাই তাদের পার্থক্য বুঝার চেষ্টা করছি

**আমরা কেন আপনাকে এই গবেষণায় অংশগ্রহনে আমন্ত্রণ জানাচ্ছি?**

যেহেতু আপনারা স্বাস্থ্যস্মত অভ্যাস, পানি বিশুদ্ধকরন ও স্যানিটেশন ইনটারভেনশনের সবচেয়ে নিকটবর্তী স্থানে বসবাস করেন তাই আমরা আপনার শিশুর স্বাস্থ্যের মূল্যায়ন করতে আগ্রহী আমরা। আমরা আপনাকে এই গবেষণায় অংশগ্রহণের জন্য আমন্ত্রন জানাচ্ছি।

**গবেষণায় অংশগ্রহণকারীর কাছে প্রত্যাশা কী ?**

আপনি যদি অংশগ্রহণ করতে রাজি থাকেন তবে একজন মাঠ গবেষনাকারী এই উদ্দেশ্যে আপনার বাড়িতে দুইবার পরিদর্শনে আসবে। এই গবেষনায় অংশগ্রহণ করলে আপনার এক-দুই ঘনটা সময় ব্যয় হবে। প্রথম পরিদর্শনে আজকে যা করতে ইচ্ছুক , তাতে আমরা আপনার ঘর.বাড়ি ও আপনার ঘরের কিছু অভ্যাসের সর্ম্পকে কিছু প্রশ্ন জিঙ্গাসা করব। আমরা আপনাকে কিছু প্রারম্ভিক তথ্য , আপনার পানি, স্যানিটেশন এবং স্বাস্থ্যসম্মত অভ্যাস এবং পাঁচ বছরের নিচের শিশুদেও সর্ম্পকে কিছু প্রশ্ন জিঙ্গাসা করব। আপনার অনুমতি ক্রমে ,আমরা আপনার স্যানিটেশন সুবিধাগুলো পযবেক্ষন করব। আমরা মা অথবা শিশুর পরিচর্যাকারীদের তাদের শিশুরা অসুস্থ হয়েছিল কিনা সে সম্পর্কে জিঙ্গাসা করব।

আমরা আপনার শিশুকে ১টা খেলনার বল দিব এবং আপনার বাড়ির পায়খানা ও রান্নাঘরের আশেপাশে মাছি ধরার ফাঁদ বাধঁব/সেট করব। আপনি আপনার শিশুকে খাওয়ার জন্য যে পানি দেন, সেই পানি থেকে আমাদেরকে একগস্নাস পানি দিতে অনুরোধ করব। আপনার দেওয়া এই পানি থেকে আমরা নমুনা হিসাবে সামান্য পানি সংগ্রহ করব।
দ্বিতীয়বার পরিদর্শনের জন্য আমরা আগামীকাল আবার আপনার বাড়িতে আসব। আমরা পানি দিয়ে আপনার শিশুকে দেওয়া খেলনা বলগুলো ধোব/ধৌত করব এবং বল পানিটুকু সংরৰন করব। আপনার শিশু খেলনা বলগুলো দিয়ে খেলতেও পারে আবার নাও খেলতে পারে। এটা সম্পূর্ন তার ইচ্ছার উপর নির্ভর করে। দ্বিতীয় পরিদর্শনের সময় আমরা আপনার বাড়ির পায়খানা ও রান্নাঘরের আশেপাশে বেধেঁ রাখা ফাঁদে/ ফ্লাই ট্রেপে আটকা পড়া মাছির সংখ্যা গননা করব। আমরা আপনাকে কিছু প্রশ্নও জিজ্ঞাসা করব। প্রত্যেকবার পরিদর্শনকালে আমরা আপনার খানাতে প্রায় ১ ঘনটা সময় কাটাব। আমরা আরও আপনার পাঁচ বছরের নিচের শিশুদেও পায়খানার নমুনা সংগ্রহ করব। আমরা আপনাদের শিশুদের কোন কৃমির সংক্রমন আছে কিনা তা পরীক্ষা করব কিন্তু এই পরীক্ষা ঢাকায় করা হবে এবং এর ফলাফলা জানাতে পারব না। পায়খানার নমুনার পরীক্ষার ফলাফল যাই হোক না কেন আমরা আপনাদেরকে কৃমির ঔষধ প্রদান করব।
একজন মাঠকর্মী আপনাকে পায়খানা সংগ্রহ করার সামগ্রী দিয়ে যাবে এবং কিভাবে আপনার শিশুর পায়খানা সংগ্রহ করতে হবে তা দেখিয়ে দেবে। পরের দিন সকালে যদি মাঠকর্মী পৌছানোর পূর্বে শিশু পায়খানা করে, তবে আপনি আপনার শিশুর পায়খানা সংগ্রহ করবেন। পায়খানা সংগ্রহ করার জন্য আপনাকে একটি প্লাষ্টিক শিট দেয়া হবে, যেখানে শিশু পায়খানা করবে এবং আপনি একটি প্লাষ্টিক (ছেনি) স্কুপ ব্যবহার করে উপর থেকে অল্প একটু সদ্য (এইমাত্র করা) পায়খানা বোতলে ভরবেন। আমাদের মাঠকর্মী যখন অন্যান্য নমুনা সংগ্রহ করার জন্য আসবে, তখন এই বোতলটি আপনার কাছ থেকে নিয়ে নেবে।

## ঝুঁকি এবং সুবিধা

এই গবেষণায় অংশগ্রহনে আপনার এবং আপনার শিশুর ঝুঁকির সম্ভাবনা নুন্যতম। আপনার পরিচয়সংক্রান্ত তথ্যসমূহ গোপন না থাকার সামান্য সম্ভাবনা রয়েছে। কিন্তু আপনার শিশুর অংশগ্রহণ, শিশুদের ডায়রিয়া রোগ সম্পর্কে জ্ঞানলাভে আমাদেরকে সহায়তা করবে।

## নিজস্বতা, গোপনীয়তা ও বিশ্বাসযোগ্যতা:

আমরা আপনার/ আপনাদের দেওয়া সমস্ত তথ্যের গোপনীয়তা বজায় রাখার জন্য সর্বোচ্চ চেষ্টা করব। আপনাদের দেওয়া কিছু তথ্য আইসিডিডিআর,বি র গবেষকদের কাছে থাকবে যার ফলে আপনাকে সনাক্ত করা সম্ভব, যেমন আপনার নাম এবং ফোন নম্বর। তারা আপনাদের দেওয়া তথ্য থেকে আপনার সনাক্তকরণ মুলক তথ্য সমুহ বাদ দিয়ে অন্যান্য গবেষকদের সাথে শেয়ার করবে (যেমন আমেরিকার ক্যালিফোর্নিয়া বার্কলে বিশ্ববিদ্যালয়ের গবেষক), যার ফলে কোন গবেষকই আপনাকে চিহ্নিত করতে পারবে না। গবেষণার সব রেকর্ড তালাবদ্ধ করে রাখা হবে ও কমপিউটারে তথ্যাবলী পাসওআরড দিয়ে সংরক্ষন করা হবে।

অন্যান্য ব্যক্তি অথবা গবেষণার সাথে সংশ্লিষ্ট দল আপনার তথ্য দেখতে পারবে যদি গবেষণার কারণে প্রয়োজন মনে করে। এরা হলেন- আমেরিকার ক্যালিফোর্নিয়া বার্কলে বিশ্ববিদ্যালয়ের গবেষক ও গবেষণার স্পন্সর (বিল ও মেলিন্ডা গেটস ফাউন্ডেশন)। আইসিডিডিআর,বি র গবেষকদের ছাড়া কেউ আপনাকে এবং আপনার শিশুকে সনাক্ত করতে পারবে না। আমরা শুধুমাত্র গবেষণার কাজে এই তথ্য ব্যবহার করব এবং আপনার নাম অথবা পরিচয়ের কোন সূত্র এই গবেষনা-সংক্রান্ত কোন রির্পোট/ প্রকাশনায় উল্লেখ করা হবে না। আপনার কাছ থেকে প্রাপ্ত সকল তথ্য ও নমুনার সর্বোচ্চ গোপনীয়তা রক্ষার জন্য আমরা সর্বাত্মক চেষ্টা করব, তবুও ভুলবশতঃ অনাকাঙ্খিত কোন ঘটনার কারনে গোপনীয়তা রক্ষা নিশ্চিত করতে না পারলে সেজন্য মার্জনা প্রার্থনা করছি।

#### ভবিষ্যতে তথ্যের ব্যবহার:

এই গবেষণায় সংগৃহীত তথ্য প্রয়োজনে অন্যান্য গবেষকের সাথে বিনিময় হতে পারে, কিন্তু তথ্যের গোপণীয়তা ও নিজস্বতা কঠোরভাবে রক্ষা করা হবে। ভবিষ্যতে, সংগ্রহকৃত রক্ত ও পায়খানার নমুণার অতিরিক্ত কোন পরীক্ষা করতে পারি। এই নমুণা আইসিডিডিআর,বি তে ২০ বছর পর্যন্ত সংরক্ষণ করা হতে পারে যাতে করে আমরা আপনার কোন সংক্রমন ছিল কিনা তা জানার জন্য এবং আপনার শরীরে কোন কোন জীবাণু বসবাস করত তা বের করার জন্য উন্নত ধরনের পরীক্ষা করতে পারি। এর জন্য পরবর্তীতে আপনার কাছ থেকে কোন অনুমতি নেওয়া হবে না এবং আপনি এই পরীক্ষার কোন ফলাফলও পাবেন না। আইসিডিডিআর,বি একজন সিনিয়র বিজ্ঞানী ঠিক করবেন কে সংরক্ষণকৃত নমূণা ব্যবহার করতে পারবে এবং কোন কোন পরীক্ষা করতে পারবে। এই নমুনা ভবিষ্যতে পরীক্ষার জন্য রেখে দেওয়ার ব্যপারে যদি আপনি কখনও আপনার মতামত পরিবর্তন করেন তাহলে আপনি আমাদের সাথে যোগাযোগ করতে পারেন এবং আমরা আপনার নমুণা ফেলে দিব।

#### স্বেচ্ছা অংশগ্রহণ :

এ গবেষণায় আপনার শিশুর অংশগ্রহণ সেচ্ছামূলক এবং আপনার শিশুর অংশগ্রহণ করা বা না করা সম্পূর্ণভাবে আপনার ইচ্ছার উপর নির্ভর করছে। আপনি ইচ্ছা করলে যে কোন সময়ে আপনার অনুমতি ফিরিয়ে নিতে পারবেন এবং তাহলে আমরা আপনার ও আপনার শিশুর কাছ থেকে নেওয়া নমুণা নষ্ট করে ফেলব এবং আপনার দেওয়া সকল তথ্যও মুছে ফেলব। এই গবেষণায় অংশগ্রহণ না করার অধিকার আপনার আছে এবং এ জন্য ভবিষ্যতে আপনার পরিবারের সদস্যদের কলেরা হাসপাতালে স্বাস্থ্যসেবা গ্রহণের উপর কোন প্রভাব পড়বে না।

## ক্ষতিপূরণ :

এই গবেষণায় অংশগ্রহণের জন্য আপনার সরাসরি কোন ধরনের অর্থনৈতিক খরচ/ ব্যয় হবে না এবং একইভাবে আপনি এই গবেষণায় অংশগ্রহনের জন্য সরাসরি কোন ধরণের অর্থনৈতিক সহায়তাও পাবেন না।

## যোগাযোগ:

আপনার কোনো প্রশ্ন থাকলে আমাকে জিজ্ঞাসা করতে পারেন। যদি গবেষণা সম্পর্কিত অতিরিক্ত কোন প্রশ্ন থাকে তাহলে আপনি আবু নাসের , (সেন্টার ফর কমিউনিকেবল ডিজিজ, আইসিডিডিআর,বি: মহাখালি, ঢাকা-১২১২) এর সাথে ৮৮১৯৪১৯-২০, ৮৮৬০৫২৩-৩২ (এক্স-১২০) টেলিফোন নম্বরে সরাসরি যোগাযোগ করতে পারেন। এই গবেষণায় আপনার অধিকার-সংক্রান্ত কোন প্রশ্ন থাকলে অথবা গবেষনার কারণে আপনার কোন ক্ষতি হতে পারে বলে যদি

মনে করেন, তাহলে আপনি ৯৮৮৬৪৯৮ অথবা ৯৮৪০৫২৩ (এক্স- ৩২০৬) টেলিফোন নম্বরে অথবা সরাসরি এম এ সালাম খান, আইসিডিডিআরবি সেক্রেটারিয়েট এর সাথে যোগাযোগ করতে পারেন।

আপনি যদি এই গবেষণায় অংশগ্রহণ করতে আগ্রহী হন তাহলে নিম্নের নির্ধারিত স্থানে স্বাক্ষর অথবা বাম বৃদ্ধাঙ্গুলীর ছাপ দিন।

আপনার সহযোগিতার জন্য ধন্যবাদ।

মা নমুণা দিতে রাজী হয়েছেঃ হ্যাঁ _______ না _______

........................................... ..................................

উত্তরদাতার স্বাক্ষর/বাম বৃদ্ধাঙ্গুলির ছাপ তারিখ

........................................... ..................................

স্বাক্ষীর স্বাক্ষর/বাম বৃদ্ধাঙ্গুলির ছাপ তারিখ

........................................... ..................................

পি আই/ গবেষকের প্রতিনিধির স্বাক্ষর তারিখ

PROTOCOL
Biomedical
Non-Exempt
Berkeley

---------------------------------------------------------------------------------
---

**Protocol Title:** Measuring the benefits of sanitation, water quality, handwashing and nutrition interventions for improving health and development in rural Bangladesh

**Protocol Status:** APPROVED

**Date Submitted:** 10/26/2017

**Approval Period:** **10/30/2017-11/03/2018**

**Important Note:** **This Print View may not reflect all comments and contingencies for approval. Please check the comments section of the online protocol. Questions that appear to not have been answered may not have been required for this submission. Please see the system application for more details.**

*** * * Attached Document * * ***

| Document Name | Created Date |
|---|---|
| WASH Benefits Bangladesh Spillover recruitment script.pdf | 10/26/2017 |

**WASH Benefits Bangladesh (CPHS # 2011-09-3652)**
**Recruitment Script for Spillover Compounds**

Hello (Assalamualaikum/Nomoshkar). My name is ________ and I work with the ICDDR,B (Cholera Hospital) in Dhaka. We are interested in conducting research on diarrheal diseases in children to learn how it affects their physical and mental development. We are looking to invite compounds with at least one child under five years old to participate in this research. Are there any children under five years old who live in this household?

If YES → Proceed with informed consent process.

If NO → Ok, we will try another compound. Thank you for your time.

-------------------------------------------------------------------------------------

| | |
|---|---|
| **Protocol Title:** | Measuring the benefits of sanitation, water quality, handwashing and nutrition interventions for improving health and development in rural Bangladesh |
| **Protocol Status:** | APPROVED |
| **Date Submitted:** | 10/26/2017 |
| **Approval Period:** | **10/30/2017-11/03/2018** |
| **Important Note:** | This Print View may not reflect all comments and contingencies for approval. Please check the comments section of the online protocol. Questions that appear to not have been answered may not have been required for this submission. Please see the system application for more details. |

*** * * Attached Document * * ***

| Document Name | Created Date |
|---|---|
| 2011-09-3652_IIA_Wagner.pdf | 10/26/2017 |

### Institutional Review Board (IRB)/Independent Ethics Committee (IEC) Authorization Agreement

**Name of Institution or Organization Providing IRB Review** (Institution/Organization A):
_*University of California, Berkeley* ______________________________

UCB IRB Registration #: *IRB00000455 & IRB00005610* UCB FWA #: *FWA00006252*

**Name of Institution Relying on the Designated IRB** (Institution B): *Wagner College*

FWA #: *None*

The Officials signing below agree that (*Name of Institution B)* may rely on the designated IRB for review and continuing oversight of its human subjects research described below.

This agreement is limited to the following specific protocol(s):

Name of UCB Principal Investigator: *John M. Colford, Jr., M.D., Ph.D.*

Name of Investigator at Relying Institution: *Mohammed Alauddin, Ph.D.*

Name of Research Project(s): *Measuring the benefits of sanitation, water quality, handwashing and nutrition interventions for improving health and development in rural Bangladesh*

ePrototcol #(s): *2011-09-3652*

Sponsor or Funding Agency: *Bill & Melinda Gates Foundation, National Institutes of Health, Stanford University*

Award Number(s), if any: *20130310, 028064-002, 20131960, 20141262, 20140633*

The review performed by the designated IRB will meet the human subject protection requirements of Institution B's OHRP-approved FWA. The IRB at Institution/Organization A will follow written procedures for reporting its findings and actions to the Human Protections Administrator at Institution B. Relevant minutes of IRB meetings will be made available to Institution B upon request. Institution B remains responsible for ensuring compliance with the IRB's determinations and with the Terms of its OHRP-approved FWA. This document must be kept on file by both parties and provided to OHRP upon request.

Signature of Signatory Official (Institution/Organization A):

[signature] Date: 7/18/14

Print Full Name: *Graham Fleming* Institutional Title: *Vice Chancellor for Research*

Signature of Signatory Official (Institution B):

[signature] Date: 7/16/2014

Print Full Name: *William E. Mea* Institutional Title: *Vice President for Finance & Business*

------------------------------------------------------------------------------------
---

**Protocol Title:** Measuring the benefits of sanitation, water quality, handwashing and nutrition interventions for improving health and development in rural Bangladesh

**Protocol Status:** APPROVED

**Date Submitted:** 10/26/2017

**Approval Period:** **10/30/2017-11/03/2018**

**Important Note:** **This Print View may not reflect all comments and contingencies for approval. Please check the comments section of the online protocol.**
**Questions that appear to not have been answered may not have been required for this submission. Please see the system application for more details.**

*** * * Attached Document * * ***

| Document Name | Created Date |
|---|---|
| 2011-09-3652_Colford_1j-Consent-EE-v21-English.pdf | 10/26/2017 |

**Appendix 1j: English consent form for environmental enteropathy and parasitic assessment**

Protocol Title: Effect of hand washing, water treatment, sanitation and nutritional supplement interventions on child health and development in rural Bangladesh

Principal Investigator's name: Dr. Leanne Unicomb

**Purpose of the research**

Hello/Assalamualaikum/Namaste. My name is ________ and I work with the ICDDR,B (Cholera Hospital) in Dhaka. We are conducting research on diarrheal diseases and through this particular study we want to learn about the health impact of diarrheal diseases in children. Malnutrition in children is associated with infections of our intestines that can change our use of nutrients. It is possible that genes in our body, made up of DNA, make some children more susceptible to malnutrition than others. This DNA is inherited from our parents and we also carry genes from many friendly microbes that live on and in our body. We are interested in learning if the exposure of a child to diarrheal disease has long term effects. We also are interested in learning how certain foods, nutrition, or life experiences in general may interact with diarrhea or other illnesses.

**Why are we inviting you to participate in the study?**

We are interested in enrolling this compound because we collected information on your household earlier and we are interested in conducting additional testing to evaluate your child's digestive tract. We are interested in evaluating your child's blood, stool, saliva, hair and urine for markers of nutritional status, infections and health. We are also interested in evaluating your saliva, hair, blood, and urine for markers of infections, stress and health.

**What is expected from the participants of the research study?**

Participation in this study will only span a few hours. To achieve the aim of the project, if you agree to be in this study, we will collect a blood, stool, and urine sample from your child. With your permission, we might also collect 5 saliva samples and 3-4 strands of hair from your child and a urine sample, a blood sample, 4 saliva samples, and 3-4 strands of hair from you. From the blood sample, we will measure nutritional markers, indicators of factors children inherit from their parents, and we will be able to understand whether your child has been exposed to infection. The urine sample will help us understand whether there has been a long term physical effect as a result of diarrhea. We will also collect a stool sample from your child with your help.

If you agree to participate, a field research person will visit your household twice for this purpose. Today a field member will weigh the child using a scale, measure his/her height using a height board, measure arm and head circumference using a tape measure and collect general health measures of blood pressure, sweat, and heart rate. The blood pressure and heart rate results from your child will be provided to you after the testing. The field representative will also collect general health measures of blood pressure, sweat, and heart rate. Your blood pressure and heart rate results will be provided to you after the testing. The field representative will also cut

and collect 3-4 strands of hair from you and your child. With your permission, we will collect a small amount of blood (equivalent to 1.5 teaspoons) from a vein in your child's arm to determine their blood group and genes (DNA) and test for infections that they may have had in the past, and measure their nutritional status. One drop of blood will be used to test your child for anemia, a condition of lower than normal red blood cells. The field member can provide you with the results of this test during their visit. We will record how your child responds to these procedures. If you agree, we would like to videotape your child during the blood-draw. We will use this information to better understand how these procedures affect child behavior. We will also ask you about how your child reacts to new situations, and what helps your child feel comfortable in new situations, which will take 10-15 minutes. This will help us understand your child's reaction to the different procedures we are administering for this project. We might also collect a small amount (10ml) of blood from your vein. We might also collect 3 saliva samples each from both you and your child before, during, and after the blood draw using a soft sponge placed under the tongue. The field member will deliver a stool collection kit and instruct you how to collect stool from your child. You will be instructed to collect your child's stool on the following morning, if the child defecates before their arrival, by having your child defecate in a clean diaper and use a plastic scoop to collect a small amount of fresh stool from the top of the pile into a container. The field person will collect this container when they come to collect the other specimens on the following day. Total participation time today will be approximately 3 hours and 15 minutes.

Tomorrow, we will ask you to collect your first urine sample of the morning immediately after you wake up. The main procedure will involve a second saliva collection, collecting your child's urine for 1 hour, feeding your child sugar syrup and then collecting their urine sample over a period of 5 hours. We intend to test the urine for the syrup to help us understand the health impacts of diarrhea in children. For a urine sample, due to the age of your child, we will be using a special urine collection bag to collect the urine from your child and we will demonstrate how it is used.

You/the mother will be requested to not feed your child for at least one hour before we feed him/her the syrup. During this fasting period, we will collect your child's urine for 1 hour by attaching the urine collection bag with a drainage tube (show sample) to the child. We will also collect two additional saliva samples from your child and one additional saliva sample from you using a soft sponge. We will then give a dose of the sugar syrup to the child and attach the urine collection bag with a drainage tube (show sample) to the child immediately after feeding him/her the syrup. We will encourage the child to drink water 30 minutes after taking the syrup to help urination. The field representative will remove the urine from the bag, whenever the child urinates. This collection will take place for 6 hours after which the bag will be removed from the child. During the 6-hour period of urine collection, you will be asked the quantities of foods (and the ingredients) you fed the child in the previous day and night. You will also be asked about your personal life experiences and health. You will be asked about perceptions regarding social norms that may affect a child's health. From the stool sample, we will examine whether your child has any worm infections, but this analysis will be done in Dhaka. We will offer your child deworming medication regardless of the stool sample result. Total participation time tomorrow will be approximately 7 hours.

Later, at the laboratory, we will measure your blood, saliva, hair, and urine samples and your child's blood, stool, saliva, hair, and urine samples for markers of nutritional status, infections, and health. You will not receive the results of any laboratory tests.

**Study time**: Study participation will take a total of approximately *10 hours and 15 minutes over 2 days.*

**Risks & Benefits**

Although we will try to protect your identity there is some possibility that confidentiality could be compromised. The syrup is a natural sugar solution that tastes pleasant. The blood will be collected by a trained professional. You and your child may feel some momentary pain during the blood collection. Your child may also feel some discomfort due to the presence of urine collection bag for 6 hours. Some of the questions I would like to ask you may seem private or personal since they touch on your life and health. You may feel uncomfortable talking about some of the topics. You do not have to answer any question or take part in the discussion/interview/survey if you don't wish to do so, and that is also fine. You do not have to give us any reason for not responding to any question, or for refusing to take part in the interview. All your answers will be kept as confidential as possible, and we anticipate that the risks from participating in this survey will be very minimal. There is no direct benefit for participating in this study, but your child's participation will help us to gain knowledge on the health impact of diarrheal diseases and how certain foods, nutrition, and life experiences in general may interact with diarrhea or other illnesses.

**Confidentiality**

All data and specimens collected will be kept confidential as allowed by the law of this country. The samples and the test results will be coded without your name or your child's name or personal information, and stored separately for analysis by the researchers. None of these researchers will be able to identify you or your child as the sample donors. Other persons who may have access to your test results include research groups that oversee the safety of the study including institutes such as the University of California Berkeley and the Bill and Melinda Gates Foundation, USA. Because of the potential need to release information to these parties, and the genetic information we are collecting is potentially identifying, we cannot guarantee absolute confidentiality. However, we will ensure every effort that we can to preserve your confidentiality. There is no expiration date to this permission. This information will be coded and it is unlikely that anyone will be able to trace it to you and your child, protecting your confidentiality.

All videotapes will be identified by a number only; no recordings will identify you or your child by name. These will be viewed only by trained personnel for coding of your child's response to the procedure. The videos will not be viewed by any other person. The videos will be stored in a locked cabinet accessible by study personnel only. These will never be made available to any persons not participating in the study.

**Future use of information**

The blood, urine, saliva, hair, and stool samples may be stored until the end of the study, so they can be analyzed in the lab at the same time. If you agree, the information, along with some of the blood, urine, saliva, hair and stool collected will be stored at ICDDR,B for up to 20 years so that we can conduct advanced tests on these samples to understand the infections that you may have had, and to find out what microbes lived in your body. No further consent will be taken from you, and you will not receive the results of these tests. A senior scientist at ICDDR,B will decide who can use the stored samples, and what tests will be done. If you change your mind about having your specimen saved for further testing, you may contact us and the sample will be discarded and not saved. The videotapes will be stored indefinitely.

**Right not to participate and withdraw**

Taking part in the study is completely voluntary. You may choose not to allow your child to participate in this study. You can drop out of this study at any time, even in the middle of the sample/urine collection. You have the right to refuse participation in this study, which will not affect your family's treatment at the Cholera Hospital in the future.

**Principle of compensation**

You need not pay us to take part in this study, and similarly we will not pay you money for attending in the study.

**Persons to contact:**

If you have any questions, you can ask me any time. If you have additional questions about the survey, you may contact:

If you have questions about your rights as a participant of a research study, or if you think some harm has been done to you because of the survey, you may contact or meet IRB Secretariat, M.A. Salam Khan:

M. A. Salam Khan, IRB Secretariat, phone: 9886498 or PABX 8860523-32 ext. 3206

If you agree to participate and you allow your child to participate, please indicate that by checking the boxes of the activities that you will agree to below:

**Urine Collection |__| mother |__| child**

**Venous Blood Collection |__| mother |__| child**

**Stool Collection |__| child**

**Saliva sample collection |__| mother |__| child**

**Hair sample collection |__| mother |__| child**

**Urine Long-term storage |__| mother |__| child**

**Venous Blood Long-term storage |__| mother |__| child**

**Stool Long-term storage |__| child**

**Saliva sample Long-term storage |__| mother |__| child**

**Hair sample Long-term storage |__| mother |__| child**

If you agree to our proposal of enrolling your household in our study, please indicate that by putting your signature or your left thumb impression at the specified space below

Thank you for your cooperation

| | |
|---|---|
| ______________________________ | ________________ |
| Signature or left thumb impression of Guardian | Date |
| ______________________________ | ________________ |
| Signature or left thumb impression of the witness | Date |
| ______________________________ | ________________ |
| Signature of the PI or his/her representative | Date |

PROTOCOL
Biomedical
Non-Exempt
Berkeley

Protocol # 2011-09-3652
Date Printed: 07/31/2018

---------------------------------------------------------------------------------
---

| | |
|---|---|
| **Protocol Title:** | Measuring the benefits of sanitation, water quality, handwashing and nutrition interventions for improving health and development in rural Bangladesh |
| **Protocol Status:** | APPROVED |
| **Date Submitted:** | 10/26/2017 |
| **Approval Period:** | **10/30/2017-11/03/2018** |
| **Important Note:** | **This Print View may not reflect all comments and contingencies for approval. Please check the comments section of the online protocol. Questions that appear to not have been answered may not have been required for this submission. Please see the system application for more details.** |

*** * * Attached Document * * ***

| Document Name | Created Date |
|---|---|
| 2011-09-3652_Colford_1j-Consent-EE-v21-English.pdf | 10/26/2017 |

**Appendix 1j: English consent form for environmental enteropathy and parasitic assessment**

Protocol Title: Effect of hand washing, water treatment, sanitation and nutritional supplement interventions on child health and development in rural Bangladesh

Principal Investigator's name: Dr. Leanne Unicomb

**Purpose of the research**

Hello/Assalamualaikum/Namaste. My name is ________ and I work with the ICDDR,B (Cholera Hospital) in Dhaka. We are conducting research on diarrheal diseases and through this particular study we want to learn about the health impact of diarrheal diseases in children. Malnutrition in children is associated with infections of our intestines that can change our use of nutrients. It is possible that genes in our body, made up of DNA, make some children more susceptible to malnutrition than others. This DNA is inherited from our parents and we also carry genes from many friendly microbes that live on and in our body. We are interested in learning if the exposure of a child to diarrheal disease has long term effects. We also are interested in learning how certain foods, nutrition, or life experiences in general may interact with diarrhea or other illnesses.

**Why are we inviting you to participate in the study?**

We are interested in enrolling this compound because we collected information on your household earlier and we are interested in conducting additional testing to evaluate your child's digestive tract. We are interested in evaluating your child's blood, stool, saliva, hair and urine for markers of nutritional status, infections and health. We are also interested in evaluating your saliva, hair, blood, and urine for markers of infections, stress and health.

**What is expected from the participants of the research study?**

Participation in this study will only span a few hours. To achieve the aim of the project, if you agree to be in this study, we will collect a blood, stool, and urine sample from your child. With your permission, we might also collect 5 saliva samples and 3-4 strands of hair from your child and a urine sample, a blood sample, 4 saliva samples, and 3-4 strands of hair from you. From the blood sample, we will measure nutritional markers, indicators of factors children inherit from their parents, and we will be able to understand whether your child has been exposed to infection. The urine sample will help us understand whether there has been a long term physical effect as a result of diarrhea. We will also collect a stool sample from your child with your help.

If you agree to participate, a field research person will visit your household twice for this purpose. Today a field member will weigh the child using a scale, measure his/her height using a height board, measure arm and head circumference using a tape measure and collect general health measures of blood pressure, sweat, and heart rate. The blood pressure and heart rate results from your child will be provided to you after the testing. The field representative will also collect general health measures of blood pressure, sweat, and heart rate. Your blood pressure and heart rate results will be provided to you after the testing. The field representative will also cut

and collect 3-4 strands of hair from you and your child. With your permission, we will collect a small amount of blood (equivalent to 1.5 teaspoons) from a vein in your child's arm to determine their blood group and genes (DNA) and test for infections that they may have had in the past, and measure their nutritional status. One drop of blood will be used to test your child for anemia, a condition of lower than normal red blood cells. The field member can provide you with the results of this test during their visit. We will record how your child responds to these procedures. If you agree, we would like to videotape your child during the blood-draw. We will use this information to better understand how these procedures affect child behavior. We will also ask you about how your child reacts to new situations, and what helps your child feel comfortable in new situations, which will take 10-15 minutes. This will help us understand your child's reaction to the different procedures we are administering for this project. We might also collect a small amount (10ml) of blood from your vein. We might also collect 3 saliva samples each from both you and your child before, during, and after the blood draw using a soft sponge placed under the tongue. The field member will deliver a stool collection kit and instruct you how to collect stool from your child. You will be instructed to collect your child's stool on the following morning, if the child defecates before their arrival, by having your child defecate in a clean diaper and use a plastic scoop to collect a small amount of fresh stool from the top of the pile into a container. The field person will collect this container when they come to collect the other specimens on the following day. Total participation time today will be approximately 3 hours and 15 minutes.

Tomorrow, we will ask you to collect your first urine sample of the morning immediately after you wake up. The main procedure will involve a second saliva collection, collecting your child's urine for 1 hour, feeding your child sugar syrup and then collecting their urine sample over a period of 5 hours. We intend to test the urine for the syrup to help us understand the health impacts of diarrhea in children. For a urine sample, due to the age of your child, we will be using a special urine collection bag to collect the urine from your child and we will demonstrate how it is used.

You/the mother will be requested to not feed your child for at least one hour before we feed him/her the syrup. During this fasting period, we will collect your child's urine for 1 hour by attaching the urine collection bag with a drainage tube (show sample) to the child. We will also collect two additional saliva samples from your child and one additional saliva sample from you using a soft sponge. We will then give a dose of the sugar syrup to the child and attach the urine collection bag with a drainage tube (show sample) to the child immediately after feeding him/her the syrup. We will encourage the child to drink water 30 minutes after taking the syrup to help urination. The field representative will remove the urine from the bag, whenever the child urinates. This collection will take place for 6 hours after which the bag will be removed from the child. During the 6-hour period of urine collection, you will be asked the quantities of foods (and the ingredients) you fed the child in the previous day and night. You will also be asked about your personal life experiences and health. You will be asked about perceptions regarding social norms that may affect a child's health. From the stool sample, we will examine whether your child has any worm infections, but this analysis will be done in Dhaka. We will offer your child deworming medication regardless of the stool sample result. Total participation time tomorrow will be approximately 7 hours.

Later, at the laboratory, we will measure your blood, saliva, hair, and urine samples and your child's blood, stool, saliva, hair, and urine samples for markers of nutritional status, infections, and health. You will not receive the results of any laboratory tests.

**Study time**: Study participation will take a total of approximately *10 hours and 15 minutes over 2 days.*

**Risks & Benefits**

Although we will try to protect your identity there is some possibility that confidentiality could be compromised. The syrup is a natural sugar solution that tastes pleasant. The blood will be collected by a trained professional. You and your child may feel some momentary pain during the blood collection. Your child may also feel some discomfort due to the presence of urine collection bag for 6 hours. Some of the questions I would like to ask you may seem private or personal since they touch on your life and health. You may feel uncomfortable talking about some of the topics. You do not have to answer any question or take part in the discussion/interview/survey if you don't wish to do so, and that is also fine. You do not have to give us any reason for not responding to any question, or for refusing to take part in the interview. All your answers will be kept as confidential as possible, and we anticipate that the risks from participating in this survey will be very minimal. There is no direct benefit for participating in this study, but your child's participation will help us to gain knowledge on the health impact of diarrheal diseases and how certain foods, nutrition, and life experiences in general may interact with diarrhea or other illnesses.

**Confidentiality**

All data and specimens collected will be kept confidential as allowed by the law of this country. The samples and the test results will be coded without your name or your child's name or personal information, and stored separately for analysis by the researchers. None of these researchers will be able to identify you or your child as the sample donors. Other persons who may have access to your test results include research groups that oversee the safety of the study including institutes such as the University of California Berkeley and the Bill and Melinda Gates Foundation, USA. Because of the potential need to release information to these parties, and the genetic information we are collecting is potentially identifying, we cannot guarantee absolute confidentiality. However, we will ensure every effort that we can to preserve your confidentiality. There is no expiration date to this permission. This information will be coded and it is unlikely that anyone will be able to trace it to you and your child, protecting your confidentiality.

All videotapes will be identified by a number only; no recordings will identify you or your child by name. These will be viewed only by trained personnel for coding of your child's response to the procedure. The videos will not be viewed by any other person. The videos will be stored in a locked cabinet accessible by study personnel only. These will never be made available to any persons not participating in the study.

**Future use of information**

The blood, urine, saliva, hair, and stool samples may be stored until the end of the study, so they can be analyzed in the lab at the same time. If you agree, the information, along with some of the blood, urine, saliva, hair and stool collected will be stored at ICDDR,B for up to 20 years so that we can conduct advanced tests on these samples to understand the infections that you may have had, and to find out what microbes lived in your body. No further consent will be taken from you, and you will not receive the results of these tests. A senior scientist at ICDDR,B will decide who can use the stored samples, and what tests will be done. If you change your mind about having your specimen saved for further testing, you may contact us and the sample will be discarded and not saved. The videotapes will be stored indefinitely.

**Right not to participate and withdraw**

Taking part in the study is completely voluntary. You may choose not to allow your child to participate in this study. You can drop out of this study at any time, even in the middle of the sample/urine collection. You have the right to refuse participation in this study, which will not affect your family's treatment at the Cholera Hospital in the future.

**Principle of compensation**

You need not pay us to take part in this study, and similarly we will not pay you money for attending in the study.

**Persons to contact:**

If you have any questions, you can ask me any time. If you have additional questions about the survey, you may contact:

If you have questions about your rights as a participant of a research study, or if you think some harm has been done to you because of the survey, you may contact or meet IRB Secretariat, M.A. Salam Khan:

M. A. Salam Khan, IRB Secretariat, phone: 9886498 or PABX 8860523-32 ext. 3206

If you agree to participate and you allow your child to participate, please indicate that by checking the boxes of the activities that you will agree to below:

**Urine Collection |__| mother |__| child**

**Venous Blood Collection |__| mother |__| child**

**Stool Collection |__| child**

**Saliva sample collection |__| mother |__| child**

**Hair sample collection |__| mother |__| child**

**Urine Long-term storage |__| mother |__| child**

**Venous Blood Long-term storage |__| mother |__| child**

**Stool Long-term storage |__| child**

**Saliva sample Long-term storage |__| mother |__| child**

**Hair sample Long-term storage |__| mother |__| child**

If you agree to our proposal of enrolling your household in our study, please indicate that by putting your signature or your left thumb impression at the specified space below

Thank you for your cooperation

| | |
|---|---|
| ______________________________ | ____________________ |
| Signature or left thumb impression of Guardian | Date |
| ______________________________ | ____________________ |
| Signature or left thumb impression of the witness | Date |
| ______________________________ | ____________________ |
| Signature of the PI or his/her representative | Date |

PROTOCOL
Biomedical
Non-Exempt
Berkeley

Protocol # 2011-09-3652
Date Printed: 07/31/2018

-----------------------------------------------------------------------------------

| | |
|---|---|
| **Protocol Title:** | Measuring the benefits of sanitation, water quality, handwashing and nutrition interventions for improving health and development in rural Bangladesh |
| **Protocol Status:** | APPROVED |
| **Date Submitted:** | 10/26/2017 |
| **Approval Period:** | **10/30/2017-11/03/2018** |
| **Important Note:** | **This Print View may not reflect all comments and contingencies for approval. Please check the comments section of the online protocol. Questions that appear to not have been answered may not have been required for this submission. Please see the system application for more details.** |

*** * * Attached Document * * ***

| Document Name | Created Date |
|---|---|
| 2011-09-3652_Colford_1z-Consent-Microbiome-v20-English.pdf | 10/26/2017 |

**Appendix 1z: English consent for microbiome substudy**

Protocol Title: Effect of hand washing, water treatment, sanitation and nutritional supplement interventions on child health and development in rural Bangladesh

Principal Investigator's name: Dr. Leanne Unicomb

**Purpose of the research**

Hello/Assalamualaikum/Namaste. My name is ________ and I work with the ICDDR,B (Cholera Hospital) in Dhaka. We are conducting research on diarrheal diseases and through this particular study we want to learn about the health impact of diarrheal diseases in children. Malnutrition in children is associated with infections of our intestines that can change our use of nutrients. It is possible that genes in our body, made up of DNA, make some children more susceptible to malnutrition than others. This DNA is inherited from our parents and we also carry genes from many friendly microbes that live on and in our body. We are interested in learning if the exposure of a child to diarrheal disease has long term effects.

**Why are we inviting you to participate in the study?**

We are interested in enrolling this compound because we collected information on your household earlier and we are interested in conducting additional testing to evaluate your child's digestive tract.

**What is expected from the participants of the research study?**

Participation in this study will only take 15 minutes each month. To achieve the aim of this study, we will collect a stool sample from your child once a month for the next 10 months. The samples will help us understand whether your child has been exposed to infections and will also help us understand the friendly microbes that live in your child's digestive tract.

If you agree to participate, a field research person will visit your household twice per month for the next 10 months for this purpose. On the first day, a field member will deliver a stool collection kit and instruct you how to collect stool from your child. You will be instructed to collect your child's stool on the following morning, if the child defecates before the field member's arrival, by having your child defecate on a sheet of provided plastic and you will use a plastic scoop to collect a small amount of fresh stool from the top of the pile into a container. The field person will collect this container. We will examine whether your child has infections or determine which friendly microbes live in your child's digestive tract through different laboratory test, but we will not be able to share the results with you.

**Risks & Benefits**

Although we will try to protect your identity there is some possibility that confidentiality could be compromised. There is no direct benefit from participating in this study, but your child's participation will help us to gain knowledge on diarrheal disease and friendly microbes in children.

**Confidentiality**

All data and specimens collected will be kept confidential as allowed by the law of this country.

Confidentiality of the data and test results will be strictly maintained. We will use the information only for the purpose of the study, and we will not use you or your child's name in sharing and publishing the results of this study.

**Future use of information**

The stool samples may be stored until the end of the study, so they can be analyzed in the lab. If you agree, the information, along with some of the stool collected will be stored at ICDDR,B for up to 20 years so that we can conduct advanced tests on these samples to understand the infections that you may have had. No further consent will be taken from you, and you will not receive the results of these tests. A senior scientist at ICDDR,B will decide who can use the stored samples, and what tests will be done. If you change your mind about having your specimen saved for further testing, you may contact us and the sample will be discarded and not saved.

**Right not to participate and withdraw**

Taking part in the study is completely voluntary. You may choose not to allow your child to participate in this study. You can drop out of this study at any time, even in the middle of the sample collection. You have the right to refuse participation in this study, which will not affect your family's treatment at the Cholera Hospital in the future.

**Principle of compensation**

You need not pay us to take part in this study, and similarly we will not pay you money for attending in the study.

**Persons to contact:**

If you have any questions, you can ask me any time. If you have additional questions about the survey, you may contact:

Masud Parvez: phone: 01817541872, Center of Communicable Diseases, ICDDR,B, Mohakhali, Dhaka 1212.

If you have questions about your rights as a participant of a research study, or if you think some harm has been done to you because of the survey, you may contact or meet IRB Secretariat, M.A. Salam Khan:

M. A. Salam Khan, IRB Secretariat, phone: 9886498 or PABX 8860523-32 ext. 3206

If you agree to our proposal of enrolling your household in our study, please indicate that by putting your signature or your left thumb impression at the specified space below

I agree to have my child's stool samples to be stored long term [ ]

Thank you for your cooperation

_________________________________________ ____________________

Signature or left thumb impression of Guardian Date

_________________________________________ ____________________

Signature or left thumb impression of the witness Date

_________________________________________ ____________________

Signature of the PI or his/her representative Date

---------------------------------------------------------------------------------
---

**Protocol Title:** Measuring the benefits of sanitation, water quality, handwashing and nutrition interventions for improving health and development in rural Bangladesh

**Protocol Status:** APPROVED

**Date Submitted:** 10/26/2017

**Approval Period:** **10/30/2017-11/03/2018**

**Important Note:** This Print View may not reflect all comments and contingencies for approval. Please check the comments section of the online protocol.
Questions that appear to not have been answered may not have been required for this submission. Please see the system application for more details.

*** * * Attached Document * * ***

| Document Name | Created Date |
|---|---|
| 2011-09-3652_Colford_2z-Consent-Microbiome-v20-Bengali.pdf | 10/26/2017 |

icddr,b
KNOWLEDGE FOR GLOBAL LIFESAVING SOLUTIONS

### Appendix 2z: Bengali consent for microbiome substudy

Protocol Title: Effect of hand washing, water treatment, sanitation and nutritional supplement interventions on child health and development in rural Bangladesh

প্রধান গবেষকের নাম:ডঃ লিয়েন ইউনিকম্ব

## গবেষনার উদ্দেশ্য

আসসালামুআলাইকুম/ নমস্কার। আমার নাম---------------- । আমি ঢাকার আইসিডিডিআর, বি (কলেরা হাসপাতাল) তে কাজ করি। আমরা ডায়রিয়া রোগ নিয়ে গবেষণা করছি এবং এই গবেষণার মাধ্যমে আমরা শিশুদের স্বাস্থ্যের উপর ডায়রিয়া রোগের প্রভাব সম্পর্কে জানতে চাচ্ছি। শিশুদের ডায়রিয়া রোগের **ঝুঁকির** কোন দীর্ঘমেয়াদি প্রভাব রয়েছে কিনা, তা জানতে আমরা আগ্রহী।শিশুদের অপুষ্টি অন্ত্রের সংক্রমনের সাথে সম্পর্কীত যা আমাদের পুষ্টি উপাদানের ব্যবহার বদলে দিতে পারে। আমাদের দেহের ডিএনএ-র জিনের কারণে কিছু কিছু শিশুদের অন্যদের তুলনায় অপুষ্টির ঝুঁকি বেশী থাকে। এই ডিএনএ আমরা উত্তরাধিকার সূত্রে বাবা-মার কাছ থেকে পাই এবং এছাড়াও আমরা আমাদের দেহে বসবাসকারী ৰতিকর নয় এমন জীবানূর অনেক জিন বহন করি।শিশুদের ডায়রিয়া রোগের ঝুঁকির কোন দীর্ঘমেয়াদি প্রভাব রয়েছে কিনা, তা জানতে আমরা আগ্রহী।

#### আমরা কেন আপনাকে এই গবেষণায় অংশগ্রহণে আমন্ত্রণ জানাচ্ছি?

আমরা এই বাড়ীকে গবেষণায় অন্র্‌ভূক্ত করতে চাচ্ছি কারণ আমরা এর আগে আপনার খানা সম্পর্কীত তথ্য সংগ্রহ করেছি এবং আমরা আপনার শিশুর আরও কিছু পরীক্ষার মাধ্যমে শিশুটির পরিপাক নালী সর্ম্পকে জানতে আগ্রহী।

#### গবেষণায় অংশগ্রহণকারীর কাছে প্রত্যাশা কী?

এই গবেষণায় অংশগ্রহণ করলে প্রতিমাসে আনুমানিক ১৫ মিনিট সময় ব্যয় হবে।এই গবেষণার উদ্দেশ্য সফল করতে হলে প্রতিমাসেএকবার করে মোট ১০ মাস যাবত শিশুর কিছুটা পায়খানার নমুনা সংগ্রহ করতে হবে। পায়খানার নমুনা পরীক্ষাগারে পরীক্ষার মাধ্যমে আমরা বুঝতে সক্ষম হব যে, আপনারশিশুর সংক্রমনের ঝুঁকি কেমন এবং শিশুর দেহে বসবাসকারী ৰতিকর নয় এমন জীবানূ সম্পর্কে জানতে সহায়তা করবে।

আপনি অংশগ্রহণ করতে রাজী হলে, একজন মাঠ গবেষক এই কাজে প্রতিমাসেদুইবার করে মোট ১০ মাস যাবত আপনার খানায় আসবে। নমুনা সংগ্রহ করার আগের দিন একজন মাঠকর্মী আপনাকে পায়খানা সংগ্রহ করার সামগ্রী দিয়ে যাবে এবং কিভাবে আপনার শিশুর পায়খানা সংগ্রহ করতে হবে তা দেখিয়ে দেবে। পরের দিন সকালে যদি মাঠকর্মী পৌছানোর পূর্বে শিশু পায়খানা করে, তবে আপনি আপনার শিশুর পায়খানা সংগ্রহ করবেন। পায়খানা সংগ্রহ করার জন্য আপনাকে একটি পস্নাষ্টিক শিট দেয়া হবে, যেখানে শিশু পায়খানা করবে এবং আপনি একটি পস্নাষ্টিক (ছেনি) স্কুপ ব্যবহার করে উপর থেকে অল্প একটু সদ্য (এইমাত্র করা) পায়খানা বোতলে ভরবেন। আমাদের মাঠকর্মী যখন অন্যান্য নমুনা সংগ্রহ করার জন্য আসবে, তখন এই বোতলটি আপনার কাছ থেকে নিয়ে নেবে। আমরা পরীৰাগারে পরীৰার মাধ্যমে দেখবো যে আপনার শিশুর শরীরে কোন সংক্রামন আছে কিনা অথবা শিশুর দেহে বসবাসকারী অনেক ৰতিকর নয় এমন কোন জীবানু আছে কিনা । কিন্তু আমরা এর ফলাফল আপনাকে জানাবো না।

## ঝুঁকি এবং সুবিধা

এই গবেষণায় অংশগ্রহণে আপনার কোনো ঝুঁকি নেই। আপনি গবেষণায় অংশগ্রহণের জন্য সরাসরি কোন ধরণের অর্থনৈতিক বা অন্য কোন সহায়তাও পাবেন না। কিন্তু আপনার শিশুর অংশগ্রহণ, শিশুদের ডায়রিয়া রোগ সম্পর্কে এবং শিশুর দেহে বসবাসকারী ৰতিকর নয় এমন কোন জীবানু আছে কিনা সে সম্পর্কে জ্ঞানলাভে আমাদেরকে সহায়তা করবে।

## গোপনীয়তা

সকল তথ্য এবং সংগৃহীত নমুনা এই দেশের আইন অনুযায়ী গোপনীয়তার সাথে রাখা হবে। তথ্য এবং নমুনার ফলাফলের গোপনীয়তা কঠোরভাবে পালন করা হবে। আমরা শুধু গবেষণার প্রয়োজনে এই তথ্য ব্যবহার করব এবং আপনার নাম অথবা পরিচয়ের কোন সূত্র এই গবেষণা সংক্রান্ৰ কোন রির্পোটি/ প্রকাশনায় উলেস্নখ করা হবে না।

**ভবিষ্যতে তথ্যের ব্যবহার**

একই সময়ে ল্যাবে পরীক্ষা করার জন্য এই পায়খানার নমুনা গবেষণার শেষ সময় পর্যন্ৰ সংরক্ষণ করা হতে পারে। এই পায়খানার নমুনার পাশাপাশি সংগ্রহকৃত তথ্যও এই নমুণা আইসিডিডিআর,বি তে ২০ বছর পর্যন্ৰ সংরৰণ করা হতেপারে যাতে করে আমরা আপনার কোন সংক্রমন ছিল কিনা তা জানার জন্য উন্নত ধরনের পরীক্ষা করা হতে পারে। এর জন্য পরবর্তীতে আপনার কাছ থেকে কোন অনুমতি নেওয়া হবে না এবং আপনি এই পরীক্ষার কোন ফলাফলও পাবেন না। আইসিডিডিআর,বি একজন সিনিয়র বিজ্ঞানী ঠিক করবেন কে সংরক্ষণকৃত নমূণা ব্যবহার করতে পারবে এবং কোন কোন পরীক্ষা করতে পারবে। এই নমুনা ভবিষ্যতে পরীৰার জন্য রেখে দেওয়ার ব্যপারে যদি আপনি কখনও আপনার মতামত পরিবর্তন করেন তাহলে আপনি আমাদের সাথে যোগাযোগ করতে পারেন এবং আমরা আপনার নমুণা ফেলে দিব।

**স্বেচ্ছা অংশগ্রহণ**

এই গবেষণায় অংশগ্রহণ সম্পূর্ণভাবে আপনার ইচ্ছার ওপর র্নিভর করে। আপনি এই গবেষণায় আপনার শিশুকে অংশগ্রহণ নাও করতে দিতে পারেন। আপনার লিখিত অনুমতি দেয়ার পরবর্তীতেও আপনি ইচ্ছা করলে যে কোন সময়ে এমনকি নমুনা সংগ্রহের মাঝখানেও আপনার অনুমতি প্রত্যাখ্যান করতে পারেন। এই গবেষণায় অংশগ্রহণ না করার অধিকার আপনার আছে এবং এজন্য ভবিষ্যতে আপনার পরিবারের সদস্যদের আইসিডিডিআর,বি-র স্বাস্থ্যসেবা গ্রহণের উপর কোন প্রভাব পড়বে না।

**ক্ষতিপূরণ**

এই গবেষণায় অংশগ্রহণের জন্য আপনার সরাসরি কোন ধরণের অর্থনৈতিক খরচ/ ব্যয় হবে না এবং একইভাবে আপনি এই গবেষণায় অংশগ্রহণের জন্য সরাসরি কোন ধরণের অর্থনৈতিক সহায়তাও পাবেন না।

**যোগাযোগ**

আপনার কোনো প্রশ্ন থাকলে আমাকে জিজ্ঞাসা করতে পারেন। যদি গবেষণা সম্পর্কিত অতিরিক্ত কোন প্রশ্ন থাকে তাহলে আপনি **জনাব মাসুদ পারভেজ এর সাথে ০১৮১৭৫৪১৮৭২ মোবাইল নম্বরে সরাসরি যোগাযোগ করতে পারেন।** এই গবেষণায় আপনার অধিকার-সংক্রান্ৰ কোন প্রশ্ন থাকলে অথবা গবেষণার কারণে আপনার কোন ক্ষতি হতে পারে বলে যদি মনে করেন, তাহলে আপনি ৯৮৮৬৪৯৮ অথবা ৮৮৬০৫২৩ (এক্স- ৩২০৬) টেলিফোন নম্বরে অথবা সরাসরি এম এ সালাম খান, আইআরবি সেক্রেটারিয়েট এর সাথে যোগাযোগ করতে পারেন।

আপনি যদি এই গবেষণায় অংশগ্রহণ করতে আগ্রহী হন তাহলে নিম্নের নির্ধারিত স্থানে স্বাক্ষর অথবা বাম বৃদ্ধাঙ্গুলীর ছাপ দিন।

আপনার সহযোগিতার জন্য ধন্যবাদ।

............................................ ..................................

অভিভাবকের স্বাক্ষর/বাম বৃদ্ধাঙ্গুলির ছাপ তারিখ

............................................ ..................................

সাক্ষীর স্বাক্ষর/বাম বৃদ্ধাঙ্গুলির ছাপ তারিখ

............................................ ..................................

পি আই/ গবেষকের প্রতিনিধির স্বাক্ষর তারিখ

দ্বিমত:

আমি চাই না আমার শিশুর পায়খানার নমুনা দীর্ঘ সময় ধরে সংরক্ষণ করা হোক☐

---------------------------------------------------------------------------------------------

**Protocol Title:** Measuring the benefits of sanitation, water quality, handwashing and nutrition interventions for improving health and development in rural Bangladesh

**Protocol Status:** APPROVED

**Date Submitted:** 10/26/2017

**Approval Period:** **10/30/2017-11/03/2018**

**Important Note:** **This Print View may not reflect all comments and contingencies for approval. Please check the comments section of the online protocol. Questions that appear to not have been answered may not have been required for this submission. Please see the system application for more details.**

*** * * Attached Document * * ***

| Document Name | Created Date |
| --- | --- |
| Prot PR-11063_ERC approval addendum_10 June 2014.pdf | 10/26/2017 |

icddr,b

### Memorandum

10 June 2014

To: Dr Leanne Unicomb
Principal Investigator of research protocol # PR-11063
Centre for Communicable Diseases (CCD)

From: Professor Kazi Zulfiquer Mamun
Chairperson
Ethical Review Committee (ERC)

Sub: Approval of an addendum to research protocol # PR-11063

Thank you for your memo dated 3 June 2014 attaching the modified version of your addendum proposal to previously approved research protocol # PR-11063 titled **"A randomized controlled trial of the benefits of water sanitation hygiene plus nutrition interventions on child growth"** addressing the issues raised by the committee in its special meeting held on 18 May 2014 to the satisfaction of the Committee. I have the pleasure to accord approval of the addendum proposal of the above protocol.

Other terms and conditions for implementation of your research protocol, as contained in our memo dated 9 April 2012 according approval of the research protocol shall, however, remain unchanged.

Thank you.

Cc: Director, CCD

GPO Box 128, Dhaka 1000, Bangladesh.
**For Packages :** 68, Shaheed Tajuddin Ahmed Sarani, Mohakhali, Dhaka 1212, Bangladesh.

PROTOCOL
Biomedical
Non-Exempt
Berkeley

-------------------------------------------------------------------------------------

| | |
|---|---|
| **Protocol Title:** | Measuring the benefits of sanitation, water quality, handwashing and nutrition interventions for improving health and development in rural Bangladesh |
| **Protocol Status:** | APPROVED |
| **Date Submitted:** | 10/26/2017 |
| **Approval Period:** | **10/30/2017-11/03/2018** |
| **Important Note:** | **This Print View may not reflect all comments and contingencies for approval. Please check the comments section of the online protocol. Questions that appear to not have been answered may not have been required for this submission. Please see the system application for more details.** |

*** * * Attached Document * * ***

| Document Name | Created Date |
|---|---|
| PR-11063_RRC Addendum Approval_13 April 2014.pdf | 10/26/2017 |

icddr,b

### Memorandum

13 April 2014

To: Dr Leanne Unicomb
Principal Investigator of research protocol # PR-11063
Centre for Communicable Diseases (CCD)

From: Abbas Bhuiya, Ph.D.
Chairperson
Research Review Committee (RRC)

Sub: **Addendum proposal to research protocol # PR-11063**

Thank you for submitting your addendum proposal to previously approved research protocol # PR-11063 titled "**A randomized controlled trial of the benefits of water sanitation hygiene plus nutrition interventions on child growth**" for consideration of the RRC in its meeting held on 3 April 2014. This is to inform you that after review and discussion, the committee unanimously approved your addendum proposal. Accordingly, you may proceed to obtain approval of the Ethical Review Committee.

Other terms and conditions for implementation of your research protocol, as contained in our memo dated 26 November 2013 according initial approval of the research protocol shall, however, remain unchanged

Thank you.

Copy: Director, CCD.

GPO Box 128, Dhaka 1000, Bangladesh.
**For Packages :** 68, Shaheed Tajuddin Ahmed Sarani, Mohakhali, Dhaka 1212, Bangladesh.

PROTOCOL
Biomedical
Non-Exempt
Berkeley

Protocol # 2011-09-3652
Date Printed: 07/31/2018

-------------------------------------------------------------------------------------
---

**Protocol Title:** Measuring the benefits of sanitation, water quality, handwashing and nutrition interventions for improving health and development in rural Bangladesh

**Protocol Status:** APPROVED

**Date Submitted:** 10/26/2017

**Approval Period:** **10/30/2017-11/03/2018**

**Important Note:** **This Print View may not reflect all comments and contingencies for approval. Please check the comments section of the online protocol.**
**Questions that appear to not have been answered may not have been required for this submission. Please see the system application for more details.**

*** * * Attached Document * * ***

| Document Name | Created Date |
| --- | --- |
| Wash Benefits Midline & Endline Survey Questionnaires_no life experiences.pdf | 10/26/2017 |

ICDDR,B

### Wash Benefits
#### Midline and Endline Questionnaire

09/02/2014

**NOTES TO THE FIELD TEAM**

| POPULATION | DEFINITION |
|---|---|
| Target children | Children in utero at enrollment (primary study population) |
| Mothers | Mothers of target children |
| Diarrhea cohort | Children < 36 months old at enrollment, living in study compounds |
| Parasite cohort | Children 18 – 27 months old at baseline (blood spot, stool) |
| EE subsample | Subsample of Target Children for EE specimen collection |
| New births | Children born into study households after Target Children |
| All HH | All enrolled study households |
| LNS households | Enrolled study households in the LNS or LNS+WASH arms |

আগামী কয়েক মাস আমাদের এই প্রকল্পের কিছু বিষয়ে আপনাকে কিছু তথ্য দেওয়ার জন্য ফোনে যোগাযোগ রাখতে চাই । আপনার কি কোন ফোন আছে? যদি না থাকে তাহলে আপনার বাড়ীতে কারো বা প্রতিবেশীর ফোন আছে কি? তার নাম্বার কত? At some point over the next several months, we may need to get in touch with you by phone to give you information about our project. Do you have a phone? If not, does someone in your compound have a phone? Maybe a neighbor? What is the number?

**Wash Benefit Module 0 TRACKING**

| | | |
|---|---|---|
| | 001 | উত্তরদাতার/ উত্তরদাতার খানার কোন সদস্যের কী কোন ফোন নাম্বার আছে?(Is there a phone available?) 1 Yes 0 No (Skip to 005) |
| C.11 | **002** | -----------প্রাথমিক ফোন নাম্বার (Primary phone number) |
| C.12 | 003 | এই ফোনে কি আপন াকে সরাসরি পাওয়া যাবে? (Does this phone number belong to you?)<br>1 Yes(Skip to 005) 0 No |

| | | |
|---|---|---|
| C.13 | 004 | যদি আপন াকে পাওয়া না যায় তাহলে এই নাম্বারটি কার? IF NO, to whom does this number belong? (Specify relationship to respondent)<br>1 . স্বামী (HUSBAND)<br>2. শ্বশুর IN LAWS<br>3. পিতা(FATHER)<br>4.মা(MOTHER)<br>5. পরিবারের সদস্য(FAMILY MEMBER)<br>99. অন্যান্য (নাম ও উত্তরদাতার সাথে সম্পর্ক লিখুন) (OTHER) |
|  | 004.a | দ্বিতীয় কোন ফোন নাম্বার আছে কী?(Is there a phone available?)<br>1 Yes 0 No (Skip to 008) |
| C.14 | 005 | -----------দ্বিতীয় ফোন নাম্বার Secondary phone number |
| C.15 | 006 | এই ফোনে কি আপন াকে সরাসরি পাওয়া যাবে? Does this phone number belong to you?<br>1 Yes(Skip to 008) 0 No |
| C.16 | 007 | যদি আপন াকে পাওয়া না যায় তাহলে এই নাম্বারটি কার? IF NO, to whom does this number belong?<br>1. স্বামী (HUSBAND)<br>2. শ্বশুর IN LAWS<br>3. পিতা(FATHER)<br>4 .মা(MOTHER)<br>5 .পরিবারের সদস্য(FAMILY MEMBER)<br>99 .অন্যান্য (নাম ও উত্তরদাতার সাথে সম্পর্ক লিখুন) (OTHER) |
| C.17 | 008 | এই খানাটির আশেপাশে কোন উলেস্নখযোগ্য কিছু আছে কি? (Location/landmark) |
| | 009 | \|_____\| \|_____\| সবচেয়ে ছোট বাচ্চার মা কত ক্লাস পর্যন্ত পড়ালেখা শেষ করেছেন ? (জানি না = ***99***) Education of mother of the youngest child *(Years of education completed, DK=99)* |
| | 010 | \|_____\| \|_____\| সবচেয়ে ছোট বাচ্চার বাবা কত ক্লাস পর্যন্ত পড়ালেখা শেষ করেছেন ? (জানি না = ***99***) Education of father of the youngest child *(Years of education completed, DK=99)* |
| | 011 | \|_____\| \|_____\| সবচেয়ে ছোট বাচ্চার বাবার প্রধান পেশা [Main occupation of father of the youngest child]<br>পেশা কোড Occupation Code:<br>1. কৃষক [Farmer/Cultivator]<br>2. ঘরে কাজ করে [Homemaker]<br>3. কৃষি শ্রমিক [Agri-labor]<br>4. শ্রমিক (কৃষি শ্রমিক ছাড়া) [Non-agri labor]<br>5. বেতনভুক্ত কর্মচারী (সরকারী/প্রাইভেট/এন.জি.ও) [Salaried job (Govt./Private/NGO)]<br>6. রাজমিস্ত্রী [Mason (*Rajmistri*)]<br>7. কাঠ মিস্ত্রী [Carpenter]<br>8. ভ্যান/রিকশা চালক [Van/Rickshaw puller]<br>9. জেলে [Fisherman]<br>10. নৌকা চালক/মাঝি [Boatman]<br>11. কর্মকার [Blacksmith]<br>12. স্বর্ণকার [Goldsmith]<br>13. কুমার/কুম্ভকার [Potter (soil smith)]<br>14. মুচি [Shoe polish /maker]<br>15. দোকানদার [Shopkeeper]<br>16. ফেরিওয়ালা [Vendor (*Feriwala/howker*)]<br>17. ক্ষুদ্র ব্যবসায়ী (মূলধন <=১০০০০)<br>21. কুটীর শিল্প [Cottage industry]<br>22. পোলট্রি/ ব্যবসার জন্য পশু লালন-পালনকারী [Poultry /livestock rearer]<br>23. বৈদ্যুতিক মিস্ত্রী [Electrician]<br>24. হোমিওপ্যাথি ডাক্তার [Homeopath]<br>25. আধ্যাত্মিক চিকিৎসক/ কবিরাজ/ ওঝাঁ [Spiritual healer/kabiraj/ Ojha]<br>26. পেশাদার ডাক্তার/উকিল [Professional practitioner (Doctor/lawyer)]<br>27. ইমাম/ ধর্মযাজক [Imam/priest]<br>28. অবসরপ্রাপ্ত চাকুরীজীবী [Retired service holder]<br>29. ছাত্র [Student]<br>30. বেকার [Unemployed]<br>31. অক্ষম [Disabled]<br>32. কাজের লোক [Domestic maid / servant]<br>33. জমিদার (শস্য উৎপাদন অথবা অন্য কোন কাজে কৃষকদের জমি বর্গা দেয়) [Landlord (Provide land for farmers for sharecropping or others)]<br>34. বিদেশে থাকে [Staying abroad] |

| | | | |
|---|---|---|---|
| | | [Petty trader, capital <=10000]<br>18. ব্যবসায়ী (মূলধন >১০০০০) [Business, capital >10000]<br>19. দর্জি [Tailor]<br>20. ড্রাইভার [Driver] | 35. মৃত/নিখোঁজ [Died/untraced]<br>36. ভিক্ষুক (Beggar<br>37. নাপিত Haircutter<br>99. জানিনা [Don't know]<br><br>77. অন্যান্য (বর্ণনা লিখুন) [Others (specify] |
| | 012 | I_____I I_____I আপনার খানাতে/পরিবারে কত জন লোক প্রতিদিন একই হাড়িতে রান্না করে খান? How many people in this house eat from the same cooking pot every day (খানা বলতে একই হাড়িতে রান্না করে খায় এমন সদস্যদের বুঝানো হয়েছে) | |
| | 013 | I_____I আপনার খানাতে ৩৬ মাসের নীচে কতজন শিশু আছে? How many children do you have < 36 months do you have? | |
| | 014 | I_____I এই বাড়ীতে কতগুলো খানা আছে? How many HHs do you have in this bari? | |
| | 014a | I_____I এই বাড়ীত বইজ লাইন কতগুলা খানা ছিল? How many HHs in baseline do you have in this bari? | |
| | 014b | I_____I এই বাড়ীত কতগুলা খানা নতন আসছ? How many new HHs do you have in this bari? | |

A. এই বাড়ীর খানাগুলোর সেনসাস (সকল খানার) CENSUS OF HOUSEHOLDS ARE IN THIS COMPOUND (All households)

পড়ে শোনান READ: ঠিক আছে। আমি এখন আপনাকে সকল খানার সম্পর্কে আলাদাভাবে কিছু প্রশ্ন করব। প্রথমে আপনার খানা সম্পর্কে বলুন। Okay. Now I am going to ask you about each household separately. First, think about your own household.
MARK "99" IF RESPONDENT DOESN'T KNOW THE ANSWER TO ANY OF THE FOLLOWING QUESTIONS.

| | A.1 | A.2 | A.3 | A.4 | A.5 | A.6 | A.7 |
|---|---|---|---|---|---|---|---|
| খানার নাম্বার HH No. | খানা প্রধানের নাম কি? What is the name of the household head? FOR REFERENCE ONLY | এই খানাতে ৪০ বছরের উপরের বয়সের কতজন সদস্য আছে? How many adults over 40 years live in this household? | এই খানাতে (>১৮-৪০) বছরের বয়সের কতজন সদস্য আছে? How many young adults (>18-40 years) live in this household? | এই খানাতে (>৩-১৮) বছরের বয়সের কতজন সদস্য আছে? How many school-aged children (>3-18 years) live in this household? | এই খানাতে (০-৩) বছরের বয়সের কতজন সদস্য আছে? How many young children (0-3 years) live in this household? | TOTAL NUMBER OF PEOPLE IN BARI | Migration status of HH<br>1. Migration out<br>2. Migration in<br>3. No change<br>4. Mix with other |
| 1 | ONLY LOAD THE HH HEAD NAMES LIST | | | | | | |
| 2 | | | | | | | |

#### RESPONDENT IDENTIFICATION

| | | |
|---|---|---|
| মডিউল আইডি Module ID | WBM 1 | |
| | 101. | : What is your full name? প্রধান উত্তরদাতার নাম [Name of respondent: |
| | 102. | : I_____I প্রধান উত্তরদাতার পরিচয় [Status of main respondent]<br><br>1. প্রধান পরির্চযাকারী /মা [Mother of youngest child]<br>2. পরির্চযাকারী (পুরুষ) [Male caregiver]<br>3. পরির্চযাকারী (মহিলা) [Female caregiver]<br>if answer is 2 or 3 go to q001 |
| | 103. | প্রধান উত্তরদাতার বয়স (বছরে) [Age of main respondent: *(in years)*] |
| | 103a | ধর্ম **Religion**<br>মুসলিম **1. Muslim**<br>হিন্দু **2. Hindu**<br>খ্রিষ্টান **3. Christian**<br>অন্যান্য (উলেস্নখ করম্নণ) **4. Other** |
| | 104. | আপনি কি এখন গর্ভবতী ? Are you pregnant right now?<br>হ্যাঁ 1. Yes<br>না 0. No (Skip to 001) |

আমি আপনাকে আপনার গর্ভবতী হওয়া বিষয়ে কিছু ব্যক্তিগত প্রশ্ন এই গবেষনার জন্য করতে চাই ।আমি আশা করি এ বিষয়ে আপনি কিছু মনে করবেন না এবং আমাদিগকে সহায়তা করবেন। I need to ask you some personal questions regarding your pregnancy because that is our project's eligibility criteria. I hope you do not mind and will cooperate with us. [see scripts of other studies]

| | | |
|---|---|---|
| C. 108 | 105. | **উত্তরদাতার সর্বশেষ মাসিকের তারিখ (What was the date of your last menstrual period?)__/__/____**<br>88 / 88 / 88 = সর্বশেষ গর্ভধারণের পর থেকে মাসিক না হলে (No menstruation since their last pregnancy)<br>99 = জানি না (Don't know / not sure) |
| C. 109 | 106. | \|__\| গর্ভধারণের পূর্নাঙ্গ মাস গণনা করম্নন( Record number of completed months)<br>99 = জানি না (Don't know / not sure) |
| C.110 | 107. | \|___\|গর্ভধারণের মসয়কালের উৎস(Source of Pregnancy Length)<br>1 = শুধুমাত্র মায়ের নির্ণয় (Estimated by mother only)<br>2 = মায়ের এবং স্বাস্থ্য সেবাদানকারীর নির্ণয় ,আলট্রাসাউন্ড ছাড়া(Estimated by mother and a health practitioner, no ultrasound (last prenatal visit)<br>3 = মা, স্বাস্থ্য সেবাদানকারী এবং আলট্রাসাউন্ড দ্বারা নির্ণয় Estimated by mother and health practitioner, using ultrasound (last prenatal visit) |
| | 108. | __/__/__ কখন আপনার শিশু জন্মাবে? (What will be the expected DOB of your child?)<br>__/__/__ |
| | 109. | |
| | 110. | বাচ্চা জন্মদানের জন্য আপনি কি মায়ের বাড়ী যাবেন ? Do you have any paln to go to your mother's house for delivary porpoise?<br>1. Yesহ্যাঁ<br>0. No না (Skip to module 0) |
| | 111. | আপনার মায়ের বাড়ী যে গ্রামে সেই গ্রামের নাম কি, যেখানে আপনি বাচচার জন্মগ্রহনের সময় যাবেন?<br>What is the name of your mother's (maternal) village, where you might go to give birth?<br>0. Same as this একই গ্রাম<br>1. গ্রামের নাম লিখুন ______________ |

**Mother's Movement Questions**

| | | | | **Variable name** |
|---|---|---|---|---|
| **112** | সর্বশেষ পরিদর্শনের সময় পর্যন্ত গর্ভবতী অবস্থায়, ডেলিভারী বা ডেলিভারীর পরে মা বাড়ীর বাহিরে বেড়াতে গিয়েছিল কী?<br><br>Has the mother resided outside the bari since our last visit during pregnancy, delivery and after deliver? | হ্যাঁ (Yes)..........1<br>না (No).......0 skip to 001<br>DK (জানিনা)...999 skip to 001 | □ | q112 |
| **113** | যদি ১১২ এর উত্তর ১ হয়, কতবার মা বাড়ীর বাহিরে বেড়াতে গিয়েছিল?<br><br>(If 112 answer is 1), how many times the mother visited outside the bari? | | □ | q113 |
| **114** | যদি ১১২ এর উত্তর ১ হয়, প্রতিবার কতদিন করে ছিল?<br><br>(If 112 answer is 1), How long was the duration of each travel (enter the number of days for each time mother traveled)? | | □□<br>□□<br>□□<br>□□ | q114_1<br>q114_2<br>q114_3<br>↓<br>q114_10 |

#### Wash Benefit Module 1: Birth date, age & sex measurement

| | | | | | | | |
|---|---|---|---|---|---|---|---|
| | 4019.a | I_____I এই বাড়ীতে ৩৬ মাসের নীচে কতজন শিশু আছে? How many children do you have < 36 months in this bari? | | | | | |
| | 4019.b | Number of target child (র্নিদিষ্ট □□□□□ □□□□□□) | | | | | |
| | 4019.c | Number of baseline child (□□□□ □□□□□ □□□□□ □□□□□□) | | | | | |
| | 4019.d | Number of migration in child (□□□□ □□□□□ □□□□□□) | | | | | |
| শিশুর প্রদত্ত তালিকা দেখুন এবং শিশুর উপস্থিতি উলেস্নখ করম্নন (Now look at the list provided to ensure the presence of the additional enrolled children in the bari) | | | | | | | |
| | | শিশুর পরিচয় (Who is this child?) | নাম (Name) | লিঙ্গ (Sex) | জন্ম তারিখ(Birth date) | জন্ম তারিখের উৎস(Source of DOB )(C.104) | নির্বাচন করার কারণ (একাধিক উত্তর হতে পারে) Reason for enrollment (select all that apply) **[PDA to store accordingly]** |
| **C.06.** | 4001. | 1. র্নিদিষ্ট শিশু (Target child)<br>2 .একই মায়ের অন্য শিশু(Another child of same mother)<br>3. একই বাড়ীর অন্য খানার শিশু (Child from the same bari)<br>4. একই খানার অন্য মায়ের শিশু(Another child of another mother in the same HH) | | 1 পুরম্নষ M<br>0 মহিলা F | দিন/মাস/বছর DD/MM/YY | 1 = কার্ড দ্বারা নিশ্চিত হয়েছিল (Confirmed DOB by valid vaccination/health card)<br>2 = মা/আত্মীয় দ্বারা নিশ্চিত হয়েছিল (Mother/Relative remembers DOB)<br>3 = উভয় (Both 1 & 2)<br>4 = ঘটনা দ্বারা নিশ্চিত হয়েছিল (Estimated DOB with 2 and event calendar) | ☐1 ডায়রিয়া Diarrhea<br>☐2 পায়খানার নমুণা Stool specimen<br>☐3 রক্তের নমুণা Blood specimen<br>☐ 4 Target child<br>☐ 5 Missing/absent<br>☐ 6 New birth<br>☐ 7 Death<br>☐ 8 Migration in |
| | 4020_1. [For target child only] | র্নিদিষ্ট শিশু১ প্রতি মিনিটে কতবার শ্বাস-প্রশ্বাস নেয়? [Respiratory rate of index child1 (per minute)] □□ | | | | র্নিদিষ্ট শিশু১ মায়ের কত নম্বর সন্তান? [Birth order of index child1] □ | |
| **C.07.** | 4002. | 1. র্নিদিষ্ট শিশু (Target child)<br>2 .একই মায়ের অন্য শিশু(Another child of same mother)<br>3. একই বাড়ীর অন্য খানার শিশু (Child from | | 1 পুরম্নষ M<br>0 মহিলা F | দিন/মাস/বছর DD/MM/YYY | 1 = কার্ড দ্বারা নিশ্চিত হয়েছিল (Confirmed DOB by valid vaccination/health card)<br>2 = মা/আত্মীয় দ্বারা নিশ্চিত হয়েছিল (Mother/Relative remembers DOB)<br>3 = উভয় (Both 1 & 2) | ☐1 ডায়রিয়া Diarrhea<br>☐2 পায়খানার নমুণা Stool specimen<br>☐3 রক্তের নমুণা Blood specimen<br>☐ 4 Target child<br>☐ 5 Missing/absent |

| | | | | | | | |
|---|---|---|---|---|---|---|---|
| | | the same bari)<br>4. একই খানার অন্য মায়ের শিশু(Another child of another mother in the same HH) | | | | 4 = ঘটনা দ্বারা নিশ্চিত হয়েছিল (Estimated DOB with 2 and event calendar) | ☐ 6 New birth<br>☐ 7 Death<br>☐ 8 Migration in |
| | 4021_1<br>[For target child only] | নির্দিষ্ট শিশু২ প্রতি মিনিটে কতবার শ্বাস-প্রশ্বাস নেয়?<br>[Respiratory rate of index child1 (per minute)]<br>☐☐ | | | | নির্দিষ্ট শিশু২ মায়ের কত নম্বর সন্তান?<br>[Birth order of index child1]<br>☐ | |

#### Wash Benefit Module 2 DIARRHEA AND ILLNESS SYMPTOMS (Children < 36 months)

৩৬ মাসের নীচের যে সকল শিশু খানায়/বাড়ীতে বাস করে (বেইজ লাইলের জন্য) । Administer to: Children < 36 months living in a study compound at baseline. This may include the child in the household or in other households in the bari.

PDA TO OPEN CHILD ID FROM **DIARRHEA COHORT**

যদি উত্তরদাতার 0-36 মাসের শিশু না থাকে তাহলে এখানেই শেষ করুনন নং প্রশ্ন যেতে হবে | IF study compound DOES NOT HAVE A CHILD 0-36 MONTHS → **SKIP TO END**

| উত্তরদাতা শিশুর প্রধান পরিচর্যাকারী Respondent: Child's primary caregiver | | | | | | |
|---|---|---|---|---|---|---|
| | WB M 2 | আপনাকে ধন্যবাদ । আমি এখন আপনাকে আপনার শিশুর স্বাস্থ্য বিষয়ে কিছু প্রশ্ন জিজ্ঞাসা করব। দয়া করে আপনি আমাকে ঠিকমতো প্রত্যক প্রশ্নের জবাব দিবেন। যদি উত্তর না জানা থাকে তখন বলবেন জানি না। শিশুটির নাম ধরে শুরু করতে হবে। Thank you. Now I am going to ask you some questions about the health of your [child/children]. Please answer each question as accurately as you can. If you don't know the answer to a question, say "I don't know." We will begin with [NAME OF CHILD 1]. | | | | |
| | | | **A** | **B** | **C** | **D** |
| | | **শিশুটির নাম বলুন এবং তার লক্ষণ গুলো ছিল কী?Did [NAME] have [SYMPTOM] :** | আজ **Today** ১=হ্যাঁ YES ০=না NO ৯৯৯=জানি না DK | গতকাল **Yesterday** ১=হ্যাঁ YES ০=না NO ৯৯৯=জানি না DK | গত পরশুদিন **Day before Yesterday** ১=হ্যাঁ YES ০=না NO ৯৯৯=জানি না DK | গত ৭দিনে **In the last 7 days** (since this day last week) ১=হ্যাঁ YES ০=না NO ৯৯৯=জানি না না DK |
| **C.201** | 201. | জ্বর Fever | | | | |
| **C.202** | 202. | পাতলা পায়খানা Diarrhea | | | | |
| **C.203** | 203. | ২৪ ঘন্টায় ৩ বার বা তার বেশী পায়খানা করেছে কিনা? 3 or more bowel movements in 24 hours | | | | |
| **C.204** | 204. | দৈনিক পায়খানায় সংখ্যা Number of bowl movements each day | | | | |
| **C.205** | 205. | পানির মত বা নরম পায়খানা? Watery or soft stool (unformed) | | | | |
| **C.206** | 206. | পায়খানায় রক্ত Blood in the stool | | | | |

| | | | |
|---|---|---|---|
| **C.207** | 207. | শরীর বা মাথায় চুলকানি, রেশ?Skin rash (anywhere on the body) | |
| **C.208** | 208. | দীর্ঘ কাশি Constant cough | |
| **C.209** | 209. | নাক বন্ধ/নাক দিয়ে পানি পড়া?Congestion / runny nose | |
| **C.210** | 210. | শ্বাস নেওয়ার সময় শোঁ শোঁ করে আওয়াজ বা শ্বাসকষ্ট? (যে কোন ধরনের শ্বাসকষ্ট, তবে নাক বন্ধ থাকার কারণে শ্বাসকষ্ট হলে তা বাদ দিতে হবে) Panting / wheezing / difficulty breathing | |
| **C.211** | 211. | দাগ, থেঁৎলে যাওয়া বা কাটা Bruising, scrapes or cuts | |
| **C.212** | 212. | দাঁতের ব্যাথাToothache / teething | |
| **C.213** | 213. | *C.202 (Diarrhea)নং-এর* উত্তর যদি হ্যাঁ হয় তখন ডায়রিয়া অংশ শুরু করতে হবে।সময় দিনে বা সপ্তাহে রেকর্ড করুন যদি ১৪ দিনের কম হয় তখন উত্তর দিনে আনতে হবে *If answered Yes to C.202 (Diarrhea):* When did the diarrhea start?<br>Record length of time in days or weeks. If < 14 days, record the response in days. | A ☐☐ B☐<br>1 দিন আগে Days ago<br>2 সপ্তাহ আগে Weeks ago |
| | | \|____\| Is absent? | 1= Yes<br>0= No |
| | | \|____\|Is not applicable? | 1= Yes<br>0= No |

| **Wash Benefit Module 3 DEWORMING** | | | |
|---|---|---|---|
| Administer to:<br>৩৬ মাসের নীচের শিশু Children < 36 months at enrollment **[PDA to open DIARRHEA COHORT]** | | | |
| **উত্তরদাতা শিশুর প্রধান পরিচর্যাকারী বা** গর্ভবতী মা Respondent: Child's primary caregiver or pregnant mother.<br>**আমি এখন** কৃমির ঔষধ যা নেয়া হয়েছে এ বিষয়ে আপনাকে জিজ্ঞাসা করব Now I would like to ask you about any medications you might have taken for intestinal worms. | | | |
| C.301 | 301. | গত ৬ মাসের মধ্যে শিশুটি (নাম বলুন) কোন কৃমিনাশক ঔষধ খেয়েছিল কী? In the last six months, has [NAME] received a pill or drug for intestinal worms? | [1] হ্যাঁ YES<br>[2] না NO → **Skip to 305**<br>[99] জানি না/নিশ্চিত না DK/NOT SURE → **Skip to 305** |
| C.302 | 302. | কোথা থেকে কৃমিনাশক ঔষধ খেয়েছিল?Where did [NAME] receive the drug for intestinal worms? | [1] বাড়ীতে/গ্রামে AT HOME/IN VILLAGE<br>[2] হাসপাতাল/স্বাস্থ্যকেন্দ্র AT A CLINIC OR HEALTH FACILITY<br>[3] স্কুলে AT A SCHOOL<br>[99] জানি না/নিশ্চিত না DK/NOT SURE |
| C.303 | 303. | কৃমিনাশক ঔষধটি কোন বড় প্রচারনার মাধ্যমে খেয়েছিল কী?Did [NAME] receive the drug as part of a larger campaign? | [1] হ্যাঁ YES<br>[2] না NO<br>[99] জানি না/নিশ্চিত না DK/NOT SURE |
| C.304 | 304. | আনুমানিক কত সময় আগে কৃমিনাশক ঔষধ খেয়েছিল? Approximately how long ago did [NAME] receive the drug?<br><br>MARK "99" FOR MONTHS IF DK/NOT SURE | \|__\| মাস(০-৬)MONTHS (0-6)<br>\|__\|__\|সপ্তাহ WEEKS |

| | | | |
|---|---|---|---|
| C.305 | 305. | শিশুটি কী কোন ময়লা/মাটি খেয়েছে ? Has the [NAME] eaten any dirt or soil?<br>*Ask for each recall period:* | |
| | 1 | 1. আজ Today | ☐ 1 হ্যাঁ Yes ☐ 2 না No<br>☐ 99 জানি না/নিশ্চিত না Don't know |
| | 2 | 2. গতকাল yesterday | ☐ 1 হ্যাঁ Yes ☐ 2 না No<br>☐ 99 জানি না/নিশ্চিত না Don't know |
| | 3 | 3. গত পরশুদিন Day before yesterday | ☐ 1 হ্যাঁ Yes ☐ 2 না No<br>☐ 99 জানি না/নিশ্চিত না Don't know |
| | 4 | 4. গত ৭দিনে In the past 7 days (since this day last week?) | ☐ 1 হ্যাঁ Yes ☐ 2 না No<br>☐ 99 জানি না/নিশ্চিত না Don't know |
| | 306. | প্রশ্ন করুন: আনুমানিক কত দিন আগে আপনি এন্টিবায়োটিক খেয়েছেন? (উদাহরণ দিন) Ask: "How long ago did you take any antibiotics?" [Provide examples] (focusing on the mother for baseline because samples for microbiome is being tested for moms only for baseline) | \|__\|__\| দিন Days<br><br>\|__\|__\| সপ্তাহ Weeks<br><br>\|__\|__\| মাস Months<br><br>[কখনই না বললে 88 লিখুন, জানি না বললে99 লিখুন 88 in all for NEVER, 99 for DK) |
| | 307. | গত তিন মাসে আপনি কতবার এন্টিবায়োটিক খেয়েছেন? Can you tell us how many times in the last 3 months you have used antibiotics?<br><br>Confirm 3 or longer recall time | \|___\| প্রতিবারের জন্য নীচের প্রশ্নটি রিপিট করুন (loop next question for each time) |
| | 308. | প্রতিবারে আপনি কতদিন করে এন্টিবায়োটিক খেয়েছেন? Please try and recall how many days you took the antibiotics (for each time). | \|___\| দিন days<br><br>99 for DK |
| | 309. | পর্যবেক্ষণ: শিশুটি জুতা পড়েছিল কী? Observe: Is the child wearing shoes at the time of the interview? | 1. হ্যাঁ Yes<br>0. না No<br>88. প্রযোজ্য নয় Not applicable |
| | | \|____\| Is absent? | 1= Yes<br>0= No |
| | | \|____\|Is not applicable? | 1= Yes<br>0= No |

**WASH Benefits Module 4: Anthropometry**

| IDENTIFICATION | | Date: | |
|---|---|---|---|
| **0.1. Data ID:** | ☐☐☐☐☐ | | |
| **0.2. Bari ID:** | ☐ | **0.3. HOUSEHOLD ID:** | ☐ |

| 0.4. CHILD ID | ☐☐☐☐☐☐ |
|---|---|

| | | |
|---|---|---|
| **C.401 FA ID** | ## | •••• |
| **C.402 Name of FA** | Full Name | |
| **C.403**<br>**Is mother wearing heavy clothing during weight measurement?** | 1 = Light clothing<br>2 = Light clothing plus sweater<br>3 = Heavy clothing | •• |
| **C.404**<br>**Weight of Mother**<br>**Measurement #1** | Weight<br>(kg) | •••• ••••<br>• |
| **C.405**<br>**Weight of Mother**<br>**Measurement #2** | Weight<br>(kg) | •••• ••••<br>• |
| **C.406**<br>**Weight of Mother**<br>**Measurement #3**<br>(If difference between measures 1 & 2 is ≥ 0.1 kg) | Weight<br>(kg) | •••• ••••<br>• |
| **C.407**<br>**Is child wearing clothing during weight measurement?** | 0 = No Clothes<br>1 = Only Tops<br>2 = Only Pants<br>3 = Both Tops & Pants | •• |
| **C.408**<br>**Weight of Mother + Child**<br>**Measurement #1** | Weight<br>(kg) | •••• ••••<br>• |
| **C.409**<br>**Weight of Mother + Child**<br>**Measurement #2** | Weight<br>(kg) | •••• ••••<br>• |

| | | |
|---|---|---|
| **C.410**<br>**Weight of Mother + Child**<br>**Measurement #3**<br>(If difference between measures 1 & 2 is ≥ 0.1 kg) | Weight<br>(kg) | ●●●● ●●●●<br>● |
| **C.411 – C.413** **(Reserved for child weight measurement without mother (follow-up visits))** | | |
| **C.414**<br>**Length of Child**<br>**Measurement #1** | Length<br>(cm) | ●●●●●● ●●<br>● |
| **C.415**<br>**Length of Child**<br>**Measurement #2** | Length<br>(cm) | ●●●●●● ●●<br>● |
| **C.416**<br>**Length of Child**<br>**Measurement #3**<br>(If difference between measures 1 & 2 is ≥ 0.5 cm) | Length<br>(cm) | ●●●●●● ●●<br>● |
| **C.417**<br>**Length Measurement Method** | Child was:<br>1 = lying (recumbent)<br>2 = standing | ●● |
| **C.418**<br>**Head Circumference**<br>**Measurement #1** | Circumference<br>(cm) | ●●●● ●●<br>● |
| **C.419**<br>**Head Circumference**<br>**Measurement #2** | Circumference<br>(cm) | ●●●● ●●<br>● |
| **C.420**<br>**Head Circumference**<br>**Measurement #3**<br>(If difference between measures 1 & 2 is ≥ 0.5 cm) | Circumference<br>(cm) | ●●●● ●●<br>● |
| **C.421**<br>**Does the child have swollen feet (bi-pedal edema)?** | 1 = Yes (**>> Referral**)<br>2 = No | ●● |
| **C.422**<br>**Length of Mother**<br>**Measurement #1** | Length<br>(cm) | ●●●●●● ●●<br>● |

| | | |
|---|---|---|
| **C.423**<br>**Length of Mother**<br>**Measurement #2** | Length<br>(cm) | •••••• ••<br>• |
| **C.424**<br>**Length of Mother**<br>**Measurement #3**<br>(If difference between measures 1 & 2 is ≥ 0.5 cm) | Length<br>(cm) | •••••• ••<br>• |

#### Wash Benefit Module 5 Vaccination [Open for target child and new births]

**Vaccination** (টিকাদান)

V101. ☐ Do you have a card where [NAME'S] vaccinations are written down? [*If YES*: May I see it please?] (আপনার শিশুর (নাম বলুন) কোন টিকা কার্ড আছে কী? হ্যাঁ হলে বলুন: আমি কী টিকা কার্ডটি দেখতে পারি?)

1 Yes, seen (হ্যাঁ, দেখাতে পেরেছে) **(Skip to V104)** (**V104** নং প্রশ্নে চলে যান)

2 Yes, not seen (হ্যাঁ, দেখাতে পারে নাই)

3 No card (কোন টিকা কার্ড নাই)

V102. ☐ Did [NAME] ever receive any vaccinations to prevent him/her from getting diseases, including vaccinations received in a national immunization campaign? (আপনার শিশু [নাম বলুন] রোগ থেকে রক্ষা পাওয়ার জন্য কখনও কোন টিকা বা জাতীয় টিকা দিবসে কোন টিকা নিয়েছিল কী?)

1 Yes (হ্যাঁ)

2 No (না) (Skip to C601)

99 Don't know / not sure (জানি না/নিশ্চিত না) (Skip to C601)

V103. *For children without a vaccination card* (যে শিশুর কোন টিকা কার্ড নেই)

Please tell me if [NAME] received any of the following vaccinations: (আপনার শিশু [নাম বলুন] নিম্নলিখিত কোন টিকা নিয়ে থাকলে আমাকে তা বলুন)

| | | |
|---|---|---|
| 1 | BCG vaccination against tuberculosis that is an injection in the arm or shoulder that usually causes a scar? (যক্ষা প্রতিরোধের জন্য বিসিজি-র টিকা দেওয়ার কারণে বাহু বা কাধে ক্ষত চিহ্ন আছে কী?) | ☐ 1. Yes (হ্যাঁ)<br>☐ 0. No (না)<br>☐ 88. N/A (প্রযোজ্য নয়)<br>☐ 99. Don't Know (জানি না) |
| 2 | Polio vaccine, that is, (pink) drops in the mouth? (মুখে কোন পোলিও টিকার [ গোলাপী রং] ফোঁটা দেওয়া হয়েছিল কী?) | ☐ 1. Yes (হ্যাঁ)<br>☐ 0. No (না) (skip to 103.4)<br>☐ 88. N/A (প্রযোজ্য নয়) (skip to 103.4)☐ 99. Don't Know (জানি না) (skip to 103.4) |
| 3 | How many times was the polio vaccine received? (কতবার পোলিও টিকা দেওয়া হয়েছিল?) | ☐ 9 = DK (জানি না) |
| 4 | A Penta (DPT + Hib vaccination + Hepatitis-B vaccination ) vaccination, that is, an injection given in the thigh or buttocks, sometimes at the same time as the polio drops? (উরুতে বা পাছাতে কোন ডিপিটি-র টিকা দেওয়া হয়েছিল কী, যা অনেক সময় পোলিও টিকার সাথে দেওয়া হয়?) | ☐ 1. Yes (হ্যাঁ)<br>☐ 0. No (না) (skip to 103.6)<br>☐ 88. N/A (প্রযোজ্য নয়) (skip to |

| | | |
|---|---|---|
| | | 103.6)<br>☐ 99. Don't Know (জানি না) (skip to 103.6) |
| 5 | How many times was the Penta vaccine received? (কতবার ডিপিটি-র টিকা দেওয়া হয়েছিল?) | ☐ 9 = DK(জানি না) |
| 6. | A MR (measles-rubella) injection, that is, a shot in the arm at the age of 9 months or older to prevent him/her from getting measles and rubella? (হাম প্রতিরোধের জন্য ৯ মাস বয়সে হামের টিকা বাহুতে দেওয়া হয়েছে কী?) | ☐ 1. Yes (হ্যাঁ)<br>☐ 0. No (না)<br>☐ 88. N/A (প্রযোজ্য নয়)<br>☐ 99. Don't Know (জানি না) |
| 7 | A measles injection, that is, a shot in the arm at the age of 15 months or older to prevent him/her from getting measles? (হাম প্রতিরোধের জন্য ১৫ মাস বয়সে হামের টিকা বাহুতে দেওয়া হয়েছে কী?) | ☐ 1. Yes (হ্যাঁ)<br>☐ 0. No (না)<br>☐ 88. N/A (প্রযোজ্য নয়)<br>☐ 99. Don't Know (জানি না) |
| 11 | Within the last six months, has [NAME] received a vitamin A dose (like this / any of these)?<br>SHOW COMMON TYPES OF AMPULES/CAPSULES/SYRUPS (গত ছয় মাসের মধ্যে শিশুটি (নাম বলুন) এই রকম/অনেকটা এই রকমের কোন 'ভিটামিন এ' ক্যাপসুল খেয়েছিল কী? (ভিটামিন এ ক্যাপসুল দেখান) | ☐ 1. Yes (হ্যাঁ)<br>☐ 0. No (না)<br>☐ 88. N/A (প্রযোজ্য নয়)<br>☐ 99. Don't Know (জানি না) |

V104. *For children with a vaccination card:* (যে শিশুর টিকা কার্ড আছে)

Copy the vaccination date for each vaccine from the card. (টিকা কার্ড থেকে প্রতিটি টিকা প্রদানের তারিখ লিখুন)

Record '000' if timely vaccination was not given.

Record "888" in the DAY column if card shows that a vaccination was given, but no date is was recorded. (যদি টিকা কার্ডে উলেস্নখ থাকে টিকা দিয়েছে কিন্তু কোন তারিখ না থাকলে দিনের কলামে [**DAY column**] "888"রেকর্ড করম্নন)

Record "999" in the DAY column if vaccination not given due to Not Applicability (N/A) . প্রযোজ্য নয় এই কারণে টিকা না দিলে দিনের কলামে [**DAY column**] **"999"**]রেকর্ড করম্নন )

| | | DAY | MONTH | YEAR |
|---|---|---|---|---|
| 1 | BCG | I__I__I | I__I__I | I__I__I |
| 2 | POLIO 0 | I__I__I | I__I__I | I__I__I |
| 3 | POLIO 1 | I__I__I | I__I__I | I__I__I |
| 4 | POLIO 2 | I__I__I | I__I__I | I__I__I |
| 5 | POLIO 3 | I__I__I | I__I__I | I__I__I |
| 6 | Penta 1 | I__I__I | I__I__I | I__I__I |
| 7 | Penta 2 | I__I__I | I__I__I | I__I__I |
| 8 | Penta 3 | I__I__I | I__I__I | I__I__I |
| 9 | MR | I__I__I | I__I__I | I__I__I |
| 10 | MEASLES | I__I__I | I__I__I | I__I__I |
| 11 | VITAMIN A (with measles vaccine) | I__I__I | I__I__I | I__I__I |

#### Wash Benefit Module 6 CHILD FOOD FREQUENCY

[Open for target child]

Administer to:Target Children(নির্দ্দিষ্ট শিশুর জন্য প্রযোজ্য)

Respondent: Child'sprimarycaregiver.( উত্তরদাতা শিশুর প্রধান পরিচর্চাকারী)

**ভূমিকা**

আমি এখন আপনার শিশুর (নাম) খাদ্যগ্রহণ সংক্রান্ত বিষয়ে কিছু প্রশ্ন করতে চাই। প্রথমে আমার জানা দরকার যে গতকাল এবং গত ৭ দিনের মধ্যে শিশুর (নাম) খাদ্যগ্রহণ সম্পর্কে বলতে পারবেন।

**Introduction** Now I would like to ask you some questions about feeding [NAME]. First I need to know if you will be able to tell me about feeding [NAME] yesterday and over the last week.

**C.601.** আপনি কি জানেন শিশু (নাম) গতকাল কি কি খেয়েছে? Do you know what [NAME] consumed yesterday?

- ☐ 1 হ্যাঁ (৬০৩ নং প্রশ্নে যান) Yes [skip C.603]
- ☐ 2 না No

**C.602.** যদি না হয় তবে এমন কেউ কি আছে যে জানে শিশু কি খেয়েছিল এবং যিনি আমাদের প্রশ্নসমূহের উত্তর দিয়ে সহায়তা করবে? *If No,* Is there someone else who knows what the child ate, who can sit with us today and help answer questions?

- ☐ 1 হ্যাঁ Yes
- ☐ 2 না No

**C.603.** শিশুর সাথে উত্তরদাতার সম্পর্ক Respondent relationship to the participating child

- ☐ 1 মা Mother
- ☐ 2 বাবা Father
- ☐ 3 বোন Sister
- ☐ 4 দাদী Grandmother
- ☐ 77 অন্যান্য (নির্দ্দিষ্ট করুন) _________________

  Other (specify) ________________

শিশু যে খাবার খেয়েছে যেমন বুকের দুধ, তরল খাবার এবং অন্যান্য খাবার **Breastfeeding, Liquids, and Foods Eaten by the Child**

**C.604.** জন্মের কতক্ষণ পর শিশুকে (নাম) বুকের দুধ দিয়েছিলেন? **(উত্তর গুলো পড়ে শোনান)** How long after the birth did you first put [name] to the breast?

- ☐ 0 প্রথম আধা ঘন্টার মধ্যে within the first half hour
- ☐ 1 প্রথম এক ঘন্টার মধ্যে Within the first hour
- ☐ 2 এক ঘন্টার বেশী কিন্তু ২৪ ঘন্টার মধ্যে More than one hour but less than 24 hours
- ☐ 3 ২৪ ঘন্টার বেশী More than 24 hours
- ☐ 88 কখনো না/প্রযোজ্য নয় (৬০৮ নং প্রশ্নে যান) Never breastfed / Not Applicable (**Skip to C.608**)
- ☐ 99 জানি না/ মনে নেই Don't know / not sure

**C.604a** জন্মের পরপরই শিশুকে (নাম) শাল দুধ দিয়ে কি বুকের দুধ খাওয়ানো শুরু করেছিলেন? When [NAME] was born, did you start breastfeeding him/her immediately with the very first breast milk (colostrum)?

- ☐ 1 হ্যাঁ Yes

☐ 2 না No

**C.605.** শিশুটি কি এখনো বুকের দুধ পান করছে, নাকি সে সম্পূর্নভাবে বারতি খাবার খাচ্ছে?( যদি শিশু বুকের দুধ খায় তাহলে ১ কোড করুন)
Is the baby still breastfeeding, or is he/she completely weaned?

☐ 1 এখনো বুকের দুধ পান করে ( ৬০৭ নং প্রশ্নে যান) **Still**, breastfeeding (**Skip to C.607)**

☐ 2 বাড়তি খাবার খাচ্ছে. কোন বুকের দুধ পাচ্ছে না (weaned, not receiving any breast milk)

**C.606.** শিশুটির (নাম) যখন সর্বশেষ বুকের দুধ পান করেছিল,তখন তার বয়স কত ছিল ? How old was [NAME] the last time he/she was breastfed?

৯৯= জানি না / নিশ্চিত না 99 Don't know / not sure

☐☐ দিন

☐☐ মাস

☐☐ Skip to B.607x

**C.607.** এখন আমাকে বলুন শিশুটি (নাম) গত ২৪ ঘন্টা কতবার বুকের দুধ পান করেছে ? Now I would like you to tell me how many times [NAME] breastfed yesterday.

☐☐ বার TIMES

**B. 607x** আপনি কি কখনো শিশুকে (নাম) গোসলের পানির ফোটা খাইয়েছেন ?

**Have you ever given the bath water drop to [Name]?**

**1 হ্যাঁ Yes**

**0 না No**

**C. 607a** বুকের দুধ ছাড়া শিশুকে (নাম)সর্বপ্রথম কি খাওয়ানো হয়েছে ? What was the very first thing that [Name] consumed apart from breast milk?

☐ ১ মধু Honey

☐ ২ চিনির পানি Sugary water

☐ ৩ তাল মিছরি Tal misree ( Crystalline sugar cube from palm sugar)

☐ ৪ শুধু পানি Plain water

☐ ৫ কিছুই না ( ৬০৭ সি নং প্রশ্নে যান) Nothing (**Skip to C.60 7c)**

**☐ 6 সুজি বা অন্য কোন বাড়তি খাবার Suji or any other complementary food**

☐ ৭৭ অন্যান্য (নির্দিষ্ট করুন) ________________ others (specify) ________________

**C. 607b** শিশুটি কোন বয়সে এই খাবার খেয়েছিল ? At what age did [Name] consume this item?

☐☐ **দিন** [যদি মা দিন বলতে পারে ] DAYS [if mother mentioned days] অথবা OR

☐☐ মাস [যদি মা মাস বলতে পারে ] MONTHS [if mother mentioned months]

**C. 607c** গত সপ্তাহে কি কোন দিন শিশুর (নাম) ক্ষুধামন্দা হয়েছিল? In the past week, were there any days that [NAME] lost his/her appetite?

| 1. উত্তর Response | 2. কতদিন How many days? |
| --- | --- |
| ☐ ১ হ্যাঁ Yes | ☐ দিন Days |

| |
|---|
| ☐ ২ না (৬০৮ নং প্রশ্নে যান ) No [skip to 608]<br>☐ ৯৯ জানি না (৬০৮ নং প্রশ্নে যান )<br>99 Don't Know [skip to 608] |

এখন গতকাল সারাদিন আর সারারাতে (শেষ ২৪ ঘন্টা, সাক্ষাৎকারের সময় থেকে) আপনার শিশু [নাম] কি কি তরল খাবার খেয়েছে আমি সে সম্পর্কে কিছু জানতে চাইবো। আপনার শিশু যে তরল খাবার খেয়েছে আমি সে সম্পর্কে জানতে চাই, তা অন্য যে কোন খাবারের সাথে মিশিয়ে হোক না কেন ।

তালিকা থেকে প্রশ্নগুলো পড়ুন এবং সঠিক ঘরে টিক চিহ্ন দিন

Now I would like to ask you about liquids that [NAME] may have had yesterday during the day or at night (last 24hours, starting from the time interview). I am interested in whether your child had the item even if it was combined with other foods.

*For each item on the list, read the question below and tick the appropriate box.*

**C.608.** কোন পানীয় পান করেছে কিনা? (তালিকা থেকে প্রত্যেকটি প্রশ্ন পড়ে শোনান )

Did [NAME] drink/have any [ITEM FROM LIST]? *Read question 14 times, once for each item*

| খাবার এর নাম **Food Item** | এই পানীয় টা খেয়েছে/ পান করেছে কিনা?<br>**Drink / take / eat this item?** | গতকাল দিনে বা রাতে কত বার (নাম) এই খাবারটি খেয়েছে/ পান করেছে ? (তালিকা থেকে)<br>**How many times yesterday during the day or night did [NAME] consume [ITEM FROM LIST]?** |
|---|---|---|
| 1. পানি Water? | ☐ ১ হ্যাঁ Yes<br>☐ ২ না No<br>☐ ৯৯ জানি না<br>99 Don't Know | ☐☐ বার Times |
| 2. চিনির পানি Sugar water? | ☐ ১ হ্যাঁ Yes<br>☐ ২ না No<br>☐ ৯৯ জানি না<br>99 Don't Know | ☐☐ বার Times |
| 3. প্রাণিজ দুধ, কৌটার দুধ, গুড়া দুধ?<br>Milk, including any animal fresh milk, milk in tin or box, or powdered milk? | ☐ ১ হ্যাঁ Yes<br>☐ ২ না No<br>☐ ৯৯ জানি না<br>99 Don't Know | ☐☐ বার Times |
| 4. শিশু খাদ্য যেমন ন্যান, ল্যাকটোজেন, বায়োমিল অথবা মাই বয়? Infant formula such as NAN or Lactogen or Biomeal or My Boy? | ☐ ১ হ্যাঁ Yes<br>☐ ২ না No<br>☐ ৯৯ জানি না<br>99 Don't Know | ☐☐ বার Times |
| 5. দুধ চা? Tea made with milk? | ☐ ১ হ্যাঁ Yes<br>☐ ২ না No<br>☐ ৯৯ জানি না<br>99 Don't Know | ☐☐ বার Times |

| | | | |
|---|---|---|---|
| 6. | রং চা? Tea made without milk? | ☐ ১ হ্যাঁ Yes<br>☐ ২ না No<br>☐ ৯৯ জানি না<br>99 Don't Know | ☐☐ বার Times |
| 7. | দই? Yogurt? | ☐ ১ হ্যাঁ Yes<br>☐ ২ না No<br>☐ ৯৯ জানি না<br>99 Don't Know | ☐☐ বার Times |
| 8. | দুধ বা পানি দিয়ে ফুটানো শস্য জাতীয় খাবার যেমন সুজি, চালের গুড়া ইত্যাদি ?<br>Thin Porridge or Barley? | ☐ ১ হ্যাঁ Yes<br>☐ ২ না No<br>☐ ৯৯ জানি না<br>99 Don't Know | ☐☐ বার Times |
| 9. | ফলের রস? Fruit juice | ☐ ১ হ্যাঁ Yes<br>☐ ২ না No<br>☐ ৯৯ জানি না<br>99 Don't Know | ☐☐ বার Times |
| 10. | পানির মত পাতলা স্যুপ [কোন শক্ত উপাদান ছাড়া]<br>Clear broth [soup without solid ingredient] | ☐ ১ হ্যাঁ Yes<br>☐ ২ না No<br>☐ ৯৯ জানি না<br>99 Don't Know | ☐☐ বার Times |
| 11. | ডাবের পানি (অন্য কিছুর সাথে না মিশিয়ে)<br>Coconut water (plain) | ☐ ১ হ্যাঁ Yes<br>☐ ২ না No<br>☐ ৯৯ জানি না<br>99 Don't Know | ☐☐ বার Times |
| 12. | হরলিকস বা অন্য কোন সম্পুরক পানিয় Horlicks (or any other supplementary drinks) | ☐ ১ হ্যাঁ Yes<br>☐ ২ না No<br>☐ ৯৯ জানি না<br>99 Don't Know | ☐☐ বার Times |
| 13. | Others: Specify_______________ | ☐ ১ হ্যাঁ Yes<br>☐ ২ না No<br>☐ ৯৯ জানি না<br>99 Don't Know | ☐☐ বার Times |

এর পর গতকাল সারাদিন আর রাতে (শেষ ২৪ ঘন টা, সাক্ষাৎকারের সময় থেকে) আপনার শিশু [নাম]কি কি খেয়েছে আমি সে সম্পর্কে কিছু প্রশ্ন জিজ্ঞাসা করবো। আপনার শিশু যা কিছু খেয়েছে,আমি সে সম্পর্কে সব কিছুই জানতে চাই, তা বাসায় বা অন্য যে কোন জায়গায় হোক না কেন ।

নির্দেশনার জন্য আলাদা পৃষ্ঠা ব্যবহার করুন,যা মা কেমনে করতে সাহায্য করবে। মাকে বলতে দিন শিশুটি কি খেয়েছিল। শিশু টি যা খেয়েছিল , প্রতিটি খাবার ( অথবা উপাদান) এ গোল চিহ্ন দিন এবং হ্যাঁ তে টিক চিহ্ন দিন

Next I would like to ask you some questions about the foods that [NAME] ate yesterday during the day or at night (last 24hours, starting from the time interview). I would like to know everything that [NAME] ate, whether at home or someplace else. *Use the separate page of instructions, with questions to help the mother remember. Do not read the list below. Let the mother tell you what the child ate. Circle each food (or ingredient) that the child ate, and tick "[√] 1 Yes" for that food group.*

<u>নিম্নের তালিকাটি পড়ে শুনাবেন না</u>

**C.609.**

| |
|---|
| শিশু গতকাল যে খাবার/খাদ্য তৈরীর উপাদান খেয়েছে(ঝোল অথবা সিদ্ধ যা ই হোক না কেন)<br>Foods/ingredients in recipes (may be in a sauce or porridge) eaten by the child |

| | | |
|---|---|---|
| | yesterday | |
| 1 | খিচুড়ী Khichuri<br>( খিচুড়ী কি কি দিয়ে রান্না হয়েছে জেনে নিয়ে ফুড গ্রপ অনুযায়ী নীচে কোড করুন)<br>(tick ‘yes’ and ask for the ingredients and code below according to food group) | ☐ ১ হ্যাঁ Yes<br>☐ ২ না No |
| 2 | পানিতে বা দুধে সিদ্ধ করা শস্য জাতীয় খাবার যেমন: সুজি,ভাত, রুটি, নুডুলস, অন্যান্য শস্য জাতীয় দানাদার খাদ্য<br>Porridge Rice Bread/ Roti Noodles Other foods made by grain | ☐ ১ হ্যাঁ Yes<br>☐ ২ না No |
| 3 | মিষ্টি কুমড়া, গাজর, অন্যান্য হলুদ সবজী<br>Pumpkin Carrots Other yellow vegetable | ☐ ১ হ্যাঁ Yes<br>☐ ২ না No |
| 4 | আলু সাদা মিষ্টি আলু<br>Potato White sweet potato | ☐ ১ হ্যাঁ Yes<br>☐ ২ নার No |
| 5 | মিষ্টি কুমড়া শাক, সরিষা শাক, মটর শুটি শাক, পুইঁ শাক, অন্যান্য গাঢ় সবুজ শাক<br>Pumpkin Mustard Bean Pigeon pea/ Motorshuti Other dark green<br>leaves leaves leaves leaves<br>leaves | ☐ ১ হ্যাঁ Yes<br>☐ ২ না No |
| 6 | পাকা আম , পাকা পেঁপেঁ<br>Ripe mango Ripe papaya | ☐ ১ হ্যাঁ Yes<br>☐ ২ না No |
| 7 | কলা, আনারস, পেয়ারা, আপেল, আঙ্গুর, কমলা অন্যান্য ফল<br>Banana Pineapple Guava Apple Grape Orange Other fruit | ☐ ১ হ্যাঁ Yes<br>☐ ২ না No |
| 8 | টমেটো, পিঁয়াজ , মাশরুম, ঢেড়স, তাজা সীম/মটর শুটি অন্যান্য সবজি<br>Tomato Onion Mushroom Lady’s finger Fresh bean / Motorshuti other Vegetable | ☐ ১ হ্যাঁ Yes<br>☐২ না No |
| 9 | কলিজা গিলা<br>Liver Kidney Heart | ☐ ১ হ্যাঁ Yes<br>☐ ২ না No |
| 10 | যে কোন মাংস ,পশু পাখিসহ<br>Any type of meat / flesh, including from birds and animals | ☐ ১ হ্যাঁ Yes<br>☐ ২ না No |
| 11 | যে কোন ধরণের ডিম<br>Any type of egg | ☐ ১ হ্যাঁ Yes<br>☐ ২ না No |
| 12 | তাজা মাছ, শুটকি মাছ, অন্যান্য মাছ / সামুদ্রিক খাবার<br>Fresh fish Dried fish Other fish / seafood | ☐ ১ হ্যাঁ Yes<br>☐ ২ না No |
| 13 | সীম ,ডাল,সয়া, চীনাবাদাম, কেশর, ভারীচীনাবাদাম অন্যান্য যে কোন ডাল বা বাদাম জাতীয় খাদ্য<br>Beans Peas / Soya Groundnut Cashew Pounded Any other<br>Lentils<br>groundnut legume or nut | ☐ ১ হ্যাঁ Yes<br>☐ ২ না No |
| 14 | পনির দই অন্যান্য দুধ জাতীয় খাদ্য | ☐ ১ হ্যাঁ Yes |

| | | |
|---|---|---|
| | Cheese Yogurt Other milk products | ☐ ২ না No |
| 15 | উদ্ভিজ্জ তেল (ডালডা) পশুর চর্বি এক ধরনের মাখন<br>Vegetable oil Animal fat Margarine | ☐ ১ হ্যাঁ Yes<br>☐ ২ না No |
| 16 | চকলেট মিষ্টি/ ক্যান্ডি/ পিঠা মিষ্টি বিস্কুট<br>Chocolate Sweets / candies Cake Cookies / sweet biscuits Sugar | ☐ ১ হ্যাঁ Yes<br>☐ ২ না No |
| 17 | স্বাদ বৃদ্ধি কারক রসুন মসলা<br>Seasonings Garlic Spices Salt | ☐ ১ হ্যাঁ Yes<br>☐ ২ না No |
| 18 | চিংড়ি, কাকড়া<br>Prawns Crab | ☐ ১ হ্যাঁ Yes<br>☐ ২ না No |
| 19 | যদি খাদ্য তালিকায় না থাকে তাহলে নিচে খাবারের নাম লিখুন।<br>*If not on list above, write food(s) here and at bottom* | ☐ ১ হ্যাঁ Yes<br>☐ ২ না No |

**C.610.**

আপনি গতকাল সারাদিন আর রাতে (শেষ ২৪ ঘন্টা, সাক্ষাৎকারের সময় থেকে) আপনার শিশু [নাম] কি কি খেয়েছে তা বললেন [পিছনের পৃষ্ঠার গোল করা খাবারে নাম গুলো পড়ুন] এছাড়া আপনার শিশু [নাম] নাস্তাসহ আর কি কিছু খেয়েছে ?

You mentioned that [NAME] ate [read back circled foods on previous page] yesterday during the day or at night (last 24hours, starting from the time interview).

Did [NAME] have any other food at all, including snacks?

☐ ১ হ্যাঁ Yes

☐ ২ না No

যদি হ্যা হয়, তাহলে পূর্বের পৃষ্ঠার প্রশ্ন গুলো আবার করুন এবং গোল চিহ্ন দিন । মনে করা শেষ হলে, যদি সেই দলের কোন খাবার বা খাদ্য উপাদানে গোল করা না হয় তাহলে না তে টিক চিহ্ন দিন।

যদি C.609এবং C.610 এর সব উত্তর না হয় তাহলে C.611 নং প্রশ্নে যান ।

*If "yes", use the same probing questions and circle on the list on previous page.* ***At the end of the recall tick "[√] 2 No" if no food or ingredient is circled for that group.***

If C.609 and C.610 are all 'No' skip to C.611

**C. 610a**

গতকাল সারাদিন আর সারারাতে (শেষ ২৪ ঘন্টা, সাক্ষাৎকারের সময় থেকে) আপনার শিশু [নাম] কতবার শক্ত, আধা শক্ত আথবা নরম খাবার খেয়েছে তরল খাবার বাদে?

নোট: সকল ধরনের দুধ বা পানি দিয়ে ফুটানো শস্য জাতীয় খাবার ।

How many times did [NAME] eat solid, semi-solid, or soft foods other than liquids yesterday during the day or night (last 24hours, starting from the time interview)? *Note: include all forms of porridge, including thin porridge.*

☐☐ বার Times

আমি এখন আপনাকে আপনার শিশু [নাম] গত ৭দিনে [সাক্ষ্যাৎকারের দিন থেকে] কি কি খাবার খেয়েছে সে বিষয়ে কিছু প্রশ্ন করতে চাই।প্রতিটি খাবারএর কথা আমি জানতে চাইবো, গত ৭দিনের মধ্যে কতদিন আপনার শিশু নিম্নলিখিত খাবার গুলো খেয়েছেন তা বলুন।

Now I would like to ask you some questions about foods [NAME] ate in the last 7 days, since last [INTERVIEW DAY]. For each food I ask about, please tell me how many days in the last 7 days you think the child ate that food.

যদি শিশুটি (নাম) খাবার খেয়ে থাকে, এমনকি এটি যদি অন্যান্য খাবারের সংঙ্গে সংযুক্ত থাকে, উদাহরণস্বরূপ,যদি শিশুটি (নাম) মুরগির মাংসের ঝোল খায় যা পিয়াজ, টমেটো, মুরগির মাংস দিয়ে তৈরী তাহলে আমি যখন আপনাকে প্রশ্ন করবো মাংস সম্পর্কে তখন আপনি "হ্যাঁ" বলবেন আবার যখন আমি আপনাকে প্রশ্ন করবো সবজি সম্পর্কে তখন আপনি আবার "হ্যাঁ" বলবেন যদি আপনার শিশু (নাম) দুটো খাবার ইখেয়ে থাকে, কিন্তু যদি মুরগির মাংস আর সবজি কোনটাই না খেয়ে থাকে তাহলে "হ্যাঁ" বলবেন না কারণ সে এগুলা খায় নি।

I would like to know if [NAME] had the food, even if it was combined with other foods. For example, if [NAME] ate a sauce or relish made with chicken, onions, and tomatoes, you should say "yes" when I ask about meat, and again "yes" when I ask about vegetables. However, if [NAME] only had the broth, not the chicken or vegetables, do not say "yes" because they did not eat it.

তালিকার প্রতিটি খাবারের জন্য, প্রশ্নটি পরুন এবং উওর দাতা যে কয়দিন বলে সেই নাম্বার টি খালি ঘরে লিখুন (০-৭)

*For each item on the list, read the question below and fill in the number of days the respondent says (0-7).*

**C.611.**

গত সাত দিনে আপনার শিশু [নাম] কয়দিন এই খাবার খেয়েছে [ তালিকার খাবার]? **<u>(তালিকা থেকে প্রত্যেকটি প্রশ্ন পড়ে শোনান )</u>**

How many days in the last 7 days did [NAME] have [ITEM FROM LIST]?

| | খাবারগুলো (উপাদান অনুযায়ী ভাগ করা হয়েছে) যা শিশু গত সাত দিনে খেয়েছে Foods (in groups) eaten by the child in the last seven days | শিশুটি যে কয় দিন খাবার খেয়েছে =(০-৭)<br>Number of days food was eaten by child (0-7)<br>খেয়েছে,কিন্তু কয়দিন খেয়েছে তা জানেনা = ৬৬<br>Eaten, don't know how many days = **66**<br>খেয়েছে কি খায়নি জানেনা = ৯৯<br>Don't know if eaten or not = **99** |
|---|---|---|
| 1 | খিচুড়ী ( খিচুড়ী কি কি দিয়ে রান্না হয়েছে জেনে নিয়ে ফুড গ্রুপ অনুযায়ী নীচে কোড করুন), ভাত, রুটি<br>Khichuri (tick 'yes' and ask for the ingredients and code below according to food group), rice, bread? | I__I__I |
| 2 | মিষ্টি কুমড়া, গাজর ?<br>Pumpkin, carrots? | I__I__I |
| 3 | আলু, মিষ্টি আলু অথবা অন্যান্য যে কোন সাদা রং এর আলু জাতীয় খাদ্য?<br>Potato, sweet potatoes or any other white colored root or tuber? | I__I__I |
| 4 | গাঢ় সবুজ শাক যেমন মিষ্টি কুমড়া শাক, সরিষা শাক দিয়ে তৈরী ঝোল বা কোন মজাদার খাবার?<br>Any sauce or relish made with dark green leaves such as pumpkin leaves or mustard leaves? | I__I__I |
| 5 | পাকা আম বা পাকা পেঁপে?<br>Ripe mango or ripe papaya,? | I__I__I |
| 6 | অন্যান্য যে কোন ফল যেমন পেয়ারা ,আপেল,আঙ্গুর, কলা কমলা ?<br>Any other fruit such as, guava, apple, grape or banana, orange | I__I__I |
| 7 | অন্যান্য যে কোন সবজি যেমন টমেটো,পিঁয়াজ,মাশরুম,সীম,মটরশুটি ?<br>Any other vegetable such as tomato, onions, mushroom, beans/ green peas or any other? | I__I__I |
| 8 | যে কোন মাংস ,পাখি বা প্রাণীর মাংস সহ?<br>Any type of meat, including from birds or from animals? | I__I__I |
| 9 | যে কোন ধরণের ডিম ?<br>Any type of egg? | I__I__I |
| 10 | যে কোন ধরণের শুটকি মাছ অথবা তাজামাছ?<br>Any type of dried fish or fresh fish? | I__I__I |
| 11 | সীম বীচি, মটর,ডাল,চীনাবাদাম বা অন্যান্য বাদাম?<br>Any dishes made with beans, peas, lentils, groundnut, or other nuts,? | I__I__I |

| | | |
|---|---|---|
| 12 | যে কোন দুধ, পনির, দই বা অন্যান্য দুধের তৈরী খাদ্য?<br>Any milk, cheese, yogurt, or foods/drinks made with milk? | I__I__I |
| 13 | উদ্ভিজ্জ তেল, পশুর চর্বি , ঘি, মাখন বা অন্য যে কোন খাবার যা এই গুলো দিয়ে তৈরী ?<br>Vegetable oil, fat from animals, ghee, butter or any foods made with these? | I__I__I |
| 14 | মিষ্টি খাবার যেমন: চকলেট, মিষ্টি/ ক্যান্ডি, পিঠা অথবা মিষ্টি বিস্কুট<br>Sweet foods such as chocolate, sweets/candies, cake or cookies/sweet biscuits, mishti? | I__I__I |

আমি এখন আপনার কাছে আপনার শিশুর [নাম] শিশু খাদ্য এবং কিছু বিশেষ খাবার যা শিশুরা মাঝে মাঝে খায় সে বিষয়ে কিছু প্রশ্ন করতে চাই।যদি আপনি এই খাবার সম্পর্কে আগেই বলে থাকেন, তবে আনুগ্রহ করে আবার বলুন যাতে আমি এই বিশেষ খাবার গুলো নির্ভুল ভাবে লিখতে পারি ।

Now I would like to ask you about infant formula and about some special foods that are sometimes given to infants and small children. Even if you already told me about the food, please tell me again so I can be sure to write down these special foods.

## C.611a

#### আমি এখন জানতে চাই আপনার শিশু [নাম] গত ৭দিনের মধ্যে নিম্নলিখিত খাবারগুলো যা বাজার থেকে কেনা হয়েছে, সেগুলো কতদিন খেয়েছে ।

Now I would like to ask you during last 7 days how many days [Name] has eaten any of these foods items bought from the market.

| Sl no. | Variable | ফুড গ্রুপ<br>Food categories | তালিকা থেকে প্রশ্নগুলো পড়ুন এবং সঠিক (নিকটতম খাদ্য) ঘরে টিক চিহ্ন দিন<br>Name of the specific food item<br>(Select the closest food item, check box for "yes" or "no") | ০-৭ দিন<br>Days 0-7 | Values (options) |
|---|---|---|---|---|---|
| 1 | 611a_1 | কোমল পাণীয়<br>Soft drinks | কোলা, সেভেন আপ, ফানটা<br>Cola, 7 up, Fanta | | **0 to 7** |
| 2 | 611a_2 | ফলের রস<br>Fruit juice | আম, কমলা, লেবু, লিচ<br>Mango, Orange, Lemon, Lichi | | **0 to 7** |
| 3 | 611a_3 | বোতলজাত/ক্যান দুধ<br>Bottled or canned milk | চকলেট দুধ, মিষ্টি দুধ (টেট্রা প্যাক)<br>Chocolate milk, Sweetened milk (tetra pack) | | **0 to 7** |
| 4 | 611a_4 | দুগ্ধ জাতীয় খাবার<br>Milk products | আইসক্রিম, কুলফি, দই, মাঠা<br>Ice cream, kulfi, yogurt, matha | | **0 to 7** |

| | | | | | |
|---|---|---|---|---|---|
| 5 | 611a_5 | মিষ্টি খাবার<br>Sweet snacks/ sweet meat | চকলেট, ক্যান্ডি, ওয়েফার,লিচু, শণপাপড়ি, হাওয়াই মিঠাই, জিলাপি, মুড়ি-মোয়া, মুরলি, সন্দেশ, মিষ্টি<br>chocolate, candy, wafer, lichi, soan papri, hawai mithai, jilabi, murir moa, muroli, shondesh/mishti | | **0 to 7** |
| 6 | 611a_6 | নোনতা/ মশলাদার খাবার<br>Savory snacks | চিপস, চানাচুর, ডালভাজা, নিমকি, সিংগাড়া, সমুচা<br>chips, chanachur, dal vaja, nimki, singara, samosa | | **0 to 7** |
| 7 | 611a_7 | আচার<br>Pickles | আম, জলপাই, তেতুল, বরই, চালতা<br>Mango, Olive, Tamarinds, Boroi, Chalta | | **0 to 7** |

**C.612.** গত সাত দিনের মধ্যে কয়দিন আপনার শিশু [নাম] এই খাবার খেয়েছে [সাক্ষাৎকারের দিন থেকে] [তালিকার খাবার]? On how many days in the last 7 days, since last [INTERVIEW DAY], did [NAME] have any [ITEM FROM LIST]?

| | | |
|---|---|---|
| | শিশু খাদ্য এবং বিশেষ খাবার যা শিশু গত সাত দিনে খেয়েছে<br>Infant formula and special foods eaten by the child in the last seven days | শিশুটি যে কয় দিন খাবার খেয়েছে =(০-৭)<br>Number of days food was eaten by child (0-7)<br>খেয়েছে,কিন্তু কয়দিন খেয়েছে তা জানেনা = ৬৬<br>Eaten, don't know how many days = **66**<br>খেয়েছে কি খায়নি জানেনা = ৯৯<br>Don't know if eaten or not = **99** |
| 1 | শিশু খাদ্য যেমন ল্যাকটোজেন অথবা নান অথবা বায়োমিল,মাইবয় অন্যান্য?<br>Infant formula such as Lactogen or NAN or Biomeal, My Boy, or others? | I__I__I |
| 1.1 | অন্যান্য হলে ,নির্দিষ্ট করুন? If others, What type?<br>____________________________________ | |
| 2 | পানিতে বা দুধে সিদ্ধ করা শস্য জাতীয় খাবার যেমন: সুজি অথবা অন্যান্য খাবার যা ভূট্টার তৈরী, যেগুলো দোকানে কিনতে পাওয়া যায়? | I__I__I |

| | | |
|---|---|---|
| | Porridge or other food made with corn/ vutta, of the type bought in stores? | |
| 3 | অথবা শিশুদের খাদ্য শস্য যেমন সেরেলাক?<br>Other baby cereal such as Cerelac or other? | I__I__I |
| 3.1 | অন্যান্য হলে ,নির্দিষ্ট করুন? If others, What type?<br>_______________ | |
| 4 | পুষ্টিকণা, মনিমিক্স মিশ্রিত খাবার [গুড়া বা মাইক্রোনিউট্রিয়েন্ড দানা যা বাজারে পাওয়া যায়]?<br>Foods to which you added pushtikona/ monimix/ other [a powder or micronutrient sprinkles available in the market]? | I__I__I |
| 4.1 | অন্যান্য হলে ,নির্দিষ্ট করুন? If other, what type?<br>_______________ | |
| 5 | সোনামনি যা আমাদের থেকে পেয়েছেন?<br>Shonamoni you received from us? | I__I__I<br>যদি ওয়াশ-বেনিফিট এ অন্তর্ভূক্ত হওয়ার পর পর হয় তাহলে "**00**" কোড করুন ।<br>যদি শিশুটির বয়স ৬ মাসের বেশী হয় এবং সে ওয়াশ-বেনিফিট থেকে কোন LNS না পেয়ে থাকে তাহলে "**88**" কোড করুন ।<br>[Fill in "**00**" at enrollment.<br>Fill in "**88**" if infant does not receive LNS from WASH Benefits] |
| 6 | অন্য যে কোন (এল এন এস) বা সম্পূরক পুষ্টি/পুষ্টি প্যাকেট?<br>Any other [Lipid-based nutrient supplement (LNS)]? | I__I__I |
| 6.1 | যদি হ্যা হয়, দেখাতে বলুন এবং নাম লিখুন:<br>_______________<br>If Yes, ask to see and write name/type: | |

এখন আমি আপনার কাছে ভিটামিন/মিনারেল এর বড়ি অথবা ড্রপস সর্ম্পকে জানতে চাইবো ।Now I would like to ask you about vitamin/mineral pills or drops.

**C.613.** গত সাত দিনে কয়দিন আপনার শিশু [নাম] ভিটামিন/মিনারেল এর বড়ি অথবা ড্রপস নিয়েছে? On how many days did [NAME] have any vitamin/mineral pills or drops in the last 7 days?

১ = (০ - ৭) দিন I__I__I days

1= (0 – 7) DAYS I__I__I দিন

৬৬ = খেয়েছে,কিন্তু কয়দিন খেয়েছে তা জানেনা [C.615 নং প্রশ্নে যান]

66 Child had, but number of days not known [skip to C.615]

৯৯ = খেয়েছে কি খায়নি জানেনা [C.615 নং প্রশ্নে যান]

99 Don't know if child had or not [skip to C.615]

2= No, not eaten, **২= খায়নি** [C.615 নং প্রশ্নে যান]

**C.614.** 👁 **পর্যবেক্ষণ:** বড়ি অথবা ড্রপস এর নামের , তথ্যের উৎস কি? Observation:
What is the source of the information on the pill or drop name?

- ☐ 1 তথ্য পাওয়া যায় নি Information not available
- ☐ 2 তথ্য সংগ্রহকারী মোড়ক দেখেছে Data collector saw package
- ☐ 3 উওর দাতা মনেকরে নাম বলতে পেরেছে Respondent remembered and told name

c. 614_1 যদি শিশু ভিটামিন/মিনারেল এর বড়ি অথবা ড্রপস নিয়ে থাকে : কি ধরনের? If baby was given vitamin / mineral drops or pills: What type?

|__|__|

___________________________________________________________

উত্তর দাতাকে মোড়ক দেখাতে বলুন এবং উপরের লাইনে নাম লিখুন।

*Ask the respondent to show the package and write the name on the line above.*

**C.615.**
শিশু কি কখনো ময়লা বা মাটি খিয়েছে? Has the child eaten any dirt or soil?

মনে করার জন্য প্রতিটি সময় আলাদা ভাবে জানতে চান: *Ask for each recall period:*

| | | | |
|---|---|---|---|
| 1 | আজ Today | ☐ 1 Yes | ☐ 2 No | ☐ 99 Don't know |
| 2 | গতকাল Yesterday | ☐ 1 Yes | ☐ 2 No | ☐ 99 Don't know |
| 3 | গত কালের আগের দিন Day before yesterday | ☐ 1 Yes | ☐ 2 No | ☐ 99 Don't know |
| 4 | গত সাত দিন (আজকের দিন থেকে গত এক সপ্তাহ?) In the past 7 days (since this day last week?) | ☐ 1 Yes | ☐ 2 No | ☐ 99 Don't know |

#### Wash Benefit Module 7 HANDWASHING

| | WBM 7 | | |
|---|---|---|---|
| C.700 | 700. | এই প্রশ্নের আগে উত্তরদাতা যে কোন সময় তার হাত ধুয়েছিল কিনা তা পর্যবেক্ষণ করুন। RECORD WHETHER THE RESPONDENT HAS WASHED HER HANDS AT ANY TIME BEFORE THIS QUESTION | [1] উত্তরদাতা হাত ধুয়েছিল OBSERVED RESPONDENT WASHING HANDS<br>[2] উত্তরদাতা হাতধোয় নাই DID NOT OBSERVE RESPONDENT WASHING HANDS |
| C.701 | 701. | আপনাকে ধন্যবাদ । এখন আমি আপনার হাতগুলো দেখতে চাই। আশা করি আপনি কিছু মনে করবেন না। আপনি কি আমাকে দেখাবেন? READ: Thank you. Now, I would like to do a quick inspection of your hands. I hope you don't mind. Can you please show me your hands? | 👁 MOTHER<br><br>**বাম হাত Left Hand**<br>A|__| হাতের নখ FINGERNAILS<br>B|__| হাতের তালু PALMS<br>C|__| আঙ্গুলের সম্মুখভাগ FINGER PADS |

| | | | |
|---|---|---|---|
| | | দুই হাতই দেখতে হবে(প্রথমে এক হাত)। এরপর নিচের উলেস্নখিত হাতের অবস্থার কোড দেখে রেকর্ড করতে হবে। BOTH HANDS SHOULD BE SHOWN (NOT JUST ONE HAND). USE APPEARANCE CODES BELOW TO RECORD THE DESCRIPTION THAT BEST DESCRIBES THE DEFINITIONS THE LEVEL OF CLEANLINESS.<br>APPEARANCE CODES:<br>[1] ময়লা স্পষ্টভাবে দেখা যাচ্ছিল VISIBLE DIRT (DIRT/MUD/SOIL/ASH OR ANY OTHER MATERIAL IS VISIBLE)<br>[2] ময়লা স্পষ্টভাবে দেখা নাগেলেও অপরিচ্ছন্নভাব ছিল UNCLEAN APPEARANCE (NO DIRT IS VISIBLE ON THIS PART OF THE HAND BUT, IN GENERAL, THIS PART OF THE HAND APPEARS UNCLEAN)<br>[3] পরিষ্কার ছিল CLEAN (OBSERVED PART OF THE HAND IS CLEAN AS WOULD APPEAR AFTER SOMEONE WASHES HANDS OR TAKES A BATH)<br>৮৮. পর্যবেক্ষণ করা সম্ভব হয়নি/প্রত্যাখান N/A; OBSERVATION NOT POSSIBLE/REFUSED | **ডান হাত Right Hand**<br>D\|__\| হাতের নখ FINGERNAILS<br>E\|__\| হাতের তালু PALMS<br>F\|__\| আঙ্গুলের সম্মুখভাগ FINGER PADS |
| **C.701** | 702. | ASK: দয়া করে আমাকে(শিশুর নাম ধরে) হাতগুলো দেখাও । Please show me [NAME]'s hands.<br><br>উত্তরদাতার সবচেয়ে বড় বাচ্চার (০-৩৬ মাস) হাত দেখুন । এই বয়সের বাচ্চা না থাকলে স্কিপ করু। TO SEE THE HANDS OF THE RESPONDENT'S OLDEST CHILD 0-36 MONTHS (SEE C.1) IF RESPONDENT DOESN'T HAVE A CHILD 0-36 MONTHS → **SKIP TO NEXT PAGE**.<br><br>APPEARANCE CODES:<br>[1] ময়লা স্পষ্টভাবে দেখা যাচ্ছিল(**ময়লা/কাঁদা/মাটি/ছাই/এছাড়া অন্য কিছু**)VISIBLE DIRT (DIRT/MUD/SOIL/ASH OR ANY OTHER MATERIAL IS VISIBLE)<br>[2] ময়লা স্পষ্টভাবে দেখা না গেলেও অপরিচ্ছন্নভাব ছিল(হাতে **ময়লা দেখা না** | ৩সবচেয়ে বড় বাচ্চার ০-৩৬ মাস OLDEST CHILD 0-36 MONTHS<br><br>**বাম হাত Left Hand**<br>A\|__\| হাতের নখ FINGERNAILS<br>B\|__\| হাতের তালু PALMS<br>C\|__\| আঙ্গুলের সম্মুখভাগ FINGER PADS<br><br>**ডান হাত Right Hand**<br>D\|__\| হাতের নখ FINGERNAILS<br>E\|__\| হাতের তালু PALMS<br>F\|__\| আঙ্গুলের সম্মুখভাগ FINGER PADS |

| | | | |
|---|---|---|---|
| | | গেলেও অপরিচ্ছন্ন ছিল)UNCLEAN APPEARANCE (NO DIRT IS VISIBLE ON THIS PART OF THE HAND BUT, IN GENERAL, THIS PART OF THE HAND APPEARS UNCLEAN)<br>[3] পরিষ্কার ছিল(হাতধোয়ার পর/গোসলের পর হাত দেখলে পরিষ্কার হবে) CLEAN (OBSERVED PART OF THE HAND IS CLEAN AS WOULD APPEAR AFTER SOMEONE WASHES HANDS OR TAKES A BATH)<br>[88] পর্যবেক্ষণ করা সম্ভব হয়নি/প্রত্যাখান/০-৩৬ মাসের বাচ্চা নাই N/A; OBSERVATION NOT POSSIBLE/REFUSED/ NO CHILD 0-36 MONTHS | |
| C.702 a | 703. | পর্যবেক্ষনঃ প্রাথমিক/ প্রধান হাত ধোয়ার স্থানটি কোথায় তা রেকর্ড করুন **[Observation: Record the location where the primary hand washing station is located?]** | 1. ঘরের ভিতরে/ কাছে (≤৬ কদম) In/near main house (≤6 feet to entrance)<br>2. পায়খানার ভিতরে/ কাছে (≤৬ কদম) In/near latrine (≤6 feet to entrance)<br>3. রান্নার স্থানের ভিতরে/ কাছে (≤৬ কদম) In/near cooking area (≤6 feet to entrance)<br>4. ঘর, পায়খানা, রান্নার স্থানে থেকে >৬ কদম >6 feet away from main house, latrine and cooking area<br>**5.** নির্দিষ্ট কোন জায়গায় নাই No specific place 714 নং প্রশ্নে চলে যান (skips to 714) |
| C.702 c | 704. | পর্যবেক্ষনঃ প্রাথমিক/ প্রধান হাত ধোয়ার স্থানটিতে কোন কোন উপাদান আছে? (আপনি যদি দেখে থাকেন তাহলে "1" কোড করুন আর না দেখলে "0" কোড করুন) [Observation: Which of the following are present at the primary hand washing station? (If you observe the listed item, write "1" for "yes" in the box below. If you do not observe the listed item, write "0" for "no" in the box below.)] *1= হ্যাঁ [Yes], 0= না [No]* | MATERIALS PRESENT<br>[1] পানি WATER<br>[2] গোসল/হাত ধোয়ার সাবান (লাক্স, লাইফবয়) BAR SOAP (Body/hand Bar)<br>[3] অন্য যে কোন ধরনের সাবান (হুইল) BAR SOAP (other)<br>[4] গুড়া পাউডার POWDERED SOAP<br>[5] সাবান-পানি SOAPY WATER<br>[6] তরল সাবান LIQUID SOAP<br>[7] ছাই Ash<br>[8] মাটি /বালু Mud/Sand<br>[9] আইসিডিডিআর, বি-র দেওয়া ট্যাপসহ বালতি ICDDR,B drum with tap<br>[10] হাত ধোয়ার স্থানটি ভিজা ছিল/ব্যবহারের চিহ্ন ছিল Moisture below handwashing station<br>[11] বালতি Bucket<br>[12] বেসিন, জগ, মগ, বদনা Basin/Jug<br>[13] কিছুই নাই NONE OF THE ABOVE<br>**[777]** অন্যান্য OTHER (SPECIFY): _____________ |
| | 705. | **প্রধান হাত ধোয়ার স্থানটি সাবান জাতীয় যেকোন ধরনের উপাদান (যা হাত ধোয়ার কাজে ব্যবহার করা হয়/ সাবান পানি ও পানি একসাথে উপস্থিত আছে কী?** [Soap/soapy water and water present | 1. হ্যাঁ [Yes]<br>0. না [No] |

| | | | |
|---|---|---|---|
| | | together in PHWS] | |
| C.702 c | 706. | প্রাথমিক/ **প্রধান** হাত ধোয়ার স্থানটি রান্নাঘর থেকে কত কদম দূরে? [Observation: The primary hand washing station is away from the kitchen---Count in steps, allow in continuous numbers] | \|____\| |
| C.702 d | 707. | প্রাথমিক/ **প্রধান** হাত ধোয়ার স্থানটি পায়খানা থেকে কত কদম দূরে? [Observation: The primary hand washing station is away from the toilet---Count in steps, allow in continuous numbers] | \|____\| |
| C.703 | 708. | উত্তরদাতাকে প্রশ্ন করুনঃ আপনার হাতধোয়ার জন্য প্রাথমিক/ **প্রধান** স্থান ছাড়া অন্যকোন জায়গা আছে কি? [Ask the respondent: "Is there anywhere else you wash your hands?" (Record code in box)]: | 1. হ্যাঁ [Yes]<br>0. না [No] **714** নং প্রশ্নে চলে যান (skips to 714) |
| C.703 a | 709. | পর্যবেক্ষনঃ দ্বিতীয় প্রধান হাত ধোয়ার স্থানটি কোথায় তা রেকর্ড করুন [Observation: Record the location of the secondary hand washing station]. | 1. ঘরের ভিতরে/ কাছে (≤৬ কদম) In/near main house (≤6 feet to entrance)<br>2. পায়খানার ভিতরে/ কাছে (≤৬ কদম) In/near latrine (≤6 feet to entrance)<br>3. রান্নার স্থানে ভিতরে/ কাছে (≤৬ কদম) In/near cooking area (≤6 feet to entrance)<br>4. ঘর, পায়খানা, রান্নার স্থানে থেকে >৬ কদম >6 feet away from main house, latrine and cooking area |
| C.703 c | 710. | পর্যবেক্ষনঃ দ্বিতীয় প্রধান হাত ধোয়ার স্থানটিতে কোন কোন উপাদান আছে? (আপনি যদি দেখে থাকেন তাহলে "1" কোড করুন আর না দেখলে "0" কোড করুন) [Observation: Which of the following are present at the secondary hand washing station? (If you observe the listed item, write "1" for "yes" in the box below. If you do not observe the listed item, write "0" for "no" in the box below.) ] | *1= হ্যাঁ [Yes], 0= না [No]*<br>[1] **পানি** WATER<br>[2] **গোসল/হাত ধোয়ার সাবান (লাক্স, লাইফবয়)** BAR SOAP (Body/hand Bar)<br>[3] **অন্য যে কোন ধরনের সাবান (হুইল)** BAR SOAP (other)<br>[4] **গুড়া পাউডার** POWDERED SOAP<br>[5] **সাবান-পানি** SOAPY WATER<br>[6] **তরল সাবান** LIQUID SOAP<br>[7] **ছাই** Ash<br>[8] **মাটি /বালু** Mud/Sand<br>[9] **আইসিডিডিআর, বি-র দেওয়া ট্যাপসহ বালতি** ICDDR,B drum with tap<br>[10] হাত ধোয়ার স্থানটি ভিজা ছিল/ব্যবহারের চিহ্ন ছিল Moisture below handwashing station<br>[11] (**বালতি** Bucket<br>[12] **বেসিন, জগ, মগ, বদনা** Basin/Jug<br>[13] **কিছুই নাই** NONE OF THE ABOVE<br>৭৭৭. **অন্যান্য** OTHER (SPECIFY): _____________ |
| | 711. | **দ্বিতীয় প্রধান হাত ধোয়ার স্থানটিতে সাবান জাতীয় যেকোন ধরনের উপাদান (যা হাত ধোয়ার কাজে ব্যবহার করা হয়/সাবান পানি ও পানি একসাথে উপস্থিত আছে কী?** [Soap/soapy water and water present together in SHWS] | 1. হ্যাঁ [Yes]<br>0. না [No] |

| | | | |
|---|---|---|---|
| C.703 c | 712. | দ্বিতীয় প্রধান হাত ধোয়ার স্থানটি রান্নাঘর থেকে কত কদম দূরে? [Observation:T he secondary hand washing station is away from the kitchen---Count in steps, allow in continuous numbers]] | \|____\| |
| C.703 d | 713. | দ্বিতীয় প্রধান হাত ধোয়ার স্থানটি পায়খানা থেকে কত কদম দূরে? [Observation: The secondary hand washing station is away from the toilet---Count in steps, allow in continuous numbers] | \|____\| |
| INDICATOR 4: HANDWASHING সাবানের উপস্থিতিতে হাতধোয়া দেখানো DEMONSTRATION, SOAP PRESENCE | | | |
| আপনাকে ধন্যবাদ। আপনি দয়া করে আমাকে পায়খানা থেকে আসার পর যেখানে বেশীরভাগ সময় হাত ধোন সেই স্থানটি দেখান(নোট ;পূর্বে পর্যবেক্ষীত স্থানও হতে পারে। এরপর উত্তরদাতা কর্তক চিহ্নিত স্থানে যেতে হবে এবং রেকর্ড করতে হবে।Thank you. Please show me where you most often wash your hands after defecation. (Note: this may be the same place you already observed. Go to the place identified by the respondent and record location). | | | |
| C.704 a | 714. | 👁 হাতধোয়ার স্থান পর্যবেক্ষণ করুন(একটিকে গোলাকার করুন)। OBSERVE AND RECORDLOCATION OF HANDWASHING (CIRCLE ONE) | 1. ঘরের ভিতরে/ কাছে (≤৬ কদম) In/near main house (≤6 feet to entrance)<br>2. পায়খানার ভিতরে/ কাছে (≤৬ কদম) In/near latrine (≤6 feet to entrance)<br>3. রান্নার স্থানে ভিতরে/ কাছে (≤৬ কদম) In/near cooking area (≤6 feet to entrance)<br>4. ঘর, পায়খানা, রান্নার স্থানে থেকে >৬ কদম >6 feet away from main house, latrine <u>and</u> cooking area<br>**5.** নির্দিষ্ট কোন জায়গায় নাই |
| **সময় গনণার জন্য ষ্টপ ওয়াচ প্রস্তুতকরন PREPARE YOUR STOP WATCH FOR TIMING.** | | | |
| C.705 | 715. | পায়খানা থেকে আসার পর যেভাবে আপনি আপনার হাতধোন একইভাবে আমাকে এখন তা করে দেখান। Now, can you demonstrate how you normally clean your hands <u>after defecation</u>? Try in the same manner as you would if I wasn't here. TIME HOW LONG IT TAKES THE RESPONDENT TO BRING MATERIALS TO THAT PLACE (START AT THE MOMENT YOU FINISH ASKING THE QUESTION). WHEN RESPONDENT STARTS CLEANING HER HANDS, TIME DURATION OF RUBBING HANDS TOGETHER. | [1] দেখানো হয়েছে DEMONSTRATED<br><br>[2] দেখানো হয়নি COULD NOT DEMONSTRATE<br>IF NOT → **RECORD REASON:**<br>**________________________________________AND <u>SKIP TO 721</u>** |
| C.705 b | 716. | 👁পর্যবেক্ষণ করুন এবং হাতধোয়ার সাবান আনতে কতটুকু সময় লেগেছে তা রেকর্ড করুন। যদি হাতধোয়ার স্থানে সাবান থাকে তাহলে ০০:০০ বসান। যদি সাবান ব্যবহার না করে থাকে তাহলে ৮৮:৮৮ বসান । OBSERVE AND RECORD TIME TO PRODUCE (Bangladesh: <u>SOAP)</u> FOR CLEANING | \|____:____\| মি.mm. সে.ss |

| | | | |
|---|---|---|---|
| | | HANDS<br>MARK “00:00” IF SOAP IS ALREADY PRESENT AT HANDWASHING PLACE. (Bangladesh ONLY) MARK “88:88” IF SOAP IS N T USED | |
| C.705 c | 717. | 👁 পর্যবেক্ষণ এবং হাতঘষার সময় রেকর্ড করুন \| উত্তরদাতা যখন হাত পরিস্কারের জন্য দুহাত ঘষবে সেই সময় গননা করতে হবে ।<br>OBSERVE AND RECORD LENGTH OF TIME RESPONDENT SPENDS RUBBING HER HANDS TOGETHER: | \|____:____\| মি.mm. সে.ss |
| C.705 d | 718. | 👁 পর্যবেক্ষণ এবং হাত ধোয়ার সকল উপাদান রেকর্ড করুন<br>OBSERVE AND RECORD ALL MATERIALS USED FOR HAND CLEANSING (CIRCLE ALL THAT APPLY) 1 হ্যাঁ Yes 0 না No | 1. পানি WATER<br>2. বার সাবান BAR SOAP (Body/hand BAR)<br>3. বার সাবান (অন্যান্য)BAR SOAP (other)<br>4. গুড়া পাউডার POWDERED SOAP<br>5. সাবান-পানি SOAPY WATER<br>6. কাপড়/পাতা CLOTH OR LEAVES<br>7. বেসিন/কলসBASIN/PITCHER<br>8. ছাই ASH<br>9. মাটি/বালু MUD<br>88. অন্যান্য উল্লেখ করুন OTHER (SPECIFY): ___________________________ |
| C.705 e | 719. | 👁 পর্যবেক্ষণ এবং ২ হাত ধোয়া রেকর্ড করুন \|OBSERVE AND RECORD WHETHER BOTH HAN S WERE CLEANED | [1] হ্যাঁ, ২-হাত ধুয়েছে YES, BOTH HANDS CLEANED<br>[2] না ,১-হাত ধুয়েছে NO, ONLY ONE HAND CLEANED |
| C.705 f | 720. | 👁 পর্যবেক্ষণ এবং হাত শুকানো রেকর্ড করুন OBSERVE AND RECORD HOW RESPONDENT DRIES HANDS<br>1 Yes 0 No | 1. পরিধেয় কাপড় DRIES BY WIPING HER HANDS ON HER CLOTHES<br>2.এক হাত অন্য হাতের সাথে ঘষে DRIES BY WIPING HER HANDS ON ANOTHER<br>3. পরিধেয় ছাড়া অন্য পরিষ্কার কাপড় Clean CLOTH DRIES BY WIPING HER HANDS ON<br>৪. পরিধেয় ছাড়া অন্য অপরিষ্কার কাপড় Unclean CLOTH DRIES BY WIPING HER HANDS ON<br>5. কাপড় ছাড়া অন্য জিনিস যেমন: টিস্যু, কাগজ ইত্যাদি MATERIAL (NOT CLOTH)<br>5. বাতাসে/হাত ঝাঁকিয়ে বাতাসে শুকানো হয়েছে DRIES BY SHAKING HER HANDS IN THE AIR<br>6. হাত শুকানো হয় নাই NOT DRIED<br>88. অন্যান্য উল্লেখ করুন OTHER, SPECIFY________________________ |
| C.705 g | 721. | **যদি উত্তরদাতা হাতধোয়া না দেখান অথবা সাবান ব্যবহার না করে তখন তাকে তার খানায় হাতধোয়ার জন্য সাবান আছে কিনা জানতে হবে এবং তা আনতে বলতে হবে?**IF RESPONDENT COULD NOT DEMONSTRATE, OR DID NOT USE SOAP, ASK:<br>Do you have soap in your | \|____:____\| মি.mm. সে.ss |

| | | | |
|---|---|---|---|
| | | house that you use for handwashing? Can you bring it to me?<br><br>👁পর্যবেক্ষণ করুন এবং হাতধোয়ার সময় রেকর্ড করুন। যদি 716 তে উত্তর/সময় উল্লেখ থাকে তাহলে ৮৮:৮৮ বসান। যদি খানাতেসাবান না থাকে তাহলে ৯৯:৯৯ বসান ।<br>OBSERVE AND RECORD TIME TO PRODUCE SOAP FOR WASHING HANDS MARK "88:88" IF ALREADY TIMED IN C.704a MARK "99:99" IF OBSERVATION NOT POSSIBLE OR SOAP NOT AVAILABLE | |
| C. 706a | 722. | 👁 পর্যবেক্ষণ করুন : উত্তরদাতার হাতধোয়া দেখানোর সময়ে এ বিষয়ে কোন প্রতিক্রিয়া দেখতে পেয়েছিলেন কী?OBSERVE: WAS THERE ANY OBVIOUS REACTIVITY DURING THE COURSE OF HIS DEMONSTRATION? | [1] *হ্যাঁ YES*<br>[0] *না (৭২৩ নং এ চলে যাবে)* NO → skip to 723 |
| b | 1 | অধিকক্ষণ যাবৎ হাত ধুয়েছে কিনা<br>Longer time spent cleaning/rubbing hands | [1] *হ্যাঁ* YES<br>[0] *না* NO |
| c | 2 | সাবান দিয়ে হাত ধুয়েছে কিনা অন্যথায় Using soap where they otherwise would not | [1] *হ্যাঁ* YES<br>[0] *না* NO |
| d | 3 | অন্যান্য (লিখুন) Other, (BRIEFLY explain)_______________ | [1] *হ্যাঁ* YES<br>[0] *না* NO |

| | | | | | |
|---|---|---|---|---|---|
| | 723. | প্রশ্ন করুন ; আপনি কখন কখন সাবান দিয়ে হাত ধোন । ASK: "Please tell me about all of the times you wash your hands with soap."<br><br>A কলা ম (লিড ছাড়া)-এ যখন উত্তরদাতা যে কাজ করবে তা উল্লেখ করতে হবে । ১নং গোলাকার হবে যদি নির্দিষ্ট বিষয়ে উত্তর পাওয়া যায়। CIRCLE "1" IN COLUMN A IF CRITICAL TIME IS MENTIONED WITHOUT PROMPTING.<br><br>উত্তরদাতার সকল কাজ উল্লেখ করার পর অন্য আর কোন কাজে সাবান দিয়ে হাত ধোন কিনা এ ব্যপারে জিজ্ঞাসা করুন? AFTER THE RESPONDENT FINISHES NAMING ALL THE TIMES ASK "IS THERE ANY OTHER TIME YOU WASH YOUR HANDS WITH SOAP?" AND STOP WHEN THE RESPONDENT SAYS THERE IS NO OTHER TIME.<br><br>B কলাম-এ লিড দেওয়া ছাড়া প্রতিবার ব্যবহার করে কিনা তা উল্লেখ করুন । EACH TIME MENTIONED WITHOUT PROMPTING ASK COLUMN B.<br><br>C.707A-G নং প্রশ্নের উত্তর যদি A কলা ম [1] না হয় | A.<br>লিড দেওয়া ছাড়া হাতধোয়া উল্লেখ করুন<br>MENTIONED WITH NO PROMPTING | B.<br>লিড দেওয়া ছাড়া শুধুমাত্র হাতধোয়ার সংখ্যা/বার উল্লেখ করুন । সাবান ও পানি দিয়ে কত বার আপনার হাতধোন তা বলুন । Only for times mentioned with NO PROMPTING, ask: How often do you wash your hands with soap and water [insert event]?<br><br>[1] সবসময়/প্রতিবার ALWAYS<br>[2] মাঝে মাঝে/ অর্ধেকের কম বার SOMETIMES<br>[3] খুব কম সময়/ দু-এক বার RARELY | C.<br>লিড দেওয়ার পর শুধুমাত্র হাতধোয়ার সংখ্যা/বার উল্লেখ করুন । সাবান ও পানি দিয়ে কত বার আপনার হাতধোন তা বলুন । REPLIED "AFTER BEING PROMPTED<br><br>[1] সবসময়/প্রতিবার ALWAYS<br>[2] মাঝে মাঝে/ |

| | | | | | |
|---|---|---|---|---|---|
| | | তখন সাবান ও পানি দিয়ে আপনি আপনার হাতধোন কিনা তা জিজ্ঞাসা করুন। যদি উত্তরদাতা হ্যাঁ বলে তখন C কলা ম-এ হাতধোয়ার সংখ্যা/বার উল্লেখ করুন । FOR QUESTIONS C.707A-G WHERE [1] IS NOT CIRCLED IN COLUMN A, PROMPT BY ASKING: Do you usually wash your hands with soap and water [CRITICAL TIME]? IF THE RESPONDENT SAYS "YES" THEN IMMEDIATELY ASK HOW OFTEN AND MARK APPROPRIATE ANSWER CHOICE IN COLUMN C. | | | অর্ধেকের কম বার SOMETIMES<br>[3] খুব কম সময়/ দু-এক বার RARELY<br>[4] না NO<br>[88] প্রযোজ্য নয় Not applicable |
| a | 1 | খাবার তৈরী করার আগে BEFORE PREPARING FOOD | | | |
| | 2 | মাংস/মাছ কাটার পর AFTER HANDLING MEAT/FISH | | | |
| | 3 | ফল/সব্জি কাটার আগে BEFORE CUTTING FRUITS AND VEGETABLES | | | |
| | 4 | ভর্তা/ম্যাসড খাবার তৈরী করার আগে BEFORE MAKING MASHED FOOD (BHORTA) | | | |
| b | 5 | খাবারের পূর্বে BEFORE EATING | | | |
| c | 6 | খাবারের পরে AFTER EATING | | | |
| d | 7 | শিশুকে খাওয়ানোর আগে BEFORE FEEDING A CHILD | | | |
| e | 8 | বাচ্চাকে সৌচানোর পর AFTER CLEANING A CHILD'S ANUS | | | |
| f | 9 | বাচ্চার পায়খানা ফেলার পর AFTER DISPOSING OF CHILDREN'S FECES | | | |
| g | 10 | পায়খানার পর AFTER TDEFECATION | | | |
| h | 11 | কম্পাউন্ড/বাড়ীর বাহির থেকে ফেরার পর AFTER RETURNING FROM OUTSIDE THE COMPOUND | | | |
| m | 12 | রুগ্ন ব্যক্তিকে ধরার পর AFTER TOUCHING A SICK PERSON | | | |
| o | 13 | পশু-পাখি ধরার পর AFTER HANDLING LIVESTOCK | | | |
| | 14 | গোবর ধরার পর AFTER HANDLING COW-DUNG (OR ANY ANIMAL FECES) | | | |
| j | 15 | অন্যান্য (উল্লেখ করুন) OTHER (SPECIFY): ______________________ | | | |
| i | 16 | উত্তরদাতা কখনই সাবান দিয়ে হাতধোয় নি RESPONDENT NEVER WASHES HANDS WITH SOAP<br>Skip to module 8 | | | |
| | 17 | উঠান ঝাড়ু দেওয়া/ময়লা পরিষ্কারের পর After weaping courtyard | | | |
| | 18 | রান্নার পর After cooking | | | |
| | 19 | থালা-বাসন ধোয়ার পর After washing dishes | | | |

#### Wash Benefit Module 8 SANITATION (All households)

| | |
|---|---|
| SAY | এই গবেষণায় আপনার অংশ গ্রহনের জন্য আপনাকে ধন্যবাদ। ষ্টাডির এই অংশটি কিছটা স্পর্শকাতর । আমি আপনাকে আপনার/এই বড়ীর পয়ঃ ব্যবস্থাপনা/পয়ঃ নিষ্কাশন ও তার অনুশীলন এবং কিভাবে তা করেন এ ব্যপারে জানতে চাই \| আমি কিছুটা পর্যবেক্ষণ করতে চাই। আপনার স্বাচ্ছন্দবোধ হবে কি ? যদি না হয় তাহলে উত্তরদাতাকে ইহার গুরুত্ব বোঝাতে চেষ্টা করুন।Thank you so much for your participation so far. The next part of the survey is a bit sensitive. I would like to ask you some questions about the sanitation practices of people in your compound, including how you usually dispose of your children's feces. I would also like to make some observations. Are you comfortable with this? IF NOT, EXPLAIN THE IMPORTANCE OF THIS MODULE AND ENCOURAGE RESPONDENT TO PROCEED. |
| খোলা-পায়খানা(সকল খানার ) OPEN DEFECATION (ADMINISTER TO ALL HOUSEHOLDS) | |

| | | | Group A | B | C | D | E |
|---|---|---|---|---|---|---|---|
| | WBM 8 | | **Men** পুরুষ | **Women** মহিলা | **Children** **<3 years** <৩ বছরের বাচ্চা | **Children** **3-<8 years** ৩-<৮ বছরের বাচ্চা | **Children** **8-15** ৮-১৫ ছরের বাচ্চা |
| **C.801.** | 801. | এই খানার কোন সদস্যরা খোলা জায়গাতে পায়খানা করে কী? [Do [GROUP] in this household ever practice open defecation?<br>1 প্রতিদিন Daily<br>2 মাঝেমাঝে Occasionally<br>3 কখনই না Never **(Skip to 806)**<br>88. প্রযোজ্য নয় Not applicable (**806** নং প্রশ্নে যান) **(Skip to 806)**<br>99 জানি না (**806** নং প্রশ্নে যান) Don't Know **(Skip to 806)** | ☐ | ☐ | ☐ | ☐ | ☐ |
| **C.802.** | 802. | সাধারনত: একই জায়গাতে প্রতিবার যান কী? [Do [GROUP] go to more or less the same area every time?<br>1 হ্যাঁ Yes<br>2 না No (**805** নং প্রশ্নে যান) **(Skip to 805)**<br>99 জানি না Don't Know / Not Sure (**805** নং প্রশ্নে যান) **(Skip to 805)** | ☐ | ☐ | ☐ | ☐ | ☐ |
| **C.803.** | 803. | আপনার খানা থেকে উক্তস্থানে যেতে (শুধু যাওয়া) কতটুকু সময় লাগে (মিনিট) [How long does it take to walk (one way) from your house to the most commonly visited place? | I_____I মিনিট MINUTES 99 জানি না Don't Know / Not Sure | | | | |
| **C.804.** | 804. | এই জায়গাটি ঐ গ্রামের মধ্যেই কী? [Is that place within the village?<br>1হ্যাঁ Yes<br>2 না No<br>99 জানি না Don't Know / Not Sure<br>888: প্রযোজ্য নয় N/A | I__I | I__I | I__I | I__I | I__I |

| | | | | | | | |
|---|---|---|---|---|---|---|---|
| C804E | **804a** | Is that place within the compound? এই জায়গাটা কি এই বাড়ীর মধ্যেই অবস্থিত ?<br>1 হ্যাঁ Yes<br>2 না No<br>99 জানি না Don't Know / Not Sure<br>888: প্রযোজ্য নয় N/A | I__I | I__I | I__I | I__I | I__I |

#### Open Defecation Questions

| | | | **Men** পুরুষ | **Women** মহিলা | **Children <3 years** <৩ বছরের বাচ্চা | **Children 3-<8 years** ৩-<৮ বছরের বাচ্চা | **Children 8-16** ৮-১৫ ছরের বাচ্চা |
|---|---|---|---|---|---|---|---|
| **C.804a** | 805. | আপনার খানার লোকজনের খোলা জায়গায় পায়খানা করার প্রধান কারন কি কি? উত্তর শুনানো যাবে না। What are the main reasons that [GROUP] in your household practice open defecation? ☝DO NOT READ RESPONSES MARK ALL THAT APPLY | | | | | |
| | 1 | 1 কোনও কিছুই নাই /পায়খানা নাই No choice (nothing else is available) | ☐ | ☐ | ☐ | ☐ | ☐ |
| | 2 | 2 কোথায় শিশু পায়খানা করবে তা নিয়ন্ত্রন করতে পারে না Cannot control where young children defecate | | | ☐ | ☐ | ☐ |
| | 3 | 3 গোপনীয়তা Privacy | ☐ | ☐ | ☐ | ☐ | ☐ |
| | 4 | 4 অভ্যাস/নিয়মিত কাজ Habit / Routine | ☐ | ☐ | ☐ | ☐ | ☐ |
| | 5 | 5 পায়খানা থেকে বাঁশঝাড় পছন্দ Prefer to use the bush rather than a toilet | ☐ | ☐ | ☐ | ☐ | ☐ |
| | 6 | 6 কাজে/স্কুলে পায়খানা/টয়লেট সহজে পাওয়া না গেলে Toilet not available at work / school | ☐ | ☐ | ☐ | ☐ | ☐ |
| | 7 | ভাগের পায়খানা/টয়লেট পছন্দ না হলে Choose not to share toilets with in-laws / extended family (or cannot) | ☐ | ☐ | ☐ | ☐ | ☐ |

| | | | | | | | |
|---|---|---|---|---|---|---|---|
| | 8 | 8 সুবিধাজনক Convenience | ☐ | ☐ | ☐ | ☐ | ☐ |
| | 9 | 9 নিরাপদ Safety | ☐ | ☐ | ☐ | ☐ | ☐ |
| | 10 | 10 আরামদায়ক Comfort | ☐ | ☐ | ☐ | ☐ | ☐ |
| | 11 | 11 অসুস্থতা/ পাতলা পায়খানা Sickness/diarrhoea | ☐ | ☐ | ☐ | ☐ | ☐ |
| | 12 | 12 পায়খানা/টয়লেট ভরে গেছে Latrine overflowed | ☐ | ☐ | ☐ | ☐ | ☐ |
| | 13 | 13 পায়খানা/টয়লেট ভেঙে গেছে(উপরের অংশ/সস্নাব) Latrine broken (superstructure and /or slab) | ☐ | ☐ | ☐ | ☐ | ☐ |
| | 14 | 14 পায়খানা/টয়লেটে যেতে ভয় পায় Fear of latrine | ☐ | ☐ | ☐ | ☐ | ☐ |
| | 15 | 15 পায়খানা/টয়লেট কিভাবে ব্যবহার করতে হয় তা জানে না Don't know how to use the latrine | ☐ | ☐ | ☐ | ☐ | ☐ |
| | 16 | 16 বাচ্চা খুব ছোট হওয়ায় পায়খানা/টয়লেট ব্যবহার করে না Too young to use latrine | ☐ | ☐ | | | |
| | 17 | বয়স্ক (খুবই বৃদ্ধ) Old age (too old) | | | | | |
| | 18 | অন্যান্য (উল্লেখ করুন) OTHER (SPECIFY): | | | | | |
| | | | <৩ বছরের বাচ্চা **Children < 3 years** | ৩-<৮ বছরের বাচ্চা **Children 3 – <8 years** | ৮-১৫ বছরের বাচ্চা **Children 8 – 15 years** | পুরুষ **Men** | মহিলা **Women** |
| **C.804e** | 806. | ~~পায়খানা থাকা সত্ত্বেও~~ এই বাড়ীর অন্য কোন পরিবার খোলা জায়গায় পায়খানা করে কিনা তা আপনি জানেন কী? Do you know of other households in the bari/compound whose [GROUP] practice OD, ~~even if they might have a toilet or latrine at their house~~?<br>1 হ্যাঁ, প্রায়ই Yes, Often<br>2 হ্যাঁ, মাঝে মাঝে Yes, Sometimes<br>3 না, কখনই না No, Never **(Skip to 807a)** | ☐ | ☐ | ☐ | ☐ | ☐ |

| | | | | | | | |
|---|---|---|---|---|---|---|---|
| | | 88. প্রযোজ্য নয় Not applicable (**807a** নং প্রশ্নে যান) **(Skip to 807a)**<br>99 জানি না/নিশ্চিত না Don't Know / Not Sure | | | | | |
| **C.804f** | 807. | আপনি শিশুদের পায়খানা উঠানে/উঠানের চার-পাশে/আশেপাশে /নদীতে ফেলে দিতে দেখেছেন কী? Do you see that children's stools are disposed in the yard / surrounding / community in your neighborhood / river?<br>1 হ্যাঁ, প্রায়ই Yes, Often<br>2 হ্যাঁ, মাঝে মাঝে Yes, Sometimes<br>3 না, কখনই না No, Never<br>888. কখনই ফেলে দিতে হয় না, প্রযোজ্য নয় Not applicable<br>99 জানি না/নিশ্চিত না Don't Know / Not Sure | | | | □ | □ |
| **C.804b** | 807a | সর্বশেষ যে সময় এই [গ্রুপ] এর কারো ডায়রিয়া হয়েছিল, তারা তখন কোথায় পায়খানা করেছিল ?The last time somebody in [GROUP] was sick with diarrhea, where did they defecate?<br>1. পায়খানায়/ টয়লেটে In the toilet / latrine **(Skip: Next Group)**<br>2 এই বাড়ীর মধ্যেই বালতি অথবা অন্যান্য অপরিকল্পিত / সহসা কনটেইনারে In a bucket or other improvised container within the compound<br>3 এই বাড়ীর মধ্যেই পটিতে Child potty within compound<br>4 এই বাড়ীর মধ্যেই খোলা পায়খানা Open | □ | □ | □ | □ | □ |

| | | | | | | | |
|---|---|---|---|---|---|---|---|
| | | defecation – within compound<br>5 (specify)_____<br>**অন্যান্য (উল্লেখ করুন)**<br>6. বাড়ির বাহিরে খোলা জায়গায়(Open defecation outside compound)<br>7. বিছানায় অথবা কাঁথাতে (In Bed or in covering)<br>8.□□□□ □□□□□□□□ □□□□<br>88 প্রযোজ্য নয়Not Applicable **(Skip: Next Group)**_____<br>99 জানি না Don't know / not sure **(Skip: Next Group)**<br>Open defecation – in the bush / forest / field **(Skip: Next Group)**<br>77 Other | | | | | |
| **C.804c** | 807b | এই পায়খানা কি করা হয়েছিল ? What was done with the feces?<br>1. সেখানেই ফেলে রাখা হয়েছিল Left there<br>2. টয়লেটের মধ্যে ফেলা/ধোয়া হয়েছিল Put / rinsed into toilet or latrine<br>3. ড্রেনে/ খালের মধ্যে ফেলা/ধোয়া হয়েছিল Put / rinsed into drain or ditch<br>4.বাশ ঝাড়/ জঙ্গলে/ মাঠে ফেলে দেয়া হয়েছিল Thrown into the bush / forest / field<br>5. ময়লা ফেলার জায়গায় ফেলে দেয়া হয়েছিল Thrown into garbage | ☐ | ☐ | ☐ | ☐ | ☐ |

| | | | | | | | |
|---|---|---|---|---|---|---|---|
| | | 6.শিশুদের পায়খানা ফেলার যে নির্দিষ্ট জায়গা সেখানে ফেলে দেয়া হয়েছিল Thrown into a specific pit for child's feces<br>7. পুড়িয়ে ফেলা হয়েছিল Buried<br>77. অন্যান্য (উল্লেখ করুন) Other (specify)__________<br>99. Don't know / not sure | | | | | |
| **C.804d** | 807c | পায়খানা করার কতক্ষণ পর কেউ পরিস্কার করেছিল ?How long after defecation did somebody dispose of the feces?<br>A ☐☐ HOURS<br>B ☐☐ MINUTES | A __ __<br>B __ __ | A __ __<br>B __ __ | A __ __<br>B __ __ | A __ __<br>B __ __ | A __ __<br>B __ __ |

#### Sanitation Facility

Administer to: All study households

| | | | |
|---|---|---|---|
| **C.805.** | 808. | আপনার খানায় কোন টয়লেট/পায়খানা আছে কি?<br>আমি কি তা দেখতে পারি?<br>Does your household have a toilet facility that is in use? Can I see it? | 1 হ্যাঁ আছে,পর্যবেক্ষন করতে পারি Yes have toilet, can observe<br>2 হ্যাঁ আছে,পর্যবেক্ষন করতে দেওয়া হয়নি/প্রত্যাখান (৮১২ নং এ যান ) Yes have toilet, refused observation **(Skip to 812)**<br>3 হ্যাঁ আছে,পর্যবেক্ষন করা যায়নি (৮১২ নং এ যান ) Yes have toilet, cannot observe **(Skip to 812)**<br>4 কোন পায়খানা নাই No toilet facility **(Skip to MODULE 9)** |
| **C.806.** | 809. | 👁 পর্যবেক্ষন; ধরণ,অবস্থা এবং পুনঃব্যবহার অবস্থা Observation: Note the type, condition and apparent use of the toilet:<br>1 হ্যাঁ Yes<br>2 না No<br>88 প্রযোজ্য নয়/ পর্যবেক্ষন করতে পারেনি/বলতে পারেনি Not Applicable / Could not observe / cannot tell | |
| বর্হিরাংশ পর্যবেক্ষণ **Exterior observation** | | | |
| 1 | 1 | পায়খানার চারপাশে ৩টি ইটের/ টিনের/মাটির | *(1= হ্যাঁ [Yes], 0= না [No], প্রযোজ্য নয় [not applicable] =888)]* |

| | | | |
|---|---|---|---|
| | | দেয়াল At least 3 walls around the toilet | |
| 2 | 2 | পায়খানার চারপাশে ৩টি বাঁশের বেড়ার দেয়াল Bamboo fences around the toilet | *(1= হ্যাঁ [Yes], 0= না [No],* **প্রযোজ্য নয়** *[*not applicable] =888)] |
| 3 | 3 | পায়খানার চারপাশে দরজা/পর্দা দিয়ে ঘেরা Door/curtain or walls that guarantee privacy around the toilet | *(1= হ্যাঁ [Yes], 0= না [No],* **প্রযোজ্য নয়** *[*not applicable] =888)] |
| 4 | 4 | পায়খানার ছাদ Roof over toilet | *(1= হ্যাঁ [Yes], 0= না [No],* **প্রযোজ্য নয়** *[*not applicable] =888)] |
| 5 | 5 | বাতাস বের হওয়ার পথ Ventilation pipe | *(1= হ্যাঁ [Yes], 0= না [No],* **প্রযোজ্য নয়** *[*not applicable] =888)] |
| 6 | 6 | পায়খানায় যাবার রাস্তা দেখে বুঝা যাচ্ছে নিয়মিত ব্যবহার করা হয় (পরিষ্কার, জীর্ণ ইত্যাদি ) Path to the toilet suggests regular use (is clear, well-worn, without grass or any barriers etc.) | *(1= হ্যাঁ [Yes], 0= না [No],* **প্রযোজ্য নয়** *[*not applicable] =888)] |
| ভিতরের অংশ পর্যবেক্ষণ **Interior observation** | | | |
| 7 | 7 | টয়লেটে স্লাব আছে Toilet has a slab | *(1= হ্যাঁ [Yes], 0= না [No], 2= Yes but not functional* **প্রযোজ্য নয়** *[*not applicable] =888)] |
| 8 | 8 | গর্তের পাশে পাদানী আছে Raised footing around hole | *(1= হ্যাঁ [Yes], 0= না [No],* **প্রযোজ্য নয়** *[*not applicable] =888)] |
| 9 | 9 | পানি ঢেলে ফ্লাস করা যায় Flush or poor Flush: | *(1= হ্যাঁ [Yes], 0= না [No],* **প্রযোজ্য নয়** *[*not applicable] =888)] [If 0/888 skip to 12] |
| 9 a | 9a | যদি পানি ঢেলে ফ্লাস করা যায় If Flush or Pour Flush:<br><br>**পানির সীলের অবস্থা** Water seal condition: | [ 1 ] ওয়াটার সিল ভাল আছে/কাজ করে Functional water seal<br>[ 2 ] ওয়াটার সিল ভাঙ্গা Broken water seal<br>[ 3 ] ওয়াটার সিল নেই No water seal<br>[ 4 ] পায়খানা ভরে উপছে পড়ছে Overflowing with feces<br>[ 88 ] **পায়খানাতে পানির সীল আছে/পর্যবেক্ষণ করা সম্ভব হয়নি** N/A / could not observe / cannot tell |
| 9 b | 9b | যদি পানি ঢেলে ফ্লাস করা যায় If Flush or Pour Flush:<br><br>পানি ঢাললে/দিলে Flushes to:<br><br>(খানার সদস্যকে জিজ্ঞাসা/যাচাই করুন *Ask / probe household members if necessary)* | ১. টয়লেটিতে পয়ঃনিষ্কাশন পাইপের সাথে সংযোগ করে দেয়া [Piped sewer system]<br>২. টয়লেটিতে সেপটিক ট্যাংক বসানো আছে [Septic tank]<br>৩. সেপটিক ট্যাংক নাই কিন্তু ফ্লাস করে বা পানি ঢেলে পায়খানা দুরে বা পিটের মধ্যে সরিয়ে দেয়া যায় [Flush to pit latrine ]<br>৪. ফ্লাশ-টয়লেট অথবা পানি ঢেলে ফ্লাস করা টয়লেট যা কোন খাল, ড্রেন, নদী ইত্যাদির সাথে সংযোজন করার ফলে অস্বাস্থ্যকর অবস্থার সৃষ্টি করে থাকে [Flush or pour flush toilet connected to somewhere else (canal, ditch, river, etc.)]<br>৫. Offset pit<br>৬. Onsite pit directly under |
| | 9c | পিটের/ ট্যাংকির বাইরে বা ভিতরের চর্তুদিকে পর্যবেক্ষণ করুন [Observe the onsite or off site pit/tank in all direction] (1= হ্যাঁ [yes], ☐= না | 1. \|__\|পিটের উপরিভাগ মাটি থেকে উপরে দেখা যাচ্ছে কি? [Is the top of the pit visible (above the ground)]?<br>2. \|__\|সংযোগ নলের ছিদ্রের/ভাংগার কারণে পাইপের ভিতরে বা বাইরে ময়লা দেখা যাচ্ছে কি? [Waste/faeces visible in or around the pipe, because of Leakage in the connecting pipe?] |

| | | | |
|---|---|---|---|
| | | [No], 999= জানিনা [DK], 888= প্রযোজ্য নয় [Not applicable]) | 3. \|__\|পিট/ট্যাংকির ছিদ্রের কারণে ভিতরে বা বাইরে ময়লা দেখা যাচ্ছে [Waste/faeces visible because of leakage in the pit/tank?]<br>Skip Note: 9c.3 নং প্রশ্নের উত্তর 1 হলে 10 নং প্রশ্নে চলে যান।<br>[If the answer to *question* 9c.3 is 1 go to question 10]<br>4. \|__\|ময়লা দেখা যাচ্ছে না কিন্তু পিট/ট্যাংকি ভাংগা যা দিয়ে মশা মাছি আশা যাওয়া করতে পারবে [No visible waste but broken pit/tank that may allow flies coming out of the toilet?]<br>Skip Note: 9c.4 নং প্রশ্নের উত্তর 1 হলে 10 নং প্রশ্নে চলে যান।<br>[If the answer to *question* 9c.4 is 1 go to question 10]<br>5. \|__\|ময়লা দেখা যাচ্ছে না কিন্তু পিট/ট্যাংকিতে ফাটল/ ভাংগন দেখা যাচ্ছে যা দিয়ে মশা মাছি আশা যাওয়া করতে পারবে না [No visible waste but crack in the pit/tank?]<br>6. পিট/ ট্যাংকিতে কোন ফাটল / ছিদ্র নাই (No crack/ leakage in pit/tank) |
| 1<br>0 | 10 | মেঝের প্রধান উপাদান Main material of the floor (*select 1*) | [ 1 ] মাটি Mud<br>[ 2 ] কাঠ Wood<br>[ 3 ] সিমেন্ট Cement<br>[ 4 ] টাইলস/ইটTile / brick<br>[ 5 ] প্লাষ্টিক Plastic<br>[ 88 ] প্রযোজ্য নয়/পর্যবেক্ষণ সম্ভব হয় নি N/A / could not observe / cannot tell |
| 1<br>1 | 11 | Bucket toilet | *(1= হ্যাঁ [Yes], 0= না [No],* প্রযোজ্য নয় *[*not applicable] =888)] |
| 1<br>2 | 12 | ঝুলন্ত পায়খানা Hanging toilet | [ 1 Yes ] [ 2 No ] [ 88 N/A] |
| 1<br>3 | 13 | ব্যবহৃত হচ্ছে বুঝা যায় এমন Latrine appears to be in use (by your best judgment) | *(1= হ্যাঁ [Yes], 0= না [No],* প্রযোজ্য নয় *[*not applicable] =888)] |
| 1<br>4 | 14 | পায়খানা/টয়লেটে পায়খানার গন্ধ রয়েছে Odor of feces in the latrine/bathroom | *(1= হ্যাঁ [Yes], 0= না [No],* প্রযোজ্য নয় *[*not applicable] =888)] |
| 1<br>5 | 15 | পায়খানা/টয়লেটে প্রস্রাবের গন্ধ রয়েছে Odor of urine in the latrine/bathroom | *(1= হ্যাঁ [Yes], 0= না [No],* প্রযোজ্য নয় *[*not applicable] =888)] |
| 1<br>6 | 16 | স্লাব অথবা মেঝেতে পায়খানা দেখতে পাওয়া গেছে Stool is visible on the slab or floor | *(1= হ্যাঁ [Yes], 0= না [No],* প্রযোজ্য নয় *[*not applicable] =888)] |
| 1 | 17 | ড্রপ হোল(মল-মূত্র প্রবেশ গর্ত) ঢাকা Drop hole is | *(1= হ্যাঁ [Yes], 0= না [No],* প্রযোজ্য নয় *[*not applicable] =888)][If |

| | | | |
|---|---|---|---|
| 7 | | covered | 0/888 skip to 18] |
| 1<br>7<br>a | 17<br>a | যদি ড্রপ হোল (মল-মূত্র প্রবেশ গর্ত)ঢাকা থাকে তাহলে মাছি আসা/যাওয়া করতে পারে কি If yes, cover is correctly placed over the drop hole so that flies cannot enter/exit | *(1= হ্যাঁ [Yes], 0= না [No],* **প্রযোজ্য নয়** *[*not applicable] =888)] |
| 1<br>8 | 18 | **মাছির উপস্থিতি রয়েছে** Flies present | *(1= হ্যাঁ [Yes], 0= না [No],* **প্রযোজ্য নয়** *[*not applicable] =888)] |
| 1<br>9 | 19 | **সাধারন বৈশিষ্ট General Characteristics**<br>একটি অথবা দুই পিট/গর্ত টয়লেট Single or double pit latrine | [ 1 ] একটি পিট/গত Single pit<br>[ 2 ] দুই পিট/গর্ত Double pit<br>[ 88 ] প্রযোজ্য নয়/পর্যবেক্ষণ সম্ভব হয় নি /বলতে পারে নিN/A / could not observe / cannot tell |
| 2<br>0 | 20 | কমপোষ্টিং টয়লেট Composting toilet | *(1= হ্যাঁ [Yes], 0= না [No],* **প্রযোজ্য নয়** *[*not applicable] =888)] |
| **C.807.** | 810. | **পর্যবেক্ষন করুন** Observation: টয়লেট-এর গর্ত মলে ভরেছে কিনা-আলোর সাহায্যে পর্যবেৰণ করম্নন Fullness of the pit – shine a light into the pit to see if solid waste is… | 1 **শুকনা ময়লা/পায়খানা ৩ ফুটের চেয়ে বেশী দূরে (>৩ ফুট)** Very far from surface (>1 meter)<br>2 **শুকনা ময়লা/পায়খানা ৩ ফুটের মধ্যে (<৩ ফুট)** Within 1 meter<br>3 **পায়খানার খুব কাছাকাছি বা পায়খানা ভরে গেছে** Very close to surface or full<br>88 **পায়খানাতে পানির সীল আছে/পর্যবেৰণ করা সম্ভব হয়নি** Water seal latrine / non direct pit latrine / could not observe |
| C. 808 | 811. | **পর্যবেক্ষন করুন** Observation: : সৌচ কাজের জন্য টয়লেট-এর ভেতরে এবং কাছে **কী কী উপাদান রয়েছে** What materials for anal cleansing are present inside or immediately outside the latrine? | *(1= হ্যাঁ [Yes], 0= না [No],* **প্রযোজ্য নয়** *[*not applicable] =888)]<br>1.*পাতা/ঘাস* Leaves/grass<br>2.*কাঠি* Twigs / sticks<br>3. *কাপড়* Rag or cloth<br>4. *পাথর* Stones<br>5. *স্বাস্থ সম্মত কাগজ*Hygienic (toilet) paper<br>6. *পানির পাত্র/বদনা/মগ* Water container / vessel<br>7. *পানির ট্যাপ* Water tap<br>8. *সাবান* Soap<br>9. *ছাই অথবা মাটি* Ash or soil for cleansing<br>10. *খবরের কাগজ* Newspaper<br>11. *কিছুই নাই* Nothing |
| C. 809 | 812. | **আপনার খানার কে কে এই পায়খানা/টয়লেটটি পায়খানার জন্য ব্যবহার করে?** Please tell me about who in your family uses the latrine for defecation. | |
| 1 | 1 | < **3** বাচ্চা Children <3 | 1. সব সময় [Always]<br>2. প্রায়ই [Usually]<br>3. মাঝেমাঝে [Sometime]<br>4. কখনই না [Never]<br>888. প্রযোজ্য নয় [ N/A] |
| 2 | 2 | ৩-<৮ বাচ্চা Children 3-<8 | 1. সব সময় [Always]<br>2. প্রায়ই [Usually]<br>3. মাঝেমাঝে [Sometime]<br>4. কখনই না [Never]<br>888. প্রযোজ্য নয় [ N/A] |

| | | | |
|---|---|---|---|
| 3 | 3 | ৮-১৫ বাচ্চা Children 8-15 | 1. সব সময় [Always]<br>2. প্রায়ই [Usually]<br>3. মাঝেমাঝে [Sometime]<br>4. কখনই না [Never]<br>888. প্রযোজ্য নয় [ N/A] |
| 4 | 4 | পুরুষ Men | 1. সব সময় [Always]<br>2. প্রায়ই [Usually]<br>3. মাঝেমাঝে [Sometime]<br>4. কখনই না [Never]<br>888. প্রযোজ্য নয় [ N/A] |
| 5 | 5 | মহিলা Women | 1. সব সময় [Always]<br>2. প্রায়ই [Usually]<br>3. মাঝেমাঝে [Sometime]<br>4. কখনই না [Never]<br>888. প্রযোজ্য নয় [ N/A] |
| C. 810 | 813. | অন্য কোন খানা কি এই পায়খানা ব্যবহার করে? Do you share this toilet with other households? | *1= হ্যাঁ [Yes]*<br>*0= না [No].......skip to 815* |
| C.811 | 814. | কতগুলো খানা মিলে এই পায়খানাটি ব্যবহার করে ? How many HHs use it? | (Don't know=99)<br>I_____I I_____I |
| C.812 | 815. | এইখানার শিশুসহ আপনারা কতজন এই পায়খানাটি ব্যবহার করেন? How many people including children in your household use this toilet? | I_____I I_____I |
| C. 813 | 816. | পায়খানার মালিকানা ? (প্রশ্ন করুন) [**Ask**: Who owns the toilet facility?] | শুধুমাত্র ঐ খানার জন্য [Only for the household] 1<br>কয়েকঘর মিলে/ অংশীদার [Shared] 2<br>অন্য কেউ [Someone else] 3<br>পাবলিক [Public] 4<br>প্রযোজ্য নয় [Not applicable]................................................888 |
| C. 814 | 817. | এই স্থানে এই পায়খানাটি কত বছর ধরে আছে? (জানি না=999) [How long have you had the present latrine in this place? [In years] (Don't know = 99)] | A. \|_____\|\|_____\| বৎসর Years<br>B. \|_____\|\|_____\| মাস Months |

#### Wash Benefit Module 9 Child defecation and feces disposal practice

Enter Child ID |_____|____|____|____|____|_____|

| শিশুর পায়খানা ব্যবস্থাপনা এবং মল নিষ্কাশন অভ্যাস **Child defecation and feces disposal** | | | |
|---|---|---|---|
| Administer to: All study households ( সকল খানায় হবে) | | | |
| C.912 | 901. | \|_____\|\|_____\| কত বছর বয়স থেকে একটি শিশু সাধারনতঃ টয়লেট/পায়খানা ব্যবহার করা শুরু করে (মাসে)? [At what age (in months) does a child start using a latrine, if at all?] | |
| | 901_1 | আপনার খানায় <৩ বছরের যে শিশুটি আছে তার বয়স কত (মাসে)? (একাধিক শিশু থাকলে সবচেয়ে ছোট বাচ্চার ক্ষেত্রে জিজ্ঞেস করুন) [Age of the youngest child (<3 years)] | ........................... মাস [Months] |
| C.902 | 902. | \|_____\| আপনার (<□ বছর) শিশু | 1. আজ [Today]<br>2. গতকাল [Yesterday] |

| | | | |
|---|---|---|---|
| | | সর্বশেষ কখন পায়খানা করেছে? (একাধিক শিশু থাকলে সবচেয়ে ছোট বাচ্চার ক্ষেত্রে জিজ্ঞেস করুন) [When was the last time your youngest child/infant (<3 years) defecated?] | 3. ২ দিন বা তার আগে [Before 2 or more days back]<br>4. কখনই না/ মনে করতে পারছি না [Never/can't remember]....... **906** নং প্রশ্নে চলে যান (skips to 906)<br>5. বলতে রাজি না [Refused] ..........**906** নং প্রশ্নে চলে যান (skips to 906)<br>88. প্রযোজ্য নয় [Not applicable]......... **908** নং প্রশ্নে চলে যান (skips to 908) |
| **C.903.** | 903. | \|_____\| আপনার শিশু সর্বশেষ কোথায় পায়খানা করেছে? [Where did the child defecate the last time?]<br>**Note:** উত্তর পড়ে শোনাবেন না, তাকে নির্দিষ্ট করে বলতে বলুন । এ বিষয়ে যতক্ষণ পর্যন্ত কিছু বলার থাকে ততক্ষণ পর্যন্ত তাকে বলতে উৎসাহিত করুন<br>[Don't read the answer, encourage by asking if there is anything else until he/she mentions there in nothing else and check all mentioned.] | 1. পটি (উঠানে) [Potty (in the courtyard)]<br>2. পটি (ঘরের ভিতরে) [Potty (inside the house)]<br>3. উঠানে (পটি ছাড়া) [Courtyard (without potty)]<br>4. ঘরের ভিতরে (পটি ছাড়া) [Inside the house (without potty)]<br>5. উঠানের বাহিরে খোলা জায়গাতে [Open space outside the front yard<br>6. ঝোপ-ঝাড়ে/জঙ্গলে Bush/jungle<br>7. পায়খানায়/টয়লেটে [In toilet] ......**906** নং প্রশ্নে চলে যান (Skip to 906)<br>8. কাথা/ন্যাপি/ডায়পার Katha/nappy/diaper<br>77. অন্যান্য (বর্ণনা লিখুন) [Other (Please describe)]....................<br>99. জানি না [Don't Know]............. **906** নং প্রশ্নে চলে যান (Skip to 906) |
| **C.904.** | 904. | \|_____\| সেই পায়খানা কি করা হয়েছিল? [What was done to the feces?] | <u>**হ্যাঁ [Yes].....1, না [No]......0**</u><br>1. যেখানে পায়খানা করেছিল সেখানেই ফেলে রাখা হয়েছিল [It is left there] **905.a** নং প্রশ্নে চলে যান (Skip to 905a)<br>2. \|_____\| টয়লেট/পায়খানার ভিতরে ফেলা/ধোয়া হয়েছিল [Put/rinsed into toilet or latrine]<br>3. \|_____\| ড্রেনে/নর্দমার ভিতরে ফেলা/ধোয়া হয়েছিল [Put/rinsed into drain or ditch]<br>4. \|_____\|ঝোপ-ঝাড়ে/জঙ্গলে ফেলা হয়েছিল [Thrown into Bush/jungle<br>5. \|_____\| ময়লা আবর্জনার মধ্যে ফেলা হয়েছিল [Thrown into garbage]<br>6. \|_____\| নির্দিষ্ট গর্তে ফেলা হয়েছিল [Thrown into a specific pit for child's feces]<br>7. \|_____\| মাটির নীচে পুতে ফেলা হয়েছিল [Buried]<br>8. \|_____\|পুকুর/ টিউবওয়েলের পানিতে ধোয়া হয়েছিল Rinsed into Tubewell or pond<br>77. \|_____\| অন্যান্য (বর্ণনা লিখুন) [Other (specify)]....<br>99. \|_____\| জানি না [DK]**906** নং প্রশ্নে চলে যান (Skip to 906) |
| **C.905.** | 905. | আপনি কিভাবে পায়খানা পরিষ্কার করেন? উত্তর পড়ে শুনানো যাবে না \|হ্যাঁ হলে ১ বসান ।did you handle the feces? ☟<u>Do Not Read Responses. Mark All that Apply</u> (1 = Yes) | 1. \|_____\|নগ্ন/খালি হাতে Hands only (bare hands)<br>2. \|_____\|হাতে এবং কাপড়/পাতা/কাগজ Hands and cloth / paper / leaves<br>3. \|_____\| পায়খানা ফেলার জন্য কোন উপাদান ব্যবহার করে Scrap material to scoop fece<br>4. \|_____\|পটি Potty<br>5. \|_____\|স্থানীয় কৃষিকাজ-এর হাতিয়ার Local agricultural hoe/instrument<br>6. \|_____\|সেনি স্কুপ Sani-scoop<br>7. \|_____\|কোন কিছুই করা হয় না Did nothing<br>77 \|_____\|অন্যান্য (উল্লেখ করুন)Others (specify)___________<br>99 \|_____\|জানি না Don't know / not sure |

| | | | |
|---|---|---|---|
| C.905.b | 905.a | পায়খানা করার পর কতক্ষণ এই পায়খানা এখানে পরে ছিল? How long after defecation did you dispose of the feces? | \|____\|Hour ঘন্টা \|____\|Minute মিনিট<br>OR<br>\|____\| দিন days |
| **C.906.** | 906. | আপনার ৩ বছরের ছোট বাচ্চার পায়খানা কে কে ফেলে? (যতজন ফেলে সবাইকে কোড করুন) [Who disposes your under 3 child's feces? | **হ্যাঁ [Yes].....1, না [No]......0**<br><br>1. \|_____\| মা [Mother]<br>2. \|_____\| বাবা [Father]<br>3. \|_____\| বোন [Sister]<br>4. \|_____\| ভাই [Brother]<br>5. \|_____\| খালা/মামী/চাচী/ফুফু/দাদী [Aunt/grandmother]<br>6. \|_____\| খালু/মামা/চাচা/ফুফা/দাদা [Uncle/grantfather]<br>7. \|_____\| কেউ পায়খানা ফেলে না [Nobody dispose]<br>77. \|_____\| অন্যান্য (নির্দিষ্ট করে লিখুন) Others (Specify) ....................<br><br>888. \|_____\|প্রযোজ্য নয় [Not applicable] |
| | 907. | ৩ বছরের ছোট বাচ্চার পায়খানা ফেলার স্থানটি রান্না ঘর থেকে কতটুকু দূরে? [How far away is this disposal site to the kitchen?] | \|_____\| কদম দূরে? [steps]<br>**888.** প্রযোজ্য নয় [Not applicable] |
| **C.907.** | 908. | \|_____\| আপনার খানায় ৩-<৮ বছরের অন্য কোন শিশু আছে কি? [Is there any other child between the age of 3-<8 years?] | 1 হ্যাঁ [Yes]<br>0. না [No].......... **913** নং প্রশ্নে চলে যান (skips to 913) |
| **C.908.** | 909. | আপনার খানায় ৩-<৮ বছরের যে শিশুটি আছে তার বয়স কত (মাসে)? (একাধিক শিশু থাকলে সবচেয়ে বড় বাচ্চার ক্ষেত্রে জিজ্ঞেস করুন) [Age of the oldest child (aged 3-8)] | .......................... মাস [Months] |
| **C.909.** | 910. | \|_____\| আপনার শিশু (৩-৮ বছরের) সর্বশেষ কোথায় পায়খানা করেছে? (একাধিক শিশু থাকলে সবচেয়ে বড় বাচ্চার ক্ষেত্রে জিজ্ঞেস করুন) [Where did your oldest child (aged 3-8) defecate the last time?] | 1. পটি (উঠানে) [Potty (in the courtyard)]<br>2. পটি (ঘরের ভিতরে) [Potty (inside the house)]<br>3. উঠানে (পটি ছাড়া) [Courtyard (without potty)]<br>4. ঘরের ভিতরে (পটি ছাড়া) [Inside the house (without potty)]<br>5. উঠানের বাহিরে খোলা জায়গাতে [Open space outside the front yard<br>6. ঝোপ-ঝাড়ে/জঙ্গলে Bush/jungle<br>7. পায়খানায়/টয়লেটে [In toilet] ...**913** নং প্রশ্নে চলে যান (Skip to |

| | | | |
|---|---|---|---|
| | | | 91<br>3)<br>8. কাথা/ন্যাপি/ডায়পার Katha/nappy/diaper<br>77. অন্যান্য (বর্ণনা লিখুন) [Other (Please describe)]....................<br>99. জানি না [Don't Know]............. **913** নং প্রশ্নে চলে যান (Skip to 913) |
| **C.910.** | 911. | \|_____\|সেই পায়খানা কি করা হয়েছিল? [What is done to the feces?] | **হ্যাঁ [Yes].....1, না [No]......0**<br>1. যেখানে পায়খানা করেছিল সেখানেই ফেলে রাখা হয়েছিল [It is left there]**912.a** নং প্রশ্নে চলে যান (Skip to 912.a)<br>2. \|_____\| টয়লেট/পায়খানার ভিতরে ফেলা/ধোয়া হয়েছিল [Put/rinsed into toilet or latrine]<br>3. \|_____\| ড্রেনে/নর্দমার ভিতরে ফেলা/ধোয়া হয়েছিল [Put/rinsed into drain or ditch]<br>4. \|_____\|ঝোপ-ঝাড়ে/জঙ্গলে ফেলা হয়েছিল [Thrown into Bush/jungle<br>5. \|_____\| ময়লা আবর্জনার মধ্যে ফেলা হয়েছিল [Thrown into garbage]<br>6. \|_____\| নির্দিষ্ট গর্তে ফেলা হয়েছিল [Thrown into a specific pit for child's feces]<br>7. \|_____\| মাটির নীচে পুতে ফেলা হয়েছিল [Buried]<br>77. \|_____\| অন্যান্য (বর্ণনা লিখুন) [Other (specify)]................<br>99. \|_____\| জানি না [DK]**913** নং প্রশ্নে চলে যান (Skip to 913) |
| **C.910.** | 912. | আপনি কিভাবে পায়খানা হ্যান্ডলিং (ব্যবস্থাপনা )করেন? How did you handle the feces? ☟ Do Not Read Responses. Mark All that Apply (1 = Yes) | ☐ 1 নগ্ন/খালি হাতে Hands only (bare hands)<br>☐ 2 হাতে এবং কাপড়/পাতা/কাগজ Hands and cloth / paper / leaves<br>☐ 3 ফেলানো সামগ্রী মাটিতে পুঁতে Scrap material to scoop feces<br>☐ 4 পটি Potty<br>☐ 5 স্থানীয় কৃষিকাজ-এর হাতিয়ার Local agricultural hoe/instrument<br>☐ 6 সেনি স্কুপ Sani-scoop<br>☐ 7 কোন কিছুই করা হয় না Did nothing<br>☐ 77 অন্যান্য (উলেস্নখ করুন) Others (specify)___________<br>☐ 99 জানি না Don't know / not sure |
| C.910.b | 912.a | পায়খানা করার পর কতৰণ এই পায়খানা এখানে পরে ছিল? How long after defecation did you dispose of the feces? | \|____\|Hour ঘন্টা<br>\|____\|Minute মিনিট<br>OR \|____\| দিন DAYS |

| | | | |
|---|---|---|---|
| **Child Potty Use**<br>**( শিশুর পটি ব্যবহার সম্পর্কীত তথ্য)** | | | |
| Administer to: All study households ( সকল খানায় হবে) | | | |
| C.8 15 | 913. | Does your household have a potty that children use for defecation?<br>আপনার খানাতে কি শিশুদের পায়খানা করার পটি আছে? | 1 Yes হ্যাঁ<br>2 No না **(Skip to 919)** (919 নং প্রশ্নে যান)<br>99 Don't know / not sure জানি না **(Skip to 919)** (919 নং প্রশ্নে যান) |
| C.8 16 | 914. | In the last week, how often did your child use the potty? গত এক সপ্তাহের মধ্যে, শিশুটি কতবার পটি ব্যবহার করছে?<br>[ *If multiple children use the potty, ask about the youngest child* ][যদি একাধিক শিশু ব্যবহার করে থাকে তবে যে সবচেয়ে বেশী ব্যবহার করে সেই শিশুর জন্য প্রশ্ন করম্নন] | 1 Every time প্রতিবার<br>2 More than half of all defecation events, but not every time যতবার পায়খানা করেছে তার ৫০ ভাগ বা তার বেশী কিন্তু প্রতিবার নয়<br>3 যতবার পায়খানা করেছে তার ৫০ ভাগের কম কিন্তু মাঝেমাঝে Less than half of all defecation events<br>4 আগে করেছি কিন্তু এখন আর করি না Used to use it, but no longer use it **(Skip to 919)** (919 নং প্রশ্নে যান)<br>5 কখনই না Never **(Skip to 919)** (919 নং প্রশ্নে যান)<br>99 জানি না Don't know / not sure **(Skip to 919)** (919 নং প্রশ্নে যান) |
| C.8 17 | 915. | What is the age of the child (or children) who is using the potty? যে শিশুটি (বা শিশুরা) পটি ব্যবহার করছে তার বয়স কত? | <u>Mark All that Apply.</u> Yes = 1 হ্যাঁ = 1, না = 0<br>1 ☐ < 1 year ১ বছরের কম<br>2 ☐ 1- < 3 years ১-<৩ বছর<br>3 ☐ 3- < 5 years ৩-<৫ বছর<br>4 ☐ >= 5 years >=৫ বছর |
| C.8 18 | 916. | Ask the respondent to identify the place where the child typically uses the potty. Then ask her to retrieve the potty<br>Could I please see the potty?<br>👁 Observation: *Record how long it takes to produce the potty.*<br>আমি কি পটিটি দেখতে পারি<br>👁 আমি কি পটিটি দেখতে পারি (পটিটি আনতে কতটুকু সময় লেগেছে তা সেকেন্ডে গণনা করম্নন) | ☐☐:☐☐ MINUTES : SECONDS (99:99 if could not measure) মিনিট : সেকেন্ড (পটি দেখাতে না পারলে 999 কোড করুন) |

| | | | |
|---|---|---|---|
| C.8<br>19 | 917. | 👁 Observation: Potty condition.<br>পটিটির অবস্থা কি তা পর্যবেৰণ করম্নন (একাধিক উত্তর হতে পারে) | Mark All that Apply. Yes = 1 হ্যাঁ = 1, না = 0<br>1.Easily accessible when needed by the child প্রয়োজনের সময় বাচ্চা যেন পায় এমন কাছাকাছি জায়গাতে রয়েছে<br>2.Easily accessible when needed by the mother প্রয়োজনের সময় মা যেন পায় এমন কাছাকাছি জায়গাতে রয়েছে<br>3.Visible signs of feces inside / on the potty / removable pot ভিতরে/ পটিতে পায়খানার চিহ্ন আছে<br>4.Potty was covered with the lid পটি লিড দিয়ে ঢাকা ছিল<br>5. Potty was covered with anything other than the lid পটি লিড ছাড়া অন্য কিছু দিয়ে ঢাকা ছিল<br>6.Dry শুকনা<br>7.Broken/or missing parts(tray) ভেঙ্গে গেছে<br>8.Covered in dust / signs of non-use পটি ধুলায় ঢাকা/ ব্যবহার না করার চিহ্ন রয়েছে<br>9.Cannot produce a potty পটি দেখাতে পারে নাই |
| C.8<br>20 | 918. | ☐Where do you usually dispose of feces from the potty? সাধারনতঃ এই পটি থেকে আপনি পায়খানা কোথায় ফেলেন? | 1 Latrine পায়খানায়/টয়লেটে<br>2 Open Pit / separate pit for child or animal fecesখোলা গর্তে / শিশুর বা পশুপাখির পায়খানা ফেলার গর্তে<br>3 Bury it / Covered Pit মাটির নীচে পুতে ফেলা হয়েছিল/ ঢাকা গর্তে<br>4 Undefined open site near the compound (including open garbage disposal sites / dumps) উঠানের কাছে খোলা জায়গা<br>5 Bush / forest / fieldঝোপ-ঝাড়ে/জঙ্গলে/মাঠে<br>6 Nearby water (pond, canal, river) পানির উৎসের কাছে (পুকুর/সেচ নালা/ নদী)<br>77 Other (specify) অন্যান্য (লিখুন)______________ |
| **Sani Scoop Use**<br>**(সেনিস্কুপ ব্যবহার)** | | | |
| Administer to: All households ( সকল খানায় হবে) | | | |
| C.8<br>21 | 919. | ☐Does your household have a dedicated tool [sani scoop] to clean up feces around your household?<br>আপনার খানার চারপাশের পায়খানা পরিষ্কার করার জন্য আপনার খানায় কোন নির্দিষ্ট কিছু আছে কি? | 1 Yes, icddrb sani scoop *হ্যাঁ আইসিডিডি আরবির সেনিস্কুপ*<br>2. Yes, other household scoop *হ্যাঁ অন্যখানার সেনিস্কুপ*<br>3 No না (1001 নং প্রশ্নে যান) **(Skip to 1001)**<br>4 Own Household Scoop (□□□□□ □□□□□ □□□□□□□□□□)<br>99 Don't know / not sure জানি না (1001 নং প্রশ্নে যান) **(Skip to 1001)** |
| C.8<br>22 | 920. | How often do you use the [sani scoop]?<br>আপনি কখন কখন | 1 Multiple times per day প্রতিদিন কয়েকবার<br>2 Once per day প্রতিদিন একবার<br>3 A few times each week প্রতি সপ্তাহে কয়েকবার |

| | | | |
|---|---|---|---|
| | | পায়খানা পরিষ্কার করার জন্য কোদাল/সেনিস্কুপ ব্যবহার করেন? | 4 Less than once per week সপ্তাহে একবারের কম<br>5 Used to use it, but no longer use it প্রথমদিকে ব্যবহার করেছে কিন্তু এখন আর করে না (1001 নং প্রশ্নে যান) **(Skip to 1001)**<br>6 Never কখনই না (1001 নং প্রশ্নে যান) **(Skip to 1001)** |
| C.8 23 | 921. | What do you use the [sani scoop] for? ☟ Do Not Read Responses. আপনি কি কি কাজে কোদাল/সেনিস্কুপ ব্যবহার করেন? (একাধিক উত্তর হতে পারে) ☟ উত্তর পড়ে শোনাবেন না | Mark All that Apply. Yes = 1 হ্যাঁ = 1, না = 0<br>1 ☐ Clean up animal feces জীবজন্তুর পায়খানা ফেলার কাজে<br>2 ☐ Clean up child feces বাচ্চার পায়খানা ফেলার কাজে<br>3 ☐ Clean up garbage ময়লা আবর্জনা ফেলার কাজে<br>4 ☐ Take the scoop to the field (for work) কৃষিকাজে/মাঠে ইহা নিয়ে যাই<br>5 ☐ Digging / gardening গর্ত তৈরীর কাজে<br>77 ☐ Other (specify) অন্যান্য (বর্ণনা লিখুন)______ |
| C.8 24 | 922. | Could I please see the [sani scoop]? 👁 Observation: Sani scoop condition. আমি কি কোদাল/সেনিস্কুপটি দেখতে পারি? (একাধিক উত্তর হতে পারে) 👁 পর্যবেৰণ: সেনিস্কুপটির অবস্থা পর্যবেৰণ করম্নন | Mark All that Apply. Yes = 1 হ্যাঁ = 1, না = 0<br>1 ☐ Visible signs of feces on the sani scoop কোদাল/সেনিস্কুপে পায়খানার চিহ্ন আছে<br>2 ☐ Dry শুকনা<br>3 ☐ Broken and needs repair ভেঙ্গে গেছে এবং মেরামত করা দরকার<br>4 ☐ Easily accessible when needed by an adult প্রয়োজনের সময় বড়রা যেন পায় এমন কাছাকাছি জায়গাতে রয়েছে<br>5 ☐ Signs that the sani scoop is not used কোদাল/সেনিস্কুপ ব্যবহার না করার চিহ্ন রয়েছে<br>6 ☐ Cannot produce a sani scoop কোদাল/সেনিস্কুপ দেখাতে পারে নাই |
| C.8 25 | 923. | I'd like to ask a few more questions about the types of feces you pick up with the [sani scoop] and how you dispose of the different types of feces. [Ask about each type of feces separately.]. কোদাল/সেনিস্কুপ দিয়ে আপনি | Where do you dispose the feces? পায়খানা কোথায় ফেলা হয়েছিল?<br>☟ Do Not Read Responses. উত্তর পড়ে শোনাবেন না<br>Mark All that Apply (একাধিক উত্তর হতে পারে) 1 = হ্যাঁ Yes, না = 0,<br>88 Not applicable প্রযোজ্য নয়<br>Columns: (blank) / A / B / C / D / E / F<br>Feces পায়খানা / Use Sani Scoop for disposal? পায়খানা ফেলার জন্য কোদাল/সেনিস্কুপ ব্যবহার / Latrine পায়খানায় / Open Pit খোলা গর্তে / Bush/Farm ঝোপে ঝাড়ে/খোলা জায়গায় / Surface Water পানিতে / Dig Hole and cover গর্ত করা হয় এবং ঢেকে রাখা হয় |

| | | | | |
|---|---|---|---|---|
| | | যেসব পায়খানা ফেলেন এবং কিভাবে ফেলেন সেসম্পর্কে জানার জন্য এখন আমি আপনাকে আরো কিছু প্রশ্ন করতে চাই। (প্রতিটি পায়খানার ধরনের কথা উলেস্নখ করে কোড করম্নন) (শিশু/পশুপাখির পায়খানা ফেলার কাজে ব্যবহার না করলে প্রযোজ্য নয় ঘরটিতে টিক দিন) | | করে? |
| | | | 1 Child (<8 years) শিশুর (<৮ বছর) | ☐ |
| | | | 2 Cow গোবর | ☐ |
| | | | 3 Poultry / pigeons হাঁস-মুরগীর | ☐ |
| | | | 4 Goat ছাগলের | ☐ |
| | | | 5 Pig শুকরের | ☐ |
| | | | 6 Dog or cat কুকুর বা বিড়ালের | ☐ |
| | | | 88 Not applicable | ☐ |

#### Wash Benefit Module 10 Water Access

আপনাকে ধন্যবাদ। আমি আপনাকে এবং আপনার খানার সদস্যদের বাড়ীতে খাবার পানি বিষয়ে কিছু প্রশ্ন করতে চাই।

Thank you very much. Now, I would like to ask you some questions about the water you and your family drink at home.

| WATER TREATMENT, STORAGE, AND QUALITY (All households) | | | |
|---|---|---|---|
| SAY | Thank you very much. Now, I would like to ask you some questions about the water you and your family drink at home. | | |
| C.1001 | 1001. | আপনি কিভাবে খাবার পানি সংরক্ষন করেন ? [How do you store drinking water?<br><br>প্রশ্ন করম্নন এবং সম্ভব হলে পর্যবেক্ষন করম্নন ASK THE QESTION AND OBSERVE IF POSSIBLE | [1] প্লাষ্টিক অথবা ধাতব পাত্রে IN PLASTIC OR METAL CONTAINERS<br>[2] মাটির পাত্রে IN CLAY POTS<br>[3] ছাদে পানির ট্যাংক ROOF TANK OR CISTERN<br>[4] পানি সংরক্ষণ করা হয় না DO NOT STORE WATER<br>[5] icddrb Topaz *আইসিডিডিআরবির টোপাস* |
| C.1002 | 1002. | এই খানায় ০-<৩৬ মাসের কোন শিশু আছে কি যে পানি পান করে ?Is there a child 0-<36 months in this household who drinks water? | [1] হ্যাঁ YES<br>[2] না NO<br>[88] Not applicable প্রযোজ্য নয় |
| C.1003 | 1003. | **IF YES → প্রশ্ন করম্নন এবং পর্যবেক্ষন করম্ননঃ যদি আপনার শিশু (<৩ বছর) এখন পানি খেতে চাইত তাহলে আপনি কিভাবে তাকে পানি দিতেন তা আমাকে দেখান** ASK: If your child 0-3 years wanted a | |

| | | | | | |
|---|---|---|---|---|---|
| | | drink of water right now, could you show me how you would give it to him/her?<br>**IF NO** → ASK: **যদি আপনি এখন পানি খেতে চাইতেন তাহলে আপনি কিভাবে পানি নিতেন তা আমাকে দেখান** If <u>you</u> wanted a drink of water right now, could you show me how you would get it?<br><br>উত্তরদাতার পানি দেওয়ার প্রক্রিয়াকে পর্যবেক্ষণ/অনুসরণ করুন । যদি অন্য ঘরে পানি সংরক্ষিত এবং সেখানে প্রবেশাধিকার থাকে তাহলে প্রক্রিয়াকে পর্যবেক্ষণ/অনুসরণ করুন । অন্যথায় ৯৯ কে গোলাকার করতে হবে। ATTEMPT TO FOLLOW RESPONDENT WHEN SHE GOES TO RETRIEVE THE WATER. IF WATER IS STORED IN ANOTHER ROOM, ASK HER IF IT IS OKAY TO ENTER THAT ROOM. CIRCLE "99" IF YOU CANNOT OBSERVE. | | | |
| | | 👁 উত্তরদাতা পানি নেওয়ার আগে কি করেছিল ? WHAT DID RESPONDENT DO <u>BEFORE TAKING THE WATER</u>? | **হ্যাঁ** YES | **না** NO | **জানি না** D/K |
| 2 | 2 | **গ্লাস বা পাত্রটিতে পানি ঢালার পূর্বে গ্লাস/পাত্রটি খাবার পানি দিয়ে ধুয়েছিল** RINSED GLASS/CONTAINER WITH DRINKING WATER BEFORE FILLING | [1] | [2] | [99] |
| 3 | 3 | **পানি ঢালার পূর্বে হাত ধুয়েছিল** WASHED HANDS WITH WATER BEFORE DRINKING WATER WAS OBTAINED | [1] | [2] | [99] |
| 4 | 4 | **পানি ঢালার পূর্বে সাবান দিয়ে হাত ধুয়েছিল** WASHED HANDS WITH SOAP BEFORE DRINKING WATER OBTAINED | [1] | [2] | [99] |
| | | 👁 উত্তরদাতা কোথা থেকে পানি সংগ্রহ করেছিল? <u>FROM WHERE</u> DID THE RESPONDENT TAKE THE WATER? | **হ্যাঁ** YES | **না** NO | **জানি না** D/K |
| 5 | 5 | সরাসরি পানির উৎস থেকে BROUGHT DIRECTLY FROM WATER SOURCE | [1] | [2] | [99] |
| 6 | 6 | সরাসরি খোলা পানির পাত্র থেকে BROUGHT IRECTLY FROM uncovered STORAGE CONTAINER | [1] | [2] | [99] |
| 7 | 7 | সরাসরি পানির ফ্লিটার থেকে BROUGHT DIRECTLY FROM WATER FILTER | [1] | [2] | [99] |
| | 8 | | [1] | [2] | [99] |
| | 8a | | | | |
| | 8b | সরাসরি ঢেকে রাখা পানির পাত্র থেকে STORED WATER WAS COVERED | [1] | [2] | |
| | 8c | *আইসিডিডিআরবির টোপাস* থেকে From icddrb topaz | | | [99] |
| | | *আইসিডিডিআরবির টোপাস* থেকে পরিশোধনকৃত পানি এনেছিল Brought treated water *আইসিডিডিআরবির টোপাস* থেকে (অনুরোধ করার পর) from icddrb topaz (umprompted) | [1]<br>[1] | [2]<br>[2] | [99]<br>[99] |
| | | *আইসিডিডিআরবির টোপাস* থেকে পরিশোধনকৃত পানি এনেছিল (অনুরোধ করার পর) Brought untreated water directly from source or other container (unprompted) | [1] | [2] | [99] |
| | | 👁 উত্তরদাতা কিভাবে গ্লাসে/কাপে/পাত্রে পানি দিয়েছিল? <u>HOW DID SHE GET THE WATER</u> INTO THE CUP? | **হ্যাঁ** YES | **না** NO | **জানি না** D/K |
| 9 | 9 | **গ্লাস বা পাত্রটিতে ঢালা পানির ভিতরে তার হাত বা হাতের আঙ্গুল লেগেছিল কি?** HANDS TOUCHED / CONTACTED THE DRINKING WATER | [1] | [2] | [9 ] |
| 10 | 10 | **গ্লাস বা পাত্রটিতে পানি ভরার সময় তা পানির পাত্রের ভিতর ডুবিয়ে পানি ভরা হয়েছিল** CONTAINER/GLASS DIPPED INTO WATER CONTAINER | [1] | [2] | [99] |
| 11 | 11 | **গ্লাসে পানি ভরার জন্য লম্বা হাতলযুক্ত কোন চামচ/মগ ব্যবহার করা হয়েছিল** | [1] | [2] | [99] |

| | | | | | |
|---|---|---|---|---|---|
| | | LADLE USED TO OBTAIN WATE | | | |
| 12 | 12 | পানির পাত্র থেকে সরাসরি পানি ঢেলেছিল WATER POURED FROM CONTAINER | [1] | [2] | [99] |
| 13 | 13 | সরাসরি টিউবওয়েল/ পানির উৎস থেকে এনেছে WATER POURED FROM TAP | [1] | [2] | [99] |

| | | | | |
|---|---|---|---|---|
| C.100 4 | 1004. | আপনি/এ খানার অন্যকেউ কতক্ষণ আগে এই সংরক্ষণকৃত পানি সংগ্রহ করেছেন? How long ago did you or somebody in your home collect this water? ( 99 = Don't know জানি না; 87 = No Water কোন পানি নাই, 88 Not applicable প্রযোজ্য নয় ) If 88, skip to 1007 | A.\|__\|__\| ঘন্টা HOURS<br>B.\|__\|__\| দিন DAYS | |
| C.100 5 | 1005. | এই খাবার পানি নিরাপদ করার জন্য কোন কিছু করেছেন কি? Have you done anything to make this water less cloudy or safer to drink? | [1] হ্যাঁ YES<br>[2] না NO<br>[99] জানি না DON'T KNOW/NOT SURE | → SKIP to 1007<br>→ SKIP to 1007 |
| C.100 6 | 1006. | কোন পদ্বতি কি আপনি ব্যবহার করেন? What method(s) did you use? উত্তর পড়ে শুনাবেন না \|উত্তরদাতাকে উত্তর বলতে সাহায্য করা যাবে তার উত্তর দেয়া শেষ হলে ।DO NOT READ RESPONSES. PROBE UNTIL RESPONDENT IS FINISHED. CIRCLE ALL THAT APPLY. | | |
| 1 | 1 | আকুয়াট্যাবস_Aquatabs | [1] হ্যাঁ Yes [0] না No. If yes ='1' then skip to 1022 | |
| 2 | 2 | ক্লোরিন (তরল) মিশিয়ে নেই (ওয়াটার গার্ড নয়) Waterguard / bottled chlorine | [[1] হ্যাঁ Yes [0] না No | |
| 3 | 3 | পানি ফুটিয়ে নেই Boil | [1] হ্যাঁ Yes [0] না No | |
| 4 | 4 | কাপড়ে ছেকে নেই Strain it through clothor other material | [1] হ্যাঁ Yes [0] না No | |
| 5 | 5 | পানির ফিল্টার ব্যবহার করে (সিরামিক, বালু ইত্যাদি) Water filter [ceramic, sand, composite] | [1] হ্যাঁ Yes [0] না No | |
| 6 | 6 | সূর্যের আলোতে জীবানুমুক্তকরণ Solar disinfection (SODIS) | [1] হ্যাঁ Yes [0] না No | |
| 7 | 7 | থিতিয়ে নেই Let it stand and settle | [1] হ্যাঁ Yes [0] না No | |
| 8 | 8 | স্যান্ড ফিল্টার এর মাধ্যমে বিশুদ্ধ করে নেই Biosand filter | [1] হ্যাঁ Yes [0] না No | |
| 9 | 9 | ফিটকিরি যোগ করে Coagulant (alum) | [1] হ্যাঁ Yes [0] না No | |
| 10 | 10 | পার PUR (flocculant + disinfectant) | [1] হ্যাঁ Yes [0] না No | |

| | | | | |
|---|---|---|---|---|
| C.1007 | 1007. | আপনি কখনও আপনার খাবার পানি পরিশোধন করার জন্য কিছু করেছিলেন কি? Do you ever treat your drinking water or do anything to make it less cloudy? | [1] হ্যা YES<br>[0] না NO | → **1010** নং এ চলে যান **SKIP TO 1010** |
| C.1008 | 1008. | **সর্বশেষ কখন আপনি** আপনার খাবার পানি পরিশোধন করেছিলেন ? When was the last time you treated your water or did anything to make it less cloudy?<br><br>উত্তর পড়ে শুনানো যাবে না DO NOT READ RESPONSES | [1] আজ TODAY<br>[2] গতকালYESTERDAY<br>[3] গত সপ্তাহের মধ্যে WITHIN THE PAST WEEK<br>[4] গত দুই সপ্তাহের মধ্যে WITHIN THE PAST 2 WEEKS<br>[5] গত মাসের মধ্যে WITHIN THE PAST MONTH<br>[6] গত বছরের মধ্যে WITHIN THE PAST YEAR<br>[99] জানি না/নিশ্চিত না DON'T KNOW/NOT SURE | |

| | | | |
|---|---|---|---|
| C.1009 | 1009. | কোন কোন উপায়ে আপনি পানি পরিশোধন করে থাকেন ? What are all the ways you treat your drinking water? উত্তর পড়ে শুনানো যাবে না । উত্তর দেয়া শেষ হলে উত্তর পেতে সাহায্য করা যেতে পারে ।DO NOT READ RESPONSES. PROBE UNTIL RESPONDENT IS FINISHED. CIRCLE ALL THAT APPLY. | |
| 1 | 1 | আকুয়াট্যাবস_Aquatabs | [1] হ্যাঁ Yes [0] না No |
| 2 | 2 | ক্লোরিন (তরল) মিশিয়ে নেই (ওয়াটার গার্ড নয়) Waterguard / bottled chlorine | [[1] হ্যাঁ Yes [0] না No |
| 3 | 3 | পানি ফুটিয়ে নেই Boil | [1] হ্যাঁ Yes [0] না No |
| 4 | 4 | কাপড়ে ছেকে নেই Strain it through clothor other material | [1] হ্যাঁ Yes [0] না No |
| 5 | 5 | পানির ফিল্টার ব্যবহার করে (সিরামিক, বালু ইত্যাদি) Water filter [ceramic, sand, composite] | [1] হ্যাঁ Yes [0] না No |
| 6 | 6 | সূর্যের আলোতে জীবানুমুক্তকরণ Solar disinfection (SODIS) | [1] হ্যাঁ Yes [0] না No |
| 7 | 7 | থিতিয়ে নেই Let it stand and settle | [1] হ্যাঁ Yes [0] না No |
| 8 | 8 | স্যান্ড ফিল্টার এর মাধ্যমে বিশুদ্ধ করে নেই Biosand filter | [1] হ্যাঁ Yes [0] না No |
| 9 | 9 | ফিটকিরি যোগ করে Coagulant (alum) | [1] হ্যাঁ Yes [0] না No |
| 10 | 10 | PUR (flocculant + disinfectant) | [1] হ্যাঁ Yes [0] না No |

র উৎস **DRINKING WATER SOURCES (ALL HOUSEHOLDS)**

উত্তর দাতাকে তার **পানির প্রধান এবং দ্বিতীয় উৎস বিষয়ে প্রশ্ন করুন** ASK THE FOLLOWING QUESTIONS ABOUT THE RESPONDENT'S PRIMARY AND SECONDARY WATER SOURCES.

| | | | |
|---|---|---|---|
| C.1016 | 1010. | **আপনার খানায় ব্যবহৃত পানির প্রধান উৎস কি** ? What type of | 1 **অগভীর টিউবওয়েল** Shallow tubewell |

| | | | |
|---|---|---|---|
| | | water source does your household collect most of the water that you use from? | 2 গভীর টিউবওয়েল Deep tubewell<br>3 ঘরের ভিতর ট্যাপ বা পাইপের পানি Piped into home<br>4 উঠানে ট্যাপ বা পাইপের পানি Piped into yard<br>5 পাতকূয়া (উঠানে) Borewell in yard<br>6 পাবলিক ট্যাপ Public tap<br>7 পাবলিক পাতকূয়া Public borewell<br>8 অগভীর পাকা পাতকূয়া Shallow well with concrete reinforcement<br>9 অগভীর কাঁচা পাতকূয়া Shallow well (no concrete)<br>10 নদী/বাধ/লেক/পুকুর/সেচ নালা থেকে সরাসরি সংগৃহীত পানি River, lake, stream<br>11 অন্যান্য (লিখুন) Other: _____________ |
| C.1017 | 1011. | আপনি আমাকে বলবেন আপনার বাড়ী থেকে প্রধান পানির উৎসে একবার যেতে কতক্ষণ সময় লাগে? Can you tell me how long it takes you to walk one-way to this PRIMARY water source from your home? | \|__\|ঘন্টা Hours<br>\|__\|মিনিট Min |
| C.1018 | 1012. | আপনি আমাকে বলবেন কোন কোন কাজের জন্য এই উৎসের পানি ব্যবহার করেন Can you please tell me whether you use water from this source for… | |
| 1 | 1 | পানি পান Drinking | [1] হ্যাঁ Yes [0] না No |
| 2 | 2 | রান্না-বান্না/বাসন ধোয়া Cooking/ washing dishes | 1] হ্যাঁ Yes [0] না No |
| 3 | 3 | কাপড় ধোয়া Laundry | 1] হ্যাঁ Yes [0] না No |
| 4 | 4 | গোসল/হাতধোয়া Bathing/hand washing | 1] হ্যাঁ Yes [0] না No |
| 5 | 5 | কৃষিকাজ/বাগান Irrigate crop land or garden | 1] হ্যাঁ Yes [0] না No |
| 6 | 6 | পশু-পালন Water for livestock | 1] হ্যাঁ Yes [0] না No |
| C.1019 | 1013. | আপনার খানাকে এই পানির উৎস ব্যবহারের জন্য কি কোন প্রকার টাকা দিতে হয়? your household pay any money to use this source? | [1] হ্যাঁ Yes [0] না No |
| C.1020 | 1014. | আপনার খানায় পানি ব্যবহারের জন্য কি ২য় কোন উৎস ব্যবহার করতে হয়? যদি হয় তাহলে কি ধরনের ? Does your household currently use a secondary water source? If so, what type is it? | 1. অগভীর টিউবওয়েল Shallow tubewell<br>2. গভীর টিউবওয়েল Deep tubewell<br>3. ঘরের ভিতর ট্যাপ বা পাইপের পানি Piped into home<br>4. উঠানে ট্যাপ বা পাইপের পানি Piped into yard<br>5. গভীর পাতকূয়া (উঠানে)Borewell in yard<br>6. পাবলিক ট্যাপ Public tap<br>7. পাবলিক গভীর পাতকূয়া Public borewell<br>8. অগভীর পাকা পাতকূয়া Shallow well with concrete reinforcement<br>9. অগভীর কাঁচা পাতকূয়া Shallow well (no concrete) |

| | | | |
|---|---|---|---|
| | | | 10. নদী/বাধ/লেক/পুকুর/সেচ নালা থেকে সরাসরি সংগৃহীত পানি River, lake, stream<br>11. অন্যান্য (লিখুন) Other: ____________<br>88] ২য় কোন পানির উৎস ব্যবহার করতে হয় না Don't use any secondary water source (Skip to 1017.a) |
| C1021 | 1015. | আপনি আমাকে বলবেন আপনার বাড়ী থেকে ২য় পানির উৎসে একবার যেতে কতক্ষণ সময় লাগে? Can you tell me how long it takes you to walk one-way to this SECONDARY water source from your home? | \|__\|ঘন্টা Hours<br>\|__\|মিনিট Min |
| C1022 | 1016. | আপনি আমাকে বলবেন কোন কোন কাজের জন্য এই উৎসের পানি ব্যবহার করেন Can you please tell me whether you use water from this source for… | |
| 1 | 1 | পানি পান Drinking | [1] হ্যাঁ Yes [0] না No |
| 2 | 2 | রান্না-বান্না/বাসন ধোয়া Cooking/ washing dishes | 1] হ্যাঁ Yes [0] না No |
| 3 | 3 | কাপড় ধোয়া Laundry | 1] হ্যাঁ Yes [0] না No |
| 4 | 4 | গোসল/হাতধোয়া Bathing/hand washing | 1] হ্যাঁ Yes [0] না No |
| 5 | 5 | কৃষিকাজ/বাগান Irrigate crop land or garden | 1] হ্যাঁ Yes [0] না No |
| 6 | 6 | পশু-পালন Water for livestock | 1] হ্যাঁ Yes [0] না No |
| C1023 | 1017. | আপনার খানাকে এই পানির উৎস ব্যবহারের জন্য কি কোন প্রকার টাকা দিতে হয়? your household pay any money to use this source? | 1] হ্যাঁ Yes [0] না No |
| | 1017.a | আপনার খানার খাবার পানির প্রধান উৎস কি? What is your main drinking water source? | 1. অগভীর টিউবওয়েল Shallow tubewell<br>2. গভীর টিউবওয়েল Deep tubewell<br>3. ঘরের ভিতর ট্যাপ বা পাইপের পানি Piped into home<br>4. উঠানে ট্যাপ বা পাইপের পানি Piped into yard<br>5. গভীর পাতকূয়া (উঠানে)Borewell in yard<br>6. পাবলিক ট্যাপ Public tap<br>7. পাবলিক গভীর পাতকূয়া Public borewell<br>8. অগভীর পাকা পাতকূয়া Shallow well with concrete reinforcement<br>9. অগভীর কাঁচা পাতকূয়া Shallow well (no concrete)<br>10. নদী/বাধ/লেক/পুকুর/সেচ নালা থেকে সরাসরি সংগৃহীত পানি River, lake, stream<br>11. অন্যান্য (লিখুন) Other: ____________<br>88] Not applicable |

| | | | |
|---|---|---|---|
| C1024 | 1018. | আপনার খানা কি কখনও বৃষ্টির পানি সংগ্রহ করত ? Does your household collect rainwater? | 1] হ্যাঁ Yes [0] না No |
| C1025 | 1019. | আপনার খানা কি বর্তমানে বৃষ্টির পানি সংগ্রহ করে ? Is your household currently collecting rainwater? | 1] হ্যাঁ Yes [0] না No if no, skip to 1301 |
| C1026 | 1020. | আপনি আমাকে বলবেন কি আপনার খানা কি কি কাজে বৃষ্টির পানি ব্যবহার করে ? Can you please tell me whether you use rainwater for | |
| 1 | 1 | পানি পান Drinking | [1] হ্যাঁ Yes [0] না No |
| 2 | 2 | রান্না-বান্না/বাসন ধোয়া Cooking/ washing dishes | 1] হ্যাঁ Yes [0] না No |
| 3 | 3 | কাপড় ধোয়া Laundry | 1] হ্যাঁ Yes [0] না No |
| 4 | 4 | গোসল/হাতধোয়া Bathing/hand washing | 1] হ্যাঁ Yes [0] না No |
| 5 | 5 | কৃষিকাজ/বাগান Irrigate crop land or garden | 1] হ্যাঁ Yes [0] না No |
| 6 | 6 | পশু-পালন Water for livestock | 1] হ্যাঁ Yes [0] না No |
| **Skip to 1301 (Module 13 MEASURES FOR SPILLOVER)** | | | |

| | | | |
|---|---|---|---|
| ~~C.1027.~~ | ~~1021.~~ | | |
| **C.1010.** | 1022. | আপনার খানায় কত ঘন ঘন খাবার পানিতে আকুয়াট্যাবস মেশানো হয়? How frequently do you treat your water using [Aquatabs]? | Do Not Read Responses.<br>1 যতবার পানি সংগ্রহ করা হয়/সবসময় Every time they collect water<br>2 মাঝেমাঝে Sometimes / occasionally<br>3 প্রথমদিকে দিয়েছিলাম কিন্তু এখন আর দেই না Treated water in the beginning [of the program] but not any more<br>4 আকুয়াট্যাবস দিয়ে কখনও পানি পরিশোধন করি নাই Never treated water with [Aquatabs] **(Skip to 1026)**<br>99 জানি না Don't know / not sure |
| **C.1011** | 1023. | আপনার খানায় সংরক্ষনকৃত খাবার পানি কি কুয়াট্যাবস দ্বারা পরিশোধন করা হয়েছে? Is the drinking water stored in your household today treated with [Aquatabs]? | 1 হ্যাঁ, সবটুকু পানি Yes, all of it [TOPAZ]<br>2 হ্যাঁ, কিছু পানি Yes, some of it<br>3 না Not treated<br>4 খানায় কোন পানি নাই No water in the house[skip to 1026]<br>99 জানি না Don't know |
| **C.1012** | 1024. | পর্যবেক্ষণ করুন: আকুয়াট্যাবস দ্বারা পরিশোধিত | 1 হ্যাঁ Yes |

| | | | |
|---|---|---|---|
| | | পানির পাত্র দেখাতে বলুন: পানির পাত্র ঢাকা ছিল কী?<br>Ask to see the treated water. Observe: Is the water covered? | 2 না No<br>3 খানায় পাত্রে কোন পানি নাই No water stored in the house<br>88 প্রযোজ্য নয়/প্রত্যাখ্যান Not applicable / refused |
| C.1013 | 1025. | পানি পরিশোধন করার জন্য কত সময় আগে আকুয়াট্যাবস ব্যবহার করা হয়েছিল?<br>Approximately how long ago did you treat the water with [Aquatabs]? | ঘিন্টা HH: মিনিট MM ☐☐:☐☐ ( 9 9 : 9 9 Don't know ) |
| ক্লোরিন অবশেষ পরীক্ষা **Residual Chlorine Test**<br>সকল খানার জন্য প্রযোজ্য Administer to: All study households<br>*Ask to collect a water sample from the source identified in C.1003. Explain that some (but not all) of the samples may be tested to see if there is any chlorine in the water.* | | | |
| | 1026. | ক্লোরিণের জন্য আমি আপনার খাবার পানি পরীৰা করতে পারি কী?<br>May I collect a small sample of your drinking water? | 1 হ্যাঁ Yes<br>2 না/প্রত্যাখান No / refused<br>3 খানায় কোন পানি নাই No drinking water available to test |
| | 1027. | ক্লোরিন অবশেষ পরীক্ষার জন্য সংরক্ষনকৃত পানি যা নির্দ্দিষ্ট বাচ্চা খায় তার নমুণা সংগ্রহ করম্নন। ভিন্ন ভিন্ন অবস্থানেই খানার সংরক্ষিত খাবার পানির ক্লারিন অবশেষ পরীক্ষা করুন । শুধুমাত্র সেই খানার পানি পরীক্ষা করতে হবে যারা বলবে সংরক্ষিত খাবার পানিতে ক্লারিন মিশিনো হয়েছে(C.1009 = 1, 2, or 11)। যদি এই খানায় সংরক্ষিত খাবার পানিতে ক্লারিন না মেশানো হয়, তাহলে পানি পরীক্ষা থেকে বিরত থাকুন এবং ৮৮ কোড করুন।<br>Collect a small water sample from stored drinking water for target children.<br>Test for free residual chlorinein a discrete location after you leave the household.<br>Only test samples collect from households who report to have treated their water with some form of chlorine (C.1009 = 1, 2, or 11).If the household did not report treating the water with chlorine, discard the sample without testing it and record | ৮৮৮ ক্লোরিন মেশানো হয়েছে কিনা এ বিষয়ে রিপোর্ট দিতে পারে নি<br>8 88 Did not report that the water was treated with chlorine<br>**999** পরীক্ষা করা হয় নাই<br>9 9 9 Could not test<br>ক্লোরিন অবশেষের পরিমান Level of Free Residual Chlorine: ☐☐.☐ মি.গ্রাম/লিটার mg / L |

| | | |
|---|---|---|
| | | 8 88 in the field below. |
| Now complete 1007 to 1020 if aquatabs is used | | |

#### Wash Benefit Module 13 MEASURES FOR SPILLOVER

সকল খানার জন্য প্রযোজ্য Administer to: All study households

| | | | |
|---|---|---|---|
| C.1301. | 1301. | আপনার সবচেয়ে কাছের হেলথ ফ্যাসিলিটি (স্বাস্থ্যসেবা পাওয়ার সুযোগ)-এর নাম কি? যদি না জানে/নিশ্চিত না হয় তবে ৯৯ বসান। খানার কেউ ডাক্তার হলে বা ডাক্তার খানায় এসে চিকিৎসা করলে ৫৫ লিখুন। What is the name of your nearest health facility?<br>Enter "99" if Don't know / not sure | |
| C.1301.a | 1302. | গত ১ বছরে আপনি/আপনার শিশু মোটামুটি কতবার যে কোন অসুখে এই সুযোগ গ্রহন করেছিলেন? In the past year, (since this time last year) approximately how many times did you visit this health facility for any reason (including with your children)? | \|__\|__\| বার TIMES |
| C.1301.b | 1303. | আপনি সেখানে কিভাবে যান? What is your primary mode of travel to the health facility? | ১. পায়ে হেঁটে<br>2. রিকশা/ভ্যান/নৌকা/ সাইকেল<br>3. ব্যাটারি চালিত অটোরিকশা<br>4. ট্যাম্পু/লেগুনা/ সিএনজি/ বাস/মোটরসাইকেল<br>5. প্রযোজ্য নয় |
| C.1301.c | 1303.a | আনুমানিক সেখানে যেতে কতক্ষণ সময় লাগে(আপনার খানা থেকে শুধু যেতে কত মিনিট সময় লাগে) ? যদি না জানে/নিশ্চিত না হয় তবে ৯৯ বসান। Approximately how long does it take to travel there (in minutes, one-way, from where we are now)?<br><br>Enter "999" if Don't know / not sure | \|__\|__\|__\| মিনিটে MINUTES |
| C.1302. | 1304. | আপনার কাছের প্রাক- প্রাইমারী স্কুলের -এর নাম কি ? যেখানে আপনার শিশু যেতে পারে/বড় হলেও যেতে পারবে । যদি না জানে/নিশ্চিত না হয় তবে ৯৯, যদি না থাকে তবে ৮৮ বসান ।What is the name of the ECD center [preschool] where your children go (or will go when they are old enough)? | |
| C.1303. | 1305. | আপনার কাছের প্রাইমারী স্কুলের -এর নাম কি ? যেখানে আপনার শিশু যেতে পারে/বড় হলেও যেতে পারবে । What is the name of the primary school where your children go (or will go when they are old enough)? | |
| C.1304. | 1306. | আপনার কাছের বাজার -এর নাম কি যেখানে আপনি সবসময়ই কেনা-বেচা করতে যান? । What is the name of the market that you most often go to when you buy or sell things? | |
| C.1304.a | 1307. | কতবার আপনি সেখানে যান? How often do you go to that market? | 0 কখনই না Never<br>1 সপ্তাহে একবার অথবা তার অধিক Once per week or more<br>2 প্রতি ২ সপ্তাহে একবার Once every 2 weeks<br>3 প্রতি ৩ সপ্তাহে একবার Once every 3 weeks<br>4 প্রতি ৪ সপ্তাহে একবার(প্রতি মাসে) Once every 4 weeks (every month)<br>5 প্রতি মাসে একবারের কম Less than once every month<br>99 জানি না/নিশ্চিত না Don't know / not sure |

| | | | |
|---|---|---|---|
| | 1308. | আপনি সেখানে কিভাবে যান? What is your primary mode of travel to the health facility? | 1. পায়ে হেঁটে<br>2. রিকশা/ভ্যান/নৌকা / সাইকেল<br>3. ব্যাটারি চালিত অটোরিকশা<br>4. ট্যাম্পু/লেগুনা/ সিএনজি/ বাস/মোটরসাইকেল<br>5. প্রযোজ্য নয় |
| C.1304.b | 1308 .a | আনুমানিক সেখানে যেতে কতক্ষণ সময় লাগে(আপনার খানা থেকে শুধু যেতে কত মিনিট সময় লাগে) ? যদি না জানে/নিশ্চিত না হয় তবে ৯৯ বসান। Approximately how long does it take to travel there (in minutes, one-way, from where we are now)?<br><br>Enter "999" if Don't know / not sure | \|__\|__\|__\| মিনিটে MINUTES |
| C.1305. | 1309. | আপনার কাছের মসজিদ/মন্দির/গীর্জা/প্যাগোডা -এর নাম কি? যেখানে আপনি প্রার্থনা করতে যান । What is the name of the church/mosque/temple (if any) that you attend? | |
| C.1305.a | 1310. | কতবার আপনি সেখানে যান? How often do you go to church? | ০. কখনই না Never<br>1 সপ্তাহে একবার অথবা তার অধিক Once per week or more<br>2 প্রতি ২ সপ্তাহে একবার Once every 2 weeks<br>3 প্রতি ৩ সপ্তাহে একবার Once every 3 weeks<br>4 প্রতি ৪ সপ্তাহে একবার(প্রতি মাসে) Once every 4 weeks (every month)<br>5 প্রতি মাসে একবারের কম Less than once every month<br>99 জানি না/নিশ্চিত না Don't know / not sure |
| | 1311. | আপনি সেখানে কিভাবে যান? What is your primary mode of travel to the health facility? | 1. পায়ে হেঁটে<br>2. রিকশা/ভ্যান/নৌকা<br>3. ব্যাটারি চালিত অটোরিকশা<br>4. ট্যাম্পু/লেগুনা/ সিএনজি/ বাস<br>5. প্রযোজ্য নয় |
| C.1305.b | 1312 .a | আনুমানিক সেখানে যেতে কতক্ষণ সময় লাগে(আপনার খানা থেকে শুধু যেতে কত মিনিট সময় লাগে) ? যদি না জানে/নিশ্চিত না হয় তবে ৯৯ বসান। Approximately how long does it take to travel there (in minutes, one-way, from where we are now)?<br><br>Enter "999" if Don't know / not sure | \|__\|__\|__\| মিনিটে MINUTES |
| C.1306. | 1312. | | |

#### Wash Benefit Module 14 LNS UPTAKE

| Adherence to Sonamoni | | | |
|---|---|---|---|
| **N.1401** | Is this a nutrition intervention arm?<br>এঁটা কি একটা পুষ্টি ইনটারভেনশন আরম ? | 1. Yes হ্যাঁ<br>0. No না | If No, skip to 1801 |

| | | | |
|---|---|---|---|
| **N.1402** | Did they receive LNS yet?<br>আপনি কি LNS পেয়েছেন? | 1. Yes হ্যাঁ<br>2. No,child is below 6 months<br>না, শিশুর বয়স ৬ মাসের কম<br>3. No, as they were from home for last one month<br>না, গত ১ মাস ধরে শিশুটি বাড়ীর বাইরে<br>777. Others, specify_________<br>অন্যান্য উলেস্নখ করম্নন -------- | Stop the intervie w here. |
| **N.1403** | If yes, please show me the items you received?<br>Record your observation.<br>DO NOT READ THE RESPONSES. MULTIPLE RESPONSES ALLOWED.<br>যদি হ্যাঁ হয়,তাহলে আপনার বাচ্চার জন্য কোন কোন উপকরণ পেয়েছেন তা আমাকে দেখান ?আপনার পর্যবেক্ষণ লিপিবদ্ধ করম্নন ।একাধিক উত্তর আসতে পারে, উত্তর পড়ে শুনানো যাবে না | Items (Yes=1 হ্যাঁ , No=0 না)<br>1. Sonamoni<br>সোনামনি<br>2. Plastic container for Sonamoni<br>সোনামনি রাখার জন্য পস্নাস্টিকের পাত্র | |
| **N.1404** | How many days ago were the Sonamoni sachets distributed?<br>(FRAs ask household and see the records )<br>কত দিন আগে সোনামনি স্যাশেট গুলো সরবরাহ করা হয়েছে? প্রশ্ন করম্নন এবং রেকর্ড দেখুন । | I__II__I days দিন | |
| **N.1405** | How many unused Sonamoni sachets you had at the time of last distribution?<br>(FRAs calculate and put the number)<br>শেষ বিতরনের দিন আপনার কাছে কতগুলো অব্যবহৃত এল এন এস স্যাশেট ছিল? গনণা করম্নন এবং রেকর্ড করম্নন। | I__II__I Sachets স্যাশেট<br>Not sure 99 | |
| **N.1406** | How many Sonamoni sachets did you receive during last distribution? (FRAs ask household & see the records)<br>আপনি কতগুলো সোনামনি স্যাশেট শেষ বিতরনের দিন পেয়েছিলেন? প্রশ্ন করম্নন এবং রেকর্ড দেখুন । | I__I__I Sachets স্যাশেট<br>Not sure 99 | |
| **N.1407** | How many Sonamoni sachets are unused at present?<br>(FRAs count and put the number)<br>এখন ও কতগুলো অব্যবহৃত স্যাশেট আছে? গননা করম্নন এবং নম্বরটি লিখুন। | I__I__I Sachets স্যাশেট | |
| N.1407a | Did the child ever ate Sonamoni?<br>শিশুটি কি কখনো সোনামণি খেয়েছে ? | 1. Yes হ্যাঁ<br>0. No না | If 0 then skip to 1414 |
| **N.1408** | During the past week, how many days did you feed Sonamoni to your child?"<br>গত সপ্তাহে কতদিন আপনি শিশুটিকে সোনামণি খাইয়েছেন? | \|__\| days দিন | If 0 then skip to 1414 |
| **N.1409** | How much supplement did you feed most of the days during the last seven days?<br>~~DO NOT READ THE RESPONSES. SINGLE RESPONSE ALLOWED~~<br>গত সপ্তাহের বেশীরভাগ দিন ~~গড়ে~~ কয়টা করে সোনামণি আপনি শিশুটিকে ~~১ দিনে~~ খাইয়েছেন? | \|___\|. \|__ \| sachets স্যাশেট | |

| | | | |
|---|---|---|---|
| **N.1410** | If yes, how did you primarily feed the Sonamoni to your Child during last seven days?<br>DO NOT READ THE RESPONSES. SINGLE RESPONSES ALLOWED.<br>যদি হ্যাঁ হয়, তাহলে গত সপ্তাহের বেশীরভাগ সময় আপনি কিভাবে আপনার শিশুকে সোনামনি খাইয়েছিলেন ? উত্তর পড়ে শুনানো যাবে না। | Supplementation (Yes=1 হ্যাঁ , No=0 না)<br>1. Eaten alone<br>স্যাশেট থেকে সরাসরি নিয়ে খাওয়ানো হয়েছিল<br>2. Mixed with rice<br>ভাতের সাথে মিশিয়ে খাওয়ানো হয়েছিল<br>3. Mixed with other food<br>অন্যান্য খাবারের সাথে মিশিয়ে খাওয়ানো হয়েছিল | |
| **N.1411** | What did you use to feed the child last time you fed Sonamoni?<br>শেষবার আপনি কি দিয়ে শিশুটিকে সোনামণি খাইয়েছেন? | 1. Spoon চামচ<br>2. Mother's fingers<br>মায়ের আঙুল<br>3. Directly from sachet<br>স্যাশেট থেকে সরাসরি নিয়ে<br>4. Child fed him/herself by hand<br>শিশুটি নিজের হাত দিয়ে খেয়েছে | If 2 or 4 then ask N.1412 otherwise skip to N.1414 |
| **N.1412** | Did you wash your hands last time you fed Sonamoni to your child?<br>Or<br>Did your child washed hands last time s/he ate Sonamoni by hands?<br>শেষবার যখন শিশুকে সোনামণি খাইয়েছেন তখন কি আপনি হাত ধুয়েছিলেন অথবা শেষবার যখন শিশুটি নিজ হাতে সোনামণি খেয়েছে তখন কি হাত ধুয়েছে? | 1. Yes হ্যাঁ<br>0. No না | If no, skip to N.1414 |
| **N.1413** | If yes, what did you use to wash your hands last time you fed Sonamoni to your child?<br>DO NOT READ THE RESPONSES. MULTIPLE RESPONSES ALLOWED.<br>যদি উত্তর হ্যাঁ হয়, তাহলে আপনি/ আপনার শিশু কি দিয়ে হাত ধুয়েছিলেন ? একাধিক উত্তর আসতে পারে, উত্তর পড়ে শুনানো যাবে না । | Used (Yes=1 হ্যাঁ , No=0 না)<br>1. Bar soap বার সাবান<br>২. Detergent গুড়া পাউডার<br>৩. Liquid soap তরল সাবান<br>৪. Soapy water সাবান-পানি<br>৫. Ash ছাই<br>৬. Mud মাটি<br>৭. Only water শুধু পানি | |
| **N.1414** | Did anybody else ever share Sonamoni with the target child?<br>নির্দিষ্ট শিশুর সাথে অন্যকেহ সোনামনি খেয়েছিল কি? | 1. Yes হ্যাঁ<br>0. No না | If no, skip to 1416 |
| **N.1415** | If yes, who?<br>DO NOT READ THE RESPONSES. MULTIPLE RESPONSES ALLOWED.<br>যদি উত্তর হ্যাঁ হয়, তাহলে কে কে সোনামনি খেয়েছিল? একাধিক উত্তর আসতে পারে, উত্তর পড়ে শুনানো যাবে না | Relationship (Yes=1 হ্যাঁ , No=0 না)<br>1. Sibling<br>একই মায়ের অন্য শিশু<br>2. Other child/children<br>অন্য শিশু/রা<br>3. Adult relative(s)<br>বয়স্ক আত্মীয়<br>4. Other adult(s)<br>অন্যান্য বয়স্ক | |

| | | |
|---|---|---|
| **N.1416** | If no to N.1407a or N.1408 then why Sonamoni was not eaten?<br>DO NOT READ THE RESPONSES. MULTIPLE RESPONSES ALLOWED.<br>যদি N.1407a অথবা N.1408এর উত্তর না হয়, তাহলে কেন সোনামনি খায়নি? একাধিক উত্তর আসতে পারে, উত্তর পড়ে শুনানো যাবে না | Reasons (Yes=1 হ্যাঁ , No=0 না)<br>1. Child didn't like শিশু পছন্দ করে না<br>2. Not sure about the product দ্রব্যটি সম্মন্ধে জানি না<br>3. Peer pressure not to take অন্যরা খেতে নিষেধ করেছে<br>4. Child was sick শিশুটি অসুস্থ ছিল<br>5. Away from home শিশুটি বাড়ীতে ছিল না<br>6. Gave supplement to another person অন্যকে এই সম্পূরক খাবার দিয়ে দেওয়া হয়েছে<br>777. Others (Specify) অন্যান্য উলেস্নখ করুনন -------- |
| **N.1417** | Were any of the Sonamoni sachets damaged or opened prior or during distribution?<br>সোনামনি স্যাশেট গুলো সরবরাহের সময় কি নষ্ট অথবা খোলা ছিল? | 1. Yes হ্যাঁ<br>0. No না |
| **N.1418** | What shortcomings or problems are there in your opinion with Sonamoni?<br><br>DO NOT READ THE RESPONSES. MULTIPLE RESPONSES ALLOWED.<br>আপনার মতে কোন ত্রুটিবিচ্যুতি আছে কি? উত্তর পড়ে শোনানো যাবে না। একাধিক উত্তর হতে পারে। | Shortcoming/problems (Yes=1 হ্যাঁ , No=0 না)<br>1. Irregular supply অনিয়মিত সরবরাহ<br>2. Taste is not good স্বাদ ভাল না<br>3. Child vomit after swallow খাওয়ার পরে বাচ্চা বমি করে<br>4. Color of the food changes when mixed with Sonamoni খাবারের সাথে মিশালে রং পরিবর্তিত হয়<br>5. Limited supply পরিমিত সরবরাহ<br>6. Bad smell পঁচা গন্ধ<br>7. Causes indigestion হজম হয় না<br>8. Loose stool পাতলা পায়খানা<br>9. Allergic Reaction (skin rash) এলারজিক প্রতিক্রিয়া<br>10. Child doesn't like to eat Sonamoni শিশু **সোনামনি** খেতে পছন্দ করে না<br>11. Difficult to take out of the sachet স্যাশেট থেকে বের করা কষ্টসাধ্য<br>12. No Problem কোন সমস্যা নাই<br>13. Increased appetite ক্ষুধা বৃদ্ধি<br>14. Child won't eat other food because they are eating Sonamoni বাচ্চারা অন্যান্য খাবার খায় না কারণ |

| | | |
|---|---|---|
|  |  | তারা সোনামণি খাচ্ছে<br>77. Others, ( Specify )<br>অন্যান্য উলেস্নখ করম্নন -------- |
| **N.1419** | What is that you liked about the supplement? DO NOT READ THE RESPONSES. MULTIPLE RESPONSES ALLOWED.<br>আপনি কি কি কারণে এই সম্পূরক খাবার পছন্দ করেন? উত্তর পড়ে শোনানো যাবে না। একাধিক উত্তর হতে পারে। | Benefits (Yes=1 হ্যাঁ , No=0 না)<br>1. Increased appetite<br>ক্ষুধা বৃদ্ধি<br>2. Weight gain<br>ওজন বাড়ে<br>3. Increased energy<br>শক্তি বৃদ্ধি<br>4. Easy to feed<br>খাওয়ানো সহজ<br>5. Child liked it<br>শিশু এটি পছন্দ করে<br>6. Child remains playful<br>শিশু খেলাধূলা করে<br>7. Good taste<br>ভাল স্বাদ<br>8. Good smell<br>ভাল গন্ধ<br>9. Don't have to feed other foods because Sonamoni meets the child's needs<br>অন্যান্য খাবার খাওয়ানোর প্রয়োজন হয় না কারণ সোনামণি বাচ্চার চাহিদা পূরণ করে<br>777. Other, Specify ___________<br>অন্যান্য উলেস্নখ করম্নন -------- |

#### Wash Benefit Module 15: Environmental Sampling (Water, Hand, Sentinel Toy and Fly Density)

#### DAY 1 SURVEY

##### SECTION 1. IDENTIFICATION AND DEMOGRAPHICS

1.1 Enter the FRAemployee ID(এফআর এ এমপস্নয়ী আইডি নম্বর লিখুন)

1.2Please enter the current day[আজকের তারিখ (সংখ্যায়) লিখুন]

1.3Please enter the current month[বর্তমান মাসের নাম (সংখ্যায়) লিখুন]

1.4Enter five-digit participant ID
[অংশগ্রহনকারীর আইডি নম্বর লিখুন (৫ সংখ্যার আইডি ওয়াশ বেনিফিট স্টাডি প্রদত্ত তালিকা থেকে)]

1.5 Name of respondent (target child's mother)
[প্রধান উত্তরদাতার (টার্গেট শিশুর মা) নাম লিখুন]ঃ

1.6 Name of target child
[টার্গেট শিশুর নাম লিখুন]ঃ

##### SECTION 2. DELIVER TOY BALL AND FLY TAPE

PROMPT: Please deliver the toy ball to the target child or respondent.

[অনুগ্রহপূর্বক টার্গেট শিশু বা পরিচর্যাকারীকে খেলনা বল বিতরন করুন]

2.1 Who did you give the ball to?
[খেলনা বলটি আপনি কাকে দিয়েছেন?]

1 = Target child(টার্গেট শিশু)
2 = Respondent(পরিচর্যাকারী)
3 = Other caregiver(অন্যান্য পরিচর্যাকারী)
4 = Ball not delivered(খেলনা বল দেওয়া হয়নি)
77 = Other (specify)[অন্যান্য (নির্দিষ্ট করে লিখুন)]

2.2 (if question 2.1 is not 4) Record the time that the toy ball was delivered (24H format, HH:MM).
[(যদি 2.1নং প্রশ্নের উত্তর 4 না হয়) খেলনা বলটি যে সময়ে বিতরন করা হয়েছে, সেই সময়টা লিপিবদ্ধ করুন। (24ঘন্টা হিসাবে, ঘন্টাঃ মিনিট)]

2.3 (ifquestion2.1 is 4) Why was the toy ball not delivered?
[(যদি 2.1নং প্রশ্নের উত্তর 4 হয়) কেন খেলনা বল বিতরন করা হয়নি?]

1 = Respondent/caregiver refused(উত্তরদাতা/ পরিচর্যাকারী প্রত্যাখ্যান করেছে)
2 = Did not have a toy ball to deliver(বিতরন করার জন্য খেলনা বল ছিল না)
77 = Other (specify)[অন্যান্য (নির্দিষ্ট করে লিখুন)]

PROMPT: Can you please show me the area where you prepare food?
(অনুগ্রহপূর্বক আপনাদের খাবার তৈরির স্থান/ রান্নাঘরটি আমাকে দেখাবেন কি?)

PROMPT: Hang fly tape as close as possible to the food preparation area. If there is more than one food preparation area, hang the strips near where food was most recently prepared. Do NOT hang the strips over or near to a cooking fire. Ask the respondents to leave the tape undisturbed.

(মাছি ধরার ফাঁদ যতটা সম্ভব খাবার তৈরির স্থান/ রান্নাঘরের কাছাকাছি ঝোলাতে হবে। যদি কোন খানায় একাধিক খাবার তৈরির স্থান/ রান্নাঘর থাকে, তাহলে সর্বশেষ যে স্থান/ রান্নাঘরে খাবার তৈরি করা হয়েছে সেখানে মাছি ধরার ফাঁদ ঝোলাতে হবে। রান্নাঘরে আগুনের কাছাকাছি বা উপরে মাছি ধরার ফাঁদ ঝোলাবেন না। মাছি ধরার ফাঁদ ঝোলানোর পরে উত্তরদাতাকে ইহা নড়াচড়া না করে স্থিরভাবে যথাস্থানে রাখতে অনুরোধ করুন।)

2.4 Record the time that the fly tapewas hung at the food preperation area (24H format, HH:MM)
[খাবার তৈরির স্থানে মাছি ধরার ফাঁদ ঝোলানোর সময় লিপিবদ্ধ করুন (24 ঘন্টা হিসাবে, ঘন্টা: মিনিট)]

PROMPT: Can you please show me your primary latrinearea?
(অনুগ্রহপূর্বক আপনাদের প্রাথমিকভাবে ব্যবহৃত টয়লেটের স্থানটি/টয়লেটটি আমাকে দেখাবেন কি?)

PROMPT: Hang fly tape as close as possible to the primary latrine area. If there is more than one primary latrine area, hang the strips near the latrine that the respondent used most recently.Ask the respondents to leave the tape undisturbed.

(মাছি ধরার ফাঁদ যতটা সম্ভব প্রাথমিকভাবে ব্যবহৃত টয়লেটের কাছাকাছি ঝোলাতে হবে। যদি কোন খানায় একাধিক টয়লেট থাকে, তাহলে টার্গেট শিশুর মাসর্বশেষ যে টয়লেটটি ব্যবহারকরেছেসেখানে মাছি ধরার ফাঁদ ঝোলাতে হবে। মাছি ধরার ফাঁদ ঝোলানোর পরে উত্তরদাতাকে ইহা নড়াচড়া না করে স্থিরভাবে যথাস্থানে রাখতে অনুরোধ করুন।)

2.5 Record the time that the fly tapewas hung at the primary latrine area (24H format, HH:MM)
[টয়লেটের আশেপাশেমাছি ধরার ফাঁদ ঝোলানোর সময় লিপিবদ্ধ করুন। (24 ঘন্টা হিসাবে, ঘন্টা: মিনিট)]

**SECTION 3. HAND RINSE SAMPLING**

| | |
|---|---|
| 3.1 | Record whether the respondent has washed her or target child's hands at any time before this question since you arrived at the household Select all that apply.<br><br>(উক্ত খানায় আসার পর থেকে এই প্রশ্নটি করার আগেপর্যন্ত যেকোন সময়ে আপনি উত্তরদাতাকে বা টার্গেট শিশুকে হাত ধুতে দেখেছেন কি? )(নিচে প্রদত্ত অপশনগুলো থেকে যতগুলো প্রযোজ্য তা নির্বাচন করুন) |

| | |
|---|---|
| | [1] Observed respondent washing hands<br>(উত্তরদাতাকে হাত ধুতে দেখা গেছে)<br>[2] Did not observe handwashing directly but respondent had wet hands<br>(উত্তরদাতাকে সরাসরি হাত ধুতে দেখা যায়নি তবে তার হাত ভেজা দেখা গেছে)<br>[3] Did not observe respondent washing hands or with wet hands<br>(উত্তরদাতাকে হাত ধুতে দেখা যায়নি অথবা হাত ভেজাও দেখা যায়নি)<br>[4] Observed respondent washing/wiping target child's hands<br>(উত্তরদাতাকে টার্গেট শিশুর হাত ধুয়ে/ মুছে দিতে দেখা গেছে)<br>[5] Did not observe handwashing directly but target child had wet hands<br>(উত্তরদাতাকে সরাসরি টার্গেট শিশুর হাত ধুয়ে দিতে দেখা যায়নি কিন্তু টার্গেট শিশুর হাত ভেজা দেখা গেছে)<br>[6] Did not observe respondent washing/wiping target child's hands or child with wet hands(উত্তরদাতাকে টার্গেট শিশুর হাত ধুয়ে/মুছে দিতে দেখা যায়নি অথবা টার্গেট শিশুর হাতও ভেজা দেখা যায়নি)<br>[99] Could not observe<br>(পর্যবেৰণ করা সম্ভব হয়নি) |

<table>
<tr><td rowspan="4">3.2</td><td rowspan="4">READ: Thank you. Now, I would like to do a quick inspection of your hands. I hope you don't mind. Can you please show me your hands?<br><br>উত্তরদাতাকে পড়ে শোনানঃ আপনাকে ধন্যবাদ, এখন আমি আপনার হাতগুলো একটু দেখতে চাচ্ছি। আমি আশা করছি এতে আপনি কিছু মনে করবেন না। আপনি অনুগ্রহপূর্বক আমাকে আপনার হাতগুলো দেখাবেন কি?<br><br>BOTH HANDS SHOULD BE SHOWN (NOT JUST ONE HAND). USE APPEARANCE CODES BELOW TO RECORD THE DESCRIPTION THAT BEST DESCRIBES THE LEVEL OF CLEANLINESS.<br><br>দুইটি হাতই দেখতে হবে (যে কোন একটি দেখালে চলবে না)। এরপর নিচের উলেস্নখিত হাতের অবস্থা কোড দেখে রেকর্ড করতে হবে।<br><br>ASK: Please show me [TARGET CHILD NAME]'s hands.DO NOT TOUCH HANDS.<br><br>প্রশ্ন করুননঃ দয়া করে আমাকে [শিশুর নাম ধরে] হাতগুলো দেখান। দয়া করে বাচ্চার হাত স্পর্শ করবেন না।</td><td colspan="2">MOTHER/CAREGIVER(মা/পরিচর্যাকারী)</td></tr>
<tr><td>Left Hand<br>(বাম হাত)<br><br>A|__|FINGERNAILS<br>(হাতের নখ)<br><br>B|__| PALMS<br>(হাতের তালু)<br><br>C|__|FINGER PADS<br>(আঙ্গুলের সম্মুখভাগ)</td><td>Right Hand<br>(ডান হাত)<br><br>D|__|FINGERNAILS<br>(হাতের নখ)<br><br>E|__| PALMS<br>(হাতের তালু)<br><br>F|__|FINGER PADS<br>(আঙ্গুলের সম্মুখভাগ)</td></tr>
<tr><td colspan="2">TARGET CHILD(টার্গেট শিশু)</td></tr>
<tr><td>Left Hand<br>(বাম হাত)<br><br>G|__|FINGERNAILS<br>(হাতের নখ)<br><br>H|__| PALMS<br>(হাতের তালু)<br><br>I |__|FINGER PADS<br>(আঙ্গুলের সম্মুখভাগ)</td><td>Right Hand<br>(ডান হাত)<br><br>J |__|FINGERNAILS<br>(হাতের নখ)<br><br>K |__| PALMS<br>(হাতের তালু)<br><br>L |__|FINGER PADS<br>(আঙ্গুলের সম্মুখভাগ)</td></tr>
</table>

APPEARANCE CODES:

1 = VISIBLE DIRT (DIRT/MUD/SOIL/ASH OR ANY OTHER MATERIAL IS VISIBLE)
ময়লা স্পষ্টভাবে দেখা যাচ্ছিল (ময়লা/ কাঁদা/ মাটি/ছাই অথবাঅন্য কোন উপাদানু)

2 = UNCLEAN APPEARANCE (NO DIRT IS VISIBLE ON THIS PART OF THE HAND BUT, IN GENERAL, THIS PART OF THE HAND APPEARS UNCLEAN)
ময়লাস্পষ্টভাবে দেখা না গেলেও অপরিচ্ছন্নভাব ছিল (হাতে ময়লা না দেখা গেলেও অপরিচ্ছন্ন ছিল)

3 = CLEAN (OBSERVED PART OF THE HAND IS CLEAN AS WOULD APPEAR AFTER SOMEONE WASHES HANDS OR TAKES A BATH)

পরিষ্কার ছিল (হাত ধোয়ার পর/ গোসলের পর পর হাত দেখলে যেমনপরিষ্কার হয়)

88 = N/A(প্রযোজ্য নয়)

99 = OBSERVATION NOT POSSIBLE/REFUSED/TARGET CHILD NOT AVAILABLE
(পর্যবেৰণ করা সম্ভব হয়নি/ প্রত্যাখ্যান/ নির্দিষ্ট শিশু উপস্থিত ছিল না)

PROMPT: Collect a hand rinse sample from the target child by rinsing both hands in same Whirlpak bag.
দয়া করে নির্দিষ্ট শিশুর উভয় হাতধোয়া পানির নমুনা একই ব্যাগে সংগ্রহ করম্নন।

3.3 Please enter the 5-digit unique numerical ID.
যে ব্যাগে নমুনা সংগ্রহ করা হবে অনুগ্রহপূর্বক তার গায়ে লেখা5সংখ্যার ইউনিক আইডি নম্বরটি এন্ট্রি করম্ননঃ

PROMPT: Please label the whirlpak with the following label: H.[PID].[DAY].[MONTH]
দয়া করে হুইলপ্যাক ব্যাগে এই ক্রমানুসারে লেবেল বসানঃ H.[খানাআইডি নম্বর].[দিন]. [মাস]

3.4 Has a child hand rinse sample been collected successfully? Select all that apply.
নমুনা হিসাবে নির্দিষ্ট শিশুর উভয় হাতধোয়া পানি সঠিকভাবে সংগ্রহ করা হয়েছে কি? (নিচে প্রদত্ত অপশনগুলো থেকে যতগুলো প্রযোজ্য তা নির্বাচন করম্নন।)

1 = Yes, both hands rinsed(হ্যাঁ, উভয় হাত ধুয়েছে)
2 = No, only one hand rinsed(না, শুধু এক হাত ধুয়েছে)
3 = No, some sample was spilled(না, কিছু নমুনা/পানি ঝরে পড়েছে)
4 = No, the inside of the bag was contaminated(না, নমুনা সংগ্রহের ব্যাগের ভিতরটা দূষিত হয়ে গেছে)
5 = No, sample not collected(না, নমুনা সংগ্রহ করা হয়নি)

3.5 (if 3.4 is not 5) Record time that sample was collected (24H format, HH:MM).
(যদি 3.4এর উত্তর 5না হয়) নমুনা সংগ্রহের সময় লিপিবদ্ধ করম্নন। [24 ঘনটা হিসাবে (ঘনটাঃ মিনিট)]

#### SECTION 4. WATER SAMPLING

If (target child's name) wanted a drink of water right now, could you show me how you would give it to him / her? [If target child it too young to drink water, ask: If your child < 3 years wanted a drink of water right now, could you show me how you would give it to him / her?] [If the mother has no children < 3 years, ask: If you wanted a drink of water right now, could you show me how you would get it?]

Ask the question and observe.

যদি আপনার শিশু (টার্গেট শিশুর নাম বলুন) এখন পানি খেতে চাইত তাহলে আপনি কিভাবে তাকে পানি দিতেন দয়া করে তা আমাকে দেখান। যদি টার্গেট শিশুটি খুব বেশি ছোট হওয়ার কারনে পানি পান না করে থাকে, সেৰেত্রে ঐ টার্গেট মায়ের <৩ wkï এখন পানি খেতে চাইলে তাকেযেভাবেপানি দিতেন দয়া করে তা আমাকে দেখান। যদি মায়ের <৩ বয়সের শিশু না থাকে, তাহলে মাকে জিজ্ঞাসা করম্নন, এখন আপনি পানি খেতে চাইলে যেভাবে পানি নিতেন দয়া করে তা আমাকে দেখান। (প্রশ্ন করম্নন এবং পর্যবেৰন করম্নন)

| | | | | |
|---|---|---|---|---|
| 4.1 | IS WATER FOR DRINKING CURRENTLY AVAILABLE<br><br>(বর্তমানে কি কোন খাবার পানি আছে?) | [1] YES(হ্যাঁ)<br>[2] NO → **SKIP TO SECTION 5**(না)→ **সেকশন 5 এ চলে যান** | | |
| | WHAT DID RESPONDENT DO BEFORE TAKING THE WATER?<br><br>(এইপানিটা খাওয়ার পূর্বে উত্তরদাতা কি করেন?) | YES (হ্যাঁ) | NO (না) | D/K (জানি না) |
| 4.2 | Rinsed glass / cup with drinking water before filling<br>(পানি ভরার পূর্বে গস্নাস/পেয়ালাটি খাওয়ার পানি দিয়ে ধুয়ে নেয়) | [1] | [2] | [99] |

| | | | | |
|---|---|---|---|---|
| 4.3 | Washed hands with water before drinking water was obtained<br>(খাওয়ার পানি ধরার আগে দুই হাত পানি দিয়ে ধুয়ে নিয়েছিল) | [1] | [2] | [99] |
| 4.4 | Washed hands with soap before drinking water obtained<br>(খাওয়ার পানি ধরার আগে দুই হাত সাবান দিয়ে ধুয়ে নিয়েছিল) | [1] | [2] | [99] |
| | FROM WHERE DID THE RESPONDENT TAKE THE WATER?<br>(উত্তরদাতা খাবার পানিটা কোথা থেকে সংগ্রহ করেছিল/নিয়েছিল?) | YES(হ্যাঁ) | NO(না) | D/K(জানি না) |
| 4.5 | Brought directly from the water source<br>(সরাসরি পানির উৎস থেকে নিয়েছিল) | [1] | [2] | [99] |
| 4.6 | Brought directly from water stored in a container<br>(সরাসরি সংরৰনকৃত পানির পাত্র থেকে নিয়েছিল) | [1] | [2] | [99] |
| 4.7 | Brought directly from water filter<br>(সরাসরি ফিল্টার থেকে নিয়েছিল) | [1] | [2] | [99] |
| 4.8 | Stored water was covered<br>(সংরৰনকৃত পানিটা ঢেকে রাখা ছিল) | [1] | [2] | [99] |
| | HOW DID SHE GET THE WATER INTO THE GLASS/CUP?<br>(উত্তরদাতা কিভাবে গস্নাসে/কাপে পানি দিয়েছিল?) | YES (হ্যাঁ) | NO(না) | D/K(জানি না) |
| 4.9 | Hands touched / contacted the drinking water<br>(গস্নাস বা পাত্রটিতে ঢালা পানির ভিতরে তার হাত বা হাতের আঙ্গুল লেগেছিল) | [1] | [2] | [99] |
| 4.10 | Glass/cup dipped into water container<br>(গস্নাস বা পেয়ালাটিতে পানি ভরার সময় তা পানির পাত্রের ভিতর ডুবিয়ে পানি ভরা হয়েছিল) | [1] | [2] | [99] |
| 4.11 | Ladle used to obtain water<br>(গস্নাসে পানি ভরার জন্য লম্বা হাতলযুক্ত কোন চামচ/মগ ব্যবহার করা হয়েছিল) | [1] | [2] | [99] |
| 4.12 | Water poured from container<br>(পানির পাত্র থেকে সরাসরি পানি ঢেলেছিল) | [1] | [2] | [99] |
| 4.13 | Water poured from tap / handpump<br>(সরাসরি ট্যাপ/টিউবওয়েল/ পানির উৎস থেকেঢেলেছিল) | [1] | [2] | [99] |

4.14 (obs) (if 4.6 is 1) What is the type of the container?
[(পর্যবেৰণ) (4.6 নং প্রশ্নের উত্তর 1 হলে) সংরৰণের পাত্রটি কি ধরনের?]

1= Kolshi(কলসি)
2 = Jug(জগ)
3 = Topaz (provided by icddrb)(আইসিডিডিআরবি'র পাত্র-টোপাজ)
4 = Bucket(বালতি)
5 = Other wide mouth container(প্রসস্ব মুখের অন্য কোন পাত্র)
6 = Other narrow mouth container(সরম্ন মুখের অন্য কোন পাত্র)
99 =Could not observe(পর্যবেৰণ করা সম্ভব হয়নি)

4.15 (if 4.6 is 1) How long ago did you or somebody in your home collect this water? (99=Don't know)
(যদি 4.6 এর উত্তর 1 হয়) কতৰন আগে আপনি বা আপনার খানার কোন সদস্য এই খাবার পানিসংগ্রহ করেছেন? (99=জানি না)

☐☐ Minutes(মিনিট)
☐☐ Hours (ঘন্টা)

☐☐ Days (দিন)

4.16 (if 4.6 is 1) Have you done anything to make this water less cloudy or safer to drink?
(যদি 4.6 এর উত্তর 1 হয়) এই খাবার পানি নিরাপদ করার জন্য আপনি কোন কিছু করেছেন কি?
1 = Yes(হ্যাঁ)
2 = No(না)
99 = Don't know / not sure(জানি না/ নিশ্চিত না)

4. 17 (if 4.16 is 1) How was this water treated?
(যদি 4.16 এর উত্তর 1 হয়) কিভাবে এই খাবার পানি নিরাপদ করা হয়?

Do Not Read Responses. Mark all that apply.

উত্তরগুলো পড়ে শুনাবেন না ।( নিচে প্রদত্ত অপশনগুলো থেকে যতগুলো প্রযোজ্য তা নির্বাচন করুন ।)

| | | |
|---|---|---|
| 1 | [1] Yes(হ্যাঁ) [2] No(না) | Aquatabs (আকুয়াট্যাবস) |
| 2 | [1] Yes [2] No | Waterguard / bottled chlorine/halotab[ক্লোরিন (তরল)/হ্যালোট্যাব মিশিয়ে নিই] |
| 3 | [1] Yes [2] No | Boil (পানি ফুটিয়ে নিই) |
| 4 | [1] Yes [2] No | Strain it through cloth or other material (কাপড়েবা অন্য কোন উপাদান দিয়ে ছেকে নিই) |
| 5 | [1] Yes [2] No | Water filter [ceramic, sand, composite] [পানির ফিল্টার ব্যবহার করে (সিরামিক, বালু ইত্যাদি)] |
| 6 | [1] Yes [2] No | Solar disinfection (SODIS) [সূর্যের আলোতে জীবানুমুক্তকরণ] |
| 7 | [1] Yes [2] No | Let it stand and settle [থিতিয়ে নেই] |
| 8 | [1] Yes [2] No | Biosand filter [বায়োস্যান্ডফিল্টার এর মাধ্যমে বিশুদ্ধ করে নিই] |
| 9 | [1] Yes [2] No | Lifestraw Family Filter (Vestergaard Frandsen distributed) [Kenya only][লাইফস্ট্রফ্যামিলি ফিল্টার (শুধুমাত্র কেনিয়ার জন্য প্রযোজ্য)] |
| 10 | [1] Yes [2] No | Coagulant (alum) [ফিটকিরি যোগ করে] |
| 11 | [1] Yes [2] No | PUR (flocculant + disinfectant) [পার] |
| 12 | [1] Yes [2] No | Other (specify) [অন্যান্য (নির্দিষ্ট করে লিখুন)] |

4.18(if 4.16 is 1)Approximately how long ago did you treat the water?
(যদি 4.16এর উত্তর 1=হ্যাঁ হয়) আনুমানিক কতক্ষন আগে আপনি এই খাবার পানি নিরাপদ করেছেন?
(99=জানি না)

☐☐ Minutes(মিনিট)
☐☐ Hours (ঘন্টা)
☐☐ Days (দিন)

PROMPT: Please collect a drinking water sample by pouring the water from glass provided by participant.
দয়া করে উত্তরদাতার দেওয়া খাওয়ারপানি পরীক্ষার জন্য নমুনা হিসাবে হুইল প্যাক ব্যাগে সংগ্রহ করুন।

PROMPT: (if 4.17 is 1=Y or 2=Y or 11=Y)Please use special bag.
(যদি 4.17এর উত্তর 1=হ্যাঁ অথবা 2=হ্যাঁ অথবা 11= হ্যাঁ হয়) দয়া করে বিশেষহুইল প্যাক ব্যাগে সংগ্রহ করুন

4.19 Please enter the 5-digit unique numerical ID.
যে ব্যাগে নমুনা সংগ্রহ করা হবে অনুগ্রহপূর্বক তার গায়ে লেখা 5সংখ্যার ইউনিক আইডি নম্বরটি এনিট্রি করুননঃ

PROMPT: Please label the whirlpak with the following label: W.[PID].[DAY].[MONTH]
দয়া করে হুইলপ্যাক ব্যাগে এই ক্রমানুসারে লেবেল বসানঃ W. [খানা আইডি নম্বর]. [দিন]. [মাস]

4.20 Has a drinking water sample been collected for microbiological testing?
পরীৰা করার জন্য নমুনা হিসাবে খাবার পানি সংগ্রহ করা হয়েছে কি?
1 = Yes(হ্যাঁ)
2 = No(না)

4.21 (if 4.20 is 1) Record time that sample was collected (24H format, HH:MM)
(যদি 4.20এর উত্তর 1 হয়) নমুনা সংগ্রহের সময় লিপিবদ্ধ করুনন। (24ঘন্টা হিসাবে, ঘন্টাঃমিনিট)

PROMPT: (if 4.17 is 1=Y, 2=Y or 11=Y) Please fill a plastic tube with stored water for chlorine testing.
(যদি 4.17এর উত্তর 1=হ্যাঁ, ২= হ্যাঁ অথবা 11= হ্যাঁ হয়) দয়া করে সংরৰনকৃত পানি পরীৰার জন্য নমুনা হিসাবে পস্নাষ্টিক টিউবের মধ্যে সংগ্রহ করুনন।

4.22 (if 4.17 is 1=Y, 2=Y or 11=Y) Has sample from storage container been collected for chlorine testing?
(যদি 4.17এর উত্তর 1=হ্যাঁ, ২= হ্যাঁ অথবা 11= হ্যাঁ হয়) পানি সংরৰনের মুল পাত্র থেকে ক্লোরিন পরীৰা করার জন্য নমুনা হিসাবে পানি সংগ্রহ করা হয়েছে কি?
1 = Yes(হ্যাঁ)
2 = No(না)

4.23 (if 4.22 is 1) Record time that sample was collected (24H format, HH:MM)
[(যদি4.22এর উত্তর 1হয়) নমুনা সংগ্রহের সময় লিপিবদ্ধ করুনন। [24 ঘন্টা হিসাবে, (ঘন্টাঃ মিনিট)]

4.24 (ask/observe) What is the source of thisobserved drinking water that the respondent provided?
[(প্রশ্ন/পর্যবেৰণ) উত্তরদাতারদেওয়া (আপনার পর্যবেৰণকৃত) খাবার পানির প্রধান উৎস কি?]

1 = Tubewell(টিউবওয়েল)
2 = Unprotected spring(অসংরৰিত ঝরনার পানি)
3 = Protected spring(সংরৰিত ঝরনার পানি)
4 = Unprotected dug well(অসংরৰিত পাতকুয়া)
5 = Protected dug well(সংরৰিত পাতকুয়া)
6 = Rainwater collection(বৃষ্টির পানি সংগ্রহ)
7 = Cart with small tank/drum(ছোট ট্যাংকযুক্ত কার্ট)
8 = Tanker truck(ট্যাংকার ট্রাক)
9 = Surface water(river, dam, lake, pond, stream, canal, irrigation channel)[ভূ-পৃষ্ঠের উপরিভাগের পানি(নদী/বাধ/লেক/পুকুর/সেচ নালাথেকে সংরৰিত পানি]
10 = Piped water into dwelling(ঘরের ভিতর ট্যাপ বা পাইপের পানি)
11 = Piped water into yard/plot(উঠানে ট্যাপ বা পাইপের পানি)
77 = Other(specify)[অন্যান্য (নির্দিষ্ট করে লিখুন)]
99 = DK(জানি না)

4.25 (ask/obs) (if 4.24 is 1) Didthe respondent prime the tubewell prior to retrieving the glass of water?
(প্রশ্ন করুনন/পর্যবেৰণ) (যদি 4.24 এর উত্তর 1 হয়) উত্তরদাতা গস্নাসে পানি ভরার জন্য টিউবওয়েলের পানি উত্তোলনে কোন অতিরিক্ত পানি দিয়েছিল কি?

1 = Yes(হ্যাঁ)
2 = No(না)
99 = Could not observe(পর্যবেৰন করা সম্ভব হয়নি)

4.26 (ask) (if 4.25 is 1) What kind of water was used to prime the tubewell?
(প্রশ্ন করুনন) (যদি 4.25 এর উত্তর 1 হয়) টিউবওয়েলের পানি উত্তোলনে কোন ধরনের পানি ব্যবহার করা হয়েছিল?

1= Stored water from same tubewell(একই টিউবওয়েলের সংরক্ষিত পানি)
2 = Water from another tubewell(অন্যান্য টিউবওয়েলের পানি)
3 = Pond water(পুকুরের পানি)
77 = Other(অন্যান্য)

4.27 (ask) (if 4.24 is 1)What is the depth of the tubewell? (ft) (99 = Don't know)
(প্রশ্ন করুন) (4.24 নং প্রশ্নের উত্তর 1হলে) টিউবওয়েলের গভীরতা কত?(ফুট) (99=জানিনা)

4.28 (obs) (if 4.24 is 1) Are there any materials attached to the tubewell mouth?
(পর্যবেক্ষণ) (4.24এর উত্তর 1 হলে) নমুনা সংগ্রহের সময় টিউবওয়েলের মুখে কোন বস্তু লাগানো / আটকানো ছিল কি?

1 = No materials arecovering the tubewell mouth(টিউবওয়েলের মুখে কোন বস্তু লাগানো/ আটকানো নাই)
2 = Cloth (কাপড় )
3 = Plastic(প্লাষ্টিক)
4 =Bamboo(বাঁশ)
77 = Other(অন্যান্য)
99 = Could not observe(পর্যবেক্ষন করা সম্ভব হয়নি)

4.29 (obs) (if 4.24 is 1)Does the tubewell have a platform?
(পর্যবেক্ষণ) (4.24 নং প্রশ্নের উত্তর 1হলে) টিউবওয়েলের কি প্লাটফর্ম আছে?
1 = Yes(হ্যাঁ)
2 = No(না)
99 = Could not observe(পর্যবেক্ষন করা সম্ভব হয়নি)

4.30 (obs) (if. 4.29 is 1) Is the platform intact?
(পর্যবেক্ষণ) (4.29 নং প্রশ্নের উত্তর 1 হলে) টিউবওয়েলের প্লাটফর্মটি কি অক্ষত/ অটুট আছে?
1 = Yes(হ্যাঁ)
2 = No(না)
99 = Could not observe(পর্যবেক্ষন করা সম্ভব হয়নি)

4.31 (obs) (if 4.24 is 1) Is faulty drainage allowing ponding within 2 meters of the tubewell?
(পর্যবেক্ষণ) (4.24 নং প্রশ্নের উত্তর 1 হলে) ত্রুটিপূর্ন নিষ্কাশন ব্যবস্থার মাধ্যমে টিউবওয়েলের ২ মিটারের মধ্যে কোনগর্তেপানি ঢুকে জমে থাকে কি?
1 = Yes(হ্যাঁ)
2 = No(না)
99 = Could not observe(পর্যবেক্ষন করা সম্ভব হয়নি)

4.32 (obs) (if 4.24 is 1)Is there a latrine within 10 meters of the tubewell?
(পর্যবেক্ষণ) (4.24 নং প্রশ্নের উত্তর 1হলে) টিউবওয়েলের আশেপাশে ১০ মিটারের মধ্যে কোন পায়খানা আছে কি?
1 = Yes(হ্যাঁ)
2 = No(না)
99 = Could not observe(পর্যবেক্ষন করা সম্ভব হয়নি)

4.33 (obs) (if 4.32 is 1) How many steps is the closest latrine <10 m from the tubewell?
(পর্যবেক্ষণ) (যদি 4.32 নং প্রশ্নের উত্তর 1হয়) < ১০ মিটারের ভিতরে সবচেয়ে কাছের টয়লেটটি টিউবওয়েল থেকে কত কদম দুরে অবস্থিত?

4.34 (obs) (if 4.32 is 1) How many latrines are present within 10 meters of the tubewell?
(পর্যবেক্ষণ) (যদি 4.32 নং প্রশ্নের উত্তর 1হয়) টিউবওয়েলের আশেপাশে ১০ মিটারের মধ্যে কতগুলো পায়খানা আছে?

4.35 (obs) (if 4.24 is 1)Is there a pond within 10 meters of the tubewell?
(পর্যবেক্ষণ) (যদি 4.24 নং প্রশ্নের উত্তর 1হয়) টিউবওয়েলের আশেপাশে ১০ মিটারের মধ্যে কোন পুকুর আছে কি?
1 = Yes(হ্যাঁ)
2 = No(না)
99 = Could not observe(পর্যবেক্ষন করা সম্ভব হয়নি)

**SECTION 5. RESIDUAL CHLORINE MEASUREMENT**

PROMPT: Please leave the compound before conducting the chlorine test.
ক্লোরিন পরীৰর পূর্বে উক্ত বাড়ি ত্যাগ করম্নন।

PROMPT: (if 4.22 is 1) Measure the free chlorine residual in the stored water sample.
(যদি 4.22 এর উত্তর 1 হয়) সংরৰনকৃত পানির নমুনার মধ্যে মুক্তক্লোরিন আছে কিনা তা পরিমাপ করম্নন।

5.1 (record) (if 4.22 is 1) Free chlorine: _____._____ mg/L
(লিখুন) (যদি 4.22এর উত্তর 1হয়) মুক্ত ক্লোরিনঃ____ . ____ mg/L

5.2 (record) (if 4.22 is 1) Record time that chlorine was measured (24H format, HH:MM)
(লিখুন) (যদি 4.22এর উত্তর 1হয়) ক্লোরিন পরিমাপের সময় লিপিবদ্ধ করম্নন। 24 ঘনটা হিসাবে, (ঘনটাঃ মিনিট)

#### DAY 2 SURVEY

**SECTION 1: IDENTIFICATION AND DEMOGRAPHICS**

1.1 Enter the FRA employee ID(এফআরএ এমপস্নয়ী আইডি নম্বর লিখুন)

1.2 Enter the current day[আজকের তারিখ (সংখ্যায়) লিখুন]

1.3 Enter the current month[বর্তমান মাসের নাম (সংখ্যায়) লিখুন]

1.4 Enter five-digit participant ID
[অংশগ্রহনকারীর আইডি নম্বর লিখুন (৫ সংখ্যার আইডি ওয়াশ বেনিফিট স্টাডি প্রদত্ত তালিকা থেকে)]

1.5 Name of respondent (target child's mother)
[প্রধান উত্তরদাতার (টার্গেট শিশুর মা) নাম লিখুন]ঃ

1.6 Name of target child
[টার্গেট শিশুর নাম লিখুন]ঃ

**SECTION 2. SENTINEL TOY SAMPLING**

2.1(ask) Did you receive a toy ball from our team yesterday?
(প্রশ্ন করম্নন) আপনি গতকাল আমাদের কাছ থেকে একটি খেলনা বল পেয়েছেন কি?
1 = Yes(হ্যাঁ)
2 = No → Skip to Section 3 (না→সেকশন 3 এ চলে যান)

PROMPT: Could you please show me where the ball that was delivered to your household yesterday is currently located?(গতকাল আপনার খানায় যে খেলনা বলটি দেওয়া হয়েছিল বর্তমানে সেটা কোন স্থানে রাখা আছে অনুগ্রহপূর্বক আমাকে সেস্থানটা দেখাতে পারেন কি?)

2.2 (ask) May I now rinse the toy ball that was delivered to your household yesterday?
(প্রশ্ন করম্নন) গতকাল আপনার খানায় যে খেলনা বলটি দেওয়া হয়েছিল সেটা আমি ধুতে পারি কি?

1 = Yes(হ্যাঁ)
2 = Ball was lost → Skip to Section 3(বলটি হারিয়ে গেছে→ সেকশন 3-এ যান)
3 = Ball was given away to another household → Skip to Section 3(বলটি অন্য খানায় দিয়ে দেওয়া হয়েছে→ সেকশন 3-এ যান)
4= Refused → Skip to Section 3(প্রত্যাখ্যান করেছে→ সেকশন 3-এ যান)
77 = Could not retrieve the ball for other reason (specify) → Skip to Section 3[অন্যকোন কারনে বলটি খুঁজে পাওয়া যায়নি (নির্দিষ্ট করম্নন)→ সেকশন 3-এ যান]

2.3 (obs) Where is the ball located?
(পর্যবেৰণ) বলটি কোথায় (স্থান) পাওয়া গিয়াছে?

1 = Outside: on dirt floor [ঘরের বাইরে:ময়লাযুক্তমেঝেতেরাখা]
2 = Outside: on concrete/cement/wood[ঘরের বাইরে:কংক্রিট/ পাকাঁ/কাঠের উপর রাখা]
3 = Outside: in container[ঘরের বাইরে: কোন পাত্রেরমধ্যে রাখা]
4 = Outside: in another home[ঘরের বাইরে:অন্য কোন ঘরে রাখা]
5 = Inside: on dirt floor [ঘরের ভিতরে:ময়লাযুক্ত মেঝেতে রাখা]
6 = Inside: on concrete/cement/wood floor [ঘরের ভিতরে: কংক্রিট/ পাকাঁ/কাঠের ফ্লোর বা মাটিতে রাখা]
7 = Inside: on surface other than ground, not in container (e.g. on bed, on table, etc.) [ঘরের ভিতরেঃ মাটি বা পাত্র ছাড়া অন্য কোন কিছুর মধ্যে রাখা (যেমন-টেবিল, বিছানা ইত্যাদি)]
8= Inside: in storage container/cabinet[ঘরের ভিতরেঃ কোন পাত্রে বা কেবিনেটে রাখা]
9 = In child's hands(বাচ্চার হাতে ছিল)
77 = Other (specify) [অন্য জায়গায়(নির্দিষ্ট করে লিখুন)]
2.4.a (ask)In your opinion, how much did (target child's name) play with the toy ball over the past 24 hours? (read each choice)
(প্রশ্ন করম্নন)আপনার মতে, আপনার বাচ্চা (টার্গেট শিশুর নাম বলুন)গত24ঘন্টায়কত বার আমাদের দেয়া এই খেলনা বল দিয়ে খেলা করেছে? (প্রতিটি অপশন পড়ে শোনাতে হবে)

1= Several times (4 or more times)দিনে বেশকয়েকবার (৪ বার বা তার বেশী)]
2= Few times (2-3 times) দিনে বেশ কয়েকবার (২-৩ বার)]
3 = Only once since he/she got the ball (বাচ্চা বলটি পাওয়ার পর মাত্র একবার খেলেছে)
4 = Never [কখনো না]
99 = DK/Not sure[জানি না/নিশ্চিত না]

2.4.b In your opinion, how much time total did (target child's name) spend playing with the toy ball over the past 24 hours?
(আপনার মতে, গত 24 ঘন্টায় সর্বমোট কতটুকু সময় আপনার শিশু (টার্গেট শিশুর নাম বলুন) এই খেলনা বলটি দিয়ে খেলা করেছে?(99=জানি না)

☐☐ Minutes(মিনিট)
☐☐ Hours (ঘন্টা)

2.5.a (ask) In your opinion, how much did any of the other children in the household or bari play with the toy ball in the past 24 hours? (read each choice)
[(প্রশ্ন করম্নন) আপনার মতে, গত 24ঘন্টায় কতবার আপনার খানার বা বাড়ির অন্যান্য শিশু খেলনা বলটি দিয়ে খেলা করেছে? (প্রতিটি অপশন পড়ে শোনাতে হবে ।)]

1= Several times (4 or more times)(দিনে বেশ কয়েক বার (4 বার বা তার অধিক বার)]
2 = Few times (2-3 times) [দিনে বেশ কয়েক বার ২-৩ বার)]
3 = Only once since he/she got the ball (বাচ্চা বলটি পাওয়ার পর মাত্র একবার খেলেছে)
4 = Never [কখনো না]
99 = DK/Not sure [জানি না/নিশ্চিত না]

2.5.b In your opinion, how much time total did other children in the household or bari spend playing with the toy ball over the past 24 hours?
(আপনার মতে, গত 24 ঘন্টায় সর্বমোট কতটুকু সময় আপনার খানার বা বাড়ির অন্যান্য শিশু এই খেলনা বলটি দিয়ে খেলা করেছে?(99=জানি না)

☐☐ Minutes(মিনিট)
☐☐ Hours (ঘন্টা)

2.6(ask) Did children from other baris play with the toy ball in the past 24 hours? (read each choice)
[(প্রশ্ন করম্নন) গত 24ঘন্টায় (আপনার খানা বা বাড়ি ব্যতীত) অন্য বাড়ির শিশুরা এই খেলনা বলটি দিয়ে খেলা করেছে কি? (প্রতিটি অপশন পড়ে শোনাতে হবে ।)]

1 = They played with it within this bari(তারা খেলনা বলটি দিয়ে এই বাড়িতেই খেলা করেছে)
2 = They played with it outside this bari (তারা খেলনা বলটি দিয়ে এই বাড়ির বাইরে খেলা করেছে)
3 = No(না)

99 = DK/Not sure[জানি না/নিশ্চিত না]

2.7(ask)Did the children play with the ball mostly inside the home (indoors), mostly outside the home (outdoors) or equal amount inside and outside the home?
(প্রশ্ন) বাচ্চারা আমাদের দেয়া এই বল নিয়ে সাধারনত বেশীর ভাগ সময় কোথায় খেলা করেছিল: ঘরের ভেতরে, ঘরের বাইরে অথবা এই দুই জায়গাতেই সমান সময় খেলা করেছিল?

1 = Mostly inside the home [বেশীর ভাগ সময় ঘরের ভেতরে খেলা করেছিল]
2 = Mostly outside the home [বেশীর ভাগ সময় ঘরের বাইরে খেলা করেছিল]
3 = Equal amounts inside/outside home [ঘরের ভিতরে এবং বাহিরে দুই জায়গাতেই সমান সময় খেলা করেছিল]
4 = Children did not play with ball[শিশুরা বলটি দিয়ে খেলা করেনি]
99 = DK/Not sure [জানি না]
2.8 (ask) How did the child play with the ball? Read each choice.
(প্রশ্ন করুন) বল নিয়ে শিশুটি কিভাবে খেলা করেছিল? (প্রতিটি অপশন পড়ে শোনাতে হবে)

1 = With hands(হাতে খেলা করেছিল)
2 = With feet(পায়ে খেলা করেছিল)
3 =With hands and feet(হাত এবং পায়ে খেলা করেছিল)
4 = Children did not play with ball[শিশুরা বলটি দিয়ে খেলা করেনি]
99=DK/Not sure[জানি না/নিশ্চিত না]

2.9 (ask) Did you do anything to clean the ball?
(প্রশ্ন করুন) বলটি পরিস্কার করার জন্য আপনি কিছু করেছেন কি?
1 = Yes(হ্যাঁ)
2 = No(না)

2.10 (ask) (if 2.9 is 1) What did you do to clean the ball? Select all that apply.Read each choice.
(প্রশ্ন করুন) (যদি 2.9এর উত্তর 1 হয়) বলটি পরিষ্কার করার জন্য আপনি কি করেছেন? (নিচে প্রদত্ত অপশনগুলো থেকে যতগুলো প্রযোজ্য তা নির্বাচন করুন) (প্রতিটি অপশন পড়ে শোনাতে হবে)

1 = Washed with water only(শুধু পানি দিয়ে ধুয়েছেন)
2 = Washed with water and soap(পানি ও সাবান দিয়ে ধুয়েছেন)
3 = Wiped with towel(টাওয়াল দিয়ে মুছেছেন)
4 = Wiped on clothes(কাপড় দিয়ে মুছেছেন)
77 = Other (specify)[অন্যান্য (নির্দিষ্ট করে লিখুন)]

2.11 (ask) (if 2.9 is 1) How long ago did you clean the ball? (99=DK)
[(প্রশ্ন করুন)(যদি 2.9 এর উত্তর 1 হয়) কতক্ষন আগে আপনি এই বলটি পরিষ্কার করেছেন? (99= জানি না)]

☐☐ Minutes(মিনিট)
☐☐ Hours (ঘন্টা)

2.12 (obs) Have children played with the ball since you arrived at the household? Select all that apply.
[(পর্যবেক্ষন) আপনি এই খানায় আসার পর থেকে বাচ্চারা কি বলটি দিয়ে খেলা করেছে? (নিচে প্রদত্ত অপশনগুলো থেকে যতগুলো প্রযোজ্য তা নির্বাচন করুন)]

1 = Target child has played with ball(টার্গেট শিশু বলটি দিয়ে খেলা করেছে)
2 = Other children in the household or bari have played with ball(উক্ত খানার বা বাড়ির অন্যান্য শিশুরা বলটি দিয়ে খেলা করেছে)
3 = Children from other baris have played with ball(অন্য বাড়ির শিশুরা বলটি দিয়ে খেলা করেছে)
4= No children have played with ball (কোন শিশুই বলটি দিয়ে খেলা করেনি)

PROMPT: Please label the whirlpak with the following label: T.[PID].[DAY].[MONTH]
দয়া করে হুইলপ্যাক ব্যাগে এই ক্রমানুসারে লেবেল বসানঃ T.[খানা আইডি নম্বর]. [দিন]. [মাস]

2.13 (obs) Toy appearance (পর্যবেক্ষন) যেঅবস্থায় খেলনাটি পাওয়া গেছে:
1=Unused[অব্যবহৃত]
2 =Used, clean appearance[ব্যবহৃত এবং পরিষ্কার অবস্থায়]

3 =Used, unclean appearance[ব্যবহৃত এবং অপরিষ্কার অবস্থায়]
4 =Used, visibly dirty[ব্যবহৃত এবং,খেলনাটিতে দৃশ্যমান ময়লা]
5 =Used, visibly wet(ব্যবহৃত, দৃশ্যত ভেজা)

2.14 (obs) Toy condition[(পর্যবেৰন) খেলনার অবস্থা]
1 = Good condition(ভাল অবস্থায় ছিল)
2 = Bulging (from being stepped on, sat on, heat applied, etc)[চাপ খাওয়া (পায়ের নিচে পড়ে, বলের উপর বসার কারনে, বলের গায়ে তাপ লাগার কারনে, ইত্যাদি]
3 = Deflated (ball has lost air)[চুপসে যাওয়া (বলের ভিতর থেকে বাতাস বের হয়ে যাওয়ার কারনে]
4 = Burst/Torn (ball has visible hole)[ফুটে গেছে/টুটা (বলের মধ্যে গর্ত দেখা যাচ্ছে)]
PROMPT: Collect a toy rinse sample by rinsing the toy in the Whirlpak bag.
হুইলপ্যাক ব্যাগে খেলনা ধোয়া পানির নমুনা সংগ্রহ

2.15 Please enter the 5-digit unique numerical ID.
যে ব্যাগে নমুনা সংগ্রহ করা হবে অনুগ্রহপূর্বক তার গায়ে লেখা 5 সংখ্যার ইউনিক আইডি নম্বরটি এন্ট্রি করুনঃ

2.16 Has a toy rinse sample been collected successfully?
নমুনা হিসাবে খেলনা ধোয়া পানি সঠিকভাবে সংগ্রহ করা হয়েছে কি?

1 = Yes, toy rinsed successfully(হ্যাঁ, খেলনাটি সফলভাবে/ভালভাবে ধুয়েছে)
2 = No, some sample was spilled(না, কিছু নমুনা/পানি ঝরে পড়েছে)
3 = No, the inside of the bag was contaminated(না, নমুনা সংগ্রহের ব্যাগের ভিতরটা দূষিত হয়ে গেছে)
4 = No, sample not collected(না, নমুনা সংগ্রহ করা হয়নি)

2.17 (if 2.16 is not 4) Record time that sample was collected (24H format, HH:MM).
[(যদি 2.16 এর উত্তর 4না হয়) নমুনা সংগ্রহের সময় লিপিবদ্ধ করুন। [24 ঘন্টা হিসাবে, (ঘন্টাঃ মিনিট)]

#### SECTION 3. FLY DENSITY

3.1 Observe fly activity in the food preparation area. Estimate the total number of flies seen.
খাবার তৈরির স্থানে মাছির কর্মকান্ড পর্যবেৰণ করুন। উক্ত স্থানের আশেপাশে কতগুলো মাছি মুক্তভাবে বিচরন করতে দেখা গেছে তার আনুমানিক হিসাব দিন।

1 = None(কোন মাছি দেখা যায়নি)
2 = Low (1-5)(সামান্য পরিমানে)(1-5)
3 = Moderate (6-10)(পরিমানে মাঝারি ধরনের)(6-10)
4 = High(পরিমানে বেশি)(>10)

3.2 Was the fly tape hung under a roof (protected from rain)?
(যে স্থানে ফ্লাইটেপটি বাধাঁ হয়েছে তার উপরে কি ছাদ আছে?(বৃষ্টি থেকে সুরৰিত কিনা?)
1 = Yes(হ্যাঁ)
2 = No(না)

3.3 Is the food prep area inside or outside?
(খাবার তৈরির এই স্থানটি কি ভিতরে না বাহিরে?)

1 = Walls and roof(দেয়াল এবং ছাদ উভয়ই রয়েছে)
2 = Walls but no roof(দেয়াল আছে কিন্তু ছাদ নাই)
3 = Roof but no walls(ছাদ আছে কিন্তু দেয়াল নাই)
4 = No roof and no walls(দেয়াল এবং ছাদ কোনটাই নাই)

3.4 (record) How many steps are the strips hung from the food preparation area?
(লিখুন) [খাবার তৈরির স্থান থেকে কত কদম দুরে (পায়ের ধাপ গননা করুন) ফ্লাইটেপ বাধা হয়েছিল?]

3.5 (obs) Was the fly tape in the food preparation area tampered with or did it fall down?
[(পর্যবেৰণ)(খাবার তৈরির স্থানে বাধাঁ ফ্লাইটেপটি ৰতিগ্রস্ত হয়েছে বা নিচে পড়ে গেছে কি?]

3.6 (record) What is the total number of flies counted on all strips?

(লিখুন) (ফ্লাইটেপে সব মিলিয়ে কতগুলো মাছি ধরা পড়েছে তার সংখ্যা লিপিবদ্ধ করুন।)

3.7(record) (if 3.6 is not 0) Write the number of each species observed.
(লিখুন) (যদি 3.6এর উত্তর 0 নাহয়)(যত প্রজাতির মাছি দেখা গেছে তার সংখ্যা লিখুনঃ)

Musca domestica(মুস্কা ডমেস্টিকা) ______
Lesser house fly(লেজার হাউস ফ্লাই) ______
Blow/bottle fly(বেম্লা/ বোতল ফ্লাই) ______
Flesh fly/sarcophaga(ফ্লেশ ফ্লাই/সারকোফাগা) ______
Other(অন্যান্য) ______
Cannot distinguish(চেনা যায়নি) ______

3.8 Record the time you counted the flies on fly tape at the food preparation area (24H format, HH:MM)
(খাবার তৈরির স্থানে ফ্লাইটেপে আটক মাছি গননার সময় লিপিবদ্ধ করুন: (24 ঘনটা হিসাবে, ঘনটাঃ মিনিট)

3.9 Observe fly activity in the latrine area. Estimate the total number of flies seen.
টয়লেটের মধ্যে মাছির কর্মকান্ড পর্যবেক্ষণ করুন। উক্ত স্থানে কতগুলো মাছি মুক্তভাবে বিচরন করতে দেখা গেছে তার আনুমানিক হিসাব দিন।

1 = None(কোন মাছি দেখা যায়নি)
2 = Low (1-5)(সামান্য পরিমানে)(1-5)
3 = Moderate (6-10)(পরিমানে মাঝারি ধরনের)(6-10)
4 = High(পরিমানে বেশি)(>10)

3.10 Was the fly tape hung under a roof (protected from rain)?
(যে স্থানে ফ্লাইটেপটি বাধাঁ হয়েছে তার উপরে কি ছাদ আছে?(বৃষ্টি থেকে সুরক্ষিত কিনা?)
1 = Yes(হ্যাঁ)
2 = No(না)

3.11 (record) How many steps are the strips hung from the latrine area?
(লিপিবদ্ধ করুন) টয়লেট থেকে কত কিদম দুরে (পায়ের ধাপ গনণা করুন) ফ্লাইটেপ বাধাঁ ছিল?

3.12 (obs) Was the fly tape in the latrine area tampered with or did it fall down?
(পর্যবেক্ষণ)(টয়লেটের আশেপাশে বাধাঁ ফ্লাইটেপটি ক্ষতিগ্রস্থ হয়েছে বা নিচে পড়ে গেছে কি?)

3.13 (record) What is the total number of flies counted on all strips?
(লিপিবদ্ধ করুন) (ফ্লাইটেপে সব মিলিয়ে কতগুলো মাছি ধরা পড়েছে তার সংখ্যা লিপিবদ্ধ করুন।

3.14(record) (if 3.13 is not 0) Write the number of each species observed.
(লিপিবদ্ধ করুন) (যদি 3.13এর উত্তর 0 নাহয়)(যত প্রজাতির মাছি দেখা গেছে তার সংখ্যা লিখুনঃ)

Musca domestica(মুস্কা ডমেস্টিকা) ______
Lesser house fly(লেজার হাউস ফ্লাই) ______
Blow/bottle fly(বেম্লা/ বোতল ফ্লাই) ______
Flesh fly/sarcophaga(ফ্লেশ ফ্লাই/সারকোফাগা) ______
Other(অন্যান্য) ______
Cannot distinguish(চেনা যায়নি) ______

3.15 Record the time you counted the flies on the fly tape at the latrine area (24H format, HH:MM)
(টয়লেটের আশেপাশে/স্থানে ফ্লাইটেপে আটক মাছির মাছি গননার সময় লিপিবদ্ধ করুন: (24 ঘনটা হিসাবে, ঘনটাঃ মিনিট)

3.16 (ask) "When was the most recent time it rained?" (জিজ্ঞাসা করুনঃ সর্বশেষ কখন বৃষ্টি হয়েছিল?)

| | |
|---|---|
| [1] Currently rainingএখনও বৃষ্টি হচেছ | [4]Rained the day before yesterdayগত পরশু বৃষ্টি হয়েছিল |
| [2] Rained earlier todayআজ সকালে বৃষ্টি হয়েছিল | [5]Rained in the last weekগত সপ্তাহে বৃষ্টি হয়েছিল |
| [3] Rained yesterdayগতকাল বৃষ্টি হয়েছিল | [6] Did not rain in the last week. গত সপ্তাহের মধ্যে বৃষ্টি হয়নি |

#### Wash Benefit Module 40: Geographical Identification and target groups

নির্দিষ্টকরন তথ্য গুরুত্বের সাথে পূরন করুন FILL THE FOLLOWING IDENTIFICATION INFORMATION FOR THE RESPONDENT VERY CAREFULLY.

| | | |
|---|---|---|
| Module ID | WBM 40 | |
| | 4001. | এফ.আর.এ কোড (FRA code): ÿÿ |
| C.10<br>1 | 4002. | তথ্য সংগ্রহের তারিখ [Date of data collection]: __/__/____ |
| | dataid | \|__\|__\|__\|__\|__\| |
| | 4003. | |
| | 4004. | |
| | 4005. | |
| | 4006. | জেলার নাম(DISTRICT NAME): |
| | 4007. | উপজেলার নাম (UPAZILA NAME): |
| | 4008. | ইউনিয়নের নাম [Union name _____________ |
| | 4009. | খানার ঠিকানা Household Address: |
| | 4010. | খানা প্রধানের নাম [Name of Household head]_____________________________ |
| | 4011. | খানা প্রধানের বাবা/স্বামীর নাম [Father's/Husband's Name of HH head] _______________ |
| | 4012. | বাড়ির নাম [Bari Name]: ________________________________________ |
| | 4013. | গ্রামের নাম [Village Name]: _____________________________________ |
| | 4014. | Cluste No [1-720] |
| | 4015. | বাড়ী নাম্বার [Bari No. [1-8] |
| | 4016. | **HH ID:** [1-8] |
| | 4017. | |
| | 4017<br>a | এই খানার অবস্থা কি?<br>What is the status of this households?<br>নির্দিষ্ট শিশু জন্মগ্রহণ করেছে 1. Index child born and present<br>নির্দিষ্ট শিশু মৃত্যুবরণ করেছে 2. Index child dead<br>খানা অন্য কোথাও চলে গেছে 3. Household migrated out (skip to 4017e) ) |
| | 4017<br>b | |
| | 4017<br>c. | নির্দিষ্ট শিশু মৃত্যুর কারণ কি? Why is the reason for the child's death?<br>গর্ভপাত (ইচ্ছাকৃত) 1. Abortion<br>গর্ভপাত (অনিচ্ছাকৃত) 2. Miscarriage<br>মৃত সন্তান প্রসব 3. Still birth<br>বাচ্চা অসুস্থতার কারণে মারা গেছে 4. Child died of illness<br>অন্যান্য (উল্লেখ করুণ) [Others (specify)] 5. Other Specify_______________ |
| | 4017<br>d. | কখন এই ঘটনা ঘটেছে? When did this incident happen?<br>__/__/_____ 1. DD/MM/YYYY<br>__ মাস আগে 2. \|__\|__\| months ago<br>মনে নাই ৯৯. Don't remember<br>2nd child (□□□ □□□□)<br>__/__/_____ 1. DD/MM/YYYY<br>__ মাস আগে 2. \|__\|__\| months ago<br>মনে নাই ৯৯. Don't remember |
| | 4017<br>e. | কখন তারা চলে গেছে ?If 4017a=3, When did the household leave?<br>__ মাস আগে 1. \|___\|___\| months ago |
| | 4017<br>f. | আপনি কি জানেন তারা কোথায় গেছে ?Do you know where they went? 1 Yes 0 No (skip to 4019a) |
| | 4017<br>g. | জেলার নাম District Name_________ |
| | 4017i<br>. | উপজেলার নাম Upazilla Name_________ |
| | 4017j | গ্রামের নাম Village Name__________ |

| | | |
|---|---|---|
| | . | |
| | 4017 k. | বাড়ির নাম Bari Name____________ |
|  | 4017 l | -----------প্রাথমিক ফোন নাম্বার (Primary phone number) (Put 00 if not available) (Skip to end) |
| | 4018. | |
| | 4019. | |

#### Wash Benefit Module 41 Household Assets

আপনার সময়ের জন্য আপনাকে ধন্যবাদ।আমি আপনার খানা সম্মন্ধে জানতে চাই যেখানে আপনি বসবাস করেন। প্রথমেই আমি আপনার [illegible] কী দিয়ে তৈরী সেটা দেখতে চাই । তাহলে আমি কি ইহা দেখতে পারি? Thank you very much for your time. Now I would talk abo[illegible] the house that you live in. First, I would like to observe the material your house is made of. Can I take a look at your house?

OBSERVE MAIN MATERIAL OF THE HOUSE:

| | | |
|---|---|---|
| | 4101. | ছাদ তৈরীতে কি কি উপাদান ব্যবহার করা হয়েছ ? (পরির্দশন করে যাচাই করুন) [Main material of the roof]<br>কাঁচা (বাঁশ/খড়) [Kaccha (bamboo / thatch)] 1<br>টিন [Tin] 2<br>সিমেন্ট/ কংক্রিট/ টালি [Cement / concrete / tiled]…3 |
| | 4102. | দেয়াল তৈরীতে কি কি উপাদান ব্যবহার করা হয়েছ ? (পরির্দশন করে যাচাই করুন) [Main material of the walls]<br>পাট/বাঁশ/মাটি (কাঁচা) [Jute / bamboo / mud (*kaccha*)] .. 1<br>কাঠ [Wood] ..................................................... 2<br>ইট/সিমেন্ট [Brick / cement] ............................. 3<br>টিন [Tin].................................................. 4 |
| | 4103. | মেঝে তৈরীতে কি কি উপাদান ব্যবহার করা হয়েছ ? (পরির্দশন করে যাচাই করুন) [*Main material of the floor]*<br>মাটি/বাঁশ (কাঁচা) [Earth / bamboo (kaccha)] ..................1<br>কাঠ [Wood] 2<br>ইট/ সিমেন্ট [Cement / concrete] ...................................3 |
| Thank you. Now I would like to ask you some questions about your household assets. | | |
| | 4104. | খানার ঘরটিতে কতগুলো কক্ষ আছে? (রান্নাঘর ও বাথরুম বাদে) [How many rooms the households have (exclude bathroom and Kitchen)?]. |
| | 4105. | আপনার খানায় নিম্নলিখিত জিনিসগুলো আছে কী? [How many does the households have<br>হ্যাঁ (Yes)....1, না (No)....0, জানি না (DK).....999<br>a. \|_____\|বিদ্যুৎ [Electricity]<br>b. \|_____\|\|_____\|আলমারী/ওয়ারড্রব (সংখ্যা) [Number of Almirah or wardrob[illegible]<br>c. \|_____\|\|_____\|টেবিল (সংখ্যা) [Number of tables]<br>d. \|_____\|\|_____\|চেয়ার/বেঞ্চ (সংখ্যা) [Number of chair or bench]<br>e. \|_____\|\|_____\|হাত ঘড়ি/দেয়াল ঘড়ি (সংখ্যা) [Number of watch or clock]<br>f. \|_____\|\|_____\|খাট (সংখ্যা) [Number of khat]<br>g. \|_____\|\|_____\|চৌকি (সংখ্যা) [Number of chouki ]<br>h. \|_____\|রেডিও [A radio that is working]<br>i. \|_____\|টেলিভিশন (সাদা/কালো) [A B/W television that is working]<br>j. \|_____\|টেলিভিশন (রঙিন) [A color television that is working]<br>k. _____\|রেফ্রিজারেটর [Refrigerator]<br>l. \|_____\|সাইকেল (খেলনা সাইকেল নয়) A bicycle (used for commercial purposes not toy for children)<br>m. \|_____\|মটর সাইকেল [A motorcycle]<br>n. \|_____\|সেলাই মেশিন [A sewing machine]<br>o. \|_____\|\|_____\|মোবাইল ফোন (সংখ্যা) [Number of Mobile phones]<br>p. \|_____\|ল্যান্ড ফোন [A land phone] |

| | | |
|---|---|---|
| | 4106. | আপনার খানায় রান্নার জন্য প্রধানত কি ধরনের জ্বালানী ব্যবহার করা হয় ? [What type of fuel does your household mainly use for cooking?]<br>কাঠ [Wood] ........ 01<br>শস্যের অবশিষ্টাংশ/ঘাস [Crop residue / grass ] ........ 02<br>শুকনা গোবর [Dung cakes ] ........ 03<br>কয়লা [Coal / coke / lignite ] ........ 04<br>কাঠ কয়লা [Charcoal ] ........ 05<br>কেরোসিন [Kerosene ] ........ 06<br>বিদ্যুৎ [Electricity ] ........ 07<br>তরল গ্যাস/প্রাকৃতিক গ্যাস [Liquid gas / gas ] ........ 08<br>বায়ো-গ্যাস [Bio-gas] ........ 09<br>অন্যান্য [Other] ........ 77<br>(বর্ণণা লিখুন) [*Specify other]* ______________________<br>জানি না [Don't know ] ........ 99 |
| | 4107. | I_____I আপনার খানার মোট মাসিক আয় কত? (সকল আয়ের উৎসের যোগফল যেমনঃ বেতন, ঘর ভাড়া, কৃষিখাত ইত্যাদি) For the purpose of our research, would you please tell us your total monthly household income? (Please sum up your income from all sources like wage, rent, agriculture etc.) |
| | Q4107a | **Text value of q4107** |
| | 4108. | I____I বসতবাড়ি মালিকানা [What is the ownership status of the house where your household is currently living?]<br>নিজের বাড়ি [Self-owned] ........ 1<br>ভাড়া বাড়ি [Rental] ........ 2<br>সরকারী জমি [Govt. land] ........ 3.<br>কোন জমিদারের/জোতদারের বাড়িতে থাকে [Owned by a landlord] ........ 4<br>বাড়ি ভাড়া না দিয়ে অন্যের বাড়িতে থাকে [Living in someone's house without giving rent] .......5 |
| | 4109. | খানার সদস্যদের বসতবাড়িতে কোন জমি আছে কি?<br>1. হ্যাঁ<br>0. না (skip to 4111) |
| | 4110. | I____I I____I I____I I____I I____I উত্তর হ্যাঁ হলে, খানার সদস্যদের বসতবাড়িতে মোট কতটুকু জমি আছে (ডেসিমেল)? (How much homestead land (decimal) does your household own?]<br>বলতে রাজি হয়নি [Refused] ........ 88<br>জানি না [Don't know] ........ 99 |
| | 4111. | খানার সদস্যদের বসতবাড়ি ছাড়া অন্য কোন জমি আছে কি?<br>1. হ্যাঁ<br>০. না (skip to 4113) |
| | 4112. | I____I I____I I____I I____I I____I উত্তর হ্যাঁ হলে, খানার সদস্যদের বসতবাড়িতে মোট কতটুকু জমি আছে (ডেসিমেল)? (How much homestead land (decimal) does your household own?]<br>বলতে রাজি হয়নি [Refused] ........ 88<br>জানি না [Don't know] ........ 99 |
| | 4113. | আপনার মতে সামাজিক প্রেক্ষাপটে আপনার খানাটির অবস্থান কি রকম [How would you describe your economic status?]<br>ধনী [Dhoni] ........ 1<br>উচ্চ মধ্যবিত্ত [Uchho modho bitto] ........ 2<br>মধ্যবিত্ত [Modho Bitto] ........ 3<br>দরিদ্র [Doridro or Nimno motho bitto] ........ 4<br>হতদরিদ্র [Hotodoridro] ........ 5 |

| | | | | |
|---|---|---|---|---|
| | 4114. | আমি পশুর নামের তালিকা আপনাকে পড়ে শুনাতে চাচ্ছি ।এ গুলো কোনটি আপনার কম্পাউন্ড/াবাড়ীতে কতগুলো আছে এবং নিজের কয়টা আছে তা আমাকে বলুন । যদি না থাকে তাহলে ০০ বসান । আর যদি অজানা থাকে তাহলে ৯৯ বসান I'm going to read you a list of animals. Please tell me how many of each stays in your compound, and how many belong to your household. IF NONE, ENTER "00", IF UNKNOWN, ENTER "99" | কম্পাউন্ড/াবাড়ী COMPOUND | খানা HOUSEHOLD |
| | 1 | গরু CATTLE | \|__\|__\| | \|__\|__\| |
| | 2 | ছাগল GOATS | \|__\|__\| | \|__\|__\| |
| | 3 | মুরগী/হাঁস CHICKENS | \|__\|__\| | \|__\|__\| |
| | 4 | কবুতর (Pigeon) | \|__\|__\| | \|__\|__\| |
| | 77 | অন্যান্য [Other]............... (বর্নণা লিখুন) [*Specify other]* ___________________ | | |
| | 4115. | কতবার গরম্ন/ছাগল তাদের ঘরের বাহিরে বাড়ীতে/উঠানে ছেড়ে দেন? How often do the cattleroam free in the compound?<br>উত্তরগুলো পড়ুন READ RESPONSES.<br>[1] সবসময় ছেড়ে দেন ALWAYS ROAM FREE<br>[2] মাঝে মাঝে ছেড়ে দেন SOMETIMES ROAM FREE<br>[3] কখনও ছেড়ে দেয়া হয় না NEVER ROAM FREE<br>4. প্রযোজ্য নয়; গরম্ন/ছাগল ছাড়া শুধুমাত্র মুরগী/হাঁস থাকলে N/A; ONLY HAVE CHICKENS | | |
| | 4116. | মুরগী অথবা অন্যান্য পাখি জাতীয় কোন কিছু আপনার প্রধান থাকার ঘরে প্রবেশ করে কি? Do chickens or other poultry ever go inside your main house?<br>READ RESPONSES.<br>[1] সবসময় ALWAYS<br>[2] মাঝে মাঝে SOMETIMES<br>[3] কখনও না NEVER<br>4. প্রযোজ্য নয়; মুরগী/হাঁস না থাকলে N/A; NO CHICKENS | | |

#### Wash Benefit Module 42 Feces Observed in and Around the Compound

সকল খানার জন্য প্রযোজ্য Administer to: All study households

👁 পর্যবেক্ষণ Observation: For the following: প্রত্যেক জায়গায় মানুষের মল/পায়খানার স্তুপের সংখ্যা পর্যবেক্ষণ করুন (১০টি পর্যন্ত ) *Record the number of piles of human feces you observe in each area (up to 10 piles)*

55 অধিক সংখ্যক (স্তুপের সংখ্যা ১০টির উপরে) Too numerous to count (more than 10 piles)

99 বলা যাচ্ছে না/ পর্যবেক্ষণ করা সম্ভব হয়নি Cannot tell / could not observe

👁উঠানে Within the courtyard

| | | | |
|---|---|---|---|
| C.826. | 4201. | উঠানে মানুষের মল/পায়খানা পড়ে আছে যা উন্মুক্ত পায়খানা হিসেবে বিবেচিত হবে **Human** feces within the **courtyard** that could be considered open defecation<br><br>55 অধিক সংখ্যক (স্তুপের সংখ্যা ১০টির উপরে) Too numerous to count (more than 10 piles)<br>99 বলা যাচ্ছে না/ পর্যবেক্ষণ করা সম্ভব হয়নি Cannot tell / could not observe | ☐☐ |
| C.827. | 4202. | উঠানে পশুর/পাখির মল/পায়খানা পড়ে আছে (নির্দিষ্টভাবে উল্লেখ করুন) **Animal** feces present within the **courtyard** (mark all that apply) | ☐1 পাখি বিশেষ(মুরগী/হাঁস/কবুতর )Poultry (chicken, duck, pigeon)<br>☐2 গরু/মহিষ Cow / Buffalo<br>☐3 ছাগল/ভেড়া Goat / Sheep |

| | | | |
|---|---|---|---|
| | | 55 অধিক সংখ্যক (স্তুপের সংখ্যা ১০টির উপরে) Too numerous to count (more than 10 piles)<br>99 বলা যাচ্ছে না/ পর্যবেক্ষণ করা সম্ভব হয়নি Cannot tell / could not observe | ☐4শুকর Pig<br>☐5 কুকুর অথবা বিড়াল Dog or Cat<br>☐8 অন্যান্য Other<br>☐9 কোনও মল/পায়খানা পাওয়া যায়নি NO FECES OBSERVED<br>☐99 পর্যবেক্ষণ করা সম্ভব হয়নি COULD NOT OBSERVE |
| | 4202.a | অন্যকোন খানা এই উঠান ব্যবহার করে কী? Do any other household share this **courtyard** | ১. হ্যাঁ<br>০. না |
| 👁 নির্দিষ্ট শিশু বেশীরভাগ সময় যেখানে সময় কাটায় তা নিরূপন ও যাচাই করুন । বেইজ লাইনে নির্দিষ্ট শিশু থাকবে না সেজন্য গর্ভবতী মা যেখানে বেশীরভাগ সময় কাটায় তা নিরূপন করুন In the area where the target child spends the most time<br>*Probe and identify where the target child spends the most time. At baseline, the target child will not be present, so the search should be defined as the area where the pregnant mother spends the most time.* | | | |
| C.828. | 4203. | মানুষের মল/পায়খানা পড়ে আছে যেখানে নির্দিষ্ট শিশু (বেইজলাইনে গর্ভবতী মা) বেশীরভাগ সময় কাটায়। ইহা উম্মুক্ত পায়খানা হিসেবে বিবেচিত হবে **Human** fecesin the **area where the target child spends the most time** that could be considered open defecation<br>55 অধিক সংখ্যক (স্তুপের সংখ্যা ১০টির উপরে) Too numerous to count (more than 10 piles)<br>99 বলা যাচ্ছে না/ পর্যবেক্ষণ করা সম্ভব হয়নি Cannot tell / could not observe | ☐☐ |
| C.829. | 4204. | পশুর/পাখির মল/পায়খানা পড়ে আছে যেখানে নির্দিষ্ট শিশু (বেইজলাইনে গর্ভবতী মা)বেশীরভাগ সময় সময় কাটায় আছে । (নির্দ্দিষ্টভাবে উল্লেখ করুন) **Animal** feces in the **area where the target child spends the most time**<br>(mark all that apply)<br>55 অধিক সংখ্যক (স্তুপের সংখ্যা ১০টির উপরে) Too numerous to count (more than 10 piles)<br>99 বলা যাচ্ছে না/ পর্যবেক্ষণ করা সম্ভব হয়নি Cannot tell / could not observe | ☐1 পাখি বিশেষ(মুরগী/হাঁস/কবুতর )Poultry (chicken, duck, pigeon)<br>☐2 গরু/মহিষ Cow / Buffalo<br>☐3 ছাগল/ভেড়া Goat / Sheep<br>☐4শুকর Pig<br>☐5 কুকুর অথবা বিড়াল Dog or Cat<br>☐8 অন্যান্য Other<br>☐9 কোনও মল/পায়খানা পাওয়া যায়নি NO FECES OBSERVED<br>☐99 পর্যবেক্ষণ করা সম্ভব হয়নি COULD NOT OBSERVE |
| | 4205. | | |
| 👁 নির্দ্দিষ্ট খানার পিছনে Behind the study house | | | |
| C.830. | 4206. | খানার পিছনে মানুষের মল/পায়খানা পড়ে আছে/থাকলে। ইহা উম্মুক্ত পায়খানা হিসেবে বিবেচিত হবে **Human** fecesbehindthe **house** that could be considered open defecation<br>55 অধিক সংখ্যক (স্তুপের সংখ্যা ১০টির উপরে) Too numerous to count (more than 10 piles)<br>99 বলা যাচ্ছে না/ পর্যবেক্ষণ করা সম্ভব হয়নি Cannot tell / could not observe | ☐☐ |
| C.831. | 4207. | খানার পিছনে পশুর মল/পায়খানা পড়ে আছে/থাকলে। (নির্দ্দিষ্টভাবে উল্লেখ করুন) **Animal** feces **behind the house**<br>(mark all that apply)<br>55 অধিক সংখ্যক (স্তুপের সংখ্যা ১০টির উপরে) Too numerous to count (more than 10 piles)<br>99 বলা যাচ্ছে না/ পর্যবেক্ষণ করা সম্ভব হয়নি Cannot tell / could not observe | ☐1 পাখি বিশেষ(মুরগী/হাঁস/কবুতর )Poultry (chicken, duck, pigeon)<br>☐2 গরু/মহিষ Cow / Buffalo<br>☐3 ছাগল/ভেড়া Goat / Sheep<br>☐4শুকর Pig<br>☐5 কুকুর অথবা বিড়াল Dog or Cat<br>☐8 অন্যান্য Other<br>☐9 কোনও মল/পায়খানা পাওয়া যায়নি NO FECES OBSERVED<br>☐99 পর্যবেক্ষণ করা সম্ভব হয়নি COULD NOT |

| | | | OBSERVE |
|---|---|---|---|

**Wash Benefit Module 22 5 HOURS STRUCTURED OBSERVATIONS**
HHID: ____________________

*Before you begin the observation ask about the people that are currently present in the household. This is so you know how to code each person that is present during the observation.* [PDA will save each entry as Mother, Father according to the codes in col 1.]

Target Household Person Index:

| 1. Person | 1. Sex<br>1.M<br>0.F | 1. Enter the age (in years) | 4.Does the child:<br>1. Crawl<br>2. Walk<br>0. None of the above |
|---|---|---|---|
| 1. Pregnant woman/Mother/ primary care giver<br>2. Father/ male care giver<br>3. Other adult(If 1-3, Skip col. 3)<br>4. Index child<br>5. Other Child | | | |

*Next, before you begin the observation take a look around the compound and locate possible hand washing stations.* [PDA will save each entry as HWS 1, HWS 2 etc.]

Hand washing station index:

| Hand washing station | Location | Hand cleansing agents present (LOOK AT HW) |
|---|---|---|
| HWS1<br>HWS2<br>HWS3 | 1…In/near main house (≤6 ft to entrance)<br>2…In/near latrine (≤6 ft to entrance)<br>3…In/near cooking area (≤6 ft to entrance)<br>4… >6 ft away from main house, latrine and cooking area | [allow multiple options]WATER (present directly from pump/tubewell/water source)<br>[1] WATER (present directly from pump/tubewell/water source)<br>[2] WATER (water present in a container or dispenser, not directly from water source )<br>[3] BAR SOAP (Body/hand Bar)<br>[4] BAR SOAP (other)<br>[5] POWDERED SOAP<br>[6] SOAPY WATER (icddr,b provided<br>[7] LIQUID SOAP<br>[8] Ash<br>[9] Mud/Sand<br>[10] ICDDR,B provided 40L bucket with tap<br>[11] Soapy water (not icddr,b provided)<br>[12] (B) ICDDR,B provided 16L bucket with tap<br>[13] (B) Other bucket<br>[14] (B) Basin<br>[15] Jug<br>[16] Bowl<br>[17] None of the above |

Determine an appropriate location so that child defecation, feeding events and hand washing event can be detected. You are allowed to move during this observation. Always keep the primary care giver in the line of vision. There might be more than one event going on at the same time. Focus on one event at a time. Record details about each observation according to the guide. We will focus on the target household for this observation. If you have notes or comments about related events please record them in paper with the HH ID, during these 5 hours.

Start time: --------/--------hh/mm (24 hour format)

End time: --------/---------hh/mm (24 hour format)

| 0. Who are you observing? [PDA: open HH person list] | 1. Which event are you observing? | 2. PDA to auto-record time at each observation | 3. Open sections within each event |
|---|---|---|---|
| 1. Open target index list<br>2. Other household<br>- Adult, M<br>- Adult, F<br>- < 5 Child, M<br>- < 5 Child, F | 1) Toilet use / defecation | --------/---------hh/mm | 1 Defecation/ Feces Handling/ Disposal<br>2 Hand washing |
| | 1) Other contact with feces | | 1 Defecation/ Feces Handling/ Disposal<br>3 Hand washing |

| | | | |
|---|---|---|---|
| | 1) Food handling<br>1. Mashing food (bhorta)<br>2. Handling raw meat/fish<br>3. Cutting fruits<br>4. Cutting vegetables to be cooked<br>5. Cutting VEGETABLE FOR salad | | 1 Hand washing |
| | 1) Eating | | 1. Other LNS event<br><br>2 Hand washing |
| | 1) Eating (Index child) | | 1 Hand washing<br><br>2 Infant feeding |
| | 1) Washing hands | | 1 Hand washing |
| | 1) Water event | | 1 Water |

Defecation/ Feces Handling/ Disposal Record end time [allow option]

| 1. Whose feces is it? | 2. Location of fecal contact | 3. Person handling feces: | 4. Feces disposed using (allow multiple options): | 5. Disposal site: | 6. Hardware cleaning |
|---|---|---|---|---|---|
| 1. [From person list]<br>2. Cow dung<br>3. Goat feces<br>4. Chicken feces<br>5. Other animal feces (if not known) | 1… Inside toilet (skip col. 7)<br>2… In the courtyard<br>3… Inside the household<br>4… Outside the household but not in the courtyard (beside/behind)<br>5… Potty<br>6… Bush/Jungle/field<br>1. Other | Open list<br>99. No one (skip to end) | 1. Hands only<br>2. Cloth/paper/leaves<br>3. Scrap material to scoop feces<br>4. Potty [do col.6, or else skip]<br>5. Local agricultural hoe<br>6. Sani scoop [do col.6, or else skip]<br>7. Did nothing<br>8. Other<br>99. Could not observe | 1. Improved Latrine<br>2. Unimproved latrine<br>3. Open Pit / separate pit for child or animal feces<br>4. Bury it / Covered Pit<br>5. Undefined open site near the compound<br>6. Garbage disposal sites / dumps)<br>7. Bush / forest / field<br>8. Nearby water (pond, canal, river)<br>99. Could not observe | 1. Cleaned at tube well<br>2. Cleaned in the latrine<br>3. Cleaned at nearby water source (pond)<br>4. Cleaned in the courtyard using self poured water<br>5. Did not clean during this event observation |

Hand washing Record end time [allow option]

| 1. Were both hands cleansed? | 2. When were the hands cleansed? | 2. Hand cleansing materials: | 3. How were the hands washed? [multiple options] | 4. How were hands dried? | 5. Hand washing station index number | 6.Location if not at indexed HWS |
|---|---|---|---|---|---|---|
| **1....** Yes<br>**2....** No [skip to end]<br>99.Could not observe [skip to end] | (mostly applicable for food preparation times)<br>**1.** Before<br>2. After | **1....** Water only<br>**2....** Bar soap<br>**3....** Powdered Soap and water<br>**4....** Soapy water and rinse water<br>**5....** Soapy water only<br>**6....** Sanitizer<br>**7....** Other, Specify<br>9.... Could not observe | 1 Y 0 N<br>**1.** palms<br>**2.** back of hands<br>**3.** between fingers of hands<br>**4.** under fingernails | **1....** Not Dried<br>**2....** Air dried<br>**3....** Towel/Cloth<br>**4....** Clothing<br>99.Could not observe | [PDA open HW list to choose from] skip col 6<br>(put 99 if at a place not indexed) | *Record first option that fits*<br>1...In/near main house (≤6 ft to entrance)<br>2...In/near latrine (≤6 ft to entrance)<br>3...In/near cooking area (≤6 ft to entrance)<br>4... >6 ft away from main house, latrine and cooking area |

Water Event

| 1. What are you observing? | Where did respondent take the water from? | What type of storage container is it?<br><br>[multiple options] | Treatment/treated | Handling<br><br>[multiple response] |
|---|---|---|---|---|
| 1. Water collection<br>2. Water treatment<br>3. Water handling<br>4. Water storage<br>5. Drinking event | 1. BROUGHT DIRECTLY FROM WATER SOURCE<br>2. BROUGHT DIRECTLY FROM STORAGE CONTAINER<br>3. BROUGHT DIRECTLY FROM WATER FILTER<br>4. STORED WATER WAS COVERED | 1. 5L icddrb Kolshi<br>2. <5l container<br>3. >5L container<br>4. Hari (wide container)<br>5. Jug<br>6. Glass | 1. Aquatabs<br>2. Boiling<br>3. Straining<br>4. Did not treat<br>5. Could not determine | 1. rinsed glass/container with drinking water before filling<br>2. washed hands with water (no soap) before drinking water was obtained<br>3. washed hands with soap before drinking water obtained<br>4. hands touched / contacted the drinking water during retrieval<br>5. container/glass dipped into water container<br>6. ladle used to obtain water<br>7. water poured from container<br>8. water poured from tap |

Infant Feeding/Eating Record end time [allow option]

| 1. Type of food (allow multiple options) | 2. Person feeding (allow multiple options) | 3. How was LNS served: | 4. *How was LNS fed to child:* | 5. Consumption: |
|---|---|---|---|---|
| **1.** Breastfed<br>**2.** LNS (fill Col 3,4,5 else skip)<br>**3.** Water/sugar water/honey water<br>**4.** Milk<br>**5.** Formula<br>**6.** Boiled rice<br>**7.** Khichuri (rice, lentil, potato, tomato, brinjal, beans)<br>**8.** Semolina/Porridge /Grains<br>**9.** Lentil<br>**10.** Eggs<br>**11.** Juice<br>**12.** Other liquids (ex. soup)<br>**13.** leafy vegetable<br>**14.** orange or yellow fruits/vegetables<br>**15.** other fruits/veg<br>**16.** Cheese/yogurt<br>**17.** Meat/fish/eggs<br>**18.** Packaged food (biscuits, chips, candy)<br>99. Other foods/drinks not observed | **1.** Child (self)<br>2. Primary Caregiver, F<br>**3.** Primary caregiver, M<br>**4.** Secondary Caregiver F<br>**5.** Secondary caregiver, M<br>**6.** <6 year Child, F<br>**7.** <6 year Child, M<br>8. ≥7-15 year Child, F<br>**9.** ≥7-15 year Child, M<br>**10.** Other Adult, F<br>16. Other Adult, M | 1. Mixed into food with a spoon<br>2. Mixed into food by hand.<br>3. Fed directly from the packet | 1. By hand<br>2. By spoon<br>3. Direct from packet | 1. Consumed full packet<br>2. Partial and thrown away<br>3. Partial and eaten later |

Other LNS event

| 1. Did they consume LNS?<br><br>**1....** Yes<br>**2....** No [skip to end]<br>99.Could not observe [skip to end] | 2. Who consumed or took the LNS? | 3.How much was consumed? |
|---|---|---|
| | **1.** [From person list] | **1.** Just a taste<br>**2.** Partial packet consumed<br>**3.** Whole packet consumed<br>4. Whole packet taken, consumption not observed |

Additional questions at the end of observation time: 1 Y 0 N

1. Did you feed LNS to your child today before I arrived?
2. Are you planning to feed LNS to your child later this afternoon or this evening?

#### Wash Benefit module: Sustainability Instrument for Endline

| Water পানি | | | |
|---|---|---|---|
| (OBSERVE): Types of water container and covering status?<br>পর্যবেক্ষণ: পানির পাত্র এবং ঢাকনা বিষয়ক প্রশ্নাবলী | | 1=Yes হ্যাঁ<br><br>0=No না | 1= Water present and completely uncovered<br>পাত্রে পানি আছে এবং কোন ঢাকনা নাই<br>2= Water present and partially covered<br>পাত্রে পানি আছে এবং আংশিক ঢাকা আছে<br>3= Water present and completely covered<br>পাত্রে পানি আছে এবং সম্পূর্ণ ঢাকা আছে<br>4= No water present<br>পাত্রে পানি নাই |
| 1.1 | Intervention container<br>আইসিডিডিআরবি'র পাত্র/টোপাস | □ | □ |
| 1.2 | Bucket বালতি | □ | □ |
| 1.3 | Jug জগ | □ | □ |
| 1.4 | Matka মটকা | □ | □ |
| 1.5 | Cooking pot রান্নার পাত্র | □ | □ |
| 1.6 | Jerry can জেরিকেন | □ | □ |
| 1.7 | *Hari*(Wide-mouthed container)<br>হাড়ি ( প্রশস্ব মুখবিশিষ্ট পাত্র) | □ | □ |
| 1.8 | Other wide-mouthed containe<br>অন্যান্য প্রশস্ব মুখবিশিষ্ট পাত্র | □ | □ |
| 1.9 | *Kalash* (narrow-mouthed container)<br>কলস (অপ্রশস্ব মুখবিশিষ্ট পাত্র) | □ | □ |
| 1.10 | Other narrow-mouthed container<br>অন্যান্য অপ্রশস্ব মুখবিশিষ্ট পাত্র | □ | □ |
| 1.11 | Other (Specify)<br>অন্যান্য পাত্র (নির্দিষ্ট করুন) | □ | □ |

| What benefits come from safely storing drinking water? [Note: Don't read the answer.]<br>নিরাপদভাবে সংরক্ষকৃত খাবার পানির সুবিধা কি কি? [নোট: উত্তরগুলো পড়ে শোনানো যাবে না] | | | |
|---|---|---|---|
| 2.1 | Less diarrhea ডায়রিয়া কম হয় | Yes =1 হ্যাঁ =১<br>No=0 না= ০ | □ |
| 2.2 | Less illness (type of illness not specifie)<br>অসুখ কম হয় (নির্দিষ্ট কোন অসুখের নাম বলেনি) | Yes =1 হ্যাঁ =১<br>No=0 না= ০ | □ |
| 2.3 | Less germs জীবানু কম হয় | Yes =1 হ্যাঁ =১<br>No=0 না= ০ | □ |
| 2.4 | Water isn't contaminated (germs not mentioned)<br>পানি দুষিত হয়না | Yes =1 হ্যাঁ =১<br>No=0 না= ০ | □ |
| 2.5 | Water tastes better পানির স্বাদ ভালো থাকে | Yes =1 হ্যাঁ =১<br>No=0 না= ০ | □ |
| 2.6 | Water looks better পানি ভালো দেখায় | Yes =1 হ্যাঁ =১<br>No=0 না= ০ | □ |
| 2.7 | No benefits কোন সুবিধা নাই | Yes =1 হ্যাঁ =১<br>No=0 না= ০ | □ |
| 2.8 | Don't know জানিনা | Yes =1 হ্যাঁ =১<br>No=0 না= ০ | □ |
| 2.9 | Other (specify) অন্যান্য (নির্দিষ্ট করুন) | Yes =1 হ্যাঁ =১<br>No=0 না= ০ | □ |

| What kind of storage container do you prefer to use?<br>পানি সংরক্ষণের জন্য কোন ধরনের পাত্র ব্যবহার করতে পছন্দ করেন? | | |
|---|---|---|
| 1 | ¨ | Topaz টোপাস |
| 2 | ¨ | Bucket বালতি |
| 3 | ¨ | Jug জগ |

| | | |
|---|---|---|
| 4 | ¨ | Matka মটকা |
| 5 | ¨ | Cooking pot রান্নার পাত্র |
| 6 | ¨ | Hari (wide-mouthed container) হাড়ি ( প্রশস্ব মুখবিশিষ্ট পাত্র) |
| 7 | ¨ | Other wide-mouthed container অন্যান্য প্রশস্ব মুখবিশিষ্ট পাত্র |
| 8 | ¨ | Kalash (narrow-mouthed container) কলস (অপ্রশস্ব মুখবিশিষ্ট পাত্র) |
| 9 | ¨ | Other narrow-mouthed container অন্যান্য অপ্রশস্ব মুখবিশিষ্ট পাত্র |
| 10 | ¨ | Other (specify) অন্যান্য পাত্র (নির্দিষ্ট করুন) |

What would you do if your topaz breaks? Do Not Read Responses. Mark All that Apply. 1 = Yes
যদি টোপাস ভেঙ্গে যায় আপনি কী করবেন? উত্তর পড়ে শোনানো যাবেনা। একাধিক উত্তর। ১=হ্যাঁ

| | | |
|---|---|---|
| 1 | ¨ | Try to repair it মেরামত করার চেষ্টা করবো |
| 2 | ¨ | Call CHP সিএইচপি'কে ডাকবো |
| 3 | ¨ | Buy a wide-mouthed storage container প্রশস্ব মুখবিশিষ্ট পাত্র কিনবো |
| 4 | ¨ | Buy a narrow-mouthed storage container অপ্রশস্ব মুখবিশিষ্ট পাত্র কিনবো |
| 5 | ¨ | Buy a storage container with a lid and a tap ঢাকনা ও ট্যাপ বিশিষ্ট সংরৰণ পাত্র কিনবো |
| 6 | ¨ | Use a container present in the home (without a lid or tap)<br>বাড়ীতে আছে এমন কোন পাত্র ব্যবহার করবো (ঢাকনা ও ট্যাপ ছাড়া) |
| 7 | ¨ | Use a container present in the home and cover it<br>বাড়ীতে আছে এমন কোন পাত্র ব্যবহার করবো এবং ঢাকনা দিয়ে ঢেকে রাখবো |
| 8 | ¨ | Do not buy a storage container and get water straight from tap/tube well for use<br>কোন পাত্র কিনবো না, সরাসরি ট্যাপ বা টিউবওয়েল থেকে পানি ব্যবহার করবো |
| 9 | ¨ | Use a kolshi কলসি ব্যবহার করবো |
| 999 | ¨ | Don't know জানিনা |

| |
|---|
| How much do you think it would cost to repair your topaz if it broke? (open ended)<br>যদি টোপাস ভেঙ্গে যায় এটি মেরামতের জন্য কত টাকা খরচ হতে পারে বলে আপনি মনে করেন? (খোলা প্রশ্ন) |

| |
|---|
| At what price would you buy a new safe water storage container? (open ended)<br>কত টাকা দামে আপনি নিরাপদ পানির জন্য নতুন একটা সংরৰণ পাত্র কিনবেন? (খোলা প্রশ্ন) |

How does stored water get contaminated? Do Not Read Responses. Mark All that Apply. 1=Yes
কীভাবে সংরৰিত পানি দূষিত হতে পারে? **উত্তর পড়ে শোনানো যাবেনা। একাধিক উত্তর।** ১=হ্যাঁ

| | | |
|---|---|---|
| 1 | ¨ | Reaching into container with hand<br>সংরৰণ পাত্রের ভেতর হাত পৌছালে |
| 2 | ¨ | Reaching into container with a ladle হাতলযুক্ত কোন কিছু সংরৰণ পাত্রের ভেতর পৌছালে |
| 3 | ¨ | No lid on container সংরৰিত পাত্রের ঢাকনা না থাকলে |
| 4 | ¨ | Animals contaminate stored water পশুপাখির মাধ্যমে সংরৰিত পানি দুষিত হতে পারে |
| 5 | ¨ | Children contaminate stored water শিশুদের মাধ্যমে সংরৰিত পানি দুষিত হতে পারে |
| 6 | ¨ | Stored water becomes contaminated if you store it too long<br>যদি দীর্ঘ সময় পানি সংরৰণ করা হয়, পানি দূষিত হতে পারে |
| 7 | ¨ | Other অন্যান্য |
| 999 | ¨ | Don't know জানিনা |

| | |
|---|---|
| If you do not store your drinking water in a topaz...<br>যদি আপনি খাবার পানি টোপাসে সংরৰণ না করেন... | |
| How likely is it that you will get diarrhea?<br>আপনি ডায়রিয়ায় আক্রান্ত হবেন বলে মনে করেন | [1 Very unlikely] সম্ভাবনা খুবই কম<br>[2 Unlikely] সম্ভাবনা কম<br>[3 Likely] সম্ভবত<br>[4 Very likely] খুব সম্ভবত |
| How likely is it that your child will get diarrhea?<br>আপনার শিশু ডায়রিয়ায় আক্রান্ত হবেন বলে মনে করেন | [1 Very unlikely] সম্ভাবনা খুবই কম<br>[2 Unlikely] সম্ভাবনা কম<br>[3 Likely] সম্ভবত<br>[4 Very likely] খুব সম্ভবত |

| | |
|---|---|
| It is a problem when my neighbors do not safely store their drinking water.<br>আমার প্রতিবেশীরা নিরাপদভাবে খাবার পানি সংরৰণ করেনা এটি একটি সমস্যা.... | [1 Strongly disagree]<br>দৃঢ়ভাবে ভিন্নমত পোষণ করছি<br>[2 Disagree] ভিন্নমত পোষণ করছি<br>[3 Agree ] একমত<br>[4 Strongly agree]<br>দৃঢ়ভাবে একমত পোষণ করছি |
| If my child gets diarrhea it is very serious<br>যদি আমার শিশুর ডায়রিয়া হয় এটা খুবই গুরুত্বর বিষয়.... | [1 Strongly disagree]<br>দৃঢ়ভাবে ভিন্নমত পোষণ করছি<br>[2 Disagree] ভিন্নমত পোষণ করছি<br>[3 Agree ] একমত<br>[4 Strongly agree]<br>দৃঢ়ভাবে একমত পোষণ করছি |
| My child's health has not improved after drinking treated water.<br>বিশুদ্ধ খাবার পানি খাওয়ার পরও আমার শিশুর স্বাস্থ্যের উন্নতি হয়নি... | [1 Strongly disagree]<br>দৃঢ়ভাবে ভিন্নমত পোষণ করছি<br>[2 Disagree] ভিন্নমত পোষণ করছি<br>[3 Agree ] একমত<br>[4 Strongly agree]<br>দৃঢ়ভাবে একমত পোষণ করছি |

| | |
|---|---|
| How confident are you that you can clean your water storage container before using it to store drinking water?<br>পানীয় জল সংরৰণ করতে পানীয় জলের সংরৰণ পাত্র ব্যবহার করার পূর্বে আপনি আপনার সংরৰণ পাত্র পরিষ্কার করেন এ ব্যাপারে আপনি কতটা আত্মবিশ্বাসী? | [ 1 Highly confident ]<br>অত্যন্ত আত্মবিশ্বাসী<br>[ 2 Confident ] আত্মবিশ্বাসী<br>[ 3 Less confident ]<br>কম আত্মবিশ্বাসী<br>[ 4 Not confident ]<br>আত্মবিশ্বাসী নই |

| | |
|---|---|
| Do you share your clean drinking water with your neighbors?<br>আপনি আপনার প্রতিবেশীর সাথে পরিষ্কার পানীয় জল শেয়ার করেন? | 1 Yes হ্যাঁ<br>2 No না |

| | |
|---|---|
| Do you find the topaz an asset for your household?<br>আপনি কী টোপাসকে আপনার খানার একটা সম্পদ মনে করেন? | 1 Yes হ্যাঁ<br>2 No না<br>999 Don't know জানিনা |

| | |
|---|---|
| Has your household social status changed by being known as drinking treated water?<br>বিশুদ্ধ খাবার পানির পরিচিতি কী আপনার খানার সামাজিক অবস্থানকে পরিবর্তিত করেছে? | 1 Our social status increased<br>আমাদের সামাজিক মর্যাদা বৃদ্ধি পেয়েছে<br>2 Our social status decreased<br>আমাদের সামাজিক মর্যাদা কমেছে<br>3 No change in social status<br>আমাদের সামাজিক মর্যাদার কোন পরিবর্তন ঘটেনি |

| | |
|---|---|
| Has your household social status changed by being known as storing your drinking water in a topaz? | 1 Our social status increased<br>আমাদের সামাজিক মর্যাদা বৃদ্ধি পেয়েছে<br>2 Our social status decreased |

| | |
|---|---|
| টোপাসে বিশুদ্ধ খাবার পানি সংরক্ষণের পরিচিতি কী আপনার খানার সামাজিক অবস্থানকে পরিবর্তিত করেছে | আমাদের সামাজিক মর্যাদা কমেছে<br>3 No change in social status<br>আমাদের সামাজিক মর্যাদার কোন পরিবর্তন ঘটেনি |

| | |
|---|---|
| How likely is it that you will continue to store your water in a container with a lid and tap?<br>ঢাকনা ও ট্যাপবিশিষ্ট পাত্রে আপনার পানি সংরক্ষণ করার প্রক্রিয়া বহাল রাখার সম্ভাবনা কেমন? | 1 Very unlikely] সম্ভাবনা খুবই কম<br>[2 Unlikely] সম্ভাবনা কম<br>[3 Likely] সম্ভবত<br>[4 Very likely] খুব সম্ভবত |

### Handwashing

What would you do if your handwashing station breaks? Do Not Read Responses. Mark All that Apply. 1 = Yes
যদি আপনার হাত ধোয়ার স্থানের/যন্ত্রপাতি কোন কিছু ভেঙ্গে যায় আপনি কী করবেন? **উত্তর পড়ে শোনানো যাবেনা। একাধিক উত্তর।** ১=হ্যাঁ

| | | |
|---|---|---|
| 1 | ¨ | Try to repair it মেরামত করার চেষ্টা করবো |
| 2 | ¨ | Call CHP সিএইচপি'কে ডাকবো |
| 3 | ¨ | Buy parts to make my own station আমার নিজের ষ্টেশনের তৈরী করার জন্য যন্ত্রপাতি কিনবো |
| 4 | ¨ | Replace it with a bucket and jug বালতি ও জগ দিয়ে এর প্রতিস্থাপন করবো |
| 5 | ¨ | Stop using it ব্যবহার বন্ধ করবো |
| 6 | ¨ | Other (specify) অন্যান্য (নির্দিষ্ট করুন) |
| 999 | ¨ | Don't know জানিনা |

| | |
|---|---|
| ¨Did the lidto your handwashing station ever break?<br>আপনার হাত ধোয়ার স্থানের/যন্ত্রপাতির ঢাকনা কখনো ভেঙ্গে গিয়েছিল? | 1 Yes হ্যাঁ<br>2 No না |

| | |
|---|---|
| ¨Did the tap to your handwashing station ever break?<br>আপনার হাত ধোয়ার স্থানের/ যন্ত্রপাতির ট্যাপ কখনো ভেঙ্গে গিয়েছিল? | 1 Yes হ্যাঁ<br>2 No না |

| | |
|---|---|
| ¨ Is your handwashing station kept in the shade?<br>আপনার হাত ধোয়ার স্থান/ যন্ত্রপাতি কী ছায়ায় রাখা হয়? | 1 Yes হ্যাঁ<br>2 No না |

| | |
|---|---|
| If you never wash your hands…<br>যদি আপনি কখনো আপনার হাত না ধোন.... | |
| How likely is it that you will get diarrhea?<br>আপনার ডায়রিয়ায় আক্রান্ত হবার সম্ভাবনা কেমন? | 1 Very unlikely] সম্ভাবনা খুবই কম<br>[2 Unlikely] সম্ভাবনা কম<br>[3 Likely] সম্ভবত<br>[4 Very likely] খুব সম্ভবত |
| How likely is it that your child will get diarrhea?<br>আপনার বাচ্চার ডায়রিয়ায় আক্রান্ত হবার সম্ভাবনা কেমন? | 1 Very unlikely] সম্ভাবনা খুবই কম<br>[2 Unlikely] সম্ভাবনা কম<br>[3 Likely] সম্ভবত<br>[4 Very likely] খুব সম্ভবত |

| | |
|---|---|
| It is a problem when my neighbors do not wash their hands with soapy water.<br>এটা একটা সমস্যা যখন আমার প্রতিবেশীরা সাবান-পানি দিয়ে হাত ধোয় না। | 1 Strongly disagree]<br>দৃঢ়ভাবে ভিন্নমত পোষণ করছি<br>[2 Disagree] ভিন্নমত পোষণ করছি<br>[3 Agree ] একমত<br>[4 Strongly agree]<br>দৃঢ়ভাবে একমত পোষণ করছি |

Ask husband or head of household স্বামী বা খানার প্রধানকে জিজ্ঞাসা করুন

If your handwashing station breaks: যদি আপনার হাত ধোয়ার স্থান/যন্ত্রপাতি ভেঙ্গে যায়:

1. At what price would you buy a new tap? (open ended)

| |
|---|
| ১. কত দামে আপনি একটি নতুন ট্যাপ কিনবেন? (খোলা প্রশ্ন) |
| 2. At what price would you buy a new bucket? (open ended)<br>২. কত দামে আপনি একটি নতুন বালতি কিনবেন? (খোলা প্রশ্ন) |
| 3. At what price would you buy a new stool? (open ended)<br>৩. কত দামে আপনি একটি নতুন টুল কিনবেন? (খোলা প্রশ্ন) |
| 4. At what price would you buy a new bowl? (open ended)<br>৪. কত দামে আপনি একটি নতুন গামলা কিনবেন? (খোলা প্রশ্ন) |
| 5. At what price would you buy a new covering? (open ended)<br>৫. কত দামে আপনি একটি নতুন ঢাকনা কিনবেন? (খোলা প্রশ্ন) |

**If your handwashing station breaks: Without help from a CHP how confident are you that…**
যদি আপনার হাত ধোয়ার স্থান/ যন্ত্রপাতি ভেঙ্গে যায়: আপনি কতটা আত্মবিশ্বাসী যে একজন সিএইচপির সাহায্য ছাড়াই......

| | | |
|---|---|---|
| 1 | You know which parts to buy for the handwashing station?<br>আপনি জানেন হাত ধোয়ার স্থানের জন্য কোন যন্ত্র বা অংশটি কিনতে হবে? | [ 1 Highly confident ]<br>অত্যন্ত আত্মবিশ্বাসী<br>[ 2 Confident ] আত্মবিশ্বাসী<br>[ 3 Less confident ]<br>কম আত্মবিশ্বাসী<br>[ 4 Not confident ]<br>আত্মবিশ্বাসী নই |
| 2 | You can fit a new tap?<br>আপনি একটি নতুন কল/ট্যাপ লাগাতে পারেন? | [ 1 Highly confident ]<br>অত্যন্ত আত্মবিশ্বাসী<br>[ 2 Confident ] আত্মবিশ্বাসী<br>[ 3 Less confident ]<br>কম আত্মবিশ্বাসী<br>[ 4 Not confident ]<br>আত্মবিশ্বাসী নই |
| 3 | You can buy a new bucket?<br>আপনি একটি নতুন বালতি কিনতে পারেন? | [ 1 Highly confident ]<br>অত্যন্ত আত্মবিশ্বাসী<br>[ 2 Confident ] আত্মবিশ্বাসী<br>[ 3 Less confident ]<br>কম আত্মবিশ্বাসী<br>[ 4 Not confident ]<br>আত্মবিশ্বাসী নই |
| 4 | You can buy a new stool?<br>আপনি একটি নতুন টুল কিনতে পারেন? | [ 1 Highly confident ]<br>অত্যন্ত আত্মবিশ্বাসী<br>[ 2 Confident ] আত্মবিশ্বাসী<br>[ 3 Less confident ]<br>কম আত্মবিশ্বাসী<br>[ 4 Not confident ]<br>আত্মবিশ্বাসী নই |
| 5 | You can buy a new bowl?<br>আপনি একটি নতুন গামলা কিনতে পারেন? | [ 1 Highly confident ]<br>অত্যন্ত আত্মবিশ্বাসী<br>[ 2 Confident ] আত্মবিশ্বাসী<br>[ 3 Less confident ]<br>কম আত্মবিশ্বাসী<br>[ 4 Not confident ]<br>আত্মবিশ্বাসী নই |
| 6 | You can make a new covering?<br>আপনি একটি নতুন ঢাকনা তৈরী পারেন? | [ 1 Highly confident ]<br>অত্যন্ত আত্মবিশ্বাসী<br>[ 2 Confident ] আত্মবিশ্বাসী<br>[ 3 Less confident ]<br>কম আত্মবিশ্বাসী<br>[ 4 Not confident ] |

| | | |
|---|---|---|
| | | আত্মবিশ্বাসী নই |
| 7 | If your soapy water bottle breaks, how confident are you that you can make your own soapy water bottle?<br>যদি আপনার সাবান-পানির বোতল ভেঙ্গে যায়, আপনি আপনার নিজের সাবান-পানির বোতল তৈরী করতে পারবেন এ ব্যাপারে আপনি কতটা আত্মবিশ্বাসী ? | [ 1 Highly confident ] অত্যন্ব আত্মবিশ্বাসী<br>[ 2 Confident ] আত্মবিশ্বাসী<br>[ 3 Less confident ] কম আত্মবিশ্বাসী<br>[ 4 Not confident ] আত্মবিশ্বাসী নই |
| 8 | How confident are you that you can continue washing your hands with soapy water after defecation?<br>এ ব্যাপারে আপনি কতটা আত্মবিশ্বাসী যে পায়খানা করার পর আপনি আপনার হাত সাবান-পানি দিয়ে ধোয়া চালিয়ে যাবেন? | [ 1 Highly confident ] অত্যন্ব আত্মবিশ্বাসী<br>[ 2 Confident ] আত্মবিশ্বাসী<br>[ 3 Less confident ] কম আত্মবিশ্বাসী<br>[ 4 Not confident ] আত্মবিশ্বাসী নই |
| 9 | How confident are you that you can continue washing your hands with soapy water after cleaning a child's bottom?<br>এ ব্যাপারে আপনি কতটা আত্মবিশ্বাসী যে বাচ্চাকে শৌচানের পর আপনি আপনার হাত সাবান-পানি দিয়ে ধোয়া চালিয়ে যাবেন? | [ 1 Highly confident ] অত্যন্ব আত্মবিশ্বাসী<br>[ 2 Confident ] আত্মবিশ্বাসী<br>[ 3 Less confident ] কম আত্মবিশ্বাসী<br>[ 4 Not confident ] আত্মবিশ্বাসী নই |
| 10 | How confident are you that you can continue washing your hands with soapy water before preparing a meal?<br>এ ব্যাপারে আপনি কতটা আত্মবিশ্বাসী যে খাবার তৈরী করার পূর্বে আপনি আপনার হাত সাবান-পানি দিয়ে ধোয়া চালিয়ে যাবেন? | [ 1 Highly confident ] অত্যন্ব আত্মবিশ্বাসী<br>[ 2 Confident ] আত্মবিশ্বাসী<br>[ 3 Less confident ] কম আত্মবিশ্বাসী<br>[ 4 Not confident ] আত্মবিশ্বাসী নই |
| 11 | How confident are you that you can continue washing your hands with soapy water before eating a meal?<br>এ ব্যাপারে আপনি কতটা আত্মবিশ্বাসী যে খাবার খাওয়ার পূর্বে আপনি আপনার হাত সাবান-পানি দিয়ে ধোয়া চালিয়ে যাবেন? | [ 1 Highly confident ] অত্যন্ব আত্মবিশ্বাসী<br>[ 2 Confident ] আত্মবিশ্বাসী<br>[ 3 Less confident ] কম আত্মবিশ্বাসী<br>[ 4 Not confident ] আত্মবিশ্বাসী নই |

| | |
|---|---|
| How likely is it that you will recommend using a handwashing station to your neighbors?<br>এটির সম্ভাবনা কী রকম যে আপনি আপনার প্রতিবেশীর কাছে হাত ধোয়ার স্থান ব্যবহার করার কথা বলবেন? | 1 Very unlikely] সম্ভাবনা খুবই কম<br>[2 Unlikely] সম্ভাবনা কম<br>[3 Likely] সম্ভবত<br>[4 Very likely] খুব সম্ভবত |

| | |
|---|---|
| How likely is it that you will always wash your hands with soap?<br>এটির সম্ভাবনা কী রকম যে আপনি আপনার আপনার হাত সবসময় সাবান দিয়ে ধৌত করবেন? | 1 Very unlikely] সম্ভাবনা খুবই কম<br>[2 Unlikely] সম্ভাবনা কম<br>[3 Likely] সম্ভবত<br>[4 Very likely] খুব সম্ভবত |
| When you run out of soap, how likely is it that your household will buysoap to make soapy water?<br>যখন আপনার সাবান শেষ হয়ে যেতে থাকে এটির সম্ভাবনা কী রকম যে আপনার খানা সাবান-পানি তৈরী করার জন্য সাবান কিনবেন? | 1 Very unlikely] সম্ভাবনা খুবই কম<br>[2 Unlikely] সম্ভাবনা কম<br>[3 Likely] সম্ভবত<br>[4 Very likely] খুব সম্ভবত |

If you plan on washing your hands with soap, what kind of soap will you use?Do Not Read Responses. Mark All that Apply. 1 = Yes
যদি আপনি পরিকল্পনা করে থাকেন যে আপনি সাবান দিয়ে হাত ধৌত করবেন তবে কোন্ ধরনের সাবান ব্যবহার করবেন? **উত্তর পড়ে শোনানো যাবেনা। একাধিক উত্তর**। ১=হ্যাঁ

| | | |
|---|---|---|
| 1 | ¨ | Liquid body/hand soap তরল গায়ে মাখা/হাত ধোয়া সাবান |
| 2 | ¨ | Powdered detergent গুঁড়া পাউডার |

| | | |
|---|---|---|
| 3 | ¨ | Bar soap বার সাবান |
| 4 | ¨ | Dish soap থালা বাসন ধোয়া সাবান |
| 5 | ¨ | Make soapy water সাবান-পানি তৈরী করবো |
| 6 | ¨ | Other অন্যান্য |
| 999 | ¨ | Don't know জানিনা |

| | |
|---|---|
| If my child gets diarrhea it is very serious.<br>যদি আমার শিশুর ডায়রিয়া হয় এটা খুবই গুরম্নতর বিষয়...... | 1 Strongly disagree]<br>দৃঢ়ভাবে ভিন্নমত পোষণ করছি<br>[2 Disagree] ভিন্নমত পোষণ করছি<br>[3 Agree ] একমত<br>[4 Strongly agree]<br>দৃঢ়ভাবে একমত পোষণ করছি |

| | |
|---|---|
| My child's health has not improved even though I wash my hands with soapy water.<br>আমার শিশুর স্বাস্থ্যের কোন উন্নতি হয়নি এমনকি যখন আমি আমার হাত সাবান-পানি দিয়ে ধৌত করতাম তখনও..... | 1 Strongly disagree]<br>দৃঢ়ভাবে ভিন্নমত পোষণ করছি<br>[2 Disagree] ভিন্নমত পোষণ করছি<br>[3 Agree ] একমত<br>[4 Strongly agree]<br>দৃঢ়ভাবে একমত পোষণ করছি |

| | |
|---|---|
| Do you share your handwashing station with your neighbors?<br>আপনি কী আপনার হাত ধোয়ার জিনিসপত্র আপনার প্রতিবেশীর সাথে শেয়ার করবেন? | 1 Yes হ্যাঁ<br>2 No না |

| | |
|---|---|
| Do you find the handwashing station an asset for your household?<br>আপনি কী হাত ধোয়ার জিনিসপত্রকে আপনার খানার একটা সম্পদ মনে করেন? | 1 Yes হ্যাঁ<br>2 No না<br>999 Don't know জানিনা |

| | |
|---|---|
| Has your household social status changed by being known as having a handwashing station?<br>আপনার হাত ধোয়ার জিনিসপত্র আছে এই পরিচিতি কী আপনার খানার সামাজিক অবস্থানকে পরিবর্তিত করেছে? | 1 Our social status increased<br>আমাদের সামাজিক মর্যাদা বৃদ্ধি পেয়েছে<br>2 Our social status decreased<br>আমাদের সামাজিক মর্যাদা কমেছে<br>3 No change in social status<br>আমাদের সামাজিক মর্যাদার কোন পরিবর্তন ঘটেনি |

#### Sanitation
**পয়ঃ ব্যবস্থাপনা**

**How confident are you that…**
**আপনি কতটা আত্মবিশ্বাসী যে.....**

| | | |
|---|---|---|
| 1 | Your household members can empty a full pit?<br>আপনার খানার সদস্যরা একটা পুরো গর্ত খালি করতে পারবে? | [ 1 Highly confident ]<br>অত্যন্ম্ব আত্মবিশ্বাসী<br>[ 2 Confident ] আত্মবিশ্বাসী<br>[ 3 Less confident ]<br>কম আত্মবিশ্বাসী<br>[ 4 Not confident ]<br>আত্মবিশ্বাসী নই |
| 2 | Your household members can switch to another pit?<br>আপনার খানার সদস্যরা অন্য গর্তে স্থানান্ম্বর করতে পারবে? | [ 1 Highly confident ]<br>অত্যন্ম্ব আত্মবিশ্বাসী<br>[ 2 Confident ] আত্মবিশ্বাসী<br>[ 3 Less confident ]<br>কম আত্মবিশ্বাসী<br>[ 4 Not confident ]<br>আত্মবিশ্বাসী নই |
| 3 | The head of your household can hire someone to help switch pits?<br>আপনার খানা প্রধান গর্ত স্থানান্ম্বর করতে অন্য কাউকে ভাড়া করতে পারবে? | [ 1 Highly confident ]<br>অত্যন্ম্ব আত্মবিশ্বাসী |

| | | |
|---|---|---|
| | | [ 2 Confident ] আত্মবিশ্বাসী<br>[ 3 Less confident ]<br>কম আত্মবিশ্বাসী<br>[ 4 Not confident ]<br>আত্মবিশ্বাসী নই |
| 4 | You can keep the latrine slab clean?<br>আপনি পায়খানার স্স্লাব পরিষ্কার রাখতে পারবেন? | [ 1 Highly confident ]<br>অত্যন্ত আত্মবিশ্বাসী<br>[ 2 Confident ] আত্মবিশ্বাসী<br>[ 3 Less confident ]<br>কম আত্মবিশ্বাসী<br>[ 4 Not confident ]<br>আত্মবিশ্বাসী নই |
| 5 | You can buy a new potty?<br>আপনি একটি নতুন পটি কিনতে পারবেন? | [ 1 Highly confident ]<br>অত্যন্ত আত্মবিশ্বাসী<br>[ 2 Confident ] আত্মবিশ্বাসী<br>[ 3 Less confident ]<br>কম আত্মবিশ্বাসী<br>[ 4 Not confident ]<br>আত্মবিশ্বাসী নই |
| 6 | You can help your child use the potty until he/she is old enough to use the latrine?<br>আপনি আপনার শিশুকে পটি ব্যবহারে সাহায্য করতে পারবেন যতদিন পর্যন্ত না আপনার শিশু পায়খানা ব্যবহারের উপযোগী হয়? | [ 1 Highly confident ]<br>অত্যন্ত আত্মবিশ্বাসী<br>[ 2 Confident ] আত্মবিশ্বাসী<br>[ 3 Less confident ]<br>কম আত্মবিশ্বাসী<br>[ 4 Not confident ]<br>আত্মবিশ্বাসী নই |
| 7 | You can teach your next child how to use the potty?<br>আপনি আপনার পরবর্তী শিশুকে পটি কীভাবে ব্যবহার করতে হয় এ ব্যাপারে শিক্ষা দিবেন? | [ 1 Highly confident ]<br>অত্যন্ত আত্মবিশ্বাসী<br>[ 2 Confident ] আত্মবিশ্বাসী<br>[ 3 Less confident ]<br>কম আত্মবিশ্বাসী<br>[ 4 Not confident ]<br>আত্মবিশ্বাসী নই |
| 8 | You can continue to use the saniscoop as to remove feces from your courtyard area?<br>আপনি আপনার উঠান থেকে মল পরিষ্কার করতে সেনিস্কুপ ব্যবহার অব্যাহত রাখবেন? | [ 1 Highly confident ]<br>অত্যন্ত আত্মবিশ্বাসী<br>[ 2 Confident ] আত্মবিশ্বাসী<br>[ 3 Less confident ]<br>কম আত্মবিশ্বাসী<br>[ 4 Not confident ]<br>আত্মবিশ্বাসী নই |
| 9 | You can dispose of your child's feces into the latrine?<br>আপনি আপনার শিশুর মল পায়খানায় ফেলবেন? | [ 1 Highly confident ]<br>অত্যন্ত আত্মবিশ্বাসী<br>[ 2 Confident ] আত্মবিশ্বাসী<br>[ 3 Less confident ]<br>কম আত্মবিশ্বাসী<br>[ 4 Not confident ]<br>আত্মবিশ্বাসী নই |

| If your family does not use an improved latrine…<br>যদি আপনার পরিবার উন্নত পায়খানা ব্যবহার না করে....... | |
|---|---|
| How likely is it that you will get diarrhea?<br>আপনার ডায়রিয়ায় আক্রান্ত হবার সম্ভাবনা কী রকম? | 1 Very unlikely] সম্ভাবনা খুবই কম<br>[2 Unlikely] সম্ভাবনা কম<br>[3 Likely] সম্ভবত<br>[4 Very likely] খুব সম্ভবত |
| How likely is it that your child will get diarrhea?<br>আপনার শিশুর ডায়রিয়ায় আক্রান্ত হবার সম্ভাবনা কী রকম? | 1 Very unlikely] সম্ভাবনা খুবই কম<br>[2 Unlikely] সম্ভাবনা কম<br>[3 Likely] সম্ভবত<br>[4 Very likely] খুব সম্ভবত |

| | |
|---|---|
| It is a problem when my neighbors do not defecate in an improved latrine.<br>আমার প্রতিবেশীরা উন্নত পায়খানা ব্যবহার করেনা এটি একটি সমস্যা..... | 1 Strongly disagree] দৃঢ়ভাবে ভিন্নমত পোষণ করছি<br>[2 Disagree] ভিন্নমত পোষণ করছি<br>[3 Agree ] একমত<br>[4 Strongly agree] দৃঢ়ভাবে একমত পোষণ করছি |

| | |
|---|---|
| If you do not remove feces from your courtyard area…<br>যদি আপনি আপনার উঠান থেকে মল অপসারন না করেন.... | |
| How likely is it that you will get diarrhea?<br>আপনার ডায়রিয়ায় আক্রান্ত হবার সম্ভাবনা কী রকম? | 1 Very unlikely] সম্ভাবনা খুবই কম<br>[2 Unlikely] সম্ভাবনা কম<br>[3 Likely] সম্ভবত<br>[4 Very likely] খুব সম্ভবত |
| How likely is it that your child will get diarrhea?<br>আপনার শিশুর ডায়রিয়ায় আক্রান্ত হবার সম্ভাবনা কী রকম? | 1 Very unlikely] সম্ভাবনা খুবই কম<br>[2 Unlikely] সম্ভাবনা কম<br>[3 Likely] সম্ভবত<br>[4 Very likely] খুব সম্ভবত |

| | |
|---|---|
| It is a problem when my neighbors do not remove feces from their courtyard areas.<br>আমার প্রতিবেশীরা তাদের উঠান থেকে মল অপসারন করে না এটি একটি সমস্যা... | 1 Strongly disagree] দৃঢ়ভাবে ভিন্নমত পোষণ করছি<br>[2 Disagree] ভিন্নমত পোষণ করছি<br>[3 Agree ] একমত<br>[4 Strongly agree] দৃঢ়ভাবে একমত পোষণ করছি |

| | |
|---|---|
| If your child openly defecates on the ground…<br>যদি আপনার শিশু প্রকাশ্যে মাটিতে মল ত্যাগ করে.... | |
| How likely is it that you will get diarrhea?<br>আপনার ডায়রিয়ায় আক্রান্ত হবার সম্ভাবনা কী রকম? | 1 Very unlikely] সম্ভাবনা খুবই কম<br>[2 Unlikely] সম্ভাবনা কম<br>[3 Likely] সম্ভবত<br>[4 Very likely] খুব সম্ভবত |
| How likely is it that your child will get diarrhea?<br>আপনার শিশুর ডায়রিয়ায় আক্রান্ত হবার সম্ভাবনা কী রকম? | 1 Very unlikely] সম্ভাবনা খুবই কম<br>[2 Unlikely] সম্ভাবনা কম<br>[3 Likely] সম্ভবত<br>[4 Very likely] খুব সম্ভবত |

| | |
|---|---|
| It is a problem when my neighbors' children openly defecate on the ground.<br>আমার প্রতিবেশীর সন্তানেরা প্রকাশ্যে মাটিতে মল ত্যাগ করে এটি একটি সমস্যা। | 1 Strongly disagree] দৃঢ়ভাবে ভিন্নমত পোষণ করছি<br>[2 Disagree] ভিন্নমত পোষণ করছি<br>[3 Agree ] একমত<br>[4 Strongly agree] দৃঢ়ভাবে একমত পোষণ করছি |
| If my child gets diarrhea it is very serious.<br>যদি আমার শিশুর ডায়রিয়া হয় এটা খুবই গুরুতর বিষয়...... | 1 Strongly disagree] দৃঢ়ভাবে ভিন্নমত পোষণ করছি<br>[2 Disagree] ভিন্নমত পোষণ করছি<br>[3 Agree ] একমত<br>[4 Strongly agree] দৃঢ়ভাবে একমত পোষণ করছি |
| After using an improved latrine, potty, and saniscoop my child's health has not improved.<br>উন্নত পায়খানা, পটি এবং সেনিস্কুপ ব্যবহার করার পর আমার শিশুর স্বাস্থ্যের উন্নতি হয়নি | 1 Strongly disagree] দৃঢ়ভাবে ভিন্নমত পোষণ করছি<br>[2 Disagree] ভিন্নমত পোষণ করছি<br>[3 Agree ] একমত<br>[4 Strongly agree] দৃঢ়ভাবে একমত পোষণ করছি |

| |
|---|
| What would you do if your saniscoop breaks and there is feces on the ground in the courtyard area? Do Not Read Responses. Mark All that Apply. 1 = Yes |

| যদি আপনার সেনিস্কুপ ভেঙ্গে যায় এবং আপনার উঠানে মল পড়ে থাকে তখন আপনি কী করবেন? **উত্তর পড়ে শোনানো যাবেনা। একাধিক উত্তর।** ১=হ্যাঁ | | |
|---|---|---|
| 1 | ¨ | Try to repair the saniscoop and use it সেনিস্কুপ মেরামত করার চেষ্টা করবো এবং ব্যবহার করবো |
| 2 | ¨ | Call a CHP সিএইচপি'কে ডাকবো |
| 3 | ¨ | Pick up feces with straw খড় ব্যবহার করে মল তুলে ফেলবো |
| 4 | ¨ | Pick up feces with leaves পাতা ব্যবহার করে মল তুলে ফেলবো |
| 5 | ¨ | Use a seni to scoop up the feces মল ছেঁচে তুলে ফেলতে ছেনি ব্যবহার করবো |
| 6 | ¨ | Nothing, leave the feces on the ground কিছুই করবো না, মল উঠানে ফেলে রাখবো |
| 7 | ¨ | Other অন্যান্য |
| 999 | ¨ | Don't know জানিনা |

| | |
|---|---|
| How likely is it that you will share what you learned about child potties with other family members or neighbors? শিশুদের পটি ব্যবহার সম্বন্ধে আপনি যা শিখেছেন তা অন্য পরিবারের সদস্য বা প্রতিবেশীদের কাছে শেয়ার করবেন এর সম্ভাবনা কেমন? | 1 Very unlikely] সম্ভাবনা খুবই কম<br>[2 Unlikely] সম্ভাবনা কম<br>[3 Likely] সম্ভবত<br>[4 Very likely] খুব সম্ভবত |

| Was there ever a time when you were home but could not use your latrine? Do Not Read Responses. Mark All that Apply. 1 = Yes<br>কখনও এমন সময় ছিল যখন আপনি বাড়ীতে কিন্তু আপনি পায়খানা ব্যবহার করেননি? **উত্তর পড়ে শোনানো যাবেনা। একাধিক উত্তর।** ১=হ্যাঁ | | |
|---|---|---|
| 1 | ¨ | It was broken (mark broken features on question C.617)<br>এটি ভেঙ্গে গিয়েছিল (ভাঙ্গা বৈশিষ্ট্য প্রশ্ন C.617 চিহ্নিত করুন) |
| 2 | ¨ | It was too dirty এটি খুবই নোংরা ছিল |
| 3 | ¨ | The pit was full পিট ভর্তি হয়ে গিয়েছিল |
| 4 | ¨ | I could not walk up to the raised slab আমি উঁচু স্লাবে উঠতে পারিনা |
| 5 | ¨ | There was not enough space inside the latrine for me to comfortably use it<br>সহজে ব্যবহার করার মত পায়খানার ভেতরে যথেষ্ট জায়গা ছিলনা |
| 6 | ¨ | It was too dark পায়খানার ভেতরে খুব অন্ধকার ছিল |
| 7 | ¨ | It was too far away পায়খানা অনেক দূরে ছিল |
| 8 | ¨ | There was a queue সেখানে লাইন ছিল |
| 9 | ¨ | I have always been able to use my latrine আমি সবসময় আমার পায়খানা ব্যবহার করতে সৰম হয়েছি |
| 10 | ¨ | Other (specify) অন্যান্য (নির্দিষ্ট করুন) |
| 999 | ¨ | Don't know জানিনা |

| | | |
|---|---|---|
| ¨If you were unable to use your latrine, how long were you unable to use your latrine?<br><br>¨ যদি আপনি আপনার আপনার পায়খানা ব্যবহার করতে সৰম না হয়ে থাকেন তবে আপনি কতদিন পায়খানা ব্যবহার করতে পারেননি? | 1 | Less than 24 hours<br>২৪ঘন্টার কম |
| | 2 | 1-2 days<br>১-২ দিন |
| | 3 | 3-6 days<br>৩-৬দিন |
| | 4 | 1-2 weeks<br>১-২সপ্তাহ |
| | 5 | 2-4 weeks<br>২-৪সপ্তাহ |

¨What did you do to solve the problem? (free response)
¨এই সমস্যা সমাধান করার জন্য আপনি কী করেছিলেন? (খোলা উত্তর)

| | | |
|---|---|---|
| Is there enough space in your latrine to<br>আপনার পায়খানার ভেতরে যথেষ্ঠ জায়গা আছে কী...... | | |
| 1 | Store water?<br>পানি ভরে রাখার জন্য? | [ 1 Yes ] হ্যাঁ<br>[ 2 No ] না<br>[ 88 N/A] জানিনা |
| 2 | Store items for cleaning the latrine?<br>পায়খানা পরিষ্কার করার সামগ্রী রাখার জন্য ? | [ 1 Yes ] হ্যাঁ<br>[ 2 No ] না<br>[ 88 N/A] জানিনা |
| 3 | Comfortably use the latrine?<br>স্বাচ্ছন্দে পায়খানা ব্যবহার করার জন্য? | [ 1 Yes ] হ্যাঁ<br>[ 2 No ] না<br>[ 88 N/A] জানিনা |

| | |
|---|---|
| ¨ Do you feel safe going to your latrine at night?<br>¨ আপনি রাতের বেলায় পায়খানায় যেতে নিরাপদ বোধ করেন? | [ 1 Yes ] হ্যাঁ<br>[ 2 No ] না<br>[ 88 N/A] জানিনা |

| | |
|---|---|
| ¨Can you smell the pit contents?<br>¨ আপনি কী পিটের বিষয়বস্তুর গন্ধ পান? | [ 1 Yes ] হ্যাঁ<br>[ 2 No ] না<br>[ 88 N/A] জানিনা |

| | |
|---|---|
| Do you find the improved latrine an asset for your household?<br>¨ আপনি কী উন্নত পায়খানাকে আপনার খানার একটা সম্পদ মনে করেন? | [ 1 Yes ] হ্যাঁ<br>[ 2 No ] না<br>[ 88 N/A] জানিনা |

| | |
|---|---|
| Do you find the potty an asset for your household?<br>¨ আপনি কী পটিকে আপনার খানার একটা সম্পদ মনে করেন? | [ 1 Yes ] হ্যাঁ<br>[ 2 No ] না<br>[ 88 N/A] জানিনা |

| | |
|---|---|
| Do you find the saniscoop an asset for your household?<br>¨ আপনি কী সেনিস্কুপকে আপনার খানার একটা সম্পদ মনে করেন? | [ 1 Yes ] হ্যাঁ<br>[ 2 No ] না<br>[ 88 N/A] জানিনা |

| | |
|---|---|
| Has your household social status changed by being known as having an improved latrine?<br>আপনার একটি উন্নত পায়খানা আছে এই পরিচিতি কী আপনার খানার সামাজিক অবস্থানকে পরিবর্তিত করেছে? | 1 Our social status increased<br>আমাদের সামাজিক মর্যাদা বৃদ্ধি পেয়েছে<br>2 Our social status decreased<br>আমাদের সামাজিক মর্যাদা কমেছে<br>3 No change in social status<br>আমাদের সামাজিক মর্যাদার কোন পরিবর্তন ঘটেনি |

| | |
|---|---|
| Has your household social status changed by being known as having a child potty?<br>আপনার একটি পটি আছে এই পরিচিতি কী আপনার খানার সামাজিক অবস্থানকে পরিবর্তিত করেছে? | 1 Our social status increased<br>আমাদের সামাজিক মর্যাদা বৃদ্ধি পেয়েছে<br>2 Our social status decreased |

| | |
|---|---|
| | আমাদের সামাজিক মর্যাদা কমেছে<br>3 No change in social status<br>আমাদের সামাজিক মর্যাদার কোন পরিবর্তন ঘটেনি |

| | |
|---|---|
| Has your household social status changed by being known as having a saniscoop?<br>আপনার একটি সেনিস্কুপ আছে এই পরিচিতি কী আপনার খানার সামাজিক অবস্থানকে পরিবর্তিত করেছে? | 1 Our social status increased<br>আমাদের সামাজিক মর্যাদা বৃদ্ধি পেয়েছে<br>2 Our social status decreased<br>আমাদের সামাজিক মর্যাদা কমেছে<br>3 No change in social status<br>আমাদের সামাজিক মর্যাদার কোন পরিবর্তন ঘটেনি |

Ask head of household খানা প্রধানকে প্রশ্ন করুন
If your household has emptied the pit…
যদি আপনার খানা গর্ত খালি করে থাকে….

| | |
|---|---|
| ¨Does your household use the pit contents?<br>¨ আপনার খানা কী পিটের বিষয়বস্তু ব্যবহার করে? | [ 1 Yes ] হ্যাঁ<br>[ 2 No ] না<br>[ 88 N/A] জানিনা |

| |
|---|
| ¨ If yes, how does your household use the pit contents? (free response)<br>যদি হ্যাঁ হয়, কীভাবে আপনার খানা পিট বিষয়বস্তু ব্যবহার করে? (খোলা প্রশ্ন) |

| |
|---|
| ¨ If no, where does your household dispose of the pit contents? (free response)<br>যদি না হয়, আপনার খানা পিট বিষয়বস্তুকে কোথায় ফেলেছিল? (খোলা প্রশ্ন) |

**Additional questions for Toilet Details**
**পায়খানার খুঁটিনাটি বিষয়ে আরো কিছু প্রশ্ন**

**C.617.**

| **Condition of toilet পায়খানার অবস্থা** | | |
|---|---|---|
| 1 | Broken wall around toilet<br>পায়খানার চারপাশের দেয়াল ভাঙ্গা | [ 1 Yes ] হ্যাঁ<br>[ 2 No ] না<br>[ 88 N/A] জানিনা |
| 2 | Broken door/curtain or walls that guarantee privacy around the toilet<br>ভাঙ্গা দরজা/পর্দা বা দেয়াল যা পায়খানার গোপনীয়তার নিশ্চয়তা দেয় | [ 1 Yes ] হ্যাঁ<br>[ 2 No ] না<br>[ 88 N/A] জানিনা |
| 3 | Broken roof over toilet<br>পায়খানার ছাদ ভাঙ্গা | [ 1 Yes ] হ্যাঁ<br>[ 2 No ] না<br>[ 88 N/A] জানিনা |
| 4 | Broken ventilation pipe<br>বায়ু চলাচলের নল ভাঙ্গা | [ 1 Yes ] হ্যাঁ<br>[ 2 No ] না<br>[ 88 N/A] জানিনা |
| 5 | Broken toilet slab<br>পায়খানার স্ল্যাব ভাঙ্গা | [ 1 Yes ] হ্যাঁ<br>[ 2 No ] না<br>[ 88 N/A] জানিনা |
| 6 | Broken delivery pipe<br>ডেলিভারী পাইপ ভাঙ্গা | [ 1 Yes ] হ্যাঁ<br>[ 2 No ] না<br>[ 88 N/A] জানিনা |

| | | |
|---|---|---|
| 7 | Broken footrest in the squatting slab<br>পাদানি ভাঙ্গা | [ 1 Yes ] হ্যাঁ<br>[ 2 No ] না<br>[ 88 N/A] জানিনা |
| 8 | Cracked slab<br>চিড়ধরা স্স্নাব | [ 1 Yes ] হ্যাঁ<br>[ 2 No ] না<br>[ 88 N/A] জানিনা |
| 9 | Broken siphon<br>ভাঙ্গা সাইফোন | [ 1 Yes ] হ্যাঁ<br>[ 2 No ] না<br>[ 88 N/A] জানিনা |
| 10 | Broken or cracked raised soil support of 2nd pit (pit not being used currently)<br>দ্বিতীয় পিটের চারপাশের মাটিতে ফাটল ধরেছে বা সরে গেছে (পিট বর্তমানে ব্যবহার করা হচ্ছে না) | [ 1 Yes ] হ্যাঁ<br>[ 2 No ] না<br>[ 88 N/A] জানিনা |
| 11 | Broken or damaged timber/bamboo bar supporting the wall, door and roof.<br>দেয়াল, দরজা এবং ছাদের জন্য ব্যবহৃত কাঠ/বাঁশের দন্ড ভেঙ্গে গেছে বা নষ্ট হয়ে গেছে | [ 1 Yes ] হ্যাঁ<br>[ 2 No ] না<br>[ 88 N/A] জানিনা |
| 12 | Corrugated iron sheet has been disconnected from bars<br>দন্ডথেকে ঢেউটিন খুলে গেছে | [ 1 Yes ] হ্যাঁ<br>[ 2 No ] না<br>[ 88 N/A] জানিনা |
| 13 | Latrine structure has been tilted aside<br>পায়খানাটি হেলে পড়েছে | [ 1 Yes ] হ্যাঁ<br>[ 2 No ] না<br>[ 88 N/A] জানিনা |

### Nutrition

| | | |
|---|---|---|
| 1 | How confident are you that you can continue to wash your hands with soap while preparing food for your child?<br>এ ব্যাপারে আপনি কতটা আত্মবিশ্বাসী যে আপনি আপনার হাত সাবান দিয়ে ধোয়া চালিয়ে যেতে পারেন যখন আপনি আপনার শিশুর জন্য খাবার তৈরী করেন ? | [ 1 Highly confident ] অত্যন্ত আত্মবিশ্বাসী<br>[ 2 Confident ] আত্মবিশ্বাসী<br>[ 3 Less confident ] কম আত্মবিশ্বাসী<br>[ 4 Not confident ] আত্মবিশ্বাসী নই |
| 2 | How confident are you that you can wash your hands with soap before feeding your child?<br>এ ব্যাপারে আপনি কতটা আত্মবিশ্বাসী যে আপনি আপনার শিশুকে খাওয়ানোর পূর্বে আপনার হাত সাবান দিয়ে ধুতে পারেন ? | [ 1 Highly confident ] অত্যন্ত আত্মবিশ্বাসী<br>[ 2 Confident ] আত্মবিশ্বাসী<br>[ 3 Less confident ] কম আত্মবিশ্বাসী<br>[ 4 Not confident ] আত্মবিশ্বাসী নই |
| 3 | How confident are you that you can include a diverse diet for your children?<br>এ ব্যাপারে আপনি কতটা আত্মবিশ্বাসী যে আপনি আপনার শিশুর জন্য বিভিন্ন ধরনের খাবার অন্তর্ভূক্ত করতে পারেন ? | [ 1 Highly confident ] অত্যন্ত আত্মবিশ্বাসী<br>[ 2 Confident ] আত্মবিশ্বাসী<br>[ 3 Less confident ] কম আত্মবিশ্বাসী<br>[ 4 Not confident ] আত্মবিশ্বাসী নই |
| 4 | How likely is it that you will share what you learned from the CHP with other family members or neighbors?<br>আপনি সিএইচপি'র কাছ থেকে যা শিখেছেন কা অন্য পরিবার বা প্রতিবেশীদের কাছে শেয়ার করবেন এর সম্ভাবনা কী রকম? | 1 Very unlikely] সম্ভাবনা খুবই<br>[2 Unlikely] সম্ভাবনা কম<br>[3 Likely] সম্ভবত<br>[4 Very likely] খুব সম্ভবত |

| | | |
|---|---|---|
| **For your next child:**<br>**পরবর্তী শিশুর জন্য:** | | |
| 1 | How likely is it that you will add Pushtikona/ MoniMix/ other nutrient sprinkle available at the market to your child's meal? | 1 Very unlikely] সম্ভাবনা খুবই কম<br>[2 Unlikely] সম্ভাবনা কম |

| | | |
|---|---|---|
| | পুষ্টিকনা/মনিমিক্স/বা অন্য যেসব পুষ্টিকনা বাজারে পাওয়া যায় তা আপনি আপনার শিশুর খাবারের তালিকায় যোগ করবেন এটার সম্ভাবনা কী রকম? | [3 Likely] সম্ভবত<br>[4 Very likely] খুব সম্ভবত |
| 2 | How confident are you that you or your head of household can buy Pushtikona/ MoniMix/ other nutrient sprinkle available at the market?<br>পুষ্টিকনা/মনিমিক্স/বা অন্য যেসব পুষ্টিকনা বাজারে পাওয়া যায় তা আপনি বা আপনার খানা প্রধান কিনতে পারবেন এ ব্যাপারে আপনি কতটা আশাবাদী ? | [ 1 Highly confident ]<br>অত্যন্ত আত্মবিশ্বাসী<br>[ 2 Confident ] আত্মবিশ্বাসী<br>[ 3 Less confident ]<br>কম আত্মবিশ্বাসী<br>[ 4 Not confident ]<br>আত্মবিশ্বাসী নই |
| 3 | How confident are you that you know when to start adding foods other than breast milk to your child's diet?<br>আপনি এ ব্যাপারে কতটা আত্মবিশ্বাসী যে আপনি জানেন কখন আপনার বাচ্চার খাবারে বুকের দুধ ছাড়া অন্য খাবার যোগ করতে হবে? | [ 1 Highly confident ]<br>অত্যন্ত আত্মবিশ্বাসী<br>[ 2 Confident ] আত্মবিশ্বাসী<br>[ 3 Less confident ]<br>কম আত্মবিশ্বাসী<br>[ 4 Not confident ]<br>আত্মবিশ্বাসী নই |
| 4 | How confident are you that you know how to select from the different food categories to prepare complementary food?<br>আপনি এ ব্যাপারে কতটা আত্মবিশ্বাসী যে আপনি জানেন পরিপূরক খাদ্য তৈরী করার জন্য কীভাবে বিভিন্ন বিভাগ থেকে খাদ্য নির্বাচন করতে হয়? | [ 1 Highly confident ]<br>অত্যন্ত আত্মবিশ্বাসী<br>[ 2 Confident ] আত্মবিশ্বাসী<br>[ 3 Less confident ]<br>কম আত্মবিশ্বাসী<br>[ 4 Not confident ]<br>আত্মবিশ্বাসী নই |
| 5 | How confident are you that you can select the correct amount of food to prepare for your child at different ages?<br>আপনার বাচ্চার জন্য বয়স অনুপাতে সঠিক পরিমান প্রস্তুত খাদ্য নির্ধারন করতে পারেন এ ব্যাপারে আপনি কতটা আত্মবিশ্বাসী? | [ 1 Highly confident ]<br>অত্যন্ত আত্মবিশ্বাসী<br>[ 2 Confident ] আত্মবিশ্বাসী<br>[ 3 Less confident ]<br>কম আত্মবিশ্বাসী<br>[ 4 Not confident ]<br>আত্মবিশ্বাসী নই |

| | |
|---|---|
| If my child gets diarrhea it is very serious.<br>যদি আমার শিশুর ডায়রিয়া হয় এটা খুবই গুরুতর বিষয়...... | 1 Strongly disagree]<br>দৃঢ়ভাবে ভিন্নমত পোষণ করছি<br>[2 Disagree] ভিন্নমত পোষণ করছি<br>[3 Agree ] একমত<br>[4 Strongly agree]<br>দৃঢ়ভাবে একমত পোষণ করছি |
| After taking LNS my child's health has not improved.<br>সোনামনি খাওয়ানোর পরেও আমার শিশুর স্বাস্থ্যের উন্নতি হয়নি। | 1 Strongly disagree]<br>দৃঢ়ভাবে ভিন্নমত পোষণ করছি<br>[2 Disagree] ভিন্নমত পোষণ করছি<br>[3 Agree ] একমত<br>[4 Strongly agree]<br>দৃঢ়ভাবে একমত পোষণ করছি |

| | |
|---|---|
| Do you find the LNS an asset for your household?<br>আপনি সোনামনিকে আপনার খানার সম্পদ হিসেবে দেখেন? | [ 1 Yes ] হ্যাঁ<br>[ 2 No ] না<br>[ 88 N/A] জানিনা |
| Has your household social status changed by being known as feeding your child LNS?<br>আপনি আপনার শিশুকে সোনামনি খাওয়ান এই পরিচিতি কী আপনার খানার সামাজিক অবস্থানকে পরিবর্তন করেছে? | 1 Our social status increased<br>আমাদের সামাজিক মর্যাদা বৃদ্ধি পেয়েছে<br>2 Our social status decreased<br>আমাদের সামাজিক মর্যাদা কমেছে<br>3 No change in social status<br>আমাদের সামাজিক মর্যাদার কোন পরিবর্তন ঘটেনি |

**Wash Benefit Module 12. Home care environment**

#### Measure of the Home Environment

**Now I am interested to know about your home environment and behavior of you and others towards the development of your child.**
□□□ □□□□□ □□□□□ □□□□□□ □□□ □□□□□□□ □□□□□/□□ □□□□ □□□□□□□□□ □□□□□ □□□ □□□□□ □□□□□□□□ □□□ □□□□ □□□□ □□□□ □□□□□ □□□□□ □

**Learning Stimulation**

1.a. How many children's books or picture books do you have for (name)? (If the score is 0, skip Qs- 1.b. go Qs- 2) □□

□□□□□ বাচ্চা□□ (□□□□□ □□□) □□□□□□ □□ □□□ □□□□ □□□□ □□ (□□□□□ □□ □□ □□□□) □□□□□ □□□□□□ □□□?

হ্যাঁ =1, না=0, □□□□ □□ =99

**1.b**. □□□□ □□ □□□□□□□□

□□

**2.** I am interested in learning about the things that (name) plays with when he/she is at home.
"□□□□□ □□□□ (বাচ্চার নাম) □□□□□□ □□□□□□□□□ □□□□ □□ □□□□□ □□□□□ □□□□ □□□□□□ □□ □□□□□ □□□□□ □□ □□□ □□□□□ □□□ □□□□□□",

হ্যাঁ =1, না=0, □□□□ □□ =99

| | | |
|---|---|---|
| a | Homemade toys (such as dolls, cars, or other toys made at home)? ঘরে বানানো এমন কোন খেলনা আছে □□ ? যেমনঃ পুতুল, □□□□ □□ □□□□□□□□ □□□□□ □□ □□□ □□□□ \| | □□ |
| b | Toys from a shop or manufactured toys? দোকানের কেনা খেলনা আছে যা দিয়ে সে খেলতে পারে? *FCI | □□ |
| c | Household objects (such as bowls or pots) or objects found outside (such as sticks, rocks, animal shells or leaves)? □□□□ □□□□□ (□□□□ □□ □□□ )□□ ঘরের □□□□□ যা যা জিনিস পাওয়া যায় (যেমন লাঠি, পাথর,পশুপাখি, শামুক,গাছের পাতা) যা দিয়ে সে ইচ্ছামত খেলতে পারে? | □□ |

| | | |
|---|---|---|
| d | Things which play music or that you use to make music, for example, plastic toy phones for children, toy radio, dolls that sing, drum<br><br>[illegible]? *FCI | □□ |
| e | Things for drawing or writing, for example, pencils, pens, chalk<br><br>[illegible]? *FCI | □□ |
| f | Things for pretending, for example, dolls to play house, tins to pretend to cook<br><br>[illegible] *FCI | □□ |

**3:** Household books, magazines and newspapers ([illegible]):

| No | Question | Code |
|---|---|---|
| a . | How many books do you have in your home right now?<br><br>[illegible] ? *FCI | হ্যাঁ =1<br><br>না=0<br><br>□□□ [illegible]<br><br>[illegible]=99 |
| b. | How many magazines and newspapers do you have in your home right now?<br><br>[illegible] ? *FCI | হ্যাঁ =1<br><br>না=0<br><br>□□□ [illegible]<br><br>[illegible]=99 |

**4:** Sometimes adults taking care of children have to leave the house to go shopping, wash clothes, or for other reasons and have to leave young children. (If the score is 0, skip Qs- 1.b. go Qs- 5)

□□□□□□□□ □□□□□□ □□□□□□□□□□ □□□□□□ □□□ □□□□□ □□□□□□ □□□□□□ □□□□□ □□□□, □□□□□ □□□□□ □□□ □□ □□□□□ □□□□□ □□□□□ □□□□□ □□□□, □□□ □□□□□□ □□□□□ □□ □□□□□□ □□□ □□□□□ □□□□□?

On how many days in the past week was (name) □□ □□□□□□□□ □□□ □□ □□□ □□□□□□?

□□ Days (□□□) হ্যাঁ =1, না=0,

| | | |
|---|---|---|
| a | Left alone for more than an hour?<br>□□ □□□□□□□ □□□□□ □□□ □□□□ □□□□□□□□□? | □□ |
| b | Left in the care of another child, that is, someone less than 10 years old, for more than an hour?<br>□□ □□□□□□□ □□□□□ □□□ □□ □□□ □□ □□□ □□□□□□ □□□□□ □□□□□ □□□□ ? | □□ |

**5.** In the past 3 days, did you or any household member over 15 years of age engage in any of the following activities with (name): গত তিন দিনে আপনি বা বাড়ীর বড় কেউ ( ১৫ বছরের আধিক ব্যক্তি) বাচ্চার সাথে নিম্নলিখিত কাজগুলো করেছেন? হ্যাঁ =1, না=0 I___I

If YES, ask: Who engaged in this activity with (name)? প্রতি কাজের জন্যে হ্যাঁ উত্তরে গোল দাগ দিন

Circle all that apply.

| | Mother/□ □ | Father/□ □□□ | Others/□ □□□ □□□ | Total |
|---|---|---|---|---|
| a. Read books to or looked at picture books with (name)? বাচ্চাকে বই পড়ে শুনিয়েছেন বা ছবির বই দেখিয়েছেন? | | | | |
| b. Told stories to (name)? বাচ্চাকে গল্প বলেছেন। | | | | |
| c. Sang songs to (name) or with (name), including lullabies? বাচ্চার সাথে গান গেয়েছেন ? | | | | |

| | |
|---|---|
| d. | Took (name) outside the home, compound, yard or enclosure?<br>বাচ্চাকে বাড়ীর বাইরে নিয়ে গিয়েছেন ? |
| e. | Played with (name)?<br>বাচ্চার সাথে খেলনা দিয়ে খেলেছেন ? |
| f. | Named, counted, or drew things to or with (name)? বাচ্চাকে সময় দিয়ে কিছু শিখিয়েছেন । (যেমন কোন কিছুর নাম, সংখ্যা গননা, আকাঁ আকিঁ) |

**Spanking ( □□□□ □□ □□□□□ □□□ )**

6. Sometimes kids mind pretty well and sometimes they don't. About how many times

have you spanked your child in the past week?

□□□□□□□□ □□□□ □□□□ □□□ □□□ □□□ □□□□ □□□□ □□□□ □□□□ □□□□ □□□ □□ □□□□□□□ □□□ □□ □□□ □□□□ □□□□□ □□□□□□ □□ □□□□□□□?

বাচ্চা যদি এমন কিছু করে যা আপনি পছন্দ করেননা তখন সাধারনত আপনি কি করেন? □□

যদি মা বলে মার দেই তবে জিজ্ঞাসা করুন □□ □□□□□□□ □□□ □□□□□ □□□□□□□ ?

না=0, □/□ □□□ =1, □-□ □□□=2 □ □□□ □□ □□□ □□□□=3, □□□□ □□=99

**Teaching**

**7.** Some parents spend time teaching their children new skills while other parents believe children learn best on their own. Which of the following best describes your attitude?

□□□□ □□□□ □□□□ □□ □□□□□ □□□□□□□□ □□ □□□□ □□□ □□ □□□□□ □□□□□□ □□□□ □□□ □□□ □□□□ □□□ □□□ □□□ □□□□ □□□□□□ □□□□ □□□□□ □□□ □□□□□ □□□□□. □□□□ □□□□□ □□□□□ □□□□□ □□□□□ □□□□□□ □□□□ □□□□ □□□□ □□□□?

Parent always teaches □□□□-□□□□□□□□□□ □□□□□ □□□□□□□ □□□ □□ □□□□□□ □□□□□□ □□ =1 □□

Parent usually teaches □□□□-□□□□□□□□□ □□□□ □□□□ □□□□□□□ □□□ □□ □□□□□□ □□□□□□ □□ =2

Usually learn on own □□□□□□ □□□□□□ □□□□ □□□□□ □□□ □□ □□□□□□ □□□□ =3

Always learn on own [illegible] =4

Don't know / not sure [illegible] =99

**Other**

**8.** Children seem to demand attention when their parents are busy, doing housework, for example. How often do you talk to your child while you are working?

[illegible]-[illegible], [illegible] ?

Always talking [illegible]=1

Often talking [illegible]= 2

Sometimes [illegible]=3

Rarely talk [illegible]=4

Never talk [illegible]=5

Don't know / not sure [illegible]=99

**9.** Does your child have a regular feeding schedule?

[illegible]?

হ্যাঁ =1, না=0

**10.** Does your child see his/her father or father figure on a daily basis?

[illegible]?

হ্যাঁ =1, না=0

**Wash Benefit Module 19: Maternal Depression**

<u>মানসিক অনুভূতির প্রশ্নাবলী</u>

**CESD**

#### The response should be 0-7

Sometimes we feel sad and unhappy and other times we feel good. I now want to ask you some questions about how you've been feeling this past week. Explain about the past week (e.g. today is Monday so I want you to tell me how you have been feeling in the past week, from last Monday until this morning.

মাঝে মাঝে আমাদের মনটা খুব ভাল থকে আবার কখনো কখনো নিজেকে খুব অসুখী মনে হয়। আমাদের মধ্যে কিছু কিছু মানুষ আছে যারা সব সময় খুব হাসি খুশী থাকে আবার কেউ কেউ সারাক্ষন মন খারাপ করে থাকে। আমারা তাই বিভিন্ন মায়েদের সাথে কথা বলে দেখতে চাই যাতে আমরা এ ব্যাপারে কিছু করতে পারি। এজন্য আপনার সহযোগিতা আমাদের একান্ত দরকার। আমরা আপনার সাথে যে কথা গুলো বলবো সে গুলো সম্পর্কে গোপন রাখা হবে। এগুলো শুধু একটা গবেষণার কাজে ব্যবহার করা হবে। আমরা অনেক আগের কথা মনে রাখতে পারি না। কিন্তু খুব কাছাকাছি সময়ের কথা মনে রাখতে পারি। সে জন্য গত এক সপ্তাহে আপনার মনমানসিকতা কেমন ছিল সে ব্যাপারে আপনার কাছেকিছু জানতে চাইব। **(N.B.** এক সপ্তাহ বলতে আজ যদি সোমবার হয় তবে গত সোমবার সকাল থেকে আজকে সকাল পর্যন্ত সময়ে আপনার মন মানসিকতা কেমন ছিল বা আপনি কেমন বোধ করেছেন বুঝিয়ে বলুন)।

**Q1.** Did anything unusual happen that bothered you during the last week?

If yes how many days? _____________ days

১. আমরা ছোটখাটো বিষয়ে সাধারণত বিরক্ত হইনা তাই না? গত সপ্তাহে এমন কোন দিন গেছে যে ছোটখাটো বিষয়েও আপনি বিরক্ত হয়েছেন? (যদি হ্যাঁ হয় তাহলে কতদিন) /___/ days

**Q2.** Sometimes we don't feel hungry. How many days did you not want to eat anything?_____ days

২. মাঝে মাঝে আমাদের খেতে ইচ্ছে করে না। গত ১ সপ্তাহে এমন কোন দিন গেছে যে আপনার খেতে ইচ্ছে করেনি? (যদি হ্যাঁ হয় তাহলে কতদিন) /___/ days

**Q3.** How many days did you feel so down that nobody could cheer you up? _____________ days

৩. গত সপ্তাহে এমন কোন দিন গেছে যে আপনার নিজেকে এত দুঃখী মনে হয়েছে যে কেউই আপনার দুঃখ কমাতে পারবে না বলে মনে হয়েছে? (যদি হ্যাঁ হয় তাহলে কতদিন) /___/ days

**Q4**. Sometimes we feel that we're no good and other times we feel that we're just as good as everyone else.

How many days in the last week did you feel that you were just as good as other people?_________ days

৪. মাঝে মাঝে আমাদের মনে হয় যে আমাকে দিয়ে ভালো কিছু হবে না, আবার একেক সময় মনে হয় সে আমিও ভলো কিছু করতে পারি বা আমিও কারও চেয়ে কম না।

গত ১ সপ্তাহে আপনার কি কখনো মনে হয়েছে অন্যরা যা পারে আপনিও তা পারেন ? (যদি হ্যাঁ হয় তাহলে কতদিন) /___/ days

**Q5.** How many days could you not do anything attentively? ____________ days

৫. গত ১ সপ্তাহে এমন কোনদিন গেছে যে আপনি কোন কাজেই মন বসাতে পারেন নি? (যদি হ্যাঁ হয় তাহলে কতদিন) /___/ days

**Q6.** How many days did you feel depressed? ____________ days

৬. গত ১ সপ্তাহে এমন কোন দিন গেছে যে আপনি খুব মন মরা বা হতাশ ছিলেন?

(যদি হ্যাঁ হয় তাহলে কতদিন) /___/ days

**Q7.** How many days did you feel that easy work was just too difficult to do? ____________ days

৭ আমাদের জীবনে অনেক কাজ আছে সহজ আবার অনেক কাজ আছে কঠিন। গত ১ সপ্তাহে কি আপনার এমন কোন দিন গেছে যে সহজ কাজও চেষ্টা করে করতে হয়েছে বা কষ্ট কর মনে হয়েছে? /___/ days (যদি হ্যাঁ হয় তাহলে কতদিন)

**Q8.** How many days did you think that the future looks good? ____________ days

৮. গত সপ্তাহে আপনার কখনো কি মনে হয়েছে যে আপনার সামনের দিন গুলিতে ভালো কিছু হওয়ার আশা আছে? /___/ days

**Q9.** How many days did you think your life had been a failure? ____________ days

৯. গত সপ্তাহে এমন কোন দিন আপনার মনে হয়েছে যে আপনি জীবনে কিছুই করতে পারেননি? /___/ days (যদি হ্যাঁ হয় তাহলে কতদিন)

**Q10.** How many days did you feel scared that something bad was going to happen, even though it was not likely? ____________ days

১০. গত ১ সপ্তাহে এমন কোনদিন গেছে যে খারাপ কিছু ঘটতে পারে বলে আশংকা করেছেন বা মনে কোন ভয় কাজ করেছে? (যদি হ্যাঁ হয় তাহলে কতদিন) /___/ days

**Q11.** How many nights did you not sleep well and toss and turn? ____________ nights

১১ গত ১ সপ্তাহে এমন কোন রাত গেছে যে আপনার রাতে ভালো ঘুম হয়নি, শুধু এপাশ ও পাশ করেছেন ?

(যদি হ্যাঁ হয় তাহলে কতদিন) /___/ nights

**Q12.** How many days did you feel happy? ____________ days

১২ গত সপ্তাহে কতদিন আপনার নিজেকে সুখী মনে হয়েছে ? /___/ days

**Q13.** How many days did you have no interest in talking to anyone? ____________ days

১৩ গত ১ সপ্তাহে আপনার এমন কোন দিন গেছে যে কারও সাথে কথা বলতে একেবারেই ইচ্ছে করেনি?

(যদি হ্যাঁ হয় তাহলে কতদিন) /___/ days

**Q14.** How many days did you feel lonely? ____________ days

১৪. গত ১ সপ্তাহে আপনার কখনও একা একা বা নিঃসঙ্গ লেগেছে? (যদি হ্যাঁ হয় তাহলে কতদিন) /___/ days

**Q15.** How many days did you feel that no one cares about you? ____________ days

১৫ গত সপ্তাহে এমন কোন দিন আপনার মনে হয়েছে যে কেউ আপনার খেয়াল করে না বা, আপনার কথা ভাবে না ? (যদি হ্যাঁ হয় তাহলে কতদিন) /___/ days

**Q16.** How many days did you enjoy yourself? ____________ days

১৬. গত সপ্তাহে কতদিন আপনার মনে ফূর্তি ফূর্তি ভাব ছিলো? /___/ days

**Q17.** How many days did you feel like crying? ____________ days

১৭. গত সপ্তাহে কতদিন আপনার খুব কাঁদতে ইচ্ছে করেছে ? /___/ days

**Q18.** How many days did you feel so sad that you had no interest in anything? ________ days

১৮. গত ১ সপ্তাহে এমন কোন দিন গেছে যে আপনার এতই মন খারাপ হয়েছে যার জন্য কোন কিছুতেই আপনি আনন্দ পাননি বা আপনার ভালো লাগেনি ? (যদি হ্যাঁ হয় তাহলে কতদিন) /___/ days

**Q19.** How many days did you feel that people don’t like you? ____________ days

১৯. গত সপ্তাহে আপনার কখনো মনে হয়েছে যে আপনাকে কেউ পছন্দ করে না ? (যদি হ্যাঁ হয় তাহলে কতদিন) /___/ days

**Q20.** How many days could you not get going and didn't feel like moving? _________ days

২০. গত সপ্তাহে আপনার এমন কোন দিন গেছে যে আপনার কোন কিছু করতে ইচ্ছে করেনি ? (যদি হ্যাঁ হয় তাহলে কতদিন) /___/ days

***Comments:***

**Wash Benefit Module 16. Child development**

#### MacArthur Adapted Communicative Development Inventory

##### Words and Gestures (MWG)

**Score : YES=1, NO=0**

| **Very easy** | **Un (Yes=1, no=0)** | **Say (Yes=1, no=0)** |
|---|---|---|
| 1.এদিকে আস/Come here | | |
| 2.এদিকে তাকাও/ Look at me | | |
| 3. **উঠ /** Wake up | | |
| 4. আম্মু আব্বু যাবে /Go to mom or dad | | |
| 5. ঝুনঝুনি /Rattle | | |
| 6. পানি / মাম Water | | |
| 7. দুধ, /Milk | | |
| 8. গ্লাস/Glass | | |

| |
|---|
| 9. বালিশ/Pillow |
| 10. চাঁদ /Moon |
| 11. মা/Mommy |
| 12. বাবা /Daddy |
| 13. **দাদা/দাদী** Grand father or mother |
| 14. বাবু, Baby |
| 15. উকিটুক্কা /Peack-a-boo |
| 16. হাততালি /Clap hand |
| 17. খাওয়া/To eat |
| 18. কামড় দেওয়া/ To bite |
| 19. বসা/ To sit |
| 20. টা-টা/ Bye bye |
| Total very easy |

| **Easy** | **Un (Yes=1, no=0)** | **Say (Yes=1, no=0)** |
|---|---|---|
| 21. ব্যাথা পেয়েছ /Are you hurt | | |
| 22. এটা মাকে দাও/ Give it to mom | | |
| 23. আদর কর জড়িয়ে ধর /Love/hug me | | |
| 24. বাবু হিসু করবে /Want to pass urine | | |
| 25.ধরোনা/Don't touch | | |
| 26.মিয়াও/ Meow | | |
| 27. মুরগী/Chicken | | |
| 28. গাড়ি/Car | | |
| 29. বল/ Ball | | |
| 30. ভাত/ Rice | | |
| 31. বিস্কুট/ Biscuit | | |
| 32. মুড়ি/ Puff.rice | | |
| 33.প্যান্ট/ Pant | | |
| 34. আয়না/ Mirror | | |

| |
|---|
| 35. ময়লা/ Its dirty |
| 36. দাঁতমাজা/Brush teeth |
| 37. বাড়িমারা /To bang |
| 38. খেলা করা/To play |
| 39. হাটা/To walk |
| 40. নাচা/To dance |
| Total Easy |

| **Moderate** | **Un (Yes=1, no=0)** | **Say (Yes=1, no=0)** |
|---|---|---|
| 41. সাপ /Snake | | |
| 42. খিচুড়ি /Khichuri | | |
| 43. মাংশ/ Meat | | |
| 44. কাঁঠাল/Jackfruit | | |
| 45. নুডুলস /Noodles | | |
| 46. চোখ/ Eye | | |
| 47. নাক /Nose | | |
| 48. দাড়ি/ Beard | | |
| 49.রান্নাঘর /Kitchen | | |
| 50. টেবিল /Table | | |
| 51. গোয়াল ঘর/Cowshed | | |
| 52. জগ/Jug | | |
| 53. কলসি /Pitcher | | |
| 54. কেচি/ Scissor | | |
| 55. বাক্স/Box | | |
| 56. দোকান/ Shop | | |
| 57. গাছ/Tree | | |
| 58. স্কুলঘর/ School room | | |
| 59.খোলা/To open | | |
| 60. লেখা /To write | | |
| Total Moderate | | |

| **Difficult words** | **Un (Yes=1, no=0)** | **Say (Yes=1, no=0)** |
|---|---|---|
| 61. সাবধান/ Be careful | | |
| 62. পুতুল খেলা/ Play doll | | |
| 63. সবগুলো, All | | |

| |
|---|
| 64. ছড়া-কবিতা বলা/ Tell ryhmes |
| 65. পাওয়া/ To get |
| 66. চিড়া/ Dreid rice |
| 67. শিয়ালের ডাক/ Hua hu |
| 68. বাঘ/ Tiger |
| 69. কাধ/ ঘাড়, Shoulder |
| 70. একটু পরে/ Little bit later |
| 71. লুকোচুরি/Hide & seek |
| 72. লাটিম/Top (Toy) |
| 73. বৃষ্টি/ Rain |
| 74. আলমারি/ সোকেস, Almira/ showcase |
| 75. ভ্যান গাড়ি/ Van |
| 76. খুন্টি/ Khunti |
| 77. ঔটা/ That |
| 78. আঁকা/ To draw |
| 79. রেডিও/Radio |
| 80. পাশে/ Beside |
| Total score |

| **Very difficult words** | **Un**<br><br>**(Yes=1, no=0)** | **Say**<br><br>**(Yes=1, no=0)** |
|---|---|---|
| 81. জ্যাকেট/Jacket | | |
| 82. কেন /Why | | |
| 83. মহিলা/ Women | | |
| 84. মাঠ/ Field | | |
| 85. ভিতরে/Inside | | |
| 86. মানুষ/Person | | |
| 87. ভ্রু/ Brow | | |
| 88. বাগান/ Garden | | |
| 89. গুটি/ Block | | |
| 90. কেমন/ How | | |
| 91. ফিতা/ Lace | | |
| 92. পরিষ্কার/ Clean | | |
| 93. চি চি/ Chi chi | | |
| 94. দুপুর/Noon | | |

| |
|---|
| 95. তার / Her, His |
| 96. মহিষ/ Buffelo |
| 97. ঘটি-বদনি/Small pitcer |
| 98. রেলগাড়ি/Train |
| 99. গাজর/ Carrot |
| 100. একই/ Same |
| Total very difficult |
| **100.1. Total Scores** |

#### Ages and Stages Qquestionnaire-3

#### *18-19 Months (17 months 16 days through 19 months 15 days)*

***25.7 Communication:***

| **Sl** | **18-19 Months 25.7.23-25.7.29** | **Yes=2** | **Sometimes=1** | **Not yet=0** | **Not applicable=-97** | **Score** |
|---|---|---|---|---|---|---|
| 25.7.1 | Does your child imitate a two-word sentence? For example, when you say a two-word phrase, such as “Mama eat,” “Daddy play,” “Go home,” or “What’s this?” does your child say both words back to you? (MARK “YES” EVEN IF THE CAREGIVER RESPONDS THAT THE WORDS ARE DIFFICULT TO UNDERSTAND.) আপনি ২টি শব্দ দিয়ে □□□ বললে ওকি শুনে শুনে সেই কথাটি বলে? (যেমন মা যায়,বাবা খেলে, বাড়ী যাও,) (ওর কথা বুঝতে কিছুটা অসুবিধা হলেও হবে ।) (Probe- আপনি ওর সামনে কোন কথা বললে ও কি তা অনুকরণ করে?)বাচ্চার বলা বাক্যের একটি উদাহরণ ... 25.7.23 | | | | | |
| 25.7.2 | Does your child say eight or more words in addition to “Mama” or “Dada”? (IF “YES,” ASK THE CAREGIVER TO NAME THE WORDS THE CHILD CAN SAY AND KEEP COUNT TO BE SURE THERE ARE AT LEAST EIGHT. DO NOT SUGGEST OR OFFER WORDS) ওকি ”মা, বাবা” ছাড়া আরো ৮টি বা তার বেশী শব্দ বলতে পারে? (যদি হ্যাঁ হয় তাহলে নামগুলো জিজ্ঞেস করতে হবে এবং গুনতে হবে যেন ৮টি হয়। মাকে উদাহরন দেয়া যাবে না।) (Probe- আপনার বাচ্চা কি কি শব্দ বলতে পারে?) 25.7.24 | | | | | |
| 25.7.3 | Does he point towards a picture in the book, scratch or try to take the picture?ওকি বইয়ের ছবিতে হাত দিয়ে দেখায়, ছবিতে চাপড় দেয় অথবা | | | | | |

| | |
|---|---|
| | ছবিটি নিতে চেষ্টা করে? |
| 25.7.4 | Does he point towards things he want? ও কিছু চাইলে কি সেটা আঙুল দিয়ে দেখায়? |
| 25.7.5 | If you point to a picture of a ball (kitty, cup, hat, etc.) and ask your child, "What is this?" does your child correctly name at least one picture? (HE NEEDS TO NAME ONLY ONE PICTURE CORRECTLY.)কোন □□□□ ছবি দেখিয়ে যদি জিজ্ঞেস করেন " এটি কি" তাহলে কি সে সঠিকভাবে বলতে পারে? (Probe- □□□□□, □□□, □□□□ ইত্যাদি অন্তত একটি ছবি দেখাতে পারলেও হবে।) (Test) 25.7.27 |
| 25.7.6 | Without giving him clues by pointing or using gestures, can your child carry out at least three of these kinds of directions? (PLEASE ASK ABOUT THESE DIRECTIONS ONLY. OBJECTS IN BRACKETS CAN BE SUBSTITUTED WITH THOSE MORE FAMILIAR TO THE CHILD.)<br><br>a. "Put the [TOY] on the table." b. "Close the door."<br>c. "Bring me a [TOWEL]."<br><br>d. "Find your [COAT]." e. "Take my hand." f. "Get your [BOOK]." ওকি নীচের কাজগুলোর মধ্যে কমপক্ষে তিনটি কাজ ইশারা ছাড়া শুধু মুখে বললেই করতে পারে?(Probe- না দেখিয়ে দিয়ে শুধু মুখে বললে ও ছোটখাট কি কি কাজ করতে পারে?) (Test)<br><br>ক. খেলনাটি টেবিলের উপর রাখ ঘ. তোমার জামা খুজে আন<br><br>খ. দরজাটা বন্ধ কর ঙ. আমার হাতটা ধর<br><br>গ. গামছা টা আমাকে দাও চ. তোমার বইটা আন 25.7.28 |
| 25.7.7 | When you ask her to point to her nose, eyes, hair, feet, ears, and so forth, does your child correctly point to at least seven body parts? She can point to part of herself, you, or a doll. ওর চোখ, কান, নাক, চুল, পা ইত্যাদি দেখাতে বললে ওকি কমপক্ষে ৭টি অঙ্গ দেখাতে পারে? (তার নিজের, আপনার অথবা পুতুলের অঙ্গ দেখালেও হবে) (Probe- ও কি কি অঙ্গ দেখাতে পারে? ) (Test) 25.7.29 |
| 25.7.8 | Does your child say two or three words that represent different ideas together, such as "See dog," "Mommy come home," or "Kitty gone"? (DO NOT COUNT WORD COMBINATIONS THAT |

| | |
|---|---|
| | EXPRESS ONE IDEA, SUCH AS “BYE-BYE,” “ALL GONE,” “ALL RIGHT,” AND “WHAT’S THAT?”) Please give an example of your child’s word combinations:ওকি নিজে নিজেই ২ বা ৩ শব্দ দিয়ে কোন কথা বলে? কি কি বলে? (শব্দগুলোতে ভিন্ন ভিন্ন ধারনা থাকতে হবে, যেমন- কুকুরটা দেখ, মা এখানে আস। এক জাতীয় শব্দ হলে হবে না যেমন- টা টা)<br><br>বাচ্চার বলা বাক্যের একটি উদাহরণ ... 25.7.25 |
| 25.7.9 | Without showing him first, does your child point to the correct picture when you say, “Show me the kitty” or ask, “Where is the dog?” (HE NEEDS TO IDENTIFY ONLY ONE PICTURE CORRECTLY.)বাচ্চাকে বলে না দিয়ে বইয়ের কোন ছবির নাম মুখে বললে সেকি ঠিকমত দেখাতে পারে? (যেমন- বিড়ালটি দেখাও , কুকুরটিকোথায় -অন্তত একটি ছবি দেখাতে পারলেও হবে।) (Test) 25.7.26 |
| 25.7.10 | Can he say words like “Me”, “My”, “Mine” and “You” correctly? (At least 2 should be correct)ওকি ”আমি”, ”আমাকে”, ”আমার” এবং ”তুমি” বা এই ধরনের শব্দ বলতে পারে? (কমপক্ষে ২টি সঠিকভাবে বলতে হবে) |
| 25.7.11 | If asked to bring things he knows/uses from inside/outside the house, can he bring that? (Clothing, Ball, Glass etc that are not present at that room)ওকে ওর চেনা/ ব্যবহারের কোন জিনিস অন্য ঘর/বাইরে থেকে নিয়ে আসতে বললে ওকি তা নিয়ে আসতে পারে? (জামা, বল, গ্লাস ইত্যাদি যা এই ঘরে নেই।) |
| 25.7.12 | Can he speak 15 or more words other then “Ma-Baba”? (If yes, then ask those words and count for at least 15. No example should be given to the mother) (Probe- What are the words your child can say?)ওকি ”মা, বাবা” ছাড়া আরো ১৫টি বা তার বেশী শব্দ বলতে পারে? (যদি হ্যাঁ হয় তাহলে নামগুলো জিজ্ঞেস করতে হবে এবং গুনতে হবে যেন ১৫টি হয়। মাকে উদাহরন দেয়া যাবে না।) (Probe- আপনার বাচ্চা কি কি শব্দ বলতে পারে?) |

#### Motor milestone survey form

**খানা নং** (Household ID)

**Will be filled according to ChildID 6-24 months**

Note-1: FRO-র সহায়তায় FRA এই পরিমাপ নিবে। FRAs will take this measurement with the help of FRO.

Note 2: একই খানাতে ৬-২৪ মাস বয়সী একাধিক শিশু থাকলে প্রত্যেকের জন্য আলাদা **প্রশ্নপত্র** পূরণ করুন। Please fill up as many separate questionnaires as many <36 months children are in a household.

**Section 1: Motor milestone development**

The data collector will observe some of the activities of the child listed here and may need to ask the mother/caregiver for assistance. Answer any questions the mother/caregiver has. Observe each item and note in the column next. Clarify items as follows [তথ্য সংগ্রকারী লিস্টে উলেস্নখিত শিশুর কিছু আচরন পর্যবেৰণ করবেন এবং এৰেত্রে মা/পরিচর্যাকারীর সহযোগিতার প্রয়োজন হতে পারে। মা/পরিচর্যাকারীর কোন প্রশ্ন থাকলে উত্তর দিন। প্রত্যেকটি আচরন পর্যবেৰণ করম্নন এবং পরবর্তী কলামে উলেস্নখ করম্নন। আচরনগুলোকে নিম্নলিখিতভাবে ব্যাখ্যা করম্নন]

**No (inability)**, the child tried but failed to perform the test item because it surpassed his or her developmental level **[না (অৰমতা),** শিশুটি চেষ্টা করেছিল কিন্তু পরীৰণীয় আইটেম করে দেখাতে পারে নাই কারণ ইহা তার ক্রম-বিকাশের লেভেলের চেয়ে বেশী]

**Yes**, the child performed the test item according to the specific criteria **[হ্যাঁ,** শিশুটি পরীৰণীয় আইটেম সুনির্দিষ্ট ক্রাইটেরিয়া/নির্ণায়ক অনুযায়ী করে দেখাতে পেরেছে ]

**Refused**, the child was calm and alert but just refused to cooperate **[প্রত্যাখ্যান,** শিশুটি শান্ত এবং সজাগ থাকা স্বত্ত্বেও পরীৰণীয় আইটেম করে দেখাতে অস্বীকার করেছে]

**Unable to test**, the child could not be tested on this milestone because his or her emotional state (drowsiness, fussiness or crying) was interfering with testing, the child was sick or child's caregiver was distraught **[পরীৰণে অৰম,** বাচ্চার মাইলস্টোন পরীৰা করা সম্ভব হয় নাই কারণ তার আবেগ (ঘুমঘুম ভাব, অনিচ্ছা অথবা কান্না) পরীৰণে বাঁধা প্রদান করছিল, বাচ্চা অসুস্থ ছিল অথবা বাচ্চার পরিচর্যাকারী তাড়াহুড়া করছিল]

**Date of achievement**, the date that CHP entered in her record form. CHP records the date as caregiver reported and cross checked during her visit to the household. অর্জনের তারিখ, *সি.এইচ.পি-র রেকর্ড ফরম থেকে প্রাপ্ত তারিখ। পরিচর্যাকারী যে তারিখের কথা উলেস্নখ করবে সি.এইচ.পি তা রেকর্ড করবে এবং খানাতে পরিদর্শনের সময় ক্রসচেক করবে।*

| **পরীৰণীয় আইটেম Test item** | **(A) পরীৰকের রিপোর্ট Examiner Report** | **(B) অর্জনের তারিখ Date of Achievement** |
|---|---|---|
| | 0= No (inability) [না (অৰমতা )]<br>1= Yes [হ্যাঁ] | *Only enter dates for milestones achieved for the first time from the CHP's record form. সি.এইচ.পি-র* |

| | | | | |
|---|---|---|---|---|
| | 222= Unable to test [পরীৰণে অৰম]<br>666= Refused [প্রত্যাখ্যান] | *রেকর্ড ফরম থেকে মাইলস্টোন অর্জনের প্রথম তারিখটি লিপিবদ্ধ করম্নন* | | |
| | Skip column B in baseline and go to next question. | D | M | Y |
| M 101. Sitting without support -10 secs [কারো/কোন কিছুর সহায়তা ছাড়া বসতে পারে (১০ সেকেন্ড)] | 0 1 222 666 | — — | — — | — — |
| M 102. Hands-and-knees crawling -3 steps [হাত ও হাটুর সাহায্যে হামাগুড়ি দিতে পারে (আগে/পরে ৩ ধাপ)] | 0 1 222 666 | — — | — — | — — |
| M 103. Standing with assistance -10 secs [কারো/কোন কিছুর সহায়তায় দাঁড়িয়ে থাকতে পারে (১০ সেকেন্ড)] | 0 1 222 666 | — — | — — | — — |
| M 104. Walking with assistance -5 steps [কারো/কোন কিছুর সহায়তায় হাটঁতে পারে (৫ পা/ ধাপ)] | 0 1 222 666 | — — | — — | — — |
| M 105. Standing alone assistance [কারো/কোন কিছুর সহায়তা ছাড়া একা একা দাড়াঁতে পারে] | 0 1 222 666 | — — | — — | — — |
| M 106. Walking alone crawling- 5 steps [কারো/কোন কিছুর সহায়তায় ছাড়া একা একা হাঁটতে পারে (৫ পা/ ধাপ)] | 0 1 222 666 | — — | — — | — — |
| M 107. Squats without support [কারো/কোন কিছুর সহায়তা ছাড়া হাঁটু ভাজ করে দাঁড়ানো অবস্থা থেকে গোঁড়ালীর উপর বসতে পারে] | 0 1 222 666 | — — | — — | — — |

| | | | | |
|---|---|---|---|---|
| M 108. Balances on right foot with support- 2 secs [কারো/কোন কিছুর সহায়তায় শুধু ডান পায়ের পাতার উপরে দাঁড়িয়ে থাকতে পারে (২ সেকেন্ড)] | 0 1 222 666 | — — | — — | — — |
| M 109. Balances on left foot with support- 2 secs [কারো/কোন কিছুর সহায়তায় শুধু বাম পায়ের পাতার উপরে দাঁড়িয়ে থাকতে পারে (২ সেকেন্ড)] | 0 1 222 666 | — — | — — | — — |
| M 110. Jumps forward - 4 inches [লাফিয়ে চার ইঞ্চি সামনে যেতে পারে] | 0 1 222 666 | — — | — — | — — |
| M 111. Balances on right foot for 2 seconds alone [ডান পায়ের পাতার উপরে একা একা দুই সেকেন্ড দাঁড়িয়ে থাকতে পারে] | 0 1 222 666 | — — | — — | — — |
| M 112. Balances on left foot for 2 seconds alone [বাম পায়ের পাতার উপরে একা একা দুই সেকেন্ড দাঁড়িয়ে থাকতে পারে] | 0 1 222 666 | — — | — — | — — |

#### Tower Test

The child is invited to help the tester build a tower with wooden blocks.

The child is scored in her/his ability to take turns with the tester.

[For this one we need to set a constant number of blocks and trials (make sure to record it, for example 8-10 blocks and two trials). Record the number of times the child respect turns with the tester].

[illegible], “[illegible], [illegible]
[illegible]|” [illegible], “[illegible]|
[illegible]
[illegible]” [illegible]
[illegible]?[illegible]
[illegible]

[illegible] " [illegible] ", [illegible] | [illegible] ,[illegible]

At the beginning, the tester will offer some wooden blocks to the child and will ask "Let's play a game now", then she will say "let's make a tree with these blocks. First I will place a block, then you have to put one the next on the top of it". The tester needs to ensure whether the child understood the instruction clearly and following the rules accordingly during play. Remind the child "Remember you can place the block only when your turn comes after my turn". If the child does not follow the instruction or becomes eager to place all the blocks by him, he will not get any score. If the child can make a tower with eight blocks, s/he will get 4 turns. Tester will record the number of correct turns, where the child followed instructions.

[illegible] ([illegible]) : [illegible] /Correct turns: The child followed instruction and waited until the tester placed her block on the top.

[illegible] ([illegible]) : [illegible] /Wrong turn: If the child breaks the rule and wants to place block before his turn.

| [illegible] | |
|---|---|
| [illegible] (No of correct turns) | \|__\|__\| |
| [illegible] (No of correct turns) | \|__\|__\| |

**A not B test**

The tester presents the child with a board with two shallow wells, one on the right and one on the left. Then s/he hides a treat in one of the wells and covers both wells with opaque cups so that the child cannot see the treat. After distracting the child with a song for 5 seconds, the tester prompts the child to find the treat. The treat is hidden in the same place until the child searches in the correct place on two consecutive trials, after which the treat is hidden in the opposite well. Ten trials are given.The total score represents the number of trials (out of 10) in which the child searched in the correct place for the snack.

□□□□□□□□□□□□□ □□□□□□□ □□□□ □□□□□□ □□ □□□ □ □□ □□□□ □□□□ □□□□ □□□□□ □ □□ □□□□□□□□□, □□□ □□□□□□ □□□ □□□□ □□□ □□□□□□□ □□□□ □□□□□ □□□ □□□□ □□□□□ □□□ □□□□□ □□□□□ □□□□□□□□ □□ □□□□ □□□□ □□□□□ □□ □□□ □□□□□ □□ □□□□ □ □□□ □□□□□ □□□□□□□ □□□ □□□ □□□□ □□□□ □□□□ □□□□ □□□□□□□ Xv□□□ n□□ □□□ □□□□□□□ □□ □□□□□□□□ bv □□□□□ □□□□□ □□□□ □□□□ □□□□ □□□. □□ □□□□ □□□ □□ □□□□ □□□ □□□ □□□□□□ □□ □□□□□□□□ □□□□□ □□□□□□ □□□□ □□□. □□ □□□□□□□□□□□□□ □□□□□□□□□ □□□ □□□□□□□ □□□ □□□□ □□□□□□ □□□□□□□ □□□ □□□□ □□□□□□ □□□ hZ□□□□□ bv □□□□□□ □□ □□ □□□ □□□ □□□□ □□□□ □□□□ □□□□, □□□□ □□□□ □□□□ □□□□□ □□ □□□□□□ □□□ □ □□□□□ □□ □□□ □□□□□□□ □□□□□ □□ □□□□□□□ □□□□□, □□ □□ □□□□□ □□□□□ □□□□ □□□□

□□□ □□□□□□□□□□ /Score Codes : □□□□ □□□□□□□□□□ Correct Trial- n¨vu (Yes)=1, bv (No)=0

□□□□□□□□□ 1 (Trial 1)……………………………………………………………… |__|

□□□□□□□□□ □ (Trial 1) ………………………………………………………………|__|

□□□□□□□□□ □ (Trial 1) ………………………………………………………………|__|

□□□□□□□□□ □ (Trial 1) ………………………………………………………………|__|

□□□□□□□□□ □ (Trial 1) ………………………………………………………………|__|

□□□□□□□□□ □ (Trial 1) ………………………………………………………………|__|

□□□□□□□□□ □ (Trial 1) ………………………………………………………………|__|

□□□□□□□□□ □ (Trial 1) ………………………………………………………………|__|

□□□□□□□□□ □ (Trial 1) ………………………………………………………………|__|

□□□□□□□□□ □□(Trial 1) ………………………………………………………………|__|

Total Score ……………………………………………………………………|__|__|

**25.8 Gross Motor Skills**

| SI | 18-19 Months 25.8.24-25.8.31 | Yes=2 | Sometimes=1 | Not yet=0 | Not applicable = -97 | Score |
|---|---|---|---|---|---|---|
| 25.8.1 | Does your child walk well and seldom fall? একটু আধটু পড়ে গেলেও ওকি ভালোভাবে হাঁটতে পারে? 25.8.24 | | | | | |
| 25.8.2 | Does your child climb on an object such as a chair (OR ROCK, ETC.) to reach something she wants?<br>কোন কিছু করতে বা ধরতে চাইলে ওকি একা একা চেয়ারের উপর উঠতে পারে? (যেমন- কোন খেলনা তাকের উপর থেকে নেয়ার জন্য।) 25.8.25 | | | | | |
| 25.8.3 | When you show him how to kick a large ball [OR OTHER OBJECT], does your child try to kick the ball by moving his leg forward or by walking into it? (IF CHILD ALREADY KICKS A BALL OR OTHER OBJECT, CHECK "YES" FOR THIS ITEM.)<br>একটি বড় বলকে কিভাবে লাথি দিতে হয় যখন ওকে দেখান তখন কি ও এগিয়ে গিয়ে/পা সামনে বাড়িয়ে বলটি লাথি মারে/মারার চেষ্টা করে?(যদি বাচ্চা আগে থেকে কোন জিনিস বা বলকে লাথি মারে তাহলে উত্তর 'হ্যাঁ' হবে।) 25.8.26<br>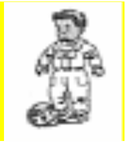 | | | | | |
| 25.8.4 | Does your child walk down stairs (AT LEAST ONE STEP) if you hold onto one of her hands?<br>ওকি ওর এক হাত দিয়ে আপনার হাত ধরে সিঁড়ি (কমপক্ষে সিঁড়ির ১ ধাপ) দিয়ে নামতে পারে? 25.8.27 | | | | | |
| 25.8.5 | Does your child run fairly well, stopping herself without bumping into things or falling কোন কিছুর সাথে ধাক্কা না খেয়ে বা পড়ে না | | | | | |

| | |
|---|---|
| | গিয়ে ওকি ভালভাবে দৌঁড় দিয়ে নিজে নিজেই থামতে পারে? 25.8.28<br>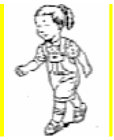 |
| 25.8.6 | Does your child walk **either up or down at least two steps** by himself? (CHECK "YES" EVEN IF HE HOLDS ONTO THE WALL OR RAILING.<br>ওকি নিজে নিজে কমপক্ষে সিঁড়ির দুই ধাপ উঠতে অথবা নামতে পারে? ( দেয়াল বা রেলিং ধরে রাখলেও হ্যাঁ স্কোর পাবে।) 25.8.29<br>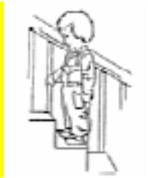 |
| 25.8.7 | Does your child jump with both feet leaving the floor at the same time? □□□ □□□ □□ □□□□ □□□ □□□ □ □□□□□ □□□□ □□□ □□□□ □□□□ ? 25.8.30<br>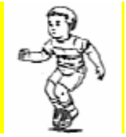 |
| 25.8.8 | Without holding onto anything for support, does your child kick a ball [OR SOME OTHER OBJECT] by swinging his leg forward? □□□ □□□□ □□□□ □□ □□□ □□□□□ □□□□ □□ □□ □□□□ □□□□ □□□□ □□□□ □□□□ ? (Test) 25.8.31<br>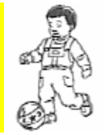 |
| 25.8.9 | Can he, with both feet, jump forward at least 3 inches (Show the measurement with hand)? (Probe- Can the child jump forward?) □□□ □□□□<br>□□□□ /□□□ □□ □□□□ □□□ □□□ □□□□□ □<br>□□□□□ □□□□□□ □□□□ □□□ □□□□ □□□□ ? |

| | |
|---|---|
| | 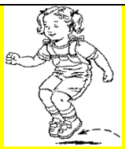 |
| 25.8.10 | Can he, without holding anything, fold his knees, pick up anything from the floor and stand up again (By himself)? ওকি কোন কিছু না ধরে হাঁটু ভাজ করে বসে মেঝে থেকে কোন খেলনা তুলে আবার (নিজে নিজে) দাঁড়িয়ে যায়? |
| 25.8.11 | Can he climb up steps? (Taking support of wall or railing, right foot on one step, then left on the other-in this way) □□□ □□ □□ □□□□ □□□□□□□ □□□□ □□□□ □□□□ □□□□? (দেয়াল বা রেলিং ধরে □□□ □□ □□ □□□□□□□ □□□ □□ □□ □□□□□□□ □□□□□□ ) |
| 25.8.12 | Without holding anything, can he swing his feet from back to front and kick the ball?ওকি কোন কিছু না ধরে □□□ □□ □□□□ □□□□ □□□□□ □□□ (swing) □□□ □□□□ □□□□□ □□□□? |

#### 25.9 Personal- Social

| SI | 18-19 Months 25.9.23-25.9.30 | Yes=2 | Sometimes=1 | Not yet=0 | Not applicable= -97 | Score |
|---|---|---|---|---|---|---|
| 25.9.1 | While looking at himself in the mirror, does your child offer a toy to his own image? □□□□□ □□□□□□ □□□□□ □□□ □□□ □□□□□□□ □□□□□ □□□□ □□□□□ □□□□□□□ □□□□ □□□? (□□□□□ □□□□ □□□□□ □□□□ □□□□□□ 'হ্যাঁ' □□□□□ □□□□ ) (Test) 25.9.23 | | | | | |
| 25.9.2 | Does your child come to you when he needs your help with something, such as [WINDING UP A TOY] or opening something [A BOTTLE, BANANA, ORANGE, SOME OTHER KIND OF FOOD]? □□□ □□□ □□□□ □□□□□□□ □ □□ □□□□ □□□□□ □□□□ □□□□□ □□□□ □□□□□□□□□ □□□□ □□□? □□□□: | | | | | |

| | |
|---|---|
| | □□□□, □□□, □□□□, □□ □□□□ □□□□ □□□□□? 25.9.24 |
| 25.9.3 | Does your child copy the activities you do, such as [WASH DISHES, COOK, PREPARE FOOD, CARE FOR ANIMALS, ETC. OR…] wipe up a spill, sweep, shave, or comb hair? [NOTE: CHILDREN DO NOT HAVE TO DO THE ACTIVITIES PERFECTLY. THE POINT IS THEY TRY TO COPY THEM.]আপনি যা করেন ওকি তা অনুকরন/নকল করে? যেমন- ছলকে পড়ে যাওয়া পানি মোছা, ঝাড়ু দেয়া, সেভ করা অথবা চুল আঁচড়ানো। 25.9.25 |
| 25.9.4 | Does your child drink without help from a cup or glass, putting it down again with little spilling? [CHILD CAN DRINK FROM A CUP WITHOUT SPILLING TOO MUCH]<br><br>ওকি নিজেই গ্লাস বা কাপ থেকে দুধ খেয়ে তা নিচে রাখতে পারে? (সামান্য একটু পড়ে গেলেও হবে) 25.9.26 |
| 25.9.5 | Is he able to feed himself with a spoon or by his hands (without spilling)? (If the answer is yes, ask what does he eat?)ও কি নিজের হাতে বা চামচ দিয়ে খাবার (না ফেলে) খেতে পারে? (যদি হ্যাঁ বলে, তাহলে জিজ্ঞেস করুন কি কি খায়? ) |
| 25.9.6 | If you do any of the following gestures, does your child copy at least one of them? (GESTURES MUST BE THESE EXACTLY; DO NOT SUBSTITUTE OTHER GESTURES.)<br><br>a. Open and close your mouth.<br><br>b. Blink your eyes.<br><br>c. Pull on your earlobe.<br><br>d. Pat your cheek<br><br>এই কাজগুলি আপনি করে দেখালে ওকি করতে পারে? (কমপক্ষে ১টি পারতে হবে) (Test)<br><br>ক. মুখ হা এবং বন্ধ করা খ. চোখ পিট পিট করা<br><br>গ. কানের লতি টানা ঘ. গালে চাপড় দেওয়া<br>25.9.29 |

| | |
|---|---|
| 25.9.7 | Does your child push a little shopping cart, stroller, or wagon, [OR OTHER OBJECT WITH WHEELS] steering it around objects and backing out of corners if he cannot turn?<br>ও কি কোন ছোট ঠেলার গাড়ি বিভিন্ন জিনিসের চারদিকে ঘুরিয়ে খেলতে পারে এবং কোন কোণাতে ঘুরাতে না পারলে কি সে গাড়ীটা পেছনে টেনে নিয়ে আসে? (Test) 25.9.30 |
| 25.9.8 | Is he able to feed himself with a spoon (some spilling acceptable)?খাবার পড়ে গেলেও (ফেলে ছেড়ে) ও কি চামচ দিয়ে নিজে নিজে খেতে পারে? |
| 25.9.9 | Does he hug human or animal like stuff toys?ও কি মানুষ/ প্রানী জাতীয় নরম পুতুল জড়িয়ে ধরে খেলে? |
| 25.9.10 | Does he try to do the same when you take off his shoes, socks, caps?জুতা, মোজা, টুপি খোলার সময় ওকি আপনার সাথে সাথে খুলতে চেষ্টা করে? |
| 25.9.11 | To grab your attention or to show you something, does he pull your hand or clothing?আপনি যাতে ওর দিকে খেয়াল করেন অথবা যদি আপনাকে কিছু দেখাতে চায় তখন কি আপনার হাত বা জামা ধরে টানে? |
| 25.9.12 | When playing with either a stuffed animal or doll [OR ITEM REPRESENTING THIS], does your child pretend to rock it, feed it, change its diapers, put it to bed, and so forth? যখন ও নরম পুতুল বা প্রানী জাতীয় খেলনা নিয়ে খেলে তখন কি সে পুতুলকে দোলানোর, খাওয়ানোর, কাপড় বদলানোর অথবা ঘুম পাড়ানো ইত্যাদির অভিনয় করে খেলে? 25.9.27 |

**Wash benefit Module 21. Maternal intelligence**

**DIGITS BACKWARD**

READ EACH DIGIT SPAN **ONLY ONCE** AT AN EVEN RATE OF 1 DIGIT PER SECOND.

READ PART A OF QUESTION; PAUSE FOR RESPONSE, THEN SCORE.

READ PART B OF QUESTION; PAUSE FOR RESPONSE, THEN SCORE.

IF MOTHER DOES NOT RESPOND, DO **NOT** ENCOURAGE FURTHER.

STOP WHEN CHILD MISSES PART A **AND** PART B OF ANY **ONE** QUESTION.

1. [illegible]

2. [illegible] A [illegible]

3. [illegible] B [illegible]

4. [illegible]

5. [illegible] A [illegible] B [illegible]

Now I am going to say some numbers, but when I stop, I want you to say them backward. For example, if I say 9-2-7, what would you say? I said 9-2-7, so to say it backward you would say 7-2-9. Now try these numbers. Remember, you are to say them backward: 3-6-5.

[illegible]

| | Correct ([illegible])=1 | Wrong ([illegible])=0 |
|---|---|---|
| 1A. 2–5<br>1A. 2 - 5 | | |
| 1B. ৬-৩ | | |

| |
|---|
| 1B. 6 - 3 |
| *IF BOTH A & B ARE CODED WRONG, STOP TEST.*<br>□□□ 1A □□□ 1B □□□□□□ □□□ □□ □□□ □□□□□□ □□ □□□□□ □□□□□ □□□ |
| 2A. ৫-৭-৪<br>2A. 5 - 7 - 4 |
| 2B. ২-৫-৯<br>2B. 2 - 5 – 9 |
| *IF BOTH A & B ARE CODED WRONG, STOP TEST.*<br>□□□ 2A □□□ 2B □□□□□□ □□□ □□ □□□ □□□□□□ □□ □□□□□ □□□□□ □□□ |
| 3A. ৭-২-৯-৬<br>3A. 7 - 2 - 9 - 6 |
| 3B. ৮-৪-৯-৩<br>3B. 8 - 4 - 9 – 3 |
| *IF BOTH A & B ARE CODED WRONG, STOP TEST.*<br>□□□ 3A □□□ 3B □□□□□□ □□□ □□ □□□ □□□□□□ □□ □□□□□ □□□□□ □□□ |
| 4A. ৪-১-৩-৫-৭<br>4A. 4 - 1 - 3 - 5 – 7 |
| 4B. ৯-৭-৮-৫-২<br>4B. 9 - 7 - 8 - 5 – 2 |

| |
|---|
| *IF BOTH A & B ARE CODED WRONG, STOP TEST.*<br>□□□ 4A □□□ 4B □□□□□□ □□□ □□ □□□ □□□□□□ □□ □□□□□ □□□□□ □□□ |
| 5A. ১-৬-৫-২-৯-৮<br>5A. 1 - 6 - 5 - 2 - 9 – 8 |
| 5B. ৩-৬-৭-১-৯-৪ |

| |
|---|
| 5B. 3 - 6 - 7 - 1 - 9 – 4 |
| *IF BOTH A & B ARE CODED WRONG, STOP TEST.*<br><br>□□□ 5A □□□ 5B □□□□□□ □□□ □□ □□□ □□□□□□ □□ □□□□□ □□□□□ □□□ |
| 6A. ৮-৫-৯-২-৩-৪-২<br><br>6A. 8 - 5 - 9 - 2 - 3 - 4 – 2 |
| 6B. ৪-৫-৭-৯-২-৮-১<br><br>6B. 4 - 5 - 7 - 9 - 2 - 8 – 1 |
| *IF BOTH A & B ARE CODED WRONG, STOP TEST.*<br><br>□□□ 6A □□□ 6B □□□□□□ □□□ □□ □□□ □□□□□□ □□ □□□□□ □□□□□ □□□ |
| 7A. ৬-৯-১-৬-৩-২-৫-৮<br><br>7A. 6 - 9 - 1 - 6 - 3 - 2 - 5 – 8 |
| 7B. ৩-১-৭-৯-৫-৪-৮-২<br><br>7B. 3 - 1 - 7 - 9 - 5 - 4 - 8 – 2 |

**Wash Benefit Module 20. Environmental enteropathy subsample**

EE Sample Collection Form – Section 1: Child Identification & Medical History

**Cluster ID |__|__|__| Mother ID |__|_**

| | | |
|---|---|---|
| 1. Cluster ID and Mother ID | ### and ## | \|__\|__\|__\| **and** \|__\|__\| |
| 2. ID of MT/FRA/FRO | ## | \|__\|__\| |
| 3. Name of MT/FRA/FRO (choose 1): | 1. Balay Chand Sikder<br>2. Mary Marget<br>3. ParimolSarker<br>4. Dipali Rani Das<br>5. SaniaAkter Dolly<br>6. SyedaFatemaKhatun<br>7.Md. AltafurRahman<br>8. AmalSarker<br>9. Nasrin Sultana<br>10. Md. NurulAlam<br>11. Fatema Begum Sumi<br>12. SyedaLuthfaFamida | |

| | | |
|---|---|---|
| | 13. Shahjahan Ali<br>16. NitaiHalder | |
| 4. Date of Data Collection | DD/MM/YYYY | \|__\|__\| / \|__\|__\| / \|__\|__\|__\|_ |
| 5. Consent<br><br>Note: If consent given, skip next question. If no consent, answer Q6 and skip to end. | Y/N | \|__\| |
| 6. Why Was Consent Not Given? | 1. Migration<br>2. Household head Is not home<br>3. Primary caregiver Is not home<br>4. Child is not home<br>5. Child is sick<br>6. Household head refused<br>7. Primary caregiver refused<br>8. Too busy<br>9. No reason given<br>10. Other | \|__\|__\| |
| 7. Verify Primary Caregiver's Identity: Does Caregiver Match Household and Cluster ID? | Y/N | \|__\| |

| |
|---|
| Note: If no, then skip rest of form and alert SFRO that our team is in the wrong location. |

EE Sample Collection Form – Section 1: Child Identification & Medical History

**Cluster ID |__|__|__| Mother ID |__|_**

<table>
<tr><td>8. Identification of Primary Caregiver</td><td>1. Child’s Mother<br>2. Child’s Maternal Grandparents<br>3. Child’s Paternal Grandparents<br>4. Child’s Maternal Aunt/Uncle<br>5. Child’s Paternal Aunt/Uncle<br>6. Child’s Father (And His New Partner/Wife)<br>7. Other: Specify</td><td>|__|</td></tr>
<tr><td>9. Mobile Number of Household Head</td><td>###########<br>99. Don’t know/not available</td><td>|__|__|__|__|__|__|__|__|__|__|__<br>|__|__|</td></tr>
<tr><td>10. Mobile Number of Primary Caregiver</td><td>###########<br>99. Don’t know/not available</td><td>|__|__|__|__|__|__|__|__|__|__|__<br>|__|__|</td></tr>
<tr><td>11. Number of Eligible Children</td><td># of Eligible Children</td><td>|__|</td></tr>
<tr><td>12. Child Full Name</td><td></td><td></td></tr>
<tr><td>13. Child Nickname</td><td></td><td></td></tr>
<tr><td>14. Gender of Child</td><td>1. Male<br>2. Female</td><td>|__|</td></tr>
</table>

| | | |
|---|---|---|
| 15. Date of Birth | DD/MM/YYYY | \|__\|__\| / \|__\|__\| / \|__\|__\|__\|__\| |
| 16. Source of Date of Birth (DOB) | 1. Confirmed DOB by valid vaccination card<br>2. Mother/Relative remember DOB<br>3. Both 1 & 2<br>99. Don't know | \|__\|__\| |
| 17. Age | ### Days | \|__\|__\|__\| Days |
| 18. ASK: "How long ago did [CHILD NAME] take any antibiotics? (Provide examples from the 'List of Common Antibiotics and Medicine' in the EE Sampling Handbook) | ## Days<br>## Months<br>88. Never<br>99. Don't know | \|__\|__\| Days<br>\|__\|__\| Months<br>\|__\|__\| |
| 19. ASK: "Can you tell us how many times in the last 3 months this child has used antibiotics?"<br>Note: If answer to Q19 is 0, 88, or 99, then skip to Q22. If answer to Q19 is>0, then answer Q20 and Q21. | ## Times<br>88. Never<br>99. Don't know | \|__\|__\| Times<br>\|__\|__\| |

EE Sample Collection Form – Section 1: Child Identification & Medical History

**Cluster ID |__|__|__| Mother ID |__|_**

| | | |
|---|---|---|
| | 20. ASK: "Please try and recall the name of each antibiotic this child used in the last 3 months (90 days before collection date) | 21. A this antibiotic?" (For Each Episode) |
| | 1. Cotrimoxazole<br>2. Amoxycillin<br>3. Flucloxacillin<br>4. Ciprofloxacin<br>5. Erythromycin<br>6. Azythromycin<br>7. Nalidixic acid<br>8. Doxycycline<br>9. Betapen (Penicillin)<br>10. Chloramphenicol<br>11. Metronidazole<br>12. Other: Specify<br>99. Don't know | ## Days<br>99. Don't Know |
| Episode 1 | \|__\|__\| (choose from list above) | \|__\|__\| Days |

| | | |
|---|---|---|
| Episode 2 | \|__\|__\| (choose from list above) | \|__\|__\| Days |
| Episode 3 | \|__\|__\| (choose from list above) | \|__\|__\| Days |
| Episode 4 | \|__\|__\| (choose from list above) | \|__\|__\| Days |
| Episode 5 | \|__\|__\| (choose from list above) | \|__\|__\| Days |
| 22. ASK: “Has this child had malaria in the last 1 month?” | 1. Yes, diagnosed in a clinic/by a physician<br>2. Yes, not diagnosed in a clinic/ by a physician<br>3. No<br>4. Not applicable<br>99. Don’t know | \|__\|__\| |
| 23. ASK: “Has this child had malaria in the last 6 months?” | 1. Yes, diagnosed in a clinic/by a physician<br>2. Yes, not diagnosed in a clinic/by a physician<br>3. No | \|__\|__\| |

| | |
|---|---|
| | 4. Not applicable<br><br>99. Don’t know |

EE Sample Collection Form – Section 1: Child Identification & Medical History

**Cluster ID |__|__|__| Mother ID |__|_**

| | | |
|---|---|---|
| 24. ASK: FOR CHILDREN 6 MONTHS AND OVER "In the last 6 months, has [CHILD NAME] received a pill or drug for intestinal worms?" (Provide examples from the 'List of Common Antibiotics and Medicine' in the EE Sampling Handbook)<u>Note:</u> If child is <6 months old, enter 3 and skip to end. If answer to Q24 is 2 or 99, then skip Q25-27 and answer Q 28-29. | 1. Yes<br>2. No<br>3. Not applicable<br>99. Don't know | \|__\|__\| |
| 25. ASK: "Where did [CHILD NAME] receive the drug for intestinal worms?" | 1. At Home / In Village<br>2. At A Clinic or Health Facility<br>3. At A School<br>4. From icddr,b<br>99. Don't know | \|__\|__\| |
| 26. ASK: "Approximately how long ago did [CHILD NAME] receive the deworming drug?" | ## Days<br>## Months<br>99. Don't know | \|__\|__\| Days<br>\|__\|__\| Months<br>\|__\|__\| |

| | | |
|---|---|---|
| 27. ASK: "Please try and recall the name of the deworming drug [CHILD NAME] took." (Provide examples from the 'List of Common Antibiotics and Medicine' in the EE Sampling Handbook) | 1. Albendazole<br>2. Mebendazole<br>3. Other: Specify<br>99. Don't know | \|__\|__\| |
| 28. ASK: "Does [CHILD NAME] typically wear shoes/socks/sandals?" | 1. Yes<br>2. No<br>99. Don't know | \|__\|__\| |
| 29. OBSERVE: Is child wearing shoes/socks/sandals? | 1. Yes<br>2. No<br>99. Don't know | \|__\|__\| |

EE Sample Collection Form – Section 1: Child Identification & Medical History

**Cluster ID |__|__|__| Mother ID |__|__|**

**ASK: "Thank you. Now I am going to ask you some questions about the health of your [child/children]. Please answer each question as accurately as you can. If you don't know the answer to a question, say "I don't know". We will begin with [NAME OF CHILD]."**

| ASK: "Did [NAME OF CHILD] have [SYMPTOM]: | A | B | C | D | E |
|---|---|---|---|---|---|
| | **Today**<br><br>1. Yes<br>2. No<br>99. Don't know | **Yesterday**<br><br>1. Yes<br>2. No<br>99. Don't know | **Day before Yesterday**<br><br>1. Yes<br>2. No<br>99. Don't know | **In the last 7 days** (since this day last week)<br><br>1. Yes<br>2. No<br>99. Don't know | **In the last 2 weeks** (since this day 2 weeks ago)<br><br>1. Yes<br>2. No<br>99. Don't know |
| **30. Fever** | | | | | |
| **31. Diarrhea** | | | | | |
| **32. 3 or more bowel movements in 24 hours** | | | | | |
| **33. Number of bowl movements each day** | \|__\|__\|times | \|__\|__\|times | \|__\|__\|times | | |
| **34. Watery or soft stool (unformed)** | | | | | |
| **35. Blood in the stool** | | | | | |
| **36. Skin rash (anywhere on the body)** | | | | | |
| **37. Constant cough** | | | | | |

| |
|---|
| **38. Congestion / runny nose** |
| **39. Panting / wheezing / difficulty breathing** |
| **40. Bruising, scrapes or cuts** |
| **41. Toothache / teething** |

| | | | |
|---|---|---|---|
| 42. ASK: "How long ago did [CHILD NAME] have diarrhea?"<br><br>Note: Q42 will be asked if Q31 turns out yes for column D and/or E (ie, Q42 will not be asked for col A, B and C of Q31. Also validate Q42 by number of days. For example, if Q42 is asked for col D, the validation is not more than 7 | ## Days<br><br>77. Not applicable<br><br>99. Don't know | D<br><br>**In the last 7 days** (since this day last week)<br><br>\|__\|__\| Days | E<br><br>**In the last 2 weeks** (since this day 2 weeks ago)<br><br>\|__\|__\| Days |
| 43. ASK: "During that specific episode of diarrhea, how many consecutive days did it last?"<br><br>Note:Q43 is linked to Q42, i.e., Q43 will be asked only when Q42 is asked. Also validate Q43 by number of days. For example, if Q43 is asked for col D, [illegible] | ## Days<br><br>77. Not applicable<br><br>99. Don't know | D<br><br>**In the last 7 days** (since this day last week)<br><br>\|__\|__\| Days | E<br><br>**In the last 2 weeks** (since this day 2 weeks ago)<br><br>\|__\|__\| Days |

Cluster ID |__|__|__| Mother ID |__|__|

EE Sample Collection Form – Section 2: Anthropometry

<table>
<tr><td>1. Cluster ID and Mother ID</td><td>### and ##</td><td>|__|__|__| and |__|__|</td></tr>
<tr><td>2. ID of MT/FRA/FRO</td><td>##</td><td>|__|__|</td></tr>
<tr><td>3. Name of MT/FRA/FRO (choose 1):</td><td>4. Dipali Rani Das<br>5. SaniaAkter Dolly</td><td></td></tr>
<tr><td>4. Date of Data Collection</td><td>DD/MM/YYYY</td><td>|__|__| / |__|__| / |__|__|__|__|</td></tr>
<tr><td>5. Number of Eligible Children</td><td># of Eligible Children</td><td>|__|</td></tr>
<tr><td>6. Child Full Name</td><td></td><td></td></tr>
<tr><td>6A. Type of Scale Used for Weight Measurement</td><td>1. Seca 383 (Infant Scale)<br>2. Seca 874 (Mother-Infant Scale)<br>3. Seca 354 (Infant Scale New)</td><td>|__|</td></tr>
<tr><td>7. Weight of Child – Measurement #1</td><td>Weight (kg)##.###</td><td>|__|__|.|__|__|__| kg</td></tr>
</table>

| | | |
|---|---|---|
| 8. Weight of Child – Measurement #2 | Weight (kg) ##.### | \|__\|__\|.\|__\|__\|__\| kg |
| 9. Weight of Child – Measurement #3 | Weight (kg) ##.### | \|__\|__\|.\|__\|__\|__\| kg |
| 10. **Median** Weight of The Child | Weight (kg) ##.### | \|__\|__\|.\|__\|__\|__\| kg |
| 11. Was Child Wearing Clothing During Weight Measurement?<br>(Multiple answers) | 1. No clothes<br>2. Underpants/shorts<br>3. Shirt<br>4. Pants/trousers<br>5. Dress/frock<br>6. Other: Specify | \|__\|<br>\|__\|<br>\|__\|<br>\|__\| |

EE Sample Collection Form – Section 2: Anthropometry

**Cluster ID |__|__|__| Mother ID |__|__|**

| | | |
|---|---|---|
| 12. Length or Height?<br><br>(If ≥ 2 years of age, measure the child's height; if < 2 years of age, measure the child's length) | 1. Length measured<br><br>2. Height measured | \|__\| |
| 13. Length/Height of Child – Measurement #1 | Length/Height (cm)<br><br>###.# | \|__\|__\|__\|.\|__\| cm |
| 14. Length/Height of Child – Measurement #2 | Length/Height (cm)<br><br>###.# | \|__\|__\|__\|.\|__\| cm |
| 15. Length/Height of Child – Measurement #3 | Length/Height (cm)<br><br>###.# | \|__\|__\|__\|.\|__\| cm |
| 16. Head Circumference of Child – Measurement #1 | Circumference (cm)<br><br>##.# | \|__\|__\|.\|__\| cm |
| 17. Head Circumference of Child – Measurement #2 | Circumference (cm)<br><br>##.# | \|__\|__\|.\|__\| cm |
| 18. Head Circumference of Child – Measurement #3 | Circumference (cm)<br><br>##.# | \|__\|__\|.\|__\| cm |

| | | |
|---|---|---|
| 19. Mid Upper Arm Circumference of Child – Measurement #1 | Circumference (cm)<br>##.# | \|__\|__\|.\|__\| cm |
| 20. Mid Upper Arm Circumference of Child – Measurement #2 | Circumference (cm)<br>##.# | \|__\|__\|.\|__\| cm |
| 21. Mid Upper Arm Circumference of Child – Measurement #3 | Circumference (cm)<br>##.# | \|__\|__\|.\|__\| cm |
| 22. Anthropometry Data Obtained?<br>Note: If Q22 is 1, skip to end. | 1. All<br>2. Partial<br>3. None | \|__\| |
| 23. Why Was Anthropometry Data Not Collected? | 1. Child Did Not Cooperate<br>2. Household Head Refused<br>3. Primary Caregiver Refused<br>4. Other | \|__\| |

EE Sample Collection Form – Section 3: Blood Sample Collection

**Cluster ID |__|__|__| Mother ID |__|__|**

| | | |
|---|---|---|
| 1. Cluster ID and Mother ID | ### and ## | \|__\|__\|__\| and \|__\|__\| |
| 2. ID of MT/FRA/FRO | ## | \|__\|__\| |
| 3. Name of MT/FRA/FRO (choose 1): | 1. Balay Chand Sikder<br>2. Mary Marget<br>3. ParimolSarker<br>14. Md. Shariful Islam<br>15. Shimul Das<br>16. NitaiHalder | |
| 4. Date of Sample Collection | DD/MM/YYYY | \|__\|__\| / \|__\|__\| / \|__\|__\|__\|__\| |
| 5. Number of Eligible Children | # of Eligible Children | \|__\| |

| | | |
|---|---|---|
| 6. Child Full Name | | |
| 7. Date of Child's Last Meal/Snack/Breastfeed | DD/MM/YYYY | \|__\|__\| / \|__\|__\| / \|__\|__\|__\|__\| |
| 8. Time of Child's Last Meal/Snack/Breastfeed | 24-hour scale<br>Hours :Minutes | \|__\|__\|:\|__\|__\| |
| 9. Enter Information About Child's Last Meal/Snack/Breastfeed | 1. Breast Milk Only<br>2. Snack Only<br>3. Meal Only<br>4. Meal and Snack Only<br>5. Snack and Breast Milk Only<br>6. Meal and Breast Milk Only<br>7. Meal, Snack, and Breast Milk | \|__\| |
| 10. Blood Collection Tube Sample Obtained?<br><br>Note:<br>If no blood draw, then Q14 is asked and rest of form skipped. | 1. all (5 ml)<br>2. partial (< 5 ml)<br>3.none (0 ml) | \|__\| |

EE Sample Collection Form – Section 3: Blood Sample Collection

**Cluster ID |__|__|__| Mother ID |__|__|**

| | | |
|---|---|---|
| 11. Estimate of Blood Volume Collected in Tube | # ml | \|__\|.\|__\|ml |
| 12. Blood Spots Samples Obtained? | 1. all (6 spots)<br>2. partial (< 6 spots)<br>3. none (0 spots) | \|__\| |
| 13. Number of FULLY SaturatedSpots Obtained | # Spots | \|__\| Spots |
| 13b. Anemia Measurement | ###.# | \|__\|\|__\|.\|__\| g/dl |
| 14. Why Was Sample Not Obtained?<br><br>Note:<br><br>If Q10 or Q 12 are 2 or 3, then Q14 is asked. If Q10 or 12 is 1, then Q 14 is skipped. | 1. Parent not available<br>2. Parent refused<br>3. Child not available<br>4. Volume insufficient<br>5. Vein visibility<br>6. Other: Specify | \|__\| |
| 15. Blood Collection: End Time | 24-hour scale<br>Hours : Minutes | \|__\|__\|:\|__\|__\| |

| Samples: | | |
|---|---|---|
| Note:This is the time when sample has been placed in cold box (for tube) or room temperature box (for spots). | Hours : Minutes | |
| 16b. Cooler box Temperature Data Logger ID | #### | \|__\|__\|__\|__\| |

| | | |
|---|---|---|
| 17. Centrifugation Start Time<br><br>Note: Only for child venous blood samples (types B and X). | 24-hour scale<br><br>Hours : Minutes | \|__\|__\|:\|__\|__\| |
| 18. Duration of Centrifugation<br><br>Note: Only for child venous blood samples (type B). | Minutes<br><br>## | \|__\|__\| |
| 19. Plasma Sample Hemolyzed?<br><br>(Plasma is red, not yellow)<br><br>Note: Only for child venous blood samples (type B). | Y/N | \|__\| |
| 20. PBMC Processing: Start Time | 24-hour scale<br><br>Hours : Minutes | \|__\|__\|:\|__\|__\| |
| 21. PBMC Processing: Number of Cells | | |

<table>
<tr><td>22. Sample ID<br><br>Note:<br><br>IDs for all aliquots of blood samples expected in a HH are automatically generated when cluster ID and mother ID are entered in Q1.</td><td>23. Random ID<br><br>Note:<br><br>Random ID that is linked to the specific sample ID is retrieved from barcode ID database and appears automatically.</td><td>24. Sample Type<br><br>Note:<br><br>Field staff selects sample type from dropdown menu.</td><td>25. Aliquots</td><td>26. Samples H & B -80$^0$C Freezer Storage: Start Time</td><td>27. PBMC -80$^0$CFreezer Storage: Start Time</td><td>28. PBMC Liquid Nitrogen Storage: Start Date</td><td>29. PBMC Liquid Nitrogen Storage: Start Time</td></tr>
<tr><td>Cluster (###) + Mother ID (##) + Baseline (B) + Eligible Child (#) + Sample Type + Aliquot (#)</td><td></td><td>H (Whole)<br><br>B (Plasma)<br><br>X (PBMC)<br><br>P (Filter)</td><td></td><td>24-hour scale<br><br>Hours : Minutes</td><td>24-hour scale<br><br>Hours : Minutes</td><td>DD/MM/YYYY</td><td>24-hour scale<br><br>Hours : Minut</td></tr>
<tr><td>|__|__|__|__|__|E|__|H01</td><td>|__|__|__|__|__|__|</td><td>H</td><td>No Aliquot<br><br>Partial Aliquot<br>Full Aliquot</td><td></td><td></td><td></td><td></td></tr>
<tr><td>|__|__|__|__|__|E|__|H02</td><td>|__|__|__|__|__|__|</td><td>H</td><td>No Aliquot<br><br>Partial Aliquot<br>Full Aliquot</td><td>|__|__|:|__|__|</td><td></td><td></td><td></td></tr>
<tr><td>|__|__|__|__|__|E|__|B01</td><td>|__|__|__|__|__|__|</td><td>B</td><td>No Aliquot<br><br>Partial Aliquot<br>Full Aliquot</td><td></td><td></td><td></td><td></td></tr>
<tr><td>|__|__|__|__|__|E|__|B02</td><td>|__|__|__|__|__|__|</td><td>B</td><td>No Aliquot<br><br>Partial Aliquot</td><td></td><td></td><td></td><td></td></tr>
</table>

<table>
<tr><td></td><td></td><td></td><td>Full Aliquot</td><td rowspan="5"></td><td></td><td></td><td></td></tr>
<tr><td>|__|__|__|__|__|E|__|B03</td><td>|__|__|__|__|__|__|</td><td>B</td><td>No Aliquot<br>Partial Aliquot<br>Full Aliquot</td><td></td><td></td><td></td></tr>
<tr><td>|__|__|__|__|__|E|__|B04</td><td>|__|__|__|__|__|__|</td><td>B</td><td>No Aliquot<br>Partial Aliquot<br>Full Aliquot</td><td></td><td></td><td></td></tr>
<tr><td>|__|__|__|__|__|E|__|B05<br>(Optional Aliquot)</td><td>|__|__|__|__|__|__|</td><td>B</td><td>No Aliquot<br>Partial Aliquot<br>Full Aliquot</td><td></td><td></td><td></td></tr>
<tr><td>|__|__|__|__|__|E|__|B06<br>(Optional Aliquot)</td><td>|__|__|__|__|__|__|</td><td>B</td><td>No Aliquot<br>Partial Aliquot<br>Full Aliquot</td><td></td><td></td><td></td></tr>
<tr><td>|__|__|__|__|__|E|__|B07<br>(Optional Aliquot)</td><td>|__|__|__|__|__|__|</td><td>B</td><td>No Aliquot<br>Partial Aliquot<br>Full Aliquot</td><td></td><td></td><td></td><td></td></tr>
<tr><td>|__|__|__|__|__|E|__|X01</td><td>|__|__|__|__|__|__|</td><td>X</td><td>No Aliquot<br>Partial Aliquot<br>Full Aliquot</td><td></td><td rowspan="2">|__|__|:|__|__|</td><td rowspan="2">|__|__| / |__|__|<br>/ |__|__|__|__|</td><td rowspan="2">|__|__|:|__|_</td></tr>
<tr><td>|__|__|__|__|__|E|__|X02</td><td>|__|__|__|__|__|__|</td><td>X</td><td>No Aliquot<br>Partial Aliquot<br>Full Aliquot</td><td></td></tr>
<tr><td>|__|__|__|__|__|E|__|P01</td><td>|__|__|__|__|__|__|</td><td>P</td><td>No Aliquot<br>Partial Aliquot<br>Full Aliquot</td><td></td><td></td><td></td><td></td></tr>
</table>

EE Sample Collection Form – Section 4: Stool Sample Collection

**Cluster ID |__|__|__|   Mother ID |__|__|**

<table>
<tr><td>1. Cluster ID and Mother ID</td><td>### and ##</td><td>|__|__|__| and |__|__|</td></tr>
<tr><td>2. ID of MT/FRA/FRO</td><td>##</td><td>|__|__|</td></tr>
<tr><td>3. Name of MT/FRA/FRO (choose 1):</td><td>4. Dipali Rani Das<br>5. SaniaAkter Dolly<br>6. SyedaFatemaKhatun<br>7.Md. AltafurRahman<br>8. AmalSarker<br>9. Nasrin Sultana<br>10. Md. NurulAlam<br>11. Fatema Begum Sumi<br>12. SyedaLuthfaFamida<br>13. Shahjahan Ali</td><td></td></tr>
<tr><td>4. Date of Sample Collection (when FRA picks up tube)</td><td>DD/MM/YYYY</td><td>|__|__| / |__|__| / |__|__|__|__|</td></tr>
<tr><td>5. Number of Eligible Children</td><td># of Eligible Children</td><td>|__|</td></tr>
<tr><td>6. Child Full Name</td><td></td><td></td></tr>
</table>

| Samples: | | | |
|---|---|---|---|
| 7. Sample ID<br><br>Cluster (###) + Mother ID (##) + Baseline (B) + Eligible Child (#) + Sample Type + Aliquot (#) | 8. Random ID<br><br>Note:<br><br>Random ID that is linked to the specific sample ID is retrieved from barcode ID database and appears automatically. | 9. Sample type<br><br>Note:<br><br>Field staff selects sample type from dropdown menu. | 10. Aliquots<br><br>Note:<br><br>If sample has been fully collected, next question is skipped. |
| \|__\|__\|__\|__\|__\|E\|__\|S01 | \|__\|__\|__\|__\|__\|__\| | S | No Aliquot<br>Partial Aliquot<br>Full Aliquot |
| \|__\|__\|__\|__\|__\|E\|__\|S02 | \|__\|__\|__\|__\|__\|__\| | S | No Aliquot<br>Partial Aliquot<br>Full Aliquot |
| \|__\|__\|__\|__\|__\|E\|__\|S03 | \|__\|__\|__\|__\|__\|__\| | S | No Aliquot<br>Partial Aliquot<br>Full Aliquot |
| \|__\|__\|__\|__\|__\|E\|__\|S04 | \|__\|__\|__\|__\|__\|__\| | S | No Aliquot<br>Partial Aliquot<br>Full Aliquot |
| \|__\|__\|__\|__\|__\|E\|__\|S05 | \|__\|__\|__\|__\|__\|__\| | S | No Aliquot |

| | | | |
|---|---|---|---|
| | | | Partial Aliquot<br><br>Full Aliquot |

**Cluster ID |__|__|__| Mother ID |__|__**

EE Sample Collection Form – Section 4: Stool Sample Collection

| | | |
|---|---|---|
| 11. Why Was It Not Fully Collected?<br><br>Note:In Q10, If Partial Aliquot collected, Q11 is asked. In Q10, if No Aliquotcollected,then<br><br>Q11 is asked and skip to end. | 1. Parents not available<br><br>2. Parents refused<br><br>3. Subject not available<br><br>4. Child did not defecate<br><br>5. Defecation volume insufficient<br><br>6. Other: Specify | \|__\| |
| 12. Date of Defecation<br><br>Note:This is only applicable for stool samples as stool collection containers are left in<br><br>householdsthe day before sample collection. | DD/MM/YYYY | \|__\|__\| / \|__\|__\| / \|__\|__\|__\|__\| |
| 13. Time of Defecation<br><br>Note:Exact or approximate time is entered. | 24-hour scale<br><br>Hours: Minutes | \|__\|__\|:\|__\|__\| |
| 14. FRA Present During Defecation? | 1. Yes | \|__\|__\| |

| | | |
|---|---|---|
| Note:FRA presence or absence during defecation is indicated. | 2. No | |
| 15. Cold Chain Start Time<br><br>Note: This is the time when sample has been placed in cold box. | 24-hour scale<br><br>Hours : Minutes | \|__\|__\|:\|__\|__\| |
| 16. OBSERVE: Consistency of Collected Stool Sample | 1. Normal stool<br><br>(Formed, soft, semi-solid, moist)<br><br>2. Diarrheal stool<br><br>(Unformed, watery)<br><br>3. Constipated stool<br><br>(Formed, hard, dry) | \|__\| |
| 17. OBSERVE: Color of Collected Stool Sample | 1. Yellow<br><br>2. Brown<br><br>3. Black<br><br>4. Green<br><br>5. White<br><br>6. Red<br><br>7. Other: Specify | \|__\| |

#### EE Sample Collection Form – Section 4: Stool Sample Collection

**Cluster ID |__|__|__| Mother ID |__|__**

| | | |
|---|---|---|
| 18. OBSERVE: Any Abnormal Characteristics of Collected Stool Sample | 1. Yes, Mucus<br>2. Yes, Blood<br>3. Yes, Worms<br>4. Yes, Other: Specify<br>5. No | \|__\| |
| 19. ASK: “Does [CHILD NAME] currently have diarrhea?”<br>(Diarrhea = 3 or more unformed stools in a 24-hour period)<br>Note: If Q19 is 1, then ask next question and skip to end. If Q19 is 2 or 99, then skip to Q21. | 1. Yes<br>2. No<br>99. Don’t know | \|__\|__\| |
| 20. ASK: How many consecutive days has this child had diarrhea?”<br>Note: Exact or approximate number of days is entered. | ## Days | \|__\|__\| Days |
| 21. ASK: “How long ago did [CHILD NAME] have diarrhea?”<br>Note: Exact or approximate number of days is entered. If Q21 answer is >0 and NOT 88 or 99, then Q22 is asked. If Q21 answer is 0, 88, or 99, then skip to end. | ## Days<br>88. Never<br>99. Don’t know | \|__\|__\| Days |

| | | |
|---|---|---|
| 22. ASK: “During that specific episode of diarrhea, how many consecutive days did it last?”<br><br>Note: Exact or approximate number of days is entered. | ## Days<br><br>99. Don’t know | \|__\|__\| Days |
| 23. ASK: Where was stool sample collected from? | 1. Diaper provided<br><br>2. Katha (blanket/cloth)<br><br>3. Bed sheet<br><br>4. Potty<br><br>5. Other: Specify<br><br>77. Not applicable | \|__\| |
| 24. ASK: Was the stool in contact with urine (in the diaper/potty/katha/etc.)? | 1. Yes<br><br>2. No<br><br>99. Don’t know | \|__\|__\| |
| 25. Cooler box Temperature Data Logger ID | #### | \|__\|__\|__\|__\| |

EE Sample Collection Form – Section 5: Urine Sample Collection

**Cluster ID |__|__|__| Mother ID |__|__|**

| | | |
|---|---|---|
| 1. Cluster ID and Mother ID | ### and ## | \|__\|__\|__\| and \|__\|__\| |
| 2. ID of MT/FRA/FRO | ## | \|__\|__\| |
| 3. Name of MT/FRA/FRO (choose 1): | 4. Dipali Rani Das<br>5. SaniaAkter Dolly<br>6. SyedaFatemaKhatun<br>7.Md. AltafurRahman<br>8. AmalSarker<br>9. Nasrin Sultana<br>10. Md. NurulAlam<br>11. Fatema Begum Sumi<br>12. SyedaLuthfaFamida<br>13. Shahjahan Ali<br>14. Md. Shariful Islam | |
| 4. Date of Sample Collection | DD/MM/YYYY | \|__\|__\| / \|__\|__\| / \|__\|__\|__\|__\| |

| | | |
|---|---|---|
| 5. Number of Eligible Children | # of Eligible Children | \|__\| |
| 6. Child Full Name | | |
| 7. Consent<br><br>Note: If yes, then skip next question. | Y/N | \|__\| |
| 8. Why Was Consent Not Given?<br><br>Note:If Q7 is no, then answer this question and skip to end. | 1. Household head/primary caregiver not available<br>2. Household head/ primary caregiver refused<br>3. Subject not available<br>4. Genital skin disease<br>5. Other: Specify | \|__\| |
| 9. Volume of Lactulose-MannitolFed to Child | Volume (ml) = average weight X 2 (maximum of 20 ml)<br>##.# | \|__\|__\|.\|__\| ml |
| 10. Lactulose-Mannitol Batch Number | ### | \|__\|__\|__\| |
| 11. Fasting Start Time | 24-hour scale<br>Hours : Minutes | \|__\|__\|:\|__\|__\| |
| 12. Fasting End Time | 24-hour scale<br>Hours : Minutes | \|__\|__\|:\|__\|__\| |

| | | |
|---|---|---|
| 13. Urine Collection Start Time | 24-hour scale<br><br>Hours : Minutes | |__|__|:|__|__|

**EE Sample Collection Form – Section 5: Urine Sample Collection**

**Cluster ID |__|__|__| Mother ID |__|__|**

| 14. Urination episode<br><br>Note:Urine episode number is automatically generated starting from 1. Q14 and the next questions (Q15, Q16, Q17, Q18) will loop continuously until end of urine collection time. | 15. Time of Collection | 16. Volume of Each Urination | 17. Estimated Volume Lost | 18. Stool in Bag? |
|---|---|---|---|---|
| **Urination Episode**<br><br>**#** | 24-hour scale<br><br>Hours : Minutes | Volume (ml)<br><br>### | 1. None<br>2. < ½<br>3. ½<br>4. > ½<br>5. All | Y/N |
| **1** | \|__\|__\|:\|__\|__\| | \|__\|__\|__\|ml | \|__\| | \|__\| |
| **2** | \|__\|__\|:\|__\|__\| | \|__\|__\|__\| ml | \|__\| | \|__\| |
| **3** | \|__\|__\|:\|__\|__\| | \|__\|__\|__\| ml | \|__\| | \|__\| |

| | | | | |
|---|---|---|---|---|
| **4** | \|__\|__\|:\|__\|__\| | \|__\|__\|__\| ml | \|__\| | \|__\| |
| **5** | \|__\|__\|:\|__\|__\| | \|__\|__\|__\| ml | \|__\| | \|__\| |
| **6** | \|__\|__\|:\|__\|__\| | \|__\|__\|__\| ml | \|__\| | \|__\| |
| **7** | \|__\|__\|:\|__\|__\| | \|__\|__\|__\| ml | \|__\| | \|__\| |
| **8** | \|__\|__\|:\|__\|__\| | \|__\|__\|__\| ml | \|__\| | \|__\| |
| **9** | \|__\|__\|:\|__\|__\| | \|__\|__\|__\| ml | \|__\| | \|__\| |
| **10** | \|__\|__\|:\|__\|__\| | \|__\|__\|__\| ml | \|__\| | \|__\| |

EE Sample Collection Form – Section 5: Urine Sample Collection

**Cluster ID |__|__|__| Mother ID |__|__|**

| | | |
|---|---|---|
| 19. Urine Collection End Time for 2-Hour Urine Sample | 24-hour scale<br>Hours : Minutes | |__|__|:|__|__|
| 20. Total Urine Volume After 2 Hours | Volume (ml)<br>### | |__|__|__| ml |
| 21. Total Number of Thimerasol Drops Added After 2 Hours | Number of drops<br>## | |__|__| Drops |
| 22. Number of Loose Stools During 2-Hour Collection | # loose stools | |__| Loose Stools |
| 23. Total Volume of All Aliquots From 2-Hour Urine Collection | Volume (ml)<br>### | |__|__| ml |
| 24. Urine Collection End Time for 5-Hour Urine Sample | 24-hour scale<br>Hours: Minutes | |__|__|:|__|__|
| 25. Total Urine Volume After 5 Hours | Volume (ml)<br>### | |__|__|__| ml |

| | | |
|---|---|---|
| 26. Total Number of Thimerasol Drops Added After 5 Hours | Number of drops<br>## | \|__\|__\| Drops |
| 27.Number of Loose Stools During 5-Hour Collection | # of loose stools | \|__\|Loose Stools |
| 28. Total Volume of All Aliquots From 5-Hour Urine Collection | Volume (ml)<br>### | \|__\|__\| ml |
| 29. Did Child Eat During First 1-Hour Fasting Period?<br>(Before LM dose) | Y/N | \|__\| |
| 30. Did Child Eat During Last 0.5-Hour Fasting Period?<br>(Immediately after LM dose) | Y/N | \|__\| |

**Cluster ID |__|__|__| Mother ID |__|__**

<table>
<tr><td colspan="6">Samples:</td></tr>
<tr>
<td>31. Sample ID<br><br>Note:<br><br>IDs for all aliquots of urine samples expected in a HH are automatically generated when cluster ID and mother ID are entered in Q1.(Aliquots 01-06 after 2 hrs urine collection. Aliquots 07-12 after 5 hrs urine collection)</td>
<td>32. Random ID<br><br>Note:<br><br>Random ID that is linked to the specific sample ID is retrieved from barcode ID database and appears automatically</td>
<td>33. Sample Type<br><br>Note:<br><br>Field staff selects sample type from dropdown menu.</td>
<td>34. Aliquots<br><br>Note:<br><br>If sample has been fully collected, next question is skipped. For sample type U, full aliquot = 2 ml per cryovial.</td>
<td>35. Why was it not fully collected?<br><br>Note:<br><br>If Q34 is No Aliquot or Partial Aliquot, then Q 35 is asked.<br><br>1. Child did not urinate<br>2. Urination volume insufficient<br>3. Leakage<br>4. Partial: Parent Refused<br>5. Other: Specify</td>
<td>36. Cold Chain Start Time<br><br>Note:<br><br>This is the time when sample has been placed in cold box.<br><br>24-hour scale<br>Hours : Minutes</td>
</tr>
<tr>
<td>|__|__|__|__|__|E|__|U01</td>
<td>|__|__|__|__|__|__|</td>
<td>U</td>
<td>No Aliquot<br>Partial Aliquot<br>Full Aliquot</td>
<td rowspan="3">|__| 2 Hours</td>
<td rowspan="3">|__|__|:|__|__|</td>
</tr>
<tr>
<td>|__|__|__|__|__|E|__|U02</td>
<td>|__|__|__|__|__|__|</td>
<td>U</td>
<td>No Aliquot<br>Partial Aliquot<br>Full Aliquot</td>
</tr>
<tr>
<td>|__|__|__|__|__|E|__|U03</td>
<td>|__|__|__|__|__|__|</td>
<td>U</td>
<td>No Aliquot</td>
</tr>
</table>

| | | | | | |
|---|---|---|---|---|---|
| | | | Partial Aliquot<br>Full Aliquot | | |
| \|__\|__\|__\|__\|__\|E\|__\|U04 | \|__\|__\|__\|__\|__\|__\| | U | No Aliquot<br>Partial Aliquot<br>Full Aliquot | | |
| \|__\|__\|__\|__\|__\|E\|__\|U05 | \|__\|__\|__\|__\|__\|__\| | U | No Aliquot<br>Partial Aliquot<br>Full Aliquot | | |
| \|__\|__\|__\|__\|__\|E\|__\|U06 | \|__\|__\|__\|__\|__\|__\| | U | No Aliquot<br>Partial Aliquot<br>Full Aliquot | | |
| \|__\|__\|__\|__\|__\|E\|__\|U07 | \|__\|__\|__\|__\|__\|__\| | U | No Aliquot<br>Partial Aliquot<br>Full Aliquot | | |
| \|__\|__\|__\|__\|__\|E\|__\|U08 | \|__\|__\|__\|__\|__\|__\| | U | No Aliquot<br>Partial Aliquot<br>Full Aliquot | \|__\| 5 Hours | \|__\|__\|:\|__\|__\| |
| \|__\|__\|__\|__\|__\|E\|__\|U09 | \|__\|__\|__\|__\|__\|__\| | U | No Aliquot<br>Partial Aliquot<br>Full Aliquot | | |

| | | | |
|---|---|---|---|
| \|__\|__\|__\|__\|__\|E\|__\|U10 | \|__\|__\|__\|__\|__\|__\| | U | No Aliquot<br>Partial Aliquot<br>Full Aliquot |
| \|__\|__\|__\|__\|__\|E\|__\|U11 | \|__\|__\|__\|__\|__\|__\| | U | No Aliquot<br>Partial Aliquot<br>Full Aliquot |
| \|__\|__\|__\|__\|__\|E\|__\|U12 | \|__\|__\|__\|__\|__\|__\| | U | No Aliquot<br>Partial Aliquot<br>Full Aliquot |

| | | |
|---|---|---|
| 37. Why was urine collection stopped before 5 hours?<br><br>Note:<br>Skip if both 2 and 5 hour urine collection completed. | 1. Baby crying a lot<br>2. Baby developed a rash at U-bag attachment site<br>3. Other parent came home and refused<br>4. Another family member arrived and encouraged mother to refuse<br>5. Mother and child left to visit another place<br>6. Other: Specify<br>77. Not applicable | \|__\| |
| 38. Cooler box Temperature Data Logger ID | #### | \|__\|__\|__\|__\| |

#### EE Sample Collection Form – Section 5A: Pre-LM Urine Sample Collection

**Cluster ID |__|__|__| Mother ID |__|__|**

| | | |
|---|---|---|
| 39. Cluster ID and Mother ID | ### and ## | \|__\|__\|__\| **and** \|__\|__\| |
| 40. Date of Sample Collection | DD/MM/YYYY | \|__\|__\| / \|__\|__\| / \|__\|__\|__\|__\| |
| 41. **Pre-LM** Urine Collection Start Time | 24-hour scale<br>Hours : Minutes | \|__\|__\|:\|__\|__\| |

| 42. **Pre-LM**Urination episode<br><br>Note:Urine episode number is automatically generated starting from 1. Q42 and the next questions (Q43, Q44, Q45, Q46) will loop | 43. Time of Collection | 44. Volume of Each Urination | 45. Estimated Volume Lost | 46. Stool in Bag? |
|---|---|---|---|---|

| continuously until end of urine collection time. | | | | |
|---|---|---|---|---|
| **Urination Episode**<br><br>**#** | 24-hour scale<br><br>Hours : Minutes | Volume (ml)<br><br>### | 1. None<br><br>2. < ½<br><br>3. ½<br><br>4. > ½<br><br>5. All | Y/N |
| **1** | \|__\|__\|:\|__\|__\| | \|__\|__\|__\| ml | \|__\| | \|__\| |
| **2** | \|__\|__\|:\|__\|__\| | \|__\|__\|__\| ml | \|__\| | \|__\| |
| **3** | \|__\|__\|:\|__\|__\| | \|__\|__\|__\| ml | \|__\| | \|__\| |
| **4** | \|__\|__\|:\|__\|__\| | \|__\|__\|__\| ml | \|__\| | \|__\| |
| **5** | \|__\|__\|:\|__\|__\| | \|__\|__\|__\| ml | \|__\| | \|__\| |

**Cluster ID |__|__|__| Mother ID |__|__|**

| | | |
|---|---|---|
| 47. **Pre-LM** Urine Collection End Time | 24-hour scale<br><br>Hours : Minutes | |__|__|:|__|__|
| 48. Total Urine Volume of **Pre-LM** Urine | Volume (ml)<br><br>### | |__|__|__| ml |
| 49. Total Number of Thimerasol Drops Added to **Pre-LM** Urine | Number of drops<br><br>## | |__|__| Drops |
| 50. Number of Loose Stools During **Pre-LM** Urine Collection | # loose stools | |__| Loose Stools |
| 51. Total Volume of All Aliquots From **Pre-LM** Urine Collection | Volume (ml)<br><br>### | |__|__| ml |

| Samples: | | | | | |
|---|---|---|---|---|---|
| 52. Sample ID<br><br>Note:<br><br>IDs for all aliquots of urine samples expected in a HH are automatically generated when cluster ID and mother ID are entered in Q1.<br><br>**(Aliquots 13-18 after Pre-LM urine collection)** | 53. Random ID<br><br>Note:<br><br>Random ID that is linked to the specific sample ID is retrieved from barcode ID database and appears automatically | 54. Sample Type<br><br>Note:<br><br>Field staff selects sample type from dropdown menu. | 55. Aliquots<br><br>Note:<br><br>If sample has been fully collected, next question is skipped. For sample type U, full aliquot = 2 | 56. Why was Pre-LM urine not fully collected?<br><br>Note:<br><br>If Q55 is No Aliquot or Partial Aliquot, then Q56 is asked.<br><br>1. Child did not urinate | 57. Cold Chain Start Time<br><br>Note:<br><br>This is the time when sample has been placed in cold box. |

| | | | | | |
|---|---|---|---|---|---|
| | | | ml per cryovial. | 2. Urination volume insufficient<br>3. Leakage<br>4. Partial: Parent Refused<br>5. Other: Specify | 24-hour scale<br>Hours : Minutes |
| \|__\|__\|__\|__\|__\|E\|__\|U13 | \|__\|__\|__\|__\|__\|__\| | U | No Aliquot<br>Partial Aliquot<br>Full Aliquot | \|__\| Pre-LM Urine | \|__\|__\|:\|__\|__\| |
| \|__\|__\|__\|__\|__\|E\|__\|U14 | \|__\|__\|__\|__\|__\|__\| | U | No Aliquot<br>Partial Aliquot<br>Full Aliquot | | |
| \|__\|__\|__\|__\|__\|E\|__\|U15 | \|__\|__\|__\|__\|__\|__\| | U | No Aliquot<br>Partial Aliquot<br>Full Aliquot | | |
| \|__\|__\|__\|__\|__\|E\|__\|U16 | \|__\|__\|__\|__\|__\|__\| | U | No Aliquot<br>Partial Aliquot<br>Full Aliquot | | |
| \|__\|__\|__\|__\|__\|E\|__\|U17 | \|__\|__\|__\|__\|__\|__\| | U | No Aliquot<br>Partial Aliquot<br>Full Aliquot | | |

| | | | |
|---|---|---|---|
| \|__\|__\|__\|__\|__\|E\|__\|U18 | \|__\|__\|__\|__\|__\|__\| | U | No Aliquot<br><br>Partial Aliquot<br><br>Full Aliquot |

#### WASH Benefits Common Module 50. Lead exposure in depth interview guide

**Note: Ask questions to the mother and the person in the household with the most knowledge regarding the exposure**

Questions for mother

1. How long have youlived in the current location?
2. How much of the rice that you eat comes from the fields that your household farm?
3. During which season do you purchase rice that is grown from somewhere else?
4. How much of the vegetables that you eat come from the fields your household farm?
5. Which vegetables do you purchase in the local market?
6. During which season do you purchase these vegetables?
7. How much of the fruits that you eat come from your household farm?
8. Which fruits do you purchase in the local market?
9. During which season do you purchase these fruits?
10. Do you or anyone in your household work with paint, solder, industrial waste or batteries?
    a) If yes, how often do you work with it?
    b) Are any of the batteries the type used in cars, trucks or motorcycle that contain acid?
    c) How are these batteries handled?
11. Do you or anyone in your household work inmanufacturing or recycling goods? If yes, explain. Detail any contact with metals, solvents, chemicals, exhaust.
12. How close is the nearest road that motor vehicles use regularly to your home?
13. What foods do your purchase that come in a metal can? How often do you consume each of these foods?
14. How often do you wear kohl (makeup)? From where do you obtain the makeup? Probe details on whether it is a manufactured product or made. If it is a manufactured product, record the brand name of the product.
15. Does your house have tin roofs or tin walls? How long have you had them?
16. What material is your cooking pot made from?
17. What materials are your dinner plates made from? Are they painted?
18. What material is your water pot made from?

Questions for person most knowledgeable regarding household farming

19. How many harvests of rice does your land produce each year?
20. What other crops do you grow on your land (including small gardens)?
21. How often did you apply fertilizer on your land in the last year?
22. How do you decide how much fertilizer to use?
23. Did you use the same type of fertilizer for each application? If there were different types, probe and explicate the differences, e.g. different types of fertilizers for different times of year or for different crop?
24. What are the names of the types of fertilizers that you used? If there is more than one name, describe the roles of the different types.
25. How do you decide which fertilizer to use?
26. How do you decide how much fertilizer to use?
27. What packaging does the fertilizer come in?
28. Do you have to mix or otherwise prepare the fertilizer? If yes,
    a) Who is involved? (e.g. other family members?)
    b) Where is it mixed?
    c) Please describe the process.
29. What do you do with the leftover fertilizer packaging? Probe: Any re-use the packaging/container?
30. Who applies the fertilizer?
31. Describe the process of applying the fertilizer.
32. What do you do with leftover fertilizer?

33. How often did you apply pesticide on your land in the last year?
34. Did you use the same type of pesticide for each application? If there were different types, probe and explicate the differences, e.g. different types of pesticides for different times of year? for different crops? for different pests?
35. What are the names of the types of pesticides that you used? If there is more than one name, describe the roles of the different types.
36. Did you use pesticides before you saw any pests or do you always wait until you see pests? Why do you follow this practice?
37. How do you decide which pesticide to use?
38. How do you decide how much pesticide to use?
39. What packaging does the pesticide come in?
40. Do you have to mix or otherwise prepare the pesticide? If yes,
    a) Who is involved? (e.g. other family members?)
    b) Where is it mixed?

c) Please describe the process.

41. What do you do with the leftover pesticide packaging? Probe: Any re-use the packaging/container?
42. Who applies the pesticide?
43. Describe the process of applying the pesticide.
44. What do you do with leftover pesticide?

45. How often did you apply herbicide on your land in the last year?
46. If he/she applied herbicide, why did you apply it?
47. Did you use the same type of herbicide for each application? If there were different types, probe and explicate the differences, e.g. different types of herbicides for different times of year? for different crops? for different pests?
48. What are the names of the types of herbicides that you used? If there is more than one name, describe the roles of the different types.
49. Did you use herbicides before you saw any weeds or do you always wait until you see weeds? Why do you follow this practice?
50. How do you decide which herbicide to use?
51. How do you decide how much herbicide to use?
52. What packaging does the herbicide come in?
53. Do you have to mix or otherwise prepare the herbicide? If yes,
    a) Who is involved? (e.g. other family members?)
    b) Where is it mixed?
    c) Please describe the process.
54. What do you do with the leftover herbicide packaging? Probe: Any re-use the packaging/container?
55. Who applies the herbicide?
56. Describe the process of applying the herbicide?
57. What do you do with leftover herbicide?

58. How often did you apply fungicide on your crops in the last year?
59. If he/she applied fungicide, why did you apply it?
60. Did you use the same type of fungicide for each application? If there were different types, probe and explicate the differences, e.g. different types of fungicides for different times of year? for different crops? for different pests?
61. What are the names of the types of fungicides that you used? If there is more than one name, describe the roles of the different types.
62. Did you use fungicides before you saw any fungus or do you always wait until you see fungus? Why do you follow this practice?

63. How do you decide which fungicide to use?
64. How do you decide how much fungicide to use?
65. What packaging does the fungicide come in?
66. Do you have to mix or otherwise prepare the fungicide? If yes,
    a) Who is involved? (e.g. other family members?)
    b) Where is it mixed?
    c) Please describe the process.
67. What do you do with the leftover fungicide packaging? Probe: Any re-use the packaging/container?
68. Who applies the fungicide?
69. Describe the process of applying the fungicide?
70. What do you do with leftover fungicide?

71. Did you apply any other chemicals to crops/lands in the last year that we have not already discussed? If yes, explicate what else he/she used, how often, its name, and why it was used.

**ওয়াশ বেনিফিট মডিউল ৫০ (Module 50)**

**সীসার প্রভাব সম্পর্কিত নিবিড় সাক্ষাৎকারের গাইডলাইন**

[ নোট: মা এবং পরিবারের সবচেয়ে তথ্য সমৃদ্ধ ব্যাক্তিকে এক্সপোজার বিষয়ক প্রশ্নগুলো করুন ]

**মায়ের জন্য প্রশ্নাবলী**

১. বর্তমান জায়গায় আপনি কতদিন ধরে বসবাস করছেন?

২. আপনারা যে ভাত খান তার কি পরিমাণ আপনাদের গৃহস্থালী খামার থেকে আসে?

৩. আপনি সাধারণত কোন ঋতু/সময়ে চাল কেনেন যা অন্য কোন জায়গায় উৎপাদন করা হয়?

৪. আপনার খাবারের কি পরিমাণ সবজি আপনার নিজের খামার/গৃহস্থালী থেকে আসে?

৫. স্থানীয় বাজার থেকে আপনি কি ধরণের সবজি কেনেন?

৬. এই সব সবজি আপনি কোন ঋতু/সময়ে কেনেন?

৭. আপনার খাবারের কি পরিমাণ ফলমূল আপনার নিজের খামার/গৃহস্থালী থেকে আসে?

৮. স্থানীয় বাজার থেকে আপনি কি কি ফলমূল কেনেন?

৯. কোন ঋতুতে/সময়ে আপনি এইসব ফল কেনেন?

১০. আপনি অথবা আপনার পরিবারের কেউ কি রং, ঝালাই, শিল্পবর্জ্য অথবা ব্যাটারির কাজ করে?

ক) যদি হ্যাঁ হয়, তাহলে কত ঘনঘন এগুলো করেন?

খ) এখানে কি কোন এসিডযুক্ত ব্যাটারী আছে যা গাড়ি,ট্রাক অথবা মোটর সাইকেলে ব্যবহৃত হয়।

গ) সে সব ব্যাটারিগুলো কিভাবে নাড়াচাড়া করেন?

১১. আপনি অথবা আপনার পরিবারের কেউ কি পণ্য প্রক্রিয়াজাতকরণ এবং পূর্ণ প্রক্রিয়াজাতকরণের কাজের সাথে যুক্ত আছে? যদি থাকে দয়া করে বর্ণনা করুন। ধাতু,দ্রাবক/তরল পদার্থ, রাসায়নিক, ধোয়া নির্গম নল প্রভৃতির সাথে যে কোন ধরণের স্পর্শ বিস্তারিতভাবে বর্ণনা করুন।

১২. নিয়মিত মোটরযান চলাচলকারী রাস্তা আপনার বাড়ী থেকে কত কাছে?

১৩. ধাতব পাত্রে প্রক্রিয়াজাত কি কি ধরণের খাবার আপনি কেনেন? আপনি কত ঘনঘন এই সব খাবার খান?

১৪. আপনি সাধারণত কখন কখন সুরমা (প্রসাধন) ব্যাবহার করেন? কোথা থেকে আপনি এই প্রসাধন নেন? এটা কোন প্রক্রিয়াজাতকৃত অথবা তৈরীকৃত পণ্য কিনা তা বিস্তারিত জানতে চেস্টা করুন। এটা যদি প্রক্রিয়াজাত পণ্য হয় তাহলে ব্রান্ড নাম সংগ্রহ করুন ।

১৫. আপনার ঘরে কি কোন টিনের ছাদ বা টিনের বেড়া আছে? কত দিন ধরে আছে?

১৬. আপনার রান্নার হাড়ি পাতিল কি দিয়ে তৈরী?

১৭. আপনার খাবার থালা বাসন কি দিয়ে তৈরী/কিসের তৈরী? এগুলো কি রং করা?

১৮. আপনার পানির পাত্র, কি দিয়ে তৈরী/ কিসের তৈরী?

**গৃহস্থালী খামার সম্পর্কে সবচেয়ে তথ্যসমৃদ্ধ ব্যাক্তির জন্য প্রশ্ন**

১৯. প্রতি বছর আপনার জমি থেকে কত ধরণের ধান সংগ্রহ/তোলা হয়?

২০. আপনার জমিতে অন্য আর কি কি ধরণের ফসল আপনি চাষ করেন (ছোট বাগান সহ)?

২১. গতবছর আপনার জমিতে আপনি কত ঘনঘন রাসায়নিক সার ব্যবহার করেছেন?

২২. কি পরিমান রাসায়নিক সার ব্যবহার করতে হবে সেটা আপনি কিভাবে ঠিক করেন?

২৩. প্রতিক্ষেত্রে আপনি কি একই ধরণের রাসায়নিক সার ব্যবহার করেন? যদি ভিন্ন ধরণের হয় তাহলে পার্থক্যগুলো বিশদভাবে জানার চেস্টা করুন। যেমন, বিভিন্ন ফসলের জন্য বছরের বিভিন্ন সময়ে বিভিন্ন ধরণের রাসায়নিক সার?

২৪. যে যে ধরণের রাসায়নিক সার আপনি ব্যবহার করেছেন, সেগুলোর নাম কি? যদি একাধিক ধরণের থাকে তাহলে তাদের প্রত্যেকটির ভূমিকা বর্ণনা করুন।

২৫. কোন ধরণের রাসায়নিক সার ব্যবহার করা হবে সেটা আপনি কিভাবে নির্ধারণ করেন?

২৬. কি পরিমাণ রাসায়নিক সার ব্যবহার করা হবে সেটা আপনি কিভাবে নির্ধারণ করেন?

২৭. কি ধরণের মোড়কে রাসায়নিক সারগুলো থাকে?

২৮. অপনাকে কি কখনো রাসায়নিক সার মিশিয়ে বা অন্য কোন উপায়ে প্রস্তুত করতে হয়? যদি হয়,

ক) কে সেটা করে/কারা কারা যুক্ত থাকে (যেমন, পরিবারের অন্যান্য সদস্য?)

খ) কোথায় সেগুলো মেশানো হয়?

গ) দয়া করে সার মেশানোর পদ্ধতিটি বর্ণনা করুন।

২৯. সারের পরিত্যাক্ত মোড়কগুলো দিয়ে আপনি কি করেন? প্রোব: মোড়ক/ধারণ পাত্রের কোন প্রকার পুনঃব্যবহার?

৩০. কে এই রাসায়নিক সার প্রয়োগ করে?

৩১. এই রাসায়নিক সার প্রয়োগের পদ্ধতি বর্ণনা করুন।

৩২. পরিত্যাক্ত রাসায়নিক সার দিয়ে আপনি কি করেন?

৩৩. গতবছর কতবার/কতদিন পর পর আপনি জমিতে কীটনাশক ব্যবহার করেছেন?

৩৪. প্রতিক্ষেত্রে আপনি কি একই ধরণের কীটনাশক ব্যবহার করছেন? যদি ভিন্ন ধরণের হয় তাহলে পার্থক্যগুলো বিশদভাবে জানার চেস্টা করুন করুন। যেমন, বছরের বিভিন্ন সময়ে বিভিন্ন ধরণের কীটনাশক, বিভিন্ন ফসলের জন্য, বিভিন্ন পোকামাকড়ের জন্য।

৩৫. আপনি যে যে ধরণের কীটনাশক ব্যবহার করেছেন সেগুলোর নাম কি? যদি একাধিক ধরণের থাকে তাহলে তাদের প্রত্যেকটির ভূমিকা বর্ণনা করুন।

৩৬. পোকা মাকড় দেখার আগে কি আপনি কীটনাশক ব্যবহার করেছেন অথবা আপনি সবসময়ই পোকা-মাকড় দেখা পর্যন্ত অপেক্ষা করেন? আপনি কেন এই রীতি মেনে চলেন?

৩৭. কোন ধরণের কীটনাশক ব্যবহার করা হবে এটা আপনি কিভাবে নির্ধারণ করেন?

৩৮. কি পরিমাণ কীটনাশক ব্যবহার করা হবে এটা আপনি কিভাবে নির্ধারণ করেন?

৩৯. কি ধরণের মোড়কে কীটনাশকগুলো থাকে?

৪০. আপনাকে কি কখনও কীটনাশক মিশিয়ে বা অন্য কোন উপায়ে প্রস্তুত করতে হয়? যদি হয়,

ক) কে সেটা করে/কারা কারা এ কাজের সাথে যুক্ত থাকে (যেমন, পরিবারের অন্যান্য সদস্য?)

খ) কোথায় এ মিশ্রণের কাজটি করা হয়?

গ) দয়া করে পদ্ধতিটি বর্ণনা করুন।

৪১. পরিত্যাক্ত কীটনাশকের মোড়কগুলো দিয়ে আপনি কি করেন? প্রোব: মোড়ক/ধারণ পাত্রের কোন প্রকার পুনঃব্যবহার?

৪২. কে এই কীটনাশক প্রয়োগ করেন?

৪৩. কীটনাশক প্রয়োগের পদ্ধতি বর্ণনা করুন।

৪৪. অবশিষ্ট কীটনাশক দিয়ে আপনি কি করেন?

৪৫. গত বছরে কত ঘনঘন আপনি জমিতে আগাছা নাশক ব্যাবহার করেছেন?

৪৬. সে যদি আগাছানাশক ব্যাবহার করে তাহলে কেন এটা করেছে?

৪৭. আপনি কি প্রতিক্ষেত্রেই একই ধরণের আগাছানাশক প্রয়োগ করেছেন যদি ভিন্ন ধরণের হয় তাহলে পার্থক্যগুলো বিশদভাবে জানার চেস্টা করুন করুন। যেমন: বছরে বিভিন্ন সময়ে, বিভিন্ন ধরণের আগাছানাশক বিভিন্ন ফসলের জন্য? বিভিন্ন পোকামাকড়ের জন্য?

৪৮. আপনি যেসব আগাছানাশক ব্যবহার করেছেন সেগুলোর নাম কি? যদি একাধিক ধরণের থাকে তাহলে তাদের প্রত্যেকটির ভূমিকা বর্ণনা করুন।

৪৯. কোন আগাছা দেখার আগেই কি আপনি আগাছানাশক ব্যবহার করেছেন? অথবা আগাছা দেখা পর্যন্ত অপেক্ষা করেন? আপনি কেন এই রীতি অনুসরণ করেন?

৫০. কোন ধরণের আগাছানাশক ব্যবহার করা হবে এটা আপনি কিভাবে নির্ধারণ করেন?

৫১. কি পরিমাণ আগাছানাশক ব্যবহার করা হবে এটা আপনি কিভাবে নির্ধারণ করেন?

৫২. কি ধরণের মোড়কে এই আগাছানাশকগুলো থাকে?

৫৩. আপনাকে কি কখনও আগাছানাশক মিশিয়ে বা অন্য কোন উপায়ে প্রস্তুত করতে হয়? যদি হ্যাঁ হয়,

ক) কে সেটা করে/ কারা কারা এ কাজের সাথে যুক্ত থাকে (যেমন, পরিবারের অন্যান্য সদস্য?)

খ) কোথায় মেশানো হয়?

গ) দয়া করে পদ্ধতিটি বর্ণনা করুন।

৫৪. পরিত্যাক্ত আগাছানাশকের মোড়কগুলো দিয়ে আপনি কি করেন? মোড়ক/ধারণ পাত্রের পুনঃব্যবহার প্রোব করুন ।

৫৫. কে এই আগাছানাশক প্রয়োগ করেন?

৫৬. আগাছানাশক প্রয়োগ পদ্ধতি বর্ণনা করুন।

৫৭. অবশিষ্ট আগাছানাশক দিয়ে আপনি কি করেন?

৫৮. গতবছর কতবার/কতদিন পর পর আপনার জমিতে ছত্রাকনাশক ব্যবহার করেছেন?

৫৯. সে যদি ছত্রাকনাশক ব্যবহার করে থাকে তাহলে কেন করেছে?

৬০. প্রতিক্ষেত্রে কি আপনি একই ধরণের ছত্রাকনাশক ব্যবহার করছেন? যদি ভিন্ন ধরণের হয় তাহলে পার্থক্যগুলো বিশদভাবে জানার চেস্টা করুন করুন। প্রোবঃ বছরে বিভিন্ন সময়ে, বিভিন্ন ধরণের ছত্রাকনাশক বিভিন্ন ফসলের জন্য? বিভিন্ন পোকামাকড়ের জন্য?

৬১. আপনি যেসব ছত্রাকনাশক ব্যবহার করেছেন সেগুলোর নাম কি? যদি একাধিক ধরণের থাকে তাহলে তাদের প্রত্যেকটির ভূমিকা বর্ণনা করুন।

৬২. কোন ছত্রাক দেখার আগেই কি আপনি ছত্রাকনাশক ব্যবহার করেছেন? অথবা ছত্রাক দেখা পর্যন্ত অপেক্ষা করেন? আপনি কেন এই রীতি অনুসরণ করেন?

৬৩. কোন ধরণের ছত্রাকনাশক ব্যবহার করা হবে এটা আপনি কিভাবে নির্ধারণ করেন?

৬৪. কি পরিমাণ ছত্রাকনাশক ব্যবহার করা হবে এটা আপনি কিভাবে নির্ধারণ করেন?

৬৫. কি ধরণের মোড়কে এই ছত্রাকনাশকগুলো থাকে?

৬৬. আপনাকে কি কখনও ছত্রাকনাশক মিশিয়ে বা অন্য কোন উপায়ে প্রস্তুত করতে হয়? যদি হ্যাঁ হয়,

ক) কে সেটা করে/কারা কারা এ কাজের সাথে যুক্ত থাকে (যেমন, পরিবারের অন্যান্য সদস্য?)

খ) কোথায় মেশানো হয়?

গ) দয়া করে পদ্ধতিটি বর্ণনা করুন

৬৭. পরিত্যাক্ত ছত্রাকনাশকের মোড়কগুলো দিয়ে আপনি কি করেন? মোড়ক/ধারণ পাত্রের পুনঃব্যবহার প্রোব করুন।

৬৮. কে এই ছত্রাকনাশক প্রয়োগ করেন?

৬৯. ছত্রাকনাশকের প্রয়োগ পদ্ধতি বর্ণনা করুন।

৭০. অবশিষ্ট ছত্রাকনাশক দিয়ে আপনি কি করেন?

৭১. আপনি কি গত বছরে জমিতে/ফসলে অন্য কোন রাসায়নিক প্রয়োগ করেছেন যা আমরা ইতিমধ্যে আলোচনা করিনি। যদি হ্যাঁ হয়, তাহলে সে কি ব্যবহার করেছে, কতবার, এর নাম এবং কেন এটা ব্যবহার করা হয়েছে বিস্তারিত বর্ণনা করুন।

#### WASH Benefits Module 51. Agrochemicals salesman in-depth interview guide

1. How long have you worked in the current location?
2. How many types of fertilizer do you sell?
3. What names do you and farmers use to refer to the different types of fertilizers?
4. Why do farmers purchase different types of fertilizers?
5. Where do you secure each type of these fertilizers?
6. Where are each type of these fertilizers manufactured?
7. Which fertilizer is your biggest seller? Why?

8. How many types of pesticide do you sell?
9. What names do you and farmers use to refer to the different types of pesticides?
10. Why do farmers purchase different types of pesticides?
11. Where do you secure each type of these pesticides?
12. Where are each type of these pesticides manufactured?
13. Which pesticide is your biggest seller? Why?

14. How many types of herbicide do you sell?
15. What names do you and farmers use to refer to the different types of herbicides?
16. Why do farmers purchase different types of herbicides?
17. Where do you secure each type of these herbicides?
18. Where are each type of these herbicides manufactured?
19. Which herbicide is your biggest seller? Why?

20. How many types of fungicide do you sell?
21. What names do you and farmers use to refer to the different types of fungicides?
22. Why do farmers purchase different types of fungicides?
23. Where do you secure each type of these fungicides?
24. Where are each type of these fungicides manufactured?
25. Which fungicide is your biggest seller? Why?

**ওয়াশ বেনিফিট মডিউল ৫১ (Module 51)**

কৃষি-রাসায়নিক বিক্রেতার জন্য নিবিড় সাক্ষাৎকার গাইডলাইন

১. আপনি কতদিন ধরে এই এলাকায় কাজ করছেন?
২. আপনি কত ধরনের সার বিক্রি করেন?
৩. বিভিন্ন ধরনের সারকে বোঝাতে আপনি এবং অন্যান্য কৃষকরা কি কি নাম ব্যবহার করে থাকেন?
৪. কেন কৃষকরা বিভিন্ন ধরনের সার ক্রয় করে?
৫. এই সারগুলোর প্রত্যেকটিকে আপনারা কোথায় থেকে সংগ্রহ করেন?
৬. এই সারগুলো প্রত্যেকটি কোথায় কোথায় উৎপাদিত/প্রক্রিয়াজাতকরণ হয়?
৭. কোন সারটি সবচেয়ে বেশী বিক্রি হয়? কেন?
৮. কত ধরনের কীটনাশক আপনি বিক্রি করেন?
৯. বিভিন্ন ধরনের কীটনাশক বোঝাতে আপনি এবং অন্যান্য কৃষকরা কি কি নাম ব্যবহার করেন?
১০. কেন কৃষকরা বিভিন্ন ধরনের কীটনাশক ক্রয় করে?
১১. এই কীটনাশকগুলোর প্রত্যেকটিকে আপনারা কোথায় থেকে সংগ্রহ করেন?
১২. এই কীটনাশকগুলোর প্রত্যেকটি কোথায় কোথায় উৎপাদিত/প্রক্রিয়াজাতকরণ হয়?
১৩. কোন কীটনাশকটি সবচেয়ে বেশী বিক্রি হয়? কেন?

১৪. কত ধরনের আগাছানাশক আপনি বিক্রি করেন?
১৫. বিভিন্ন ধরনের আগাছানাশক বোঝাতে আপনি এবং অন্যান্য কৃষকরা কি কি নাম ব্যবহার করেন?
১৬. কেন কৃষকরা বিভিন্ন ধরনের আগাছানাশক ক্রয় করে?
১৭. এই আগাছানাশকগুলোর প্রত্যেকটিকে আপনারা কোথায় থেকে সংগ্রহ করেন?
১৮. এই আগাছানাশকগুলোর প্রত্যেকটি কোথায় কোথায় উৎপাদিত/প্রক্রিয়াজাতকরণ হয়?
১৯. কোন আগাছানাশকটি সবচেয়ে বেশী বিক্রি হয়? কেন?

২০. কত ধরনের ছত্রাকনাশক আপনি বিক্রি করেন?
২১. বিভিন্ন ধরনের ছত্রাকনাশক বোঝাতে আপনি এবং অন্যান্য কৃষকরা কি কি নাম ব্যবহার করেন?
২২. কেন কৃষকরা বিভিন্ন ধরনের ছত্রাকনাশক ক্রয় করে?
২৩. এই ছত্রাকনাশকগুলোর প্রত্যেকটিকে আপনারা কোথায় থেকে সংগ্রহ করেন?
২৪. এই ছত্রাকনাশকগুলোর প্রত্যেকটি কোথায় কোথায় উৎপাদিত/প্রক্রিয়াজাতকরণ হয়?
২৫. কোন ছত্রাকনাশকটি সবচেয়ে বেশী বিক্রি হয়? কেন?

#### WASH Benefits Common Module 52. Lead exposure case-control qustionnaire (সীসার প্রভাব সম্পর্কীত প্রশ্নপত্র)

Interview Start Time (সাৰাৎকার শুরম্নর সময়):

#### Section 1: Identification and Demographics (সেকসন-১: খানা পরিচিতি ও ভৌগলিক অবস্থান)

1.1 Enter participant ID (five-digit ID from WASH Benefits): [খানা আইডি নম্বর ৫ সংখ্যার আইডি ওয়াশ বেনিফিট স্টাডি প্রদত্ত তালিকা থেকে]____________________________________________________

1.2 What is your name? (Name of respondent should be the target child's mother): [আপনার নাম কি (অবশ্যই টার্গেট শিশুর মা)]____________________________________________________

1.3 How many people in total live in this compound? (আপনার/আপনাদের বাড়িতে মোট কতজন লোক বসবাস করে?)______________________________________________________________________

1.4 In your household who is the point person for cultivating the land? (আপনার খানাতে কে সবচেয়ে গুরুত্বপূর্ন ব্যক্তি যে আপনাদের চাষাবাদের জমি সম্পর্কে ভাল ধারনা রাখেন?)

1) Husband (উত্তরদাতার স্বামী)
2) Father of respondent (উত্তরদাতার বাবা/শ্বশুর)
3) Uncle (উত্তরদাতার চাচা)
4) Brother-in-law (দেবর)
5) Agricultural labor (কৃষিশ্রমিক)
77) Other : Specify (অন্যান্য: নির্দ্দিষ্ট করে লিখুন)

1.5 Age of the point person (in years) (আপনার খানার যে ব্যক্তি আপনাদের চাষাবাদের জমি সম্পর্কে সবচেয়ে ভাল ধারনা রাখেন তার বয়স কত?) (বছরে)_______________________________________________________

1.6 Year of education of the point person (in year, 0-16) [আপনার খানার যে ব্যক্তি আপনাদের চাষাবাদের জমি সম্পর্কে সবচেয়ে ভাল ধারনা রাখেন তার শিৰ বছর?) (বছরে, ০-১৬)]______________________________

1.7 Obtain informed consent. Did you obtain a signed or fingerprinted consent form? সম্মতিপত্র সংগ্রহ করুন। উত্তরদাতার কাছ থেকে স্বাৰর/ আঙ্গুলের ছাপসম্বলিত সম্মতিপত্র সংগ্রহ করেছেন কি?

0=No (না)
1= Yes (হ্যাঁ )

1.8 Interviewer name & code [সাৰাৎকার গ্রহনকারী/ইন্টারভিউয়ারের নাম ও কোড বসান]__________________

1.9 Date of data collection (dd/mm/yyyy) [তথ্য সংগ্রহের তারিখ (দিন/মাস/বছর)]____________________

#### Section-2:Questions for mother (সেকসন-২: মায়ের জন্য প্রশ্নসমূহ)

Note: Ask questions to the mother and the person in the household with the most knowledge regarding the exposure [নোট: টার্গেট শিশুর মা এবং উক্ত খানার অন্যকোন সদস্যকে নিচের প্রশ্নগুলো করুন, যার বিষয়গুলো সম্পর্কে সম্যক ধারনা আছে।]

2.1 How long have you lived in the current location? [বর্তমান এই খানাতে আপনি কতদিন (বছর) যাবত বসবাস করছেন?]________

2.2 Before you lived here, how many far away was your previous residence located? [আপনি এই খানাতে বসবাস করার আগে, আপনার পূর্বের বাসস্থানটি এখান থেকে কত দুরে (অবস্থিত) ছিল?]

1) Less than 1 kilometer (1 কিলোমিটারের কম)
2) 1 or more but less than 5 kilometer ( 1 বা তার বেশি কিন্তু 5 কিলোমিটারের কম)
3) 5 or more but less than 10 kilometer (5 বা তার বেশি কিন্তু 10 কিলোমিটারের কম )
4) 10 or more but less than 20 kilometer (10 বা তার বেশি কিন্তু 20 কিলোমিটারের কম )
5) 20 kilometers or more (20 কিলোমিটার বা তার বেশি)

2.3 How much of the rice that you eat comes from the fields that your household farm? [আপনি/ আপনার খানা সদস্যরা যে পরিমান চাল খান তার কতটুকু আপনার/আপনাদের পরিবারের চাষের জমি থেকে আসে?]

1) None (একটুও না)
2) Some but less than half (সামান্য, অর্ধেকেরও কম)
3) About half (প্রায় অর্ধেক)
4) More than half, but not all (অর্ধেকেরও বেশী, কিন্তু সবটুকু না)
5) All (সবটুকু)

2.4 How much of the vegetables that you eat come from the fields your household farm? [আপনি/ আপনার খানা সদস্যরা যে পরিমান শাকসব্জি খান তার কতটুকু আপনার/আপনাদের পরিবারের চাষের জমি থেকে আসে?]

1) None (একটুও না)
2) Some but less than half (সামান্য, অর্ধেকেরও কম)
3) About half (প্রায় অর্ধেক)
4) More than half, but not all (অর্ধেকেরও বেশী, কিন্তু সবটুকু না)
5) All (সবটুকু)

2.5 How much of the fruits that you eat come from your household farm? [আপনি/ আপনার খানা সদস্যরা যে পরিমান ফলমূল খান তার কতটুকু আপনার/আপনাদের পরিবারের বাগান থেকে আসে?]

1) None (একটুও না)
2) Some but less than half (সামান্য, অর্ধেকেরও কম)
3) About half (প্রায় অর্ধেক)
4) More than half, but not all (অর্ধেকেরও বেশী, কিন্তু সবটুকু না)
5) All (সবটুকু)

2.6 Do you work at least a few times per year with any of these materials? (answer each separately Yes = 1, No = 0) [আপনি কি নিচের উপাদানগুলোর যেকোনটি নিয়ে বছরে অন্ততঃ কয়েকবার কাজ করেন? (প্রত্যেকটি অপশনের ক্ষেত্রে আলাদাভাবে উত্তর দিন। হ্যাঁ=1, না=0)]

1) **Paint (রং)** ___
2) **Solder (ঝালাই)** ___
3) **Industrial waste (শিল্প বর্জ্য)** ___
4) **Truck, car or motorcycle batteries (ট্রাক, গাড়ি বা মোটর সাইকেলের ব্যাটারি)** ___
5) **Manufacturing metals (নির্মান শিল্পের উপাদান)** ___
6) **Solvents (দ্রাবক )** ___
7) **Chemicals ( রাসায়নিক পদার্থসমূহ)** ___
8) **Exhaust fumes (নির্গত ধোঁয়া)** _____
9) **Recycling goods (পূর্নব্যবহারযোগ্য পণ্য)** _____

2.7 Does anyone in your household work at least a few times per year with any of these materials? (answer each separately Yes = 1, No = 0) [ আপনার পরিবারের কোন সদস্য কি নিচের উপাদানগুলোর যেকোনটি নিয়ে বছরে অন্ৰতঃ কয়েকবার কাজ করে? (প্রত্যেকটি অপশনের ৰেত্রে আলাদাভাবে উত্তর দিন। হ্যাঁ=1, না=0)]

1) **Paint (রং)** ___
2) **Solder (ঝালাই)** ___
3) **Industrial waste (শিল্প বর্জ্য)** ___
4) **Truck, car or motorcycle batteries (ট্রাক, গাড়ি বা মোটর সাইকেলের ব্যাটারি)** ___
5) **Manufacturing metals (নির্মান শিল্পের উপাদান)** ___
6) **Solvents (দ্রাবক )** ___
7) **Chemicals ( রাসায়নিক পদার্থসমূহ)** ___
8) **Exhaust fumes (নির্গত ধোঁয়া)** _____
9) **Recycling goods (পূর্নব্যবহারযোগ্য পণ্য)** _____

2.8 How close is the nearest road that motor vehicles use regularly to your home? [নিয়মিতভাবে মটরযান চলাচলের রাস্ৰাটি আপনার বাড়ি থেকে কতটা কাছাকাছি অবস্থিত?]

1) < 50 meters (50 মিটারের কম)
2) 51 – 200 meters (51 থেকে 200 মিটারের মধ্যে)
3) 201 – 500 meters (201 থেকে 500 মিটারের মধ্যে)
4) 501 – 1,000 meters (501 থেকে 1,000 মিটারের মধ্যে)
5) > 1,000 meters (1,000 মিটারের বেশি )

2.9 How often do you consume food that comes from a metal can? [তদিন পর পর আপনি ধাতবপাত্রে প্রক্রিয়াজাত করা খাবার কিনে থাকেন?]

1) Never (কখনোই না)

2) A few times per year (বছরে কয়েক বার)
3) About once per month (প্রায় প্রতি মাসে এক বার)
4) About once per week (প্রায় প্রতি সপ্তাহে এক বার)
5) Most days (অধিকাংশ দিনই)

2.10 How often do you wear kohl (makeup)/ Sindoor that you purchase from a street vendor or a store? [আপনি কি প্রায়ই দোকান থেকে কেনা সুর্মা (মেকআপ)/ সিদুঁর ব্যবহার করেন?]

1) Never (কখনোই না)
2) A few times per year (বছরে কয়েক বার)
3) About once per month (প্রায় প্রতি মাসে এক বার)
4) About once per week (প্রায় প্রতি সপ্তাহে এক বার)
5) Most days (অধিকাংশ দিনই)

2.11 Does your house have tin roofs? [আপনার ঘরের ছাদ কি টিনের তৈরি?]

**1) Yes (হ্যাঁ)**
**0) No (না)**

2.12 If yes, how long have you had a tin roof? [উত্তর হ্যাঁ হলে, কতদিন (বছর) থেকে আপনার ঘরের ছাদ টিনের তৈরি?]_______

2.13 Does your house have tin walls? [আপনার ঘরের দেয়াল কি টিনের তৈরি?]
1) In all rooms (সব ঘরে)
2) In at least one room (কমপৰে একটি ঘরে)
3) In zero room (কোন ঘরে নাই) [Skip to 2.15]

2.14 How long have you had tin walls? [কতদিন (বছর) থেকে আপনার ঘরের দেয়াল টিনের তৈরি?]_______

2.15 What materials are your cooking pot made from? (answer each separately Yes = 1, No = 0) [আপনার রান্নার হাঁড়ি-পাতিল কোন উপাদান দিয়ে তৈরি? (প্রত্যেকটি অপশনের ৰেত্রে আলাদাভাবে উত্তর দিন । হ্যাঁ=1, না=0)]

1) Aluminum (অ্যালুমিনিয়াম)
2) Steel (স্টীল)
3) Unpainted clay / ceramic (রংবিহীন কাদামাটি/ সিরামিক)
4) Painted clay / ceramic (রং করা কাদামাটি/ সিরামিক)
5) Cast iron (ঢালাই লোহা)

2.16 What materials are your plates, serving dishes and utensils made from? (answer each separately Yes = 1, No = 0) [আপনার খাবারের থালা, খাবার পরিবেশনের ডিশ এবং হাঁড়ি-পাতিল, পানি রাখার/সংরক্ষণের/ধারণের বা পান করার পাত্র কিসের তৈরি? (প্রত্যেকটি অপশনের ক্ষেত্রে আলাদাভাবে উত্তর দিন । হ্যাঁ=1, না=0)]

1) Melamine (মেলামাইন)
2) Steel (স্টীল)
3) Aluminum (অ্যালুমিনিয়াম)
4) Unpainted clay / ceramic (রংবিহীন কাদামাটি/ সিরামিক)
5) Painted clay / ceramic (রং করা কাদামাটি/ সিরামিক)
6) Cast iron (ঢালাই লোহা)
7) Plastic (পস্নাষ্টিক)

2.17 How often do you wear a bracelet? [কতদিন পর পর আপনি আপনার হাতে ব্রেসলেট/চুরি পরেন?]

1) Every day (প্রতি দিন/ নিয়মিত/সবসময়)
2) A few days per week (সপ্তাহে কয়েকদিন)
3) Once a week (সপ্তাহে একদিন)
4) Less than once per week (প্রতি সপ্তাহে একদিনেরও কম)
5) Never (কখনোই না) [Skip to 2.17a]

2.18 What materials is your bracelet(s) made from?(answer each separately Yes = 1, No = 0) [আপনার ব্রেসলেট/চুরি কিসের তৈরি? (প্রত্যেকটি অপশনের ক্ষেত্রে আলাদাভাবে উত্তর দিন । হ্যাঁ=1, না=0)]

1) City gold (সিটি গোল্ড)
2) Gold (গোল্ড/স্বর্ন)
3) Sliver (সিলভার/ রূপা)
4) Copper(তামা)
5) Brass (পিতল)
77) Other (অন্যান্য) __________
99) Don't know (জানিনা)

2.17 a. How often do you wear a ear rings? [কতদিন পর পর আপনি আপনার কানে দুল পরেন?]

1) Every day (প্রতি দিন/ নিয়মিত/সবসময়)
2) A few days per week (সপ্তাহে কয়েকদিন)
3) Once a week (সপ্তাহে একদিন)
4) Less than once per week (প্রতি সপ্তাহে একদিনেরও কম)
5) Never (কখনোই না) [Skip to 2.17b]

2.18 a What materials is your ear rings made from?(answer each separately Yes = 1, No = 0) [আপনার কানের দুল কিসের তৈরি? (প্রত্যেকটি অপশনের ক্ষেত্রে আলাদাভাবে উত্তর দিন । হ্যাঁ=1, না=0)]

1) City gold (সিটি গোল্ড)
2) Gold (গোল্ড/স্বর্ন)
3) Sliver (সিলভার/ রূপা)

4) Copper(তামা)
5) Brass (পিতল)
77) Other (অন্যান্য) __________
99) Don't know (জানিনা)

2.17 b. How often do you wear a necklace? [কতদিন পর পর আপনি আপনার গলায় হার পরেন?]

1) Every day (প্রতি দিন/ নিয়মিত/সবসময়)
2) A few days per week (সপ্তাহে কয়েকদিন)
3) Once a week (সপ্তাহে একদিন)
4) Less than once per week (প্রতি সপ্তাহে একদিনেরও কম)
5) Never (কখনোই না) [Skip to 2.19]

2.18 b What materials is your necklace made from?(answer each separately Yes = 1, No = 0)
[আপনার গলার হার কিসের তৈরি? (প্রত্যেকটি অপশনের ৰেত্রে আলাদাভাবে উত্তর দিন । হ্যাঁ=1, না=0)]

1) City gold (সিটি গোল্ড)
2) Gold (গোল্ড/স্বর্ন)
3) Sliver (সিলভার/ রূপা)
4) Copper(তামা)
5) Brass (পিতল)
77) Other (অন্যান্য) __________
99) Don't know (জানিনা)

2.19 How often do you wear a nose pin? [আপনি কতদিন পর পর নাকফুল পরেছেন?]

1) Every day (প্রতি দিন/ নিয়মিত/সবসময়)
2) A few days per week (সপ্তাহে কয়েকদিন)
3) Once a week (সপ্তাহে একদিন)
4) Less than once per week (প্রতি সপ্তাহে একদিনেরও কম)
5) Never (কখনোই না) [Skip to 2.21]

2.20 What materials is your nose pin(s) made from(answer each separately Yes = 1, No = 0)
[আপনার নাকফুল কিসের তৈরি? (প্রত্যেকটি অপশনের ৰেত্রে আলাদাভাবে উত্তর দিন । হ্যাঁ=1, না=0)]

1) City gold (সিটি গোল্ড)
2) Gold (গোল্ড/স্বর্ন)
3) Sliver (সিলভার/ রূপা)
4) Copper(তামা)
77) Other (অন্যান্য) __________

99) Don't know (জানিনা)

2.21 How often did you wear an amulet during your recent pregnancy? [সর্বশেষ গর্ভবতী থাকাকালীন সময়ে আপনি কত দিন পর পর মাদুলী/কবচ পরেছেন?]

1) Every day (প্রতি দিন/ নিয়মিত/সবসময়)
2) A few days per week (সপ্তাহে কয়েকদিন)
3) Once a week (সপ্তাহে একদিন)
4) Less than once per week (প্রতি সপ্তাহে একদিনেরও কম)
5) Never (কখনোই না) [Skip to 2.23]

2.22 What materials was your amulet(s) made from? (answer each separately Yes = 1, No = 0) [আপনার মাদুলী/কবচ কিসের তৈরি? (প্রত্যেকটি অপশনের ৰেত্রে আলাদাভাবে উত্তর দিন । হ্যাঁ=1, না=0)]

1) City gold (সিটি গোল্ড)
2) Gold (গোল্ড/স্বর্ন)
3) Sliver (সিলভার/ রূপা)
4) Copper(তামা)
5) Brass (পিতল)
6) Austodhatu (Eight metals) (অষ্টধাতু)
7) Iron (লোহা)
77) Other (অন্যান্য) __________
99) Don't know (জানিনা)

2.23 Do you cultivate fish from ponds on your land? [আপনার/আপনাদের পুকুরে মাছের চাষ করেন কি?]

1) Yes (হ্যাঁ)
0) No (না)
99) Don't know (জানি না)

2.24 How many days per week does your family eat fish? [আপনার খানার সদস্যরা সপ্তাহে কয়দিন মাছ খায়?]

1) Less then 1 day per week (সপ্তাহে 1 দিনের কম)
2) 1 day per week (সপ্তাহে ১ দিন)
3) 2 to 4 days per week (সপ্তাহে 2 থেকে 4 দিন)
4) More than 4 days per week (সপ্তাহে 4 দিনের বেশি)
99) Don't know (জানি না)

2.25 How many kilometers away is the closest brick kiln to your agricultural field? [আপনার কৃষিজমি থেকে সবচেয়ে কাছের ইটের ভাটাটি কত কিলোমিটার দুরে অবস্থিত?]___________

2.26 In the last year have you applied any paint to any surface? [গত বছরে আপনাদের কোন তল/পৃষ্ঠ মেঝে/দেয়াল/ছাদ) রং করেছেন কি?]

1) Yes (হ্যাঁ)
0) No (না) [Skip to 2.30]
99) Don't know (জানি না) [Skip to 2.30]

2.27 Which of the following items in your household are painted? (answer each 1 = yes, 0 = no) [আপনার খানাতে নিচের কোনটি রং করা হয়েছে? (প্রত্যেকটি অপশনের ৰেত্রে আলাদাভাবে উত্তর দিন । হ্যাঁ=1, না=0)]

1) Walls (দেয়াল)
2) Food containers (খাবার রাখার পাত্র)
3) Water containers (পানি রাখার পাত্র)
4) Furniture (আসবাব-পত্র)
77) Other (অন্যান্য) _________

2.28 Which of the following items in your household did someone in the household paint? (answer each 1 = yes, 0 = no) [নিচের কোনটি আপনার খানার সদস্য রং করেছে? (প্রত্যেকটি অপশনের ৰেত্রে আলাদাভাবে উত্তর দিন। হ্যাঁ=1, না=0)]

1) Walls (দেয়াল)
2) Food containers (খাবার রাখার পাত্র)
3) Water containers (পানি রাখার পাত্র)
4) Furniture (আসবাব-পত্র)
77) Other (অন্যান্য) _________

2.29 Which of the following items in your household is there paint that is chipped and sometimes flakes off? (answer each 1 = yes, 0 = no) [আপনার খানাতে নিচের কোনটি রং করার সময় রংয়ের ছোট টুকরা/ফোটা পড়েছিল এবং (মেঝেতে) আটকে ছিল? (প্রত্যেকটি অপশনের ৰেত্রে আলাদাভাবে উত্তর দিন। হ্যাঁ=1, না=0)]

1) Walls (দেয়াল)
2) Food containers (খাবার রাখার পাত্র)

3) Water containers (পানি রাখার পাত্র)
4) Furniture (আসবাব-পত্র)
77) Other (অন্যান্য) _________

2.30 Do you grow any spices on your land? [আপনি কি আপনার জমিতে মসলা উৎপাদন করেন?]

1) Yes (হ্যাঁ)
0) No (না)

2.31 (If 2.30 is yes) How far away is the closest spice ground to your household? [(যদি ২.৩০ এর উত্তর হাঁ হয়) সবচেয়ে কাছের জমিটি আপনার বাসস্থান থেকে কত দুরে অবস্থিত?]

1) < 50 meters (50 মিটারের কম)
2) 51 – 200 meters (51 থেকে 200 মিটারের মধ্যে)
3) 201 – 500 meters (201 থেকে 500 মিটারের মধ্যে)
4) 501 – 1,000 meters (501 থেকে 1,000 মিটারের মধ্যে)
5) > 1,000 meters (1,000 মিটারের বেশি )

2.32 How do you consume/purchase spices? (answer each 1 = yes, 0 = no) [আপনি কোন ধরণের মসলা কেনেন বা ব্যবহার করেন? (প্রত্যেকটি অপশনের ৰেত্রে আলাদাভাবে উত্তর দিন। হ্যাঁ=1, না=0)]

1) Raw spices from local vendor (স্থানীয় দেকান থেকে খোলা মসলা)
2) Packaged spices (প্যাকেট মসলা)
3) Process from local grinding mills (স্থানীয় মসলা গুড়া করার মিল থেকে ভাঙ্গিয়ে নেই)
4) Process in own home (বাড়িতেই প্রক্রিয়/প্রস্ত্বুত করে নেই)
77) Other (অন্যান্য) _________

2.33 (If 2.32 is 2) How often do you purchase packaged spice? [(যদি ২.৩২ এর উত্তর ২ হয়) আপনি কত ঘন ঘন প্যাকেট মসলা কেনেন?]

1) All time (নিয়মিত/সবসময়)
2) Most of the times (অধিকাংশ সময়)
3) Sometimes (কখনো কখনো)
99) Don't knড়(জানি না)

2.34 How do you purchase turmeric? (answer each 1 = yes, 0 = no) [আপনি কোন ধরণের হলুদ কেনেন বা ব্যবহার করেন? (প্রত্যেকটি অপশনের ৰেত্রে আলাদাভাবে উত্তর দিন। হ্যাঁ=1, না=0)]

1) Raw turmeric from local vendor (স্থানীয় দেকান থেকে খোলা হলুদ)
2) Packaged turmeric (প্যাকেট হলুদ)
3) Process from local grinding mills (স্থানীয় হলুদ গুড়া করার মিল থেকে ভাঙ্গিয়ে নেই)
4) Process in own home (বাড়িতেই প্রক্রিয়/প্রস্ত্বুত করে নেই)
99) Don't know (জানি না)

2.35 (If 2.34 is 2) I will read the list of available turmeric in the market. Please tell me what branded turmeric did you purchased? (answer each 1 = yes, 0 = no) [(যদি ২.৩৫ এর উত্তর ২ হয়) আমি আপনাকে প্রচলিত প্যাকেট হলুদের একটি তালিকা পড়ে শোনাব। দয়া করে আমাকে বলবেন, এসব প্যাকেট হলুদের মধ্যে কোনটি কোনটি আপনি কেনেন? (প্রত্যেকটি অপশনের ৰেত্রে আলাদাভাবে উত্তর দিন। হ্যাঁ=1, না=0)]

1) ACI (এসিআই)
2) Radhuni (রাধুনী)
3) BD food ( বিডি ফুড)
4) Pran (প্রাণ)
5) Kishwan (কিষাণ)
6) Rupchanda (রুপচাদা)
7) Fresh (ফ্রেশ)

8) Rani (রানী)
9) Non brand packet (নাম ছাড়া প্যাকেট)
77) Other (অন্যান্য) __________

2.36 (If 2.34 is 2) How often do you purchase packaged turmeric? [(যদি ২.৩৪ এর উত্তর ২ হয়) আপনি কত ঘন ঘন প্যাকেট হলুদ কেনেন?]

1) All time (নিয়মিত/সবসময়)
2) Most of the times (অধিকাংশ সময়)
3) Sometimes (কখনো কখনো)
99) Don't know (জানি না)

2.37 Do you grind any crops/spices from mills? [আপনি কি কোন শস্য/মসলা মিল থেকে গুড়া করে নেন?]

1) Yes (হ্যাঁ)
0) No (না) [Skip to Section-3]

2.38 I will read the list of crops/spices. Please tell me what type of crops/spices do you usually grind? (Answer each separately Yes = 1, No = 0) আমি আপনাকে শস্য/মসলার একটি তালিকা পড়ে শোনাব। দয়া করে আমাকে বলবেন, এগুলোর মধ্যে সাধারণত কোনটি কোনটি আপনি মিল থেকে **গুড়া করে** নেন? (প্রত্যেকটি অপশনের ক্ষেত্রে আলাদাভাবে উত্তর দিন। হ্যাঁ=1, না=0)]

1) Rice (চাল)
2) Wheat (গম)
3) Coriander (ধনে)
4) Turmeric (হলুদ)
5) Cumin (জিরা)
6) Red chili pepper (মরিচ)
**77) Other: Specify (অন্যান্য: নির্দিষ্ট করে লিখুন) ________**

2.39 How often do you grind the crops/spice from mills? (.............times in a year) [please answer seperately] [আপনি এক বছরে কতবার শস্য/মসলা মিল থেকে গুড়া করে নেন? (দয়া করে প্রত্যেকটি অপশনের ক্ষেত্রে আলাদাভাবে উত্তর দিন।]

1) Rice (চাল)
2) Wheat (গম)
3) Coriander (ধনে)
4) Turmeric (হলুদ)

5) Cumin (জিরা)
6) Red chili pepper (মরিচ)
**77) Other: Specify (অন্যান্যঃ নির্দিষ্ট করে লিখুন) ________**

2.40 How far away is the closest grinding mills to your household? [সবচেয়ে কাছের শস্য/মসলা গুড়া করার মিলটি আপনার বাসস্থান থেকে কত দুরে অবস্থিত?]

1) Less than 1 kilometer (1 কিলোমিটারের কম)
2) 1 or more but less than 3 kilometer ( 1 বা তার বেশি কিন্তু 3 কিলোমিটারের কম)
3) 3 or more but less than 5 kilometer ( 3 বা তার বেশি কিন্তু 5 কিলোমিটারের কম)
4) 5 or more but less than 10 kilometer ( 5 বা তার বেশি কিন্তু 10 কিলোমিটারের কম)
5) 10 or more kilometer ( 10 বা তার বেশি কিলোমিটার)

2.41 How long ago the grinding mills established that you mostly use? .............in years [আপনি শস্য/মসলা গুড়া করার জন্য সাধারণত যে মিলটি ব্যবহার করেন তা কত বছর আগে স্থাপিত হয়েছিল?]

#### Section-3: Questions for person most knowledgeable regarding household farming (সেকসন-৩ঃ পারিবারিক চাষাবাদ সম্বন্ধে জ্ঞানবুদ্ধিসম্পন্ন ব্যক্তির জন্য প্রশ্নাবলী)

3.1 How many harvests of rice does your land produce each year? [আপনার জমিতে প্রতিবছর কয় ধরনের ধান উৎপাদন করা হয়?] ____

3.2 What other crops do you grow on your land (including small gardens)? (answer each separately Yes = 1, No = 0). [আপনার জমিতে ধান ছাড়া আর কি কি শস্য উৎপাদন করে থাকেন (ছোট বাগানসহ) ? (প্রত্যেকটি শস্যের ৰেত্রে আলাদাভাবে উত্তর দিন । হ্যাঁ=1, না=0)]

1) Lentils (মসুর ডাল)
2) Chick peas (chola)( ছোলা)
3) Eggplant (বেগুন)
4) Tomato (টমেটো)
5) Cucumbers(শসা)
6) Lady fingers (ঢেঁড়স)
7) Beans (সীম/মটরশুঁটি)
8) Red spinach(লালশাঁক)
*9) Kolmi shak (কলমীশাঁক)*

10) Water guard (লাউ)
11) Pumpkin (কুমড়া)
12) Snake gord (চিঁচিঙ্গা)
13) Onions (পেঁয়াজ)
14) Carrots (গাজর)
15) Radish (মূলা)
16) Potatoes (আলু)
17) Mango (আম)
18) Komla (কমলা)
19) Lychee (লিচু)
20) Jackfruit (কাঁঠাল)
21) Banana (কলা)
22) Garlic (রসুন)
23) Ginger (আদা)
24) Turmeric (হলুদ)
25) Spinach (পালং শাঁক)
26) Guava (পেয়ারা)
27) Papaya (পেঁপে)
28) Data (ডাটা শাঁক)
77) Other : Specify (অন্যান্যঃ নির্দ্দিষ্ট করে লিখুন)

#### Section-4: Fertilizer Questions (সেকসন-৪: সার সংক্রান্ত প্রশ্নাবলী )

4.1 How many times did you apply fertilizer on your land in the last year? [গত বছর আপনার জমিতে কতবার সার প্রয়োগ করেছেন?] _____

4.2 I will read a list of available fertilizers. Please tell me how many times you applied each of these types of fertilizer on the land you used for growing rice in the last year. [আমি আপনাকে প্রচলিত সারের একটি তালিকা পড়ে শোনাব। দয়া করে আপনি আমাকে বলবেন, এসব সারের মধ্যে কোন কোন সার গত বছর আপনি কতবার আপনার জমিতে ধান উৎপাদনের জন্য প্রয়োগ/ব্যবহার করেছেন?]

1) Urea (ইউরিয়া)
2) Triple super phosphate (TSP)[ ট্রিপল সুপার ফসফেট (টিএসপি)]
3) Diammonium phosphate(DSP) [ ডায়ামনিয়াম ফসফেট (ডিএসপি)]
4) Muriate of potash (MOP) [পটাশ]
5) Gypsum (জিপসাম)
6) Boron (বোরন)
7) Vitamin (ভিটামিন)

8) Cow dung (গোবর সার)
9) Oil cake (খৈল)
77) Other: Specify (অন্যান্য: নির্দিষ্ট করে লিখুন) ________

4.3 I will again read the list of available fertilizers. Please tell me how many times you applied each of these types of fertilizer on the land you used for growing other crops in the last year. [আমি আপনাকে পুনরায় প্রচলিত সারের তালিকাটি পড়ে শোনাব। দয়া করে আপনি আমাকে বলবেন, এসব সারের মধ্যে কোন কোন সার গত বছর আপনি কতবার আপনার জমিতে অন্যান্য শস্য (ধান ছাড়া) উৎপাদনের জন্য প্রয়োগ/ব্যবহার করেছেন?]

1) Urea (ইউরিয়া)
2) Triple super phosphate (TSP)[ ট্রিপল সুপার ফসফেট (টিএসপি)]
3) Diammonium phosphate(DSP) [ ডায়ামনিয়াম ফসফেট (ডিএসপি)]
4) Muriate of potash (MOP) [পটাশ]
5) Gypsum (জিপসাম)
6) Boron (বোরন)
7) Vitamin (ভিটামিন)
8) Cow dung (গোবর সার)
9) Oil cake (খৈল)
77) Other: Specify (অন্যান্য: নির্দিষ্ট করে লিখুন) ________

4.4 Who handled the fertilizer while preparing to apply it? (answer each separately Yes = 1, No = 0) [সারগুলো জমিতে প্রয়োগ করার পূর্বে কে এগুলো প্রস্তুত করেন/নাড়াচাড়া করেন? (প্রত্যেকটি অপশনের ৰেত্রে আলাদাভাবে উত্তর দিন । হ্যাঁ=1, না=0)]

**1) Father of the youngest child in the household (সবচেয়ে ছোট শিশুর বাবা)**
**2) Mother of the youngest child in the household (সবচেয়ে ছোট শিশুর মা)**
**3) Sibling of the youngest child in the household (সবচেয়ে ছোট শিশুর সহোদর)**
**4) Grandparent of the youngest child in the household (সবচেয়ে ছোট শিশুর দাদা-দাদী)**
**5) Cousin of the youngest child in the household (সবচেয়ে ছোট শিশুর চাচাতো ভাই-বোন)**
**6) Uncle of the youngest child in the household (সবচেয়ে ছোট শিশুর চাচা/মামা)**
**7) Aunt of the youngest child in the household (সবচেয়ে ছোট শিশুর চাচী/মামী)**

8) **Day labour (দিন মজুর)**
77) **Other: Specify (অন্যান্য: নির্দিষ্ট করে লিখুন) ________**

4.5 Where was the fertilizer prepared for application? (answer each separately Yes = 1, No = 0) [সারগুলো প্রয়োগ করার/ছিটানোর জন্য কোথায় প্রস্তুত করা হয়েছিল? (প্রত্যেকটি অপশনের ক্ষেত্রে আলাদাভাবে উত্তর দিন । হ্যাঁ=1, না=0)]

1) **Inside the house (ঘরের ভিতরে)**
2) **In the courtyard (আঙিনায়/উঠানে)**
3) **Near the house (বাড়ির কাছে)**
4) **In the field (জমিতে)**
77) **Other: Specify (অন্যান্য: নির্দিষ্ট করে লিখুন) ________**

4.6 Who applied the fertilizer to the land? (answer each separately Yes = 1, No = 0) [কে সারগুলো জমিতে দিয়েছিল? (প্রত্যেকটি অপশনের ক্ষেত্রে আলাদাভাবে উত্তর দিন । হ্যাঁ=1, না=0)]

1) **Father of the youngest child in the household (সবচেয়ে ছোট শিশুর বাবা)**
2) **Mother of the youngest child in the household (সবচেয়ে ছোট শিশুর মা)**
3) **Sibling of the youngest child in the household (সবচেয়ে ছোট শিশুর সহোদর)**
4) **Grandparent of the youngest child in the household (সবচেয়ে ছোট শিশুর দাদা-দাদী)**
5) **Cousin of the youngest child in the household (সবচেয়ে ছোট শিশুর চাচাতো ভাই-বোন)**
6) **Uncle of the youngest child in the household (সবচেয়ে ছোট শিশুর চাচা/মামা)**
7) **Aunt of the youngest child in the household (সবচেয়ে ছোট শিশুর চাচী/মামী)**
8) **Day labor (দিন মজুর)**
77) **Other: Specify (অন্যান্য: নির্দিষ্ট করে লিখুন) ________**

4.7 How did they apply the fertilizer? (answer each separately Yes = 1, No = 0) [সারগুলো কিভাবে জমিতে দিয়েছিল? ]

1) With their hands (হাতের সাহায্যে)
2) With a tool (যন্ত্রের সাহায্যে)
3) Mixed with water and sprayed (পানিতে মিশিয়েছে এবং ছিটিয়ে দিয়েছে)

**77) Other: Specify (অন্যান্য: নির্দিষ্ট করে লিখুন) ________**

4.8 What did you do with the leftover fertilizer? [জমিতে ব্যবহারের পর অবশিষ্ট সারগুলো আপনি কি করেছিলেন? ]

1) Did not have any (কোন সার অবশিষ্ট ছিল না)
2) Stored it in the house (বাড়িতে সংরক্ষন করেছিলাম)
3) Stored it outside of the house (বাড়ির বাইরে সংরক্ষন করেছিলাম)
4) Gave it away or sold it (অন্য কাউকে দিয়ে দিয়েছিলাম অথবা বিক্রি করে দিয়েছিলাম)
5) Apply in another land (উক্ত খানার অন্যকোন জমিতে প্রয়োগ করে)
6) Use in the pond (পুকুরে প্রয়োগ করে)
77) Other: Specify (অন্যান্য: নির্দিষ্ট করে লিখুন) ________

**4.9 What did you do with the leftover fertilizer packaging? (answer each separately Yes = 1, No = 0) জমিতে ব্যবহারের পর সারের অবশিষ্ট প্যাকেটগুলো আপনি কি করেছিলেন? (প্রত্যেকটি অপশনের ক্ষেত্রে আলাদাভাবে উত্তর দিন । হ্যাঁ=1, না=0) [What did you do with the leftover fertilzer packaging? (answer each separately Yes = 1, No = 0]**

1) Stored it inside the house (বাড়িতে সংরক্ষন করেছিলাম)
2) Stored it outside the house (বাড়ির বাইরে সংরক্ষন করেছিলাম)
3) Used it to store food (খাদ্য মজুদ রাখার কাজে ব্যবহার করেছিলাম)
4) Used it to store water (পানি সংরক্ষনের কাজে ব্যবহার করেছিলাম)
5) Used it to store household goods (গৃহস্থালী সামগ্রী সংরক্ষনের কাজে ব্যবহার করেছিলাম)
6) Let the children play with it (শিশুদের খেলতে দিয়েছিলাম)
7) Use as shopping bag (সপিং/ বাজারের ব্যাগ হিসাবে ব্যবহার করে)
8) Discarded it (ফেলে দিয়েছিলাম)
9) Used it to store leftover fertilizer (বাড়িতে সার সহ সংরক্ষন করেছিলাম)
10) Washed the packet before use (ধুয়ে ব্যবহার করেছিলাম)

**Section-5: Pesticide Questions (সেকসন-৫:কীটনাশক বিষয়ক প্রশ্নাবলী)**

**Prompt: Now I would like to ask you few question about your insect control practice, if your land/crops infected with insects what kind of control measure you had taken to keep your crops/lands safe-**

**[প্রমপট: এখন আমি অপনাকে কীটপতঙ্গ/ পোকা-মাকড় নিয়ন্ত্রন অভ্যাস সম্পর্কে কিছু প্রশ্ন করতে চাই, যদি আপনার ফসলের জমিতে কীটপতঙ্গ/ পোকা-মাকড় আক্রমন করে থাকে তবে তা থেকে আপনার জমি/ ফসল রক্ষা করতে আপনি কি পদক্ষেপ নিয়েছিলেন-]**

5. Did you apply pesticide on your land in the last year? (আপনি কি গত বছর আপনার জমিতে কীটনাশক প্রয়োগ করেছেন?]

0=No (না) [Skip to Section-6]
1= Yes (হ্যাঁ )

5.1 How many times did you apply pesticide on your land in the last year? [গত বছর আপনার জমিতে কতবার কীটনাশক প্রয়োগ করেছেন?] _____

5.2 I will read a list of available pesticides. Please tell me how many times you applied each of these types of pesticide on the land you used for growing rice in the last year. [আমি আপনাকে প্রচলিত কীটনাশকের একটি তালিকা পড়ে শোনাব। দয়া করে আপনি আমাকে বলবেন, এসব কীটনাশকের মধ্যে কোন কোন কীটনাশক গত বছর আপনি কতবার আপনার জমিতে ধান উৎপাদনের জন্য প্রয়োগ/ব্যবহার করেছেন?]

1) Basudin (বাসুডিন)
2) Diazinon (ডায়াজিনন)
3) Bristorly (ব্রিসটরলি)
4) Sunfuran (সানফরান)
5) Karate (ক্যারাটে)
6) Cidial (সিডিয়াল)
7) Brifur (ব্রাইফার)
8) Furadan (ফুরাডান)
9) Ripcord (রিপকর্ড)
10) Virtako (ভিরট্যাকো)
11) Voliom flexi (ভলিওম ফ্লেক্সি)
12) Proclaim (প্রক্লেইম)
13) Actara (এ্যাকটারা)
14) Platinum (পস্নাটিনাম)
15) Cruiser (ক্রুইজার)
16) Vertimec (ভারটিমেক)
17) Tilt (টিল্ট)

18) Amistar (অ্যামিস্টার)
19) Amistar Top (অ্যামিস্টার টপ)
**77) Other: Specify (অন্যান্য: নির্দ্দিষ্ট করে লিখুন) ________**

5.3 I will again read the list of available pesticides. Please tell me how many times you applied each of these types of pesticide on the land you used for growing other crops in the last year. [আমি আপনাকে পূনরায় প্রচলিত কীটনাশকের তালিকাটি পড়ে শোনাব। দয়া করে আপনি আমাকে বলবেন, এসব কীটনাশকের মধ্যে কোন কোন কীটনাশক গত বছর আপনি কতবার আপনার জমিতে অন্যান্য শস্য (ধান ছাড়া) উৎপাদনের জন্য প্রয়োগ/ব্যবহার করেছেন?]

1) Basudin (বাসুডিন)
2) Diazinon (ডায়াজিনন)
3) Bristorly (ব্রিসটরলি)
4) Sunfuran (সানফরান)
5) Karate (ক্যারাটে)
6) Cidial (সিডিয়াল)
7) Brifur (ব্রাইফার)
8) Furadan (ফুরাডান)
9) Ripcord (রিপকর্ড)
10) Virtako (ভারট্যাকো)
11) Voliom flexi (ভলিওম ফ্লেক্সি)
12) Proclaim (প্রক্লেইম)
13) Actara (এ্যাকটারা)
14) Platinum (পস্নাটিনাম)
15) Cruiser (ক্রুইজার)
16) Vertimec (ভারটিমেক)
17) Tilt (টিল্ট)
18) Amistar (অ্যামিস্টার)
19) Amistar Top (অ্যামিস্টার টপ)
**77) Other: Specify (অন্যান্য: নির্দ্দিষ্ট করে লিখুন) ________**

5.4 Who handled the pesticide while preparing to apply it? answer each separately Yes = 1, No = 0) [কীটনাশকগুলো জমিতে প্রয়োগ করার পূর্বে কে এগুলো প্রস্তুত করেন/নাড়াচাড়া করেন? (প্রত্যেকটি অপশনের ৰেত্রে আলাদাভাবে উত্তর দিন । হ্যাঁ=1, না=0)]

**1) Father of the youngest child in the household (সবচেয়ে ছোট শিশুর বাবা)**
**2) Mother of the youngest child in the household (সবচেয়ে ছোট শিশুর মা)**
**3) Sibling of the youngest child in the household (সবচেয়ে ছোট শিশুর সহোদর)**

4) **Grandparent of the youngest child in the household (সবচেয়ে ছোট শিশুর দাদা-দাদী)**
5) **Cousin of the youngest child in the household (সবচেয়ে ছোট শিশুর চাচাতো ভাই-বোন)**
6) **Uncle of the youngest child in the household (সবচেয়ে ছোট শিশুর চাচা/মামা)**
7) **Aunt of the youngest child in the household (সবচেয়ে ছোট শিশুর চাচী/মামী)**
8) **Day labor (দিন মজুর)**
77) **Other: Specify (অন্যান্য: নির্দ্দিষ্ট করে লিখুন) _______**

---

5.5 Where was the pesticide prepared for application? (answer each separately Yes = 1, No = 0) [কীটনাশকগুলো প্রয়োগ করার/ছিটানোর জন্য কোথায় প্রস্তুত করা হয়েছিল? (প্রত্যেকটি অপশনের ক্ষেত্রে আলাদাভাবে উত্তর দিন । হ্যাঁ=1, না=0)]

1) **Inside the house (ঘরের ভিতরে)**
2) **In the courtyard (আঙিনায়/উঠানে)**
3) **Near the house (বাড়ির কাছে)**
4) **In the field (জমিতে)**
77) **Other: Specify (অন্যান্য: নির্দ্দিষ্ট করে লিখুন) _______**

---

5.6 Who applied the pesticide to the land? (answer each separately Yes = 1, No = 0) [কে কীটনাশকগুলো জমিতে দিয়েছিল? (প্রত্যেকটি অপশনের ক্ষেত্রে আলাদাভাবে উত্তর দিন । হ্যাঁ=1, না=0)]

1) **Father of the youngest child in the household (সবচেয়ে ছোট শিশুর বাবা)**
2) **Mother of the youngest child in the household (সবচেয়ে ছোট শিশুর মা)**
3) **Sibling of the youngest child in the household (সবচেয়ে ছোট শিশুর সহোদর)**
4) **Grandparent of the youngest child in the household (সবচেয়ে ছোট শিশুর দাদা-দাদী)**
5) **Cousin of the youngest child in the household (সবচেয়ে ছোট শিশুর চাচাতো ভাই-বোন)**
6) **Uncle of the youngest child in the household (সবচেয়ে ছোট শিশুর চাচা/মামা)**
7) **Aunt of the youngest child in the household (সবচেয়ে ছোট শিশুর চাচী/মামী)**
8) **Day labor (দিন মজুর)**
77) **Other: Specify (অন্যান্য: নির্দ্দিষ্ট করে লিখুন) _______**

---

5.7 How did they apply the pesticide? (answer each separately Yes = 1, No = 0) [কীটনাশকগুলো কিভাবে জমিতে দিয়েছিল? (প্রত্যেকটি অপশনের ৰেত্রে আলাদাভাবে উত্তর দিন । হ্যাঁ=1, না=0)]

1) With their hands (হাতের সাহায্যে)
2) With a tool (যন্ত্রের সাহায্যে)
3) Mixed with water and sprayed (পানিতে মিশিয়েছে এবং ছিটিয়ে দিয়েছে)
4) Mixed with fertilizer and sprayed (সারের সাথে মিশিয়েছে এবং ছিটিয়ে দিয়েছে)
77) **Other: Specify (অন্যান্য: নির্দ্দিষ্ট করে লিখুন) ________**

---

5.8 What did you do with the leftover pesticide? [জমিতে ব্যবহারের পর অবশিষ্ট কীটনাশকগুলো আপনি কি করেছিলেন?]

1) **অবশিষ্ট ছিল না (Did not have any)**
2) **বাড়িতে সংরৰন করেছিলাম (Stored it in the house)**
3) **কাউকে দান করে দিয়েছিলাম বা বিক্রি করে দিয়েছিলাম (Gave it away or sold it)**
77) **Other: Specify (অন্যান্য: নির্দ্দিষ্ট করে লিখুন) ________**

---

5.9 What did you do with the leftover pesticide packaging/bottles? (answer each separately Yes = 1, No = 0) [জমিতে ব্যবহারের পর কীটনাশকের অবশিষ্ট প্যাকেটগুলো/বোতলগুলি আপনি কি করেছিলেন? (প্রত্যেকটি অপশনের ৰেত্রে আলাদাভাবে উত্তর দিন । হ্যাঁ=1, না=0)]

1) **Stored it at home (বাড়িতে সংরৰন করেছিলাম)**
2) **Used it to store food (খাদ্য মজুদ রাখার কাজে ব্যবহার করেছিলাম)**
3) **Used it to store water (পানি সংরৰনের কাজে ব্যবহার করেছিলাম)**
4) **Used it to store household goods (গৃহস্থালী সামগ্রী সংরৰনের কাজে ব্যবহার করেছিলাম)**
5) **Let the children play with it (শিশুদের খেলতে দিয়েছিলাম)**
6) **Buried under the ground (মাটির নিচে পুতে ফেলেছিলাম)**
7) **Hanged on the field with stick (লাঠির সাহায্যে জমিতে ঝুলিয়ে রেখেছিলাম)**
8) **Discarded it (ফেলে দিয়েছিলাম)**
9) **Used it to store leftover pesticide (বাড়িতে কীটনাশক সহ সংরৰন করেছিলাম)**
10) **Washed the packet/bottle before use (ধুয়ে ব্যবহার করেছিলাম)**

**Section-6: Herbicide Questions (সেকসন-৬:আগাছানাশক সংক্রান্ত প্রশ্নাবলী)**

**Prompt: please ask the respondents in details about grass/herbs control measere. if your cultivated land have additional grass or herbs that might affect your main crops then what kind of control measure you usually take to keep your crops/lands safe.**
**প্রমপট:অনুগ্রহ করে রেসপনডেনটকে আগাছা নিয়ন্ত্রন কৌশল সম্পর্কে বিস্তারিত প্রশ্ন করুন। যদি আপনার ফসলের জমি আগাছা দ্বারা আক্রান্ত হয় যা প্রধান ফসলকে ক্ষতিগ্রস্থ করতে পারে তবে তা থেকে আপনার জমি/ ফসল রক্ষা করতে আপনি সাধারনত কি পদক্ষেপ গ্রহন করে থাকেন।**

6. Did you apply herbicide on your land in the last year? (আপনি কি গত বছর আপনার জমিতে আগাছানাশক প্রয়োগ করেছেন?]

0=No (না) [Skip to Section-7]
1= Yes (হ্যাঁ )

6.1 How many times did you apply herbicide on your land in the last year? [গত বছর আপনার জমিতে কতবার আগাছানাশক প্রয়োগ করেছেন?] _____

6.2 I will read a list of available herbicides. Please tell me how many times you applied each of these types of herbicide on the land you used for growing rice in the last year. [আমি আপনাকে প্রচলিত আগাছানাশকের একটি তালিকা পড়ে শোনাব। দয়া করে আপনি আমাকে বলবেন, এসব আগাছানাশকের মধ্যে কোন কোন আগাছানাশক গত বছর আপনি কতবার আপনার জমিতে ধান উৎপাদনের জন্য প্রয়োগ/ব্যবহার করেছেন?]

1) Superhit (সুপারহিট)
2) Rifit (রিফিট)
3) Aimchlor 5G (এইমক্লোর ৫জি)
4) Marshal (মার্সাল)
5) Pirilason (পিরিলেসন)
6) Logran (লোগ্রান)
7) Gramoxon (গ্র্যামোক্সন)
77) **Other: Specify (অন্যান্য: নির্দিষ্ট করে লিখুন) ________**

6.3 I will again read the list of available herbicides. Please tell me how many times you applied each of these types of herbicide on the land you used for growing other crops in the last year. [আমি আপনাকে পূনরায় প্রচলিত আগাছানাশকের তালিকাটি পড়ে শোনাব। দয়া করে আপনি আমাকে বলবেন, এসব আগাছানাশকের মধ্যে কোন কোন আগাছানাশক গত বছর আপনি কতবার আপনার জমিতে অন্যান্য শস্য (ধান ছাড়া) উৎপাদনের জন্য প্রয়োগ/ব্যবহার করেছেন?]

1) Superhit (সুপারহিট)
2) Rifit (রিফিট)

3) Aimchlor 5G (এইমক্লোর ৫জি)
4) Marshal (মার্সাল)
5) Pirilason (পিরিলেসন)
6) Logran (লোগ্রান)
7) Gramoxon (গ্র্যামোক্সন)
77) **Other: Specify (অন্যান্য: নির্দিষ্ট করে লিখুন) ________**

6.4 Who handled the herbicide while preparing to apply it? (answer each separately Yes = 1, No = 0) [আগাছানাশক গুলো জমিতে প্রয়োগ করার পূর্বে কে এগুলো প্রস্তুত করেন/নাড়াচাড়া করেন? (প্রত্যেকটি অপশনের ৰেত্রে আলাদাভাবে উত্তর দিন । হ্যাঁ=1, না=0)]

1) **Father of the youngest child in the household (সবচেয়ে ছোট শিশুর বাবা)**
2) **Mother of the youngest child in the household (সবচেয়ে ছোট শিশুর মা)**
3) **Sibling of the youngest child in the household (সবচেয়ে ছোট শিশুর সহোদর)**
4) **Grandparent of the youngest child in the household (সবচেয়ে ছোট শিশুর দাদা-দাদী)**
5) **Cousin of the youngest child in the household (সবচেয়ে ছোট শিশুর চাচাতো ভাই-বোন)**
6) **Uncle of the youngest child in the household (সবচেয়ে ছোট শিশুর চাচা/মামা)**
7) **Aunt of the youngest child in the household (সবচেয়ে ছোট শিশুর চাচী/মামী)**
8) **Day labor (দিন মজুর)**
77) **Other: Specify (অন্যান্য: নির্দিষ্ট করে লিখুন) ________**

6.5 Where was the herbicide prepared for application? (answer each separately Yes = 1, No = 0) [আগাছানাশক গুলো প্রয়োগ করার/ছিটানোর জন্য কোথায় প্রস্তুত করা হয়েছিল? (প্রত্যেকটি অপশনের ৰেত্রে আলাদাভাবে উত্তর দিন । হ্যাঁ=1, না=0)]

1) **ঘরের ভিতরে (Inside the house)**
2) **আঙিনায়/উঠানে (In the courtyard)**
3) **বাড়ির কাছে (Near the house)**
4) **জমিতে (In the field)**
77) **Other: Specify (অন্যান্য: নির্দিষ্ট করে লিখুন) ________**

6.6 Who applied the herbicide to the land? (answer each separately Yes = 1, No = 0) [কে আগাছানাশক গুলো জমিতে দিয়েছিল? (প্রত্যেকটি অপশনের ৰেত্রে আলাদাভাবে উত্তর দিন । হ্যাঁ=1, না=0)]

**1) Father of the youngest child in the household (সবচেয়ে ছোট শিশুর বাবা)**
**2) Mother of the youngest child in the household (সবচেয়ে ছোট শিশুর মা)**
**3) Sibling of the youngest child in the household (সবচেয়ে ছোট শিশুর সহোদর)**
**4) Grandparent of the youngest child in the household (সবচেয়ে ছোট শিশুর দাদা-দাদী)**
**5) Cousin of the youngest child in the household (সবচেয়ে ছোট শিশুর চাচাতো ভাই-বোন)**
**6) Uncle of the youngest child in the household (সবচেয়ে ছোট শিশুর চাচা/মামা)**
**7) Aunt of the youngest child in the household (সবচেয়ে ছোট শিশুর চাচী/মামী)**
**8) Day labor (দিন মজুর)**
**77) Other: Specify (অন্যান্য: নির্দ্দিষ্ট করে লিখুন) ________**

6.7 How did they apply the herbicide? (answer each separately Yes = 1, No = 0) [আগাছানাশকগুলো কিভাবে জমিতে দিয়েছিল? (প্রত্যেকটি অপশনের ৰেত্রে আলাদাভাবে উত্তর দিন । হ্যাঁ=1, না=0)]

1) With their hands (হাতের সাহায্যে)
2) With a tool (যন্ত্রের সাহায্যে)
3) Mixed with water and sprayed (পানিতে মিশিয়েছে এবং ছিটিয়ে দিয়েছে)
4) Mixed with fertilizer and sprayed (সারের সাথে মিশিয়েছে এবং ছিটিয়ে দিয়েছে)
**77) Other: Specify (অন্যান্য: নির্দ্দিষ্ট করে লিখুন) ________**

6.8 What did you do with the leftover herbicide? [জমিতে ব্যবহারের পর অবশিষ্ট আগাছানাশক গুলো আপনি কি করেছিলেন?]

1) Did not have any (অবশিষ্ট ছিল না)
2) Stored it in the house (বাড়িতে সংরৰন করেছিলাম)
3) Stored it outside the house (বাড়ির বাইরে সংরৰন করেছিলাম)
4) Gave it away or sold it (কাউকে দান করে দিয়েছিলাম বা বিক্রি করে দিয়েছিলাম)

**77) Other: Specify (অন্যান্য: নির্দিষ্ট করে লিখুন) _______**

6.9 What did you do with the leftover herbicide packaging? (answer each separately Yes = 1, No = 0) [জমিতে ব্যবহারের পর আগাছানাশকের অবশিষ্ট প্যাকেটগুলো/বোতলগুলি আপনি কি করেছিলেন? (প্রত্যেকটি অপশনের ৰেত্রে আলাদাভাবে উত্তর দিন ।হ্যাঁ=1, না=0)]

1) Stored it in the house (বাড়িতে সংরৰন করেছিলাম)
2) Stored it away from the house (বাড়ির বাইরে সংরৰন করেছিলাম)
3) Used it to store food খাদ্য মজুদ রাখার কাজে ব্যবহার করেছিলাম
4) Used it to store water পানি সংরৰনের কাজে ব্যবহার করেছিলাম
5) Used it to store household goods গৃহস্থালী সামগ্রী সংরৰনের কাজে ব্যবহার করেছিলাম
6) Let the children play with it শিশুদের খেলতে দিয়েছিলাম
7) Burried in the ground (মাটির নিচে পুতে ফেলেছিলাম)
8) Hang in the field with stick লাঠির সাহায্যে জমিতে ঝুলিয়ে রেখেছিলাম)
9) Discarded it ফেলে দিয়েছিলাম
**10) Used it to store leftover herbicide (বাড়িতে আগাছানাশক সহ সংরৰন করেছিলাম)**
**11) Washed the packet/bottle before use (ধুয়ে ব্যবহার করেছিলাম)**

**Section-7: Fungicide Questions (সেকসন-৭: ছত্রাকনাশক ঔষধ সংক্রান্ৰ প্রশ্নাবলী)**

**Prompt: please ask the respondents in details when you notice wilting, mold, rust, blotches, scabs or decaying tissue on your plants or crops then what measure usually you take to keep them safe**
**প্রমপট:অনুগ্রহ করে রেসপনডেনটকে বিস্ৰারিত প্রস্ন করুন, যদি আপনার ফসল নির্জিব হয়ে ঝুলেপড়া, ছাতাপড়া, মারিচার মত/ ফোটা ফোটা দাগপড়া বা টিসুর ক্ষয় জনিত কোন রোগ দ্বরা অক্রান্ৰ হয় যা ফসলকে ক্ষতিগ্রস্থ করতে পারে তবে তা থেকে আপনার জমি/ ফসল রক্ষা করতে আপনি সাধারনত কি পদক্ষেপ গ্রহন করে থাকেন।**

7. Did you apply fungicides on your land in the last year? (আপনি কি গত বছর আপনার জমিতে ছত্রাকনাশক প্রয়োগ করেছেন?]

0=No (না) [Skip to Section-8]
1= Yes (হ্যাঁ )

7.1 How many times did you apply fungicides (chemicals to treat white spots on plant leaves) on your land in the last year? [গত বছর কতবার আপনি আপনার জমিতে ছত্রাকনাশক ঔষধ প্রয়োগ করেছেন?]_____

7.2 I will read a list of available fungicides. Please tell me how many times you applied each of these types of fungicide on the land you used for growing rice in the last year. [আমি আপনাকে প্রচলিত ছত্রাকনাশক ঔষধের একটি তালিকা পড়ে শোনাব। দয়া করে আপনি আমাকে বলবেন, এসব (ছত্রাকনাশক) ঔষধের মধ্যে কোন কোন ধরনের ঔষধ গত বছর আপনি কতবার আপনার জমিতে ধান উৎপাদনের জন্য প্রয়োগ/ব্যবহার করেছেন?]

1) Ridomol gold (রিডমল গোল্ড)
2) Bavistin (ব্যাভিস্টিন)
3) Thiovit (থিওভিট)
4) Salfotox (সালফোটক্স)
5) Haysulf (হেইসাল্ফ)
6) Jazz (জাজ)
7) Hinosum (হিনোসাম)
8) Tilt (টিল্ট)
9) Y-then (ইউ-থেন)
10) Amkojim (এমকোজিম)
77) **Other: Specify (অন্যান্য: নির্দ্দিষ্ট করে লিখুন)** _______

7.3 I will again read the list of available fungicides. Please tell me how many times you applied each of these types of fungicide on the land you used for growing other crops in the last year. [আমি আপনাকে পুনরায় প্রচলিত ছত্রাকনাশক ঔষধের তালিকাটি পড়ে শোনাব। দয়া করে আপনি আমাকে বলবেন, এসব (ছত্রাকনাশক) ঔষধের মধ্যে কোন ধরনের ঔষধ গত বছর আপনি কতবার আপনার জমিতে অন্যান্য শস্য (ধান ছাড়া) উৎপাদনের জন্য প্রয়োগ/ব্যবহার করেছেন?]

1) Ridomol gold (রিডমল গোল্ড)
2) Bavistin (ব্যাভিস্টিন)
3) Thiovit (থিওভিট)
4) Salfotox (সালফোটক্স)
5) Haysulf (হেইসাল্ফ)
6) Jazz (জাজ)
7) Hinosum (হিনোসাম)
8) Tilt (টিল্ট)
9) Y-then (ইউ-থেন)
10) Amkojim (এমকোজিম)
77) **Other: Specify (অন্যান্য: নির্দ্দিষ্ট করে লিখুন)** _______

7.4 Who handled the fungicide while preparing to apply it? (answer each separately Yes = 1, No = 0) [ছত্রাকনাশক ঔষধগুলো জমিতে প্রয়োগ করার পূর্বে কে এগুলো প্রস্তুত করেন/নাড়াচাড়া করেন? (প্রত্যেকটি অপশনের ৰেত্রে আলাদাভাবে উত্তর দিন । হ্যাঁ=1, না=0)]

1) **Father of the youngest child in the household (সবচেয়ে ছোট শিশুর বাবা)**
2) **Mother of the youngest child in the household (সবচেয়ে ছোট শিশুর মা)**
3) **Sibling of the youngest child in the household (সবচেয়ে ছোট শিশুর সহোদর)**
4) **Grandparent of the youngest child in the household (সবচেয়ে ছোট শিশুর দাদা-দাদী)**
5) **Cousin of the youngest child in the household (সবচেয়ে ছোট শিশুর চাচাতো ভাই-বোন)**
6) **Uncle of the youngest child in the household (সবচেয়ে ছোট শিশুর চাচা/মামা)**
7) **Aunt of the youngest child in the household (সবচেয়ে ছোট শিশুর চাচী/মামী)**
8) **Day labor (দিন মজুর)**
77) **Other: Specify (অন্যান্য: নির্দিষ্ট করে লিখুন) ________**

7.5 Where was the fungicide prepared for application(answer each separately Yes = 1, No = 0) [ছত্রাকনাশক ঔষধগুলো প্রয়োগ করার/ছিটানোর জন্য কোথায় প্রস্তুত করা হয়েছিল? (প্রত্যেকটি অপশনের ক্ষেত্রে আলাদাভাবে উত্তর দিন । হ্যাঁ=1, না=0)]

1) **Inside the house (ঘরের ভিতরে)**
2) **(In the courtyard (আঙিনায়/উঠানে)**
3) **Near the house (বাড়ির কাছে)**
4) **In the field (জমিতে)**
77) **Other: Specify (অন্যান্য: নির্দিষ্ট করে লিখুন) ________**

7.6 Who applied the fungicide to the land? (answer each separately Yes = 1, No = 0) [কে ছত্রাকনাশক ঔষধগুলো জমিতে দিয়েছিল? (প্রত্যেকটি অপশনের ক্ষেত্রে আলাদাভাবে উত্তর দিন । হ্যাঁ=1, না=0)]

1) **Father of the youngest child in the household (সবচেয়ে ছোট শিশুর বাবা)**
2) **Mother of the youngest child in the household (সবচেয়ে ছোট শিশুর মা)**
3) **Sibling of the youngest child in the household (সবচেয়ে ছোট শিশুর সহোদর)**
4) **Grandparent of the youngest child in the household (সবচেয়ে ছোট শিশুর দাদা-দাদী)**

**5) Cousin of the youngest child in the household (সবচেয়ে ছোট শিশুর চাচাতো ভাই-বোন)**
**6) Uncle of the youngest child in the household (সবচেয়ে ছোট শিশুর চাচা/মামা)**
**7) Aunt of the youngest child in the household (সবচেয়ে ছোট শিশুর চাচী/মামী)**
**8) Day labor (দিন মজুর)**
**77) Other: Specify (অন্যান্য: নির্দিষ্ট করে লিখুন) ________**

7.7 How did they apply the fungicide? (answer each separately Yes = 1, No = 0) [ছত্রাকনাশক ঔষধগুলো কিভাবে জমিতে দিয়েছিল? (প্রত্যেকটি অপশনের ক্ষেত্রে আলাদাভাবে উত্তর দিন । হ্যাঁ=1, না=0)]

1) With their hands (হাতের সাহায্যে)
2) With a tool (যন্ত্রের সাহায্যে)
3) Mixed with water and sprayed (পানিতে মিশিয়েছে এবং ছিটিয়ে দিয়েছে)
4) Mixed with fertilizer and sprayed (সারের সাথে মিশিয়েছে এবং ছিটিয়ে দিয়েছে)
**77) Other: Specify (অন্যান্য: নির্দিষ্ট করে লিখুন) ________**

7.8 What did you do with the leftover fungicide? [জমিতে ব্যবহারের পর অবশিষ্ট ছত্রাকনাশক ঔষধগুলো আপনি কি করেছিলেন?]

1) Did not have any (অবশিষ্ট ছিল না)
2) Stored it in the house (বাড়িতে সংরক্ষন করেছিলাম)
3) Stored it outside the house (বাড়ির বাইরে সংরক্ষন করেছিলাম)
4) Gave it away or sold it (কাউকে দান করে দিয়েছিলাম বা বিক্রি করে দিয়েছিলাম)
**77) Other: Specify (অন্যান্য: নির্দিষ্ট করে লিখুন) ________**

7.9 What did you do with the leftover fungicide packaging? (answer each separately Yes = 1, No = 0) [জমিতে ব্যবহারের পর ছত্রাকনাশক ঔষধের অবশিষ্ট প্যাকেটগুলো আপনি কি করেছিলেন? (প্রত্যেকটি অপশনের ক্ষেত্রে আলাদাভাবে উত্তর দিন । হ্যাঁ=1, না=0)]

1) Stored it inside the house (বাড়িতে সংরক্ষন করেছিলাম)
2) Stored it outside the house (বাড়ির বাইরে সংরক্ষন করেছিলাম)
3) Used it to store food (খাদ্য মজুদ রাখার কাজে ব্যবহার করেছিলাম)
4) Used it to store water (পানি সংরক্ষনের কাজে ব্যবহার করেছিলাম)

5) Used it to store household goods (গৃহস্থালী সামগ্রী সংৰক্ষনের কাজে ব্যবহার করেছিলাম)
6) Let the children play with it (শিশুদের খেলতে দিয়েছিলাম)
7) Buried under the ground (মাটির নিচে পুতে ফেলেছিলাম)
8) Hang in the field with stick (লাঠির সাহায্যে জমিতে ঝুলিয়ে রেখেছিলাম)
9) Discarded it (ফেলে দিয়েছিলাম)
10) **Used it to store leftover fungicide (বাড়িতে ছত্রাকনাশক ঔষধ সহ সংৰক্ষন করেছিলাম)**
11) **Washed the packet/bottle before use (ধুয়ে ব্যবহার করেছিলাম)**

#### Section 8: Sample collection (সেকসন-৮: নমুনা সংগ্রহ)

Rice sample collection (harvest in own land): [চালের নমুনা সংগ্রহ, নিজেদের জমিতে উৎপাদিত)]

8.1 Has a rice sample been collected successfully? (উক্ত খানা থেকে সঠিকভাবে চালের নমুনা সংগ্রহ করা হয়েছে কি?)

0=No (না)

1=Yes (হ্যাঁ)

8.2 PROMPT: Please label the bag with the following label: দয়া করে ব্যাগে এই ক্রমানুসারে লেবেল বসানঃ RS.[খানাআইডি নম্বর].[দিন]. [মাস]

RS.[PID].[DAY].[MONTH]

8.3 (record) (if 8.1 is 0) Why has the rice sample not been collected? (□□□□□) (□□□ ৮.১ □□ □□□□□ ০ □□) □□□□□ □□□□□□ □□□ □□□□□ □□□□□□ □□□ □□□□?

1) No stored rice available at time of interview (সাৰাৎকার গ্রহনের সময় খানাতে চাল ছিল না)
2) Stored rice available but not appropriate for sampling (not from own field) (খানাতে চাল সংরৰন করা আছে কিন্তু সেগুলো তাদের নিজেদের জমিতে উৎপাদিত নয়)
3) Respondent refused (চালের নমুনা দিতে অস্বীকৃতি জানিয়েছেন/প্রত্যাখ্যান করেছেন)
77) Other (□□□□□□□□□)

Soil sample collection: (মাটির নমুনা সংগ্রহ)

8.4 Has a core soil sample been collected successfully? [উক্ত খানার চাষের জমি থেকে সঠিকভাবে মাটির (অন্ৰঃস্থল) নমুনা সংগ্রহ করা হয়েছে কি?]

0=No (না)

1=Yes (হ্যাঁ)

8.5 PROMPT: Please label the bag with the following label: দয়া করে ব্যাগে এই ক্রমানুসারে লেবেল বসানঃ CS [খানাআইডি নম্বর].[দিন]. [মাস]

1) CS1. [PID].[DAY].[MONTH]
2) CS2. [PID].[DAY].[MONTH]
3) CS3. [PID].[DAY].[MONTH]

8.6 (record) (if 8.4 is 0) Why has the core soil sample not been collected? (যদি ৮.৪ এর উত্তর ০ হয়) কেন মাটির (অন্ৰঃস্থল থেকে) নমুনা সংগ্রহ করা হয়নি?

1) Full of standing water (জমিতে পানি জমে ছিল)
2) Respondent refused (মাটির নমুনা দিতে অস্বীকৃতি জানিয়েছেন/প্রত্যাখ্যান করেছেন)
77) Other (□□□□□□□□□)

8.7 Has a scrapping soil sample been collected successfully? **[**উক্ত খানার চাষের জমি থেকে সঠিকভাবে মাটির (মাটি চেঁছে) নমুনা সংগ্রহ করা হয়েছে কি?]

0=No (না)

1=Yes (হ্যাঁ)

8.8 PROMPT: Please label the bag with the following label: দয়া করে ব্যাগে এই ক্রমানুসারে লেবেল বসানঃ SS.[খানাআইডি নম্বর].[দিন]. [মাস]

1) SS1. [PID].[DAY].[MONTH]
2) SS2. [PID].[DAY].[MONTH]
3) SS3. [PID].[DAY].[MONTH]

8.9 (record) (if 8.7 is 0) Why has the scrapping soil sample not been collected? [(যদি ৮.৭ এর উত্তর ০ হয়) কেন মাটির নমুনা (মাটি চেঁছে) সংগ্রহ করা হয়নি? ]

1) Full of standing water (জমিতে পানি জমে ছিল)
2) Respondent refused (মাটির নমুনা দিতে অস্বীকৃতি জানিয়েছেন/প্রত্যাখ্যান করেছেন)
77) Other (□□□□□□□□□)

8.10 GIS coordinates of the agricultural land [চাষের জমির জিআইএস কোঅর্ডিনেট লিপিবদ্ধ করম্নন].

Latitude [ ]

Longitude [ ]

8.10.a GIS coordinates of the fertilizer/ pesticide shop [সার/ কীটনাশকের দোকানের জিআইএস কোঅর্ডিনেট লিপিবদ্ধ করুন].

Latitude [ ]

Longitude [ ]

8.11 (2nd visit) (if 8.4 is 0) PROMPT: Please label the bag with the following label: দয়া করে ব্যাগে এই ক্রমানুসারে লেবেল বসানঃ CS [খানাআইডি নম্বর].[দিন]. [মাস]

1) CS1. [PID].[DAY].[MONTH]
2) CS2. [PID].[DAY].[MONTH]
3) CS3. [PID].[DAY].[MONTH]

8.12 (2nd visit) (if 8.7 is 0) PROMPT: Please label the bag with the following label: দয়া করে ব্যাগে এই ক্রমানুসারে লেবেল বসানঃ SS.[খানাআইডি নম্বর].[দিন]. [মাস]

1) SS1. [PID].[DAY].[MONTH]
2) SS2. [PID].[DAY].[MONTH]
3) SS3. [PID].[DAY].[MONTH]

Interview End Time: (সাক্ষাৎকার গ্রহনের শেষ সময়)

***Thank you.***

**Name & signature of FRA:** **Checked by FRO:**

### Wash Benefit Module 22: Household Latrine survey

Note: Ask these Questions to any adult family member

(এই প্রশ্নগুলো ল্যাট্রিন সম্পর্কে তথ্য দিতে পারবে খানার এমন কোন প্রাপ্ত বয়স্ক সদস্যকে জিজ্ঞাস করতে হবে)

---

**Section-1: Household unique identifier**

1.1 বাড়ী নং (Bari ID): .. .. .. .. .. .. .. ... .. .. .. .. .. .. .. .. .. .. .. .. ... .. .. .. .. .. .. .. .. .. .. .. .. .. .. . .. [ | | ]

1.2 খানা নং(Household ID): ............................................................................................. [ | | | | ]

**(Please follow the specific code sheet)**

1.3 ইউনিয়নের নাম (Union name): ............................................................................................. [ | | ]

1.4 **ঠিকানা** (Address): খানা প্রধানের নাম:.............................................................................................

(Name of household head)

খানা প্রধানের বাবার/স্বামীর নাম: .............................................................................................

(Father's/ husband's name)

সাক্ষাতকার প্রদানকারীর নাম:.............................................................................................

(Name of respondent)

সাক্ষাতকার প্রদানকারীর সাথে খানা প্রধানের সম্পর্ক:

(Relation with HH head)

1.বাবা (father) 2.মা (mother) 3.বড়ভাই (elder brother) 4. দাদা ((grand-father))

5. দাদী (grand-mother) 6. চাচা (uncle) 7. চাচী (unty) 8. মামা (maternal uncle)

77. অন্যান্য:

বাড়ীর নাম:............................................................................................. [ | | ]

(Bari Name)

গ্রাম:.............................................................................................

(Village)

বাড়ীর অবস্থান (বিস্ৰারিত লিখুন):.............................................................................................

Location (specify)

[ | | | ]

Cluster ID |__|__|__| Mother ID |__|__|

1.5 সাক্ষাতকার গ্রহণকারীর নাম এবং কোড (Interviewer name & code): ..................................................

1.6 তথ্য সংগ্রহের তারিখ (Date of data collection/observation/spot check): |__|__|__|__|__|__|

**Section 2: Respondent and household demographics**

| খানার সদস্যদের নাম [**List of household members (name)]**<br>**(খানার সবচেয়ে ছোট বাচ্চাকে দিয়ে শুরু করতে হবে)** | বয়স<br>[Age] | লিঙ্গ [Sex]<br>1. Male<br>2. Female | শিক্ষা [Education]<br>99-জানিনা 88-**প্রযোজ্য নয়** | পেশা<br>[Occupations] | ল্যাট্রিন ব্যবহার<br>(Latrine user) |
|---|---|---|---|---|---|
| **1.** | | | | | |
| **2.** | | | | | |
| **3.** | | | | | |
| **4.** | | | | | |
| **5.** | | | | | |
| **6.** | | | | | |
| **7.** | | | | | |
| **8.** | | | | | |
| **9.** | | | | | |
| **10.** | | | | | |
| **11.** | | | | | |
| **12.** | | | | | |
| **13.** | | | | | |
| **14.** | | | | | |
| **15.** | | | | | |

Occupation Code:

**38.** কৃষক [Farmer/Cultivator]
**39.** ঘরে কাজ করে [Homemaker]
**40.** কৃষি শ্রমিক [Agri-labor]
**41.** শ্রমিক (কৃষি শ্রমিক ছাড়া) [Non-agri labor]
**42.** বেতনভুক্ত কর্মচারী (সরকারী/প্রাইভেট/এন.জি.ও) [Salaried job (Govt./Private/NGO)]
**43.** রাজমিস্ত্রী [Mason (*Rajmistri*)]
**44.** কাঠ মিস্ত্রী [Carpenter]
**45.** ভ্যান/রিকশা চালক [Van/Rickshaw puller]
**46.** জেলে [Fisherman]
**47.** নৌকা চালক/মাঝি [Boatman]
**48.** কর্মকার [Blacksmith]
**49.** স্বর্ণকার [Goldsmith]
**50.** কুমার/কুম্ভকার [Potter (soil smith)]
**51.** মুচি [Shoe polish /maker]
**52.** দোকানদার [Shopkeeper]
**53.** ফেরিওয়ালা [Vendor (*Feriwala/howker*)]
**54.** ক্ষুদ্র ব্যবসায়ী (মূলধন <=১০০০০) [Petty trader, capital <=10000]
**55.** ব্যবসায়ী(মূলধন >১০০০০) [Business, capital >10000]
**56.** দর্জি [Tailor]
**57.** ড্রাইভার [Driver]
**58.** কুটীর শিল্প [Cottage industry]
**59.** পোলট্রি/ ব্যবসার জন্য পশু লালন-পালনকারী [Poultry /livestock reare
**60.** বিদ্যুতিক মিস্ত্রী [Electrician]
**61.** হোমিওপ্যাথি ডাক্তার [Homeopath]
**62.** আধ্যাতিক চিকিৎসক/ কবিরাজ/ ওঝাঁ [Spiritual healer/kabiraj/ Ojha]
**63.** পেশাদার ডাক্তার/উকিল [Professional practitioner (Doctor/lawyer)
**64.** ইমাম/ ধর্মযাজক [Imam/priest]
**65.** অবসরপ্রাপ্ত চাকুরীজীবী [Retired service holder]
**66.** ছাত্র [Student]
**67.** বেকার [Unemployed]
**68.** অক্ষম [Disabled]
**69.** কাজের লোক [Domestic maid / servant]
**70.** জমিদার (শস্য উৎপাদন অথবা অন্য কোন কাজে কৃষকদের জমি বর্গা দেয়) [Landlord (Provide land for farmers for sharecropping or othe
**71.** বিদেশে থাকে [Staying abroad]
**72.** মৃত/নিখোঁজ [Died/untraced]
99. জানিনা [Don't know]

88. প্রযোজ্য নয় [Not applicable]

77. অন্যান্য (বর্ণনা লিখুন) [Others (specify)

..........................................................................

**Self-reported data**

**Section-3: Access to latrine**

1. আপনার/আপনার পরিবারের সদস্যদের কি ল্যাট্রিন ব্যবহারের সুযোগ আছে [Do you/family members have access to a latrine]?

   1. হ্যাঁ [Yes] 2. না [No]

   (যদি উত্তর হ্যাঁ হয় তাহলে প্রশ্ন-২ এ যান অথবা 'না' হয় তাহলে প্রশ্ন-২ এ যান তারপর সেকশন-৫ এ যান) [if answer is 'no', then ask ques-2 and skip to section-5]

2. আপনার/আপনার পরিবারের সদস্যরা কোথায় পায়খানা করেন [Where do you/family members defecate]?

   1. উঠানে [yard] 2.উঠানের বাহিরে খোলা জায়গাতে [Open space outside the front yard]

   3. ঝোপ-ঝাড়ে/জঙ্গলে [Bush/jungle] 4.ল্যাট্রিন /টয়লেটে [In toilet]

   77. অন্যান্য (বর্ণনা লিখুন) [Other (Please describe)]................................

3. ল্যাট্রিন কি আপনার [Do you own the latrine (that you use)]?

   1. হ্যাঁ [Yes] 2.না [No] (যদি উত্তর হ্যাঁ হয় তাহলে প্রশ্ন-৪ এ যান অথবা না হয় তাহলে প্রশ্ন-৬ এ যান )[if answer is 'No' skip to **quest-6**]

4. ল্যাট্রিনের মালিকানার ধরন (Type of ownership)?

1. একক মালিকানা (self own) 2. যৌথ মালিকানা (jointly own)

(যদি উত্তর একক মালিকানা হয় তাহলে **সেকশন-৪** এ যান) [if self own, go to **section-4**]

5. যদি ল্যাট্রিনটি যৌথমালিকানায় হয় তাহলে অংশীদার কে?

1. আত্মীয় (একই বাড়ীর মধ্যে) [relative] 2. প্রতিবেশী [neighbor] 77.**অন্যান্য** [Other]................................

6. আপনি/আপনার পরিবারের সদস্যরা কি অন্য খানা/বাড়ীর ল্যাট্রিন ব্যবহার করেন [Do you/family members use another household's latrine]?

1. হ্যাঁ [Yes] 2. না [No]

(যদি উত্তর হ্যাঁ হয় তাহলে ৭ নং প্রশ্ন জিজ্ঞাসা করুন এবং সেকশন-৫ এ যান, যতি উত্তর না হয় তাহলে সেকশন-৪ এ যান)

(If answer is 'yes', ask question-7 and skip to section-5, if answer is 'no', skip to section-4)

7. কোন খানার ল্যাট্রিন আপনি/আপনার পরিবারের সদস্যরা শেয়ার করেন (বাচ্চারা সহ) (আত্মীয় না প্রতিবেশীর ল্যাট্রিন তা জানতে হবে)
[Which household do you share this latrine (including children)]?

1. আত্মীয় (একই বাড়ীর মধ্যে) [relative] 2. প্রতিবেশী [neighbor] 77.**অন্যান্য** [Other]................................

**Section-4: Duration of latrine use and pit emptying**

8. আপনি/আপনার পরিবারের সদস্যরা কত মাস ধরে এই ল্যাট্রিন ব্যবহার করছেন [How long have you been using this latrine]?
(দিন/বছর যেতে তা মাসে লিখতে হবে)

|__|__| MM

9. এখন যে ল্যাট্রিন ব্যবহার করছেন তার কোন পিট/গর্ত কি কখনও খালি করা হয়েছে ? [Did you ever empty a pit of existing latrine]

1. হ্যাঁ [Yes] 2. না [No]

( যদি উত্তর হ্যাঁ হয় তাহলে প্রশ্ন- ১০ এ যান অথবা না হয় তাহলে প্রশ্ন-১৫ তে চলে যান) [if 'no', skip to question-15]

10. কত বার এই ল্যাট্রিনের পিট খালি করা হয়েছে? [How many times did you empty pit of existing latrine] |__|__|

11. সর্ব শেষ কবে এই ল্যাট্রিনের পিট খালি করা হয়েছিল? [When did you last empty the pit of existing latrine] MM |__|__|

**(যদি যৌথ মালিকানায় হয় তাহলে প্রশ্নঃ ১২ করতে হবে)** [if answer is Jointly own then ask to **quest-12**]

12. সর্ব শেষ কার খরচে এই ল্যাট্রিনের পিট খালি করা হয়েছিল? [Who bear the cost for last empty the pit]

1. নিজের 2.প্রতিবেশী 3.আত্মীয়

13. কি ভাবে পিট খালি করা হয়েছিল [How did you empty]?

1. নিজে/পরিবারের সদস্য [By yourself] 2. সুইপার ভাড়া করে [By hiring sweeper] 77. অন্যান্য [Other].. .. .. .. .. .. ..

14. যদি সুইপার ভাড়া করা হয় তাহলে সর্ব শেষ পিট খালি করতে কত টাকা খরচ হয়েছিল [If, emptied hiring a sweeper, how much does it cost for empty the pit last time]? টাকা [|__|__|__|__|

15. গত এক বছরে আপনার/আপনাদের ল্যাট্রিন কি মেরামত করা হয়েছে [Did you ever repair latrine structure]?

1. হ্যাঁ [Yes] 2. না [No]

(যদি উত্তর হ্যাঁ হয় তাহলে প্রশ্ন-১৬ অথবা না হয়, তাহলে সেকশন-৫ এ চলে যান) [if answer is 'no', skip to secktio-5]

16. গত এক বছরে আপনার/আপনাদের ল্যাট্রিন কতবার মেরামত করা হয়েছে [How many times did you repair the structure (within last one year)? |__|__|

17. কোন অংশ সব চেয়ে বেশী মেরামত করেছেন এবং কত বার [Which part did you repair most]?

1. দেয়াল/ বেড়া [Walls/fences] |__|__|

2. দরজা [Door] |__|__|

3. ছাদ [Roof] |__|__|

77. অন্যান্য [Other].. … … … … … … … … … … … … … … … … … … … … … … … …

18. সর্ব শেষ কোন অংশ মেরামত করেছেন [Which part did you repair last]?

1. দেয়াল/ বেড়া [Walls/fences]
2. দরজা [Door]
3. ছাদ [Roof]
77. অন্যান্য [Other-describe] . . . . . . . . . . . . . . . . . . . . . . . . . . . . . . . . . . . . . . . .

19. কে মেরামত করেছিল [Who repaired it]?

1. নিজে/পরিবারের সদস্য [By yourself] 2. শ্রমিক ভাড়া করে [hire daily labor]

77. অন্যান [other]............................

20. সর্ব শেষ মেরামতে কত খরচ হয়েছিল [How much did it cost to repair (last time repaired)]?

|__|__|__|__| টাকা [Taka]

**Section-5: consent on latrine upgrade**

21. যদি আমরা গবেষণার প্রয়োজনে নতুন ল্যাট্রিন স্থাপন করে দিতে চাই, আপনি কি তাতে রাজী হবেন [If we want to set up a new latrine for research purpose, will you agree]?

1. হ্যাঁ [Yes]  2. না [No]

[যদি উত্তর হ্যাঁ হয় তাহলে প্রশ্ন-২৩ এ যান অথবা না হলে প্রশ্ন-২২ এ যান, যদি ল্যাট্রিন না থাকে তহলে প্রশ্ন ২১ ও ২২ দুটোই প্রযোজ্য হবে] [if answer is 'No' skip to **quest-22**]

22. যদি উত্তর (২১ নং প্রশ্নের) না হয় তাহলে কারন বর্ণনা করুন (বিস্তারিত)ঃ … … … … … … … … … … … … … … … … … … … … … … … …

… … … … … … … … … … … … … … … … …...... … … … … … … … … … … … … … … … … … …...... … … … … … … … … … … … … … … … …

… … … … … … … … … … … … … … … … …...... … … … … … … … … … … … … … … … … … …...... … … … … … … … … … … … … … … … …

23. যদি আমরা আপনার ল্যাট্রিন গবেষণার প্রয়োজনে সংস্কার করতে চাই, আপনি কি তাতে রাজী হবেন [If we want to upgrade your existing latrine for research purpose, will you agree]?

1. হ্যাঁ [Yes]  2. না [No] যদি উত্তর হ্যাঁ হয় তাহলে প্রশ্ন-২৫ এ যান,উত্তর না হলে প্রশ্ন-২৪ এ যান] [if answer is 'No' skip to **quest-24**]

24. যদি উত্তর (২৩ নং প্রশ্নের) না হয় তাহলে কারন বর্ণনা করুন (বিস্তারিত)ঃ … … … … … … … … … … … … … …...... … … … … … … … … … … … … … … … …

…... … … … … … … … … … … … … … … … … …...... … … … … … … … … … … … … … … … …

25. ল্যাট্রিন স্থাপনের জন্য আপনার কতটুকু জায়গা আছে (আত্মীয়-স্বজন ও প্রতিবেশী সহ) (নিজের মালিকানায় নাও হতে পারে কিন্তু আত্মীয়/প্রতিবেশীর জায়গায় পায়খানা স্থাপনের অনুমতি আছে) [How much land do you have access to install a latrine]? [Including neighbor, relative and kin]

(ডেসিমেল)[decimal] |__|__|

26. ল্যাট্রিন স্থাপনের জন্য আপনার নিজের মালিকানাধীন কতটুকু জায়গা আছে [How much land do you own to install a latrine]? |__|__|

(ডেসিমেল) [decimal]

(যদি ল্যাট্রিন স্থাপনের জন্য নিজের জায়গা থাকে তাহলে প্রশ্ন-৩১ এ যেতে হবে, জায়গা না থাকলে প্রশ্ন-২৭ এ যেতে হবে) [if HH own land for latrine installation then skip to quest-31]

27. আপনার আত্মীয়-স্বজন এবং প্রতিবেশীরা কি (তাদের জায়গায়) ল্যাট্রিন স্থাপনের অনুমতি দেবেন

[Would your neighbors and relatives (kin group) permit you to install a latrine in their land]

1. হ্যাঁ [Yes]  2. না [No]

[যদি উত্তর হ্যাঁ হয় তাহলে প্রশ্ন-৩১ এ যান. উত্তর না হলে প্রশ্ন ২৮ এ যান] [if answer is 'No' skip to **quest-28**]

28. যদি উত্তর (২৭ নং প্রশ্নের) না হয় তাহলে কারন বর্ণনা করুন (বিস্তারিত) [Describe if answer is no to quest-27]:

… … … … … … … … … … … … … … … … …...... … … … … … … … … … … … … … … … … … …...... … … … … … … … … … … … … … … … …

… … … … … … … … … … … … … … … … …… … … … … … … … … … … … … … … … …… … … … … … … … … … … … … … … … …

29. আপনার আত্মীয়-স্বজন এবং প্রতিবেশীরা কি (তাদের জায়গায়) স্থাপিত ল্যাট্রিন কম পক্ষে দুই (২) বছর ব্যবহারের অনুমতি দেবেন
[Would your neighbors and relatives (kin group) permit you to use that latrine at least 2 year]?

1. হ্যাঁ [Yes] 2. না [No]

[যদি উত্তর হ্যাঁ হয় তাহলে প্রশ্ন-৩১ এ যান, না হলে প্রশ্ন ৩০ যান] [if answer is 'No' skip to **quest-30**]

30. যদি উত্তর (২৯ নং প্রশ্নের) না হয় তাহলে কারন বর্ণনা করুন (বিস্তারিত [Describe if answer is no to quest-29]… … … … … … … …

… … … … … … … … … … … … … … … … …… … … … … … … … … … … … … … … … …… … … … … … … … … … … … … … … … …

… … … … … … … … … … … … … … … … …… … … … … … … … … … … … … … … … …… … … … … … … … … … … … … … … … …

31. যে জায়গায় নতুন ল্যাট্রিন /পিট স্থানপন করা হবে তার নিকট বর্তী পানির উৎসগুলো বিস্তারিত বর্ননা করুন (পনির উৎসের নাম ও দূরত্ব):

[nearest water sources and distance from proposed duel pit latrine place]

| পানির উৎসের নাম<br>(Source of water) | দূরত্ব (প্রস্তাবিত নতুন ল্যাট্রিন/পিট স্থাপনের জায়গা)<br>(Distance) | গভীরতা (টিউব-ওয়েল হলে)<br>(Depth-if tub-well) |
|---|---|---|

**Section-6: Observation of household latrines (by FRA or data collector)**

32. খানায় কতগুলো ল্যাট্রিন ব্যবহৃত হচ্ছে (নিয়মিত/অনিয়মিত) [How many latrines is in-use? (regular/ irregular)] |__|__|

33. খানায় যে ল্যাট্রিনগুলো ব্যবহৃত হচ্ছে (নিয়মিত/অনিয়মিত) তার ধরন [Type of latrines is in-use in HH? (regular/ irregular)]

ল্যাট্রিন-১ [Latrine: 1] ধরন[Type:] . . . . . . . . . . . . . . . . . . . . . . . . . . . . . . . . . . . . . . . . . .

ল্যাট্রিন-২ [Latrine: 2] ধরন[Type:] . . . . . . . . . . . . . . . . . . . . . . . . . . . . . . . . . . . . . . . . . .

স্বাস্থ্য সম্মত-টয়েলেট (Improved sanitation facilities)-

ফ্লাশ-টয়লেট অথবা পানি ঢেলে ফ্লাস করা টয়লেট [Flush or pour flush toilet flushed to]:

টয়লেটটিতে পয়ঃনিষ্কাশন পাইপের সাথে সংযোগ করে দেয়া [Piped sewer system] ................................01

টয়লেটটিতে সেপটিক ট্যাংক বসানো আছে [Septic tank]......................................................................02

সেপটিক ট্যাংক নাই কিন্তু ফ্লাস করে বা পানি ঢেলে পায়খানা দুরে পিটের মধ্যে সরিয়ে দেয়া যায় ...........................03

[Flush to pit latrine (Off set)]

পিট-টয়লেট (স্লাব এবং ওয়াটার সিল আছে) [Pit latrine with slab & water seal]...............................................04

পিট-টয়লেট (স্লাব আছে কিন্তু ওয়াটার সিল নেই তবে ঢাকনা দেয়ার ব্যবস্থা আছে) .............................................................05

[Pit latrine with slab & no water seal but with a lid]

পিট-টয়লেট (স্স্লাব এবং ফ্ল্যপ আছে কিন্তু ওয়াটার সিল নেই)
[Pit latrine with slab and flap, no water seal]...................................................45

**বায়ু চলাচল উপযোগী উন্নত ল্যাট্রিন** [Ventilated Improved Pit (VIP) latrine]………………………46

কমপোষ্টিং টয়লেট **(পায়খানা এবং প্রস্রাব করার জন্য আলাদা আলাদা ঘর এবং সাথে আলাদা কোথাও পানির ব্যাবস্থা আছে)**

[Composting toilet, *(Composting toilet ensure separation of urine, water and excreta)*]..................06

**(পায়খানা এবং প্রস্রাব করার জন্য আলাদা আলাদা ঘর এবং সাথে আলাদা কোথাও পানির ব্যাবস্থা আছে)**

ফ্লাশ-টয়েলেট অথবা পানি ঢেলে ফ্লাস করা টয়লেট যা কোন খাল, ড্রেন, নদী ইত্যাদির সাথে সংযোজন

করার ফলে অস্বাস্থ্যকর অবস্থার সৃষ্টি করে থাকে ........................................................................................07

[Flush or pour flush toilet connected to somewhere else (canal, ditch, river, etc.)]

পিট/গর্ত পায়খানা, স্লাব নেই এবং যেখান থেকে মশা/মাছি যাওয়া আসা করতে পারে এবং দূর্গন্ধ ছড়ায় ......................................08

[Pit latrine without slab/open pit]

পিট/গর্ত পায়খানা স্লাব আছে তবে,ওয়াটার সিল নেই অথবা ওয়াটার সিল ভাঙ্গা এবং কোন ঢাকনাও নেই...................................09

[Pit latrine with slab & no water seal/broken water seal and no lid]

ঝুলন্ত পায়খানা [Hanging toilet/latrine]....................................................................................................10

<u>খোলা পায়খানা/ টয়লেট (**Open defecation**):</u>

কোন পায়খানা নেই/জঙ্গলে/ঝোপে ঝাড়ে/ খোলা জায়গায় [No facility/bush/field] ....................................................11

**অন্যান্য (নির্দিষ্ট করে লিখুন)** [Others: Specify] .......................................................................................77

৩৪. ল্যাট্রিনের উপরি কাঠামোর বিস্তারিত বর্ননা [ structure of latrine]

ল্যাট্রিন-১ [Latrine: 1]

1.দেয়াল/ বেড়া [Walls/fences] . . . . . . . . . . . . . . . . . . . . . . . . . . . . . . . . . . . . . . . . . . . . . . . . . . . . . .

2.দরজা [Door] . . . . . . . . . . . . . . . . . . . . . . . . . . . . . . . . . . . . . . . . . . . . . . . . . . . . . . . . . . . .

3.ছাদ [Roof] . . . . . . . . . . . . . . . . . . . . . . . . . . . . . . . . . . . . . . . . . . . . . . . . . . . . . . . . . . . . .

ল্যাট্রিন-২ [Latrine: 2]

1.দেয়াল/ বেড়া [Walls/fences] . . . . . . . . . . . . . . . . . . . . . . . . . . . . . . . . . . . . . . . . . . . . . . . . . . . . .

2.দরজা [Door] . . . . . . . . . . . . . . . . . . . . . . . . . . . . . . . . . . . . . . . . . . . . . . . . . . . . . . . . . . . .

3.ছাদ [Roof] . . . . . . . . . . . . . . . . . . . . . . . . . . . . . . . . . . . . . . . . . . . . . . . . . . . . . . . . . . . .

35. নিয়মিত খাবার পানির উৎস (রেসপনডেন্ট চিহ্নিত করবেন) [Source of regular drinking water (as defined by the respondents)]?

1. **অগভীর টিউবওয়েল (২৫০ ফিটের কম)**[Shallow tube well] (<250 feet)

2. **গভীর টিউবওয়েল (২৫০ ফিটের বেশী**) [Deep tube well]. (250+ feet)

3. **নদী/বাধ/লেক/পুকুর/সেচ নালাথেকে জীবানুমুক্তকরন পানি**

[Pathogen treatment plant (Pond Sand Filter)]: River/dam/lake/ponds/stream/canal/irrigation channel.]

4. **নদী/বাধ/লেক/পুকুর/সেচ নালাথেকে সরাসরি সংগৃহীত পানি**

[Directly from River/dam/lake/ponds/stream/canal/irrigation channel]

77. অন্যান্য [Other] (বর্ণনা করুন): . . . . . . . . . . . . . . . . . . . . . . .

36. ল্যাট্রিন (সবচেয়ে বেশী ব্যবহৃত হয় যে ল্যাট্রিনটা) ও খাবার পানির উৎসের মধ্যে দূরত্ব (আনুমানিক) [Distance (approximately) between latrine (mostly in use) and drinking water source (tube-well)]?

**(নিজের/আত্মীয়/প্রতিবেশীর মালিকানার খাবার পানির উৎসের দূরুত্ব)**

<৩০ ক[illegible]ps] (৩০ কদমের বেশী হলে মাপার দরকার নাই) > ৩০ এর বেশী (>more than 30 [illegible]s]

37. বর্তমান ল্যাট্রিনে কয়টি পিট আছে (বর্তমানে ব্যবহৃত হচ্ছে বা খানার সদস্যরা যে ল্যাট্রিন বেশী ব্যবহার করে) [Number of pit in existing latrine? (currently in use, mostly used by the household)]

38. পিটে কতটি রিং ব্যবহৃত হয়েছে (প্রশ্ন করতে হবে) [Number of rings used in the pit? (need to ask)]

39. রিং এর বর্তমান অবস্থা কি (যে রিং গুলো দেখা যায়) [What is the current condition of the rings]?

পিট-১ [Pit-1]: 1. ভাল [Functional] 2. ভাঙ্গা [Broken]

পিট-২ [Pit-2]: 1. ভাল [Functional] 2. ভাঙ্গা [Broken]

40. যদি রিং ভাঙ্গা হয়, তাহলে তার বিস্তারিত বর্ণনা করুন [If ring is broken, describe the situation]?

পিট-১ [Pit-1]: . . . . . . . . . . . . . . . . . . . . . . . . . . . . . . . . . . . . . . . . . . . . . . . . . . . . . . . . . . . .

পিট-২ [Pit-2]: . . . . . . . . . . . . . . . . . . . . . . . . . . . . . . . . . . . . . . . . . . . . . . . . . . . . . . . . . . . .

41. পিট থেকে কি পায়খানা বাইরে বের হচ্ছে [Does feces spill out from the pit]?

1. হ্যাঁ [Yes] 2. না [No]

(যদি উত্তর হ্যাঁ হয় তাহলে প্রশ্ন-৪২ এ যান অথবা 'না' হলে ৪৫ নং প্রশ্নে চলে যান) [if answer is 'yes' then describe (question-42), otherwise skip to question-45]

42. পায়খানা বের হয়ে কোথায় পরছে [Where the spilled feces get to]? … … … … … … … … … … … … … … … … … … … … … … … … …
… … … … … … … … … … … … … … … … … … …… … … … … … … … … … … … … … … … … … … … … …… … … … … … … … … … … … … … … … … … …

… … … … … … … … … … … … … … … … … … …… … … … … … … … … … … … … … … … … … … … … …… … … … … … … … … … … … … … … … … … …

43. পিটের চার পাশে পায়খানা দেখা যাচ্ছে কি [Is there feces visible around the pit]?

1. হ্যাঁ [Yes] 2. না [No]

44. পায়খানা কি ভাবে পিট এর বাইরে বের হচ্ছে তা বর্ণনা করুন [Describe how does feces come out from the pit]? . … … … … … …
… … … … … … … … … … … … … … … … … … …… … … … … … … … … … … … … … … … … … … … … …… … … … … … … … … … … … … … … … … … …

… … … … … … … … … … … … … … … … … … …… … … … … … … … … … … … … … … … … … … … … …… … … … … … … … … … … … … … … … … … …

45. ল্যাট্রিনের স্লাবের [Floor] বর্তমান অবস্থা কি [What is the current condition of latrine slab]?

1. ভাল [Functional] 2. ভাঙ্গা [Broken]

46. যদি স্লাব [Floor] ভাঙ্গা হয় বা খারাপ হয় তাহলে তার বর্ণনা করুন [If slab is damaged or broken describe it]? … … … … … … …

… … … … … … … … … … … … … … … … … … …… … … … … … … … … … … … … … … … … … … … … …… … … … … … … … … … … … … … … … … … …

… … … … … … … … … … … … … … … … … … …… … … … … … … … … … … … … … … … … … … … … …… … … … … … … … … … … … … … … … … … …

47. স্লাবে [Floor] কি পায়খানা দেখা যাচ্ছে [Are there any visible feces on the slab]?

1. হ্যাঁ [Yes] 2. না [No]

(যদি উত্তর হ্যাঁ হয় তাহলে প্রশ্ন-৪৮ এ যান অথবা 'না' হয় তাহলে ৪৯ নং প্রশ্নে চলে যান) [if answer is 'no', skip to the question-48]

48. যদি হ্যাঁ হয়, তার বর্ণনা করুন (কোন অংশে দেখা যাচ্ছে, পরিমান) [If yes, describe it? (which part it appears, quantity… … … …

… … … … … … … … … … … … … … … … … … …… … … … … … … … … … … … … … … … … … … … … …… … … … … … … … … … … … … … … … … … …

… … … … … … … … … … … … … … … … … … …… … … … … … … … … … … … … … … … … … … … … …… … … … … … … … … … … … … … … … … … …

49. প্যানে পায়খানা দেখা যাচ্ছে কি [Is there any feces appear on the pan]?

1. হ্যাঁ [Yes] 2. না [No] 3. নাই [No pan]

( যদি উত্তর হ্যাঁ হয় তাহলে প্রশ্ন- ৫০ এ যান অথবা 'না'হলে ৫১ নং প্রশ্নে চলে যান) [if answer is 'no', skip to the question-50]

50. যদি হ্যাঁ হয়, তার বর্ণনা করুন [If yes, describe it]? . . . . . . . . . . . . . . . . . . . . . . . . . . . . . . . . . . . . . . . . . . . . . . . . . . . . . . . .
. . . . . . . . . . . . . . . . . . . . . . . . . . . . . . . . . . . . . . . . . . . . . . . . . . . . . . . . .
. . . . . . . . . . . . . . . . . . . . . . . . . . . . . . . . . . . . . . . . . . . . . . . . . . . . . . . . . . . . . . . . . . . . . . . . .

51. ল্যাট্রিনে কি ওয়াটার সিল/সাইফুন দেখা যাচ্ছে [Is there is any water seal/siphon appear in the latrine]?

1. হ্যাঁ [Yes] 2. না [No] 3. নাই [No water seal]

(যদি দেখা সম্ভব হয় দেখতে হবে, প্যান পায়খানায় পরিপূর্ন থাকলে দেখা নাও যেতে পারে। পরিপূর্ণ থাকলে ল্যাট্রিনে পানি ঢেলে দেখতে হবে। যদি উত্তর হ্যাঁ হয় তাহলে পরবর্তী প্রশ্নে চলে যান) [if it is possible to watch, if latrine is overflow it may not be visible. If the answer is 'yes' proceed]

52. ওয়াটার সিল/সাইফুন এর বর্তমান অবস্থা কি [What is the current condition of the water seal]?

1. ভাল [Functional] 2.ভাঙ্গা [Broken]

53. যদি ভাঙ্গা হয়, তা বর্ণনা করুন (এর মধ্যে পানি নেই, এর মধ্য দিয়ে পিটের পায়খানা সরাসরি দেখা যাচ্ছে, গর্ত দেখা যাচ্ছে) [If broken, describe it (no water in it, pit content appears through the water seal, big hollow appears etc.)]? . . . . . . . . . . . . . . . . . . . . . . . . . . . . . . . . .
. . . . . . . . . . . . . . . . . . . . . . . . . . . . . . . . . . . . . . . . . . . . . . . . . . . . . . . . . . . . . . . . . . . . . . . . . . . . . . . . . . . . . . . . . . . . . . . . . . .
. . . . . . . . . . . . . . . . . . . . . . . . . . . . . . . . . . . . . . . .

**Interviewer instruction sheet**

১. যদি কোন খানায় ল্যাট্রিন না থাকে তাকে নতুন ল্যাট্রিনের জন্য বিবেচিত হবে (**নতুন ল্যাট্রিন হচ্ছে উপরিকাঠাম সহ সমস্ৰ উপাদান নতুন** )।
২. যদি ল্যাট্রিনের উপরি কাঠামোর শুধু ছাদ/দরজা/বেড়া না থাকে তাহলে তা মেড়ামত করে দেয়া জন্য বিবেচিত হবে।
৩. যদি ল্যাট্রিনের উপরি কাঠামো নস্ট হয়ে যায় (ব্যবহারের অনুপযগি, একই সাথে উপর দিয়ে পানি পরে, দরজা ভাঙ্গা, বেড়া ভাঙ্গা) তাহলে নতুন উপরি কাঠামো দেয়া হবে জন্য বিবেচিত হবে।
৪. কোন ল্যাট্রিনের একটা পিট/গর্ত থাকলে তা ডুয়েল পিট ল্যাট্রিনে রূপান্ৰর করে দেয়া জন্য বিবেচিত হবে (যদি প্রথম পিট/বর্তমান পিট ভাল থাকে)।
৫. কোন ল্যট্রিনের বর্তমান পিট (রিং ও স্লাব ভাঙ্গা/ রিং ভাঙ্গা এবং পায়খান বাইরে পরছে) তাহলে তাকে নতুন ল্যাট্রিন দেয়া জন্য বিবেচিত হবে।
৬. যদি কোন ল্যাট্রিনে ৩ টার কম (১ বা ২ টা) রিং থাকে তাহলে নতুন ল্যাট্রিনের জন্য বিবেচিত হবে।
৭. সব ল্যাট্রিনের সাইফুন দেয়ার জন্য বিবেচনা করা হবে।

WASH Benefits Module 53
In-Depth Interview Guidelines

**In-depth environmental assessment (assessment of exposure pathways to fecal contamination, association with diarrhea and sanitation coverage in Bangladesh)**

**Drinking water sources, handling and storage practices**

- Can you tell me about how you collect and store drinking water?

- What is your primary/main source of drinking water? (probe- type,tubewell, pond and others,their conditions, secondary and other sources, seasonal impacts. Emphasize tubewell and pond water). Observe the source and describe objectively.
- Who from your household is responsible to collect drinking water? (probe- primary and secondary collectors, when and why?)
- How is the collected water stored and for how long?Note the common storage containers. Give me an example of typical collection and storage of water and usage by the household members.
- How can these water sources or stored water be contaminated by human and/or animal feces? What are the possible pathways that could contaminate the water by humanfeces including children feces and/or animal feces(draw a map of assumed paths of fecal flow)?
- How do young children in your household get water for drinking?
- What, if any, are the difficulties associated with collecting and storing safe water?

**Availability, treating, scarcityand consumption of water**

- Can you tell me if your household ever experiences drinking water scarcity? What do you do in those times?
- Do members of your household consume water differently? How?
- Water treating or purifying behaviors:Do you do anything to your drinking water to make it safer to drink? What are the common water treatment methods in your household?
- Tell me about typical consumption of water by each of your household members including children by age group?
- Tell me yesterday's practices and consumption (note date, temperature, season etc.)

**Contamination of hands and handwashing practices**

- How frequently and when do you wash your hands? probe- key times (before having different types of food, after using toilet and cleaning child's anus). Illustrate a typical day's practices (may be yesterday or today).
- Do you think either/both of your hands could be contaminated by human feces? Probe- handwashing during these key times(draw a map from which activities fecal contamination may occur to hands)
- What agent do you use to wash your hands? Probe- why and when with what?
- How often do you clean your young children’s hands? When and how?

- How often do your young children wash their own hands? When and how?
- Do you think your children's hands could be contaminated by human feces?
- Do you find handwashing with soap difficult or easy? Why?

**Contamination of soil**

- Does your household do anything to keep your courtyard clean? Probe – what is done, how often?
- How could courtyard soil be contaminated by human feces? Probe- open defecation by children and adults, during day/night or during rain. Locate spots and describe.
- Where do young children in your compound defecate? When and why? Who disposes of those feces, how and where? Is there any variation in these practices? What are the differences and why? Observe feces disposal sites. Probe- each sites.
- Is there any child potty available in households in this household?
- Tell me about how often your children come into contact with soil and feces?
    - Child and animal feces
    - Type of animal, children from neighboring households
- How frequently do your children (by age group) spend time in courtyard or verandah and for how long? Who else accompanies them? Do children play alone or in groups? Observe- floor materials (concrete or mud), is there any child potty in the household available?
- How much time do your children spend on thecourtyard ground and why? Who watches over children's behavior when they play in courtyard? Probe- absence and present of primary and secondary caregivers.
- What do the caregivers do when a child puts something from courtyard ground/soil into their mouth?
- What do the caregivers think when a child puts something from soil into their mouth?

**Contamination of food**

- Please tell me about food preparation and preservation processes followed by your household? What is your role in preparing and storing daily food? Probe- main food and snacks, occasional/seasonal foods.
- How do you store those foods and for how long? Probe- when and why?

- Who is responsible for maintaining food storage and hygiene in your household? Probe- social and economic power structure in the household (mother/father-in-laws, husbands).
- How do prepared and stored food gets into contact with hands? What do you do to keep your food safe? Probe- hand hygiene before preparing food, heating and reheating before serving food to child, child consumption of food and feeding methods. Who feeds, when and how?

**Free roaming domestic animals**

- How many domestic animals do you have? Make a count. Identify the majority.
- Who takes care of animals and animal sheds?
- When and why are animal feces visible in compound/inside household and for how long?
- How can a child come into contact with animals?
- Is there any chance that a child can put animal feces from the courtyard ground into their mouth? Probe- which animals, when and where?
- What do parents do if a child puts animal feces in their mouth? Please describe.

**Contamination via flies**

- What do you think about the presence of different flies in your house and surrounding environment? Ask specifically about type (size, color, other characteristics).
- Why are these flies found around/inside the household? Where do they come from? When is the fly density high and why?
- Is there a chance that these flies can land on food? How frequently? Probe- time and season.
- Do you feel that flies pose any health risks to your household? Why or why not?

**Availability and type of toilet facility**

- Please tell me about your household’s sanitation practices. Where do the household members usually defecate? Probe- practice open defecation, individual or shared latrine.
- How many latrines do you have? Probe- type and other facilities available for each toilet.Map the compound marking toilets and distances from drinking water source, pond or other water sources and from living room, kitchen etc.
- Who is responsible for cleaning and maintaining the toilet? Probe- man, women, specific roles, frequency.

- What are the hardware available in this household to dispose of feces from courtyard or inside the house? Probe- child and animal feces, adult human feces, feral animal feces.
- What happens during different seasons of the year? Probe- when latrine pit is full, feces float over pan, during rainy season, when latrine is broken.
- Where do the feces from the latrinein your household end up? In ditches, ponds, drains, small rivers, other water bodies.
- What do you find difficult about managing sanitation in your household? Please describe.
- What is easy about managing sanitation in your household? Please describe.

#### WASH Benefits Module 53: In-Depth Interview Guidelines in Bengali for Environmental Assessment

**পরিবেশগত নিবিড় মূল্যায়ন (বাংলাদেশে ডায়রিয়া এবং স্যানিটেশন সুবিধার অন্তর্ভূক্তি, মলদুষণের গমনাগমনের মূল্যায়ন)**

**পানীয় জলের উৎসসমূহ, পরিচালনা এবং সংরৰণ চর্চাসমূহ**

- আপনি কি বলতে পারেন কিভাবে পানি সংগ্রহ ও সংরৰণ করেন?
- আপনার প্রাথমিক/প্রধান পান করার পানির উৎস কী কী? (জিজ্ঞাসা করম্নন- ধরণ, টিউবওয়েল, পুকুর বা অন্যান্য, সেগুলির ভৌত অবস্থা, দ্বিতীয় ও অন্যান্য উৎসসমূহ, মৌসুমী প্রভাবসমূহ। টিউবওয়েল ও পুকুরের পানির উপর গুরম্নত্ব বেশী দিন)। এই উৎসগুলি পর্যবেৰণ করম্নন এবং নৈর্ব্যক্তিকভাবে বর্ণনা করম্নন।আপনার গৃহস্থালীতে পান করার পানি সংগ্রহ করা কার দায়িত্ব? (জিজ্ঞাসা করম্নন- প্রাথমিক ও দ্বিতীয় সংগ্রহকারী, কখন ও কেন?)
- সংগৃহীত পানি কিভাবে সংরৰণ করা হয় এবং কত সময়ের জন্য? সাধারণভাবে ব্যবহার করা হয় এমন সংরৰণ পাত্রগুলি লিপিবদ্ধ করম্নন। আপনার পরিবারের সদস্যগণ কর্তৃক সাধারণ প্রাত্যহিক একটি দিনে পানি সংগ্রহ, সংরৰণ ও ব্যবহারের উদাহরণ দিন।
- এই পানির উৎসগুলি বা সংরৰণকৃত পানি কিভাবে মানুষ বা পশু-পাখির মল দ্বারা দুষিত হতে পারে? এই সম্ভাব্য দুষণের গমনাগমন পথগুলি কী কী যা প্রাপ্ত বয়স্ক মানুষ, শিশু বা পশু-পাখির মলের দ্বারা দুষিত হয়?
- আপনার পরিবারের শিশুরা কিভাবে খাবার পানি পায়?
- যদি থেকে থাকে তাহলে নিরাপদ পানি সংগ্রহ এবং সংরৰণের সাথে সংশিস্নষ্ট অসুবিধাগুলি কী কী?

**সহজপ্রাপ্যতা, পানি নিরাপদকরণ, পানি স্বল্পতা এবং পানির ব্যবহার**

- আমাকে কি বলতে পারেন আপনার পরিবার কখনো পানির স্বল্পতার মুখোমুখি হয়েছেন কি না? ঐ সময়গুলিতে আপনি কী করেছেন?
- আপনার পরিবারের সদস্যদের পানির ব্যবহারের পরিমাণে ভিন্নতা রয়েছে কি? কিভাবে ও কেন?
- পানি নিরাপদ বা বিশুদ্ধকরণ আচরণঃ আপনি কি আপনার পানি পানের জন্য নিরাপদ করতে কোন কিছু করেন? আপনার গৃহস্থালীতে পানি বিশুদ্ধকরণের সাধারণ পদ্ধতিগুলি কী কী?
- শিশুদের বয়সভেদসহ আপনার পরিবারের সকল সদস্যদের সাধারণভাবে পানি ব্যবহার সম্পর্কে বলুন।
- গতকালকের পানি ব্যবহার ও এ সংক্রান্ত চর্চা সম্পর্কে বলুন (তারিখ, ঐদিনের তাপমাত্রা, তখনকার সিজন ইত্যাদি লিপিবদ্ধ করম্নন)

**হাতে জীবানুর সংক্রমণ এবং হাত ধোয়ার চর্চা**

- কত ঘনঘন এবং কিভাবে আপনি হাত ধোন? জিজ্ঞাসা করুন- প্রধান প্রধান সময়গুলি (বিভিন্ন ধরণের খাবার খাওয়ার আগে, টয়লেট ব্যবহারের পর এবং বাচ্চাকে শৌচানোর পর)। কোন একটি সাধারণ দিনের হাত ধোয়ার চর্চাগুলি বর্ণনা করুন (হতে পারে গতকাল বা আজ)।
- আপনি কি মনে করেন যে কোন একটি বা উভয় হাতই মানুষের মল দ্বারা দুষিত হতে পারে? জিজ্ঞাসা করুন- ঐ প্রধান প্রধান সময়গুলিতে হাত ধোয়া (একটি চিত্র অংকন করুন যেসব কর্মকান্ড থেকে হাতে মলের সংক্রমন ঘটতে পারে)।
- হাত ধোয়ার জন্য আপনি কী কী ব্যবহার করেন? জিজ্ঞাসা করুন- কেন এবং কখন কখন কী কী দিয়ে?
- কতবার আপনি আপনার শিশুর হাত পরিস্কার করেন/ধোন? কখন কখন এবং কিভাবে?
- কতবার আপনার শিশুরা তাদের নিজেদের হাত ধোয়? কখন কখন এবং কিভাবে?
- আপনি কি মনে করেন আপনার শিশুর হাত মানুষের মল দ্বারা দুষিত হতে পারে?
- সাবান দিয়ে হাত ধোয়া আপনার কাছে সহজ না কঠিন মনে হয়? কেন?

**মাটি দুষণ**

- আপনার উঠানটিকে পরিস্কার রাখার জন্য আপনার পরিবার কী কী করে? জিজ্ঞাসা করুন- কী কী করা হয়, কিভাবে এবং কত সময় পর পর?
- উঠান কিভাবে মানুষের মল দ্বারা দুষিত হতে পারে? জিজ্ঞাসা করুন- শিশু এবং বয়স্কগণ, দিনে বা রাতে বা বৃষ্টির সময় খোলা জায়গায় মলত্যাগ করা। স্থানগুলি চিহ্নিত করুন এবং বর্ণনা করুন।
- আপনার বাড়ীতে ছোট ছোট শিশুরা কোথায় পায়খানা করে? কখন কখন এবং কোথায়? ছোট শিশুদের মল কে কোথায় এবং কিভাবে অপসারণ করে? এই চর্চাগুলিতে কি কোন পার্থক্য হয়? পার্থক্যগুলি কী কী এবং কেন হয়? যেখানে যেখানে মল ফেলা হয় সেই স্থানগুলি পর্যবেক্ষণ করুন এবং প্রত্যেক স্থানগুলি নিয়ে আরও প্রশ্ন জিজ্ঞাসা করুন।
- এই পরিবারে কি কোন শিশুদের মলত্যাগ করার জন্য পটি রয়েছে?
- আমাকে বলুন কত বার আপনার শিশুটি মাটি এবং মলের সংস্পর্শে আসে?
  - শিশুদের এবং পশু-পাখির মল
  - মলের ধরণ, পার্শ্ববর্তী বাড়ীর শিশুদের মল
- কত সময় পর পর আপনার শিশুরা (বয়সদল ভেদে) উঠান বা বারান্দায় সময় কাটায় এবং কত সময় ধরে? তাদেরকে কে সঙ্গ দেয়? শিশুরা কি একা না অনেকে মিলে খেলাধুলা করে? পর্যবেক্ষণ করুন- ঘরের বা বারান্দার মেঝের ধরণ (পাকা বা মাটি), ঘরে শিশুদের মলত্যাগ করার জন্য কোন ধরণের পটি আছে কি না।
- আপনার শিশু কত সময় উঠানে সময় কাটায় এবং কেন? কে তাদের আচার-আচরণ লক্ষ্য করে যখন তারা উঠানের মাটিতে খেলাধুলা করে? জিজ্ঞাসা করুন- মায়ের বা অন্য যত্নগ্রহণকারীর উপস্থিতি এবং অনুপস্থিতিতে।
- যখন শিশুরা উঠানের মাটি থেকে তাদের মুখে কোন কিছু দেয় তখন মায়েরা কী করেন?
- বাচ্চারা যখন মাটি থেকে কোন কিছু মুখে দেয় তখন মায়েরা কী ভাবেন বা মনে করেন?

**খাদ্য দূষণ**

- দয়া করে আমাকে আপনাদের পরিবারের খাবার তৈরী এবং সংরক্ষণ প্রক্রিয়া সম্পর্কে বলুন। প্রতিদিনকার খাবার তৈরী এবং রক্ষণাবেক্ষণে আপনার ভূমিকা কী? জিজ্ঞাসা করুন- প্রধান প্রধান খাবার, নাস্তা, মৌসুমি খাবার।
- এই খাবারগুলি কিভাবে ও কতক্ষণের জন্য সংরক্ষণ করেন? জিজ্ঞাসা করুন- কখন কখন ও কেন?
- আপনার পরিবারে খাবার সংরক্ষণ ও তা স্বাস্থ্যসম্মত রাখার দায়িত্ব কার? জিজ্ঞাসা করুন- পরিবারের সামাজিক ও অর্থনৈতিক ক্ষমতা (শ্বশুর-শ্বাশুড়ী, স্বামী)
- তৈরীকৃত ও সংরক্ষিত খাবারগুলি কিভাবে হাতের সংস্পর্শে আসে? আপনার পরিবারের জন্য খাবার নিরাপদ রাখার জন্য কী কী করেন? জিজ্ঞাসা করুন- খাবার তৈরীর পূর্বে হাত ধোয়া, বাচ্চাকে খাওয়ানোর আগে বার বার খাবার গরম করা, শিশুদের খাবারের পরিমাণ এবং খাওয়ানোর পদ্ধতি। কে খাওয়ায়, কখন কখন ও কেন?

**উন্মুক্তভাবে থাকা গৃহপালিত পশু-প্রাণীসমূহ**

- আপনার কতগুলি গৃহপালিত পশু রয়েছে? সংখ্যা গণনা করুন, সবচেয়ে বেশী কোনগুলি তা চিহ্নিত করুন।
- পশু-পাখী ও খোয়ার বা গোয়ালঘর কে দেখাশোনা করে
- কখন কখন এবং কেন পশু-পাখীর মল আপনার বাড়ীর উঠানে ও আশে পাশে দেখা যায় এবং কত সময় ধরে?
- একটি বাচ্চা শিশু কিভাবে পশু-পাখীর সংস্পর্শে আসতে পারে?
- এমন কি কোন সম্ভাবনা আছে যে শিশুরা উঠানের মাটি থেকে পশু-পাখির কোন মল মুখে দিতে পারে?
- বাচ্চারা যদি পশু-পাখির মল মুখে দেয় তবে মা-বাবারা কী করেন? বর্ণনা করুন।

**মশা-মাছির মাধ্যমে দূষণ/সংক্রমণ**

- আপনার বাড়ীর ভেতর এবং আশে পাশে বিভিন্ন ধরণের মাছির উপস্থিতি সম্পর্কে আপনার কী মনে হয়? সুনির্দিষ্টভাবে ধরণগুলি সম্পর্কে জিজ্ঞাসা করুন (আকৃতি, রং, অন্যান্য বৈশিষ্ট্যসমূহ)
- এই ধরণের মশা-মাছিগুলি আপনার ঘরের ভেতর বা আশে পাশে কেন পাওয়া যায়? এগুলি কোথা থেকে আসে? কখন কখন মশা-মাছির উপদ্রব বেশী হয় ও কেন?
- এমনি কি কোন সুযোগ আছে যে মশা-মাছি খাবারের উপর বসে? কত ঘন ঘন এমন হয়? জিজ্ঞাসা করুন- সময় ও ঋতু।
- আপনি কি মনে করেন যে মাছিরা আপনার পরিবারের জন্য কোন স্বাস্থ্য ঝুকি বয়ে আনে? কেন অথবা কেন নয়?

**টয়লেট সুবিধাদির ধরণ ও সহজলভ্যতা**

- আপনার পরিবারের স্বাস্থ্যসম্মত চর্চা সম্পর্কে বলুন। তারা কোথায় সচরাচর পায়খানা/মলত্যাগ করেন? জিজ্ঞাসা করুন- খোলা জায়গায়, ব্যক্তিগত/নিজস্ব বা অন্যের ল্যাট্রিনে।
- আপনার কয়টি ল্যাট্রিন আছে? জিজ্ঞাস করুন- প্রতিটি ল্যাট্রিনের ধরন ও অন্যান্য সুবিধাসমূহ। এই বাড়ীটির একটি চিত্র অংকন করুন যেখানে ল্যাট্রিন/টয়লেট সুবিধাদি এবং খাবার পানির উৎস, পুকুর বা অন্যান্য উৎস, থাকার ঘর ও রান্না ঘর থেকে তার দূরত্ব চিহ্নিত করুন।
- টয়লেট ব্যবস্থাপনা ও পরিস্কার রাখার দায়িত্ব কার? জিজ্ঞাসা করুন- পুরুষ, নারী, নির্দিষ্ট ভূমিকা, কত সময় পর পর?
- উঠান থেকে অপসারণ করার জন্য এই পরিবারের কোন কোন যন্ত্র-পাতি রয়েছে। জিজ্ঞসা করুন- শিশু ও পশুর মল, বয়স্ক দের ও অপোষা প্রাণীদের।
- বছরের বিভিন্ন মৌসুমে কী হয়? জিজ্ঞাস করুন- ল্যাট্রিন যখন ভরে যায়, প্যানের উপর মল ভাসতে থাকে, বা ল্যাট্রিন যখন ভেঙ্গে যায়।
- ল্যাট্রিন থেকে মল কোথায় গিয়ে শেষ পৌছে? নালায়, পুকুরে, ডোবায়, ছোট নদী, গাঙ্গে, বা অন্যান্য কোন পানিতে।
- আপনার পরিবারের সদস্যদের মলত্যাগ ও পরিবেশ মলমুক্ত রাখার জন্য কী কী অসুবিধার মুখোমুখি হন বর্ণনা করুন।
- এ ধরণের স্যানিটেশন কাজে কোন বিষয়গুলি সহজ বলে মনে হয় ও কেন? বর্ণনা করুন।

**WASH Benefits Module 54**
Focus Group Discussion Guidelines

**In-depth environmental assessment (assessment of exposure pathways to fecal contamination, association with diarrhea and sanitation coverage in Bangladesh)**

**Drinking water sources, handling and storage practices**

- What are the primary/main sources of drinking water in this area? Probe- type,tubewell, pond and others,their conditions, secondary and other sources, seasonal impacts. Emphasize tube-well and pond water. Observe the source and describe objectively.
- What are the common drinking water collection and storage methods? Rank these methods and probe all methods for clarification and understanding.
- Who from the households is usually responsible to collect and store drinking water? (probe- primary and secondary collectors, when and why?
- How can these water sources or stored water be contaminated by human and/or animal feces? What are the possible pathways that could contaminate the water by human feces including children feces and/or animal feces?(draw a map of assumed paths of fecal flow, check with the maps drawn from in-depth interviewsforupdate/modification)

**Availability, treating, scarcityand consumption of water**

- When is there drinking water scarcity in this area?
- What do the household members do at that time?
- How much difference is there in collecting, storing and consuming water?
- What are the common or popular water treating or purifying methods in this area? Describe to me the most popular or common method.
- Are there certain times of the year when treating or purifying water is more or less common?
- Tell me about typical consumption of water by household members. Probe- men, women, children by age group?

**Contamination of hands and handwashing practices**

- How frequently and when do people wash their hands? Probe- three key times (before having different types of food, after using toilet and cleaning child's anus). Describe a typical person's handwashing practices (could be a caregiver/homemaker).

- Do people think that either/both of their hands could be contaminated by human feces? Probe- handwashing during these key times. (draw a map from which activities fecal contamination may occur to hands)

- Do people wash the left or right hands more often? When and why?

- What agentsare available in this area that people use to wash their hands? Probe- all agents, rank on availability and frequency of use, why and when with what?

- How frequently do young children have their hands washed by their caregivers?

- How often do young children wash their own hands?

- Do people think that young children's hands could be contaminated by human and/or animal feces? Who from the household can encourage or influence handwashing? Probe- Grandfather, father-in-law, husband, school going children?

**Contamination of courtyard, soil of surrounding household environment**

- How could courtyard soil become contaminated by human feces? Probe- open defecation by children and adults, elderly, sick, during day/night or during rain, locatespots and describe. Ask for an example that they can explain. Draw a map with sources of feces and contamination.

- Where do young children defecate? When and why? Who disposes of those feces, how and where?Probe- common, fixed, temporary sites.

- Is there any variation in feces disposal practices? What are the differences and why? Observe feces disposal sites. Probe- each site.

- Are there child potties available in households in this community?

- In which place do young children spend most of their time other than lap of adults? Describe the places

- How frequently dochildren (by age group) spend time in courtyard or verandah and for how long?

- Who else accompanies them? Do children play alone or in groups in household or courtyard?

- Tell me about how often children from this community come into contact with soil and feces.
    - Courtyard or veranda or household floor soil

- Child and animal feces
- Type of animals, children from neighboring households

- How much time do children spend on the courtyard ground and why? by age group
  - <3 years
  - 3-6 years
  - 6+ years
- Who watches over children's behavior when they play in courtyard? Probe- absence and present of primary and secondary caregivers.
- What are the materials that young children can put into their mouth when playing or resting on the ground? List those materials, rank and describe.
- What do the caregivers do when a child puts something from courtyard ground/soil into their mouth?
- What do the caregivers think when a child puts something from soil into their mouth?

**Contamination of Food**

- Tell me about the food preparation methods are used in this area?
- What are the common food storing methods in this area? Rank and identify the main method and explain.
- Who plays the most important role in preparing and storing daily food? Who designates these people and why? Probe- main food and snacks, occasional/seasonal foods.
- Who from the household is responsible for maintaining food storage and hygiene in your area? Probe- social and economic power structure in the household (mother/father-in-laws, husbands).
- Who decides food preparation methods and why?
- What are the ways hands contact prepared and stored food? Whose hands get frequent contact? When, how and why? Explain at least one way.
- When is food for young children prepared during the day? How long is food stored?
- What do people do to keep their food safe? Probe- hand hygiene before preparing food, heating and reheating before serving food to child, child consumption of food and feeding methods. Who feeds, when and how?

##### Free roaming domestic animals

- How many domestic animals doesa typical household have in your area? Make a count. Identify the majority. Make at least 3 categories. Probe to cover all categories.
- Who takes care of animals and animal sheds and disposes of animal feces?
- When and why are animal feces visible in compound/inside household and for how long?
- How can a child come into contact with animals? Which animals do they like? Do parents allow children to touch animals?
- When are the chances that a child can put animal feces from the courtyard ground into their mouth? Probe- which animals, when and where?
- What do parents do if a child puts animal feces in their mouth? Why or why not?
- Are animals allowed in the house? In food preparation areas?

##### Contamination via flies

- What do people think of flies visible around their household environment?
- What are the types of flies you see around your household environment?
- Which one is most dangerous and why? Describe the most dangerous (size, color and other characteristics).
- What are the reasons for presence of different flies in house, courtyard and surrounding environment? Ask specifically type (size, color, other characteristics).
- Why are these flies found around/inside the household? Where do they come from? When is the fly density high and why?
- Is there a chance that these flies can land on food? How frequently? Probe- time and season.

##### Availability and type of toilet facility

- Where do the people in this area usually defecate? Probe- practice open defecation, individual or shared latrine.
- How many latrines does a household usually own? Probe- type and other facilities available for each toilet.Map the compound marking toilets and distances from drinking water source, pond or other water sources and from living room, kitchen etc.

- Who is responsible for cleaning and maintaining the toilet? Probe- man, women, specific roles, frequency.
- What are the hardware available in this community to dispose of feces from courtyard or inside the house? Probe- child and animal feces, adult human feces, feral animal feces?
- How does sanitation change during the different seasons of the year? Probe- when latrine pit is full, feces floats over pan, during rainy season, when latrine is broken.
- Where do the feces from the latrines end up? In ditches, ponds, drains, small rivers, other water bodies?
- What is easy about managing sanitation in your household? Please describe.

#### Module 54 – Focus Group Discussion Guideliens in Bengali for environmental assessment

**পরিবেশগত নিবিড় মূল্যায়ন (বাংলাদেশে ডায়রিয়া এবং স্যানিটেশন সুবিধার অন্তর্ভূক্তি, মলদুষণের গমনাগমনের মূল্যায়ন)**

**পানীয় জলের উৎসসমূহ, পরিচালনা এবং সংরৰণ চর্চাসমূহ**

- এই এলাকার খাবার পানির প্রধান প্রধান উৎসগুলি কী কী? জিজ্ঞাসা করম্নন - ধরণ, টিউবওয়েল, পুকুর বা অন্যান্য, সেগুলির ভৌত অবস্থা, দ্বিতীয় ও অন্যান্য উৎসসমূহ, মৌসুমী প্রভাবসমূহ। টিউবওয়েল ও পুকুরের পানির উপর গুরম্নত্ব বেশী দিন)। এই উৎসগুলি পর্যবেৰণ করম্নন এবং নৈর্ব্যক্তিকভাবে বর্ণনা করম্নন।
- গৃহস্থালীতে সাধারণত খাবার পানি সংগ্রহ করা কার দায়িত্ব? (জিজ্ঞাসা করম্নন- প্রাথমিক ও দ্বিতীয় সংগ্রহকারী, কখন ও কেন?)
- পানি সংগ্রহ ও রৰণাবেৰণের সাধারণ পদ্ধতিগুলি কী কী? এই পদ্ধতিগুলি পর্যায়ক্রমে লিখুন এবং পরিস্কারভাবে বোঝার জন্য আরও প্রয়োজনীয় প্রশ্ন করম্নন।
- আপনার গৃহস্থালীতে পান করার পানি সংগ্রহ করা কার দায়িত্ব? (জিজ্ঞাসা করম্নন- প্রাথমিক ও দ্বিতীয় সংগ্রহকারী, কখন ও কেন?)
- এই পানির উৎসগুলি বা সংরৰণকৃত পানি কিভাবে মানুষ বা পশু-পাখির মল দ্বারা দুষিত হতে পারে? এই সম্ভাব্য দুষণের গমনাগমন পথগুলি কী কী যা প্রাপ্ত বয়স্ক মানুষ, শিশু বা পশু-পাখির মলের দ্বারা দুষিত হয়? (অনুমানকৃত বা সম্ভাব্য এই মল দূষণের একটি গমনাগমন চিত্র অংকন করম্নন, নিবিড় সাৰাৎকার থেকে প্রাপ্ত চিত্রের সাথে চেক করম্নন, অতঃপর পরিবর্তন করম্নন)

**সহজপ্রাপ্যতা, পানি নিরাপদকরণ, পানি স্বল্পতা এবং পানির ব্যবহার**

- এই এলাকায় কখন কখন পানির স্বল্পতা দেখা দেয়?
- এই সময় এখানকার মানুষজন কী করেন?
- এই সময়ে অন্যান্য সময়ের সাথে পানি সংগ্রহ, সংরৰণ এবং ব্যবহারে কী ধরণের পার্থক্য দেখা যায়?
- আপনার গৃহস্থালীতে পানি বিশুদ্ধকরণের সাধারণ পদ্ধতিগুলি কী কী? সাধারণ বা জনপ্রিয় পদ্ধতিগুলি বর্ণনা করম্নন।

- কোন নির্দিষ্ট সময় কি রয়েছে যখন পানি নিরাপদকরণ বা বিশুদ্ধকরণ কম বা বেশী হয়ে থাকে?
- শিশুদের বয়সভেদসহ আপনার পরিবারের সকল সদস্যদের সাধারণভাবে পানি ব্যবহার সম্পর্কে বলুন।

#### হাতে জীবানুর সংক্রমণ এবং হাত ধোয়ার চর্চা

- কত ঘনঘন এবং কিভাবে আপনি হাত ধোন? জিজ্ঞাসা করুন্ন- প্রধান প্রধান সময়গুলি (বিভিন্ন ধরণের খাবার খাওয়ার আগে, টয়লেট ব্যবহারের পর এবং বাচ্চাকে শৌচানোর পর)। কোন একটি সাধারণ দিনের হাত ধোয়ার চর্চাগুলি বর্ণনা করুন্ন (হতে পারে গতকাল বা আজ)।
- মানুষ কি মনে করে যে তাদের কোন একটি বা উভয় হাতই মানুষের মল দ্বারা দুষিত হতে পারে? জিজ্ঞাসা করুন্ন- ঐ প্রধান প্রধান সময়গুলিতে হাত ধোয়া (একটি চিত্র অংকন করুন্ন যেসব কর্মকান্ড থেকে হাতে মলের সংক্রমন ঘটতে পারে)।
- মানুষেরা কি বাম বা ডান হাত বেশী বেশী ধোয়? কখন কখন এবং কেন?
- হাত ধোয়ার জন্য এখানে কোন কোন উপাদানগুলি সহজে পাওয়া যায়? জিজ্ঞাসা করুন্ন- সব ধরণের উপাদান, প্রাপ্তি ভেদে ও ঘন ঘন ব্যবহারে উপরি পর্যায়ে সাজান? কেন এবং কখন  কখন?
- কতবার আপনি আপনার শিশুর হাত পরিস্কার করেন/ধোন? কখন কখন এবং কিভাবে?
- কতবার আপনার শিশুরা তাদের নিজেদের হাত ধোয়? কখন কখন এবং কিভাবে?
- আপনি কি মনে করেন আপনার শিশুর হাত মানুষের মল দ্বারা দুষিত হতে পারে? পরিবারের কোন ব্যক্তি হাত ধোয়াকে উৎসাহিত বা প্রভাবিত করতে পারেন? জিজ্ঞসা করুন্ন- দাদা, শ্বশুর, স্বামী, স্কুলে যায় এমন বাচ্চারা।

## উঠানের ও ঘরের আশেপাশের মাটি দুষণ

- বাড়ীর উঠান কিভাবে মানুষের মল দ্বারা দুষিত হতে পারে? জিজ্ঞাসা করুন্ন- শিশু এবং বয়স্ক গণ, অসুস্থ্য ব্যক্তি, দিনে বা রাতে বা বৃষ্টির সময় খোলা জায়গায় মলত্যাগ করা। স্থানগুলি চিহ্নিত করুন্ন এবং বর্ণনা করুন্ন। তারা ব্যক্ত করতে পারে এমন একটি উদাহরণ দিতে বলুন। মলের উৎস ও তার সংক্রমনের একটি চিত্র অংকন করুন্ন।
- আপনার বাড়ীতে ছোট ছোট শিশুরা কোথায় পায়খানা করে? কখন কখন এবং কোথায়? জিজ্ঞাসা করুন্ন- সাধারণ, নির্দিষ্ট ও অস্থায়ী স্থানগুলি সম্পর্কে।
- ছোট শিশুদের মল কে কোথায় এবং কিভাবে অপসারণ করে? এই চর্চাগুলিতে কি কোন পার্থক্য হয়? পার্থক্যগুলি কী কী এবং কেন হয়? যেখানে যেখানে মল ফেলা হয় সেই স্থানগুলি পর্যবেৰণ করুন্ন এবং প্রত্যেক স্থানগুলি নিয়ে আরও প্রশ্ন জিজ্ঞাসা করুন্ন।
- এই পরিবারে কি কোন শিশুদের মলত্যাগ করার জন্য পটি রয়েছে?
- ছোট ছোট শিশুরা তাদের মায়ের/বয়স্ক দের কোল ছাড়া আর কোথায় কোথায় অধিকাংশ সময় কাটায়? স্থানগুলির বর্ণনা করুন্ন
- শিশুরা কত সময় উঠানে সময় কাটায় বা বারান দায় এবং কেন?
- তাদেরকে কে সঙ্গ দেয়? শিশুরা কি একা না অনেকে মিলে খেলাধুলা করে?
- বলুন কিভাবে আপনাদের বাচ্চারা মাটি ও মলের সংস্পর্শে আসে।
  - উঠান বা বারান দা বা ঘরের ভেতরের মাটি
  - বাচ্চাদের এবং পশু-পাখির মল
  - পশু-পাখির ধরন, প্রতিবেশীদের বাচ্চাদের
- কত সময় বাচ্চারা উঠানের মাটিতে সময় কাটায় এবং কেন? দলভেদে
  - <৩ বছর
  - <৩ -৬ বছর
  - ৬+বছর

- ক তাদের আচার-আচরণ লৰ্য করে যখন তারা উঠানের মাটিতে খেলাধুলা করে? জিজ্ঞাসা করম্নন- মায়ের বা অন্য যত্নগ্রহণকারীর উপস্থিতি এবং অনুপস্থিতিতে।
- খেলাধুলা বা সময় কাটানোর বেলায় বাচ্চারা কোন কোন জিনিস মুখে দিতে পারে? জিনিসগুলির নাম লিখুন, উপরি-পর্যায়ক্রমে সাজান ও বর্ণনা করম্নন।
- যখন শিশুরা উঠানের মাটি থেকে তাদের মুখে কোন কিছু দেয় তখন মায়েরা কী করেন?
- বাচ্চারা যখন মাটি থেকে কোন কিছু মুখে দেয় তখন মায়েরা কী ভাবেন বা মনে করেন?

**খাদ্য দূষণ**

- দয়া করে আমাকে এই এলাকার খাবার তৈরী এবং সংরৰণ প্রক্রিয়া সম্পর্কে বলুন।
- এই এলাকার সাধারণ খাবার সংরৰণ পদ্ধতিগুলি কী কী? প্রধান পদ্ধতিগুলি উপরি-পর্যায়ক্রমে সাজান এবং ব্যাখ্যা করম্নন।
- প্রতিদিনকার খাবার তৈরী ও সংরৰনে কে সবচেয়ে বেশী ভূমিকা পালন করে? কে তাদেরকে সেই দায়িত্ব দেয় ও কেন? জিজ্ঞাসা করম্নন- প্রধান প্রধান খাবার, নাস্ৰা, মৌসুমি খাবার।
- এই অঞ্চলে সাধারণত খাবার সংরৰণ ও তা স্বাস্থ্যসম্মত রাখার দায়িত্ব কার? জিজ্ঞাসা করম্নন- পরিবারের সামাজিক ও অর্থনৈতিক ৰমতা (শ্বশুর-শ্বাশুড়ী, স্বামী)
- খাবার তৈরীর পদ্ধতি সম্পর্কে কে সিদ্ধান্ৰ নেয় এবং কেন?
- তৈরীকৃত ও সংরৰিত খাবারগুলি কিভাবে হাতের সংস্পর্শে আসে? কার হাতে বার বার ধরা হয়?
- খাবার নিরাপদ রাখার জন্য কী কী করা হয়? জিজ্ঞাসা করম্নন- খাবার তৈরীর পূর্বে হাত ধোয়া, বাচ্চাকে খাওয়ানোর আগে বার বার খাবার গরম করা, শিশুদের খাবারের পরিমাণ এবং খাওয়ানোর পদ্ধতি। কে খাওয়ায়, কখন কখন ও কেন?
- বাচ্চাদের জন্য খাবার দিনের কোন সময়ে তৈরী করা হয়? কতৰণ সংরৰণ করা হয়?

**উন্মুক্তভাবে থাকা গৃহপালিত পশু-প্রাণীসমূহ**

- সাধারণত আপনাদের এলাকায় একটি সাধারণ গৃহস্থালীতে কতগুলি গৃহপালিত পশু-পাখি থাকে? গণনা করম্নন। সবচেয়ে বেশী কোনগুলি তা চিহ্নিত করম্নন। কমপৰে ৩ টি শ্রেণী করম্নন। সবগুলি শ্রেণী সম্পর্কে জিজ্ঞাসা করম্নন।
- পশু-পাখী ও খোয়ার বা গোয়ালঘর কে দেখাশোনা করে
- কখন কখন এবং কেন পশু-পাখীর মল আপনার বাড়ীর উঠানে ও আশে পাশে দেখা যায় এবং কত সময় ধরে?
- একটি বাচ্চা শিশু কিভাবে পশু-পাখীর সংস্পর্শে আসতে পারে? কোন প্রাণী তারা পছন্দ করে? মা-বাবারা কি বাচ্চাদেরকে পশু-পাখির সংস্পর্শে আসতে দেয়?
- এমন কি কোন সম্ভাবনা আছে যে শিশুরা উঠানের মাটি থেকে পশু-পাখির কোন মল মুখে দিতে পারে? কোন প্রাণীর, কখন ও কোথায়?
- বাচ্চারা যদি পশু-পাখির মল মুখে দেয় তবে মা-বাবারা কী করেন? কেন বা কেন নয়?
- বাড়ীতে কি পশুপাখি আসতে দেয়া হয়? রান্না ঘরে বা যেখানে খাবার তৈরী হয়?

**মশা-মাছির মাধ্যমে দূষণ/সংক্রমণ**

- আপনার বাড়ীর ভেতর এবং আশে পাশে বিভিন্ন ধরণের মাছির উপস্থিতি সম্পর্কে আপনার কী মনে হয়? সুনির্দিষ্টভাবে ধরণগুলি সম্পর্কে জিজ্ঞাসা করম্নন (আকৃতি, রং, অন্যান্য বৈশিষ্ট্যসমূহ)
- কোন ধরণের মাছি বাড়ীর চারপাশে দেখা যায়? কোনগুলি বেশী বিপদজনক ও কেন? বেশী বিপদজনকগুলি বর্ণনা করম্নন (আকৃতি, রং, অন্যান্য বৈশিষ্ট্যসমূহ)
- এই ধরণের মশা-মাছিগুলি আপনার ঘরের ভেতর বা আশে পাশে কেন পাওয়া যায়? এগুলি কোথা থেকে আসে? কখন কখন মশা-মাছির উপদ্রব বেশী হয় ও কেন?

- এমনি কি কোন সুযোগ আছে যে মশা-মাছি খাবারের উপর বসে? কত ঘন ঘন এমন হয়? জিজ্ঞাসা করুন- সময় ও ঋতু।
- আপনি কি মনে করেন যে মাছিরা আপনার পরিবারের জন্য কোন স্বাস্থ্য ঝুকি বয়ে আনে? কেন অথবা কেন নয়?

**টয়লেট সুবিধাদির ধরণ ও সহজলভ্যতা**

- এ এলাকার মানুষজন সচরাচর কোথায় সচরাচর পায়খানা/মলত্যাগ করেন? জিজ্ঞাসা করুন- খোলা জায়গায়, ব্যক্তিগত/নিজস্ব বা অন্যের ল্যাট্রিনে।
- একটি পরিবারের কয়টি ল্যাট্রিন আছে? জিজ্ঞাস করুন- প্রতিটি ল্যাট্রিনের ধরন ও অন্যান্য সুবিধাসমূহ। এই বাড়ীটির একটি চিত্র অংকন করুন যেখানে ল্যাট্রিন/টয়লেট সুবিধাদি এবং খাবার পানির উৎস, পুকুর বা অন্যান্য উৎস, থাকার ঘর ও রান্না ঘর থেকে তার দূরত্ব চিহ্নিত করুন।
- টয়লেট ব্যবস্থাপনা ও পরিস্কার রাখার দায়িত্ব কার? জিজ্ঞাসা করুন- পুরুষ, নারী, নির্দিষ্ট ভূমিকা, কত সময় পর পর?
- উঠান থেকে অপসারণ করার জন্য এই পরিবারের কোন কোন যন্ত্র-পাতি রয়েছে। জিজ্ঞসা করুন- শিশু ও পশুর মল, বয়স্ক দের ও অপোষা প্রাণীদের।
- বছরের বিভিন্ন মৌসুমে কী হয়? জিজ্ঞাস করুন- ল্যাট্রিন যখন ভরে যায়, প্যানের উপর মল ভাসতে থাকে, বা ল্যাট্রিন যখন ভেঙ্গে যায়।
- ল্যাট্রিন থেকে মল কোথায় গিয়ে শেষ পৌছে? নালায়, পুকুরে, ডোবায়, ছোট নদী, গাঙ্গে, বা অন্যান্য কোন পানিতে।
- পরিবারের সদস্যরা মলত্যাগ ও পরিবেশ মলমুক্ত রাখার জন্য কী কী অসুবিধার মুখোমুখি হন বর্ণনা করুন।

PROTOCOL
Biomedical
Non-Exempt
Berkeley

Protocol # 2011-09-3652
Date Printed: 07/31/2018

-----------------------------------------------------------------------------------
---

**Protocol Title:** Measuring the benefits of sanitation, water quality, handwashing and nutrition interventions for improving health and development in rural Bangladesh

**Protocol Status:** APPROVED

**Date Submitted:** 10/26/2017

**Approval Period:** **10/30/2017-11/03/2018**

**Important Note:** **This Print View may not reflect all comments and contingencies for approval. Please check the comments section of the online protocol. Questions that appear to not have been answered may not have been required for this submission. Please see the system application for more details.**

*** * * Attached Document * * ***

| Document Name | Created Date |
|---|---|
| 2011-09-3652_Colford_1aa-Consent-Spillover-qualitative-v21-English.pdf | 10/26/2017 |

icddr,b
KNOWLEDGE FOR GLOBAL LIFESAVING SOLUTIONS

**Appendix 1aa: English consent form for in-depth interviews about spillovers**

**International Centre for Diarrhoeal Disease Research, Bangladesh (ICDDR,B)**

Protocol Title: Effect of hand washing, water treatment, sanitation and nutritional supplement interventions on child health and development in rural Bangladesh

Principal Investigator's name: Dr. Leanne Unicomb

**Spillover substudy**

#### Purpose of the research

Hello (Assalamualaikum/Nomoshkar). My name is ________ and I work with the ICDDR,B (Cholera Hospital) in Dhaka. We are interested in conducting research on diarrheal diseases in children to learn how it affects their physical and mental development. Through this research we want to learn about the health benefits of some simple interventions to improve the sanitation, water quality, hygiene practices or nutritional status for children under five. We want to understand whether children who live in the same communities as children receiving these interventions benefit from them as well. This research is being done in collaboration with scientists at the University of California, Berkeley in the United States.

#### Why are we inviting you to participate in the study?

Because you are participating in research with ICDDR,B (Cholera Hospital) or because you live near participants in this research, we are interested in asking you some questions about who you know in this village and where you get information about health.

#### What is expected from the participants of the research study?

If you decide to enroll in this study, we will ask you a series of questions about who you know in this village and where you get information about health. It is very difficult to write all the conversation in a note pad, so, if you agree, we would like to record this sessions in a tape recorder. We would also like to take some notes from this session. This session will last about an hour.

#### Risks & Benefits

There are minimal risks involved in this study. There is a slight risk of breach of confidentiality. Your participation will help us to gain knowledge on diarrheal disease in children. There is no anticipated direct benefit to you for participating in this study. However, it is hoped that the information gained from the study will help improve our knowledge of how diarrhea affects young children.

#### Confidentiality

All data and specimens collected will be kept confidential to the greatest extent possible. Researchers at ICDDR,B will have access to some of your personal information that could identify you, such as your name. They will remove the identifiable information before sharing it with researchers at partner institutions, such as UC Berkeley. Your responses and audio recordings will be kept in a locked cabinet, and computer-based data, will be stored in an encrypted format on a password-protected server.

Other persons who may have access to your responses and audio recordings include research groups that oversee the safety of the study including institutes such as the University of California Berkeley and the Bill and Melinda Gates Foundation, USA. None of the researchers outside of ICDDR,B will be able to identify you or your compound members from information you provide. We will use the information only for the purpose of the study. We will not use your name when we share and publish the results of this study. We expect the steps we take will keep all of your information confidential, but it is possible that because of mistakes or unforeseen events, it could become compromised.

**Future use of information**

The information collected from this study may be shared with other researchers if needed, but we will strictly maintain your confidentiality and privacy. We will store the information we collect for a long time after the end of the study.

**Right not to participate and withdraw**

Taking part in the study is completely voluntary. You can drop out of this study at any time, even in the middle of an interview, without penalty or loss of benefits to which you are otherwise entitled. If you wish, we will withdraw your permission and remove your data from the database. You have the right to refuse participation in this study, which will not affect your family's treatment at the Cholera Hospital in the future.

**Principle of compensation**

You need not pay us to take part in this study, and similarly we will not pay you money for attending in the study.

**Persons to contact:**

If you have any question, you can ask me any time. If you have additional questions about the survey, you may contact:

Masud Parvez: phone: 01817541872, Center of Communicable Diseases, ICDDR,B, Mohakhali, Dhaka 1212.

If you have questions about your rights as a participant of a research study, or if you think some harm has been done to you because of the survey, you may contact or meet M.A. Salam Khan, IRB Secretariat:

M. A. Salam Khan, IRB Secretariat, phone: 9886498 or PABX 8860523-32 ext. 3206

If you agree to participate in our study, please indicate that by putting your signature or your left thumb impression at the specified space below.

Thank you for your cooperation

________________________________________ ____________________

Signature or left thumb impression of Participant Date

________________________________________ ____________________

Signature or left thumb impression of the witness Date

________________________________________ ____________________

Signature of the PI or his/her representative Date

PROTOCOL
Biomedical
Non-Exempt
Berkeley

Protocol # 2011-09-3652
Date Printed: 07/31/2018

---------------------------------------------------------------------------------------

| | |
|---|---|
| **Protocol Title:** | Measuring the benefits of sanitation, water quality, handwashing and nutrition interventions for improving health and development in rural Bangladesh |
| **Protocol Status:** | APPROVED |
| **Date Submitted:** | 10/26/2017 |
| **Approval Period:** | **10/30/2017-11/03/2018** |
| **Important Note:** | **This Print View may not reflect all comments and contingencies for approval. Please check the comments section of the online protocol. Questions that appear to not have been answered may not have been required for this submission. Please see the system application for more details.** |

*** * * Attached Document * * ***

| Document Name | Created Date |
|---|---|
| 2011-09-3652_Colford_2aa-Consent-Spillover-qualitative-Bengali.pdf | 10/26/2017 |

icddr,b
KNOWLEDGE FOR GLOBAL LIFESAVING SOLUTIONS

**Appendix 2aa: English consent form for in-depth interviews about spillovers**
**International Centre for Diarrhoeal Disease Research, Bangladesh (ICDDR,B)**

Protocol Title: Effect of hand washing, water treatment, sanitation and nutritional supplement interventions on child health and development in rural Bangladesh

**Spillover substudy**

<u>cavb M‡el‡Ki bvg</u>: W: wj‡qb BDwbK¤

M‡elYvi D‡Ïk¨

AvmmvjvgAvjvBKg/ bg¯‹vi| Avgvi bvg---------------- | Avwg XvKvi AvBwmwWwWAvi, we (K‡jiv nvmcvZvj) †Z KvR Kwi| Avgiv wkï‡`i Wvqwiqv †ivM Ges Bnv wKfv‡e Zv‡`i kvixwiK I gvbwmK eŵ×‡K cfweZ K‡i Zv wb‡q M‡elYv Ki‡Z AvMnx| Avgiv GB M‡elYvi gva¨‡g wKQ mvavib B›Uvi‡fbkb †hgb- cvqLvbvi Dbqb, cwbi ¸bMZgvb Ges ¯^v¯'m¤§Z Af¨vm ev cvP eQ‡ii †QvU ev"Pv‡`i cwóMZ Ae¯'v BZ¨w`i Dbq‡bi gva¨‡g ¯^v¯'MZ mweav m¤ú‡K Rvb‡Z Pvw"Q| Avgiv ‡evSvi †Póv KiwQ GKB GjvKvq cvkvcvwk emevmKvix Ab¨vb¨ wkïiv mvavib B›Uvi‡fbkb cvIqv wkï‡`i g‡Zv GKB mweav jvf K‡i wKbv|

Avgiv †Kb Avcbv‡K GB M‡elYvq AskMn‡Y AvgšY Rvbvw"Q?
Avgiv GB Lvbv‡K M‡elYvq AšÍf³ Ki‡Z Pvw"Q KviY Avcwb AvB, wm, wW, wW, Avi, we (K‡jiv nvmcvZvj) -i D³ M‡elYvq B‡Zvg‡a¨ AskMnb K‡i‡Qb A_ev GB M‡elYvq AšÍf³ †Kvb AskMnbKvixi KvQvKwQ emevm Ki‡Qb| Avcwb GB Mv‡gi Kv‡K Kv‡K wP‡bb Ges Avcwb †Kv_v †_‡K ¯^v¯'welqK Z_¨ cvb- Gme wel‡q Avgiv Avcbv‡K wKQ ck wRÁvmv Kie|

M‡elYvq AskMnYKvixi Kv‡Q cZ¨vkv Kx?
Avcwb GB M‡elYvq AskMnY Ki‡Z m¤§Z n‡j, Avcwb GB Mv‡gi Kv‡K Kv‡K wP‡bb  Ges Avcwb †Kv_v †_‡K ¯^v¯'welqK Z_¨ cvb- Gme wel‡q Avgiv Avcbv‡K KwZcq ck wRÁvmv Kie| †h‡nZ Avcbvi mv‡_ Av‡jvPbvi mKj welq LvZvq wj‡L ivLv LeB `i/n, ZvB Avcwb m¤§Z _vK‡j Avgiv‡`i GB Av‡jvPbv cewU †Uc‡iKW‡i avib K‡i ivL‡Z PvB| Avgiv GB Av‡jvPbvi wKQ welq wjwce× K‡i I ivL‡Z PvB| GB Av‡jvPbwU‡Z cvq GK N›Uv mgq e¨q n‡e|

SwK Ges mweav
GB M‡elYvq AskMn‡Y Avcbvi SwK bb¨Zg| Z‡e Avcbvi cwiPqmsµvšÍ Z_¨mgn †Mvcb bv _vKvi mvgvb¨ m¤vebv i‡q‡Q| wKš' Avcbvi wkïi AskMnY, wkï‡`i Wvqwiqv †ivM m¤ú‡K Ávbjv‡f Avgv‡`i‡K mnvqZv Ki‡e|

†MvcbxqZv
Avgiv Avcbvi/ Avcbv‡`i †`Iqv mg¯Í Z_¨ I bgbvi †MvcbxqZv eRvq ivLvi Rb¨ m‡ev"P †Póv Kie| Avcbvi/Avcbv‡`i †`Iqv wKQ Z_¨ AvBwmwWwWAvi,we-i <u>M‡elK‡</u>`i Kv‡Q _vK‡e hvi d‡j Avcbv‡K mbv³ Kiv m¤e, ‡hgb Avcbvi bvg| Zviv Avcbvi/Avcbv‡`i †`Iqv Z_¨ ‡_‡K Avcbvi/Avcbv‡`i  mbv³KiY gjK Z_¨ mgn ev` w`‡q Ab¨vb¨ M‡elK‡`i mv‡_ ‡kqvi Ki‡e (‡hgb Av‡gwiKvi K¨wj‡dvwbqv evK‡j wekwe`¨vj‡qi <u>M‡elK)</u>, hvi d‡j ‡Kvb M‡elKB Avcbv‡K wPwýZ Ki‡Z cvi‡e bv | Avcbvi/Avcbv‡`i †`Iqv mg¯Í DËi I †iKWKZ Z_¨ Zvjve× K‡i ivLv n‡e Ges KgwcDUv‡i Z_¨vejx cvmIqvW w`‡q msi¶b Kiv n‡e |

Ab¨vb¨ e¨w³ A_ev M‡elYvi mv‡_ mswkøó `j M‡elYvi wbivcËvi c‡qvR‡b Avcbvi ‡`Iqv mg¯Í DËi I †iKWKZ Z_¨ ‡`L‡Z cv‡i | Giv n‡jb- Av‡gwiKvi K¨wj‡dvwbqv evK‡j wekwe`¨vj‡qi <u>M‡elK</u> I M‡elYvi ¯úÝi (wej I ‡gwjÛv ‡MUm dvD‡Ûkb) | Avcbvi †`Iqv Z_¨ †_‡K AvBwmwWwWAvi,we i <u>M‡elK‡</u>`i Qvov †KD Avcbv‡K Ges Avcbvi ewoi Ab¨ †Kvb m`m¨‡K mbv³ Ki‡Z cvi‡e bv | Avgiv ïagvÎ M‡elYvi Kv‡R GB Z_¨ e¨envi Kie Ges Avcbvi bvg A_ev cwiP‡qi †Kvb mÎ GB M‡elbv-msµvšÍ †Kvb wi‡cvU/ cKvkbvq D‡jøL Kiv n‡e bv| Avcbvi KvQ ‡_‡K cvß mKj Z‡_¨i m‡ev"P

‡MvcbxqZv i¶vi Rb¨ Avgiv mevZ¥K ‡Póv Kie, Zv m‡Ë¡I fjekZt AbvKvw•LZ †Kvb NUbvi Kvi‡b ‡MvcbxqZv i¶v wbwðZ Ki‡Z bv cvi‡j ‡mRb¨ gvRbv cv_bv KiwQ|

fwel¨‡Z Z‡_¨i e¨envi
GB M‡elYvq msMnxZ Z_¨ c‡qvR‡b Ab¨vb¨ M‡el‡Ki mv‡_ wewbgq n‡Z cv‡i, wKš Z‡_¨i †MvcYxqZv K‡Vvi fv‡e cvjb Kiv n‡e|

‡¯”Qv AskMnY

G M‡elYvq Avcbvi AskMnY m¤úb ‡m”QvgjK Ges Avcwb/Avcbviv wkïi AskMnY Kiv ev bv Kiv m¯úYfv‡e Avcbvi B”Qvi Dci wbfi Ki‡Q| Avcwb B”Qv Ki‡j ‡h †Kvb mg‡q, GgbwK Av‡jvPbvi gvSc‡_I Avcbvi AskMn‡bi AbgwZ cZ¨vL¨vb Ki‡Z cvi‡eb| Avcwb PvB‡j Avgiv Avcbvi M‡elYvq AskMn‡bi AbgwZ ewZj Ges Avcbvi †`Iqv mKj Z_¨I g‡Q †dje| GB M‡elYvq AskMnY bv Kivi AwaKvi Avcbvi Av‡Q Ges G Rb¨ fwel¨‡Z Avcbvi cwiev‡ii m`m¨‡`i K‡jiv nvmcvZv‡j ¯v¯'¨‡mev Mn‡Yi Dci †Kvb cfve co‡e bv|

¶wZciY
GB M‡elYvq AskMn‡Yi Rb¨ Avcbvi mivmwi †Kvb ai‡Yi A_‰bwZK LiP/ e¨q n‡e bv Ges GKBfv‡e Avcwb GB M‡elYvq AskMn‡Yi Rb¨ mivmwi †Kvb ai‡Yi A_‰bwZK mnvqZvI cv‡eb bv|

‡hvMv‡hvM
Avcbvi †Kv‡bv ck _vK‡j Avgv‡K wRÁvmv Ki‡Z cv‡ib| hw` M‡elYv m¤úwKZ AwZwi³ †Kvb ck _v‡K Zvn‡j Avcwb gvm` cvi‡fR (wmwmwW, AvBwmwWwWAvi,we, gnvLwj, XvKv-1212) Gi mv‡_ 01817541872 ‡gvevBj b¤‡i mivmwi †hvMv‡hvM Ki‡Z cv‡ib| GB M‡elYvq Avcbvi AwaKvi-msµvš Í †Kvb ck _vK‡j A_ev M‡elYvi Kvi‡Y Avcbvi †Kvb ¶wZ n‡Z cv‡i e‡j hw` g‡b K‡ib, Zvn‡j Avcwb 9886498 A_ev 8860523 (G· - 3206) †Uwj‡dvb b¤‡i A_ev mivmwi Gg G mvjvg Lvb, AvBAviwe †m‡µUwi‡qU Gi mv‡_ †hvMv‡hvM Ki‡Z cv‡ib|
Avcwb hw` GB M‡elYvq AskMnY Ki‡Z AvMnx nb Zvn‡j wb‡gi wbawiZ ¯v‡b ¯v¶i A_ev evg e×v½jxi Qvc w`b|
Avcbvi mn‡hwMZvi Rb¨ ab¨ev`|

| | |
|---|---|
| ........................................... | ................................. |
| AskMnbKvixi ¯v¶i/evg e×v½wji Qvc | ZwiL |
| ........................................... | ................................. |
| mv¶xi ¯v¶i/evg e×v½wji Qvc | ZwiL |
| ........................................... | ................................. |
| wc AvB/ M‡el‡Ki cwZwbwai ¯v¶i | ZwiL |

wØgZ:
Awg PvB bv Avgvi wkïi i³ Ges cvqLvbvi bgbv `xN mgq a‡i msi¶Y Kiv †nvK ☐

PROTOCOL
Biomedical
Non-Exempt
Berkeley

Protocol # 2011-09-3652
Date Printed: 07/31/2018

---------------------------------------------------------------------------------
---

**Protocol Title:** Measuring the benefits of sanitation, water quality, handwashing and nutrition interventions for improving health and development in rural Bangladesh

**Protocol Status:** APPROVED

**Date Submitted:** 10/26/2017

**Approval Period:** **10/30/2017-11/03/2018**

**Important Note:** **This Print View may not reflect all comments and contingencies for approval. Please check the comments section of the online protocol.
Questions that appear to not have been answered may not have been required for this submission. Please see the system application for more details.**

*** * * Attached Document * * ***

| Document Name | Created Date |
|---|---|
| EE Collection Form v7_12.11.2014 (Paper).pdf | 10/26/2017 |

#### EE Sample Collection Form – Section 1: Child Identification & Medical History

**Cluster ID |__|__|__| Mother ID |__|__|**

| | | |
|---|---|---|
| 1. Cluster ID and Mother ID | ### and ## | \|__\|__\|__\| **and** \|__\|__\| |
| 2. ID of MT/FRA/FRO | ## | \|__\|__\| |
| 3. Name of MT/FRA/FRO (choose 1): | 1. Balay Chand Sikder<br>2. Mary Marget<br>3. ParimolSarker<br>4. Dipali Rani Das<br>5. SaniaAkter Dolly<br>6. SyedaFatemaKhatun<br>7.Md. AltafurRahman<br>8. AmalSarker<br>9. Nasrin Sultana<br>10. Md. NurulAlam<br>11. Fatema Begum Sumi<br>12. SyedaLuthfaFamida<br>13. Shahjahan Ali<br>16. NitaiHalder | |
| 4. Date of Data Collection | DD/MM/YYYY | \|__\|__\| / \|__\|__\| / \|__\|__\|__\|__\| |
| 5. Consent<br><br>Note: If consent given, skip next question. If no consent, answer Q6 and skip to end. | Y/N | \|__\| |
| 6. Why Was Consent Not Given? | 1. Migration<br>2. Household head Is not home<br>3. Primary caregiver Is not home<br>4. Child is not home<br>5. Child is sick<br>6. Household head refused<br>7. Primary caregiver refused<br>8. Too busy<br>9. No reason given<br>10. Other | \|__\|__\| |
| 7. Verify Primary Caregiver's Identity: Does Caregiver Match Household and Cluster ID?<br><br>Note: If no, then skip rest of form and alert SFRO that our team is in the wrong location. | Y/N | \|__\| |

**EE Sample Collection Form – Section 1: Child Identification & Medical History**

**Cluster ID |__|__|__| Mother ID |__|__|**

| | | |
|---|---|---|
| 8. Identification of Primary Caregiver | 1. Child's Mother<br>2. Child's Maternal Grandparents<br>3. Child's Paternal Grandparents<br>4. Child's Maternal Aunt/Uncle<br>5. Child's Paternal Aunt/Uncle<br>6. Child's Father (And His New Partner/Wife)<br>7. Other: Specify | |__| |
| 9. Mobile Number of Household Head | ###########<br>99. Don't know/not available | |__|__|__|__|__|__|__|__|__|__|__|<br>|__|__|
| 10. Mobile Number of Primary Caregiver | ###########<br>99. Don't know/not available | |__|__|__|__|__|__|__|__|__|__|__|<br>|__|__|
| 11. Number of Eligible Children | # of Eligible Children | |__| |
| 12. Child Full Name | | |
| 13. Child Nickname | | |
| 14. Gender of Child | 1. Male<br>2. Female | |__| |
| 15. Date of Birth | DD/MM/YYYY | |__|__| / |__|__| / |__|__|__|__| |
| 16. Source of Date of Birth (DOB) | 1. Confirmed DOB by valid vaccination card<br>2. Mother/Relative remember DOB<br>3. Both 1 & 2<br>99. Don't know | |__|__| |
| 17. Age | ### Days | |__|__|__| Days |
| 18. ASK: "How long ago did [CHILD NAME] take any antibiotics? (Provide examples from the 'List of Common Antibiotics and Medicine' in the EE Sampling Handbook) | ## Days<br>## Months<br>88. Never<br>99. Don't know | |__|__| Days<br>|__|__| Months<br>|__|__| |
| 19. ASK: "Can you tell us how many times in the last 3 months this child has used antibiotics?"<br>Note: If answer to Q19 is 0, 88, or 99, then skip to Q22. If answer to Q19 is > 0, then answer Q20 and Q21. | ## Times<br>88. Never<br>99. Don't know | |__|__| Times<br>|__|__| |

**EE Sample Collection Form – Section 1: Child Identification & Medical History**

**Cluster ID |__|__|__| Mother ID |__|__|**

| | | |
|---|---|---|
| | 20. ASK: "Please try and recall the name of each antibiotic this child used in the last 3 months." (90 days before collection date) | 21. ASK: "How many total days did this child use this antibiotic?" (For Each Episode) |
| | 1. Cotrimoxazole<br>2. Amoxycillin<br>3. Flucloxacillin<br>4. Ciprofloxacin<br>5. Erythromycin<br>6. Azythromycin<br>7. Nalidixic acid<br>8. Doxycycline<br>9. Betapen (Penicillin)<br>10. Chloramphenicol<br>11. Metronidazole<br>12. Other: Specify<br>99. Don't know | ## Days<br>99. Don't Know |
| Episode 1 | \|__\|__\| (choose from list above) | \|__\|__\| Days |
| Episode 2 | \|__\|__\| (choose from list above) | \|__\|__\| Days |
| Episode 3 | \|__\|__\| (choose from list above) | \|__\|__\| Days |
| Episode 4 | \|__\|__\| (choose from list above) | \|__\|__\| Days |
| Episode 5 | \|__\|__\| (choose from list above) | \|__\|__\| Days |
| 22. ASK: "Has this child had malaria in the last 1 month?" | 1. Yes, diagnosed in a clinic/by a physician<br>2. Yes, not diagnosed in a clinic/ by a physician<br>3. No<br>4. Not applicable<br>99. Don't know | \|__\|__\| |
| 23. ASK: "Has this child had malaria in the last 6 months?" | 1. Yes, diagnosed in a clinic/by a physician<br>2. Yes, not diagnosed in a clinic/by a physician<br>3. No<br>4. Not applicable<br>99. Don't know | \|__\|__\| |

#### EE Sample Collection Form – Section 1: Child Identification & Medical History

**Cluster ID |__|__|__| Mother ID |__|__|**

| | | |
|---|---|---|
| 24. ASK: FOR CHILDREN 6 MONTHS AND OVER "In the last 6 months, has [CHILD NAME] received a pill or drug for intestinal worms?" (Provide examples from the 'List of Common Antibiotics and Medicine' in the EE Sampling Handbook)<br><br>Note: If child is <6 months old, enter 3 and skip to end. If answer to Q24 is 2 or 99, then skip Q25-27 and answer Q 28-29. | 1. Yes<br>2. No<br>3. Not applicable<br>99. Don't know | \|__\|__\| |
| 25. ASK: "Where did [CHILD NAME] receive the drug for intestinal worms?" | 1. At Home / In Village<br>2. At A Clinic or Health Facility<br>3. At A School<br>4. From icddr,b<br>99. Don't know | \|__\|__\| |
| 26. ASK: "Approximately how long ago did [CHILD NAME] receive the deworming drug?" | ## Days<br>## Months<br>99. Don't know | \|__\|__\| Days<br><br>\|__\|__\| Months<br><br>\|__\|__\| |
| 27. ASK: "Please try and recall the name of the deworming drug [CHILD NAME] took." (Provide examples from the 'List of Common Antibiotics and Medicine' in the EE Sampling Handbook) | 1. Albendazole<br>2. Mebendazole<br>3. Other: Specify<br>99. Don't know | \|__\|__\| |
| 28. ASK: "Does [CHILD NAME] typically wear shoes/socks/sandals?" | 1. Yes<br>2. No<br>99. Don't know | \|__\|__\| |
| 29. OBSERVE: Is child wearing shoes/socks/sandals? | 1. Yes<br>2. No<br>99. Don't know | \|__\|__\| |

**Cluster ID |__|__|__| Mother ID |__|__|**

#### EE Sample Collection Form – Section 1: Child Identification & Medical History

**ASK: "Thank you. Now I am going to ask you some questions about the health of your [child/children]. Please answer each question as accurately as you can. If you don't know the answer to a question, say "I don't know". We will begin with [NAME OF CHILD]."**

| ASK: "Did [NAME OF CHILD] have [SYMPTOM]: | A | B | C | D | E |
|---|---|---|---|---|---|
| | **Today**<br>1. Yes<br>2. No<br>99. Don't know | **Yesterday**<br>1. Yes<br>2. No<br>99. Don't know | **Day before Yesterday**<br>1. Yes<br>2. No<br>99. Don't know | **In the last 7 days** (since this day last week)<br>1. Yes<br>2. No<br>99. Don't know | **In the last 2 weeks** (since this day 2 weeks ago)<br>1. Yes<br>2. No<br>99. Don't know |
| 30. Fever | | | | | |
| 31. Diarrhea | | | | | |
| 32. 3 or more bowel movements in 24 hours | | | | | |
| 33. Number of bowl movements each day | \|__\|__\|times | \|__\|__\|times | \|__\|__\|times | | |
| 34. Watery or soft stool (unformed) | | | | | |
| 35. Blood in the stool | | | | | |
| 36. Skin rash (anywhere on the body) | | | | | |
| 37. Constant cough | | | | | |
| 38. Congestion / runny nose | | | | | |
| 39. Panting / wheezing / difficulty breathing | | | | | |
| 40. Bruising, scrapes or cuts | | | | | |
| 41. Toothache / teething | | | | | |

| | | | |
|---|---|---|---|
| 42. ASK: "How long ago did [CHILD NAME] have diarrhea?"<br><br>Note: Q42 will be asked if Q31 turns out yes for column D and/or E (ie, Q42 will not be asked for col A, B and C of Q31. Also validate Q42 by number of days. For example, if Q42 is asked for col D, the validation is not more than 7 days and if Q42 is asked col E, the validation is not more than 14 days. | ## Days<br>77. Not applicable<br>99. Don't know | D<br><br>**In the last 7 days** (since this day last week)<br><br>\|__\|__\| Days | E<br><br>**In the last 2 weeks** (since this day 2 weeks ago)<br><br>\|__\|__\| Days |
| 43. ASK: "During that specific episode of diarrhea, how many consecutive days did it last?"<br><br>Note: Q43 is linked to Q42, i.e., Q43 will be asked only when Q42 is asked. Also validate Q43 by number of days. For example, if Q43 is asked for col D, the validation is not more than 7 days and if Q43 is asked col E, the validation is not more than 14 days. | ## Days<br>77. Not applicable<br>99. Don't know | D<br><br>**In the last 7 days** (since this day last week)<br><br>\|__\|__\| Days | E<br><br>**In the last 2 weeks** (since this day 2 weeks ago)<br><br>\|__\|__\| Days |

#### EE Sample Collection Form – Section 2: Anthropometry

**Cluster ID |__|__|__| Mother ID |__|__|**

| | | |
|---|---|---|
| 1. Cluster ID and Mother ID | ### and ## | \|__\|__\|__\| **and** \|__\|__\| |
| 2. ID of MT/FRA/FRO | ## | \|__\|__\| |
| 3. Name of MT/FRA/FRO (choose 1): | 4. Dipali Rani Das<br>5. SaniaAkter Dolly | |
| 4. Date of Data Collection | DD/MM/YYYY | \|__\|__\| **/** \|__\|__\| **/** \|__\|__\|__\|__\| |
| 5. Number of Eligible Children | # of Eligible Children | \|__\| |
| 6. Child Full Name | | |
| 6A. Type of Scale Used for Weight Measurement | 1. Seca 383 (Infant Scale)<br>2. Seca 874 (Mother-Infant Scale)<br>3. Seca 354 (Infant Scale New) | \|__\| |
| 7. Weight of Child – Measurement #1 | Weight (kg)##.### | \|__\|__\|.\|__\|__\|__\| kg |
| 8. Weight of Child – Measurement #2 | Weight (kg) ##.### | \|__\|__\|.\|__\|__\|__\| kg |
| 9. Weight of Child – Measurement #3 | Weight (kg) ##.### | \|__\|__\|.\|__\|__\|__\| kg |
| 10. **Median** Weight of The Child | Weight (kg) ##.### | \|__\|__\|.\|__\|__\|__\| kg |
| 11. Was Child Wearing Clothing During Weight Measurement? (Multiple answers) | 1. No clothes<br>2. Underpants/shorts<br>3. Shirt<br>4. Pants/trousers<br>5. Dress/frock<br>6. Other: Specify | \|__\|<br>\|__\|<br>\|__\|<br>\|__\| |

**Cluster ID |__|__|__| Mother ID |__|__|**

#### EE Sample Collection Form – Section 2: Anthropometry

| | | |
|---|---|---|
| 12. Length or Height?<br>(If ≥ 2 years of age, measure the child's height; if < 2 years of age, measure the child's length) | 1. Length measured<br>2. Height measured | \|__\| |
| 13. Length/Height of Child – Measurement #1 | Length/Height (cm)<br>###.# | \|__\|__\|__\|.\|__\| cm |
| 14. Length/Height of Child – Measurement #2 | Length/Height (cm)<br>###.# | \|__\|__\|__\|.\|__\| cm |
| 15. Length/Height of Child – Measurement #3 | Length/Height (cm)<br>###.# | \|__\|__\|__\|.\|__\| cm |
| 16. Head Circumference of Child – Measurement #1 | Circumference (cm)<br>##.# | \|__\|__\|.\|__\| cm |
| 17. Head Circumference of Child – Measurement #2 | Circumference (cm)<br>##.# | \|__\|__\|.\|__\| cm |
| 18. Head Circumference of Child – Measurement #3 | Circumference (cm)<br>##.# | \|__\|__\|.\|__\| cm |
| 19. Mid Upper Arm Circumference of Child – Measurement #1 | Circumference (cm)<br>##.# | \|__\|__\|.\|__\| cm |
| 20. Mid Upper Arm Circumference of Child – Measurement #2 | Circumference (cm)<br>##.# | \|__\|__\|.\|__\| cm |
| 21. Mid Upper Arm Circumference of Child – Measurement #3 | Circumference (cm)<br>##.# | \|__\|__\|.\|__\| cm |
| 22. Anthropometry Data Obtained?<br>Note: If Q22 is 1, skip to end. | 1. All<br>2. Partial<br>3. None | \|__\| |
| 23. Why Was Anthropometry Data Not Collected? | 1. Child Did Not Cooperate<br>2. Household Head Refused<br>3. Primary Caregiver Refused<br>4. Other | \|__\| |

**Cluster ID |__|__|__| Mother ID |__|__|**

#### EE Sample Collection Form – Section 3: Blood Sample Collection

| | | |
|---|---|---|
| 1. Cluster ID and Mother ID | ### and ## | \|__\|__\|__\| **and** \|__\|__\| |
| 2. ID of MT/FRA/FRO | ## | \|__\|__\| |
| 3. Name of MT/FRA/FRO (choose 1): | 1. Balay Chand Sikder<br>2. Mary Marget<br>3. ParimolSarker<br>14. Md. Shariful Islam<br>15. Shimul Das<br>16. NitaiHalder | |
| 4. Date of Sample Collection | DD/MM/YYYY | \|__\|__\| / \|__\|__\| / \|__\|__\|__\|__\| |
| 5. Number of Eligible Children | # of Eligible Children | \|__\| |
| 6. Child Full Name | | |
| 7. Date of Child's Last Meal/Snack/Breastfeed | DD/MM/YYYY | \|__\|__\| / \|__\|__\| / \|__\|__\|__\|__\| |
| 8. Time of Child's Last Meal/Snack/Breastfeed | 24-hour scale<br>Hours :Minutes | \|__\|__\|:\|__\|__\| |
| 9. Enter Information About Child's Last Meal/Snack/Breastfeed | 1. Breast Milk Only<br>2. Snack Only<br>3. Meal Only<br>4. Meal and Snack Only<br>5. Snack and Breast Milk Only<br>6. Meal and Breast Milk Only<br>7. Meal, Snack, and Breast Milk | \|__\| |
| 10. Blood Collection Tube Sample Obtained?<br><br>Note: If no blood draw, then Q14 is asked and rest of form skipped. | 1. all (5 ml)<br>2. partial (< 5 ml)<br>3.none (0 ml) | \|__\| |

**EE Sample Collection Form – Section 3: Blood Sample Collection**

**Cluster ID |__|__|__| Mother ID |__|__|**

<table>
<tr><td>11. Estimate of Blood Volume Collected in Tube</td><td># ml</td><td>|__|.|__|ml</td></tr>
<tr><td>12. Blood Spots Samples Obtained?</td><td>1. all (6 spots)<br>2. partial (< 6 spots)<br>3. none (0 spots)</td><td>|__|</td></tr>
<tr><td>13. Number of FULLY SaturatedSpots Obtained</td><td># Spots</td><td>|__| Spots</td></tr>
<tr><td>13b. Anemia Measurement</td><td>###.#</td><td>|__||__|.|__| g/dl</td></tr>
<tr><td>14. Why Was Sample Not Obtained?<br><u>Note</u>: If Q10 or Q 12 are 2 or 3, then Q14 is asked. If Q10 or 12 is 1, then Q 14 is skipped.</td><td>1. Parent not available<br>2. Parent refused<br>3. Child not available<br>4. Volume insufficient<br>5. Vein visibility<br>6. Other: Specify</td><td>|__|</td></tr>
<tr><td>15. Blood Collection: End Time</td><td>24-hour scale<br>Hours : Minutes</td><td>|__|__|:|__|__|</td></tr>
<tr><td>16. Cold Chain Start Time<br><u>Note</u>: This is the time when sample has been placed in cold box (for tube) or room temperature box (for spots).</td><td>24-hour scale<br>Hours : Minutes</td><td>|__|__|:|__|__|</td></tr>
<tr><td>16b. Cooler box Temperature Data Logger ID</td><td>####</td><td>|__|__|__|__|</td></tr>
</table>

<table>
<tr><td>17. Centrifugation Start Time<br><u>Note</u>: Only for child venous blood samples (types B and X).</td><td>24-hour scale<br>Hours : Minutes</td><td>|__|__|:|__|__|</td></tr>
<tr><td>18. Duration of Centrifugation<br><u>Note</u>: Only for child venous blood samples (type B).</td><td>Minutes<br>##</td><td>|__|__|</td></tr>
<tr><td>19. Plasma Sample Hemolyzed?<br>(Plasma is red, not yellow)<br><u>Note</u>: Only for child venous blood samples (type B).</td><td>Y/N</td><td>|__|</td></tr>
<tr><td>20. PBMC Processing: Start Time</td><td>24-hour scale<br>Hours : Minutes</td><td>|__|__|:|__|__|</td></tr>
<tr><td>21. PBMC Processing: Number of Cells</td><td></td><td></td></tr>
</table>

#### EE Sample Collection Form – Section 3: Blood Sample Collection

**Cluster ID |__|__|__| Mother ID |__|__|**

| Samples: | | | | | | | |
|---|---|---|---|---|---|---|---|
| 22. Sample ID<br>Note:<br>IDs for all aliquots of blood samples expected in a HH are automatically generated when cluster ID and mother ID are entered in Q1. | 23. Random ID<br>Note:<br>Random ID that is linked to the specific sample ID is retrieved from barcode ID database and appears automatically. | 24. Sample Type<br>Note:<br>Field staff selects sample type from dropdown menu. | 25.Aliquots | 26. Samples H & B -80ºC Freezer Storage: Start Time | 27. PBMC -80ºC Freezer Storage: Start Time | 28. PBMC Liquid Nitrogen Storage: Start Date | 29. PBMC Liquid Nitrogen Storage: Start Time |
| Cluster (###) + Mother ID (##) + Baseline (B) + Eligible Child (#) + Sample Type + Aliquot (#) | | H (Whole)<br>B (Plasma)<br>X (PBMC)<br>P (Filter) | | 24-hour scale<br>Hours : Minutes | 24-hour scale<br>Hours : Minutes | DD/MM/YYYY | 24-hour scale<br>Hours : Minutes |
| \|__\|__\|__\|__\|__\|E\|__\|H01 | \|__\|__\|__\|__\|__\|__\| | H | No Aliquot<br>Partial Aliquot<br>Full Aliquot | | | | |
| \|__\|__\|__\|__\|__\|E\|__\|H02 | \|__\|__\|__\|__\|__\|__\| | H | No Aliquot<br>Partial Aliquot<br>Full Aliquot | | | | |
| \|__\|__\|__\|__\|__\|E\|__\|B01 | \|__\|__\|__\|__\|__\|__\| | B | No Aliquot<br>Partial Aliquot<br>Full Aliquot | | | | |
| \|__\|__\|__\|__\|__\|E\|__\|B02 | \|__\|__\|__\|__\|__\|__\| | B | No Aliquot<br>Partial Aliquot<br>Full Aliquot | \|__\|__\|:\|__\|__\| | | | |
| \|__\|__\|__\|__\|__\|E\|__\|B03 | \|__\|__\|__\|__\|__\|__\| | B | No Aliquot<br>Partial Aliquot<br>Full Aliquot | | | | |
| \|__\|__\|__\|__\|__\|E\|__\|B04 | \|__\|__\|__\|__\|__\|__\| | B | No Aliquot<br>Partial Aliquot<br>Full Aliquot | | | | |
| \|__\|__\|__\|__\|__\|E\|__\|B05<br>***(Optional Aliquot)*** | \|__\|__\|__\|__\|__\|__\| | B | No Aliquot<br>Partial Aliquot<br>Full Aliquot | | | | |
| \|__\|__\|__\|__\|__\|E\|__\|B06<br>***(Optional Aliquot)*** | \|__\|__\|__\|__\|__\|__\| | B | No Aliquot<br>Partial Aliquot<br>Full Aliquot | | | | |
| \|__\|__\|__\|__\|__\|E\|__\|B07<br>***(Optional Aliquot)*** | \|__\|__\|__\|__\|__\|__\| | B | No Aliquot<br>Partial Aliquot<br>Full Aliquot | | | | |
| \|__\|__\|__\|__\|__\|E\|__\|X01 | \|__\|__\|__\|__\|__\|__\| | X | No Aliquot<br>Partial Aliquot<br>Full Aliquot | | \|__\|__\|:\|__\|__\| | \|__\|__\| / \|__\|__\| / \|__\|__\|__\|__\| | \|__\|__\|:\|__\|__\| |
| \|__\|__\|__\|__\|__\|E\|__\|X02 | \|__\|__\|__\|__\|__\|__\| | X | No Aliquot<br>Partial Aliquot<br>Full Aliquot | | | | |
| \|__\|__\|__\|__\|__\|E\|__\|P01 | \|__\|__\|__\|__\|__\|__\| | P | No Aliquot<br>Partial Aliquot<br>Full Aliquot | | | | |

**Cluster ID |__|__|__| Mother ID |__|__|**

#### EE Sample Collection Form – Section 4: Stool Sample Collection

| | | |
|---|---|---|
| 1. Cluster ID and Mother ID | ### and ## | \|__\|__\|__\| **and** \|__\|__\| |
| 2. ID of MT/FRA/FRO | ## | \|__\|__\| |
| 3. Name of MT/FRA/FRO (choose 1): | 4. Dipali Rani Das<br>5. SaniaAkter Dolly<br>6. SyedaFatemaKhatun<br>7.Md. AltafurRahman<br>8. AmalSarker<br>9. Nasrin Sultana<br>10. Md. NurulAlam<br>11. Fatema Begum Sumi<br>12. SyedaLuthfaFamida<br>13. Shahjahan Ali | |
| 4. Date of Sample Collection (when FRA picks up tube) | DD/MM/YYYY | \|__\|__\| / \|__\|__\| / \|__\|__\|__\|__\| |
| 5. Number of Eligible Children | # of Eligible Children | \|__\| |
| 6. Child Full Name | | |

| Samples: | | | |
|---|---|---|---|
| 7. Sample ID<br><br>Cluster (###) + Mother ID (##) + Baseline (B) + Eligible Child (#) + Sample Type + Aliquot (#) | 8. Random ID<br><br>Note:<br>Random ID that is linked to the specific sample ID is retrieved from barcode ID database and appears automatically. | 9. Sample type<br><br>Note:<br>Field staff selects sample type from dropdown menu. | 10. Aliquots<br><br>Note:<br>If sample has been fully collected, next question is skipped. |
| \|__\|__\|__\|__\|__\|E\|__\|S01 | \|__\|__\|__\|__\|__\|__\| | S | No Aliquot<br>Partial Aliquot<br>Full Aliquot |
| \|__\|__\|__\|__\|__\|E\|__\|S02 | \|__\|__\|__\|__\|__\|__\| | S | No Aliquot<br>Partial Aliquot<br>Full Aliquot |
| \|__\|__\|__\|__\|__\|E\|__\|S03 | \|__\|__\|__\|__\|__\|__\| | S | No Aliquot<br>Partial Aliquot<br>Full Aliquot |
| \|__\|__\|__\|__\|__\|E\|__\|S04 | \|__\|__\|__\|__\|__\|__\| | S | No Aliquot<br>Partial Aliquot<br>Full Aliquot |
| \|__\|__\|__\|__\|__\|E\|__\|S05 | \|__\|__\|__\|__\|__\|__\| | S | No Aliquot<br>Partial Aliquot<br>Full Aliquot |

#### EE Sample Collection Form – Section 4: Stool Sample Collection

**Cluster ID |__|__|__| Mother ID |__|__|**

| | | |
|---|---|---|
| 11. Why Was It Not Fully Collected?<br><br>Note: In Q10, If Partial Aliquot collected, Q11 is asked. In Q10, if No Aliquot collected, then Q11 is asked and skip to end. | 1. Parents not available<br>2. Parents refused<br>3. Subject not available<br>4. Child did not defecate<br>5. Defecation volume insufficient<br>6. Other: Specify | \|__\| |
| 12. Date of Defecation<br><br>Note: This is only applicable for stool samples as stool collection containers are left in households the day before sample collection. | DD/MM/YYYY | \|__\|__\| / \|__\|__\| / \|__\|__\|__\|__\| |
| 13. Time of Defecation<br><br>Note: Exact or approximate time is entered. | 24-hour scale<br>Hours: Minutes | \|__\|__\|:\|__\|__\| |
| 14. FRA Present During Defecation?<br><br>Note: FRA presence or absence during defecation is indicated. | 1. Yes<br>2. No | \|__\|__\| |
| 15. Cold Chain Start Time<br><br>Note: This is the time when sample has been placed in cold box. | 24-hour scale<br>Hours : Minutes | \|__\|__\|:\|__\|__\| |
| 16. OBSERVE: Consistency of Collected Stool Sample | 1. Normal stool<br>(Formed, soft, semi-solid, moist)<br>2. Diarrheal stool<br>(Unformed, watery)<br>3. Constipated stool<br>(Formed, hard, dry) | \|__\| |
| 17. OBSERVE: Color of Collected Stool Sample | 1. Yellow<br>2. Brown<br>3. Black<br>4. Green<br>5. White<br>6. Red<br>7. Other: Specify | \|__\| |

**Cluster ID |__|__|__| Mother ID |__|__|**

#### EE Sample Collection Form – Section 4: Stool Sample Collection

| | | |
|---|---|---|
| 18. OBSERVE: Any Abnormal Characteristics of Collected Stool Sample | 1. Yes, Mucus<br>2. Yes, Blood<br>3. Yes, Worms<br>4. Yes, Other: Specify<br>5. No | \|__\| |
| 19. ASK: "Does [CHILD NAME] currently have diarrhea?"<br>(Diarrhea = 3 or more unformed stools in a 24-hour period)<br>Note: If Q19 is 1, then ask next question and skip to end. If Q19 is 2 or 99, then skip to Q21. | 1. Yes<br>2. No<br>99. Don't know | \|__\|__\| |
| 20. ASK: How many consecutive days has this child had diarrhea?"<br>Note: Exact or approximate number of days is entered. | ## Days | \|__\|__\| Days |
| 21. ASK: "How long ago did [CHILD NAME] have diarrhea?"<br>Note: Exact or approximate number of days is entered. If Q21 answer is >0 and NOT 88 or 99, then Q22 is asked. If Q21 answer is 0, 88, or 99, then skip to end. | ## Days<br>88. Never<br>99. Don't know | \|__\|__\| Days |
| 22. ASK: "During that specific episode of diarrhea, how many consecutive days did it last?"<br>Note: Exact or approximate number of days is entered. | ## Days<br>99. Don't know | \|__\|__\| Days |
| 23. ASK: Where was stool sample collected from? | 1. Diaper provided<br>2. Katha (blanket/cloth)<br>3. Bed sheet<br>4. Potty<br>5. Other: Specify<br>77. Not applicable | \|__\| |
| 24. ASK: Was the stool in contact with urine (in the diaper/potty/katha/etc.)? | 1. Yes<br>2. No<br>99. Don't know | \|__\|__\| |
| 25. Cooler box Temperature Data Logger ID | #### | \|__\|__\|__\|__\| |

**Cluster ID |__|__|__| Mother ID |__|__|**

#### EE Sample Collection Form – Section 5: Urine Sample Collection

| | | |
|---|---|---|
| 1. Cluster ID and Mother ID | ### and ## | \|__\|__\|__\| **and** \|__\|__\| |
| 2. ID of MT/FRA/FRO | ## | \|__\|__\| |
| 3. Name of MT/FRA/FRO (choose 1): | 4. Dipali Rani Das<br>5. SaniaAkter Dolly<br>6. SyedaFatemaKhatun<br>7.Md. AltafurRahman<br>8. AmalSarker<br>9. Nasrin Sultana<br>10. Md. NurulAIam<br>11. Fatema Begum Sumi<br>12. SyedaLuthfaFamida<br>13. Shahjahan Ali<br>14. Md. Shariful Islam | |
| 4. Date of Sample Collection | DD/MM/YYYY | \|__\|__\| / \|__\|__\| / \|__\|__\|__\|__\| |
| 5. Number of Eligible Children | # of Eligible Children | \|__\| |
| 6. Child Full Name | | |
| 7. Consent<br><br>Note: If yes, then skip next question. | Y/N | \|__\| |
| 8. Why Was Consent Not Given?<br><br>Note: If Q7 is no, then answer this question and skip to end. | 1. Household head/primary caregiver not available<br>2. Household head/ primary caregiver refused<br>3. Subject not available<br>4. Genital skin disease<br>5. Other: Specify | \|__\| |
| 9. Volume of Lactulose-MannitolFed to Child | Volume (ml) = average weight X 2 (maximum of 20 ml)<br>##.# | \|__\|__\|.\|__\| ml |
| 10. Lactulose-Mannitol Batch Number | ### | \|__\|__\|__\| |
| 11. Fasting Start Time | 24-hour scale<br>Hours : Minutes | \|__\|__\|:\|__\|__\| |
| 12. Fasting End Time | 24-hour scale<br>Hours : Minutes | \|__\|__\|:\|__\|__\| |
| 13. Urine Collection Start Time | 24-hour scale<br>Hours : Minutes | \|__\|__\|:\|__\|__\| |

**Cluster ID |__|__|__| Mother ID |__|__|**

#### EE Sample Collection Form – Section 5: Urine Sample Collection

| 14. Urination episode<br><br>Note:<br>Urine episode number is automatically generated starting from 1. Q14 and the next questions (Q15, Q16, Q17, Q18) will loop continuously until end of urine collection time. | 15. Time of Collection | 16. Volume of Each Urination | 17. Estimated Volume Lost | 18. Stool in Bag? |
|---|---|---|---|---|
| **Urination Episode #** | 24-hour scale<br>Hours : Minutes | Volume (ml)<br>### | 1. None<br>2. < ½<br>3. ½<br>4. > ½<br>5. All | Y/N |
| **1** | \|__\|__\|:\|__\|__\| | \|__\|__\|__\|ml | \|__\| | \|__\| |
| **2** | \|__\|__\|:\|__\|__\| | \|__\|__\|__\| ml | \|__\| | \|__\| |
| **3** | \|__\|__\|:\|__\|__\| | \|__\|__\|__\| ml | \|__\| | \|__\| |
| **4** | \|__\|__\|:\|__\|__\| | \|__\|__\|__\| ml | \|__\| | \|__\| |
| **5** | \|__\|__\|:\|__\|__\| | \|__\|__\|__\| ml | \|__\| | \|__\| |
| **6** | \|__\|__\|:\|__\|__\| | \|__\|__\|__\| ml | \|__\| | \|__\| |
| **7** | \|__\|__\|:\|__\|__\| | \|__\|__\|__\| ml | \|__\| | \|__\| |
| **8** | \|__\|__\|:\|__\|__\| | \|__\|__\|__\| ml | \|__\| | \|__\| |
| **9** | \|__\|__\|:\|__\|__\| | \|__\|__\|__\| ml | \|__\| | \|__\| |
| **10** | \|__\|__\|:\|__\|__\| | \|__\|__\|__\| ml | \|__\| | \|__\| |

**Cluster ID |__|__|__| Mother ID |__|__|**

#### EE Sample Collection Form – Section 5: Urine Sample Collection

| | | |
|---|---|---|
| 19. Urine Collection End Time for 2-Hour Urine Sample | 24-hour scale<br>Hours : Minutes | \|__\|__\|:\|__\|__\| |
| 20. Total Urine Volume After 2 Hours | Volume (ml)<br>### | \|__\|__\|__\| ml |
| 21. Total Number of Thimerasol Drops Added After 2 Hours | Number of drops<br>## | \|__\|__\| Drops |
| 22. Number of Loose Stools During 2-Hour Collection | # loose stools | \|__\| Loose Stools |
| 23. Total Volume of All Aliquots From 2-Hour Urine Collection | Volume (ml)<br>### | \|__\|__\| ml |
| 24. Urine Collection End Time for 5-Hour Urine Sample | 24-hour scale<br>Hours: Minutes | \|__\|__\|:\|__\|__\| |
| 25. Total Urine Volume After 5 Hours | Volume (ml)<br>### | \|__\|__\|__\| ml |
| 26. Total Number of Thimerasol Drops Added After 5 Hours | Number of drops<br>## | \|__\|__\| Drops |
| 27.Number of Loose Stools During 5-Hour Collection | # of loose stools | \|__\|Loose Stools |
| 28. Total Volume of All Aliquots From 5-Hour Urine Collection | Volume (ml)<br>### | \|__\|__\| ml |
| 29. Did Child Eat During First 1-Hour Fasting Period?<br>(Before LM dose) | Y/N | \|__\| |
| 30. Did Child Eat During Last 0.5-Hour Fasting Period?<br>(Immediately after LM dose) | Y/N | \|__\| |

### EE Sample Collection Form – Section 5: Urine Sample Collection

**Cluster ID |__|__|__| Mother ID |__|__|**

| Samples: | | | | | |
|---|---|---|---|---|---|
| 31. Sample ID<br>Note:<br>IDs for all aliquots of urine samples expected in a HH are automatically generated when cluster ID and mother ID are entered in Q1. **(Aliquots 01-06 after 2 hrs urine collection. Aliquots 07-12 after 5 hrs urine collection)** | 32. Random ID<br>Note:<br>Random ID that is linked to the specific sample ID is retrieved from barcode ID database and appears automatically | 33. Sample Type<br>Note:<br>Field staff selects sample type from dropdown menu. | 34. Aliquots<br>Note:<br>If sample has been fully collected, next question is skipped. For sample type U, full aliquot = 2 ml per cryovial. | 35. Why was it not fully collected?<br>Note:<br>If Q34 is No Aliquot or Partial Aliquot, then Q 35 is asked.<br>1. Child did not urinate<br>2. Urination volume insufficient<br>3. Leakage<br>4. Partial: Parent Refused<br>5. Other: Specify | 36. Cold Chain Start Time<br>Note:<br>This is the time when sample has been placed in cold box.<br>24-hour scale<br>Hours : Minutes |
| \|__\|__\|__\|__\|__\|E\|__\|U01 | \|__\|__\|__\|__\|__\|__\| | U | No Aliquot<br>Partial Aliquot<br>Full Aliquot | \|__\| 2 Hours | \|__\|__\|:\|__\|__\| |
| \|__\|__\|__\|__\|__\|E\|__\|U02 | \|__\|__\|__\|__\|__\|__\| | U | No Aliquot<br>Partial Aliquot<br>Full Aliquot | | |
| \|__\|__\|__\|__\|__\|E\|__\|U03 | \|__\|__\|__\|__\|__\|__\| | U | No Aliquot<br>Partial Aliquot<br>Full Aliquot | | |
| \|__\|__\|__\|__\|__\|E\|__\|U04 | \|__\|__\|__\|__\|__\|__\| | U | No Aliquot<br>Partial Aliquot<br>Full Aliquot | | |
| \|__\|__\|__\|__\|__\|E\|__\|U05 | \|__\|__\|__\|__\|__\|__\| | U | No Aliquot<br>Partial Aliquot<br>Full Aliquot | | |
| \|__\|__\|__\|__\|__\|E\|__\|U06 | \|__\|__\|__\|__\|__\|__\| | U | No Aliquot<br>Partial Aliquot<br>Full Aliquot | | |
| \|__\|__\|__\|__\|__\|E\|__\|U07 | \|__\|__\|__\|__\|__\|__\| | U | No Aliquot<br>Partial Aliquot<br>Full Aliquot | \|__\| 5 Hours | \|__\|__\|:\|__\|__\| |
| \|__\|__\|__\|__\|__\|E\|__\|U08 | \|__\|__\|__\|__\|__\|__\| | U | No Aliquot<br>Partial Aliquot<br>Full Aliquot | | |
| \|__\|__\|__\|__\|__\|E\|__\|U09 | \|__\|__\|__\|__\|__\|__\| | U | No Aliquot<br>Partial Aliquot<br>Full Aliquot | | |
| \|__\|__\|__\|__\|__\|E\|__\|U10 | \|__\|__\|__\|__\|__\|__\| | U | No Aliquot<br>Partial Aliquot<br>Full Aliquot | | |
| \|__\|__\|__\|__\|__\|E\|__\|U11 | \|__\|__\|__\|__\|__\|__\| | U | No Aliquot<br>Partial Aliquot<br>Full Aliquot | | |
| \|__\|__\|__\|__\|__\|E\|__\|U12 | \|__\|__\|__\|__\|__\|__\| | U | No Aliquot<br>Partial Aliquot<br>Full Aliquot | | |

| | | |
|---|---|---|
| 37. Why was urine collection stopped before 5 hours?<br><br>Note: Skip if both 2 and 5 hour urine collection completed. | 1. Baby crying a lot<br>2. Baby developed a rash at U-bag attachment site<br>3. Other parent came home and refused<br>4. Another family member arrived and encouraged mother to refuse<br>5. Mother and child left to visit another place<br>6. Other: Specify<br>77. Not applicable | \|__\| |
| 38. Cooler box Temperature Data Logger ID | #### | \|__\|__\|__\|__\| |

**Cluster ID |__|__|__| Mother ID |__|__|**

#### EE Sample Collection Form – Section 5A: Pre-LM Urine Sample Collection

| | | |
|---|---|---|
| 39. Cluster ID and Mother ID | ### and ## | \|__\|__\|__\| **and** \|__\|__\| |
| 40. Date of Sample Collection | DD/MM/YYYY | \|__\|__\| / \|__\|__\| / \|__\|__\|__\|__\| |
| 41. **Pre-LM** Urine Collection Start Time | 24-hour scale<br>Hours : Minutes | \|__\|__\|:\|__\|__\| |

| 42. **Pre-LM**Urination episode<br><br>Note:<br>Urine episode number is automatically generated starting from 1. Q42 and the next questions (Q43, Q44, Q45, Q46) will loop continuously until end of urine collection time. | 43. Time of Collection | 44. Volume of Each Urination | 45. Estimated Volume Lost | 46. Stool in Bag? |
|---|---|---|---|---|
| **Urination Episode #** | 24-hour scale<br>Hours : Minutes | Volume (ml)<br>### | 1. None<br>2. < ½<br>3. ½<br>4. > ½<br>5. All | Y/N |
| **1** | \|__\|__\|:\|__\|__\| | \|__\|__\|__\| ml | \|__\| | \|__\| |
| **2** | \|__\|__\|:\|__\|__\| | \|__\|__\|__\| ml | \|__\| | \|__\| |
| **3** | \|__\|__\|:\|__\|__\| | \|__\|__\|__\| ml | \|__\| | \|__\| |
| **4** | \|__\|__\|:\|__\|__\| | \|__\|__\|__\| ml | \|__\| | \|__\| |
| **5** | \|__\|__\|:\|__\|__\| | \|__\|__\|__\| ml | \|__\| | \|__\| |

#### EE Sample Collection Form – Section 5A: Pre-LM Urine Sample Collection

**Cluster ID |__|__|__| Mother ID |__|__|**

| | | |
|---|---|---|
| 47. **Pre-LM** Urine Collection End Time | 24-hour scale<br>Hours : Minutes | \|__\|__\|:\|__\|__\| |
| 48. Total Urine Volume of **Pre-LM** Urine | Volume (ml)<br>### | \|__\|__\|__\| ml |
| 49. Total Number of Thimerasol Drops Added to **Pre-LM** Urine | Number of drops<br>## | \|__\|__\| Drops |
| 50. Number of Loose Stools During **Pre-LM** Urine Collection | # loose stools | \|__\| Loose Stools |
| 51. Total Volume of All Aliquots From **Pre-LM** Urine Collection | Volume (ml)<br>### | \|__\|__\| ml |

Samples:

| 52. Sample ID<br>Note:<br>IDs for all aliquots of urine samples expected in a HH are automatically generated when cluster ID and mother ID are entered in Q1.<br>**(Aliquots 13-18 after Pre-LM urine collection)** | 53. Random ID<br>Note:<br>Random ID that is linked to the specific sample ID is retrieved from barcode ID database and appears automatically. | 54. Sample Type<br>Note:<br>Field staff selects sample type from dropdown menu. | 55. Aliquots<br>Note:<br>If sample has been fully collected, next question is skipped.<br>For sample type U, full aliquot = 2 ml per cryovial. | 56. Why was Pre-LM urine not fully collected?<br>Note:<br>If Q55 is No Aliquot or Partial Aliquot, then Q56 is asked.<br>1. Child did not urinate<br>2. Urination volume insufficient<br>3. Leakage<br>4. Partial: Parent Refused<br>5. Other: Specify | 57. Cold Chain Start Time<br>Note:<br>This is the time when sample has been placed in cold box.<br>24-hour scale<br>Hours : Minutes |
|---|---|---|---|---|---|
| \|__\|__\|__\|__\|__\|E\|__\|U13 | \|__\|__\|__\|__\|__\|__\| | U | No Aliquot<br>Partial Aliquot<br>Full Aliquot | \|__\| Pre-LM Urine | \|__\|__\|:\|__\|__\| |
| \|__\|__\|__\|__\|__\|E\|__\|U14 | \|__\|__\|__\|__\|__\|__\| | U | No Aliquot<br>Partial Aliquot<br>Full Aliquot | | |
| \|__\|__\|__\|__\|__\|E\|__\|U15 | \|__\|__\|__\|__\|__\|__\| | U | No Aliquot<br>Partial Aliquot<br>Full Aliquot | | |
| \|__\|__\|__\|__\|__\|E\|__\|U16 | \|__\|__\|__\|__\|__\|__\| | U | No Aliquot<br>Partial Aliquot<br>Full Aliquot | | |
| \|__\|__\|__\|__\|__\|E\|__\|U17 | \|__\|__\|__\|__\|__\|__\| | U | No Aliquot<br>Partial Aliquot<br>Full Aliquot | | |
| \|__\|__\|__\|__\|__\|E\|__\|U18 | \|__\|__\|__\|__\|__\|__\| | U | No Aliquot<br>Partial Aliquot<br>Full Aliquot | | |

PROTOCOL
Biomedical
Non-Exempt
Berkeley

Protocol # 2011-09-3652
Date Printed: 07/31/2018

---

| | |
|---|---|
| **Protocol Title:** | Measuring the benefits of sanitation, water quality, handwashing and nutrition interventions for improving health and development in rural Bangladesh |
| **Protocol Status:** | APPROVED |
| **Date Submitted:** | 10/26/2017 |
| **Approval Period:** | **10/30/2017-11/03/2018** |
| **Important Note:** | **This Print View may not reflect all comments and contingencies for approval. Please check the comments section of the online protocol. Questions that appear to not have been answered may not have been required for this submission. Please see the system application for more details.** |

*** * * Attached Document * * ***

| Document Name | Created Date |
|---|---|
| WASH Benefits Bangladesh Parasites Recruitment script_v1.pdf | 10/26/2017 |

**WASH Benefits Bangladesh Endline Parasites Assessment**
**Recruitment Script**

Hello (Assalamualaikum/Nomoshkar). My name is ________ and I work with the ICDDR,B (Cholera Hospital) in Dhaka. We are interested in conducting research to learn how to prevent parasite infections. We are inviting compounds already enrolled in the WASH Benefits Study to participate. If you agree to participate, we would like to ask two of your children under five years and one child 5-12 years old to provide a stool sample as part of this research. Are you interested in participating in this study?

If YES, Proceed with informed consent process.

If NO, Ok, thank you for your time.

---------------------------------------------------------------------------------------

**Protocol Title:** Measuring the benefits of sanitation, water quality, handwashing and nutrition interventions for improving health and development in rural Bangladesh

**Protocol Status:** APPROVED

**Date Submitted:** 10/26/2017

**Approval Period:** 10/30/2017-11/03/2018

**Important Note:** This Print View may not reflect all comments and contingencies for approval. Please check the comments section of the online protocol.
Questions that appear to not have been answered may not have been required for this submission. Please see the system application for more details.

*** * * Attached Document * * ***

| Document Name | Created Date |
| --- | --- |
| Appendix 1i-2 Parasite Assent Form.pdf | 10/26/2017 |

Appendix 1i-2: English assent for parasitic assessment of children aged 7-12 years at endline in the same compound

Protocol Title: Effect of hand washing, water treatment, sanitation and nutritional supplement interventions on child health and development in rural Bangladesh

Principal Investigator's Name: Dr. Leanne Unicomb

Hello/Assalamualaikum/Namaste. My name is _______________, and I work with the ICDDR,B (Cholera Hospital) in Dhaka. We are conducting research on health and worm infections. We are conducting this research with scientists at the University of California, Berkeley in the United States. You are being invited to participate in this study because your bari is already participating in the study.

**What is a research study?**

A research study is when people like me collect a lot of information about a certain thing to find out more about it.

This letter tells you about the study so you can decide if you want to be in it. Before you decide, you can talk about it with your parents or anyone else you like. If you have any questions about the research, just ask me.

**Why are we doing this study?**

The purpose of this study is to learn how certain hygiene, sanitation, and nutrition behaviors and practices affect health and worm infections in children. Worm infections are common in rural Bangladesh, especially in children.

**Why are we talking to you about this study?**

We are inviting thousands of other children your age in Bangladesh to participate in this study. We're inviting you to be in the study because your bari is already participating in the study.

**What will happen if you are in this study?**
If you agree to be in this study and your parents say it's okay, we may ask you to:

- Let us take a blood sample from your finger.
  We will also ask you to let us prick your finger to collect blood. You will experience a momentary pinch and about seven drops of blood will be collected through our

trained field staff. We will collect seven drops of blood to test for infections. This part will take about 10 minutes.

If you don't want to give a blood sample, you don't have to.

- Collect a poop sample.
  We will give you a kit to collect stool with today and we will come back tomorrow. We will ask you to poop on a sheet of plastic and use a plastic scoop to collect a small amount of fresh poop from the top of the pile into a container. Your parents can help you if needed. The field person will collect this container when they return. This part will take about 10 minutes.

  If you don't want to give a poop sample, you don't have to.

- Offer you deworming medication.
  We will offer you deworming medication. This part will take about 5 minutes

  If you don't want to take deworming medication, you don't have to.

Total time: Altogether, the whole study will take about 25 minutes of your time.

Study place: We will collect these samples at your bari.

**Will you get healthier if you are in the study?**
We will offer you deworming medication in this study. If you have certain worm infections, the medication may help make you feel better by treating your infection.

**Will any part of the study be uncomfortable or hurt?**

- Blood drawing: Getting your finger pricked can hurt for a few seconds from the needle stick going in, like when you get a shot at the doctor's office. Afterwards, you might get a little bruise. Sometimes an infection can develop there, but that hardly ever happens.

- Deworming medication: The medication also may make you feel sick for a short time. There is a very small chance that you will have stomach pain, diarrhea, nausea, vomiting, dizziness, itchiness or skin rash.

**Will you get paid for being in the study?**

You will not be paid for being in this study.

**Do you have to be in the study?**

No, you don't! Research is something you do only if you want to. Nothing bad will happen if you don't want to be in the study. Just tell us. And remember, you can always change your mind later if you don't want to be in the study any more.

**Do you have any questions?**

You can ask questions about this study at any time, now or later. You can talk to me, or your parents, or someone else if you like.

You can contact Masud Parvez: phone: 01817541872, Center of Communicable Diseases, ICDDR,B, Mohakhali, Dhaka 1212.

****************************************

**ASSENT OF CHILD (7-12 years old)**

If you decide to participate, and your parents agree, we will give you a copy of this form to keep. That way you can look at it later if you want to.

***Do you agree to be in this research study?***

_______________________________________ _______________
Child's Name/Signature (*printed or written by child*)* Date
*or by field worker if child cannot write*

_______________________________________ _______________
Signature of Investigator/Person Obtaining Assent Date

****************************************

**If verbal assent only is being obtained:*

Investigator or Person Conducting Assent Discussion: Initial here if child cannot sign, to document that child received this information and gave assent verbally: ______

PROTOCOL
Biomedical
Non-Exempt
Berkeley

Protocol # 2011-09-3652
Date Printed: 07/31/2018

---------------------------------------------------------------------------------
---

**Protocol Title:** Measuring the benefits of sanitation, water quality, handwashing and nutrition interventions for improving health and development in rural Bangladesh

**Protocol Status:** APPROVED

**Date Submitted:** 10/26/2017

**Approval Period:** **10/30/2017-11/03/2018**

**Important Note:** **This Print View may not reflect all comments and contingencies for approval. Please check the comments section of the online protocol.**
**Questions that appear to not have been answered may not have been required for this submission. Please see the system application for more details.**

*** * * Attached Document * * ***

| Document Name | Created Date |
| --- | --- |
| Spillover study qualitative instrument_v1.pdf | 10/26/2017 |

**WASH Benefits Spillover Study**
**In-Depth Interview Instrument for Spillover Compounds**

Date: ____________
Cluster ID: ___________
WASH Benefits Compound ID: ___________
Spillover Compound ID: ___________

Participant age (years): __________
Participant sex: M F

---

We would like to know more about where you learn about health. We are interested in learning more about your perceptions and opinions regarding water, sanitation, and hygiene.

**Social relationships** (name generator)
First, I would like to ask you about people you know in this village.

1. Who are the 4 non-relatives in this village whom you speak to the most?

2. In your free time, whose house in this village do you visit the most? When was the last time you saw this person?

3. Who from this village visits your house the most in his or her free time? When was the last time you saw this person?

4. If you needed to borrow kerosene or rice, to whom in your village would you go? When was the last time you saw this person?

5. Who in your village would come to you if he/she needed to borrow kerosene or rice? When was the last time you saw this person?

6. If you had to go to the hospital, is there anyone outside of your compound who you would ask to watch your children? Who? When was the last time you saw this person?

7. If you had to make a difficult personal decision, is there anyone outside of your compound and family who you would ask for advice? Who? When was the last time you saw this person? Whose advice are you most likely to follow?

8. Who comes to you for advice? When was the last time you saw this person?

**Environmental Contamination and Hygiene**
Now I would like to ask you about what you notice about contamination of the environment near your compound and in your village. Environmental contamination includes presence of human or animal feces, sewage, or garbage on the ground or near houses.

9. Can you describe to me the level of contamination <u>in your compound</u> these days?

    a. PROBE: Do any of your compound members defecate in the open? How often does this happen? Where does it happen?

10. Can you describe to me the level of contamination near your compound these days?
    a. PROBE: Do you see your neighbors defecating in the open? How often does this happen? Where does it happen?

11. Can you describe to me the level of contamination in your village these days?
    b. PROBE: Do you see people in your village defecating in the open? How often does this happen? Where does it happen?

12. Now I would like you to think about how things were two years ago.
    a. Two years ago, what was the level of contamination in your compound?
        a. PROBE: Did people defecate in the open more or less then?
    b. Two years ago, did you see your neighbors defecating in the open near your compound?
        a. PROBE: Did they defecate in the open more or less then?
    c. Two years ago, did you see people in your village defecating in the open?
        a. PROBE: Did people defecate in the open more or less then?

13. Do you think there has been a decrease in open defecation or increase in cleanliness and hygiene practices in your villages in the last 2 years? If yes, how did it happen? If no, what are reasons behind this?

14. What do you do to maintain hygiene in and near your compound?
    a. What motivates you to maintain hygiene in and near your compound?

15. What should people do to maintain hygiene in and near their compounds?

16. What practices do you see your neighbors do to maintain hygiene in and near your compound?

**Perception of health in the village**

Now I would like to ask you about health of children in your village.

17. Tell me about the last conversation you had about health.
    a. PROBE: Who did you talk to? What did you talk about? When was this?

18. When was the last time one of your children was sick? What did you do?
    a. PROBE: Is there anything you wanted to do but couldn’t because you didn’t have the time, resources, or ability to do it?
    b. PROBE: Did you do anything differently afterwards to prevent him or her from getting sick again?

19. Is there anything you regularly do to prevent your children from getting sick?

20. What are things you think people should do to keep their children from getting sick?

21. What are things that <u>you see other people</u> in your village do to prevent their children from getting sick?

22. These days, how healthy are children in your village?
    a. PROBE: Do many children get sick often? What kind of illnesses do they have?

23. Who has the healthiest children in your village?
    a. PROBE to determine whether this person is a WASH Benefits recipient without mentioning the WASH Benefits recipient's name

24. (If answer to previous question is not their own children) How healthy are your children compared to these children?

25. Why do you think their children are healthier than other children? (PROBE for preventive practices, such as handwashing, storing drinking water safely, etc. Also probe for whether these children are possibly from wealthier families.)

26. Have you adopted any practices that these people use to prevent their children from getting sick to protect your own children's health? Which practices?

27. Are there things you would like to do to protect your children's health that you cannot do for some reason (e.g., because you would need to purchase something you can't afford)? If so, what are these things?

28. What health practices do you <u>see</u> your neighbors doing?

29. What water, sanitation, and hygiene practices do you <u>see</u> your neighbors doing?

30. Do you ever see your neighbors doing health, sanitation or hygiene practices that you do not currently do but that you would like to do?
    a. If so, which practices are these?
    b. If so, why don't you currently practice these things?

**Knowledge of WASH Benefits**

31. Are there any activities going on in your village to promote children's health? If so, please tell me about them. (Do not mention WASH Benefits)

32. (If in the previous question they are do mention WASH Benefits:) Are you familiar with the Cholera Hospital activities in this village? What activities are they conducting?

33. Have you seen someone from Cholera Hospital come to this village to conduct these activities?
    a. Do you ever talk to someone from Cholera Hospital? What do you talk about?
    b. Who in your village do you talk to after the Cholera Hospital staff come to your village?

34. Do you know anyone who is participating in the Cholera Hospital activities?

    a. How well do you know them?
    b. How often do you talk to them?
    c. What do you think about their children's health?
    d. Do they ever teach you about the things they learn from the Cholera Hospital staff?

35. Have you changed any of your health and hygiene behaviors to be more similar to the behaviors of people participating in Cholera Hospital activities? (PROBE about handwashing at key times, reduced open defecation, safe disposal of child feces, safe water storage)
    a. What motivated you to change these behaviors?

**WASH Benefits Spillover Study**
**In-Depth Interview Instrument for WASH Benefits Compounds**

Date: ____________
Cluster ID: __________
WASH Benefits Compound ID: __________
Spillover Compound ID: __________

Participant age (years): _________
Participant sex: M F

---

We would like to know more about where you learn about health. We are interested in learning more about your perceptions and opinions regarding water, sanitation, and hygiene.

**Social relationships** (name generator)
First, I would like to ask you about people you know in this village.

36. Who are the 4 non-relatives in this village whom you speak to the most?

37. In your free time, whose house in this village do you visit the most? When was the last time you saw this person?

38. Who from this village visits your house the most in his or her free time? When was the last time you saw this person?

39. If you needed to borrow kerosene or rice, to whom in your village would you go? When was the last time you saw this person?

40. Who in your village would come to you if he/she needed to borrow kerosene or rice? When was the last time you saw this person?

41. If you had to go to the hospital, is there anyone outside of your compound who you would ask to watch your children? Who? When was the last time you saw this person?

42. If you had to make a difficult personal decision, is there anyone outside of your compound and family who you would ask for advice? Who? When was the last time you saw this person? Whose advice are you most likely to follow?

43. Who comes to you for advice? When was the last time you saw this person?

**Environmental Contamination and Hygiene**

Now I would like to ask you about what you notice about contamination of the environment near your compound and in your village. Environmental contamination includes presence of human or animal feces, sewage, or garbage on the ground or near houses.

44. Can you describe to me the level of contamination in your compound these days?
    c. PROBE: Do any of your compound members defecate in the open? How often does this happen? Where does it happen?

45. Can you describe to me the level of contamination near your compound these days?
    b. PROBE: Do you see your neighbors defecating in the open? How often does this happen? Where does it happen?

46. Can you describe to me the level of contamination in your village these days?
    d. PROBE: Do you see people in your village defecating in the open? How often does this happen? Where does it happen?

47. Now I would like you to think about how things were two years ago.
    d. Two years ago, what was the level of contamination in your compound?
        a. PROBE: Did people defecate in the open more or less then?
    e. Two years ago, did you see your neighbors defecating in the open near your compound?
        a. PROBE: Did they defecate in the open more or less then?
    f. Two years ago, did you see people in your village defecating in the open?
        a. PROBE: Did people defecate in the open more or less then?

48. Do you think there has been a decrease in open defecation or increase in cleanliness and hygiene practices in your villages in the last 2 years? If yes, how did it happen? If no, what are reasons behind this?

49. What do you do to maintain hygiene in and near your compound?
    a. What motivates you to maintain hygiene in and near your compound?

50. What should people do to maintain hygiene in and near their compounds?

51. What practices do you see your neighbors do to maintain hygiene in and near your compound?

**Perception of health in the village**

Now I would like to ask you about health of children in your village.

52. Tell me about the last conversation you had about health.

    a. PROBE: Who did you talk to? What did you talk about? When was this?

53. When was the last time one of your children was sick? What did you do?
    c. PROBE: Is there anything you wanted to do but couldn't because you didn't have the time, resources, or ability to do it?
    d. PROBE: Did you do anything differently afterwards to prevent him or her from getting sick again?

54. Is there anything <u>you regularly do</u> to prevent your children from getting sick?

55. What are things you think <u>people should do</u> to keep their children from getting sick?

56. What are things that <u>you see other people</u> in your village do to prevent their children from getting sick?

57. These days, how healthy are children in your village?
    a. PROBE: Do many children get sick often? What kind of illnesses do they have?

58. Who has the healthiest children in your village?
    a. PROBE to determine whether this person is a WASH Benefits recipient without mentioning the WASH Benefits recipient's name.

59. (If answer to previous question is not their own children) How healthy are your children compared to these children?

60. Why do you think their children are healthier than other children? (PROBE for preventive practices, such as handwashing, storing drinking water safely, etc. Also probe for whether these children are possibly from wealthier families.)

61. Have you adopted any practices that these people use to prevent their children from getting sick to protect your own children's health? Which practices?

62. Are there things you would like to do to protect your children's health that you cannot do for some reason (e.g., because you would need to purchase something you can't afford)? If so, what are these things?

63. What health practices do you <u>see</u> your neighbors doing?

64. What water, sanitation, and hygiene practices do you <u>see</u> your neighbors doing?

65. Do you ever see your neighbors doing health, sanitation or hygiene practices that you do not currently do but that you would like to do?
    a. If so, which practices are these?
    b. If so, why don't you currently practice these things?

**Experience in WASH Benefits**

66. Tell me about your experience participating in the Cholera Hospital activities in your village. What have you liked about it? What have you not liked about it?

67. Since you started participating in the Cholera Hospital activities, how has the health of your children been?

68. When someone from the Cholera Hospital comes to visit you, do your neighbors ever come listen to what you are discussing? Do they ever participate in the discussion?

69. Do you ever talk to your neighbors about what you learned from Cholera Hospital promoter?
    e. What do you talk about?
    f. How often do you talk to them?
    g. What do you think about their children's health?

70. Have you ever taught your neighbors about the things you learn from the Cholera Hospital staff?

71. Have your neighbors changed any of their health and hygiene behaviors to be more similar to the behaviors of people participating in Cholera Hospital activities? (PROBE about handwashing at key times, reduced open defecation, safe disposal of child feces, safe water storage)

PROTOCOL
Biomedical
Non-Exempt
Berkeley

Protocol # 2011-09-3652
Date Printed: 07/31/2018

---------------------------------------------------------------------------------
---

| | |
|---|---|
| **Protocol Title:** | Measuring the benefits of sanitation, water quality, handwashing and nutrition interventions for improving health and development in rural Bangladesh |
| **Protocol Status:** | APPROVED |
| **Date Submitted:** | 10/26/2017 |
| **Approval Period:** | **10/30/2017-11/03/2018** |
| **Important Note:** | **This Print View may not reflect all comments and contingencies for approval. Please check the comments section of the online protocol. Questions that appear to not have been answered may not have been required for this submission. Please see the system application for more details.** |

*** * * Attached Document * * ***

| Document Name | Created Date |
|---|---|
| Environmental Enteropathy Supplemental questionnaires.docx | 10/26/2017 |

OBSERVATION CODES FOR USE BEFORE, DURING AND AFTER EE BLOOD DRAW:

| **Reactivity Code** | **Observed Behavior** |
|---|---|
| **0-No reactivity** | No negative or positive facial affect and/or vocalizations |
| **1-Mild negative reactivity** | Infant is fussy, whiny, frowning, furrowed brow, crinkled nose, mouth slightly open or pressed lips |
| **2-Moderate negative reactivity** | Crying, wide squared mouth, eyes can be open or partially opened. |
| **3-High negative reactivity** | Screams, wails, eyes are partially closed or completely closed, may have wide or open mouth. Can include breath-holding, tears, face may change colors. Same facial expression as moderate, but with greater intensity. |
| **4 -Positive reactivity** | Smiling, bright eyes, raised cheeks, can be closed mouth or slightly parted, may be accompanied by positive vocalization |
| **5- Recovery** | Child's recovery from stressful event, self-soothing behaviors and extent comforted by caregiver |

TEMPERAMENT MEASURE (FEAR AND SOOTHABILITY).

**Fear:**

| **Items** |
|---|
| 49. *when this child tries to do something new (as exploring a new object), shows a little fear at the beginning* |
| 50. *when you take this child to a new place at first he looks afraid* |
| 51. *this child gets upset if there are changes in his activities inside or outside the house* |
| 53. *this child gets uncomfortable or cries the first time he is put to bed for a nap at a new place* |
| 55. *when this child listens to loud sounds (screams, radio, a honk of a car, dogs barking, etc.), he gets startled, screams or cries* |
| 56. *this child cries if he listens to other children cry* |
| 57. *when the child is being bathed and the temperature of the water of the bath tub is changed, this child gets startled* |
| 58. *if during the day this child gets upset with something, at night when he is sleeping he will cry, complain or wake up often* |
| 59. *when strangers caress him, this child cries* |
| 60. *the first reaction of this child is to get a hold of something strongly if he gets scared.* |
| Total |

**Soothability:**

| **Items** |
| --- |
| 101. *when this child is upset because his face is being washed, he calms down if he is talked to, singed to, or given something to play with* |
| 102. *when this child is upset because his diaper is being changed, he calms down if he is talked to, singed to, or given something to play with* |
| 103. *if this child is crying because he is hungry he calms down if he is held in arms or talked to* |
| 104. *if this child is upset or frightened by a sudden noise, he calms down on his own* |
| 105. *when this child is upset or frightened by the presence of a stranger, he calms down on his own* |
| 107. *when he is upset, this child calms down if he is rocked or held in arms* |
| 108. *when this child is upset because a toy is taken away from him, he calms down by himself* |
| 109. *this child stops crying if a cracker, bread or other food is given to him* |
| 111. *when this child is upset, he calms down if a toy is given to him* |
| 112. *when this child is angry, he calms down easily if he is taken for a ride* (or carried) |
| Total |

PROTOCOL
Biomedical
Non-Exempt
Berkeley

Protocol # 2011-09-3652
Date Printed: 07/31/2018

---------------------------------------------------------------------------------

| | |
|---|---|
| **Protocol Title:** | Measuring the benefits of sanitation, water quality, handwashing and nutrition interventions for improving health and development in rural Bangladesh |
| **Protocol Status:** | APPROVED |
| **Date Submitted:** | 10/26/2017 |
| **Approval Period:** | **10/30/2017-11/03/2018** |
| **Important Note:** | **This Print View may not reflect all comments and contingencies for approval. Please check the comments section of the online protocol. Questions that appear to not have been answered may not have been required for this submission. Please see the system application for more details.** |

*** * * Attached Document * * ***

| Document Name | Created Date |
|---|---|
| References.pdf | 10/26/2017 |

Adu-Afarwuah, S., A. Lartey, K. H. Brown, S. Zlotkin, A. Briend, K. G. Dewey 2008. Home fortification of complementary foods with micronutrient supplements is well accepted and has positive effects on infant iron status in Ghana. Am J Clin Nutr 87 (4): 929--938.

Adu-Afarwuah, S., A. Lartey, K. H. Brown, S. Zlotkin, A. Briend, K. G. Dewey 2007. Randomized comparison of 3 types of micronutrient supplements for home fortification of complementary foods in Ghana: effects on growth and motor development. Am J Clin Nutr 86 (2): 412--420.

Advisory Committee on Childhood Lead Poisoning Prevention, 2012. Low Level Lead Exposure Harms Children: A Renewed Call for Primary Prevention Centers for Disease Control and Prevention, Atlanta.

Albert MJ, Faruque ASG, Faruque SM, Sack RB, Mahalanabis D. Case-control study of enteropathogens associated with childhood diarrhea in Dhaka, Bangladesh. Journal of clinical microbiology. 1999;37(11):3458–64.

Alderman, H., J. Hoddinott, B. Kinsey 2006. Long term consequences of early childhood malnutrition. Oxf Econ Pap 58 (3): 450--474.

*Allen, L. & Gillespie, S. (2001) What Works? A Review of the Efficacy and Effectiveness of Nutrition Interventions,* Asian Development Bank, Manila, Phillipines.

Anderson RM, May RM (1992) Infectious Diseases of Humans: Dynamics and Control. London: Oxford University Press. 772 p.

Anderson RM, Medley GF (1985) Community control of helminth infections of man by mass and selective chemotherapy. Parasitology 90: 629–660.

Anoop, S., Saravanan, B., Joseph, A., Cherian, A. & Jacob, K. S. (2004) Maternal depression and low maternal intelligence as risk factors for malnutrition in children: a community based case-control study from South India. *Arch Dis Child*,**89**, 325-329.

Arnold, B. F., et al. (2013). "Cluster-randomised controlled trials of individual and combined water, sanitation, hygiene and nutritional interventions in rural Bangladesh and Kenya: the WASH Benefits study design and rationale." BMJ open**3**(8): e003476.

Arnold, B. F., J. M. Colford 2007. Treating water with chlorine at point-of-use to improve water quality and reduce child diarrhea in developing countries: a systematic review and meta-analysis. Am J Trop Med Hyg 76 (2): 354--364.

Arnold, B. F., R. S. Khush, P. Ramaswamy, A. G. London, P. Rajkumar, P. Ramaprabha, N. Durairaj, A. E. Hubbard, K. Balakrishnan, J. M. Colford 2010. Causal inference methods to study nonrandomized, preexisting development interventions. Proc Natl Acad Sci U S A 107 (52): 22605 - 22610.

Aubertin, D. and J. Smart (1940). *The Fauna of British India. Diptera volume 6: Calliphoridae*. London, Taylor and Francis.

Baddeley, A. (1992) Working memory. Science,**255**, 556-559.

Baddeley, A., Meeks Gardner, J. & Grantham-Mcgregor, S. (1995) Cross-cultural cognition: Developing tests for developing countries. *Applied Cognitive Psychology*,**9**, S173-S195.

Bansal S, Pourbohloul B, Meyers LA (2006) A Comparative Analysis of Influenza Vaccination Programs. PLoS Med 3: e387. doi:10.1371/journal.pmed.0030387.

Baqui AH, et al. (1991) Methodological issues in diarrhoeal diseases epidemiology: definition of diarrhoeal episodes. Int J Epidemiol 20: 1057-1063.

Barham, T. & Calimeris, L. (2008) Long-term effects of family planning and child health interventions on adolescent cognition: Evidence from Matlab in Bangladesh.Working paper. University of Colorado: Health and Society program.

Barreto, M. L., B. Genser, A. Strina, M. G. Teixeira, S. Cairncross, A. M. Assis, R. F. Rego, C. A. Teles, M. S. Prado, S. M. Matos, D. N. Santos, L. A. dos Santos 2007. Effect of city-wide sanitation programme on reduction in rate of childhood diarrhoea in northeast Brazil: assessment by two cohort studies. Lancet 370 (9599): 1622--1628.

Basáñez M-G, McCarthy JS, French MD, Yang G-J, Walker M, et al. (2012) A Research Agenda for Helminth Diseases of Humans: Modelling for Control and Elimination. PLoS Negl Trop Dis 6: e1548. doi:10.1371/journal.pntd.0001548.

Bellinger, D.C., Stiles, K.M., Needleman, H.L., 1992. Low-level lead exposure, intelligence and academic achievement: a long-term follow-up study. Pediatrics 90, 855-861.

Bergkvist, C., Kippler, M., Hamadani, J.D., Grander, M., Tofail, F., Berglund, M., Vahter, M., 2010. Assessment of early-life lead exposure in rural Bangladesh. Environmental research 110, 718-724.

Berkman, D. S., A. G. Lescano, R. H. Gilman, S. L. Lopez, M. M. Black 2002. Effects of stunting, diarrhoeal disease, and parasitic infection during infancy on cognition in late childhood: a follow-up study. Lancet 359 (9306): 564--571.

Bhutta, Z. A., T. Ahmed, R. E. Black, S. Cousens, K. Dewey, E. Giugliani, B. A. Haider, B. Kirkwood, S. S. Morris, H. P. S. Sachdev, M. Shekar, Maternal, C. U. S. Group 2008. What works? Interventions for maternal and child undernutrition and survival. Lancet 371 (9610): 417--440.

Black RE, Mersen MH, Abdul Alim AR, Yunus M. Incidence and severity of rotavirus and Escherichia coli diarrhoea in rural Bangladesh: implications for vaccine development. The Lancet. 1981;317(8212):141–3.

Black, R. E., L. H. Allen, Z. A. Bhutta, L. E. Caulfield, M. de Onis, M. Ezzati, C. Mathers, J. Rivera, Maternal, C. U. S. Group 2008. Maternal and child undernutrition: global and regional exposures and health consequences. Lancet 371 (9608): 243--260.

Blair, Clancy; Granger, Douglas A.; Kivlighan, Katie T.; Mills-Koonce, Roger; Willoughby, Michael; Greenberg, Mark T.; Hibel, Leah C.; Fortunato, Christine K. Family Life Project Investigators US

Developmental Psychology, Vol 44(4), Jul 2008, 1095-1109

Blair, Clancy. "Stress and the Development of Self-Regulation in Context." *Child Development Perspectives* 4.3 (2010): 181-188.

Boehm AB, Grant SB, Kim JH, Mowbray SL, McGee CD, Clark CD, Foley DM, Wellman DE. Decadal and Shorter Period Variability of Surf Zone Water Quality at Huntington Beach, California. Environmental Science & Technology. 2002;36(18):3885–92.

Boissiere, M., J. B. Knight, R. H. Sabot 1985. Earnings, schooling, ability, and cognitive skills. Am Econ Rev 75 (5): 1016--1030.

Boyce, A. B. Review of the Ages & Stages Questionnaires: A Parent-Completed, Child-Monitoring System, Second Edition (1995-1999) Mental Measurements Yearbook.

Bradley, R. & Corwyn, R. (2005) Caring for children around the world: A view from HOME. *International Journal of Behavioral Development*,**29**, 468-478.

Bradley, R., Corwyn, R., Mcadoo, H. & Garcia Coll, C. (2001) The home environments of children in the United States part I: Variations by age, ethnicity, and poverty status. *Child Development*,**72**, 1844-1867.

Bricker D, Squires J. Ages and Stages Questionnaires: A Parent Completed, Child Monitoring System, 2nd Ed. Baltimore, MD: Paul Brookes; 1999.

Bryan, J., S. Osendarp, D. Hughes, E. Calvaresi, K. Baghurst, J.-W. van Klinken 2004. Nutrients for cognitive development in school-aged children. Nutr Rev 62 (8): 295--306.

Campbell, D. I., M. Elia, P. G. Lunn 2003. Growth faltering in rural Gambian infants is associated with impaired small intestinal barrier function, leading to endotoxemia and systemic inflammation. J Nutr 133 (5): 1332--1338.

Chan MS, Guyatt HL, Bundy DA, Medley GF (1994) The development and validation of an age-structured model for the evaluation of disease control strategies for intestinal helminths. Parasitology 109: 389–396.

Checkley, W., G. Buckley, R. H. Gilman, A. M. Assis, R. L. Guerrant, S. S. Morris, K. Mølbak, P. Valentiner-Branth, C. F. Lanata, R. E. Black, C. Malnutrition, I. Network 2008. Multi-country analysis of the effects of diarrhoea on childhood stunting. Int J Epidemiol 37 (4): 816--830.

Checkley, W., L. D. Epstein, R. H. Gilman, L. Cabrera, R. E. Black 2003. Effects of acute diarrhea on linear growth in Peruvian children. Am J Epidemiol 157 (2): 166--175.

Clasen T, Edmondson P. Sodium dichloroisocyanurate (NaDCC) tablets as an alternative to sodium hypochlorite for the routine treatment of drinking water at the household level. Int J Hyg Environ Health. 2006;209(2):173-181.

Clasen T, Fabini D, Boisson S, Taneja J, Song J, Aichinger E, Bui A, Dadashi S, Schmidt W-P, Burt Z, Nelson KL. Making Sanitation Count: Developing and Testing a Device for Assessing Latrine Use in Low-Income Settings. Environmental Science & Technology. 2012;46:3295-303.

Clasen T, Saeed T, Boisson S, Edmondson P, Shipin O (2007). Household-based chlorination of drinking water using sodium dichloroisocyanurate (NaDCC) tablets: a randomized, controlled trial to assess microbiological effectiveness in Bangladesh. Am J. Trop. Med. & Hyg. 76(1):187-92

Clasen, T., W.-P. Schmidt, I. Roberts, S. Cairncross, T. Rabie 2006. Interventions to improve water quality for preventing diarrhoea. Cochrane Database Syst Rev 3: CD004794.

Clasen, T., W.-P. Schmidt, S. Cairncross, T. Rabie, I. Roberts 2007. Interventions to improve water quality for preventing diarrhoea: systematic review and meta-analysis. Bmj 334 (7597): 782.

Cogill, B. Anthropometric indicators measurement guide. Food and Nutrition Technical Assistance Project, Academy for Educational Development 2003

Crimmins, E. M., C. E. Finch 2006. Infection, inflammation, height, and longevity. Proc Natl Acad Sci U S A 103 (2): 498--503.

Daniels, D. L., S. N. Cousens, L. N. Makoae, R. G. Feachem 1990. A case-control study of the impact of improved sanitation on diarrhoea morbidity in Lesotho. Bull World Health Organ 68 (4): 455--463.

de Onis, M.; Onyango, A. W.; den Broeck, J. V.; Chumlea, W. C. & Martorell, R. Measurement and standardization protocols for anthropometry used in the construction of a new international growth reference. Food Nutr Bull, 2004, 25, S27-S36

Dewey, K. (2003).Guiding principles for complementary feeding of the breastfed child., Pan-American Health Organization (PAHO).

Dewey, K. G., S. Adu-Afarwuah 2008. Systematic review of the efficacy and effectiveness of complementary feeding interventions in developing countries. Matern Child Nutr 4 Suppl 1: 24--85.

Diallo, M. O., D. R. Hopkins, M. S. Kane, S. Niandou, A. Amadou, B. Kadri, A. Amza, P. M. Emerson, J. A. Zingeser 2007. Household latrine use, maintenance and acceptability in rural Zinder, Niger. Int J Environ Health Res 17 (6): 443--452.

Ejemot, R. I., J. E. Ehiri, M. M. Meremikwu, J. A. Critchley 2008. Hand washing for preventing diarrhoea. Cochrane Database Syst Rev (1): CD004265.

Emerson, P. M., S. W. Lindsay, et al. (2004). Role of flies and provision of latrines in trachoma control: cluster-randomised controlled trial. The *Lancet*363(9415): 1093-1098.

Ertem, I. O., Forsyth, B. W. C., Avni-Singer, A. J., Damour, L. K. & Cicchetti, D. V. (1997) Development of a Supplement to the HOME Scale for Children Living in Impoverished Urban Environments. *Journal of Developmental & Behavioral Pediatrics*,**18**, 322-328.

Espy, K. A.; Kaufmann, P. M.; McDiarmid, M. D.; and Glisky, M. L., "Executive functioning in preschool children: Performance on ANot-B and other delayed response format tasks" (1999). *Developmental Cognitive Neuroscience Laboratory - Faculty and Staff, Publications.* Paper 22. http://digitalcommons.unl.edu/dcnlfacpub/22

Esrey, S. A. 1996. Water, waste, and well-being: a multicountry study. Am J Epidemiol 143 (6): 608--623.

Esrey, S. A., J. B. Potash, L. Roberts, C. Shiff 1991. Effects of improved water supply and sanitation on ascariasis, diarrhoea, dracunculiasis, hookworm infection, schistosomiasis, and trachoma. Bull World Health Organ 69 (5): 609--621.

Ezzati, M., S. V. Hoorn, A. Rodgers, A. D. Lopez, C. D. Mathers, C. J. Murray 2003. Estimates of global and regional potential health gains from reducing multiple major risk factors. Lancet 362 (9380): 271--280.

Feikin DR, et al. (2010) Evaluation of the optimal recall period for disease symptoms in home-based morbidity surveillance in rural and urban Kenya. Int J Epidemiol 39: 450-458.

Fenson, L., Dale, P.S., Reznick, J.S., Bates, E., Thal, D., & Pethick, S. (1994). Variability in early communicative development. Monographs of the Society for Research in Child Development, 59(5), Serial No. 242).

Ferdous, T., Cederholm, T., Kabir, Z. N., Hamadani, J. D. & Wahlin, Å. (2010) Nutritional Status and Cognitive Function in Community-Living Rural Bangladeshi Older Adults: Data from the Poverty and Health in Ageing Project. *Journal of the American Geriatrics Society*,**58**, 919-924.

Ferguson AS, Layton AC, Mailloux BJ, Culligan PJ, Williams DE, Smartt AE, Sayler GS, Feighery J, McKay LD, Knappett PSK, Alexandrova E, Arbit T, Emch M, Escamilla V, Ahmed KM, Alam MJ, Streatfield PK, Yunus M, van Geen A. Comparison of fecal indicators with pathogenic bacteria and rotavirus in groundwater. Sci. Total Environ. 2012;431:314–22.

Fernald, Lia CH, et al. "Socioeconomic gradients in child development in very young children: Evidence from India, Indonesia, Peru, and Senegal."*Proceedings of the National Academy of Sciences* 109.Supplement 2 (2012): 17273-17280.

Fewtrell, L., R. B. Kaufmann, D. Kay, W. Enanoria, L. Haller, J. M. Colford 2005. Water, sanitation, and hygiene interventions to reduce diarrhoea in less developed countries: a systematic review and meta-analysis. Lancet Infect Dis 5 (1): 42--52.

Gerson, C. D., T. H. Kent, J. R. Saha, N. Siddiqi, J. Lindenbaum 1971. Recovery of small-intestinal structure and function after residence in the tropics. II. Studies in Indians and Pakistanis living in New York City. Ann Intern Med 75 (1): 41--48.

Gorter AC, Sandiford P, Pauw J, Morales P, Perez RM, Alberts H. Hygiene behaviour in rural Nicaragua in relation to diarrhoea. Int J Epidemiol. 1998;27(6):1090–100.

Goto, R., C. G. N. Mascie-Taylor, P. G. Lunn 2009. Impact of anti-Giardia and anthelminthic treatment on infant growth and intestinal permeability in rural Bangladesh: a randomised double-blind controlled study. Trans R Soc Trop Med Hyg 103 (5): 520--529.

Goto, R., C. G. N. Mascie-Taylor, P. G. Lunn 2009. Impact of intestinal permeability, inflammation status and parasitic infections on infant growth faltering in rural Bangladesh. Br J Nutr 101 (10): 1509--1516.

Grantham-McGregor, S., Y. B. Cheung, S. Cueto, P. Glewwe, L. Richter, B. Strupp, I. C. D. S. Group 2007. Developmental potential in the first 5 years for children in developing countries. Lancet 369 (9555): 60--70.

Guerrant, R. L., D. I. Guerrant, S. R. Moore, A. A. Lima, P. D. Patrick, J. B. Schorling 1999. Association of early childhood diarrhea and cryptosporidiosis with impaired physical fitness and cognitive function four-seven years later in a poor urban community in northeast Brazil. Am J Trop Med Hyg 61 (5): 707--713.

Guerrant, R. L., J. B. Schorling, J. F. McAuliffe, M. A. de Souza 1992. Diarrhea as a cause and an effect of malnutrition: diarrhea prevents catch-up growth and malnutrition increases diarrhea frequency and duration. Am J Trop Med Hyg 47 (1 Pt 2): 28--35.

Guerrant, R. L., R. B. Oriá, M. O. B. Oriá, S. R. Moore, A. A. M. Lima 2008. Malnutrition as an enteric infectious disease with long-term effects on child development. Nutr Rev 66 (9): 487--505.

Haghighi, P., P. L. Wolf 1997. Tropical sprue and subclinical enteropathy: a vision for the nineties. Crit Rev Clin Lab Sci 34 (4): 313--341.

Halloran ME, Longini IM, Cowart DM, Nizam A (2002) Community interventions and the epidemic prevention potential.Vaccine 20: 3254–3262.doi:10.1016/S0264-410X(02)00316-X.

Hamadani, J. D., Baker-Henningham, H., Tofail, F., Mehrin, F., Huda, S. N. & Grantham-Mcgregor, S. M. (2010) Validity and reliability of mothers' reports of language development in 1-year-old children in a large-scale survey in Bangladesh. *Food & Nutrition Bulletin*,**31**, 198-206.

Hamadani, J. D., Tofail, F., Cole, T. & Grantham-Mcgregor, S. (2013) The relation between age of attainment of motor milestones and future cognitive and motor development in Bangladeshi children. *Maternal & Child Nutrition*,**9**, 89-104.

Hamadani, J., Tofail, F., Hilaly, A., Huda, S., Engle, P. & Grantham-Mcgregor, S. (2010) The use of indicators of family care and their relation to child development in Bangladesh. *J Health Population Nutr*,**28**, 23-33.

Haque, A; Russell, N. Phenotypic and genotypic characterisation of Bacillus cereus isolates from Bangladeshi rice. International Journal of Food Microbiology. 2005;98,23-34.

Hoddinott, J., J. A. Maluccio, J. R. Behrman, R. Flores, R. Martorell 2008. Effect of a nutrition intervention during early childhood on economic productivity in Guatemalan adults. Lancet 371 (9610): 411--416.

Huda, S. N., Grantham-Mcgregor, S. M. & Tomkins, A. (2001) Cognitive and motor functions of iodine-deficient but euthyroid children in Bangladesh do not benefit from iodized poppy seed oil (Lipiodol). *The Journal of Nutrition*,**131**, 72-77.

Huda, S. N., Grantham-Mcgregor, S. M., Rahman, K. M. & Tomkins, A. (1999) Biochemical hypothyroidism secondary to iodine deficiency is associated with poor school achievement and cognition in Bangladeshi children. *The Journal of Nutrition*,**129**, 980-987.

Humphrey, J. H. 2009. Child undernutrition, tropical enteropathy, toilets, and handwashing. Lancet 374 (9694): 1032--1035.

Humphrey, J. H. 2009. Child undernutrition, tropical enteropathy, toilets, and handwashing. Lancet 374 (9694): 1032--1035.

Imai K, King G, Nall C(2009) The Essential Role of Pair Matching in Cluster-Randomized Experiments, with Application to the Mexican Universal Health Insurance Evaluation Statistical Science 24: 29-72.

Jensen PK, Jayasinghe G, van der Hoek W, Cairncross S, Dalsgaard A. Is there an association between bacteriological drinking water quality and childhood diarrhoea in developing countries? Trop. Med. Int. Health. 2004;9(11):1210–5.

Jukes, M., Nokes, C., Alcock, K. J., Lambo, J. K., Kihamia, C., Ngorosho, N., Mbise, A., Lorri, W., Yona, E. & Mwanri, L. (2002) Heavy schistosomiasis associated with poor short-term memory and slower reaction times in Tanzanian schoolchildren. *Tropical Medicine and International Health*,**7**, 104-117

Kabir, Z. N. & Herlitz, A. (2000) The Bangla Adaptation of Mini,Äêmental State Examination (BAMSE): an instrument to assess cognitive function in illiterate and literate individuals. *International Journal of Geriatric Psychiatry*,**15**, 441-450.

Kaplan, L. M., M. McGuckin 1986. Increasing handwashing compliance with more accessible sinks. Infect Control 7 (8): 408--410.

Kariger, P. K., Stoltzfus, R. J., Olney, D., Sazawal, S., Black, R., Tielsch, J. M., Frongillo, E. A., Khalfan, S. S. & Pollitt, E. (2005) Iron Deficiency and Physical Growth Predict Attainment of Walking but Not Crawling in Poorly Nourished Zanzibari Infants. *J. Nutr.*,**135**, 814-819.

Kariger, P., Frongillo, E. A., Engle, P., Britto, P. M. R., Sywulka, S. M. & Menon, P. (2012) Indicators of family care for development for use in multicountry surveys. *Journal of Health, Population, & Nutrition*,**30**, 472-486.

Knappett PSK, McKay LD, Layton A, Williams DE, Alam MJ, Huq MR, Mey J, Feighery JE, Culligan PJ, Mailloux BJ, Zhuang J, Escamilla V, Emch M, Perfect E, Sayler GS, Ahmed KM, van Geen A. Implications of fecal bacteria input from latrine-polluted ponds for wells in sandy aquifers. Environmental Science & Technology. 2011;46(3):1361–70.

Kosek M, Haque R, Lima A, et al. Fecal markers of intestinal inflammation and permeability associated with the subsequent acquisition of linear growth deficits in infants. *Am J Trop Med Hyg*. 2013 Feb;88(2):390-6.

Kremer, M., E. Miguel, S. Mullainathan, A. P. Zwane, C. Null 2009. Making water safe: Price, persuasion, peers, promoters, or product design?. Working Paper.

Law, J. & Roy, P. (2008) Parental Report of Infant Language Skills: A Review of the Development and Application of the Communicative Development Inventories. *Child and Adolescent Mental Health*,**13**, 198-206.

Leber J, Rahman KM, Ahmed B, Mailloux B, van Geen A. Contrasting Influence of Geology on E. coli and Arsenic in Aquifers of Bangaldesh. Ground Water. 2011;49(01):111-23.

Levy K, Hubbard AE, Nelson KL, Eisenberg JNS. Drivers of water quality variability in northern coastal Ecuador. Environmental Science & Technology. 2009;43(6):1788–97.

Lorntz, B., A. M. Soares, S. R. Moore, R. Pinkerton, B. Gansneder, V. E. Bovbjerg, H. Guyatt, A. M. Lima, R. L. Guerrant 2006. Early childhood diarrhea predicts impaired school performance. Pediatr Infect Dis J 25 (6): 513--520.

Luby, S. P., A. K. Halder, C. Tronchet, S. Akhter, A. Bhuiya, R. B. Johnston 2009. Household characteristics associated with handwashing with soap in rural Bangladesh. Am J Trop Med Hyg 81 (5): 882--887.

Lunn, P. G. 2000. The impact of infection and nutrition on gut function and growth in childhood. Proc Nutr Soc 59 (1): 147--154.

Lunn, P. G., C. A. Northrop-Clewes, R. M. Downes 1991. Intestinal permeability, mucosal injury, and growth faltering in Gambian infants. The Lancet 338 (8772): 907--910.

Magalhães RJS, Clements ACA, Patil AP, Gething PW, Brooker S (2011) The Applications of Model-Based Geostatistics in Helminth Epidemiology and Control.Adv Parasitol 74: 267–296.doi:10.1016/B978-0-12-385897-9.00005-7.

McKay, S., E. Gaudier, D. I. Campbell, A. M. Prentice, R. Albers 2010. Environmental enteropathy: new targets for nutritional interventions. Int Health 2 (3): 172--180.

Medley GF, Guyatt HL, Bundy DA (1993) A quantitative framework for evaluating the effect of community treatment on the morbidity due to ascariasis.Parasitology 106: 211–221.

Mitchell, A. J. (2009) A meta-analysis of the accuracy of the mini-mental state examination in the detection of dementia and mild cognitive impairment. *Journal of psychiatric research*,**43**, 411-431.

Mitra, A.K., Haque, A., Islam, M., Bashar, S.A., 2009. Lead poisoning: an alarming public health problem in Bangladesh. Int J Environ Res Public Health 6, 84-95.Meegan M, Morley D, Chavasse D. Fly traps. The Lancet. 1997;349(9055):886.

MM Black, AH Baqui, K Zaman, S El Arifeen, RE Black Maternal depressive symptoms and infant growth in rural Bangladesh. Am J Clin Nutr, 89 (2009), pp. 951S–957S

Muhibbullah, M., Momotaz, S., Chowdhury, A., 2005. Use of Agrochemical Fertilizers and Their Impact on Soil, Water and Human Health in the Khamargao Village of Mymensingh District, Bangladesh. Journal of Agronomy 4, 109-115.

Nahar, B., Hossain, M., Hamadani, J., Ahmed, T., Huda, S., Grantham-Mcgregor, S. & Persson, L. (2012) Effects of a community-based approach of food and psychosocial stimulation on growth and development of severely malnourished children in Bangladesh: a randomised trial. *European Journal of Clinical Nutrition*,**66**, 701-709.

Nandi, B. C., Ed. (2002). *The Fauna of India and Adjacent countries. Diptera volume 10: Sarcophagidae*. Kolkata, Zoological survey of India.

National Scientific Council on the Developing Child. (2005/2014). Excessive Stress Disrupts the Architecture of the Developing Brain: Working Paper 3. Updated Edition.

Niehaus, M. D., S. R. Moore, P. D. Patrick, L. L. Derr, B. Lorntz, A. A. Lima, R. L. Guerrant 2002. Early childhood diarrhea is associated with diminished cognitive function 4 to 7 years later in children in a northeast Brazilian shantytown. Am J Trop Med Hyg 66 (5): 590--593.

O'Brien KL, Millar EV, Zell ER, Bronsdon M, Weatherholtz R, et al. (2007) Effect of Pneumococcal Conjugate Vaccine on Nasopharyngeal Colonization Among Immunized and Unimmunized Children in a Community-Randomized Trial. J Infect Dis 196: 1211–1220. doi:10.1086/521833.

Pedersen, D. M., S. Keithly, K. Brady 1986. Effects of an observer on conformity to handwashing norm. Percept Mot Skills 62 (1): 169--170.

Petri, W. A., R. L. Guerrant, M. Miller, H. J. Binder, M. M. Levine, R. Dillingham 2008. Enteric infections, diarrhea, and their impact on function and development. J Clin Invest 118 (4): 1277--1290.

Phuka, J. C., K. Maleta, C. Thakwalakwa, Y. B. Cheung, A. Briend, M. J. Manary, P. Ashorn 2009. Postintervention growth of Malawian children who received 12-mo dietary complementation with a lipid-based nutrient supplement or maize-soy flour. Am J Clin Nutr 89 (1): 382--390.

Phuka, J. C., K. Maleta, C. Thakwalakwa, Y. B. Cheung, A. Briend, M. J. Manary, P. Ashorn 2008. Complementary feeding with fortified spread and incidence of severe stunting in 6- to 18-month-old rural Malawians. Arch Pediatr Adolesc Med 162 (7): 619--626.

Pickering AJ, Julian TR, Mamuya S, Boehm AB, Davis J. Bacterial hand contamination among Tanzanian mothers varies temporally and following household activities. Trop. Med. Int. Health. 2011;16(2):233–9.

Rabie, T., V. A. Curtis 2006. Handwashing and risk of respiratory infections: a quantitative systematic review. Trop Med Int Health 11 (3): 258--267.

Radloff, L. (1977) The CES-D Scale: A self-report depression scale for research in the general population. *Applied Psychological Measurement*,**1**, 385-401.

Ram PK, Jahid I, Halder AK, Nygren B, Islam MS, Granger SP, Molyneaux JW, Luby SP. Variability in Hand Contamination Based on Serial Measurements: Implications for Assessment of Hand-Cleansing Behavior and Disease Risk. Am. J. Trop. Med. Hyg. 2011;84(4):510–6.

Ram, P. K.; Halder, A. K.; Granger, S. P.; Jones, T.; Hall, P.; Hitchcock, D.; Wright, R.; Nygren, B.; Islam, M. S.; Molyneaux, J. W. & Luby, S. P. Is structured observation a valid technique to measure handwashing behavior? Use of acceleration sensors embedded in soap to assess reactivity to structured observation. Am J Trop Med Hyg, 2010, 83, 1070-1076

Ramakrishnan, U., P. Nguyen, R. Martorell 2009. Effects of micronutrients on growth of children under 5 y of age: meta-analyses of single and multiple nutrient interventions. Am J Clin Nutr 89 (1): 191--203.

Rosales, F. J., J. S. Reznick, S. H. Zeisel 2009. Understanding the role of nutrition in the brain and behavioral development of toddlers and preschool children: identifying and addressing methodological barriers. Nutr Neurosci 12 (5): 190--202.

Ruel MT, Arimond M. Spot-check Observational Method for Assessing Hygiene Practices: Review of Experience and Implications for Programmes. J Health Popul Nutr. 2002;20(1):65–76.

Sallis, J. F., N. Owen, E. B. Fisher 2008. Ecological Models of Health Behavior. 4th Edited by K Glanz, B K Rimer and K Viswanath. San Francisco: Josey-Bass.

Santamaría J, Toranzos GA. Enteric pathogens and soil: a short review. Int Microbiol. 2003;6(1):5–9.

Schmidt, W.-P., S. Cairncross 2009. Household Water Treatment in Poor Populations: Is There Enough Evidence for Scaling up Now?. Environ Sci Technol 43 (4): 986--992.

Shonkoff, J. P., Boyce, W. T., & McEwen, B. S. (2009). Neuroscience, molecular biology, and the childhood roots of health disparities: Building a new framework for health promotion and disease prevention. Journal of the American Medical Association, 301(21), 2252-2259.

Siegel, E. H., Stoltzfus, R. J., Kariger, P. K., Katz, J., Khatry, S. K., Leclerq, S. C., Pollitt, E. & Tielsch, J. M. (2005) Growth Indices, Anemia, and Diet Independently Predict Motor Milestone Acquisition of Infants in South Central Nepal. *J. Nutr.*,**135**, 2840-2844.

Sigman, M. (1995) Nutrition and Child Development: More Food for Thought. *Current Directions in Psychological Science*,**4**, 52-55.

Simms, V. M., P. Makalo, R. L. Bailey, P. M. Emerson 2005. Sustainability and acceptability of latrine provision in The Gambia. Trans R Soc Trop Med Hyg 99 (8): 631--637.

Stoeckel DM, Harwood VJ. Performance, Design, and Analysis in Microbial Source Tracking Studies. Applied and Environmental Microbiology. 2007;73(8):2405–15.

Tarleton, J. L., R. Haque, D. Mondal, J. Shu, B. M. Farr, W. A. Petri 2006. Cognitive effects of diarrhea, malnutrition, and Entamoeba histolytica infection on school age children in Dhaka, Bangladesh. Am J Trop Med Hyg 74 (3): 475--481.

TD Wachs, MM Black, PL Engle. Maternal depression: a global threat to children's health, development, and behavior and to human rights. Child Development Perspectives, 3 (2009), pp. 51–59

The Early Childhood Outcome Measures. *Instrument Crosswalks*. Retrieved November, 2006, from http://www.fpg.unc.edu/~eco/pdfs/ASQ crosswalk_5-9-06.pdf

The International Lipid-based Nutrient Supplements Project. http://ilins.org/, accessed September 28, 2011., iLiNS, Bill and Melinda Gates Foundation through the University of California, Davis(2011).

Thomas, G., D. J. Clain, A. C. Wicks 1976. Tropical enteropathy in Rhodesia. Gut 17 (11): 888--894.

Tofail F. Effect of food and micronutrient supplementation during pregnancy on subsequent development of infants in Bangladesh: a randomized trial. London, Institute of Child Health, University College London, 2006. 271 p. (Monograph of PhD thesis)

Unicef (2011).Program Guide for Infant and Young Child Feeding Practices., Unicef.

US department of health and wellness: Administration for children and families. *Child Development Instruments.* Retrieved November, 2006, from http://www.acf.hhs.gov/programs/opre/ehs/perf_measures/reports/resources_measuring/res_meas_cdi.html

Van Emden, F., Ed. (1965). *The Fauna of British India. Diptera volume 7: Muscidae*. London, Taylor and Francis.

van Geen A, Ahmed KM, Akita Y, Alam MJ, Culligan PJ, Emch M, Escamilla V, Feighery J, Ferguson AS, Knappett P, Layton AC, Mailloux BJ, McKay LD, Mey JL, Serre ML, Streatfield PK, Wu J, Yunus M. Fecal Contamination of Shallow Tubewells in Bangladesh Inversely Related to Arsenic. Environmental Science & Technology. 2011;45:1199–205.

Victora, C. G., L. Adair, C. Fall, P. C. Hallal, R. Martorell, L. Richter, H. S. Sachdev, Maternal, C. U. S. Group 2008. Maternal and child undernutrition: consequences for adult health and human capital. Lancet 371 (9609): 340--357.

Victora, C. G., M. de Onis, P. C. Hallal, M. Blossner, R. Shrimpton 2010. Worldwide timing of growth faltering: revisiting implications for interventions. Pediatrics 125 (3): e473--e480.

Walker, S. P., Chang, S. M., Powell, C. A., Simonoff, E. & Grantham-Mcgregor, S. M. (2007) Early Childhood Stunting Is Associated with Poor Psychological Functioning in Late Adolescence and Effects Are Reduced by Psychosocial Stimulation. *J. Nutr.*,**137**, 2464-2469.

Walker, S. P., T. D. Wachs, J. M. Gardner, B. Lozoff, G. A. Wasserman, E. Pollitt, J. A. Carter, I. C. D. S. Group 2007. Child development: risk factors for adverse outcomes in developing countries. Lancet 369 (9556): 145--157.

Walker, S. P., Wachs, T. D., Grantham-Mcgregor, S., Black, M. M., Nelson, C. A., Huffman, S. L., Baker-Henningham, H., Chang, S. M., Hamadani, J. D. & Lozoff, B. (2011) Inequality in early childhood: risk and protective factors for early child development. *The Lancet*,**378**, 1325-1338.

Walker, S. P., Wachs, T. D., Meeks Gardner, J., Lozoff, B., Wasserman, G. A., Pollitt, E. & Carter, J. A. (2007) Child development: risk factors for adverse outcomes in developing countries. *Lancet*,**369**, 145-157.

Warrell DA. Guidelines for the management of snake-bites. New Delhi: World Health Organization, 2010.

Wasserman GA, Liu X, Parvez F, Factor-Litvak P, Ahsan H, Levy D, Kline J, van Geen A, Mey J, Slavkovich V, Siddique AB, Islam T, Graziano JH. Arsenic and manganese exposure and children's intellectual function. Neurotoxicology. 2011 Aug;32(4):450-7. doi: 10.1016/j.neuro.2011.03.009. Epub 2011 Mar 29.

Wechsler, D. (1974) *Manual for the Wechsler intelligence scale for children, revised,* Psychological Corporation.

Wechsler, D. (1997) *WAIS-III: Wechsler adult intelligence scale,* Psychological Corporation San Antonio.

WHO (2006).WHO Child Growth Standards: Length/height-for-age, weight-for-age, weight-for-length, weight-for-height and body mass index-for-age: Methods and developments., World Health Organization, Geneva, 312.

WHO World Health Organization and United Nations Children's Fund Joint Monitoring Programme for Water Supply and Sanitation (JMP). Progress on Drinking Water and Sanitation: Special Focus on Sanitation., WHO, UNICEF, UNICEF, New York and WHO, Geneva(2008).

WHO. (2010). Indicators for assessing infant and young child feeding practices: Part II Measurement. Retrieved August 31, 2013, from http://www.who.int/nutrition/publications/infantfeeding/9789241599290/en/

Wijnhoven, T. M. A., De Onis, M., Onyango, A. W., Wang, T., Bjoerneboe, G.-E. A., Bhandari, N., Lartey, A. & Al Rashidi, B. (2004) Assessment of gross motor development in the WHO Multicentre Growth Reference Study. *Food and Nutrition Bulletin*,**25**, S37-S45.

Zhou, G; Liu, H; He, J; Yuan, Y; Yuan Z. The occurrence of Bacillus cereus, B. thuringiensis and B. mycoides in Chinese pasteurized full fat milk. International Journal of Food Microbiology. 2008;121,195-200.

Zwane AP, Zinman J, Dusen EV, Pariente W, Null C, et al. (2011) Being surveyed can change later behavior and related parameter estimates. Proc Natl Acad Sci U S A 108: 1821–1826. doi:10.1073/pnas.1000776108

PROTOCOL
Biomedical
Non-Exempt
Berkeley

---------------------------------------------------------------------------------------

**Protocol Title:** Measuring the benefits of sanitation, water quality, handwashing and nutrition interventions for improving health and development in rural Bangladesh

**Protocol Status:** APPROVED

**Date Submitted:** 10/26/2017

**Approval Period:** **10/30/2017-11/03/2018**

**Important Note:** **This Print View may not reflect all comments and contingencies for approval. Please check the comments section of the online protocol. Questions that appear to not have been answered may not have been required for this submission. Please see the system application for more details.**

*** * * Attached Document * * ***

| Document Name | Created Date |
|---|---|
| 2011-09-3652_Colford_1j-Consent-EE-v21-English_Tracked Changes.docx | 10/26/2017 |

**Appendix 1j: English consent form for environmental enteropathy and parasitic assessment**

Protocol Title: Effect of hand washing, water treatment, sanitation and nutritional supplement interventions on child health and development in rural Bangladesh

Principal Investigator's name: Dr. Leanne Unicomb

**Purpose of the research**
Hello/Assalamualaikum/Namaste. My name is ________ and I work with the ICDDR,B (Cholera Hospital) in Dhaka. We are conducting research on diarrheal diseases and through this particular study we want to learn about the health impact of diarrheal diseases in children. Malnutrition in children is associated with infections of our intestines that can change our use of nutrients. It is possible that genes in our body, made up of DNA, make some children more susceptible to malnutrition than others. This DNA is inherited from our parents and we also carry genes from many friendly microbes that live on and in our body. We are interested in learning if the exposure of a child to diarrheal disease has long term effects. We also are interested in learning how certain foods, nutrition, or life experiences in general may interact with diarrhea or other illnesses.

**Why are we inviting you to participate in the study?**

We are interested in enrolling this compound because we collected information on your household earlier and we are interested in conducting additional testing to evaluate your child's digestive tract. We are interested in evaluating your child's blood, stool, saliva, hair and urine for markers of nutritional status, infections and health. We are also interested in evaluating your saliva, hair, blood, and urine for markers of infections, stress and health.

**What is expected from the participants of the research study?**

Participation in this study will only span a few hours. To achieve the aim of the project, if you agree to be in this study, we will collect a blood, stool, and urine sample from your child. With your permission, we might also collect 5 saliva samples and 3-4 strands of hair from your child and a urine sample, a blood sample, 4 saliva samples, and 3-4 strands of hair from you. From the blood sample, we will measure nutritional markers, indicators of factors children inherit from their parents, and we will be able to understand whether your child has been exposed to infection. The urine sample will help us understand whether there has been a long term physical effect as a result of diarrhea. We will also collect a stool sample from your child with your help.

If you agree to participate, a field research person will visit your household twice for this purpose. Today a field member will weigh the child using a scale, measure his/her height using a height board, measure arm and head circumference using a tape measure and collect general health measures of blood pressure, sweat, and heart rate. The blood pressure and heart rate results from your child will be provided to you after the testing. The field representative will also collect general health measures of blood pressure, sweat, and heart rate. Your blood pressure and heart rate results will be provided to you after the testing. The field representative will also cut

and collect 3-4 strands of hair from you and your child. With your permission, we will collect a small amount of blood (equivalent to 1.5 teaspoons) from a vein in your child's arm to determine their blood group and genes (DNA) and test for infections that they may have had in the past, and measure their nutritional status. One drop of blood will be used to test your child for anemia, a condition of lower than normal red blood cells. The field member can provide you with the results of this test during their visit. We will record how your child responds to these procedures. If you agree, we would like to videotape your child during the blood-draw. We will use this information to better understand how these procedures affect child behavior. We will also ask you about how your child reacts to new situations, and what helps your child feel comfortable in new situations, which will take 10-15 minutes. This will help us understand your child's reaction to the different procedures we are administering for this project. We might also collect a small amount (10ml) of blood from your vein. We might also collect 3 saliva samples each from both you and your child before, during, and after the blood draw using a soft sponge placed under the tongue. The field member will deliver a stool collection kit and instruct you how to collect stool from your child. You will be instructed to collect your child's stool on the following morning, if the child defecates before their arrival, by having your child defecate in a clean diaper and use a plastic scoop to collect a small amount of fresh stool from the top of the pile into a container. The field person will collect this container when they come to collect the other specimens on the following day. Total participation time today will be approximately 3 hours and 15 minutes.

Tomorrow, we will ask you to collect your first urine sample of the morning immediately after you wake up. The main procedure will involve a second saliva collection, collecting your child's urine for 1 hour, feeding your child sugar syrup and then collecting their urine sample over a period of 5 hours. We intend to test the urine for the syrup to help us understand the health impacts of diarrhea in children. For a urine sample, due to the age of your child, we will be using a special urine collection bag to collect the urine from your child and we will demonstrate how it is used.

You/the mother will be requested to not feed your child for at least one hour before we feed him/her the syrup. During this fasting period, we will collect your child's urine for 1 hour by attaching the urine collection bag with a drainage tube (show sample) to the child. We will also collect two additional saliva samples from your child and one additional saliva sample from you using a soft sponge. We will then give a dose of the sugar syrup to the child and attach the urine collection bag with a drainage tube (show sample) to the child immediately after feeding him/her the syrup. We will encourage the child to drink water 30 minutes after taking the syrup to help urination. The field representative will remove the urine from the bag, whenever the child urinates. This collection will take place for 6 hours after which the bag will be removed from the child. During the 6-hour period of urine collection, you will be asked the quantities of foods (and the ingredients) you fed the child in the previous day and night. You will also be asked about your personal life experiences and health. You will be asked about perceptions regarding social norms that may affect a child's health. From the stool sample, we will examine whether your child has any worm infections, but this analysis will be done in Dhaka. We will offer your child deworming medication regardless of the stool sample result. Total participation time tomorrow will be approximately 7 hours.

Later, at the laboratory, we will measure your blood, saliva, hair, and urine samples and your child's blood, stool, saliva, hair, and urine samples for markers of nutritional status, infections, and health. You will not receive the results of any laboratory tests.

**Study time**: Study participation will take a total of approximately *10 hours and 15 minutes over 2 days.*

**Risks & Benefits**

Although we will try to protect your identity there is some possibility that confidentiality could be compromised. The syrup is a natural sugar solution that tastes pleasant. The blood will be collected by a trained professional. You and your child may feel some momentary pain during the blood collection. Your child may also feel some discomfort due to the presence of urine collection bag for 6 hours. Some of the questions I would like to ask you may seem private or personal since they touch on your life and health. You may feel uncomfortable talking about some of the topics. You do not have to answer any question or take part in the discussion/interview/survey if you don't wish to do so, and that is also fine. You do not have to give us any reason for not responding to any question, or for refusing to take part in the interview. All your answers will be kept as confidential as possible, and we anticipate that the risks from participating in this survey will be very minimal. There is no direct benefit for participating in this study, but your child's participation will help us to gain knowledge on the health impact of diarrheal diseases and how certain foods, nutrition, and life experiences in general may interact with diarrhea or other illnesses.

**Confidentiality**

All data and specimens collected will be kept confidential as allowed by the law of this country. The samples and the test results will be coded without your name or your child's name or personal information, and stored separately for analysis by the researchers. None of these researchers will be able to identify you or your child as the sample donors. Other persons who may have access to your test results include research groups that oversee the safety of the study including institutes such as the University of California Berkeley and the Bill and Melinda Gates Foundation, USA. Because of the potential need to release information to these parties, and the genetic information we are collecting is potentially identifying, we cannot guarantee absolute confidentiality. However, we will ensure every effort that we can to preserve your confidentiality. There is no expiration date to this permission. This information will be coded and it is unlikely that anyone will be able to trace it to you and your child, protecting your confidentiality.

All videotapes will be identified by a number only; no recordings will identify you or your child by name. These will be viewed only by trained personnel for coding of your child's response to the procedure. The videos will not be viewed by any other person. The videos will be stored in a locked cabinet accessible by study personnel only. These will never be made available to any persons not participating in the study.

**Future use of information**

The blood, urine, saliva, hair, and stool samples may be stored until the end of the study, so they can be analyzed in the lab at the same time. If you agree, the information, along with some of the blood, urine, saliva, hair and stool collected will be stored at ICDDR,B for up to 20 years so that we can conduct advanced tests on these samples to understand the infections that you may have had, and to find out what microbes lived in your body. No further consent will be taken from you, and you will not receive the results of these tests. A senior scientist at ICDDR,B will decide who can use the stored samples, and what tests will be done. If you change your mind about having your specimen saved for further testing, you may contact us and the sample will be discarded and not saved. The videotapes will be stored indefinitely.

**Right not to participate and withdraw**

Taking part in the study is completely voluntary. You may choose not to allow your child to participate in this study. You can drop out of this study at any time, even in the middle of the sample/urine collection. You have the right to refuse participation in this study, which will not affect your family's treatment at the Cholera Hospital in the future.

**Principle of compensation**

You need not pay us to take part in this study, and similarly we will not pay you money for attending in the study.

**Persons to contact:**

If you have any questions, you can ask me any time. If you have additional questions about the survey, you may contact:

If you have questions about your rights as a participant of a research study, or if you think some harm has been done to you because of the survey, you may contact or meet IRB Secretariat, M.A. Salam Khan:

M. A. Salam Khan, IRB Secretariat, phone: 9886498 or PABX 8860523-32 ext. 3206

If you agree to participate and you allow your child to participate, please indicate that by checking the boxes of the activities that you will agree to below:

**Urine Collection |__| mother |__| child**

**Venous Blood Collection |__| mother |__| child**

**Stool Collection |__| child**

**Saliva sample collection |__| mother |__| child**

**Hair sample collection |__| mother |__| child**

**Urine Long-term storage |__| mother |__| child**

**Venous Blood Long-term storage |__| mother |__| child**

**Stool Long-term storage |__| child**

**Saliva sample Long-term storage |__| mother |__| child**

**Hair sample Long-term storage |__| mother |__| child**

If you agree to our proposal of enrolling your household in our study, please indicate that by putting your signature or your left thumb impression at the specified space below

Thank you for your cooperation

_______________________________________ _____________________

Signature or left thumb impression of Guardian Date

_______________________________________ _____________________

Signature or left thumb impression of the witness Date

_______________________________________ _____________________

Signature of the PI or his/her representative Date

PROTOCOL
Biomedical
Non-Exempt
Berkeley

Protocol # 2011-09-3652
Date Printed: 07/31/2018

--------------------------------------------------------------------------------

**Protocol Title:** Measuring the benefits of sanitation, water quality, handwashing and nutrition interventions for improving health and development in rural Bangladesh

**Protocol Status:** APPROVED

**Date Submitted:** 10/26/2017

**Approval Period:** **10/30/2017-11/03/2018**

**Important Note:** **This Print View may not reflect all comments and contingencies for approval. Please check the comments section of the online protocol. Questions that appear to not have been answered may not have been required for this submission. Please see the system application for more details.**

*** * * Attached Document * * ***

| Document Name | Created Date |
|---|---|
| icddr,b PR-11063_Continuation Approval_24 Feb 2015_9 April 2015-8 April 2016.pdf | 10/26/2017 |

icddr,b

### Approval for continuation of research activity

This is to certify that icddr,b research protocol # PR-11063 titled 'A randomized controlled trial of the benefits of water sanitation hygiene plus nutrition interventions on child growth' PI – Dr Leanne Unicomb, Centre for Communicable Diseases (CCD) had been approved by Ethical Review Committee (ERC) on 9 April 2012.

The ERC undertakes annual/periodic review of all ERC-approved protocols for reappraisal. The ERC approval for implementation of any research protocol is not, however, affected unless any unanticipated problems involving risks to the study participants or any serious or continuing noncompliance of the ERC Guidelines are detected in the implementation of the study, during the review period.

The review undertaken as of 24 February 2015 to oversee the implementation of the above protocol reveals no Adverse Event (AE) or Serious Adverse Event (SAE) or unanticipated problems involving risks to the study participants or any serious or continuing noncompliance of the ERC Guidelines. Therefore, the ERC is pleased to **approve** the protocol for continuation of its activity for next one year starting **from 9 April 2015 to 8 April 2016**.

The continuing review application must be submitted to the IRB Secretariat for this study to continue beyond **8 April 2016**. All necessary materials for continuing review must be reviewed with sufficient time for review and issuing continued approval before the expiration date. Failure to initiate a continuing review application in a timely fashion may result in discontinuation of study activities until approval can be renewed. Performing study activities, including data analysis, beyond the expiration date results in noncompliance of federal regulations.

Other terms and conditions for implementation of your research protocol, as contained in our memo dated 9 April 2012 according initial approval of the research protocol shall, however, remain unchanged.

Professor Kazi Zulfiquer Mamun
*MBBS, M.Trop. Med, PhD.*
Chairperson
Ethical Review Committee of icddr,b
24 February 2015

Cc: Coordination Manager, Grants RAS (GR-00741)
Financial Controller
(Budget code # 4112258, 11112836, 11112837, 11112838)

GPO Box 128, Dhaka 1000, Bangladesh.
**For Packages :** 68, Shaheed Tajuddin Ahmed Sarani, Mohakhali, Dhaka 1212, Bangladesh.

PROTOCOL
Biomedical
Non-Exempt
Berkeley

Protocol # 2011-09-3652
Date Printed: 07/31/2018

-----------------------------------------------------------------------------------

| | |
|---|---|
| **Protocol Title:** | Measuring the benefits of sanitation, water quality, handwashing and nutrition interventions for improving health and development in rural Bangladesh |
| **Protocol Status:** | APPROVED |
| **Date Submitted:** | 10/26/2017 |
| **Approval Period:** | **10/30/2017-11/03/2018** |
| **Important Note:** | **This Print View may not reflect all comments and contingencies for approval. Please check the comments section of the online protocol. Questions that appear to not have been answered may not have been required for this submission. Please see the system application for more details.** |

*** * * Attached Document * * ***

| Document Name | Created Date |
|---|---|
| Stanford WASH Benefits IRB continuation approval Nov. 2015.pdf | 10/26/2017 |

### STANFORD UNIVERSITY

*Stanford, CA 94305 [Mail Code 5579]*

Ronald L. Ariagno, M.D. (650) 724-7541

CHAIR, PANEL ON MEDICAL HUMAN SUBJECTS (650) 725-8815

#### Certification of Human Subjects Approvals

**Date:** November 24, 2015

**To:** Stephen P Luby, M.D., Medicine - Med/Infectious Diseases
Suzanne Mueller N/A, Amy Janel Pickering PhD, Firdaus Dhabhar Ph.D.

**From:** Ronald L. Ariagno, M.D., Administrative Panel on Human Subjects in Medical Research

**Protocol** Randomized Controlled Trial of the Benefits of Water Sanitation Hygiene Plus Nutrition Interventions on Child Growth.

**Protocol ID:** 25863 **IRB Number:** 351 (Panel: 4)

The IRB approved human subjects involvement in your research project on 11/24/2015. **'Prior to subject recruitment and enrollment, if this is: a Cancer-related study, you must obtain Cancer Center Scientific Review Committee (SRC) approval;** a CTRU study, you must obtain CTRU approval; a VA study, you must obtain VA R and D Committee approval; and if a contract is involved, it must be signed.'

The expiration date of this approval is 10/31/2016 at Midnight. If this project is to continue beyond that date, you must submit an updated protocol in advance for the IRB's re-approval. If this protocol is used in conjunction with any other human use it must be re-approved. Proposed changes to approved research must be reviewed and approved prospectively by the IRB. No changes may be initiated without prior approval by the IRB, except where necessary to eliminate apparent immediate hazards to subjects. (Any such exceptions must be reported to the IRB within 10 working days.) Unanticipated problems involving risks to participants or others and other events or information, as defined and listed in the Report Form, must be submitted promptly to the IRB. (See Events and Information that Require Prompt Reporting to the IRB at http://humansubjects.stanford.edu.)

All continuing projects and activities must be reviewed and re-approved on or before Midnight of the expiration date. The approval period will be less than one year if so determined by the IRB. It is your responsibility to resubmit the project to the IRB for continuing review and to report the completion of the protocol to the IRB within 30 days.

Please remember that all data, including all signed consent form documents, must be retained for a minimum of three years past the completion of this research. Additional requirements may be imposed by your funding agency, your department, or other entities. (See Policy 1.9 on Retention of and Access to Research Data at http://doresearch.stanford.edu/policies/research-policy-handbook)

This institution is in compliance with requirements for protection of human subjects, including 45 CFR 46, 21 CFR 50 and 56, and 38 CFR 16.

Includes: Personnel changes: Removed Rossanna Gonzalez-Ayala and added Firdaus Dhabhar
Revised study target to 16,900.
Removed inactive funding SPO 112788.
Waiver of Individual Authorization under 45 CFR 164.512(i)(2)(ii)(A),(B),(C), pursuant to information provided in the HIPAA section of the protocol application.

Ronald L. Ariagno, M.D., Chair

**Approval Period:** 11/24/2015 THROUGH 10/31/2016

**Review Type:** REGULAR - CONTINUING REVIEW

**Funding:** University of California Berkeley, Prime Bill and Melinda Gates Foundation , SPO: 108704

**Assurance Number:** FWA00000935 (SU)

### STANFORD UNIVERSITY

*Stanford, CA 94305 [Mail Code 5579]*

Ronald L. Ariagno, M.D. (650) 724-7541
CHAIR, PANEL ON MEDICAL HUMAN SUBJECTS (650) 725-8815

#### Certification of Human Subjects Approvals

**Date:** November 24, 2015

**To:** Stephen P Luby, M.D., Medicine - Med/Infectious Diseases
Suzanne Mueller N/A, Amy Janel Pickering PhD, Firdaus Dhabhar Ph.D.

**From:** Ronald L. Ariagno, M.D., Administrative Panel on Human Subjects in Medical Research

**Protocol** Randomized Controlled Trial of the Benefits of Water Sanitation Hygiene Plus Nutrition Interventions on Child Growth.

**Protocol ID:** 25863 **IRB Number:** 351 (Panel: 4)

The IRB approved human subjects involvement in your research project on 11/24/2015. **'Prior to subject recruitment and enrollment, if this is: a Cancer-related study, you must obtain Cancer Center Scientific Review Committee (SRC) approval;** a CTRU study, you must obtain CTRU approval; a VA study, you must obtain VA R and D Committee approval; and if a contract is involved, it must be signed.'

The expiration date of this approval is 10/31/2016 at Midnight. If this project is to continue beyond that date, you must submit an updated protocol in advance for the IRB's re-approval. If this protocol is used in conjunction with any other human use it must be re-approved. Proposed changes to approved research must be reviewed and approved prospectively by the IRB. No changes may be initiated without prior approval by the IRB, except where necessary to eliminate apparent immediate hazards to subjects. (Any such exceptions must be reported to the IRB within 10 working days.) Unanticipated problems involving risks to participants or others and other events or information, as defined and listed in the Report Form, must be submitted promptly to the IRB. (See Events and Information that Require Prompt Reporting to the IRB at http://humansubjects.stanford.edu.)

All continuing projects and activities must be reviewed and re-approved on or before Midnight of the expiration date. The approval period will be less than one year if so determined by the IRB. It is your responsibility to resubmit the project to the IRB for continuing review and to report the completion of the protocol to the IRB within 30 days.

Please remember that all data, including all signed consent form documents, must be retained for a minimum of three years past the completion of this research. Additional requirements may be imposed by your funding agency, your department, or other entities. (See Policy 1.9 on Retention of and Access to Research Data at http://doresearch.stanford.edu/policies/research-policy-handbook)

This institution is in compliance with requirements for protection of human subjects, including 45 CFR 46, 21 CFR 50 and 56, and 38 CFR 16.

Includes: Personnel changes: Removed Rossanna Gonzalez-Ayala and added Firdaus Dhabhar
Revised study target to 16,900.
Removed inactive funding SPO 112788.
Waiver of Individual Authorization under 45 CFR 164.512(i)(2)(ii)(A),(B),(C), pursuant to information provided in the HIPAA section of the protocol application.

Ronald L. Ariagno, M.D., Chair

**Approval Period:** 11/24/2015 THROUGH 10/31/2016

**Review Type:** REGULAR - CONTINUING REVIEW

**Funding:** Woods Institute for the Environment , SPO: 112476

**Assurance Number:** FWA00000935 (SU)

LELAND STANFORD JUNIOR UNIVERSITY
DIE LUFT DER FREIHEIT WEHT
1891

### STANFORD UNIVERSITY

*Stanford, CA 94305 [Mail Code 5579]*

Ronald L. Ariagno, M.D. (650) 724-7541
CHAIR, PANEL ON MEDICAL HUMAN SUBJECTS (650) 725-8815

#### Certification of Human Subjects Approvals

**Date:** November 24, 2015

**To:** Stephen P Luby, M.D., Medicine - Med/Infectious Diseases
Suzanne Mueller N/A, Amy Janel Pickering PhD, Firdaus Dhabhar Ph.D.

**From:** Ronald L. Ariagno, M.D., Administrative Panel on Human Subjects in Medical Research

**Protocol** Randomized Controlled Trial of the Benefits of Water Sanitation Hygiene Plus Nutrition Interventions on Child Growth.

**Protocol ID:** 25863 **IRB Number:** 351 (Panel: 4)

The IRB approved human subjects involvement in your research project on 11/24/2015. **'Prior to subject recruitment and enrollment, if this is: a Cancer-related study, you must obtain Cancer Center Scientific Review Committee (SRC) approval;** a CTRU study, you must obtain CTRU approval; a VA study, you must obtain VA R and D Committee approval; and if a contract is involved, it must be signed.'

The expiration date of this approval is 10/31/2016 at Midnight. If this project is to continue beyond that date, you must submit an updated protocol in advance for the IRB's re-approval. If this protocol is used in conjunction with any other human use it must be re-approved. Proposed changes to approved research must be reviewed and approved prospectively by the IRB. No changes may be initiated without prior approval by the IRB, except where necessary to eliminate apparent immediate hazards to subjects. (Any such exceptions must be reported to the IRB within 10 working days.) Unanticipated problems involving risks to participants or others and other events or information, as defined and listed in the Report Form, must be submitted promptly to the IRB. (See Events and Information that Require Prompt Reporting to the IRB at http://humansubjects.stanford.edu.)

All continuing projects and activities must be reviewed and re-approved on or before Midnight of the expiration date. The approval period will be less than one year if so determined by the IRB. It is your responsibility to resubmit the project to the IRB for continuing review and to report the completion of the protocol to the IRB within 30 days.

Please remember that all data, including all signed consent form documents, must be retained for a minimum of three years past the completion of this research. Additional requirements may be imposed by your funding agency, your department, or other entities. (See Policy 1.9 on Retention of and Access to Research Data at http://doresearch.stanford.edu/policies/research-policy-handbook)

This institution is in compliance with requirements for protection of human subjects, including 45 CFR 46, 21 CFR 50 and 56, and 38 CFR 16.

Includes: Personnel changes: Removed Rossanna Gonzalez-Ayala and added Firdaus Dhabhar
Revised study target to 16,900.
Removed inactive funding SPO 112788.
Waiver of Individual Authorization under 45 CFR 164.512(i)(2)(ii)(A),(B),(C), pursuant to information provided in the HIPAA section of the protocol application.

Ronald L. Ariagno, M.D., Chair

**Approval Period:** 11/24/2015 THROUGH 10/31/2016

**Review Type:** REGULAR - CONTINUING REVIEW

**Funding:** University of California Berkeley, Prime NIH , SPO: 110499

**Assurance Number:** FWA00000935 (SU)

LELAND STANFORD JUNIOR UNIVERSITY
DIE LUFT DER FREIHEIT WEHT
1891

### STANFORD UNIVERSITY

*Stanford, CA 94305 [Mail Code 5579]*

Ronald L. Ariagno, M.D. (650) 724-7541

CHAIR, PANEL ON MEDICAL HUMAN SUBJECTS (650) 725-8815

#### Certification of Human Subjects Approvals

**Date:** November 24, 2015

**To:** Stephen P Luby, M.D., Medicine - Med/Infectious Diseases
Suzanne Mueller N/A, Amy Janel Pickering PhD, Firdaus Dhabhar Ph.D.

**From:** Ronald L. Ariagno, M.D., Administrative Panel on Human Subjects in Medical Research

**Protocol** Randomized Controlled Trial of the Benefits of Water Sanitation Hygiene Plus Nutrition Interventions on Child Growth.

**Protocol ID:** 25863 **IRB Number:** 351 (Panel: 4)

The IRB approved human subjects involvement in your research project on 11/24/2015. **'Prior to subject recruitment and enrollment, if this is: a Cancer-related study, you must obtain Cancer Center Scientific Review Committee (SRC) approval;** a CTRU study, you must obtain CTRU approval; a VA study, you must obtain VA R and D Committee approval; and if a contract is involved, it must be signed.'

The expiration date of this approval is 10/31/2016 at Midnight. If this project is to continue beyond that date, you must submit an updated protocol in advance for the IRB's re-approval. If this protocol is used in conjunction with any other human use it must be re-approved. Proposed changes to approved research must be reviewed and approved prospectively by the IRB. No changes may be initiated without prior approval by the IRB, except where necessary to eliminate apparent immediate hazards to subjects. (Any such exceptions must be reported to the IRB within 10 working days.) Unanticipated problems involving risks to participants or others and other events or information, as defined and listed in the Report Form, must be submitted promptly to the IRB. (See Events and Information that Require Prompt Reporting to the IRB at http://humansubjects.stanford.edu.)

All continuing projects and activities must be reviewed and re-approved on or before Midnight of the expiration date. The approval period will be less than one year if so determined by the IRB. It is your responsibility to resubmit the project to the IRB for continuing review and to report the completion of the protocol to the IRB within 30 days.

Please remember that all data, including all signed consent form documents, must be retained for a minimum of three years past the completion of this research. Additional requirements may be imposed by your funding agency, your department, or other entities. (See Policy 1.9 on Retention of and Access to Research Data at http://doresearch.stanford.edu/policies/research-policy-handbook)

This institution is in compliance with requirements for protection of human subjects, including 45 CFR 46, 21 CFR 50 and 56, and 38 CFR 16.

Includes: Personnel changes: Removed Rossanna Gonzalez-Ayala and added Firdaus Dhabhar
Revised study target to 16,900.
Removed inactive funding SPO 112788.
Waiver of Individual Authorization under 45 CFR 164.512(i)(2)(ii)(A),(B),(C), pursuant to information provided in the HIPAA section of the protocol application.

Ronald L. Ariagno, M.D., Chair

**Approval Period:** 11/24/2015 THROUGH 10/31/2016

**Review Type:** REGULAR - CONTINUING REVIEW

**Funding:** University of California Berkeley, Prime NIH , SPO: 108161

**Assurance Number:** FWA00000935 (SU)

LELAND STANFORD JUNIOR UNIVERSITY
DIE LUFT DER FREIHEIT WEHT
1891

PROTOCOL
Biomedical
Non-Exempt
Berkeley

Protocol # 2011-09-3652
Date Printed: 07/31/2018

---------------------------------------------------------------------------------

| | |
|---|---|
| **Protocol Title:** | Measuring the benefits of sanitation, water quality, handwashing and nutrition interventions for improving health and development in rural Bangladesh |
| **Protocol Status:** | APPROVED |
| **Date Submitted:** | 10/26/2017 |
| **Approval Period:** | **10/30/2017-11/03/2018** |
| **Important Note:** | **This Print View may not reflect all comments and contingencies for approval. Please check the comments section of the online protocol. Questions that appear to not have been answered may not have been required for this submission. Please see the system application for more details.** |

*** * * Attached Document * * ***

| Document Name | Created Date |
|---|---|
| PR-11063 ERC approval addendum 09Apr2016.pdf | 10/26/2017 |

icddr,b

### Approval for continuation of research activity

This is to certify that icddr,b research protocol # PR-11063 titled "A randomized controlled trial of the benefits of water sanitation hygiene plus nutrition interventions on child growth": PI – Dr Md Mahbubur Rahman of the Infectious Diseases Division (IDD) had been approved by Ethical Review Committee (ERC) on 09 April 2012.

The ERC undertakes annual/periodic review of all ERC-approved protocols for reappraisal. The ERC approval for implementation of any research protocol is not, however, affected unless any unanticipated problems involving risks to the study participants or any serious or continuing noncompliance of the ERC Guidelines are detected in the implementation of the study, during the review period.

The review undertaken as of 21 March 2016 to oversee the implementation of the above protocol reveals no Adverse Event (AE) or Serious Adverse Event (SAE) or unanticipated problems involving risks to the study participants or any serious or continuing noncompliance of the ERC Guidelines. Therefore, the ERC is pleased to **approve** the protocol for continuation of its activity for next one year starting from **09 April 2016** to **08 April 2017**

The continuing review application must be submitted to the IRB Secretariat for this study to continue beyond 08 April 2017. All necessary materials for continuing review must be reviewed with sufficient time for review and issuing continued approval before the expiration date. Failure to initiate a continuing review application in a timely fashion may result in discontinuation of study activities until approval can be renewed. Performing study activities, including data analysis, beyond the expiration date results in noncompliance of federal regulations.

Other terms and conditions for implementation of your research protocol, as contained in our memo dated 09 April 2012 according initial approval of the research protocol shall, however, remain unchanged.

[signature]

Professor Kazi Zulfiquer Mamun
*MBBS, M.Trop. Med, PhD.*
Chairperson
Ethical Review Committee of icddr,b
22 March 2016

Cc: Coordination Manager, Grants, RA (GR-00741)
Financial Controller
(Budget Code # 04112258, 11112836, 11112837, 11112838)

GPO Box 128, Dhaka 1000, Bangladesh.
**For Packages :** 68, Shaheed Tajuddin Ahmed Sarani, Mohakhali, Dhaka 1212, Bangladesh.

PROTOCOL
Biomedical
Non-Exempt
Berkeley

Protocol # 2011-09-3652
Date Printed: 07/31/2018

---------------------------------------------------------------------------------------

**Protocol Title:** Measuring the benefits of sanitation, water quality, handwashing and nutrition interventions for improving health and development in rural Bangladesh

**Protocol Status:** APPROVED

**Date Submitted:** 10/26/2017

**Approval Period:** **10/30/2017-11/03/2018**

**Important Note:** **This Print View may not reflect all comments and contingencies for approval. Please check the comments section of the online protocol. Questions that appear to not have been answered may not have been required for this submission. Please see the system application for more details.**

*** * * Attached Document * * ***

| Document Name | Created Date |
| --- | --- |
| PR#11063 (WASHB) Continuation approval upto April 8, 2018.pdf | 10/26/2017 |

icddr,b

### Approval for continuation of research activity

This is to certify that icddr,b research protocol # PR-11063 titled "A randomized controlled trial of the benefits of water sanitation hygiene plus nutrition interventions on child growth": PI – Dr Md Mahbubur Rahman of the Infectious Diseases Division (IDD) had been approved by Ethical Review Committee (ERC) on 09 April 2012.

The ERC undertakes annual/periodic review of all ERC-approved protocols for reappraisal. The ERC approval for implementation of any research protocol is not, however, affected unless any unanticipated problems involving risks to the study participants or any serious or continuing noncompliance of the ERC Guidelines are detected in the implementation of the study, during the review period.

The review undertaken as of 24 January 2017 to oversee the implementation of the above protocol reveals no Adverse Event (AE) or Serious Adverse Event (SAE) or unanticipated problems involving risks to the study participants or any serious or continuing noncompliance of the ERC Guidelines. Therefore, the ERC is pleased to **approve** the protocol for continuation of its activity for next one year starting from **09 April 2017** to **08 April 2018**.

The continuing review application must be submitted to the IRB Secretariat for this study to continue beyond 08 April 2018. All necessary materials for continuing review must be reviewed with sufficient time for review and issuing continued approval before the expiration date. Failure to initiate a continuing review application in a timely fashion may result in discontinuation of study activities until approval can be renewed. Performing study activities, including data analysis, beyond the expiration date results in noncompliance of federal regulations.

Other terms and conditions for implementation of your research protocol, as contained in our memo dated 09 April 2012 according initial approval of the research protocol shall, however, remain unchanged.

Professor Kazi Zulfiquer Mamun
*MBBS, M.Trop. Med, PhD.*
Chairperson
Ethical Review Committee of icddr,b
12 January 2017

Cc: Coordination Manager, Grants, RA (GR-00741)
Controller, Finance
(Budget Code # 04112258, 11112836, 11112837, 11112838)

GPO Box 128, Dhaka 1000, Bangladesh.
**For Packages :** 68, Shaheed Tajuddin Ahmed Sarani, Mohakhali, Dhaka 1212, Bangladesh.

PROTOCOL
Biomedical
Non-Exempt
Berkeley

Protocol # 2011-09-3652
Date Printed: 07/31/2018

-----------------------------------------------------------------------------------
---

**Protocol Title:** Measuring the benefits of sanitation, water quality, handwashing and nutrition interventions for improving health and development in rural Bangladesh

**Protocol Status:** APPROVED

**Date Submitted:** 10/26/2017

**Approval Period:** 10/30/2017-11/03/2018

**Important Note:** This Print View may not reflect all comments and contingencies for approval. Please check the comments section of the online protocol.
Questions that appear to not have been answered may not have been required for this submission. Please see the system application for more details.

*** Attached Document ***

| Document Name | Created Date |
| --- | --- |
| Stanford IRB ApprovalLetter-25863 through Oct 2017.pdf | 10/26/2017 |

### STANFORD UNIVERSITY

*Stanford, CA 94305 [Mail Code 5579]*

Ronald L. Ariagno, M.D. (650) 724-7541
CHAIR, PANEL ON MEDICAL HUMAN SUBJECTS (650) 725-8815

#### Certification of Human Subjects Approvals

**Date:** October 31, 2016

**To:** Stephen P Luby, M.D., Medicine - Med/Infectious Diseases
Suzanne Mueller N/A, Amy Janel Pickering PhD

**From:** Ronald L. Ariagno, M.D., Administrative Panel on Human Subjects in Medical Research

**eProtocol Title:** Randomized Controlled Trial of the Benefits of Water Sanitation Hygiene Plus Nutrition Interventions on Child Growth.

**eProtocol #:** 25863 **IRB 4 (Registration #:** 351)

The IRB approved human subjects involvement in your research project on 10/31/2016. **'Prior to subject recruitment and enrollment, if this is: a Cancer-related study, you must obtain Cancer Center Scientific Review Committee (SRC) approval;** a CTRU study, you must obtain CTRU approval; a VA study, you must obtain VA R and D Committee approval; and if a contract is involved, it must be signed.'

The expiration date of this approval is 10/31/2017 at Midnight. If this research is to continue beyond that date, it is your responsibility to submit a Continuing Review application in eProtocol. Research activities must be reviewed and re-approved on or before midnight of the expiration date. The approval period may be less than one year if so determined by the IRB. Proposed changes to approved research must be reviewed and approved prospectively by the IRB. No changes may be initiated without prior approval by the IRB, except where necessary to eliminate apparent immediate hazards to subjects. (Any such exceptions must be reported to the IRB within 10 working days.) Unanticipated problems involving risks to participants or others and other events or information, as defined and listed in the Report Form, must be submitted promptly to the IRB. (See Events and Information that Require Prompt Reporting to the IRB at http://humansubjects.stanford.edu.) Upon completion, you must report to the IRB within 30 days.

Please remember that all data, including all signed consent form documents, must be retained for a minimum of three years past the completion of this research. Additional requirements may be imposed by your funding agency, your department, HIPAA, or other entities. (See Policy 1.9 on Retention of and Access to Research Data at http://doresearch.stanford.edu/policies/research-policy-handbook)

This institution is in compliance with requirements for protection of human subjects, including 45 CFR 46, 21 CFR 50 and 56, and 38 CFR 16.

Includes: Personnel change: removed Fridus Dhabhar

Waiver of Individual Authorization under 45 CFR 164.512(i)(2)(ii)(A),(B),(C), pursuant to information provided in the HIPAA section of the protocol application.

Ronald L. Ariagno, M.D., Chair

**Approval Period:** 10/31/2016 THROUGH 10/31/2017

**Review Type:** REGULAR - CONTINUING REVIEW

**Funding:** University of California Berkeley, Prime Bill and Melinda Gates Foundation , SPO: 108704

**Assurance #:** FWA00000935 (SU)

LELAND STANFORD JUNIOR UNIVERSITY
DIE LUFT DER FREIHEIT WEHT
1891

### STANFORD UNIVERSITY

*Stanford, CA 94305 [Mail Code 5579]*

Ronald L. Ariagno, M.D. (650) 724-7541

CHAIR, PANEL ON MEDICAL HUMAN SUBJECTS (650) 725-8815

#### Certification of Human Subjects Approvals

**Date:** October 31, 2016

**To:** Stephen P Luby, M.D., Medicine - Med/Infectious Diseases
Suzanne Mueller N/A, Amy Janel Pickering PhD

**From:** Ronald L. Ariagno, M.D., Administrative Panel on Human Subjects in Medical Research

**eProtocol Title:** Randomized Controlled Trial of the Benefits of Water Sanitation Hygiene Plus Nutrition Interventions on Child Growth.

**eProtocol #:** 25863 **IRB 4** **(Registration #:** 351)

The IRB approved human subjects involvement in your research project on 10/31/2016. **'Prior to subject recruitment and enrollment, if this is: a Cancer-related study, you must obtain Cancer Center Scientific Review Committee (SRC) approval;** a CTRU study, you must obtain CTRU approval; a VA study, you must obtain VA R and D Committee approval; and if a contract is involved, it must be signed.'

The expiration date of this approval is 10/31/2017 at Midnight. If this research is to continue beyond that date, it is your responsibility to submit a Continuing Review application in eProtocol. Research activities must be reviewed and re-approved on or before midnight of the expiration date. The approval period may be less than one year if so determined by the IRB. Proposed changes to approved research must be reviewed and approved prospectively by the IRB. No changes may be initiated without prior approval by the IRB, except where necessary to eliminate apparent immediate hazards to subjects. (Any such exceptions must be reported to the IRB within 10 working days.) Unanticipated problems involving risks to participants or others and other events or information, as defined and listed in the Report Form, must be submitted promptly to the IRB. (See Events and Information that Require Prompt Reporting to the IRB at http://humansubjects.stanford.edu.) Upon completion, you must report to the IRB within 30 days.

Please remember that all data, including all signed consent form documents, must be retained for a minimum of three years past the completion of this research. Additional requirements may be imposed by your funding agency, your department, HIPAA, or other entities. (See Policy 1.9 on Retention of and Access to Research Data at http://doresearch.stanford.edu/policies/research-policy-handbook)

This institution is in compliance with requirements for protection of human subjects, including 45 CFR 46, 21 CFR 50 and 56, and 38 CFR 16.

Includes: Personnel change: removed Fridus Dhabhar

Waiver of Individual Authorization under 45 CFR 164.512(i)(2)(ii)(A),(B),(C), pursuant to information provided in the HIPAA section of the protocol application.

Ronald L. Ariagno, M.D., Chair

**Approval Period:** 10/31/2016 THROUGH 10/31/2017

**Review Type:** REGULAR - CONTINUING REVIEW

**Funding:** Woods Institute for the Environment , SPO: 112476

**Assurance #:** FWA00000935 (SU)

LELAND STANFORD JUNIOR UNIVERSITY
DIE LUFT DER FREIHEIT WEHT
1891

### STANFORD UNIVERSITY

*Stanford, CA 94305 [Mail Code 5579]*

Ronald L. Ariagno, M.D. (650) 724-7541

CHAIR, PANEL ON MEDICAL HUMAN SUBJECTS (650) 725-8815

#### Certification of Human Subjects Approvals

**Date:** October 31, 2016

**To:** Stephen P Luby, M.D., Medicine - Med/Infectious Diseases
Suzanne Mueller N/A, Amy Janel Pickering PhD

**From:** Ronald L. Ariagno, M.D., Administrative Panel on Human Subjects in Medical Research

**eProtocol Title:** Randomized Controlled Trial of the Benefits of Water Sanitation Hygiene Plus Nutrition Interventions on Child Growth.

**eProtocol #:** 25863 **IRB 4 (Registration #:** 351)

The IRB approved human subjects involvement in your research project on 10/31/2016. **'Prior to subject recruitment and enrollment, if this is: a Cancer-related study, you must obtain Cancer Center Scientific Review Committee (SRC) approval;** a CTRU study, you must obtain CTRU approval; a VA study, you must obtain VA R and D Committee approval; and if a contract is involved, it must be signed.'

The expiration date of this approval is 10/31/2017 at Midnight. If this research is to continue beyond that date, it is your responsibility to submit a Continuing Review application in eProtocol. Research activities must be reviewed and re-approved on or before midnight of the expiration date. The approval period may be less than one year if so determined by the IRB. Proposed changes to approved research must be reviewed and approved prospectively by the IRB. No changes may be initiated without prior approval by the IRB, except where necessary to eliminate apparent immediate hazards to subjects. (Any such exceptions must be reported to the IRB within 10 working days.) Unanticipated problems involving risks to participants or others and other events or information, as defined and listed in the Report Form, must be submitted promptly to the IRB. (See Events and Information that Require Prompt Reporting to the IRB at http://humansubjects.stanford.edu.) Upon completion, you must report to the IRB within 30 days.

Please remember that all data, including all signed consent form documents, must be retained for a minimum of three years past the completion of this research. Additional requirements may be imposed by your funding agency, your department, HIPAA, or other entities. (See Policy 1.9 on Retention of and Access to Research Data at http://doresearch.stanford.edu/policies/research-policy-handbook)

This institution is in compliance with requirements for protection of human subjects, including 45 CFR 46, 21 CFR 50 and 56, and 38 CFR 16.

Includes: Personnel change: removed Fridus Dhabhar

Waiver of Individual Authorization under 45 CFR 164.512(i)(2)(ii)(A),(B),(C), pursuant to information provided in the HIPAA section of the protocol application.

Ronald L. Ariagno, M.D., Chair

**Approval Period:** 10/31/2016 THROUGH 10/31/2017

**Review Type:** REGULAR - CONTINUING REVIEW

**Funding:** University of California Berkeley, Prime NIH , SPO: 110499

**Assurance #:** FWA00000935 (SU)

LELAND STANFORD JUNIOR UNIVERSITY
DIE LUFT DER FREIHEIT WEHT
1891

**STANFORD UNIVERSITY**

*Stanford, CA 94305 [Mail Code 5579]*

Ronald L. Ariagno, M.D. (650) 724-7541

CHAIR, PANEL ON MEDICAL HUMAN SUBJECTS (650) 725-8815

### Certification of Human Subjects Approvals

**Date:** October 31, 2016

**To:** Stephen P Luby, M.D., Medicine - Med/Infectious Diseases
Suzanne Mueller N/A, Amy Janel Pickering PhD

**From:** Ronald L. Ariagno, M.D., Administrative Panel on Human Subjects in Medical Research

**eProtocol Title:** Randomized Controlled Trial of the Benefits of Water Sanitation Hygiene Plus Nutrition Interventions on Child Growth.

**eProtocol #:** 25863 **IRB 4 (Registration #:** 351)

The IRB approved human subjects involvement in your research project on 10/31/2016. **'Prior to subject recruitment and enrollment, if this is: a Cancer-related study, you must obtain Cancer Center Scientific Review Committee (SRC) approval;** a CTRU study, you must obtain CTRU approval; a VA study, you must obtain VA R and D Committee approval; and if a contract is involved, it must be signed.'

The expiration date of this approval is 10/31/2017 at Midnight. If this research is to continue beyond that date, it is your responsibility to submit a Continuing Review application in eProtocol. Research activities must be reviewed and re-approved on or before midnight of the expiration date. The approval period may be less than one year if so determined by the IRB. Proposed changes to approved research must be reviewed and approved prospectively by the IRB. No changes may be initiated without prior approval by the IRB, except where necessary to eliminate apparent immediate hazards to subjects. (Any such exceptions must be reported to the IRB within 10 working days.) Unanticipated problems involving risks to participants or others and other events or information, as defined and listed in the Report Form, must be submitted promptly to the IRB. (See Events and Information that Require Prompt Reporting to the IRB at http://humansubjects.stanford.edu.) Upon completion, you must report to the IRB within 30 days.

Please remember that all data, including all signed consent form documents, must be retained for a minimum of three years past the completion of this research. Additional requirements may be imposed by your funding agency, your department, HIPAA, or other entities. (See Policy 1.9 on Retention of and Access to Research Data at http://doresearch.stanford.edu/policies/research-policy-handbook)

This institution is in compliance with requirements for protection of human subjects, including 45 CFR 46, 21 CFR 50 and 56, and 38 CFR 16.

Includes: Personnel change: removed Fridus Dhabhar

Waiver of Individual Authorization under 45 CFR 164.512(i)(2)(ii)(A),(B),(C), pursuant to information provided in the HIPAA section of the protocol application.

Ronald L. Ariagno, M.D., Chair

**Approval Period:** 10/31/2016 THROUGH 10/31/2017

**Review Type:** REGULAR - CONTINUING REVIEW

**Funding:** University of California Berkeley, Prime NIH , SPO: 108161

**Assurance #:** FWA00000935 (SU)

LELAND STANFORD JUNIOR UNIVERSITY
DIE LUFT DER FREIHEIT WEHT
1891

**Text S2. CONSORT checklist**

| Item | Description | Reported in Section |
|---|---|---|
| **Title and Abstract** | | |
| 1a | Identification as a randomized trial in the title; Identification as a cluster randomized trial in the title | Abstract |
| 1b | Structured summary of trial design, methods, results, and conclusions | Abstract |
| **Introduction** | | |
| Background and Objectives | | |
| 2a | Scientific background and explanation of rationale; Rationale for using a cluster design | Introduction |
| 2b | Specific objectives or hypotheses; Whether objectives pertain to the cluster level, the individual participant level, or both | Introduction |
| **Methods** | | |
| Trial Design | | |
| 3a | Description of trial design (such as parallel, factorial) including allocation ratio; Definition of cluster and description of how the design features apply to the clusters | Methods (Randomization and masking section) |
| 3b | Important changes to methods after trial commencement (such as eligibility criteria), with reasons | N/A |
| Participants | | |
| 4a | Eligibility criteria for participants; Eligibility criteria for clusters | Methods (Randomization and masking section) |
| 4b | Settings and locations where the data were collected | Methods (Study setting section) |
| Interventions | | |
| 5 | The interventions for each group with sufficient details to allow replication, including how and when they were actually administered; Whether interventions pertain to the cluster level, the individual participant level, or both | Methods (Interventions section) |
| Outcomes | | |
| 6a | Completely defined pre-specified primary and secondary outcome measures, including how and when they were assessed; Whether outcome measures pertain to the cluster level, the individual participant level, or both | Methods (Outcome assessment section) |

| Item | Description | Reported in Section |
|---|---|---|
| 6b | Any changes to trial outcomes after the trial commenced, with reasons | N/A |
| Sample Size | | |
| 7a | How sample size was determined; Method of calculation, number of cluster(s) (and whether equal or unequal cluster sizes are assumed), cluster size, a coefficient of intracluster correlation (ICC or k), and an indication of its uncertainty | Methods (Statistical analysis section) |
| 7b | When applicable, explanation of any interim analyses and stopping guidelines | N/A |
| **Randomization** | | |
| Sequence Generation | | |
| 8a | Method used to generate the random allocation sequence | Methods (Randomization and masking section) |
| 8b | Type of randomization; details of any restriction (such as blocking and block size); Details of stratification or matching if used | Methods (Randomization and masking section) |
| Allocation Concealment Mechanism | | |
| 9 | Mechanism used to implement the random allocation sequence (such as sequentially numbered containers), describing any steps taken to conceal the sequence until interventions were assigned; Specification that allocation was based on clusters rather than individuals and whether allocation concealment (if any) was at the cluster level, the individual participant level, or both | Methods (Randomization and masking section) |
| Implementation | | |
| 10a | Who generated the random allocation sequence, who enrolled clusters, and who assigned clusters to interventions | Methods (Randomization and masking section) |
| 10b | Mechanism by which individual participants were included in clusters for the purposes of the trial (such as complete enumeration, random sampling) | Methods (Randomization and masking section) |
| 10c | From whom consent was sought (representatives of the cluster, or individual cluster members, or both) and whether consent was sought before or after randomization | Methods (Ethics section) |
| Blinding | | |
| 11a | If done, who was blinded after assignment to interventions (for example, participants, care providers, those assessing outcomes) and how | Methods (Randomization and masking section) |

| Item | Description | Reported in Section |
|---|---|---|
| 11b | If relevant, description of the similarity of interventions | N/A |
| Statistical Methods | | |
| 12a | Statistical methods used to compare groups for primary and secondary outcomes; How clustering was taken into account | Methods (Statistical analysis section) |
| 12b | Methods for additional analyses, such as subgroup analyses and adjusted analyses | Methods (Statistical analyses section) |
| **Results** | | |
| Participant Flow | | |
| 13a | For each group, the numbers of participants/clusters who were randomly assigned, received intended treatment, and were analyzed for the primary outcome | Results (Enrollment section), Figure 1 (CONSORT diagram) |
| 13b | For each group, losses and exclusions after randomization, together with reasons, for both clusters and individual cluster members | Results (Enrollment section), Figure 1 (CONSORT diagram) |
| Recruitment | | |
| 14a | Dates defining the periods of recruitment and follow-up | Results (Enrollment section) |
| 14b | Why the trial ended or was stopped | N/A |
| Baseline Data | | |
| 15 | A table showing baseline demographic and clinical characteristics for each group; Baseline characteristics for the individual and cluster levels as applicable for each group | Table 1 |
| Numbers Analysed | | |
| 16 | For each group, number of participants/clusters (denominator) included in each analysis and whether the analysis was by the original assigned groups | Methods (Statistical analyses section), Figure 3, Figure 4 |
| Outcomes and Estimation | | |
| 17a | For each primary and secondary outcome, results for each group, and the estimated effect size and its precision (such as 95% confidence interval); Results at the individual and cluster levels as applicable and a coefficient of intracluster correlation (ICC or k) for each primary outcome | Results (Interventions vs. control section, Combined vs. single interventions section), Figure 3, Figure 4 |
| 17b | For binary outcome, presentation of both absolute and relative effect sizes is recommended | Results (Interventions vs. control section, Combined vs. single interventions section), Figure 3, Figure 4 |

| Item | Description | Reported in Section |
|---|---|---|
| Ancillary Analyses | | |
| 18 | Results of any other analyses performed, including subgroup analyses and adjusted analyses, distinguishing pre-specified from exploratory | Results (Other effects section, Subgroup analyses section) |
| Harms | | |
| 19 | All important harms or unintended effects in each group | N/A |
| **Discussion** | | |
| Limitations | | |
| 20 | Trial limitations, addressing sources of potential bias, imprecision and, if relevant, multiplicity of analyses | Discussion (Limitations section) |
| Generalisability | | |
| 21 | Generalisability (external validity, applicability) of the trial findings; Generalisability to clusters and/or individual participants (as relevant) | Discussion (Limitations section) |
| Interpretation | | |
| 22 | Interpretation consistent with results, balancing benefits and harms, and considering other relevant evidence | Discussion |
| **Other Information** | | |
| Registration | | |
| 23 | Registration number and name of trial registry | Methods (Registration section) |
| Protocol | | |
| 24 | Where the full trial protocol can be accessed, if available | Methods (Registration section) |
| Funding | | |
| 25 | Sources of funding and other support (such as supply of drugs), role of funders | Acknowledgements |

#### Text S3. Intervention promotion and user adherence

WASH Benefits included an extensive promotion effort to ensure user adherence with the interventions. The trial hired and trained local women to serve as community health promoters; these were hired specifically for WASH Benefits and were not involved in other ongoing programs (e.g., MDA) in the study area. Community health promoters were instructed to visit participants in intervention arms in their home at least bi-weekly throughout the trial and visited households six times per month on average. During these visits, they delivered and replenished the supply of intervention products, demonstrated their correct use, resolved hardware problems and encouraged their consistent use through discussion with participants. The targeted intervention adherence behaviors were: (1) children <36 months drinking chlorinated and safely stored water, (2) use of latrines for defecation and disposal of human and animal feces, (3) handwashing with soap at specified critical times (before food preparation, and after defecation and contact with feces) and (4) consumption of provided nutritional supplements and adoption of the promoted age-appropriate feeding behaviors. Control arm participants were not visited by community health promoters.

Intervention adherence was measured throughout the trial with multiple methods. Field staff conducted (1) unannounced spot check observations and a structured questionnaire in a random subset of 224 households per month over the course of 20 months starting after approximately two months of intervention, and (2) structured observations in a random subset of 324 households after approximately 15 months of intervention. The spot check observations assessed whether the provided intervention products were available, functional and had visible signs of use while the structured questionnaire recorded reported behaviors (e.g., water treatment, latrine use), and the structured observations included direct monitoring of the promoted behaviors. Stored water samples were tested for residual free chlorine in households reporting chlorine water treatment to validate the self-reported practices. Further details of the adherence assessment methods are described elsewhere [1].

All assessments showed high intervention adherence, both in terms of the observed status of the hardware and observed practices. In all arms with a sanitation component, 95-98% of households had a latrine with a functional water seal compared to 23% of controls [1]. Stool was observed on the latrine slab/floor or immediately outside the latrine in 24-38% of sanitation households vs. 62% of control households [1]. During structured observations, 94-97% of adults in sanitation arms were observed to defecate in a hygienic latrine compared to 40% of adults in the control arm [1]. In contrast, child defecation practices were similar between arms; in sanitation arms, 37-54% of young children were observed to defecate in a latrine or potty vs. 32% of control children [1]. In arms with a handwashing component, 77-85% of households had observed water and soap at the latrine and food preparation areas compared to 21% of controls [1]. During structured observations, 67-74% of participants in handwashing arms washed hands after latrine use vs. 29% of controls [1]. However, handwashing before food handling was infrequent (<10%) in all arms [1]. In all arms with a water treatment component, 50-70% of households had detectable residual free chlorine in their stored water; no household had chlorinated water in the control arm [1]. In 56-59% of households with the nutrition intervention, caregivers were observed to feed ≥1 nutritional supplement package to index children during five hours of structured observation [1]. Further details of adherence are described elsewhere [1].

**Text S4. Quality assurance for Kato-Katz**

A team of four laboratory technicians was trained at the parasitology laboratory of the International Centre for Diarrhoeal Disease Research, Bangladesh (icddr,b) for STH enumeration by Kato-Katz. 10% of Kato-Katz slides were counted by two technicians, and 5% were counted by a senior parasitologist for quality assurance. We assessed interrater agreement between two individual technicians as well as between a given technician and the senior parasitologist by calculating the kappa statistic for slides categorized as positive [2].

Technician 1 vs. 2

| ***A. lumbricoides*** | Not detected by technician 2 | Detected by technician 2 | Total |
|---|---|---|---|
| Not detected by technician 1 | 850 | 2 | 852 |
| Detected by technician 1 | 1 | 939 | 940 |
| Total | 851 | 941 | 1792 |

Kappa = 0.997

| **Hookworm** | Not detected by technician 2 | Detected by technician 2 | Total |
|---|---|---|---|
| Not detected by technician 1 | 1652 | 0 | 1652 |
| Detected by technician 1 | 2 | 138 | 140 |
| Total | 1654 | 138 | 1792 |

Kappa = 0.992

| ***T. trichiura*** | Not detected by technician 2 | Detected by technician 2 | Total |
|---|---|---|---|
| Not detected by technician 1 | 1645 | 1 | 1646 |
| Detected by technician 1 | 1 | 147 | 148 |
| Total | 1646 | 148 | 1794 |

Kappa = 0.993

Technician vs. senior parasitologist

| ***A. lumbricoides*** | Not detected by senior | Detected by senior | Total |
|---|---|---|---|
| Not detected by technician | 405 | 5 | 410 |
| Detected by technician | 25 | 291 | 316 |
| Total | 430 | 296 | 726 |

Kappa = 0.915

| **Hookworm** | Not detected by senior | Detected by senior | Total |
|---|---|---|---|
| Not detected by technician | 689 | 0 | 689 |
| Detected by technician | 30 | 4 | 34 |
| Total | 719 | 4 | 723 |

Kappa = 0.203

| ***T. trichiura*** | Not detected by senior | Detected by senior | Total |
|---|---|---|---|
| Not detected by technician | 664 | 2 | 666 |
| Detected by technician | 12 | 46 | 58 |
| Total | 676 | 48 | 724 |

Kappa = 0.858

#### Text S5. Bias in effect estimates with imperfect sensitivity and specificity under non-differential classification

Definitions:

| | |
|---|---|
| P1 | True probability of disease in intervention group |
| P0 | True probability of disease in control group |
| P1* | Measured probability of disease in intervention group |
| P0* | Measured probability of disease in control group |
| PR | True prevalence ratio between intervention vs. control (P1/P0) |
| PD | True prevalence difference between intervention vs. control (P1-P0) |
| PR* | Measured prevalence ratio between intervention vs. control (P1*/P0*) |
| PD* | Measured prevalence difference between intervention vs. control (P1*-P0*) |
| x1 | Probability of false negatives in intervention group |
| x0 | Probability of false negatives in control group |
| y1 | Probability of false positives in intervention group |
| y0 | Probability of false positives in control group |

$P1^* = P1 - P1\ x1 + (1-P1)\ y1$

$P0^* = P0 - P0\ x0 + (1-P0)\ y0$

Assumptions under non-differential misclassfication:

$x1 = x0 = x$

$y1 = y0 = y$

$P1^* = P1 - P1\ x + (1-P1)\ y$

$P0^* = P0 - P0\ x + (1-P0)\ y$

Scenario 1: Imperfect sensitivity, perfect specificity ($x > 0$, $y = 0$)

$$PR^* = \frac{P1 - P1\ x}{P0 - P0\ x} = \frac{P1\ (1 - x)}{P0\ (1 - x)} = \frac{P1}{P0} = PR$$

$PD^* = P1 - P1\ x - (P0 - P0\ x) = P1\ (1 - x) - P0\ (1 - x) = (1 - x)\ (P1 - P0) = (1 - x)\ PD$

Scenario 2: Perfect sensitivity, imperfect specificity (x = 0, y > 0)

$$PR^* = \frac{P1 + (1 - P1)\ y}{P0 + (1 - P0)\ y} = \frac{P1 + y - P1\ Y}{P0 + y - P0\ y} = \frac{P1\ (1 - y + y/P1)}{P0\ (1 - y + y/p0)} = \frac{P1}{P0} \cdot \frac{1 - y + y/P1}{1 - y + y/P0} = PR \cdot \frac{1 - y + y/P1}{1 - y + y/P0}$$

when P1 < P0, PR* > PR
when P1 > P0, PR* < PR

$$PD^* = P1 + (1 - P1)\ y - [\ P0 + (1 - P0)\ y\ ] = P1 + y - P1\ y - (P0 + y - P0y\ ) = P1\ (1 - y) - P0\ (1 - y) = (1 - y)\ (P1 - P0) = (1-y)\ PD$$

Scenario 3: Imperfect sensitivity, imperfect sensitivity (x > 0, y > 0)

$$PR^* = \frac{P1 - P1\ x + (1 - P1)\ y}{P0 - P0\ x + (1 - P0)\ y} = \frac{P1 - P1\ x + y - P1\ y}{P0 - P0\ x + y - P0\ y} = \frac{P1\ (1 - x - y + y/P1)}{P0\ (1 - x - y + y/P0)} = \frac{P1}{P0} \cdot \frac{1 - x - y + y/P1}{1 - x - y + y/P0} = PR \cdot \frac{1 - x - y + y/P1}{1 - x - y + y/P0}$$

when P1 < P0, PR* > PR
when P1 > P0, PR* < PR

$$PD^* = P1 - P1\ x + (1 - P1)\ y - [\ P0 - P0\ x + (1 - P0)\ y\ ] = P1 - P1\ x + y - P1\ y - (P0 - P0\ x + y - P0\ y) = P1\ (1 - x - y\ ) - P0\ (1 - x - y) = (1 - x - y\ )\ (P1 - P0) = (1 - x - y)\ PD$$

**Supplemental Information References**

1. Parvez SM, Azad R, Rahman M, Unicomb L, Ram PK, Naser AM, et al. Achieving optimal technology and behavioral uptake of single and combined interventions of water, sanitation hygiene and nutrition, in an efficacy trial (WASH benefits) in rural Bangladesh. Trials. 2018;19. doi:10.1186/s13063-018-2710-8

2. McHugh ML. Interrater reliability: the kappa statistic. Biochemia medica : Biochemia medica. 2012;22: 276–282.
