## Supplemental Tables for "Effects of water, sanitation, handwashing and nutritional interventions on soil-transmitted helminth infections in young children: a cluster-randomized controlled trial in rural Bangladesh"

### Supplemental Information Tables

Table S1: Enrollment characteristics of individuals with missing vs. observed outcomes
Table S2: Infection prevalence, all interventions vs. control
Table S3: Infection prevalence, combined vs. individual WSH interventions
Table S4: Infection prevalence, combined N+WSH vs. WSH and nutrition interventions
Table S5: Moderate/heavy infection prevalence, all interventions vs. control
Table S6: Moderate/heavy infection prevalence, combined vs. individual WSH interventions
Table S7: Moderate/heavy infection prevalence, combined N+WSH vs. WSH and nutrition interventions
Table S8: Fecal egg count reduction, all interventions vs. control
Table S9: Fecal egg count reduction, combined vs. individual WSH interventions
Table S10: Fecal egg count reduction, combined nutrition plus WSH vs. WSH and nutrition interventions

Table S1: Enrollment characteristics of individuals with missing vs. observed outcomes

| No. of individuals: | Missing (N=2824) | Observed (N=7187) |
|---|---|---|
| Index child, % | 16.3 | 17.7 |
| **Maternal** | | |
| Age, mean | 23.1 | 24.3 |
| Years of education, mean | 5.9 | 5.7 |
| **Paternal** | | |
| Years of education, mean | 4.9 | 4.7 |
| Works in agriculture, % | 28.1 | 33.1 |
| **Household** | | |
| Number of persons, mean | 4.6 | 4.8 |
| Has electricity, % | 56.0 | 59.3 |
| Has a cement floor, % | 12.2 | 9.6 |
| Acres of agricultural land owned, mean | 0.1 | 0.1 |
| **Drinking water** | | |
| Shallow tubewell primary water source, % | 72.0 | 74.0 |
| Stored water observed at home, % | 50.6 | 46.7 |
| Reported treating water yesterday, % | 0.0 | 0.2 |
| **Sanitation** | | |
| Daily defecating in the open, % | | |
| Adult men | 7.4 | 8.1 |
| Adult women | 3.6 | 4.9 |
| Children: 8-<15 years | 9.2 | 11.0 |
| Children: 3-<8 years | 36.7 | 38.0 |
| Children: 0-<3 years | 86.2 | 83.7 |
| Latrine, % | | |
| Owned | 51.7 | 51.5 |
| Concete slab | 90.4 | 88.8 |
| Functional water seal | 25.8 | 23.7 |
| Visible stool on slab or floor | 47.8 | 48.2 |
| Owned a potty, % | 5.7 | 4.5 |
| Human feces observed in, % | | |
| House | 7.8 | 9.3 |
| Child's play area | 1.4 | 1.2 |
| **Handwashing** | | |
| Has within 6 steps of latrine, % | | |
| Water | 12.6 | 10.0 |
| Soap | 7.8 | 5.1 |
| Has within 6 steps of kitchen, % | | |
| Water | 8.3 | 7.5 |
| Soap | 2.2 | 2.3 |

Table S2: Infection prevalence, all interventions vs. control

| Arm | N | Prevalence | Prevalence ratio | | | Prevalence difference | | |
|---|---|---|---|---|---|---|---|---|
| | | | Unadjusted | Adjusted [a] | IPCW [b] | Unadjusted | Adjusted [a] | IPCW [b] |
| **Ascaris** | | | | | | | | |
| Control | 1530 | 36.8% | | | | | | |
| Water | 971 | 35.8% | 0.97 (0.86, 1.11) | 0.98 (0.86, 1.11) | 0.97 (0.86, 1.10) | -0.96 (-5.54, 3.62) | -0.78 (-5.36, 3.80) | -0.93 (-5.32, 3.45) |
| Sanitation | 972 | 35.9% | 0.98 (0.88, 1.09) | 0.97 (0.87, 1.07) | 0.98 (0.88, 1.09) | -0.89 (-4.81, 3.03) | -1.28 (-5.13, 2.57) | -0.84 (-4.74, 3.07) |
| Handwashing | 977 | 40.5% | 1.10 (0.98, 1.24) | 1.10 (0.99, 1.23) | 1.10 (0.98, 1.24) | 3.73 (-0.79, 8.26) | 3.76 (-0.44, 7.96) | 3.71 (-0.84, 8.26) |
| WSH | 941 | 34.3% | 0.93 (0.83, 1.05) | 0.92 (0.82, 1.03) | 0.93 (0.83, 1.04) | -2.47 (-6.54, 1.60) | -2.91 (-6.88, 1.07) | -2.51 (-6.52, 1.50) |
| Nutrition | 863 | 40.3% | 1.10 (0.98, 1.23) | 1.12 (1.00, 1.25) | 1.11 (0.99, 1.23) | 3.53 (-1.06, 8.11) | 4.36 (0.05, 8.67) | 3.87 (-0.49, 8.23) |
| Nutrition + WSH | 933 | 33.0% | 0.90 (0.80, 1.01) | 0.90 (0.81, 1.01) | 0.90 (0.80, 1.00) | -3.79 (-7.74, 0.17) | -3.57 (-7.44, 0.30) | -3.86 (-7.71, -0.01) |
| **Hookworm** | | | | | | | | |
| Control | 1530 | 9.2% | | | | | | |
| Water | 971 | 6.4% | 0.69 (0.50, 0.95) | 0.73 (0.54, 0.99) | 0.71 (0.52, 0.98) | -2.83 (-5.16, -0.50) | -2.46 (-4.74, -0.18) | -2.67 (-5.11, -0.23) |
| Sanitation | 972 | 7.0% | 0.76 (0.54, 1.06) | 0.79 (0.58, 1.08) | 0.77 (0.56, 1.06) | -2.22 (-4.82, 0.38) | -1.89 (-4.30, 0.53) | -2.13 (-4.60, 0.34) |
| Handwashing | 977 | 8.3% | 0.90 (0.66, 1.22) | 0.92 (0.69, 1.21) | 0.88 (0.66, 1.18) | -0.93 (-3.56, 1.71) | -0.77 (-3.20, 1.66) | -1.10 (-3.58, 1.37) |
| WSH | 941 | 6.6% | 0.71 (0.52, 0.99) | 0.71 (0.53, 0.96) | 0.70 (0.51, 0.97) | -2.63 (-4.95, -0.31) | -2.65 (-4.83, -0.46) | -2.79 (-5.07, -0.52) |
| Nutrition | 863 | 9.5% | 1.03 (0.74, 1.43) | 1.02 (0.75, 1.40) | 1.02 (0.75, 1.38) | 0.29 (-2.77, 3.34) | 0.23 (-2.69, 3.14) | 0.19 (-2.70, 3.07) |
| Nutrition + WSH | 933 | 6.2% | 0.67 (0.50, 0.91) | 0.69 (0.52, 0.90) | 0.69 (0.52, 0.93) | -3.00 (-5.14, -0.85) | -2.89 (-4.88, -0.91) | -2.84 (-4.93, -0.75) |
| **Trichuris** | | | | | | | | |
| Control | 1530 | 7.5% | | | | | | |
| Water | 971 | 7.1% | 0.95 (0.68, 1.32) | 0.99 (0.72, 1.35) | 0.94 (0.67, 1.32) | -0.41 (-2.80, 1.98) | -0.10 (-2.39, 2.20) | -0.43 (-2.86, 2.00) |
| Sanitation | 972 | 5.3% | 0.71 (0.52, 0.98) | 0.71 (0.52, 0.96) | 0.71 (0.52, 0.98) | -2.17 (-4.10, -0.24) | -2.20 (-4.03, -0.38) | -2.19 (-4.09, -0.29) |
| Handwashing | 977 | 6.0% | 0.80 (0.59, 1.10) | 0.77 (0.57, 1.04) | 0.77 (0.57, 1.05) | -1.48 (-3.54, 0.58) | -1.74 (-3.68, 0.20) | -1.75 (-3.74, 0.24) |
| WSH | 941 | 6.4% | 0.85 (0.59, 1.22) | 0.85 (0.59, 1.21) | 0.83 (0.55, 1.25) | -1.14 (-3.50, 1.22) | -1.16 (-3.51, 1.20) | -1.30 (-3.95, 1.36) |
| Nutrition | 863 | 7.2% | 0.96 (0.69, 1.33) | 0.94 (0.68, 1.28) | 0.92 (0.66, 1.30) | -0.33 (-2.73, 2.07) | -0.48 (-2.76, 1.81) | -0.58 (-3.03, 1.87) |
| Nutrition + WSH | 933 | 9.1% | 1.21 (0.89, 1.65) | 1.23 (0.91, 1.67) | 1.22 (0.88, 1.69) | 1.59 (-1.13, 4.32) | 1.75 (-0.92, 4.42) | 1.64 (-1.27, 4.54) |
| **Any STH** | | | | | | | | |
| Control | 1530 | 43.4% | | | | | | |
| Water | 971 | 42.5% | 0.98 (0.87, 1.10) | 1.00 (0.89, 1.12) | 0.98 (0.88, 1.10) | -0.87 (-5.83, 4.10) | -0.11 (-4.99, 4.77) | -0.76 (-5.53, 4.00) |
| Sanitation | 972 | 40.6% | 0.94 (0.84, 1.04) | 0.94 (0.84, 1.04) | 0.94 (0.85, 1.04) | -2.76 (-7.11, 1.59) | -2.73 (-7.02, 1.56) | -2.76 (-6.99, 1.46) |
| Handwashing | 977 | 46.3% | 1.07 (0.96, 1.19) | 1.07 (0.97, 1.18) | 1.07 (0.96, 1.19) | 2.87 (-1.99, 7.72) | 2.93 (-1.51, 7.37) | 2.96 (-1.81, 7.74) |
| WSH | 941 | 39.3% | 0.91 (0.81, 1.01) | 0.89 (0.80, 0.99) | 0.90 (0.80, 1.00) | -4.08 (-8.53, 0.37) | -4.68 (-8.96, -0.41) | -4.56 (-9.03, -0.08) |
| Nutrition | 863 | 45.1% | 1.04 (0.93, 1.16) | 1.06 (0.96, 1.18) | 1.05 (0.94, 1.16) | 1.68 (-3.27, 6.62) | 2.66 (-2.02, 7.34) | 2.04 (-2.69, 6.76) |
| Nutrition + WSH | 933 | 38.8% | 0.89 (0.81, 0.99) | 0.90 (0.82, 0.99) | 0.89 (0.81, 0.98) | -4.60 (-8.56, -0.64) | -4.27 (-8.10, -0.44) | -4.74 (-8.54, -0.93) |

[a] Adjustment covariates considered include ID of the lab staff member who performed the Kato-Katz analysis, month of measurement, child age, sex and birthorder, mother's age, height and education, household food insecurity, number of children <18 years in household, number of individuals in compound, distance to the household's drinking water source, housing materials and assets. The adjusted model for each outcome includes covariates associated with the outcome at $p<0.2$ level in bivariate analysis.

[b] Inverse probability of censoring weighting. Adjustment covariates considered include the variables above except for ID of the lab staff member who performed the Kato-Katz analysis, month of measurement, child age, sex and birth order since this information is not available for individuals lost to follow-up. An indicator variable distinguishing index vs. non-index child status was included as a proxy for age.

Table S3: Infection prevalence, combined vs. individual WSH interventions

| Arm | N | Prevalence | Prevalence ratio | | | Prevalence difference | | |
|---|---|---|---|---|---|---|---|---|
| | | | Unadjusted | Adjusted [a] | IPCW [b] | Unadjusted | Adjusted [a] | IPCW [b] |
| **Ascaris** | | | | | | | | |
| WSH | 941 | 34.3% | | | | | | |
| Water | 971 | 35.8% | 0.96 (0.83, 1.10) | 0.95 (0.83, 1.08) | 0.95 (0.83, 1.09) | -1.51 (-6.36, 3.33) | -1.92 (-6.65, 2.82) | -1.68 (-6.36, 3.01) |
| Sanitation | 972 | 35.9% | 0.96 (0.83, 1.10) | 0.97 (0.84, 1.12) | 0.95 (0.83, 1.09) | -1.58 (-6.61, 3.45) | -0.95 (-5.95, 4.05) | -1.85 (-6.63, 2.92) |
| Handwashing | 977 | 40.5% | 0.85 (0.75, 0.96) | 0.84 (0.75, 0.96) | 0.84 (0.74, 0.96) | -6.21 (-10.96, -1.46) | -6.32 (-10.93, -1.71) | -6.50 (-11.29, -1.72) |
| **Hookworm** | | | | | | | | |
| WSH | 941 | 6.6% | | | | | | |
| Water | 971 | 6.4% | 1.03 (0.68, 1.57) | 0.98 (0.65, 1.47) | 1.00 (0.66, 1.52) | 0.20 (-2.52, 2.93) | -0.14 (-2.76, 2.49) | 0.01 (-2.74, 2.76) |
| Sanitation | 972 | 7.0% | 0.94 (0.62, 1.44) | 0.92 (0.61, 1.39) | 0.90 (0.60, 1.36) | -0.41 (-3.29, 2.47) | -0.57 (-3.34, 2.19) | -0.72 (-3.50, 2.07) |
| Handwashing | 977 | 8.3% | 0.79 (0.54, 1.16) | 0.76 (0.52, 1.11) | 0.80 (0.54, 1.18) | -1.70 (-4.46, 1.05) | -2.05 (-4.78, 0.68) | -1.62 (-4.46, 1.22) |
| **Trichuris** | | | | | | | | |
| WSH | 941 | 6.4% | | | | | | |
| Water | 971 | 7.1% | 0.90 (0.60, 1.33) | 0.87 (0.59, 1.28) | 0.87 (0.60, 1.28) | -0.73 (-3.33, 1.87) | -0.95 (-3.47, 1.57) | -0.93 (-3.48, 1.63) |
| Sanitation | 972 | 5.3% | 1.19 (0.76, 1.87) | 1.19 (0.77, 1.84) | 1.18 (0.75, 1.85) | 1.03 (-1.74, 3.79) | 1.03 (-1.66, 3.71) | 0.98 (-1.76, 3.72) |
| Handwashing | 977 | 6.0% | 1.06 (0.66, 1.68) | 1.08 (0.69, 1.68) | 1.06 (0.70, 1.61) | 0.34 (-2.59, 3.27) | 0.46 (-2.38, 3.29) | 0.36 (-2.30, 3.03) |
| **Any STH** | | | | | | | | |
| WSH | 941 | 39.3% | | | | | | |
| Water | 971 | 42.5% | 0.92 (0.81, 1.05) | 0.91 (0.80, 1.03) | 0.91 (0.80, 1.03) | -3.21 (-8.48, 2.05) | -3.99 (-9.06, 1.09) | -3.87 (-8.90, 1.17) |
| Sanitation | 972 | 40.6% | 0.97 (0.85, 1.10) | 0.97 (0.85, 1.11) | 0.96 (0.85, 1.09) | -1.32 (-6.55, 3.92) | -1.13 (-6.46, 4.20) | -1.68 (-6.68, 3.32) |
| Handwashing | 977 | 46.3% | 0.85 (0.76, 0.95) | 0.85 (0.76, 0.95) | 0.84 (0.75, 0.94) | -6.94 (-11.68, -2.20) | -7.18 (-11.88, -2.48) | -7.49 (-12.29, -2.69) |

[a] Adjustment covariates considered include ID of the lab staff member who performed the Kato-Katz analysis, month of measurement, child age, sex and birthorder, mother's age, height and education, household food insecurity, number of children <18 years in household, number of individuals in compound, distance to the household's drinking water source, housing materials and assets. The adjusted model for each outcome includes covariates associated with the outcome at p<0.2 level in bivariate analysis.

[b] Inverse probability of censoring weighting. Adjustment covariates considered include the variables above except for ID of the lab staff member who performed the Kato-Katz analysis, month of measurement, child age, sex and birth order since this information is not available for individuals lost to follow-up. An indicator variable distinguishing index vs. non-index child status was included as a proxy for age.

Table S4: Infection prevalence, combined nutrition plus WSH vs. WSH and nutrition interventions

| | | | Prevalence ratio | | | Prevalence difference | | |
|---|---|---|---|---|---|---|---|---|
| Arm | N | Prevalence | Unadjusted | Adjusted [a] | IPCW [b] | Unadjusted | Adjusted [a] | IPCW [b] |
| **Ascaris** | | | | | | | | |
| Nutrition + WSH | 933 | 33.0% | | | | | | |
| WSH | 941 | 34.3% | 0.96 (0.83, 1.11) | 0.99 (0.86, 1.13) | 0.98 (0.85, 1.12) | -1.31 (-6.15, 3.52) | -0.48 (-5.05, 4.08) | -0.84 (-5.41, 3.74) |
| Nutrition | 863 | 40.3% | 0.82 (0.72, 0.93) | 0.81 (0.71, 0.91) | 0.82 (0.72, 0.93) | -7.31 (-12.06, -2.57) | -7.81 (-12.38, -3.24) | -7.38 (-12.00, -2.77) |
| **Hookworm** | | | | | | | | |
| Nutrition + WSH | 933 | 6.2% | | | | | | |
| WSH | 941 | 6.6% | 0.94 (0.67, 1.32) | 0.98 (0.71, 1.37) | 0.99 (0.71, 1.37) | -0.37 (-2.55, 1.81) | -0.10 (-2.23, 2.02) | -0.07 (-2.22, 2.07) |
| Nutrition | 863 | 9.5% | 0.65 (0.47, 0.91) | 0.67 (0.49, 0.90) | 0.67 (0.50, 0.91) | -3.29 (-5.96, -0.61) | -3.12 (-5.61, -0.64) | -3.12 (-5.57, -0.68) |
| **Trichuris** | | | | | | | | |
| Nutrition + WSH | 933 | 9.1% | | | | | | |
| WSH | 941 | 6.4% | 1.43 (0.91, 2.25) | 1.48 (0.96, 2.28) | 1.44 (0.93, 2.24) | 2.73 (-0.61, 6.08) | 3.04 (-0.13, 6.21) | 2.82 (-0.44, 6.09) |
| Nutrition | 863 | 7.2% | 1.27 (0.86, 1.87) | 1.26 (0.87, 1.83) | 1.26 (0.89, 1.78) | 1.93 (-1.31, 5.16) | 1.90 (-1.20, 5.00) | 1.89 (-0.98, 4.76) |
| **Any STH** | | | | | | | | |
| Nutrition + WSH | 933 | 38.8% | | | | | | |
| WSH | 941 | 39.3% | 0.99 (0.87, 1.12) | 1.01 (0.89, 1.13) | 0.99 (0.88, 1.12) | -0.52 (-5.49, 4.45) | 0.24 (-4.42, 4.89) | -0.25 (-4.97, 4.47) |
| Nutrition | 863 | 45.1% | 0.86 (0.77, 0.97) | 0.86 (0.77, 0.96) | 0.86 (0.77, 0.96) | -6.28 (-11.15, -1.40) | -6.43 (-11.15, -1.70) | -6.41 (-11.13, -1.68) |

[a] Adjustment covariates considered include ID of the lab staff member who performed the Kato-Katz analysis, month of measurement, child age, sex and birthorder, mother's age, height and education, household food insecurity, number of children <18 years in household, number of individuals in compound, distance to the household's drinking water source, housing materials and assets. The adjusted model for each outcome includes covariates associated with the outcome at p<0.2 level in bivariate analysis.

[b] Inverse probability of censoring weighting. Adjustment covariates considered include the variables above except for ID of the lab staff member who performed the Kato-Katz analysis, month of measurement, child age, sex and birth order since this information is not available for individuals lost to follow-up. An indicator variable distinguishing index vs. non-index child status was included as a proxy for age.

Table S5: Moderate/heavy infection prevalence, all interventions vs. control

| Arm | N | Prevalence | Prevalence ratio: Unadjusted | Prevalence ratio: Adjusted [a] | Prevalence ratio: IPCW [b] | Prevalence difference: Unadjusted | Prevalence difference: Adjusted [a] | Prevalence difference: IPCW [b] |
|---|---|---|---|---|---|---|---|---|
| **Ascaris** | | | | | | | | |
| Control | 1530 | 4.2% | | | | | | |
| Water | 971 | 4.1% | 0.97 (0.65, 1.44) | 1.01 (0.69, 1.47) | 0.96 (0.66, 1.40) | -0.13 (-1.76, 1.51) | 0.04 (-1.55, 1.63) | -0.17 (-1.75, 1.42) |
| Sanitation | 972 | 3.9% | 0.92 (0.63, 1.35) | 0.95 (0.67, 1.34) | 0.91 (0.61, 1.35) | -0.34 (-1.89, 1.21) | -0.22 (-1.64, 1.19) | -0.38 (-1.94, 1.18) |
| Handwashing | 977 | 5.6% | 1.33 (0.94, 1.86) | 1.38 (1.01, 1.90) | 1.29 (0.92, 1.83) | 1.38 (-0.43, 3.19) | 1.60 (-0.11, 3.30) | 1.26 (-0.56, 3.09) |
| WSH | 941 | 3.3% | 0.78 (0.49, 1.22) | 0.73 (0.45, 1.18) | 0.72 (0.44, 1.17) | -0.95 (-2.54, 0.63) | -1.15 (-2.76, 0.47) | -1.21 (-2.85, 0.43) |
| Nutrition | 863 | 4.2% | 0.98 (0.67, 1.45) | 1.04 (0.72, 1.49) | 1.01 (0.68, 1.50) | -0.08 (-1.71, 1.56) | 0.15 (-1.40, 1.70) | 0.05 (-1.64, 1.74) |
| Nutrition + WSH | 933 | 6.1% | 1.44 (0.96, 2.16) | 1.49 (1.03, 2.15) | 1.50 (1.01, 2.22) | 1.86 (-0.45, 4.17) | 2.05 (-0.09, 4.20) | 2.10 (-0.24, 4.45) |
| **Hookworm** | | | | | | | | |
| Control | 1530 | 0.1% | | | | | | |
| Water | 971 | 0.0% | – [c] | – [c] | – [c] | -0.13 (-0.39, 0.13) | -0.13 (-0.40, 0.14) | -0.14 (-0.36, 0.09) |
| Sanitation | 972 | 0.2% | 1.57 (0.18, 13.83) | 1.57 (0.17, 14.61) | 1.53 (0.20, 11.43) | 0.08 (-0.26, 0.41) | 0.08 (-0.27, 0.42) | 0.07 (-0.26, 0.39) |
| Handwashing | 977 | 0.1% | 0.78 (0.06, 9.46) | 0.79 (0.06, 10.29) | 0.77 (0.07, 8.01) | -0.03 (-0.32, 0.27) | -0.03 (-0.34, 0.28) | -0.03 (-0.32, 0.26) |
| WSH | 941 | 0.2% | 1.63 (0.13, 19.90) | 1.64 (0.13, 20.30) | 1.62 (0.18, 14.25) | 0.08 (-0.35, 0.52) | 0.08 (-0.35, 0.52) | 0.08 (-0.31, 0.47) |
| Nutrition | 863 | 0.3% | 2.66 (0.21, 33.94) | 2.74 (0.21, 35.30) | 2.77 (0.32, 24.20) | 0.22 (-0.29, 0.72) | 0.23 (-0.27, 0.72) | 0.23 (-0.24, 0.71) |
| Nutrition + WSH | 933 | 0.0% | – [c] | – [c] | – [c] | -0.13 (-0.41, 0.15) | -0.13 (-0.42, 0.16) | -0.13 (-0.34, 0.07) |
| **Trichuris** | | | | | | | | |
| Control | 1530 | 0.4% | | | | | | |
| Water | 971 | 0.4% | 1.05 (0.27, 4.14) | 1.07 (0.28, 4.15) | 1.02 (0.25, 4.23) | 0.02 (-0.53, 0.57) | 0.03 (-0.53, 0.58) | 0.01 (-0.56, 0.58) |
| Sanitation | 972 | 0.2% | 0.52 (0.08, 3.63) | 0.53 (0.08, 3.47) | 0.53 (0.08, 3.52) | -0.19 (-0.69, 0.31) | -0.18 (-0.67, 0.30) | -0.18 (-0.68, 0.32) |
| Handwashing | 977 | 0.1% | 0.26 (0.03, 2.04) | 0.26 (0.04, 1.98) | 0.27 (0.03, 2.34) | -0.29 (-0.70, 0.12) | -0.29 (-0.68, 0.11) | -0.29 (-0.71, 0.14) |
| WSH | 941 | 0.7% | 1.90 (0.53, 6.81) | 1.87 (0.52, 6.69) | 1.82 (0.46, 7.31) | 0.35 (-0.40, 1.11) | 0.34 (-0.42, 1.11) | 0.34 (-0.50, 1.17) |
| Nutrition | 863 | 0.5% | 1.18 (0.34, 4.13) | 1.13 (0.33, 3.91) | 1.11 (0.27, 4.55) | 0.07 (-0.46, 0.60) | 0.05 (-0.48, 0.58) | 0.04 (-0.55, 0.63) |
| Nutrition + WSH | 933 | 1.1% | 2.73 (0.90, 8.31) | 2.71 (0.91, 8.07) | 2.64 (0.78, 8.94) | 0.68 (-0.11, 1.47) | 0.68 (-0.11, 1.46) | 0.64 (-0.23, 1.52) |
| **Any STH** | | | | | | | | |
| Control | 1530 | 4.5% | | | | | | |
| Water | 971 | 4.5% | 1.00 (0.71, 1.43) | 1.05 (0.75, 1.47) | 1.00 (0.71, 1.41) | 0.02 (-1.57, 1.62) | 0.23 (-1.32, 1.77) | -0.01 (-1.59, 1.57) |
| Sanitation | 972 | 4.2% | 0.94 (0.64, 1.37) | 0.97 (0.69, 1.36) | 0.92 (0.63, 1.34) | -0.29 (-1.93, 1.35) | -0.15 (-1.63, 1.33) | -0.36 (-1.98, 1.26) |
| Handwashing | 977 | 5.7% | 1.27 (0.90, 1.80) | 1.31 (0.95, 1.82) | 1.24 (0.87, 1.76) | 1.22 (-0.66, 3.10) | 1.40 (-0.39, 3.19) | 1.08 (-0.83, 2.98) |
| WSH | 941 | 3.8% | 0.85 (0.55, 1.31) | 0.82 (0.52, 1.28) | 0.80 (0.50, 1.28) | -0.68 (-2.41, 1.04) | -0.83 (-2.58, 0.92) | -0.91 (-2.71, 0.90) |
| Nutrition | 863 | 4.8% | 1.05 (0.71, 1.57) | 1.12 (0.77, 1.62) | 1.08 (0.72, 1.62) | 0.24 (-1.62, 2.10) | 0.51 (-1.26, 2.28) | 0.37 (-1.56, 2.30) |
| Nutrition + WSH | 933 | 6.6% | 1.47 (0.97, 2.25) | 1.51 (1.04, 2.21) | 1.53 (1.02, 2.29) | 2.14 (-0.44, 4.71) | 2.30 (-0.06, 4.67) | 2.37 (-0.21, 4.95) |

[a] Adjustment covariates considered include ID of the lab staff member who performed the Kato-Katz analysis, month of measurement, child age, sex and birthorder, mother's age, height and education, household food insecurity, number of children <18 years in household, number of individuals in compound, distance to the household's drinking water source, housing materials and assets. The adjusted model for each outcome includes covariates associated with the outcome at p<0.2 level in bivariate analysis.

[b] Inverse probability of censoring weighting. Adjustment covariates considered include the variables above except for ID of the lab staff member who performed the Kato-Katz analysis, month of measurement, child age, sex and birth order since this information is not available for individuals lost to follow-up. An indicator variable distinguishing index vs. non-index child status was included as a proxy for age.

[c] Could not calculate due to sparse data.

Table S6: Moderate/heavy infection prevalence, combined vs. individual WSH interventions

| | | | Prevalence ratio | | | Prevalence difference | | |
|---|---|---|---|---|---|---|---|---|
| Arm | N | Prevalence | Unadjusted | Adjusted [a] | IPCW [b] | Unadjusted | Adjusted [a] | IPCW [b] |
| **Ascaris** | | | | | | | | |
| WSH | 941 | 3.3% | | | | | | |
| Water | 971 | 4.1% | 0.80 (0.47, 1.36) | 0.74 (0.42, 1.30) | 0.77 (0.44, 1.34) | -0.83 (-2.75, 1.10) | -1.10 (-3.10, 0.90) | -0.97 (-2.96, 1.01) |
| Sanitation | 972 | 3.9% | 0.84 (0.47, 1.50) | 0.80 (0.45, 1.42) | 0.79 (0.44, 1.44) | -0.62 (-2.67, 1.44) | -0.80 (-2.80, 1.20) | -0.80 (-2.84, 1.24) |
| Handwashing | 977 | 5.6% | 0.59 (0.34, 1.00) | 0.54 (0.32, 0.92) | 0.56 (0.34, 0.93) | -2.34 (-4.56, -0.11) | -2.62 (-4.74, -0.50) | -2.49 (-4.57, -0.42) |
| **Hookworm** | | | | | | | | |
| WSH | 941 | 0.2% | | | | | | |
| Water | 971 | 0.0% | – [c] | – [c] | – [c] | 0.21 (-0.06, 0.49) | 0.41 (0.09, 0.72) | 0.23 (-0.09, 0.56) |
| Sanitation | 972 | 0.2% | 1.03 (0.11, 9.49) | 1.20 (0.16, 9.04) | 1.10 (0.11, 10.70) | 0.01 (-0.46, 0.47) | 0.04 (-0.40, 0.48) | 0.02 (-0.47, 0.51) |
| Handwashing | 977 | 0.1% | 2.08 (0.22, 19.55) | 2.24 (0.25, 20.09) | 2.10 (0.21, 20.53) | 0.11 (-0.26, 0.48) | 0.12 (-0.25, 0.50) | 0.11 (-0.26, 0.49) |
| **Trichuris** | | | | | | | | |
| WSH | 941 | 0.7% | | | | | | |
| Water | 971 | 0.4% | 1.81 (0.41, 8.04) | 1.83 (0.42, 7.87) | 1.82 (0.42, 7.87) | 0.33 (-0.57, 1.23) | 0.34 (-0.55, 1.24) | 0.34 (-0.54, 1.22) |
| Sanitation | 972 | 0.2% | 3.62 (0.55, 23.77) | 3.64 (0.57, 23.40) | 3.70 (0.42, 32.64) | 0.54 (-0.30, 1.38) | 0.54 (-0.30, 1.38) | 0.55 (-0.30, 1.41) |
| Handwashing | 977 | 0.1% | 7.27 (0.75, 70.31) | 7.35 (0.77, 69.90) | 7.41 (0.97, 56.68) | 0.64 (-0.14, 1.43) | 0.66 (-0.12, 1.44) | 0.67 (-0.03, 1.37) |
| **Any STH** | | | | | | | | |
| WSH | 941 | 3.8% | | | | | | |
| Water | 971 | 4.5% | 0.84 (0.50, 1.41) | 0.78 (0.45, 1.35) | 0.81 (0.48, 1.37) | -0.71 (-2.81, 1.40) | -1.00 (-3.16, 1.17) | -0.88 (-3.02, 1.26) |
| Sanitation | 972 | 4.2% | 0.91 (0.54, 1.54) | 0.87 (0.52, 1.47) | 0.86 (0.51, 1.47) | -0.39 (-2.50, 1.71) | -0.53 (-2.56, 1.49) | -0.57 (-2.64, 1.49) |
| Handwashing | 977 | 5.7% | 0.67 (0.40, 1.12) | 0.62 (0.37, 1.05) | 0.64 (0.39, 1.04) | -1.91 (-4.25, 0.44) | -2.17 (-4.43, 0.08) | -2.07 (-4.27, 0.12) |

[a] Adjustment covariates considered include ID of the lab staff member who performed the Kato-Katz analysis, month of measurement, child age, sex and birthorder, mother's age, height and education, household food insecurity, number of children <18 years in household, number of individuals in compound, distance to the household's drinking water source, housing materials and assets. The adjusted model for each outcome includes covariates associated with the outcome at p<0.2 level in bivariate analysis.

[b] Inverse probability of censoring weighting. Adjustment covariates considered include the variables above except for ID of the lab staff member who performed the Kato-Katz analysis, month of measurement, child age, sex and birth order since this information is not available for individuals lost to follow-up. An indicator variable distinguishing index vs. non-index child status was included as a proxy for age.

[c] Could not calculate due to sparse data.

Table S7: Moderate/heavy infection prevalence, combined nutrition plus WSH vs. WSH and nutrition interventions

| | | | Prevalence ratio | | | Prevalence difference | | |
|---|---|---|---|---|---|---|---|---|
| Arm | N | Prevalence | Unadjusted | Adjusted [a] | IPCW [b] | Unadjusted | Adjusted [a] | IPCW [b] |
| **Ascaris** | | | | | | | | |
| Nutrition + WSH | 933 | 6.1% | | | | | | |
| WSH | 941 | 3.3% | 1.85 (1.09, 3.16) | 2.02 (1.21, 3.39) | 2.02 (1.18, 3.47) | 2.81 (0.40, 5.23) | 3.15 (0.90, 5.41) | 3.16 (0.77, 5.55) |
| Nutrition | 863 | 4.2% | 1.46 (0.89, 2.42) | 1.42 (0.88, 2.29) | 1.46 (0.87, 2.45) | 1.94 (-0.67, 4.55) | 1.79 (-0.69, 4.26) | 1.96 (-0.76, 4.68) |
| **Hookworm** | | | | | | | | |
| Nutrition + WSH | 933 | 0.0% | | | | | | |
| WSH | 941 | 0.2% | – [c] | – [c] | – [c] | -0.21 (-0.50, 0.08) | -0.26 (-0.52, 0.00) | -0.22 (-0.54, 0.09) |
| Nutrition | 863 | 0.3% | – [c] | – [c] | – [c] | -0.35 (-0.74, 0.05) | -0.36 (-0.74, 0.02) | -0.35 (-0.72, 0.03) |
| **Trichuris** | | | | | | | | |
| Nutrition + WSH | 933 | 1.1% | | | | | | |
| WSH | 941 | 0.7% | 1.44 (0.50, 4.17) | 1.39 (0.50, 3.86) | 1.41 (0.54, 3.68) | 0.33 (-0.59, 1.25) | 0.30 (-0.58, 1.18) | 0.31 (-0.53, 1.15) |
| Nutrition | 863 | 0.5% | 2.31 (0.95, 5.65) | 2.26 (0.94, 5.44) | 2.27 (0.90, 5.72) | 0.61 (-0.08, 1.29) | 0.59 (-0.07, 1.25) | 0.60 (-0.13, 1.33) |
| **Any STH** | | | | | | | | |
| Nutrition + WSH | 933 | 6.6% | | | | | | |
| WSH | 941 | 3.8% | 1.74 (1.06, 2.84) | 1.88 (1.17, 3.01) | 1.87 (1.14, 3.07) | 2.82 (0.30, 5.34) | 3.17 (0.84, 5.51) | 3.15 (0.66, 5.65) |
| Nutrition | 863 | 4.8% | 1.40 (0.85, 2.29) | 1.33 (0.83, 2.11) | 1.39 (0.85, 2.29) | 1.89 (-0.98, 4.77) | 1.60 (-1.09, 4.29) | 1.89 (-1.01, 4.80) |

[a] Adjustment covariates considered include ID of the lab staff member who performed the Kato-Katz analysis, month of measurement, child age, sex and birthorder, mother's age, height and education, household food insecurity, number of children <18 years in household, number of individuals in compound, distance to the household's drinking water source, housing materials and assets. The adjusted model for each outcome includes covariates associated with the outcome at p<0.2 level in bivariate analysis.

[b] Inverse probability of censoring weighting. Adjustment covariates considered include the variables above except for ID of the lab staff member who performed the Kato-Katz analysis, month of measurement, child age, sex and birth order since this information is not available for individuals lost to follow-up. An indicator variable distinguishing index vs. non-index child status was included as a proxy for age.

[c] Could not calculate due to sparse data.

Table S8: Fecal egg count reduction, all interventions vs. control

| | | | Geometric FECR [a] | | | Arithmetic FECR [a] | | |
|---|---|---|---|---|---|---|---|---|
| Arm | N | Geo-mean | Unadjusted | Adjusted [b] | IPCW [c] | Unadjusted | Adjusted [c] | IPCW [c] |
| **Ascaris** | | | | | | | | |
| Control | 1530 | 5.2 | | | | | | |
| Water | 971 | 5.0 | -0.04 (-0.30, 0.21) | -0.02 (-0.27, 0.24) | -0.04 (-0.28, 0.20) | -0.22 (-0.67, 0.23) | -0.21 (-0.65, 0.23) | -0.24 (-0.65, 0.16) |
| Sanitation | 972 | 4.8 | -0.08 (-0.27, 0.12) | -0.06 (-0.26, 0.13) | -0.08 (-0.28, 0.12) | -0.29 (-0.66, 0.07) | -0.28 (-0.63, 0.07) | -0.29 (-0.67, 0.08) |
| Handwashing | 977 | 7.6 | 0.38 (0.01, 0.75) | 0.40 (0.04, 0.76) | 0.37 (0.00, 0.74) | 0.44 (-0.31, 1.20) | 0.40 (-0.31, 1.10) | 0.33 (-0.38, 1.03) |
| WSH | 941 | 4.4 | -0.14 (-0.34, 0.07) | -0.15 (-0.35, 0.05) | -0.15 (-0.36, 0.05) | -0.17 (-0.71, 0.37) | -0.23 (-0.78, 0.31) | -0.25 (-0.80, 0.31) |
| Nutrition | 863 | 6.3 | 0.16 (-0.12, 0.45) | 0.23 (-0.06, 0.52) | 0.18 (-0.10, 0.46) | 0.22 (-0.70, 1.13) | 0.25 (-0.67, 1.16) | 0.21 (-0.66, 1.07) |
| Nutrition + WSH | 933 | 5.1 | -0.02 (-0.28, 0.24) | 0.02 (-0.24, 0.28) | 0.00 (-0.26, 0.27) | 0.73 (-0.27, 1.73) | 0.80 (-0.19, 1.80) | 0.79 (-0.21, 1.79) |
| **Hookworm** | | | | | | | | |
| Control | 1530 | 0.6 | | | | | | |
| Water | 971 | 0.4 | -0.13 (-0.23, -0.03) | -0.11 (-0.21, -0.01) | -0.12 (-0.23, -0.02) | -0.41 (-0.78, -0.04) | -0.38 (-0.79, 0.04) | -0.43 (-0.79, -0.06) |
| Sanitation | 972 | 0.4 | -0.09 (-0.22, 0.03) | -0.08 (-0.19, 0.04) | -0.09 (-0.21, 0.03) | -0.09 (-0.77, 0.60) | -0.09 (-0.79, 0.62) | -0.12 (-0.76, 0.52) |
| Handwashing | 977 | 0.5 | -0.04 (-0.16, 0.09) | -0.03 (-0.15, 0.09) | -0.04 (-0.16, 0.08) | -0.13 (-0.76, 0.51) | -0.11 (-0.79, 0.57) | -0.12 (-0.73, 0.49) |
| WSH | 941 | 0.4 | -0.11 (-0.22, 0.00) | -0.11 (-0.21, -0.00) | -0.11 (-0.22, -0.00) | -0.06 (-0.81, 0.69) | -0.10 (-0.84, 0.63) | -0.08 (-0.82, 0.66) |
| Nutrition | 863 | 0.6 | 0.02 (-0.14, 0.17) | 0.02 (-0.13, 0.17) | 0.02 (-0.13, 0.16) | 0.63 (-0.95, 2.20) | 0.61 (-0.97, 2.19) | 0.61 (-0.84, 2.06) |
| Nutrition + WSH | 933 | 0.3 | -0.14 (-0.23, -0.04) | -0.14 (-0.23, -0.05) | -0.13 (-0.23, -0.04) | -0.55 (-0.88, -0.22) | -0.56 (-0.90, -0.21) | -0.55 (-0.85, -0.24) |
| **Trichuris** | | | | | | | | |
| Control | 1530 | 0.4 | | | | | | |
| Water | 971 | 0.4 | -0.00 (-0.13, 0.12) | 0.01 (-0.11, 0.13) | -0.00 (-0.13, 0.12) | 0.12 (-0.63, 0.87) | 0.16 (-0.60, 0.92) | 0.12 (-0.65, 0.89) |
| Sanitation | 972 | 0.3 | -0.10 (-0.19, -0.01) | -0.10 (-0.18, -0.01) | -0.10 (-0.19, -0.01) | -0.31 (-0.96, 0.33) | -0.32 (-0.94, 0.31) | -0.33 (-0.94, 0.28) |
| Handwashing | 977 | 0.3 | -0.09 (-0.18, 0.00) | -0.10 (-0.19, -0.01) | -0.10 (-0.19, -0.01) | -0.51 (-0.86, -0.16) | -0.54 (-0.86, -0.21) | -0.54 (-0.89, -0.18) |
| WSH | 941 | 0.4 | -0.03 (-0.17, 0.10) | -0.03 (-0.16, 0.11) | -0.04 (-0.19, 0.11) | 0.59 (-0.81, 1.99) | 0.59 (-0.82, 2.01) | 0.57 (-1.00, 2.15) |
| Nutrition | 863 | 0.4 | -0.02 (-0.14, 0.10) | -0.02 (-0.13, 0.09) | -0.03 (-0.15, 0.10) | -0.05 (-0.71, 0.61) | -0.06 (-0.69, 0.57) | -0.09 (-0.78, 0.60) |
| Nutrition + WSH | 933 | 0.6 | 0.10 (-0.06, 0.26) | 0.11 (-0.05, 0.27) | 0.10 (-0.07, 0.27) | 1.34 (-0.31, 3.00) | 1.31 (-0.36, 2.98) | 1.30 (-0.74, 3.35) |

[a] Faecal egg count reduction (FECR) defined as egg ratio (ER) - 1, where ER is the ratio of mean eggs per gram between arms.
[b] Adjustment covariates considered include ID of the lab staff member who performed the Kato-Katz analysis, month of measurement, child age, sex and birthorder, mother's age, height and education, household food insecurity, number of children <18 years in household, number of individuals in compound, distance to the household's drinking water source, housing materials and assets. The adjusted model for each outcome includes covariates associated with the outcome at p<0.2 level in bivariate analysis.
[c] Inverse probability of censoring weighting. Adjustment covariates considered include the variables above except for ID of the lab staff member who performed the Kato-Katz analysis, month of measurement, child age, sex and birth order since this information is not available for individuals lost to follow-up. An indicator variable distinguishing index vs. non-index child status was included as a proxy for age.

Table S9: Fecal egg count reduction, combined vs. individual WSH interventions

| Arm | N | Geo-mean | Geometric FECR [a] | | | Arithmetic FECR [a] | | |
|---|---|---|---|---|---|---|---|---|
| | | | Unadjusted | Adjusted [b] | IPCW [c] | Unadjusted | Adjusted [c] | IPCW [c] |
| **Ascaris** | | | | | | | | |
| WSH | 941 | 4.4 | | | | | | |
| Water | 971 | 5.0 | -0.10 (-0.36, 0.17) | -0.14 (-0.39, 0.11) | -0.13 (-0.38, 0.12) | 0.06 (-0.69, 0.81) | -0.00 (-0.74, 0.74) | 0.01 (-0.75, 0.78) |
| Sanitation | 972 | 4.8 | -0.06 (-0.34, 0.21) | -0.06 (-0.33, 0.21) | -0.07 (-0.33, 0.19) | 0.17 (-0.69, 1.04) | 0.10 (-0.73, 0.93) | 0.12 (-0.83, 1.08) |
| Handwashing | 977 | 7.6 | -0.37 (-0.57, -0.18) | -0.38 (-0.57, -0.20) | -0.39 (-0.57, -0.21) | -0.43 (-0.88, 0.03) | -0.43 (-0.90, 0.03) | -0.43 (-0.86, 0.00) |
| **Hookworm** | | | | | | | | |
| WSH | 941 | 0.4 | | | | | | |
| Water | 971 | 0.4 | 0.02 (-0.12, 0.16) | 0.00 (-0.13, 0.14) | 0.01 (-0.13, 0.16) | 0.60 (-0.57, 1.78) | 0.54 (-0.61, 1.69) | 0.59 (-0.69, 1.87) |
| Sanitation | 972 | 0.4 | -0.02 (-0.17, 0.14) | -0.02 (-0.17, 0.12) | -0.03 (-0.18, 0.12) | 0.03 (-0.93, 1.00) | -0.01 (-0.94, 0.91) | 0.00 (-0.97, 0.97) |
| Handwashing | 977 | 0.5 | -0.07 (-0.20, 0.06) | -0.09 (-0.22, 0.04) | -0.07 (-0.21, 0.06) | 0.08 (-0.73, 0.89) | 0.02 (-0.79, 0.82) | 0.04 (-0.79, 0.86) |
| **Trichuris** | | | | | | | | |
| WSH | 941 | 0.4 | | | | | | |
| Water | 971 | 0.4 | -0.03 (-0.17, 0.12) | -0.04 (-0.18, 0.10) | -0.04 (-0.18, 0.10) | 0.41 (-0.99, 1.82) | 0.42 (-0.98, 1.83) | 0.41 (-0.95, 1.77) |
| Sanitation | 972 | 0.3 | 0.08 (-0.09, 0.24) | 0.07 (-0.09, 0.24) | 0.07 (-0.09, 0.24) | 1.32 (-1.30, 3.93) | 1.38 (-1.27, 4.02) | 1.35 (-1.58, 4.28) |
| Handwashing | 977 | 0.3 | 0.06 (-0.12, 0.24) | 0.07 (-0.10, 0.25) | 0.07 (-0.09, 0.23) | 2.24 (-1.29, 5.77) | 2.46 (-1.31, 6.23) | 2.44 (-0.91, 5.78) |

[a] Faecal egg count reduction (FECR) defined as egg ratio (ER) - 1, where ER is the ratio of mean eggs per gram between arms.

[b] Adjustment covariates considered include ID of the lab staff member who performed the Kato-Katz analysis, month of measurement, child age, sex and birthorder, mother's age, height and education, household food insecurity, number of children <18 years in household, number of individuals in compound, distance to the household's drinking water source, housing materials and assets. The adjusted model for each outcome includes covariates associated with the outcome at p<0.2 level in bivariate analysis.

[c] Inverse probability of censoring weighting. Adjustment covariates considered include the variables above except for ID of the lab staff member who performed the Kato-Katz analysis, month of measurement, child age, sex and birth order since this information is not available for individuals lost to follow-up. An indicator variable distinguishing index vs. non-index child status was included as a proxy for age.

Table S10: Fecal egg count reduction, combined nutrition plus WSH vs. WSH and nutrition interventions

| Arm | N | Geo-mean | Geometric FECR [a] Unadjusted | Geometric FECR [a] Adjusted [b] | Geometric FECR [a] IPCW [c] | Arithmetic FECR [a] Unadjusted | Arithmetic FECR [a] Adjusted [b] | Arithmetic FECR [a] IPCW [c] |
|---|---|---|---|---|---|---|---|---|
| **Ascaris** | | | | | | | | |
| Nutrition + WSH | 933 | 5.1 | | | | | | |
| WSH | 941 | 4.4 | 0.14 (-0.21, 0.49) | 0.21 (-0.14, 0.55) | 0.18 (-0.16, 0.53) | 1.09 (-0.50, 2.68) | 1.35 (-0.41, 3.12) | 1.33 (-0.48, 3.14) |
| Nutrition | 863 | 6.3 | -0.16 (-0.41, 0.10) | -0.18 (-0.42, 0.07) | -0.17 (-0.42, 0.08) | 0.42 (-0.60, 1.45) | 0.41 (-0.59, 1.41) | 0.43 (-0.57, 1.42) |
| **Hookworm** | | | | | | | | |
| Nutrition + WSH | 933 | 0.3 | | | | | | |
| WSH | 941 | 0.4 | -0.03 (-0.14, 0.07) | -0.03 (-0.13, 0.07) | -0.03 (-0.13, 0.08) | -0.52 (-0.81, -0.23) | -0.51 (-0.81, -0.22) | -0.51 (-0.81, -0.22) |
| Nutrition | 863 | 0.6 | -0.15 (-0.27, -0.04) | -0.14 (-0.26, -0.03) | -0.15 (-0.26, -0.04) | -0.72 (-0.94, -0.51) | -0.72 (-0.94, -0.51) | -0.72 (-0.93, -0.52) |
| **Trichuris** | | | | | | | | |
| Nutrition + WSH | 933 | 0.6 | | | | | | |
| WSH | 941 | 0.4 | 0.14 (-0.06, 0.34) | 0.16 (-0.03, 0.35) | 0.14 (-0.05, 0.34) | 0.48 (-1.01, 1.96) | 0.48 (-0.98, 1.94) | 0.46 (-0.93, 1.85) |
| Nutrition | 863 | 0.4 | 0.13 (-0.06, 0.31) | 0.13 (-0.04, 0.30) | 0.13 (-0.04, 0.29) | 1.46 (-0.53, 3.45) | 1.42 (-0.50, 3.35) | 1.45 (-0.76, 3.65) |

[a] Faecal egg count reduction defined as egg ratio (ER) - 1, where ER is the ratio of mean eggs per gram between arms.

[b] Adjustment covariates considered include ID of the lab staff member who performed the Kato-Katz analysis, month of measurement, child age, sex and birthorder, mother's age, height and education, household food insecurity, number of children <18 years in household, number of individuals in compound, distance to the household's drinking water source, housing materials and assets. The adjusted model for each outcome includes covariates associated with the outcome at p<0.2 level in bivariate analysis.

[c] Inverse probability of censoring weighting. Adjustment covariates considered include the variables above except for ID of the lab staff member who performed the Kato-Katz analysis, month of measurement, child age, sex and birth order since this information is not available for individuals lost to follow-up. An indicator variable distinguishing index vs. non-index child status was included as a proxy for age.
